# Single paternal Dexamethasone challenge programs offspring metabolism and reveals circRNAs as novel candidates in RNA-mediated inheritance

**DOI:** 10.1101/2021.02.10.429888

**Authors:** Katharina Gapp, Guillermo Parada, Fridolin Gross, Alberto Corcoba, Evelyn Grau, Martin Hemberg, Johannes Bohacek, Eric A. Miska

## Abstract

Single traumatic events that elicit an exaggerated stress response can lead to the development of neuropsychiatric conditions. Studies in mice suggests germline RNA as a mediator of effects of chronic environmental exposures to the progeny. The effects of an acute paternal stress exposure on the germline and their potential consequences on offspring remain unknown. We find that acute administration of an agonist for the stress-sensitive Glucocorticoid receptor, using the common corticosteroid Dexamethasone, affects the RNA payload of post-meiotic transcriptionally silent, mature sperm as soon as 3 hours post exposure. It further impacts early embryonic transcriptional trajectories, as determined by single embryo sequencing, and metabolism in the offspring. Importantly, we show persistent regulation of tRNA fragments in sperm and the descendant 2-cell- embryos, suggesting actual transmission from sperm to embryo. Lastly, we unravel environmentally induced alterations in the previously underconsidered class of sperm circRNAs, and their targets in the early embryo, highlighting this class as a novel candidate in RNA-mediated inheritance.

## Introduction

Acute stress elicits a complex but well-studied cascade of neuroendocrine responses regulated by the hypothalamic pituitary adrenal axis. It involves the release of neuropeptides in the brain that induce the secretion of corticosteroid hormones from the adrenals. These hormones in turn activate mainly two types of nuclear glucocorticoid receptors (GR) (*1*) expressed throughout the body, which then regulate gene expression and enable physiological and behavioral adjustments (*2*). In vulnerable individuals, this response is excessive and it can lead to long lasting maladaptations with consequences on psychological and metabolic health (*3*).

It is also known that parental experiences can compromise the health of their progeny both in humans (*4–6*) and in animal models (7, 8, 17–26, 9, 27, 10–16). Research on the underlying mechanism of such transmission has found changes in germline epigenetic make- up, in particular DNA methylation, histone post translational modifications (PTMs), histone positioning and RNA (*28*). These epigenetic regulators are responsive to the environment and have been implicated in a variety of environmentally induced diseases (*29*). Altered modifications must circumvent epigenetic reprogramming events in zygote and, depending on the timing of exposure, during germline development (*28, 30*). In the male germline, RNA is excluded from reprogramming and therefore a promising candidate for transgenerational information delivery (*28, 31*). Several studies carried out in *D. melangolaster* and *C. elegans* reported on transgenerational inheritance of induced traits and provided firm evidence for the involvement of small RNAs in the mechanism of transmission (*32–34*). In mammals, a causal implication in the transmission of environmentally induced effects across generations has also solely been demonstrated for sperm RNA (7, 24, 27, 35). Such RNA differs substantially from somatic RNA since it mainly consists of small RNA, predominantly tRNA- derived small fragments (tsRNAs), but also miRNAs, piRNAs and circRNAs, among others (*31*)(*28, 36*). CircRNAs, comprise a very stable class of RNA that has recently been observed to be present in high amounts in testis but also to some extent in sperm (*37*). Some have been shown to act as miRNA sponges, thereby competing with mRNA targets, while also regulating the expression of their host genes (*37*). Hence, circRNAs have a strong potential for amplifying an inherited signal, which makes circRNAs exceptionally interesting candidates for epigenetic germline inheritance. To date, the involvement of circRNAs in soma-to-germline signalling has not yet been investigated.

tsRNAs and miRNAs are crucial regulators of early embryonic development and players in non-genetic inheritance (7, 10, 24, 27, 35, 38, 39). They have been reported to be acquired through exosomal uptake during epididymal transfer from caput to cauda epididymis (16, 44). This might explain their responsiveness to environmental perturbations, despite mature sperm’s presumably transcriptionally silent state caused by tightly packed chromatin. Sperm RNA can indeed change in response to chronic stress or by chronic treatments that mimic stress exposure, such as repeated injection of GR agonists (7, 23, 41, 42). Uptake of epididysomal miRNA was sufficient to replicate a chronic stress induced effect on stress response in offspring mice(*43*). Surprisingly, acute stress has also recently been shown to affect offspring weight and glucose metabolism in mice (*44*) and some of these effects were germline dependent (*45*). Together these related lines of evidence led us to hypothesize that acute GR activation has an intergenerational effect on offspring phenotype and that the transmission potentially implicates changes in the germline. The male germline cells - including mature sperm (*46, 47*) and their surrounding Sertoli cells (*48*) – as well es the epididymal epithelial cells (*49*) express GRs that mediate the effects of glucocorticoids on transcription. Dexamethasone (Dex) a specific GR agonist is known to directly activate GR in the rat epididymis (*50*). It is unknown whether acute stress affects sperm RNA, and if so, whether uptake via epididymisomes is involved in establishing germline changes that are relevant for offspring phenotypic alterations.

Here we investigate the impact of acute GR agonist administration on the germline RNA payload including circRNAs, at various time points post administration and interrogate the fate of altered sperm RNA. We further test germ-line dependency of transmitted metabolic effects and dissect the underlying molecular trajectories during early embryonic development using single cell sequencing of in vitro fertilization (IVF) derived embryos. Identifying a readout of transgenerational risk load at the level of the paternal sperm epigenome could pave the way for future studies aiming at a prevention of the transmission of the effects of acute GR activation to the offspring.

## Results

### Effects of acute Dex injection on the germline small RNA payload

Two reports have suggested that a single foot shock could elicit effects on offspring phenotype (*44, 45*).To examine potential epigenetic mediators of such acute stressful impacts we investigated sperm RNA of males 2 weeks after a single activation of the GR (Figure 1A). This timeline was chosen to mimic the timing at which breeding occurred when effects on offspring had been observed in a previous study (*45*). We injected the specific GR agonist Dex once intraperitoneally into 8 adult males. This drug is in frequent clinical use, now also as an apparently effective treatment for patients suffering from lower respiratory tract infection as a consequence of Covid-19 virus (*51–53*). A sperm population was harvested from each animal and RNA was extracted for ultra-deep small RNA sequencing, resulting in 16 libraries representing one injected male each (8 vehicle and 8 Dex-injected). Purity of the sperm samples was confirmed by inspecting RNA size profiles generated on the bioanalyzer to be absent of ribosomal RNA peaks, that would indicate contamination by somatic cells (Supplementary Fig. 1A). Reaching an average of 55.4 Million sequencing reads while also using randomized adaptors for 3′ ligation put us in a position to reduce PCR biases (*54*) and accurately quantify less abundant miRNAs that are by far outnumbered in sperm by other small RNAs e.g. tsRNAs(*55*). Our data showed an average of 60% mappable reads across all libraries, including 34% of multimappers. We detected an expected dominant prevalence of reads mapping to tsRNAs and abundant miRNAs in all samples (Supplementary Fig. 2A). Differential gene expression analysis, using DEseq2 (*56*), revealed that a single acute activation of GR receptors induced changes in tsRNAs and miRNAs collected 14 days post injection (Figure 1B,C, FDR q<0.05), as has been observed in response to chronic environmental stress previously (*7, 42*). Interestingly, tsRNA-Gly-GCC, a tsRNA previously associated with the effects of nutritional challenge (*24*), was among the most strongly altered tsRNAs.

**Figure 1.**
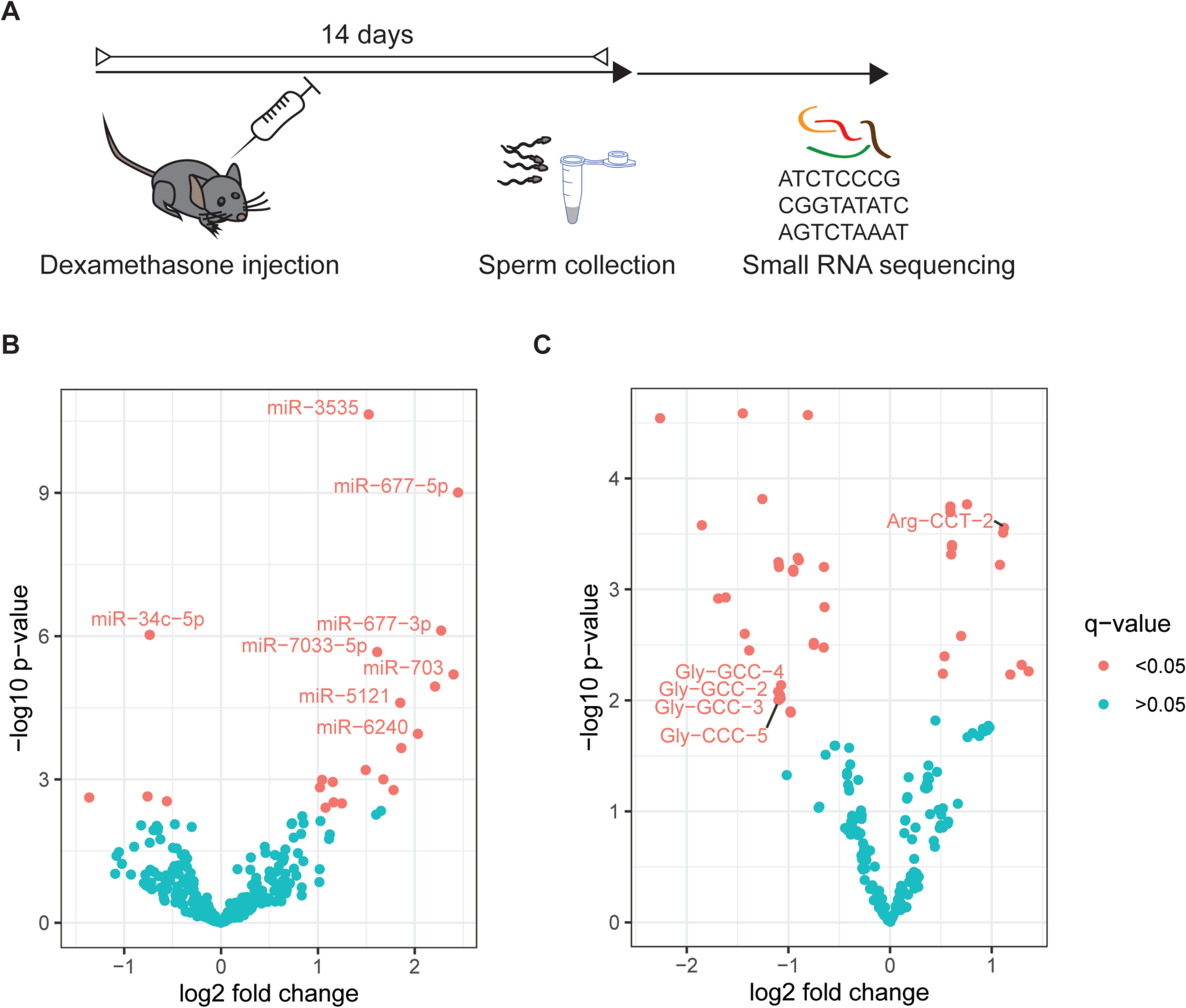
Effects of Dex on small RNA payload of sperm cells residing in testis at time of administration. (A) Experimental design depicting a time window of two weeks between injection of Dex and sperm collection for molecular analysis. (B) Volcano plot depicting fold changes and significance level of miRNAs and tsRNAs (C) in mature sperm 14 days post injection of Dex (n=8) versus vehicle (n=8) as assessed by small RNA sequencing.

Some recent publications have suggested that sperm miRNAs and tsRNAs are acquired during epididymal transit from caput to cauda (*24, 40, 57*). Further, it was shown that chronic nutritional-challenge-induced change in sperm tsRNAs is acquired by uptake of distinct sets of tsRNAs (*24*). To investigate whether the changes in small RNA after acute stress are due to uptake during transit from caput to cauda epididymis we investigated the mature sperm small RNA payload at two time points, 3 hours and 7 days following injection (Figure 2A, Supplementary Fig. 1C). Cells collected from cauda 7 days after injection had already exited testis, but have had time to pass through the entire epididymal tract before collection. Cells collected 3 hours post injection will most likely not have passed through the corpus epididiymis, and already resided in cauda epididymis at the time of injection where sperm resides up to 5 days (*58, 59*). Importantly, spontaneous ejaculation regularly voids cauda epididymis of sperm, even in the absence of a mating partner(*60*), excluding the retention of “old” mature sperm in cauda for prolonged periods of time. The cells collected 7 days after exposure therefore represent a mixture of cells that might have already resided in the cauda and those cells that indeed passed through the corpus epididymis, yet the spontaneous ejaculation ensures that the sample predominantly contains the latter.

**Figure 2.**
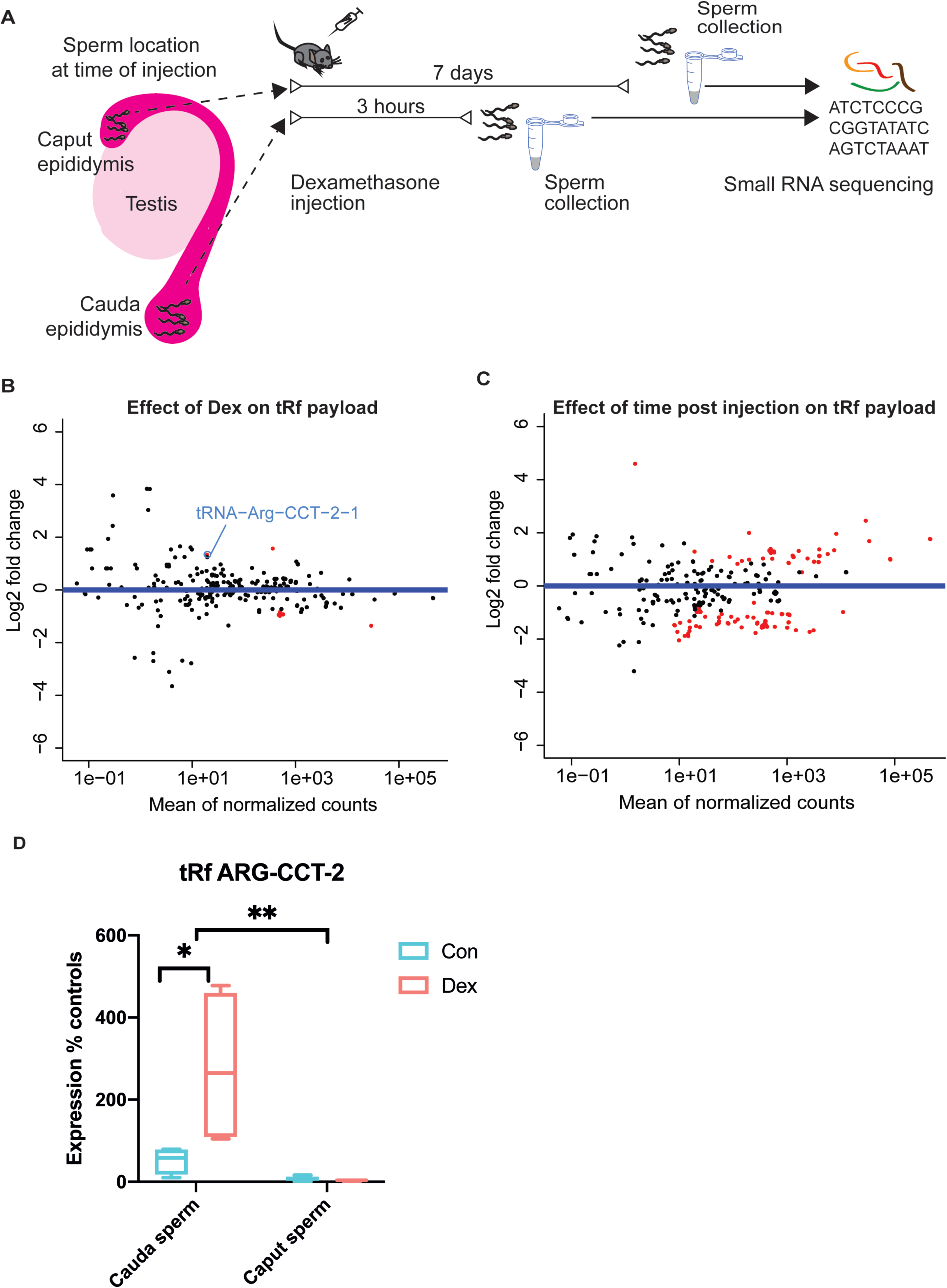
Effect of Dex on sperm cells at different time points post Dex administration. (A) Experimental design showing the location of sperm at time of injection and timing of sperm harvest. MA (log-intensity ratios (M-values) versus log-intensity averages (A-values) plots depicting (B) effect of Dex (log2 fold changes control versus dexamethasone), (C) of time post injection (log2 fold changes 7 days versus 3 hours) (7 days Dex n =4 and controls n =4, 3 hours Dex n =3 and controls n =4). TsRNAs are indicated by sequence identity for display only, each dot represents one small RNA. MA plot depicts log2 fold changes on the y axis and the expression level on the x axis (the higher the expression the further to the right). Statistically significantly changed small RNAs are highlighted in red. (D) Relative expression of ArgCCT-2 as obtained by q-RT-PCR (cauda: Dex n=4, controls n=4, caput: Dex n=4, controls n=5; interaction F(1, 13)=6.34, p=0.0257, treatment F(1, 13)=5.97, p=0.0040, site of collection (F(1,13)=12.15, p=0.0296; cauda control versus cauda Dex t(13)=3.42, p=0.0274, cauda Dex verus caput Dex t(13)=4.137, p=0.007). Whiskers display minimum and maximum.

The collected samples were confirmed for their purity (Supplementary Fig. 1) and again processed separately to represent sperm from one animal per library. The resulting libraries were analysed jointly as to test for (1) effects of Dex injection independent of sampling time post injeciton (2) effects of sampling time post injection independent of Dex treatment and (3) effects depending on both Dex injection and the sampling time post injection (interaction). We report an average of 64% mappable reads including 46% of multimappers and observe that tsRNAs were significantly affected by sampling time post injection independent of treatment. This demonstrates the fluctuation of tsRNAs over time in response to external signals such as injections, or potentially due to uncontrollable external inputs from the animal husbandry (Figure 2C). Interaction between treatment and time was statistically significant for 27 tsRNA mapping loci including Gly-GCC-6-1, showing all upregulation after 7 days but either no regulation (26 tsRNAs) or downregulation (Thr-TGT1-1) after 3 hours (Supplementary Table2 sheet 3, q<0.05). This finding is consistent with the dominating view that tsRNAs are acquired during epididymal transit from caput to cauda epididymis. However, most tsRNAs that showed a significant change in response to treatment not after 3 hours but after 7 days (interaction between treatment and time post injection, Supplementary Table 2 sheet 3, q<0.05) were not persistently altered in the dataset of 14 days after injection (Supplementary Fig. 3). This indicates that on the one hand changes in sperm RNA are dynamic and many do not persist for prolonged time. On the other hand, this suggests that potentially relevant small RNA changes mostly require either sperm to reside in testis at exposure time or rely on a prolonged residency in the exposed organism. Interestingly, tsRNA-Leu-CAA and tsRNA-Arg-CCT, (Figure 2B) were persistently affected 3 hours and 7 days post exposure, that necessarily requires a mode of rapid acquisition of tsRNA-changes in cauda epididymis. While the change in tsRNA-Leu-CAA was temporary and did not persist, strikingly tsRNA-Arg-CCT-2 deregulation persisted until 14 days post injection (Figure 1C). To additionally validate the Dex induced change of tsRNA-Arg-CCT-2 independent of epididymal transit from caput to cauda we replicated the effect observed in mature sperm sampled from cauda epididymis 3 hours post injection using q-PCR (Figure 2D, Supplementary table 2). Additionally, we sampled caput sperm 3 hours post injection and measured tsRNA-Arg-CCT-2 levels. Overall 2-way ANOVA revealed a significant interaction between sperm sampling location (caput versus cauda) and treatment (vehicle versus Dex). Post hoc tests confirm a significant increase in tsRNA-Arg-CCT-2 levels in response to Dex in cauda but not in caput sperm and a significant increase in Arg-CCT-2 levels between cauda and caput sperm independent of treatment.

The behaviour of miRNAs differed considerably from tsRNAs. As would be expected if epididymal transit was required for miRNA changes to be implemented, we observe no group effect of treatment on miRNAs (Supplementary Fig. 4A, Supplementary table 2) across 3 hours and 7 days post injection. Further, we detected no effect of time post injection on sperm miRNA payload (Supplementary Fig. 4B, Supplementary table 2) confirming the absence of an effect of injection on miRNAs per se. However, we neither detected an interaction between Dex and time post injection (Supplementary Fig. 4C, Supplementary table 2) in miRNAs 7 days and 3 hours post injection. Importantly, when inspecting those miRNAs that were significantly altered 14 days after injection, no alterations were apparent 3 hours or 7 days post injection (Supplementary Fig. 4D), indicating that changes in miRNAs occur more slowly or require sperm cells to reside in testis at the time of injection.

### Effects of acute GR activation on in vivo offspring metabolic phenotype

Based on the two reports on effects of single foot shock on offspring weight and the impact of a single GR activation on germline small RNA payload, we hypothesized that this acute impact on the receptor is sufficient to elicit transgenerational effects. We thus injected Dex once intraperitoneally, then harvested sperm 14 days post injection, and performed *IVF* using naïve oocytes to generate offspring for phenotyping (Figure 3A). Dex treatment did not affect sperm count, fertility rate or resulting litter-sizes (Supplemenatary Fig.10-12).

**Figure 3.**
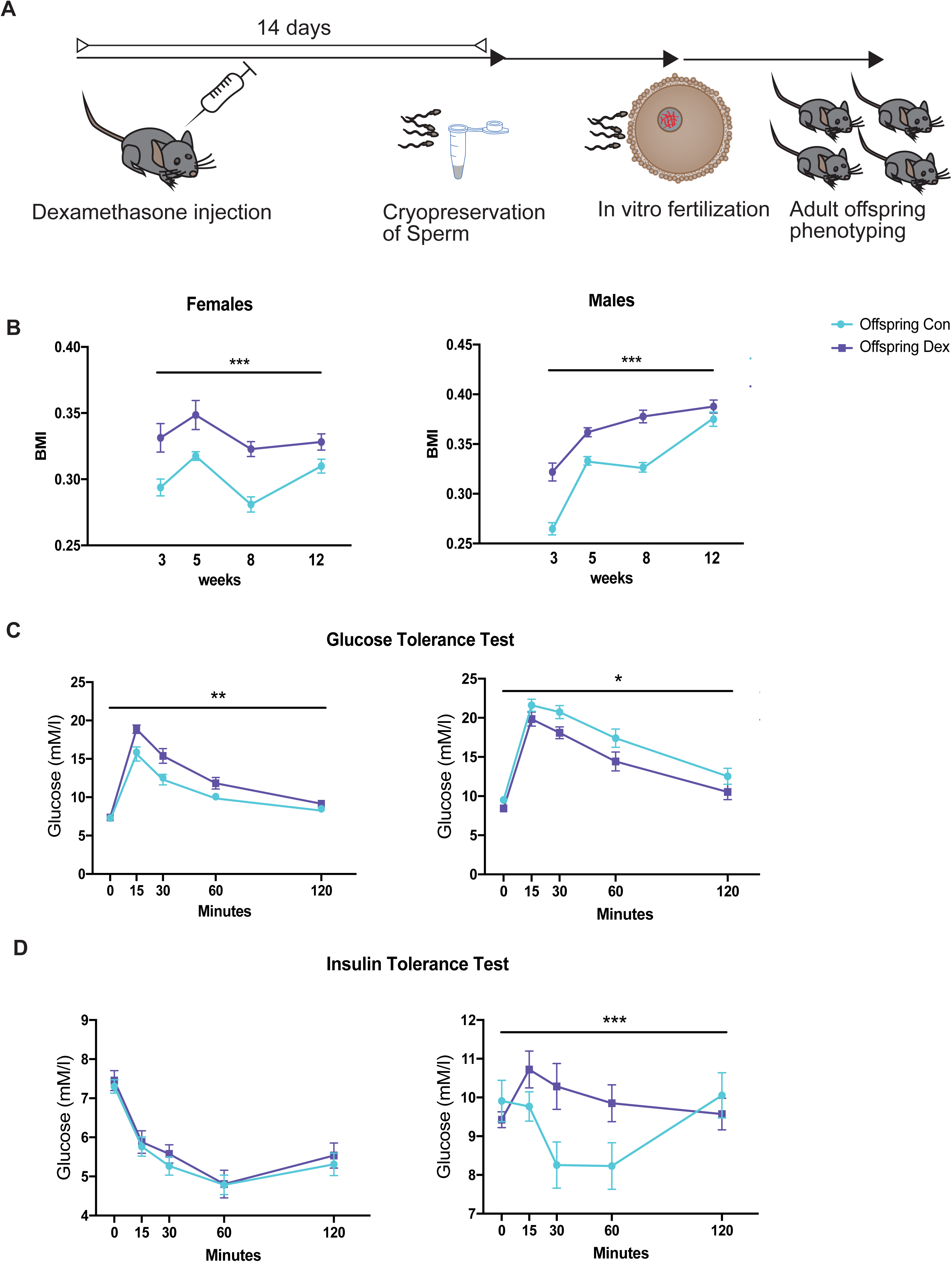
Effect of Dex on metabolic phenotype in the offspring (A) Experimental design depicting timeline between injection, sperm harvest, in vitro fertilization and phenotyping. (B) Impact of Dex on male and female adult offspring (B) Body mass index (males vehicle offspring n=21, Dex offspring n=22, females vehicle offspring n=17, Dex offspring n=17) (C) glucose tolerance (males vehicle offspring n=12, Dex offspring n=12, females vehicle offspring n=12, Dex offspring n=12) and (D) insulin tolerance (males vehicle offspring n=9, Dex offspring n=8, females vehicle offspring n=12, Dex offspring n=12). Error bars display SEM. Detailed statistical results are depicted in Supplementary Fig. 5, raw data are provided in supplementary table 3).

The weight and size of pups was measured every 2 to 4 weeks starting at weaning (3 weeks of age) until adulthood (12 weeks of age) and body mass index (BMI) was calculated as a ratio of weight and squared length. Overall ANOVA of the resulting offspring showed a significant effect of treatment ((F1.71)= 76.55, p<0.0001), time post injection ((F2.087,144.7)=41.99, p<0.0001) and sex (F (1, 71) = 76.55, p<0.0001) on BMI, and a significant interaction between time and sex (F (3, 208) = 33.75, p<0.0001) and time and treatment (F (3, 208) = 5.834, p=0.0008) (Figure 3B, Supplementary Fig. 5C, Supplementary table 3). These results show that while males had generally higher BMI, both male and female offspring of Dex injected fathers had a higher BMI.

To further explore potential causes and consequences of altered BMI, adult animals were additionally tested for their glucose tolerance following glucose injection. Overall ANOVA analysis of blood glucose levels revealed a significant effect of sex (F (1, 44) = 54.80, p<0.0001) and time post injection (F (2.593, 114.1) = 196.6, p<0.0001) and significant interactions between sex and time post injection (F (4, 176) = 6.115, P=0.0001), and sex and treatment (F (1, 44) = 15.62, P=0.0003) (Supplementary Fig. 5C). Follow up repeated measurements ANOVA analysis separated by sex showed a significant effect of treatment, time and interaction in females (treatment: F (1, 22) = 12.35, p=0.0020; time: F (4, 88) = 110.1, p<0.0001; interaction: F (4, 88) = 2.835, p=0.0291) and significant effects of treatment and time but no interaction in males (treatment: F (1, 22) = 6.019, p=0.0225; time: F (4, 88) = 96.36, p<0.0001; F (4, 88) = 0.5401, p=0.7067; Figure 3C). These data hence demonstrate a sex-dependent effect of paternal Dex injection on glucose tolerance, with impaired tolerance in females and decreased glucose levels in males in response to glucose challenge.

In addition, blood glucose levels were assessed during the insulin tolerance test. Overall ANOVA analysis showed significant effects of sex (F (1, 37) = 162.6, P<0.0001) and time (F (3.314, 122.6) = 23.85, P<0.0001) and revealed a significant interaction between sex and time (F (4, 148) = 12.49, P<0.0001), time and treatment (F (4, 148) = 5.380, P=0.0005) and time and treatment and sex (F (4, 148) = 5.392, P=0.0004) (Supplementary Fig. 5C). Follow-up repeated measurements ANOVA separated by sex showed a significant effect of time (F (2.982, 65.60) = 44.73, p<0.0001), yet no significant effect of treatment (F (1, 22) = 0.3465, p=0.5621) nor an interaction between time and treatment (F (4, 88) = 0.1373, p=0.9681) in females (Figure 3D). In males we observe no effect of treatment F (1, 15) = 1.467, p=0.2446 yet detected a significant effect of time post injection time: F (2.914, 43.71) = 4.538, p=0.0079, and a significant interaction between treatment and time post injection F (4, 60) = 7.003, p=0.0001, Figure 3D). These results indicate sex and time dependent effects of paternal Dex on insulin tolerance. They further show no change in insulin tolerance in female descendants of fathers injected with Dex, whereas impaired insulin tolerance in male progeny.

Lastly, we explored a potential reflection of altered BMI in tissue composition by necropsy and weighing the dissected organs and fat pads. Overall ANOVA of necropsy weights revealed a significant effect of sex (F (1, 140) = 28.27, P<0.0001), tissue (F (4, 140) = 232.7, P<0.0001) and a significant interaction between sex and tissue (F (4, 140) = 3.379, P=0.0113) yet no effect of treatment (F (1, 140) = 0.2587, P=0.6118), or interaction between treatment and sex (F (1, 140) = 0.0004794, P=0.9826) or treatment and tissue (F (4, 140) = 0.1635, P=0.9565) on tissue weight (Supplementary Fig. 5A,B,C). This confirms sex dependency, yet no effect of paternal Dex injection on tissue weight in both sexes.

### Effects of acute Dex on offspring early embryonic small RNA

The small quantity of paternal RNAs in the zygote relative to the large pool of maternal RNAs poses serious obstacles to their accurate quantification (36). While initial reports on small RNA transmission relied on comparative sequencing or microarrays analyses of unfertilized oocytes and fertilized zygotes (*61*), today we are aware that such comparisons can be deceiving, as they rely heavily on both assessment method (eg. microarray restricted to a selective set versus unbiased genome-wide sequencing) and sequencing depth (*54, 62*). An example are inconsistent results regarding miRNAs that are exclusively supplied from the sperm, such as miR-34c, -99a, -214 (*63, 64*). Alternative approaches have used indirect measures, e.g. assessing mRNA targets of paternally derived small RNAs (*24, 64–66*). We attempted to directly examine the relative difference between the small RNA landscape in early embryos resulting from IVF of naïve oocytes with sperm from either Dex or vehicle injected males (Figure 4A). We used small-RNA sequencing to compare 2-cell embryos derived from Dex treated or control fathers. We detected an average of 29 % mappable reads including 21% multimappers. While we only detected subtle changes in miRNAs of Dex exposed progeny (Supplementary Fig. 6), we observed downregulation of several tsRNAs from 6 different genomic locations (q<0.1) (Figure 4B). Strikingly, two of the downregulated tsRNAs (Gly-GCC at several genomic loci and Gly-CCC) were consistently downregulated in sperm 14 days post Dex injection. This likely indicates a direct delivery of these tsRNAs in control conditions, yet absence or reduced delivery of this sperm RNA cargo of Dex treated males to the oocytes they fertilize.

**Figure 4.**
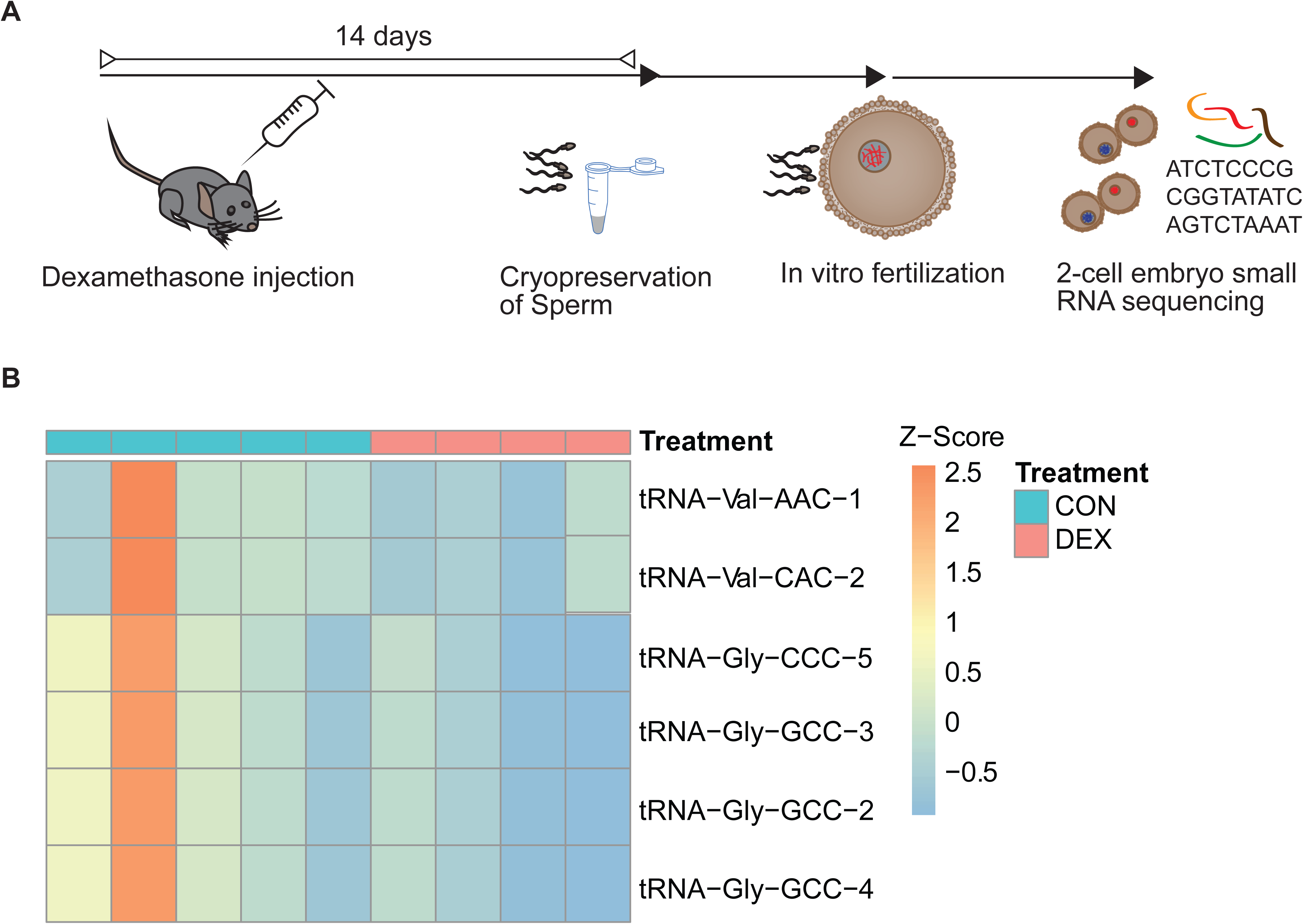
Effect of paternal Dex injection on embryonic offspring small RNA. (A) Experimental design depicting timeline between injection, sperm harvest, in vitro fertilization and small RNA sequencing at 2-cell stage. (B) Volcano plot showing effect of paternal Dex on small RNA tsRNAs (vehicle embryonic offspring n=5, Dex embryonic offspring n=4). TsRNAs are grouped by sequence identity for display only.

### Effects of acute Dex administration on offspring early embryonic transcriptome

If sperm RNA was directly impacting the zygotic mRNA pool, or affecting early embryonic gene expression we hypothesized this to be apparent in the 2 cell embryo’s transcriptome (Figure 5A). To examine the effect of paternal Dex on early embryonic RNA content we subjected 2-cell embryos to the Smartseq single cell sequencing protocol (Supplementary Fig. 7). After performing quality control and filtering the sequenced 2 cell embryo data on criteria such as minimal read count/embryo (Supplementary Fig. 7A), we carried out unsupervised clustering based on their gene expression profiles using SC3 (*67*). We identified two distinct clusters (C1 and C2), which were composed by a balanced mixture of treated and control cells. (Figure 5B). Since the resolution of single cell experiments allows characterizing distinctive transcriptomic profiles within early cell division stages, we used scmap (*68*), to project each 2 cell embryo gene expression profile onto a reference dataset of single cells from 2 cell embryo states previously reported by Deng *et al.* (*69*) (Figure 5C.) Most of the 2 cell embryos belonging to cluster C1 projected to the late 2 cell stage, whereas embryos from C2 exclusively projected to cells from the mid 2 cell stage. This shows that the two clusters identified through unsupervised clustering correspond to 2-cell embryos in the mid and late 2-cell stage respectively.

**Figure 5.**
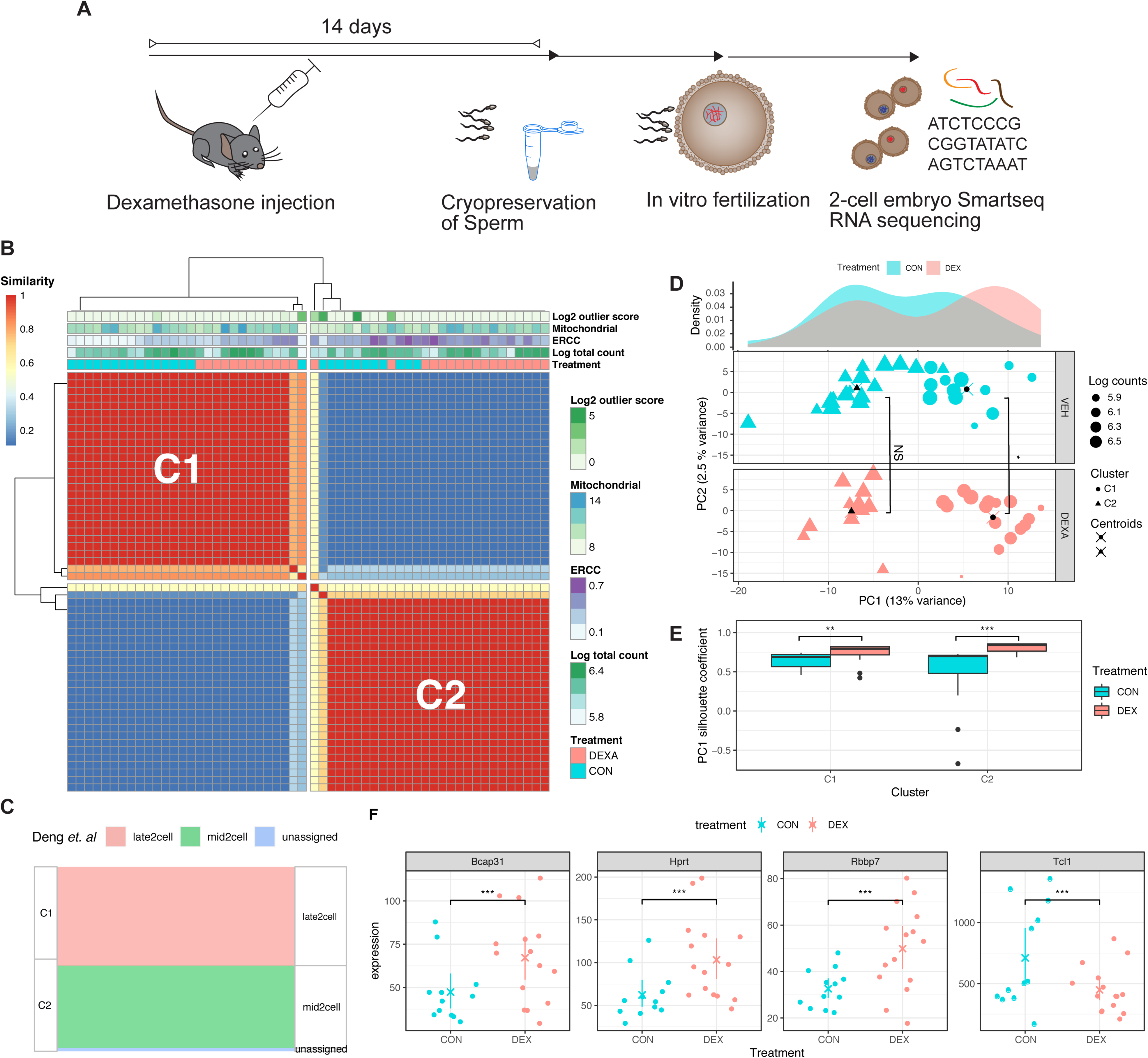
Effect of paternal Dex injection on embryonic offspring long RNA transcriptome (A) Experimental design depicting timeline between injection, sperm harvest, in vitro fertilization and Smartseq2 sequencing at 2-cell stage. (B) Consensus matrix representing the similarity between cells as reported by SC3. Similarity 0 indicates that a given pair of embryos were never assigned to the same cluster, whereas similarity 1 means that a pair of embryos were always assigned to the same cluster. (C) Sankey diagram showing projection of the obtained clusters (C1 and C2) into clusters reported by Deng *et al.* for single cells obtained from 2 cell embryos. (D) Principal component analysis of two-cell embryos gene expression. The top panel indicates the density of 2 cell embryos along PC1 grouped by condition; control (red) and Dex treatment (blue). The other two bottom panels show the distribution of 2 cell embryos across PC1 and PC2 for control (red) and treated (blue) groups. The cluster membership of each embryo is denoted by the point shapes (C1 cycles; C2 triangles) and centroids of each cluster is indicated with a black dot overlaid with an x. Wilcox tests were performed to assess differences on PC1 values of C1 and C2 clusters between the treated and control groups. NS denotes non-significant change for C2 cluster, while * indicates a significant difference for C1 cluster (p-value< 0.05). (E) Silhouette coefficient comparison between treatment and control, statistical significance was assessed with Wilcox test (** p- value < 0.01; *** p-value < 0.005) (F) Selection of differentially expressed genes as determined by Monocle within C1 corresponding to late 2-cell embryo stage (*** adjusted p- value < 0.005).

Principal component analysis (PCA) revealed a prominent separation between C1 and C2 along the PC1 axis, suggesting a correlation between PC1 and developmental transitions between mid and late single cell embryos (Supplementary Fig. 7B.) Interestingly, 2 cell embryo offspring of males injected with Dex exhibited a significant shift of the C1 cluster across PC1 (two-sided Wilcox test p< 0.03), while the C2 clusters did not show significant differences across PC1 between treatment and control groups (Figure 5D.) These results suggest that the effect of paternal Dex treatment on the transcriptome only becomes apparent at the late 2 cell embryo stage. To further explore this hypothesis, we calculated the silhouette coefficient (*70*) on PC1, as a measure of distance between C1 and C2 clusters, for the control and treatment group. We observed a significant increase of PC1 silhouette coefficient between treatment and controls for both C1 (one-sided Wilcoxon test p-value <0.005) and C2 (one-sided Wilcoxon test p-value < 2×10^-5^.) This confirms that Dex treatment affects embryonic gene expression, promoting altered late 2 cell embryo stages since the divergence from mid 2 cell embryos is significantly bigger in Dex offspring compared to control offspring (Figure 5E.)

Accordingly, differential gene expression analysis using Monocle2 (*71*) focused on late 2-cell embryos (cluster C1), revealed significant gene expression changes between offspring of males injected with Dex and controls across 38 genes, some of which were already apparent to a less significant extent during mid-2 cell embryos (cluster C1; e.g. Tcl1; Supplementary Fig. 8A,B.) In line with a potentially altered developmental trajectory becoming apparent in cluster 1, the late 2-cell stage includes several affected genes that are involved in early embryonic development. For example, Bcap31 (B-cell receptor-associated protein 31) is an important element for endoplasmatic reticulum and Golgi apparatus function, and Bcap31 mutations lead to developmental diseases with metabolic disturbances (*72*). This is reminiscent of the metabolic phenotype observed in the adult offspring of Dex injected fathers. Hprt (Hypoxanthine-guanine phosphoribosyltransferase) is crucial for cell cycle division, and Tcl1 (T-cell leukemia/lymphoma) regulates cell proliferation (*73, 74*). Hence, an upregulation of Hprt, and a concomitant down-regulation of Tcl1 might indicate that cell fate decisions later during development may be affected. Another differentially expressed gene is Rbbp7 (RB binding protein 7), which is part of many histone deacetylase complexes such as Nurd and PRC2/EED-EZH2, and thus plays an essential role in chromatin mediated gene regulation (*75*). Interestingly, several forms of PRC mutations in humans lead to different kinds of overgrowth phenotypes (*76*), an abnormality reminiscent of the increased BMI observed in Dex-offspring (Figure 3B.)

### Effects of Dex administration on a novel candidate for sperm RNA mediated inheritance

Despite the observed changes in sperm tsRNAs following acute Dex injection, we did not find an obvious causal connection to the altered 2-cell embryonic transcripts. This prompted us to investigate whether other germline changes might be more crucial for the offspring *in vivo* alterations in our model. We previously showed that chronic stress exposure also led to changes in sperm long RNAs that contributed functionally to the transmission of effects to the offspring (*79*), yet the fact that sperm RNA is stable through transmission and that the minute amounts of transmitted paternal RNA can elicit major changes in the embryo remains puzzling. Therefore, we evaluated the impact of Dex injection on the highly stable class of circRNAs in male sperm. CircRNAs were previously detected in swine(*80*) and human sperm (*81*) and suggested to have functional implications in epigenetic regulation. They have been attributed a critical role post transcriptional cessation in the male germline(*82*). Using Circexplorer in combination with EdgeR, analysis of sperm long RNA sequencing of males treated with Dex and controls revealed significant upregulation of two circRNAs (Figure 6A, q<0.1, Supplementary Fig. 1B.) while we also observed several significant changes in the sperm long RNA protein coding transcripts following acute Dex treatment (Figure 6B,C, Supplementary Fig. 1B, Supplementary Fig. 2B, Supplementary table 5). Both circRNAs are hosted in genes relevant for immune function (Taspase 1: Tasp1 and DENN Domain Containing 1B Dennd1b), yet the host genes did not show differential abundance of the protein coding host- gene transcript (Supplementary table 5). CircAtlas(*83*) revealed several potential miRNA sponge-targets to be captured by the altered circRNAs. Some of these miRNAs are common sponge-targets of both circRNAs such as mir3110-5p, mir706, mir1955 (Supplementary Fig. 9). Diamine acetyltransferase 1 (Sat1), one of 3110-5p’s high confidence miRNA-targets, as predicted by TargetScan(*84*), is indeed significantly upregulated in the embryos composing cluster 1 (later developmental stage). MiRNA-target- upregulation is expected if mir3110-5p was downregulated through circRNA-mediated- sponging and highlights a potential effective contribution of increased circRNA in sperm to embryonic pathway regulation. This is the first report of a change induced by environmental exposure in this compelling class of RNA in sperm.

**Figure 6.**
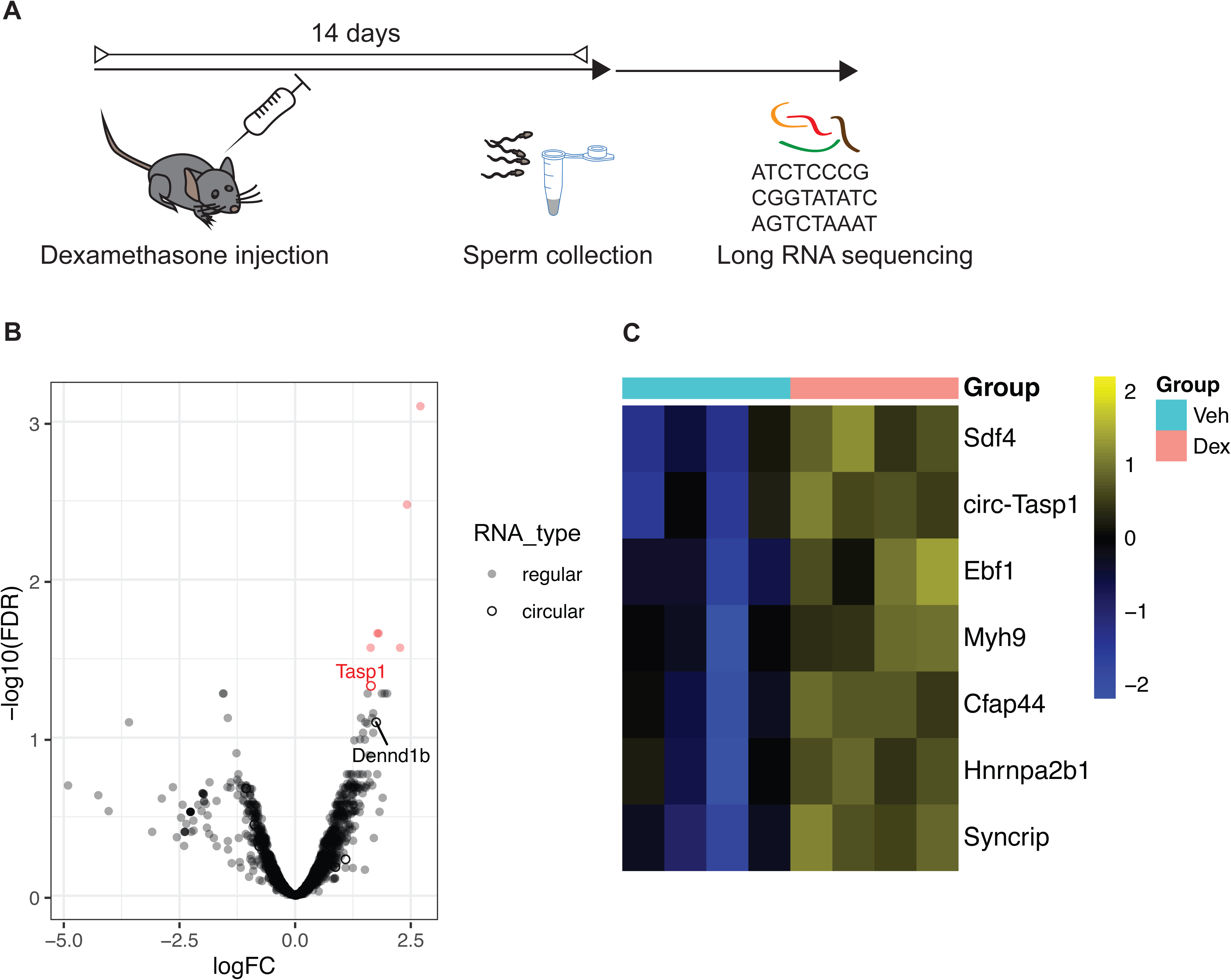
Effects of Dex on long RNA payload of sperm cells residing in testis at time of administration. (A) Experimental design depicting a time window of two weeks between injection of Dex and sperm collection for molecular analysis. (B) Volcano plot depicting fold changes and significance level of long RNA in mature sperm 14 days post injection of Dex (n=4) versus vehicle (n=4) as assessed by small RNA sequencing. (C) Heatmap showing significantly differentially expressed long RNA transcripts of the same experiment (multiple comparison corrected, q<0.05).

## Discussion

By generating offspring using assisted reproductive techniques (IVF), we circumvent potential confounding variables such as transmission via RNA contained in seminal exosomes (*85*) and affected maternal care by altered mating behavior (*86*) and prove germline dependence (*30*). Consistent with the significant changes of miRNAs and tsRNAs in the germline 2 weeks post GR activation, previous studies including our own have observed regulation of mouse sperm small RNA in a variety of contexts (7, 23, 24, 27, 35, 38, 42, 87–89).

Especially relevant specifically for our analysis, sperm RNA sequencing after drinking water administration of corticosterone for 4 weeks followed by mating, led to strong downregulation of tsRNA-GluCTC and tsRNA CysGCA, two of our top down-regulated tsRNAs, indicating that these tsRNAs are responding similarly to acute and chronic insults. At the same time this chronic manipulation elicits changes of several miRNAs, e.g. 34c and 471 (*41*), albeit in the opposite direction of what we find in response to acute Dex treatment. These discrepancies may arise either by the Dex induced short-term suppression of internal corticosteroid (*90*) or due to adaptations in response to chronic administration.

While four (23, 41, 91, 92) out of five (*93*) previous studies did report phenotypic effects following chronic paternal Dex exposure, only two assessed sperm small RNAs to associate the alterations to the sperm RNA payload (*91, 93*) yielding conflicting outcomes.

These differences might be due to inconsistent life stages (adulthood versus gestational), sperm collection (swim up, somatic lysis, or no purification) and/or dosage of exposure. Depending on the dosage and timing, the complex autoregulation of the GR can lead to GR downregulation after prolonged activation (*94*). Acute exposures have the advantage of avoiding such long-term feedback regulation, and hence provide an elegant approach for studying the signaling pathways leading to germline changes.

Mature sperm tsRNAs and miRNAs have been shown to be acquired during epididymal transit (*24, 40*) and miRNAs are necessary for early embryonic development under certain circumstances (*57, 63*). Furthermore, a recent publication suggests that chronic stress induced sperm miRNAs are taken up primarily from epididymisomes originating from the caput epididymis or the proximal epididymal tract (*43*). Chronic nutritional manipulation with effects on offspring also report the necessity of epididymal transit to acquire tsRNA changes in sperm(*24*). Harvesting mature sperm 3 hours after exposure yields a population enriched for cells that had been exposed while already in the cauda epididymis, where spermatids reside for roughly 5 days (*59*). These cells have not traveled through the epididymis nor have they had a chance to potentially take up small RNA from caput-derived epididymisomes after Dex administration. As expected, we detect no changes in miRNAs in these samples. We do however detect changes in tsRNAs 3 hours post Dex, some of which even persist 14 days post injection. These results for the first time show rapid acquisition of changes in vivo and corroborate previous in vitro findings that show that incubation with epididymisomes can alter sperm RNA payload(*24*). Our acute intervention for the first time assesses effects on germline payload already after a short interval. Whereas chronic interventions - based on their experimental design - do not assess changes in mature sperm soon after the first intervention. Studies aiming at the elucidation of the origin of sperm RNA changes might benefit from acute interventions to circumvent confounders such as dynamic exosomal RNA supply as a result of cumulative interventive strain on animals.

An additional option for sperm RNA alterations in transcriptionally inert sperms was suggested in a recent study that found mitochondrial tRNA cleavage in the T-loop in response to a one-week high sugar diet in humans(*95*). In line with this observation, Dex injection could trigger oxidative stress(*96*) that provokes such cleavage to increase tsRNA levels(*97*). A role for oxidative stress in sperm RNA dynamics is further supported by a recent study in boar sperm that found seasonal differences in sperm small and long RNA associated with changed abundance of transcripts mapping to oxidative stress-, DNA damage- and autophagy- related genes (*98*). Such potentially oxidative stress mediated mechanism does not explain though a rapid decrease of tsRNAs 3 hours post Dex injection. Importantly, we show alterations in sperm tsRNAs that persist in the oocyte concomitant with changes in early embryonic gene expression and a metabolic phenotype in adulthood. tsRNAs and tRNA-Gly derived fragments in particular are known to induce chromatin structure mediated gene regulation and to regulate cell differentiation in various contexts (*99, 100*). Hence, we propose that the transmitted reduction in key tsRNAs such as Gly-GCC and Gly-CCC explains in part the observed perturbations during late two cell embryo developmental stage. This might reflect an accelerated developmental transcriptional program in the preimplantation embryo of Dex injected males, ultimately resulting in aberrant BMI and glucose metabolism later in life.

Additionally, we have discovered alterations in circRNA abundance in mature sperm that might also impact the developmental program in the early embryo. CircRNAs have the potential to be translated into proteins via back-splicing(*101*). Accordingly, they are crucial contributors to spermiogenesis post transcriptional cessation, since they provide a stable alternative to linear mRNA templates for protein translation(*82*). Transmitted sperm-circRNAs could likewise contribute to translation post fertilization, yet the unconventional lattice-state of ribosomes preventing normal rates of translation post transcription(*102*) accompanied by a rapid increase in proteins of the ubiquitine/proteasome pathway(*103*) make this unlikely. Never the less a study on human sperm detected abundant levels of circRNA with predicted regulatory function of early developmental genes in sperm heads, suggesting transmission and function post fertilization(*81*). By sponging miRNAs that regulate early embryonic transcripts, circRNA could amplify minute signals from paternal environment, such as might be the case for the gene transcript Sat1, that displays increased expression in Dex offspring in 2-cell embryos from cluster 1.

Besides altered RNA identity, nucleic acid modifications especially of RNA but also DNA methylation and chromatin accessibility might contribute further to the effects of Dex injections on offspring metabolism. While detection of changes in each player should be subject of further investigation and might reveal a glimpse of their potential implication, proof of the individual relative causal contribution is extremely challenging since they likely require tight interaction to unravel their orchestrated effects.

Finally, it might be useful to consider testing the translatability of our findings to humans. Here we investigated the effects of a single Dex administration soon after the injection in mice, mimicking a single GR activation such as elicited by treatment of an acute asthma exacerbation (*104*). The recent report that Dex can reduce the number of deaths associated with the Covid-19 pandemic (*51*), further prompts the re-evaluation of the impact of prolonged Dex treatment on offspring phenotype. From a clinical perspective, additional consideration is warranted for consequences on offspring health when extended time has elapsed between treatment and time of conception. Such designs may pave the way for the extrapolation of our findings.

We conclude that acute Dex treatment can induce germline epimodifications in the form of small and long non-coding RNA, which likely are relevant in the transmission of the effects of single traumatic events on offspring well-being. Our data suggest that sperm small RNAs are not solely regulated through epididysomal uptake during transition from caput to cauda epididymis. This expands the interpretation from chronic dietary and stress exposures(*24, 43*), where uptake of tsRNAs and miRNAs via epididymisomes has been suggested to lead to differential sperm payload, yet required sperm to transit from caput to cauda to bring about the changes. A persistent detection of significant fold changes of the exact same sperm small RNA in the embryo suggests functional implication in the information transfer from father to offspring. Together with potentially transmitted miRNA sponges in the form of circRNAs, this likely contributes to a slight developmental acceleration of gene expression programs in the early embryo and ultimately manifests in a metabolic phenotype. Future studies may aim at testing the causal contribution of specific sperm RNAs to the transmission of effects of acute impacts. Certainly, continuous methodological refinement will help dissect the relative implication and the interplay of the distinct germline modifications such as DNA- methylation, histone-PTMs and chromatin architecture in this highly complex process.

## Conflict of interests

The authors declare no competing interests. E.A.M. is a founder and Director of STORM Therapeutics Ltd. STORM Therapeutics had no role in the design of the study and collection, analysis, and interpretation of data as well as in writing the manuscript.

## Materials and Methods

### Animals

C57Bl/6 mice were obtained from the Sanger Research support facility in-house-breeding colony. They were housed in a temperature and humidity-controlled facility in individually ventilated cages under a non-reversed light-dark cycle (Sanger Research support facility) or a reversed-light-dark cycle (ETH EPIC). Standard chow (LabDiet(r) 5021-3 supplied by IPS) and water were provided *ad libitum after weaning* unless stated otherwise (e.g. oocyte donors). Breeding colony was provided SAFE R03-10 breeding diet, supplied by SAFE diets. Experimental procedures were performed during the animals’ inactive cycle at Sanger. Age and weight matched (margin of one week) males were used in each experimental group receiving Dex injections. Animals used for Dex injection followed by sperm sequencing were all sexually mature (14 and 7 days or 3 hours post treatment were 13, 11 and 9 weeks of age respectively) at the time of sperm collection.

C57Bl/6 males used for sperm sequencing 14 days post Dex injection and q-PCR experiments/validation 3 hours post Dex were obtained from the ETH’s EPIC in house breeding colony in Zürich and were 14-18 weeks old. These mice were fed chow #3734 by Kliba/Granovit.

IVF oocyte donor females and embryo recipients were fed SAFE R03-10 breeding diet, supplied by SAFE diets until 10 days post embryo transfer. Until this time embryo recipients were housed in pairs after which they were split into single housing. IVF offspring was weaned at PND21 and assigned to cages avoiding littermate cohousing. Offspring phenotyping was carried out between 3.5 to 4 months and necropsy at 4.5 months of age in balanced (offspring controls, offspring treatment) and age matched groups (all animals had an age spread of 3 days). Animals were housed in groups of 4-5 mice/cage in the Sager Institute barrier research support facility (all animals apart from animals for q-RT-PCR experiments) and ETHZ’s EPIC facility (animals for q-RT-PCR).

All experiments were approved by the UK home office (project license P176396F2) and Cantonal commission for animal experimentation Zürich (project license ZH222/19).

### Dex treatment and sample collection

Age matched males with an age spread of 1 week were randomly assigned to control and treatment groups. Males were injected with either 2mg/kg of Dex in 10 % DMSO, 0.9% saline or vehicle (10%DMSO in 0.9% saline). Males used for sperm collection did not undergo any metabolic testing. They were sacrificed 2 weeks, 7 days and 3 hours after Dex or vehicle treatment. Cauda epididymis and vas deference were dissected and placed in M2 medium. After allowing sperm to diffuse into M2 medium, cells were pelleted by short centrifugation and washed with PBS. For sperm RNA sequencing and q-PCR, mature sperm cells were separated from potential somatic contamination by somatic lysis, followed by 2 washes with PBS (*105*). Sperm counts and fertilization rate appeared unaffected post Dex injection (Supplementary Fig. 10&11).

### Superovulation, in vitro fertilization and embryo culture

12 randomly selected, C57BL/6 females were superovulated at 26-31 days of age with Card Hyperova (Cosmo Bio, KYD-010-EX-X5), followed by 7.5 IU human chorionic gonadotrophin (HCG) 48 hours later.

Cumulus-oocyte complexes (COCs) were released from the ampulla of the oviduct 16-17 hours after HCG administration, and preincubated in high calcium HTF with Glutathione medium for 30-60 minutes (in CO2 incubator at 37 deg C, 5% CO2 in air) before insemination. Frozen sperm used for insemination was pooled from 2 males that had been injected with Dex or vehicle 14 days prior to cryopreservation. Thawed sperm was preincubated for 30 minutes in TYH (with Methyl-b-cyclodextrin, Sigma C4555) medium at 37 deg C, 5% CO2 in air, before being added to the COC complexes for fertilisation. 4 hours after insemination, the presumptive zygotes were washed through several drops of KSOM (Millipore, MR-121-D) and incubated overnight in KSOM.

For in vivo offspring, 14-20 x 2 cell embryos from overnight culture in 6 individual IVF dishes /group were implanted into 0.5 dpc pseudo-pregnant F1 females (6 females/group). Each dish contained oocytes from one female with the exception of 2 dishes (out of 6) in the Dex group that contained oocytes of the same female, since one female failed to super-ovulate. For molecular (single) embryo gene expression analysis at the two cell stage, 2-cell-embryos from overnight culture were frozen, and after thawing briefly cultured in preincubated KSOM until/during plating into 96 well plates. The females used to generate these embryos were superovulated with PMSG. The IVF Protocol is based on EMMA Harwell’s protocol (adapted from Takeo & Nakagata 2011(*106*)), and the Sperm Freeze Protocol is based on Ostermeier G.C. et al (2008)(*107*). Resulting litter sizes did not differ between Vehicle and Dex injected offspring (Supplementary Fig. 12)

### Sperm and embryo RNA extraction

Total RNA was prepared from adult mouse sperm using Trizol (Thermo Scientific 15596026) and Directzol (Zymo R2080). Total RNA was prepared from zygotes using the Trizol LS protocol. Quantity and purity of RNA were determined by Agilent 2100 Bioanalyser (Agilent Technologies) and Qubit fluorometer (Life Technologies). Absence of prominent ribosomal peaks indicated absence of somatic cell contamination.

### Sperm RNA sequencing (RNAseq)

Sequencing was done using an Illumina Genome Analyzer HiSeq 2500 (Illumina) in Rapid run mode for long 100bp and small 50 bp RNA sequencing runs respectively.

Libraries for long RNA sequencing were prepared using the TruSeq Stranded Total RNA kit according to the manufacturer’s instructions with indices diluted at 1:3. 200 ng of total sperm RNA was subjected to removal of rRNA using Ribozero gold kit. Approximately 100ng of sperm RNA and total RNA of several 2-cell zygotes was subjected to TruSeq or Nextflex (sperm 14 days post injection) small RNA library preparation following the manufacturer’s recommendations with the following modifications: adaptors were diluted 1:4 and PCR cycles were augmented to 18 and 22 (Nextflex) PCR cycles respectively. When library preparation of samples was split across days groups were balanced to circumvent batch effects.

### Single embryo seq

2 cell embryos were generated using the same conditions as indicated for in vivo offspring yet followed by embryo cryopreservation until processing for library preparation. They were thawed and those that appeared intact (34 controls and 37 Dex) pipetted into wells of 2 96 well culture plates containing lysis buffer and stored at -80°C before processing according to the Smartseq 2 protocol and manufacturer’s recommendations (Nextera). Libraries contained a 1:19 Million dilution of External RNA Controls Consortium (ERCC) spike-ins (4456740 Ambion) and were amplified for 18 PCR cycles. Sequencing was performed on a HiSeq V4 under paired end 75bp mode.

### Insulin and Glucose tolerance test

Animals were fasted 4 hours to establish a shared baseline glucose level. They received a single injection of insulin (insulin: 1mU/g body weight) (Actrapid Novo Nordisk), glucose (2mg/g body weight) or vehicle (saline) intraperitoneally. Blood samples were taken from lateral tail vein in adult animals to assess blood glucose level using an Accuckeck aviva device.

### Body mass index

Animal lengths were measured using a standard ruler and weighed for assessing body weight. Body mass index was calculated using the following formula: weight (g)/(length (cm)^2).

### Necropsy

Organs were dissected after sacrifice and weighed immediately on a scale using “g” as a unit with an accuracy of 2 decimals (accurate down to 10 mg).

### q-RT-PCR

5ng/sample RNA isolated from sperm was reverse transcribed (RT) using the miCURY LNA RT kit (Qiagen #339340). Quantitative RT-PCR (qRT-PCR) was performed using SYBR green based detection in a Biorad thermal cycler with MiRCURY LNA-based small RNA probes designed against 5’end of tRNA ArgCCT-2 (5‘GCCCCAGUGGCCUAAUGGAUAAGGCACUGGCC3’) with a polyA tail directed reverse miRCURY primer (Qiagen # 339317). U6 was used as an internal control (Qiagen # 339306).

### Bioinformatic and statistical data analysis

#### Sperm RNA sequencing

Each sequencing library represented sperm harvested from a single male. Sequencing quality was assessed with FastQC (*108*) and MultiQC (*109*). Adapters were removed from the 3’ ends with cutadap t(*110*) (version 1.14) and resulting sequences with 14 nucleotides of length or less were discarded. All other reads were aligned end to end (no soft clipping) to the ENSEMBL *Mus musculus* genome (release 75) (*111*) with STAR (*112*). No mismatches were allowed. Featurecounts was used to match the alignments against the miRbase(*113*) annotation (version 21) and obtain a matrix of miRNA counts. We applied fractional counts whenever alignment occurred at multiple genomic locations. Differential expression was analyzed using DESeq2 (*56*). Quantification of tRNA fragments was performed as above, but all CCA-3 trinucleotides were trimmed after adapter removal, sequences with 15 nucleotides or less were subsequently discarded and GtRNAdb (*114*) annotation (GRCm38/mm10) was used to obtain the count matrix.

For the data set collected 14 days after Dex injection, library preparation included the insertion of 2 random tetranucleotides between read and adapters. By including only unique sequences in the analysis we removed duplicates due to PCR amplification.

Long RNAseq libraries were pre-processed with trimmomatic (*115*) to remove adapters. Reads were aligned to the genome using STAR (*112*) and quantified using featurecounts (*116*) Circular RNAs (*117*) based on junction reads as detected by STAR. Differential expression analysis was performed on the combined set of counts for circular and non-circular RNAs using edgeR (*118*). Robust estimation of dispersion was used to avoid spurious significance due to outliers.

### 2-cell single embryo sequencing analysis

Reads from 2 cell embryos were mapped to the mouse reference genome (mm10) and ERCC spike-ins using STAR (*112*). Resultant alignments were processed to quantify the expression of annotated genes by GENCODE (vM11) and ERCC spike-ins using featureCounts (*116*). To filter low-quality sequenced embryos we only considered those which had a total read count of at least 0.5 million reads with less than 15% and 10% their read counts mapping to mitochondrial genes and ERCC spike-ins respectively. After these filters were applied a total of 56 embryos (29 controls and 27 treated) remained. We clustered their gene expression profiles using SC3 (*67*) obtaining two main clusters (C1 and C2). Using scmap (*68*), we projected the gene expression profiles for the two cell embryos onto an index containing expression profiles from zygotic, early/mid/late 2 cell embryos and 4 cell embryo cells reported by Deng *et al.* (*69*). We performed PCA analyses using scater (*119*) (runPCA function) and we calculated the PC1 silhouette coefficient using in-house R scripts. To perform differential gene expression analyses we normalized the read counts of each embryo as FPKM and we used Census (*71*) algorithm to convert these values into relative transcripts counts. We computed the obtained ‘Census counts’ using Monocle (v 2.99.2), assuming a negative binomial distribution and a lower detection limit of 0.5. We performed differential gene expression analyses between the total treated and control embryos, and also between the treated and control embryos inside of C1 and C2 clusters.

### Remaining statistical analyses

Sample size for in vivo offspring phenotyping was estimated based on previous work on similar models (*44, 45*). 3-Way repeated measures ANOVA was used to assess statistical significance for BMI, GTT and ITT measurements. Necropsy data were analysed using 3- way ANOVA followed by multiple t-tests and corrected for multiple comparisons using the Benjamini-Hochberg method. Normality was assessed with the Kolmogorov Smirnov test and met in all necropsy data. Homogeneity of variances was assessed and met in all necropsy data unless gonadal WAT. All t-tests did not assume homogeneity of variances (applied Welchs correction). All statistics of behavioural, metabolic tests and q-RT-PCR were computed with Prism. Outliers were removed from q-PCR results using Prism^s inbuild ROUT method and are depicted in the supplementary table 3 containing raw data with a star. Q-RT-PCR results were anaylysed with a 2-Way Anova followed by posthoc tests to compare individual groups applying the Bonferroni correction for multiple comparisons. All reported replicates were biological replicates. Significance was set at *p* < 0.05 for all tests.

### Availability of Data

The datasets supporting the conclusions of this article are included within the article (supplementary tables). All sequencing data were deposited to Gene Omnibus (accession number: xxx) and ENA (accession number: <u>xxxx</u>). All code is available on Github (https://github.com/ETHZ-INS/Sperm-RNA-Dex)

## Supplementary Material

This article contains supplementary Figures and tables. Supplementary figures are compiled in one word document.

## Supplementary table legend

*Supplementary Table 1*

Sheet 1: List of normalized miRNA read counts of sperm harvested 14 days post Dex and vehicle injection.

Sheet 2: Deseq2 results of a comparison between miRNA from sperm harvested 14 days post Dex and vehicle injection.

Sheet 3: List of normalized tsRNA read counts of sperm harvested 14 days post Dex and vehicle injection.

Sheet 4: Deseq2 results of a comparison between tsRNAs from sperm harvested 14 days post Dex and vehicle injection.

*Supplementary table 2*

Sheet 1: Deseq2 results of the comparison of tsRNAs in sperm harvested 3 hours and 7 days post Dex with tsRNAs in sperm harvested 3 hours and 7 days post vehicle injection. Sheet 2: Deseq2 results of the comparison between tsRNAs from sperm harvested post Dex and vehicle injection at 3 hours with tsRNAs in sperm harvested 7 days post Dex and vehicle injection.

Sheet 3: Deseq2 results of the interaction between treatment and time for tsRNAs from sperm harvested 3 hours and 7 days post Dex and vehicle injection.

Sheet 4: Deseq2 results of the comparison of miRNAs in sperm harvested 3 hours and 7 days post Dex with miRNAs in sperm harvested 3 hours and 7 days post vehicle injection. Sheet 5: Deseq2 results of the comparison between miRNAs from sperm harvested post Dex and vehicle injection at 3 hours with miRNAs in sperm harvested 7 days post Dex and vehicle injection.

Sheet 6: Deseq2 results of the interaction between treatment and time for miRNAs from sperm harvested 3 hours and 7 days post Dex and vehicle injection.

Sheet 7: Raw values of qRT-PCR analysis for tsRNA-Arg-CCT-2 in caput and cauda sperm sampled 3 hours post Dex and vehicle injection.

*Supplementary table 3*

Raw data for BMI (sheet 1), GTT (sheet 2), ITT (sheet 3) and necropsy weights (sheet 4) of adult offspring animals resulting from IVF of wildtype oocytes and sperm harvested 14 days post Dex and vehicle injection.

*Supplementary table 4*

Sheet 1: Monocle output list on significantly differentially regulated genes between 2-cell embryos resulting from IVF of wildtype oocytes and sperm harvested 14 days post Dex and vehicle injection.

*Supplementary table 5*

Sheet 1: List of normalized long RNA seq counts of sperm harvested 14 days post Dex and vehicle injection.

Sheet 2: EdgeR results of a comparison between long RNA reads from sperm harvested 14 days post Dex and vehicle injection.

## Acknowledgements

We thank Wayo Matsushima and Pierre-Luc Germain for valuable advice on bulk sequencing analysis and the Sanger RSF for help with animal care, especially the teams of Michael Woods and Francesca Flack. We also thank Sergio Mompart Barrenechea and Jasmine Kaur for help with the q-RT-PCR.

KG was funded by the Swiss National Science Foundation early postdoc and advanced postdoc mobility a SPARK and Novartis foundation grant. Some of this work was supported by Cancer Research UK (C13474/A18583, C6946/A14492) and Wellcome (104640/Z/14/Z, 092096/Z/10/Z) to EAM. GP and MH were supported by a core grant from the Wellcome Trust. The lab of JB is currently funded by the ETH Zurich, SNSF Project Grant 310030_172889/1, ETH Research Grant ETH-20 19-1, Kurt und Senta Herrmann-Stiftung, and Botnar Research Center for Child Health. For the purpose of Open Access, the author has applied a CC BY public copyright license to any Author Accepted Manuscript version arising from this submission.

## Authoŕs contributions

KG performed animal exposures, collected samples and prepared sequencing libraries. EG performed IVF and embryo culture. GP analysed single cell sequencing data. MH supervised the single-embryo sequencing analysis. AC and FG analyzed bulk sequencing data. KG, JB and EAM designed the study, interpreted the results and wrote the manuscript with input from the other authors.

## Notes

### Competing Interest Statement

The authors have declared no competing interest.

### Summary of Updates

The revised version has clarified the relation of our findings in relation to previous findings. This is reflected in changes to the abstract but also to the discussion.

