## Supplementary Figures for "Single paternal Dexamethasone challenge programs offspring metabolism and reveals circRNAs as novel candidates in RNA-mediated inheritance"

**Supplementary Information**

**Supplementary Figure 1**

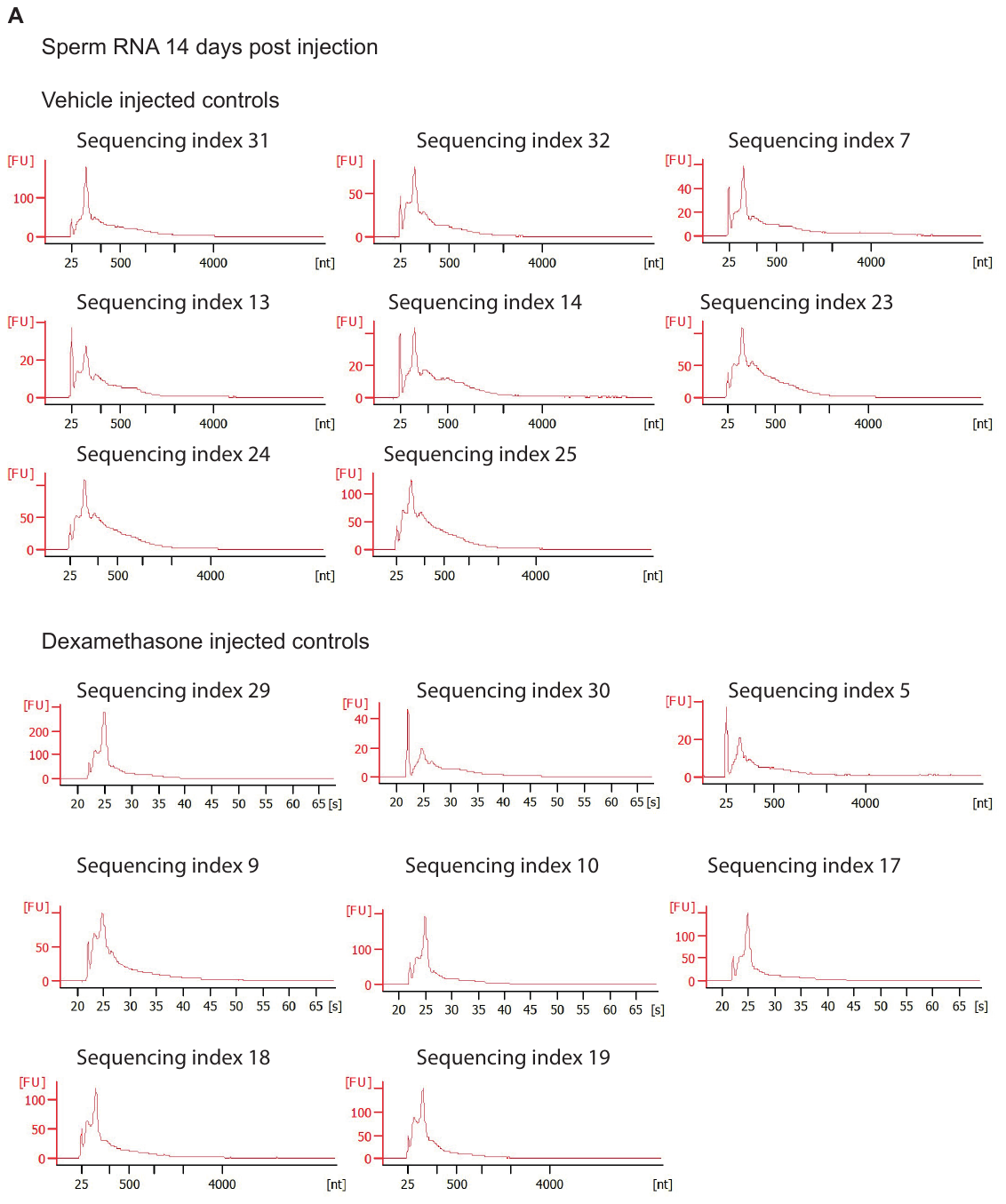

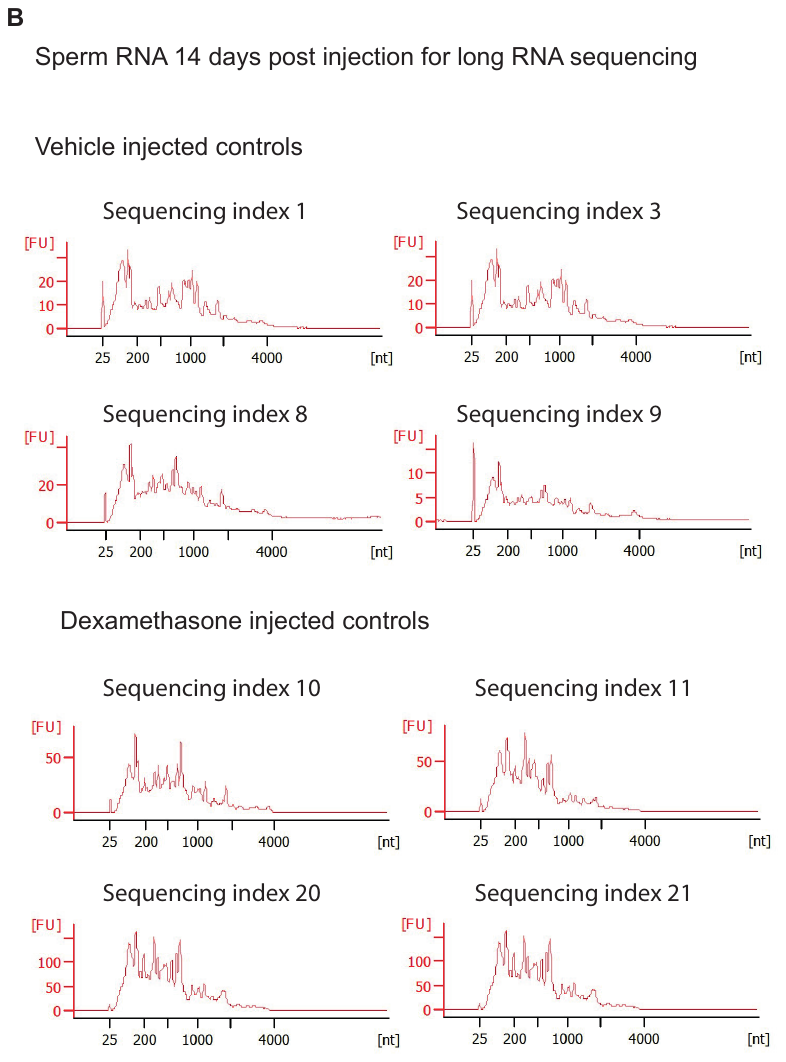

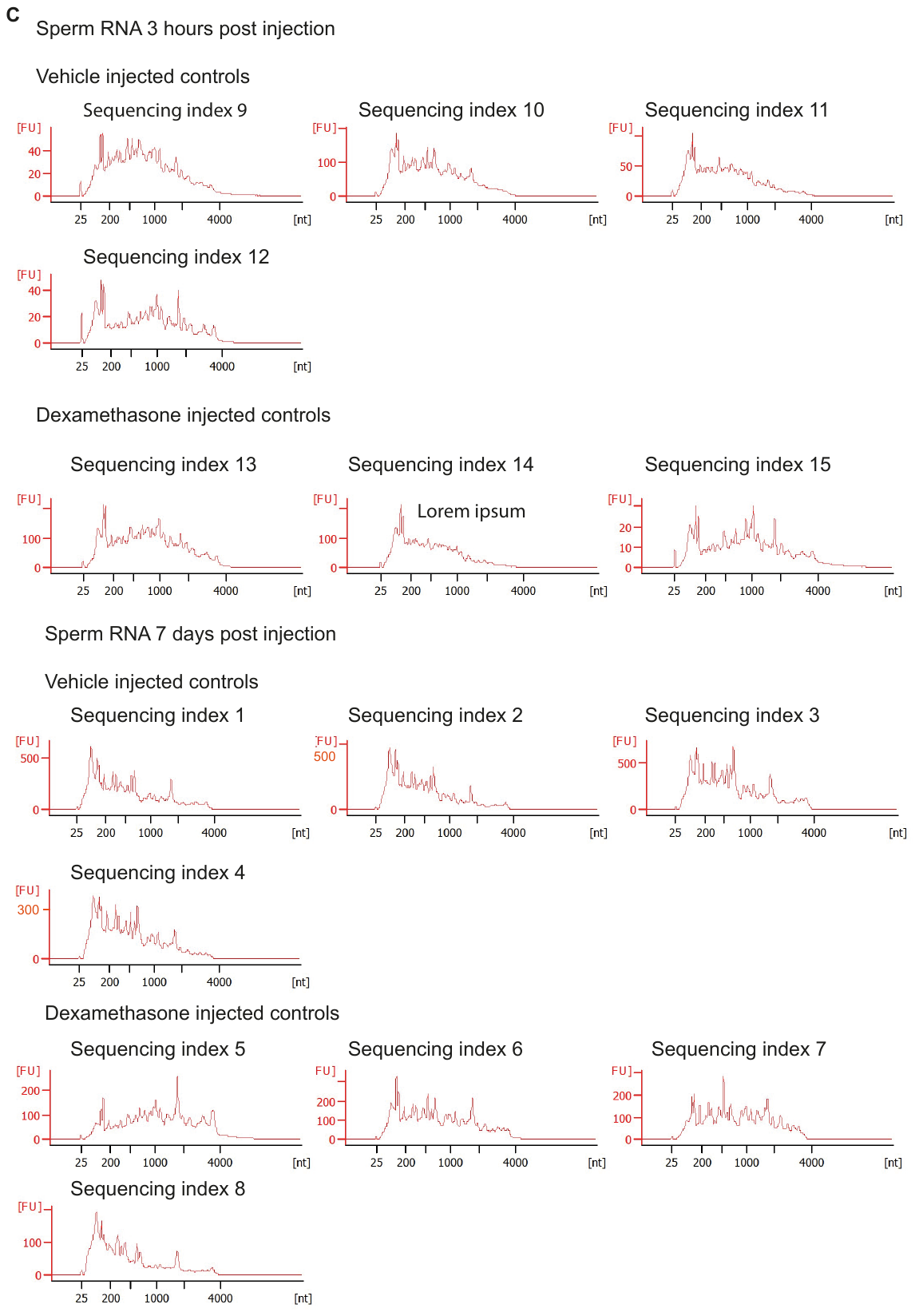

Size profile of sperm from males (A) 14 days post vehicle and dexamethasone injection used for small RNA sequencing (B) 14 days post vehicle and dexamethasone injection used for long RNA sequencing (C) 3 hours and 7 days post st vehicle and dexamethasone injection used for small RNA sequencing, as determined by Bioanalyzer pico kit. Absence of ribsosmal peak 28s (4.7kb) is indicative of a high degree of purity in sperm cells. Nt= nucleotides. S= seconds (22s equals approximately 25 nt, 40s equals approximately 1900nt).

**Supplementary Figure 2**

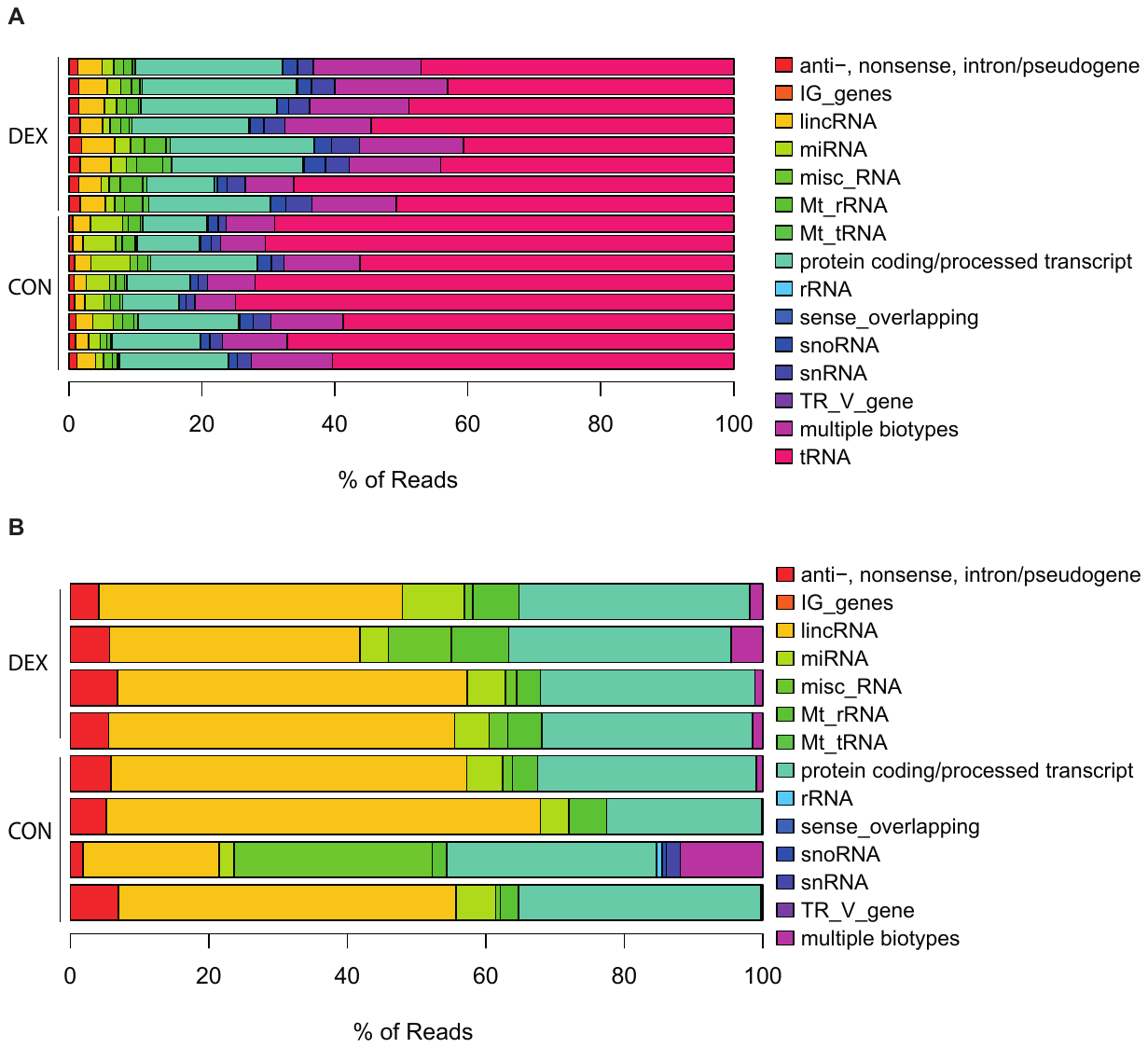

Relative percentage of RNA Ensemble biotypes and tRNA matching reads in (A) small RNA sequencing libraries of 8 sperm samples from control males and 8 sperm samples 14 days post dexamethasone injection. (B) long RNA sequencing libraries of 4 sperm samples from control males and 4 sperm samples 14 days post dexamethasone injection.

**Supplementary Figure 3**

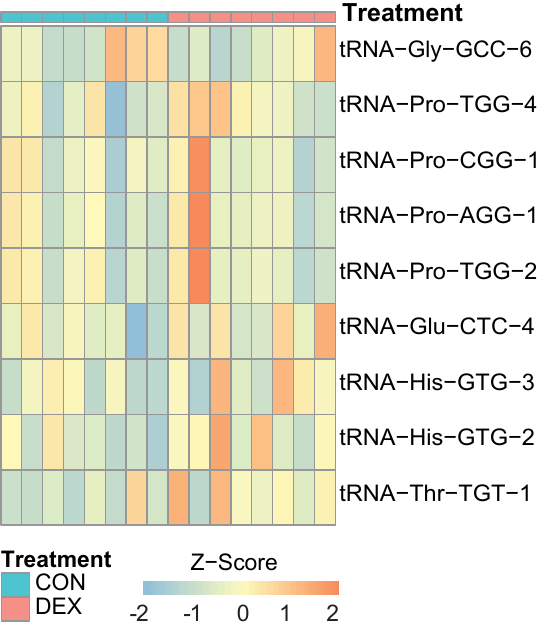

Effect of Dex on sperm cells at different time points post Dex administration. Heatmap showing significantly interacting tRFs (3 hours and 7 days post dex analysis) in the data obtained form sperm sequencing 14 days post injection (Dex=8 and controls=8). TRFs are grouped by sequence identity for display only.

**Supplementary Figure 4**

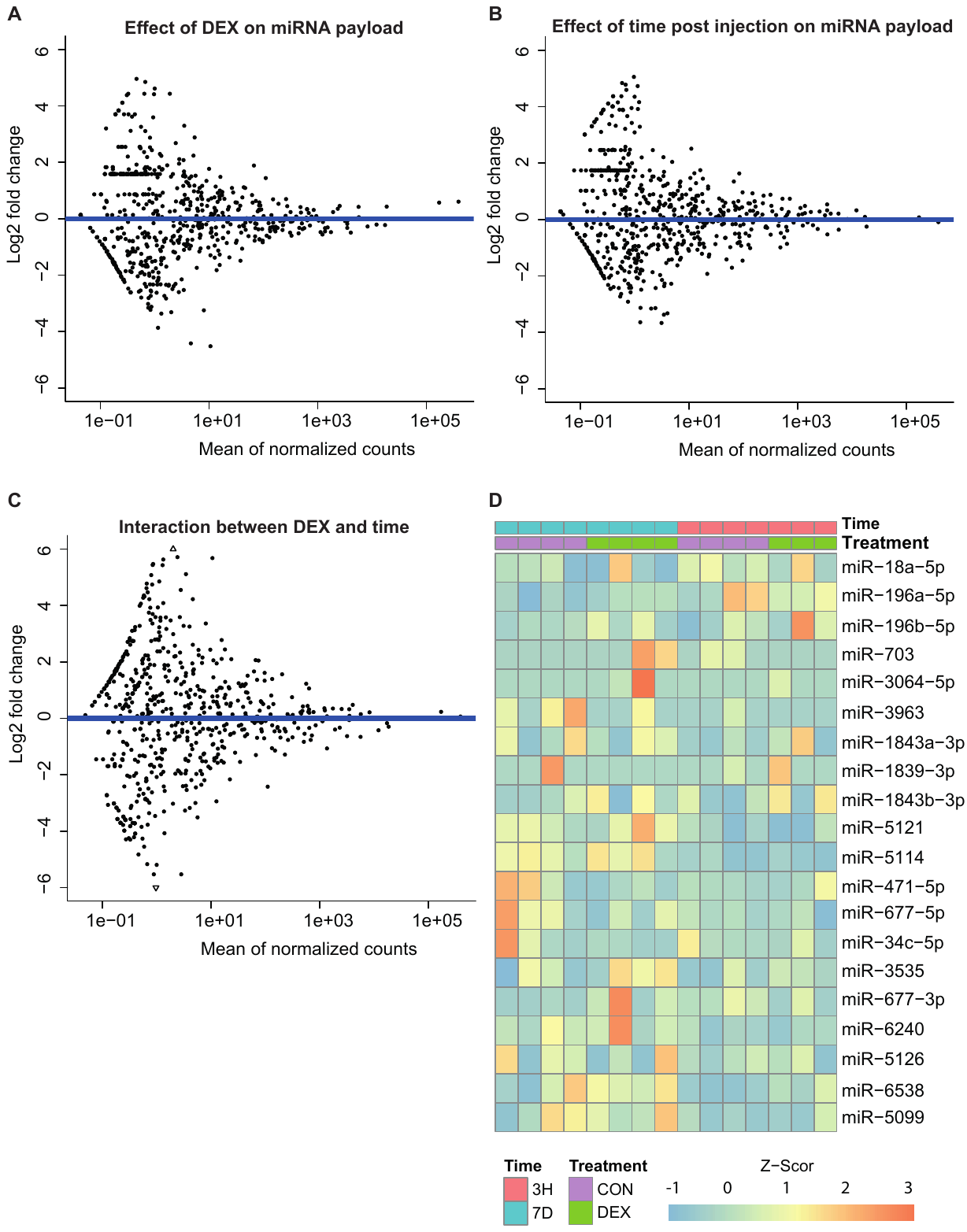

Mature sperm miRNA payload as determined by next generation sequencing . Deseq2 analysis of sperm samples collected 3 hours and 14 days post dexamethasone injection did not reveal any significant effects of (A) treatment (dexamethasone injection), (B) time (time elapsed since injection) nor an (C) interaction between the two. (D) Heat map depicting those miRNAs that are significantly affected in the data obtained from 14 days post injection, at 3 hours and 7 days post dexamethasone injection.

**Supplementary Figure 5**

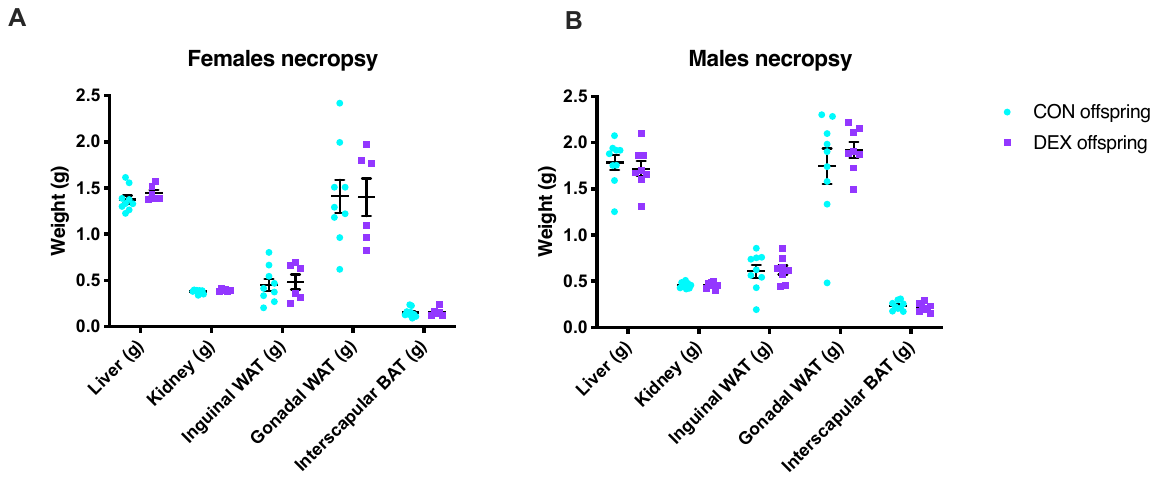

**C**
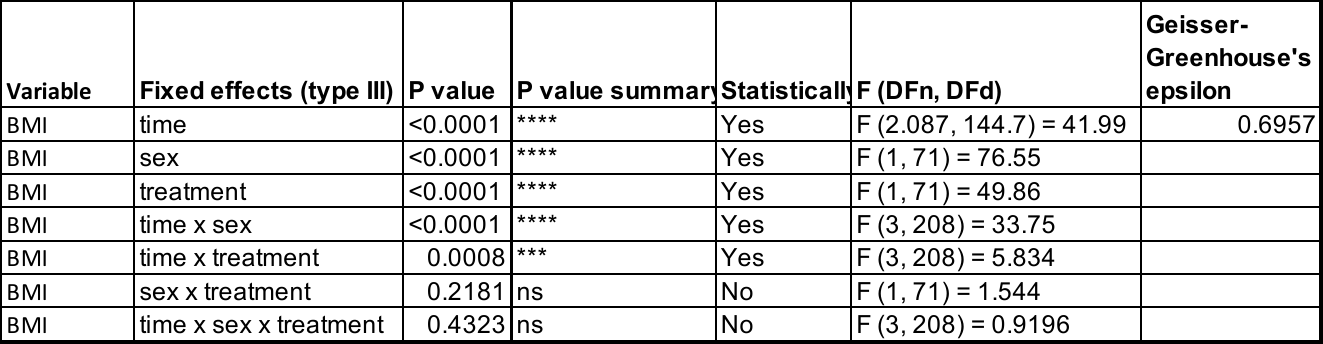

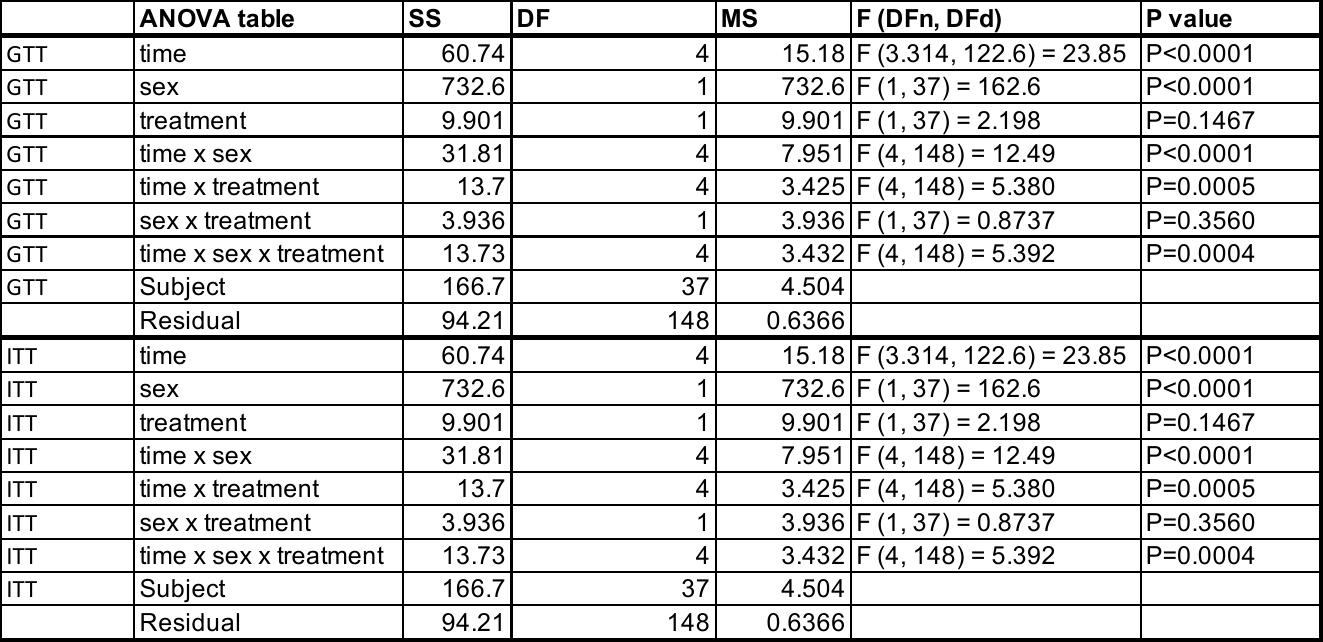

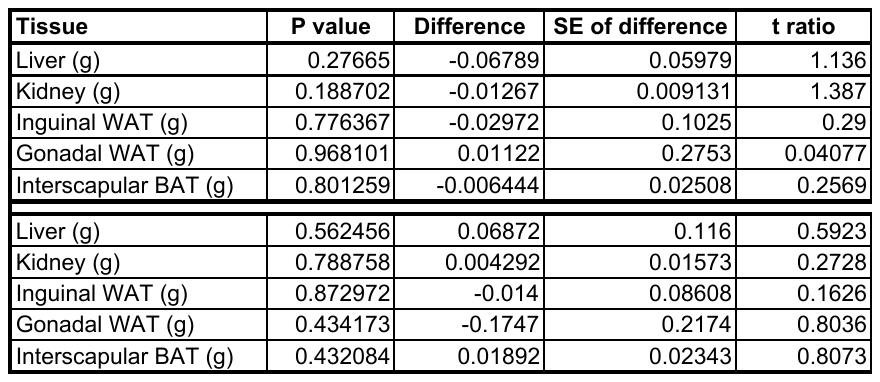

Necropsy followed by weighing of dissected tissue from offspring of control injected and dexamethasone injected males. Comparison of the weight of liver, kidney, inguinal and gonadal white adipose tissue (WAT) as well as interscapular brown adipose tissue (BAT) did not reveal significant differences between (A) female and (B) male offspring of control and dexamethasone injected males. (C) Statistical values obtained by overall ANOVAs (BMI, GTT, ITT) and Multiple t-tests corrected for multiple comparisons (necropsy). All Data besides male gonadal WAT showed equal distribution of variances. Graphs show scattered dot plots with standard error of the mean. ITT = insulin tolerance test, GTT= Glucose tolerance test, BMI= Body mass index, WAT = white adipose tissue. BAT= brown adipose tissue.

**Supplementary Figure 6**

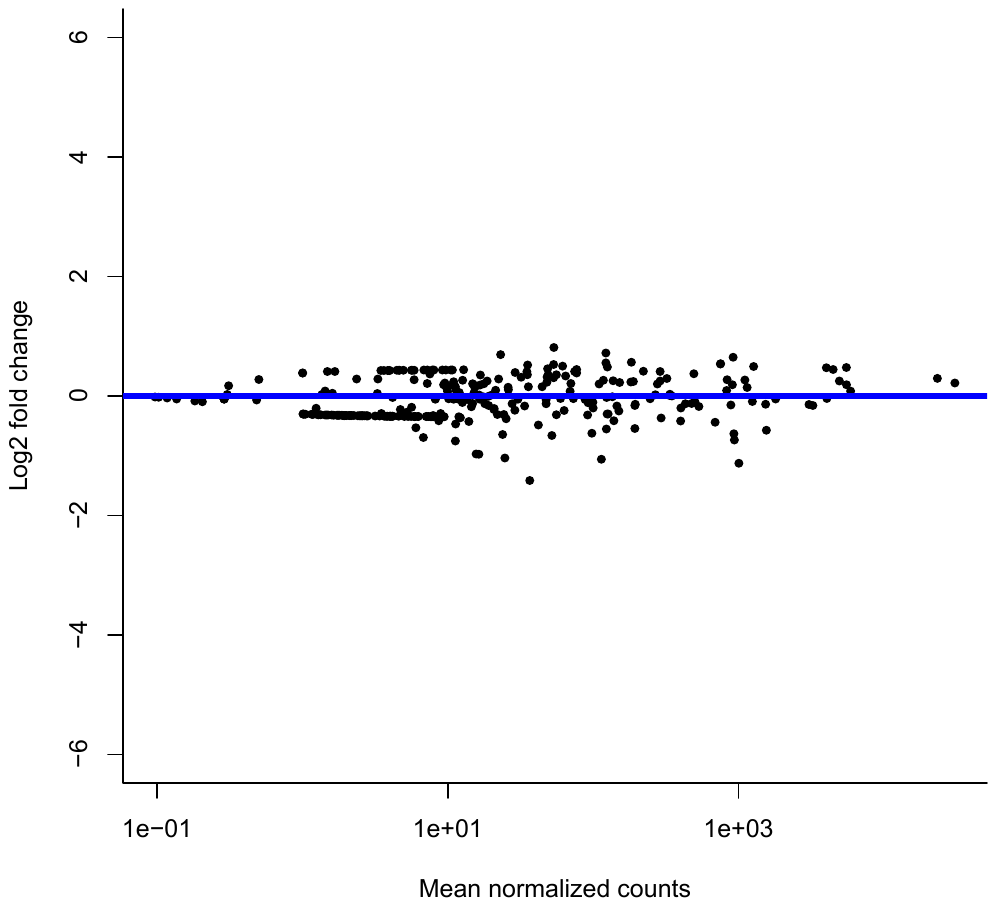

Small RNA sequencing of 2-cell embryos as assessed by bulk small RNA sequencing. MA plot showing absence of statistically significant fold-changes in miRNAs.

**Supplementary Figure 7**

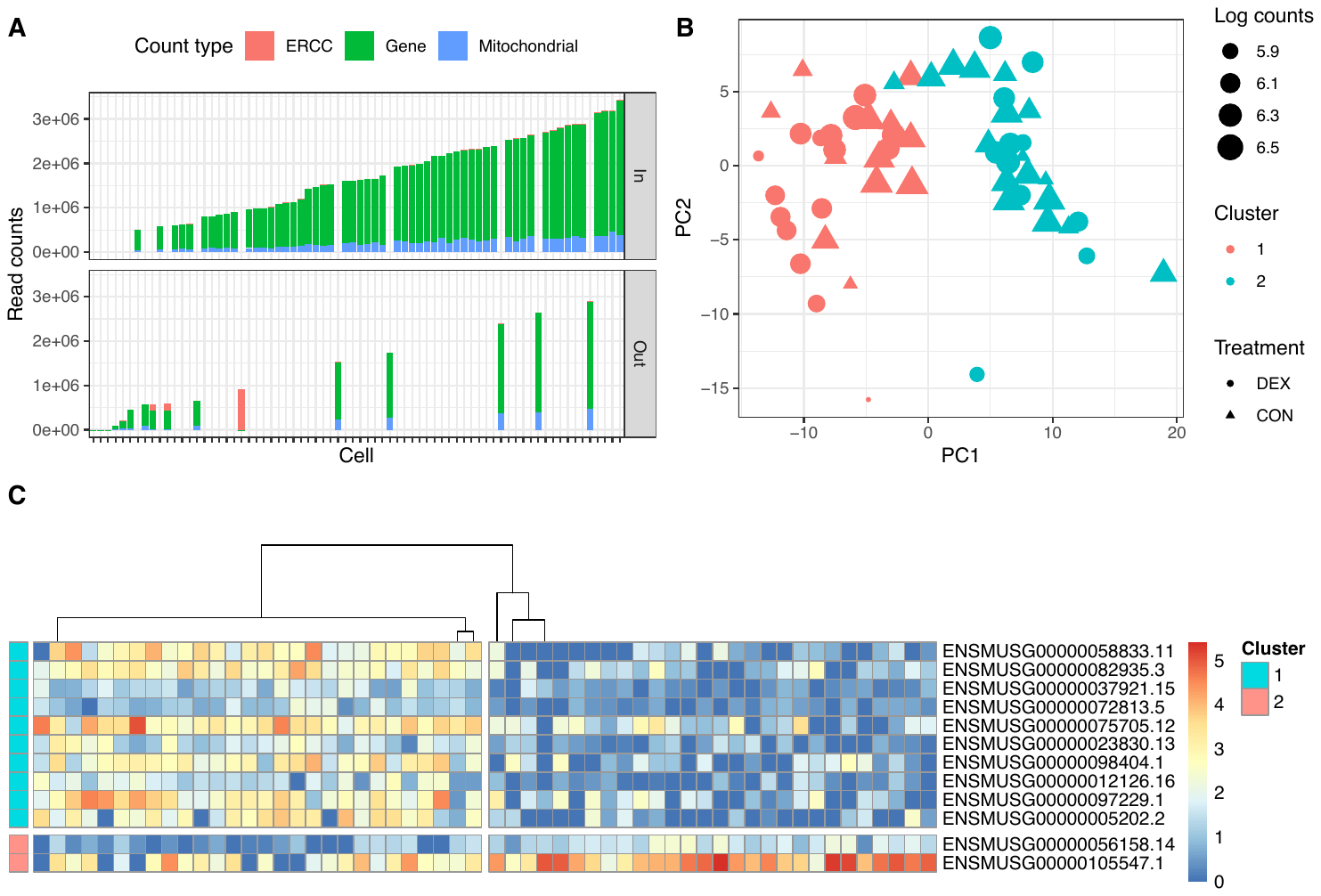

Single- 2-cell embryo sequencing using the Smartseq method. (A) Result of the quality filters implemented to select the 2 cell embryos that were used for downstream analyses. Libraries (embryos/cells) that contained a suboptimal number of mitochondrial (>15%) or ERCC mapping reads (>10%) or yielded less than 500 000 reads were excluded (displayed in Out). Most libraries showed a high number of gene mapping reads and were retained and processed for further analysis (displayed in In). (B) PCA results shown as a single panel, revealing segregation of embryos for the first two principle components (matching almost perfectly cluster 1 (red) and cluster 2(blue)) and additionally depicting read count number (size of circle) and assignment to treatments (offspring embryos of control fathers = CON, dots and fathers who were injected with dexamethasone 14 days prior sperm harvest = DEX, triangles). PC2 is attributed to technical factors such as read counts. (C) Marker genes identified by sc3 for C1 and C2 clusters.

**­­Supplementary Figure 8**

**A**

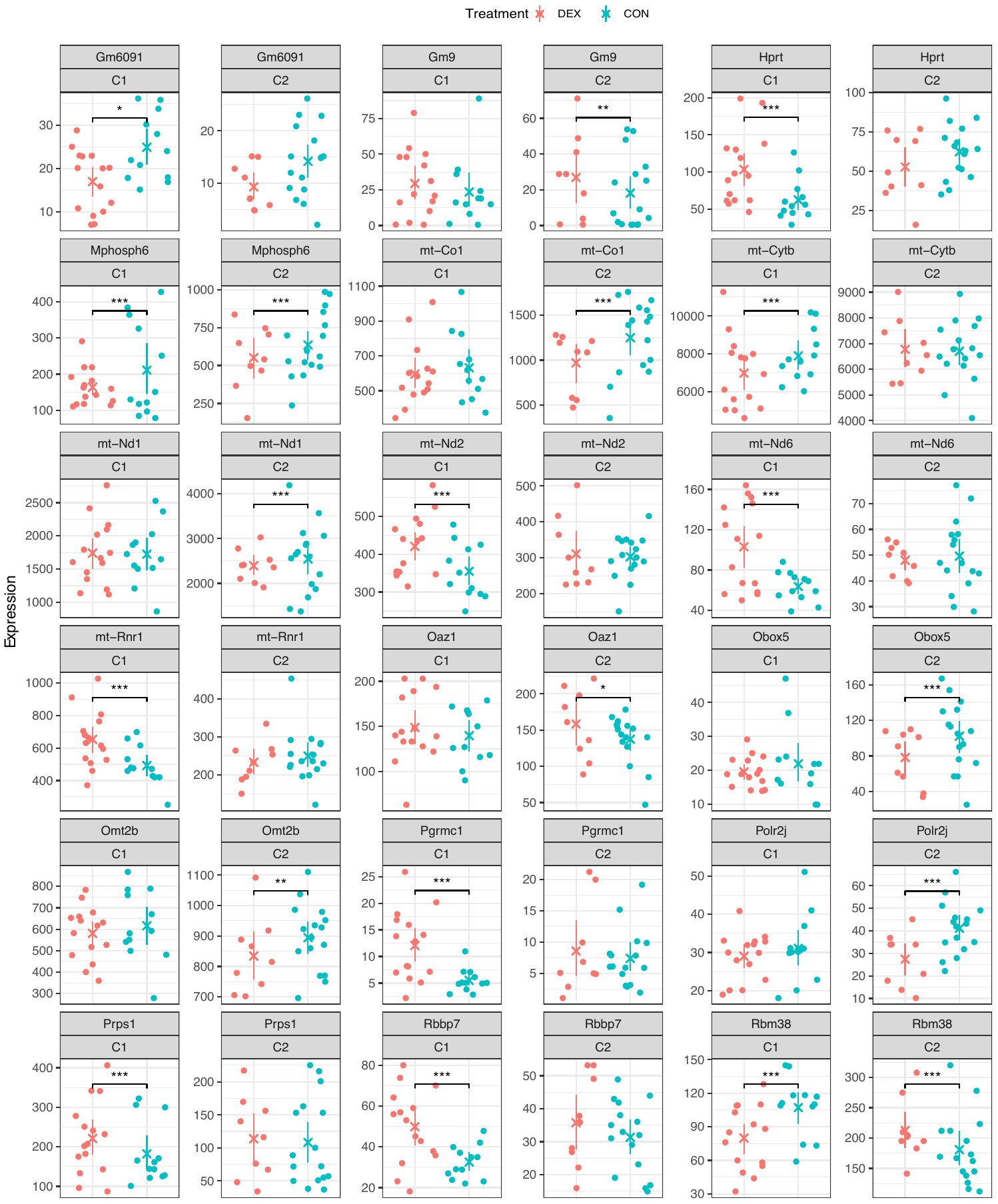

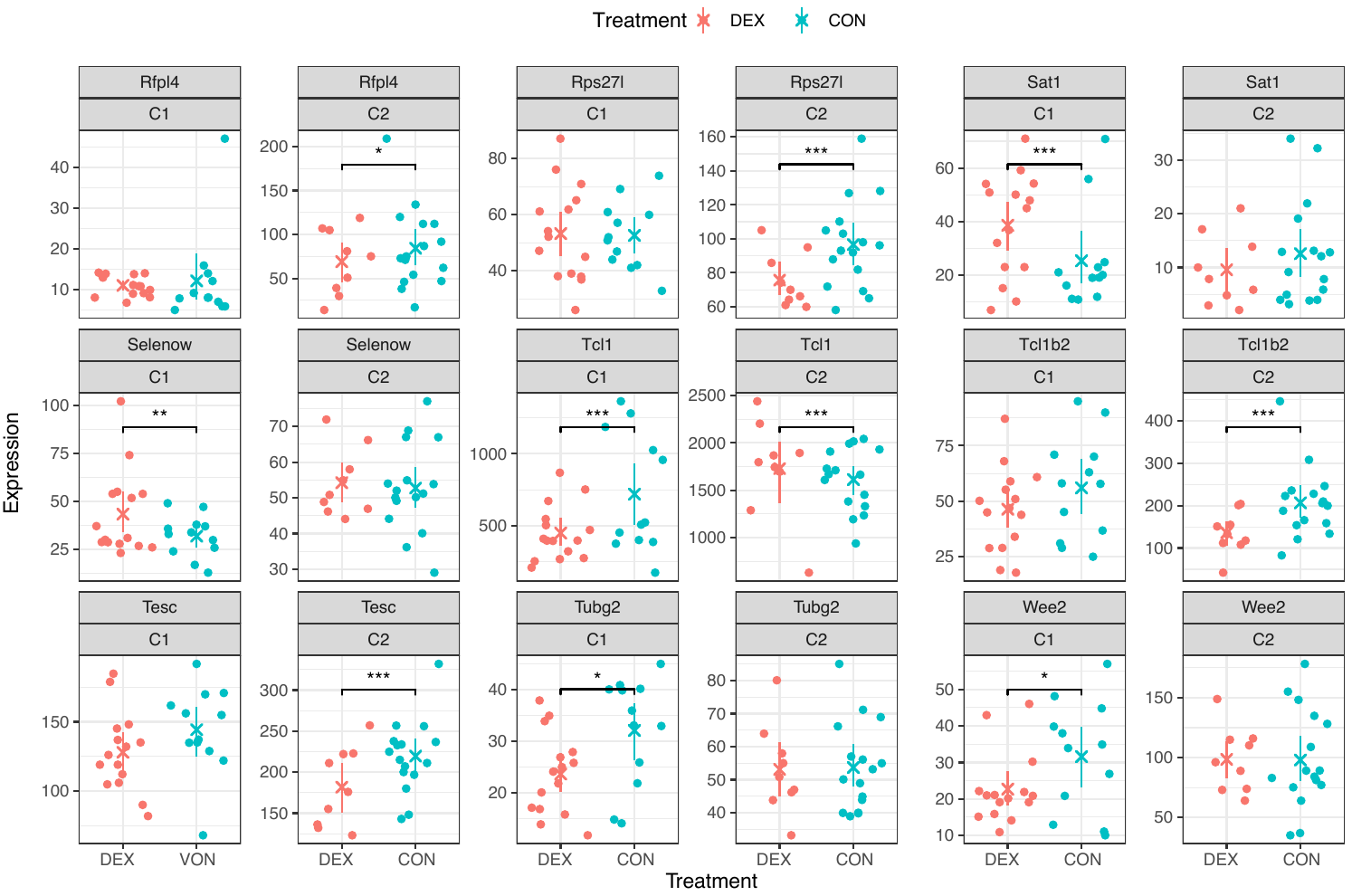

**B**

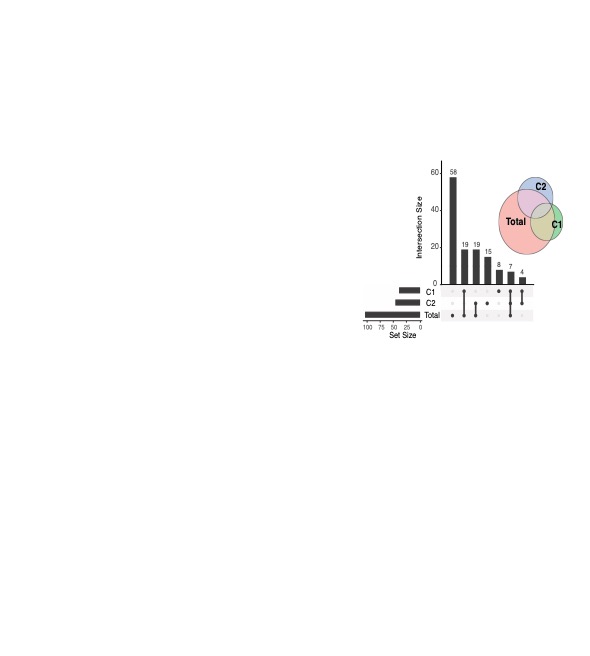

Differential gene expression analysis of 2-cell embryonic offspring of dex injected and control males using Monocle. (A) Single embryo sequencing reveals several statistically differentially expressed genes in cluster 1 and/or cluster 2. (B) Significantly affected genes overlap partially between clusters 1 and 2.

**Supplementary Figure 9**

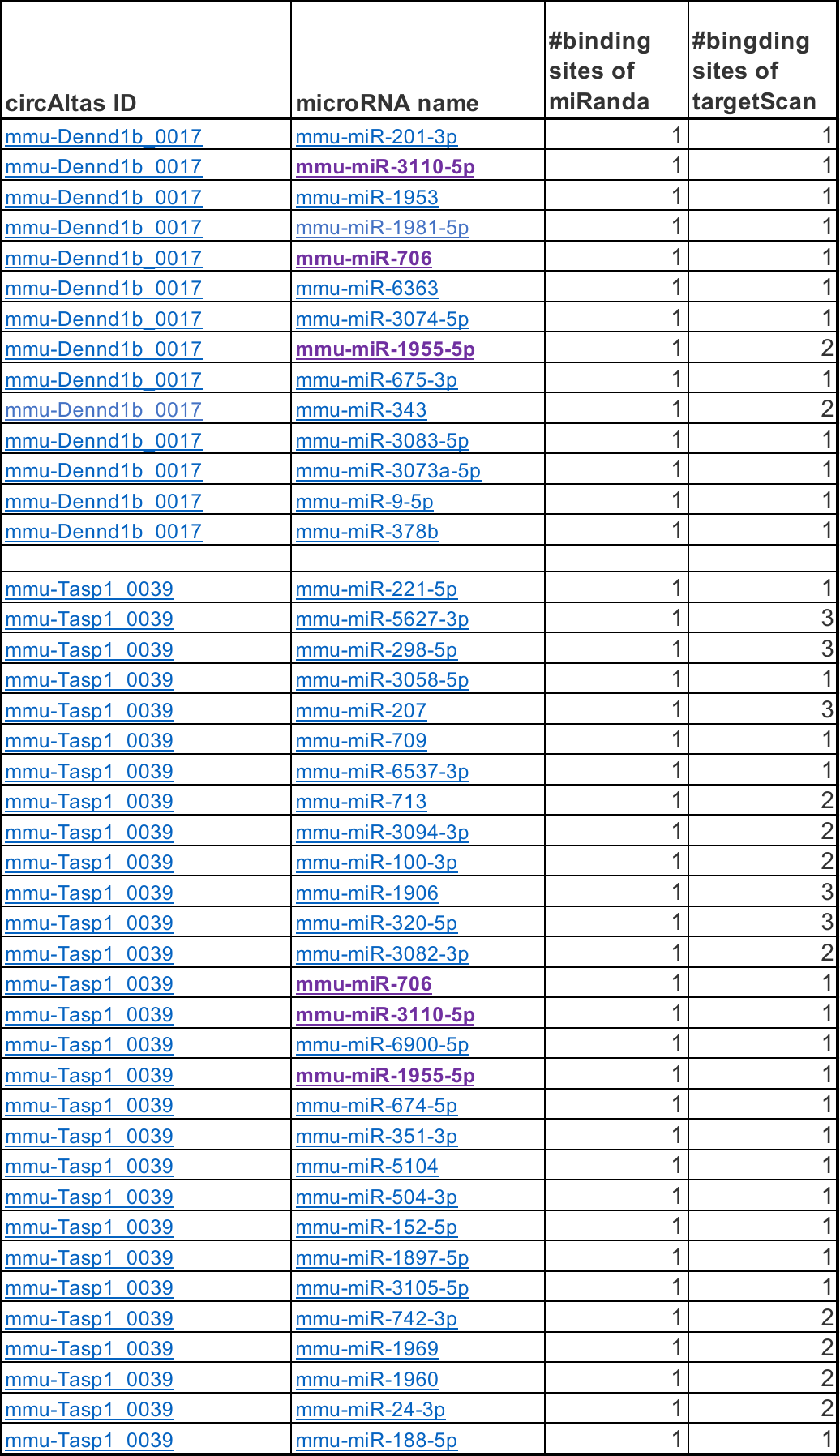

CircRNA targets as predicted by CircAtlas. Each column displays results from miRanda and targetScan.

miRNAs that are targeted by both circRNAs are highlighted in purple.

**Supplementary Figure 10**

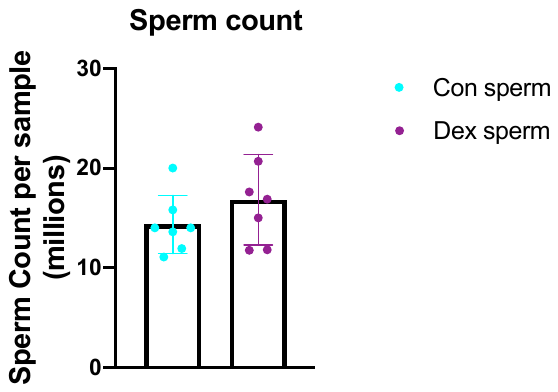

Sperm cell count 14 days post vehicle (n=7) or Dex (n=7) injection did not reveal any difference in sperm number (t(12)=1.221, p=0.25) suggesting no impairing effect of Dex on spermatogenesis.

**Supplementary Figure 11**

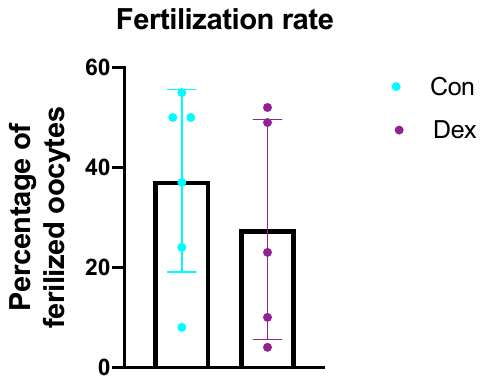

Count of fertilized oocytes (as of the appearance of the second pronucleus) over total available oocytes of 6 replicates of cryopreserved sperm from a pool of 2 dex injected versus 2 vehicle injected males did reveal no significant difference in the fertilization rate between dex and vehicle sperm.

**Supplementary Figure 12**

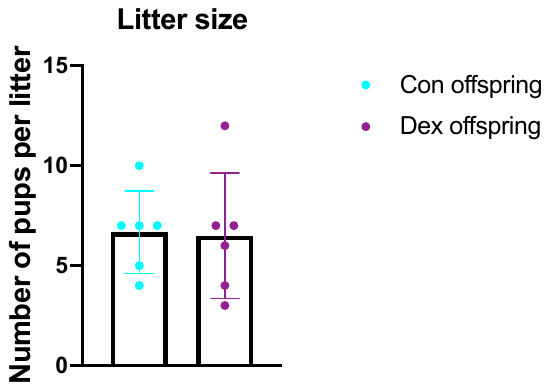

Number of pups per litter was similar in offspring resulting from in vitro fertilization with sperm from fathers that were injected with Vehicle (n=6) or Dex (n=6) (t(12)=1.122, p=0.25).
