## Supplementary Table 1 for "Single paternal Dexamethasone challenge programs offspring metabolism and reveals circRNAs as novel candidates in RNA-mediated inheritance"

| FGCZ6510_Free_miRNA_dedup_count | | | FGCZ6510_Free_miRNA_dedup_count | | | FGCZ6510_Free_miRNA_dedup_count | | | FGCZ6510_F |
|---|---|---|---|---|---|---|---|---|---|
| | CON.11 | CON.12 | CON.13 | CON.14 | CON.15 | CON.16 | CON.17 | CON.18 | DEX.3 |
| miR-3535 | 94.6751851 | 110.689369 | 83.225827 | 43.9455155 | 55.0620579 | 81.3113014 | 41.1167327 | 64.0963362 | 188.769212 |
| miR-677-5p | 44.5530283 | 38.2629916 | 33.4535187 | 14.6485052 | 14.71956 | 7.5890548 | 7.08909184 | 12.2088259 | 159.96352 |
| miR-677-3p | 58.4758496 | 21.8645666 | 11.8311225 | 7.64269835 | 2.72584445 | 20.598863 | 9.92472857 | 1.52610324 | 66.8046886 |
| miR-34c-5p | 735.124967 | 1127.39172 | 693.956528 | 650.266252 | 731.61665 | 660.247767 | 725.923004 | 799.678099 | 495.212737 |
| miR-7033-5p | 11.1382571 | 20.4980312 | 17.9506686 | 18.4698544 | 10.3582089 | 10.8415069 | 7.08909184 | 0 | 41.0634324 |
| miR-703 | 11.1382571 | 8.19921248 | 2.44781844 | 2.54756612 | 1.09033778 | 6.50490411 | 0 | 0 | 27.5799173 |
| miR-5126 | 22.2765142 | 15.0318896 | 5.71157636 | 15.2853967 | 7.08719557 | 1.08415069 | 2.83563673 | 3.05220649 | 43.5149806 |
| miR-5121 | 25.0610784 | 64.2271645 | 87.3055244 | 75.790092 | 42.5231734 | 16.2622603 | 9.92472857 | 10.6827227 | 117.674314 |
| miR-6240 | 19.4919499 | 36.8964562 | 44.8766714 | 47.7668647 | 27.8036134 | 6.50490411 | 2.83563673 | 3.05220649 | 117.061427 |
| miR-5114 | 12.5305392 | 12.2988187 | 4.48766714 | 0.63689153 | 2.72584445 | 4.33660274 | 0 | 1.52610324 | 14.7092892 |
| miR-1839-3p | 6.96141067 | 10.9322833 | 3.26375792 | 3.82134918 | 3.27101334 | 2.16830137 | 2.83563673 | 0 | 14.7092892 |
| miR-1949 | 11.1382571 | 31.4303145 | 6.93548559 | 3.82134918 | 2.18067556 | 6.50490411 | 0 | 7.63051622 | 41.6763195 |
| miR-196a-5p | 8.35369281 | 21.8645666 | 20.8064568 | 17.1960713 | 27.2584445 | 31.4403699 | 9.92472857 | 10.6827227 | 44.7407547 |
| miR-3064-5p | 38.9838998 | 34.1633854 | 15.0948804 | 5.09513224 | 12.5388845 | 21.6830137 | 17.0138204 | 13.7349292 | 32.4830137 |
| miR-196b-5p | 38.9838998 | 31.4303145 | 30.1897608 | 34.3921426 | 56.1523956 | 22.7671644 | 14.1781837 | 13.7349292 | 93.7717188 |
| miR-6538 | 801.954509 | 493.319284 | 207.248628 | 343.921426 | 96.4948935 | 83.4796028 | 58.1305531 | 36.6264778 | 848.235679 |
| miR-18a-5p | 22.2765142 | 31.4303145 | 30.1897608 | 43.9455155 | 32.1649645 | 15.1781096 | 24.1029122 | 35.1003746 | 18.3866115 |
| miR-3963 | 2.78456427 | 6.83267707 | 11.015183 | 19.7436374 | 7.63236446 | 32.5245206 | 24.1029122 | 7.63051622 | 3.67732231 |
| miR-471-5p | 69.6141067 | 128.454329 | 127.694529 | 125.467631 | 91.5883735 | 97.5735617 | 121.93238 | 111.405537 | 79.0624296 |
| miR-5099 | 2.78456427 | 2.73307083 | 5.30360662 | 5.09513224 | 2.72584445 | 4.33660274 | 2.83563673 | 7.63051622 | 14.7092892 |
| miR-1843b-3p | 6.96141067 | 5.46614166 | 2.0398487 | 3.82134918 | 4.90652001 | 2.16830137 | 2.83563673 | 3.05220649 | 9.80619282 |
| miR-1843a-3p | 23.6687963 | 21.8645666 | 7.75142507 | 3.18445765 | 8.17753335 | 13.0098082 | 11.3425469 | 3.05220649 | 28.1928043 |
| miR-690 | 2.78456427 | 2.73307083 | 2.85578818 | 1.27378306 | 0.54516889 | 1.08415069 | 0 | 0 | 4.29020936 |
| miR-6516-3p | 12.5305392 | 15.0318896 | 4.0796974 | 0.63689153 | 1.09033778 | 1.08415069 | 0 | 3.05220649 | 18.9994986 |
| miR-199a-3p | 15.3151035 | 17.7649604 | 28.1499121 | 24.2018781 | 17.9905734 | 15.1781096 | 11.3425469 | 7.63051622 | 37.9989972 |
| miR-3068-3p | 583.366214 | 1078.19644 | 362.685099 | 146.485052 | 185.902591 | 319.824452 | 127.603653 | 219.758867 | 836.590825 |
| miR-1a-3p | 38.9838998 | 50.5618103 | 37.5332161 | 30.5707934 | 46.3393556 | 56.3758356 | 56.7127347 | 27.4698584 | 69.2562368 |
| miR-152-3p | 23.6687963 | 39.629527 | 45.2846412 | 32.481468 | 31.0746267 | 36.8611233 | 25.5207306 | 10.6827227 | 62.5144792 |

| | | | | | | | | | |
|---|---|---|---|---|---|---|---|---|---|
| miR-143-3p | 1800.2208 | 2286.21375 | 2454.75393 | 3418.19684 | 2502.87037 | 1731.38864 | 1770.85514 | 862.248332 | 3139.20748 |
| miR-7a-5p | 167.073856 | 244.609839 | 221.119599 | 182.150977 | 203.893165 | 328.497658 | 317.591314 | 363.212572 | 155.673311 |
| let-7d-3p | 43.1607462 | 64.2271645 | 44.4687017 | 43.9455155 | 49.0652001 | 126.84563 | 143.199655 | 94.6184011 | 36.160336 |
| miR-465c-3p | 275.671863 | 184.482281 | 151.356774 | 141.38992 | 271.494107 | 249.354658 | 262.296398 | 378.473604 | 91.3201706 |
| miR-465a-5p | 225.549706 | 258.275193 | 281.499121 | 243.292564 | 245.326 | 251.522959 | 262.296398 | 335.742713 | 187.543438 |
| miR-465b-3p | 278.456427 | 183.115745 | 153.804592 | 142.663703 | 273.129614 | 250.438808 | 262.296398 | 381.525811 | 93.1588318 |
| miR-1892 | 1.39228213 | 0 | 1.22390922 | 0 | 0.54516889 | 0 | 0 | 0 | 1.83866115 |
| miR-7657-5p | 1.39228213 | 0 | 0 | 0 | 0 | 0 | 0 | 0 | 2.4515482 |
| miR-465a-3p | 139.228213 | 91.5578727 | 76.6983112 | 71.3318513 | 136.837391 | 124.677329 | 130.43929 | 190.762905 | 46.5794159 |
| miR-880-3p | 47.3375926 | 32.7968499 | 59.1556124 | 83.4327903 | 41.9780045 | 29.2720685 | 35.4454592 | 42.7308908 | 31.8701267 |
| miR-199b-3p | 6.96141067 | 9.5657479 | 14.2789409 | 11.4640475 | 8.72270224 | 7.5890548 | 5.67127347 | 4.57830973 | 18.3866115 |
| miR-382-5p | 4.1768464 | 1.36653541 | 2.0398487 | 3.82134918 | 6.54202668 | 3.25245206 | 2.83563673 | 0 | 7.96753166 |
| let-7b-5p | 309.086634 | 347.099995 | 139.525651 | 126.741414 | 293.300863 | 434.744425 | 526.010614 | 241.124312 | 460.891062 |
| miR-3084-3p | 43.1607462 | 24.5976375 | 17.1347291 | 7.64269835 | 5.4516889 | 8.67320548 | 1.41781837 | 3.05220649 | 33.0959008 |
| miR-9-3p | 13.9228213 | 27.3307083 | 19.1745778 | 16.5591798 | 22.8970934 | 17.346411 | 15.596002 | 18.3132389 | 11.644854 |
| miR-671-5p | 0 | 2.73307083 | 0.40796974 | 0 | 0.54516889 | 0 | 0 | 3.05220649 | 0.61288705 |
| miR-466b-3p | 25.0610784 | 13.6653541 | 19.5825475 | 17.1960713 | 23.4422623 | 8.67320548 | 12.7603653 | 21.3654454 | 13.4835151 |
| miR-362-3p | 9.74597494 | 15.0318896 | 13.8709712 | 11.4640475 | 8.17753335 | 8.67320548 | 12.7603653 | 15.2610324 | 12.8706281 |
| miR-222-3p | 107.205724 | 124.354723 | 100.360556 | 164.318015 | 133.566378 | 70.4697945 | 75.1443735 | 77.8312654 | 187.543438 |
| miR-194-5p | 48.7298747 | 47.8287395 | 65.6831282 | 49.6775393 | 38.1618223 | 39.0294247 | 51.0414612 | 65.6224395 | 39.8376583 |
| miR-340-3p | 29.2379248 | 38.2629916 | 48.1404294 | 60.5046953 | 49.0652001 | 81.3113014 | 58.1305531 | 68.6746459 | 33.0959008 |
| miR-144-3p | 5.56912854 | 1.36653541 | 5.30360662 | 5.09513224 | 2.72584445 | 2.16830137 | 1.41781837 | 0 | 0.61288705 |
| miR-145a-3p | 93.282903 | 97.0240144 | 59.9715518 | 80.2483327 | 79.049489 | 56.3758356 | 52.4592796 | 21.3654454 | 140.351135 |
| miR-34b-5p | 6.96141067 | 12.2988187 | 7.75142507 | 6.36891529 | 9.26787113 | 3.25245206 | 5.67127347 | 10.6827227 | 4.90309641 |
| miR-26b-5p | 307.694352 | 675.068495 | 571.973576 | 456.014335 | 636.757263 | 646.153808 | 504.743339 | 529.557825 | 960.394009 |
| miR-224-5p | 1.39228213 | 5.46614166 | 6.52751585 | 6.36891529 | 15.8098978 | 13.0098082 | 2.83563673 | 10.6827227 | 23.902595 |
| miR-743b-3p | 370.347048 | 308.837004 | 370.844494 | 250.935263 | 303.113903 | 389.210096 | 347.3655 | 436.465528 | 244.541933 |
| miR-465c-5p | 501.221568 | 441.390939 | 348.406158 | 352.201016 | 406.150823 | 666.752671 | 557.202618 | 663.854911 | 341.378087 |
| miR-10a-3p | 20.884232 | 40.9960624 | 33.4535187 | 28.0232273 | 40.8876667 | 46.6184795 | 8.5069102 | 25.9437551 | 51.4825123 |
| miR-296-5p | 8.35369281 | 5.46614166 | 5.30360662 | 12.1009391 | 10.3582089 | 3.25245206 | 2.83563673 | 6.10441297 | 4.29020936 |

| | | | | | | | | | |
|---|---|---|---|---|---|---|---|---|---|
| miR-320-3p | 29.2379248 | 30.0637791 | 13.4630014 | 17.1960713 | 27.2584445 | 19.5147123 | 18.4316388 | 16.7871357 | 34.3216749 |
| miR-26a-5p | 2024.37822 | 2287.58028 | 1984.77279 | 1883.28825 | 2448.35348 | 2025.19348 | 1813.38969 | 1561.20362 | 3292.42924 |
| miR-434-3p | 2.78456427 | 6.83267707 | 11.4231527 | 13.3747221 | 5.99685779 | 7.5890548 | 15.596002 | 7.63051622 | 19.6123856 |
| miR-340-5p | 239.472527 | 198.147635 | 199.497203 | 159.859774 | 192.444618 | 300.30974 | 221.179665 | 329.6383 | 161.189294 |
| miR-185-5p | 2.78456427 | 2.73307083 | 4.89563688 | 3.82134918 | 5.4516889 | 9.75735617 | 1.41781837 | 3.05220649 | 6.12887051 |
| miR-1291 | 12.5305392 | 25.9641729 | 18.3586383 | 12.7378306 | 11.4485467 | 18.4305616 | 11.3425469 | 7.63051622 | 25.7412561 |
| miR-743a-3p | 534.63634 | 471.454718 | 281.499121 | 270.6789 | 410.512174 | 417.398014 | 307.666586 | 561.605993 | 367.119344 |
| miR-31-3p | 2.78456427 | 0 | 0.81593948 | 0 | 0.54516889 | 0 | 0 | 0 | 3.06443526 |
| miR-293-5p | 2.78456427 | 2.73307083 | 8.56736455 | 11.4640475 | 3.81618223 | 4.33660274 | 5.67127347 | 10.6827227 | 0 |
| miR-7672-5p | 0 | 0 | 0.40796974 | 0 | 1.09033778 | 0 | 0 | 0 | 0 |
| miR-7670-3p | 1.39228213 | 1.36653541 | 0.40796974 | 0 | 0 | 1.08415069 | 0 | 0 | 1.2257741 |
| miR-871-3p | 161.504728 | 131.1874 | 168.083533 | 122.283174 | 158.098978 | 195.147123 | 259.460761 | 299.116236 | 95.61038 |
| miR-23a-5p | 1.39228213 | 1.36653541 | 1.63187896 | 1.27378306 | 2.18067556 | 0 | 2.83563673 | 0 | 3.06443526 |
| miR-130b-5p | 48.7298747 | 45.0956687 | 48.5483991 | 66.8736106 | 40.8876667 | 117.088274 | 76.5621918 | 57.9919232 | 52.0953993 |
| miR-449a-5p | 37.5916176 | 40.9960624 | 42.8368227 | 40.1241664 | 59.9685779 | 22.7671644 | 29.7741857 | 28.9959616 | 28.1928043 |
| miR-29c-3p | 4.1768464 | 13.6653541 | 30.1897608 | 24.8387696 | 21.8067556 | 8.67320548 | 14.1781837 | 15.2610324 | 31.8701267 |
| miR-3064-3p | 1.39228213 | 1.36653541 | 1.22390922 | 0.63689153 | 0 | 0 | 0 | 1.52610324 | 3.06443526 |
| miR-466n-3p | 2.78456427 | 4.09960624 | 1.22390922 | 3.82134918 | 2.18067556 | 3.25245206 | 1.41781837 | 3.05220649 | 0.61288705 |
| miR-34b-3p | 1784.9057 | 1048.13266 | 1595.97762 | 1889.02028 | 1281.14689 | 1103.6654 | 1552.51111 | 1637.50878 | 1220.25812 |
| miR-149-5p | 2.78456427 | 2.73307083 | 1.63187896 | 5.09513224 | 1.63550667 | 1.08415069 | 9.92472857 | 1.52610324 | 1.2257741 |
| miR-7652-5p | 0 | 0 | 0 | 0 | 0 | 0 | 0 | 0 | 0.61288705 |
| miR-1306-3p | 0 | 0 | 0.40796974 | 1.27378306 | 0 | 0 | 0 | 1.52610324 | 1.2257741 |
| miR-145a-5p | 137.835931 | 83.3586603 | 77.5142507 | 150.306401 | 92.6787113 | 21.6830137 | 38.2810959 | 39.6786843 | 136.060925 |
| miR-23a-3p | 389.838998 | 307.470468 | 296.594001 | 343.284534 | 366.898663 | 366.442932 | 345.947682 | 293.011823 | 349.958506 |
| miR-133a-3p | 6.96141067 | 1.36653541 | 1.63187896 | 4.45824071 | 4.90652001 | 2.16830137 | 0 | 0 | 8.58041871 |
| miR-30a-3p | 83.5369281 | 117.522046 | 138.709712 | 100.628862 | 121.027494 | 186.473918 | 148.870929 | 119.036053 | 160.576407 |
| miR-365-3p | 20.884232 | 15.0318896 | 18.7666081 | 22.2912035 | 17.9905734 | 29.2720685 | 32.6098224 | 18.3132389 | 10.4190799 |
| miR-466c-3p | 9.74597494 | 5.46614166 | 7.75142507 | 7.00580682 | 9.26787113 | 3.25245206 | 4.2534551 | 7.63051622 | 4.90309641 |
| miR-7043-3p | 0 | 0 | 0 | 1.27378306 | 1.63550667 | 2.16830137 | 2.83563673 | 1.52610324 | 0 |
| miR-7073-5p | 0 | 0 | 0 | 0 | 0 | 1.08415069 | 0 | 0 | 1.83866115 |

| | | | | | | | | | |
|---|---|---|---|---|---|---|---|---|---|
| miR-375-3p | 839.546127 | 576.677945 | 343.510521 | 454.740552 | 636.757263 | 759.98963 | 626.675718 | 799.678099 | 346.281184 |
| miR-330-5p | 1.39228213 | 1.36653541 | 2.0398487 | 1.91067459 | 1.09033778 | 2.16830137 | 0 | 0 | 4.29020936 |
| let-7c-5p | 1807.18221 | 2584.11847 | 1012.17293 | 935.593657 | 2206.84367 | 3338.09996 | 3598.42302 | 1961.04267 | 2580.25448 |
| miR-669a-3p | 20.884232 | 12.2988187 | 16.3187896 | 40.1241664 | 22.3519245 | 5.42075343 | 2.83563673 | 18.3132389 | 6.12887051 |
| miR-181b-5p | 9.74597494 | 17.7649604 | 7.34345533 | 15.2853967 | 18.5357423 | 13.0098082 | 2.83563673 | 10.6827227 | 25.7412561 |
| miR-744-3p | 0 | 0 | 2.44781844 | 1.91067459 | 0.54516889 | 0 | 0 | 0 | 4.29020936 |
| miR-425-5p | 911.944798 | 502.885032 | 743.728837 | 722.871886 | 597.505103 | 624.470795 | 666.374633 | 731.003453 | 449.859095 |
| miR-22-3p | 311.871198 | 370.331097 | 550.35118 | 833.691012 | 626.944223 | 294.888986 | 401.242598 | 384.578017 | 189.994986 |
| miR-205-5p | 55.6912854 | 24.5976375 | 38.7571253 | 60.5046953 | 64.8750979 | 30.3562192 | 17.0138204 | 30.5220649 | 69.8691238 |
| miR-676-5p | 0 | 4.09960624 | 3.26375792 | 7.64269835 | 3.81618223 | 8.67320548 | 4.2534551 | 12.2088259 | 0.61288705 |
| miR-672-3p | 0 | 1.36653541 | 0.81593948 | 0 | 0.54516889 | 0 | 0 | 0 | 1.83866115 |
| miR-192-5p | 11.1382571 | 32.7968499 | 24.4781844 | 26.1125527 | 16.3550667 | 16.2622603 | 29.7741857 | 27.4698584 | 14.7092892 |
| miR-1306-5p | 11.1382571 | 6.83267707 | 7.34345533 | 10.1902645 | 5.99685779 | 23.8513151 | 12.7603653 | 22.8915486 | 9.80619282 |
| miR-297a-3p | 2.78456427 | 2.73307083 | 3.67172766 | 6.36891529 | 3.27101334 | 2.16830137 | 2.83563673 | 6.10441297 | 1.2257741 |
| miR-196a-1-3p | 0 | 2.73307083 | 1.63187896 | 0.63689153 | 0.54516889 | 0 | 0 | 0 | 1.83866115 |
| miR-3089-3p | 0 | 0 | 0.40796974 | 0 | 0 | 0 | 0 | 0 | 1.83866115 |
| miR-883b-5p | 5.56912854 | 1.36653541 | 2.0398487 | 1.91067459 | 1.09033778 | 4.33660274 | 0 | 6.10441297 | 2.4515482 |
| miR-501-3p | 29.2379248 | 17.7649604 | 15.0948804 | 5.73202376 | 9.81304002 | 20.598863 | 22.6850939 | 30.5220649 | 7.96753166 |
| miR-187-3p | 0 | 5.46614166 | 4.48766714 | 3.18445765 | 3.27101334 | 4.33660274 | 0 | 3.05220649 | 6.12887051 |
| miR-1902 | 1.39228213 | 0 | 0 | 0 | 0.54516889 | 0 | 1.41781837 | 0 | 3.06443526 |
| miR-7b-5p | 1.39228213 | 0 | 1.63187896 | 1.91067459 | 1.09033778 | 4.33660274 | 1.41781837 | 6.10441297 | 0.61288705 |
| miR-741-3p | 186.565806 | 184.482281 | 170.939321 | 123.556957 | 197.896307 | 242.849753 | 172.973841 | 283.855203 | 161.189294 |
| miR-15b-3p | 0 | 0 | 0.40796974 | 0 | 0.54516889 | 0 | 0 | 0 | 1.2257741 |
| miR-466d-3p | 9.74597494 | 2.73307083 | 8.15939481 | 8.27958988 | 12.5388845 | 4.33660274 | 7.08909184 | 6.10441297 | 6.12887051 |
| miR-881-3p | 484.514183 | 690.100384 | 633.984976 | 648.355577 | 420.325214 | 944.295247 | 686.22409 | 938.553494 | 437.601354 |
| miR-30d-3p | 0 | 2.73307083 | 0.40796974 | 2.54756612 | 3.27101334 | 0 | 4.2534551 | 1.52610324 | 0.61288705 |
| miR-200b-5p | 33.4147712 | 36.8964562 | 20.398487 | 28.0232273 | 22.8970934 | 36.8611233 | 36.8632776 | 32.0481681 | 34.9345619 |
| miR-128-3p | 359.208791 | 285.605902 | 287.210697 | 264.309985 | 239.874312 | 490.03611 | 462.208788 | 523.453412 | 323.604363 |
| miR-99b-3p | 0 | 2.73307083 | 1.63187896 | 0.63689153 | 1.63550667 | 1.08415069 | 0 | 1.52610324 | 1.83866115 |
| miR-546 | 4.1768464 | 2.73307083 | 0 | 0 | 0.54516889 | 0 | 2.83563673 | 0 | 2.4515482 |

| | | | | | | | | | |
|---|---|---|---|---|---|---|---|---|---|
| miR-379-5p | 25.0610784 | 31.4303145 | 33.4535187 | 36.3028172 | 48.5200312 | 46.6184795 | 34.0276408 | 19.8393422 | 25.1283691 |
| miR-7238-5p | 1.39228213 | 0 | 0 | 0 | 0 | 1.08415069 | 0 | 0 | 0 |
| miR-7219-5p | 1.39228213 | 0 | 0 | 0 | 0 | 0 | 0 | 0 | 1.83866115 |
| miR-664-3p | 118.343981 | 65.5936999 | 25.2941239 | 20.3805289 | 32.1649645 | 18.4305616 | 7.08909184 | 27.4698584 | 71.0948979 |
| miR-106b-5p | 5.56912854 | 5.46614166 | 9.79127377 | 4.45824071 | 3.27101334 | 2.16830137 | 4.2534551 | 3.05220649 | 11.644854 |
| miR-743b-5p | 16.7073856 | 10.9322833 | 15.0948804 | 12.7378306 | 7.08719557 | 19.5147123 | 5.67127347 | 24.4176519 | 6.12887051 |
| miR-297c-3p | 1.39228213 | 1.36653541 | 1.63187896 | 3.18445765 | 1.63550667 | 1.08415069 | 1.41781837 | 3.05220649 | 0.61288705 |
| miR-297b-3p | 1.39228213 | 1.36653541 | 1.63187896 | 3.18445765 | 1.63550667 | 1.08415069 | 1.41781837 | 3.05220649 | 0.61288705 |
| miR-484 | 11.1382571 | 4.09960624 | 7.75142507 | 7.00580682 | 6.54202668 | 3.25245206 | 7.08909184 | 12.2088259 | 10.4190799 |
| miR-465b-5p | 278.456427 | 322.502358 | 212.552235 | 191.70435 | 241.509818 | 407.640658 | 301.995312 | 376.947501 | 264.767206 |
| miR-130c | 0 | 0 | 0 | 0 | 0 | 0 | 0 | 0 | 0 |
| miR-19a-3p | 9.74597494 | 21.8645666 | 14.6869107 | 11.4640475 | 6.54202668 | 6.50490411 | 5.67127347 | 13.7349292 | 12.8706281 |
| miR-494-3p | 0 | 5.46614166 | 0.40796974 | 1.91067459 | 1.63550667 | 1.08415069 | 0 | 1.52610324 | 1.2257741 |
| miR-664-5p | 18.0996677 | 5.46614166 | 0.81593948 | 1.91067459 | 3.81618223 | 6.50490411 | 0 | 0 | 4.90309641 |
| miR-30e-5p | 338.324559 | 602.642118 | 687.021043 | 684.021503 | 484.655143 | 430.407822 | 433.85242 | 387.630224 | 416.150308 |
| miR-499-5p | 4.1768464 | 9.5657479 | 18.7666081 | 22.2912035 | 6.54202668 | 6.50490411 | 2.83563673 | 10.6827227 | 6.74175756 |
| miR-470-5p | 226.941988 | 165.350785 | 112.599648 | 120.372499 | 170.092694 | 242.849753 | 224.015302 | 280.802997 | 87.6428483 |
| miR-741-5p | 5.56912854 | 9.5657479 | 11.4231527 | 15.9222882 | 8.72270224 | 4.33660274 | 5.67127347 | 0 | 6.74175756 |
| miR-200a-5p | 40.3761819 | 35.5299208 | 29.3738213 | 43.308624 | 23.9874312 | 35.7769726 | 25.5207306 | 22.8915486 | 28.8056914 |
| miR-29b-1-5p | 2.78456427 | 1.36653541 | 2.85578818 | 2.54756612 | 1.63550667 | 9.75735617 | 4.2534551 | 4.57830973 | 4.29020936 |
| miR-871-5p | 140.620496 | 120.255116 | 124.838741 | 128.015197 | 117.75648 | 92.1528082 | 85.069102 | 180.080183 | 135.448038 |
| miR-410-5p | 0 | 0 | 0 | 0 | 0 | 0 | 0 | 0 | 1.83866115 |
| miR-139-5p | 15.3151035 | 12.2988187 | 22.030366 | 20.3805289 | 16.9002356 | 20.598863 | 18.4316388 | 10.6827227 | 16.5479504 |
| miR-199a-5p | 6.96141067 | 17.7649604 | 31.0057003 | 14.6485052 | 24.5326 | 7.5890548 | 5.67127347 | 4.57830973 | 23.902595 |
| miR-467c-5p | 26.4533606 | 20.4980312 | 32.2296095 | 35.0290341 | 28.3487823 | 17.346411 | 25.5207306 | 30.5220649 | 26.3541432 |
| miR-433-3p | 1.39228213 | 0 | 0 | 0 | 0 | 0 | 0 | 0 | 0 |
| let-7d-5p | 417.68464 | 459.155899 | 228.463055 | 224.185818 | 441.586801 | 453.174986 | 496.236429 | 444.096044 | 503.793156 |
| miR-1247-5p | 11.1382571 | 1.36653541 | 4.89563688 | 15.2853967 | 10.9033778 | 4.33660274 | 9.92472857 | 3.05220649 | 0.61288705 |
| miR-7229-3p | 6.96141067 | 5.46614166 | 1.22390922 | 1.27378306 | 1.63550667 | 0 | 0 | 0 | 2.4515482 |
| miR-155-5p | 18.0996677 | 35.5299208 | 13.8709712 | 16.5591798 | 17.4454045 | 13.0098082 | 12.7603653 | 30.5220649 | 24.515482 |

| | | | | | | | | | |
|---|---|---|---|---|---|---|---|---|---|
| miR-124-3p | 22.2765142 | 13.6653541 | 34.6774279 | 39.4872748 | 23.9874312 | 24.9354658 | 29.7741857 | 18.3132389 | 10.4190799 |
| miR-7241-3p | 8.35369281 | 16.398425 | 4.0796974 | 5.09513224 | 5.99685779 | 4.33660274 | 7.08909184 | 1.52610324 | 2.4515482 |
| miR-8094 | 18.0996677 | 6.83267707 | 1.22390922 | 0 | 0.54516889 | 2.16830137 | 2.83563673 | 10.6827227 | 8.58041871 |
| miR-3066-3p | 0 | 0 | 0 | 0.63689153 | 0 | 0 | 0 | 0 | 0 |
| miR-5100 | 6.96141067 | 6.83267707 | 5.71157636 | 12.7378306 | 5.99685779 | 3.25245206 | 2.83563673 | 1.52610324 | 6.74175756 |
| miR-382-3p | 0 | 0 | 2.0398487 | 0.63689153 | 1.63550667 | 1.08415069 | 1.41781837 | 0 | 0 |
| miR-409-5p | 1.39228213 | 1.36653541 | 1.22390922 | 0.63689153 | 0.54516889 | 2.16830137 | 7.08909184 | 4.57830973 | 0 |
| miR-8104 | 1.39228213 | 0 | 0.40796974 | 0.63689153 | 0 | 0 | 0 | 0 | 1.2257741 |
| miR-674-3p | 12.5305392 | 5.46614166 | 11.4231527 | 3.18445765 | 3.81618223 | 10.8415069 | 17.0138204 | 7.63051622 | 3.06443526 |
| miR-7653-5p | 1.39228213 | 0 | 0.40796974 | 1.27378306 | 1.63550667 | 0 | 0 | 0 | 2.4515482 |
| miR-202-5p | 4.1768464 | 12.2988187 | 4.89563688 | 17.1960713 | 4.36135112 | 6.50490411 | 0 | 3.05220649 | 3.06443526 |
| miR-7672-3p | 0 | 0 | 0.40796974 | 0 | 0.54516889 | 0 | 0 | 0 | 1.2257741 |
| miR-34c-3p | 182.38896 | 80.6255894 | 108.111981 | 156.675316 | 146.650431 | 100.826014 | 94.9938306 | 117.50995 | 88.2557353 |
| miR-471-3p | 15.3151035 | 15.0318896 | 21.2144265 | 15.2853967 | 15.2647289 | 5.42075343 | 7.08909184 | 9.15661946 | 10.4190799 |
| miR-93-3p | 2.78456427 | 1.36653541 | 1.63187896 | 3.82134918 | 2.72584445 | 0 | 2.83563673 | 3.05220649 | 1.2257741 |
| miR-3082-3p | 1.39228213 | 0 | 0 | 0 | 0 | 0 | 0 | 0 | 0.61288705 |
| let-7c-1-3p | 1.39228213 | 9.5657479 | 4.89563688 | 5.09513224 | 2.18067556 | 3.25245206 | 1.41781837 | 0 | 7.35464461 |
| miR-143-5p | 0 | 2.73307083 | 2.85578818 | 8.91648141 | 2.18067556 | 1.08415069 | 5.67127347 | 3.05220649 | 11.0319669 |
| miR-92a-3p | 2344.60311 | 1221.68266 | 970.560012 | 1285.884 | 1298.04713 | 1152.45218 | 1409.31146 | 1436.06315 | 867.235177 |
| miR-127-3p | 1.39228213 | 2.73307083 | 5.30360662 | 8.91648141 | 8.17753335 | 4.33660274 | 1.41781837 | 6.10441297 | 6.12887051 |
| miR-34a-5p | 4.1768464 | 8.19921248 | 5.71157636 | 6.36891529 | 5.99685779 | 8.67320548 | 4.2534551 | 9.15661946 | 7.96753166 |
| miR-98-5p | 58.4758496 | 94.2909436 | 82.0019178 | 50.9513224 | 111.214454 | 191.894671 | 138.9462 | 108.35333 | 106.02946 |
| miR-5122 | 1.39228213 | 0 | 0 | 0 | 0 | 0 | 0 | 0 | 1.2257741 |
| miR-598-3p | 0 | 1.36653541 | 1.63187896 | 1.91067459 | 0.54516889 | 4.33660274 | 2.83563673 | 0 | 0.61288705 |
| miR-5113 | 1.39228213 | 0 | 0 | 0 | 0 | 0 | 0 | 0 | 0.61288705 |
| miR-8112 | 0 | 1.36653541 | 0 | 0.63689153 | 0.54516889 | 0 | 0 | 0 | 0 |
| miR-6900-3p | 0 | 0 | 0 | 0 | 0 | 0 | 0 | 0 | 0 |
| miR-22-5p | 2.78456427 | 2.73307083 | 6.93548559 | 3.82134918 | 8.17753335 | 7.5890548 | 1.41781837 | 1.52610324 | 12.257741 |
| miR-7233-5p | 1.39228213 | 2.73307083 | 0 | 0 | 0.54516889 | 0 | 2.83563673 | 0 | 0 |
| miR-301a-3p | 0 | 1.36653541 | 1.63187896 | 5.09513224 | 1.09033778 | 0 | 0 | 0 | 0.61288705 |

| | | | | | | | | | |
|---|---|---|---|---|---|---|---|---|---|
| miR-463-5p | 44.5530283 | 50.5618103 | 60.7874913 | 82.1590073 | 65.9654357 | 11.9256575 | 29.7741857 | 53.4136135 | 58.2242698 |
| miR-200c-5p | 1.39228213 | 0 | 0.81593948 | 0.63689153 | 0 | 0 | 0 | 0 | 0 |
| miR-692 | 2.78456427 | 1.36653541 | 1.22390922 | 1.27378306 | 0 | 0 | 0 | 0 | 1.2257741 |
| miR-210-3p | 0 | 0 | 1.63187896 | 0.63689153 | 4.90652001 | 1.08415069 | 0 | 1.52610324 | 3.67732231 |
| miR-682 | 2.78456427 | 0 | 1.63187896 | 0.63689153 | 0.54516889 | 0 | 1.41781837 | 0 | 2.4515482 |
| miR-669o-3p | 1.39228213 | 1.36653541 | 1.63187896 | 3.82134918 | 2.18067556 | 0 | 0 | 1.52610324 | 0.61288705 |
| miR-29c-5p | 2.78456427 | 2.73307083 | 4.89563688 | 5.09513224 | 3.27101334 | 0 | 0 | 0 | 4.90309641 |
| miR-1893 | 1.39228213 | 0 | 0.81593948 | 0 | 0.54516889 | 0 | 0 | 0 | 1.2257741 |
| miR-497a-5p | 1.39228213 | 0 | 2.0398487 | 6.36891529 | 3.27101334 | 0 | 2.83563673 | 0 | 6.74175756 |
| miR-7230-5p | 12.5305392 | 2.73307083 | 6.11954611 | 1.91067459 | 7.08719557 | 3.25245206 | 0 | 9.15661946 | 2.4515482 |
| miR-483-3p | 6.96141067 | 0 | 2.85578818 | 1.91067459 | 1.63550667 | 0 | 0 | 1.52610324 | 0.61288705 |
| miR-7232-5p | 5.56912854 | 2.73307083 | 0.81593948 | 0.63689153 | 1.63550667 | 0 | 0 | 0 | 1.83866115 |
| miR-106b-3p | 43.1607462 | 38.2629916 | 39.9810346 | 56.0464546 | 39.2521601 | 74.8063973 | 69.4731 | 73.2529557 | 35.547449 |
| miR-98-3p | 0 | 1.36653541 | 0 | 1.91067459 | 1.09033778 | 1.08415069 | 4.2534551 | 0 | 0 |
| miR-184-3p | 334.147712 | 274.673618 | 159.516168 | 182.150977 | 275.310289 | 332.83426 | 330.35168 | 312.851165 | 155.673311 |
| miR-6516-5p | 8.35369281 | 2.73307083 | 0.81593948 | 0 | 0 | 1.08415069 | 4.2534551 | 1.52610324 | 3.06443526 |
| let-7e-5p | 47.3375926 | 51.9283457 | 15.0948804 | 21.0174205 | 49.0652001 | 39.0294247 | 56.7127347 | 35.1003746 | 54.5469475 |
| miR-7210-5p | 25.0610784 | 24.5976375 | 10.6072132 | 8.91648141 | 22.8970934 | 6.50490411 | 11.3425469 | 9.15661946 | 18.3866115 |
| miR-9-5p | 32.0224891 | 73.7929124 | 50.1802781 | 35.6659256 | 43.6135112 | 75.890548 | 59.5483714 | 45.7830973 | 40.4505454 |
| miR-181c-5p | 9.74597494 | 12.2988187 | 15.5028501 | 32.481468 | 19.0809111 | 6.50490411 | 9.92472857 | 9.15661946 | 7.96753166 |
| miR-541-5p | 6.96141067 | 10.9322833 | 10.6072132 | 5.09513224 | 19.62608 | 10.8415069 | 12.7603653 | 12.2088259 | 11.0319669 |
| miR-181b-1-3p | 0 | 1.36653541 | 0 | 0.63689153 | 0 | 0 | 0 | 0 | 0.61288705 |
| miR-872-3p | 1.39228213 | 1.36653541 | 2.0398487 | 0.63689153 | 0.54516889 | 0 | 1.41781837 | 0 | 3.67732231 |
| miR-101a-3p | 19.4919499 | 54.6614166 | 51.812157 | 63.0522614 | 40.3424978 | 48.7867808 | 53.877098 | 53.4136135 | 60.6758181 |
| miR-331-3p | 5.56912854 | 4.09960624 | 1.63187896 | 1.91067459 | 4.36135112 | 0 | 0 | 1.52610324 | 3.06443526 |
| miR-33-3p | 1.39228213 | 0 | 1.63187896 | 0.63689153 | 1.63550667 | 0 | 0 | 0 | 0 |
| miR-466e-3p | 2.78456427 | 1.36653541 | 2.85578818 | 2.54756612 | 3.27101334 | 1.08415069 | 1.41781837 | 3.05220649 | 1.83866115 |
| miR-7670-5p | 0 | 0 | 0.81593948 | 0 | 0 | 0 | 0 | 0 | 1.2257741 |
| miR-181a-5p | 82.1446459 | 68.3267707 | 77.5142507 | 187.24611 | 119.391987 | 69.3856439 | 43.9523694 | 68.6746459 | 99.9005893 |
| miR-148a-3p | 967.636083 | 1395.23266 | 1950.9113 | 1420.26811 | 1215.72662 | 2374.29 | 1894.20534 | 1260.56128 | 1464.80005 |

| | | | | | | | | | |
|---|---|---|---|---|---|---|---|---|---|
| miR-3066-5p | 4.1768464 | 0 | 1.63187896 | 1.27378306 | 2.72584445 | 8.67320548 | 1.41781837 | 1.52610324 | 0.61288705 |
| miR-191-5p | 3175.79555 | 1902.2173 | 2416.81274 | 1918.31729 | 2315.87744 | 2172.63797 | 2410.29122 | 2893.49175 | 2364.51824 |
| miR-101b-3p | 30.630207 | 58.7610228 | 47.3244899 | 41.3979494 | 38.7069912 | 49.8709315 | 43.9523694 | 28.9959616 | 47.80519 |
| miR-21a-3p | 1.39228213 | 1.36653541 | 6.93548559 | 0.63689153 | 1.63550667 | 1.08415069 | 1.41781837 | 4.57830973 | 4.29020936 |
| miR-7210-3p | 8.35369281 | 2.73307083 | 5.71157636 | 3.82134918 | 4.36135112 | 0 | 5.67127347 | 1.52610324 | 8.58041871 |
| miR-874-3p | 0 | 0 | 1.22390922 | 0 | 0.54516889 | 0 | 0 | 0 | 1.83866115 |
| miR-5125 | 5.56912854 | 2.73307083 | 0.40796974 | 0 | 0 | 0 | 0 | 1.52610324 | 3.67732231 |
| miR-27b-5p | 11.1382571 | 9.5657479 | 1.63187896 | 3.82134918 | 5.4516889 | 5.42075343 | 0 | 1.52610324 | 8.58041871 |
| miR-409-3p | 8.35369281 | 2.73307083 | 4.89563688 | 8.27958988 | 4.36135112 | 9.75735617 | 15.596002 | 12.2088259 | 4.90309641 |
| miR-147-5p | 9.74597494 | 2.73307083 | 4.89563688 | 1.27378306 | 1.09033778 | 5.42075343 | 5.67127347 | 3.05220649 | 1.83866115 |
| miR-466p-3p | 2.78456427 | 1.36653541 | 2.44781844 | 2.54756612 | 3.27101334 | 1.08415069 | 1.41781837 | 3.05220649 | 1.83866115 |
| miR-466a-3p | 2.78456427 | 1.36653541 | 2.44781844 | 2.54756612 | 3.27101334 | 1.08415069 | 1.41781837 | 3.05220649 | 1.83866115 |
| miR-223-5p | 15.3151035 | 13.6653541 | 6.11954611 | 7.64269835 | 4.36135112 | 3.25245206 | 2.83563673 | 9.15661946 | 3.06443526 |
| miR-6917-3p | 0 | 1.36653541 | 0.40796974 | 0 | 0.54516889 | 1.08415069 | 0 | 0 | 1.2257741 |
| miR-744-5p | 11.1382571 | 0 | 2.44781844 | 3.82134918 | 4.36135112 | 8.67320548 | 14.1781837 | 12.2088259 | 7.35464461 |
| miR-7654-5p | 0 | 0 | 1.22390922 | 0 | 0 | 0 | 0 | 0 | 3.06443526 |
| miR-6340 | 4.1768464 | 0 | 0.40796974 | 2.54756612 | 0 | 0 | 0 | 0 | 0 |
| miR-451a | 25.0610784 | 20.4980312 | 36.7172766 | 44.5824071 | 13.0840534 | 35.7769726 | 32.6098224 | 18.3132389 | 22.0639338 |
| miR-7241-5p | 2.78456427 | 0 | 0.81593948 | 0.63689153 | 1.63550667 | 0 | 0 | 0 | 0 |
| miR-148b-5p | 0 | 0 | 0.81593948 | 2.54756612 | 0 | 0 | 0 | 0 | 0 |
| miR-7653-3p | 2.78456427 | 0 | 0 | 0 | 0 | 0 | 0 | 0 | 0 |
| miR-674-5p | 1.39228213 | 1.36653541 | 0.40796974 | 1.91067459 | 3.27101334 | 3.25245206 | 0 | 0 | 1.2257741 |
| miR-132-3p | 1.39228213 | 1.36653541 | 3.67172766 | 3.18445765 | 3.81618223 | 0 | 0 | 3.05220649 | 0 |
| miR-31-5p | 76.5755174 | 117.522046 | 129.734377 | 91.0754887 | 130.295365 | 96.489411 | 72.3087367 | 53.4136135 | 109.093895 |
| miR-7657-3p | 0 | 0 | 0 | 0 | 0 | 0 | 0 | 0 | 1.2257741 |
| miR-543-3p | 0 | 1.36653541 | 0 | 0 | 0.54516889 | 0 | 2.83563673 | 0 | 1.2257741 |
| miR-132-5p | 2.78456427 | 2.73307083 | 1.22390922 | 7.64269835 | 4.90652001 | 0 | 11.3425469 | 3.05220649 | 2.4515482 |
| miR-23b-5p | 1.39228213 | 1.36653541 | 3.26375792 | 0.63689153 | 2.72584445 | 1.08415069 | 0 | 0 | 2.4515482 |
| miR-342-3p | 61.2604139 | 24.5976375 | 32.2296095 | 53.4988885 | 44.703849 | 32.5245206 | 26.938549 | 51.8875103 | 38.6118842 |
| miR-511-5p | 0 | 0 | 0.40796974 | 1.27378306 | 0 | 1.08415069 | 0 | 0 | 0 |

| | | | | | | | | | |
|---|---|---|---|---|---|---|---|---|---|
| miR-190b-5p | 2.78456427 | 15.0318896 | 6.52751585 | 10.1902645 | 7.63236446 | 23.8513151 | 17.0138204 | 6.10441297 | 11.0319669 |
| miR-30b-3p | 1.39228213 | 2.73307083 | 1.63187896 | 4.45824071 | 2.18067556 | 4.33660274 | 1.41781837 | 0 | 4.29020936 |
| miR-196b-3p | 4.1768464 | 2.73307083 | 5.71157636 | 2.54756612 | 1.63550667 | 1.08415069 | 2.83563673 | 0 | 5.51598346 |
| miR-1948-5p | 0 | 0 | 0 | 0 | 0 | 0 | 0 | 0 | 0.61288705 |
| miR-10b-5p | 5493.9453 | 7152.44636 | 3966.28182 | 4362.07008 | 5779.88057 | 11268.6622 | 9617.06199 | 7511.48016 | 5399.53492 |
| let-7e-3p | 0 | 0 | 0.40796974 | 0 | 0 | 0 | 0 | 0 | 0.61288705 |
| miR-193a-5p | 0 | 0 | 0 | 0 | 0.54516889 | 0 | 0 | 0 | 0 |
| miR-345-3p | 4.1768464 | 8.19921248 | 4.48766714 | 7.64269835 | 0 | 0 | 1.41781837 | 0 | 2.4515482 |
| miR-1843b-5p | 2.78456427 | 2.73307083 | 6.52751585 | 11.4640475 | 5.4516889 | 6.50490411 | 8.5069102 | 3.05220649 | 7.35464461 |
| miR-99b-5p | 221.372859 | 110.689369 | 95.0569495 | 132.473438 | 133.566378 | 177.800712 | 172.973841 | 74.7790589 | 123.803184 |
| miR-26a-2-3p | 1.39228213 | 2.73307083 | 2.85578818 | 1.91067459 | 0.54516889 | 0 | 0 | 0 | 1.2257741 |
| miR-16-1-3p | 1.39228213 | 1.36653541 | 2.0398487 | 0.63689153 | 0 | 1.08415069 | 2.83563673 | 0 | 0 |
| miR-214-3p | 5.56912854 | 6.83267707 | 2.44781844 | 3.82134918 | 3.27101334 | 0 | 0 | 3.05220649 | 4.29020936 |
| miR-6962-3p | 0 | 0 | 0.40796974 | 0 | 0 | 0 | 0 | 0 | 1.2257741 |
| miR-5116 | 0 | 0 | 0.40796974 | 0 | 0 | 0 | 0 | 0 | 0.61288705 |
| miR-30d-5p | 2205.3749 | 1365.16888 | 1424.22236 | 1632.98988 | 1487.22073 | 2248.52852 | 2047.32972 | 2116.7052 | 1412.09177 |
| miR-142a-5p | 126.697674 | 86.0917311 | 128.510468 | 316.53509 | 89.9528668 | 179.969014 | 121.93238 | 103.775021 | 63.1273663 |
| miR-30c-5p | 1038.64247 | 848.618492 | 1229.21283 | 1166.14839 | 743.065197 | 1652.24564 | 1386.62636 | 1872.52868 | 1149.16322 |
| miR-465d-5p | 0 | 0 | 0.40796974 | 0 | 0.54516889 | 3.25245206 | 4.2534551 | 3.05220649 | 0.61288705 |
| miR-1249-3p | 4.1768464 | 1.36653541 | 4.48766714 | 4.45824071 | 7.08719557 | 14.0939589 | 1.41781837 | 1.52610324 | 1.83866115 |
| miR-872-5p | 48.7298747 | 86.0917311 | 138.301742 | 128.652089 | 98.675569 | 62.8807397 | 83.6512837 | 82.4095751 | 53.3211734 |
| let-7i-5p | 277.064145 | 390.829128 | 352.485856 | 397.420314 | 387.069912 | 563.758356 | 459.373151 | 405.943463 | 391.634826 |
| miR-127-5p | 0 | 2.73307083 | 1.63187896 | 0.63689153 | 0 | 0 | 1.41781837 | 3.05220649 | 0 |
| miR-378a-3p | 9.74597494 | 25.9641729 | 11.8311225 | 21.654312 | 18.5357423 | 41.197726 | 34.0276408 | 24.4176519 | 21.4510468 |
| miR-450a-5p | 0 | 4.09960624 | 0.81593948 | 2.54756612 | 0 | 0 | 1.41781837 | 3.05220649 | 1.2257741 |
| miR-669o-5p | 5.56912854 | 5.46614166 | 6.11954611 | 2.54756612 | 10.3582089 | 8.67320548 | 2.83563673 | 3.05220649 | 8.58041871 |
| miR-878-5p | 27.8456427 | 36.8964562 | 64.459219 | 43.9455155 | 42.5231734 | 57.4599863 | 41.1167327 | 42.7308908 | 44.7407547 |
| miR-125a-3p | 0 | 0 | 2.44781844 | 1.27378306 | 1.09033778 | 0 | 0 | 3.05220649 | 1.2257741 |
| miR-25-3p | 700.317914 | 612.207865 | 515.265782 | 523.524837 | 450.854672 | 624.470795 | 499.072065 | 576.867026 | 541.792153 |
| miR-431-5p | 1.39228213 | 0 | 0 | 0.63689153 | 0.54516889 | 0 | 0 | 0 | 0.61288705 |

| | | | | | | | | | |
|---|---|---|---|---|---|---|---|---|---|
| miR-345-5p | 5.56912854 | 2.73307083 | 4.0796974 | 4.45824071 | 8.17753335 | 2.16830137 | 0 | 1.52610324 | 6.12887051 |
| miR-339-3p | 4.1768464 | 2.73307083 | 1.22390922 | 1.27378306 | 2.72584445 | 1.08415069 | 4.2534551 | 0 | 0.61288705 |
| miR-92b-3p | 1.39228213 | 0 | 1.22390922 | 3.18445765 | 2.18067556 | 1.08415069 | 1.41781837 | 0 | 1.2257741 |
| miR-378c | 0 | 0 | 0.40796974 | 0 | 0.54516889 | 0 | 0 | 0 | 0 |
| miR-218-5p | 8.35369281 | 13.6653541 | 15.0948804 | 15.9222882 | 9.26787113 | 20.598863 | 8.5069102 | 1.52610324 | 12.8706281 |
| miR-7217-5p | 18.0996677 | 20.4980312 | 21.2144265 | 14.0116136 | 21.8067556 | 14.0939589 | 24.1029122 | 13.7349292 | 16.5479504 |
| miR-1955-5p | 0 | 0 | 0.40796974 | 0.63689153 | 0 | 4.33660274 | 0 | 0 | 0 |
| miR-25-5p | 0 | 1.36653541 | 0.81593948 | 0 | 0.54516889 | 0 | 4.2534551 | 0 | 0 |
| miR-15a-5p | 22.2765142 | 16.398425 | 24.0702147 | 19.1067459 | 23.9874312 | 7.5890548 | 8.5069102 | 15.2610324 | 17.7737245 |
| miR-1930-5p | 0 | 0 | 0 | 0 | 0 | 0 | 0 | 0 | 0 |
| miR-221-5p | 4.1768464 | 8.19921248 | 3.67172766 | 3.18445765 | 10.9033778 | 8.67320548 | 4.2534551 | 6.10441297 | 7.35464461 |
| miR-191-3p | 9.74597494 | 16.398425 | 10.6072132 | 14.0116136 | 15.2647289 | 13.0098082 | 28.3563673 | 12.2088259 | 10.4190799 |
| miR-467a-3p | 20.884232 | 8.19921248 | 7.75142507 | 14.6485052 | 15.8098978 | 3.25245206 | 0 | 24.4176519 | 9.80619282 |
| miR-1981-5p | 2.78456427 | 1.36653541 | 3.67172766 | 2.54756612 | 4.90652001 | 7.5890548 | 5.67127347 | 3.05220649 | 1.83866115 |
| miR-7073-3p | 1.39228213 | 0 | 0.40796974 | 0 | 0 | 0 | 0 | 0 | 1.2257741 |
| miR-16-5p | 1019.15052 | 923.77794 | 1116.20521 | 1143.85719 | 1106.69285 | 807.69226 | 966.952127 | 990.441005 | 826.784632 |
| miR-186-3p | 0 | 0 | 0.40796974 | 0 | 0.54516889 | 0 | 0 | 0 | 0.61288705 |
| miR-7648-3p | 0 | 0 | 0 | 0 | 0 | 0 | 0 | 0 | 0 |
| miR-134-5p | 1.39228213 | 0 | 0.40796974 | 0 | 0 | 0 | 0 | 0 | 0 |
| miR-7658-5p | 8.35369281 | 0 | 0 | 0 | 0 | 0 | 2.83563673 | 0 | 1.2257741 |
| let-7f-5p | 2043.87017 | 3716.97633 | 2315.63625 | 1490.96307 | 3294.4556 | 5610.4798 | 5244.51014 | 4007.54712 | 3404.58757 |
| miR-7021-3p | 1.39228213 | 0 | 0 | 0 | 0 | 0 | 0 | 0 | 1.83866115 |
| miR-10b-3p | 18.0996677 | 24.5976375 | 23.2542752 | 20.3805289 | 32.1649645 | 27.1037671 | 9.92472857 | 25.9437551 | 23.902595 |
| miR-29b-3p | 20.884232 | 12.2988187 | 37.1252464 | 47.7668647 | 34.8908089 | 21.6830137 | 14.1781837 | 7.63051622 | 32.4830137 |
| miR-221-3p | 77.9677995 | 151.685431 | 86.0816152 | 157.949099 | 163.550667 | 85.6479041 | 70.8909184 | 88.5139881 | 200.414066 |
| miR-678 | 0 | 0 | 0 | 0 | 0 | 0 | 0 | 0 | 0 |
| miR-335-5p | 6.96141067 | 10.9322833 | 9.79127377 | 14.0116136 | 7.63236446 | 3.25245206 | 8.5069102 | 15.2610324 | 12.257741 |
| miR-669f-3p | 2.78456427 | 0 | 0 | 1.27378306 | 0.54516889 | 0 | 0 | 0 | 0 |
| miR-10a-5p | 6989.25631 | 10221.6849 | 5527.98998 | 6067.02871 | 8276.75408 | 17123.0759 | 14738.2219 | 10177.5825 | 7694.79693 |
| miR-7236-3p | 0 | 0 | 0.40796974 | 0 | 0 | 0 | 0 | 0 | 1.83866115 |

| | | | | | | | | | |
|---|---|---|---|---|---|---|---|---|---|
| miR-429-3p | 169.85842 | 311.570074 | 359.013372 | 510.787007 | 322.194814 | 356.685575 | 395.571324 | 276.224687 | 316.862605 |
| miR-151-5p | 263.141323 | 256.908658 | 241.926056 | 192.341242 | 282.942654 | 212.493534 | 282.145855 | 322.007784 | 240.864611 |
| miR-1981-3p | 0 | 0 | 1.63187896 | 0.63689153 | 2.18067556 | 2.16830137 | 1.41781837 | 3.05220649 | 1.83866115 |
| miR-201-5p | 0 | 0 | 0.40796974 | 0 | 1.09033778 | 4.33660274 | 0 | 0 | 0 |
| miR-701-5p | 1.39228213 | 1.36653541 | 0.81593948 | 1.91067459 | 1.63550667 | 1.08415069 | 1.41781837 | 3.05220649 | 1.83866115 |
| let-7b-3p | 8.35369281 | 1.36653541 | 1.63187896 | 1.27378306 | 2.18067556 | 0 | 2.83563673 | 1.52610324 | 3.67732231 |
| miR-141-5p | 2.78456427 | 6.83267707 | 2.44781844 | 0.63689153 | 1.09033778 | 0 | 1.41781837 | 0 | 6.12887051 |
| miR-211-5p | 1.39228213 | 0 | 0 | 0.63689153 | 1.63550667 | 1.08415069 | 0 | 0 | 0 |
| miR-146b-5p | 41.768464 | 30.0637791 | 22.4383357 | 26.7494442 | 32.1649645 | 43.3660274 | 39.6989143 | 35.1003746 | 26.3541432 |
| miR-3068-5p | 0 | 0 | 2.0398487 | 5.09513224 | 3.27101334 | 0 | 2.83563673 | 0 | 3.06443526 |
| miR-379-3p | 0 | 1.36653541 | 0 | 1.27378306 | 0 | 2.16830137 | 0 | 1.52610324 | 0 |
| let-7a-5p | 2110.69972 | 3208.62515 | 1446.6607 | 1155.95813 | 3001.15474 | 4471.03743 | 4494.48422 | 3566.50328 | 2697.9288 |
| miR-146a-5p | 161.504728 | 109.322833 | 98.3207074 | 189.793676 | 123.208169 | 202.736178 | 136.110563 | 152.610324 | 96.223267 |
| miR-32-3p | 2.78456427 | 5.46614166 | 2.0398487 | 4.45824071 | 4.90652001 | 2.16830137 | 1.41781837 | 0 | 3.06443526 |
| miR-881-5p | 32.0224891 | 51.9283457 | 35.0853977 | 29.9339019 | 45.7941867 | 19.5147123 | 8.5069102 | 21.3654454 | 47.80519 |
| miR-20a-5p | 51.514439 | 50.5618103 | 35.4933674 | 36.9397087 | 32.7101334 | 27.1037671 | 36.8632776 | 51.8875103 | 61.9015922 |
| miR-3058-5p | 0 | 0 | 0.40796974 | 0 | 0 | 0 | 0 | 0 | 0 |
| miR-204-5p | 12.5305392 | 6.83267707 | 22.030366 | 23.5649866 | 14.71956 | 7.5890548 | 11.3425469 | 15.2610324 | 7.35464461 |
| miR-107-3p | 2.78456427 | 1.36653541 | 2.44781844 | 1.91067459 | 2.72584445 | 5.42075343 | 4.2534551 | 3.05220649 | 4.29020936 |
| miR-3475-3p | 2.78456427 | 5.46614166 | 0.81593948 | 1.27378306 | 2.18067556 | 2.16830137 | 1.41781837 | 0 | 0.61288705 |
| miR-686 | 0 | 0 | 0 | 0 | 0 | 0 | 0 | 0 | 1.2257741 |
| miR-30a-5p | 1566.3174 | 1855.75509 | 1935.40845 | 2153.96715 | 1614.79025 | 2901.18723 | 2322.38649 | 1927.4684 | 1694.01981 |
| miR-295-5p | 0 | 0 | 0 | 0 | 0 | 0 | 0 | 0 | 0.61288705 |
| miR-195a-5p | 23.6687963 | 25.9641729 | 35.0853977 | 39.4872748 | 32.1649645 | 14.0939589 | 19.8494571 | 16.7871357 | 20.2252727 |
| miR-17-5p | 6.96141067 | 1.36653541 | 6.52751585 | 3.18445765 | 3.27101334 | 4.33660274 | 0 | 3.05220649 | 6.74175756 |
| miR-467b-5p | 1.39228213 | 1.36653541 | 3.26375792 | 6.36891529 | 2.72584445 | 1.08415069 | 1.41781837 | 1.52610324 | 1.83866115 |
| miR-194-2-3p | 0 | 0 | 0 | 0 | 0.54516889 | 0 | 2.83563673 | 0 | 0 |
| miR-215-3p | 0 | 0 | 0.40796974 | 1.27378306 | 0 | 0 | 0 | 3.05220649 | 0 |
| miR-125b-1-3p | 8.35369281 | 4.09960624 | 0.40796974 | 0 | 2.18067556 | 0 | 0 | 1.52610324 | 1.2257741 |
| miR-181d-5p | 13.9228213 | 5.46614166 | 6.52751585 | 8.91648141 | 9.26787113 | 3.25245206 | 7.08909184 | 4.57830973 | 8.58041871 |

| | | | | | | | | | |
|---|---|---|---|---|---|---|---|---|---|
| miR-28a-5p | 34.8070534 | 28.6972437 | 26.5180331 | 21.654312 | 37.6166534 | 31.4403699 | 56.7127347 | 28.9959616 | 30.6443526 |
| miR-130b-3p | 1.39228213 | 2.73307083 | 2.0398487 | 1.27378306 | 1.63550667 | 2.16830137 | 2.83563673 | 0 | 1.83866115 |
| miR-652-3p | 18.0996677 | 19.1314958 | 13.4630014 | 25.4756612 | 21.2615867 | 16.2622603 | 22.6850939 | 18.3132389 | 15.3221763 |
| miR-672-5p | 33.4147712 | 40.9960624 | 17.9506686 | 18.4698544 | 29.43912 | 49.8709315 | 41.1167327 | 41.2047876 | 33.0959008 |
| miR-361-5p | 13.9228213 | 9.5657479 | 8.97533429 | 15.9222882 | 12.5388845 | 15.1781096 | 24.1029122 | 9.15661946 | 26.3541432 |
| miR-467d-5p | 13.9228213 | 2.73307083 | 20.398487 | 24.2018781 | 10.9033778 | 6.50490411 | 7.08909184 | 10.6827227 | 7.35464461 |
| miR-3110-5p | 1.39228213 | 0 | 0 | 0 | 0 | 0 | 0 | 0 | 1.2257741 |
| miR-300-3p | 1.39228213 | 0 | 0.81593948 | 0.63689153 | 0 | 0 | 0 | 0 | 0 |
| miR-135a-5p | 1.39228213 | 12.2988187 | 9.79127377 | 15.2853967 | 17.9905734 | 6.50490411 | 7.08909184 | 9.15661946 | 15.3221763 |
| miR-376b-5p | 0 | 0 | 0.40796974 | 0.63689153 | 1.63550667 | 0 | 0 | 0 | 0 |
| miR-582-5p | 0 | 0 | 0 | 0 | 0 | 2.16830137 | 2.83563673 | 0 | 0 |
| miR-669m-3p | 0 | 0 | 0 | 0.63689153 | 0 | 0 | 4.2534551 | 0 | 0 |
| miR-362-5p | 1.39228213 | 0 | 1.63187896 | 1.27378306 | 1.63550667 | 3.25245206 | 1.41781837 | 4.57830973 | 4.29020936 |
| miR-467e-5p | 40.3761819 | 61.4940936 | 85.2656757 | 78.3376581 | 85.5915157 | 45.5343288 | 53.877098 | 50.361407 | 60.062931 |
| miR-467a-5p | 16.7073856 | 9.5657479 | 30.5977305 | 63.6891529 | 27.8036134 | 6.50490411 | 9.92472857 | 21.3654454 | 15.9350633 |
| miR-100-5p | 98.8520315 | 68.3267707 | 44.8766714 | 49.6775393 | 55.6072268 | 114.919973 | 104.918559 | 70.2007492 | 77.2237684 |
| miR-135a-2-3p | 0 | 2.73307083 | 0.40796974 | 0 | 0 | 0 | 0 | 0 | 0 |
| miR-339-5p | 4.1768464 | 0 | 5.30360662 | 7.00580682 | 3.27101334 | 8.67320548 | 4.2534551 | 1.52610324 | 5.51598346 |
| miR-742-3p | 0 | 1.36653541 | 2.0398487 | 1.27378306 | 0 | 0 | 0 | 0 | 1.2257741 |
| miR-7663-5p | 0 | 0 | 0 | 0 | 0 | 0 | 0 | 0 | 1.2257741 |
| miR-152-5p | 0 | 0 | 0.81593948 | 0 | 0 | 0 | 0 | 0 | 0 |
| miR-342-5p | 4.1768464 | 1.36653541 | 0.40796974 | 0 | 0.54516889 | 3.25245206 | 0 | 0 | 1.2257741 |
| miR-125b-2-3p | 2.78456427 | 4.09960624 | 6.93548559 | 7.64269835 | 3.81618223 | 3.25245206 | 2.83563673 | 10.6827227 | 4.90309641 |
| miR-323-3p | 1.39228213 | 1.36653541 | 0.40796974 | 0 | 0.54516889 | 0 | 4.2534551 | 0 | 1.83866115 |
| miR-187-5p | 2.78456427 | 0 | 1.63187896 | 0.63689153 | 1.09033778 | 0 | 0 | 0 | 0.61288705 |
| miR-1938 | 0 | 0 | 0.40796974 | 0.63689153 | 0 | 1.08415069 | 0 | 0 | 0 |
| miR-1191b-3p | 0 | 0 | 0 | 0 | 0 | 0 | 0 | 0 | 0 |
| miR-200a-3p | 66.8295425 | 150.318896 | 236.622449 | 236.923649 | 121.572662 | 177.800712 | 161.631294 | 134.297085 | 221.252225 |
| miR-669a-3-3p | 0 | 0 | 0 | 1.27378306 | 0 | 0 | 0 | 3.05220649 | 0 |
| miR-369-3p | 8.35369281 | 17.7649604 | 12.2390922 | 9.55337294 | 8.17753335 | 14.0939589 | 5.67127347 | 10.6827227 | 10.4190799 |

| | | | | | | | | | |
|---|---|---|---|---|---|---|---|---|---|
| miR-144-5p | 5.56912854 | 0 | 4.0796974 | 3.18445765 | 0 | 8.67320548 | 5.67127347 | 4.57830973 | 2.4515482 |
| miR-700-3p | 0 | 0 | 0.81593948 | 2.54756612 | 1.09033778 | 1.08415069 | 0 | 1.52610324 | 1.83866115 |
| miR-485-5p | 1.39228213 | 0 | 1.63187896 | 1.91067459 | 1.63550667 | 2.16830137 | 0 | 0 | 0 |
| miR-329-5p | 2.78456427 | 0 | 0 | 1.27378306 | 0 | 0 | 0 | 0 | 0 |
| miR-668-5p | 0 | 0 | 0.40796974 | 0 | 0.54516889 | 0 | 0 | 0 | 0.61288705 |
| miR-653-5p | 0 | 0 | 0 | 1.27378306 | 0.54516889 | 2.16830137 | 0 | 0 | 0.61288705 |
| miR-135b-5p | 0 | 0 | 0.81593948 | 0.63689153 | 1.09033778 | 0 | 0 | 0 | 0.61288705 |
| miR-26b-3p | 11.1382571 | 0 | 3.26375792 | 1.27378306 | 1.09033778 | 0 | 0 | 1.52610324 | 4.29020936 |
| miR-21a-5p | 4084.95578 | 5650.62394 | 5611.21581 | 5039.08578 | 4953.9497 | 5482.55002 | 5915.13823 | 4686.66306 | 5056.31817 |
| miR-3057-5p | 0 | 0 | 0.40796974 | 0 | 0 | 0 | 0 | 0 | 0 |
| miR-495-3p | 0 | 1.36653541 | 0.81593948 | 0 | 0.54516889 | 3.25245206 | 0 | 0 | 1.83866115 |
| miR-8101 | 0 | 0 | 0 | 0 | 0 | 0 | 0 | 0 | 0.61288705 |
| miR-99a-5p | 728.163556 | 888.248019 | 849.392999 | 1055.32926 | 849.37313 | 1049.45786 | 1122.91215 | 674.537633 | 801.656263 |
| miR-185-3p | 0 | 0 | 0 | 0 | 0 | 0 | 0 | 0 | 0 |
| miR-182-3p | 0 | 2.73307083 | 2.44781844 | 0 | 0.54516889 | 0 | 0 | 0 | 1.83866115 |
| miR-708-3p | 0 | 1.36653541 | 1.22390922 | 0 | 1.09033778 | 0 | 0 | 0 | 0 |
| miR-6368 | 0 | 0 | 0 | 0 | 0 | 0 | 0 | 0 | 0 |
| miR-17-3p | 2.78456427 | 2.73307083 | 3.67172766 | 2.54756612 | 2.18067556 | 0 | 4.2534551 | 7.63051622 | 3.06443526 |
| miR-328-3p | 15.3151035 | 17.7649604 | 10.1992435 | 19.1067459 | 13.0840534 | 11.9256575 | 36.8632776 | 16.7871357 | 10.4190799 |
| miR-6993-3p | 0 | 0 | 0.40796974 | 0 | 0 | 0 | 5.67127347 | 0 | 0 |
| miR-101b-5p | 0 | 0 | 0 | 0 | 0.54516889 | 0 | 0 | 0 | 0 |
| miR-181a-1-3p | 0 | 0 | 0.81593948 | 2.54756612 | 0 | 3.25245206 | 1.41781837 | 0 | 0.61288705 |
| miR-669h-5p | 0 | 0 | 0 | 0 | 0 | 0 | 0 | 0 | 0 |
| miR-7233-3p | 0 | 6.83267707 | 2.44781844 | 3.18445765 | 5.99685779 | 1.08415069 | 0 | 1.52610324 | 4.29020936 |
| miR-411-5p | 4.1768464 | 6.83267707 | 12.647062 | 9.55337294 | 10.3582089 | 24.9354658 | 12.7603653 | 10.6827227 | 11.644854 |
| miR-7079-5p | 0 | 0 | 0 | 0 | 0 | 0 | 0 | 0 | 0 |
| miR-7242-3p | 0 | 0 | 1.22390922 | 0 | 1.63550667 | 0 | 2.83563673 | 0 | 0.61288705 |
| miR-203-3p | 33.4147712 | 68.3267707 | 51.4041873 | 40.1241664 | 70.8719557 | 49.8709315 | 60.9661898 | 47.3092005 | 52.0953993 |
| miR-322-3p | 1.39228213 | 4.09960624 | 1.22390922 | 4.45824071 | 1.63550667 | 2.16830137 | 0 | 0 | 3.06443526 |
| miR-463-3p | 4.1768464 | 10.9322833 | 7.75142507 | 6.36891529 | 6.54202668 | 0 | 5.67127347 | 3.05220649 | 1.83866115 |

| | | | | | | | | | |
|---|---|---|---|---|---|---|---|---|---|
| miR-504-5p | 0 | 0 | 0 | 1.27378306 | 0.54516889 | 2.16830137 | 0 | 1.52610324 | 0 |
| miR-215-5p | 0 | 1.36653541 | 0.81593948 | 0 | 0 | 0 | 0 | 1.52610324 | 0 |
| miR-7036a-5p | 0 | 0 | 0.40796974 | 0 | 0 | 0 | 0 | 0 | 0 |
| miR-702-3p | 0 | 0 | 0.40796974 | 0 | 0 | 0 | 0 | 0 | 0 |
| miR-7222-5p | 0 | 1.36653541 | 0 | 0.63689153 | 1.09033778 | 0 | 0 | 0 | 0 |
| miR-2137 | 0 | 0 | 0.81593948 | 0.63689153 | 0.54516889 | 0 | 0 | 0 | 0.61288705 |
| miR-125a-5p | 473.375926 | 352.566137 | 259.468755 | 391.051399 | 376.166534 | 474.858 | 621.004445 | 552.449374 | 498.89006 |
| miR-7221-5p | 0 | 0 | 0 | 0.63689153 | 1.09033778 | 0 | 0 | 0 | 1.2257741 |
| miR-429-5p | 0 | 0 | 1.22390922 | 0 | 1.09033778 | 0 | 0 | 0 | 0.61288705 |
| miR-7223-3p | 0 | 1.36653541 | 0 | 1.27378306 | 0.54516889 | 0 | 0 | 0 | 0.61288705 |
| miR-1894-5p | 0 | 0 | 0.40796974 | 0 | 0 | 0 | 0 | 0 | 1.2257741 |
| miR-150-3p | 1.39228213 | 0 | 0 | 1.91067459 | 0.54516889 | 0 | 2.83563673 | 4.57830973 | 0 |
| miR-7026-3p | 0 | 0 | 0 | 0 | 0 | 0 | 0 | 0 | 0.61288705 |
| miR-30b-5p | 128.089956 | 129.820864 | 301.897608 | 298.702127 | 159.189316 | 140.939589 | 138.9462 | 198.393422 | 174.67281 |
| miR-210-5p | 0 | 0 | 0.40796974 | 0 | 0 | 2.16830137 | 0 | 0 | 0 |
| miR-128-1-5p | 2.78456427 | 0 | 0 | 0.63689153 | 0 | 0 | 2.83563673 | 1.52610324 | 0 |
| miR-7214-3p | 4.1768464 | 5.46614166 | 2.0398487 | 3.82134918 | 3.81618223 | 5.42075343 | 1.41781837 | 9.15661946 | 2.4515482 |
| miR-7648-5p | 1.39228213 | 0 | 0 | 0 | 0 | 0 | 0 | 0 | 0 |
| miR-369-5p | 0 | 0 | 1.63187896 | 2.54756612 | 3.81618223 | 5.42075343 | 0 | 0 | 2.4515482 |
| miR-676-3p | 34.8070534 | 31.4303145 | 11.8311225 | 17.8329628 | 17.9905734 | 33.6086712 | 24.1029122 | 19.8393422 | 22.0639338 |
| miR-6928-5p | 0 | 0 | 0 | 0 | 0 | 0 | 0 | 0 | 0.61288705 |
| miR-203-5p | 2.78456427 | 2.73307083 | 0.40796974 | 0.63689153 | 0 | 3.25245206 | 0 | 0 | 0 |
| let-7a-1-3p | 0 | 6.83267707 | 0.81593948 | 2.54756612 | 1.09033778 | 1.08415069 | 0 | 0 | 2.4515482 |
| miR-6539 | 0 | 0 | 0 | 0 | 0 | 0 | 0 | 0 | 0.61288705 |
| miR-1198-3p | 0 | 0 | 0 | 0 | 0 | 0 | 0 | 0 | 0 |
| miR-378b | 0 | 0 | 0 | 0 | 0 | 0 | 0 | 0 | 0 |
| miR-5119 | 0 | 0 | 0 | 0 | 0 | 0 | 0 | 0 | 0 |
| miR-217-5p | 0 | 0 | 0.81593948 | 0 | 0.54516889 | 0 | 0 | 0 | 0 |
| miR-140-5p | 12.5305392 | 4.09960624 | 12.2390922 | 8.91648141 | 7.08719557 | 5.42075343 | 4.2534551 | 4.57830973 | 4.90309641 |
| miR-669l-5p | 2.78456427 | 2.73307083 | 6.52751585 | 5.09513224 | 6.54202668 | 1.08415069 | 2.83563673 | 1.52610324 | 3.06443526 |

| | | | | | | | | | |
|---|---|---|---|---|---|---|---|---|---|
| miR-8108 | 0 | 0 | 2.0398487 | 1.91067459 | 0 | 4.33660274 | 0 | 0 | 0 |
| miR-27a-3p | 44.5530283 | 65.5936999 | 95.4649192 | 151.580184 | 76.8688135 | 52.0392329 | 39.6989143 | 33.5742713 | 73.5464461 |
| miR-423-5p | 80.7523638 | 21.8645666 | 17.1347291 | 17.1960713 | 26.7132756 | 7.5890548 | 31.1920041 | 53.4136135 | 14.7092892 |
| miR-96-5p | 8.35369281 | 23.231102 | 33.045549 | 36.3028172 | 37.0714845 | 13.0098082 | 31.1920041 | 32.0481681 | 17.1608374 |
| miR-669p-5p | 2.78456427 | 2.73307083 | 2.44781844 | 3.18445765 | 3.27101334 | 2.16830137 | 1.41781837 | 1.52610324 | 2.4515482 |
| miR-7237-5p | 0 | 0 | 0.40796974 | 0 | 0 | 0 | 0 | 0 | 0 |
| miR-6905-3p | 0 | 0 | 0 | 0 | 0 | 0 | 0 | 0 | 1.83866115 |
| miR-298-5p | 2.78456427 | 0 | 0.40796974 | 0.63689153 | 1.63550667 | 1.08415069 | 1.41781837 | 0 | 1.83866115 |
| miR-148a-5p | 25.0610784 | 15.0318896 | 18.3586383 | 7.64269835 | 11.4485467 | 8.67320548 | 7.08909184 | 1.52610324 | 10.4190799 |
| miR-743a-5p | 1.39228213 | 2.73307083 | 7.75142507 | 8.91648141 | 9.26787113 | 8.67320548 | 7.08909184 | 3.05220649 | 12.8706281 |
| miR-883a-3p | 36.1993355 | 49.1952749 | 56.7077939 | 71.3318513 | 49.610369 | 23.8513151 | 51.0414612 | 35.1003746 | 42.9020936 |
| miR-376b-3p | 0 | 1.36653541 | 0.40796974 | 1.91067459 | 1.09033778 | 0 | 0 | 0 | 0.61288705 |
| miR-377-3p | 2.78456427 | 0 | 0 | 0.63689153 | 0 | 0 | 0 | 0 | 0 |
| miR-103-3p | 211.626884 | 159.884643 | 104.848223 | 84.0696819 | 155.918302 | 192.978822 | 199.91239 | 207.550041 | 209.607371 |
| miR-1193-3p | 0 | 1.36653541 | 0.40796974 | 0 | 0 | 0 | 2.83563673 | 0 | 0.61288705 |
| miR-6910-5p | 4.1768464 | 0 | 0 | 0 | 0 | 0 | 0 | 0 | 0 |
| miR-6904-5p | 0 | 0 | 0 | 0 | 0 | 0 | 0 | 0 | 0 |
| miR-27a-5p | 5.56912854 | 1.36653541 | 3.67172766 | 4.45824071 | 3.81618223 | 3.25245206 | 0 | 3.05220649 | 4.90309641 |
| miR-455-3p | 1.39228213 | 1.36653541 | 4.48766714 | 3.18445765 | 6.54202668 | 2.16830137 | 1.41781837 | 0 | 4.90309641 |
| miR-615-3p | 4.1768464 | 5.46614166 | 1.63187896 | 1.27378306 | 2.18067556 | 0 | 5.67127347 | 1.52610324 | 0 |
| miR-7036b-3p | 0 | 0 | 0 | 0 | 0 | 0 | 0 | 0 | 1.2257741 |
| miR-3101-3p | 0 | 0 | 0 | 0 | 0 | 0 | 0 | 0 | 0 |
| miR-182-5p | 729.555838 | 875.9492 | 818.387299 | 557.91698 | 889.170459 | 1854.98182 | 1500.05183 | 1053.01124 | 799.817602 |
| miR-5106 | 0 | 0 | 0.40796974 | 0 | 0 | 0 | 0 | 0 | 0 |
| miR-6236 | 0 | 0 | 0 | 0 | 0 | 0 | 0 | 0 | 1.2257741 |
| miR-667-3p | 0 | 2.73307083 | 0 | 0 | 0 | 0 | 0 | 1.52610324 | 0.61288705 |
| miR-7230-3p | 4.1768464 | 4.09960624 | 2.85578818 | 3.82134918 | 5.4516889 | 0 | 0 | 0 | 1.83866115 |
| miR-16-2-3p | 0 | 6.83267707 | 2.44781844 | 5.73202376 | 5.4516889 | 1.08415069 | 1.41781837 | 6.10441297 | 1.83866115 |
| miR-5046 | 0 | 0 | 0.81593948 | 0.63689153 | 0 | 0 | 0 | 0 | 0.61288705 |
| miR-93-5p | 430.21518 | 355.299208 | 245.597784 | 208.26353 | 278.036134 | 259.112014 | 218.344029 | 289.959616 | 351.18428 |

| | | | | | | | | | |
|---|---|---|---|---|---|---|---|---|---|
| miR-6951-5p | 0 | 0 | 0.40796974 | 0 | 0.54516889 | 0 | 1.41781837 | 0 | 0 |
| miR-582-3p | 0 | 1.36653541 | 0 | 0 | 0 | 0 | 1.41781837 | 0 | 0 |
| miR-6909-5p | 0 | 0 | 0 | 0 | 0 | 0 | 0 | 0 | 1.83866115 |
| miR-6990-5p | 0 | 0 | 0 | 0 | 0 | 0 | 0 | 0 | 1.2257741 |
| miR-324-5p | 1.39228213 | 0 | 2.85578818 | 1.91067459 | 2.72584445 | 0 | 4.2534551 | 0 | 1.2257741 |
| miR-5136 | 0 | 0 | 0 | 0 | 0 | 0 | 0 | 0 | 0 |
| miR-301b-3p | 0 | 0 | 0.40796974 | 0.63689153 | 0 | 0 | 0 | 3.05220649 | 0 |
| miR-190a-5p | 0 | 5.46614166 | 3.26375792 | 1.91067459 | 2.18067556 | 0 | 0 | 1.52610324 | 1.83866115 |
| miR-140-3p | 215.803731 | 144.852754 | 165.635715 | 184.061652 | 141.743911 | 277.542575 | 214.090573 | 241.124312 | 146.480005 |
| let-7g-5p | 1887.93457 | 2349.07438 | 1895.01944 | 1630.44232 | 2023.66692 | 3500.72256 | 3319.1128 | 2919.4355 | 2542.25549 |
| miR-129-5p | 0 | 0 | 0 | 0.63689153 | 0 | 0 | 2.83563673 | 0 | 0 |
| miR-501-5p | 0 | 0 | 0 | 0.63689153 | 1.09033778 | 0 | 0 | 3.05220649 | 0 |
| miR-27b-3p | 197.704063 | 278.773224 | 345.55037 | 331.820487 | 287.304005 | 300.30974 | 300.577494 | 230.44159 | 294.798672 |
| miR-145b | 0 | 0 | 0.40796974 | 2.54756612 | 0.54516889 | 0 | 0 | 0 | 0.61288705 |
| miR-8092 | 4.1768464 | 0 | 0 | 0 | 0 | 0 | 0 | 0 | 0 |
| miR-361-3p | 6.96141067 | 10.9322833 | 6.93548559 | 5.73202376 | 8.72270224 | 3.25245206 | 9.92472857 | 7.63051622 | 4.90309641 |
| miR-351-5p | 4.1768464 | 4.09960624 | 2.44781844 | 3.18445765 | 5.4516889 | 3.25245206 | 0 | 1.52610324 | 3.06443526 |
| miR-468-5p | 5.56912854 | 10.9322833 | 4.48766714 | 5.73202376 | 4.90652001 | 1.08415069 | 0 | 3.05220649 | 7.35464461 |
| miR-137-3p | 0 | 0 | 0 | 0 | 0 | 0 | 0 | 0 | 0 |
| miR-455-5p | 0 | 0 | 0.81593948 | 1.27378306 | 0 | 0 | 0 | 3.05220649 | 0 |
| miR-293-3p | 2.78456427 | 6.83267707 | 2.44781844 | 1.27378306 | 2.18067556 | 0 | 0 | 3.05220649 | 0.61288705 |
| miR-23b-3p | 97.4597494 | 133.920471 | 126.06265 | 119.098716 | 137.927729 | 123.593178 | 96.411649 | 103.775021 | 124.416071 |
| miR-1198-5p | 11.1382571 | 2.73307083 | 3.67172766 | 5.73202376 | 4.36135112 | 4.33660274 | 11.3425469 | 9.15661946 | 5.51598346 |
| miR-669a-5p | 13.9228213 | 16.398425 | 15.9108199 | 17.8329628 | 21.2615867 | 13.0098082 | 11.3425469 | 10.6827227 | 15.3221763 |
| miR-6418-3p | 0 | 0 | 0 | 0 | 0 | 0 | 2.83563673 | 0 | 0.61288705 |
| miR-26a-1-3p | 0 | 2.73307083 | 0 | 0 | 0 | 0 | 0 | 0 | 0.61288705 |
| miR-384-5p | 0 | 0 | 0.40796974 | 0 | 0 | 0 | 1.41781837 | 1.52610324 | 0 |
| let-7j | 1.39228213 | 0 | 0 | 0.63689153 | 0 | 2.16830137 | 0 | 1.52610324 | 1.83866115 |
| miR-138-2-3p | 0 | 1.36653541 | 0.81593948 | 0 | 0.54516889 | 0 | 0 | 0 | 0 |
| miR-421-3p | 2.78456427 | 0 | 4.0796974 | 1.91067459 | 3.81618223 | 5.42075343 | 1.41781837 | 1.52610324 | 2.4515482 |

| | | | | | | | | | |
|---|---|---|---|---|---|---|---|---|---|
| miR-760-3p | 0 | 1.36653541 | 0 | 0 | 0 | 0 | 1.41781837 | 0 | 0 |
| miR-3088-3p | 1.39228213 | 1.36653541 | 0.40796974 | 0 | 0 | 0 | 0 | 0 | 0.61288705 |
| miR-202-3p | 0 | 0 | 0 | 0.63689153 | 1.09033778 | 0 | 0 | 3.05220649 | 0 |
| miR-6911-3p | 2.78456427 | 0 | 0 | 1.27378306 | 0 | 0 | 0 | 0 | 0 |
| miR-212-5p | 0 | 2.73307083 | 0.81593948 | 4.45824071 | 2.72584445 | 0 | 2.83563673 | 6.10441297 | 0 |
| miR-29b-2-5p | 4.1768464 | 0 | 0 | 1.27378306 | 0.54516889 | 0 | 0 | 1.52610324 | 1.83866115 |
| miR-1941-3p | 0 | 0 | 1.22390922 | 0.63689153 | 0.54516889 | 0 | 0 | 0 | 0 |
| miR-96-3p | 0 | 0 | 0.81593948 | 1.27378306 | 1.09033778 | 0 | 0 | 0 | 0.61288705 |
| miR-468-3p | 2.78456427 | 1.36653541 | 1.63187896 | 1.91067459 | 2.18067556 | 0 | 8.5069102 | 3.05220649 | 1.83866115 |
| miR-154-3p | 0 | 0 | 1.22390922 | 0.63689153 | 0.54516889 | 0 | 1.41781837 | 0 | 0 |
| miR-6922-3p | 0 | 0 | 0 | 0 | 0 | 0 | 0 | 0 | 0 |
| miR-7084-3p | 0 | 0 | 0 | 0 | 0 | 0 | 0 | 0 | 0 |
| miR-1983 | 2.78456427 | 5.46614166 | 6.11954611 | 9.55337294 | 6.54202668 | 5.42075343 | 4.2534551 | 0 | 5.51598346 |
| miR-3100-5p | 0 | 1.36653541 | 0 | 0 | 0.54516889 | 0 | 0 | 0 | 0 |
| miR-322-5p | 4.1768464 | 1.36653541 | 2.0398487 | 0.63689153 | 3.27101334 | 0 | 0 | 0 | 4.29020936 |
| miR-195a-3p | 1.39228213 | 0 | 0.81593948 | 1.27378306 | 0.54516889 | 0 | 0 | 0 | 0 |
| miR-7022-3p | 0 | 0 | 0 | 0 | 0 | 0 | 0 | 0 | 0.61288705 |
| miR-1668 | 0 | 0 | 0 | 0 | 0 | 0 | 0 | 0 | 0 |
| miR-8097 | 2.78456427 | 0 | 0.40796974 | 0 | 0 | 0 | 0 | 0 | 0 |
| miR-3105-3p | 0 | 0 | 0.40796974 | 0.63689153 | 0 | 0 | 1.41781837 | 0 | 0.61288705 |
| miR-7222-3p | 1.39228213 | 0 | 0.40796974 | 0.63689153 | 0.54516889 | 0 | 0 | 0 | 0 |
| miR-301a-5p | 4.1768464 | 1.36653541 | 1.63187896 | 2.54756612 | 1.09033778 | 0 | 0 | 4.57830973 | 5.51598346 |
| miR-350-3p | 5.56912854 | 0 | 8.97533429 | 19.1067459 | 7.63236446 | 4.33660274 | 1.41781837 | 13.7349292 | 5.51598346 |
| miR-186-5p | 27.8456427 | 31.4303145 | 68.1309466 | 91.0754887 | 39.2521601 | 82.3954521 | 69.4731 | 65.6224395 | 42.9020936 |
| miR-6926-3p | 0 | 0 | 0 | 0 | 0 | 0 | 0 | 0 | 0 |
| miR-7219-3p | 0 | 0 | 0 | 0 | 0 | 0 | 0 | 0 | 0 |
| miR-20a-3p | 1.39228213 | 0 | 0 | 0 | 0 | 0 | 1.41781837 | 1.52610324 | 1.83866115 |
| miR-6928-3p | 0 | 0 | 0 | 0 | 1.09033778 | 0 | 0 | 0 | 1.2257741 |
| miR-32-5p | 2.78456427 | 6.83267707 | 10.1992435 | 13.3747221 | 8.72270224 | 1.08415069 | 0 | 0 | 3.67732231 |
| miR-133a-5p | 0 | 2.73307083 | 0.40796974 | 0 | 0 | 0 | 0 | 0 | 0 |

| | | | | | | | | | |
|---|---|---|---|---|---|---|---|---|---|
| let-7f-1-3p | 0 | 0 | 0 | 1.27378306 | 0 | 0 | 0 | 0 | 1.83866115 |
| miR-547-3p | 0 | 0 | 0.40796974 | 1.91067459 | 0 | 0 | 0 | 1.52610324 | 1.2257741 |
| miR-6955-5p | 0 | 0 | 0 | 0 | 0 | 0 | 0 | 0 | 0 |
| miR-7068-5p | 0 | 0 | 0 | 0 | 0 | 0 | 0 | 0 | 0 |
| miR-505-3p | 0 | 0 | 0.81593948 | 2.54756612 | 1.09033778 | 0 | 0 | 0 | 1.2257741 |
| miR-615-5p | 0 | 0 | 0.40796974 | 2.54756612 | 1.09033778 | 0 | 0 | 0 | 1.83866115 |
| miR-6942-3p | 0 | 0 | 0 | 0 | 0 | 0 | 0 | 0 | 0 |
| miR-671-3p | 6.96141067 | 6.83267707 | 0.81593948 | 7.64269835 | 5.99685779 | 5.42075343 | 8.5069102 | 0 | 7.35464461 |
| miR-222-5p | 1.39228213 | 0 | 3.67172766 | 2.54756612 | 0.54516889 | 3.25245206 | 4.2534551 | 1.52610324 | 0 |
| miR-423-3p | 83.5369281 | 19.1314958 | 52.2201268 | 49.6775393 | 49.610369 | 39.0294247 | 45.3701878 | 38.1525811 | 24.515482 |
| miR-1896 | 0 | 0 | 0 | 0 | 0 | 0 | 0 | 0 | 0 |
| miR-5124a | 0 | 0 | 0 | 0 | 0 | 0 | 0 | 0 | 0 |
| miR-7667-3p | 0 | 0 | 0 | 0 | 0 | 0 | 0 | 0 | 0 |
| miR-6371 | 0 | 0 | 0 | 0 | 0 | 0 | 0 | 0 | 0 |
| miR-330-3p | 0 | 0 | 1.63187896 | 0.63689153 | 0 | 0 | 1.41781837 | 0 | 0.61288705 |
| miR-6992-5p | 0 | 0 | 0 | 0 | 0 | 0 | 0 | 0 | 0.61288705 |
| miR-181b-2-3p | 0 | 0 | 0 | 0 | 0 | 0 | 0 | 0 | 0.61288705 |
| miR-5625-3p | 0 | 0 | 0 | 0 | 0 | 0 | 0 | 0 | 0.61288705 |
| miR-7650-5p | 0 | 0 | 0 | 0 | 0 | 0 | 0 | 0 | 0.61288705 |
| miR-150-5p | 320.224891 | 76.5259832 | 157.06835 | 513.334573 | 105.217596 | 208.156932 | 163.049112 | 228.915486 | 67.4175756 |
| miR-6405 | 0 | 0 | 0 | 0 | 0 | 0 | 0 | 0 | 1.2257741 |
| miR-1982-3p | 0 | 0 | 0 | 0 | 0 | 0 | 0 | 0 | 0 |
| miR-6934-3p | 0 | 0 | 0 | 0 | 0 | 0 | 0 | 0 | 0 |
| miR-7008-3p | 0 | 0 | 0 | 0 | 0 | 0 | 0 | 0 | 0 |
| miR-7227-3p | 0 | 0 | 0 | 0 | 0 | 0 | 0 | 0 | 0 |
| miR-328-5p | 0 | 0 | 0 | 0 | 0 | 0 | 0 | 0 | 0 |
| miR-532-3p | 0 | 0 | 1.22390922 | 1.27378306 | 0.54516889 | 2.16830137 | 4.2534551 | 1.52610324 | 1.2257741 |
| miR-7234-3p | 8.35369281 | 15.0318896 | 3.26375792 | 2.54756612 | 10.3582089 | 5.42075343 | 2.83563673 | 4.57830973 | 14.0964022 |
| miR-883a-5p | 0 | 2.73307083 | 3.26375792 | 1.91067459 | 2.18067556 | 0 | 0 | 4.57830973 | 4.90309641 |
| miR-3618-5p | 0 | 1.36653541 | 0 | 0 | 0 | 0 | 0 | 0 | 1.83866115 |

| | | | | | | | | | |
|---|---|---|---|---|---|---|---|---|---|
| miR-122-5p | 0 | 0 | 0.81593948 | 0.63689153 | 0 | 2.16830137 | 0 | 0 | 0 |
| miR-3968 | 0 | 0 | 0 | 0 | 0 | 0 | 0 | 0 | 0 |
| miR-673-5p | 0 | 0 | 0 | 0 | 0 | 0 | 0 | 0 | 0.61288705 |
| miR-679-5p | 0 | 0 | 0 | 0 | 0 | 0 | 0 | 0 | 0 |
| miR-496a-3p | 0 | 0 | 0 | 0 | 0 | 0 | 0 | 0 | 0 |
| miR-6992-3p | 0 | 0 | 0 | 0 | 0 | 0 | 0 | 0 | 0.61288705 |
| miR-7013-3p | 0 | 0 | 0 | 0 | 0 | 0 | 0 | 0 | 0 |
| let-7a-2-3p | 0 | 0 | 0 | 0 | 0 | 0 | 0 | 0 | 0 |
| miR-7646-3p | 0 | 0 | 0 | 0 | 0 | 0 | 0 | 0 | 0 |
| let-7g-3p | 1.39228213 | 0 | 0 | 1.27378306 | 0.54516889 | 0 | 0 | 0 | 1.2257741 |
| miR-29a-5p | 0 | 0 | 0.40796974 | 0.63689153 | 1.09033778 | 0 | 0 | 3.05220649 | 1.2257741 |
| miR-344-3p | 0 | 0 | 0.40796974 | 0 | 0.54516889 | 0 | 0 | 0 | 0 |
| miR-1933-3p | 0 | 0 | 0 | 0 | 0 | 0 | 0 | 0 | 0 |
| miR-378a-5p | 4.1768464 | 0 | 0.40796974 | 0 | 0.54516889 | 2.16830137 | 0 | 0 | 2.4515482 |
| miR-7092-3p | 1.39228213 | 0 | 0 | 0 | 0 | 0 | 1.41781837 | 0 | 0 |
| miR-1191a | 0 | 0 | 0 | 1.27378306 | 0.54516889 | 0 | 0 | 0 | 0 |
| miR-99a-3p | 2.78456427 | 0 | 0.40796974 | 0.63689153 | 0.54516889 | 0 | 0 | 0 | 0.61288705 |
| miR-3079-5p | 0 | 1.36653541 | 0 | 0 | 0 | 0 | 0 | 3.05220649 | 0 |
| miR-411-3p | 0 | 0 | 1.63187896 | 0.63689153 | 2.18067556 | 0 | 1.41781837 | 0 | 0.61288705 |
| miR-465d-3p | 0 | 0 | 0 | 0 | 0 | 1.08415069 | 1.41781837 | 0 | 0 |
| miR-363-3p | 0 | 0 | 0.40796974 | 1.91067459 | 0 | 0 | 0 | 0 | 0 |
| miR-30c-1-3p | 0 | 0 | 0 | 0 | 1.63550667 | 1.08415069 | 1.41781837 | 0 | 1.2257741 |
| miR-503-5p | 0 | 1.36653541 | 1.22390922 | 0.63689153 | 1.63550667 | 3.25245206 | 0 | 0 | 0.61288705 |
| miR-8114 | 6.96141067 | 4.09960624 | 0.81593948 | 1.27378306 | 0 | 0 | 0 | 3.05220649 | 0.61288705 |
| miR-6902-5p | 0 | 0 | 0 | 0 | 0 | 2.16830137 | 0 | 0 | 0 |
| miR-6983-5p | 0 | 0 | 0 | 0 | 0 | 0 | 2.83563673 | 0 | 0 |
| miR-467e-3p | 0 | 0 | 0 | 0.63689153 | 0 | 0 | 1.41781837 | 0 | 0 |
| miR-1955-3p | 0 | 0 | 0 | 0 | 0 | 2.16830137 | 0 | 0 | 0 |
| miR-7049-5p | 0 | 0 | 0 | 0.63689153 | 0 | 0 | 1.41781837 | 0 | 0 |
| miR-7082-3p | 0 | 0 | 0 | 0 | 0 | 2.16830137 | 0 | 0 | 0 |

| | | | | | | | | | |
|---|---|---|---|---|---|---|---|---|---|
| miR-6967-5p | 0 | 0 | 0 | 1.27378306 | 0 | 0 | 0 | 0 | 0 |
| miR-5107-3p | 0 | 0 | 0 | 1.27378306 | 0 | 0 | 0 | 0 | 0 |
| miR-483-5p | 0 | 0 | 0 | 0 | 0 | 0 | 0 | 3.05220649 | 0 |
| miR-449c-3p | 2.78456427 | 0 | 0 | 0 | 0 | 0 | 0 | 0 | 0 |
| miR-7019-5p | 2.78456427 | 0 | 0 | 0 | 0 | 0 | 0 | 0 | 0 |
| miR-6920-3p | 0 | 2.73307083 | 0 | 0 | 0 | 0 | 0 | 0 | 0 |
| miR-6959-3p | 0 | 2.73307083 | 0 | 0 | 0 | 0 | 0 | 0 | 0 |
| miR-3082-5p | 4.1768464 | 0 | 0.40796974 | 0 | 0 | 2.16830137 | 0 | 0 | 0 |
| miR-183-3p | 8.35369281 | 1.36653541 | 2.85578818 | 2.54756612 | 4.90652001 | 1.08415069 | 0 | 1.52610324 | 1.83866115 |
| miR-1247-3p | 0 | 0 | 0.40796974 | 0 | 1.09033778 | 1.08415069 | 0 | 0 | 1.83866115 |
| miR-6907-3p | 1.39228213 | 0 | 0 | 0 | 0 | 0 | 0 | 0 | 0 |
| miR-7234-5p | 4.1768464 | 1.36653541 | 1.63187896 | 1.91067459 | 1.63550667 | 0 | 0 | 0 | 1.2257741 |
| miR-7116-3p | 0 | 0 | 0 | 0.63689153 | 0.54516889 | 0 | 0 | 0 | 0 |
| miR-3070-5p | 1.39228213 | 0 | 0 | 0 | 0.54516889 | 0 | 0 | 0 | 0 |
| miR-2139 | 1.39228213 | 0 | 0 | 0 | 0.54516889 | 0 | 0 | 0 | 0 |
| miR-214-5p | 0 | 0 | 1.22390922 | 0.63689153 | 0 | 0 | 0 | 0 | 0.61288705 |
| miR-5626-5p | 0 | 0 | 0 | 0 | 0 | 2.16830137 | 0 | 0 | 0.61288705 |
| miR-207 | 0 | 0 | 0 | 0 | 0 | 3.25245206 | 0 | 0 | 0 |
| miR-6238 | 1.39228213 | 0 | 0 | 0 | 0 | 0 | 0 | 0 | 0.61288705 |
| miR-6897-5p | 0 | 0 | 0 | 0 | 1.09033778 | 0 | 0 | 0 | 0 |
| miR-3084-5p | 0 | 0 | 0.81593948 | 0 | 0.54516889 | 1.08415069 | 0 | 3.05220649 | 1.83866115 |
| miR-125b-5p | 955.105544 | 755.694084 | 727.818017 | 780.829015 | 654.747837 | 771.915288 | 917.328484 | 668.43322 | 881.944466 |
| miR-3062-5p | 0 | 0 | 1.22390922 | 0 | 2.18067556 | 2.16830137 | 0 | 0 | 0.61288705 |
| miR-5132-5p | 2.78456427 | 0 | 0 | 0 | 0 | 0 | 0 | 0 | 0.61288705 |
| miR-7688-5p | 0 | 0 | 0.81593948 | 0 | 0.54516889 | 2.16830137 | 0 | 0 | 1.2257741 |
| miR-7007-3p | 2.78456427 | 0 | 0 | 0.63689153 | 0.54516889 | 0 | 0 | 0 | 0.61288705 |
| miR-7015-3p | 0 | 0 | 0.81593948 | 0 | 0 | 0 | 1.41781837 | 0 | 0 |
| miR-1843a-5p | 1.39228213 | 2.73307083 | 7.34345533 | 7.64269835 | 7.08719557 | 6.50490411 | 5.67127347 | 7.63051622 | 8.58041871 |
| miR-466c-5p | 0 | 0 | 0.81593948 | 0 | 0 | 0 | 0 | 1.52610324 | 0 |
| miR-487b-3p | 0 | 0 | 0.40796974 | 0 | 2.72584445 | 2.16830137 | 2.83563673 | 0 | 1.2257741 |

| | | | | | | | | | |
|---|---|---|---|---|---|---|---|---|---|
| miR-7661-3p | 0 | 4.09960624 | 0.40796974 | 0 | 0 | 0 | 0 | 0 | 2.4515482 |
| miR-183-5p | 445.530283 | 550.713772 | 419.800863 | 349.016558 | 621.492534 | 1242.43669 | 917.328484 | 767.629931 | 409.40855 |
| miR-485-3p | 2.78456427 | 2.73307083 | 0 | 0 | 1.63550667 | 0 | 0 | 0 | 0 |
| miR-7001-3p | 0 | 0 | 0 | 0 | 0 | 0 | 0 | 1.52610324 | 0 |
| miR-8098 | 1.39228213 | 0 | 0 | 0 | 0 | 0 | 0 | 0 | 0.61288705 |
| miR-539-5p | 0 | 0 | 0 | 0 | 0 | 0 | 0 | 3.05220649 | 0 |
| miR-1903 | 0 | 1.36653541 | 0 | 0 | 0 | 0 | 0 | 0 | 0.61288705 |
| miR-7225-3p | 0 | 0 | 0.40796974 | 0 | 0 | 1.08415069 | 0 | 0 | 0 |
| miR-363-5p | 0 | 1.36653541 | 0.40796974 | 0 | 0 | 0 | 0 | 0 | 0 |
| miR-6935-3p | 1.39228213 | 0 | 0.40796974 | 0 | 0 | 0 | 0 | 0 | 0 |
| miR-3091-3p | 1.39228213 | 0 | 0.40796974 | 0 | 0 | 0 | 0 | 0 | 0 |
| miR-1933-5p | 0 | 0 | 0 | 0 | 0 | 1.08415069 | 0 | 0 | 0 |
| miR-3108-5p | 0 | 0 | 0 | 0 | 0.54516889 | 0 | 0 | 0 | 0 |
| miR-3474 | 0 | 0 | 0.40796974 | 0 | 0 | 0 | 0 | 0 | 0.61288705 |
| miR-450b-3p | 0 | 0 | 0.81593948 | 1.27378306 | 0 | 0 | 0 | 0 | 0 |
| miR-381-3p | 0 | 0 | 0.40796974 | 0.63689153 | 1.63550667 | 0 | 0 | 0 | 0.61288705 |
| miR-376c-3p | 1.39228213 | 0 | 1.22390922 | 0.63689153 | 3.81618223 | 0 | 0 | 0 | 0 |
| miR-199b-5p | 1.39228213 | 2.73307083 | 2.44781844 | 1.27378306 | 3.81618223 | 1.08415069 | 0 | 1.52610324 | 1.2257741 |
| miR-6948-3p | 0 | 0 | 0 | 0.63689153 | 0 | 0 | 0 | 0 | 0 |
| miR-3094-3p | 0 | 0 | 0 | 0 | 0 | 1.08415069 | 0 | 0 | 0.61288705 |
| miR-212-3p | 0 | 0 | 0.81593948 | 0 | 0.54516889 | 0 | 0 | 0 | 0 |
| miR-7217-3p | 5.56912854 | 0 | 4.48766714 | 1.91067459 | 6.54202668 | 2.16830137 | 5.67127347 | 10.6827227 | 1.83866115 |
| miR-335-3p | 1.39228213 | 2.73307083 | 0.40796974 | 4.45824071 | 1.09033778 | 0 | 1.41781837 | 1.52610324 | 4.29020936 |
| miR-28c | 0 | 0 | 0.40796974 | 0.63689153 | 0 | 2.16830137 | 0 | 0 | 0.61288705 |
| miR-665-3p | 0 | 0 | 0 | 0 | 0 | 0 | 1.41781837 | 0 | 0.61288705 |
| miR-3108-3p | 0 | 1.36653541 | 0 | 0 | 0 | 0 | 0 | 0 | 0.61288705 |
| miR-200b-3p | 1285.07641 | 1730.03383 | 1083.56763 | 944.510138 | 1108.87352 | 2591.12014 | 2512.37415 | 1866.42427 | 1497.89595 |
| miR-6932-5p | 0 | 0 | 0.81593948 | 0 | 0 | 0 | 0 | 0 | 0 |
| miR-3081-3p | 0 | 0 | 0.81593948 | 0 | 0 | 0 | 0 | 0 | 0 |
| miR-3102-5p | 0 | 0 | 0.81593948 | 0 | 0 | 0 | 0 | 0 | 0 |

| | | | | | | | | | |
|---|---|---|---|---|---|---|---|---|---|
| miR-3086-5p | 0 | 0 | 0.40796974 | 0 | 0 | 0 | 0 | 0 | 0 |
| miR-6975-3p | 0 | 0 | 0 | 1.27378306 | 0 | 0 | 1.41781837 | 0 | 0 |
| miR-7079-3p | 0 | 0 | 0 | 1.27378306 | 0 | 0 | 0 | 0 | 1.2257741 |
| miR-7218-3p | 1.39228213 | 2.73307083 | 0 | 2.54756612 | 0.54516889 | 0 | 0 | 0 | 1.83866115 |
| miR-1906 | 0 | 0 | 0.40796974 | 0 | 0 | 0 | 0 | 0 | 0 |
| miR-7118-3p | 0 | 0 | 0.40796974 | 0 | 0 | 0 | 0 | 0 | 0 |
| miR-205-3p | 0 | 0 | 1.22390922 | 1.27378306 | 0 | 0 | 0 | 0 | 1.2257741 |
| miR-19b-3p | 18.0996677 | 45.0956687 | 46.1005807 | 41.3979494 | 20.7164178 | 7.5890548 | 7.08909184 | 13.7349292 | 31.2572396 |
| miR-376a-3p | 0 | 0 | 0.40796974 | 1.27378306 | 0.54516889 | 0 | 0 | 0 | 2.4515482 |
| miR-92a-1-5p | 4.1768464 | 4.09960624 | 0.81593948 | 1.91067459 | 1.09033778 | 0 | 1.41781837 | 1.52610324 | 1.83866115 |
| miR-200c-3p | 2240.18195 | 1959.61178 | 1131.30009 | 1052.7817 | 2091.81303 | 2180.22703 | 2345.07158 | 2055.66107 | 1875.43438 |
| miR-324-3p | 0 | 0 | 0.81593948 | 2.54756612 | 0 | 0 | 0 | 0 | 0 |
| miR-719 | 1.39228213 | 1.36653541 | 0 | 0 | 0 | 0 | 0 | 0 | 0 |
| miR-3058-3p | 0 | 1.36653541 | 0 | 0 | 0 | 0 | 0 | 0 | 0 |
| miR-681 | 4.1768464 | 0 | 0 | 1.27378306 | 0 | 0 | 0 | 0 | 1.83866115 |
| miR-29a-3p | 714.240735 | 896.447232 | 878.766821 | 1022.8478 | 743.610366 | 697.108891 | 849.273202 | 892.770397 | 729.335591 |
| miR-505-5p | 0 | 0 | 0 | 0 | 0.54516889 | 0 | 0 | 0 | 1.2257741 |
| miR-467d-3p | 1.39228213 | 1.36653541 | 0.81593948 | 1.27378306 | 1.63550667 | 0 | 0 | 3.05220649 | 1.2257741 |
| miR-28a-3p | 6.96141067 | 10.9322833 | 3.26375792 | 6.36891529 | 8.17753335 | 15.1781096 | 7.08909184 | 7.63051622 | 3.67732231 |
| miR-350-5p | 0 | 0 | 0.81593948 | 1.91067459 | 0.54516889 | 0 | 0 | 0 | 0 |
| miR-935 | 0 | 0 | 0.40796974 | 0 | 0 | 0 | 0 | 3.05220649 | 0 |
| miR-6946-5p | 0 | 1.36653541 | 0 | 0.63689153 | 0 | 0 | 0 | 0 | 0 |
| miR-7a-1-3p | 2.78456427 | 2.73307083 | 1.63187896 | 1.27378306 | 2.18067556 | 0 | 1.41781837 | 3.05220649 | 4.29020936 |
| miR-7059-3p | 1.39228213 | 0 | 0 | 0.63689153 | 0 | 0 | 0 | 0 | 0.61288705 |
| miR-181a-2-3p | 0 | 2.73307083 | 0 | 0.63689153 | 2.18067556 | 2.16830137 | 0 | 0 | 3.06443526 |
| let-7i-3p | 0 | 1.36653541 | 0.81593948 | 2.54756612 | 2.18067556 | 0 | 4.2534551 | 0 | 0.61288705 |
| miR-574-3p | 4.1768464 | 6.83267707 | 7.34345533 | 7.00580682 | 9.81304002 | 1.08415069 | 5.67127347 | 3.05220649 | 6.74175756 |
| miR-470-3p | 0 | 0 | 0.40796974 | 0.63689153 | 0.54516889 | 0 | 0 | 0 | 0.61288705 |
| miR-669b-5p | 0 | 0 | 0.40796974 | 0.63689153 | 0.54516889 | 0 | 0 | 0 | 0 |
| miR-669m-5p | 0 | 0 | 0.40796974 | 0.63689153 | 0.54516889 | 0 | 0 | 0 | 0 |

| | | | | | | | | | |
|---|---|---|---|---|---|---|---|---|---|
| miR-138-5p | 0 | 0 | 0.40796974 | 0 | 1.09033778 | 0 | 1.41781837 | 0 | 0.61288705 |
| miR-7226-3p | 0 | 0 | 0.40796974 | 0 | 0 | 0 | 0 | 0 | 0 |
| miR-3960 | 2.78456427 | 0 | 0 | 0 | 0 | 0 | 0 | 0 | 0 |
| miR-412-5p | 0 | 1.36653541 | 0 | 0 | 0.54516889 | 0 | 0 | 0 | 0 |
| miR-331-5p | 0 | 0 | 1.63187896 | 1.91067459 | 0.54516889 | 0 | 0 | 0 | 0.61288705 |
| miR-3087-3p | 0 | 0 | 0 | 0 | 0 | 0 | 0 | 3.05220649 | 0.61288705 |
| miR-7232-3p | 4.1768464 | 4.09960624 | 0.81593948 | 1.27378306 | 1.63550667 | 1.08415069 | 0 | 3.05220649 | 2.4515482 |
| miR-139-3p | 0 | 0 | 0.81593948 | 0.63689153 | 0.54516889 | 6.50490411 | 0 | 3.05220649 | 0.61288705 |
| let-7c-2-3p | 0 | 6.83267707 | 0.81593948 | 2.54756612 | 1.09033778 | 1.08415069 | 0 | 0 | 4.90309641 |
| miR-7042-3p | 0 | 0 | 0 | 0 | 0 | 0 | 2.83563673 | 0 | 0.61288705 |
| miR-1966-3p | 0 | 0 | 0 | 0 | 0 | 0 | 2.83563673 | 0 | 0 |
| miR-295-3p | 1.39228213 | 0 | 0.40796974 | 2.54756612 | 0.54516889 | 1.08415069 | 0 | 0 | 0 |
| miR-7221-3p | 0 | 1.36653541 | 0 | 0 | 1.63550667 | 0 | 1.41781837 | 0 | 1.2257741 |
| miR-329-3p | 0 | 0 | 0.40796974 | 1.27378306 | 0 | 0 | 1.41781837 | 0 | 1.2257741 |
| miR-370-3p | 0 | 0 | 0.81593948 | 0.63689153 | 0 | 0 | 0 | 0 | 0 |
| miR-493-3p | 0 | 0 | 0 | 0.63689153 | 0.54516889 | 0 | 0 | 0 | 0.61288705 |
| miR-511-3p | 0 | 0 | 1.22390922 | 0 | 0 | 0 | 0 | 0 | 0.61288705 |
| miR-669d-5p | 1.39228213 | 1.36653541 | 3.67172766 | 2.54756612 | 2.18067556 | 1.08415069 | 1.41781837 | 3.05220649 | 4.29020936 |
| miR-153-3p | 0 | 0 | 0.40796974 | 0 | 0 | 1.08415069 | 0 | 0 | 0 |
| miR-669c-5p | 80.7523638 | 72.4263769 | 26.9260029 | 23.5649866 | 54.516889 | 79.143 | 65.2196449 | 94.6184011 | 68.0304627 |
| miR-742-5p | 2.78456427 | 4.09960624 | 7.75142507 | 7.00580682 | 4.36135112 | 4.33660274 | 1.41781837 | 1.52610324 | 1.83866115 |
| miR-877-5p | 4.1768464 | 0 | 0 | 0 | 0.54516889 | 0 | 0 | 0 | 0 |
| miR-883b-3p | 1.39228213 | 0 | 4.89563688 | 9.55337294 | 2.72584445 | 2.16830137 | 0 | 1.52610324 | 2.4515482 |
| miR-297b-5p | 0 | 0 | 0.40796974 | 0 | 1.09033778 | 0 | 0 | 0 | 0 |
| miR-151-3p | 23.6687963 | 36.8964562 | 13.8709712 | 18.4698544 | 23.9874312 | 56.3758356 | 17.0138204 | 22.8915486 | 12.257741 |
| miR-1968-5p | 0 | 0 | 0 | 0 | 1.09033778 | 0 | 0 | 0 | 0 |
| miR-540-3p | 1.39228213 | 0 | 0.40796974 | 0 | 0 | 0 | 0 | 0 | 0.61288705 |
| miR-5131 | 1.39228213 | 0 | 0.40796974 | 0 | 0 | 0 | 0 | 0 | 0.61288705 |
| miR-669p-3p | 1.39228213 | 0 | 0.40796974 | 0 | 0 | 0 | 0 | 0 | 0.61288705 |
| miR-532-5p | 18.0996677 | 12.2988187 | 13.4630014 | 8.91648141 | 9.26787113 | 18.4305616 | 9.92472857 | 18.3132389 | 16.5479504 |

| | | | | | | | | | |
|---|---|---|---|---|---|---|---|---|---|
| miR-7214-5p | 4.1768464 | 13.6653541 | 4.89563688 | 7.00580682 | 10.9033778 | 4.33660274 | 11.3425469 | 7.63051622 | 11.644854 |
| miR-7229-5p | 4.1768464 | 8.19921248 | 1.63187896 | 1.27378306 | 1.63550667 | 2.16830137 | 0 | 1.52610324 | 2.4515482 |
| miR-223-3p | 23.6687963 | 32.7968499 | 39.5730648 | 54.7726715 | 32.1649645 | 19.5147123 | 28.3563673 | 15.2610324 | 26.9670302 |
| miR-326-3p | 1.39228213 | 0 | 0.81593948 | 0.63689153 | 0 | 1.08415069 | 0 | 0 | 0 |
| miR-148b-3p | 34.8070534 | 30.0637791 | 62.0114005 | 80.2483327 | 34.8908089 | 62.8807397 | 31.1920041 | 56.46582 | 26.3541432 |
| miR-7658-3p | 0 | 0 | 0.40796974 | 0 | 0.54516889 | 0 | 0 | 0 | 0 |
| miR-7242-5p | 2.78456427 | 2.73307083 | 0.40796974 | 0.63689153 | 4.36135112 | 0 | 0 | 0 | 0 |
| miR-1927 | 1.39228213 | 0 | 0.40796974 | 0.63689153 | 0 | 0 | 0 | 0 | 1.2257741 |
| miR-425-3p | 6.96141067 | 2.73307083 | 5.30360662 | 1.27378306 | 4.90652001 | 4.33660274 | 0 | 0 | 3.67732231 |
| miR-7a-2-3p | 1.39228213 | 0 | 0.81593948 | 0.63689153 | 0 | 0 | 0 | 0 | 0 |
| miR-7068-3p | 2.78456427 | 1.36653541 | 0.40796974 | 1.27378306 | 0.54516889 | 0 | 1.41781837 | 0 | 0.61288705 |
| miR-206-3p | 0 | 0 | 0 | 0 | 0.54516889 | 1.08415069 | 2.83563673 | 0 | 0.61288705 |
| miR-193b-3p | 0 | 0 | 0 | 1.27378306 | 0 | 0 | 0 | 3.05220649 | 0 |
| miR-130a-3p | 2.78456427 | 5.46614166 | 8.15939481 | 5.73202376 | 9.81304002 | 2.16830137 | 4.2534551 | 3.05220649 | 6.12887051 |
| miR-294-3p | 0 | 1.36653541 | 1.22390922 | 0 | 1.63550667 | 0 | 4.2534551 | 4.57830973 | 1.2257741 |
| miR-878-3p | 26.4533606 | 12.2988187 | 46.9165201 | 45.8561901 | 31.6197956 | 4.33660274 | 5.67127347 | 19.8393422 | 16.5479504 |
| miR-5101 | 0 | 0 | 0.40796974 | 0 | 0.54516889 | 0 | 0 | 0 | 0 |
| miR-3060-3p | 1.39228213 | 0 | 0.40796974 | 0 | 0 | 0 | 0 | 0 | 0 |
| miR-7240-5p | 0 | 0 | 0.81593948 | 0.63689153 | 0 | 0 | 0 | 0 | 0 |
| miR-6546-5p | 0 | 0 | 0.81593948 | 0 | 0 | 0 | 0 | 0 | 0.61288705 |
| miR-365-2-5p | 1.39228213 | 0 | 0.40796974 | 0.63689153 | 1.09033778 | 2.16830137 | 0 | 0 | 1.2257741 |
| miR-18a-3p | 1.39228213 | 0 | 0.81593948 | 1.91067459 | 0.54516889 | 1.08415069 | 0 | 0 | 1.2257741 |
| miR-30c-2-3p | 29.2379248 | 23.231102 | 14.2789409 | 14.0116136 | 28.8939512 | 30.3562192 | 18.4316388 | 16.7871357 | 15.9350633 |
| miR-1839-5p | 58.4758496 | 62.860629 | 57.5237334 | 34.3921426 | 70.3267868 | 80.2271507 | 76.5621918 | 93.0922978 | 60.062931 |
| miR-3069-3p | 0 | 2.73307083 | 0.40796974 | 0 | 0 | 0 | 0 | 0 | 0.61288705 |
| miR-7223-5p | 2.78456427 | 0 | 0.40796974 | 0 | 1.63550667 | 1.08415069 | 0 | 0 | 1.83866115 |
| miR-668-3p | 0 | 0 | 0 | 0.63689153 | 1.09033778 | 3.25245206 | 0 | 0 | 0.61288705 |
| miR-3620-3p | 0 | 0 | 0.40796974 | 0 | 0 | 0 | 0 | 0 | 0 |
| miR-370-5p | 0 | 0 | 0.40796974 | 0 | 0 | 0 | 0 | 0 | 0 |
| miR-666-3p | 0 | 0 | 0.40796974 | 0 | 0 | 0 | 0 | 0 | 0 |

| | | | | | | | | | |
|---|---|---|---|---|---|---|---|---|---|
| miR-6357 | 0 | 0 | 0.40796974 | 0 | 0 | 0 | 0 | 0 | 0 |
| miR-7075-3p | 0 | 0 | 0.40796974 | 0 | 0 | 0 | 0 | 0 | 0 |
| miR-1929-5p | 0 | 0 | 0.40796974 | 0 | 0 | 0 | 0 | 0 | 0 |
| miR-6989-3p | 0 | 0 | 0.40796974 | 0 | 0 | 0 | 0 | 0 | 0 |
| miR-7215-5p | 0 | 0 | 0.40796974 | 0 | 0 | 0 | 0 | 0 | 0.61288705 |
| miR-541-3p | 0 | 0 | 0.40796974 | 0 | 0 | 0 | 0 | 0 | 0.61288705 |
| miR-6964-3p | 0 | 0 | 0.40796974 | 0 | 0 | 0 | 0 | 0 | 0.61288705 |
| miR-7026-5p | 0 | 0 | 0.40796974 | 0 | 0 | 0 | 0 | 0 | 0 |
| miR-7647-3p | 0 | 0 | 0.40796974 | 0 | 0 | 0 | 0 | 0 | 0.61288705 |
| miR-224-3p | 0 | 0 | 0.40796974 | 0 | 0 | 0 | 0 | 0 | 0.61288705 |
| let-7f-2-3p | 0 | 0 | 0 | 1.27378306 | 0.54516889 | 0 | 0 | 3.05220649 | 1.2257741 |
| miR-7006-3p | 1.39228213 | 0 | 0 | 0 | 0 | 0 | 0 | 0 | 0 |
| miR-3103-3p | 1.39228213 | 0 | 0 | 0 | 0 | 0 | 0 | 0 | 0 |
| miR-434-5p | 0 | 1.36653541 | 1.22390922 | 1.91067459 | 1.63550667 | 0 | 0 | 4.57830973 | 1.2257741 |
| miR-141-3p | 19.4919499 | 39.629527 | 69.3548559 | 109.545343 | 65.9654357 | 20.598863 | 43.9523694 | 33.5742713 | 53.3211734 |
| miR-3620-5p | 0 | 0 | 0 | 0 | 0.54516889 | 0 | 0 | 0 | 0 |
| miR-761 | 0 | 1.36653541 | 0 | 0 | 0 | 0 | 0 | 0 | 0 |
| miR-1898 | 0 | 1.36653541 | 0 | 0 | 0 | 0 | 0 | 0 | 0 |
| miR-669e-5p | 0 | 1.36653541 | 0 | 0 | 0 | 0 | 0 | 0 | 0 |
| miR-291a-5p | 0 | 0 | 0 | 0 | 0 | 0 | 0 | 1.52610324 | 0 |
| miR-7046-3p | 0 | 0 | 0 | 0 | 0 | 0 | 0 | 1.52610324 | 0 |
| miR-5129-3p | 0 | 0 | 0 | 0 | 0 | 0 | 1.41781837 | 0 | 0 |
| miR-7019-3p | 0 | 0 | 0 | 0.63689153 | 0 | 0 | 0 | 0 | 0 |
| miR-7059-5p | 0 | 0 | 0 | 0 | 0 | 1.08415069 | 0 | 0 | 0 |
| miR-7060-3p | 0 | 1.36653541 | 0 | 0 | 0 | 0 | 0 | 0 | 0 |
| miR-7650-3p | 1.39228213 | 0 | 0 | 0 | 0 | 0 | 0 | 0 | 0 |
| miR-1249-5p | 0 | 0 | 0 | 0.63689153 | 0 | 0 | 0 | 0 | 0 |
| miR-3083-5p | 0 | 0 | 0 | 0.63689153 | 0 | 0 | 0 | 0 | 0 |
| miR-296-3p | 0 | 0 | 0 | 0 | 0.54516889 | 0 | 0 | 0 | 0 |
| miR-7070-5p | 0 | 0 | 0 | 0.63689153 | 0 | 0 | 0 | 0 | 0 |

| | | | | | | | | | |
|---|---|---|---|---|---|---|---|---|---|
| miR-6899-3p | 0 | 1.36653541 | 0 | 0 | 0 | 0 | 0 | 0 | 0 |
| miR-1982-5p | 0 | 0 | 0 | 0.63689153 | 0 | 0 | 0 | 0 | 0.61288705 |
| miR-103-1-5p | 0 | 0 | 0 | 0.63689153 | 0 | 0 | 0 | 0 | 0 |
| miR-7652-3p | 0 | 0 | 0 | 0 | 0.54516889 | 0 | 0 | 0 | 0 |
| miR-1936 | 0 | 0 | 0 | 0 | 0.54516889 | 0 | 0 | 0 | 0 |
| miR-666-5p | 0 | 0 | 0 | 0 | 0.54516889 | 0 | 0 | 0 | 0.61288705 |
| miR-7118-5p | 0 | 0 | 0 | 0.63689153 | 0 | 0 | 0 | 0 | 0 |
| miR-877-3p | 0 | 0 | 0 | 0.63689153 | 0 | 0 | 0 | 0 | 0.61288705 |
| miR-6988-3p | 0 | 1.36653541 | 0 | 0 | 0 | 0 | 0 | 0 | 0.61288705 |
| miR-3970 | 0 | 1.36653541 | 0 | 0 | 0 | 0 | 0 | 0 | 0 |
| miR-708-5p | 0 | 0 | 0 | 0 | 0.54516889 | 0 | 0 | 0 | 0.61288705 |
| miR-7060-5p | 0 | 1.36653541 | 0 | 0 | 0 | 0 | 0 | 0 | 0.61288705 |
| miR-5130 | 1.39228213 | 0 | 0 | 0.63689153 | 0 | 0 | 0 | 0 | 0.61288705 |
| miR-6946-3p | 0 | 0 | 0.40796974 | 0.63689153 | 0 | 0 | 0 | 3.05220649 | 1.2257741 |
| miR-1964-3p | 0 | 0 | 0 | 0 | 0 | 0 | 0 | 3.05220649 | 0 |
| miR-193a-3p | 0 | 0 | 0 | 1.27378306 | 0.54516889 | 0 | 0 | 0 | 1.2257741 |
| miR-8118 | 2.78456427 | 0 | 0 | 0 | 0 | 0 | 0 | 0 | 0.61288705 |
| miR-147-3p | 0 | 0 | 0 | 1.27378306 | 0.54516889 | 0 | 0 | 0 | 0 |
| miR-1934-5p | 1.39228213 | 0 | 0 | 0.63689153 | 0 | 0 | 0 | 1.52610324 | 0.61288705 |
| miR-500-3p | 0 | 1.36653541 | 0.40796974 | 1.27378306 | 2.18067556 | 0 | 0 | 1.52610324 | 4.29020936 |
| miR-1969 | 0 | 0 | 1.22390922 | 0 | 0 | 0 | 0 | 0 | 0 |
| miR-452-5p | 0 | 0 | 0 | 0 | 0 | 0 | 0 | 3.05220649 | 0 |
| miR-130a-5p | 0 | 0 | 0 | 0 | 0 | 0 | 2.83563673 | 0 | 0 |
| miR-542-3p | 0 | 0 | 0 | 0.63689153 | 0 | 3.25245206 | 0 | 0 | 0.61288705 |
| miR-15b-5p | 552.736007 | 254.175587 | 379.003889 | 351.564124 | 318.923801 | 199.483726 | 293.488402 | 405.943463 | 291.734236 |

| ree_miRNA_dedup_count | FGCZ6510_Free_miRNA_dedup_count | | | | | |
|---|---|---|---|---|---|---|
| DEX.4 | DEX.5 | DEX.6 | DEX.7 | DEX.8 | DEX.9 | DEX.10 |
| 178.892386 | 305.740062 | 151.689991 | 217.537745 | 203.293939 | 135.875649 | 258.777239 |
| 208.299354 | 135.220738 | 111.643833 | 145.025163 | 29.0419913 | 63.1702577 | 90.7752618 |
| 107.825548 | 83.0874415 | 90.4072344 | 120.854303 | 39.414131 | 59.5945828 | 69.0975874 |
| 487.665547 | 309.541449 | 503.004009 | 575.012051 | 399.32738 | 495.826929 | 410.52096 |
| 14.7034838 | 52.6763518 | 58.2489564 | 35.6202155 | 39.414131 | 30.989183 | 25.7422384 |
| 9.80232254 | 9.23193794 | 7.88787951 | 17.8101078 | 25.9303493 | 15.4945915 | 46.0650583 |
| 129.880774 | 19.5499862 | 47.934037 | 31.8037639 | 32.1536332 | 21.4540498 | 12.1936919 |
| 154.38658 | 261.752593 | 144.408871 | 310.404735 | 69.4933362 | 83.4324159 | 70.452442 |
| 306.32258 | 95.0346553 | 67.3503558 | 68.6961299 | 67.4189083 | 48.8675579 | 21.6776745 |
| 29.4069676 | 5.9736069 | 9.10139943 | 19.0822583 | 24.8931354 | 11.9189166 | 21.6776745 |
| 14.7034838 | 7.60277242 | 6.06759962 | 22.89871 | 11.4093537 | 7.15134993 | 8.12912793 |
| 7.35174191 | 19.0069311 | 12.1351992 | 43.2531189 | 28.0047773 | 21.4540498 | 40.6456396 |
| 14.7034838 | 36.9277518 | 26.6974383 | 71.2404311 | 32.1536332 | 27.4135081 | 55.5490408 |
| 22.0552257 | 30.4110897 | 32.765038 | 59.7910761 | 43.5629869 | 36.9486413 | 85.3558432 |
| 85.7703223 | 67.3388415 | 28.5177182 | 77.6011838 | 67.4189083 | 23.8378331 | 56.9038955 |
| 1666.39483 | 447.477463 | 1277.83648 | 718.765063 | 1010.24641 | 858.161992 | 459.295728 |
| 9.80232254 | 18.4638759 | 20.6298387 | 24.1708605 | 15.5582096 | 10.7270249 | 18.9679652 |
| 4.90116127 | 7.60277242 | 4.8540797 | 2.54430111 | 7.26049782 | 7.15134993 | 5.41941862 |
| 29.4069676 | 77.6568898 | 69.7773956 | 89.0505388 | 66.3816943 | 101.310791 | 73.1621513 |
| 7.35174191 | 10.3180483 | 9.70815939 | 7.63290333 | 12.4465677 | 5.95945828 | 4.06456396 |
| 17.1540645 | 3.80138621 | 8.49463947 | 13.9936561 | 5.18606987 | 8.34324159 | 9.48398258 |
| 29.4069676 | 17.9208207 | 10.3149194 | 27.9873122 | 17.6326376 | 29.7972914 | 28.4519477 |
| 0 | 5.43055173 | 3.64055977 | 6.36075277 | 5.18606987 | 5.95945828 | 5.41941862 |
| 2.45058064 | 15.7486 | 8.49463947 | 16.5379572 | 18.6698515 | 15.4945915 | 8.12912793 |
| 17.1540645 | 51.5902414 | 20.6298387 | 29.2594628 | 48.7490568 | 20.2621581 | 24.3873838 |
| 423.95045 | 673.388415 | 360.415418 | 1034.2584 | 1000.91148 | 748.50796 | 1082.52887 |
| 49.0116127 | 89.6041036 | 33.9785579 | 63.6075277 | 113.056323 | 34.564858 | 116.5175 |
| 22.0552257 | 129.247131 | 60.0692363 | 45.79742 | 43.5629869 | 32.1810747 | 44.7102036 |

| | | | | | | |
|---|---|---|---|---|---|---|
| 2553.50502 | 5705.33765 | 1914.32768 | 2836.89574 | 4941.28737 | 1946.35907 | 4820.57286 |
| 129.880774 | 185.724869 | 185.668548 | 207.36054 | 163.879808 | 196.662123 | 222.196163 |
| 46.5610321 | 18.4638759 | 58.8557163 | 30.5316133 | 49.7862707 | 64.3621494 | 37.9359303 |
| 205.848773 | 65.709676 | 186.275308 | 122.126453 | 183.586873 | 184.743207 | 172.066541 |
| 183.793548 | 212.334573 | 227.534986 | 199.727637 | 185.661301 | 264.599948 | 238.454419 |
| 210.749935 | 66.7957863 | 187.488828 | 127.215055 | 186.698515 | 185.935098 | 172.066541 |
| 9.80232254 | 0.54305517 | 1.21351992 | 1.27215055 | 1.03721397 | 7.15134993 | 1.35485465 |
| 0 | 0.54305517 | 0 | 2.54430111 | 2.07442795 | 4.76756662 | 1.35485465 |
| 105.374967 | 33.6694207 | 93.4410342 | 63.6075277 | 93.3492576 | 92.9675491 | 86.7106979 |
| 63.7150965 | 20.6360966 | 29.7312381 | 27.9873122 | 20.7442795 | 29.7972914 | 16.2582559 |
| 9.80232254 | 26.0666483 | 10.3149194 | 13.9936561 | 24.8931354 | 9.53513324 | 12.1936919 |
| 4.90116127 | 9.77499312 | 3.64055977 | 6.36075277 | 6.22328384 | 5.95945828 | 10.8388372 |
| 754.778836 | 232.427614 | 415.023814 | 274.78452 | 684.561223 | 425.505321 | 814.267647 |
| 44.1104514 | 21.7222069 | 12.1351992 | 49.6138716 | 32.1536332 | 17.8783748 | 29.8068024 |
| 4.90116127 | 15.2055448 | 12.1351992 | 21.6265594 | 9.33492576 | 11.9189166 | 8.12912793 |
| 7.35174191 | 1.62916552 | 2.42703985 | 2.54430111 | 6.22328384 | 0 | 5.41941862 |
| 17.1540645 | 4.88749656 | 12.1351992 | 19.0822583 | 7.26049782 | 8.34324159 | 6.77427327 |
| 12.2529032 | 10.3180483 | 3.64055977 | 2.54430111 | 2.07442795 | 4.76756662 | 1.35485465 |
| 154.38658 | 164.545717 | 153.51027 | 111.949249 | 116.167965 | 106.078357 | 178.840814 |
| 17.1540645 | 60.2791242 | 30.9447581 | 39.4366672 | 31.1164192 | 33.3729664 | 25.7422384 |
| 41.6598708 | 47.2458001 | 35.1920778 | 43.2531189 | 28.0047773 | 45.2918829 | 43.3553489 |
| 0 | 2.17222069 | 0.60675996 | 1.27215055 | 0 | 0 | 2.70970931 |
| 93.1220642 | 138.479069 | 44.2934772 | 104.316345 | 144.172742 | 45.2918829 | 146.324303 |
| 4.90116127 | 2.17222069 | 1.82027989 | 12.7215055 | 0 | 1.19189166 | 4.06456396 |
| 308.77316 | 929.167401 | 584.309844 | 889.233238 | 753.017345 | 616.207986 | 905.042909 |
| 12.2529032 | 11.4041586 | 9.70815939 | 11.449355 | 17.6326376 | 10.7270249 | 13.5485465 |
| 142.133677 | 223.195676 | 261.513544 | 230.25925 | 357.838821 | 369.486413 | 303.487443 |
| 455.807998 | 275.872028 | 341.605859 | 231.531401 | 408.662306 | 524.432328 | 395.617559 |
| 26.956387 | 46.1596897 | 32.765038 | 71.2404311 | 45.6374148 | 34.564858 | 55.5490408 |
| 4.90116127 | 1.08611035 | 5.46083966 | 7.63290333 | 3.11164192 | 0 | 0 |

| | | | | | | |
|---|---|---|---|---|---|---|
| 66.1656772 | 13.5763793 | 26.0906784 | 19.0822583 | 44.6002009 | 26.2216164 | 39.290785 |
| 1568.37161 | 2921.63683 | 2153.99787 | 2974.288 | 2356.55015 | 2046.47797 | 2524.09422 |
| 22.0552257 | 16.2916552 | 10.9216793 | 7.63290333 | 12.4465677 | 13.1108082 | 13.5485465 |
| 225.453419 | 172.14849 | 133.487192 | 155.202368 | 212.628865 | 225.267523 | 161.227704 |
| 4.90116127 | 6.51666208 | 8.49463947 | 1.27215055 | 14.5209956 | 9.53513324 | 12.1936919 |
| 2.45058064 | 38.5569173 | 12.7419592 | 24.1708605 | 30.0792052 | 26.2216164 | 18.9679652 |
| 296.520257 | 200.930414 | 318.54898 | 245.525057 | 398.290166 | 384.981005 | 285.874332 |
| 2.45058064 | 1.08611035 | 0 | 5.08860222 | 0 | 1.19189166 | 4.06456396 |
| 0 | 5.9736069 | 2.42703985 | 5.08860222 | 2.07442795 | 4.76756662 | 4.06456396 |
| 0 | 1.62916552 | 0 | 2.54430111 | 4.14885589 | 3.57567497 | 1.35485465 |
| 0 | 2.17222069 | 1.21351992 | 1.27215055 | 1.03721397 | 3.57567497 | 1.35485465 |
| 122.529032 | 100.465207 | 121.958752 | 115.7657 | 162.842594 | 206.197256 | 192.389361 |
| 7.35174191 | 1.08611035 | 1.82027989 | 10.1772044 | 4.14885589 | 2.38378331 | 0 |
| 46.5610321 | 34.2124759 | 52.7881167 | 33.0759144 | 36.3024891 | 64.3621494 | 48.7747676 |
| 12.2529032 | 39.6430276 | 29.7312381 | 17.8101078 | 42.5257729 | 23.8378331 | 18.9679652 |
| 17.1540645 | 36.9277518 | 18.2027989 | 36.8923661 | 17.6326376 | 26.2216164 | 23.0325291 |
| 0 | 2.17222069 | 1.82027989 | 1.27215055 | 1.03721397 | 2.38378331 | 4.06456396 |
| 0 | 3.25833104 | 1.82027989 | 0 | 0 | 0 | 0 |
| 1257.14787 | 627.77178 | 1502.94443 | 1396.82131 | 1017.50691 | 1473.17809 | 821.041921 |
| 0 | 1.08611035 | 0 | 2.54430111 | 3.11164192 | 1.19189166 | 1.35485465 |
| 0 | 1.08611035 | 0.60675996 | 1.27215055 | 0 | 2.38378331 | 1.35485465 |
| 0 | 1.62916552 | 1.21351992 | 2.54430111 | 1.03721397 | 2.38378331 | 1.35485465 |
| 426.401031 | 117.299917 | 66.7435958 | 125.942905 | 88.1631878 | 53.6351245 | 78.58157 |
| 443.555095 | 369.820573 | 362.235697 | 358.746456 | 415.922803 | 405.243163 | 360.391338 |
| 17.1540645 | 5.43055173 | 4.24731974 | 6.36075277 | 3.11164192 | 2.38378331 | 1.35485465 |
| 98.0232254 | 178.122097 | 137.127751 | 153.930217 | 172.17752 | 152.562132 | 193.744216 |
| 31.8575483 | 7.05971725 | 18.8095588 | 13.9936561 | 14.5209956 | 14.3026999 | 20.3228198 |
| 7.35174191 | 1.62916552 | 4.8540797 | 7.63290333 | 3.11164192 | 3.57567497 | 2.70970931 |
| 0 | 0 | 0 | 0 | 0 | 0 | 1.35485465 |
| 0 | 0 | 0 | 1.27215055 | 2.07442795 | 2.38378331 | 1.35485465 |

| | | | | | | |
|---|---|---|---|---|---|---|
| 1007.18864 | 223.738731 | 447.182092 | 304.043983 | 454.29972 | 483.908012 | 407.811251 |
| 7.35174191 | 1.62916552 | 4.24731974 | 5.08860222 | 2.07442795 | 0 | 1.35485465 |
| 2982.35663 | 1859.96397 | 2833.56902 | 1938.75745 | 4523.29014 | 2698.44271 | 5263.61033 |
| 12.2529032 | 7.60277242 | 9.10139943 | 12.7215055 | 13.4837817 | 17.8783748 | 10.8388372 |
| 26.956387 | 13.0333242 | 15.775759 | 12.7215055 | 28.0047773 | 11.9189166 | 10.8388372 |
| 0 | 1.62916552 | 2.42703985 | 3.81645166 | 5.18606987 | 1.19189166 | 0 |
| 967.979351 | 440.417745 | 580.669284 | 618.265169 | 395.178524 | 560.189078 | 429.488925 |
| 401.895224 | 447.477463 | 393.180455 | 634.803127 | 175.289162 | 212.156715 | 250.648111 |
| 200.947612 | 27.6958138 | 42.4731974 | 73.7847322 | 53.9351266 | 23.8378331 | 32.5165117 |
| 0 | 1.08611035 | 6.67435958 | 1.27215055 | 3.11164192 | 0 | 6.77427327 |
| 0 | 0 | 0.60675996 | 1.27215055 | 1.03721397 | 5.95945828 | 1.35485465 |
| 7.35174191 | 21.7222069 | 21.2365987 | 17.8101078 | 16.5954236 | 19.0702665 | 10.8388372 |
| 17.1540645 | 0.54305517 | 6.67435958 | 2.54430111 | 9.33492576 | 10.7270249 | 4.06456396 |
| 2.45058064 | 1.62916552 | 1.21351992 | 5.08860222 | 3.11164192 | 2.38378331 | 0 |
| 4.90116127 | 2.17222069 | 1.21351992 | 1.27215055 | 1.03721397 | 3.57567497 | 1.35485465 |
| 0 | 1.08611035 | 0 | 0 | 1.03721397 | 1.19189166 | 2.70970931 |
| 0 | 0.54305517 | 0.60675996 | 0 | 2.07442795 | 2.38378331 | 0 |
| 17.1540645 | 9.77499312 | 12.1351992 | 7.63290333 | 16.5954236 | 19.0702665 | 10.8388372 |
| 2.45058064 | 6.51666208 | 6.67435958 | 6.36075277 | 2.07442795 | 3.57567497 | 8.12912793 |
| 4.90116127 | 0.54305517 | 0.60675996 | 1.27215055 | 0 | 1.19189166 | 0 |
| 2.45058064 | 1.62916552 | 0 | 0 | 0 | 0 | 2.70970931 |
| 83.3197416 | 106.981869 | 162.61167 | 145.025163 | 179.438017 | 202.621581 | 199.163634 |
| 0 | 1.62916552 | 0.60675996 | 2.54430111 | 0 | 2.38378331 | 0 |
| 4.90116127 | 2.17222069 | 6.06759962 | 5.08860222 | 4.14885589 | 4.76756662 | 6.77427327 |
| 333.278967 | 664.156477 | 396.821015 | 582.644954 | 496.825493 | 750.891743 | 667.943345 |
| 0 | 1.08611035 | 1.21351992 | 0 | 1.03721397 | 0 | 0 |
| 73.5174191 | 27.1527587 | 43.6867173 | 26.7151616 | 39.414131 | 30.989183 | 44.7102036 |
| 296.520257 | 174.320711 | 239.063425 | 202.271938 | 304.940908 | 423.121538 | 329.229681 |
| 9.80232254 | 2.71527587 | 2.42703985 | 2.54430111 | 3.11164192 | 0 | 1.35485465 |
| 0 | 1.08611035 | 0.60675996 | 5.08860222 | 4.14885589 | 2.38378331 | 6.77427327 |

| | | | | | | |
|---|---|---|---|---|---|---|
| 34.3081289 | 77.1138346 | 37.6191177 | 41.9809683 | 57.0467685 | 16.6864832 | 101.614099 |
| 4.90116127 | 0.54305517 | 1.21351992 | 1.27215055 | 0 | 0 | 2.70970931 |
| 0 | 0.54305517 | 0 | 0 | 1.03721397 | 2.38378331 | 1.35485465 |
| 85.7703223 | 16.2916552 | 49.1475569 | 94.139141 | 78.828262 | 59.5945828 | 40.6456396 |
| 4.90116127 | 10.3180483 | 7.88787951 | 6.36075277 | 3.11164192 | 4.76756662 | 10.8388372 |
| 4.90116127 | 8.1458276 | 13.3487192 | 3.81645166 | 12.4465677 | 14.3026999 | 12.1936919 |
| 0 | 0.54305517 | 0.60675996 | 2.54430111 | 1.03721397 | 1.19189166 | 0 |
| 0 | 0.54305517 | 0.60675996 | 2.54430111 | 1.03721397 | 1.19189166 | 0 |
| 24.5058064 | 5.9736069 | 12.1351992 | 10.1772044 | 11.4093537 | 10.7270249 | 4.06456396 |
| 171.540645 | 229.712338 | 213.579507 | 162.835271 | 290.419913 | 296.781022 | 276.39035 |
| 0 | 0 | 0 | 1.27215055 | 3.11164192 | 1.19189166 | 2.70970931 |
| 7.35174191 | 10.8611035 | 3.64055977 | 8.90505388 | 6.22328384 | 5.95945828 | 2.70970931 |
| 0 | 0 | 0 | 1.27215055 | 1.03721397 | 0 | 0 |
| 34.3081289 | 1.08611035 | 7.28111955 | 7.63290333 | 7.26049782 | 9.53513324 | 4.06456396 |
| 303.871999 | 638.632884 | 436.260413 | 461.790651 | 291.457127 | 344.456688 | 420.004943 |
| 2.45058064 | 8.1458276 | 4.24731974 | 12.7215055 | 0 | 8.34324159 | 8.12912793 |
| 286.717934 | 79.2860553 | 138.341271 | 90.3226894 | 174.251948 | 176.399965 | 149.034012 |
| 0 | 7.05971725 | 6.67435958 | 8.90505388 | 3.11164192 | 1.19189166 | 0 |
| 34.3081289 | 67.3388415 | 36.4055977 | 43.2531189 | 33.1908472 | 27.4135081 | 51.4844769 |
| 0 | 4.88749656 | 4.8540797 | 5.08860222 | 13.4837817 | 5.95945828 | 6.77427327 |
| 127.430193 | 61.3652346 | 117.104673 | 119.582152 | 86.0887598 | 95.3513324 | 71.8072967 |
| 0 | 0.54305517 | 1.21351992 | 0 | 3.11164192 | 0 | 0 |
| 31.8575483 | 21.7222069 | 25.4839184 | 27.9873122 | 31.1164192 | 21.4540498 | 8.12912793 |
| 14.7034838 | 28.7819242 | 19.4163188 | 45.79742 | 18.6698515 | 5.95945828 | 18.9679652 |
| 22.0552257 | 15.7486 | 20.0230788 | 35.6202155 | 19.7070655 | 19.0702665 | 18.9679652 |
| 0 | 0 | 0.60675996 | 5.08860222 | 0 | 1.19189166 | 1.35485465 |
| 563.633546 | 249.80538 | 484.80121 | 357.474306 | 678.337939 | 410.010729 | 700.459856 |
| 14.7034838 | 1.08611035 | 6.06759962 | 6.36075277 | 2.07442795 | 5.95945828 | 4.06456396 |
| 0 | 0.54305517 | 0 | 1.27215055 | 1.03721397 | 0 | 0 |
| 12.2529032 | 25.5235931 | 23.6636385 | 22.89871 | 29.0419913 | 25.0297248 | 25.7422384 |

| | | | | | | |
|---|---|---|---|---|---|---|
| 46.5610321 | 11.4041586 | 23.6636385 | 10.1772044 | 14.5209956 | 32.1810747 | 13.5485465 |
| 0 | 2.71527587 | 3.64055977 | 0 | 13.4837817 | 4.76756662 | 2.70970931 |
| 4.90116127 | 7.60277242 | 4.24731974 | 7.63290333 | 7.26049782 | 16.6864832 | 12.1936919 |
| 4.90116127 | 0 | 0.60675996 | 0 | 1.03721397 | 0 | 2.70970931 |
| 4.90116127 | 3.25833104 | 4.8540797 | 2.54430111 | 4.14885589 | 2.38378331 | 1.35485465 |
| 0 | 0.54305517 | 0 | 0 | 1.03721397 | 0 | 0 |
| 0 | 0.54305517 | 0.60675996 | 1.27215055 | 1.03721397 | 1.19189166 | 4.06456396 |
| 4.90116127 | 0.54305517 | 0 | 0 | 2.07442795 | 0 | 1.35485465 |
| 4.90116127 | 3.25833104 | 6.06759962 | 10.1772044 | 4.14885589 | 8.34324159 | 9.48398258 |
| 2.45058064 | 0 | 0.60675996 | 1.27215055 | 2.07442795 | 0 | 5.41941862 |
| 0 | 4.34444139 | 9.70815939 | 2.54430111 | 4.14885589 | 2.38378331 | 4.06456396 |
| 2.45058064 | 1.08611035 | 1.21351992 | 0 | 1.03721397 | 0 | 0 |
| 132.331354 | 46.7027449 | 104.969473 | 124.670754 | 116.167965 | 121.572949 | 86.7106979 |
| 2.45058064 | 7.05971725 | 12.7419592 | 19.0822583 | 6.22328384 | 11.9189166 | 6.77427327 |
| 0 | 1.08611035 | 1.21351992 | 0 | 3.11164192 | 1.19189166 | 1.35485465 |
| 0 | 1.08611035 | 1.21351992 | 1.27215055 | 0 | 1.19189166 | 0 |
| 2.45058064 | 7.60277242 | 3.64055977 | 5.08860222 | 10.3721397 | 3.57567497 | 2.70970931 |
| 7.35174191 | 7.60277242 | 3.03379981 | 3.81645166 | 5.18606987 | 2.38378331 | 2.70970931 |
| 2747.10089 | 666.871753 | 1048.48121 | 868.878829 | 915.859939 | 958.280891 | 720.782676 |
| 17.1540645 | 10.3180483 | 3.64055977 | 5.08860222 | 11.4093537 | 4.76756662 | 5.41941862 |
| 9.80232254 | 10.8611035 | 7.28111955 | 3.81645166 | 6.22328384 | 13.1108082 | 10.8388372 |
| 49.0116127 | 118.386028 | 121.351992 | 166.651723 | 149.358812 | 150.178349 | 196.453925 |
| 0 | 0 | 1.21351992 | 2.54430111 | 0 | 1.19189166 | 0 |
| 2.45058064 | 0 | 0.60675996 | 1.27215055 | 1.03721397 | 0 | 1.35485465 |
| 2.45058064 | 0 | 1.21351992 | 0 | 1.03721397 | 0 | 1.35485465 |
| 7.35174191 | 0.54305517 | 0 | 2.54430111 | 1.03721397 | 0 | 0 |
| 0 | 0.54305517 | 0.60675996 | 0 | 0 | 3.57567497 | 1.35485465 |
| 4.90116127 | 4.34444139 | 5.46083966 | 13.9936561 | 3.11164192 | 9.53513324 | 2.70970931 |
| 4.90116127 | 0.54305517 | 0 | 2.54430111 | 4.14885589 | 3.57567497 | 1.35485465 |
| 0 | 1.08611035 | 0 | 0 | 0 | 0 | 1.35485465 |

| | | | | | | |
|---|---|---|---|---|---|---|
| 17.1540645 | 36.3846966 | 38.8326376 | 76.3290333 | 25.9303493 | 28.6053997 | 14.9034012 |
| 2.45058064 | 0 | 1.82027989 | 3.81645166 | 2.07442795 | 0 | 0 |
| 2.45058064 | 1.62916552 | 1.21351992 | 2.54430111 | 1.03721397 | 2.38378331 | 1.35485465 |
| 4.90116127 | 1.62916552 | 0.60675996 | 1.27215055 | 3.11164192 | 4.76756662 | 2.70970931 |
| 2.45058064 | 1.08611035 | 1.21351992 | 1.27215055 | 2.07442795 | 1.19189166 | 2.70970931 |
| 0 | 0.54305517 | 0.60675996 | 1.27215055 | 1.03721397 | 1.19189166 | 1.35485465 |
| 4.90116127 | 3.25833104 | 4.8540797 | 7.63290333 | 5.18606987 | 1.19189166 | 2.70970931 |
| 4.90116127 | 0.54305517 | 1.82027989 | 1.27215055 | 0 | 0 | 0 |
| 2.45058064 | 3.80138621 | 3.03379981 | 1.27215055 | 1.03721397 | 7.15134993 | 4.06456396 |
| 4.90116127 | 2.17222069 | 2.42703985 | 5.08860222 | 5.18606987 | 4.76756662 | 2.70970931 |
| 4.90116127 | 0 | 0.60675996 | 1.27215055 | 1.03721397 | 1.19189166 | 0 |
| 0 | 0 | 1.21351992 | 0 | 0 | 0 | 0 |
| 56.3633546 | 44.5305242 | 41.8664374 | 17.8101078 | 47.7118428 | 66.7459327 | 48.7747676 |
| 0 | 1.62916552 | 0.60675996 | 0 | 0 | 0 | 0 |
| 245.058064 | 117.842973 | 200.230788 | 176.828927 | 360.950463 | 300.356697 | 280.454913 |
| 0 | 4.88749656 | 3.03379981 | 10.1772044 | 2.07442795 | 1.19189166 | 4.06456396 |
| 78.4185804 | 25.5235931 | 34.5853178 | 20.3544089 | 85.0515458 | 33.3729664 | 77.2267153 |
| 7.35174191 | 2.17222069 | 12.7419592 | 7.63290333 | 20.7442795 | 8.34324159 | 6.77427327 |
| 31.8575483 | 45.0735794 | 26.0906784 | 35.6202155 | 53.9351266 | 61.9783661 | 51.4844769 |
| 7.35174191 | 14.6624897 | 12.1351992 | 20.3544089 | 9.33492576 | 7.15134993 | 10.8388372 |
| 17.1540645 | 15.2055448 | 10.9216793 | 5.08860222 | 17.6326376 | 21.4540498 | 18.9679652 |
| 0 | 2.17222069 | 0 | 0 | 2.07442795 | 0 | 1.35485465 |
| 0 | 2.17222069 | 4.8540797 | 0 | 1.03721397 | 1.19189166 | 0 |
| 34.3081289 | 110.2402 | 35.1920778 | 21.6265594 | 58.0839825 | 57.2107995 | 105.678663 |
| 2.45058064 | 2.17222069 | 1.82027989 | 0 | 0 | 0 | 0 |
| 0 | 0 | 0 | 0 | 1.03721397 | 0 | 0 |
| 2.45058064 | 0.54305517 | 1.82027989 | 2.54430111 | 1.03721397 | 1.19189166 | 1.35485465 |
| 0 | 0.54305517 | 0.60675996 | 0 | 2.07442795 | 1.19189166 | 0 |
| 154.38658 | 116.756862 | 89.8004744 | 122.126453 | 106.833039 | 119.189166 | 69.0975874 |
| 1012.0898 | 3063.37423 | 1658.88174 | 1667.78938 | 1416.83429 | 1897.49152 | 2590.4821 |

| | | | | | | |
|---|---|---|---|---|---|---|
| 4.90116127 | 1.08611035 | 1.21351992 | 1.27215055 | 2.07442795 | 2.38378331 | 1.35485465 |
| 2215.32489 | 1524.89893 | 2396.70185 | 2170.28885 | 2057.83252 | 2581.63733 | 2067.5082 |
| 44.1104514 | 76.0277242 | 41.8664374 | 26.7151616 | 59.1211965 | 46.4837746 | 50.1296222 |
| 7.35174191 | 4.88749656 | 1.82027989 | 5.08860222 | 5.18606987 | 1.19189166 | 2.70970931 |
| 4.90116127 | 5.9736069 | 4.24731974 | 8.90505388 | 7.26049782 | 3.57567497 | 1.35485465 |
| 4.90116127 | 0.54305517 | 0.60675996 | 1.27215055 | 0 | 0 | 0 |
| 2.45058064 | 0.54305517 | 3.64055977 | 1.27215055 | 0 | 3.57567497 | 1.35485465 |
| 2.45058064 | 3.25833104 | 4.24731974 | 10.1772044 | 9.33492576 | 5.95945828 | 8.12912793 |
| 17.1540645 | 4.34444139 | 7.28111955 | 3.81645166 | 6.22328384 | 2.38378331 | 5.41941862 |
| 2.45058064 | 2.17222069 | 3.03379981 | 0 | 2.07442795 | 2.38378331 | 8.12912793 |
| 2.45058064 | 0.54305517 | 1.82027989 | 2.54430111 | 1.03721397 | 1.19189166 | 1.35485465 |
| 2.45058064 | 0.54305517 | 1.82027989 | 2.54430111 | 1.03721397 | 1.19189166 | 1.35485465 |
| 9.80232254 | 12.490269 | 7.28111955 | 20.3544089 | 10.3721397 | 8.34324159 | 9.48398258 |
| 0 | 1.08611035 | 1.82027989 | 1.27215055 | 0 | 1.19189166 | 0 |
| 9.80232254 | 6.51666208 | 10.3149194 | 7.63290333 | 8.29771179 | 7.15134993 | 14.9034012 |
| 0 | 0.54305517 | 0 | 0 | 2.07442795 | 1.19189166 | 0 |
| 0 | 0 | 0.60675996 | 0 | 1.03721397 | 0 | 0 |
| 14.7034838 | 24.4374828 | 20.6298387 | 43.2531189 | 18.6698515 | 15.4945915 | 28.4519477 |
| 2.45058064 | 0 | 0 | 0 | 1.03721397 | 0 | 0 |
| 0 | 1.62916552 | 1.82027989 | 2.54430111 | 0 | 1.19189166 | 2.70970931 |
| 2.45058064 | 0 | 1.82027989 | 3.81645166 | 0 | 0 | 0 |
| 2.45058064 | 2.17222069 | 2.42703985 | 0 | 6.22328384 | 5.95945828 | 0 |
| 0 | 1.08611035 | 1.21351992 | 3.81645166 | 1.03721397 | 3.57567497 | 0 |
| 66.1656772 | 82.0013311 | 174.746869 | 169.196024 | 85.0515458 | 91.7756575 | 121.936919 |
| 0 | 1.08611035 | 0 | 0 | 0 | 0 | 2.70970931 |
| 0 | 1.08611035 | 0.60675996 | 0 | 0 | 2.38378331 | 2.70970931 |
| 4.90116127 | 5.43055173 | 1.21351992 | 1.27215055 | 1.03721397 | 2.38378331 | 4.06456396 |
| 4.90116127 | 1.08611035 | 1.82027989 | 5.08860222 | 2.07442795 | 2.38378331 | 1.35485465 |
| 39.2092902 | 32.0402552 | 40.6529175 | 47.0695705 | 26.9675633 | 35.7567497 | 23.0325291 |
| 0 | 2.17222069 | 0.60675996 | 0 | 2.07442795 | 0 | 2.70970931 |

| | | | | | | |
|---|---|---|---|---|---|---|
| 0 | 6.51666208 | 5.46083966 | 15.2658067 | 8.29771179 | 15.4945915 | 1.35485465 |
| 2.45058064 | 3.80138621 | 3.03379981 | 5.08860222 | 5.18606987 | 0 | 2.70970931 |
| 2.45058064 | 7.60277242 | 0 | 11.449355 | 2.07442795 | 2.38378331 | 1.35485465 |
| 0 | 0.54305517 | 0 | 0 | 0 | 0 | 4.06456396 |
| 6457.27998 | 5487.57252 | 4268.55633 | 3738.85048 | 5584.36003 | 6946.34457 | 9543.59619 |
| 0 | 0.54305517 | 0 | 1.27215055 | 0 | 2.38378331 | 0 |
| 0 | 0 | 1.21351992 | 1.27215055 | 1.03721397 | 0 | 1.35485465 |
| 4.90116127 | 4.88749656 | 3.64055977 | 5.08860222 | 7.26049782 | 4.76756662 | 6.77427327 |
| 12.2529032 | 11.9472138 | 6.06759962 | 6.36075277 | 8.29771179 | 5.95945828 | 5.41941862 |
| 406.796386 | 161.287386 | 86.1599146 | 133.575808 | 162.842594 | 109.654032 | 170.711686 |
| 0 | 2.71527587 | 2.42703985 | 2.54430111 | 3.11164192 | 1.19189166 | 2.70970931 |
| 0 | 1.08611035 | 0 | 1.27215055 | 1.03721397 | 0 | 1.35485465 |
| 7.35174191 | 4.34444139 | 4.24731974 | 5.08860222 | 7.26049782 | 2.38378331 | 0 |
| 0 | 1.08611035 | 0.60675996 | 0 | 0 | 0 | 1.35485465 |
| 0 | 0 | 0.60675996 | 0 | 2.07442795 | 0 | 1.35485465 |
| 2943.14734 | 1418.46011 | 1359.14232 | 1158.92916 | 1313.11289 | 1588.79158 | 1707.11686 |
| 73.5174191 | 218.30818 | 85.5531547 | 81.4176355 | 122.391249 | 226.459415 | 67.7427327 |
| 776.834062 | 1108.37561 | 1159.51829 | 1078.78367 | 913.785511 | 1379.01865 | 1261.36968 |
| 0 | 0.54305517 | 1.21351992 | 0 | 0 | 2.38378331 | 0 |
| 4.90116127 | 2.17222069 | 7.28111955 | 2.54430111 | 3.11164192 | 4.76756662 | 2.70970931 |
| 41.6598708 | 117.842973 | 100.115394 | 138.66441 | 46.6746288 | 64.3621494 | 50.1296222 |
| 259.761547 | 455.62329 | 411.383254 | 534.303233 | 517.569773 | 414.778296 | 597.490903 |
| 0 | 1.08611035 | 0.60675996 | 1.27215055 | 1.03721397 | 0 | 0 |
| 17.1540645 | 43.4444139 | 25.4839184 | 20.3544089 | 35.2652751 | 29.7972914 | 23.0325291 |
| 0 | 2.17222069 | 2.42703985 | 1.27215055 | 1.03721397 | 3.57567497 | 5.41941862 |
| 4.90116127 | 5.43055173 | 10.9216793 | 3.81645166 | 6.22328384 | 4.76756662 | 13.5485465 |
| 24.5058064 | 65.1666208 | 40.6529175 | 44.5252694 | 50.8234847 | 73.8972826 | 63.6781688 |
| 0 | 1.08611035 | 1.21351992 | 3.81645166 | 2.07442795 | 1.19189166 | 4.06456396 |
| 622.447482 | 656.010649 | 575.815204 | 623.353772 | 516.532559 | 648.389061 | 606.974885 |
| 2.45058064 | 1.62916552 | 0.60675996 | 0 | 1.03721397 | 0 | 0 |

| | | | | | | |
|---|---|---|---|---|---|---|
| 14.7034838 | 4.34444139 | 5.46083966 | 5.08860222 | 4.14885589 | 2.38378331 | 2.70970931 |
| 2.45058064 | 0.54305517 | 0.60675996 | 1.27215055 | 2.07442795 | 1.19189166 | 4.06456396 |
| 0 | 1.08611035 | 0.60675996 | 1.27215055 | 1.03721397 | 1.19189166 | 0 |
| 0 | 0.54305517 | 0.60675996 | 1.27215055 | 0 | 1.19189166 | 1.35485465 |
| 9.80232254 | 30.9541449 | 13.9554791 | 13.9936561 | 13.4837817 | 9.53513324 | 9.48398258 |
| 12.2529032 | 11.4041586 | 21.8433586 | 11.449355 | 22.8187074 | 13.1108082 | 16.2582559 |
| 0 | 0 | 0 | 0 | 1.03721397 | 0 | 0 |
| 4.90116127 | 0.54305517 | 1.82027989 | 0 | 1.03721397 | 0 | 5.41941862 |
| 4.90116127 | 22.8083173 | 18.2027989 | 19.0822583 | 7.26049782 | 13.1108082 | 6.77427327 |
| 0 | 1.08611035 | 1.82027989 | 1.27215055 | 0 | 0 | 0 |
| 4.90116127 | 5.9736069 | 6.67435958 | 7.63290333 | 8.29771179 | 11.9189166 | 8.12912793 |
| 4.90116127 | 13.5763793 | 11.5284393 | 20.3544089 | 9.33492576 | 14.3026999 | 13.5485465 |
| 22.0552257 | 2.71527587 | 6.67435958 | 10.1772044 | 6.22328384 | 10.7270249 | 8.12912793 |
| 2.45058064 | 3.25833104 | 2.42703985 | 1.27215055 | 3.11164192 | 3.57567497 | 6.77427327 |
| 0 | 0 | 0 | 1.27215055 | 1.03721397 | 0 | 1.35485465 |
| 757.229417 | 916.677132 | 1078.81921 | 1571.10593 | 702.19386 | 777.113359 | 726.202095 |
| 0 | 0 | 0 | 1.27215055 | 0 | 3.57567497 | 0 |
| 0 | 0 | 0 | 1.27215055 | 0 | 0 | 4.06456396 |
| 0 | 0.54305517 | 0.60675996 | 2.54430111 | 1.03721397 | 0 | 0 |
| 2.45058064 | 0.54305517 | 0 | 0 | 0 | 0 | 1.35485465 |
| 1553.66812 | 3234.97967 | 3621.75021 | 3531.48994 | 5679.78372 | 4589.97477 | 6951.75923 |
| 0 | 0 | 0 | 1.27215055 | 0 | 0 | 1.35485465 |
| 7.35174191 | 29.8680345 | 31.551518 | 27.9873122 | 29.0419913 | 20.2621581 | 33.8713664 |
| 26.956387 | 56.477738 | 25.4839184 | 58.5189255 | 21.7814934 | 15.4945915 | 9.48398258 |
| 122.529032 | 139.565179 | 129.239872 | 75.0568827 | 98.5353275 | 97.7351157 | 139.550029 |
| 0 | 0.54305517 | 0.60675996 | 1.27215055 | 1.03721397 | 0 | 0 |
| 2.45058064 | 8.68888277 | 4.8540797 | 6.36075277 | 9.33492576 | 9.53513324 | 6.77427327 |
| 0 | 0 | 0 | 0 | 0 | 0 | 0 |
| 8033.00332 | 7684.77376 | 7154.30671 | 6187.7403 | 8030.11058 | 10774.7006 | 14079.6496 |
| 0 | 0 | 0 | 0 | 0 | 1.19189166 | 1.35485465 |

| | | | | | | |
|---|---|---|---|---|---|---|
| 191.14529 | 342.667814 | 379.224976 | 503.77162 | 179.438017 | 220.499956 | 253.35782 |
| 360.235353 | 145.538786 | 292.458302 | 178.101078 | 214.703293 | 235.994548 | 215.42189 |
| 7.35174191 | 1.08611035 | 1.82027989 | 0 | 1.03721397 | 5.95945828 | 0 |
| 0 | 1.08611035 | 0 | 0 | 0 | 0 | 0 |
| 0 | 0.54305517 | 0.60675996 | 1.27215055 | 1.03721397 | 2.38378331 | 0 |
| 0 | 2.17222069 | 0.60675996 | 3.81645166 | 3.11164192 | 4.76756662 | 6.77427327 |
| 0 | 3.25833104 | 1.82027989 | 8.90505388 | 0 | 0 | 1.35485465 |
| 2.45058064 | 0 | 0.60675996 | 0 | 0 | 0 | 0 |
| 36.7587095 | 37.4708069 | 36.4055977 | 27.9873122 | 41.4885589 | 39.3324246 | 50.1296222 |
| 4.90116127 | 3.25833104 | 2.42703985 | 5.08860222 | 0 | 1.19189166 | 2.70970931 |
| 0 | 0.54305517 | 0.60675996 | 1.27215055 | 0 | 0 | 0 |
| 2725.04567 | 1964.23056 | 3059.28373 | 2282.23809 | 4954.77115 | 3766.37763 | 5640.25993 |
| 95.5726448 | 201.473469 | 165.03871 | 216.265594 | 206.405581 | 177.591857 | 138.195175 |
| 0 | 0.54305517 | 2.42703985 | 6.36075277 | 4.14885589 | 0 | 0 |
| 26.956387 | 17.3777655 | 25.4839184 | 33.0759144 | 26.9675633 | 20.2621581 | 10.8388372 |
| 19.6046451 | 35.8416414 | 33.9785579 | 26.7151616 | 34.2280611 | 28.6053997 | 35.226221 |
| 0 | 1.62916552 | 0 | 1.27215055 | 1.03721397 | 0 | 0 |
| 22.0552257 | 9.77499312 | 13.3487192 | 24.1708605 | 3.11164192 | 5.95945828 | 16.2582559 |
| 4.90116127 | 2.17222069 | 1.82027989 | 1.27215055 | 1.03721397 | 1.19189166 | 1.35485465 |
| 0 | 1.08611035 | 0.60675996 | 1.27215055 | 0 | 3.57567497 | 4.06456396 |
| 0 | 0 | 0 | 0 | 0 | 1.19189166 | 1.35485465 |
| 1639.43845 | 2826.60218 | 1626.72346 | 1434.98583 | 1331.78274 | 1812.86721 | 2560.6753 |
| 0 | 0 | 0.60675996 | 0 | 0 | 2.38378331 | 0 |
| 34.3081289 | 35.2985863 | 34.5853178 | 45.79742 | 24.8931354 | 29.7972914 | 17.6131105 |
| 7.35174191 | 4.34444139 | 4.24731974 | 3.81645166 | 2.07442795 | 4.76756662 | 5.41941862 |
| 2.45058064 | 2.71527587 | 2.42703985 | 3.81645166 | 1.03721397 | 1.19189166 | 0 |
| 2.45058064 | 0 | 0 | 1.27215055 | 1.03721397 | 2.38378331 | 0 |
| 0 | 0.54305517 | 0 | 0 | 0 | 0 | 0 |
| 0 | 2.17222069 | 0 | 3.81645166 | 1.03721397 | 0 | 1.35485465 |
| 9.80232254 | 14.6624897 | 4.24731974 | 11.449355 | 5.18606987 | 8.34324159 | 8.12912793 |

| | | | | | | |
|---|---|---|---|---|---|---|
| 53.912774 | 24.4374828 | 31.551518 | 35.6202155 | 25.9303493 | 14.3026999 | 25.7422384 |
| 4.90116127 | 2.17222069 | 2.42703985 | 2.54430111 | 1.03721397 | 1.19189166 | 4.06456396 |
| 49.0116127 | 11.9472138 | 18.2027989 | 17.8101078 | 9.33492576 | 7.15134993 | 16.2582559 |
| 34.3081289 | 22.8083173 | 37.6191177 | 20.3544089 | 47.7118428 | 52.4432328 | 54.1941862 |
| 14.7034838 | 15.2055448 | 20.0230788 | 10.1772044 | 13.4837817 | 8.34324159 | 10.8388372 |
| 22.0552257 | 7.60277242 | 10.9216793 | 17.8101078 | 8.29771179 | 7.15134993 | 8.12912793 |
| 0 | 0 | 0 | 1.27215055 | 0 | 0 | 1.35485465 |
| 0 | 1.62916552 | 1.21351992 | 0 | 1.03721397 | 0 | 1.35485465 |
| 12.2529032 | 11.9472138 | 10.9216793 | 16.5379572 | 7.26049782 | 10.7270249 | 12.1936919 |
| 0 | 3.25833104 | 0.60675996 | 2.54430111 | 0 | 0 | 0 |
| 0 | 0 | 0.60675996 | 0 | 0 | 0 | 0 |
| 0 | 0 | 0 | 0 | 0 | 0 | 0 |
| 7.35174191 | 2.17222069 | 1.21351992 | 0 | 1.03721397 | 0 | 5.41941862 |
| 17.1540645 | 59.1930139 | 70.9909156 | 87.7783883 | 40.451345 | 57.2107995 | 58.2587501 |
| 17.1540645 | 25.5235931 | 26.0906784 | 40.7088177 | 10.3721397 | 15.4945915 | 6.77427327 |
| 147.034838 | 99.9221519 | 40.0461575 | 48.3417211 | 87.1259738 | 76.2810659 | 105.678663 |
| 0 | 0.54305517 | 0 | 1.27215055 | 2.07442795 | 0 | 1.35485465 |
| 14.7034838 | 4.34444139 | 3.64055977 | 10.1772044 | 2.07442795 | 4.76756662 | 2.70970931 |
| 0 | 0 | 0 | 0 | 0 | 0 | 0 |
| 0 | 0.54305517 | 0 | 0 | 0 | 1.19189166 | 0 |
| 0 | 1.08611035 | 0.60675996 | 0 | 0 | 2.38378331 | 0 |
| 0 | 1.62916552 | 1.21351992 | 1.27215055 | 4.14885589 | 2.38378331 | 0 |
| 2.45058064 | 10.8611035 | 6.06759962 | 11.449355 | 3.11164192 | 1.19189166 | 9.48398258 |
| 2.45058064 | 0.54305517 | 0 | 0 | 0 | 0 | 0 |
| 0 | 1.08611035 | 0 | 1.27215055 | 2.07442795 | 0 | 5.41941862 |
| 0 | 2.17222069 | 0 | 1.27215055 | 1.03721397 | 0 | 0 |
| 2.45058064 | 0.54305517 | 0 | 1.27215055 | 0 | 0 | 0 |
| 100.473806 | 229.712338 | 184.455028 | 221.354196 | 105.795825 | 168.056723 | 182.905378 |
| 0 | 0 | 0 | 0 | 0 | 0 | 0 |
| 12.2529032 | 16.8347104 | 5.46083966 | 6.36075277 | 8.29771179 | 7.15134993 | 8.12912793 |

| | | | | | | |
|---|---|---|---|---|---|---|
| 0 | 7.60277242 | 3.03379981 | 1.27215055 | 1.03721397 | 2.38378331 | 4.06456396 |
| 0 | 0.54305517 | 0.60675996 | 0 | 0 | 0 | 1.35485465 |
| 4.90116127 | 0.54305517 | 1.82027989 | 1.27215055 | 3.11164192 | 0 | 4.06456396 |
| 0 | 0 | 0 | 0 | 0 | 0 | 0 |
| 2.45058064 | 0.54305517 | 0 | 0 | 1.03721397 | 0 | 0 |
| 0 | 0.54305517 | 0 | 0 | 0 | 0 | 0 |
| 0 | 0.54305517 | 0 | 2.54430111 | 1.03721397 | 1.19189166 | 0 |
| 0 | 3.25833104 | 1.82027989 | 6.36075277 | 0 | 1.19189166 | 6.77427327 |
| 4376.73702 | 6843.58129 | 4634.43259 | 5734.8547 | 4464.16894 | 5291.99895 | 7028.98595 |
| 0 | 0 | 1.21351992 | 0 | 0 | 2.38378331 | 0 |
| 0 | 0.54305517 | 0 | 0 | 0 | 0 | 0 |
| 2.45058064 | 0 | 0.60675996 | 0 | 0 | 0 | 0 |
| 1367.42399 | 994.877077 | 705.055076 | 941.39141 | 495.788279 | 682.953919 | 769.557444 |
| 0 | 0 | 0.60675996 | 1.27215055 | 1.03721397 | 0 | 0 |
| 0 | 2.17222069 | 0 | 2.54430111 | 1.03721397 | 0 | 1.35485465 |
| 0 | 1.08611035 | 0.60675996 | 3.81645166 | 0 | 0 | 1.35485465 |
| 4.90116127 | 0 | 0.60675996 | 0 | 0 | 0 | 0 |
| 7.35174191 | 1.08611035 | 0.60675996 | 0 | 4.14885589 | 2.38378331 | 4.06456396 |
| 51.4621934 | 6.51666208 | 15.775759 | 10.1772044 | 14.5209956 | 9.53513324 | 12.1936919 |
| 0 | 0.54305517 | 1.21351992 | 0 | 0 | 0 | 0 |
| 0 | 1.08611035 | 0.60675996 | 1.27215055 | 0 | 0 | 0 |
| 0 | 1.08611035 | 0 | 0 | 3.11164192 | 0 | 0 |
| 0 | 0 | 0 | 0 | 2.07442795 | 0 | 1.35485465 |
| 4.90116127 | 0 | 2.42703985 | 0 | 1.03721397 | 1.19189166 | 4.06456396 |
| 29.4069676 | 17.9208207 | 10.3149194 | 5.08860222 | 16.5954236 | 3.57567497 | 14.9034012 |
| 7.35174191 | 0 | 0 | 0 | 0 | 0 | 0 |
| 0 | 0.54305517 | 0 | 0 | 1.03721397 | 0 | 1.35485465 |
| 44.1104514 | 68.968007 | 43.0799573 | 50.8860222 | 53.9351266 | 65.554041 | 71.8072967 |
| 2.45058064 | 3.80138621 | 2.42703985 | 1.27215055 | 1.03721397 | 4.76756662 | 0 |
| 7.35174191 | 4.88749656 | 2.42703985 | 7.63290333 | 6.22328384 | 7.15134993 | 4.06456396 |

| | | | | | | |
|---|---|---|---|---|---|---|
| 0 | 0 | 0 | 0 | 0 | 2.38378331 | 1.35485465 |
| 0 | 0 | 0.60675996 | 3.81645166 | 0 | 0 | 1.35485465 |
| 4.90116127 | 0 | 0 | 0 | 1.03721397 | 0 | 0 |
| 4.90116127 | 0 | 0 | 1.27215055 | 0 | 0 | 0 |
| 0 | 0 | 0 | 1.27215055 | 0 | 0 | 0 |
| 7.35174191 | 0 | 0.60675996 | 0 | 0 | 0 | 0 |
| 637.150965 | 259.580373 | 434.440133 | 326.942693 | 336.057327 | 338.49723 | 440.327763 |
| 0 | 0 | 0 | 0 | 2.07442795 | 1.19189166 | 0 |
| 0 | 1.08611035 | 0 | 0 | 1.03721397 | 1.19189166 | 1.35485465 |
| 0 | 0 | 0 | 0 | 0 | 0 | 0 |
| 0 | 1.62916552 | 0 | 0 | 0 | 0 | 0 |
| 0 | 0.54305517 | 0 | 0 | 4.14885589 | 1.19189166 | 1.35485465 |
| 0 | 0 | 0.60675996 | 1.27215055 | 0 | 0 | 0 |
| 112.726709 | 232.970669 | 201.444307 | 203.544089 | 154.544882 | 185.935098 | 123.291774 |
| 2.45058064 | 0 | 0 | 2.54430111 | 0 | 0 | 1.35485465 |
| 0 | 0 | 0 | 1.27215055 | 1.03721397 | 1.19189166 | 1.35485465 |
| 4.90116127 | 1.62916552 | 1.82027989 | 2.54430111 | 8.29771179 | 3.57567497 | 5.41941862 |
| 0 | 0 | 0 | 0 | 1.03721397 | 0 | 2.70970931 |
| 4.90116127 | 1.08611035 | 2.42703985 | 2.54430111 | 2.07442795 | 3.57567497 | 1.35485465 |
| 39.2092902 | 15.2055448 | 22.4501186 | 15.2658067 | 31.1164192 | 14.3026999 | 14.9034012 |
| 0 | 0 | 0.60675996 | 0 | 1.03721397 | 0 | 0 |
| 0 | 0.54305517 | 1.82027989 | 3.81645166 | 1.03721397 | 3.57567497 | 1.35485465 |
| 0 | 2.17222069 | 2.42703985 | 2.54430111 | 2.07442795 | 0 | 2.70970931 |
| 0 | 0.54305517 | 0 | 0 | 1.03721397 | 0 | 0 |
| 0 | 0.54305517 | 0 | 0 | 0 | 0 | 2.70970931 |
| 0 | 0.54305517 | 0.60675996 | 0 | 1.03721397 | 0 | 0 |
| 0 | 0.54305517 | 0.60675996 | 0 | 1.03721397 | 0 | 0 |
| 0 | 0.54305517 | 0 | 0 | 1.03721397 | 1.19189166 | 1.35485465 |
| 12.2529032 | 7.05971725 | 9.70815939 | 12.7215055 | 5.18606987 | 2.38378331 | 1.35485465 |
| 0 | 4.34444139 | 4.8540797 | 10.1772044 | 5.18606987 | 1.19189166 | 8.12912793 |

| | | | | | | |
|---|---|---|---|---|---|---|
| 0 | 0.54305517 | 1.82027989 | 1.27215055 | 1.03721397 | 0 | 1.35485465 |
| 68.6162578 | 128.704076 | 76.4517552 | 103.044195 | 67.4189083 | 53.6351245 | 46.0650583 |
| 88.2209029 | 2.71527587 | 20.6298387 | 12.7215055 | 26.9675633 | 38.140533 | 20.3228198 |
| 14.7034838 | 45.6166345 | 32.765038 | 26.7151616 | 14.5209956 | 15.4945915 | 25.7422384 |
| 0 | 2.17222069 | 4.8540797 | 7.63290333 | 3.11164192 | 1.19189166 | 1.35485465 |
| 0 | 0.54305517 | 0 | 0 | 1.03721397 | 0 | 1.35485465 |
| 0 | 0.54305517 | 0 | 0 | 0 | 0 | 0 |
| 0 | 0 | 1.82027989 | 2.54430111 | 0 | 1.19189166 | 2.70970931 |
| 12.2529032 | 5.9736069 | 9.10139943 | 17.8101078 | 16.5954236 | 7.15134993 | 8.12912793 |
| 0 | 4.88749656 | 3.64055977 | 6.36075277 | 3.11164192 | 5.95945828 | 5.41941862 |
| 24.5058064 | 40.729138 | 57.0354364 | 48.3417211 | 37.339703 | 67.9378244 | 28.4519477 |
| 0 | 1.08611035 | 0 | 0 | 1.03721397 | 0 | 0 |
| 0 | 0 | 0 | 0 | 0 | 0 | 0 |
| 176.441806 | 109.697145 | 168.679269 | 100.499894 | 154.544882 | 143.026999 | 168.001977 |
| 0 | 0 | 0.60675996 | 0 | 0 | 0 | 0 |
| 0 | 0 | 0 | 0 | 0 | 0 | 0 |
| 0 | 2.17222069 | 0 | 0 | 0 | 0 | 0 |
| 4.90116127 | 2.17222069 | 1.21351992 | 3.81645166 | 5.18606987 | 4.76756662 | 5.41941862 |
| 0 | 2.71527587 | 2.42703985 | 2.54430111 | 6.22328384 | 1.19189166 | 6.77427327 |
| 12.2529032 | 0.54305517 | 3.03379981 | 0 | 1.03721397 | 2.38378331 | 1.35485465 |
| 0 | 0 | 0 | 0 | 0 | 0 | 1.35485465 |
| 0 | 0 | 1.21351992 | 0 | 0 | 0 | 1.35485465 |
| 637.150965 | 1497.20311 | 1018.14322 | 1078.78367 | 958.385712 | 1119.18626 | 1755.89163 |
| 0 | 0 | 0.60675996 | 0 | 0 | 0 | 2.70970931 |
| 0 | 0 | 0 | 0 | 0 | 1.19189166 | 0 |
| 0 | 0 | 0 | 0 | 0 | 0 | 0 |
| 0 | 1.08611035 | 3.03379981 | 2.54430111 | 3.11164192 | 1.19189166 | 5.41941862 |
| 2.45058064 | 2.71527587 | 4.24731974 | 5.08860222 | 6.22328384 | 0 | 2.70970931 |
| 0 | 1.62916552 | 0 | 0 | 1.03721397 | 0 | 0 |
| 333.278967 | 220.4804 | 241.490465 | 250.613659 | 282.122201 | 281.286431 | 224.905873 |

| | | | | | | |
|---|---|---|---|---|---|---|
| 0 | 0 | 0 | 0 | 0 | 0 | 0 |
| 0 | 2.17222069 | 0.60675996 | 0 | 0 | 0 | 0 |
| 0 | 0 | 0 | 0 | 0 | 0 | 0 |
| 0 | 0 | 0.60675996 | 0 | 0 | 0 | 0 |
| 2.45058064 | 1.08611035 | 1.82027989 | 2.54430111 | 1.03721397 | 1.19189166 | 1.35485465 |
| 0 | 0 | 1.82027989 | 0 | 0 | 0 | 0 |
| 0 | 0 | 0.60675996 | 0 | 0 | 1.19189166 | 0 |
| 2.45058064 | 2.71527587 | 0.60675996 | 0 | 0 | 3.57567497 | 1.35485465 |
| 200.947612 | 368.191407 | 123.172272 | 147.569464 | 213.666079 | 261.024273 | 208.647617 |
| 1296.35716 | 2135.29294 | 2654.57483 | 2292.4153 | 2989.25067 | 2889.14537 | 3765.14108 |
| 0 | 0 | 0 | 0 | 0 | 0 | 1.35485465 |
| 0 | 1.08611035 | 0 | 0 | 1.03721397 | 0 | 0 |
| 188.694709 | 415.437207 | 297.919141 | 302.771832 | 332.945686 | 237.186439 | 288.584041 |
| 0 | 0.54305517 | 0 | 1.27215055 | 0 | 0 | 0 |
| 0 | 0 | 0 | 1.27215055 | 0 | 0 | 0 |
| 19.6046451 | 2.71527587 | 7.88787951 | 1.27215055 | 4.14885589 | 9.53513324 | 9.48398258 |
| 14.7034838 | 1.08611035 | 3.64055977 | 0 | 2.07442795 | 0 | 0 |
| 0 | 4.34444139 | 6.06759962 | 3.81645166 | 3.11164192 | 2.38378331 | 1.35485465 |
| 0 | 1.08611035 | 0.60675996 | 0 | 0 | 0 | 0 |
| 0 | 0 | 0 | 1.27215055 | 2.07442795 | 0 | 0 |
| 2.45058064 | 1.62916552 | 1.21351992 | 5.08860222 | 3.11164192 | 8.34324159 | 0 |
| 120.078451 | 119.472138 | 126.206072 | 160.29097 | 124.465677 | 95.3513324 | 105.678663 |
| 9.80232254 | 3.25833104 | 6.67435958 | 3.81645166 | 13.4837817 | 4.76756662 | 9.48398258 |
| 2.45058064 | 11.4041586 | 27.9109583 | 44.5252694 | 15.5582096 | 9.53513324 | 4.06456396 |
| 0 | 1.08611035 | 0 | 1.27215055 | 0 | 0 | 0 |
| 0 | 1.08611035 | 0 | 1.27215055 | 0 | 0 | 0 |
| 0 | 0 | 0.60675996 | 0 | 0 | 0 | 0 |
| 0 | 0 | 0 | 0 | 0 | 0 | 1.35485465 |
| 0 | 0 | 1.82027989 | 0 | 0 | 0 | 2.70970931 |
| 0 | 1.62916552 | 3.03379981 | 6.36075277 | 2.07442795 | 2.38378331 | 8.12912793 |

| | | | | | | |
|---|---|---|---|---|---|---|
| 0 | 0 | 0 | 1.27215055 | 1.03721397 | 0 | 1.35485465 |
| 0 | 0 | 0 | 0 | 0 | 0 | 0 |
| 0 | 0 | 0.60675996 | 0 | 0 | 1.19189166 | 1.35485465 |
| 2.45058064 | 1.08611035 | 1.21351992 | 0 | 1.03721397 | 0 | 0 |
| 0 | 1.08611035 | 3.64055977 | 2.54430111 | 0 | 3.57567497 | 5.41941862 |
| 0 | 1.08611035 | 1.21351992 | 1.27215055 | 0 | 2.38378331 | 0 |
| 0 | 0.54305517 | 0 | 1.27215055 | 0 | 0 | 0 |
| 0 | 0.54305517 | 0 | 0 | 0 | 1.19189166 | 0 |
| 4.90116127 | 1.62916552 | 1.21351992 | 2.54430111 | 4.14885589 | 0 | 2.70970931 |
| 0 | 0 | 1.82027989 | 0 | 0 | 0 | 0 |
| 2.45058064 | 0 | 0 | 0 | 1.03721397 | 0 | 0 |
| 2.45058064 | 0 | 0 | 0 | 1.03721397 | 0 | 0 |
| 7.35174191 | 5.43055173 | 9.70815939 | 2.54430111 | 4.14885589 | 1.19189166 | 1.35485465 |
| 0 | 0 | 0 | 0 | 1.03721397 | 2.38378331 | 0 |
| 0 | 2.17222069 | 0.60675996 | 1.27215055 | 0 | 0 | 0 |
| 4.90116127 | 0 | 0 | 0 | 0 | 0 | 1.35485465 |
| 2.45058064 | 0 | 0 | 0 | 0 | 0 | 0 |
| 2.45058064 | 0 | 0.60675996 | 0 | 0 | 0 | 0 |
| 0 | 0 | 0 | 0 | 0 | 0 | 1.35485465 |
| 0 | 0 | 0 | 0 | 0 | 0 | 0 |
| 0 | 0 | 0 | 0 | 2.07442795 | 0 | 0 |
| 0 | 1.08611035 | 1.82027989 | 1.27215055 | 1.03721397 | 0 | 0 |
| 19.6046451 | 4.34444139 | 7.88787951 | 6.36075277 | 3.11164192 | 10.7270249 | 2.70970931 |
| 49.0116127 | 89.6041036 | 50.9678368 | 49.6138716 | 42.5257729 | 64.3621494 | 62.3233141 |
| 0 | 0 | 0 | 1.27215055 | 0 | 0 | 1.35485465 |
| 0 | 0 | 0 | 1.27215055 | 0 | 0 | 1.35485465 |
| 0 | 0.54305517 | 0 | 0 | 0 | 0 | 1.35485465 |
| 0 | 0.54305517 | 0.60675996 | 0 | 0 | 0 | 0 |
| 2.45058064 | 11.4041586 | 7.88787951 | 17.8101078 | 3.11164192 | 4.76756662 | 0 |
| 0 | 0.54305517 | 0 | 0 | 0 | 0 | 0 |

| | | | | | | |
|---|---|---|---|---|---|---|
| 0 | 0 | 0 | 1.27215055 | 0 | 0 | 0 |
| 0 | 0.54305517 | 0 | 0 | 0 | 0 | 0 |
| 4.90116127 | 0 | 0 | 0 | 0 | 0 | 0 |
| 4.90116127 | 0 | 0 | 0 | 0 | 0 | 0 |
| 0 | 1.08611035 | 0.60675996 | 1.27215055 | 1.03721397 | 0 | 1.35485465 |
| 0 | 0.54305517 | 0 | 0 | 0 | 0 | 0 |
| 0 | 0 | 0 | 1.27215055 | 1.03721397 | 0 | 0 |
| 4.90116127 | 2.71527587 | 1.82027989 | 3.81645166 | 6.22328384 | 10.7270249 | 9.48398258 |
| 0 | 1.62916552 | 1.82027989 | 1.27215055 | 4.14885589 | 1.19189166 | 5.41941862 |
| 73.5174191 | 32.5833104 | 61.8895161 | 35.6202155 | 58.0839825 | 42.9080996 | 35.226221 |
| 0 | 0 | 0 | 0 | 0 | 2.38378331 | 0 |
| 0 | 0 | 0 | 0 | 0 | 0 | 2.70970931 |
| 0 | 0 | 0 | 2.54430111 | 0 | 0 | 0 |
| 0 | 0 | 0 | 0 | 0 | 0 | 2.70970931 |
| 0 | 0 | 0 | 0 | 1.03721397 | 0 | 1.35485465 |
| 0 | 0 | 0 | 0 | 1.03721397 | 0 | 0 |
| 0 | 0 | 0 | 0 | 0 | 1.19189166 | 0 |
| 0 | 0 | 0 | 1.27215055 | 0 | 0 | 0 |
| 0 | 0 | 0 | 1.27215055 | 0 | 0 | 0 |
| 350.433031 | 115.127697 | 216.006547 | 73.7847322 | 252.042996 | 498.210712 | 62.3233141 |
| 0 | 0 | 0 | 0 | 0 | 0 | 0 |
| 0 | 0 | 0.60675996 | 1.27215055 | 0 | 0 | 0 |
| 0 | 0 | 0.60675996 | 1.27215055 | 0 | 0 | 0 |
| 0 | 0 | 0.60675996 | 1.27215055 | 0 | 0 | 0 |
| 0 | 0 | 0.60675996 | 0 | 0 | 0 | 1.35485465 |
| 0 | 0 | 0.60675996 | 0 | 1.03721397 | 0 | 0 |
| 0 | 0 | 2.42703985 | 1.27215055 | 2.07442795 | 0 | 1.35485465 |
| 2.45058064 | 2.71527587 | 7.28111955 | 3.81645166 | 5.18606987 | 5.95945828 | 2.70970931 |
| 0 | 0.54305517 | 0.60675996 | 1.27215055 | 0 | 3.57567497 | 1.35485465 |
| 0 | 0 | 0 | 0 | 0 | 0 | 0 |

| | | | | | | |
|---|---|---|---|---|---|---|
| 0 | 0 | 0.60675996 | 1.27215055 | 1.03721397 | 0 | 0 |
| 0 | 0.54305517 | 0 | 1.27215055 | 0 | 0 | 0 |
| 0 | 0.54305517 | 0 | 0 | 0 | 0 | 0 |
| 0 | 0.54305517 | 0 | 0 | 1.03721397 | 0 | 0 |
| 0 | 0.54305517 | 0 | 1.27215055 | 0 | 0 | 0 |
| 0 | 0.54305517 | 0 | 0 | 0 | 0 | 0 |
| 0 | 0.54305517 | 0 | 1.27215055 | 0 | 0 | 0 |
| 0 | 0.54305517 | 0 | 0 | 1.03721397 | 0 | 0 |
| 0 | 0.54305517 | 0 | 0 | 1.03721397 | 0 | 0 |
| 0 | 0.54305517 | 0 | 1.27215055 | 1.03721397 | 0 | 0 |
| 0 | 0.54305517 | 1.21351992 | 2.54430111 | 0 | 0 | 0 |
| 0 | 0.54305517 | 0.60675996 | 0 | 1.03721397 | 0 | 0 |
| 0 | 1.08611035 | 0 | 0 | 0 | 0 | 0 |
| 0 | 0.54305517 | 0 | 1.27215055 | 1.03721397 | 2.38378331 | 0 |
| 0 | 0 | 0 | 0 | 0 | 0 | 0 |
| 0 | 0 | 0.60675996 | 0 | 0 | 0 | 0 |
| 4.90116127 | 0 | 0 | 0 | 1.03721397 | 1.19189166 | 0 |
| 0 | 1.08611035 | 0 | 0 | 0 | 0 | 0 |
| 0 | 1.62916552 | 1.21351992 | 1.27215055 | 0 | 0 | 0 |
| 0 | 0 | 0 | 0 | 0 | 0 | 0 |
| 2.45058064 | 0 | 0 | 1.27215055 | 0 | 0 | 0 |
| 0 | 0.54305517 | 0.60675996 | 0 | 1.03721397 | 0 | 1.35485465 |
| 2.45058064 | 1.08611035 | 1.82027989 | 0 | 1.03721397 | 3.57567497 | 0 |
| 14.7034838 | 1.08611035 | 1.21351992 | 1.27215055 | 1.03721397 | 1.19189166 | 0 |
| 0 | 0 | 0 | 0 | 0 | 0 | 0 |
| 0 | 0 | 0 | 0 | 0 | 0 | 0 |
| 0 | 0 | 0 | 0 | 0 | 0 | 0 |
| 0 | 0 | 0 | 0 | 0 | 0 | 0 |
| 0 | 0 | 0 | 0 | 0 | 0 | 0 |
| 0 | 0 | 0 | 0 | 0 | 0 | 0 |

| | | | | | | |
|---|---|---|---|---|---|---|
| 0 | 0 | 0 | 0 | 0 | 0 | 0 |
| 0 | 0 | 0 | 0 | 0 | 0 | 0 |
| 0 | 0 | 0 | 0 | 0 | 0 | 0 |
| 0 | 0 | 0 | 0 | 0 | 0 | 0 |
| 0 | 0 | 0 | 0 | 0 | 0 | 0 |
| 0 | 0 | 0 | 0 | 0 | 0 | 0 |
| 0 | 0 | 0 | 0 | 0 | 0 | 0 |
| 2.45058064 | 0 | 1.21351992 | 1.27215055 | 1.03721397 | 0 | 0 |
| 4.90116127 | 2.17222069 | 1.82027989 | 5.08860222 | 1.03721397 | 2.38378331 | 4.06456396 |
| 0 | 0.54305517 | 0 | 0 | 1.03721397 | 0 | 0 |
| 0 | 1.62916552 | 0 | 0 | 0 | 0 | 0 |
| 2.45058064 | 1.08611035 | 1.82027989 | 2.54430111 | 1.03721397 | 0 | 0 |
| 0 | 0 | 0 | 0 | 0 | 0 | 0 |
| 0 | 0 | 0 | 0 | 0 | 0 | 0 |
| 0 | 0 | 0 | 0 | 0 | 0 | 0 |
| 0 | 0.54305517 | 0 | 1.27215055 | 0 | 0 | 1.35485465 |
| 2.45058064 | 0 | 0.60675996 | 0 | 0 | 0 | 0 |
| 0 | 0 | 0.60675996 | 0 | 1.03721397 | 0 | 0 |
| 2.45058064 | 0 | 0 | 0 | 0 | 0 | 0 |
| 0 | 0 | 0 | 0 | 0 | 0 | 0 |
| 0 | 0 | 1.21351992 | 1.27215055 | 1.03721397 | 0 | 0 |
| 1222.83974 | 559.346828 | 987.198458 | 639.891729 | 802.803616 | 643.621494 | 654.394798 |
| 0 | 2.17222069 | 1.21351992 | 0 | 0 | 0 | 0 |
| 0 | 0 | 0.60675996 | 0 | 0 | 0 | 1.35485465 |
| 0 | 0 | 3.03379981 | 0 | 0 | 0 | 0 |
| 0 | 0.54305517 | 0 | 0 | 1.03721397 | 0 | 0 |
| 0 | 0 | 0.60675996 | 0 | 0 | 0 | 0 |
| 2.45058064 | 7.60277242 | 6.06759962 | 5.08860222 | 8.29771179 | 7.15134993 | 2.70970931 |
| 0 | 0 | 0 | 2.54430111 | 0 | 1.19189166 | 0 |
| 2.45058064 | 1.62916552 | 0 | 0 | 1.03721397 | 0 | 1.35485465 |

| | | | | | | |
|---|---|---|---|---|---|---|
| 0 | 0 | 0 | 0 | 2.07442795 | 0 | 0 |
| 524.424256 | 948.174332 | 629.816841 | 667.879041 | 570.467685 | 710.367427 | 1047.30265 |
| 0 | 0.54305517 | 1.21351992 | 0 | 1.03721397 | 1.19189166 | 1.35485465 |
| 0 | 0 | 0.60675996 | 1.27215055 | 0 | 0 | 0 |
| 0 | 0 | 0 | 0 | 1.03721397 | 0 | 0 |
| 0 | 0.54305517 | 0.60675996 | 1.27215055 | 0 | 0 | 0 |
| 0 | 0 | 0 | 0 | 1.03721397 | 0 | 0 |
| 0 | 0 | 0 | 0 | 0 | 0 | 0 |
| 0 | 0 | 0 | 0 | 0 | 0 | 0 |
| 0 | 0 | 0 | 0 | 0 | 0 | 0 |
| 0 | 0 | 0 | 0 | 0 | 0 | 0 |
| 4.90116127 | 0 | 0 | 0 | 0 | 0 | 0 |
| 0 | 0 | 0 | 1.27215055 | 1.03721397 | 0 | 0 |
| 0 | 0 | 0 | 0 | 0 | 0 | 1.35485465 |
| 0 | 0 | 0.60675996 | 0 | 0 | 1.19189166 | 0 |
| 2.45058064 | 0 | 0.60675996 | 0 | 0 | 0 | 0 |
| 0 | 3.25833104 | 1.82027989 | 0 | 2.07442795 | 0 | 1.35485465 |
| 0 | 3.25833104 | 0.60675996 | 1.27215055 | 3.11164192 | 3.57567497 | 0 |
| 0 | 0 | 0.60675996 | 0 | 0 | 1.19189166 | 0 |
| 0 | 0 | 0.60675996 | 0 | 0 | 0 | 0 |
| 0 | 0 | 0 | 0 | 0 | 1.19189166 | 0 |
| 7.35174191 | 1.08611035 | 4.8540797 | 3.81645166 | 12.4465677 | 0 | 4.06456396 |
| 0 | 1.62916552 | 0 | 1.27215055 | 1.03721397 | 2.38378331 | 0 |
| 0 | 0.54305517 | 0 | 0 | 0 | 0 | 1.35485465 |
| 0 | 0.54305517 | 0 | 0 | 0 | 0 | 0 |
| 0 | 0.54305517 | 0 | 0 | 0 | 0 | 0 |
| 1266.95019 | 1464.6198 | 1520.54047 | 1109.31528 | 1460.39727 | 1830.74558 | 2601.32094 |
| 0 | 0 | 0 | 0 | 0 | 0 | 0 |
| 0 | 0 | 0 | 0 | 0 | 0 | 0 |
| 0 | 0 | 0 | 0 | 0 | 0 | 0 |

| | | | | | | |
|---|---|---|---|---|---|---|
| 0 | 0.54305517 | 0 | 0 | 1.03721397 | 0 | 0 |
| 0 | 0 | 0 | 1.27215055 | 0 | 0 | 1.35485465 |
| 0 | 0 | 0 | 0 | 0 | 0 | 1.35485465 |
| 0 | 1.62916552 | 1.21351992 | 2.54430111 | 0 | 0 | 0 |
| 0 | 0 | 0 | 0 | 2.07442795 | 0 | 0 |
| 0 | 0 | 0 | 0 | 2.07442795 | 0 | 0 |
| 0 | 0.54305517 | 0.60675996 | 1.27215055 | 0 | 0 | 0 |
| 24.5058064 | 32.5833104 | 16.9892789 | 30.5316133 | 13.4837817 | 32.1810747 | 31.1616571 |
| 0 | 0.54305517 | 0 | 0 | 0 | 0 | 0 |
| 0 | 3.25833104 | 0.60675996 | 3.81645166 | 2.07442795 | 2.38378331 | 0 |
| 3087.7316 | 1249.0269 | 2043.56755 | 1679.23873 | 1620.12823 | 1491.05646 | 2375.06021 |
| 4.90116127 | 0 | 0 | 1.27215055 | 0 | 0 | 0 |
| 0 | 0 | 0 | 1.27215055 | 0 | 0 | 0 |
| 0 | 0 | 1.21351992 | 0 | 0 | 0 | 0 |
| 0 | 0.54305517 | 0 | 1.27215055 | 0 | 0 | 0 |
| 1460.54606 | 976.413201 | 661.975119 | 754.385279 | 602.621319 | 694.872835 | 709.943839 |
| 0 | 0 | 0 | 0 | 0 | 0 | 0 |
| 2.45058064 | 0.54305517 | 0.60675996 | 1.27215055 | 1.03721397 | 1.19189166 | 1.35485465 |
| 2.45058064 | 11.4041586 | 8.49463947 | 3.81645166 | 6.22328384 | 4.76756662 | 18.9679652 |
| 0 | 1.62916552 | 0 | 0 | 0 | 0 | 1.35485465 |
| 0 | 1.08611035 | 0 | 0 | 0 | 0 | 0 |
| 0 | 0 | 0 | 0 | 1.03721397 | 0 | 0 |
| 0 | 0.54305517 | 2.42703985 | 0 | 2.07442795 | 1.19189166 | 1.35485465 |
| 0 | 1.08611035 | 0 | 0 | 0 | 0 | 0 |
| 0 | 1.08611035 | 0.60675996 | 0 | 1.03721397 | 0 | 0 |
| 4.90116127 | 1.62916552 | 1.21351992 | 2.54430111 | 0 | 0 | 1.35485465 |
| 4.90116127 | 4.34444139 | 4.8540797 | 10.1772044 | 9.33492576 | 5.95945828 | 4.06456396 |
| 0 | 0 | 0 | 1.27215055 | 0 | 1.19189166 | 0 |
| 0 | 0 | 0.60675996 | 1.27215055 | 1.03721397 | 0 | 0 |
| 0 | 0 | 0.60675996 | 1.27215055 | 1.03721397 | 0 | 0 |

| | | | | | | |
|---|---|---|---|---|---|---|
| 0 | 0.54305517 | 0 | 0 | 0 | 2.38378331 | 0 |
| 0 | 1.08611035 | 0 | 0 | 0 | 0 | 0 |
| 0 | 0 | 0 | 0 | 0 | 0 | 1.35485465 |
| 0 | 0 | 0 | 0 | 1.03721397 | 0 | 0 |
| 0 | 0.54305517 | 0.60675996 | 0 | 0 | 0 | 2.70970931 |
| 0 | 0 | 0 | 0 | 0 | 0 | 0 |
| 0 | 1.08611035 | 0.60675996 | 1.27215055 | 4.14885589 | 5.95945828 | 0 |
| 0 | 0 | 3.03379981 | 1.27215055 | 0 | 1.19189166 | 4.06456396 |
| 0 | 2.17222069 | 2.42703985 | 0 | 2.07442795 | 0 | 0 |
| 0 | 0 | 0 | 0 | 0 | 0 | 0 |
| 0 | 0.54305517 | 0 | 0 | 0 | 0 | 0 |
| 0 | 0.54305517 | 0.60675996 | 2.54430111 | 2.07442795 | 0 | 0 |
| 0 | 0 | 0 | 1.27215055 | 2.07442795 | 0 | 0 |
| 0 | 0 | 0 | 1.27215055 | 1.03721397 | 0 | 0 |
| 2.45058064 | 0.54305517 | 0.60675996 | 0 | 0 | 0 | 0 |
| 0 | 0 | 0 | 0 | 0 | 0 | 0 |
| 0 | 0.54305517 | 0.60675996 | 0 | 0 | 0 | 0 |
| 0 | 2.71527587 | 3.03379981 | 0 | 4.14885589 | 0 | 0 |
| 2.45058064 | 0 | 0 | 0 | 0 | 0 | 0 |
| 63.7150965 | 19.0069311 | 46.7205171 | 27.9873122 | 106.833039 | 97.7351157 | 77.2267153 |
| 14.7034838 | 5.43055173 | 3.03379981 | 5.08860222 | 3.11164192 | 4.76756662 | 1.35485465 |
| 0 | 0 | 0 | 0 | 2.07442795 | 2.38378331 | 0 |
| 9.80232254 | 2.17222069 | 3.03379981 | 0 | 2.07442795 | 1.19189166 | 5.41941862 |
| 0 | 0.54305517 | 0.60675996 | 0 | 0 | 0 | 1.35485465 |
| 58.8139353 | 31.4972 | 24.8771584 | 17.8101078 | 30.0792052 | 22.6459415 | 24.3873838 |
| 0 | 0 | 0 | 0 | 1.03721397 | 0 | 0 |
| 0 | 0 | 0 | 0 | 0 | 0 | 0 |
| 0 | 0 | 0 | 0 | 0 | 0 | 0 |
| 0 | 0 | 0 | 0 | 0 | 0 | 0 |
| 12.2529032 | 13.5763793 | 10.9216793 | 5.08860222 | 12.4465677 | 16.6864832 | 14.9034012 |

| | | | | | | |
|---|---|---|---|---|---|---|
| 4.90116127 | 4.88749656 | 1.21351992 | 1.27215055 | 20.7442795 | 7.15134993 | 13.5485465 |
| 4.90116127 | 0.54305517 | 1.21351992 | 0 | 5.18606987 | 2.38378331 | 5.41941862 |
| 26.956387 | 52.6763518 | 25.4839184 | 30.5316133 | 45.6374148 | 32.1810747 | 12.1936919 |
| 0 | 0.54305517 | 0.60675996 | 0 | 2.07442795 | 0 | 0 |
| 39.2092902 | 95.0346553 | 49.7543169 | 31.8037639 | 39.414131 | 47.6756662 | 56.9038955 |
| 0 | 0 | 0.60675996 | 0 | 0 | 0 | 0 |
| 0 | 0 | 1.21351992 | 1.27215055 | 3.11164192 | 0 | 5.41941862 |
| 0 | 0 | 0 | 0 | 0 | 0 | 1.35485465 |
| 4.90116127 | 3.80138621 | 3.03379981 | 3.81645166 | 0 | 5.95945828 | 1.35485465 |
| 0 | 0 | 1.21351992 | 1.27215055 | 0 | 0 | 0 |
| 0 | 0 | 0.60675996 | 3.81645166 | 2.07442795 | 0 | 0 |
| 0 | 0 | 0 | 1.27215055 | 1.03721397 | 0 | 1.35485465 |
| 2.45058064 | 1.08611035 | 0.60675996 | 0 | 0 | 0 | 0 |
| 7.35174191 | 8.68888277 | 6.06759962 | 6.36075277 | 5.18606987 | 2.38378331 | 2.70970931 |
| 0 | 0.54305517 | 3.64055977 | 1.27215055 | 1.03721397 | 2.38378331 | 1.35485465 |
| 56.3633546 | 18.4638759 | 33.3717979 | 39.4366672 | 14.5209956 | 19.0702665 | 6.77427327 |
| 0 | 0.54305517 | 0 | 0 | 0 | 1.19189166 | 0 |
| 0 | 0.54305517 | 0 | 0 | 1.03721397 | 0 | 0 |
| 0 | 0 | 0 | 1.27215055 | 0 | 1.19189166 | 0 |
| 0 | 0 | 0 | 0 | 0 | 0 | 0 |
| 0 | 0 | 0.60675996 | 1.27215055 | 1.03721397 | 0 | 1.35485465 |
| 0 | 0 | 0.60675996 | 0 | 3.11164192 | 1.19189166 | 0 |
| 19.6046451 | 9.77499312 | 23.6636385 | 13.9936561 | 36.3024891 | 27.4135081 | 31.1616571 |
| 49.0116127 | 61.9082897 | 57.0354364 | 33.0759144 | 87.1259738 | 69.129716 | 107.033518 |
| 0 | 0.54305517 | 0 | 0 | 0 | 0 | 1.35485465 |
| 0 | 0.54305517 | 0 | 0 | 3.11164192 | 0 | 0 |
| 0 | 0.54305517 | 1.21351992 | 0 | 0 | 0 | 2.70970931 |
| 2.45058064 | 0 | 0 | 0 | 0 | 0 | 0 |
| 0 | 0 | 0 | 0 | 0 | 0 | 1.35485465 |
| 0 | 0 | 0.60675996 | 0 | 0 | 0 | 0 |

| | | | | | | |
|---|---|---|---|---|---|---|
| 0 | 0 | 0.60675996 | 0 | 0 | 0 | 0 |
| 0 | 0 | 0.60675996 | 0 | 0 | 0 | 0 |
| 0 | 0 | 0 | 1.27215055 | 0 | 0 | 0 |
| 0 | 0 | 0 | 1.27215055 | 0 | 0 | 0 |
| 0 | 0 | 0 | 0 | 0 | 0 | 0 |
| 0 | 0 | 0 | 0 | 0 | 0 | 0 |
| 0 | 0 | 0 | 0 | 0 | 0 | 0 |
| 0 | 0.54305517 | 0 | 0 | 0 | 0 | 0 |
| 0 | 0 | 0 | 0 | 0 | 0 | 0 |
| 0 | 0 | 0 | 0 | 0 | 0 | 0 |
| 0 | 0.54305517 | 0.60675996 | 1.27215055 | 0 | 0 | 0 |
| 2.45058064 | 0 | 0 | 0 | 0 | 0 | 0 |
| 2.45058064 | 0 | 0 | 0 | 0 | 0 | 0 |
| 0 | 3.25833104 | 0.60675996 | 1.27215055 | 1.03721397 | 2.38378331 | 0 |
| 49.0116127 | 78.7430001 | 44.2934772 | 73.7847322 | 31.1164192 | 30.989183 | 43.3553489 |
| 2.45058064 | 0 | 0 | 0 | 0 | 0 | 0 |
| 0 | 0 | 0 | 0 | 0 | 0 | 1.35485465 |
| 0 | 0 | 0 | 0 | 1.03721397 | 0 | 0 |
| 0 | 0 | 0 | 0 | 1.03721397 | 0 | 0 |
| 0 | 0 | 0.60675996 | 0 | 0 | 0 | 0 |
| 0 | 0 | 0.60675996 | 0 | 0 | 0 | 0 |
| 0 | 0 | 0.60675996 | 0 | 0 | 0 | 0 |
| 0 | 0 | 0.60675996 | 0 | 0 | 0 | 0 |
| 0 | 0 | 0.60675996 | 0 | 0 | 0 | 0 |
| 0 | 0 | 0.60675996 | 0 | 0 | 0 | 0 |
| 0 | 0.54305517 | 0 | 0 | 0 | 0 | 0 |
| 0 | 0 | 0 | 1.27215055 | 0 | 0 | 0 |
| 0 | 0 | 0 | 1.27215055 | 0 | 0 | 0 |
| 0 | 0 | 0 | 0 | 0 | 1.19189166 | 0 |
| 0 | 0 | 0 | 0 | 1.03721397 | 0 | 0 |

| | | | | | | |
|---|---|---|---|---|---|---|
| 0 | 0.54305517 | 0 | 0 | 0 | 0 | 0 |
| 0 | 0 | 0 | 0 | 0 | 0 | 0 |
| 0 | 0.54305517 | 0 | 0 | 0 | 0 | 0 |
| 0 | 0.54305517 | 0 | 0 | 0 | 0 | 0 |
| 0 | 0.54305517 | 0 | 0 | 0 | 0 | 0 |
| 0 | 0 | 0 | 0 | 0 | 0 | 0 |
| 0 | 0.54305517 | 0 | 0 | 0 | 0 | 0 |
| 0 | 0 | 0 | 0 | 0 | 0 | 0 |
| 0 | 0 | 0 | 0 | 0 | 0 | 0 |
| 0 | 0.54305517 | 0 | 0 | 0 | 0 | 0 |
| 0 | 0 | 0 | 0 | 0 | 0 | 0 |
| 0 | 0 | 0 | 0 | 0 | 0 | 0 |
| 0 | 0 | 0 | 0 | 1.03721397 | 0 | 0 |
| 0 | 0 | 0.60675996 | 1.27215055 | 0 | 0 | 0 |
| 0 | 0 | 0 | 0 | 1.03721397 | 0 | 1.35485465 |
| 0 | 0 | 0.60675996 | 0 | 0 | 0 | 0 |
| 0 | 0 | 0 | 0 | 1.03721397 | 0 | 0 |
| 0 | 1.62916552 | 0 | 0 | 0 | 0 | 0 |
| 0 | 0.54305517 | 0 | 0 | 0 | 1.19189166 | 0 |
| 0 | 0 | 0.60675996 | 0 | 1.03721397 | 0 | 0 |
| 0 | 0 | 0.60675996 | 0 | 0 | 0 | 1.35485465 |
| 0 | 0.54305517 | 0.60675996 | 0 | 0 | 0 | 0 |
| 0 | 0.54305517 | 0.60675996 | 0 | 0 | 0 | 0 |
| 0 | 1.62916552 | 0 | 0 | 0 | 0 | 1.35485465 |
| 1014.54038 | 184.638759 | 330.684179 | 277.328821 | 236.484786 | 281.286431 | 151.743721 |

| | baseMean | log2FoldChai | lfcSE | stat | pvalue | padj |
|---|---|---|---|---|---|---|
| miR-3535 | 138.418659 | 1.52311082 | 0.22774601 | 6.68776086 | 2.27E-11 | 6.32E-09 |
| miR-677-5p | 69.7290435 | 2.4481208 | 0.4006183 | 6.11085612 | 9.91E-10 | 1.38E-07 |
| miR-677-3p | 48.2297058 | 2.27216607 | 0.45953873 | 4.94444954 | 7.64E-07 | 6.50E-05 |
| miR-34c-5p | 612.519753 | -0.7366842 | 0.15017658 | -4.9054533 | 9.32E-07 | 6.50E-05 |
| miR-7033-5p | 24.6752257 | 1.61250806 | 0.34035917 | 4.737666 | 2.16E-06 | 0.00012064 |
| miR-703 | 11.9831413 | 2.40025244 | 0.53167971 | 4.51447066 | 6.35E-06 | 0.00029516 |
| miR-5126 | 25.6780927 | 2.2075404 | 0.50286989 | 4.38988387 | 1.13E-05 | 0.00045202 |
| miR-5121 | 96.486377 | 1.85068286 | 0.43887854 | 4.21684517 | 2.48E-05 | 0.00086401 |
| miR-6240 | 61.3535994 | 2.03319299 | 0.52617764 | 3.86408096 | 0.00011151 | 0.00345676 |
| miR-5114 | 10.9566072 | 1.85899427 | 0.50314982 | 3.69471318 | 0.00022014 | 0.00614177 |
| miR-1839-3p | 7.87033995 | 1.49370073 | 0.43718664 | 3.41662023 | 0.00063404 | 0.01608147 |
| miR-1949 | 17.69808 | 1.67536106 | 0.50889122 | 3.29217913 | 0.00099414 | 0.02198243 |
| miR-196a-5p | 28.5595684 | 1.04335634 | 0.31773088 | 3.28377377 | 0.00102427 | 0.02198243 |
| miR-3064-5p | 31.3550538 | 1.15351357 | 0.35408443 | 3.25773595 | 0.00112305 | 0.02238075 |
| miR-196b-5p | 46.4368258 | 1.02130587 | 0.32091054 | 3.18252515 | 0.00145997 | 0.02715541 |
| miR-6538 | 587.974314 | 1.78064781 | 0.56598144 | 3.14612405 | 0.0016545 | 0.02885031 |
| miR-18a-5p | 23.1934484 | -0.7590746 | 0.24851063 | -3.0544956 | 0.00225439 | 0.03680873 |
| miR-3963 | 9.7298299 | -1.3605821 | 0.4477279 | -3.0388594 | 0.00237476 | 0.03680873 |
| miR-471-5p | 91.2212066 | -0.5585286 | 0.18743697 | -2.9798211 | 0.00288417 | 0.0420035 |
| FGCZ6510_Free_miRNA_dedup_pvalu | | | | FGCZ6510_Free_miRNA_dedup_pvalu | | |
| miR-5099 | 6.60223164 | 1.16205934 | 0.39187447 | 2.9653867 | 0.00302303 | 0.0420035 |
| miR-1843b-3p | 6.71966549 | 1.24541745 | 0.42194979 | 2.95157735 | 0.00316155 | 0.0420035 |
| miR-1843a-3p | 17.6097526 | 1.07867716 | 0.37360471 | 2.88721511 | 0.00388668 | 0.04929022 |
| miR-690 | 2.97272164 | 1.64944501 | 0.58129867 | 2.83751726 | 0.00454659 | 0.0551521 |
| miR-6516-3p | 8.87690997 | 1.59990245 | 0.57553277 | 2.77986333 | 0.00543818 | 0.06321882 |
| miR-199a-3p | 24.2253002 | 0.83795352 | 0.30411384 | 2.75539424 | 0.00586215 | 0.06542154 |
| miR-3068-3p | 574.023387 | 1.02827521 | 0.38431867 | 2.67557961 | 0.00746002 | 0.07985761 |
| miR-1a-3p | 57.134014 | 0.73283455 | 0.27731283 | 2.64262762 | 0.00822655 | 0.07985761 |
| miR-152-3p | 42.8338371 | 0.85304811 | 0.32302702 | 2.64079487 | 0.00827118 | 0.07985761 |
| miR-143-3p | 2792.76504 | 0.72698395 | 0.2766185 | 2.62811033 | 0.00858607 | 0.07985761 |
| miR-7a-5p | 217.199695 | -0.478412 | 0.18203862 | -2.6280797 | 0.00858684 | 0.07985761 |
| let-7d-3p | 59.5117461 | -0.8254162 | 0.31634438 | -2.6092329 | 0.00907455 | 0.08126992 |
| miR-465c-3p | 195.387288 | -0.6643412 | 0.25551187 | -2.6000403 | 0.00932128 | 0.08126992 |
| miR-465a-5p | 237.697156 | -0.302954 | 0.11740813 | -2.5803492 | 0.00987005 | 0.08344675 |
| miR-465b-3p | 197.221231 | -0.650902 | 0.25430281 | -2.559555 | 0.01048063 | 0.08600279 |
| miR-1892 | 1.71090551 | 2.4425092 | 0.96385106 | 2.53411477 | 0.01127318 | NA |
| miR-7657-5p | 0.94550224 | 2.93764918 | 1.160214 | 2.53198908 | 0.01134175 | NA |
| miR-465a-3p | 98.5770647 | -0.6514109 | 0.25783066 | -2.5265067 | 0.01152032 | 0.09008211 |
| miR-880-3p | 38.3055603 | -0.6971596 | 0.27628081 | -2.5233732 | 0.0116235 | 0.09008211 |
| miR-199b-3p | 12.1261003 | 0.82717135 | 0.33637462 | 2.45907778 | 0.01392944 | 0.10338677 |
| miR-382-5p | 4.98132957 | 1.12273555 | 0.45729231 | 2.45518138 | 0.01408135 | 0.10338677 |
| let-7b-5p | 404.992122 | 0.74924284 | 0.31209575 | 2.40068262 | 0.01636452 | 0.11706928 |
| miR-3084-3p | 21.9779482 | 1.1161417 | 0.47043137 | 2.37259197 | 0.01766376 | 0.12059435 |
| miR-9-3p | 15.3772701 | -0.6264733 | 0.26418078 | -2.371381 | 0.01772175 | 0.12059435 |
| miR-671-5p | 2.05914086 | 2.12041036 | 0.89711068 | 2.3635995 | 0.01809837 | NA |

| | | | | | | |
|---|---|---|---|---|---|---|
| miR-466b-3p | 14.4291798 | -0.7349009 | 0.3186726 | -2.3061314 | 0.02110329 | 0.14018613 |
| miR-362-3p | 9.05051934 | -0.8948766 | 0.40037448 | -2.2350989 | 0.02541086 | 0.1630634 |
| miR-222-3p | 126.642076 | 0.45116029 | 0.20227118 | 2.23047249 | 0.02571609 | 0.1630634 |
| miR-194-5p | 42.7286453 | -0.4959374 | 0.22399381 | -2.2140677 | 0.02682412 | 0.16630956 |
| miR-340-3p | 46.9016574 | -0.4578034 | 0.20799411 | -2.2010401 | 0.02773318 | 0.1682078 |
| miR-144-3p | 1.93875591 | -1.6742829 | 0.76374844 | -2.1921916 | 0.02836568 | NA |
| miR-145a-3p | 87.2579919 | 0.66413275 | 0.30411175 | 2.18384444 | 0.02897368 | 0.1719927 |
| miR-34b-5p | 5.87685053 | -1.0537335 | 0.49472258 | -2.1299482 | 0.03317589 | 0.19283488 |
| miR-26b-5p | 642.131805 | 0.46003547 | 0.21747449 | 2.11535374 | 0.03439981 | 0.19464358 |
| miR-224-5p | 10.7948938 | 0.79338517 | 0.37606159 | 2.10972138 | 0.03488236 | 0.19464358 |
| miR-743b-3p | 306.848474 | -0.3744895 | 0.17936912 | -2.0878147 | 0.03681456 | 0.20139728 |
| miR-465c-5p | 432.005517 | -0.4046218 | 0.19496821 | -2.075322 | 0.0379567 | 0.20144696 |
| miR-10a-3p | 38.1043265 | 0.557811 | 0.26921695 | 2.07197576 | 0.0382677 | 0.20144696 |
| miR-296-5p | 5.01612229 | -1.0797286 | 0.52521691 | -2.0557766 | 0.03980406 | 0.2056543 |
| miR-320-3p | 27.5813736 | 0.61300546 | 0.30075257 | 2.03823849 | 0.04152609 | 0.21065052 |
| miR-26a-5p | 2241.62536 | 0.30837274 | 0.15254266 | 2.02155088 | 0.04322277 | 0.21367857 |
| miR-434-3p | 11.6779574 | 0.65139181 | 0.32288829 | 2.01739065 | 0.04365476 | 0.21367857 |
| miR-340-5p | 205.447145 | -0.3490945 | 0.17405204 | -2.0056901 | 0.04488932 | 0.21593311 |
| miR-185-5p | 6.09231223 | 0.85170255 | 0.43714688 | 1.94832124 | 0.05137654 | 0.23805644 |
| miR-1291 | 18.5858571 | 0.63879735 | 0.32821696 | 1.94626551 | 0.05162285 | 0.23805644 |
| miR-743a-3p | 359.577587 | -0.3785245 | 0.19484105 | -1.942735 | 0.05204818 | 0.23805644 |
| miR-31-3p | 1.31824104 | 1.95487091 | 1.02547207 | 1.90631316 | 0.05660959 | NA |
| miR-293-5p | 4.65322724 | -1.0244622 | 0.54145142 | -1.8920667 | 0.05848209 | 0.2631694 |
| miR-7672-5p | 0.92194748 | 2.25516062 | 1.20677552 | 1.86874907 | 0.06165773 | NA |
| miR-7670-3p | 1.00639668 | 1.76431832 | 0.94722037 | 1.86262709 | 0.06251475 | NA |
| miR-871-3p | 163.290013 | -0.4180286 | 0.22462959 | -1.8609685 | 0.06274863 | 0.2729424 |
| miR-23a-5p | 2.54457518 | 1.30256343 | 0.70155378 | 1.85668363 | 0.06335618 | NA |
| miR-130b-5p | 54.3718721 | -0.441548 | 0.23805127 | -1.8548442 | 0.06361847 | 0.2729424 |
| miR-449a-5p | 32.2510132 | -0.4797628 | 0.25950728 | -1.8487451 | 0.06449462 | 0.2729424 |
| miR-29c-3p | 21.2952374 | 0.59769529 | 0.32338545 | 1.84824424 | 0.06456702 | 0.2729424 |
| miR-3064-3p | 1.37252307 | 1.51841668 | 0.82371371 | 1.84337916 | 0.06527367 | NA |
| miR-466n-3p | 1.72025496 | -1.450306 | 0.79267459 | -1.829636 | 0.06730438 | NA |
| miR-34b-3p | 1325.59618 | -0.3527148 | 0.19330609 | -1.8246438 | 0.06805479 | 0.28339235 |
| miR-149-5p | 2.30810683 | -1.291005 | 0.71281406 | -1.8111385 | 0.07011942 | NA |
| miR-7652-5p | 0.45728412 | 2.43188806 | 1.34695307 | 1.80547348 | 0.07100062 | NA |
| miR-1306-3p | 0.91227929 | 1.73868884 | 0.96419156 | 1.80326079 | 0.07134727 | NA |
| miR-145a-5p | 108.385313 | 0.75321365 | 0.42206003 | 1.78461261 | 0.07432417 | 0.29984322 |
| miR-23a-3p | 360.960171 | 0.17203449 | 0.09696384 | 1.77421287 | 0.07602795 | 0.29984322 |
| miR-133a-3p | 4.38226715 | 1.01853599 | 0.57411565 | 1.77409549 | 0.07604736 | 0.29984322 |
| miR-30a-3p | 141.379344 | 0.30278656 | 0.17141531 | 1.76639158 | 0.07733018 | 0.29984322 |
| miR-365-3p | 19.152857 | -0.5007815 | 0.28447719 | -1.7603574 | 0.07834722 | 0.29984322 |
| miR-466c-3p | 5.6338535 | -0.7296911 | 0.41466082 | -1.75973 | 0.0784536 | 0.29984322 |
| miR-7043-3p | 0.67463661 | -2.279817 | 1.2995973 | -1.7542488 | 0.07938789 | NA |
| miR-7073-5p | 0.62550177 | 2.31747143 | 1.32977425 | 1.74275552 | 0.08137633 | NA |
| miR-375-3p | 544.501842 | -0.4594731 | 0.26492601 | -1.734345 | 0.08285689 | 0.31239288 |
| miR-330-5p | 2.25026883 | 1.16239142 | 0.67033479 | 1.73404609 | 0.08290991 | NA |

| | | | | | | |
|---|---|---|---|---|---|---|
| let-7c-5p | 2632.73258 | 0.50093631 | 0.28989863 | 1.72797062 | 0.08399349 | 0.31245579 |
| miR-669a-3p | 14.2847503 | -0.6920128 | 0.40229125 | -1.7201786 | 0.08539998 | 0.31350783 |
| miR-181b-5p | 15.0121538 | 0.56978617 | 0.33281681 | 1.71201139 | 0.08689457 | 0.31485175 |
| miR-744-3p | 1.46528061 | 1.562142 | 0.92162937 | 1.69497854 | 0.09007949 | NA |
| miR-425-5p | 621.426982 | -0.3138912 | 0.18530007 | -1.6939617 | 0.09027257 | 0.32289803 |
| miR-22-3p | 404.958972 | -0.4833673 | 0.28801791 | -1.6782545 | 0.09329744 | 0.32294568 |
| miR-205-5p | 52.9611185 | 0.6643591 | 0.39703372 | 1.67330647 | 0.09426698 | 0.32294568 |
| miR-676-5p | 3.96807213 | -1.092792 | 0.6531194 | -1.6731888 | 0.09429014 | 0.32294568 |
| miR-672-3p | 0.9247964 | 1.83343054 | 1.09762557 | 1.67036063 | 0.09484803 | NA |
| miR-192-5p | 19.6076054 | -0.4374969 | 0.26312521 | -1.6626948 | 0.09637354 | 0.32294568 |
| miR-1306-5p | 10.1158268 | -0.7817714 | 0.47123103 | -1.6589982 | 0.09711615 | 0.32294568 |
| miR-297a-3p | 2.94004438 | -0.9280173 | 0.55957625 | -1.6584288 | 0.09723096 | 0.32294568 |
| miR-196a-1-3p | 1.43202921 | 1.36123025 | 0.8244681 | 1.65104054 | 0.0987303 | NA |
| miR-3089-3p | 0.51697226 | 2.19733975 | 1.33988772 | 1.6399432 | 0.10101698 | NA |
| miR-883b-5p | 1.90481971 | -1.2582358 | 0.76862694 | -1.6369915 | 0.10163224 | NA |
| miR-501-3p | 15.7886294 | -0.563155 | 0.34421149 | -1.6360727 | 0.10182438 | 0.33162424 |
| miR-187-3p | 4.10678409 | 0.77951891 | 0.47700925 | 1.63417985 | 0.10222109 | 0.33162424 |
| miR-1902 | 0.9334202 | 1.8087553 | 1.10892607 | 1.63108736 | 0.10287189 | NA |
| miR-7b-5p | 1.58039688 | -1.4089006 | 0.86631802 | -1.6263088 | 0.10388399 | NA |
| miR-741-3p | 175.216903 | -0.3238715 | 0.19966428 | -1.6220805 | 0.10478612 | 0.33491229 |
| miR-15b-3p | 0.58393266 | 1.96784145 | 1.21548571 | 1.61897538 | 0.10545256 | NA |
| miR-466d-3p | 6.18976079 | -0.6434648 | 0.39766047 | -1.6181261 | 0.10563542 | 0.33491229 |
| miR-881-3p | 611.032282 | -0.3284986 | 0.20560306 | -1.5977319 | 0.11010266 | 0.34515329 |
| miR-30d-3p | 1.16805685 | -1.5260516 | 0.96223563 | -1.5859437 | 0.11275208 | NA |
| miR-200b-5p | 35.5326713 | 0.36428505 | 0.23019462 | 1.58250897 | 0.11353343 | 0.34884223 |
| miR-128-3p | 325.311301 | -0.3431491 | 0.21698679 | -1.5814286 | 0.11378008 | 0.34884223 |
| miR-99b-3p | 2.06510619 | 1.11940367 | 0.70822615 | 1.58057375 | 0.11397553 | NA |
| miR-546 | 2.051916 | 1.37771337 | 0.87791551 | 1.56930064 | 0.1165779 | NA |
| miR-379-5p | 41.671937 | 0.49765272 | 0.31884934 | 1.56077701 | 0.11857637 | 0.35959574 |
| miR-7238-5p | 0.81975182 | 1.91832473 | 1.26095084 | 1.52133189 | 0.12817657 | NA |
| miR-7219-5p | 0.53436565 | 2.04977841 | 1.35274605 | 1.51527214 | 0.1297035 | NA |
| miR-664-3p | 50.6424293 | 0.65926694 | 0.43508263 | 1.51526837 | 0.12970446 | 0.38116247 |
| miR-106b-5p | 6.11628141 | 0.63885133 | 0.42280971 | 1.51096655 | 0.13079697 | 0.38116247 |
| miR-743b-5p | 11.7154502 | -0.5260191 | 0.34883049 | -1.5079506 | 0.13156718 | 0.38116247 |
| miR-297c-3p | 1.33130908 | -1.1932066 | 0.79233987 | -1.5059278 | 0.13208572 | NA |
| miR-297b-3p | 1.33130908 | -1.1932066 | 0.79233987 | -1.5059278 | 0.13208572 | NA |
| miR-484 | 9.28120819 | 0.53674452 | 0.35672394 | 1.50464957 | 0.1324142 | 0.38116247 |
| miR-465b-5p | 264.958432 | -0.2826074 | 0.18787349 | -1.5042428 | 0.13251885 | 0.38116247 |
| miR-130c | 0.51783709 | 2.561873 | 1.71053053 | 1.49770668 | 0.13420948 | NA |
| miR-19a-3p | 9.29601073 | -0.5724972 | 0.38594847 | -1.4833515 | 0.13798105 | 0.39234841 |
| miR-494-3p | 0.97285533 | -1.580337 | 1.06789553 | -1.4798611 | 0.13891032 | NA |
| miR-664-5p | 7.04281646 | 1.01597096 | 0.68899265 | 1.47457444 | 0.14032699 | 0.39234841 |
| miR-30e-5p | 460.07374 | -0.2883887 | 0.19572194 | -1.4734614 | 0.14062667 | 0.39234841 |
| miR-499-5p | 8.25906604 | -0.7180635 | 0.49116232 | -1.4619679 | 0.14375001 | 0.39348993 |
| miR-470-5p | 170.313899 | -0.393621 | 0.2693121 | -1.4615793 | 0.14385654 | 0.39348993 |
| miR-741-5p | 5.93095736 | -0.7974593 | 0.54938114 | -1.4515593 | 0.14662418 | 0.39716648 |

| | | | | | | |
|---|---|---|---|---|---|---|
| miR-200a-5p | 36.1853401 | 0.33937257 | 0.23565178 | 1.44014428 | 0.14982658 | 0.39842795 |
| miR-29b-1-5p | 4.69481142 | 0.71160677 | 0.49450956 | 1.4390152 | 0.15014622 | 0.39842795 |
| miR-871-5p | 112.685363 | -0.2833302 | 0.19797878 | -1.4311138 | 0.1523976 | 0.39842795 |
| miR-410-5p | 0.41917989 | 2.39131707 | 1.67118453 | 1.43091145 | 0.1524556 | NA |
| miR-139-5p | 20.0585506 | 0.36988741 | 0.25871616 | 1.42970356 | 0.15280212 | 0.39842795 |
| miR-199a-5p | 18.059427 | 0.58958725 | 0.41584796 | 1.41779523 | 0.15625056 | 0.4031395 |
| miR-467c-5p | 24.5934115 | -0.3231078 | 0.22858163 | -1.4135334 | 0.15749895 | 0.4031395 |
| miR-433-3p | 0.60214941 | 2.16274869 | 1.54838071 | 1.39678096 | 0.16247945 | NA |
| let-7d-5p | 444.556237 | 0.32026968 | 0.22973454 | 1.39408593 | 0.16329166 | 0.41206855 |
| miR-1247-5p | 6.36450159 | -0.773608 | 0.5557777 | -1.3919377 | 0.16394125 | 0.41206855 |
| miR-7229-3p | 1.36654495 | -1.3939655 | 1.01721885 | -1.3703693 | 0.17057166 | NA |
| miR-155-5p | 21.654104 | 0.33991086 | 0.25082847 | 1.35515263 | 0.17536893 | 0.43342562 |
| miR-124-3p | 23.0995389 | -0.4468841 | 0.32990125 | -1.3545997 | 0.17554514 | 0.43342562 |
| miR-7241-3p | 5.16525278 | -0.7382526 | 0.55313397 | -1.3346722 | 0.18198364 | 0.44369875 |
| miR-8094 | 6.96833326 | 0.83188645 | 0.62466502 | 1.3317321 | 0.18294824 | 0.44369875 |
| miR-3066-3p | 0.6182335 | 2.05513383 | 1.54557766 | 1.32968655 | 0.18362157 | NA |
| miR-5100 | 4.75260432 | -0.6234269 | 0.46976769 | -1.3270961 | 0.1844769 | 0.44369875 |
| miR-382-3p | 0.52465532 | -1.650113 | 1.24976452 | -1.3203391 | 0.18672183 | NA |
| miR-409-5p | 1.73225784 | -1.0917138 | 0.83632685 | -1.3053674 | 0.19176779 | NA |
| miR-8104 | 0.78352603 | 1.53867893 | 1.18186556 | 1.3019069 | 0.19294821 | NA |
| miR-674-3p | 7.58444554 | -0.5558133 | 0.42870194 | -1.2965029 | 0.1948023 | 0.46452855 |
| miR-7653-5p | 1.18652672 | 1.28659349 | 1.00045571 | 1.28600745 | 0.19844046 | NA |
| miR-202-5p | 5.17152346 | -0.7129873 | 0.55595948 | -1.2824448 | 0.19968667 | 0.46966246 |
| miR-7672-3p | 0.4978961 | 1.5947278 | 1.24444684 | 1.28147522 | 0.2000268 | NA |
| miR-34c-3p | 113.072734 | -0.2827167 | 0.22150177 | -1.2763634 | 0.2018271 | 0.46966246 |
| miR-471-3p | 11.2780049 | -0.4482183 | 0.35130708 | -1.275859 | 0.20200536 | 0.46966246 |
| miR-93-3p | 1.71261301 | -0.8950061 | 0.70656889 | -1.2666933 | 0.20526496 | NA |
| miR-3082-3p | 0.4230526 | 1.73387506 | 1.37073605 | 1.26492263 | 0.20589906 | NA |
| let-7c-1-3p | 4.4121518 | 0.63599422 | 0.50363904 | 1.2627977 | 0.20666189 | 0.4713694 |
| miR-143-5p | 4.35062136 | 0.67985846 | 0.54063106 | 1.25752756 | 0.20856267 | 0.4713694 |
| miR-92a-3p | 1244.50594 | -0.3406258 | 0.27125078 | -1.2557597 | 0.20920312 | 0.4713694 |
| miR-127-3p | 6.39426829 | 0.55261095 | 0.44212247 | 1.24990471 | 0.21133436 | 0.4713694 |
| miR-34a-5p | 7.65238416 | 0.42604415 | 0.34185769 | 1.24626171 | 0.21266833 | 0.4713694 |
| miR-98-5p | 118.346912 | 0.34629682 | 0.27799494 | 1.24569469 | 0.2128765 | 0.4713694 |
| miR-5122 | 0.47298556 | 1.85477115 | 1.49764024 | 1.23846242 | 0.21554465 | NA |
| miR-598-3p | 1.24755901 | -1.1052035 | 0.8991361 | -1.2291838 | 0.21900292 | NA |
| miR-5113 | 0.50383365 | 1.69921009 | 1.38280624 | 1.22881286 | 0.21914198 | NA |
| miR-8112 | 0.87655675 | 1.6143259 | 1.32448432 | 1.21883353 | 0.22290738 | NA |
| miR-6900-3p | 0.38002155 | 2.17111843 | 1.78875907 | 1.21375677 | 0.22484062 | NA |
| miR-22-5p | 5.70620647 | 0.5795917 | 0.47788291 | 1.21283203 | 0.22519405 | 0.49471763 |
| miR-7233-5p | 1.53587885 | 1.27111199 | 1.04851692 | 1.21229516 | 0.22539941 | NA |
| miR-301a-3p | 0.76485853 | -1.476529 | 1.22373229 | -1.2065784 | 0.22759454 | NA |
| miR-463-5p | 43.4690051 | -0.4233076 | 0.3559943 | -1.1890852 | 0.23440613 | 0.5109321 |
| miR-200c-5p | 0.81292833 | 1.48107623 | 1.24851094 | 1.18627413 | 0.23551406 | NA |
| miR-692 | 1.28049907 | 0.98493485 | 0.83073673 | 1.18561611 | 0.23577394 | NA |
| miR-210-3p | 2.02881387 | 0.88059016 | 0.74461402 | 1.18261292 | 0.23696259 | NA |

| | | | | | | |
|---|---|---|---|---|---|---|
| miR-682 | 1.34164129 | 0.94001304 | 0.79589069 | 1.18108309 | 0.23756972 | NA |
| miR-669o-3p | 1.15860234 | -0.99365 | 0.84627288 | -1.1741485 | 0.24033556 | NA |
| miR-29c-5p | 3.33854126 | 0.66953541 | 0.57144708 | 1.17164901 | 0.241338 | 0.52196359 |
| miR-1893 | 0.78223822 | 1.33275943 | 1.14062603 | 1.16844557 | 0.24262708 | NA |
| miR-497a-5p | 2.84128118 | 0.76110984 | 0.65331893 | 1.16498972 | 0.24402315 | NA |
| miR-7230-5p | 4.53087599 | -0.599195 | 0.51602677 | -1.1611704 | 0.24557262 | 0.52703662 |
| miR-483-3p | 1.53197174 | -1.0531814 | 0.92141829 | -1.1430003 | 0.25303848 | NA |
| miR-7232-5p | 0.90266988 | -1.3924499 | 1.21839988 | -1.1428513 | 0.25310032 | NA |
| miR-106b-3p | 49.599141 | -0.2733833 | 0.24025726 | -1.1378775 | 0.25517162 | 0.54345711 |
| miR-98-3p | 0.74631744 | -1.4324855 | 1.27531593 | -1.1232397 | 0.26133569 | NA |
| miR-184-3p | 252.452 | -0.2615194 | 0.23467155 | -1.1144063 | 0.26510494 | 0.54867102 |
| miR-6516-5p | 2.95376449 | 0.83504643 | 0.7509154 | 1.11203796 | 0.26612183 | 0.54867102 |
| let-7e-5p | 45.272878 | 0.37487052 | 0.3372894 | 1.11142099 | 0.26638718 | 0.54867102 |
| miR-7210-5p | 12.7019253 | -0.4679915 | 0.42211589 | -1.10868 | 0.26756824 | 0.54867102 |
| miR-9-5p | 47.6867293 | -0.2558525 | 0.23078987 | -1.1085951 | 0.26760491 | 0.54867102 |
| miR-181c-5p | 12.7807975 | -0.3966039 | 0.35911211 | -1.1044013 | 0.2694191 | 0.54867102 |
| miR-541-5p | 12.905583 | 0.35982137 | 0.327681 | 1.09808431 | 0.27216768 | 0.55025204 |
| miR-181b-1-3p | 0.51361358 | 1.52042529 | 1.38695099 | 1.09623577 | 0.27297561 | NA |
| miR-872-3p | 1.27070459 | 0.96282026 | 0.88767748 | 1.084651 | 0.27807626 | NA |
| miR-101a-3p | 54.2783753 | 0.33397603 | 0.30801608 | 1.08428118 | 0.27824015 | 0.55848202 |
| miR-331-3p | 1.7878912 | -0.8839197 | 0.82303847 | -1.0739713 | 0.28283557 | NA |
| miR-33-3p | 0.39586083 | -1.6299336 | 1.52693148 | -1.0674569 | 0.28576555 | NA |
| miR-466e-3p | 1.94753007 | -0.6805418 | 0.63766212 | -1.0672451 | 0.28586116 | NA |
| miR-7670-5p | 0.40361552 | 1.51147334 | 1.41769935 | 1.06614519 | 0.28635802 | NA |
| miR-181a-5p | 99.6704484 | 0.27125514 | 0.25618554 | 1.05882297 | 0.28968041 | 0.57729167 |
| miR-148a-3p | 1703.16091 | 0.24384806 | 0.23169396 | 1.05245758 | 0.29258966 | 0.57895401 |
| miR-3066-5p | 2.27027344 | -0.7264975 | 0.69105692 | -1.0512846 | 0.29312788 | NA |
| miR-191-5p | 2286.5095 | -0.1439485 | 0.13776141 | -1.0449119 | 0.29606367 | 0.58170257 |
| miR-101b-3p | 45.7437175 | 0.22077035 | 0.21308481 | 1.03606802 | 0.30017041 | 0.58467524 |
| miR-21a-3p | 3.2233113 | 0.61530168 | 0.59584745 | 1.03264968 | 0.30176787 | 0.58467524 |
| miR-7210-3p | 4.81106281 | 0.48067669 | 0.46865211 | 1.02565778 | 0.30505292 | 0.5869639 |
| miR-874-3p | 0.68317914 | 1.31549692 | 1.29636013 | 1.01476194 | 0.31021932 | NA |
| miR-5125 | 1.6719044 | 0.95104383 | 0.94195237 | 1.00965172 | 0.31266218 | NA |
| miR-27b-5p | 5.66832158 | 0.49319933 | 0.48900859 | 1.00856987 | 0.31318096 | 0.59197088 |
| miR-409-3p | 7.35694906 | -0.4212268 | 0.41828323 | -1.0070374 | 0.31391682 | 0.59197088 |
| miR-147-5p | 3.4978524 | -0.5898922 | 0.58589531 | -1.0068218 | 0.3140204 | 0.59197088 |
| miR-466p-3p | 1.92203196 | -0.6346712 | 0.64036489 | -0.9911088 | 0.32163248 | NA |
| miR-466a-3p | 1.92203196 | -0.6346712 | 0.64036489 | -0.9911088 | 0.32163248 | NA |
| miR-223-5p | 8.97129253 | 0.40802656 | 0.41283207 | 0.98835965 | 0.32297654 | 0.60402969 |
| miR-6917-3p | 0.62500195 | 1.07554043 | 1.09037194 | 0.98639775 | 0.32393797 | NA |
| miR-744-5p | 8.05018161 | 0.42691499 | 0.4335265 | 0.98474947 | 0.32474715 | 0.60402969 |
| miR-7654-5p | 0.50610745 | 1.54757974 | 1.58231047 | 0.97805063 | 0.32804929 | NA |
| miR-6340 | 0.54852226 | -1.5233122 | 1.55874705 | -0.9772671 | 0.32843693 | NA |
| miR-451a | 25.8966956 | -0.2814627 | 0.28861132 | -0.9752309 | 0.32944574 | 0.60871101 |
| miR-7241-5p | 0.58504353 | -1.3205771 | 1.35487572 | -0.9746851 | 0.3297165 | NA |
| miR-148b-5p | 0.82867832 | 1.17474933 | 1.2093434 | 0.97139433 | 0.33135195 | NA |

| | | | | | | |
|---|---|---|---|---|---|---|
| miR-7653-3p | 0.67949228 | 1.63683755 | 1.69313251 | 0.966751 | 0.33366853 | NA |
| miR-674-5p | 2.00370529 | 0.73097972 | 0.763668 | 0.95719569 | 0.33846849 | NA |
| miR-132-3p | 1.70077265 | -0.7665684 | 0.80637809 | -0.9506315 | 0.34179148 | NA |
| miR-31-5p | 104.211405 | 0.2255009 | 0.23733498 | 0.95013765 | 0.34204231 | 0.62782767 |
| miR-7657-3p | 0.31384961 | 1.95749885 | 2.06961162 | 0.94582907 | 0.34423577 | NA |
| miR-543-3p | 0.79746738 | 1.08397741 | 1.14733336 | 0.94477982 | 0.3447713 | NA |
| miR-132-5p | 3.52750056 | -0.580391 | 0.61768466 | -0.9396235 | 0.34741071 | 0.62893529 |
| miR-23b-5p | 1.97688937 | 0.64437334 | 0.68786653 | 0.93677089 | 0.34887642 | NA |
| miR-342-3p | 38.1863586 | -0.2105686 | 0.22510601 | -0.9354199 | 0.34957195 | 0.62893529 |
| miR-511-5p | 0.64556384 | 1.17224153 | 1.2542524 | 0.93461374 | 0.34998739 | NA |
| miR-190b-5p | 9.53491127 | -0.4309351 | 0.46174881 | -0.9332673 | 0.35068196 | 0.62893529 |
| miR-30b-3p | 2.79443292 | 0.52454315 | 0.56310436 | 0.93152031 | 0.35158448 | NA |
| miR-196b-3p | 3.34725695 | 0.62101203 | 0.6684249 | 0.92906777 | 0.35285396 | 0.62893529 |
| miR-1948-5p | 0.32628164 | 1.96618063 | 2.11841155 | 0.92813912 | 0.35333541 | NA |
| miR-10b-5p | 6411.12022 | -0.2176952 | 0.23483379 | -0.9270184 | 0.35391699 | 0.62893529 |
| let-7e-3p | 0.32624036 | 1.51980285 | 1.64406519 | 0.92441763 | 0.35526892 | NA |
| miR-193a-5p | 0.33893175 | 1.49873512 | 1.6288476 | 0.92011991 | 0.3575101 | NA |
| miR-345-3p | 4.10599678 | 0.55971504 | 0.61163717 | 0.91510959 | 0.3601341 | 0.63317992 |
| miR-1843b-5p | 6.91778818 | 0.34730178 | 0.38008105 | 0.91375714 | 0.36084447 | 0.63317992 |
| miR-99b-5p | 154.596475 | 0.26409628 | 0.29135988 | 0.90642638 | 0.36471021 | 0.63360364 |
| miR-26a-2-3p | 1.58516365 | 0.6644422 | 0.73525282 | 0.90369214 | 0.36615867 | NA |
| miR-16-1-3p | 0.88160467 | -0.9368962 | 1.03816338 | -0.9024555 | 0.36681495 | NA |
| miR-214-3p | 3.7475493 | 0.48266582 | 0.53513352 | 0.901954 | 0.3670813 | 0.63360364 |
| miR-6962-3p | 0.2925918 | 1.46042346 | 1.62165902 | 0.9005737 | 0.36781502 | NA |
| miR-5116 | 0.31605621 | 1.48047499 | 1.64717202 | 0.89879804 | 0.36876024 | NA |
| miR-30d-5p | 1714.27076 | -0.172775 | 0.19357377 | -0.892554 | 0.37209605 | 0.63360364 |
| miR-142a-5p | 130.748837 | -0.2960737 | 0.33196546 | -0.8918811 | 0.37245666 | 0.63360364 |
| miR-30c-5p | 1172.74605 | -0.1692559 | 0.18989557 | -0.8913105 | 0.37276264 | 0.63360364 |
| miR-465d-5p | 1.01653111 | -0.9674027 | 1.09361368 | -0.8845927 | 0.37637632 | NA |
| miR-1249-3p | 4.24629671 | -0.4702413 | 0.53331177 | -0.881738 | 0.37791851 | 0.63360364 |
| miR-872-5p | 83.8851766 | -0.2497606 | 0.28390254 | -0.8797406 | 0.37899984 | 0.63360364 |
| let-7i-5p | 426.03059 | 0.15063675 | 0.17131985 | 0.87927204 | 0.37925379 | 0.63360364 |
| miR-127-5p | 0.84213131 | -0.9536396 | 1.08701328 | -0.8773026 | 0.38032228 | NA |
| miR-378a-3p | 25.2098307 | 0.25865765 | 0.29563197 | 0.87493126 | 0.38161131 | 0.63374736 |
| miR-450a-5p | 1.81641434 | 0.67081436 | 0.76886595 | 0.87247246 | 0.38295069 | NA |
| miR-669o-5p | 6.42570623 | 0.35622107 | 0.41294082 | 0.86264436 | 0.38833303 | 0.63877724 |
| miR-878-5p | 47.8104951 | 0.1889956 | 0.21949832 | 0.86103438 | 0.38921911 | 0.63877724 |
| miR-125a-3p | 1.40855534 | 0.69420674 | 0.80708521 | 0.86014059 | 0.38971155 | NA |
| miR-25-3p | 580.868545 | 0.092424 | 0.10809492 | 0.85502627 | 0.39253661 | 0.6395048 |
| miR-431-5p | 0.55693436 | 1.02203904 | 1.20171766 | 0.85048183 | 0.39505726 | NA |
| miR-345-5p | 4.60504135 | 0.41815528 | 0.49222031 | 0.8495287 | 0.39558717 | 0.6395048 |
| miR-339-3p | 1.89296104 | -0.6080949 | 0.71724118 | -0.8478249 | 0.3965355 | NA |
| miR-92b-3p | 1.05644964 | -0.7187488 | 0.84861974 | -0.8469621 | 0.39701624 | NA |
| miR-378c | 0.37011566 | 1.16175668 | 1.37281775 | 0.84625704 | 0.39740937 | NA |
| miR-218-5p | 12.9384432 | 0.30694998 | 0.36299998 | 0.84559227 | 0.39778024 | 0.6395048 |
| miR-7217-5p | 17.077987 | -0.2135047 | 0.25305434 | -0.8437109 | 0.39883095 | 0.6395048 |

| | | | | | | |
|---|---|---|---|---|---|---|
| miR-1955-5p | 0.40116737 | -1.5303517 | 1.81776258 | -0.8418875 | 0.3998509 | NA |
| miR-25-5p | 1.29388924 | 0.91708216 | 1.09287335 | 0.8391477 | 0.40138643 | NA |
| miR-15a-5p | 15.4443855 | -0.2797337 | 0.33583969 | -0.8329381 | 0.40487962 | 0.64549379 |
| miR-1930-5p | 0.2611588 | 1.77833171 | 2.16960737 | 0.81965601 | 0.41241224 | NA |
| miR-221-5p | 6.87807047 | 0.30847757 | 0.3787936 | 0.81436849 | 0.41543388 | 0.65125818 |
| miR-191-3p | 13.5980374 | -0.2340302 | 0.28781737 | -0.8131206 | 0.41614891 | 0.65125818 |
| miR-467a-3p | 10.716942 | -0.3904557 | 0.48101678 | -0.8117298 | 0.41694669 | 0.65125818 |
| miR-1981-5p | 3.5186126 | -0.4042172 | 0.49891801 | -0.8101877 | 0.41783231 | 0.65125818 |
| miR-7073-3p | 0.41814032 | 1.16256299 | 1.44661676 | 0.8036427 | 0.42160336 | NA |
| miR-16-5p | 964.430921 | -0.1345105 | 0.16758611 | -0.8026352 | 0.42218559 | 0.65438767 |
| miR-186-3p | 0.4008657 | 1.31400521 | 1.64961091 | 0.79655464 | 0.42570973 | NA |
| miR-7648-3p | 0.33354466 | 1.98344241 | 2.50522102 | 0.79172352 | 0.4285219 | NA |
| miR-134-5p | 0.40822388 | 1.14142659 | 1.4476264 | 0.78848147 | 0.43041513 | NA |
| miR-7658-5p | 1.04772463 | -1.1519535 | 1.4626095 | -0.7876015 | 0.43092982 | NA |
| let-7f-5p | 3768.27698 | 0.23261036 | 0.29627638 | 0.78511275 | 0.43238747 | 0.66649781 |
| miR-7021-3p | 0.36612178 | 1.39556729 | 1.79690551 | 0.77665035 | 0.43736509 | NA |
| miR-10b-3p | 24.0816276 | 0.19016734 | 0.24524847 | 0.77540685 | 0.43809929 | 0.67101231 |
| miR-29b-3p | 27.6961084 | 0.30177689 | 0.39254859 | 0.76876315 | 0.44203393 | 0.67101231 |
| miR-221-3p | 117.807058 | 0.18313102 | 0.23864738 | 0.76737073 | 0.44286113 | 0.67101231 |
| miR-678 | 0.21619873 | 1.50212132 | 1.96423122 | 0.76473753 | 0.44442786 | NA |
| miR-335-5p | 8.53785686 | -0.2706191 | 0.35426732 | -0.7638838 | 0.44493648 | 0.67101231 |
| miR-669f-3p | 0.28771976 | -1.6284302 | 2.14198436 | -0.7602437 | 0.4471089 | NA |
| miR-10a-5p | 9297.54226 | -0.1841127 | 0.24345137 | -0.7562607 | 0.44949289 | 0.67423934 |
| miR-7236-3p | 0.29958608 | 1.41886502 | 1.88130508 | 0.75419188 | 0.45073401 | NA |
| miR-429-3p | 318.054586 | -0.177471 | 0.23760945 | -0.746902 | 0.45512272 | 0.67903336 |
| miR-151-5p | 246.07656 | -0.1309487 | 0.1764056 | -0.7423158 | 0.45789604 | 0.6795372 |
| miR-1981-3p | 1.88632736 | 0.57148386 | 0.7707992 | 0.7414173 | 0.45844046 | NA |
| miR-201-5p | 0.43256379 | -1.3216559 | 1.79489831 | -0.7363403 | 0.46152365 | NA |
| miR-701-5p | 1.27229612 | -0.5771833 | 0.78488886 | -0.7353695 | 0.46211449 | NA |
| let-7b-3p | 2.75590889 | 0.47022343 | 0.64010682 | 0.7346015 | 0.46258222 | NA |
| miR-141-5p | 2.29234359 | 0.61197214 | 0.83755098 | 0.73066853 | 0.46498164 | NA |
| miR-211-5p | 0.48788573 | -1.0328826 | 1.42235841 | -0.7261761 | 0.46773084 | NA |
| miR-146b-5p | 35.4548425 | 0.14759503 | 0.20424999 | 0.72261954 | 0.46991366 | 0.69368207 |
| miR-3068-5p | 2.2426751 | 0.52318767 | 0.72445822 | 0.72217784 | 0.47018515 | NA |
| miR-379-3p | 0.54729305 | -0.9517214 | 1.32659561 | -0.7174164 | 0.47311721 | NA |
| let-7a-5p | 3159.07868 | 0.20810374 | 0.29087931 | 0.71542985 | 0.47434347 | 0.69632985 |
| miR-146a-5p | 154.398342 | 0.1515662 | 0.21298697 | 0.71162192 | 0.47669893 | 0.69632985 |
| miR-32-3p | 2.48659838 | -0.4849229 | 0.68192223 | -0.7111118 | 0.47701498 | NA |
| miR-881-5p | 28.3074452 | -0.2313344 | 0.32709806 | -0.7072324 | 0.47942204 | 0.69666016 |
| miR-20a-5p | 37.4484559 | -0.1606464 | 0.22915517 | -0.7010378 | 0.48327944 | 0.69806435 |
| miR-3058-5p | 0.27165624 | 1.33386029 | 1.91051571 | 0.69816767 | 0.48507234 | NA |
| miR-204-5p | 13.4940351 | -0.2668389 | 0.38247919 | -0.6976559 | 0.48539241 | 0.69806435 |
| miR-107-3p | 2.62511464 | -0.3913529 | 0.56676415 | -0.6905039 | 0.48987736 | NA |
| miR-3475-3p | 1.70783566 | -0.5293382 | 0.76816802 | -0.6890917 | 0.49076559 | NA |
| miR-686 | 0.23578253 | 1.53260477 | 2.23867153 | 0.68460457 | 0.49359348 | NA |
| miR-30a-5p | 1950.27346 | -0.1244491 | 0.18198483 | -0.6838435 | 0.49407401 | 0.70690589 |

| | | | | | | |
|---|---|---|---|---|---|---|
| miR-295-5p | 0.2252144 | 1.53077151 | 2.26521838 | 0.67577215 | 0.49918532 | NA |
| miR-195a-5p | 28.1012138 | 0.17692452 | 0.26193571 | 0.67545015 | 0.49938981 | 0.70816968 |
| miR-17-5p | 4.21642922 | 0.31634084 | 0.4690446 | 0.67443659 | 0.50003379 | 0.70816968 |
| miR-467b-5p | 2.16390764 | -0.4260579 | 0.63522686 | -0.6707177 | 0.50240037 | NA |
| miR-194-2-3p | 0.65778338 | 0.9584527 | 1.4334753 | 0.66862171 | 0.50373682 | NA |
| miR-215-3p | 0.3298134 | -1.21708 | 1.82532081 | -0.6667759 | 0.50491527 | NA |
| miR-125b-1-3p | 1.63591017 | -0.6500715 | 0.99026973 | -0.656459 | 0.51152881 | NA |
| miR-181d-5p | 8.08881438 | 0.24116696 | 0.3687612 | 0.65399223 | 0.51311681 | 0.72065561 |
| miR-28a-5p | 31.7864995 | -0.1606582 | 0.24618277 | -0.6525972 | 0.51401601 | 0.72065561 |
| miR-130b-3p | 2.1409677 | 0.39308147 | 0.61244799 | 0.64182016 | 0.52098996 | NA |
| miR-652-3p | 18.7331529 | -0.1978084 | 0.31025345 | -0.6375703 | 0.52375347 | 0.72173299 |
| miR-672-5p | 35.937379 | 0.17445749 | 0.27367886 | 0.6374533 | 0.52382962 | 0.72173299 |
| miR-361-5p | 14.2807521 | 0.19545046 | 0.30694209 | 0.63676657 | 0.5242769 | 0.72173299 |
| miR-467d-5p | 11.6099358 | -0.2495352 | 0.39268755 | -0.6354547 | 0.52513189 | 0.72173299 |
| miR-3110-5p | 0.32781634 | 1.15231465 | 1.81560492 | 0.63467257 | 0.52564199 | NA |
| miR-300-3p | 0.5049917 | 0.8065998 | 1.27574853 | 0.63225611 | 0.52721954 | NA |
| miR-135a-5p | 11.0420065 | 0.21282102 | 0.3388285 | 0.62810838 | 0.52993294 | 0.72476123 |
| miR-376b-5p | 0.56811 | 0.89800292 | 1.44003089 | 0.62359977 | 0.53289046 | NA |
| miR-582-5p | 0.35066863 | -1.3217033 | 2.12889986 | -0.6208387 | 0.53470579 | NA |
| miR-669m-3p | 0.30564666 | -1.5712743 | 2.53725643 | -0.6192808 | 0.53573136 | NA |
| miR-362-5p | 2.29164722 | 0.4203398 | 0.6950522 | 0.60476005 | 0.54533844 | NA |
| miR-467e-5p | 59.4961354 | -0.1483025 | 0.24546834 | -0.6041615 | 0.54573628 | 0.73876324 |
| miR-467a-5p | 21.5132456 | -0.2608821 | 0.43343421 | -0.6018955 | 0.54724373 | 0.73876324 |
| miR-100-5p | 80.5646163 | 0.16285498 | 0.27115938 | 0.60058765 | 0.54811466 | 0.73876324 |
| miR-135a-2-3p | 0.52409556 | 0.85269507 | 1.42994775 | 0.59631205 | 0.55096678 | NA |
| miR-339-5p | 5.13396336 | 0.29043592 | 0.48967413 | 0.59312081 | 0.5531003 | 0.74189896 |
| miR-742-3p | 0.36912133 | -1.0659759 | 1.80449341 | -0.5907342 | 0.55469856 | NA |
| miR-7663-5p | 0.18504506 | 1.40046964 | 2.37393233 | 0.58993663 | 0.55523314 | NA |
| miR-152-5p | 0.30578707 | 1.09875802 | 1.87095211 | 0.58727212 | 0.55702096 | NA |
| miR-342-5p | 1.35138886 | 0.51760795 | 0.88921249 | 0.58209703 | 0.56050132 | NA |
| miR-125b-2-3p | 5.72303747 | 0.26178674 | 0.45554258 | 0.57467018 | 0.56551438 | 0.75492111 |
| miR-323-3p | 0.79985677 | -0.6928749 | 1.21096791 | -0.5721662 | 0.56720937 | NA |
| miR-187-5p | 1.03804169 | 0.58581859 | 1.03337787 | 0.56689678 | 0.57078431 | NA |
| miR-1938 | 0.41316232 | 0.82416237 | 1.47059427 | 0.56042811 | 0.57518746 | NA |
| miR-1191b-3p | 0.26661165 | 1.21268001 | 2.16658479 | 0.55971962 | 0.5756707 | NA |
| miR-200a-3p | 168.750113 | 0.13737342 | 0.24666217 | 0.55692942 | 0.57757565 | 0.76303526 |
| miR-669a-3-3p | 0.27037435 | -1.4253266 | 2.56611057 | -0.5554424 | 0.57859211 | NA |
| miR-369-3p | 10.0901926 | -0.1847634 | 0.33342255 | -0.554142 | 0.5794817 | 0.76303526 |
| miR-144-5p | 3.35011903 | -0.3672981 | 0.66337538 | -0.5536807 | 0.57979741 | 0.76303526 |
| miR-700-3p | 0.71296427 | -0.59081 | 1.06768879 | -0.5533542 | 0.58002092 | NA |
| miR-485-5p | 1.52821853 | 0.46520144 | 0.844134 | 0.55109904 | 0.58156578 | NA |
| miR-329-5p | 0.25364671 | -1.4259241 | 2.61136999 | -0.5460445 | 0.58503537 | NA |
| miR-668-5p | 0.34980472 | 0.8037032 | 1.47669336 | 0.54425869 | 0.58626347 | NA |
| miR-653-5p | 0.32144972 | -0.8937802 | 1.66823937 | -0.5357626 | 0.59212265 | NA |
| miR-135b-5p | 0.52953236 | 0.65726411 | 1.23007928 | 0.53432662 | 0.59311558 | NA |
| miR-26b-3p | 2.62424857 | 0.43532489 | 0.81681173 | 0.53295622 | 0.59406389 | NA |

| | | | | | | |
|---|---|---|---|---|---|---|
| miR-21a-5p | 5303.45375 | 0.068407 | 0.12969424 | 0.52744826 | 0.59788234 | 0.7821449 |
| miR-3057-5p | 0.25032956 | 1.17868651 | 2.23593372 | 0.52715628 | 0.59808507 | NA |
| miR-495-3p | 0.52261326 | -0.7608077 | 1.44518282 | -0.5264439 | 0.59857979 | NA |
| miR-8101 | 0.22938923 | 1.16841485 | 2.22618021 | 0.52485187 | 0.59968617 | NA |
| miR-99a-5p | 873.50738 | -0.0976882 | 0.18624713 | -0.5245087 | 0.59992476 | 0.7821449 |
| miR-185-3p | 0.18225778 | 1.24655945 | 2.379311 | 0.52391614 | 0.60033687 | NA |
| miR-182-3p | 0.91708186 | 0.57862079 | 1.10980547 | 0.52137136 | 0.6021081 | NA |
| miR-708-3p | 0.65905994 | 0.65002843 | 1.26108057 | 0.51545353 | 0.60623613 | NA |
| miR-6368 | 0.34424508 | 1.24726218 | 2.4212834 | 0.51512441 | 0.60646609 | NA |
| miR-17-3p | 3.03173915 | -0.3181079 | 0.61794594 | -0.5147827 | 0.60670484 | 0.78507217 |
| miR-328-3p | 16.9779311 | -0.1969728 | 0.38379836 | -0.5132195 | 0.60779781 | 0.78507217 |
| miR-6993-3p | 0.48973864 | -0.9427441 | 1.84540033 | -0.5108616 | 0.60944798 | NA |
| miR-101b-5p | 0.21938686 | 1.0069445 | 1.97423973 | 0.51004165 | 0.61002228 | NA |
| miR-181a-1-3p | 0.80277596 | -0.623656 | 1.23095616 | -0.5066435 | 0.61240495 | NA |
| miR-669h-5p | 0.21433016 | 1.38100439 | 2.72892945 | 0.50606086 | 0.61281392 | NA |
| miR-7233-3p | 2.43650906 | -0.3856927 | 0.76356862 | -0.5051186 | 0.61347552 | NA |
| miR-411-5p | 12.5873365 | 0.19171797 | 0.38195713 | 0.50193584 | 0.61571265 | 0.79163055 |
| miR-7079-5p | 0.45948387 | 1.3824957 | 2.77699575 | 0.49783861 | 0.6185978 | NA |
| miR-7242-3p | 0.57769147 | -0.6265234 | 1.26577626 | -0.4949716 | 0.62062014 | NA |
| miR-203-3p | 54.5452797 | 0.09427292 | 0.19057719 | 0.4946705 | 0.62083273 | 0.79239907 |
| miR-322-3p | 2.11238872 | 0.32865053 | 0.66574349 | 0.49365941 | 0.62154674 | NA |
| miR-463-3p | 5.37950108 | -0.2310751 | 0.46868277 | -0.493031 | 0.62199067 | 0.79239907 |
| miR-504-5p | 0.57824966 | -0.6878838 | 1.41754389 | -0.4852646 | 0.62748867 | NA |
| miR-215-5p | 0.59291528 | 0.6935578 | 1.43413082 | 0.48360846 | 0.62866377 | NA |
| miR-7036a-5p | 0.39664656 | 0.99967447 | 2.07155426 | 0.48257219 | 0.62939953 | NA |
| miR-702-3p | 0.4113301 | 0.99975524 | 2.07266477 | 0.4823526 | 0.62955549 | NA |
| miR-7222-5p | 0.2728697 | -0.9036155 | 1.88587245 | -0.4791498 | 0.63183204 | NA |
| miR-2137 | 0.6605868 | 0.70891035 | 1.47983024 | 0.47904843 | 0.63190417 | NA |
| miR-125a-5p | 423.301695 | -0.0986167 | 0.20598904 | -0.4787475 | 0.63211825 | 0.80164087 |
| miR-7221-5p | 0.38870769 | 0.77818056 | 1.62893074 | 0.47772477 | 0.63284611 | NA |
| miR-429-5p | 0.47482529 | 0.63482309 | 1.33120042 | 0.47688017 | 0.63344745 | NA |
| miR-7223-3p | 0.2373984 | -0.9321006 | 1.95702827 | -0.4762837 | 0.63387229 | NA |
| miR-1894-5p | 0.20393184 | 1.12348038 | 2.36959695 | 0.47412299 | 0.63541221 | NA |
| miR-150-3p | 1.15629559 | -0.5266911 | 1.11714182 | -0.471463 | 0.6373101 | NA |
| miR-7026-3p | 0.15573735 | 1.16893908 | 2.48012223 | 0.47132318 | 0.63740997 | NA |
| miR-30b-5p | 180.319339 | -0.107409 | 0.22924706 | -0.4685294 | 0.63940605 | 0.80721397 |
| miR-210-5p | 0.55787547 | 0.73920112 | 1.58904104 | 0.46518693 | 0.6417976 | NA |
| miR-128-1-5p | 0.78995666 | -0.5865004 | 1.26442594 | -0.4638472 | 0.64275724 | NA |
| miR-7214-3p | 4.1221763 | -0.2353514 | 0.50784235 | -0.4634339 | 0.64305339 | 0.80816169 |
| miR-7648-5p | 0.32120034 | 0.97794353 | 2.11229267 | 0.46297729 | 0.64338065 | NA |
| miR-369-5p | 2.11446869 | 0.31602917 | 0.68734399 | 0.45978313 | 0.64567189 | NA |
| miR-676-3p | 22.8725104 | -0.1325294 | 0.28873881 | -0.4589941 | 0.64623842 | 0.8085225 |
| miR-6928-5p | 0.14105381 | 1.16905029 | 2.54807047 | 0.45879826 | 0.64637905 | NA |
| miR-203-5p | 1.37265492 | 0.41649404 | 0.93550273 | 0.44520879 | 0.65616887 | NA |
| let-7a-1-3p | 1.67186989 | 0.34298746 | 0.77056279 | 0.44511293 | 0.65623815 | NA |
| miR-6539 | 0.13707226 | 1.14246868 | 2.56802022 | 0.44488305 | 0.65640427 | NA |

| | | | | | | |
|---|---|---|---|---|---|---|
| miR-1198-3p | 0.20329778 | 1.21989373 | 2.74596381 | 0.44424975 | 0.65686203 | NA |
| miR-378b | 0.13668932 | 1.13991315 | 2.57021905 | 0.44350817 | 0.65739821 | NA |
| miR-5119 | 0.13668932 | 1.13991315 | 2.57021905 | 0.44350817 | 0.65739821 | NA |
| miR-217-5p | 0.34300774 | 0.66880935 | 1.51844592 | 0.44045649 | 0.65960652 | NA |
| miR-140-5p | 7.16847016 | -0.1825035 | 0.41643578 | -0.4382511 | 0.66120425 | 0.81150433 |
| miR-669l-5p | 4.12971567 | 0.22466984 | 0.51432981 | 0.43682056 | 0.66224148 | 0.81150433 |
| miR-8108 | 0.89466752 | -0.4927403 | 1.12841324 | -0.4366666 | 0.66235319 | NA |
| miR-27a-3p | 73.5534303 | 0.13067825 | 0.30122682 | 0.4338201 | 0.66441908 | 0.81150433 |
| miR-423-5p | 30.0177129 | -0.2201301 | 0.50884757 | -0.4326053 | 0.66530156 | 0.81150433 |
| miR-96-5p | 25.4358504 | -0.1405835 | 0.32521403 | -0.4322799 | 0.66553801 | 0.81150433 |
| miR-669p-5p | 2.64389298 | 0.24053354 | 0.55875358 | 0.43048234 | 0.66684481 | NA |
| miR-7237-5p | 0.20894335 | 0.85915856 | 1.99631975 | 0.43037121 | 0.66692563 | NA |
| miR-6905-3p | 0.14885727 | 1.3120558 | 3.04996643 | 0.43018696 | 0.66705964 | NA |
| miR-298-5p | 1.12948402 | 0.3882177 | 0.90636639 | 0.42832314 | 0.66841588 | NA |
| miR-148a-5p | 11.3915157 | -0.1641307 | 0.38345374 | -0.4280325 | 0.66862745 | 0.81150433 |
| miR-743a-5p | 5.6953494 | -0.1974057 | 0.46171854 | -0.4275455 | 0.66898207 | 0.81150433 |
| miR-883a-3p | 45.0175904 | -0.1018116 | 0.24253741 | -0.4197768 | 0.67464854 | 0.81483525 |
| miR-376b-3p | 0.46948306 | -0.5435234 | 1.30578857 | -0.4162415 | 0.6772333 | NA |
| miR-377-3p | 0.21384099 | -1.1330092 | 2.72738497 | -0.4154196 | 0.67783476 | NA |
| miR-103-3p | 159.205521 | -0.0891121 | 0.21491607 | -0.4146366 | 0.67840795 | 0.81584405 |
| miR-1193-3p | 0.36436181 | -0.6789337 | 1.6466654 | -0.4123082 | 0.68011351 | NA |
| miR-6910-5p | 0.2610529 | -1.2475431 | 3.04728724 | -0.4093946 | 0.68225007 | NA |
| miR-6904-5p | 0.13576379 | 1.24633725 | 3.05102544 | 0.40849782 | 0.68290823 | NA |
| miR-27a-5p | 3.59787363 | 0.20457669 | 0.50264993 | 0.40699635 | 0.68401067 | 0.8190514 |
| miR-455-3p | 2.95864067 | 0.2368827 | 0.59238825 | 0.39987744 | 0.68924679 | 0.82179424 |
| miR-615-3p | 2.65826953 | -0.3149905 | 0.78876175 | -0.3993481 | 0.68963674 | NA |
| miR-7036b-3p | 0.1612893 | 1.17209443 | 2.93717002 | 0.3990557 | 0.68985216 | NA |
| miR-3101-3p | 0.16052341 | 1.16696512 | 2.94038217 | 0.39687532 | 0.6914594 | NA |
| miR-182-5p | 1071.47418 | 0.09980653 | 0.25230273 | 0.39558243 | 0.6924131 | 0.8220564 |
| miR-5106 | 0.23277744 | 0.89494288 | 2.26656684 | 0.39484513 | 0.69295718 | NA |
| miR-6236 | 0.15110411 | 1.17215911 | 2.99647595 | 0.39117921 | 0.69566477 | NA |
| miR-667-3p | 0.30450382 | -0.8483292 | 2.17158894 | -0.3906491 | 0.69605666 | NA |
| miR-7230-3p | 2.41444397 | -0.251592 | 0.65898387 | -0.3817877 | 0.70261883 | NA |
| miR-16-2-3p | 3.39650143 | -0.2163289 | 0.57203756 | -0.3781726 | 0.7053024 | 0.83381089 |
| miR-5046 | 0.2957561 | 0.6400526 | 1.70678974 | 0.37500378 | 0.70765766 | NA |
| miR-93-5p | 279.386861 | -0.0626387 | 0.16892905 | -0.3707991 | 0.7107872 | 0.83384934 |
| miR-6951-5p | 0.14818481 | -0.922326 | 2.50240915 | -0.3685752 | 0.71244437 | NA |
| miR-582-3p | 0.3477084 | 0.68938638 | 1.88699721 | 0.36533513 | 0.71486125 | NA |
| miR-6909-5p | 0.11491632 | 1.11354986 | 3.05330069 | 0.36470364 | 0.71533264 | NA |
| miR-6990-5p | 0.11453338 | 1.11132353 | 3.05334044 | 0.36396974 | 0.7158806 | NA |
| miR-324-5p | 1.61556568 | -0.2623808 | 0.72268837 | -0.3630621 | 0.71655848 | NA |
| miR-5136 | 0.11376749 | 1.10685814 | 3.05342033 | 0.3624978 | 0.71698007 | NA |
| miR-301b-3p | 0.36848246 | -0.5818323 | 1.60994014 | -0.3614 | 0.71780048 | NA |
| miR-190a-5p | 1.68057251 | -0.285741 | 0.80206573 | -0.3562563 | 0.72164864 | NA |
| miR-140-3p | 203.409622 | 0.08012677 | 0.22586626 | 0.35475317 | 0.72277452 | 0.83384934 |
| let-7g-5p | 2505.61508 | 0.07562399 | 0.21319097 | 0.35472415 | 0.72279626 | 0.83384934 |

| | | | | | | |
|---|---|---|---|---|---|---|
| miR-129-5p | 0.30171143 | -0.7571069 | 2.13536067 | -0.3545569 | 0.72292157 | NA |
| miR-501-5p | 0.43142251 | -0.562992 | 1.58879607 | -0.3543513 | 0.7230756 | NA |
| miR-27b-3p | 289.426169 | 0.0571856 | 0.16138243 | 0.35434836 | 0.72307783 | 0.83384934 |
| miR-145b | 0.37054985 | -0.52977 | 1.50944157 | -0.3509709 | 0.72561021 | NA |
| miR-8092 | 0.34056231 | -0.8664844 | 2.47901893 | -0.3495272 | 0.72669358 | NA |
| miR-361-3p | 7.47766385 | -0.1471449 | 0.42127677 | -0.3492832 | 0.72687672 | 0.83384934 |
| miR-351-5p | 3.04424938 | -0.2466155 | 0.70762294 | -0.3485125 | 0.72745529 | 0.83384934 |
| miR-468-5p | 4.01233732 | -0.1856224 | 0.53435855 | -0.3473743 | 0.72831017 | 0.83384934 |
| miR-137-3p | 0.10580439 | 1.05886158 | 3.05429285 | 0.34667978 | 0.7288319 | NA |
| miR-455-5p | 0.53053172 | -0.5298689 | 1.53094255 | -0.3461064 | 0.72926279 | NA |
| miR-293-3p | 2.56383523 | 0.24540747 | 0.7130205 | 0.34418011 | 0.73071084 | NA |
| miR-23b-3p | 119.638034 | 0.04603232 | 0.13411305 | 0.34323523 | 0.7314215 | 0.83384934 |
| miR-1198-5p | 6.82968617 | 0.14278384 | 0.42028188 | 0.33973353 | 0.73405721 | 0.83384934 |
| miR-669a-5p | 15.6957965 | 0.13301697 | 0.39332332 | 0.33818736 | 0.735222 | 0.83384934 |
| miR-6418-3p | 0.36292404 | 0.60381441 | 1.81243894 | 0.33315021 | 0.7390209 | NA |
| miR-26a-1-3p | 0.35651367 | 0.60362137 | 1.81573694 | 0.33243878 | 0.73955796 | NA |
| miR-384-5p | 0.24741571 | -0.6380753 | 1.92620421 | -0.3312605 | 0.74044775 | NA |
| let-7j | 0.55731838 | -0.4840923 | 1.46145898 | -0.3312391 | 0.74046391 | NA |
| miR-138-2-3p | 0.45360206 | 0.52106322 | 1.57367447 | 0.33111246 | 0.74055954 | NA |
| miR-421-3p | 2.93864994 | 0.1909812 | 0.57807817 | 0.33037262 | 0.74111843 | NA |
| miR-760-3p | 0.40303581 | 0.53124864 | 1.61468064 | 0.32901159 | 0.74214693 | NA |
| miR-3088-3p | 0.23622965 | -0.6384821 | 1.94336779 | -0.3285442 | 0.74250027 | NA |
| miR-202-3p | 0.49580888 | -0.465454 | 1.42329225 | -0.3270263 | 0.74364801 | NA |
| miR-6911-3p | 0.61536076 | 0.4414349 | 1.36855484 | 0.3225555 | 0.7470319 | NA |
| miR-212-5p | 2.24620062 | -0.2547107 | 0.7936154 | -0.3209498 | 0.7482484 | NA |
| miR-29b-2-5p | 0.95725793 | 0.33382956 | 1.04313959 | 0.32002387 | 0.74895024 | NA |
| miR-1941-3p | 0.26382346 | -0.5505644 | 1.72441038 | -0.3192769 | 0.74951653 | NA |
| miR-96-3p | 0.34549339 | -0.4657285 | 1.47047909 | -0.3167189 | 0.75145694 | NA |
| miR-468-3p | 2.52617623 | -0.2029312 | 0.6452043 | -0.3145224 | 0.75312433 | NA |
| miR-154-3p | 0.35275424 | -0.5145018 | 1.64157941 | -0.3134188 | 0.75396252 | NA |
| miR-6922-3p | 0.21798716 | 0.84108385 | 2.69334923 | 0.31228176 | 0.7548264 | NA |
| miR-7084-3p | 0.21798716 | 0.84108385 | 2.69334923 | 0.31228176 | 0.7548264 | NA |
| miR-1983 | 4.8366375 | -0.1521171 | 0.48731485 | -0.3121537 | 0.75492373 | 0.85272761 |
| miR-3100-5p | 0.33329385 | 0.57175272 | 1.84800441 | 0.30938926 | 0.75702544 | NA |
| miR-322-5p | 1.23952975 | -0.3027908 | 0.99663099 | -0.3038143 | 0.76126936 | NA |
| miR-195a-3p | 0.64269934 | -0.4313659 | 1.42304106 | -0.3031296 | 0.76179108 | NA |
| miR-7022-3p | 0.19146673 | 0.83602992 | 2.77193732 | 0.30160491 | 0.76295326 | NA |
| miR-1668 | 0.19108379 | 0.83254345 | 2.77392777 | 0.30013162 | 0.76407676 | NA |
| miR-8097 | 0.28421179 | -0.642586 | 2.14384424 | -0.2997354 | 0.76437901 | NA |
| miR-3105-3p | 0.19222292 | -0.601237 | 2.02923704 | -0.2962872 | 0.76701073 | NA |
| miR-7222-3p | 0.31604627 | -0.4810796 | 1.63772496 | -0.2937487 | 0.76894995 | NA |
| miR-301a-5p | 1.63270079 | -0.2430809 | 0.82976538 | -0.2929514 | 0.76955933 | NA |
| miR-350-3p | 7.56468761 | -0.1523857 | 0.52365744 | -0.2910026 | 0.77104932 | 0.86424084 |
| miR-186-5p | 57.9085187 | -0.0754986 | 0.25974912 | -0.2906595 | 0.77131172 | 0.86424084 |
| miR-6926-3p | 0.16418783 | 0.84200172 | 2.90213394 | 0.29013193 | 0.77171531 | NA |
| miR-7219-3p | 0.16418783 | 0.84200172 | 2.90213394 | 0.29013193 | 0.77171531 | NA |

| | | | | | | |
|---|---|---|---|---|---|---|
| miR-20a-3p | 0.50454842 | 0.4236893 | 1.46406019 | 0.28939336 | 0.77228038 | NA |
| miR-6928-3p | 0.21662044 | 0.58094418 | 2.03132695 | 0.28599245 | 0.7748839 | NA |
| miR-32-5p | 5.88170733 | 0.19295804 | 0.67587691 | 0.28549287 | 0.77526657 | 0.86517944 |
| miR-133a-5p | 0.23025598 | -0.6445981 | 2.26560146 | -0.2845152 | 0.77601559 | NA |
| let-7f-1-3p | 0.27403717 | 0.63134013 | 2.22104726 | 0.28425335 | 0.77621623 | NA |
| miR-547-3p | 0.35084855 | -0.4691938 | 1.65383792 | -0.2837 | 0.77664029 | NA |
| miR-6955-5p | 0.30632258 | 0.84227377 | 2.97317366 | 0.28329114 | 0.77695367 | NA |
| miR-7068-5p | 0.30632258 | 0.84227377 | 2.97317366 | 0.28329114 | 0.77695367 | NA |
| miR-505-3p | 0.68979419 | 0.30234759 | 1.07004923 | 0.28255484 | 0.7775181 | NA |
| miR-615-5p | 0.40172437 | -0.46463 | 1.65436583 | -0.2808508 | 0.77882481 | NA |
| miR-6942-3p | 0.14433528 | 0.84240931 | 3.01043366 | 0.27982989 | 0.77960802 | NA |
| miR-671-3p | 5.57620948 | 0.14403409 | 0.51802903 | 0.27804251 | 0.78097973 | 0.86517944 |
| miR-222-5p | 2.04190733 | -0.2008813 | 0.7240654 | -0.2774353 | 0.78144587 | NA |
| miR-423-3p | 46.3170562 | -0.0751396 | 0.2708448 | -0.2774268 | 0.7814524 | 0.86517944 |
| miR-1896 | 0.14898646 | 0.84257754 | 3.05855442 | 0.27548228 | 0.78294574 | NA |
| miR-5124a | 0.16935683 | 0.84257754 | 3.05855442 | 0.27548228 | 0.78294574 | NA |
| miR-7667-3p | 0.15901882 | 0.84257754 | 3.05855442 | 0.27548228 | 0.78294574 | NA |
| miR-6371 | 0.16935683 | 0.84257754 | 3.05855442 | 0.27548228 | 0.78294574 | NA |
| miR-330-3p | 0.41822153 | -0.3993357 | 1.4566068 | -0.2741548 | 0.78396567 | NA |
| miR-6992-5p | 0.10313131 | 0.83670734 | 3.05867796 | 0.27355196 | 0.78442896 | NA |
| miR-181b-2-3p | 0.11279867 | 0.83670734 | 3.05867796 | 0.27355196 | 0.78442896 | NA |
| miR-5625-3p | 0.11781485 | 0.83670734 | 3.05867796 | 0.27355196 | 0.78442896 | NA |
| miR-7650-5p | 0.11781485 | 0.83670734 | 3.05867796 | 0.27355196 | 0.78442896 | NA |
| miR-150-5p | 212.98997 | -0.1198591 | 0.43848088 | -0.2733508 | 0.78458355 | 0.86521269 |
| miR-6405 | 0.07661088 | 0.83317737 | 3.05875245 | 0.27239124 | 0.78532121 | NA |
| miR-1982-3p | 0.11743191 | 0.8331104 | 3.05875386 | 0.27236922 | 0.78533814 | NA |
| miR-6934-3p | 0.11743191 | 0.8331104 | 3.05875386 | 0.27236922 | 0.78533814 | NA |
| miR-7008-3p | 0.11743191 | 0.8331104 | 3.05875386 | 0.27236922 | 0.78533814 | NA |
| miR-7227-3p | 0.12260091 | 0.8331104 | 3.05875386 | 0.27236922 | 0.78533814 | NA |
| miR-328-5p | 0.10274837 | 0.8331104 | 3.05875386 | 0.27236922 | 0.78533814 | NA |
| miR-532-3p | 1.2090605 | -0.2385676 | 0.87941234 | -0.2712807 | 0.78617518 | NA |
| miR-7234-3p | 6.03780516 | -0.1320578 | 0.49546175 | -0.2665348 | 0.7898274 | 0.86756631 |
| miR-883a-5p | 1.68263002 | -0.2207348 | 0.83665819 | -0.2638292 | 0.79191156 | NA |
| miR-3618-5p | 0.20032479 | 0.73789615 | 2.79825993 | 0.26369821 | 0.79201247 | NA |
| miR-122-5p | 0.40857855 | -0.3810942 | 1.44707159 | -0.2633554 | 0.79227663 | NA |
| miR-3968 | 0.11345036 | 0.79800541 | 3.05929938 | 0.26084581 | 0.79421142 | NA |
| miR-673-5p | 0.07224639 | 0.79800541 | 3.05929938 | 0.26084581 | 0.79421142 | NA |
| miR-679-5p | 0.09876682 | 0.79800541 | 3.05929938 | 0.26084581 | 0.79421142 | NA |
| miR-496a-3p | 0.11345036 | 0.79800541 | 3.05929938 | 0.26084581 | 0.79421142 | NA |
| miR-6992-3p | 0.07224639 | 0.79800541 | 3.05929938 | 0.26084581 | 0.79421142 | NA |
| miR-7013-3p | 0.11345036 | 0.79800541 | 3.05929938 | 0.26084581 | 0.79421142 | NA |
| let-7a-2-3p | 0.09876682 | 0.79800541 | 3.05929938 | 0.26084581 | 0.79421142 | NA |
| miR-7646-3p | 0.09876682 | 0.79800541 | 3.05929938 | 0.26084581 | 0.79421142 | NA |
| let-7g-3p | 0.45558924 | 0.34554202 | 1.33113249 | 0.25958499 | 0.79518391 | NA |
| miR-29a-5p | 0.66962849 | 0.29034385 | 1.14392522 | 0.25381366 | 0.79963952 | NA |
| miR-344-3p | 0.19626048 | 0.45600841 | 1.80421387 | 0.25274632 | 0.80046425 | NA |

| | | | | | | |
|---|---|---|---|---|---|---|
| miR-1933-3p | 0.0678819 | 0.77282568 | 3.05947073 | 0.2526011 | 0.80057648 | NA |
| miR-378a-5p | 0.93662735 | 0.27198859 | 1.09650598 | 0.24805026 | 0.80409552 | NA |
| miR-7092-3p | 0.17563128 | -0.709092 | 2.86064186 | -0.2478786 | 0.80422831 | NA |
| miR-1191a | 0.15160699 | -0.6257275 | 2.53278846 | -0.2470508 | 0.80486888 | NA |
| miR-99a-3p | 0.75735927 | 0.30400744 | 1.23638427 | 0.24588427 | 0.80577182 | NA |
| miR-3079-5p | 0.34405327 | -0.5228682 | 2.14443749 | -0.2438253 | 0.80736609 | NA |
| miR-411-3p | 0.66218672 | -0.2732088 | 1.12875968 | -0.2420434 | 0.80874654 | NA |
| miR-465d-3p | 0.15637307 | -0.7099069 | 2.95173861 | -0.2405047 | 0.80993904 | NA |
| miR-363-3p | 0.37758597 | -0.4449041 | 1.85385775 | -0.2399883 | 0.81033935 | NA |
| miR-30c-1-3p | 0.55657085 | 0.2893195 | 1.22045323 | 0.23705907 | 0.81261097 | NA |
| miR-503-5p | 1.16862761 | 0.20507302 | 0.87071274 | 0.23552316 | 0.81380271 | NA |
| miR-8114 | 2.3325127 | 0.21786207 | 0.9305017 | 0.23413399 | 0.81488096 | NA |
| miR-6902-5p | 0.13551884 | -0.7107499 | 3.05572269 | -0.2325963 | 0.81607488 | NA |
| miR-6983-5p | 0.1772273 | -0.7107499 | 3.05572269 | -0.2325963 | 0.81607488 | NA |
| miR-467e-3p | 0.12841937 | -0.7107499 | 3.05572269 | -0.2325963 | 0.81607488 | NA |
| miR-1955-3p | 0.13551884 | -0.7107499 | 3.05572269 | -0.2325963 | 0.81607488 | NA |
| miR-7049-5p | 0.12841937 | -0.7107499 | 3.05572269 | -0.2325963 | 0.81607488 | NA |
| miR-7082-3p | 0.13551884 | -0.7107499 | 3.05572269 | -0.2325963 | 0.81607488 | NA |
| miR-6967-5p | 0.07961144 | -0.7107499 | 3.05572269 | -0.2325963 | 0.81607488 | NA |
| miR-5107-3p | 0.07961144 | -0.7107499 | 3.05572269 | -0.2325963 | 0.81607488 | NA |
| miR-483-5p | 0.19076291 | -0.7107499 | 3.05572269 | -0.2325963 | 0.81607488 | NA |
| miR-449c-3p | 0.17403527 | -0.7107499 | 3.05572269 | -0.2325963 | 0.81607488 | NA |
| miR-7019-5p | 0.17403527 | -0.7107499 | 3.05572269 | -0.2325963 | 0.81607488 | NA |
| miR-6920-3p | 0.17081693 | -0.7107499 | 3.05572269 | -0.2325963 | 0.81607488 | NA |
| miR-6959-3p | 0.17081693 | -0.7107499 | 3.05572269 | -0.2325963 | 0.81607488 | NA |
| miR-3082-5p | 0.79541141 | -0.305991 | 1.31653001 | -0.2324224 | 0.81620998 | NA |
| miR-183-3p | 2.87167768 | -0.1370793 | 0.59116578 | -0.2318796 | 0.81663154 | NA |
| miR-1247-3p | 0.37508678 | 0.34435669 | 1.4858814 | 0.23175247 | 0.81673027 | NA |
| miR-6907-3p | 0.18884048 | 0.65906936 | 2.84859555 | 0.23136642 | 0.81703015 | NA |
| miR-7234-5p | 1.30535638 | -0.181517 | 0.81243587 | -0.2234232 | 0.82320616 | NA |
| miR-7116-3p | 0.07387878 | -0.6774701 | 3.05612552 | -0.2216761 | 0.82456601 | NA |
| miR-3070-5p | 0.12109069 | -0.6774701 | 3.05612552 | -0.2216761 | 0.82456601 | NA |
| miR-2139 | 0.12109069 | -0.6774701 | 3.05612552 | -0.2216761 | 0.82456601 | NA |
| miR-214-5p | 0.35273426 | 0.33404918 | 1.51644601 | 0.22028426 | 0.82564978 | NA |
| miR-5626-5p | 0.36490806 | 0.38588551 | 1.7840475 | 0.21629778 | 0.82875564 | NA |
| miR-207 | 0.30602662 | -0.4719374 | 2.18882164 | -0.2156125 | 0.82928979 | NA |
| miR-6238 | 0.27848436 | 0.45887773 | 2.13011218 | 0.21542421 | 0.8294366 | NA |
| miR-6897-5p | 0.06814611 | -0.6558924 | 3.05639099 | -0.214597 | 0.83008152 | NA |
| miR-3084-5p | 0.6786882 | 0.24650508 | 1.16602283 | 0.21140674 | 0.8325699 | NA |
| miR-125b-5p | 788.994538 | 0.03486954 | 0.16524381 | 0.21101872 | 0.83287266 | 0.91089782 |
| miR-3062-5p | 0.59821961 | -0.2837689 | 1.35525901 | -0.2093835 | 0.83414888 | NA |
| miR-5132-5p | 0.33494162 | 0.38423266 | 1.83852267 | 0.2089899 | 0.83445613 | NA |
| miR-7688-5p | 0.48681148 | 0.33018534 | 1.5837885 | 0.20847818 | 0.83485562 | NA |
| miR-7007-3p | 0.38498631 | -0.3054014 | 1.47767647 | -0.2066768 | 0.83626228 | NA |
| miR-7015-3p | 0.17753236 | -0.4944555 | 2.39846101 | -0.2061553 | 0.8366696 | NA |
| miR-1843a-5p | 5.87213379 | 0.07957568 | 0.38680376 | 0.20572624 | 0.83700476 | 0.91089782 |

| | | | | | | |
|---|---|---|---|---|---|---|
| miR-466c-5p | 0.37988972 | 0.37062838 | 1.80492237 | 0.20534311 | 0.83730406 | NA |
| miR-487b-3p | 0.98970882 | -0.2132747 | 1.04734391 | -0.2036339 | 0.83863962 | NA |
| miR-7661-3p | 0.56459701 | 0.36268878 | 1.78308173 | 0.20340558 | 0.83881804 | NA |
| miR-183-5p | 676.361868 | 0.0531207 | 0.26157236 | 0.20308224 | 0.83907075 | 0.91089782 |
| miR-485-3p | 0.78085482 | -0.2405302 | 1.1876216 | -0.202531 | 0.83950163 | NA |
| miR-7001-3p | 0.21281336 | 0.4559067 | 2.274563 | 0.20043705 | 0.84113879 | NA |
| miR-8098 | 0.19014895 | 0.45921361 | 2.34663746 | 0.19569005 | 0.84485277 | NA |
| miR-539-5p | 0.34213576 | 0.35857276 | 1.8356301 | 0.19534043 | 0.84512645 | NA |
| miR-1903 | 0.18853978 | 0.45922103 | 2.35219786 | 0.19523061 | 0.84521242 | NA |
| miR-7225-3p | 0.09325753 | -0.595086 | 3.05715754 | -0.1946534 | 0.84566432 | NA |
| miR-363-5p | 0.11090657 | -0.595086 | 3.05715754 | -0.1946534 | 0.84566432 | NA |
| miR-6935-3p | 0.11251574 | -0.595086 | 3.05715754 | -0.1946534 | 0.84566432 | NA |
| miR-3091-3p | 0.11251574 | -0.595086 | 3.05715754 | -0.1946534 | 0.84566432 | NA |
| miR-1933-5p | 0.374082 | 0.4638873 | 2.38840561 | 0.19422467 | 0.84599996 | NA |
| miR-3108-5p | 0.17840834 | 0.4638965 | 2.39090086 | 0.19402582 | 0.84615566 | NA |
| miR-3474 | 0.14848197 | 0.48608115 | 2.51712207 | 0.19310988 | 0.84687291 | NA |
| miR-450b-3p | 0.24302338 | -0.3772897 | 1.96208004 | -0.1922907 | 0.84751454 | NA |
| miR-381-3p | 0.39691222 | -0.2839409 | 1.47685818 | -0.1922601 | 0.84753849 | NA |
| miR-376c-3p | 0.97357242 | 0.21013616 | 1.10030802 | 0.19097939 | 0.84854174 | NA |
| miR-199b-5p | 1.7077327 | -0.1342502 | 0.71551349 | -0.1876278 | 0.85116843 | NA |
| miR-6948-3p | 0.15222145 | 0.45604379 | 2.49917954 | 0.1824774 | 0.85520809 | NA |
| miR-3094-3p | 0.14398736 | 0.45361693 | 2.53927932 | 0.17864003 | 0.85822036 | NA |
| miR-212-3p | 0.1595625 | -0.4436013 | 2.496774 | -0.1776698 | 0.8589823 | NA |
| miR-7217-3p | 4.53062318 | -0.1065478 | 0.60868603 | -0.1750456 | 0.86104379 | 0.93112875 |
| miR-335-3p | 1.47739659 | -0.149939 | 0.85821599 | -0.1747101 | 0.86130742 | NA |
| miR-28c | 0.35774747 | -0.2578908 | 1.47986089 | -0.1742669 | 0.8616557 | NA |
| miR-665-3p | 0.16086004 | 0.42318349 | 2.46087983 | 0.17196431 | 0.86346559 | NA |
| miR-3108-3p | 0.15765485 | 0.42311799 | 2.47498296 | 0.17095794 | 0.86425684 | NA |
| miR-200b-3p | 1617.11035 | -0.0404356 | 0.23692734 | -0.1706665 | 0.86448598 | 0.93124165 |
| miR-6932-5p | 0.05099622 | -0.5194769 | 3.05786529 | -0.1698822 | 0.86510278 | NA |
| miR-3081-3p | 0.05099622 | -0.5194769 | 3.05786529 | -0.1698822 | 0.86510278 | NA |
| miR-3102-5p | 0.05099622 | -0.5194769 | 3.05786529 | -0.1698822 | 0.86510278 | NA |
| miR-3086-5p | 0.12426493 | 0.44954603 | 2.64866706 | 0.16972538 | 0.86522611 | NA |
| miR-6975-3p | 0.33241291 | -0.3099978 | 1.84076069 | -0.1684075 | 0.86626274 | NA |
| miR-7079-3p | 0.24090074 | 0.37739312 | 2.26170958 | 0.16686188 | 0.86747872 | NA |
| miR-7218-3p | 0.90273348 | 0.18322128 | 1.09974243 | 0.16660381 | 0.86768179 | NA |
| miR-1906 | 0.15514986 | 0.4936506 | 2.97404918 | 0.16598603 | 0.86816795 | NA |
| miR-7118-3p | 0.15514986 | 0.4936506 | 2.97404918 | 0.16598603 | 0.86816795 | NA |
| miR-205-3p | 0.3840895 | 0.24262475 | 1.47707796 | 0.16425995 | 0.86952651 | NA |
| miR-19b-3p | 25.7823201 | 0.06094581 | 0.37114072 | 0.16421212 | 0.86956416 | 0.93310923 |
| miR-376a-3p | 0.32634532 | 0.27893194 | 1.72664666 | 0.16154547 | 0.8716638 | NA |
| miR-92a-1-5p | 1.81348382 | 0.11556322 | 0.72117896 | 0.16024209 | 0.87269038 | NA |
| miR-200c-3p | 1904.86827 | 0.03409422 | 0.21371307 | 0.15953269 | 0.87324921 | 0.93347329 |
| miR-324-3p | 0.59605109 | -0.2847696 | 1.79333341 | -0.1587935 | 0.87383159 | NA |
| miR-719 | 0.25193551 | -0.3392105 | 2.2015185 | -0.1540802 | 0.87754648 | NA |
| miR-3058-3p | 0.16125346 | 0.45061434 | 2.93930487 | 0.15330643 | 0.87815663 | NA |

| | | | | | | |
|---|---|---|---|---|---|---|
| miR-681 | 0.56903102 | -0.2435373 | 1.59499654 | -0.1526883 | 0.87864409 | NA |
| miR-29a-3p | 830.322418 | -0.0267765 | 0.17640819 | -0.151787 | 0.87935495 | 0.93619561 |
| miR-505-5p | 0.11068394 | 0.45632603 | 3.05772643 | 0.14923704 | 0.8813666 | NA |
| miR-467d-3p | 1.20115837 | -0.1217922 | 0.81943232 | -0.14863 | 0.88184559 | NA |
| miR-28a-3p | 7.83772416 | -0.06289 | 0.42553083 | -0.1477919 | 0.88250697 | 0.93619561 |
| miR-350-5p | 0.3909877 | -0.2382074 | 1.62680545 | -0.1464265 | 0.88358472 | NA |
| miR-935 | 0.28414291 | -0.3199946 | 2.18536104 | -0.1464264 | 0.88358478 | NA |
| miR-6946-5p | 0.19004006 | -0.3420099 | 2.35235065 | -0.1453907 | 0.88440239 | NA |
| miR-7a-1-3p | 1.68471726 | -0.1047659 | 0.72356734 | -0.1447909 | 0.884876 | NA |
| miR-7059-3p | 0.23301069 | 0.28530481 | 1.97741092 | 0.144282 | 0.88527779 | NA |
| miR-181a-2-3p | 0.84459118 | -0.1633539 | 1.1371606 | -0.1436507 | 0.88577633 | NA |
| let-7i-3p | 1.46375383 | -0.12094 | 0.84443199 | -0.1432205 | 0.88611605 | NA |
| miR-574-3p | 5.95981554 | 0.05810497 | 0.40705392 | 0.14274515 | 0.88649146 | 0.93686029 |
| miR-470-3p | 0.29168496 | 0.21808065 | 1.53225079 | 0.14232699 | 0.88682173 | NA |
| miR-669b-5p | 0.28163467 | 0.21652163 | 1.53922649 | 0.14066912 | 0.88813134 | NA |
| miR-669m-5p | 0.28163467 | 0.21652163 | 1.53922649 | 0.14066912 | 0.88813134 | NA |
| miR-138-5p | 0.40349071 | 0.2020453 | 1.44806208 | 0.13952806 | 0.88903288 | NA |
| miR-7226-3p | 0.09338001 | 0.42426481 | 3.05863865 | 0.13871034 | 0.88967905 | NA |
| miR-3960 | 0.25871368 | -0.3453646 | 2.58133206 | -0.1337932 | 0.89356613 | NA |
| miR-412-5p | 0.18430739 | -0.3131297 | 2.37110134 | -0.1320609 | 0.89493615 | NA |
| miR-331-5p | 0.53500837 | -0.1711741 | 1.29704413 | -0.1319725 | 0.89500607 | NA |
| miR-3087-3p | 0.22906835 | -0.3466061 | 2.68461645 | -0.1291082 | 0.89727201 | NA |
| miR-7232-3p | 1.97893264 | 0.09439163 | 0.73347898 | 0.1286903 | 0.89760271 | NA |
| miR-139-3p | 1.35815022 | -0.1296422 | 1.01203878 | -0.1281001 | 0.89806978 | NA |
| let-7c-2-3p | 1.496716 | 0.12020086 | 0.94164543 | 0.12764981 | 0.89842611 | NA |
| miR-7042-3p | 0.21553274 | -0.3470368 | 2.72343754 | -0.127426 | 0.89860324 | NA |
| miR-1966-3p | 0.21116824 | -0.3472135 | 2.73986252 | -0.1267266 | 0.89915677 | NA |
| miR-295-3p | 0.73410511 | -0.140655 | 1.11717281 | -0.1259026 | 0.89980899 | NA |
| miR-7221-3p | 0.56201332 | 0.17086577 | 1.42697941 | 0.11973948 | 0.90468953 | NA |
| miR-329-3p | 0.41466936 | 0.17350006 | 1.45468204 | 0.11927009 | 0.90506138 | NA |
| miR-370-3p | 0.31582667 | 0.19138016 | 1.65370847 | 0.11572787 | 0.90786822 | NA |
| miR-493-3p | 0.11218422 | -0.3153937 | 2.72569425 | -0.1157113 | 0.90788132 | NA |
| miR-511-3p | 0.18666321 | 0.24465273 | 2.1323307 | 0.11473489 | 0.90865526 | NA |
| miR-669d-5p | 1.93131896 | -0.0767407 | 0.69759723 | -0.1100071 | 0.91240373 | NA |
| miR-153-3p | 0.24641882 | -0.2365938 | 2.18831069 | -0.1081171 | 0.9139028 | NA |
| miR-669c-5p | 62.7764284 | 0.03726253 | 0.35116805 | 0.10611026 | 0.91549487 | 0.95611268 |
| miR-742-5p | 4.53827749 | -0.0529939 | 0.50615029 | -0.1047 | 0.91661386 | 0.95611268 |
| miR-877-5p | 0.57376416 | 0.18801768 | 1.79671876 | 0.10464503 | 0.91665748 | NA |
| miR-883b-3p | 3.02544816 | -0.0709799 | 0.69215158 | -0.1025496 | 0.91832046 | 0.95611268 |
| miR-297b-5p | 0.25018608 | 0.17304105 | 1.73367749 | 0.09981155 | 0.92049393 | NA |
| miR-151-3p | 27.2214617 | 0.0318272 | 0.32852788 | 0.09687823 | 0.9228231 | 0.95611268 |
| miR-1968-5p | 0.13297198 | -0.2952217 | 3.05639097 | -0.0965916 | 0.92305072 | NA |
| miR-540-3p | 0.15082118 | -0.2355312 | 2.50244846 | -0.0941203 | 0.92501358 | NA |
| miR-5131 | 0.15082118 | -0.2355312 | 2.50244846 | -0.0941203 | 0.92501358 | NA |
| miR-669p-3p | 0.15082118 | -0.2355312 | 2.50244846 | -0.0941203 | 0.92501358 | NA |
| miR-532-5p | 13.196146 | -0.0269246 | 0.28690484 | -0.0938451 | 0.9252322 | 0.95611268 |

| | | | | | | |
|---|---|---|---|---|---|---|
| miR-7214-5p | 8.08250289 | 0.04550744 | 0.49267647 | 0.0923678 | 0.92640583 | 0.95611268 |
| miR-7229-5p | 2.66938679 | 0.06172964 | 0.67900339 | 0.09091213 | 0.92756241 | NA |
| miR-223-3p | 31.1709963 | 0.02642737 | 0.29533065 | 0.08948401 | 0.92869726 | 0.95611268 |
| miR-326-3p | 0.44709418 | -0.1086557 | 1.32549572 | -0.0819736 | 0.9346677 | NA |
| miR-148b-3p | 48.6693625 | -0.0233746 | 0.2882945 | -0.0810791 | 0.93537908 | 0.95945134 |
| miR-7658-3p | 0.09749366 | -0.2278047 | 2.84241014 | -0.0801449 | 0.93612201 | NA |
| miR-7242-5p | 1.37128616 | -0.0800821 | 1.06622777 | -0.0751079 | 0.94012887 | NA |
| miR-1927 | 0.31361076 | 0.11626827 | 1.65273369 | 0.07034906 | 0.94391583 | NA |
| miR-425-3p | 3.25371426 | -0.0415402 | 0.59100518 | -0.0702874 | 0.94396491 | 0.96471139 |
| miR-7a-2-3p | 0.33317398 | -0.1019637 | 1.63226822 | -0.0624675 | 0.95019055 | NA |
| miR-7068-3p | 0.9316479 | -0.0644684 | 1.03814127 | -0.0620998 | 0.95048333 | NA |
| miR-206-3p | 0.54637891 | 0.08235788 | 1.34957253 | 0.06102516 | 0.95133917 | NA |
| miR-193b-3p | 0.52934003 | -0.0904176 | 1.52990282 | -0.0591002 | 0.95287227 | NA |
| miR-130a-3p | 5.3941586 | 0.0251097 | 0.42833399 | 0.05862178 | 0.95325336 | 0.96895092 |
| miR-294-3p | 1.53219423 | 0.04709805 | 0.82473784 | 0.05710669 | 0.95446021 | NA |
| miR-878-3p | 24.8463178 | 0.02580204 | 0.45784981 | 0.05635482 | 0.95505915 | 0.96895092 |
| miR-5101 | 0.16800534 | 0.11640902 | 2.1093964 | 0.05518594 | 0.95599033 | NA |
| miR-3060-3p | 0.21128256 | 0.10963355 | 1.98911572 | 0.05511673 | 0.95604547 | NA |
| miR-7240-5p | 0.24480458 | -0.1062763 | 1.95507304 | -0.0543592 | 0.95664895 | NA |
| miR-6546-5p | 0.08930166 | -0.1588059 | 3.05786528 | -0.0519336 | 0.95858161 | NA |
| miR-365-2-5p | 0.69953349 | -0.054175 | 1.06196591 | -0.0510139 | 0.95931443 | NA |
| miR-18a-3p | 0.74276771 | 0.05376996 | 1.08094609 | 0.04974342 | 0.96032685 | NA |
| miR-30c-2-3p | 22.067386 | 0.01460597 | 0.29894552 | 0.0488583 | 0.96103222 | 0.97010151 |
| miR-1839-5p | 66.1152609 | -0.0108557 | 0.23495039 | -0.0462044 | 0.96314737 | 0.97010151 |
| miR-3069-3p | 0.35323984 | 0.06184501 | 1.62076944 | 0.03815781 | 0.96956186 | NA |
| miR-7223-5p | 0.71284685 | 0.04810322 | 1.26112785 | 0.03814302 | 0.96957365 | NA |
| miR-668-3p | 0.6286783 | 0.04865118 | 1.28190877 | 0.03795214 | 0.96972584 | NA |
| miR-3620-3p | 0.1786594 | 0.10414143 | 2.82109085 | 0.0369153 | 0.97055254 | NA |
| miR-370-5p | 0.11017652 | 0.10435535 | 3.0595507 | 0.03410806 | 0.97279098 | NA |
| miR-666-3p | 0.06342061 | 0.10435535 | 3.0595507 | 0.03410806 | 0.97279098 | NA |
| miR-6357 | 0.06342061 | 0.10435535 | 3.0595507 | 0.03410806 | 0.97279098 | NA |
| miR-7075-3p | 0.06342061 | 0.10435535 | 3.0595507 | 0.03410806 | 0.97279098 | NA |
| miR-1929-5p | 0.10500752 | 0.10435535 | 3.0595507 | 0.03410806 | 0.97279098 | NA |
| miR-6989-3p | 0.10500752 | 0.10435535 | 3.0595507 | 0.03410806 | 0.97279098 | NA |
| miR-7215-5p | 0.06380355 | 0.10435535 | 3.0595507 | 0.03410806 | 0.97279098 | NA |
| miR-541-3p | 0.06380355 | 0.10435535 | 3.0595507 | 0.03410806 | 0.97279098 | NA |
| miR-6964-3p | 0.06380355 | 0.10435535 | 3.0595507 | 0.03410806 | 0.97279098 | NA |
| miR-7026-5p | 0.05943906 | 0.10435535 | 3.0595507 | 0.03410806 | 0.97279098 | NA |
| miR-7647-3p | 0.06380355 | 0.10435535 | 3.0595507 | 0.03410806 | 0.97279098 | NA |
| miR-224-3p | 0.06380355 | 0.10435535 | 3.0595507 | 0.03410806 | 0.97279098 | NA |
| let-7f-2-3p | 0.53243114 | -0.0439155 | 1.30333561 | -0.0336947 | 0.97312064 | NA |
| miR-7006-3p | 0.24017892 | 0.07696759 | 2.57720071 | 0.0298648 | 0.97617488 | NA |
| miR-3103-3p | 0.24017892 | 0.07696759 | 2.57720071 | 0.0298648 | 0.97617488 | NA |
| miR-434-5p | 1.28118429 | 0.02453453 | 0.8389686 | 0.02924368 | 0.97667025 | NA |
| miR-141-3p | 50.4204726 | -0.0094323 | 0.33266798 | -0.0283534 | 0.97738026 | 0.98089602 |
| miR-3620-5p | 0.18723435 | 0.07652998 | 2.7806298 | 0.02752254 | 0.97804297 | NA |

| | | | | | | |
|---|---|---|---|---|---|---|
| miR-761 | 0.17008688 | 0.07638533 | 2.85909434 | 0.02671662 | 0.97868576 | NA |
| miR-1898 | 0.15023434 | 0.07619736 | 2.97158137 | 0.02564202 | 0.97954287 | NA |
| miR-669e-5p | 0.15023434 | 0.07619736 | 2.97158137 | 0.02564202 | 0.97954287 | NA |
| miR-291a-5p | 0.13330395 | 0.07606468 | 3.05935708 | 0.02486296 | 0.98016427 | NA |
| miR-7046-3p | 0.13330395 | 0.07606468 | 3.05935708 | 0.02486296 | 0.98016427 | NA |
| miR-5129-3p | 0.12653615 | 0.07606468 | 3.05935708 | 0.02486296 | 0.98016427 | NA |
| miR-7019-3p | 0.07772822 | 0.07606468 | 3.05935708 | 0.02486296 | 0.98016427 | NA |
| miR-7059-5p | 0.10568192 | 0.07606468 | 3.05935708 | 0.02486296 | 0.98016427 | NA |
| miR-7060-3p | 0.12333096 | 0.07606468 | 3.05935708 | 0.02486296 | 0.98016427 | NA |
| miR-7650-3p | 0.12095858 | 0.07606468 | 3.05935708 | 0.02486296 | 0.98016427 | NA |
| miR-1249-5p | 0.11931513 | 0.07606468 | 3.05935708 | 0.02486296 | 0.98016427 | NA |
| miR-3083-5p | 0.11931513 | 0.07606468 | 3.05935708 | 0.02486296 | 0.98016427 | NA |
| miR-296-3p | 0.10856628 | 0.07606468 | 3.05935708 | 0.02486296 | 0.98016427 | NA |
| miR-7070-5p | 0.10463159 | 0.07606468 | 3.05935708 | 0.02486296 | 0.98016427 | NA |
| miR-6899-3p | 0.11934941 | 0.07606468 | 3.05935708 | 0.02486296 | 0.98016427 | NA |
| miR-1982-5p | 0.07811116 | 0.07606468 | 3.05935708 | 0.02486296 | 0.98016427 | NA |
| miR-103-1-5p | 0.07374667 | 0.07606468 | 3.05935708 | 0.02486296 | 0.98016427 | NA |
| miR-7652-3p | 0.068014 | 0.07606468 | 3.05935708 | 0.02486296 | 0.98016427 | NA |
| miR-1936 | 0.068014 | 0.07606468 | 3.05935708 | 0.02486296 | 0.98016427 | NA |
| miR-666-5p | 0.0723785 | 0.07606468 | 3.05935708 | 0.02486296 | 0.98016427 | NA |
| miR-7118-5p | 0.07374667 | 0.07606468 | 3.05935708 | 0.02486296 | 0.98016427 | NA |
| miR-877-3p | 0.07811116 | 0.07606468 | 3.05935708 | 0.02486296 | 0.98016427 | NA |
| miR-6988-3p | 0.1237139 | 0.07606468 | 3.05935708 | 0.02486296 | 0.98016427 | NA |
| miR-3970 | 0.11934941 | 0.07606468 | 3.05935708 | 0.02486296 | 0.98016427 | NA |
| miR-708-5p | 0.0723785 | 0.07606468 | 3.05935708 | 0.02486296 | 0.98016427 | NA |
| miR-7060-5p | 0.1237139 | 0.07606468 | 3.05935708 | 0.02486296 | 0.98016427 | NA |
| miR-5130 | 0.22995467 | 0.04606805 | 1.95570607 | 0.02355571 | 0.981207 | NA |
| miR-6946-3p | 0.45010952 | 0.03272254 | 1.43939018 | 0.02273362 | 0.98186276 | NA |
| miR-1964-3p | 0.34026719 | 0.04734994 | 2.08352202 | 0.02272591 | 0.98186891 | NA |
| miR-193a-3p | 0.22821788 | 0.03895803 | 2.00940622 | 0.01938783 | 0.98453172 | NA |
| miR-8118 | 0.27716658 | 0.04196828 | 2.18163517 | 0.01923707 | 0.98465198 | NA |
| miR-147-3p | 0.21550734 | -0.0379911 | 2.37945468 | -0.0159663 | 0.98726126 | NA |
| miR-1934-5p | 0.36894442 | -0.022515 | 1.48639096 | -0.0151474 | 0.98791457 | NA |
| miR-500-3p | 0.79307814 | -0.0105511 | 1.19132054 | -0.0088566 | 0.99293352 | NA |
| miR-1969 | 0.19909524 | -0.0087037 | 2.40325867 | -0.0036216 | 0.99711038 | NA |
| miR-452-5p | 0.26262635 | 0.00674643 | 2.22413132 | 0.00303329 | 0.99757979 | NA |
| miR-130a-5p | 0.24909074 | 0.00625513 | 2.2463625 | 0.00278456 | 0.99777824 | NA |
| miR-542-3p | 0.46789068 | -0.0022112 | 1.62505096 | -0.0013607 | 0.99891433 | NA |
| miR-15b-5p | 345.23502 | 0.00012725 | 0.32764006 | 0.00038837 | 0.99969012 | 0.99969012 |

| | FGCZ6510_Free_tRNA_dedup_counts | | FGCZ6510_Free_tRNA_dedup_counts | | |
|---|---|---|---|---|---|
| | CON.11 | CON.12 | CON.13 | CON.14 | CON.15 |
| tRNA-Ala-TGC-2-1 | 168.592357 | 143.585913 | 231.8949 | 312.931948 | 224.90119 |
| tRNA-Arg-ACG-3-1 | 120.893056 | 84.1710522 | 176.084374 | 122.090874 | 140.212712 |
| tRNA-Glu-CTC-3-1 | 1701.5492 | 755.558975 | 601.369236 | 719.50641 | 604.597214 |
| tRNA-Ser-TGA-1-1 | 249.187728 | 245.581423 | 237.086577 | 252.479185 | 319.124132 |
| tRNA-Thr-TGT-2-1 | 512.356285 | 549.587459 | 631.221378 | 391.757609 | 427.368346 |
| tRNA-Ser-TGA-2-1 | 521.402704 | 330.742723 | 589.687964 | 395.906328 | 624.787844 |
| tRNA-Ser-AGA-1-1 | 519.7579 | 328.762227 | 587.957405 | 394.128306 | 623.105291 |
| tRNA-Ser-TGA-2-2 | 520.580302 | 328.762227 | 588.822684 | 394.128306 | 623.105291 |
| tRNA-Glu-TTC-3-1 | 3991.11565 | 2410.26284 | 1152.11962 | 1999.68256 | 1749.29379 |
| tRNA-Glu-TTC-3-2 | 3991.11565 | 2409.27259 | 1151.68698 | 1998.49721 | 1748.73294 |
| tRNA-Arg-CCT-2-1 | 215.469256 | 259.44489 | 436.966136 | 261.961972 | 238.922461 |
| tRNA-Arg-CCT-2-2 | 216.291658 | 259.44489 | 436.966136 | 261.961972 | 238.922461 |
| tRNA-Arg-CCT-1-1 | 215.469256 | 261.425386 | 435.235577 | 261.369297 | 240.605014 |
| tRNA-Ser-AGA-2-4 | 472.058599 | 284.201082 | 510.514891 | 330.712172 | 579.919776 |
| tRNA-Ser-AGA-2-5 | 472.058599 | 284.201082 | 510.947531 | 330.712172 | 579.919776 |
| tRNA-Ser-AGA-2-3 | 472.058599 | 284.201082 | 510.947531 | 331.304846 | 580.480627 |
| tRNA-Ser-AGA-2-6 | 472.058599 | 284.201082 | 510.947531 | 330.712172 | 580.480627 |
| tRNA-Ser-AGA-2-1 | 477.815412 | 285.19133 | 516.571848 | 332.490195 | 586.089136 |
| tRNA-Ser-AGA-2-2 | 472.058599 | 284.201082 | 511.812811 | 331.304846 | 581.602329 |
| tRNA-Cys-GCA-3-4 | 982.770081 | 1092.24318 | 849.704446 | 1027.69697 | 1278.17908 |
| tRNA-Cys-GCA-3-1 | 983.592483 | 1093.23343 | 846.675968 | 1023.54825 | 1278.17908 |
| tRNA-Cys-GCA-3-2 | 990.171697 | 1093.23343 | 851.435005 | 1028.28964 | 1285.47014 |
| tRNA-Cys-GCA-3-3 | 989.349295 | 1093.23343 | 851.435005 | 1029.47499 | 1285.47014 |
| tRNA-Pro-AGG-3-1 | 7.40161567 | 7.92198139 | 10.3833537 | 8.89011215 | 5.60850847 |
| tRNA-Lys-CTT-3-2 | 13775.2292 | 11863.1671 | 9961.52998 | 9921.95784 | 9349.38363 |
| tRNA-Arg-CCT-3-1 | 221.226068 | 262.415633 | 444.321011 | 236.476983 | 242.848417 |
| tRNA-Lys-CTT-3-4 | 13495.6126 | 11732.4544 | 9718.81908 | 9726.37537 | 9173.27646 |
| tRNA-Lys-CTT-3-1 | 13494.7902 | 11731.4642 | 9715.35797 | 9726.37537 | 9171.03306 |
| tRNA-Lys-CTT-3-7 | 13461.8941 | 11717.6007 | 9701.5135 | 9702.66841 | 9141.86881 |
| tRNA-Trp-CCA-5-1 | 148.854715 | 213.893497 | 181.70869 | 248.92314 | 262.478197 |
| tRNA-Lys-CTT-3-3 | 13461.0717 | 11705.7177 | 9690.26486 | 9695.55632 | 9137.38201 |
| tRNA-Lys-CTT-3-5 | 13449.5581 | 11702.747 | 9676.85303 | 9694.37097 | 9132.33435 |
| tRNA-Lys-CTT-3-6 | 13449.5581 | 11702.747 | 9676.42039 | 9693.77829 | 9132.33435 |
| tRNA-Cys-GCA-2-1 | 944.939601 | 1049.66253 | 760.58066 | 963.095483 | 1233.31101 |
| tRNA-Cys-GCA-1-2 | 940.00519 | 1045.70154 | 754.523704 | 945.907933 | 1224.3374 |
| tRNA-Cys-GCA-1-1 | 931.781173 | 1043.72105 | 747.601468 | 943.537237 | 1218.72889 |
| tRNA-Ala-TGC-5-1 | 18.91524 | 18.8147058 | 9.51807425 | 22.5216175 | 33.0902 |
| tRNA-Ala-TGC-5-2 | 18.91524 | 18.8147058 | 9.51807425 | 22.5216175 | 33.0902 |
| tRNA-Ala-CGC-5-1 | 1837.24549 | 1319.0099 | 466.385638 | 909.754811 | 738.640566 |
| tRNA-Gly-CCC-3-1 | 1963.89536 | 1918.10974 | 615.646348 | 963.095483 | 960.737502 |
| tRNA-Gly-CCC-4-1 | 1963.89536 | 1918.10974 | 612.61787 | 962.502809 | 959.6158 |
| tRNA-Ala-TGC-5-3 | 18.91524 | 17.8244581 | 9.08543451 | 22.5216175 | 33.0902 |
| tRNA-Thr-AGT-5-1 | 3.28960697 | 5.94148604 | 11.2486332 | 6.51941558 | 3.92595593 |
| tRNA-SeC-TCA-1-1 | 3162.9571 | 3141.06562 | 3588.74663 | 3641.38994 | 3608.51435 |

| | | | | | |
|---|---|---|---|---|---|
| tRNA-Thr-AGT-7-1 | 3.28960697 | 5.94148604 | 12.1139127 | 4.74139315 | 3.36510508 |
| tRNA-Glu-CTC-2-1 | 12960.229 | 9560.84128 | 3944.80914 | 6643.28448 | 5648.88974 |
| tRNA-His-GTG-1-1 | 30.4288644 | 49.5123837 | 61.8674826 | 38.5238193 | 24.6774373 |
| tRNA-Arg-CCT-4-1 | 457.255368 | 406.001546 | 579.30461 | 403.018418 | 486.818536 |
| tRNA-Lys-CTT-1-1 | 6.57921393 | 9.90247673 | 5.19167686 | 5.92674144 | 8.41276271 |
| tRNA-Lys-CTT-2-2 | 4033.05814 | 2871.71825 | 3131.01379 | 2568.64974 | 2592.25262 |
| tRNA-Lys-CTT-2-1 | 4007.56369 | 2861.81578 | 3108.51652 | 2560.94497 | 2579.35305 |
| tRNA-Asp-GTC-4-1 | 1637.40187 | 1274.44876 | 702.606935 | 1251.13512 | 1031.96556 |
| tRNA-Gly-GCC-5-1 | 25030.6194 | 27630.8808 | 12295.1887 | 15242.3936 | 16716.7204 |
| tRNA-Thr-TGT-3-1 | 732.759951 | 648.612226 | 840.619012 | 731.359893 | 632.078905 |
| tRNA-Asn-GTT-2-1 | 63.3249341 | 88.1320429 | 105.996736 | 82.381706 | 58.3284881 |
| tRNA-Asn-GTT-4-1 | 61.6801306 | 84.1710522 | 104.266177 | 75.8622904 | 55.5242339 |
| tRNA-Thr-TGT-3-2 | 758.254405 | 651.582969 | 868.307955 | 739.657331 | 646.100176 |
| tRNA-Asn-GTT-1-1 | 129.939475 | 139.624922 | 209.830273 | 137.500401 | 118.90038 |
| tRNA-Gly-GCC-4-1 | 47664.7601 | 50932.3988 | 27617.9903 | 31634.5751 | 33107.0255 |
| tRNA-Gly-GCC-2-8 | 42138.2204 | 47639.8253 | 25582.853 | 28746.474 | 29090.2118 |
| tRNA-Gly-GCC-2-7 | 42089.6987 | 47622.0009 | 25565.5474 | 28729.8791 | 29073.3862 |
| tRNA-Gly-GCC-2-5 | 42073.2507 | 47619.0301 | 25547.3766 | 28722.767 | 29052.0739 |
| tRNA-Gly-GCC-2-4 | 42073.2507 | 47618.0399 | 25546.5113 | 28722.767 | 29052.0739 |
| tRNA-Gly-GCC-2-1 | 42103.6795 | 47676.4645 | 25566.8453 | 28769.5883 | 29079.5556 |
| tRNA-Thr-CGT-4-1 | 12.3360261 | 11.8829721 | 13.4118319 | 10.0754604 | 15.1429729 |
| tRNA-Gly-GCC-2-2 | 43814.2752 | 49567.8375 | 26458.0832 | 29849.4406 | 30299.4062 |
| tRNA-Gly-GCC-2-6 | 43821.6768 | 49581.701 | 26453.3242 | 29860.1087 | 30315.11 |
| tRNA-Gly-GCC-2-3 | 43912.141 | 49757.9651 | 26494.8576 | 29947.2318 | 30370.6342 |
| tRNA-Gly-GCC-3-1 | 44843.9221 | 51470.1033 | 26824.9617 | 30611.6195 | 31066.6501 |
| tRNA-Gly-CCC-5-1 | 36466.938 | 43391.6628 | 22831.2643 | 26215.1627 | 26846.2475 |
| tRNA-Ala-TGC-6-1 | 8.22401741 | 9.90247673 | 7.35487555 | 10.6681346 | 23.5557356 |
| tRNA-Glu-CTC-1-4 | 36901.1661 | 35053.7774 | 18064.4396 | 26248.9451 | 23093.0336 |
| tRNA-Glu-CTC-1-7 | 36884.7181 | 35024.07 | 18056.2195 | 26197.3825 | 23064.4302 |
| tRNA-Glu-CTC-1-8 | 36928.3054 | 35102.2995 | 18017.2819 | 26228.2016 | 23067.2345 |
| tRNA-Glu-CTC-1-6 | 36911.8574 | 35117.1532 | 18023.3389 | 26232.3503 | 23073.4039 |
| tRNA-Glu-CTC-1-5 | 36911.035 | 35117.1532 | 18023.3389 | 26232.3503 | 23073.9647 |
| tRNA-Glu-CTC-1-1 | 36689.8089 | 34944.8501 | 17894.4122 | 26072.3283 | 22944.969 |
| tRNA-Glu-CTC-1-2 | 36689.8089 | 34944.8501 | 17894.4122 | 26072.3283 | 22944.969 |
| tRNA-Glu-CTC-1-3 | 36689.8089 | 34944.8501 | 17894.4122 | 26072.3283 | 22944.969 |
| tRNA-Glu-CTC-1-9 | 36943.931 | 35148.8412 | 18057.0848 | 26259.0206 | 23104.8115 |
| tRNA-Ala-AGC-4-1 | 123.360261 | 156.459132 | 171.757976 | 126.239593 | 146.382071 |
| tRNA-Asn-GTT-3-4 | 153.789126 | 185.176315 | 237.951856 | 192.619097 | 156.477386 |
| tRNA-Asn-GTT-3-5 | 153.789126 | 185.176315 | 237.951856 | 192.619097 | 156.477386 |
| tRNA-Asn-GTT-3-3 | 153.789126 | 185.176315 | 237.951856 | 194.397119 | 156.477386 |
| tRNA-Asn-GTT-3-7 | 166.947553 | 193.098296 | 249.633129 | 196.175142 | 164.890149 |
| tRNA-Asn-GTT-3-9 | 157.901134 | 185.176315 | 242.710893 | 194.397119 | 157.599088 |
| tRNA-Asn-GTT-3-6 | 158.723536 | 185.176315 | 251.363688 | 195.582467 | 162.646746 |
| tRNA-Asn-GTT-3-8 | 164.480348 | 197.059287 | 256.122725 | 207.43595 | 170.498658 |
| tRNA-Asn-GTT-3-1 | 169.414759 | 192.108049 | 253.959526 | 196.767816 | 166.572702 |
| tRNA-Asn-GTT-3-2 | 185.040392 | 212.90325 | 250.065769 | 206.250602 | 171.059508 |

| | | | | | |
|---|---|---|---|---|---|
| tRNA-Thr-TGT-1-1 | 12.3360261 | 11.8829721 | 15.1423908 | 10.0754604 | 16.8255254 |
| tRNA-Leu-TAA-2-1 | 677.659035 | 732.783278 | 836.725254 | 873.009014 | 1261.35356 |
| tRNA-Asp-GTC-2-1 | 21134.9023 | 9690.56373 | 8262.12109 | 8335.36916 | 9313.48917 |
| tRNA-Val-AAC-2-1 | 300.176636 | 637.719502 | 450.810607 | 587.340076 | 464.945353 |
| tRNA-Val-AAC-2-2 | 300.176636 | 637.719502 | 450.810607 | 587.340076 | 464.945353 |
| tRNA-Arg-CCG-1-1 | 17.2704366 | 29.7074302 | 28.9868625 | 30.8190555 | 30.2859458 |
| tRNA-Pro-AGG-2-1 | 3.28960697 | 3.96099069 | 4.32639738 | 3.55604486 | 3.92595593 |
| tRNA-Leu-TAA-5-1 | 3.28960697 | 3.96099069 | 4.32639738 | 3.55604486 | 3.92595593 |
| tRNA-Leu-TAA-5-2 | 3.28960697 | 3.96099069 | 4.32639738 | 3.55604486 | 3.92595593 |
| tRNA-Leu-TAA-5-3 | 3.28960697 | 3.96099069 | 4.32639738 | 3.55604486 | 3.92595593 |
| tRNA-Cys-GCA-5-1 | 49.3441045 | 41.5904023 | 43.6966136 | 62.2307851 | 46.5506203 |
| tRNA-Leu-TAG-3-1 | 3070.8481 | 1748.77739 | 1584.75936 | 3467.14374 | 2609.07814 |
| tRNA-Gln-TTG-5-1 | 32.8960697 | 74.2685755 | 86.095308 | 91.8644923 | 38.6987085 |
| tRNA-Arg-ACG-3-2 | 50.988908 | 63.3758511 | 86.5279477 | 79.4183353 | 79.0799695 |
| tRNA-Cys-GCA-11-1 | 6.57921393 | 5.94148604 | 10.8159935 | 6.51941558 | 3.92595593 |
| tRNA-Thr-CGT-3-1 | 13.1584279 | 9.90247673 | 13.8444716 | 7.11208972 | 7.29106102 |
| tRNA-Gly-CCC-1-1 | 9576.04588 | 9430.12859 | 8128.00277 | 9381.43902 | 6565.32002 |
| tRNA-Gly-CCC-1-2 | 9576.04588 | 9430.12859 | 8128.00277 | 9381.43902 | 6565.32002 |
| tRNA-Ala-AGC-4-2 | 143.097903 | 176.264086 | 204.638596 | 133.351682 | 163.207597 |
| tRNA-Ala-CGC-7-1 | 1.64480348 | 1.98049535 | 0.86527948 | 2.37069657 | 3.36510508 |
| tRNA-Phe-GAA-2-1 | 6.57921393 | 6.93173371 | 12.5465524 | 7.70476387 | 5.60850847 |
| tRNA-Ala-AGC-6-1 | 141.4531 | 267.366872 | 251.796328 | 193.804445 | 232.753102 |
| tRNA-Asp-GTC-1-11 | 7029.06768 | 3526.27196 | 3069.57894 | 3359.27705 | 3638.23945 |
| tRNA-Asp-GTC-1-6 | 7007.68524 | 3512.4085 | 3070.44422 | 3354.53565 | 3635.99604 |
| tRNA-Asp-GTC-1-8 | 7007.68524 | 3513.39874 | 3071.3095 | 3356.90635 | 3638.8003 |
| tRNA-Asp-GTC-1-13 | 7023.31087 | 3524.29147 | 3069.57894 | 3367.57448 | 3649.45646 |
| tRNA-Asp-GTC-1-12 | 7007.68524 | 3512.4085 | 3068.28103 | 3354.53565 | 3635.99604 |
| tRNA-Asp-GTC-1-10 | 7020.84367 | 3521.32073 | 3069.57894 | 3357.49902 | 3638.8003 |
| tRNA-Asp-GTC-1-1 | 7001.92843 | 3506.46701 | 3069.57894 | 3349.79426 | 3632.63094 |
| tRNA-Asp-GTC-1-7 | 7000.28362 | 3509.43775 | 3063.95463 | 3349.79426 | 3631.50924 |
| tRNA-Asp-GTC-1-2 | 6999.46122 | 3507.45726 | 3063.95463 | 3349.79426 | 3632.07009 |
| tRNA-Asp-GTC-1-9 | 7000.28362 | 3505.47676 | 3063.95463 | 3349.20159 | 3630.94839 |
| tRNA-Asp-GTC-1-3 | 7000.28362 | 3505.47676 | 3063.52199 | 3349.20159 | 3631.50924 |
| tRNA-Asp-GTC-1-4 | 7000.28362 | 3505.47676 | 3063.52199 | 3349.20159 | 3631.50924 |
| tRNA-Asp-GTC-1-5 | 7001.92843 | 3506.46701 | 3064.81991 | 3350.97961 | 3631.50924 |
| tRNA-Leu-TAG-1-1 | 1537.06885 | 1075.40897 | 1358.05614 | 1524.3579 | 1959.05201 |
| tRNA-Arg-TCG-2-1 | 83.0625759 | 136.654179 | 78.3077927 | 535.184752 | 92.5403898 |
| tRNA-Ala-AGC-5-2 | 76.4833619 | 111.897987 | 111.621053 | 88.9011215 | 102.635705 |
| tRNA-Lys-TTT-1-2 | 1395.61575 | 1637.86965 | 1643.59837 | 1295.58568 | 1446.99519 |
| tRNA-Ala-AGC-5-3 | 75.6609602 | 113.878482 | 112.053692 | 88.3084474 | 103.196556 |
| tRNA-Arg-TCT-2-1 | 101.977816 | 125.761454 | 170.027417 | 178.987591 | 138.530159 |
| tRNA-iMet-CAT-1-3 | 852.830606 | 1042.7308 | 881.719787 | 1002.21198 | 1092.53745 |
| tRNA-Cys-GCA-13-1 | 5.75681219 | 1.98049535 | 2.16319869 | 3.55604486 | 2.80425424 |
| tRNA-Ala-AGC-12-1 | 73.193755 | 110.907739 | 110.755773 | 88.9011215 | 102.635705 |
| tRNA-Leu-AAG-1-3 | 1287.88113 | 934.793803 | 1252.92468 | 1408.78644 | 1853.61205 |
| tRNA-His-GTG-2-4 | 3005.87836 | 1999.31005 | 3250.85499 | 2287.12952 | 2323.04421 |

| | | | | | |
|---|---|---|---|---|---|
| tRNA-His-GTG-2-5 | 3005.87836 | 1999.31005 | 3250.85499 | 2287.12952 | 2323.04421 |
| tRNA-His-GTG-2-6 | 3005.87836 | 1999.31005 | 3250.85499 | 2287.12952 | 2323.04421 |
| tRNA-His-GTG-2-7 | 3008.34557 | 1999.31005 | 3251.72027 | 2287.12952 | 2323.04421 |
| tRNA-His-GTG-2-8 | 3006.70077 | 1999.31005 | 3251.72027 | 2287.12952 | 2323.60506 |
| tRNA-His-GTG-2-2 | 3007.52317 | 1999.31005 | 3251.72027 | 2287.72219 | 2323.60506 |
| tRNA-His-GTG-2-1 | 3008.34557 | 1999.31005 | 3253.45083 | 2287.72219 | 2323.60506 |
| tRNA-His-GTG-2-3 | 3007.52317 | 2000.3003 | 3251.72027 | 2287.72219 | 2323.60506 |
| tRNA-Ala-AGC-5-1 | 77.3057637 | 112.888235 | 115.94745 | 89.4937957 | 104.879108 |
| tRNA-Ala-AGC-10-1 | 74.8385585 | 111.897987 | 111.621053 | 88.9011215 | 102.635705 |
| tRNA-Leu-AAG-1-2 | 1273.9003 | 928.852317 | 1225.23574 | 1395.74761 | 1832.86057 |
| tRNA-Leu-AAG-1-1 | 1268.14349 | 928.852317 | 1223.93782 | 1397.52563 | 1832.86057 |
| tRNA-Ser-CGA-1-1 | 37.0080784 | 29.7074302 | 43.6966136 | 48.5992798 | 55.5242339 |
| tRNA-Lys-TTT-1-5 | 1427.68942 | 1656.68436 | 1694.64986 | 1297.3637 | 1473.91603 |
| tRNA-Tyr-GTA-6-1 | 9.04641915 | 16.8342104 | 31.1500612 | 25.4849882 | 17.3863763 |
| tRNA-Ser-CGA-3-1 | 32.0736679 | 25.7464395 | 31.1500612 | 40.3018418 | 49.9157254 |
| tRNA-Lys-TTT-1-4 | 1428.51182 | 1650.74287 | 1685.13178 | 1296.77103 | 1460.45561 |
| tRNA-Ile-TAT-2-3 | 439.16253 | 546.616716 | 1103.23133 | 490.734191 | 377.45262 |
| tRNA-Gly-CCC-2-1 | 595.418861 | 836.759284 | 529.1184 | 526.887314 | 415.029627 |
| tRNA-Gly-CCC-2-2 | 595.418861 | 836.759284 | 529.1184 | 526.887314 | 415.029627 |
| tRNA-Val-CAC-4-1 | 2093.83483 | 2523.15107 | 791.730721 | 1532.65534 | 1183.95614 |
| tRNA-Lys-TTT-2-1 | 139.808296 | 181.215324 | 218.483068 | 153.502603 | 110.487617 |
| tRNA-Lys-TTT-2-2 | 139.808296 | 181.215324 | 218.483068 | 153.502603 | 110.487617 |
| tRNA-Tyr-GTA-3-2 | 96.2210037 | 120.810216 | 165.26838 | 99.5692561 | 58.3284881 |
| tRNA-iMet-CAT-1-2 | 825.691348 | 1013.02337 | 869.605874 | 980.283034 | 1071.22512 |
| tRNA-iMet-CAT-1-7 | 850.3634 | 1015.99411 | 903.351774 | 986.802449 | 1086.92894 |
| tRNA-Ala-AGC-7-1 | 106.912226 | 236.669194 | 208.099714 | 169.504805 | 208.636515 |
| tRNA-Lys-TTT-5-1 | 455.610565 | 394.118574 | 435.235577 | 312.931948 | 375.770068 |
| tRNA-iMet-CAT-1-1 | 831.44816 | 1015.99411 | 875.230191 | 981.468382 | 1071.78597 |
| tRNA-iMet-CAT-1-4 | 821.57934 | 1014.01362 | 866.144756 | 977.912337 | 1067.29916 |
| tRNA-Leu-TAA-1-1 | 99.5106107 | 153.488389 | 110.755773 | 181.950962 | 181.715675 |
| tRNA-Leu-AAG-2-1 | 397.220041 | 266.376624 | 324.912444 | 422.576664 | 445.315573 |
| tRNA-iMet-CAT-1-6 | 831.44816 | 1022.92585 | 870.038514 | 986.802449 | 1073.46852 |
| tRNA-iMet-CAT-1-5 | 846.251392 | 1036.78931 | 876.96075 | 993.914539 | 1086.36809 |
| tRNA-Lys-TTT-1-6 | 1463.0527 | 1663.61609 | 1737.91383 | 1306.25381 | 1487.37645 |
| tRNA-Pro-TGG-5-1 | 2.46720522 | 1.98049535 | 2.16319869 | 2.37069657 | 1.68255254 |
| tRNA-Lys-TTT-1-3 | 1463.8751 | 1664.60634 | 1738.77911 | 1307.43916 | 1489.059 |
| tRNA-iMet-CAT-3-1 | 705.620694 | 816.95433 | 502.727376 | 753.288837 | 874.927322 |
| tRNA-Lys-CTT-6-1 | 16.4480348 | 27.7269348 | 50.1862097 | 38.5238193 | 25.799139 |
| tRNA-Met-CAT-7-1 | 0 | 0.99024767 | 0.43263974 | 0.59267414 | 0.56085085 |
| tRNA-Met-CAT-4-1 | 333.072705 | 411.943032 | 840.186372 | 397.091676 | 377.45262 |
| tRNA-Ser-GCT-4-1 | 796.907287 | 596.129099 | 533.012158 | 641.866098 | 945.033678 |
| tRNA-Ser-GCT-4-3 | 795.262484 | 595.138852 | 530.416319 | 645.422142 | 947.837932 |
| tRNA-Arg-ACG-1-3 | 28.7840609 | 50.5026313 | 57.973725 | 64.6014817 | 65.0586983 |
| tRNA-Ser-GCT-2-1 | 792.795279 | 599.099842 | 526.522562 | 640.088075 | 941.668573 |
| tRNA-Gln-TTG-6-1 | 2.46720522 | 1.98049535 | 8.22015503 | 8.29743801 | 4.48680678 |
| tRNA-Met-CAT-5-1 | 130.761877 | 195.078792 | 382.020889 | 183.13631 | 164.329298 |

| | | | | | |
|---|---|---|---|---|---|
| tRNA-Ser-GCT-4-2 | 794.440082 | 594.148604 | 532.146878 | 643.051446 | 947.837932 |
| tRNA-Tyr-GTA-4-1 | 53.4561132 | 83.1808045 | 142.771114 | 93.0498406 | 48.2331729 |
| tRNA-Glu-CTC-4-1 | 502.487464 | 604.051081 | 459.463402 | 544.074864 | 481.210027 |
| tRNA-Met-CAT-3-1 | 379.949604 | 446.601701 | 893.40106 | 425.540035 | 418.394732 |
| tRNA-Asp-GTC-3-1 | 128.294672 | 100.015015 | 65.7612402 | 73.4915938 | 72.9106102 |
| tRNA-Tyr-GTA-3-1 | 48.5217027 | 90.1125383 | 135.416238 | 91.8644923 | 44.3072169 |
| tRNA-Ser-GCT-1-1 | 783.748859 | 589.197366 | 522.196164 | 631.197963 | 932.134108 |
| tRNA-Lys-TTT-3-1 | 5.75681219 | 2.97074302 | 13.8444716 | 10.6681346 | 5.60850847 |
| tRNA-Gly-GCC-1-1 | 12998.8819 | 12187.9684 | 7710.50542 | 8482.35234 | 8042.60115 |
| tRNA-Gly-GCC-1-2 | 12998.8819 | 12187.9684 | 7710.50542 | 8482.35234 | 8042.60115 |
| tRNA-Gly-GCC-1-3 | 12998.8819 | 12187.9684 | 7710.50542 | 8482.35234 | 8042.60115 |
| tRNA-Val-TAC-1-1 | 5732.14014 | 4589.79797 | 4028.74124 | 4213.32049 | 4571.49526 |
| tRNA-Lys-CTT-4-1 | 1.64480348 | 1.98049535 | 6.92223582 | 5.92674144 | 2.80425424 |
| tRNA-Lys-TTT-1-1 | 1421.93261 | 1659.6551 | 1666.09563 | 1294.40033 | 1471.67262 |
| tRNA-Leu-AAG-3-1 | 391.463229 | 262.415633 | 321.018686 | 413.093878 | 439.707064 |
| tRNA-Lys-CTT-10-1 | 19.7376418 | 24.7561918 | 47.5903712 | 37.3384711 | 37.0161559 |
| tRNA-Val-AAC-4-1 | 452.320958 | 961.530491 | 814.227988 | 1091.70577 | 593.941047 |
| tRNA-Val-TAC-1-3 | 5668.8152 | 4540.28558 | 3932.69522 | 4175.98202 | 4526.06634 |
| tRNA-Val-TAC-1-2 | 5667.9928 | 4540.28558 | 3932.69522 | 4176.57469 | 4526.62719 |
| tRNA-Tyr-GTA-1-2 | 22.204847 | 35.6489162 | 68.3570787 | 42.0798642 | 19.6297797 |
| tRNA-Tyr-GTA-1-1 | 26.3168557 | 35.6489162 | 68.3570787 | 42.0798642 | 19.6297797 |
| tRNA-Leu-CAG-3-1 | 91.2865933 | 94.073529 | 98.2092206 | 113.200761 | 111.609319 |
| tRNA-Tyr-GTA-1-3 | 22.204847 | 35.6489162 | 69.2223582 | 42.0798642 | 20.1906305 |
| tRNA-Arg-ACG-2-1 | 21.3824453 | 36.6391639 | 65.7612402 | 49.1919539 | 49.9157254 |
| tRNA-His-GTG-3-1 | 81.4177724 | 144.57616 | 180.410771 | 163.578064 | 72.9106102 |
| tRNA-Tyr-GTA-1-4 | 29.6064627 | 39.6099069 | 71.8181966 | 43.8578866 | 20.7514814 |
| tRNA-Tyr-GTA-5-1 | 14.8032313 | 24.7561918 | 54.0799673 | 32.5970779 | 12.3387186 |
| tRNA-Tyr-GTA-1-5 | 24.6720522 | 34.6586686 | 69.2223582 | 43.2652125 | 20.1906305 |
| tRNA-Ser-GCT-5-1 | 25.494454 | 20.7952011 | 16.8729498 | 15.4095277 | 20.1906305 |
| tRNA-Ile-AAT-1-8 | 74.0161567 | 85.1612999 | 173.488535 | 121.498199 | 118.90038 |
| tRNA-Gln-TTG-2-1 | 143.920305 | 210.922754 | 250.065769 | 233.513613 | 199.662902 |
| tRNA-Ala-CGC-3-1 | 1.64480348 | 1.98049535 | 1.29791922 | 2.37069657 | 3.36510508 |
| tRNA-Ala-CGC-3-2 | 1.64480348 | 1.98049535 | 1.29791922 | 2.37069657 | 3.36510508 |
| tRNA-Arg-ACG-1-1 | 27.1392575 | 49.5123837 | 56.6758057 | 64.6014817 | 63.9369966 |
| tRNA-Ile-AAT-1-6 | 74.0161567 | 85.1612999 | 163.537821 | 116.756806 | 117.778678 |
| tRNA-Val-CAC-1-1 | 1083.9255 | 1800.27027 | 1186.7308 | 1175.8655 | 1074.59022 |
| tRNA-Arg-ACG-1-2 | 26.3168557 | 49.5123837 | 56.243166 | 64.6014817 | 64.4978475 |
| tRNA-Ala-CGC-1-2 | 106.912226 | 90.1125383 | 112.918972 | 107.866694 | 121.704634 |
| tRNA-Ile-AAT-1-4 | 69.904148 | 84.1710522 | 159.211424 | 115.571458 | 114.974424 |
| tRNA-Ser-GCT-3-1 | 856.120213 | 654.553712 | 625.597062 | 765.734994 | 1022.99195 |
| tRNA-Ile-AAT-1-1 | 72.3713532 | 84.1710522 | 164.83574 | 116.164132 | 118.90038 |
| tRNA-Met-CAT-6-1 | 20.5600435 | 43.5708976 | 103.400897 | 48.0066056 | 31.4076475 |
| tRNA-Ala-CGC-4-1 | 1.64480348 | 0 | 0.43263974 | 0.59267414 | 1.12170169 |
| tRNA-Leu-TAG-2-1 | 403.799255 | 283.210835 | 324.912444 | 425.540035 | 452.045783 |
| tRNA-Ile-AAT-1-7 | 71.5489515 | 85.1612999 | 161.374622 | 116.164132 | 117.217827 |
| tRNA-Asn-GTT-5-1 | 0.82240174 | 0 | 0 | 0.59267414 | 1.12170169 |

| | | | | | |
|---|---|---|---|---|---|
| tRNA-Ile-AAT-3-1 | 69.0817463 | 82.1905569 | 157.048225 | 112.608087 | 112.73102 |
| tRNA-Val-AAC-1-1 | 11794.8858 | 19965.3736 | 10042.4336 | 13327.4635 | 11670.1844 |
| tRNA-Val-AAC-1-2 | 11787.4842 | 19966.3638 | 10029.4544 | 13323.3147 | 11663.4542 |
| tRNA-Pro-TGG-4-1 | 31.2512662 | 34.6586686 | 21.6319869 | 29.041033 | 37.0161559 |
| tRNA-Ala-CGC-3-3 | 1.64480348 | 1.98049535 | 1.29791922 | 2.37069657 | 3.36510508 |
| tRNA-Val-CAC-2-1 | 13562.2271 | 21879.5223 | 10707.8335 | 14436.9495 | 12348.2531 |
| tRNA-Val-CAC-2-5 | 13556.4703 | 21880.5126 | 10701.7766 | 14431.6154 | 12350.4965 |
| tRNA-Ile-AAT-1-2 | 70.7265498 | 83.1808045 | 160.076703 | 115.571458 | 114.974424 |
| tRNA-Val-CAC-2-3 | 13560.5823 | 21887.4443 | 10703.0745 | 14432.2081 | 12350.4965 |
| tRNA-Ile-AAT-1-3 | 69.904148 | 83.1808045 | 160.509343 | 116.164132 | 115.535275 |
| tRNA-Ile-AAT-1-5 | 72.3713532 | 83.1808045 | 160.076703 | 115.571458 | 116.096125 |
| tRNA-Val-CAC-2-2 | 13764.5379 | 22225.1188 | 10871.3713 | 14679.9459 | 12546.7943 |
| tRNA-Gln-TTG-1-1 | 374.192792 | 475.318883 | 734.622276 | 696.984793 | 558.046593 |
| tRNA-Val-CAC-2-4 | 13716.0162 | 22114.211 | 10829.8379 | 14647.3488 | 12515.9475 |
| tRNA-Lys-CTT-15-1 | 19.7376418 | 24.7561918 | 54.512607 | 40.3018418 | 50.4765763 |
| tRNA-Gly-ACC-1-1 | 129.117073 | 325.791484 | 281.64847 | 309.375903 | 272.573512 |
| tRNA-Val-CAC-3-1 | 346.231133 | 674.358665 | 460.761321 | 614.010413 | 446.437275 |
| tRNA-Val-AAC-3-1 | 274.682182 | 609.992567 | 425.284863 | 552.372302 | 427.929197 |
| tRNA-Ile-AAT-2-1 | 32.8960697 | 42.5806499 | 76.5772337 | 62.2307851 | 54.4025322 |
| tRNA-Ile-AAT-4-1 | 30.4288644 | 42.5806499 | 73.1161158 | 60.4527627 | 53.8416814 |
| tRNA-Cys-GCA-26-1 | 3.28960697 | 3.96099069 | 7.78751529 | 5.33406729 | 3.36510508 |
| tRNA-Cys-GCA-23-1 | 3.28960697 | 3.96099069 | 7.35487555 | 5.33406729 | 3.36510508 |
| tRNA-Arg-TCG-1-1 | 40.2976853 | 41.5904023 | 58.8390044 | 51.5626505 | 52.7199797 |
| tRNA-Arg-CCG-3-1 | 493.441045 | 501.065323 | 408.844553 | 603.342278 | 505.326614 |
| tRNA-Gly-TCC-1-6 | 9519.30016 | 9858.90583 | 4572.5694 | 5910.73923 | 8431.27079 |
| tRNA-Gly-TCC-2-1 | 9417.32234 | 9757.90057 | 4418.54965 | 5796.35313 | 8303.95765 |
| tRNA-Leu-TAA-3-1 | 0.82240174 | 1.98049535 | 3.89375765 | 3.55604486 | 1.12170169 |
| tRNA-Arg-TCT-5-1 | 0.82240174 | 9.90247673 | 2.59583843 | 3.55604486 | 2.24340339 |
| tRNA-Val-AAC-5-1 | 2912.94697 | 5502.80632 | 3182.93056 | 4001.73582 | 3269.76044 |
| tRNA-Gly-TCC-1-5 | 9455.97522 | 9835.13989 | 4487.77201 | 5878.14216 | 8383.03762 |
| tRNA-Gly-TCC-1-1 | 9455.97522 | 9835.13989 | 4486.04145 | 5878.73483 | 8384.15932 |
| tRNA-Gly-TCC-1-7 | 9457.62002 | 9840.09113 | 4490.80049 | 5879.32751 | 8385.84187 |
| tRNA-Gly-TCC-1-3 | 9470.77845 | 9849.00336 | 4491.23312 | 5885.84692 | 8399.30229 |
| tRNA-Gly-TCC-1-4 | 9470.77845 | 9849.00336 | 4491.23312 | 5885.84692 | 8399.30229 |
| tRNA-Gly-TCC-1-2 | 9472.42326 | 9849.99361 | 4492.53104 | 5887.03227 | 8401.54569 |
| tRNA-Ala-AGC-8-1 | 28.7840609 | 37.6294116 | 25.0931048 | 32.0044038 | 46.5506203 |
| tRNA-Ala-CGC-6-1 | 1.64480348 | 1.98049535 | 1.29791922 | 1.77802243 | 3.36510508 |
| tRNA-Gln-CTG-3-3 | 463.834582 | 582.265632 | 990.312361 | 855.228789 | 715.645681 |
| tRNA-Gln-CTG-3-2 | 463.834582 | 582.265632 | 990.312361 | 855.228789 | 715.645681 |
| tRNA-Gln-CTG-3-1 | 463.834582 | 584.246127 | 992.47556 | 856.414138 | 716.767383 |
| tRNA-Phe-GAA-3-1 | 7.40161567 | 4.95123837 | 9.51807425 | 5.92674144 | 4.48680678 |
| tRNA-Thr-CGT-2-1 | 126.649868 | 144.57616 | 234.923378 | 113.793436 | 144.699519 |
| tRNA-Thr-AGT-6-1 | 13.1584279 | 9.90247673 | 9.95071398 | 7.11208972 | 8.97361356 |
| tRNA-Cys-GCA-20-1 | 4.11200871 | 4.95123837 | 8.22015503 | 5.92674144 | 3.36510508 |
| tRNA-Gln-CTG-5-1 | 269.747771 | 290.142568 | 589.255324 | 541.111493 | 421.198986 |
| tRNA-Gln-CTG-4-1 | 278.79419 | 310.937769 | 619.972745 | 557.706369 | 435.781108 |

| | | | | | |
|---|---|---|---|---|---|
| tRNA-Lys-CTT-16-1 | 26.3168557 | 32.6781732 | 66.6265197 | 48.0066056 | 63.9369966 |
| tRNA-Phe-GAA-1-1 | 28.7840609 | 12.8732198 | 31.1500612 | 18.3728985 | 17.9472271 |
| tRNA-Met-CAT-1-1 | 510.711481 | 737.734517 | 1534.57315 | 621.715177 | 666.851658 |
| tRNA-Met-CAT-2-2 | 528.80432 | 581.275384 | 1038.33537 | 523.331269 | 579.919776 |
| tRNA-Cys-GCA-21-1 | 2.46720522 | 2.97074302 | 7.35487555 | 4.14871901 | 2.24340339 |
| tRNA-Met-CAT-1-2 | 523.869909 | 743.676003 | 1552.74402 | 628.234592 | 677.507824 |
| tRNA-Met-CAT-2-1 | 528.80432 | 583.255879 | 1038.33537 | 522.738595 | 579.358925 |
| tRNA-Ser-GCT-6-1 | 120.893056 | 118.829721 | 146.664871 | 112.608087 | 183.398227 |
| tRNA-Ile-TAT-2-1 | 32.0736679 | 45.551393 | 129.359282 | 62.2307851 | 43.7463661 |
| tRNA-Lys-CTT-5-1 | 0.82240174 | 0.99024767 | 2.16319869 | 1.77802243 | 1.68255254 |
| tRNA-Gly-GCC-6-1 | 6232.9828 | 6293.02396 | 3718.10591 | 3835.78706 | 4631.5063 |
| tRNA-Cys-GCA-12-1 | 25.494454 | 21.7854488 | 16.0076703 | 23.1142916 | 23.5557356 |
| tRNA-Ile-TAT-2-2 | 30.4288644 | 43.5708976 | 122.869686 | 59.2674144 | 42.6246644 |
| tRNA-Gln-CTG-2-4 | 495.90825 | 614.943805 | 1030.9805 | 878.343081 | 747.053329 |
| tRNA-Gln-CTG-2-3 | 495.90825 | 614.943805 | 1034.87425 | 879.528429 | 749.296732 |
| tRNA-Ala-CGC-2-1 | 54.2785149 | 63.3758511 | 55.3778865 | 71.7135714 | 80.762522 |
| tRNA-Gln-CTG-2-2 | 496.730652 | 615.934053 | 1034.44161 | 878.935755 | 748.17503 |
| tRNA-Gln-CTG-2-1 | 498.375455 | 614.943805 | 1037.03745 | 880.121103 | 750.418434 |
| tRNA-Cys-GCA-14-1 | 25.494454 | 21.7854488 | 16.0076703 | 22.5216175 | 24.6774373 |
| tRNA-Trp-CCA-6-1 | 15.6256331 | 28.7171825 | 42.3986944 | 37.9311452 | 21.873183 |
| tRNA-Cys-GCA-24-1 | 25.494454 | 21.7854488 | 16.8729498 | 23.1142916 | 24.1165864 |
| tRNA-Arg-TCT-4-1 | 90.4641915 | 88.1320429 | 150.125989 | 164.170738 | 130.678247 |
| tRNA-Glu-CTC-5-1 | 305.111046 | 355.498915 | 274.726234 | 332.490195 | 278.742871 |
| tRNA-Gln-CTG-6-1 | 190.797204 | 294.103559 | 389.375765 | 340.787633 | 306.224563 |
| tRNA-Val-AAC-6-1 | 0 | 1.98049535 | 0 | 0 | 0 |
| tRNA-Gln-CTG-1-1 | 202.310828 | 313.908512 | 433.505018 | 364.494598 | 335.949658 |
| tRNA-Leu-CAA-2-1 | 459.722573 | 440.660215 | 648.094328 | 466.434551 | 574.872119 |
| tRNA-Cys-GCA-19-1 | 3.28960697 | 2.97074302 | 2.59583843 | 2.37069657 | 1.12170169 |
| tRNA-Gln-CTG-7-1 | 4.11200871 | 1.98049535 | 4.32639738 | 6.51941558 | 3.92595593 |
| tRNA-Gln-TTG-3-2 | 390.640827 | 429.76749 | 709.529171 | 709.43095 | 546.268725 |
| tRNA-Ala-AGC-2-2 | 101.977816 | 76.2490708 | 114.216891 | 109.644717 | 108.244214 |
| tRNA-Gln-TTG-3-1 | 390.640827 | 429.76749 | 712.12501 | 710.023624 | 546.829576 |
| tRNA-Cys-GCA-7-1 | 74.0161567 | 71.2978325 | 117.245369 | 107.27402 | 83.0059254 |
| tRNA-iMet-CAT-4-1 | 0 | 0.99024767 | 1.29791922 | 0 | 0.56085085 |
| tRNA-Leu-CAA-1-1 | 127.47227 | 114.86873 | 119.408568 | 148.168536 | 137.408458 |
| tRNA-Lys-CTT-8-1 | 0.82240174 | 0.99024767 | 2.16319869 | 2.37069657 | 1.68255254 |
| tRNA-Ala-AGC-3-1 | 39.4752836 | 46.5416406 | 43.2639738 | 47.4139315 | 66.1804 |
| tRNA-Trp-CCA-2-1 | 844.606588 | 919.940088 | 1130.92028 | 1003.39733 | 893.996251 |
| tRNA-Ile-AAT-5-1 | 0.82240174 | 0.99024767 | 0.43263974 | 0 | 0.56085085 |
| tRNA-Pro-AGG-1-2 | 162.013143 | 158.439628 | 106.862015 | 136.907727 | 152.55143 |
| tRNA-Pro-AGG-1-5 | 164.480348 | 158.439628 | 109.025214 | 138.68575 | 156.477386 |
| tRNA-Ile-TAT-1-1 | 90.4641915 | 87.1417952 | 177.382293 | 86.530425 | 72.3497593 |
| tRNA-Arg-TCT-6-1 | 0 | 0 | 0 | 1.18534829 | 0.56085085 |
| tRNA-Pro-AGG-1-4 | 167.769955 | 159.429875 | 112.486332 | 139.871098 | 155.355685 |
| tRNA-Leu-TAG-4-1 | 0.82240174 | 0.99024767 | 1.29791922 | 1.18534829 | 1.12170169 |
| tRNA-Gln-TTG-4-1 | 0 | 0 | 1.73055895 | 1.77802243 | 0.56085085 |

| | | | | | |
|---|---|---|---|---|---|
| tRNA-Ala-AGC-2-1 | 100.333012 | 73.2783278 | 112.053692 | 110.237391 | 107.683363 |
| tRNA-Pro-AGG-1-1 | 175.171571 | 170.3226 | 122.437046 | 144.019817 | 161.525044 |
| tRNA-Arg-TCG-4-1 | 43.5872923 | 70.3075848 | 89.5564259 | 307.597881 | 75.7148644 |
| tRNA-Pro-AGG-1-6 | 174.349169 | 164.381114 | 123.734965 | 144.612491 | 162.085895 |
| tRNA-Ala-CGC-1-1 | 85.5297811 | 82.1905569 | 94.7481027 | 101.939953 | 109.926766 |
| tRNA-Pro-AGG-1-3 | 165.30275 | 160.420123 | 109.025214 | 137.500401 | 152.55143 |
| tRNA-Cys-GCA-17-1 | 7.40161567 | 5.94148604 | 9.95071398 | 7.70476387 | 5.04765763 |
| tRNA-Cys-GCA-9-1 | 26.3168557 | 22.7756965 | 17.3055895 | 23.1142916 | 24.1165864 |
| tRNA-Cys-GCA-8-1 | 49.3441045 | 52.4831267 | 98.2092206 | 85.9377508 | 62.8152949 |
| tRNA-Ile-GAT-1-1 | 0.82240174 | 1.98049535 | 0.86527948 | 1.77802243 | 1.68255254 |
| tRNA-Pro-TGG-1-1 | 152.966724 | 143.585913 | 96.0460219 | 133.351682 | 144.138668 |
| tRNA-Phe-GAA-1-5 | 13.9808296 | 11.8829721 | 20.3340677 | 14.2241794 | 12.3387186 |
| tRNA-Glu-TTC-2-1 | 20465.4673 | 26176.207 | 11066.0592 | 14000.7413 | 15761.0305 |
| tRNA-Ala-AGC-18-1 | 0 | 0 | 0.43263974 | 0 | 0 |
| tRNA-Glu-TTC-2-2 | 20459.7105 | 26161.3533 | 11077.7405 | 13990.0732 | 15782.9037 |
| tRNA-Thr-AGT-2-1 | 209.712444 | 138.634674 | 115.51481 | 124.46157 | 191.250139 |
| tRNA-Leu-TAA-4-1 | 0 | 6.93173371 | 6.05695634 | 2.37069657 | 8.41276271 |
| tRNA-Pro-TGG-2-1 | 191.619606 | 184.186067 | 125.898164 | 167.726783 | 183.398227 |
| tRNA-Leu-CAG-4-1 | 50.988908 | 79.2198139 | 89.9890656 | 125.054244 | 93.6620915 |
| tRNA-Pro-TGG-2-4 | 189.974802 | 187.15681 | 128.494002 | 166.541434 | 181.715675 |
| tRNA-Cys-GCA-22-1 | 0 | 0 | 0 | 0 | 0 |
| tRNA-Ala-TGC-4-1 | 67.4369428 | 60.4051081 | 78.7404324 | 74.084268 | 86.3710305 |
| tRNA-Cys-GCA-4-3 | 27.9616592 | 24.7561918 | 22.4972664 | 26.6703365 | 26.3599898 |
| tRNA-Cys-GCA-4-5 | 27.9616592 | 24.7561918 | 22.4972664 | 26.6703365 | 26.3599898 |
| tRNA-Cys-GCA-4-6 | 27.9616592 | 24.7561918 | 22.4972664 | 26.6703365 | 26.3599898 |
| tRNA-Cys-GCA-4-7 | 27.9616592 | 24.7561918 | 22.4972664 | 26.6703365 | 26.3599898 |
| tRNA-Cys-GCA-4-14 | 27.9616592 | 24.7561918 | 22.4972664 | 26.6703365 | 26.3599898 |
| tRNA-Cys-GCA-4-15 | 27.9616592 | 24.7561918 | 22.4972664 | 26.6703365 | 26.3599898 |
| tRNA-Cys-GCA-25-1 | 4.11200871 | 5.94148604 | 5.19167686 | 4.74139315 | 3.92595593 |
| tRNA-Phe-GAA-1-3 | 13.9808296 | 12.8732198 | 18.6035088 | 14.8168536 | 13.4604203 |
| tRNA-Ile-TAT-3-1 | 0 | 0 | 0.86527948 | 0.59267414 | 2.24340339 |
| tRNA-Pro-TGG-2-3 | 189.974802 | 184.186067 | 126.330804 | 167.726783 | 182.276525 |
| tRNA-Lys-CTT-19-1 | 0.82240174 | 0 | 0 | 0 | 0 |
| tRNA-Tyr-GTA-2-1 | 85.5297811 | 96.0540243 | 138.444716 | 74.084268 | 58.3284881 |
| tRNA-Trp-CCA-3-1 | 49.3441045 | 69.3173371 | 78.7404324 | 126.832267 | 91.979539 |
| tRNA-Cys-GCA-4-16 | 27.9616592 | 24.7561918 | 22.4972664 | 26.6703365 | 25.799139 |
| tRNA-Cys-GCA-6-1 | 27.9616592 | 24.7561918 | 22.4972664 | 26.6703365 | 25.799139 |
| tRNA-Cys-GCA-4-18 | 27.9616592 | 24.7561918 | 22.4972664 | 26.6703365 | 25.799139 |
| tRNA-Cys-GCA-4-19 | 27.9616592 | 24.7561918 | 22.4972664 | 26.6703365 | 25.799139 |
| tRNA-Cys-GCA-4-20 | 27.9616592 | 24.7561918 | 22.4972664 | 26.6703365 | 25.799139 |
| tRNA-Cys-GCA-4-24 | 27.9616592 | 24.7561918 | 22.4972664 | 26.6703365 | 25.799139 |
| tRNA-Cys-GCA-4-26 | 27.9616592 | 24.7561918 | 22.4972664 | 26.6703365 | 25.799139 |
| tRNA-Cys-GCA-4-1 | 27.9616592 | 23.7659442 | 22.4972664 | 26.6703365 | 26.3599898 |
| tRNA-Cys-GCA-4-17 | 27.9616592 | 23.7659442 | 22.4972664 | 26.6703365 | 26.3599898 |
| tRNA-Cys-GCA-4-21 | 27.9616592 | 23.7659442 | 22.4972664 | 26.6703365 | 26.3599898 |
| tRNA-Cys-GCA-10-1 | 27.9616592 | 23.7659442 | 22.4972664 | 26.6703365 | 26.3599898 |

| | | | | | |
|---|---|---|---|---|---|
| tRNA-Cys-GCA-4-27 | 27.9616592 | 23.7659442 | 22.4972664 | 26.6703365 | 26.3599898 |
| tRNA-Cys-GCA-4-29 | 27.9616592 | 23.7659442 | 22.4972664 | 26.6703365 | 26.3599898 |
| tRNA-Pro-CGG-1-3 | 178.461178 | 176.264086 | 112.053692 | 148.76121 | 163.207597 |
| tRNA-Lys-CTT-14-1 | 10.6912226 | 14.8537151 | 73.1161158 | 37.3384711 | 29.7250949 |
| tRNA-Ala-AGC-1-1 | 46.8768993 | 38.6196593 | 28.121583 | 39.7091676 | 52.7199797 |
| tRNA-Cys-GCA-18-1 | 0.82240174 | 0.99024767 | 0.43263974 | 0.59267414 | 0 |
| tRNA-Cys-GCA-4-2 | 27.9616592 | 23.7659442 | 22.4972664 | 26.6703365 | 25.799139 |
| tRNA-Cys-GCA-4-4 | 27.9616592 | 23.7659442 | 22.4972664 | 26.6703365 | 25.799139 |
| tRNA-Cys-GCA-4-8 | 27.9616592 | 23.7659442 | 22.4972664 | 26.6703365 | 25.799139 |
| tRNA-Cys-GCA-4-10 | 27.9616592 | 23.7659442 | 22.4972664 | 26.6703365 | 25.799139 |
| tRNA-Cys-GCA-4-11 | 27.9616592 | 23.7659442 | 22.4972664 | 26.6703365 | 25.799139 |
| tRNA-Cys-GCA-4-22 | 27.9616592 | 23.7659442 | 22.4972664 | 26.6703365 | 25.799139 |
| tRNA-Cys-GCA-4-23 | 27.9616592 | 23.7659442 | 22.4972664 | 26.6703365 | 25.799139 |
| tRNA-Cys-GCA-4-25 | 27.9616592 | 23.7659442 | 22.4972664 | 26.6703365 | 25.799139 |
| tRNA-Cys-GCA-4-28 | 27.9616592 | 23.7659442 | 22.4972664 | 26.6703365 | 25.799139 |
| tRNA-Leu-CAA-4-1 | 145.565108 | 120.810216 | 128.926642 | 154.687951 | 139.651861 |
| tRNA-Ala-TGC-1-1 | 41.9424888 | 25.7464395 | 36.341738 | 47.4139315 | 47.1114712 |
| tRNA-Arg-TCT-3-1 | 50.1665062 | 67.3368418 | 60.1369236 | 141.64912 | 98.1488983 |
| tRNA-Lys-CTT-13-1 | 41.9424888 | 90.1125383 | 120.706487 | 89.4937957 | 63.9369966 |
| tRNA-Lys-CTT-18-1 | 41.1200871 | 88.1320429 | 119.841208 | 88.9011215 | 63.9369966 |
| tRNA-Trp-CCA-3-2 | 49.3441045 | 73.2783278 | 80.0383516 | 129.202963 | 90.2969864 |
| tRNA-iMet-CAT-2-1 | 213.824453 | 341.635447 | 481.095389 | 387.016216 | 388.669637 |
| tRNA-Ala-TGC-3-1 | 87.1745846 | 80.2100615 | 102.968258 | 106.681346 | 111.048468 |
| tRNA-Thr-CGT-1-1 | 142.275501 | 116.849225 | 283.379029 | 75.2696162 | 125.63059 |
| tRNA-Cys-GCA-4-9 | 27.1392575 | 24.7561918 | 22.4972664 | 26.6703365 | 25.799139 |
| tRNA-Cys-GCA-4-12 | 27.1392575 | 24.7561918 | 22.4972664 | 26.6703365 | 25.799139 |
| tRNA-Cys-GCA-4-13 | 27.1392575 | 24.7561918 | 22.4972664 | 26.6703365 | 25.799139 |
| tRNA-Ser-GGA-1-1 | 0.82240174 | 0.99024767 | 2.59583843 | 1.77802243 | 1.12170169 |
| tRNA-Lys-CTT-7-1 | 2.46720522 | 2.97074302 | 5.19167686 | 4.14871901 | 3.36510508 |
| tRNA-Leu-CAG-2-3 | 1973.76418 | 1665.59659 | 1659.1734 | 2156.74121 | 2602.34793 |
| tRNA-Sup-TTA-1-1 | 0 | 0 | 0.43263974 | 0 | 0 |
| tRNA-Cys-GCA-16-1 | 5.75681219 | 4.95123837 | 9.95071398 | 10.6681346 | 6.73021017 |
| tRNA-Thr-AGT-3-1 | 148.032313 | 133.683436 | 160.076703 | 45.6359091 | 95.3446441 |
| tRNA-Pro-TGG-2-2 | 195.731614 | 187.15681 | 127.628723 | 170.690153 | 185.08078 |
| tRNA-Glu-TTC-1-1 | 16959.5687 | 18944.4282 | 8741.05328 | 11523.3634 | 12259.0778 |
| tRNA-Glu-TTC-1-2 | 16948.0551 | 18940.4672 | 8736.29424 | 11521.5854 | 12251.7868 |
| tRNA-Glu-TTC-1-4 | 17009.7352 | 19005.8236 | 8771.33806 | 11572.5553 | 12297.2157 |
| tRNA-Thr-AGT-1-1 | 154.611527 | 140.61517 | 170.460057 | 60.4527627 | 122.265485 |
| tRNA-Lys-CTT-12-1 | 0 | 0 | 0.86527948 | 1.18534829 | 0.56085085 |
| tRNA-Thr-AGT-1-2 | 153.789126 | 140.61517 | 170.027417 | 60.4527627 | 122.826336 |
| tRNA-Thr-AGT-1-3 | 153.789126 | 140.61517 | 170.460057 | 61.0454368 | 121.704634 |
| tRNA-Glu-TTC-1-3 | 16760.5475 | 18811.735 | 8683.07955 | 11457.5765 | 12165.4157 |
| tRNA-Thr-AGT-4-1 | 129.939475 | 106.946749 | 67.0591595 | 72.3062455 | 107.683363 |
| tRNA-Leu-CAG-2-1 | 1968.00737 | 1657.6746 | 1625.86014 | 2158.51923 | 2591.13092 |
| tRNA-Arg-CCG-2-1 | 55.1009167 | 70.3075848 | 48.8882904 | 53.3406729 | 108.244214 |
| tRNA-Leu-CAG-2-2 | 1940.04571 | 1649.75262 | 1567.45377 | 2134.21959 | 2555.79731 |

| | | | | | |
|---|---|---|---|---|---|
| tRNA-Phe-GAA-1-4 | 14.8032313 | 13.8634674 | 19.4687882 | 14.8168536 | 14.582122 |
| tRNA-Leu-CAG-1-4 | 1927.70968 | 1645.79163 | 1560.0989 | 2126.51483 | 2538.97179 |
| tRNA-Pro-CGG-1-1 | 188.329999 | 182.205572 | 121.139127 | 151.724581 | 169.937807 |
| tRNA-Leu-CAG-1-1 | 1923.59767 | 1638.8599 | 1551.87874 | 2129.4782 | 2538.97179 |
| tRNA-Leu-CAG-1-2 | 1923.59767 | 1638.8599 | 1551.87874 | 2129.4782 | 2538.97179 |
| tRNA-Leu-CAG-1-3 | 1923.59767 | 1638.8599 | 1551.87874 | 2129.4782 | 2538.97179 |
| tRNA-Leu-CAG-1-5 | 1926.88728 | 1641.83064 | 1554.47458 | 2132.44157 | 2535.60668 |
| tRNA-Val-CAC-7-1 | 1.64480348 | 4.95123837 | 4.32639738 | 7.70476387 | 4.48680678 |
| tRNA-Leu-CAA-3-1 | 370.903185 | 356.489162 | 544.693431 | 387.60889 | 448.680678 |
| tRNA-Ser-AGA-5-1 | 0 | 0.99024767 | 0 | 0 | 0 |
| tRNA-Pro-CGG-1-2 | 189.1524 | 186.166563 | 119.841208 | 155.8733 | 169.376956 |
| tRNA-Val-CAC-5-1 | 0.82240174 | 2.97074302 | 4.75903712 | 2.96337072 | 1.68255254 |
| tRNA-Ala-TGC-7-1 | 23.0272488 | 34.6586686 | 19.4687882 | 27.8556848 | 44.3072169 |
| tRNA-Ala-TGC-7-2 | 23.0272488 | 34.6586686 | 19.4687882 | 27.8556848 | 44.3072169 |
| tRNA-Ser-CGA-2-1 | 278.79419 | 239.639937 | 279.917911 | 349.085071 | 475.601519 |
| tRNA-Cys-GCA-15-1 | 2.46720522 | 1.98049535 | 2.16319869 | 2.96337072 | 2.80425424 |
| tRNA-Trp-CCA-1-1 | 62.5025323 | 125.761454 | 116.812729 | 168.319457 | 143.577817 |
| tRNA-Cys-GCA-28-1 | 0 | 0 | 0.86527948 | 0.59267414 | 0.56085085 |
| tRNA-Ala-TGC-8-1 | 21.3824453 | 33.6684209 | 18.6035088 | 26.6703365 | 43.7463661 |
| tRNA-Phe-GAA-1-2 | 17.2704366 | 14.8537151 | 18.170869 | 14.2241794 | 13.4604203 |
| tRNA-Cys-GCA-27-1 | 0 | 0 | 0 | 0 | 0 |
| tRNA-Val-CAC-6-1 | 333.895107 | 549.587459 | 210.262913 | 388.794238 | 286.033932 |
| tRNA-Arg-TCG-3-1 | 32.8960697 | 45.551393 | 48.023011 | 113.793436 | 53.2808305 |
| tRNA-Glu-CTC-6-1 | 1.64480348 | 2.97074302 | 3.46111791 | 3.55604486 | 1.12170169 |
| tRNA-Lys-CTT-9-1 | 0 | 0 | 0 | 0 | 0 |
| tRNA-Trp-CCA-4-1 | 25.494454 | 63.3758511 | 63.5980416 | 104.903323 | 72.3497593 |
| tRNA-Trp-CCA-4-2 | 25.494454 | 63.3758511 | 63.5980416 | 104.903323 | 72.3497593 |
| tRNA-Arg-TCT-1-1 | 64.9697376 | 84.1710522 | 112.053692 | 153.502603 | 117.778678 |
| tRNA-Arg-TCG-3-2 | 31.2512662 | 45.551393 | 47.1577315 | 113.200761 | 52.1591288 |

| FGCZ6510_Free_tRNA_dedup_counts | | | FGCZ6510_Free_tRNA_dedup_counts | | | FGCZ6510_F |
|---|---|---|---|---|---|---|
| CON.16 | CON.17 | CON.18 | DEX.3 | DEX.4 | DEX.5 | DEX.6 |
| 882.873008 | 660.132629 | 639.963207 | 106.794621 | 141.867357 | 122.077791 | 191.471937 |
| 244.317059 | 149.561299 | 163.394861 | 78.4028806 | 64.0003113 | 105.580792 | 78.6870973 |
| 5552.66042 | 6302.20369 | 9007.14174 | 1129.41824 | 1017.60495 | 389.329171 | 359.337745 |
| 677.424572 | 820.008499 | 878.24738 | 124.5069 | 352.001712 | 160.433313 | 168.914969 |
| 394.23889 | 335.2236 | 306.365365 | 1041.11732 | 476.802319 | 1026.52575 | 811.001683 |
| 466.423476 | 422.897465 | 544.649538 | 687.392698 | 558.936052 | 839.284813 | 934.278136 |
| 466.423476 | 422.897465 | 537.841419 | 686.871748 | 553.602693 | 834.748138 | 930.606071 |
| 466.423476 | 422.897465 | 544.649538 | 687.392698 | 553.602693 | 835.160563 | 930.606071 |
| 11188.6108 | 13625.55 | 17374.3203 | 2762.855 | 3432.55003 | 947.75258 | 871.328458 |
| 11188.6108 | 13625.55 | 17374.3203 | 2762.59453 | 3431.48336 | 947.340155 | 871.328458 |
| 427.554853 | 190.819588 | 142.970504 | 623.836874 | 211.201027 | 1067.76825 | 507.269488 |
| 427.554853 | 190.819588 | 142.970504 | 623.836874 | 211.201027 | 1067.76825 | 507.269488 |
| 438.660174 | 180.505016 | 156.586742 | 624.618298 | 210.134355 | 1067.76825 | 505.171165 |
| 438.660174 | 412.582893 | 510.608942 | 608.468868 | 476.802319 | 739.065545 | 883.393813 |
| 438.660174 | 412.582893 | 510.608942 | 608.468868 | 476.802319 | 739.47797 | 883.393813 |
| 438.660174 | 412.582893 | 510.608942 | 607.947918 | 476.802319 | 739.065545 | 883.393813 |
| 438.660174 | 412.582893 | 510.608942 | 608.208393 | 476.802319 | 739.47797 | 883.393813 |
| 438.660174 | 412.582893 | 510.608942 | 615.501684 | 480.002335 | 746.489195 | 891.262523 |
| 449.765494 | 412.582893 | 517.417061 | 608.729342 | 476.802319 | 739.47797 | 883.393813 |
| 1726.87739 | 2418.76721 | 1981.16269 | 507.144215 | 1062.40517 | 391.391296 | 635.267166 |
| 1688.00877 | 2413.60992 | 1967.54646 | 508.707063 | 1057.07181 | 390.566446 | 635.267166 |
| 1693.56143 | 2418.76721 | 1974.35458 | 510.009436 | 1061.3385 | 390.566446 | 637.365488 |
| 1693.56143 | 2413.60992 | 1974.35458 | 510.009436 | 1063.47184 | 390.566446 | 636.316327 |
| 11.1053208 | 10.3145723 | 6.80811923 | 15.6284812 | 18.1334215 | 19.3839736 | 18.3603227 |
| 26852.6658 | 33352.1696 | 32862.7915 | 5772.37953 | 10394.7172 | 5434.93625 | 6851.02328 |
| 427.554853 | 190.819588 | 122.546146 | 621.753077 | 216.534387 | 1010.8536 | 485.2371 |
| 26325.1631 | 32831.2837 | 32556.4261 | 5668.97108 | 10281.65 | 5373.89735 | 6748.73005 |
| 26319.6104 | 32831.2837 | 32549.618 | 5668.7106 | 10279.5167 | 5373.48493 | 6747.68089 |
| 26230.7678 | 32717.8234 | 32461.1125 | 5656.46829 | 10258.1832 | 5366.4737 | 6743.48424 |
| 288.738342 | 350.695459 | 408.487154 | 146.386774 | 197.334293 | 162.083013 | 140.063033 |
| 26214.1099 | 32722.9807 | 32461.1125 | 5654.90544 | 10260.3166 | 5363.99915 | 6739.2876 |
| 26219.6625 | 32707.5088 | 32440.6881 | 5650.2169 | 10254.9832 | 5361.11218 | 6738.23844 |
| 26219.6625 | 32707.5088 | 32440.6881 | 5649.95643 | 10254.9832 | 5360.69975 | 6737.71386 |
| 1610.27152 | 2356.87978 | 1967.54646 | 450.100258 | 1011.20492 | 331.589676 | 559.202972 |
| 1604.71886 | 2341.40792 | 1953.93022 | 443.067442 | 1002.67154 | 324.578451 | 553.432585 |
| 1599.1662 | 2320.77877 | 1933.50586 | 442.025543 | 1000.5382 | 322.103901 | 551.334262 |
| 16.6579813 | 10.3145723 | 27.2324769 | 5.99091779 | 22.400109 | 4.94909963 | 8.91787103 |
| 16.6579813 | 10.3145723 | 27.2324769 | 5.99091779 | 22.400109 | 4.94909963 | 8.91787103 |
| 5436.05456 | 5807.10422 | 7550.20422 | 1631.61344 | 1363.20663 | 647.094777 | 402.877938 |
| 8251.25339 | 6890.13431 | 8265.05674 | 1658.96328 | 1319.47308 | 539.039435 | 326.289164 |
| 8245.70073 | 6890.13431 | 8258.24862 | 1656.09806 | 1318.40641 | 538.214585 | 323.14168 |
| 16.6579813 | 10.3145723 | 27.2324769 | 5.99091779 | 22.400109 | 4.94909963 | 8.91787103 |
| 11.1053208 | 5.15728616 | 6.80811923 | 16.1494306 | 6.40003113 | 23.9206482 | 17.3111614 |
| 6224.53234 | 7106.74033 | 6338.359 | 2406.26515 | 3548.81726 | 2629.62161 | 3251.35086 |

| | | | | | | |
|---|---|---|---|---|---|---|
| 5.55266042 | 5.15728616 | 6.80811923 | 18.7541774 | 4.26668742 | 20.2088235 | 15.2128388 |
| 34359.8627 | 34914.8273 | 50897.4993 | 9370.0559 | 11197.9211 | 3677.18103 | 3209.90899 |
| 27.7633021 | 30.943717 | 13.6162385 | 75.2771844 | 29.8668119 | 126.614466 | 68.1954844 |
| 483.081457 | 257.864308 | 258.708531 | 870.506402 | 423.468726 | 1099.52497 | 552.908004 |
| 27.7633021 | 5.15728616 | 61.273073 | 3.38617092 | 6.40003113 | 4.53667467 | 4.72122584 |
| 5624.84501 | 7369.76192 | 7951.88326 | 2570.10373 | 2396.81166 | 2430.42035 | 2238.91021 |
| 5619.29235 | 7369.76192 | 7965.49949 | 2563.59186 | 2389.34495 | 2424.23397 | 2235.23815 |
| 1060.55814 | 1423.41098 | 1443.32128 | 802.262035 | 1216.00591 | 449.130792 | 516.711939 |
| 116905.713 | 95208.6598 | 116480.112 | 17602.3584 | 18877.9585 | 12080.7522 | 9689.52917 |
| 438.660174 | 458.998468 | 374.446557 | 1113.52928 | 675.203284 | 1156.02719 | 982.539556 |
| 16.6579813 | 15.4718585 | 6.80811923 | 273.237946 | 40.5335305 | 299.008103 | 119.079807 |
| 16.6579813 | 10.3145723 | 0 | 269.851775 | 39.4668586 | 295.296278 | 116.981485 |
| 449.765494 | 464.155754 | 367.638438 | 1121.86447 | 688.003346 | 1160.15144 | 990.408265 |
| 44.4212834 | 25.7864308 | 20.4243577 | 429.262283 | 82.1337328 | 493.260264 | 211.930582 |
| 160466.334 | 136389.59 | 153386.926 | 24621.6302 | 40209.2622 | 25029.2466 | 22156.7129 |
| 156535.05 | 132908.422 | 150459.435 | 20497.0136 | 37979.9181 | 23368.8236 | 20822.7043 |
| 156429.549 | 132836.22 | 150370.929 | 20484.5108 | 37947.9179 | 23362.2248 | 20815.3602 |
| 156412.892 | 132805.276 | 150364.121 | 20480.8641 | 37932.9845 | 23360.9876 | 20812.2127 |
| 156412.892 | 132805.276 | 150364.121 | 20480.8641 | 37932.9845 | 23360.9876 | 20812.2127 |
| 156490.629 | 132934.208 | 150398.162 | 20508.7349 | 37961.7846 | 23379.9591 | 20825.8518 |
| 11.1053208 | 15.4718585 | 13.6162385 | 29.4336396 | 10.6667185 | 28.8697479 | 19.409484 |
| 160183.148 | 136229.714 | 153380.118 | 21472.2308 | 39140.457 | 24109.5389 | 21388.2022 |
| 160172.043 | 136198.77 | 153414.159 | 21501.4039 | 39148.9904 | 24112.0134 | 21388.7268 |
| 160871.678 | 136575.252 | 153461.815 | 21527.9724 | 39285.5244 | 24184.1878 | 21408.6609 |
| 163159.374 | 139086.85 | 155715.303 | 21874.6642 | 40059.9282 | 24680.335 | 21652.5909 |
| 145724.02 | 124976.516 | 141969.71 | 18260.5779 | 34234.8332 | 21807.3827 | 19302.9941 |
| 11.1053208 | 15.4718585 | 20.4243577 | 4.16759498 | 11.7333904 | 4.94909963 | 6.29496779 |
| 89386.7275 | 101691.369 | 109937.509 | 14875.1884 | 38817.2555 | 15531.0995 | 17761.7762 |
| 89275.6743 | 101541.807 | 109767.306 | 14880.1374 | 38873.7891 | 15534.8113 | 17765.4483 |
| 89325.6483 | 101665.582 | 109828.579 | 14889.5145 | 38891.9225 | 15552.1332 | 17765.9728 |
| 89347.8589 | 101665.582 | 109814.963 | 14880.6584 | 38903.6559 | 15555.0202 | 17766.4974 |
| 89342.3062 | 101665.582 | 109814.963 | 14880.6584 | 38903.6559 | 15555.0202 | 17766.4974 |
| 89042.4626 | 101325.201 | 109624.336 | 14794.1808 | 38675.3881 | 15485.7328 | 17701.4494 |
| 89042.4626 | 101325.201 | 109624.336 | 14794.1808 | 38675.3881 | 15485.7328 | 17701.4494 |
| 89042.4626 | 101325.201 | 109624.336 | 14794.1808 | 38675.3881 | 15485.7328 | 17701.4494 |
| 89375.6222 | 101742.941 | 109889.852 | 14924.1576 | 38953.7895 | 15572.7544 | 17788.5298 |
| 83.2899064 | 134.08944 | 102.121788 | 193.272217 | 117.333904 | 206.62491 | 215.602647 |
| 55.5266042 | 41.2582893 | 34.0405961 | 469.375385 | 102.400498 | 530.378511 | 236.061292 |
| 55.5266042 | 41.2582893 | 34.0405961 | 469.375385 | 102.400498 | 530.378511 | 236.061292 |
| 55.5266042 | 41.2582893 | 27.2324769 | 470.417284 | 104.533842 | 530.790936 | 235.536711 |
| 55.5266042 | 51.5728616 | 47.6568346 | 474.063929 | 108.800529 | 536.15246 | 239.733357 |
| 55.5266042 | 41.2582893 | 34.0405961 | 469.896335 | 103.46717 | 531.203361 | 236.061292 |
| 77.7372459 | 41.2582893 | 34.0405961 | 473.54298 | 109.867201 | 536.15246 | 237.635034 |
| 94.3952272 | 61.8874339 | 68.0811923 | 478.491999 | 124.800607 | 536.564885 | 240.782518 |
| 55.5266042 | 56.7301478 | 54.4649538 | 471.719657 | 106.667185 | 534.915186 | 239.733357 |
| 116.605869 | 72.2020062 | 81.6974307 | 485.524816 | 118.400576 | 539.864285 | 247.602066 |

| | | | | | | |
|---|---|---|---|---|---|---|
| 11.1053208 | 25.7864308 | 13.6162385 | 30.736013 | 10.6667185 | 29.6945978 | 19.9340647 |
| 1965.64179 | 1567.81499 | 1851.80843 | 626.96257 | 897.07103 | 793.093216 | 885.492135 |
| 16469.1908 | 12681.7667 | 15059.5597 | 11682.8106 | 8850.17638 | 5854.37244 | 5789.27204 |
| 388.68623 | 474.470327 | 340.405961 | 314.132472 | 416.002023 | 324.166026 | 263.864066 |
| 388.68623 | 474.470327 | 340.405961 | 314.132472 | 416.002023 | 324.166026 | 263.864066 |
| 11.1053208 | 15.4718585 | 20.4243577 | 47.4063929 | 20.2667652 | 51.9655462 | 39.3435487 |
| 5.55266042 | 5.15728616 | 6.80811923 | 8.33518997 | 4.26668742 | 10.3106242 | 6.81954844 |
| 5.55266042 | 5.15728616 | 6.80811923 | 8.33518997 | 4.26668742 | 10.3106242 | 6.81954844 |
| 5.55266042 | 5.15728616 | 6.80811923 | 8.33518997 | 4.26668742 | 10.3106242 | 6.81954844 |
| 5.55266042 | 5.15728616 | 6.80811923 | 8.33518997 | 4.26668742 | 10.3106242 | 6.81954844 |
| 116.605869 | 113.460296 | 88.5055499 | 33.8617092 | 64.0003113 | 27.632473 | 39.3435487 |
| 1021.68952 | 1670.96072 | 1511.40247 | 1101.54745 | 2704.01315 | 803.816266 | 1223.84665 |
| 44.4212834 | 41.2582893 | 20.4243577 | 90.9056656 | 25.6001245 | 188.890636 | 82.3591619 |
| 99.9478876 | 77.3592924 | 81.6974307 | 53.9182601 | 40.5335305 | 87.8465185 | 50.8843229 |
| 11.1053208 | 5.15728616 | 13.6162385 | 11.4608862 | 5.33335927 | 10.7230492 | 14.6882582 |
| 27.7633021 | 5.15728616 | 20.4243577 | 20.3170255 | 8.53337484 | 28.8697479 | 19.9340647 |
| 4886.34117 | 7215.04334 | 6590.25941 | 6175.59434 | 8736.04249 | 4789.9036 | 3948.51854 |
| 4886.34117 | 7215.04334 | 6590.25941 | 6175.59434 | 8736.04249 | 4789.9036 | 3948.51854 |
| 94.3952272 | 139.246726 | 108.929908 | 206.556426 | 133.333982 | 214.460984 | 231.340066 |
| 5.55266042 | 5.15728616 | 6.80811923 | 0.52094937 | 2.13334371 | 0.41242497 | 1.0491613 |
| 11.1053208 | 5.15728616 | 6.80811923 | 10.1585128 | 6.40003113 | 16.4969988 | 13.6390969 |
| 105.500548 | 154.718585 | 122.546146 | 253.702345 | 132.26731 | 316.742377 | 344.124906 |
| 5463.81786 | 4285.7048 | 5426.07102 | 3830.54074 | 3720.55143 | 2237.40546 | 2512.21673 |
| 5441.60722 | 4285.7048 | 5412.45478 | 3823.24745 | 3720.55143 | 2239.05516 | 2508.54466 |
| 5430.5019 | 4280.54751 | 5412.45478 | 3824.54982 | 3719.48476 | 2239.88001 | 2510.64299 |
| 5441.60722 | 4259.91837 | 5412.45478 | 3830.80121 | 3722.68477 | 2241.52971 | 2513.79047 |
| 5430.5019 | 4280.54751 | 5412.45478 | 3823.50792 | 3719.48476 | 2238.64273 | 2508.54466 |
| 5441.60722 | 4280.54751 | 5398.83855 | 3828.45694 | 3719.48476 | 2238.23031 | 2511.16757 |
| 5447.15988 | 4265.07565 | 5392.03043 | 3822.98697 | 3712.01806 | 2237.40546 | 2508.02008 |
| 5424.94924 | 4265.07565 | 5392.03043 | 3822.20555 | 3710.95138 | 2236.58061 | 2509.59382 |
| 5424.94924 | 4265.07565 | 5392.03043 | 3822.20555 | 3710.95138 | 2236.99303 | 2508.54466 |
| 5424.94924 | 4265.07565 | 5392.03043 | 3821.94508 | 3712.01806 | 2236.99303 | 2508.02008 |
| 5424.94924 | 4265.07565 | 5392.03043 | 3822.46602 | 3712.01806 | 2236.99303 | 2508.02008 |
| 5424.94924 | 4265.07565 | 5392.03043 | 3822.46602 | 3712.01806 | 2236.99303 | 2508.02008 |
| 5424.94924 | 4265.07565 | 5392.03043 | 3822.20555 | 3712.01806 | 2237.81788 | 2510.11841 |
| 1066.1108 | 1727.69086 | 1545.44306 | 844.979883 | 1899.74257 | 749.788595 | 1509.21853 |
| 16.6579813 | 25.7864308 | 34.0405961 | 90.3847162 | 38.4001868 | 74.6489195 | 75.5396135 |
| 44.4212834 | 97.988437 | 68.0811923 | 119.818356 | 70.4003424 | 132.80084 | 136.390969 |
| 1371.50712 | 1511.08484 | 1531.82683 | 1648.28382 | 1415.47355 | 2010.1593 | 1534.92298 |
| 44.4212834 | 97.988437 | 68.0811923 | 120.07883 | 68.2669987 | 133.213265 | 135.866388 |
| 127.71119 | 159.875871 | 108.929908 | 177.904211 | 109.867201 | 272.612905 | 198.291485 |
| 588.582005 | 747.806493 | 680.811923 | 728.808173 | 980.271435 | 793.918066 | 600.644843 |
| 11.1053208 | 15.4718585 | 27.2324769 | 3.64664561 | 9.60004669 | 1.23727491 | 1.57374195 |
| 44.4212834 | 97.988437 | 68.0811923 | 117.213609 | 68.2669987 | 130.32629 | 134.817227 |
| 1043.90016 | 1614.23057 | 1518.21059 | 735.059565 | 1684.27486 | 675.964525 | 1484.03866 |
| 1843.48326 | 2135.11647 | 1531.82683 | 2882.67336 | 3019.74802 | 4354.38283 | 2348.02298 |

| | | | | | | |
|---|---|---|---|---|---|---|
| 1843.48326 | 2135.11647 | 1531.82683 | 2882.67336 | 3019.74802 | 4354.38283 | 2348.02298 |
| 1843.48326 | 2135.11647 | 1531.82683 | 2882.67336 | 3019.74802 | 4354.38283 | 2348.02298 |
| 1843.48326 | 2135.11647 | 1531.82683 | 2882.93383 | 3019.74802 | 4354.38283 | 2348.02298 |
| 1849.03592 | 2140.27376 | 1531.82683 | 2882.93383 | 3019.74802 | 4355.20768 | 2348.02298 |
| 1854.58858 | 2140.27376 | 1531.82683 | 2882.93383 | 3020.81469 | 4355.20768 | 2348.02298 |
| 1854.58858 | 2140.27376 | 1531.82683 | 2883.19431 | 3019.74802 | 4355.20768 | 2348.02298 |
| 1854.58858 | 2140.27376 | 1531.82683 | 2883.19431 | 3019.74802 | 4355.20768 | 2348.54757 |
| 55.5266042 | 97.988437 | 68.0811923 | 120.59978 | 70.4003424 | 133.62569 | 136.915549 |
| 44.4212834 | 97.988437 | 68.0811923 | 117.995033 | 68.2669987 | 130.738715 | 135.341807 |
| 1021.68952 | 1603.916 | 1497.78623 | 725.942951 | 1678.9415 | 675.139675 | 1475.12078 |
| 1010.5842 | 1588.44414 | 1497.78623 | 725.422002 | 1676.80816 | 673.9024 | 1472.49788 |
| 5.55266042 | 20.6291446 | 61.273073 | 23.9636712 | 37.3335149 | 33.8188475 | 36.1960648 |
| 1404.82309 | 1500.77027 | 1552.25118 | 1661.04708 | 1420.80691 | 2011.39658 | 1527.57885 |
| 44.4212834 | 25.7864308 | 34.0405961 | 37.2478802 | 21.3334371 | 96.5074429 | 37.2452261 |
| 0 | 15.4718585 | 34.0405961 | 17.1913293 | 29.8668119 | 25.1579231 | 25.7044518 |
| 1393.71777 | 1531.71399 | 1559.0593 | 1652.97236 | 1420.80691 | 2008.09718 | 1527.57885 |
| 477.528797 | 149.561299 | 163.394861 | 852.0127 | 166.400809 | 1037.2488 | 527.728133 |
| 2126.66894 | 1665.80343 | 1429.70504 | 450.360733 | 364.801774 | 366.645798 | 268.585292 |
| 2126.66894 | 1665.80343 | 1429.70504 | 450.360733 | 364.801774 | 366.645798 | 268.585292 |
| 1671.35079 | 3764.8189 | 2750.48017 | 2558.64285 | 3020.81469 | 569.971308 | 481.040455 |
| 427.554853 | 247.549736 | 251.900411 | 304.494909 | 115.20056 | 473.05144 | 220.323873 |
| 427.554853 | 247.549736 | 251.900411 | 304.494909 | 115.20056 | 472.639015 | 220.323873 |
| 88.8425668 | 82.5165786 | 74.8893115 | 121.641679 | 44.8002179 | 233.020108 | 99.6703233 |
| 510.844759 | 706.548204 | 639.963207 | 721.775356 | 954.67131 | 785.669567 | 592.251553 |
| 516.39742 | 722.020062 | 639.963207 | 732.715293 | 968.538044 | 790.206242 | 594.874456 |
| 94.3952272 | 144.404012 | 115.738027 | 201.607407 | 96.0004669 | 261.889856 | 295.338905 |
| 344.264946 | 134.08944 | 115.738027 | 844.198459 | 202.667652 | 640.083553 | 339.928261 |
| 527.50274 | 706.548204 | 626.346969 | 729.329122 | 971.73806 | 785.257142 | 595.399037 |
| 505.292099 | 706.548204 | 626.346969 | 720.733458 | 954.67131 | 784.019867 | 591.726972 |
| 122.158529 | 237.235163 | 136.162385 | 84.3937984 | 178.1342 | 159.196038 | 116.981485 |
| 299.843663 | 366.167317 | 367.638438 | 249.274275 | 452.268866 | 203.737935 | 403.9271 |
| 505.292099 | 711.70549 | 626.346969 | 724.901053 | 971.73806 | 788.144117 | 594.349875 |
| 505.292099 | 706.548204 | 626.346969 | 731.933869 | 982.404778 | 789.793817 | 600.644843 |
| 1454.79703 | 1562.65771 | 1593.0999 | 1668.60084 | 1431.47363 | 2020.0575 | 1534.3984 |
| 0 | 5.15728616 | 0 | 2.60474687 | 5.33335927 | 2.47454982 | 5.24580649 |
| 1454.79703 | 1562.65771 | 1599.90802 | 1669.38227 | 1431.47363 | 2021.29478 | 1535.44756 |
| 322.054305 | 546.672333 | 449.335869 | 475.887252 | 873.604249 | 388.916746 | 448.516455 |
| 11.1053208 | 10.3145723 | 6.80811923 | 40.6340511 | 23.4667808 | 76.7110443 | 39.8681293 |
| 0 | 0 | 0 | 0.52094937 | 0 | 1.23727491 | 1.0491613 |
| 660.766591 | 226.920591 | 122.546146 | 620.190229 | 144.0007 | 923.831932 | 452.7131 |
| 521.95008 | 902.525078 | 1300.35077 | 472.761556 | 988.804809 | 332.414525 | 860.312264 |
| 516.39742 | 923.154223 | 1293.54265 | 475.626778 | 993.071497 | 333.6518 | 859.787684 |
| 44.4212834 | 77.3592924 | 40.8487154 | 43.4992727 | 38.4001868 | 72.9992196 | 46.6876778 |
| 510.844759 | 902.525078 | 1279.92641 | 474.063929 | 984.538122 | 332.0021 | 851.918974 |
| 0 | 5.15728616 | 0 | 7.03281654 | 2.13334371 | 25.5703481 | 5.24580649 |
| 266.5277 | 87.6738647 | 61.273073 | 278.968389 | 64.0003113 | 450.368067 | 188.849034 |

| | | | | | | |
|---|---|---|---|---|---|---|
| 516.39742 | 912.83965 | 1279.92641 | 475.105828 | 990.938153 | 333.6518 | 860.312264 |
| 77.7372459 | 46.4155754 | 68.0811923 | 99.2408556 | 26.6667964 | 211.161584 | 88.6541297 |
| 494.186778 | 263.021594 | 381.254677 | 611.073615 | 457.602226 | 612.038655 | 444.31981 |
| 727.398516 | 263.021594 | 170.202981 | 670.461843 | 160.000778 | 968.373829 | 478.417552 |
| 116.605869 | 87.6738647 | 285.941008 | 137.530634 | 76.8003736 | 89.9086434 | 50.3597423 |
| 83.2899064 | 41.2582893 | 74.8893115 | 97.6780075 | 28.8001401 | 207.037335 | 81.3100006 |
| 483.081457 | 892.210506 | 1252.69394 | 468.593961 | 976.004747 | 326.640576 | 851.918974 |
| 5.55266042 | 0 | 20.4243577 | 12.7632596 | 2.13334371 | 19.3839736 | 11.5407743 |
| 21083.4516 | 18494.0282 | 17639.8369 | 6741.86631 | 11114.7207 | 5041.89525 | 6450.24366 |
| 21083.4516 | 18494.0282 | 17639.8369 | 6741.86631 | 11114.7207 | 5041.89525 | 6450.24366 |
| 21083.4516 | 18494.0282 | 17639.8369 | 6741.86631 | 11114.7207 | 5041.89525 | 6450.24366 |
| 3270.51699 | 5146.97159 | 4044.02282 | 3286.14865 | 6920.56699 | 2580.95546 | 2544.74073 |
| 5.55266042 | 5.15728616 | 0 | 6.25139248 | 1.06667185 | 14.8472989 | 5.77038714 |
| 1415.92841 | 1567.81499 | 1824.57595 | 1663.13087 | 1422.94025 | 2009.74688 | 1541.74253 |
| 283.185682 | 345.538173 | 326.789723 | 246.409053 | 446.935507 | 202.088235 | 402.877938 |
| 33.3159625 | 10.3145723 | 13.6162385 | 48.9692411 | 12.8000623 | 75.4737694 | 51.9334842 |
| 438.660174 | 814.851213 | 476.568346 | 388.367758 | 711.470127 | 484.186914 | 280.126067 |
| 3192.77974 | 5105.7133 | 3962.32539 | 3230.14659 | 6899.23356 | 2564.45846 | 2514.31505 |
| 3192.77974 | 5095.39873 | 3962.32539 | 3230.40706 | 6899.23356 | 2564.04604 | 2514.31505 |
| 33.3159625 | 20.6291446 | 20.4243577 | 47.1459183 | 10.6667185 | 85.7843937 | 33.5731615 |
| 33.3159625 | 20.6291446 | 20.4243577 | 47.4063929 | 11.7333904 | 86.6092436 | 33.5731615 |
| 105.500548 | 103.145723 | 74.8893115 | 92.208039 | 142.934029 | 108.467767 | 150.554646 |
| 33.3159625 | 20.6291446 | 20.4243577 | 46.6249689 | 10.6667185 | 84.5471188 | 34.0977422 |
| 38.868623 | 51.5728616 | 13.6162385 | 52.0949373 | 23.4667808 | 89.9086434 | 46.1630971 |
| 149.921831 | 72.2020062 | 68.0811923 | 159.670983 | 52.2669209 | 257.765606 | 120.128969 |
| 44.4212834 | 41.2582893 | 20.4243577 | 48.9692411 | 14.933406 | 89.4962184 | 36.7206454 |
| 27.7633021 | 10.3145723 | 6.80811923 | 36.2059814 | 5.33335927 | 65.5755702 | 24.6552905 |
| 33.3159625 | 25.7864308 | 20.4243577 | 47.4063929 | 10.6667185 | 87.4340935 | 34.0977422 |
| 33.3159625 | 46.4155754 | 68.0811923 | 25.0055699 | 34.1334994 | 22.6833733 | 14.1636775 |
| 61.0792647 | 82.5165786 | 115.738027 | 94.8127859 | 60.8002957 | 101.044118 | 95.4736781 |
| 49.9739438 | 72.2020062 | 47.6568346 | 184.155603 | 120.53392 | 392.216146 | 192.521098 |
| 0 | 5.15728616 | 0 | 0.78142406 | 3.20001556 | 0.82484994 | 1.0491613 |
| 0 | 5.15728616 | 0 | 0.78142406 | 3.20001556 | 0.82484994 | 1.0491613 |
| 38.868623 | 61.8874339 | 34.0405961 | 42.9783233 | 37.3335149 | 72.9992196 | 44.5893552 |
| 61.0792647 | 82.5165786 | 88.5055499 | 92.4685137 | 58.666952 | 99.8068426 | 94.4245168 |
| 1177.16401 | 1933.98231 | 1375.24008 | 1726.6867 | 1330.1398 | 997.656001 | 709.233037 |
| 38.868623 | 56.7301478 | 34.0405961 | 42.7178486 | 36.2668431 | 72.5867946 | 44.5893552 |
| 149.921831 | 180.505016 | 149.778623 | 85.6961719 | 120.53392 | 96.0950179 | 122.227291 |
| 61.0792647 | 72.2020062 | 95.3136692 | 90.6451909 | 54.4002646 | 99.8068426 | 92.8507749 |
| 549.713382 | 964.412512 | 1382.0482 | 688.695071 | 1065.60518 | 582.344057 | 956.835104 |
| 61.0792647 | 82.5165786 | 81.6974307 | 91.9475643 | 55.4669365 | 99.8068426 | 94.9490975 |
| 77.7372459 | 25.7864308 | 13.6162385 | 72.1514882 | 16.0000778 | 140.22449 | 41.9664519 |
| 0 | 5.15728616 | 6.80811923 | 0.52094937 | 0 | 0 | 0.52458065 |
| 294.291003 | 335.2236 | 285.941008 | 276.884592 | 464.002257 | 243.330732 | 396.582971 |
| 61.0792647 | 72.2020062 | 88.5055499 | 91.9475643 | 56.5336083 | 100.219268 | 92.8507749 |
| 0 | 0 | 0 | 0.26047469 | 0 | 0 | 0 |

| | | | | | | |
|---|---|---|---|---|---|---|
| 61.0792647 | 72.2020062 | 81.6974307 | 89.6032922 | 55.4669365 | 97.7447178 | 89.1787103 |
| 6257.8483 | 14899.3997 | 8571.42211 | 8328.6781 | 15181.9405 | 7848.03475 | 6394.63811 |
| 6241.19032 | 14889.0851 | 8564.61399 | 8326.85478 | 15179.8072 | 7847.2099 | 6390.44147 |
| 16.6579813 | 25.7864308 | 27.2324769 | 37.5083549 | 41.6002023 | 42.0673469 | 35.1469035 |
| 0 | 5.15728616 | 0 | 0.78142406 | 3.20001556 | 0.82484994 | 1.57374195 |
| 7301.74846 | 16637.4052 | 9926.23783 | 9415.37849 | 18123.8215 | 8334.69621 | 6952.26734 |
| 7296.1958 | 16637.4052 | 9926.23783 | 9414.3366 | 18123.8215 | 8333.87136 | 6953.3165 |
| 61.0792647 | 72.2020062 | 81.6974307 | 90.9056656 | 55.4669365 | 99.8068426 | 92.8507749 |
| 7290.64314 | 16637.4052 | 9926.23783 | 9417.20182 | 18127.0215 | 8335.52106 | 6956.98857 |
| 61.0792647 | 72.2020062 | 88.5055499 | 91.6870897 | 55.4669365 | 100.219268 | 92.8507749 |
| 61.0792647 | 72.2020062 | 81.6974307 | 90.6451909 | 54.4002646 | 101.044118 | 92.3261942 |
| 7346.16974 | 16828.2247 | 10007.9353 | 9557.59767 | 18448.0897 | 8437.39003 | 7028.33153 |
| 444.212834 | 417.740179 | 217.859815 | 519.647 | 571.736114 | 861.143336 | 529.826455 |
| 7318.40644 | 16740.5509 | 9973.89467 | 9536.23875 | 18365.956 | 8416.35635 | 7018.3645 |
| 22.2106417 | 10.3145723 | 6.80811923 | 56.2625323 | 11.7333904 | 75.4737694 | 51.9334842 |
| 372.028248 | 376.48189 | 265.51665 | 253.44187 | 152.534075 | 271.788055 | 245.503744 |
| 322.054305 | 340.380887 | 231.476054 | 332.626175 | 518.402521 | 329.115126 | 270.683615 |
| 260.97504 | 330.066314 | 190.627338 | 305.536807 | 391.468571 | 313.855402 | 250.22497 |
| 22.2106417 | 51.5728616 | 54.4649538 | 40.3735764 | 32.0001556 | 45.7791716 | 55.6055488 |
| 22.2106417 | 46.4155754 | 54.4649538 | 39.071203 | 28.8001401 | 44.5418967 | 55.0809681 |
| 11.1053208 | 5.15728616 | 6.80811923 | 5.99091779 | 2.13334371 | 7.83607442 | 7.86870973 |
| 11.1053208 | 5.15728616 | 6.80811923 | 5.7304431 | 2.13334371 | 7.83607442 | 7.86870973 |
| 16.6579813 | 36.1010031 | 20.4243577 | 71.1095894 | 19.2000934 | 102.281392 | 54.5563875 |
| 571.924024 | 422.897465 | 374.446557 | 355.026998 | 592.002879 | 311.793277 | 425.434906 |
| 10294.6324 | 13537.8762 | 11948.2492 | 5445.22332 | 9044.31066 | 3308.47311 | 5771.96088 |
| 10066.9734 | 13362.5284 | 11778.0463 | 5303.52509 | 8926.97675 | 3153.81374 | 5603.57049 |
| 5.55266042 | 0 | 0 | 0.78142406 | 0 | 2.06212485 | 1.0491613 |
| 0 | 0 | 0 | 2.34427218 | 0 | 3.71182473 | 1.57374195 |
| 1926.77317 | 3563.68474 | 2532.62035 | 3388.77567 | 4050.15303 | 2296.79466 | 1783.04963 |
| 10250.2111 | 13450.2023 | 11880.1681 | 5409.79876 | 9014.44385 | 3296.92521 | 5752.5514 |
| 10250.2111 | 13450.2023 | 11880.1681 | 5410.05924 | 9014.44385 | 3296.51278 | 5752.5514 |
| 10239.1058 | 13450.2023 | 11873.3599 | 5411.36161 | 9019.7772 | 3298.57491 | 5754.12514 |
| 10239.1058 | 13465.6742 | 11873.3599 | 5413.44541 | 9028.31058 | 3300.22461 | 5763.04301 |
| 10239.1058 | 13465.6742 | 11873.3599 | 5413.44541 | 9028.31058 | 3300.22461 | 5763.04301 |
| 10239.1058 | 13465.6742 | 11873.3599 | 5415.78968 | 9029.37725 | 3300.63703 | 5765.14133 |
| 27.7633021 | 15.4718585 | 34.0405961 | 31.5174371 | 28.8001401 | 40.830072 | 45.6385165 |
| 0 | 5.15728616 | 0 | 1.30237343 | 3.20001556 | 0.82484994 | 1.0491613 |
| 460.870815 | 448.683896 | 211.051696 | 628.264944 | 675.203284 | 1101.58709 | 615.333101 |
| 460.870815 | 448.683896 | 211.051696 | 628.264944 | 675.203284 | 1101.17467 | 615.333101 |
| 460.870815 | 448.683896 | 211.051696 | 628.525419 | 675.203284 | 1102.41194 | 615.857682 |
| 5.55266042 | 0 | 6.80811923 | 7.8142406 | 4.26668742 | 9.89819927 | 7.34412909 |
| 122.158529 | 46.4155754 | 81.6974307 | 223.226806 | 133.333982 | 285.398079 | 130.096001 |
| 11.1053208 | 5.15728616 | 0 | 23.4427218 | 5.33335927 | 21.4460984 | 8.39329038 |
| 11.1053208 | 5.15728616 | 6.80811923 | 6.51186716 | 2.13334371 | 8.24849939 | 8.39329038 |
| 405.344211 | 355.852745 | 129.354265 | 402.172916 | 540.80263 | 605.02743 | 387.140519 |
| 416.449532 | 371.324604 | 170.202981 | 409.726682 | 542.935974 | 626.885954 | 407.074584 |

| | | | | | | |
|---|---|---|---|---|---|---|
| 27.7633021 | 10.3145723 | 13.6162385 | 68.7653172 | 16.0000778 | 84.9595437 | 57.1792907 |
| 22.2106417 | 25.7864308 | 34.0405961 | 17.451804 | 16.0000778 | 23.0957983 | 21.5078066 |
| 977.268235 | 562.144191 | 565.073896 | 1152.60049 | 240.001167 | 1604.74556 | 583.858262 |
| 1027.24218 | 567.301478 | 319.981604 | 846.803206 | 243.201183 | 1070.2428 | 528.252714 |
| 11.1053208 | 5.15728616 | 0 | 4.94901904 | 1.06667185 | 6.59879951 | 6.29496779 |
| 977.268235 | 582.773336 | 571.882015 | 1160.93568 | 245.334527 | 1610.51951 | 589.628649 |
| 1032.79484 | 572.458764 | 319.981604 | 847.063681 | 245.334527 | 1071.06765 | 528.777294 |
| 83.2899064 | 154.718585 | 129.354265 | 130.237343 | 248.534542 | 103.931092 | 261.241163 |
| 61.0792647 | 15.4718585 | 20.4243577 | 67.4629438 | 21.3334371 | 94.445318 | 60.8513553 |
| 0 | 5.15728616 | 0 | 1.04189875 | 0 | 4.53667467 | 2.62290324 |
| 11566.1917 | 9948.405 | 9517.75068 | 3851.89966 | 5325.89257 | 2895.63571 | 4073.89332 |
| 27.7633021 | 36.1010031 | 27.2324769 | 15.6284812 | 32.0001556 | 9.89819927 | 20.983226 |
| 49.9739438 | 20.6291446 | 20.4243577 | 63.2953488 | 21.3334371 | 87.4340935 | 61.3759359 |
| 477.528797 | 469.313041 | 258.708531 | 661.605704 | 686.936675 | 1124.27047 | 622.67723 |
| 477.528797 | 469.313041 | 265.51665 | 661.866178 | 689.070018 | 1125.09532 | 623.726392 |
| 72.1845855 | 61.8874339 | 74.8893115 | 46.3644942 | 61.8669676 | 51.9655462 | 72.9167102 |
| 483.081457 | 469.313041 | 258.708531 | 662.387128 | 688.003346 | 1125.09532 | 623.201811 |
| 477.528797 | 469.313041 | 265.51665 | 663.689501 | 686.936675 | 1124.68289 | 624.775553 |
| 27.7633021 | 36.1010031 | 27.2324769 | 15.6284812 | 33.0668275 | 10.3106242 | 20.983226 |
| 16.6579813 | 15.4718585 | 13.6162385 | 45.8435448 | 9.60004669 | 51.5531212 | 25.1798712 |
| 33.3159625 | 36.1010031 | 27.2324769 | 16.9308546 | 33.0668275 | 9.89819927 | 22.0323873 |
| 155.474492 | 82.5165786 | 156.586742 | 101.324653 | 60.8002957 | 163.320288 | 109.637356 |
| 344.264946 | 211.448733 | 245.092292 | 394.879625 | 280.534698 | 335.713925 | 258.093679 |
| 66.6319251 | 103.145723 | 61.273073 | 224.00823 | 132.26731 | 546.87551 | 235.536711 |
| 0 | 0 | 0 | 0 | 0 | 0.41242497 | 0 |
| 94.3952272 | 139.246726 | 81.6974307 | 250.837123 | 158.934106 | 575.332833 | 255.470776 |
| 1632.48216 | 881.895933 | 789.74183 | 520.949373 | 404.268633 | 477.588115 | 748.576586 |
| 11.1053208 | 10.3145723 | 0 | 2.86522155 | 1.06667185 | 3.71182473 | 4.72122584 |
| 16.6579813 | 5.15728616 | 6.80811923 | 2.60474687 | 4.26668742 | 4.53667467 | 3.67206454 |
| 438.660174 | 422.897465 | 272.324769 | 483.701493 | 560.002724 | 752.263144 | 502.023681 |
| 133.26385 | 92.8311509 | 108.929908 | 106.534147 | 101.333826 | 94.0328931 | 95.9982588 |
| 438.660174 | 428.054751 | 272.324769 | 483.701493 | 560.002724 | 752.675569 | 503.072842 |
| 116.605869 | 118.617582 | 61.273073 | 83.091425 | 89.6004358 | 83.3098439 | 109.637356 |
| 0 | 0 | 0 | 0.52094937 | 1.06667185 | 1.23727491 | 0 |
| 122.158529 | 134.08944 | 88.5055499 | 102.627026 | 158.934106 | 117.128691 | 162.620001 |
| 5.55266042 | 0 | 0 | 1.30237343 | 0 | 4.53667467 | 2.62290324 |
| 49.9739438 | 87.6738647 | 34.0405961 | 43.238798 | 40.5335305 | 63.5134453 | 63.4742585 |
| 2015.61573 | 1309.95068 | 1368.43196 | 924.945612 | 882.137624 | 864.855161 | 922.737362 |
| 0 | 0 | 0 | 0.52094937 | 0 | 0.82484994 | 0.52458065 |
| 94.3952272 | 128.932154 | 108.929908 | 163.057154 | 229.334449 | 137.74994 | 135.341807 |
| 94.3952272 | 123.774868 | 122.546146 | 165.401426 | 231.467793 | 139.812065 | 137.44013 |
| 172.132473 | 51.5728616 | 170.202981 | 95.0732606 | 42.6668742 | 142.699039 | 110.161936 |
| 0 | 0 | 0 | 0.26047469 | 0 | 0.41242497 | 0.52458065 |
| 105.500548 | 128.932154 | 115.738027 | 166.964274 | 230.401121 | 139.39964 | 136.390969 |
| 0 | 0 | 0 | 1.30237343 | 1.06667185 | 2.47454982 | 1.57374195 |
| 5.55266042 | 0 | 0 | 0.26047469 | 0 | 2.06212485 | 1.0491613 |

| | | | | | | |
|---|---|---|---|---|---|---|
| 122.158529 | 82.5165786 | 102.121788 | 105.492248 | 101.333826 | 93.6204681 | 93.3753555 |
| 99.9478876 | 139.246726 | 122.546146 | 170.350445 | 240.001167 | 143.936314 | 142.161356 |
| 33.3159625 | 20.6291446 | 34.0405961 | 70.8491147 | 25.6001245 | 137.74994 | 71.3429683 |
| 105.500548 | 144.404012 | 122.546146 | 170.871394 | 240.001167 | 143.936314 | 141.636775 |
| 111.053208 | 128.932154 | 156.586742 | 78.142406 | 107.733857 | 94.0328931 | 118.555227 |
| 105.500548 | 123.774868 | 115.738027 | 163.317628 | 230.401121 | 137.74994 | 135.341807 |
| 27.7633021 | 5.15728616 | 6.80811923 | 8.59566466 | 5.33335927 | 9.07334933 | 11.0161936 |
| 33.3159625 | 36.1010031 | 27.2324769 | 16.9308546 | 33.0668275 | 11.5478991 | 23.6061292 |
| 111.053208 | 87.6738647 | 47.6568346 | 68.2443679 | 67.2003269 | 72.9992196 | 88.129549 |
| 0 | 0 | 0 | 1.82332281 | 0 | 0.82484994 | 2.62290324 |
| 177.685134 | 190.819588 | 102.121788 | 156.545287 | 202.667652 | 136.512665 | 142.161356 |
| 16.6579813 | 25.7864308 | 27.2324769 | 15.1075318 | 11.7333904 | 21.0336734 | 17.3111614 |
| 16935.6143 | 24399.1208 | 18300.2245 | 14834.2939 | 27849.7355 | 12916.3252 | 10497.9079 |
| 0 | 0 | 0 | 0 | 0 | 0.41242497 | 0 |
| 16902.2983 | 24368.1771 | 18327.457 | 14849.1409 | 27838.0021 | 12920.4494 | 10496.3342 |
| 88.8425668 | 82.5165786 | 115.738027 | 269.591301 | 119.467248 | 176.517887 | 65.0480005 |
| 0 | 5.15728616 | 0 | 5.99091779 | 2.13334371 | 5.77394957 | 5.24580649 |
| 105.500548 | 144.404012 | 122.546146 | 190.406996 | 276.26801 | 155.484214 | 153.70213 |
| 66.6319251 | 97.988437 | 115.738027 | 70.5886401 | 123.733935 | 82.0725689 | 112.260259 |
| 105.500548 | 144.404012 | 122.546146 | 190.927945 | 274.134667 | 155.896638 | 153.70213 |
| 11.1053208 | 0 | 0 | 0 | 0 | 0 | 0 |
| 105.500548 | 46.4155754 | 61.273073 | 57.304431 | 50.1335772 | 69.6998199 | 86.0312264 |
| 38.868623 | 36.1010031 | 27.2324769 | 20.0565509 | 34.1334994 | 15.6721488 | 26.7536131 |
| 38.868623 | 36.1010031 | 27.2324769 | 20.0565509 | 34.1334994 | 15.6721488 | 26.7536131 |
| 38.868623 | 36.1010031 | 27.2324769 | 20.0565509 | 34.1334994 | 15.6721488 | 26.7536131 |
| 38.868623 | 36.1010031 | 27.2324769 | 20.0565509 | 34.1334994 | 15.6721488 | 26.7536131 |
| 38.868623 | 36.1010031 | 27.2324769 | 20.0565509 | 34.1334994 | 15.6721488 | 26.7536131 |
| 38.868623 | 36.1010031 | 27.2324769 | 20.0565509 | 34.1334994 | 15.6721488 | 26.7536131 |
| 5.55266042 | 5.15728616 | 34.0405961 | 8.07471528 | 5.33335927 | 5.3615246 | 7.34412909 |
| 16.6579813 | 25.7864308 | 27.2324769 | 14.8470571 | 11.7333904 | 21.0336734 | 17.3111614 |
| 0 | 0 | 0 | 1.56284812 | 0 | 0 | 0.52458065 |
| 111.053208 | 144.404012 | 122.546146 | 190.146521 | 274.134667 | 155.484214 | 152.652969 |
| 0 | 0 | 0 | 0.26047469 | 0 | 0.41242497 | 0 |
| 161.027152 | 87.6738647 | 156.586742 | 90.3847162 | 49.0669053 | 137.337515 | 80.2608393 |
| 44.4212834 | 30.943717 | 20.4243577 | 67.7234185 | 94.9337951 | 76.7110443 | 65.0480005 |
| 38.868623 | 36.1010031 | 27.2324769 | 20.0565509 | 34.1334994 | 15.6721488 | 26.7536131 |
| 38.868623 | 36.1010031 | 27.2324769 | 20.0565509 | 34.1334994 | 15.6721488 | 26.7536131 |
| 38.868623 | 36.1010031 | 27.2324769 | 20.0565509 | 34.1334994 | 15.6721488 | 26.7536131 |
| 38.868623 | 36.1010031 | 27.2324769 | 20.0565509 | 34.1334994 | 15.6721488 | 26.7536131 |
| 38.868623 | 36.1010031 | 27.2324769 | 20.0565509 | 34.1334994 | 15.6721488 | 26.7536131 |
| 38.868623 | 36.1010031 | 27.2324769 | 20.0565509 | 34.1334994 | 15.6721488 | 26.7536131 |
| 38.868623 | 36.1010031 | 27.2324769 | 20.0565509 | 34.1334994 | 15.6721488 | 26.7536131 |
| 38.868623 | 36.1010031 | 27.2324769 | 20.0565509 | 34.1334994 | 15.6721488 | 26.7536131 |
| 38.868623 | 36.1010031 | 27.2324769 | 20.0565509 | 34.1334994 | 15.6721488 | 26.7536131 |
| 38.868623 | 36.1010031 | 27.2324769 | 20.0565509 | 34.1334994 | 15.6721488 | 26.7536131 |
| 38.868623 | 36.1010031 | 27.2324769 | 20.0565509 | 34.1334994 | 15.6721488 | 26.7536131 |

| | | | | | | |
|---|---|---|---|---|---|---|
| 38.868623 | 36.1010031 | 27.2324769 | 20.0565509 | 34.1334994 | 15.6721488 | 26.7536131 |
| 38.868623 | 36.1010031 | 27.2324769 | 20.0565509 | 34.1334994 | 15.6721488 | 26.7536131 |
| 94.3952272 | 144.404012 | 129.354265 | 174.778515 | 249.601214 | 144.348739 | 140.587614 |
| 22.2106417 | 10.3145723 | 13.6162385 | 26.568418 | 9.60004669 | 51.1406962 | 36.1960648 |
| 27.7633021 | 10.3145723 | 13.6162385 | 39.852627 | 50.1335772 | 23.5082233 | 40.9172906 |
| 0 | 0 | 0 | 0.26047469 | 0 | 0.82484994 | 1.57374195 |
| 38.868623 | 36.1010031 | 27.2324769 | 20.0565509 | 34.1334994 | 15.6721488 | 26.7536131 |
| 38.868623 | 36.1010031 | 27.2324769 | 20.0565509 | 34.1334994 | 15.6721488 | 26.7536131 |
| 38.868623 | 36.1010031 | 27.2324769 | 20.0565509 | 34.1334994 | 15.6721488 | 26.7536131 |
| 38.868623 | 36.1010031 | 27.2324769 | 20.0565509 | 34.1334994 | 15.6721488 | 26.7536131 |
| 38.868623 | 36.1010031 | 27.2324769 | 20.0565509 | 34.1334994 | 15.6721488 | 26.7536131 |
| 38.868623 | 36.1010031 | 27.2324769 | 20.0565509 | 34.1334994 | 15.6721488 | 26.7536131 |
| 38.868623 | 36.1010031 | 27.2324769 | 20.0565509 | 34.1334994 | 15.6721488 | 26.7536131 |
| 38.868623 | 36.1010031 | 27.2324769 | 20.0565509 | 34.1334994 | 15.6721488 | 26.7536131 |
| 38.868623 | 36.1010031 | 27.2324769 | 20.0565509 | 34.1334994 | 15.6721488 | 26.7536131 |
| 166.579813 | 180.505016 | 122.546146 | 109.659843 | 170.667497 | 121.252941 | 170.488711 |
| 66.6319251 | 25.7864308 | 27.2324769 | 39.3316777 | 57.6002802 | 40.005222 | 52.9826455 |
| 16.6579813 | 20.6291446 | 20.4243577 | 60.4301273 | 32.0001556 | 119.190816 | 76.0641941 |
| 427.554853 | 159.875871 | 95.3136692 | 94.2918365 | 21.3334371 | 237.969207 | 135.341807 |
| 422.002192 | 159.875871 | 95.3136692 | 92.208039 | 21.3334371 | 236.319508 | 134.817227 |
| 38.868623 | 30.943717 | 20.4243577 | 68.2443679 | 97.0671388 | 77.1234693 | 66.0971618 |
| 244.317059 | 211.448733 | 238.284173 | 382.116365 | 197.334293 | 538.62701 | 297.437228 |
| 116.605869 | 144.404012 | 129.354265 | 81.7890516 | 108.800529 | 99.8068426 | 120.128969 |
| 133.26385 | 87.6738647 | 81.6974307 | 204.733104 | 71.4670143 | 233.020108 | 108.063614 |
| 38.868623 | 36.1010031 | 34.0405961 | 20.3170255 | 35.2001712 | 16.0845738 | 27.2781937 |
| 38.868623 | 36.1010031 | 34.0405961 | 20.3170255 | 35.2001712 | 16.0845738 | 27.2781937 |
| 38.868623 | 36.1010031 | 34.0405961 | 20.3170255 | 35.2001712 | 16.0845738 | 27.2781937 |
| 0 | 10.3145723 | 0 | 1.56284812 | 0 | 2.47454982 | 1.57374195 |
| 11.1053208 | 0 | 0 | 2.34427218 | 3.20001556 | 7.01122448 | 4.72122584 |
| 549.713382 | 969.569798 | 803.358069 | 1231.00337 | 2562.1458 | 937.029531 | 1802.45911 |
| 0 | 0 | 0 | 0.26047469 | 0 | 0.41242497 | 0 |
| 16.6579813 | 25.7864308 | 6.80811923 | 9.11661403 | 4.26668742 | 9.07334933 | 9.44245168 |
| 144.369171 | 92.8311509 | 88.5055499 | 291.2107 | 66.133655 | 198.37641 | 59.2776133 |
| 116.605869 | 154.718585 | 142.970504 | 193.272217 | 281.60137 | 157.546338 | 155.275872 |
| 13109.8313 | 18726.106 | 14215.3529 | 11304.6014 | 24775.5872 | 9192.95257 | 8939.37884 |
| 13109.8313 | 18720.9488 | 14160.888 | 11295.7453 | 24773.4538 | 9190.0656 | 8935.70677 |
| 13104.2786 | 18767.3643 | 14215.3529 | 11324.9184 | 24908.9212 | 9213.98624 | 8959.3129 |
| 144.369171 | 92.8311509 | 88.5055499 | 305.536807 | 78.9337173 | 208.687035 | 68.1954844 |
| 5.55266042 | 0 | 0 | 0.52094937 | 0 | 0.82484994 | 0.52458065 |
| 144.369171 | 92.8311509 | 88.5055499 | 305.536807 | 77.8670454 | 209.09946 | 68.1954844 |
| 144.369171 | 92.8311509 | 88.5055499 | 305.536807 | 77.8670454 | 208.687035 | 68.1954844 |
| 12937.6988 | 18571.3875 | 13929.4119 | 11241.827 | 24707.3202 | 9169.03192 | 8898.98613 |
| 33.3159625 | 20.6291446 | 34.0405961 | 171.131869 | 60.8002957 | 121.665366 | 31.9994196 |
| 533.055401 | 959.255226 | 735.276876 | 1231.52432 | 2558.94578 | 938.266806 | 1809.27866 |
| 66.6319251 | 56.7301478 | 34.0405961 | 64.8581969 | 40.5335305 | 62.2761704 | 71.3429683 |
| 449.765494 | 933.468795 | 694.428161 | 1216.93774 | 2531.21231 | 924.656782 | 1791.44292 |

| | | | | | | |
|---|---|---|---|---|---|---|
| 22.2106417 | 25.7864308 | 27.2324769 | 15.8889559 | 11.7333904 | 20.6212485 | 17.8357421 |
| 416.449532 | 917.996937 | 714.852519 | 1212.77014 | 2510.94555 | 920.532532 | 1786.72169 |
| 105.500548 | 154.718585 | 163.394861 | 177.904211 | 253.867901 | 148.885414 | 145.83342 |
| 416.449532 | 887.05322 | 701.23628 | 1209.64444 | 2514.14556 | 920.532532 | 1786.72169 |
| 416.449532 | 887.05322 | 701.23628 | 1209.64444 | 2514.14556 | 920.532532 | 1786.72169 |
| 416.449532 | 887.05322 | 701.23628 | 1209.64444 | 2514.14556 | 920.532532 | 1786.72169 |
| 416.449532 | 892.210506 | 694.428161 | 1212.24919 | 2510.94555 | 920.944957 | 1786.19711 |
| 22.2106417 | 5.15728616 | 0 | 4.42806967 | 2.13334371 | 4.53667467 | 3.67206454 |
| 1188.26933 | 629.188912 | 374.446557 | 440.72317 | 370.135134 | 394.278271 | 587.530327 |
| 0 | 0 | 0 | 0.26047469 | 0 | 0.41242497 | 1.57374195 |
| 99.9478876 | 180.505016 | 136.162385 | 178.164686 | 254.934573 | 148.472989 | 143.735098 |
| 5.55266042 | 5.15728616 | 0 | 2.60474687 | 1.06667185 | 4.53667467 | 2.0983226 |
| 22.2106417 | 15.4718585 | 27.2324769 | 23.1822471 | 21.3334371 | 29.6945978 | 34.6223228 |
| 22.2106417 | 15.4718585 | 27.2324769 | 23.1822471 | 21.3334371 | 29.6945978 | 34.6223228 |
| 194.343115 | 226.920591 | 510.608942 | 255.004718 | 492.802397 | 298.595678 | 442.746068 |
| 5.55266042 | 0 | 13.6162385 | 2.60474687 | 1.06667185 | 2.88697479 | 2.62290324 |
| 44.4212834 | 87.6738647 | 34.0405961 | 108.878419 | 89.6004358 | 129.50144 | 107.014452 |
| 0 | 0 | 0 | 0.52094937 | 0 | 1.23727491 | 1.0491613 |
| 22.2106417 | 15.4718585 | 20.4243577 | 22.400823 | 20.2667652 | 29.2821728 | 34.0977422 |
| 33.3159625 | 36.1010031 | 27.2324769 | 16.1494306 | 11.7333904 | 21.8585234 | 17.8357421 |
| 0 | 0 | 0 | 0.26047469 | 0 | 0.41242497 | 0.52458065 |
| 66.6319251 | 134.08944 | 163.394861 | 413.373328 | 524.802553 | 174.455762 | 144.784259 |
| 11.1053208 | 15.4718585 | 20.4243577 | 56.0020576 | 18.1334215 | 68.05012 | 55.6055488 |
| 5.55266042 | 0 | 0 | 2.86522155 | 1.06667185 | 4.1242497 | 1.57374195 |
| 0 | 0 | 0 | 0.26047469 | 0 | 0.82484994 | 0.52458065 |
| 22.2106417 | 25.7864308 | 13.6162385 | 59.3882285 | 40.5335305 | 84.5471188 | 61.3759359 |
| 22.2106417 | 25.7864308 | 13.6162385 | 59.3882285 | 40.5335305 | 84.5471188 | 61.3759359 |
| 94.3952272 | 77.3592924 | 102.121788 | 88.5613934 | 48.0002335 | 160.433313 | 105.965291 |
| 11.1053208 | 15.4718585 | 20.4243577 | 56.523007 | 18.1334215 | 68.05012 | 55.6055488 |

ree_tRNA_dedup_counts

| DEX.7 | DEX.8 | DEX.9 | DEX.10 |
|---|---|---|---|
| 148.515266 | 155.931374 | 174.199257 | 137.137884 |
| 90.4266339 | 71.9100997 | 91.6159053 | 81.2668943 |
| 407.81813 | 501.099853 | 690.345202 | 758.067749 |
| 165.881971 | 171.070342 | 221.942757 | 153.645222 |
| 549.147174 | 787.226355 | 624.536594 | 742.830206 |
| 757.547628 | 775.872129 | 601.310027 | 731.402049 |
| 754.553369 | 773.601283 | 601.310027 | 727.592663 |
| 754.553369 | 772.844335 | 601.310027 | 727.592663 |
| 1407.30192 | 1340.55565 | 1876.19051 | 2172.61963 |
| 1407.30192 | 1340.55565 | 1876.19051 | 2172.61963 |
| 296.431681 | 751.649779 | 560.018351 | 728.862459 |
| 296.431681 | 751.649779 | 560.018351 | 728.862459 |
| 297.030532 | 753.920624 | 560.018351 | 727.592663 |
| 698.261292 | 718.344049 | 558.727986 | 700.926964 |
| 698.261292 | 718.344049 | 558.727986 | 700.926964 |
| 698.261292 | 719.100997 | 558.727986 | 700.926964 |
| 698.261292 | 718.344049 | 558.727986 | 700.926964 |
| 703.052107 | 727.42743 | 558.727986 | 703.466554 |
| 698.261292 | 720.614894 | 558.727986 | 700.926964 |
| 858.154744 | 719.857946 | 776.799648 | 1085.67492 |
| 856.358188 | 719.100997 | 779.380378 | 1085.67492 |
| 858.753596 | 722.885739 | 780.670743 | 1092.02389 |
| 858.753596 | 722.885739 | 779.380378 | 1092.02389 |
| 11.3781857 | 15.1389684 | 19.355473 | 10.1583618 |
| 7692.85125 | 8883.54663 | 10957.7784 | 13193.1724 |
| 265.890235 | 738.024708 | 526.468864 | 695.847783 |
| 7571.28432 | 8761.67794 | 10813.2576 | 12984.926 |
| 7570.08661 | 8759.40709 | 10811.9672 | 12984.926 |
| 7561.10384 | 8763.94878 | 10802.9346 | 12955.7207 |
| 154.503785 | 158.202219 | 192.264365 | 149.835836 |
| 7558.10958 | 8757.13625 | 10797.7732 | 12953.1811 |
| 7550.92335 | 8750.32371 | 10796.4828 | 12948.1019 |
| 7550.92335 | 8750.32371 | 10796.4828 | 12948.1019 |
| 824.020187 | 665.357659 | 717.442864 | 1051.39045 |
| 819.229372 | 657.031227 | 709.700675 | 1045.04147 |
| 817.432816 | 654.003433 | 707.119945 | 1036.1529 |
| 6.58737068 | 9.84032943 | 6.45182432 | 5.0791809 |
| 6.58737068 | 9.84032943 | 6.45182432 | 5.0791809 |
| 681.493439 | 827.344621 | 1058.09919 | 1221.54301 |
| 828.811002 | 938.616038 | 1494.24251 | 2130.71639 |
| 826.415594 | 938.616038 | 1494.24251 | 2126.907 |
| 6.58737068 | 9.84032943 | 7.74218918 | 5.0791809 |
| 8.38392632 | 18.166762 | 12.9036486 | 13.9677475 |
| 3130.19878 | 2461.59626 | 3263.33274 | 2831.64335 |

| | | | |
|---|---|---|---|
| 9.58163008 | 17.4098136 | 11.6132838 | 12.6979522 |
| 5527.40285 | 6904.88347 | 8103.49134 | 10962.1422 |
| 67.6702624 | 52.9863893 | 68.3893378 | 62.219966 |
| 413.806649 | 770.57349 | 580.664189 | 745.369796 |
| 2.9942594 | 6.81253576 | 2.58072973 | 2.53959045 |
| 2449.90304 | 2827.95929 | 3157.52282 | 3375.11571 |
| 2440.32141 | 2825.68844 | 3156.23246 | 3368.76673 |
| 972.535453 | 785.712458 | 592.277472 | 906.63379 |
| 16099.5339 | 21166.5486 | 27875.7522 | 39526.1857 |
| 853.363929 | 862.164248 | 700.668121 | 779.654268 |
| 101.80482 | 129.438179 | 72.2604324 | 93.9648466 |
| 99.4094121 | 127.924283 | 69.6797026 | 91.4252561 |
| 867.137522 | 869.733732 | 700.668121 | 787.273039 |
| 203.010787 | 219.515041 | 134.197946 | 163.803584 |
| 33499.7742 | 40112.9675 | 51404.2651 | 68264.1912 |
| 30268.3694 | 38223.6243 | 49773.2439 | 64922.0902 |
| 30256.9912 | 38209.2422 | 49747.4366 | 64891.6151 |
| 30250.4039 | 38201.6728 | 49747.4366 | 64875.1078 |
| 30250.4039 | 38201.6728 | 49747.4366 | 64875.1078 |
| 30282.143 | 38224.3812 | 49778.4053 | 64960.1841 |
| 16.1690008 | 20.4376073 | 15.4843784 | 21.5865188 |
| 31433.1363 | 39715.5696 | 51399.1036 | 67928.9653 |
| 31443.3168 | 39726.9238 | 51412.0073 | 67953.0914 |
| 31493.6204 | 39912.3762 | 51539.7534 | 68424.1854 |
| 32061.3319 | 40956.2081 | 52695.9203 | 70863.4621 |
| 27314.8319 | 36107.1965 | 46912.505 | 61337.4583 |
| 7.78507444 | 9.84032943 | 5.16145946 | 3.80938567 |
| 24174.4527 | 31192.3304 | 35602.457 | 45443.4315 |
| 24196.0114 | 31184.004 | 35581.8111 | 45381.2115 |
| 24184.0343 | 31229.4209 | 35626.9739 | 45466.2878 |
| 24179.2435 | 31239.2612 | 35629.5546 | 45478.9857 |
| 24179.2435 | 31239.2612 | 35629.5546 | 45478.9857 |
| 24055.2812 | 31102.2536 | 35517.2929 | 45265.6601 |
| 24055.2812 | 31102.2536 | 35517.2929 | 45265.6601 |
| 24055.2812 | 31102.2536 | 35517.2929 | 45265.6601 |
| 24236.1344 | 31268.0253 | 35666.9752 | 45546.2849 |
| 150.311822 | 254.334668 | 171.618527 | 154.915017 |
| 229.36027 | 236.167906 | 145.81123 | 173.961946 |
| 229.36027 | 236.167906 | 145.81123 | 173.961946 |
| 230.557974 | 236.167906 | 145.81123 | 175.231741 |
| 237.744196 | 237.681803 | 153.553419 | 185.390103 |
| 231.755678 | 237.681803 | 145.81123 | 173.961946 |
| 234.749937 | 239.1957 | 147.101594 | 180.310922 |
| 241.337308 | 238.438752 | 149.682324 | 181.580717 |
| 234.151085 | 239.952649 | 153.553419 | 177.771331 |
| 250.918938 | 245.251287 | 160.005243 | 193.008874 |

| | | | |
|---|---|---|---|
| 17.3667045 | 21.1945557 | 15.4843784 | 21.5865188 |
| 600.049584 | 1043.83187 | 869.705918 | 961.234985 |
| 10502.6643 | 8345.35631 | 5976.97005 | 12220.5092 |
| 482.075763 | 288.397347 | 329.04304 | 365.701025 |
| 482.075763 | 288.397347 | 329.04304 | 365.701025 |
| 27.5471865 | 40.8752146 | 29.6783919 | 38.0938567 |
| 5.38966692 | 7.56948418 | 6.45182432 | 5.0791809 |
| 5.38966692 | 7.56948418 | 6.45182432 | 5.0791809 |
| 5.38966692 | 7.56948418 | 6.45182432 | 5.0791809 |
| 5.38966692 | 7.56948418 | 6.45182432 | 5.0791809 |
| 42.5184835 | 47.6877503 | 52.9049594 | 40.6334472 |
| 1854.64427 | 1205.81883 | 997.45204 | 877.4285 |
| 70.0656699 | 111.271417 | 91.6159053 | 88.8856657 |
| 46.1115947 | 53.7433377 | 58.0664189 | 60.9501708 |
| 8.38392632 | 15.8959168 | 15.4843784 | 11.428157 |
| 9.58163008 | 19.6806589 | 15.4843784 | 24.1261093 |
| 7308.38834 | 6244.0675 | 4970.48546 | 4962.35974 |
| 7308.38834 | 6244.82445 | 4970.48546 | 4962.35974 |
| 160.492304 | 264.174998 | 180.651081 | 168.882765 |
| 1.19770376 | 1.51389684 | 0 | 1.26979522 |
| 10.7793338 | 14.3820199 | 15.4843784 | 11.428157 |
| 258.10516 | 297.480728 | 259.363338 | 213.325598 |
| 4329.10024 | 3357.82318 | 2352.33515 | 4291.90786 |
| 4321.91402 | 3351.7676 | 2353.62551 | 4286.82868 |
| 4323.71057 | 3351.7676 | 2351.04478 | 4288.09847 |
| 4329.69909 | 3355.55234 | 2353.62551 | 4289.36827 |
| 4321.91402 | 3351.7676 | 2352.33515 | 4286.82868 |
| 4329.69909 | 3357.06623 | 2351.04478 | 4289.36827 |
| 4322.51287 | 3354.03844 | 2356.20624 | 4285.55888 |
| 4320.11746 | 3351.7676 | 2351.04478 | 4285.55888 |
| 4320.11746 | 3351.7676 | 2351.04478 | 4286.82868 |
| 4319.51861 | 3351.7676 | 2351.04478 | 4285.55888 |
| 4319.51861 | 3351.7676 | 2351.04478 | 4285.55888 |
| 4319.51861 | 3351.7676 | 2351.04478 | 4285.55888 |
| 4319.51861 | 3351.7676 | 2351.04478 | 4289.36827 |
| 1145.60365 | 910.608947 | 927.772337 | 798.701196 |
| 73.6587812 | 58.2850282 | 49.0338648 | 45.7126281 |
| 107.793338 | 153.660529 | 112.261743 | 111.74198 |
| 1809.13153 | 1663.77262 | 1583.27769 | 1777.71331 |
| 110.188746 | 152.90358 | 112.261743 | 115.551365 |
| 209.598158 | 185.452362 | 169.037797 | 135.868089 |
| 687.481958 | 666.871556 | 602.600391 | 539.66297 |
| 1.19770376 | 0.75694842 | 1.29036486 | 2.53959045 |
| 105.996783 | 151.389684 | 113.552108 | 110.472184 |
| 1069.54946 | 892.442185 | 882.609567 | 783.463653 |
| 3939.84652 | 2388.17226 | 1934.25693 | 2958.62287 |

| | | | |
|---|---|---|---|
| 3939.84652 | 2388.17226 | 1934.25693 | 2958.62287 |
| 3939.84652 | 2388.17226 | 1934.25693 | 2958.62287 |
| 3940.44537 | 2389.68616 | 1934.25693 | 2961.16246 |
| 3941.04422 | 2389.68616 | 1935.5473 | 2961.16246 |
| 3940.44537 | 2389.68616 | 1935.5473 | 2961.16246 |
| 3940.44537 | 2390.4431 | 1936.83766 | 2962.43226 |
| 3940.44537 | 2389.68616 | 1935.5473 | 2961.16246 |
| 107.793338 | 153.660529 | 113.552108 | 114.28157 |
| 105.996783 | 152.90358 | 112.261743 | 109.202389 |
| 1059.96783 | 888.657443 | 877.448107 | 773.305291 |
| 1055.77586 | 887.900494 | 873.577013 | 772.035496 |
| 34.733409 | 22.7084525 | 23.2265675 | 15.2375427 |
| 1828.29479 | 1675.12685 | 1588.43915 | 1800.56963 |
| 29.3437421 | 25.7362462 | 23.2265675 | 25.3959045 |
| 29.942594 | 15.1389684 | 18.0651081 | 8.88856657 |
| 1827.69594 | 1673.61295 | 1581.98732 | 1794.22065 |
| 615.020881 | 1096.81826 | 682.603013 | 1012.02679 |
| 364.101943 | 643.406155 | 1122.61743 | 1420.90086 |
| 364.101943 | 643.406155 | 1122.61743 | 1420.90086 |
| 1355.80066 | 479.905297 | 856.80227 | 678.07065 |
| 179.056712 | 419.349424 | 330.333405 | 388.557339 |
| 179.056712 | 419.349424 | 330.333405 | 388.557339 |
| 141.329044 | 151.389684 | 162.585973 | 159.994198 |
| 683.289995 | 654.003433 | 565.17981 | 518.076451 |
| 708.441774 | 666.114608 | 578.083459 | 529.504608 |
| 226.964862 | 234.65401 | 232.265675 | 175.231741 |
| 474.889541 | 451.898206 | 326.462311 | 396.17611 |
| 695.267033 | 654.760382 | 569.050905 | 520.616042 |
| 682.092291 | 653.246485 | 570.34127 | 519.346247 |
| 109.589894 | 71.9100997 | 135.488311 | 83.8064848 |
| 340.147868 | 228.598422 | 243.878959 | 194.278669 |
| 683.289995 | 654.003433 | 569.050905 | 521.885837 |
| 692.272773 | 656.274278 | 571.631635 | 521.885837 |
| 1850.45231 | 1689.50887 | 1589.72951 | 1810.72799 |
| 2.39540752 | 3.78474209 | 6.45182432 | 2.53959045 |
| 1852.84772 | 1689.50887 | 1592.31024 | 1811.99778 |
| 534.774729 | 451.898206 | 392.270919 | 336.495734 |
| 34.733409 | 31.7918336 | 34.8398513 | 24.1261093 |
| 1.79655564 | 1.51389684 | 3.87109459 | 3.80938567 |
| 412.010093 | 765.274851 | 775.509283 | 939.648466 |
| 565.915026 | 595.718405 | 480.015729 | 472.363823 |
| 565.915026 | 597.232302 | 481.306094 | 468.554438 |
| 34.733409 | 31.0348851 | 27.0976621 | 30.4750854 |
| 565.316175 | 594.961457 | 476.144635 | 463.475257 |
| 7.78507444 | 4.54169051 | 10.3229189 | 3.80938567 |
| 168.277378 | 343.654582 | 326.462311 | 382.208362 |

| | | | |
|---|---|---|---|
| 567.711582 | 598.746199 | 480.015729 | 468.554438 |
| 97.6128564 | 111.271417 | 127.746122 | 116.821161 |
| 458.121688 | 656.274278 | 503.242297 | 744.100001 |
| 453.929725 | 805.393117 | 823.252783 | 1003.13823 |
| 106.595635 | 64.3406155 | 58.0664189 | 67.2991469 |
| 98.2117083 | 127.167334 | 113.552108 | 110.472184 |
| 561.124211 | 587.391972 | 470.983175 | 459.665871 |
| 10.7793338 | 18.166762 | 11.6132838 | 8.88856657 |
| 10224.797 | 10228.644 | 11226.1743 | 18858.9987 |
| 10224.797 | 10228.644 | 11226.1743 | 18858.9987 |
| 10224.797 | 10228.644 | 11226.1743 | 18858.9987 |
| 4770.45407 | 2403.31123 | 2610.40812 | 3350.9896 |
| 5.38966692 | 6.05558734 | 9.03255405 | 2.53959045 |
| 1824.10283 | 1675.8838 | 1603.92353 | 1795.49045 |
| 335.955905 | 223.299783 | 237.427135 | 191.739079 |
| 31.1402978 | 40.1182662 | 37.420581 | 31.7448806 |
| 1039.60686 | 370.904725 | 512.274851 | 542.202561 |
| 4709.97004 | 2388.17226 | 2597.50447 | 3324.3239 |
| 4707.57463 | 2388.17226 | 2596.21411 | 3323.0541 |
| 43.1173353 | 65.8545124 | 52.9049594 | 49.5220137 |
| 43.7161872 | 66.6114608 | 50.3242297 | 49.5220137 |
| 124.561191 | 121.868695 | 116.132838 | 76.1877134 |
| 42.5184835 | 67.3684092 | 49.0338648 | 50.791809 |
| 37.7276684 | 74.9378934 | 52.9049594 | 72.3783278 |
| 92.2231895 | 254.334668 | 181.941446 | 153.645222 |
| 45.5127429 | 72.6670481 | 63.2278783 | 59.6803755 |
| 30.5414459 | 54.5002861 | 37.420581 | 33.0146758 |
| 44.3150391 | 65.097564 | 49.0338648 | 52.0616042 |
| 18.5644083 | 15.8959168 | 11.6132838 | 11.428157 |
| 84.4381151 | 92.347707 | 100.648459 | 82.5366896 |
| 319.188052 | 168.042549 | 165.166703 | 147.296246 |
| 1.19770376 | 1.51389684 | 1.29036486 | 1.26979522 |
| 1.19770376 | 1.51389684 | 1.29036486 | 1.26979522 |
| 35.3322609 | 32.548782 | 25.8072973 | 29.2052902 |
| 77.2518925 | 89.3199133 | 99.3580945 | 77.4575087 |
| 1133.62661 | 837.941899 | 1047.77627 | 1315.50785 |
| 34.1345572 | 31.0348851 | 25.8072973 | 29.2052902 |
| 126.956599 | 97.6463459 | 104.519554 | 88.8856657 |
| 74.856485 | 88.5629649 | 94.1966351 | 76.1877134 |
| 649.155438 | 697.149493 | 536.791783 | 504.108704 |
| 76.0541887 | 90.0768618 | 96.7773648 | 77.4575087 |
| 30.5414459 | 78.7226355 | 68.3893378 | 76.1877134 |
| 0.59885188 | 0.75694842 | 2.58072973 | 0 |
| 352.723757 | 218.758093 | 259.363338 | 200.627645 |
| 76.0541887 | 88.5629649 | 95.4869999 | 78.7273039 |
| 0 | 0 | 0 | 0 |

| | | | |
|---|---|---|---|
| 71.8622256 | 86.2921197 | 94.1966351 | 76.1877134 |
| 14965.3085 | 7262.92007 | 9191.26892 | 10163.441 |
| 14956.3257 | 7259.89228 | 9184.8171 | 10158.3618 |
| 31.7391496 | 31.0348851 | 25.8072973 | 25.3959045 |
| 1.19770376 | 1.51389684 | 1.29036486 | 1.26979522 |
| 15971.3796 | 8063.7715 | 10313.8864 | 11071.3446 |
| 15964.7923 | 8066.04234 | 10311.3056 | 11070.0748 |
| 74.856485 | 88.5629649 | 94.1966351 | 76.1877134 |
| 15973.1762 | 8067.55624 | 10313.8864 | 11078.9633 |
| 75.4553369 | 88.5629649 | 94.1966351 | 79.9970991 |
| 76.6530406 | 88.5629649 | 94.1966351 | 78.7273039 |
| 16248.6481 | 8131.13991 | 10414.5348 | 11223.72 |
| 570.10699 | 532.891686 | 637.440243 | 454.58669 |
| 16230.6825 | 8126.59822 | 10361.6299 | 11155.151 |
| 37.1288166 | 36.3335241 | 34.8398513 | 29.2052902 |
| 170.073934 | 257.362462 | 296.783919 | 317.448806 |
| 523.396543 | 281.584812 | 331.62377 | 351.733277 |
| 474.290689 | 265.688895 | 298.074284 | 332.686349 |
| 34.1345572 | 48.4446988 | 52.9049594 | 41.9032424 |
| 33.5357053 | 46.1738535 | 50.3242297 | 40.6334472 |
| 4.79081504 | 8.3264326 | 11.6132838 | 7.61877134 |
| 4.79081504 | 7.56948418 | 11.6132838 | 7.61877134 |
| 43.1173353 | 52.2294408 | 41.2916756 | 36.8240615 |
| 406.620426 | 501.099853 | 445.175878 | 485.061776 |
| 6619.70868 | 9625.35608 | 8233.8182 | 13405.2282 |
| 6444.24508 | 9501.97349 | 8078.97441 | 13279.5185 |
| 4.19196316 | 1.51389684 | 0 | 1.26979522 |
| 1.79655564 | 3.02779367 | 1.29036486 | 1.26979522 |
| 4705.77807 | 1750.82169 | 2223.29866 | 2538.32065 |
| 6571.20168 | 9591.29341 | 8223.49528 | 13364.5947 |
| 6571.20168 | 9592.05035 | 8220.91455 | 13367.1343 |
| 6578.3879 | 9597.34899 | 8218.33382 | 13370.9437 |
| 6580.18446 | 9610.97406 | 8231.23747 | 13381.1021 |
| 6580.18446 | 9610.97406 | 8231.23747 | 13381.1021 |
| 6580.78331 | 9614.75881 | 8231.23747 | 13383.6417 |
| 38.3265203 | 44.6599567 | 33.5494865 | 33.0146758 |
| 1.19770376 | 1.51389684 | 1.29036486 | 1.26979522 |
| 730.000442 | 607.82958 | 725.185053 | 540.932765 |
| 730.000442 | 607.82958 | 725.185053 | 540.932765 |
| 730.000442 | 608.586528 | 725.185053 | 542.202561 |
| 8.38392632 | 8.3264326 | 9.03255405 | 6.34897612 |
| 121.566932 | 118.083953 | 131.617216 | 102.853413 |
| 10.7793338 | 6.81253576 | 6.45182432 | 6.34897612 |
| 4.79081504 | 8.3264326 | 11.6132838 | 7.61877134 |
| 303.019051 | 414.050785 | 540.662878 | 344.114506 |
| 329.368534 | 436.759237 | 561.308716 | 364.431229 |

| | | | |
|---|---|---|---|
| 48.5070023 | 40.8752146 | 38.7109459 | 33.0146758 |
| 21.5586677 | 20.4376073 | 21.9362027 | 15.2375427 |
| 631.788733 | 1082.43624 | 997.45204 | 1229.16178 |
| 537.768988 | 859.136455 | 918.739783 | 1166.94181 |
| 2.9942594 | 6.81253576 | 10.3229189 | 5.0791809 |
| 645.562326 | 1086.97793 | 1000.03277 | 1229.16178 |
| 535.972432 | 857.622558 | 918.739783 | 1166.94181 |
| 109.589894 | 155.174426 | 118.713567 | 95.2346418 |
| 57.4897805 | 77.9656871 | 80.0026216 | 68.5689421 |
| 1.79655564 | 2.27084525 | 5.16145946 | 1.26979522 |
| 5460.33144 | 6486.29099 | 7108.62003 | 11689.7348 |
| 22.7563714 | 21.1945557 | 21.9362027 | 21.5865188 |
| 51.5012617 | 74.9378934 | 76.131527 | 62.219966 |
| 749.163702 | 612.37127 | 731.636878 | 551.091127 |
| 751.559109 | 613.128219 | 731.636878 | 551.091127 |
| 61.0828917 | 71.9100997 | 60.6471486 | 58.4105803 |
| 752.157961 | 612.37127 | 732.927243 | 549.821332 |
| 752.756813 | 614.642115 | 730.346513 | 551.091127 |
| 22.7563714 | 21.1945557 | 21.9362027 | 21.5865188 |
| 26.9483346 | 33.3057304 | 28.388027 | 26.6656997 |
| 23.3552233 | 21.1945557 | 21.9362027 | 22.856314 |
| 175.463601 | 98.4032943 | 87.7448107 | 97.7742322 |
| 247.924678 | 377.717261 | 282.589905 | 375.859386 |
| 438.359576 | 196.04964 | 192.264365 | 195.548464 |
| 0 | 0 | 0 | 0 |
| 456.325132 | 218.758093 | 225.813851 | 219.674574 |
| 390.451426 | 997.658015 | 841.317891 | 723.783278 |
| 2.39540752 | 4.54169051 | 6.45182432 | 5.0791809 |
| 1.79655564 | 4.54169051 | 10.3229189 | 2.53959045 |
| 453.330873 | 526.079151 | 614.213675 | 443.158533 |
| 141.329044 | 86.2921197 | 73.5507972 | 69.8387373 |
| 452.732021 | 526.079151 | 615.50404 | 444.428328 |
| 64.676003 | 90.0768618 | 99.3580945 | 72.3783278 |
| 0 | 0 | 0 | 0 |
| 151.509526 | 149.875787 | 134.197946 | 105.393004 |
| 1.79655564 | 2.27084525 | 3.87109459 | 1.26979522 |
| 73.6587812 | 55.2572345 | 49.0338648 | 49.5220137 |
| 680.894587 | 1598.67506 | 1381.98077 | 1342.17355 |
| 0.59885188 | 0 | 0 | 0 |
| 135.939377 | 138.521561 | 99.3580945 | 119.360751 |
| 137.735932 | 140.792406 | 101.938824 | 120.630546 |
| 108.39219 | 150.632735 | 197.425824 | 165.073379 |
| 0 | 0.75694842 | 0 | 0 |
| 140.13134 | 143.820199 | 105.809919 | 123.170137 |
| 0.59885188 | 1.51389684 | 1.29036486 | 1.26979522 |
| 0.59885188 | 1.51389684 | 0 | 0 |

| | | | |
|---|---|---|---|
| 140.13134 | 86.2921197 | 68.3893378 | 64.7595564 |
| 147.317562 | 152.146632 | 112.261743 | 128.249318 |
| 108.39219 | 77.2087386 | 49.0338648 | 49.5220137 |
| 147.916414 | 152.90358 | 108.390649 | 132.058703 |
| 111.38645 | 92.347707 | 94.1966351 | 82.5366896 |
| 136.538229 | 138.521561 | 101.938824 | 119.360751 |
| 6.58737068 | 9.84032943 | 12.9036486 | 8.88856657 |
| 23.3552233 | 24.2223494 | 25.8072973 | 24.1261093 |
| 44.913891 | 70.3962029 | 81.2929864 | 52.0616042 |
| 1.19770376 | 0 | 0 | 0 |
| 148.515266 | 147.604942 | 122.584662 | 123.170137 |
| 16.7678526 | 18.9237105 | 20.6458378 | 15.2375427 |
| 21844.32 | 13876.3784 | 13117.8492 | 21290.6565 |
| 0.59885188 | 1.51389684 | 2.58072973 | 0 |
| 21854.5005 | 13892.2743 | 13124.301 | 21281.768 |
| 132.945117 | 62.0697703 | 56.776054 | 83.8064848 |
| 3.59311128 | 1.51389684 | 0 | 1.26979522 |
| 159.893452 | 153.660529 | 112.261743 | 134.598294 |
| 52.6989654 | 80.9934807 | 95.4869999 | 44.4428328 |
| 160.492304 | 153.660529 | 112.261743 | 134.598294 |
| 0 | 0.75694842 | 1.29036486 | 1.26979522 |
| 79.6473 | 76.4517902 | 68.3893378 | 68.5689421 |
| 25.7506308 | 27.2501431 | 30.9687567 | 26.6656997 |
| 25.7506308 | 27.2501431 | 30.9687567 | 26.6656997 |
| 25.7506308 | 27.2501431 | 30.9687567 | 26.6656997 |
| 25.7506308 | 27.2501431 | 30.9687567 | 26.6656997 |
| 25.7506308 | 27.2501431 | 30.9687567 | 26.6656997 |
| 25.7506308 | 27.2501431 | 30.9687567 | 26.6656997 |
| 6.58737068 | 9.08338102 | 6.45182432 | 3.80938567 |
| 16.1690008 | 18.166762 | 21.9362027 | 15.2375427 |
| 1.19770376 | 1.51389684 | 1.29036486 | 3.80938567 |
| 158.695748 | 153.660529 | 112.261743 | 132.058703 |
| 0 | 0 | 0 | 0 |
| 115.578413 | 140.792406 | 126.455757 | 157.454608 |
| 76.0541887 | 43.9030082 | 37.420581 | 36.8240615 |
| 25.7506308 | 27.2501431 | 30.9687567 | 26.6656997 |
| 25.7506308 | 27.2501431 | 30.9687567 | 26.6656997 |
| 25.7506308 | 27.2501431 | 30.9687567 | 26.6656997 |
| 25.7506308 | 27.2501431 | 30.9687567 | 26.6656997 |
| 25.7506308 | 27.2501431 | 30.9687567 | 26.6656997 |
| 25.7506308 | 27.2501431 | 30.9687567 | 26.6656997 |
| 25.7506308 | 27.2501431 | 30.9687567 | 26.6656997 |
| 25.7506308 | 27.2501431 | 30.9687567 | 26.6656997 |
| 25.7506308 | 27.2501431 | 30.9687567 | 26.6656997 |
| 25.7506308 | 27.2501431 | 30.9687567 | 26.6656997 |
| 25.7506308 | 27.2501431 | 30.9687567 | 26.6656997 |

| | | | |
|---|---|---|---|
| 25.7506308 | 27.2501431 | 30.9687567 | 26.6656997 |
| 25.7506308 | 27.2501431 | 30.9687567 | 26.6656997 |
| 145.521007 | 145.334096 | 104.519554 | 126.979522 |
| 17.3667045 | 20.4376073 | 24.5169324 | 13.9677475 |
| 42.5184835 | 30.2779367 | 11.6132838 | 16.5073379 |
| 0.59885188 | 0 | 0 | 0 |
| 25.7506308 | 27.2501431 | 30.9687567 | 26.6656997 |
| 25.7506308 | 27.2501431 | 30.9687567 | 26.6656997 |
| 25.7506308 | 27.2501431 | 30.9687567 | 26.6656997 |
| 25.7506308 | 27.2501431 | 30.9687567 | 26.6656997 |
| 25.7506308 | 27.2501431 | 30.9687567 | 26.6656997 |
| 25.7506308 | 27.2501431 | 30.9687567 | 26.6656997 |
| 25.7506308 | 27.2501431 | 30.9687567 | 26.6656997 |
| 25.7506308 | 27.2501431 | 30.9687567 | 26.6656997 |
| 25.7506308 | 27.2501431 | 30.9687567 | 26.6656997 |
| 169.475082 | 175.612033 | 149.682324 | 123.170137 |
| 53.2978173 | 36.3335241 | 25.8072973 | 30.4750854 |
| 147.916414 | 49.2016472 | 36.1302162 | 33.0146758 |
| 67.0714105 | 199.834382 | 224.523486 | 214.595393 |
| 65.8737068 | 199.077434 | 221.942757 | 212.055802 |
| 77.8507444 | 44.6599567 | 37.420581 | 38.0938567 |
| 228.162566 | 281.584812 | 237.427135 | 226.02355 |
| 117.97382 | 96.8893975 | 98.0677296 | 85.07628 |
| 116.177265 | 78.7226355 | 95.4869999 | 72.3783278 |
| 26.3494827 | 28.0070915 | 30.9687567 | 26.6656997 |
| 26.3494827 | 28.0070915 | 30.9687567 | 26.6656997 |
| 26.3494827 | 28.0070915 | 30.9687567 | 26.6656997 |
| 1.79655564 | 1.51389684 | 1.29036486 | 1.26979522 |
| 3.59311128 | 5.29863893 | 3.87109459 | 2.53959045 |
| 1780.98549 | 1188.40902 | 1165.19947 | 943.457851 |
| 0 | 0 | 0 | 0 |
| 5.9885188 | 8.3264326 | 11.6132838 | 8.88856657 |
| 131.148562 | 55.2572345 | 63.2278783 | 116.821161 |
| 163.486563 | 156.688323 | 113.552108 | 134.598294 |
| 16343.2667 | 11505.616 | 10395.1793 | 16493.3702 |
| 16337.877 | 11499.5604 | 10395.1793 | 16489.5608 |
| 16390.576 | 11525.2966 | 10413.2445 | 16523.8452 |
| 140.13134 | 59.798925 | 65.8086081 | 120.630546 |
| 0 | 1.51389684 | 2.58072973 | 0 |
| 140.13134 | 59.0419766 | 65.8086081 | 119.360751 |
| 140.13134 | 59.0419766 | 65.8086081 | 119.360751 |
| 16291.7654 | 11443.5462 | 10331.9515 | 16400.6751 |
| 77.2518925 | 21.9515041 | 20.6458378 | 35.5542663 |
| 1780.38664 | 1202.03409 | 1163.90911 | 951.076623 |
| 56.2920767 | 65.8545124 | 64.5182432 | 69.8387373 |
| 1745.65323 | 1164.18667 | 1136.81144 | 906.63379 |

| | | | |
|---|---|---|---|
| 16.7678526 | 18.9237105 | 20.6458378 | 15.2375427 |
| 1747.44979 | 1162.67277 | 1136.81144 | 905.363995 |
| 153.904933 | 157.445271 | 107.100284 | 133.328499 |
| 1740.26356 | 1158.13108 | 1136.81144 | 896.475428 |
| 1740.26356 | 1158.13108 | 1136.81144 | 896.475428 |
| 1740.26356 | 1158.13108 | 1136.81144 | 896.475428 |
| 1745.05438 | 1157.37413 | 1134.23072 | 902.824404 |
| 4.19196316 | 8.3264326 | 9.03255405 | 5.0791809 |
| 377.276684 | 741.052501 | 652.924621 | 575.217236 |
| 0 | 0.75694842 | 0 | 0 |
| 152.707229 | 152.90358 | 109.681013 | 133.328499 |
| 1.19770376 | 3.78474209 | 3.87109459 | 2.53959045 |
| 28.1460384 | 34.0626788 | 25.8072973 | 24.1261093 |
| 28.1460384 | 34.0626788 | 25.8072973 | 24.1261093 |
| 329.967386 | 244.494339 | 243.878959 | 198.088055 |
| 2.9942594 | 3.02779367 | 3.87109459 | 2.53959045 |
| 111.985302 | 86.2921197 | 101.938824 | 77.4575087 |
| 0 | 0.75694842 | 1.29036486 | 0 |
| 26.9483346 | 32.548782 | 24.5169324 | 21.5865188 |
| 17.3667045 | 18.166762 | 20.6458378 | 17.7771331 |
| 0 | 0 | 0 | 0 |
| 412.608945 | 132.465973 | 170.328162 | 147.296246 |
| 39.5242241 | 52.2294408 | 34.8398513 | 36.8240615 |
| 0.59885188 | 3.78474209 | 5.16145946 | 1.26979522 |
| 0 | 0 | 0 | 0 |
| 68.2691143 | 37.0904725 | 36.1302162 | 29.2052902 |
| 68.2691143 | 37.0904725 | 36.1302162 | 29.2052902 |
| 173.667045 | 85.5351712 | 78.7122567 | 72.3783278 |
| 38.9253722 | 51.4724924 | 34.8398513 | 38.0938567 |

| FGCZ6510_Free_tRNA_dedup_pvalue | | | | FGCZ6510_Free_tRNA_de | |
|---|---|---|---|---|---|
| | baseMean | log2FoldChai | lfcSE | stat | pvalue |
| tRNA-Ala-TGC-2-1 | 277.679415 | -1.4485942 | 0.34435998 | -4.2066275 | 2.59E-05 |
| tRNA-Arg-ACG-3-1 | 116.413494 | -0.8072572 | 0.19230003 | -4.1979047 | 2.69E-05 |
| tRNA-Glu-CTC-3-1 | 1906.1005 | -2.2628678 | 0.54085901 | -4.1838404 | 2.87E-05 |
| tRNA-Ser-TGA-1-1 | 324.846043 | -1.2581467 | 0.3323469 | -3.7856428 | 0.00015331 |
| tRNA-Thr-TGT-2-1 | 600.456646 | 0.75687273 | 0.20142145 | 3.75765698 | 0.00017151 |
| tRNA-Ser-TGA-2-1 | 611.407598 | 0.59434449 | 0.15866841 | 3.74582755 | 0.0001798 |
| tRNA-Ser-AGA-1-1 | 608.984968 | 0.59430847 | 0.15883996 | 3.74155517 | 0.00018289 |
| tRNA-Ser-TGA-2-2 | 609.526982 | 0.59155096 | 0.15912595 | 3.71750152 | 0.0002012 |
| tRNA-Glu-TTC-3-1 | 4268.88183 | -1.851725 | 0.50754418 | -3.6484016 | 0.00026388 |
| tRNA-Glu-TTC-3-2 | 4268.57504 | -1.851809 | 0.50759728 | -3.6481855 | 0.0002641 |
| tRNA-Arg-CCT-2-1 | 432.571723 | 1.12164965 | 0.30860996 | 3.63452185 | 0.0002785 |
| tRNA-Arg-CCT-2-2 | 432.623123 | 1.12108727 | 0.30852931 | 3.63364917 | 0.00027944 |
| tRNA-Arg-CCT-1-1 | 433.506919 | 1.11181529 | 0.30822982 | 3.60709844 | 0.00030964 |
| tRNA-Ser-AGA-2-4 | 557.703085 | 0.60879587 | 0.171891 | 3.54175541 | 0.00039747 |
| tRNA-Ser-AGA-2-5 | 557.755902 | 0.60871469 | 0.17193185 | 3.54044181 | 0.00039946 |
| tRNA-Ser-AGA-2-3 | 557.816971 | 0.60815827 | 0.17192233 | 3.53740121 | 0.00040409 |
| tRNA-Ser-AGA-2-6 | 557.774675 | 0.60839498 | 0.17200717 | 3.53703258 | 0.00040465 |
| tRNA-Ser-AGA-2-1 | 561.621234 | 0.61092102 | 0.17307784 | 3.52974711 | 0.00041596 |
| tRNA-Ser-AGA-2-2 | 559.229981 | 0.60158279 | 0.17240004 | 3.48945851 | 0.000484 |
| tRNA-Cys-GCA-3-4 | 1087.13101 | -0.9066827 | 0.26121297 | -3.4710479 | 0.00051843 |
| tRNA-Cys-GCA-3-1 | 1082.90758 | -0.899756 | 0.26014275 | -3.4587011 | 0.00054279 |
| tRNA-Cys-GCA-3-2 | 1086.80606 | -0.8998797 | 0.26020429 | -3.4583583 | 0.00054348 |
| tRNA-Cys-GCA-3-3 | 1086.49353 | -0.899321 | 0.26014012 | -3.4570639 | 0.0005461 |
| tRNA-Pro-AGG-3-1 | 12.2481732 | 0.95400114 | 0.27648464 | 3.45046698 | 0.00055962 |
| tRNA-Lys-CTT-3-2 | 13569.9562 | -1.0961215 | 0.31806585 | -3.4462093 | 0.00056851 |
| tRNA-Arg-CCT-3-1 | 419.301153 | 1.08035173 | 0.31476236 | 3.43227737 | 0.00059854 |
| tRNA-Lys-CTT-3-4 | 13360.2378 | -1.0932197 | 0.31906034 | -3.426373 | 0.0006117 |
| tRNA-Lys-CTT-3-1 | 13358.4571 | -1.0932049 | 0.31906805 | -3.4262437 | 0.00061199 |
| tRNA-Lys-CTT-3-7 | 13327.7229 | -1.0910416 | 0.31885561 | -3.421742 | 0.00062221 |
| tRNA-Trp-CCA-5-1 | 212.778282 | -0.6487483 | 0.18967306 | -3.4203501 | 0.00062541 |
| tRNA-Lys-CTT-3-3 | 13323.3065 | -1.091074 | 0.31899714 | -3.4203254 | 0.00062546 |
| tRNA-Lys-CTT-3-5 | 13317.1316 | -1.0911603 | 0.31906517 | -3.4198665 | 0.00062652 |
| tRNA-Lys-CTT-3-6 | 13316.9927 | -1.0911755 | 0.31907503 | -3.4198084 | 0.00062665 |
| tRNA-Cys-GCA-2-1 | 1031.03725 | -0.9516053 | 0.27955731 | -3.4039722 | 0.00066414 |
| tRNA-Cys-GCA-1-2 | 1022.83035 | -0.9559027 | 0.28119915 | -3.3993798 | 0.00067539 |
| tRNA-Cys-GCA-1-1 | 1016.84573 | -0.9525364 | 0.28086764 | -3.3914068 | 0.00069535 |
| tRNA-Ala-TGC-5-1 | 14.2050982 | -1.2264398 | 0.36781759 | -3.3343695 | 0.00085493 |
| tRNA-Ala-TGC-5-2 | 14.2050982 | -1.2264398 | 0.36781759 | -3.3343695 | 0.00085493 |
| tRNA-Ala-CGC-5-1 | 1993.60453 | -1.6171175 | 0.4987579 | -3.2422895 | 0.00118574 |
| tRNA-Gly-CCC-3-1 | 2441.50499 | -1.689942 | 0.52225683 | -3.2358448 | 0.00121283 |
| tRNA-Gly-CCC-4-1 | 2439.5542 | -1.6913223 | 0.52270076 | -3.2357373 | 0.00121329 |
| tRNA-Ala-TGC-5-3 | 14.1968155 | -1.1880056 | 0.36954007 | -3.2148221 | 0.00130525 |
| tRNA-Thr-AGT-5-1 | 10.6999487 | 1.1929342 | 0.37205688 | 3.20632214 | 0.00134443 |
| tRNA-SeC-TCA-1-1 | 3770.94571 | -0.6430741 | 0.20192215 | -3.1847623 | 0.00144873 |

| | | | | | |
|---|---|---|---|---|---|
| tRNA-Thr-AGT-7-1 | 9.79467354 | 1.21981746 | 0.39860605 | 3.06020809 | 0.00221183 |
| tRNA-Glu-CTC-2-1 | 13617.7019 | -1.4304423 | 0.47348079 | -3.0211201 | 0.00251841 |
| tRNA-His-GTG-1-1 | 51.7845717 | 0.90415981 | 0.29937161 | 3.02019227 | 0.00252614 |
| tRNA-Arg-CCT-4-1 | 549.304687 | 0.69965012 | 0.2325263 | 3.00890753 | 0.00262189 |
| tRNA-Lys-CTT-1-1 | 10.2611094 | -1.5228688 | 0.50692103 | -3.0041539 | 0.00266321 |
| tRNA-Lys-CTT-2-2 | 3599.3706 | -0.7503053 | 0.25309032 | -2.9645752 | 0.00303101 |
| tRNA-Lys-CTT-2-1 | 3592.26036 | -0.7504091 | 0.25424621 | -2.9515055 | 0.00316229 |
| tRNA-Asp-GTC-4-1 | 1004.13241 | -0.6533448 | 0.22263449 | -2.9346075 | 0.0033397 |
| tRNA-Gly-GCC-5-1 | 36776.8066 | -1.3849301 | 0.47500191 | -2.9156305 | 0.00354971 |
| tRNA-Thr-TGT-3-1 | 748.792817 | 0.53373063 | 0.18555708 | 2.87636894 | 0.00402279 |
| tRNA-Asn-GTT-2-1 | 97.9018457 | 1.29290249 | 0.45815389 | 2.821983 | 0.00477277 |
| tRNA-Asn-GTT-4-1 | 94.906968 | 1.36400712 | 0.49124111 | 2.77665507 | 0.00549214 |
| tRNA-Thr-TGT-3-2 | 758.168904 | 0.5206757 | 0.18855137 | 2.76145283 | 0.00575448 |
| tRNA-Asn-GTT-1-1 | 172.721359 | 1.18249383 | 0.42892963 | 2.75684807 | 0.00583615 |
| tRNA-Gly-GCC-4-1 | 59156.1031 | -1.0704685 | 0.39921894 | -2.681407 | 0.00733133 |
| tRNA-Gly-GCC-2-8 | 56184.7674 | -1.1007499 | 0.41716408 | -2.6386497 | 0.00832369 |
| tRNA-Gly-GCC-2-7 | 56152.0319 | -1.1005568 | 0.4171677 | -2.638164 | 0.00833563 |
| tRNA-Gly-GCC-2-5 | 56141.1535 | -1.1005441 | 0.41722376 | -2.637779 | 0.0083451 |
| tRNA-Gly-GCC-2-4 | 56141.0376 | -1.1005397 | 0.41722565 | -2.6377565 | 0.00834565 |
| tRNA-Gly-GCC-2-1 | 56183.786 | -1.1002271 | 0.41714082 | -2.6375435 | 0.00835089 |
| tRNA-Thr-CGT-4-1 | 16.568736 | 0.71566179 | 0.27290849 | 2.62235079 | 0.00873255 |
| tRNA-Gly-GCC-2-2 | 57898.0766 | -1.0863122 | 0.41491128 | -2.6181796 | 0.00884003 |
| tRNA-Gly-GCC-2-6 | 57906.4604 | -1.0859093 | 0.41485688 | -2.6175516 | 0.00885631 |
| tRNA-Gly-GCC-2-3 | 58072.991 | -1.0842226 | 0.41521216 | -2.6112496 | 0.0090212 |
| tRNA-Gly-GCC-3-1 | 59226.4515 | -1.0761676 | 0.41545657 | -2.5903252 | 0.00958853 |
| tRNA-Gly-CCC-5-1 | 52106.2063 | -1.0993663 | 0.42648438 | -2.577741 | 0.00994485 |
| tRNA-Ala-TGC-6-1 | 10.0280049 | -0.9368417 | 0.36681185 | -2.5540116 | 0.01064897 |
| tRNA-Glu-CTC-1-4 | 41485.9349 | -0.9790005 | 0.39237679 | -2.495052 | 0.01259387 |
| tRNA-Glu-CTC-1-7 | 41450.552 | -0.9771519 | 0.39224168 | -2.4911984 | 0.0127313 |
| tRNA-Glu-CTC-1-8 | 41485.587 | -0.9769554 | 0.39242095 | -2.4895597 | 0.01279014 |
| tRNA-Glu-CTC-1-6 | 41488.7115 | -0.9768603 | 0.39242927 | -2.4892647 | 0.01280076 |
| tRNA-Glu-CTC-1-5 | 41488.3481 | -0.9768413 | 0.39242618 | -2.4892357 | 0.01280181 |
| tRNA-Glu-CTC-1-1 | 41320.9754 | -0.9781447 | 0.39300053 | -2.4889144 | 0.01281338 |
| tRNA-Glu-CTC-1-2 | 41320.9754 | -0.9781447 | 0.39300053 | -2.4889144 | 0.01281338 |
| tRNA-Glu-CTC-1-3 | 41320.9754 | -0.9781447 | 0.39300053 | -2.4889144 | 0.01281338 |
| tRNA-Glu-CTC-1-9 | 41529.9223 | -0.9758726 | 0.39226093 | -2.487815 | 0.01285306 |
| tRNA-Ala-AGC-4-1 | 156.732118 | 0.44680048 | 0.18400369 | 2.42821473 | 0.01517336 |
| tRNA-Asn-GTT-3-4 | 198.772269 | 0.96824645 | 0.40558176 | 2.38730278 | 0.01697251 |
| tRNA-Asn-GTT-3-5 | 198.772269 | 0.96824645 | 0.40558176 | 2.38730278 | 0.01697251 |
| tRNA-Asn-GTT-3-3 | 198.80355 | 0.97874717 | 0.4122325 | 2.37426009 | 0.01758416 |
| tRNA-Asn-GTT-3-7 | 206.163773 | 0.91182614 | 0.3856443 | 2.36442272 | 0.01805819 |
| tRNA-Asn-GTT-3-9 | 199.903053 | 0.95691472 | 0.40669051 | 2.35293102 | 0.01862609 |
| tRNA-Asn-GTT-3-6 | 204.067795 | 0.92746457 | 0.39540329 | 2.34561673 | 0.01899563 |
| tRNA-Asn-GTT-3-8 | 213.227496 | 0.81210465 | 0.34830436 | 2.33159482 | 0.01972202 |
| tRNA-Asn-GTT-3-1 | 206.500527 | 0.8777204 | 0.38035796 | 2.30761673 | 0.02102046 |
| tRNA-Asn-GTT-3-2 | 221.025057 | 0.76067272 | 0.33048947 | 2.30165496 | 0.02135464 |

| | | | | | |
|---|---|---|---|---|---|
| tRNA-Thr-TGT-1-1 | 17.7146198 | 0.62877282 | 0.27838321 | 2.25865931 | 0.02390459 |
| tRNA-Leu-TAA-2-1 | 1027.76479 | -0.5395775 | 0.24154235 | -2.2338835 | 0.02549074 |
| tRNA-Asp-GTC-2-1 | 10635.5684 | -0.5439398 | 0.24378021 | -2.2312715 | 0.02566315 |
| tRNA-Val-AAC-2-1 | 401.746028 | -0.4013591 | 0.18125151 | -2.2143765 | 0.02680289 |
| tRNA-Val-AAC-2-2 | 401.746028 | -0.4013591 | 0.18125151 | -2.2143765 | 0.02680289 |
| tRNA-Arg-CCG-1-1 | 29.9530106 | 0.56027658 | 0.25507493 | 2.19651763 | 0.02805491 |
| tRNA-Pro-AGG-2-1 | 5.67495425 | 0.82493497 | 0.37654257 | 2.19081464 | 0.02846521 |
| tRNA-Leu-TAA-5-1 | 5.67495425 | 0.82493497 | 0.37654257 | 2.19081464 | 0.02846521 |
| tRNA-Leu-TAA-5-2 | 5.67495425 | 0.82493497 | 0.37654257 | 2.19081464 | 0.02846521 |
| tRNA-Leu-TAA-5-3 | 5.67495425 | 0.82493497 | 0.37654257 | 2.19081464 | 0.02846521 |
| tRNA-Cys-GCA-5-1 | 56.9104327 | -0.5719957 | 0.26229097 | -2.1807679 | 0.02920059 |
| tRNA-Leu-TAG-3-1 | 1715.82666 | -0.6344645 | 0.29402335 | -2.1578711 | 0.03093786 |
| tRNA-Gln-TTG-5-1 | 73.7200832 | 0.74352311 | 0.35243268 | 2.10968829 | 0.03488521 |
| tRNA-Arg-ACG-3-2 | 66.903111 | -0.4101202 | 0.19443024 | -2.1093435 | 0.03491495 |
| tRNA-Cys-GCA-11-1 | 9.81617761 | 0.69432481 | 0.32993081 | 2.10445584 | 0.0353387 |
| tRNA-Thr-CGT-3-1 | 15.6987789 | 0.70691532 | 0.33698749 | 2.09774942 | 0.03592729 |
| tRNA-Gly-CCC-1-1 | 6806.74626 | -0.3911699 | 0.18844169 | -2.0758142 | 0.03791114 |
| tRNA-Gly-CCC-1-2 | 6806.79357 | -0.3911467 | 0.18844304 | -2.0756763 | 0.0379239 |
| tRNA-Ala-AGC-4-2 | 170.189021 | 0.37769334 | 0.18272373 | 2.06701855 | 0.03873241 |
| tRNA-Ala-CGC-7-1 | 2.24010756 | -1.3943839 | 0.67728715 | -2.058778 | 0.03951551 |
| tRNA-Phe-GAA-2-1 | 10.0756267 | 0.62463485 | 0.30931444 | 2.01941701 | 0.0434439 |
| tRNA-Ala-AGC-6-1 | 221.56568 | 0.46026805 | 0.22873978 | 2.01219066 | 0.04419984 |
| tRNA-Asp-GTC-1-11 | 3901.86935 | -0.425695 | 0.21229831 | -2.0051738 | 0.04494447 |
| tRNA-Asp-GTC-1-6 | 3895.39818 | -0.4240051 | 0.21180528 | -2.0018628 | 0.04529949 |
| tRNA-Asp-GTC-1-8 | 3895.04896 | -0.4234388 | 0.21170483 | -2.0001376 | 0.04548541 |
| tRNA-Asp-GTC-1-13 | 3899.07775 | -0.4234185 | 0.21172219 | -1.9998777 | 0.04551347 |
| tRNA-Asp-GTC-1-12 | 3894.08976 | -0.4233999 | 0.21180937 | -1.9989667 | 0.04561196 |
| tRNA-Asp-GTC-1-10 | 3897.09712 | -0.4233204 | 0.21194361 | -1.9973256 | 0.04578983 |
| tRNA-Asp-GTC-1-1 | 3891.46328 | -0.4221076 | 0.21177239 | -1.9932137 | 0.04623805 |
| tRNA-Asp-GTC-1-7 | 3889.05343 | -0.4215855 | 0.21174053 | -1.9910475 | 0.04647567 |
| tRNA-Asp-GTC-1-2 | 3888.95287 | -0.4214602 | 0.21176447 | -1.990231 | 0.04656549 |
| tRNA-Asp-GTC-1-9 | 3888.67415 | -0.4214297 | 0.21180169 | -1.989737 | 0.04661991 |
| tRNA-Asp-GTC-1-3 | 3888.71472 | -0.4214066 | 0.21180741 | -1.9895745 | 0.04663783 |
| tRNA-Asp-GTC-1-4 | 3888.71472 | -0.4214066 | 0.21180741 | -1.9895745 | 0.04663783 |
| tRNA-Asp-GTC-1-5 | 3889.47617 | -0.421288 | 0.211765 | -1.9894127 | 0.04665567 |
| tRNA-Leu-TAG-1-1 | 1286.22527 | -0.4253875 | 0.21430893 | -1.9849265 | 0.04715265 |
| tRNA-Arg-TCG-2-1 | 94.2436522 | -1.0161693 | 0.51200899 | -1.9846709 | 0.0471811 |
| tRNA-Ala-AGC-5-2 | 102.93114 | 0.37083824 | 0.18816049 | 1.97086134 | 0.04873974 |
| tRNA-Lys-TTT-1-2 | 1579.80114 | 0.18429909 | 0.09369569 | 1.96699641 | 0.04918362 |
| tRNA-Ala-AGC-5-3 | 103.244998 | 0.37356303 | 0.19117038 | 1.95408426 | 0.05069125 |
| tRNA-Arg-TCT-2-1 | 160.652101 | 0.38262166 | 0.1964677 | 1.94750411 | 0.05147433 |
| tRNA-iMet-CAT-1-3 | 780.593152 | -0.3118214 | 0.16039778 | -1.9440509 | 0.05188931 |
| tRNA-Cys-GCA-13-1 | 5.74454864 | -1.2310274 | 0.64399352 | -1.9115525 | 0.05593361 |
| tRNA-Ala-AGC-12-1 | 101.807493 | 0.36337116 | 0.1904496 | 1.90796496 | 0.05639575 |
| tRNA-Leu-AAG-1-3 | 1195.10887 | -0.4111976 | 0.21641927 | -1.9000044 | 0.05743255 |
| tRNA-His-GTG-2-4 | 2637.64809 | 0.37140392 | 0.19760389 | 1.87953754 | 0.06017113 |

| | | | | | |
|---|---|---|---|---|---|
| tRNA-His-GTG-2-5 | 2637.64809 | 0.37140392 | 0.19760389 | 1.87953754 | 0.06017113 |
| tRNA-His-GTG-2-6 | 2637.64809 | 0.37140392 | 0.19760389 | 1.87953754 | 0.06017113 |
| tRNA-His-GTG-2-7 | 2638.16342 | 0.37143937 | 0.19765159 | 1.87926327 | 0.06020855 |
| tRNA-His-GTG-2-8 | 2638.93468 | 0.37086009 | 0.19747462 | 1.87801394 | 0.06037926 |
| tRNA-His-GTG-2-2 | 2639.3994 | 0.37034757 | 0.19738347 | 1.87628458 | 0.06061621 |
| tRNA-His-GTG-2-1 | 2639.71589 | 0.37029765 | 0.19738242 | 1.87604166 | 0.06064956 |
| tRNA-His-GTG-2-3 | 2639.44369 | 0.370252 | 0.19736199 | 1.87600463 | 0.06065464 |
| tRNA-Ala-AGC-5-1 | 104.558718 | 0.34331541 | 0.18379775 | 1.8678978 | 0.06177632 |
| tRNA-Ala-AGC-10-1 | 102.068274 | 0.35709873 | 0.19141073 | 1.86561502 | 0.06209525 |
| tRNA-Leu-AAG-1-2 | 1183.40699 | -0.4026622 | 0.21729646 | -1.8530545 | 0.06387455 |
| tRNA-Leu-AAG-1-1 | 1180.37836 | -0.4013987 | 0.21750682 | -1.8454533 | 0.06497162 |
| tRNA-Ser-CGA-1-1 | 33.0755365 | -0.4733016 | 0.25992331 | -1.8209278 | 0.06861783 |
| tRNA-Lys-TTT-1-5 | 1595.08798 | 0.17030232 | 0.09533104 | 1.78643106 | 0.07402949 |
| tRNA-Tyr-GTA-6-1 | 31.2616758 | 0.66517219 | 0.37256783 | 1.78537203 | 0.07420099 |
| tRNA-Ser-CGA-3-1 | 24.9159965 | -0.5494008 | 0.30923064 | -1.7766699 | 0.07562256 |
| tRNA-Lys-TTT-1-4 | 1593.31727 | 0.16831068 | 0.09511318 | 1.76958318 | 0.0767966 |
| tRNA-Ile-TAT-2-3 | 608.596358 | 0.66734809 | 0.38780427 | 1.7208374 | 0.08528033 |
| tRNA-Gly-CCC-2-1 | 820.42568 | -0.6954496 | 0.41079443 | -1.6929382 | 0.09046724 |
| tRNA-Gly-CCC-2-2 | 820.42568 | -0.6954496 | 0.41079443 | -1.6929382 | 0.09046724 |
| tRNA-Val-CAC-4-1 | 1644.56413 | -0.7043051 | 0.41873337 | -1.6819895 | 0.09257088 |
| tRNA-Lys-TTT-2-1 | 260.054348 | 0.51838064 | 0.30944848 | 1.67517593 | 0.09389972 |
| tRNA-Lys-TTT-2-2 | 260.028571 | 0.51814291 | 0.30940012 | 1.67466938 | 0.09399912 |
| tRNA-Tyr-GTA-3-2 | 118.804814 | 0.47813819 | 0.28786054 | 1.66100637 | 0.09671217 |
| tRNA-iMet-CAT-1-2 | 755.756399 | -0.2865695 | 0.17341376 | -1.6525187 | 0.09842885 |
| tRNA-iMet-CAT-1-7 | 768.143741 | -0.284933 | 0.17286378 | -1.6483093 | 0.09928923 |
| tRNA-Ala-AGC-7-1 | 188.01954 | 0.39227667 | 0.24256476 | 1.61720386 | 0.10583428 |
| tRNA-Lys-TTT-5-1 | 390.253952 | 0.50228954 | 0.31195289 | 1.61014551 | 0.10736609 |
| tRNA-iMet-CAT-1-1 | 759.858903 | -0.2786411 | 0.17337788 | -1.607132 | 0.10802542 |
| tRNA-iMet-CAT-1-4 | 753.832149 | -0.2794584 | 0.17432703 | -1.6030701 | 0.10891918 |
| tRNA-Leu-TAA-1-1 | 135.154862 | -0.3733849 | 0.2333335 | -1.6002198 | 0.10954982 |
| tRNA-Leu-AAG-2-1 | 325.385179 | -0.3236706 | 0.20338616 | -1.591409 | 0.11151755 |
| tRNA-iMet-CAT-1-6 | 758.461958 | -0.2803839 | 0.1766387 | -1.5873298 | 0.11243797 |
| tRNA-iMet-CAT-1-5 | 764.082074 | -0.2809404 | 0.17907912 | -1.5688063 | 0.11669308 |
| tRNA-Lys-TTT-1-6 | 1616.48229 | 0.14910032 | 0.09584959 | 1.55556557 | 0.11981143 |
| tRNA-Pro-TGG-5-1 | 2.91571634 | 0.77762528 | 0.50034775 | 1.55416962 | 0.12014396 |
| tRNA-Lys-TTT-1-3 | 1617.83652 | 0.1486978 | 0.09591142 | 1.55036586 | 0.12105373 |
| tRNA-iMet-CAT-3-1 | 554.621585 | -0.3614653 | 0.23328111 | -1.5494838 | 0.12126546 |
| tRNA-Lys-CTT-6-1 | 30.8177099 | 0.54140634 | 0.35115273 | 1.54179732 | 0.12312285 |
| tRNA-Met-CAT-7-1 | 1.02342067 | 1.28799838 | 0.83588185 | 1.54088568 | 0.12334461 |
| tRNA-Met-CAT-4-1 | 525.197399 | 0.57251706 | 0.37166168 | 1.5404253 | 0.12345672 |
| tRNA-Ser-GCT-4-1 | 687.880024 | -0.381911 | 0.24812174 | -1.5392082 | 0.12375348 |
| tRNA-Ser-GCT-4-3 | 688.894853 | -0.3820442 | 0.24890669 | -1.5348891 | 0.12481108 |
| tRNA-Arg-ACG-1-3 | 47.1548304 | -0.3860061 | 0.25192469 | -1.5322281 | 0.12546615 |
| tRNA-Ser-GCT-2-1 | 683.493202 | -0.3795851 | 0.24778432 | -1.5319174 | 0.12554282 |
| tRNA-Gln-TTG-6-1 | 6.06567318 | 0.86122995 | 0.5635705 | 1.52816719 | 0.12647103 |
| tRNA-Met-CAT-5-1 | 229.59939 | 0.56740407 | 0.37370041 | 1.51833943 | 0.12892885 |

| | | | | | |
|---|---|---|---|---|---|
| tRNA-Ser-GCT-4-2 | 687.239026 | -0.3760606 | 0.24775119 | -1.5178964 | 0.12904052 |
| tRNA-Tyr-GTA-4-1 | 93.2562488 | 0.49783667 | 0.33146832 | 1.50191326 | 0.13311951 |
| tRNA-Glu-CTC-4-1 | 513.532653 | 0.2492443 | 0.16877229 | 1.47680819 | 0.13972706 |
| tRNA-Met-CAT-3-1 | 567.96738 | 0.52062621 | 0.35310359 | 1.47442909 | 0.1403661 |
| tRNA-Asp-GTC-3-1 | 98.8496926 | -0.442474 | 0.30075092 | -1.4712309 | 0.14122869 |
| tRNA-Tyr-GTA-3-1 | 92.1180321 | 0.48781539 | 0.33180233 | 1.47019882 | 0.14150792 |
| tRNA-Ser-GCT-1-1 | 674.298991 | -0.3669439 | 0.24996007 | -1.46801 | 0.14210151 |
| tRNA-Lys-TTT-3-1 | 10.0059366 | 0.6033944 | 0.41228572 | 1.46353456 | 0.14332117 |
| tRNA-Gly-GCC-1-1 | 11657.9354 | -0.4162445 | 0.28779636 | -1.446316 | 0.14808859 |
| tRNA-Gly-GCC-1-2 | 11657.9354 | -0.4162445 | 0.28779636 | -1.446316 | 0.14808859 |
| tRNA-Gly-GCC-1-3 | 11657.9354 | -0.4162445 | 0.28779636 | -1.446316 | 0.14808859 |
| tRNA-Val-TAC-1-1 | 4004.03633 | -0.3230186 | 0.22490423 | -1.4362498 | 0.15093128 |
| tRNA-Lys-CTT-4-1 | 5.05885163 | 0.71841489 | 0.50688143 | 1.41732335 | 0.15638841 |
| tRNA-Lys-TTT-1-1 | 1616.1898 | 0.14076547 | 0.09951893 | 1.41445933 | 0.15722708 |
| tRNA-Leu-AAG-3-1 | 316.871544 | -0.2914198 | 0.20624986 | -1.4129453 | 0.15767182 |
| tRNA-Lys-CTT-10-1 | 34.5803867 | 0.47144028 | 0.33468296 | 1.4086175 | 0.1589483 |
| tRNA-Val-AAC-4-1 | 623.309116 | -0.3883238 | 0.27782714 | -1.3977171 | 0.16219802 |
| tRNA-Val-TAC-1-3 | 3958.29919 | -0.3151333 | 0.22623767 | -1.3929302 | 0.16364089 |
| tRNA-Val-TAC-1-2 | 3957.35601 | -0.314987 | 0.22618762 | -1.3925918 | 0.16374327 |
| tRNA-Tyr-GTA-1-2 | 40.6786852 | 0.50634944 | 0.36415992 | 1.39045904 | 0.16438953 |
| tRNA-Tyr-GTA-1-1 | 40.9936275 | 0.48420659 | 0.35495545 | 1.36413344 | 0.17252557 |
| tRNA-Leu-CAG-3-1 | 107.80187 | 0.21603376 | 0.16059615 | 1.34519887 | 0.17856102 |
| tRNA-Tyr-GTA-1-3 | 40.5853247 | 0.4869385 | 0.36347746 | 1.33966629 | 0.18035386 |
| tRNA-Arg-ACG-2-1 | 48.53316 | 0.40479169 | 0.30279563 | 1.33684789 | 0.18127229 |
| tRNA-His-GTG-3-1 | 137.817213 | 0.42832205 | 0.32277056 | 1.3270171 | 0.18450303 |
| tRNA-Tyr-GTA-1-4 | 46.4347138 | 0.43758238 | 0.33251313 | 1.31598525 | 0.18817899 |
| tRNA-Tyr-GTA-5-1 | 29.4192732 | 0.55878183 | 0.42720721 | 1.30798785 | 0.19087743 |
| tRNA-Tyr-GTA-1-5 | 41.3530433 | 0.46751875 | 0.35786686 | 1.30640416 | 0.19141515 |
| tRNA-Ser-GCT-5-1 | 25.0039612 | -0.4292065 | 0.32973662 | -1.3016647 | 0.19303103 |
| tRNA-Ile-AAT-1-8 | 96.5312681 | -0.2669022 | 0.2054736 | -1.2989612 | 0.19395723 |
| tRNA-Gln-TTG-2-1 | 181.064903 | 0.44159734 | 0.35136111 | 1.25681906 | 0.20881916 |
| tRNA-Ala-CGC-3-1 | 1.68396984 | -0.7741078 | 0.62153021 | -1.245487 | 0.21295281 |
| tRNA-Ala-CGC-3-2 | 1.68396984 | -0.7741078 | 0.62153021 | -1.245487 | 0.21295281 |
| tRNA-Arg-ACG-1-1 | 44.8410388 | -0.3204305 | 0.25778499 | -1.2430147 | 0.21386241 |
| tRNA-Ile-AAT-1-6 | 92.3816493 | -0.2535926 | 0.20410063 | -1.2424883 | 0.21405646 |
| tRNA-Val-CAC-1-1 | 1244.14605 | -0.2455377 | 0.19789302 | -1.2407598 | 0.2146945 |
| tRNA-Arg-ACG-1-2 | 44.1971233 | -0.3246914 | 0.26189559 | -1.2397743 | 0.21505888 |
| tRNA-Ala-CGC-1-2 | 116.392569 | -0.1953114 | 0.16084865 | -1.2142558 | 0.22465006 |
| tRNA-Ile-AAT-1-4 | 90.2458948 | -0.2540924 | 0.2095256 | -1.2127032 | 0.22524332 |
| tRNA-Ser-GCT-3-1 | 781.366053 | -0.2575419 | 0.21409114 | -1.2029544 | 0.22899398 |
| tRNA-Ile-AAT-1-1 | 91.5170185 | -0.2540696 | 0.2112692 | -1.2025871 | 0.22913616 |
| tRNA-Met-CAT-6-1 | 55.5168529 | 0.49911121 | 0.42237195 | 1.18168644 | 0.23733014 |
| tRNA-Ala-CGC-4-1 | 1.29620528 | -1.099632 | 0.9322918 | -1.1794934 | 0.23820178 |
| tRNA-Leu-TAG-2-1 | 326.077334 | -0.2306311 | 0.19709683 | -1.1701412 | 0.24194417 |
| tRNA-Ile-AAT-1-7 | 90.8522704 | -0.2439756 | 0.20873371 | -1.1688364 | 0.24246957 |
| tRNA-Asn-GTT-5-1 | 0.17482827 | -2.0836773 | 1.78866699 | -1.1649331 | 0.24404608 |

| | | | | | |
|---|---|---|---|---|---|
| tRNA-Ile-AAT-3-1 | 88.073168 | -0.241954 | 0.20938084 | -1.1555688 | 0.24785758 |
| tRNA-Val-AAC-1-1 | 10991.5776 | -0.2833911 | 0.24598447 | -1.1520691 | 0.24929267 |
| tRNA-Val-AAC-1-2 | 10985.5419 | -0.2830252 | 0.24619811 | -1.1495832 | 0.25031559 |
| tRNA-Pro-TGG-4-1 | 30.8485027 | 0.22636686 | 0.20197728 | 1.12075411 | 0.26239254 |
| tRNA-Ala-CGC-3-3 | 1.71675613 | -0.6871984 | 0.61351569 | -1.1200992 | 0.26267151 |
| tRNA-Val-CAC-2-1 | 12190.4202 | -0.2756293 | 0.2464566 | -1.1183685 | 0.26340962 |
| tRNA-Val-CAC-2-5 | 12188.6419 | -0.2755132 | 0.24647896 | -1.1177958 | 0.2636542 |
| tRNA-Ile-AAT-1-2 | 89.5214162 | -0.2364283 | 0.21157843 | -1.1174498 | 0.26380204 |
| tRNA-Val-CAC-2-3 | 12191.1504 | -0.2750776 | 0.24654675 | -1.1157217 | 0.26454127 |
| tRNA-Ile-AAT-1-3 | 90.3447892 | -0.2347745 | 0.21062228 | -1.1146706 | 0.26499161 |
| tRNA-Ile-AAT-1-5 | 89.9269286 | -0.2345894 | 0.21093133 | -1.1121601 | 0.26606933 |
| tRNA-Val-CAC-2-2 | 12359.9719 | -0.27517 | 0.24806178 | -1.1092802 | 0.2673093 |
| tRNA-Gln-TTG-1-1 | 537.272293 | 0.23747441 | 0.21423367 | 1.10848318 | 0.2676532 |
| tRNA-Val-CAC-2-4 | 12316.6994 | -0.2741431 | 0.24772822 | -1.1066285 | 0.26845459 |
| tRNA-Lys-CTT-15-1 | 35.1268032 | 0.41576498 | 0.37987337 | 1.09448308 | 0.27374316 |
| tRNA-Gly-ACC-1-1 | 268.591881 | -0.2347825 | 0.21497236 | -1.0921522 | 0.27476623 |
| tRNA-Val-CAC-3-1 | 398.429743 | -0.2451897 | 0.22580237 | -1.0858598 | 0.27754102 |
| tRNA-Val-AAC-3-1 | 356.484735 | -0.2459065 | 0.22734071 | -1.081665 | 0.27940139 |
| tRNA-Ile-AAT-2-1 | 46.7551024 | -0.238876 | 0.22100432 | -1.0808656 | 0.2797569 |
| tRNA-Ile-AAT-4-1 | 45.104543 | -0.242529 | 0.22545083 | -1.0757511 | 0.28203859 |
| tRNA-Cys-GCA-26-1 | 6.4366475 | 0.40669119 | 0.37860187 | 1.07419224 | 0.28273653 |
| tRNA-Cys-GCA-23-1 | 6.34601857 | 0.40412043 | 0.37842467 | 1.06790192 | 0.28556475 |
| tRNA-Arg-TCG-1-1 | 46.17519 | 0.31650718 | 0.312216 | 1.01374427 | 0.3107048 |
| tRNA-Arg-CCG-3-1 | 462.718991 | -0.1515401 | 0.15112338 | -1.0027577 | 0.31597776 |
| tRNA-Gly-TCC-1-6 | 8470.4764 | -0.2689737 | 0.27319677 | -0.984542 | 0.3248491 |
| tRNA-Gly-TCC-2-1 | 8324.63931 | -0.2735072 | 0.2781374 | -0.9833527 | 0.3254339 |
| tRNA-Leu-TAA-3-1 | 1.7372142 | -0.6652207 | 0.68051148 | -0.9775305 | 0.32830661 |
| tRNA-Arg-TCT-5-1 | 2.13340709 | -0.6253062 | 0.63981572 | -0.9773224 | 0.32840956 |
| tRNA-Val-AAC-5-1 | 3101.89065 | -0.2435493 | 0.25006037 | -0.9739618 | 0.33007548 |
| tRNA-Gly-TCC-1-5 | 8427.80954 | -0.2655289 | 0.27414326 | -0.9685771 | 0.33275622 |
| tRNA-Gly-TCC-1-1 | 8427.84377 | -0.2655154 | 0.27418166 | -0.9683923 | 0.33284848 |
| tRNA-Gly-TCC-1-7 | 8429.07515 | -0.2648671 | 0.27406379 | -0.9664431 | 0.33382249 |
| tRNA-Gly-TCC-1-3 | 8436.42661 | -0.2645993 | 0.27411221 | -0.9652955 | 0.33439684 |
| tRNA-Gly-TCC-1-4 | 8436.42661 | -0.2645993 | 0.27411221 | -0.9652955 | 0.33439684 |
| tRNA-Gly-TCC-1-2 | 8437.68952 | -0.2644414 | 0.27408212 | -0.9648253 | 0.33463233 |
| tRNA-Ala-AGC-8-1 | 33.9796352 | 0.1986795 | 0.20611787 | 0.96391206 | 0.33509002 |
| tRNA-Ala-CGC-6-1 | 1.67948704 | -0.5739172 | 0.6099308 | -0.9409547 | 0.34672809 |
| tRNA-Gln-CTG-3-3 | 647.014357 | 0.23602409 | 0.25356114 | 0.93083702 | 0.35193788 |
| tRNA-Gln-CTG-3-2 | 646.988581 | 0.23591478 | 0.2535322 | 0.93051212 | 0.35210599 |
| tRNA-Gln-CTG-3-1 | 647.644819 | 0.23500376 | 0.25379314 | 0.92596576 | 0.35446378 |
| tRNA-Phe-GAA-3-1 | 6.6287751 | 0.31057541 | 0.33779512 | 0.91941947 | 0.35787621 |
| tRNA-Thr-CGT-2-1 | 141.318142 | 0.25967713 | 0.28575539 | 0.90873925 | 0.36348778 |
| tRNA-Thr-AGT-6-1 | 9.6480043 | 0.37283174 | 0.41139991 | 0.90625139 | 0.36480281 |
| tRNA-Cys-GCA-20-1 | 6.70514239 | 0.32816164 | 0.36673644 | 0.89481601 | 0.37088545 |
| tRNA-Gln-CTG-5-1 | 408.68738 | 0.22079443 | 0.24894713 | 0.88691295 | 0.37512577 |
| tRNA-Gln-CTG-4-1 | 427.478763 | 0.20279524 | 0.23083552 | 0.87852699 | 0.3796578 |

| | | | | | |
|---|---|---|---|---|---|
| tRNA-Lys-CTT-16-1 | 42.3294582 | 0.3186448 | 0.36286409 | 0.87813814 | 0.37986876 |
| tRNA-Phe-GAA-1-1 | 21.7744152 | -0.2049424 | 0.23559506 | -0.8698925 | 0.38435913 |
| tRNA-Met-CAT-1-1 | 856.132286 | 0.28262265 | 0.32723857 | 0.86365934 | 0.38777506 |
| tRNA-Met-CAT-2-2 | 708.579895 | 0.25485653 | 0.3001144 | 0.84919794 | 0.39577116 |
| tRNA-Cys-GCA-21-1 | 4.97286915 | 0.37555592 | 0.4444095 | 0.84506726 | 0.39807329 |
| tRNA-Met-CAT-1-2 | 864.131819 | 0.27242226 | 0.32407714 | 0.84060931 | 0.40056684 |
| tRNA-Met-CAT-2-1 | 709.328002 | 0.2518121 | 0.29979534 | 0.83994668 | 0.40093828 |
| tRNA-Ser-GCT-6-1 | 142.025837 | 0.20861802 | 0.2489273 | 0.83806805 | 0.40199248 |
| tRNA-Ile-TAT-2-1 | 58.6285663 | 0.29534804 | 0.35665036 | 0.82811647 | 0.40760454 |
| tRNA-Lys-CTT-5-1 | 1.95586509 | 0.50881284 | 0.62633194 | 0.8123693 | 0.41657976 |
| tRNA-Gly-GCC-6-1 | 6414.75325 | -0.248553 | 0.30618068 | -0.8117853 | 0.41691484 |
| tRNA-Cys-GCA-12-1 | 22.9398808 | -0.2085497 | 0.25987659 | -0.8024953 | 0.42226649 |
| tRNA-Ile-TAT-2-2 | 55.5011523 | 0.2755354 | 0.3443479 | 0.80016575 | 0.42361477 |
| tRNA-Gln-CTG-2-4 | 669.533274 | 0.19260072 | 0.2415824 | 0.79724651 | 0.42530788 |
| tRNA-Gln-CTG-2-3 | 670.8802 | 0.19036796 | 0.24040018 | 0.79187944 | 0.42843097 |
| tRNA-Ala-CGC-2-1 | 63.7271322 | -0.1296169 | 0.16383049 | -0.7911646 | 0.42884794 |
| tRNA-Gln-CTG-2-2 | 670.705346 | 0.19062037 | 0.24166804 | 0.78876949 | 0.43024675 |
| tRNA-Gln-CTG-2-1 | 671.385995 | 0.18896585 | 0.24038802 | 0.78608679 | 0.43181664 |
| tRNA-Cys-GCA-14-1 | 23.0653886 | -0.2017182 | 0.2593918 | -0.7776585 | 0.43677035 |
| tRNA-Trp-CCA-6-1 | 27.4860182 | 0.2593134 | 0.33531193 | 0.77334976 | 0.43931541 |
| tRNA-Cys-GCA-24-1 | 23.7064836 | -0.1968888 | 0.25935324 | -0.759153 | 0.44776102 |
| tRNA-Arg-TCT-4-1 | 119.538597 | -0.17314 | 0.23686429 | -0.7309671 | 0.46479925 |
| tRNA-Glu-CTC-5-1 | 306.293024 | 0.10687891 | 0.1479726 | 0.72228853 | 0.4701171 |
| tRNA-Gln-CTG-6-1 | 244.578078 | 0.27677569 | 0.38996516 | 0.70974466 | 0.47786249 |
| tRNA-Val-AAC-6-1 | 0.14955752 | -1.8171713 | 2.60360582 | -0.6979441 | 0.48521217 |
| tRNA-Gln-CTG-1-1 | 270.415905 | 0.23989226 | 0.35300175 | 0.67957811 | 0.49677163 |
| tRNA-Leu-CAA-2-1 | 687.406064 | -0.1973474 | 0.29186822 | -0.6761526 | 0.49894378 |
| tRNA-Cys-GCA-19-1 | 4.03759544 | 0.33433324 | 0.50246444 | 0.66538686 | 0.50580309 |
| tRNA-Gln-CTG-7-1 | 5.23553679 | -0.2966796 | 0.45260126 | -0.6554988 | 0.51214668 |
| tRNA-Gln-TTG-3-2 | 515.893303 | 0.1265297 | 0.19420475 | 0.6515273 | 0.51470616 |
| tRNA-Ala-AGC-2-2 | 100.891715 | -0.1092612 | 0.16941163 | -0.6449448 | 0.51896293 |
| tRNA-Gln-TTG-3-1 | 516.663899 | 0.12453231 | 0.19429256 | 0.64095256 | 0.52155351 |
| tRNA-Cys-GCA-7-1 | 90.0915109 | -0.1098676 | 0.17240853 | -0.6372518 | 0.52396086 |
| tRNA-iMet-CAT-4-1 | 0.35461962 | -0.8060105 | 1.28072206 | -0.6293407 | 0.52912603 |
| tRNA-Leu-CAA-1-1 | 129.647886 | 0.09343639 | 0.14917889 | 0.62633789 | 0.53109334 |
| tRNA-Lys-CTT-8-1 | 1.95324998 | 0.37898592 | 0.60708967 | 0.62426678 | 0.53245239 |
| tRNA-Ala-AGC-3-1 | 53.2997226 | 0.12274824 | 0.19858336 | 0.61811945 | 0.5364966 |
| tRNA-Trp-CCA-2-1 | 1130.32866 | -0.1350739 | 0.22148791 | -0.6098479 | 0.54196258 |
| tRNA-Ile-AAT-5-1 | 0.32971074 | -0.6719503 | 1.10986972 | -0.6054317 | 0.5448922 |
| tRNA-Pro-AGG-1-2 | 137.980898 | 0.10958075 | 0.18290016 | 0.59912875 | 0.54908703 |
| tRNA-Pro-AGG-1-5 | 140.190231 | 0.106728 | 0.18059319 | 0.59098576 | 0.55452996 |
| tRNA-Ile-TAT-1-1 | 119.993876 | 0.17625305 | 0.30478654 | 0.57828358 | 0.56307268 |
| tRNA-Arg-TCT-6-1 | 0.23128924 | -0.7902808 | 1.36665032 | -0.5782611 | 0.56308783 |
| tRNA-Pro-AGG-1-4 | 141.948205 | 0.09667165 | 0.17364525 | 0.55671922 | 0.57771929 |
| tRNA-Leu-TAG-4-1 | 1.03174153 | 0.38308204 | 0.70109823 | 0.54640281 | 0.58478908 |
| tRNA-Gln-TTG-4-1 | 0.94416264 | -0.4962083 | 0.91166842 | -0.5442859 | 0.58624476 |

| | | | | | |
|---|---|---|---|---|---|
| tRNA-Ala-AGC-2-1 | 97.7360584 | -0.092865 | 0.17843829 | -0.5204323 | 0.60276233 |
| tRNA-Pro-AGG-1-1 | 148.227586 | 0.08905121 | 0.17224307 | 0.51700899 | 0.60514988 |
| tRNA-Arg-TCG-4-1 | 79.0280441 | -0.2369872 | 0.46462002 | -0.5100666 | 0.61000477 |
| tRNA-Pro-AGG-1-6 | 148.708084 | 0.08625778 | 0.16979634 | 0.50800732 | 0.61144821 |
| tRNA-Ala-CGC-1-1 | 103.114946 | -0.0784027 | 0.15502439 | -0.5057443 | 0.61303615 |
| tRNA-Pro-AGG-1-3 | 139.561451 | 0.09037934 | 0.17871027 | 0.50573112 | 0.61304542 |
| tRNA-Cys-GCA-17-1 | 9.25083918 | 0.16353295 | 0.32358671 | 0.50537597 | 0.61329479 |
| tRNA-Cys-GCA-9-1 | 24.558822 | -0.1254551 | 0.25013503 | -0.5015494 | 0.61598449 |
| tRNA-Cys-GCA-8-1 | 71.2757221 | -0.1060479 | 0.21160332 | -0.5011636 | 0.61625598 |
| tRNA-Ile-GAT-1-1 | 0.84984571 | -0.4215544 | 0.85125688 | -0.4952141 | 0.62044897 |
| tRNA-Pro-TGG-1-1 | 145.029843 | 0.07758996 | 0.15887246 | 0.4883789 | 0.62528148 |
| tRNA-Phe-GAA-1-5 | 17.4498973 | 0.11818907 | 0.2436291 | 0.48511886 | 0.62759203 |
| tRNA-Glu-TTC-2-1 | 17708.2457 | -0.1107356 | 0.23005731 | -0.4813391 | 0.63027555 |
| tRNA-Ala-AGC-18-1 | 0.34615895 | 0.71777192 | 1.49155436 | 0.48122411 | 0.63035723 |
| tRNA-Glu-TTC-2-2 | 17707.9053 | -0.1100845 | 0.22981949 | -0.479004 | 0.63193577 |
| tRNA-Thr-AGT-2-1 | 127.055792 | -0.1582242 | 0.33037668 | -0.4789207 | 0.63199506 |
| tRNA-Leu-TAA-4-1 | 3.40314102 | -0.3000917 | 0.63458647 | -0.4728933 | 0.63628929 |
| tRNA-Pro-TGG-2-1 | 160.097183 | 0.09058472 | 0.1945978 | 0.46549713 | 0.6415755 |
| tRNA-Leu-CAG-4-1 | 86.3468871 | -0.1119395 | 0.24130892 | -0.4638846 | 0.64273042 |
| tRNA-Pro-TGG-2-4 | 160.12548 | 0.08818184 | 0.19254439 | 0.45798189 | 0.64696545 |
| tRNA-Cys-GCA-22-1 | 0.90140183 | -1.0879981 | 2.42648583 | -0.4483843 | 0.65387588 |
| tRNA-Ala-TGC-4-1 | 71.0283377 | -0.072996 | 0.1636134 | -0.4461492 | 0.6554895 |
| tRNA-Cys-GCA-4-3 | 27.3561618 | -0.0951248 | 0.2170635 | -0.4382347 | 0.66121615 |
| tRNA-Cys-GCA-4-5 | 27.3561618 | -0.0951248 | 0.2170635 | -0.4382347 | 0.66121615 |
| tRNA-Cys-GCA-4-6 | 27.3561618 | -0.0951248 | 0.2170635 | -0.4382347 | 0.66121615 |
| tRNA-Cys-GCA-4-7 | 27.3561618 | -0.0951248 | 0.2170635 | -0.4382347 | 0.66121615 |
| tRNA-Cys-GCA-4-14 | 27.3561618 | -0.0951248 | 0.2170635 | -0.4382347 | 0.66121615 |
| tRNA-Cys-GCA-4-15 | 27.3561618 | -0.0951248 | 0.2170635 | -0.4382347 | 0.66121615 |
| tRNA-Cys-GCA-25-1 | 7.54429708 | 0.17968093 | 0.41194247 | 0.43617967 | 0.66270638 |
| tRNA-Phe-GAA-1-3 | 17.490407 | 0.10278587 | 0.23713363 | 0.43345125 | 0.66468698 |
| tRNA-Ile-TAT-3-1 | 0.85000856 | 0.41124978 | 0.95269155 | 0.43167149 | 0.66598019 |
| tRNA-Pro-TGG-2-3 | 159.84959 | 0.08041726 | 0.1922014 | 0.41840103 | 0.67565394 |
| tRNA-Lys-CTT-19-1 | 0.09345634 | -1.0021948 | 2.40395984 | -0.4168933 | 0.67675643 |
| tRNA-Tyr-GTA-2-1 | 109.691262 | 0.11019939 | 0.26493679 | 0.41594597 | 0.67744954 |
| tRNA-Trp-CCA-3-1 | 63.163821 | -0.1273626 | 0.30711915 | -0.4147009 | 0.67836086 |
| tRNA-Cys-GCA-4-16 | 27.3211086 | -0.0895832 | 0.21734444 | -0.4121716 | 0.68021364 |
| tRNA-Cys-GCA-6-1 | 27.3211086 | -0.0895832 | 0.21734444 | -0.4121716 | 0.68021364 |
| tRNA-Cys-GCA-4-18 | 27.3211086 | -0.0895832 | 0.21734444 | -0.4121716 | 0.68021364 |
| tRNA-Cys-GCA-4-19 | 27.3211086 | -0.0895832 | 0.21734444 | -0.4121716 | 0.68021364 |
| tRNA-Cys-GCA-4-20 | 27.3211086 | -0.0895832 | 0.21734444 | -0.4121716 | 0.68021364 |
| tRNA-Cys-GCA-4-24 | 27.3211086 | -0.0895832 | 0.21734444 | -0.4121716 | 0.68021364 |
| tRNA-Cys-GCA-4-26 | 27.3211086 | -0.0895832 | 0.21734444 | -0.4121716 | 0.68021364 |
| tRNA-Cys-GCA-4-1 | 27.2942713 | -0.0871343 | 0.21765721 | -0.4003282 | 0.68891478 |
| tRNA-Cys-GCA-4-17 | 27.2942713 | -0.0871343 | 0.21765721 | -0.4003282 | 0.68891478 |
| tRNA-Cys-GCA-4-21 | 27.2942713 | -0.0871343 | 0.21765721 | -0.4003282 | 0.68891478 |
| tRNA-Cys-GCA-10-1 | 27.2942713 | -0.0871343 | 0.21765721 | -0.4003282 | 0.68891478 |

| | | | | | |
|---|---|---|---|---|---|
| tRNA-Cys-GCA-4-27 | 27.2942713 | -0.0871343 | 0.21765721 | -0.4003282 | 0.68891478 |
| tRNA-Cys-GCA-4-29 | 27.2942713 | -0.0871343 | 0.21765721 | -0.4003282 | 0.68891478 |
| tRNA-Pro-CGG-1-3 | 148.660721 | 0.07602676 | 0.19041687 | 0.39926482 | 0.68969809 |
| tRNA-Lys-CTT-14-1 | 25.7287681 | -0.1700731 | 0.43277906 | -0.3929791 | 0.69433492 |
| tRNA-Ala-AGC-1-1 | 32.0668851 | -0.1254062 | 0.32200803 | -0.3894506 | 0.69694281 |
| tRNA-Cys-GCA-18-1 | 0.38099261 | -0.414638 | 1.10453598 | -0.3753957 | 0.70736621 |
| tRNA-Cys-GCA-4-2 | 27.2592182 | -0.0815628 | 0.21792109 | -0.3742766 | 0.70819857 |
| tRNA-Cys-GCA-4-4 | 27.2592182 | -0.0815628 | 0.21792109 | -0.3742766 | 0.70819857 |
| tRNA-Cys-GCA-4-8 | 27.2592182 | -0.0815628 | 0.21792109 | -0.3742766 | 0.70819857 |
| tRNA-Cys-GCA-4-10 | 27.2592182 | -0.0815628 | 0.21792109 | -0.3742766 | 0.70819857 |
| tRNA-Cys-GCA-4-11 | 27.2592182 | -0.0815628 | 0.21792109 | -0.3742766 | 0.70819857 |
| tRNA-Cys-GCA-4-22 | 27.2592182 | -0.0815628 | 0.21792109 | -0.3742766 | 0.70819857 |
| tRNA-Cys-GCA-4-23 | 27.2592182 | -0.0815628 | 0.21792109 | -0.3742766 | 0.70819857 |
| tRNA-Cys-GCA-4-25 | 27.2592182 | -0.0815628 | 0.21792109 | -0.3742766 | 0.70819857 |
| tRNA-Cys-GCA-4-28 | 27.2592182 | -0.0815628 | 0.21792109 | -0.3742766 | 0.70819857 |
| tRNA-Leu-CAA-4-1 | 146.830083 | 0.05514234 | 0.14956913 | 0.36867462 | 0.71237027 |
| tRNA-Ala-TGC-1-1 | 40.8775282 | 0.08162328 | 0.22171682 | 0.36814202 | 0.71276734 |
| tRNA-Arg-TCT-3-1 | 64.3186263 | 0.15396594 | 0.43177745 | 0.35658634 | 0.7214015 |
| tRNA-Lys-CTT-13-1 | 142.743604 | 0.16243792 | 0.4672794 | 0.34762481 | 0.72812197 |
| tRNA-Lys-CTT-18-1 | 141.421944 | 0.16207723 | 0.46727646 | 0.3468551 | 0.72870018 |
| tRNA-Trp-CCA-3-2 | 63.6846692 | -0.1081167 | 0.31214937 | -0.3463622 | 0.72907052 |
| tRNA-iMet-CAT-2-1 | 305.937754 | -0.0847863 | 0.24628903 | -0.3442554 | 0.73065426 |
| tRNA-Ala-TGC-3-1 | 105.436218 | -0.051403 | 0.14983088 | -0.3430735 | 0.73154313 |
| tRNA-Thr-CGT-1-1 | 126.630511 | -0.1101409 | 0.3270787 | -0.3367414 | 0.73631185 |
| tRNA-Cys-GCA-4-9 | 27.921463 | -0.0720858 | 0.21955651 | -0.3283244 | 0.74266639 |
| tRNA-Cys-GCA-4-12 | 27.921463 | -0.0720858 | 0.21955651 | -0.3283244 | 0.74266639 |
| tRNA-Cys-GCA-4-13 | 27.921463 | -0.0720858 | 0.21955651 | -0.3283244 | 0.74266639 |
| tRNA-Ser-GGA-1-1 | 1.81903355 | -0.2052465 | 0.62627707 | -0.3277248 | 0.74311978 |
| tRNA-Lys-CTT-7-1 | 3.86424646 | 0.14216748 | 0.44268226 | 0.32115017 | 0.74809659 |
| tRNA-Leu-CAG-2-3 | 1499.43464 | -0.0975589 | 0.30830677 | -0.3164345 | 0.75167273 |
| tRNA-Sup-TTA-1-1 | 0.06909621 | -0.8229001 | 2.60691108 | -0.315661 | 0.75225986 |
| tRNA-Cys-GCA-16-1 | 9.62659655 | -0.1031235 | 0.33335472 | -0.3093506 | 0.75705488 |
| tRNA-Thr-AGT-3-1 | 118.120756 | 0.11431378 | 0.37008809 | 0.30888263 | 0.75741081 |
| tRNA-Pro-TGG-2-2 | 164.787758 | 0.05866414 | 0.18997811 | 0.30879424 | 0.75747805 |
| tRNA-Glu-TTC-1-1 | 13964.2959 | -0.0713322 | 0.23395724 | -0.3048944 | 0.76044661 |
| tRNA-Glu-TTC-1-2 | 13956.6879 | -0.070647 | 0.23400599 | -0.3019027 | 0.76272626 |
| tRNA-Glu-TTC-1-4 | 14000.2353 | -0.0705691 | 0.23431296 | -0.3011747 | 0.76328131 |
| tRNA-Thr-AGT-1-1 | 126.364583 | 0.102131 | 0.35050188 | 0.29138504 | 0.77075686 |
| tRNA-Lys-CTT-12-1 | 0.8830716 | -0.2628352 | 0.92652511 | -0.2836784 | 0.77665684 |
| tRNA-Thr-AGT-1-2 | 126.153635 | 0.09960252 | 0.3514951 | 0.28336815 | 0.77689464 |
| tRNA-Thr-AGT-1-3 | 126.121834 | 0.09917649 | 0.35108875 | 0.28248269 | 0.77757342 |
| tRNA-Glu-TTC-1-3 | 13862.6222 | -0.0627895 | 0.23404904 | -0.268275 | 0.78848762 |
| tRNA-Thr-AGT-4-1 | 69.5575716 | -0.1176721 | 0.45888378 | -0.2564311 | 0.79761798 |
| tRNA-Leu-CAG-2-1 | 1491.51261 | -0.0767792 | 0.31357287 | -0.2448527 | 0.80657052 |
| tRNA-Arg-CCG-2-1 | 61.7999239 | -0.0451908 | 0.21042673 | -0.2147579 | 0.82995607 |
| tRNA-Leu-CAG-2-2 | 1458.90415 | -0.0676467 | 0.32695489 | -0.2068992 | 0.8360886 |

| | | | | | |
|---|---|---|---|---|---|
| tRNA-Phe-GAA-1-4 | 18.1511433 | 0.04529488 | 0.2317021 | 0.19548756 | 0.84501128 |
| tRNA-Leu-CAG-1-4 | 1451.97836 | -0.0626573 | 0.3296809 | -0.1900542 | 0.84926663 |
| tRNA-Pro-CGG-1-1 | 157.201313 | 0.03369509 | 0.18129695 | 0.1858558 | 0.85255785 |
| tRNA-Leu-CAG-1-1 | 1446.89069 | -0.0578549 | 0.33203087 | -0.1742456 | 0.86167242 |
| tRNA-Leu-CAG-1-2 | 1446.89069 | -0.0578549 | 0.33203087 | -0.1742456 | 0.86167242 |
| tRNA-Leu-CAG-1-3 | 1446.89069 | -0.0578549 | 0.33203087 | -0.1742456 | 0.86167242 |
| tRNA-Leu-CAG-1-5 | 1447.75934 | -0.0577921 | 0.33204641 | -0.1740482 | 0.86182756 |
| tRNA-Val-CAC-7-1 | 5.74263881 | -0.0723091 | 0.42919722 | -0.1684751 | 0.86620951 |
| tRNA-Leu-CAA-3-1 | 527.463631 | -0.043959 | 0.2614002 | -0.1681672 | 0.86645171 |
| tRNA-Ser-AGA-5-1 | 0.24961486 | 0.21502803 | 1.489317 | 0.1443803 | 0.88520017 |
| tRNA-Pro-CGG-1-2 | 156.934586 | 0.02349777 | 0.18598683 | 0.12634104 | 0.89946197 |
| tRNA-Val-CAC-5-1 | 2.85047491 | -0.0609675 | 0.51990615 | -0.1172664 | 0.90664899 |
| tRNA-Ala-TGC-7-1 | 27.2004571 | -0.0264614 | 0.23454317 | -0.1128211 | 0.91017242 |
| tRNA-Ala-TGC-7-2 | 27.2004571 | -0.0264614 | 0.23454317 | -0.1128211 | 0.91017242 |
| tRNA-Ser-CGA-2-1 | 316.280555 | -0.0265846 | 0.24269214 | -0.1095403 | 0.91277395 |
| tRNA-Cys-GCA-15-1 | 3.32259112 | -0.0480175 | 0.49201756 | -0.0975931 | 0.92225538 |
| tRNA-Trp-CCA-1-1 | 99.7361397 | -0.0223143 | 0.25775617 | -0.0865715 | 0.93101211 |
| tRNA-Cys-GCA-28-1 | 0.42959396 | 0.09069526 | 1.05408718 | 0.08604151 | 0.93143342 |
| tRNA-Ala-TGC-8-1 | 25.8641254 | -0.0210906 | 0.24548069 | -0.0859157 | 0.93153343 |
| tRNA-Phe-GAA-1-2 | 19.7601617 | -0.0190103 | 0.25656423 | -0.0740956 | 0.94093434 |
| tRNA-Cys-GCA-27-1 | 0.07484252 | -0.161877 | 2.55099888 | -0.0634563 | 0.94940314 |
| tRNA-Val-CAC-6-1 | 265.800319 | -0.0246232 | 0.39857905 | -0.0617776 | 0.95073998 |
| tRNA-Arg-TCG-3-1 | 43.8596876 | -0.0213441 | 0.37068714 | -0.0575797 | 0.9540834 |
| tRNA-Glu-CTC-6-1 | 2.42198782 | 0.03166303 | 0.56114582 | 0.05642567 | 0.95500271 |
| tRNA-Lys-CTT-9-1 | 0.10061908 | 0.07435459 | 2.39371401 | 0.03106244 | 0.97521975 |
| tRNA-Trp-CCA-4-1 | 50.4921654 | -0.0059935 | 0.35248541 | -0.0170035 | 0.98643381 |
| tRNA-Trp-CCA-4-2 | 50.4921654 | -0.0059935 | 0.35248541 | -0.0170035 | 0.98643381 |
| tRNA-Arg-TCT-1-1 | 101.225319 | 0.00425087 | 0.26983211 | 0.01575377 | 0.98743083 |
| tRNA-Arg-TCG-3-2 | 43.622843 | -0.0017506 | 0.37139556 | -0.0047137 | 0.99623906 |

dup_pvalue
padj
0.00257963
0.00257963
0.00257963
0.00528738
0.00528738
0.00528738
0.00528738
0.00528738
0.00528738
0.00528738
0.00528738
0.00528738
0.00528738
0.00528738
0.00528738
0.00528738
0.00528738
0.00528738
0.00528738
0.00528738
0.00528738
0.00528738
0.00528738
NA
0.00528738
0.00528738
0.00528738
0.00528738
0.00528738
0.00528738
0.00528738
0.00528738
0.00528738
0.00536338
0.00536338
0.00536411
NA
NA
0.00862074
0.00862074
0.00862074
NA
NA
0.01002967

NA
0.0169993
NA
0.0172661
NA
0.01948507
0.01985623
0.0204936
0.02129824
0.02361204
0.02741804
0.0308933
0.03151519
0.03151519
0.03881291
0.04026322
0.04026322
0.04026322
0.04026322
0.04026322
NA
0.04122765
0.04122765
0.04128347
0.04314839
0.04401819
NA
0.04957608
0.04957608
0.04957608
0.04957608
0.04957608
0.04957608
0.04957608
0.04957608
0.04957608
0.0577015
0.06277502
0.06277502
0.06415842
0.0650095
0.06617163
0.06660805
0.06826852
0.07184209
0.0720719

NA
0.08450061
0.08450061
0.08615214
0.08615214
NA
NA
NA
NA
NA
NA
0.0982732
NA
NA
NA
NA
0.11769487
0.11769487
0.11883806
NA
NA
0.12248939
0.12248939
0.12248939
0.12248939
0.12248939
0.12248939
0.12248939
0.12248939
0.12248939
0.12248939
0.12248939
0.12248939
0.12248939
0.12248939
0.12248939
0.12248939
0.12527904
0.12527904
0.12791251
0.12853316
0.12853316
NA
0.13761978
0.13761978
0.13761978

0.13761978
0.13761978
0.13761978
0.13761978
0.13761978
0.13761978
0.13761978
0.13855964
0.13855964
0.14136171
0.14262064
NA
0.16119324
NA
NA
0.16588066
0.18274357
0.19082933
0.19082933
0.19373864
0.19373864
0.19373864
0.19782034
0.19981797
0.20006038
0.21166857
0.21279462
0.21279462
0.21279462
0.21279462
0.21506957
0.21530676
0.2218812
0.22580464
NA
0.22580464
0.22580464
NA
NA
0.22730231
0.22730231
0.2274937
NA
0.2274937
NA
0.2307347

0.2307347
0.23646229
0.24437839
0.24437839
0.24437839
0.24437839
0.24437839
NA
0.2498995
0.2498995
0.2498995
0.25311457
NA
0.26117418
0.26117418
NA
0.26632941
0.26632941
0.26632941
NA
NA
0.28869147
NA
NA
0.29652272
NA
NA
NA
NA
0.3098725
0.33165396
NA
NA
NA
0.33702044
0.33702044
NA
0.35060993
NA
0.35533548
NA
NA
NA
0.37328529
NA
NA

NA
0.38183735
0.38183735
NA
NA
0.39608055
0.39608055
NA
0.39608055
NA
NA
0.39608055
0.39608055
0.39608055
NA
0.40318957
0.40505987
0.40558267
NA
NA
NA
NA
NA
0.45622457
0.46097311
0.46097311
NA
NA
0.46097311
0.46097311
0.46097311
0.46097311
0.46097311
0.46097311
0.46097311
NA
NA
0.48014453
0.48014453
0.48093075
NA
0.49070851
NA
NA
0.50390029
0.50746339

NA
NA
0.51575993
0.52381477
NA
0.52433802
0.52433802
0.52433802
NA
NA
0.54118754
NA
NA
0.5494408
0.54995516
NA
0.54995516
0.54995516
NA
NA
NA
0.58918215
0.5931384
0.60010639
NA
0.62080562
0.62080562
NA
NA
0.6374801
0.63981731
0.64008839
NA
NA
0.64884706
NA
NA
0.65914368
NA
0.66481389
0.66840665
0.67568722
NA
0.69019561
NA
NA

0.71654659
0.71654659
NA
0.71654659
0.71654659
0.71654659
NA
NA
NA
NA
0.72612198
NA
0.72612198
NA
0.72612198
0.72612198
NA
0.73400586
NA
0.73704925
NA
NA
NA
NA
NA
NA
NA
NA
NA
NA
NA
0.76531956
NA
0.76531956
NA
NA
NA
NA
NA
NA
NA
NA
NA
NA
NA
NA

NA
NA
0.77591035
NA
NA
NA
NA
NA
NA
NA
NA
NA
NA
NA
NA
0.79809117
NA
NA
0.80619039
0.80619039
NA
0.80619039
0.80619039
0.80814715
NA
NA
NA
NA
NA
0.81780141
NA
NA
0.81780141
0.81780141
0.81780141
0.81780141
0.81780141
0.82254684
NA
0.82331303
0.82331303
0.83160804
NA
0.8473698
NA
0.87497644

NA
0.88141455
0.88141455
0.88141455
0.88141455
0.88141455
0.88141455
NA
0.88279985
NA
0.91298772
NA
NA
NA
0.92302984
NA
0.93795996
NA
NA
NA
NA
0.95427433
NA
NA
NA
NA
NA
0.98743083
NA
