## Supplementary Table 2 for "Single paternal Dexamethasone challenge programs offspring metabolism and reveals circRNAs as novel candidates in RNA-mediated inheritance"

| HS692_tRNA_pvalues_TREATMENT | | | HS692_tRNA_pvalues_TREATMENT | | |
|---|---|---|---|---|---|
| | baseMean | log2FoldChai | lfcSE | stat | pvalue |
| tRNA-His-GTG-2-4 | 28998.4942 | -1.3623687 | 0.23882783 | -5.7043971 | 1.17E-08 |
| tRNA-His-GTG-2-5 | 28998.4942 | -1.3623687 | 0.23882783 | -5.7043971 | 1.17E-08 |
| tRNA-His-GTG-2-6 | 28998.4942 | -1.3623687 | 0.23882783 | -5.7043971 | 1.17E-08 |
| tRNA-His-GTG-2-7 | 28999.5299 | -1.3619111 | 0.23875975 | -5.7041065 | 1.17E-08 |
| tRNA-His-GTG-2-1 | 29001.8773 | -1.3610261 | 0.23882728 | -5.6987882 | 1.21E-08 |
| tRNA-His-GTG-2-3 | 29001.8773 | -1.3610261 | 0.23882728 | -5.6987882 | 1.21E-08 |
| tRNA-His-GTG-2-2 | 29001.8918 | -1.3610261 | 0.23882772 | -5.6987779 | 1.21E-08 |
| tRNA-His-GTG-2-8 | 29001.817 | -1.3610261 | 0.23882776 | -5.6987769 | 1.21E-08 |
| tRNA-Pro-AGG-1-6 | 498.174598 | -0.9854979 | 0.26853431 | -3.6699143 | 0.00024263 |
| tRNA-Pro-AGG-1-4 | 479.374246 | -0.9612603 | 0.27268911 | -3.5251145 | 0.0004233 |
| tRNA-Pro-AGG-1-3 | 474.29805 | -0.9522087 | 0.2717021 | -3.5046055 | 0.00045728 |
| tRNA-Pro-CGG-1-1 | 571.437924 | -0.9367117 | 0.26748252 | -3.5019547 | 0.00046186 |
| tRNA-Pro-AGG-1-2 | 472.481965 | -0.9431157 | 0.27072955 | -3.4836084 | 0.0004947 |
| tRNA-Pro-AGG-1-5 | 474.180973 | -0.9498548 | 0.27293408 | -3.480162 | 0.00050111 |
| tRNA-Pro-CGG-1-3 | 563.110855 | -0.9239633 | 0.26754986 | -3.4534248 | 0.00055352 |
| tRNA-Pro-CGG-1-2 | 574.314626 | -0.9257755 | 0.26812703 | -3.4527496 | 0.0005549 |
| tRNA-Pro-TGG-2-1 | 502.629255 | -0.863964 | 0.26231797 | -3.2935755 | 0.00098922 |
| tRNA-Pro-TGG-2-2 | 501.388162 | -0.856243 | 0.26074053 | -3.2838893 | 0.00102385 |
| tRNA-Pro-TGG-2-3 | 499.10422 | -0.8522323 | 0.261838 | -3.2548076 | 0.00113469 |
| tRNA-Pro-TGG-2-4 | 500.779936 | -0.8390404 | 0.26044046 | -3.2216208 | 0.00127468 |
| tRNA-Pro-AGG-1-1 | 498.548964 | -0.8405206 | 0.26882922 | -3.1265967 | 0.00176842 |
| tRNA-Leu-CAA-2-1 | 362.290665 | 1.5705809 | 0.50615307 | 3.10297614 | 0.00191585 |
| tRNA-Arg-CCT-2-1 | 19.3057441 | 1.3362313 | 0.44505524 | 3.00239424 | 0.00267865 |
| tRNA-Tyr-GTA-3-2 | 9.28585231 | 1.61095356 | 0.55025549 | 2.92764649 | 0.00341538 |
| tRNA-Pro-TGG-1-1 | 358.920056 | -0.7267971 | 0.25784007 | -2.8187903 | 0.0048205 |
| tRNA-Tyr-GTA-3-1 | 9.85513622 | 1.53409995 | 0.55058256 | 2.78632137 | 0.005331 |
| tRNA-Arg-CCT-2-2 | 19.5824216 | 1.24037871 | 0.44915364 | 2.76159112 | 0.00575205 |
| tRNA-Ser-GCT-1-1 | 1032.90289 | 1.07235116 | 0.38926477 | 2.75481176 | 0.00587259 |
| tRNA-Ser-GCT-3-1 | 1110.78244 | 0.98203688 | 0.36833115 | 2.66617925 | 0.00767188 |
| tRNA-Thr-TGT-1-1 | 4.06582087 | -3.6543117 | 1.38462114 | -2.6392142 | 0.00830985 |
| tRNA-Ser-GCT-4-2 | 1098.30062 | 0.98691034 | 0.3752629 | 2.62991718 | 0.00854057 |
| tRNA-Ser-GCT-4-1 | 1098.54502 | 0.98431277 | 0.37521367 | 2.62333932 | 0.00870725 |
| tRNA-Ser-GCT-4-3 | 1102.79447 | 0.9805238 | 0.3754743 | 2.61142718 | 0.00901652 |
| tRNA-Ser-GCT-2-1 | 1106.56683 | 0.96890302 | 0.37301328 | 2.59750271 | 0.00939044 |
| tRNA-Glu-TTC-1-3 | 7516.66951 | -0.6838902 | 0.27034905 | -2.5296562 | 0.01141743 |
| tRNA-Glu-TTC-1-4 | 7540.22739 | -0.6739384 | 0.26921174 | -2.5033765 | 0.01230146 |
| tRNA-Glu-TTC-1-1 | 7533.07318 | -0.6725735 | 0.26933045 | -2.4972057 | 0.01251763 |
| tRNA-Glu-TTC-1-2 | 7533.07318 | -0.6725735 | 0.26933045 | -2.4972057 | 0.01251763 |
| tRNA-His-GTG-3-1 | 196.814482 | -0.7642245 | 0.31117954 | -2.4558956 | 0.0140534 |
| tRNA-Lys-CTT-3-7 | 3829.01173 | -0.6418644 | 0.26203877 | -2.4495016 | 0.01430541 |
| tRNA-Lys-CTT-3-5 | 3828.89617 | -0.6408308 | 0.26207787 | -2.4451922 | 0.0144775 |
| tRNA-Lys-CTT-3-6 | 3828.89617 | -0.6408308 | 0.26207787 | -2.4451922 | 0.0144775 |
| tRNA-Lys-CTT-3-1 | 3852.60864 | -0.625083 | 0.26213051 | -2.3846253 | 0.01709653 |
| tRNA-Lys-CTT-3-4 | 3854.03886 | -0.6246787 | 0.26197901 | -2.3844608 | 0.01710417 |

| | | | | | |
|---|---|---|---|---|---|
| tRNA-Lys-CTT-3-3 | 3858.00155 | -0.6246076 | 0.26230142 | -2.3812588 | 0.01725358 |
| tRNA-Lys-CTT-3-2 | 3876.20711 | -0.6105629 | 0.25863853 | -2.3606806 | 0.01824143 |
| tRNA-Thr-AGT-2-1 | 50.0235375 | -1.3498356 | 0.5740309 | -2.3515033 | 0.01869772 |
| tRNA-Leu-CAA-3-1 | 144.22626 | 0.83364414 | 0.37355718 | 2.23163733 | 0.02563894 |
| tRNA-Thr-CGT-4-1 | 3.57577703 | -3.1077945 | 1.44362123 | -2.152777 | 0.03133621 |
| tRNA-Gln-TTG-6-1 | 5.84371917 | 1.65180453 | 0.79369783 | 2.08115038 | 0.03742014 |
| tRNA-Tyr-GTA-2-1 | 35.6118649 | 0.97030375 | 0.47099591 | 2.06011077 | 0.03938795 |
| tRNA-Pro-TGG-4-1 | 196.596347 | -0.8447831 | 0.41440089 | -2.0385648 | 0.04149349 |
| tRNA-Ala-CGC-5-1 | 9.54769646 | -2.7867904 | 1.3984155 | -1.99282 | 0.04628116 |
| tRNA-Glu-TTC-2-2 | 5698.90117 | -0.5113694 | 0.25781014 | -1.9835117 | 0.0473103 |
| tRNA-Glu-TTC-2-1 | 5697.21165 | -0.5128407 | 0.25857465 | -1.9833371 | 0.04732978 |
| tRNA-Thr-AGT-6-1 | 1.50954656 | 3.82726421 | 1.97064626 | 1.94213659 | 0.05212057 |
| tRNA-Tyr-GTA-1-4 | 3.89448409 | 1.54011668 | 0.80838068 | 1.90518739 | 0.05675572 |
| tRNA-Gly-GCC-6-1 | 530.397599 | -0.7203391 | 0.3860491 | -1.865926 | 0.06205172 |
| tRNA-Trp-CCA-3-1 | 38.9630914 | 0.77041778 | 0.41708799 | 1.84713491 | 0.06472759 |
| tRNA-Ala-CGC-4-1 | 1.76644092 | -2.7032341 | 1.46981958 | -1.8391605 | 0.06589159 |
| tRNA-Glu-CTC-4-1 | 6.42515408 | -2.6836511 | 1.47972379 | -1.8136163 | 0.06973683 |
| tRNA-Cys-GCA-18-1 | 1.4036703 | 3.03167053 | 1.69071796 | 1.79312611 | 0.07295276 |
| tRNA-Leu-TAA-2-1 | 3119.15501 | 0.65104272 | 0.36769169 | 1.77062125 | 0.07662371 |
| tRNA-Thr-CGT-3-1 | 1.32481075 | 3.8404165 | 2.17981098 | 1.76181171 | 0.07810112 |
| tRNA-Trp-CCA-2-1 | 1815.9669 | 0.52783028 | 0.3085327 | 1.71077579 | 0.08712251 |
| tRNA-iMet-CAT-3-1 | 1272.12042 | -0.8437164 | 0.50043998 | -1.6859492 | 0.09180558 |
| tRNA-Val-CAC-3-1 | 4033.67882 | -0.4936672 | 0.29485207 | -1.6742876 | 0.09407409 |
| tRNA-Cys-GCA-11-1 | 1.7644337 | -2.3960031 | 1.44852347 | -1.6541003 | 0.09810714 |
| tRNA-Gly-CCC-4-1 | 1065.83419 | -0.6070997 | 0.36820312 | -1.6488174 | 0.09918505 |
| tRNA-Thr-TGT-2-1 | 29.0184782 | -0.6157961 | 0.37389175 | -1.6469904 | 0.09956003 |
| tRNA-Val-AAC-3-1 | 3954.19939 | -0.4832795 | 0.29511875 | -1.6375765 | 0.10151006 |
| tRNA-Ile-AAT-2-1 | 7.08639499 | 1.24930041 | 0.76907436 | 1.62442084 | 0.10428602 |
| tRNA-Val-AAC-2-1 | 3983.63556 | -0.4738466 | 0.29416665 | -1.6108101 | 0.10722113 |
| tRNA-Val-AAC-2-2 | 3983.63556 | -0.4738466 | 0.29416665 | -1.6108101 | 0.10722113 |
| tRNA-Gly-CCC-3-1 | 1073.04047 | -0.5833597 | 0.36515212 | -1.59758 | 0.11013649 |
| tRNA-iMet-CAT-1-1 | 1324.81683 | -0.7482843 | 0.47597842 | -1.5720971 | 0.11592803 |
| tRNA-SeC-TCA-1-1 | 10932.2345 | 0.39910732 | 0.25689366 | 1.5535896 | 0.12028234 |
| tRNA-iMet-CAT-1-2 | 1316.71192 | -0.7383192 | 0.47742149 | -1.5464725 | 0.12199051 |
| tRNA-iMet-CAT-1-4 | 1316.71192 | -0.7383192 | 0.47742149 | -1.5464725 | 0.12199051 |
| tRNA-iMet-CAT-1-5 | 1316.68596 | -0.7377909 | 0.47730445 | -1.545745 | 0.12216618 |
| tRNA-iMet-CAT-1-6 | 1323.30472 | -0.7359742 | 0.47736838 | -1.5417322 | 0.12313867 |
| tRNA-iMet-CAT-1-7 | 1318.84617 | -0.7345693 | 0.47663748 | -1.5411488 | 0.12328057 |
| tRNA-iMet-CAT-1-3 | 1325.69778 | -0.7336924 | 0.4771361 | -1.5377005 | 0.12412187 |
| tRNA-Glu-CTC-3-1 | 18.9331099 | -0.9177641 | 0.60571909 | -1.5151646 | 0.12973073 |
| tRNA-Arg-CCT-1-1 | 19.0254253 | 0.72693585 | 0.48248583 | 1.50664706 | 0.13190115 |
| tRNA-Val-AAC-4-1 | 1646.3544 | -0.6586985 | 0.44492432 | -1.480473 | 0.13874705 |
| tRNA-Asn-GTT-3-6 | 24.7445466 | 0.58574179 | 0.40162707 | 1.45842208 | 0.14472424 |
| tRNA-Thr-AGT-4-1 | 13.5343955 | -1.0561924 | 0.7481653 | -1.4117099 | 0.15803539 |
| tRNA-Asn-GTT-1-1 | 15.3438029 | 0.67048176 | 0.47556535 | 1.4098625 | 0.15858029 |
| tRNA-Asn-GTT-3-7 | 27.6734014 | 0.54885941 | 0.39073176 | 1.40469617 | 0.16011165 |

| | | | | | |
|---|---|---|---|---|---|
| tRNA-Lys-CTT-2-2 | 3471.13671 | -0.3103105 | 0.22279534 | -1.3928053 | 0.16367868 |
| tRNA-Lys-CTT-2-1 | 3466.48099 | -0.3092926 | 0.22278637 | -1.3882923 | 0.16504806 |
| tRNA-Gly-CCC-1-2 | 1464.64312 | -0.3673197 | 0.2655212 | -1.3833913 | 0.16654491 |
| tRNA-Gly-CCC-1-1 | 1464.64312 | -0.3673197 | 0.2655212 | -1.3833913 | 0.16654491 |
| tRNA-Asn-GTT-3-8 | 36.4757383 | 0.6216933 | 0.44994379 | 1.38171328 | 0.16705975 |
| tRNA-Asn-GTT-3-1 | 29.9718303 | 0.53190186 | 0.3851451 | 1.38104279 | 0.1672658 |
| tRNA-Lys-TTT-2-1 | 27.0666458 | 0.81078962 | 0.58914122 | 1.37622287 | 0.16875265 |
| tRNA-Lys-TTT-2-2 | 26.9172173 | 0.81097123 | 0.59054574 | 1.37325726 | 0.16967238 |
| tRNA-Gly-CCC-2-1 | 3966.66658 | -0.5271339 | 0.3936368 | -1.3391379 | 0.18052579 |
| tRNA-Gly-CCC-2-2 | 3966.66658 | -0.5271339 | 0.3936368 | -1.3391379 | 0.18052579 |
| tRNA-Leu-TAA-3-1 | 0.76610446 | -2.5766119 | 1.93563982 | -1.3311422 | 0.18314222 |
| tRNA-Asn-GTT-3-4 | 24.0922481 | 0.57893193 | 0.43608397 | 1.32756986 | 0.18432025 |
| tRNA-Asn-GTT-3-5 | 24.0922481 | 0.57893193 | 0.43608397 | 1.32756986 | 0.18432025 |
| tRNA-Asn-GTT-3-3 | 23.967007 | 0.5780695 | 0.44111026 | 1.31048755 | 0.19003095 |
| tRNA-Val-AAC-5-1 | 33823.3676 | -0.3674765 | 0.28158617 | -1.3050231 | 0.19188498 |
| tRNA-Gln-TTG-3-2 | 184.203703 | 0.33656095 | 0.26688398 | 1.26107588 | 0.20728151 |
| tRNA-Leu-AAG-2-1 | 811.615521 | 0.35161305 | 0.27917385 | 1.25947703 | 0.20785808 |
| tRNA-Asn-GTT-3-9 | 24.368177 | 0.5491502 | 0.43623403 | 1.2588431 | 0.20808701 |
| tRNA-Gln-TTG-3-1 | 184.544006 | 0.3313856 | 0.26626138 | 1.24458756 | 0.21328339 |
| tRNA-Gln-CTG-6-1 | 107.84331 | -0.3880913 | 0.31385354 | -1.2365364 | 0.21625924 |
| tRNA-Leu-AAG-3-1 | 798.841555 | 0.34753094 | 0.28234673 | 1.23086582 | 0.21837305 |
| tRNA-Asn-GTT-3-2 | 59.2855903 | 0.56921797 | 0.46626303 | 1.22080872 | 0.22215845 |
| tRNA-Gln-TTG-5-1 | 41.1598104 | 0.54355252 | 0.44637541 | 1.21770266 | 0.22333699 |
| tRNA-Leu-AAG-1-1 | 1011.72894 | 0.30296931 | 0.2550063 | 1.1880856 | 0.23479969 |
| tRNA-Leu-AAG-1-2 | 1011.53652 | 0.30297041 | 0.25540465 | 1.18623683 | 0.23552879 |
| tRNA-Leu-AAG-1-3 | 1011.657 | 0.30297075 | 0.25553001 | 1.18565624 | 0.23575808 |
| tRNA-Leu-CAA-4-1 | 81.1825193 | 0.36287236 | 0.30752255 | 1.17998621 | 0.2380057 |
| tRNA-Ser-GCT-6-1 | 12.6153113 | 0.60050246 | 0.51583714 | 1.16413187 | 0.24437057 |
| tRNA-Val-CAC-2-5 | 459699.745 | -0.3136717 | 0.27283516 | -1.1496747 | 0.25027789 |
| tRNA-Val-CAC-2-4 | 459696.983 | -0.3136722 | 0.27283767 | -1.1496659 | 0.25028152 |
| tRNA-Val-CAC-2-1 | 459699.535 | -0.3136625 | 0.27283508 | -1.1496415 | 0.25029157 |
| tRNA-Val-CAC-2-2 | 459699.312 | -0.3136483 | 0.27283353 | -1.149596 | 0.2503103 |
| tRNA-Val-CAC-2-3 | 459702.447 | -0.3135187 | 0.27283706 | -1.1491061 | 0.25051225 |
| tRNA-Gln-CTG-1-1 | 241.967211 | -0.2984508 | 0.26391433 | -1.1308625 | 0.25811298 |
| tRNA-Val-AAC-1-2 | 454075.646 | -0.3060752 | 0.2723811 | -1.123702 | 0.26113946 |
| tRNA-Val-AAC-1-1 | 454079.297 | -0.30599 | 0.27238209 | -1.1233852 | 0.26127395 |
| tRNA-Leu-TAG-2-1 | 796.037019 | 0.2974026 | 0.27573351 | 1.07858706 | 0.28077185 |
| tRNA-Tyr-GTA-6-1 | 4.68458016 | 1.23106945 | 1.14563612 | 1.07457284 | 0.28256602 |
| tRNA-Tyr-GTA-1-2 | 3.21237119 | 0.9568298 | 0.89099354 | 1.07389084 | 0.28287161 |
| tRNA-Tyr-GTA-1-5 | 3.28708544 | 0.95703535 | 0.89263588 | 1.07214528 | 0.28365478 |
| tRNA-Leu-TAG-1-1 | 1006.84179 | 0.26706551 | 0.25019964 | 1.06740967 | 0.28578688 |
| tRNA-Gly-TCC-1-7 | 393.953649 | 0.42137553 | 0.39736666 | 1.06041995 | 0.28895359 |
| tRNA-Tyr-GTA-1-3 | 3.5270186 | 0.95844752 | 0.90411597 | 1.06009356 | 0.28910204 |
| tRNA-Gly-TCC-1-1 | 398.738556 | 0.40642328 | 0.3914181 | 1.03833543 | 0.29911392 |
| tRNA-Gly-TCC-1-6 | 427.670255 | 0.42220898 | 0.40699806 | 1.03737345 | 0.29956185 |
| tRNA-Gly-TCC-1-2 | 390.66018 | 0.41210561 | 0.39958981 | 1.0313216 | 0.30239003 |

| | | | | | |
|---|---|---|---|---|---|
| tRNA-Gly-TCC-1-3 | 390.66018 | 0.41210561 | 0.39958981 | 1.0313216 | 0.30239003 |
| tRNA-Gly-TCC-1-4 | 390.66018 | 0.41210561 | 0.39958981 | 1.0313216 | 0.30239003 |
| tRNA-Asp-GTC-2-1 | 1891.71821 | 0.22302682 | 0.21628656 | 1.03116358 | 0.30246412 |
| tRNA-Gly-TCC-1-5 | 394.166849 | 0.40855341 | 0.39941638 | 1.02287595 | 0.3063665 |
| tRNA-Tyr-GTA-1-1 | 3.24003894 | 0.89651077 | 0.88882034 | 1.0086524 | 0.31314137 |
| tRNA-Lys-TTT-1-1 | 614.218266 | 0.43493844 | 0.43615519 | 0.99721029 | 0.31866245 |
| tRNA-Arg-CCG-1-1 | 24.2194041 | -0.447937 | 0.44930016 | -0.996966 | 0.31878099 |
| tRNA-Asn-GTT-2-1 | 9.19200847 | 0.54247068 | 0.54835251 | 0.98927362 | 0.32252929 |
| tRNA-Met-CAT-4-1 | 9.82922744 | -0.7545876 | 0.77280774 | -0.9764234 | 0.32885468 |
| tRNA-Lys-TTT-1-6 | 603.189306 | 0.43009455 | 0.44294584 | 0.97098677 | 0.33155487 |
| tRNA-Lys-TTT-1-3 | 636.401634 | 0.4141849 | 0.42695833 | 0.97008272 | 0.33200526 |
| tRNA-Ala-AGC-10-1 | 66.8530965 | 0.33600689 | 0.34652326 | 0.96965176 | 0.3322201 |
| tRNA-Ala-AGC-2-2 | 85.7093237 | -0.3193602 | 0.3315709 | -0.9631731 | 0.33546066 |
| tRNA-Arg-TCT-4-1 | 12.319731 | 0.81750601 | 0.84884976 | 0.96307504 | 0.33550987 |
| tRNA-Ala-AGC-5-1 | 68.668781 | 0.33251886 | 0.34625616 | 0.96032619 | 0.33689107 |
| tRNA-Ala-AGC-5-2 | 66.9550485 | 0.33254678 | 0.34729997 | 0.95752031 | 0.3383047 |
| tRNA-Ala-AGC-5-3 | 66.9550485 | 0.33254678 | 0.34729997 | 0.95752031 | 0.3383047 |
| tRNA-Gln-TTG-1-1 | 168.678218 | 0.23855948 | 0.25393921 | 0.93943537 | 0.34750726 |
| tRNA-Lys-TTT-1-2 | 574.090964 | 0.43711394 | 0.46621019 | 0.93758983 | 0.34845524 |
| tRNA-Ala-AGC-2-1 | 83.4787422 | -0.3180111 | 0.3393722 | -0.9370571 | 0.3487292 |
| tRNA-Ile-TAT-2-3 | 2.17857793 | -1.3730326 | 1.46812145 | -0.9352309 | 0.3496693 |
| tRNA-Cys-GCA-16-1 | 19.0894496 | -0.5665824 | 0.61611063 | -0.9196115 | 0.35777583 |
| tRNA-Lys-TTT-1-5 | 604.229473 | 0.39978852 | 0.43656492 | 0.91575961 | 0.35979299 |
| tRNA-Ala-AGC-12-1 | 67.9931028 | 0.30999342 | 0.34122475 | 0.90847285 | 0.36362845 |
| tRNA-Lys-TTT-1-4 | 600.173947 | 0.3983786 | 0.4444507 | 0.89633923 | 0.37007161 |
| tRNA-His-GTG-1-1 | 169.594431 | -0.299992 | 0.33549033 | -0.8941896 | 0.37122046 |
| tRNA-Gly-TCC-2-1 | 243.066674 | 0.64640151 | 0.73890922 | 0.87480504 | 0.38168 |
| tRNA-Leu-CAA-1-1 | 79.2749203 | 0.26228593 | 0.30049487 | 0.87284659 | 0.3827467 |
| tRNA-Cys-GCA-1-1 | 3162.87586 | -0.2115895 | 0.25149782 | -0.8413174 | 0.40017012 |
| tRNA-Met-CAT-5-1 | 8.28216092 | -0.572903 | 0.69038181 | -0.829835 | 0.4066321 |
| tRNA-Cys-GCA-1-2 | 3179.62572 | -0.2089789 | 0.25222511 | -0.8285414 | 0.40736395 |
| tRNA-Gln-CTG-7-1 | 2.46529023 | -0.9955553 | 1.20855636 | -0.8237558 | 0.41007835 |
| tRNA-Cys-GCA-3-4 | 3170.81706 | -0.2047311 | 0.25035303 | -0.8177698 | 0.41348865 |
| tRNA-Trp-CCA-6-1 | 19.3788719 | 0.36457253 | 0.44680068 | 0.81596234 | 0.41452168 |
| tRNA-Cys-GCA-3-2 | 3185.66778 | -0.2044905 | 0.25061908 | -0.8159413 | 0.41453371 |
| tRNA-Cys-GCA-3-3 | 3185.20845 | -0.2042915 | 0.25062054 | -0.8151427 | 0.41499061 |
| tRNA-Lys-CTT-7-1 | 0.28965768 | 2.4290319 | 2.99666681 | 0.8105779 | 0.41760811 |
| tRNA-Gln-CTG-5-1 | 85.5342562 | 0.39012952 | 0.48299686 | 0.80772683 | 0.41924786 |
| tRNA-Arg-CCG-3-1 | 685.49171 | -0.3479006 | 0.43176805 | -0.805758 | 0.42038239 |
| tRNA-Cys-GCA-3-1 | 3172.77916 | -0.2015458 | 0.25053302 | -0.804468 | 0.42112677 |
| tRNA-Cys-GCA-2-1 | 3180.73817 | -0.2020409 | 0.25150728 | -0.8033204 | 0.42178957 |
| tRNA-Met-CAT-1-2 | 282.708567 | 0.31513183 | 0.39444301 | 0.79892866 | 0.42433178 |
| tRNA-Tyr-GTA-5-1 | 0.29380037 | 3.58420354 | 4.49578173 | 0.797237 | 0.4253134 |
| tRNA-Met-CAT-1-1 | 279.447803 | 0.31503107 | 0.40211903 | 0.78342741 | 0.43337616 |
| tRNA-Val-CAC-1-1 | 2394.51039 | -0.2410173 | 0.31533792 | -0.7643143 | 0.44467996 |
| tRNA-Glu-TTC-3-1 | 144.563688 | 0.28190372 | 0.37109608 | 0.75965155 | 0.4474629 |

| | | | | | |
|---|---|---|---|---|---|
| tRNA-Glu-TTC-3-2 | 144.563688 | 0.28190372 | 0.37109608 | 0.75965155 | 0.4474629 |
| tRNA-Tyr-GTA-4-1 | 3.5628694 | 0.89139214 | 1.17727893 | 0.75716308 | 0.44895217 |
| tRNA-Trp-CCA-3-2 | 40.7546967 | 0.29487205 | 0.39038711 | 0.75533244 | 0.45004954 |
| tRNA-Leu-CAG-3-1 | 10.3740117 | 0.35330454 | 0.50635977 | 0.69773422 | 0.48534343 |
| tRNA-Ala-TGC-1-1 | 24.514306 | -0.3275953 | 0.4716986 | -0.6945013 | 0.48736791 |
| tRNA-Ile-AAT-1-7 | 25.4197922 | 0.26103445 | 0.38634873 | 0.67564465 | 0.49926629 |
| tRNA-Arg-TCT-1-1 | 8.71325274 | -0.5311647 | 0.79434889 | -0.6686793 | 0.50370008 |
| tRNA-Thr-CGT-2-1 | 49.5938764 | 0.2869287 | 0.44024621 | 0.651746 | 0.51456504 |
| tRNA-Asp-GTC-1-10 | 404.065018 | 0.25232878 | 0.39187751 | 0.64389707 | 0.51964218 |
| tRNA-Ala-TGC-6-1 | 593.053412 | 0.23891288 | 0.3731905 | 0.64019013 | 0.522049 |
| tRNA-Met-CAT-3-1 | 30.246461 | 0.27536539 | 0.43048958 | 0.63965634 | 0.52239605 |
| tRNA-Asp-GTC-1-1 | 404.200032 | 0.24828741 | 0.39301507 | 0.63175036 | 0.52755001 |
| tRNA-Asp-GTC-1-11 | 404.495896 | 0.24429179 | 0.39133076 | 0.6242591 | 0.53245744 |
| tRNA-Asp-GTC-1-13 | 403.162405 | 0.24354079 | 0.3922992 | 0.62080368 | 0.5347288 |
| tRNA-Asp-GTC-1-3 | 402.920073 | 0.24353957 | 0.39241163 | 0.6206227 | 0.5348479 |
| tRNA-Asp-GTC-1-4 | 402.920073 | 0.24353957 | 0.39241163 | 0.6206227 | 0.5348479 |
| tRNA-Asp-GTC-1-9 | 402.920073 | 0.24353957 | 0.39241163 | 0.6206227 | 0.5348479 |
| tRNA-Asp-GTC-1-7 | 403.159636 | 0.24353891 | 0.39247172 | 0.62052602 | 0.53491153 |
| tRNA-Asp-GTC-1-2 | 403.039162 | 0.24353877 | 0.39248508 | 0.62050453 | 0.53492567 |
| tRNA-Asp-GTC-1-5 | 404.36788 | 0.24409105 | 0.39338164 | 0.62049427 | 0.53493242 |
| tRNA-Asp-GTC-1-12 | 403.134072 | 0.24173805 | 0.39241545 | 0.61602583 | 0.53787747 |
| tRNA-Asp-GTC-1-8 | 403.237111 | 0.24173754 | 0.39246298 | 0.61594993 | 0.53792757 |
| tRNA-Asp-GTC-1-6 | 403.617022 | 0.24173454 | 0.3927427 | 0.61550358 | 0.53822221 |
| tRNA-Arg-ACG-2-1 | 9.19852009 | -0.3993001 | 0.66222461 | -0.6029678 | 0.54653009 |
| tRNA-Leu-TAA-5-1 | 2.15092461 | -0.7706311 | 1.28710249 | -0.5987333 | 0.54935077 |
| tRNA-Leu-TAA-5-2 | 2.15092461 | -0.7706311 | 1.28710249 | -0.5987333 | 0.54935077 |
| tRNA-Leu-TAA-5-3 | 2.15092461 | -0.7706311 | 1.28710249 | -0.5987333 | 0.54935077 |
| tRNA-Cys-GCA-7-1 | 272.909989 | 0.13904603 | 0.23568932 | 0.58995473 | 0.555221 |
| tRNA-Gly-ACC-1-1 | 42.0915647 | -0.2006921 | 0.34478132 | -0.5820851 | 0.56050938 |
| tRNA-Lys-CTT-15-1 | 0.79335361 | 0.96089805 | 1.66211014 | 0.57811936 | 0.56318354 |
| tRNA-Ile-AAT-3-1 | 23.6073554 | 0.23471134 | 0.4064314 | 0.57749313 | 0.56360638 |
| tRNA-Glu-CTC-1-8 | 12519.5887 | -0.161039 | 0.28134914 | -0.5723813 | 0.56706366 |
| tRNA-Glu-CTC-1-4 | 12540.0237 | -0.160588 | 0.28126271 | -0.5709537 | 0.56803105 |
| tRNA-Glu-CTC-1-5 | 12503.0878 | -0.1605714 | 0.28138778 | -0.5706411 | 0.56824296 |
| tRNA-Glu-CTC-1-6 | 12503.0878 | -0.1605714 | 0.28138778 | -0.5706411 | 0.56824296 |
| tRNA-Glu-CTC-1-9 | 12504.4328 | -0.1604117 | 0.28153458 | -0.569776 | 0.56882963 |
| tRNA-Glu-CTC-1-1 | 12513.9485 | -0.160357 | 0.28159803 | -0.5694535 | 0.56904843 |
| tRNA-Glu-CTC-1-2 | 12513.9485 | -0.160357 | 0.28159803 | -0.5694535 | 0.56904843 |
| tRNA-Glu-CTC-1-3 | 12513.9485 | -0.160357 | 0.28159803 | -0.5694535 | 0.56904843 |
| tRNA-Glu-CTC-1-7 | 12547.3672 | -0.1598138 | 0.28107664 | -0.5685774 | 0.56964295 |
| tRNA-Lys-CTT-13-1 | 7.44524461 | -0.5610636 | 0.98791857 | -0.5679249 | 0.57008595 |
| tRNA-Ala-AGC-6-1 | 18.0796148 | -0.3311155 | 0.59056708 | -0.5606738 | 0.57501991 |
| tRNA-Arg-CCT-4-1 | 107.841334 | -0.1823569 | 0.3256071 | -0.560052 | 0.57544398 |
| tRNA-Val-CAC-6-1 | 8198.89709 | -0.1570676 | 0.28439006 | -0.5522963 | 0.58074536 |
| tRNA-Ser-GCT-5-1 | 13.6209205 | -0.3433073 | 0.62226697 | -0.5517043 | 0.58115098 |
| tRNA-Lys-CTT-16-1 | 1.96092478 | 0.54695404 | 1.02251966 | 0.53490809 | 0.59271341 |

| | | | | | |
|---|---|---|---|---|---|
| tRNA-Gln-CTG-4-1 | 120.62929 | -0.2104046 | 0.39787618 | -0.5288193 | 0.59693082 |
| tRNA-Arg-ACG-1-3 | 19.0353373 | -0.2725326 | 0.51599874 | -0.5281653 | 0.59738464 |
| tRNA-Cys-GCA-17-1 | 5.85840989 | -0.4024144 | 0.76403289 | -0.5266978 | 0.59840348 |
| tRNA-Leu-TAA-1-1 | 54.3850394 | 0.23226521 | 0.44791434 | 0.51854829 | 0.60407578 |
| tRNA-Ser-CGA-1-1 | 54.8551022 | -0.2082277 | 0.4043615 | -0.5149544 | 0.60658488 |
| tRNA-Trp-CCA-1-1 | 25.9166198 | 0.22509931 | 0.44283802 | 0.50831072 | 0.61123546 |
| tRNA-Ala-AGC-7-1 | 17.8601853 | -0.298462 | 0.58923947 | -0.5065208 | 0.61249111 |
| tRNA-Thr-CGT-1-1 | 20.4722709 | 0.29584239 | 0.58913634 | 0.50216287 | 0.61555296 |
| tRNA-Arg-TCG-1-1 | 61.4602617 | 0.13454696 | 0.2740041 | 0.49103995 | 0.62339819 |
| tRNA-Cys-GCA-5-1 | 283.906768 | 0.11536689 | 0.23627303 | 0.48827786 | 0.62535304 |
| tRNA-Arg-ACG-3-2 | 30.7468997 | -0.2079102 | 0.42661393 | -0.4873497 | 0.62601051 |
| tRNA-Trp-CCA-5-1 | 500.535416 | 0.16214556 | 0.33597558 | 0.48261114 | 0.62937187 |
| tRNA-Arg-TCT-2-1 | 20.4532262 | -0.2707235 | 0.56924017 | -0.4755875 | 0.63436826 |
| tRNA-Phe-GAA-1-3 | 14.6979462 | -0.2191858 | 0.46986113 | -0.4664905 | 0.64086444 |
| tRNA-Ser-CGA-3-1 | 53.5815648 | -0.1915842 | 0.41503652 | -0.461608 | 0.64436245 |
| tRNA-Ala-AGC-4-1 | 82.8674516 | 0.14361681 | 0.32159258 | 0.44657999 | 0.65517835 |
| tRNA-Ile-AAT-1-8 | 31.1021011 | 0.16167624 | 0.36562017 | 0.44219726 | 0.65834647 |
| tRNA-Val-CAC-5-1 | 0.23619037 | 1.93498431 | 4.43590052 | 0.43621003 | 0.66268435 |
| tRNA-Phe-GAA-1-1 | 22.3229563 | -0.2144657 | 0.49167242 | -0.4361963 | 0.66269435 |
| tRNA-Phe-GAA-2-1 | 0.91742194 | -0.67436 | 1.62651098 | -0.4146053 | 0.6784309 |
| tRNA-Cys-GCA-15-1 | 5.35612196 | 0.29579052 | 0.71696435 | 0.41255959 | 0.67992931 |
| tRNA-Leu-CAG-2-2 | 682.200845 | 0.12632322 | 0.30818987 | 0.40988765 | 0.68188836 |
| tRNA-Cys-GCA-4-12 | 267.618534 | 0.09921444 | 0.24417792 | 0.40632029 | 0.68450728 |
| tRNA-Phe-GAA-1-5 | 15.7564053 | -0.1839401 | 0.45539368 | -0.4039146 | 0.68627555 |
| tRNA-Cys-GCA-4-13 | 269.242604 | 0.09763215 | 0.24227795 | 0.40297581 | 0.68696602 |
| tRNA-Cys-GCA-4-9 | 269.103436 | 0.09762437 | 0.24234229 | 0.40283672 | 0.68706834 |
| tRNA-Leu-CAG-2-1 | 695.833168 | 0.12550527 | 0.31464397 | 0.39888028 | 0.68998142 |
| tRNA-Cys-GCA-4-22 | 267.388749 | 0.09795647 | 0.24578835 | 0.39853992 | 0.69023224 |
| tRNA-Cys-GCA-4-23 | 267.388749 | 0.09795647 | 0.24578835 | 0.39853992 | 0.69023224 |
| tRNA-Cys-GCA-4-25 | 267.388749 | 0.09795647 | 0.24578835 | 0.39853992 | 0.69023224 |
| tRNA-Cys-GCA-4-28 | 267.388749 | 0.09795647 | 0.24578835 | 0.39853992 | 0.69023224 |
| tRNA-Cys-GCA-24-1 | 267.526793 | 0.09795038 | 0.24583453 | 0.3984403 | 0.69030566 |
| tRNA-Cys-GCA-4-8 | 267.435139 | 0.09794967 | 0.24583996 | 0.3984286 | 0.69031428 |
| tRNA-Gly-CCC-5-1 | 81882.7096 | -0.1392963 | 0.35106706 | -0.3967797 | 0.69152992 |
| tRNA-Cys-GCA-4-15 | 267.419715 | 0.09723978 | 0.24587251 | 0.39548863 | 0.69248231 |
| tRNA-Cys-GCA-4-1 | 267.466104 | 0.097233 | 0.24592408 | 0.39537811 | 0.69256386 |
| tRNA-Cys-GCA-4-21 | 267.466104 | 0.097233 | 0.24592408 | 0.39537811 | 0.69256386 |
| tRNA-Cys-GCA-9-1 | 267.201577 | 0.09683478 | 0.24557129 | 0.39432453 | 0.69334145 |
| tRNA-Ala-AGC-4-2 | 85.6153283 | 0.13147486 | 0.33586068 | 0.39145654 | 0.69545981 |
| tRNA-Cys-GCA-4-2 | 267.317117 | 0.09589335 | 0.24537807 | 0.39079837 | 0.69594628 |
| tRNA-Cys-GCA-4-4 | 267.317117 | 0.09589335 | 0.24537807 | 0.39079837 | 0.69594628 |
| tRNA-Cys-GCA-4-10 | 267.317117 | 0.09589335 | 0.24537807 | 0.39079837 | 0.69594628 |
| tRNA-Cys-GCA-4-11 | 267.317117 | 0.09589335 | 0.24537807 | 0.39079837 | 0.69594628 |
| tRNA-Phe-GAA-1-2 | 14.5147963 | -0.1828287 | 0.47043412 | -0.3886383 | 0.69754375 |
| tRNA-Cys-GCA-14-1 | 267.406934 | 0.09518374 | 0.24540814 | 0.38785894 | 0.69812042 |
| tRNA-Cys-GCA-4-3 | 267.406934 | 0.09518374 | 0.24540814 | 0.38785894 | 0.69812042 |

| | | | | | |
|---|---|---|---|---|---|
| tRNA-Cys-GCA-4-5 | 267.406934 | 0.09518374 | 0.24540814 | 0.38785894 | 0.69812042 |
| tRNA-Cys-GCA-4-6 | 267.406934 | 0.09518374 | 0.24540814 | 0.38785894 | 0.69812042 |
| tRNA-Cys-GCA-4-7 | 267.406934 | 0.09518374 | 0.24540814 | 0.38785894 | 0.69812042 |
| tRNA-Cys-GCA-4-14 | 267.406934 | 0.09518374 | 0.24540814 | 0.38785894 | 0.69812042 |
| tRNA-Cys-GCA-12-1 | 267.348082 | 0.0951766 | 0.24546247 | 0.387744 | 0.69820549 |
| tRNA-Cys-GCA-4-16 | 267.348082 | 0.0951766 | 0.24546247 | 0.387744 | 0.69820549 |
| tRNA-Cys-GCA-6-1 | 267.348082 | 0.0951766 | 0.24546247 | 0.387744 | 0.69820549 |
| tRNA-Cys-GCA-4-17 | 267.348082 | 0.0951766 | 0.24546247 | 0.387744 | 0.69820549 |
| tRNA-Cys-GCA-4-18 | 267.348082 | 0.0951766 | 0.24546247 | 0.387744 | 0.69820549 |
| tRNA-Cys-GCA-4-19 | 267.348082 | 0.0951766 | 0.24546247 | 0.387744 | 0.69820549 |
| tRNA-Cys-GCA-4-20 | 267.348082 | 0.0951766 | 0.24546247 | 0.387744 | 0.69820549 |
| tRNA-Cys-GCA-4-24 | 267.348082 | 0.0951766 | 0.24546247 | 0.387744 | 0.69820549 |
| tRNA-Cys-GCA-10-1 | 267.348082 | 0.0951766 | 0.24546247 | 0.387744 | 0.69820549 |
| tRNA-Cys-GCA-4-26 | 267.348082 | 0.0951766 | 0.24546247 | 0.387744 | 0.69820549 |
| tRNA-Cys-GCA-4-27 | 267.348082 | 0.0951766 | 0.24546247 | 0.387744 | 0.69820549 |
| tRNA-Cys-GCA-4-29 | 267.348082 | 0.0951766 | 0.24546247 | 0.387744 | 0.69820549 |
| tRNA-Phe-GAA-3-1 | 8.07769192 | -0.2420351 | 0.62741628 | -0.3857647 | 0.69967092 |
| tRNA-Leu-CAG-1-5 | 671.968702 | 0.11944262 | 0.31458442 | 0.37968385 | 0.70418011 |
| tRNA-Lys-CTT-10-1 | 2.16766622 | 0.37914832 | 0.99889056 | 0.37956943 | 0.70426506 |
| tRNA-Leu-CAG-1-1 | 670.982084 | 0.11779261 | 0.31417175 | 0.37493063 | 0.70771206 |
| tRNA-Leu-CAG-1-2 | 670.982084 | 0.11779261 | 0.31417175 | 0.37493063 | 0.70771206 |
| tRNA-Leu-CAG-1-3 | 670.982084 | 0.11779261 | 0.31417175 | 0.37493063 | 0.70771206 |
| tRNA-Leu-CAG-2-3 | 715.525508 | 0.12044344 | 0.32190912 | 0.37415356 | 0.70829007 |
| tRNA-Arg-TCG-4-1 | 24.7520366 | 0.19637164 | 0.52488093 | 0.37412607 | 0.70831053 |
| tRNA-Gly-GCC-3-1 | 82830.6429 | -0.1293521 | 0.34629237 | -0.3735345 | 0.70875067 |
| tRNA-Ala-AGC-1-1 | 59.737632 | -0.1282055 | 0.3441865 | -0.3724885 | 0.70952918 |
| tRNA-Gly-GCC-2-8 | 82771.4561 | -0.1278637 | 0.34620737 | -0.369327 | 0.71188398 |
| tRNA-Gly-GCC-2-4 | 82742.828 | -0.127843 | 0.34623522 | -0.3692376 | 0.71195066 |
| tRNA-Gly-GCC-2-5 | 82743.8984 | -0.127835 | 0.34624134 | -0.3692077 | 0.7119729 |
| tRNA-Gly-GCC-2-6 | 82751.08 | -0.127799 | 0.3462319 | -0.369114 | 0.71204277 |
| tRNA-Gly-GCC-2-1 | 82739.3231 | -0.1276933 | 0.34623293 | -0.3688076 | 0.71227117 |
| tRNA-Gly-GCC-2-7 | 82764.1107 | -0.1276628 | 0.34617808 | -0.3687779 | 0.71229331 |
| tRNA-Gly-GCC-2-2 | 82743.4953 | -0.1276411 | 0.34625561 | -0.3686325 | 0.71240164 |
| tRNA-Gly-GCC-2-3 | 82825.9032 | -0.1270669 | 0.34581725 | -0.3674396 | 0.71329117 |
| tRNA-Leu-TAA-4-1 | 0.84691153 | 0.81276343 | 2.2139879 | 0.36710383 | 0.71354157 |
| tRNA-Gly-GCC-4-1 | 82804.0993 | -0.1268167 | 0.34587989 | -0.3666496 | 0.71388043 |
| tRNA-Leu-CAG-1-4 | 673.662652 | 0.11445342 | 0.31719151 | 0.36083381 | 0.71822369 |
| tRNA-Glu-CTC-2-1 | 367.676935 | -0.3158532 | 0.8809595 | -0.3585332 | 0.71994432 |
| tRNA-Lys-CTT-18-1 | 7.24927805 | -0.3492052 | 0.98396378 | -0.3548964 | 0.72266724 |
| tRNA-Gln-TTG-2-1 | 92.281875 | 0.10742189 | 0.3085836 | 0.34811275 | 0.72775551 |
| tRNA-Ile-AAT-1-4 | 22.5194002 | -0.1349195 | 0.39205692 | -0.3441323 | 0.73074678 |
| tRNA-Cys-GCA-26-1 | 2.44593532 | -0.3946561 | 1.15174075 | -0.3426605 | 0.73185386 |
| tRNA-Ile-TAT-3-1 | 0.11139863 | 1.53401588 | 4.54293138 | 0.33767093 | 0.73561118 |
| tRNA-Lys-CTT-4-1 | 0.11278317 | 1.53401588 | 4.54293138 | 0.33767093 | 0.73561118 |
| tRNA-Pro-GGG-1-1 | 0.09277906 | 1.53398224 | 4.54293138 | 0.33766353 | 0.73561676 |
| tRNA-Sup-TTA-1-1 | 0.10509283 | 1.53397853 | 4.54293138 | 0.33766271 | 0.73561738 |

| | | | | | |
|---|---|---|---|---|---|
| tRNA-Arg-TCG-3-1 | 23.4064501 | 0.1724804 | 0.51522003 | 0.33477038 | 0.73779831 |
| tRNA-Arg-TCG-3-2 | 23.4064501 | 0.1724804 | 0.51522003 | 0.33477038 | 0.73779831 |
| tRNA-Pro-AGG-2-1 | 1.9991773 | -0.4393631 | 1.31796202 | -0.3333655 | 0.73885838 |
| tRNA-Val-TAC-1-1 | 296.977326 | 0.09745982 | 0.30132117 | 0.32344165 | 0.74636079 |
| tRNA-Ile-TAT-2-1 | 1.06638822 | -0.5528958 | 1.7730419 | -0.3118346 | 0.75516623 |
| tRNA-Ile-TAT-2-2 | 1.06638822 | -0.5528958 | 1.7730419 | -0.3118346 | 0.75516623 |
| tRNA-Val-TAC-1-2 | 294.870895 | 0.08997042 | 0.30075896 | 0.2991446 | 0.76482972 |
| tRNA-Val-TAC-1-3 | 294.870895 | 0.08997042 | 0.30075896 | 0.2991446 | 0.76482972 |
| tRNA-Cys-GCA-19-1 | 6.12119462 | 0.22203985 | 0.74383054 | 0.29850865 | 0.76531498 |
| tRNA-Ala-CGC-1-1 | 391.59381 | 0.10338094 | 0.35440566 | 0.29170229 | 0.77051426 |
| tRNA-Gln-CTG-3-1 | 309.696451 | -0.0899803 | 0.31588947 | -0.2848473 | 0.77576112 |
| tRNA-Ala-CGC-1-2 | 394.195456 | 0.0971661 | 0.34282334 | 0.28342907 | 0.77684794 |
| tRNA-Gln-CTG-3-2 | 309.662189 | -0.0893415 | 0.31585097 | -0.2828598 | 0.77728434 |
| tRNA-Gln-CTG-3-3 | 309.634521 | -0.088832 | 0.31600538 | -0.281109 | 0.77862678 |
| tRNA-Ser-CGA-2-1 | 183.99309 | -0.0876686 | 0.31800865 | -0.27568 | 0.78279385 |
| tRNA-Thr-AGT-3-1 | 14.0743672 | -0.1606694 | 0.58900347 | -0.2727818 | 0.78502098 |
| tRNA-Met-CAT-2-2 | 238.585464 | 0.12660681 | 0.4674369 | 0.27085328 | 0.78650388 |
| tRNA-iMet-CAT-2-1 | 53.4682505 | 0.07976577 | 0.29515592 | 0.2702496 | 0.78696824 |
| tRNA-Ala-TGC-8-1 | 340.79098 | 0.09460373 | 0.35320017 | 0.26784735 | 0.78881682 |
| tRNA-Met-CAT-2-1 | 238.96371 | 0.12474268 | 0.47005397 | 0.26537948 | 0.79071714 |
| tRNA-Ala-TGC-7-1 | 341.857216 | 0.09350749 | 0.35340024 | 0.26459375 | 0.79132244 |
| tRNA-Ala-TGC-7-2 | 341.857216 | 0.09350749 | 0.35340024 | 0.26459375 | 0.79132244 |
| tRNA-Ala-AGC-8-1 | 345.193346 | 0.08964835 | 0.35153321 | 0.25502101 | 0.79870687 |
| tRNA-Leu-CAG-4-1 | 27.9424695 | 0.13375091 | 0.52474763 | 0.25488616 | 0.79881103 |
| tRNA-Arg-ACG-1-1 | 13.1807709 | -0.1309049 | 0.51430667 | -0.2545269 | 0.79908849 |
| tRNA-Ala-TGC-2-1 | 532.573921 | 0.08664677 | 0.34760673 | 0.24926667 | 0.80315451 |
| tRNA-Arg-ACG-1-2 | 12.9566281 | -0.1320844 | 0.53106184 | -0.2487175 | 0.80357932 |
| tRNA-Cys-GCA-8-1 | 44.8066896 | -0.1152615 | 0.46455709 | -0.2481106 | 0.80404882 |
| tRNA-Arg-TCT-3-1 | 2.75609899 | 0.30109379 | 1.27729195 | 0.23572824 | 0.81364355 |
| tRNA-Ala-CGC-3-1 | 259.731206 | -0.0843186 | 0.35974461 | -0.2343846 | 0.81468643 |
| tRNA-Ala-CGC-3-2 | 259.731206 | -0.0843186 | 0.35974461 | -0.2343846 | 0.81468643 |
| tRNA-Ala-CGC-3-3 | 259.731206 | -0.0843186 | 0.35974461 | -0.2343846 | 0.81468643 |
| tRNA-Ala-CGC-6-1 | 259.620535 | -0.0828777 | 0.35964242 | -0.2304446 | 0.81774629 |
| tRNA-Arg-CCG-2-1 | 92.2280519 | 0.08241255 | 0.3687872 | 0.22346912 | 0.82317042 |
| tRNA-Ala-CGC-7-1 | 260.051064 | -0.0789126 | 0.36189668 | -0.2180529 | 0.82738789 |
| tRNA-Cys-GCA-13-1 | 21.9633135 | -0.0914763 | 0.43320735 | -0.2111606 | 0.83276198 |
| tRNA-Gln-CTG-2-1 | 313.078969 | -0.0699916 | 0.33201505 | -0.2108084 | 0.83303676 |
| tRNA-Gln-CTG-2-3 | 313.626515 | -0.0685251 | 0.33378492 | -0.205297 | 0.83734006 |
| tRNA-Lys-CTT-14-1 | 0.22859632 | 0.81272282 | 4.02580661 | 0.20187826 | 0.8400119 |
| tRNA-Asn-GTT-4-1 | 0.21611973 | 0.81273317 | 4.07458491 | 0.19946404 | 0.84189977 |
| tRNA-Ser-GGA-1-1 | 0.54460155 | -0.5754139 | 2.9472249 | -0.1952392 | 0.84520569 |
| tRNA-Met-CAT-6-1 | 1.81608629 | 0.24124756 | 1.2522594 | 0.19264983 | 0.84723322 |
| tRNA-Glu-CTC-5-1 | 4.82959767 | 0.27604551 | 1.43422138 | 0.19247064 | 0.84737356 |
| tRNA-Gln-CTG-2-2 | 314.041103 | -0.0632997 | 0.33153822 | -0.1909274 | 0.84858247 |
| tRNA-Gln-CTG-2-4 | 312.494498 | -0.0628866 | 0.33209355 | -0.189364 | 0.84980753 |
| tRNA-Asp-GTC-4-1 | 215.689033 | -0.0784881 | 0.41923623 | -0.1872168 | 0.85149068 |

| | | | | | |
|---|---|---|---|---|---|
| tRNA-Lys-TTT-3-1 | 0.26740619 | 0.81266744 | 4.37843773 | 0.18560671 | 0.85275319 |
| tRNA-Thr-AGT-1-2 | 14.1257781 | -0.1035573 | 0.56266298 | -0.1840486 | 0.85397533 |
| tRNA-Thr-AGT-1-3 | 14.1257781 | -0.1035573 | 0.56266298 | -0.1840486 | 0.85397533 |
| tRNA-Thr-AGT-1-1 | 14.1257781 | -0.1035573 | 0.56266298 | -0.1840486 | 0.85397533 |
| tRNA-iMet-CAT-4-1 | 0.11553966 | 0.81273216 | 4.54293138 | 0.17890038 | 0.85801592 |
| tRNA-Lys-TTT-5-1 | 24.1426889 | -0.0778806 | 0.4374143 | -0.1780478 | 0.85868545 |
| tRNA-Ala-TGC-4-1 | 365.042126 | 0.06095763 | 0.3505693 | 0.17388184 | 0.86195833 |
| tRNA-Thr-TGT-3-1 | 51.8670138 | -0.0593899 | 0.34507016 | -0.1721096 | 0.86335139 |
| tRNA-Arg-TCG-2-1 | 63.146195 | -0.0695812 | 0.40991646 | -0.1697448 | 0.86521082 |
| tRNA-Ala-CGC-2-1 | 443.307682 | 0.04705896 | 0.28773982 | 0.16354692 | 0.87008784 |
| tRNA-Ile-AAT-1-3 | 22.9496487 | 0.05646993 | 0.37746612 | 0.14960264 | 0.88107812 |
| tRNA-Ser-TGA-1-1 | 426.511732 | -0.0514629 | 0.35013853 | -0.1469786 | 0.88314892 |
| tRNA-Ala-AGC-3-1 | 359.507349 | 0.04865254 | 0.35253502 | 0.13800768 | 0.89023435 |
| tRNA-Cys-GCA-25-1 | 0.77105215 | -0.2659149 | 1.955812 | -0.1359614 | 0.89185182 |
| tRNA-Ser-AGA-2-1 | 1069.48828 | -0.0424161 | 0.32052419 | -0.1323334 | 0.89472056 |
| tRNA-Ser-AGA-1-1 | 1069.59806 | -0.0408206 | 0.32104903 | -0.1271476 | 0.8988236 |
| tRNA-Ser-TGA-2-1 | 1071.95997 | -0.0406907 | 0.3202325 | -0.1270661 | 0.8988881 |
| tRNA-Ala-TGC-5-3 | 2624.467 | 0.04902172 | 0.3920401 | 0.12504262 | 0.90048981 |
| tRNA-Ala-TGC-5-1 | 2624.57532 | 0.04893873 | 0.3920501 | 0.12482774 | 0.90065992 |
| tRNA-Ala-TGC-5-2 | 2624.57532 | 0.04893873 | 0.3920501 | 0.12482774 | 0.90065992 |
| tRNA-Ile-AAT-1-2 | 21.8607986 | -0.0486304 | 0.38991457 | -0.1247206 | 0.90074477 |
| tRNA-Ile-AAT-1-5 | 21.8607986 | -0.0486304 | 0.38991457 | -0.1247206 | 0.90074477 |
| tRNA-Ile-AAT-1-1 | 22.4318702 | -0.0486042 | 0.38971077 | -0.1247188 | 0.90074619 |
| tRNA-Ser-AGA-2-5 | 1066.67852 | -0.0399407 | 0.32197533 | -0.1240488 | 0.90127664 |
| tRNA-Ser-TGA-2-2 | 1069.79911 | -0.0397977 | 0.32105369 | -0.1239596 | 0.90134727 |
| tRNA-Ser-AGA-2-2 | 1066.4647 | -0.0399442 | 0.32248608 | -0.1238632 | 0.90142358 |
| tRNA-Ser-AGA-2-3 | 1066.21343 | -0.0399463 | 0.32279421 | -0.1237515 | 0.90151202 |
| tRNA-Ser-AGA-2-4 | 1066.21343 | -0.0399463 | 0.32279421 | -0.1237515 | 0.90151202 |
| tRNA-Ser-AGA-2-6 | 1066.21343 | -0.0399463 | 0.32279421 | -0.1237515 | 0.90151202 |
| tRNA-Phe-GAA-1-4 | 15.4291739 | 0.05775605 | 0.4697075 | 0.12296173 | 0.9021374 |
| tRNA-Ala-TGC-3-1 | 408.032316 | 0.04124727 | 0.35034436 | 0.11773352 | 0.90627881 |
| tRNA-Lys-CTT-1-1 | 19.2527981 | 0.07120256 | 0.63866147 | 0.11148716 | 0.91123004 |
| tRNA-Ile-AAT-4-1 | 3.20242932 | 0.08432536 | 0.85900033 | 0.09816686 | 0.9217998 |
| tRNA-Asp-GTC-3-1 | 1.85075566 | 0.1143158 | 1.50440685 | 0.07598729 | 0.93942921 |
| tRNA-Ile-AAT-1-6 | 22.6951187 | 0.0265253 | 0.38758096 | 0.06843808 | 0.94543691 |
| tRNA-Pro-AGG-3-1 | 7.72797027 | -0.0448071 | 0.68270853 | -0.0656314 | 0.94767126 |
| tRNA-Val-CAC-4-1 | 473.594272 | -0.0178954 | 0.27360037 | -0.0654072 | 0.94784979 |
| tRNA-Ile-GAT-1-1 | 0.15286675 | -0.287538 | 4.50622475 | -0.0638091 | 0.94912226 |
| tRNA-Gly-GCC-5-1 | 466.407171 | -0.0240239 | 0.38571136 | -0.0622846 | 0.9503362 |
| tRNA-Lys-CTT-6-1 | 1.4433289 | 0.08526025 | 1.37768696 | 0.06188652 | 0.95065321 |
| tRNA-Ala-AGC-14-1 | 0.05863278 | 0.24391234 | 4.52695925 | 0.05387995 | 0.95703081 |
| tRNA-Arg-TCT-5-1 | 0.24748819 | 0.19751438 | 3.90976678 | 0.0505182 | 0.95970945 |
| tRNA-Leu-TAG-3-1 | 121.013674 | -0.0136005 | 0.28002128 | -0.0485695 | 0.96126237 |
| tRNA-Arg-CCT-3-1 | 21.1317531 | 0.01971894 | 0.43723552 | 0.04509912 | 0.9640283 |
| tRNA-Gly-GCC-1-1 | 1698.42143 | 0.01101208 | 0.26560147 | 0.04146093 | 0.96692844 |
| tRNA-Gly-GCC-1-2 | 1698.42143 | 0.01101208 | 0.26560147 | 0.04146093 | 0.96692844 |

| | | | | | |
|---|---|---|---|---|---|
| tRNA-Gly-GCC-1-3 | 1698.42143 | 0.01101208 | 0.26560147 | 0.04146093 | 0.96692844 |
| tRNA-Pro-TGG-3-1 | 0.08630054 | -0.1604437 | 4.5109399 | -0.0355677 | 0.97162707 |
| tRNA-Gln-TTG-4-1 | 0.42758692 | 0.09693215 | 2.72616807 | 0.03555619 | 0.97163624 |
| tRNA-Arg-ACG-3-1 | 39.6139245 | -0.0170109 | 0.49356018 | -0.0344657 | 0.97250579 |
| tRNA-Pro-TGG-5-1 | 0.08300326 | -0.1202997 | 4.51251277 | -0.0266591 | 0.9787316 |
| tRNA-Ile-TAT-1-1 | 7.23774635 | -0.0171083 | 1.02131097 | -0.0167513 | 0.98663499 |
| tRNA-Cys-GCA-23-1 | 1.97794458 | -0.0174805 | 1.19062162 | -0.0146818 | 0.98828604 |
| tRNA-Thr-TGT-3-2 | 61.3470488 | 0.00315191 | 0.33346343 | 0.00945204 | 0.99245848 |
| tRNA-Trp-CCA-4-1 | 26.4858953 | 0.00378357 | 0.43651569 | 0.00866767 | 0.99308429 |
| tRNA-Trp-CCA-4-2 | 26.4858953 | 0.00378357 | 0.43651569 | 0.00866767 | 0.99308429 |
| tRNA-Cys-GCA-20-1 | 4.07688079 | -0.0045881 | 0.87401114 | -0.0052494 | 0.99581159 |

| padj |
| ---: |
| 6.38E-07 |
| 6.38E-07 |
| 6.38E-07 |
| 6.38E-07 |
| 6.38E-07 |
| 6.38E-07 |
| 6.38E-07 |
| 6.38E-07 |
| 0.0114037 |
| 0.01467027 |
| 0.01467027 |
| 0.01467027 |
| 0.01467027 |
| 0.01467027 |
| 0.01467027 |
| 0.01467027 |
| 0.02406051 |
| 0.02406051 |
| 0.02526183 |
| 0.02695943 |
| 0.03562109 |
| 0.03683659 |
| 0.04926387 |
| 0.0601961 |
| 0.08156286 |
| 0.08673126 |
| 0.08871807 |
| 0.08871807 |
| 0.11190359 |
| 0.11509897 |
| 0.11509897 |
| 0.11509897 |
| 0.11557538 |
| 0.1168281 |
| 0.13798784 |
| 0.13934101 |
| 0.13934101 |
| 0.13934101 |
| 0.14580909 |
| 0.14580909 |
| 0.14580909 |
| 0.14580909 |
| 0.16218368 |
| 0.16218368 |

| |
|---|
| 0.16218368 |
| 0.16774189 |
| 0.16827951 |
| 0.22594318 |
| 0.27051464 |
| 0.31657441 |
| 0.32668831 |
| 0.33753355 |
| 0.36400906 |
| 0.36400906 |
| 0.36400906 |
| 0.39369642 |
| 0.42118716 |
| 0.45254959 |
| 0.46406391 |
| 0.46453568 |
| 0.48358488 |
| 0.4977261 |
| 0.51447346 |
| 0.51619956 |
| 0.56696649 |
| 0.5883903 |
| 0.59393044 |
| 0.60162701 |
| 0.60162701 |
| 0.60162701 |
| 0.60477122 |
| 0.61268039 |
| 0.61289915 |
| 0.61289915 |
| 0.62116978 |
| 0.63257289 |
| 0.63257289 |
| 0.63257289 |
| 0.63257289 |
| 0.63257289 |
| 0.63257289 |
| 0.63257289 |
| 0.63257289 |
| 0.65328691 |
| 0.65640219 |
| 0.68244189 |
| 0.70365923 |
| 0.73236141 |
| 0.73236141 |
| 0.73236141 |

| |
|---|
| 0.73236141 |
| 0.73236141 |
| 0.73236141 |
| 0.73236141 |
| 0.73236141 |
| 0.73236141 |
| 0.73236141 |
| 0.73236141 |
| 0.75696568 |
| 0.75696568 |
| 0.75696568 |
| 0.75696568 |
| 0.75696568 |
| 0.77291435 |
| 0.77302235 |
| 0.81500746 |
| 0.81500746 |
| 0.81500746 |
| 0.82769611 |
| 0.83161507 |
| 0.8321784 |
| 0.83603138 |
| 0.83603138 |
| 0.85970404 |
| 0.85970404 |
| 0.85970404 |
| 0.86048215 |
| 0.86151773 |
| 0.86151773 |
| 0.86151773 |
| 0.86151773 |
| 0.86151773 |
| 0.86151773 |
| 0.87713396 |
| 0.87713396 |
| 0.87713396 |
| 0.9194749 |
| 0.9194749 |
| 0.9194749 |
| 0.9194749 |
| 0.9194749 |
| 0.9194749 |
| 0.9194749 |
| 0.92044836 |
| 0.92044836 |
| 0.92044836 |

0.92044836
0.92044836
0.92044836
0.92566451
0.93531299
0.93531299
0.93531299
0.93531299
0.93531299
0.93531299
0.93531299
0.93531299
0.93531299
0.93531299
0.93531299
0.93531299
0.93531299
0.94210264
0.94210264
0.94210264
0.94210264
0.95718513
0.95718513
0.96134272
0.96364677
0.96364677
0.96364677
0.96364677
0.96364677
0.96364677
0.96364677
0.96364677
0.96364677
0.96364677
0.96364677
0.96364677
0.96364677
0.96364677
0.96364677
0.96364677
0.96364677
0.96364677
0.96364677
0.96364677
0.96364677
0.96364677

0.96364677
0.96364677
0.96364677
0.96364677
0.96364677
0.96364677
0.96364677
0.96364677
0.96364677
0.96364677
0.96364677
0.96364677
0.96364677
0.96364677
0.96364677
0.96364677
0.96364677
0.96364677
0.96364677
0.96364677
0.96364677
0.96364677
0.96364677
0.96364677
0.96364677
0.96364677
0.96364677
0.96364677
0.96364677
0.96364677
0.96364677
0.96364677
0.96364677
0.96364677
0.96364677
0.96364677
0.96364677
0.96364677
0.96364677
0.96364677
0.96364677
0.96364677
0.96364677
0.96364677
0.96364677
0.96364677

0.96364677
0.96364677
0.96364677
0.96364677
0.96364677
0.96364677
0.96364677
0.96364677
0.96364677
0.96364677
0.96364677
0.96364677
0.96364677
0.96364677
0.96364677
0.96364677
0.96364677
0.96364677
0.96364677
0.96364677
0.96364677
0.96364677
0.96364677
0.96364677
0.96364677
0.96364677
0.96364677
0.96364677
0.96364677
0.96364677
0.96364677
0.96364677
0.96364677
0.96364677
0.96364677
0.96364677
0.96364677
0.96364677
0.96364677
0.96364677
0.96364677
0.96364677
0.96364677
0.96364677
0.96364677
0.96364677

| |
|---|
| 0.96364677 |
| 0.96364677 |
| 0.96364677 |
| 0.96364677 |
| 0.96364677 |
| 0.96364677 |
| 0.96364677 |
| 0.96364677 |
| 0.96364677 |
| 0.96364677 |
| 0.96364677 |
| 0.96364677 |
| 0.96364677 |
| 0.96364677 |
| 0.96364677 |
| 0.96364677 |
| 0.96364677 |
| 0.96364677 |
| 0.96364677 |
| 0.96364677 |
| 0.96364677 |
| 0.96364677 |
| 0.96364677 |
| 0.96364677 |
| 0.96364677 |
| 0.96364677 |
| 0.96364677 |
| 0.96364677 |
| 0.96364677 |
| 0.96364677 |
| 0.96364677 |
| 0.96364677 |
| 0.96364677 |
| 0.96364677 |
| 0.96364677 |
| 0.96364677 |
| 0.96364677 |
| 0.96364677 |
| 0.96364677 |
| 0.96364677 |
| 0.96364677 |
| 0.96364677 |
| 0.96364677 |
| 0.96364677 |
| 0.96364677 |
| 0.96364677 |

0.96364677
0.96364677
0.96364677
0.96364677
0.96364677
0.96364677
0.96364677
0.96364677
0.96364677
0.96364677
0.96364677
0.96364677
0.96364677
0.96364677
0.96364677
0.96364677
0.96364677
0.96364677
0.96364677
0.96364677
0.96364677
0.96364677
0.96364677
0.96364677
0.96364677
0.96364677
0.96364677
0.96364677
0.96364677
0.96364677
0.96364677
0.96364677
0.96364677
0.96364677
0.96364677
0.96364677
0.96364677
0.96364677
0.96364677
0.96364677
0.96364677
0.96364677
0.96364677
0.96364677
0.96364677
0.96364677

| |
|---|
| 0.96364677 |
| 0.96364677 |
| 0.96364677 |
| 0.96364677 |
| 0.96364677 |
| 0.96364677 |
| 0.96364677 |
| 0.96364677 |
| 0.96364677 |
| 0.96364677 |
| 0.96364677 |
| 0.96364677 |
| 0.96364677 |
| 0.96364677 |
| 0.96364677 |
| 0.96364677 |
| 0.96364677 |
| 0.96364677 |
| 0.96364677 |
| 0.96364677 |
| 0.96364677 |
| 0.96364677 |
| 0.96364677 |
| 0.96364677 |
| 0.96364677 |
| 0.96364677 |
| 0.96364677 |
| 0.96364677 |
| 0.96364677 |
| 0.96364677 |
| 0.96563208 |
| 0.96846811 |
| 0.9772464 |
| 0.98887007 |
| 0.98887007 |
| 0.98887007 |
| 0.98887007 |
| 0.98887007 |
| 0.98887007 |
| 0.98887007 |
| 0.98887007 |
| 0.98887007 |
| 0.98887007 |
| 0.98887007 |
| 0.98887007 |
| 0.98887007 |

0.98887007
0.98887007
0.98887007
0.98887007
0.99281407
0.99543757
0.99543757
0.99543757
0.99543757
0.99543757
0.99581159

| HS692_tRNA_pvalues_TIME_padj05 | | | HS692_tRNA_pvalues_TIME_padj05 | | |
|---|---|---|---|---|---|
| | baseMean | log2FoldChai | lfcSE | stat | pvalue |
| tRNA-His-GTG-2-7 | 28999.5299 | 2.45371014 | 0.2200948 | 11.1484238 | 7.29E-29 |
| tRNA-His-GTG-2-4 | 28998.4942 | 2.4537478 | 0.22015774 | 11.145408 | 7.54E-29 |
| tRNA-His-GTG-2-5 | 28998.4942 | 2.4537478 | 0.22015774 | 11.145408 | 7.54E-29 |
| tRNA-His-GTG-2-6 | 28998.4942 | 2.4537478 | 0.22015774 | 11.145408 | 7.54E-29 |
| tRNA-His-GTG-2-1 | 29001.8773 | 2.45338628 | 0.22015848 | 11.143728 | 7.68E-29 |
| tRNA-His-GTG-2-3 | 29001.8773 | 2.45338628 | 0.22015848 | 11.143728 | 7.68E-29 |
| tRNA-His-GTG-2-2 | 29001.8918 | 2.45338628 | 0.22015889 | 11.1437076 | 7.69E-29 |
| tRNA-His-GTG-2-8 | 29001.817 | 2.45338628 | 0.22015893 | 11.1437057 | 7.69E-29 |
| tRNA-His-GTG-3-1 | 196.814482 | 1.99598207 | 0.22931407 | 8.70414118 | 3.20E-18 |
| tRNA-Val-CAC-6-1 | 8198.89709 | 1.96707875 | 0.2626104 | 7.49048307 | 6.86E-14 |
| tRNA-Val-AAC-1-1 | 454079.297 | 1.7721316 | 0.25216372 | 7.0277024 | 2.10E-12 |
| tRNA-Val-AAC-1-2 | 454075.646 | 1.77211979 | 0.25216281 | 7.02768097 | 2.10E-12 |
| tRNA-Val-CAC-2-2 | 459699.312 | 1.76632038 | 0.25258183 | 6.99306193 | 2.69E-12 |
| tRNA-Val-CAC-2-1 | 459699.535 | 1.76632638 | 0.25258327 | 6.99304592 | 2.69E-12 |
| tRNA-Val-CAC-2-5 | 459699.745 | 1.76632172 | 0.25258334 | 6.99302543 | 2.69E-12 |
| tRNA-Val-CAC-2-3 | 459702.447 | 1.76632139 | 0.2525851 | 6.99297538 | 2.69E-12 |
| tRNA-Val-CAC-2-4 | 459696.983 | 1.76631089 | 0.25258566 | 6.99291825 | 2.69E-12 |
| tRNA-Gly-CCC-1-2 | 1464.64312 | -1.6660624 | 0.24590258 | -6.7752944 | 1.24E-11 |
| tRNA-Gly-CCC-1-1 | 1464.64312 | -1.6660624 | 0.24590258 | -6.7752944 | 1.24E-11 |
| tRNA-Leu-TAG-1-1 | 1006.84179 | -1.5679478 | 0.23247396 | -6.7446171 | 1.53E-11 |
| tRNA-Leu-AAG-1-1 | 1011.72894 | -1.5686929 | 0.23694963 | -6.6203643 | 3.58E-11 |
| tRNA-Leu-AAG-1-2 | 1011.53652 | -1.5687038 | 0.23731716 | -6.6101575 | 3.84E-11 |
| tRNA-Leu-AAG-1-3 | 1011.657 | -1.5687073 | 0.23743282 | -6.6069521 | 3.92E-11 |
| tRNA-Val-AAC-5-1 | 33823.3676 | 1.68313985 | 0.26052971 | 6.46045267 | 1.04E-10 |
| tRNA-Pro-AGG-1-1 | 498.548964 | 1.39188484 | 0.23352419 | 5.96034535 | 2.52E-09 |
| tRNA-Pro-AGG-1-6 | 498.174598 | 1.37424836 | 0.2316316 | 5.93290526 | 2.98E-09 |
| tRNA-Pro-AGG-1-5 | 474.180973 | 1.34398722 | 0.23572569 | 5.70148822 | 1.19E-08 |
| tRNA-Pro-AGG-1-2 | 472.481965 | 1.33175338 | 0.23369358 | 5.69871627 | 1.21E-08 |
| tRNA-Pro-AGG-1-4 | 479.374246 | 1.33564124 | 0.23559489 | 5.66922836 | 1.43E-08 |
| tRNA-Pro-AGG-1-3 | 474.29805 | 1.32745207 | 0.23464464 | 5.65728701 | 1.54E-08 |
| tRNA-Pro-TGG-2-2 | 501.388162 | 1.25910603 | 0.22668411 | 5.55445204 | 2.78E-08 |
| tRNA-Pro-CGG-1-1 | 571.437924 | 1.300867 | 0.23436597 | 5.55057964 | 2.85E-08 |
| tRNA-Pro-TGG-2-1 | 502.629255 | 1.26320649 | 0.22815335 | 5.53665539 | 3.08E-08 |
| tRNA-Pro-TGG-2-4 | 500.779936 | 1.24624499 | 0.22662957 | 5.49903975 | 3.82E-08 |
| tRNA-Pro-CGG-1-3 | 563.110855 | 1.28367781 | 0.23446405 | 5.4749452 | 4.38E-08 |
| tRNA-Pro-TGG-2-3 | 499.10422 | 1.24649707 | 0.22781888 | 5.47143889 | 4.46E-08 |
| tRNA-Glu-TTC-1-4 | 7540.22739 | 1.34244609 | 0.24841157 | 5.40412069 | 6.51E-08 |
| tRNA-Glu-TTC-1-1 | 7533.07318 | 1.34275661 | 0.24852193 | 5.40297027 | 6.55E-08 |
| tRNA-Glu-TTC-1-2 | 7533.07318 | 1.34275661 | 0.24852193 | 5.40297027 | 6.55E-08 |
| tRNA-Glu-TTC-1-3 | 7516.66951 | 1.34668864 | 0.2494573 | 5.39847367 | 6.72E-08 |
| tRNA-Val-CAC-4-1 | 473.594272 | 1.31952827 | 0.24585979 | 5.36699504 | 8.01E-08 |
| tRNA-Pro-CGG-1-2 | 574.314626 | 1.26105413 | 0.23549581 | 5.35488987 | 8.56E-08 |
| tRNA-Ala-CGC-3-1 | 259.731206 | -1.772053 | 0.33482825 | -5.2924239 | 1.21E-07 |
| tRNA-Ala-CGC-3-2 | 259.731206 | -1.772053 | 0.33482825 | -5.2924239 | 1.21E-07 |

| | | | | | |
|---|---|---|---|---|---|
| tRNA-Ala-CGC-3-3 | 259.731206 | -1.772053 | 0.33482825 | -5.2924239 | 1.21E-07 |
| tRNA-Ala-CGC-6-1 | 259.620535 | -1.7706068 | 0.33473429 | -5.2895889 | 1.23E-07 |
| tRNA-Ala-CGC-7-1 | 260.051064 | -1.7715499 | 0.33682689 | -5.2595263 | 1.44E-07 |
| tRNA-Glu-TTC-2-2 | 5698.90117 | 1.2016632 | 0.23780166 | 5.05321627 | 4.34E-07 |
| tRNA-Glu-TTC-2-1 | 5697.21165 | 1.20098439 | 0.23851117 | 5.03533812 | 4.77E-07 |
| tRNA-Leu-CAA-1-1 | 79.2749203 | -1.3975763 | 0.28360761 | -4.9278519 | 8.31E-07 |
| tRNA-Leu-TAA-2-1 | 3119.15501 | -1.6690116 | 0.34074064 | -4.8981876 | 9.67E-07 |
| tRNA-Leu-CAA-4-1 | 81.1825193 | -1.420824 | 0.29144246 | -4.8751442 | 1.09E-06 |
| tRNA-Leu-TAG-3-1 | 121.013674 | -1.2610876 | 0.25950322 | -4.8596222 | 1.18E-06 |
| tRNA-Ala-TGC-8-1 | 340.79098 | -1.5703267 | 0.32828886 | -4.783369 | 1.72E-06 |
| tRNA-Ala-TGC-7-1 | 341.857216 | -1.5704857 | 0.32846703 | -4.7812582 | 1.74E-06 |
| tRNA-Ala-TGC-7-2 | 341.857216 | -1.5704857 | 0.32846703 | -4.7812582 | 1.74E-06 |
| tRNA-Ala-TGC-5-3 | 2624.467 | -1.7288803 | 0.36315621 | -4.760707 | 1.93E-06 |
| tRNA-Ala-TGC-5-1 | 2624.57532 | -1.7287572 | 0.36316538 | -4.7602478 | 1.93E-06 |
| tRNA-Ala-TGC-5-2 | 2624.57532 | -1.7287572 | 0.36316538 | -4.7602478 | 1.93E-06 |
| tRNA-Ser-TGA-1-1 | 426.511732 | -1.5356624 | 0.32478642 | -4.7282224 | 2.26E-06 |
| tRNA-Ala-TGC-4-1 | 365.042126 | -1.5370792 | 0.32560124 | -4.7207413 | 2.35E-06 |
| tRNA-Pro-TGG-1-1 | 358.920056 | 1.03126212 | 0.22247821 | 4.63533995 | 3.56E-06 |
| tRNA-Ser-AGA-2-1 | 1069.48828 | -1.3759598 | 0.29687455 | -4.6348191 | 3.57E-06 |
| tRNA-Ala-AGC-8-1 | 345.193346 | -1.5111174 | 0.32654761 | -4.6275561 | 3.70E-06 |
| tRNA-Ser-AGA-1-1 | 1069.59806 | -1.3718942 | 0.29735781 | -4.6136141 | 3.96E-06 |
| tRNA-Ser-TGA-2-1 | 1071.95997 | -1.3680391 | 0.29659824 | -4.6124316 | 3.98E-06 |
| tRNA-Ser-TGA-2-2 | 1069.79911 | -1.3714789 | 0.29736269 | -4.6121418 | 3.99E-06 |
| tRNA-Ser-AGA-2-5 | 1066.67852 | -1.3731368 | 0.29821751 | -4.6044808 | 4.13E-06 |
| tRNA-Ser-AGA-2-2 | 1066.4647 | -1.3731464 | 0.2986902 | -4.5972262 | 4.28E-06 |
| tRNA-Ser-AGA-2-3 | 1066.21343 | -1.3731522 | 0.29897536 | -4.5928607 | 4.37E-06 |
| tRNA-Ser-AGA-2-4 | 1066.21343 | -1.3731522 | 0.29897536 | -4.5928607 | 4.37E-06 |
| tRNA-Ser-AGA-2-6 | 1066.21343 | -1.3731522 | 0.29897536 | -4.5928607 | 4.37E-06 |
| tRNA-Ala-TGC-3-1 | 408.032316 | -1.4787354 | 0.32509303 | -4.5486531 | 5.40E-06 |
| tRNA-Ala-AGC-3-1 | 359.507349 | -1.4806917 | 0.32723704 | -4.5248292 | 6.04E-06 |
| tRNA-Arg-CCT-4-1 | 107.841334 | -1.3413556 | 0.30079781 | -4.4593264 | 8.22E-06 |
| tRNA-Ala-TGC-6-1 | 593.053412 | -1.537414 | 0.3463575 | -4.4388066 | 9.05E-06 |
| tRNA-Leu-CAA-3-1 | 144.22626 | -1.5643203 | 0.35300642 | -4.4314216 | 9.36E-06 |
| tRNA-Ala-CGC-1-2 | 394.195456 | -1.4050417 | 0.31812245 | -4.4166694 | 1.00E-05 |
| tRNA-Ala-CGC-1-1 | 391.59381 | -1.4480008 | 0.32894626 | -4.4019373 | 1.07E-05 |
| tRNA-Val-AAC-3-1 | 3954.19939 | 1.1380247 | 0.27207971 | 4.18268854 | 2.88E-05 |
| tRNA-Arg-TCG-1-1 | 61.4602617 | -1.0561749 | 0.25320595 | -4.1712087 | 3.03E-05 |
| tRNA-Val-CAC-3-1 | 4033.67882 | 1.13299771 | 0.27185048 | 4.16772383 | 3.08E-05 |
| tRNA-Phe-GAA-1-3 | 14.6979462 | -1.8849571 | 0.45426105 | -4.149502 | 3.33E-05 |
| tRNA-SeC-TCA-1-1 | 10932.2345 | -0.9846866 | 0.23785867 | -4.1397969 | 3.48E-05 |
| tRNA-Val-AAC-2-1 | 3983.63556 | 1.11766408 | 0.27122651 | 4.12077743 | 3.78E-05 |
| tRNA-Val-AAC-2-2 | 3983.63556 | 1.11766408 | 0.27122651 | 4.12077743 | 3.78E-05 |
| tRNA-Leu-TAG-2-1 | 796.037019 | -1.052781 | 0.25549675 | -4.120526 | 3.78E-05 |
| tRNA-Leu-AAG-2-1 | 811.615521 | -1.0605906 | 0.25875679 | -4.0987935 | 4.15E-05 |
| tRNA-Leu-AAG-3-1 | 798.841555 | -1.0666081 | 0.2616982 | -4.0757181 | 4.59E-05 |
| tRNA-Phe-GAA-1-2 | 14.5147963 | -1.8081621 | 0.45280145 | -3.9932781 | 6.52E-05 |

| | | | | | |
|---|---|---|---|---|---|
| tRNA-Ser-CGA-2-1 | 183.99309 | -1.1627363 | 0.29370752 | -3.9588236 | 7.53E-05 |
| tRNA-Cys-GCA-3-4 | 3170.81706 | 0.9112946 | 0.23074598 | 3.94934118 | 7.84E-05 |
| tRNA-Cys-GCA-1-1 | 3162.87586 | 0.91435171 | 0.23180047 | 3.94456374 | 7.99E-05 |
| tRNA-Cys-GCA-3-1 | 3172.77916 | 0.91037399 | 0.23091722 | 3.94242566 | 8.07E-05 |
| tRNA-Cys-GCA-3-3 | 3185.20845 | 0.91066963 | 0.2309994 | 3.94230307 | 8.07E-05 |
| tRNA-Cys-GCA-3-2 | 3185.66778 | 0.91047068 | 0.23099803 | 3.94146518 | 8.10E-05 |
| tRNA-Cys-GCA-2-1 | 3180.73817 | 0.91301943 | 0.23182355 | 3.93842389 | 8.20E-05 |
| tRNA-Cys-GCA-1-2 | 3179.62572 | 0.91452913 | 0.2324833 | 3.93374118 | 8.36E-05 |
| tRNA-Met-CAT-1-1 | 279.447803 | -1.4210434 | 0.373759 | -3.8020312 | 0.00014351 |
| tRNA-Gly-CCC-2-1 | 3966.66658 | 1.37975608 | 0.36340584 | 3.79673608 | 0.00014661 |
| tRNA-Gly-CCC-2-2 | 3966.66658 | 1.37975608 | 0.36340584 | 3.79673608 | 0.00014661 |
| tRNA-Lys-CTT-3-2 | 3876.20711 | 0.89958878 | 0.23816696 | 3.77713516 | 0.00015864 |
| tRNA-Lys-CTT-3-5 | 3828.89617 | 0.90906534 | 0.24131288 | 3.76716459 | 0.00016511 |
| tRNA-Lys-CTT-3-6 | 3828.89617 | 0.90906534 | 0.24131288 | 3.76716459 | 0.00016511 |
| tRNA-Met-CAT-1-2 | 282.708567 | -1.3807438 | 0.36654282 | -3.7669372 | 0.00016526 |
| tRNA-Lys-CTT-3-7 | 3829.01173 | 0.90884839 | 0.24127573 | 3.76684544 | 0.00016532 |
| tRNA-Lys-TTT-5-1 | 24.1426889 | -1.5495743 | 0.4118972 | -3.7620415 | 0.00016853 |
| tRNA-Ala-CGC-2-1 | 443.307682 | -0.9964112 | 0.26600902 | -3.7457798 | 0.00017983 |
| tRNA-Phe-GAA-1-4 | 15.4291739 | -1.6957444 | 0.45564159 | -3.7216629 | 0.00019792 |
| tRNA-Lys-CTT-3-4 | 3854.03886 | 0.89449225 | 0.24125908 | 3.70760043 | 0.00020923 |
| tRNA-Lys-CTT-3-3 | 3858.00155 | 0.89479782 | 0.24156017 | 3.70424407 | 0.00021202 |
| tRNA-Lys-CTT-3-1 | 3852.60864 | 0.89415065 | 0.24139949 | 3.70402871 | 0.0002122 |
| tRNA-Gly-TCC-1-2 | 390.66018 | -1.3630664 | 0.37134264 | -3.6706433 | 0.00024194 |
| tRNA-Gly-TCC-1-3 | 390.66018 | -1.3630664 | 0.37134264 | -3.6706433 | 0.00024194 |
| tRNA-Gly-TCC-1-4 | 390.66018 | -1.3630664 | 0.37134264 | -3.6706433 | 0.00024194 |
| tRNA-Phe-GAA-1-5 | 15.7564053 | -1.5656209 | 0.42781388 | -3.6595841 | 0.00025262 |
| tRNA-Gly-TCC-1-7 | 393.953649 | -1.3492898 | 0.36926827 | -3.6539555 | 0.00025823 |
| tRNA-Ser-CGA-3-1 | 53.5815648 | -1.4024433 | 0.38400938 | -3.6521068 | 0.0002601 |
| tRNA-Gly-TCC-1-1 | 398.738556 | -1.3242044 | 0.36366093 | -3.6413162 | 0.00027125 |
| tRNA-Val-CAC-1-1 | 2394.51039 | 1.03769834 | 0.2906513 | 3.57025181 | 0.00035664 |
| tRNA-Ser-CGA-1-1 | 54.8551022 | -1.3279051 | 0.37289106 | -3.5611073 | 0.00036929 |
| tRNA-Gly-TCC-1-5 | 394.166849 | -1.3164074 | 0.37104203 | -3.5478659 | 0.00038837 |
| tRNA-Ser-GCT-6-1 | 12.6153113 | -1.8764039 | 0.53680101 | -3.4955297 | 0.00047312 |
| tRNA-Thr-CGT-2-1 | 49.5938764 | -1.4449221 | 0.41499045 | -3.4818202 | 0.00049802 |
| tRNA-Arg-CCG-2-1 | 92.2280519 | -1.1584569 | 0.34168108 | -3.3904624 | 0.00069775 |
| tRNA-Leu-CAA-2-1 | 362.290665 | -1.5524937 | 0.47218876 | -3.2878667 | 0.0010095 |
| tRNA-Val-AAC-4-1 | 1646.3544 | 1.32118382 | 0.40973212 | 3.22450639 | 0.0012619 |
| tRNA-Ala-TGC-2-1 | 532.573921 | -1.0264435 | 0.3216777 | -3.1909065 | 0.00141827 |
| tRNA-Ser-GCT-3-1 | 1110.78244 | -1.0855243 | 0.34167637 | -3.177054 | 0.00148779 |
| tRNA-Ser-GCT-2-1 | 1106.56683 | -1.0973796 | 0.34600497 | -3.1715717 | 0.00151616 |
| tRNA-Gly-CCC-5-1 | 81882.7096 | 1.01462141 | 0.3249964 | 3.12194661 | 0.0017966 |
| tRNA-Ser-GCT-4-1 | 1098.54502 | -1.0849395 | 0.34804542 | -3.1172354 | 0.00182556 |
| tRNA-Ser-GCT-4-2 | 1098.30062 | -1.0847083 | 0.3480928 | -3.1161469 | 0.00183231 |
| tRNA-Gly-GCC-3-1 | 82830.6429 | 0.99596208 | 0.32057639 | 3.10678553 | 0.00189134 |
| tRNA-Gly-GCC-4-1 | 82804.0993 | 0.9938382 | 0.32019455 | 3.10385729 | 0.00191015 |
| tRNA-Gly-GCC-2-8 | 82771.4561 | 0.99459065 | 0.32049773 | 3.10326896 | 0.00191396 |

| | | | | | |
|---|---|---|---|---|---|
| tRNA-Gly-GCC-2-7 | 82764.1107 | 0.99448145 | 0.3204706 | 3.1031909 | 0.00191446 |
| tRNA-Gly-GCC-2-4 | 82742.828 | 0.99431129 | 0.3205235 | 3.10214784 | 0.00192122 |
| tRNA-Gly-GCC-2-5 | 82743.8984 | 0.99429798 | 0.32052917 | 3.10205142 | 0.00192185 |
| tRNA-Gly-GCC-2-6 | 82751.08 | 0.99415212 | 0.32052044 | 3.10168089 | 0.00192425 |
| tRNA-Gly-GCC-2-1 | 82739.3231 | 0.99413167 | 0.32052139 | 3.10160789 | 0.00192473 |
| tRNA-Gly-GCC-2-2 | 82743.4953 | 0.99408686 | 0.32054239 | 3.10126485 | 0.00192696 |
| tRNA-Gly-GCC-2-3 | 82825.9032 | 0.99248045 | 0.32013658 | 3.10017821 | 0.00193404 |
| tRNA-Ser-GCT-4-3 | 1102.79447 | -1.0714936 | 0.34826976 | -3.0766197 | 0.00209362 |
| tRNA-Asn-GTT-2-1 | 9.19200847 | -1.7253892 | 0.56401811 | -3.0591026 | 0.00222001 |
| tRNA-Ser-GCT-1-1 | 1032.90289 | -1.0710133 | 0.3611294 | -2.9657328 | 0.00301963 |
| tRNA-Gly-GCC-6-1 | 530.397599 | 1.02461587 | 0.3480903 | 2.94353467 | 0.00324487 |
| tRNA-Trp-CCA-3-2 | 40.7546967 | 0.94424232 | 0.32622493 | 2.89445176 | 0.00379821 |
| tRNA-Cys-GCA-16-1 | 19.0894496 | 1.29715992 | 0.44944118 | 2.88616168 | 0.00389972 |
| tRNA-Met-CAT-3-1 | 30.246461 | -1.1588702 | 0.40361682 | -2.8712138 | 0.00408899 |
| tRNA-Glu-TTC-3-1 | 144.563688 | 0.940333 | 0.3296279 | 2.85271054 | 0.00433481 |
| tRNA-Glu-TTC-3-2 | 144.563688 | 0.940333 | 0.3296279 | 2.85271054 | 0.00433481 |
| tRNA-Met-CAT-4-1 | 9.82922744 | -2.0433677 | 0.71707753 | -2.8495771 | 0.00437774 |
| tRNA-Leu-TAA-1-1 | 54.3850394 | -1.1877309 | 0.41726339 | -2.8464775 | 0.00442058 |
| tRNA-Asn-GTT-1-1 | 15.3438029 | -1.3227478 | 0.46754684 | -2.8291235 | 0.00466757 |
| tRNA-Leu-CAG-3-1 | 10.3740117 | -1.3825704 | 0.49637496 | -2.7853346 | 0.00534725 |
| tRNA-Pro-TGG-4-1 | 196.596347 | 0.99592453 | 0.35906513 | 2.77365985 | 0.00554296 |
| tRNA-Met-CAT-2-2 | 238.585464 | -1.1904347 | 0.43309588 | -2.7486633 | 0.00598388 |
| tRNA-Met-CAT-2-1 | 238.96371 | -1.1961616 | 0.43552796 | -2.7464635 | 0.00602416 |
| tRNA-Thr-CGT-1-1 | 20.4722709 | -1.5099566 | 0.56052321 | -2.6938342 | 0.00706353 |
| tRNA-Ile-AAT-1-4 | 22.5194002 | -0.9382377 | 0.35020964 | -2.6790745 | 0.0073826 |
| tRNA-Thr-AGT-6-1 | 1.50954656 | 4.60409194 | 1.72391988 | 2.67071109 | 0.00756908 |
| tRNA-Gly-TCC-1-6 | 427.670255 | -1.0019205 | 0.37729951 | -2.6555043 | 0.00791899 |
| tRNA-Gln-CTG-1-1 | 241.967211 | -0.6292222 | 0.23946845 | -2.6275789 | 0.00859949 |
| tRNA-Gly-CCC-4-1 | 1065.83419 | 0.87186824 | 0.33768053 | 2.58193226 | 0.00982489 |
| tRNA-Asp-GTC-2-1 | 1891.71821 | 0.51347062 | 0.19902558 | 2.57992274 | 0.00988224 |
| tRNA-Gly-CCC-3-1 | 1073.04047 | 0.8616697 | 0.33492832 | 2.57269887 | 0.0100909 |
| tRNA-Asn-GTT-3-6 | 24.7445466 | -0.9734001 | 0.38093421 | -2.5552973 | 0.01060971 |
| tRNA-Phe-GAA-3-1 | 8.07769192 | -1.472063 | 0.5805212 | -2.5357609 | 0.01122033 |
| tRNA-Gln-CTG-5-1 | 85.5342562 | 1.09178619 | 0.4312948 | 2.53141513 | 0.01136033 |
| tRNA-iMet-CAT-1-7 | 1318.84617 | 1.087981 | 0.43896673 | 2.47850449 | 0.01319344 |
| tRNA-iMet-CAT-1-6 | 1323.30472 | 1.08966118 | 0.43964938 | 2.47847768 | 0.01319444 |
| tRNA-Arg-CCT-3-1 | 21.1317531 | -0.9840064 | 0.39762495 | -2.47471 | 0.01333444 |
| tRNA-iMet-CAT-3-1 | 1272.12042 | 1.13900107 | 0.46075859 | 2.47201263 | 0.01343548 |
| tRNA-iMet-CAT-1-5 | 1316.68596 | 1.08646936 | 0.43958059 | 2.4716045 | 0.01345082 |
| tRNA-iMet-CAT-1-2 | 1316.71192 | 1.08594098 | 0.43968949 | 2.46979064 | 0.01351921 |
| tRNA-iMet-CAT-1-4 | 1316.71192 | 1.08594098 | 0.43968949 | 2.46979064 | 0.01351921 |
| tRNA-iMet-CAT-1-3 | 1325.69778 | 1.08393615 | 0.43944891 | 2.46658057 | 0.013641 |
| tRNA-iMet-CAT-1-1 | 1324.81683 | 1.08053723 | 0.43834492 | 2.46503881 | 0.01369984 |
| tRNA-Ile-AAT-1-3 | 22.9496487 | -0.8364728 | 0.33988063 | -2.4610781 | 0.01385202 |
| tRNA-Ile-AAT-1-2 | 21.8607986 | -0.8515955 | 0.3482968 | -2.4450281 | 0.01448408 |
| tRNA-Ile-AAT-1-5 | 21.8607986 | -0.8515955 | 0.3482968 | -2.4450281 | 0.01448408 |

| | | | | | |
|---|---|---|---|---|---|
| tRNA-Arg-CCG-1-1 | 24.2194041 | -0.9539753 | 0.39547222 | -2.4122435 | 0.01585469 |
| tRNA-Met-CAT-5-1 | 8.28216092 | -1.4891443 | 0.61851651 | -2.4076065 | 0.01605748 |
| tRNA-Gln-CTG-4-1 | 120.62929 | 0.83106425 | 0.34896417 | 2.3815174 | 0.01724148 |
| tRNA-Ile-AAT-1-6 | 22.6951187 | -0.8251492 | 0.34803208 | -2.3708997 | 0.01774484 |
| tRNA-Asn-GTT-3-3 | 23.967007 | -0.9833728 | 0.41690035 | -2.3587718 | 0.01833553 |
| tRNA-Asn-GTT-3-9 | 24.368177 | -0.9622281 | 0.41114034 | -2.3403885 | 0.01926369 |
| tRNA-Asn-GTT-3-4 | 24.0922481 | -0.949762 | 0.41148327 | -2.3081425 | 0.02099121 |
| tRNA-Asn-GTT-3-5 | 24.0922481 | -0.949762 | 0.41148327 | -2.3081425 | 0.02099121 |
| tRNA-Arg-TCT-4-1 | 12.319731 | 1.60396645 | 0.70413719 | 2.27791752 | 0.02273149 |
| tRNA-Asp-GTC-4-1 | 215.689033 | 0.84569868 | 0.3780186 | 2.23718798 | 0.02527406 |
| tRNA-Lys-CTT-6-1 | 1.4433289 | -3.2065808 | 1.50204986 | -2.1348032 | 0.03277709 |
| tRNA-Asn-GTT-3-2 | 59.2855903 | -0.9264352 | 0.43463266 | -2.1315362 | 0.033045 |
| tRNA-Gly-GCC-1-1 | 1698.42143 | 0.51538834 | 0.24456464 | 2.10737063 | 0.03508547 |
| tRNA-Gly-GCC-1-2 | 1698.42143 | 0.51538834 | 0.24456464 | 2.10737063 | 0.03508547 |
| tRNA-Gly-GCC-1-3 | 1698.42143 | 0.51538834 | 0.24456464 | 2.10737063 | 0.03508547 |
| tRNA-Ala-AGC-7-1 | 17.8601853 | 0.92417051 | 0.43969337 | 2.10185227 | 0.03556622 |
| tRNA-Ile-AAT-1-1 | 22.4318702 | -0.7082834 | 0.34451099 | -2.0559095 | 0.03979124 |
| tRNA-Ala-AGC-6-1 | 18.0796148 | 0.90482017 | 0.4407996 | 2.05267919 | 0.0401037 |
| tRNA-Ala-AGC-4-1 | 82.8674516 | -0.599721 | 0.29251207 | -2.0502435 | 0.04034068 |
| tRNA-Leu-CAG-2-2 | 682.200845 | -0.5806303 | 0.28457629 | -2.0403327 | 0.0413172 |
| tRNA-Ala-AGC-1-1 | 59.737632 | 0.57003278 | 0.28527532 | 1.99818468 | 0.04569664 |
| tRNA-Glu-CTC-1-8 | 12519.5887 | 0.51943472 | 0.26027488 | 1.99571596 | 0.04596485 |
| tRNA-Glu-CTC-1-7 | 12547.3672 | 0.51872004 | 0.26002311 | 1.99489975 | 0.04605381 |
| tRNA-Glu-CTC-1-5 | 12503.0878 | 0.51762572 | 0.26031075 | 1.98849149 | 0.04675736 |
| tRNA-Glu-CTC-1-6 | 12503.0878 | 0.51762572 | 0.26031075 | 1.98849149 | 0.04675736 |
| tRNA-Glu-CTC-1-4 | 12540.0237 | 0.51727999 | 0.26019542 | 1.98804416 | 0.04680681 |
| tRNA-Arg-CCT-2-2 | 19.5824216 | -0.8807939 | 0.44330714 | -1.9868706 | 0.04693674 |
| tRNA-Glu-CTC-1-9 | 12504.4328 | 0.5169538 | 0.26044692 | 1.98487201 | 0.04715871 |
| tRNA-His-GTG-1-1 | 169.594431 | 0.58738867 | 0.29610015 | 1.98374996 | 0.04728372 |
| tRNA-Leu-CAG-1-1 | 670.982084 | -0.5742836 | 0.2900912 | -1.9796657 | 0.04774111 |
| tRNA-Leu-CAG-1-2 | 670.982084 | -0.5742836 | 0.2900912 | -1.9796657 | 0.04774111 |
| tRNA-Leu-CAG-1-3 | 670.982084 | -0.5742836 | 0.2900912 | -1.9796657 | 0.04774111 |
| tRNA-Glu-CTC-1-1 | 12513.9485 | 0.51571008 | 0.26050605 | 1.97964722 | 0.04774318 |
| tRNA-Glu-CTC-1-2 | 12513.9485 | 0.51571008 | 0.26050605 | 1.97964722 | 0.04774318 |
| tRNA-Glu-CTC-1-3 | 12513.9485 | 0.51571008 | 0.26050605 | 1.97964722 | 0.04774318 |
| tRNA-Leu-CAG-1-4 | 673.662652 | -0.5763247 | 0.292892 | -1.9677037 | 0.04910214 |
| tRNA-Trp-CCA-3-1 | 38.9630914 | 0.72491386 | 0.36882119 | 1.96548863 | 0.04935771 |
| tRNA-Leu-CAG-1-5 | 671.968702 | -0.5706023 | 0.29047247 | -1.9643936 | 0.04948446 |
| tRNA-Leu-CAG-2-1 | 695.833168 | -0.5621221 | 0.29054667 | -1.9347048 | 0.05302653 |
| tRNA-Lys-CTT-1-1 | 19.2527981 | -1.1281413 | 0.59089819 | -1.9091974 | 0.05623663 |
| tRNA-Ala-CGC-4-1 | 1.76644092 | -2.1114194 | 1.10740002 | -1.9066456 | 0.05656649 |
| tRNA-Gln-TTG-2-1 | 92.281875 | 0.50704073 | 0.2675656 | 1.89501466 | 0.05809046 |
| tRNA-Cys-GCA-24-1 | 267.526793 | 0.40845052 | 0.22109386 | 1.84740775 | 0.06468807 |
| tRNA-Cys-GCA-9-1 | 267.201577 | 0.4078049 | 0.22083262 | 1.84666969 | 0.06479503 |
| tRNA-Cys-GCA-4-2 | 267.317117 | 0.40682414 | 0.22064867 | 1.84376431 | 0.0652175 |
| tRNA-Cys-GCA-4-4 | 267.317117 | 0.40682414 | 0.22064867 | 1.84376431 | 0.0652175 |

| | | | | | |
|---|---|---|---|---|---|
| tRNA-Cys-GCA-4-10 | 267.317117 | 0.40682414 | 0.22064867 | 1.84376431 | 0.0652175 |
| tRNA-Cys-GCA-4-11 | 267.317117 | 0.40682414 | 0.22064867 | 1.84376431 | 0.0652175 |
| tRNA-Cys-GCA-4-8 | 267.435139 | 0.4076555 | 0.221101 | 1.84375239 | 0.06521923 |
| tRNA-Cys-GCA-4-12 | 267.618534 | 0.40462202 | 0.21954615 | 1.84299299 | 0.06533003 |
| tRNA-Cys-GCA-4-22 | 267.388749 | 0.40685581 | 0.22105396 | 1.84052717 | 0.06569088 |
| tRNA-Cys-GCA-4-23 | 267.388749 | 0.40685581 | 0.22105396 | 1.84052717 | 0.06569088 |
| tRNA-Cys-GCA-4-25 | 267.388749 | 0.40685581 | 0.22105396 | 1.84052717 | 0.06569088 |
| tRNA-Cys-GCA-4-28 | 267.388749 | 0.40685581 | 0.22105396 | 1.84052717 | 0.06569088 |
| tRNA-Cys-GCA-14-1 | 267.406934 | 0.40612106 | 0.22067698 | 1.84034175 | 0.06571808 |
| tRNA-Cys-GCA-4-3 | 267.406934 | 0.40612106 | 0.22067698 | 1.84034175 | 0.06571808 |
| tRNA-Cys-GCA-4-5 | 267.406934 | 0.40612106 | 0.22067698 | 1.84034175 | 0.06571808 |
| tRNA-Cys-GCA-4-6 | 267.406934 | 0.40612106 | 0.22067698 | 1.84034175 | 0.06571808 |
| tRNA-Cys-GCA-4-7 | 267.406934 | 0.40612106 | 0.22067698 | 1.84034175 | 0.06571808 |
| tRNA-Cys-GCA-4-14 | 267.406934 | 0.40612106 | 0.22067698 | 1.84034175 | 0.06571808 |
| tRNA-Cys-GCA-12-1 | 267.348082 | 0.40612511 | 0.2207286 | 1.83992969 | 0.06577856 |
| tRNA-Cys-GCA-4-16 | 267.348082 | 0.40612511 | 0.2207286 | 1.83992969 | 0.06577856 |
| tRNA-Cys-GCA-6-1 | 267.348082 | 0.40612511 | 0.2207286 | 1.83992969 | 0.06577856 |
| tRNA-Cys-GCA-4-17 | 267.348082 | 0.40612511 | 0.2207286 | 1.83992969 | 0.06577856 |
| tRNA-Cys-GCA-4-18 | 267.348082 | 0.40612511 | 0.2207286 | 1.83992969 | 0.06577856 |
| tRNA-Cys-GCA-4-19 | 267.348082 | 0.40612511 | 0.2207286 | 1.83992969 | 0.06577856 |
| tRNA-Cys-GCA-4-20 | 267.348082 | 0.40612511 | 0.2207286 | 1.83992969 | 0.06577856 |
| tRNA-Cys-GCA-4-24 | 267.348082 | 0.40612511 | 0.2207286 | 1.83992969 | 0.06577856 |
| tRNA-Cys-GCA-10-1 | 267.348082 | 0.40612511 | 0.2207286 | 1.83992969 | 0.06577856 |
| tRNA-Cys-GCA-4-26 | 267.348082 | 0.40612511 | 0.2207286 | 1.83992969 | 0.06577856 |
| tRNA-Cys-GCA-4-27 | 267.348082 | 0.40612511 | 0.2207286 | 1.83992969 | 0.06577856 |
| tRNA-Cys-GCA-4-29 | 267.348082 | 0.40612511 | 0.2207286 | 1.83992969 | 0.06577856 |
| tRNA-Cys-GCA-4-1 | 267.466104 | 0.40695645 | 0.22118066 | 1.83992778 | 0.06577884 |
| tRNA-Cys-GCA-4-21 | 267.466104 | 0.40695645 | 0.22118066 | 1.83992778 | 0.06577884 |
| tRNA-Cys-GCA-4-13 | 269.242604 | 0.40010245 | 0.21779289 | 1.83707765 | 0.06619843 |
| tRNA-Cys-GCA-7-1 | 272.909989 | 0.38926784 | 0.21191327 | 1.83692059 | 0.06622162 |
| tRNA-Cys-GCA-4-15 | 267.419715 | 0.40615677 | 0.22113365 | 1.83670269 | 0.0662538 |
| tRNA-Ile-AAT-1-7 | 25.4197922 | -0.642616 | 0.3502591 | -1.8346874 | 0.06655202 |
| tRNA-Cys-GCA-4-9 | 269.103436 | 0.39772902 | 0.21786003 | 1.82561718 | 0.06790795 |
| tRNA-Cys-GCA-5-1 | 283.906768 | 0.38449399 | 0.2125673 | 1.80881063 | 0.07048043 |
| tRNA-Leu-CAG-2-3 | 715.525508 | -0.5365502 | 0.29725179 | -1.8050359 | 0.07106907 |
| tRNA-Arg-CCT-2-1 | 19.3057441 | -0.7844344 | 0.43990718 | -1.7831816 | 0.07455675 |
| tRNA-Ile-AAT-4-1 | 3.20242932 | -1.4769964 | 0.82960835 | -1.7803538 | 0.07501808 |
| tRNA-Ala-TGC-1-1 | 24.514306 | -0.7294921 | 0.41103965 | -1.7747487 | 0.07593938 |
| tRNA-Trp-CCA-5-1 | 500.535416 | -0.5503005 | 0.31027881 | -1.7735679 | 0.07613466 |
| tRNA-Gly-ACC-1-1 | 42.0915647 | -0.5336954 | 0.30150407 | -1.7701102 | 0.07670878 |
| tRNA-Lys-CTT-2-2 | 3471.13671 | 0.36234096 | 0.20539118 | 1.76415057 | 0.07770663 |
| tRNA-Lys-CTT-2-1 | 3466.48099 | 0.36194221 | 0.20538319 | 1.76227766 | 0.0780224 |
| tRNA-Ala-AGC-4-2 | 85.6153283 | -0.5299475 | 0.30522696 | -1.7362407 | 0.08252129 |
| tRNA-Asn-GTT-3-7 | 27.6734014 | -0.6195167 | 0.3611151 | -1.7155658 | 0.08624154 |
| tRNA-Asn-GTT-3-8 | 36.4757383 | -0.7099174 | 0.41887176 | -1.6948322 | 0.09010725 |
| tRNA-Ile-AAT-3-1 | 23.6073554 | -0.6196094 | 0.36747585 | -1.6861229 | 0.09177214 |

| | | | | | |
|---|---|---|---|---|---|
| tRNA-Tyr-GTA-3-2 | 9.28585231 | -0.9326309 | 0.57686471 | -1.6167238 | 0.10593792 |
| tRNA-Arg-TCT-1-1 | 8.71325274 | -1.0913017 | 0.68592632 | -1.5909897 | 0.1116119 |
| tRNA-Arg-CCT-1-1 | 19.0254253 | -0.7198846 | 0.45410157 | -1.5852943 | 0.11289948 |
| tRNA-Ala-AGC-5-2 | 66.9550485 | -0.4992574 | 0.31691742 | -1.5753549 | 0.11517455 |
| tRNA-Ala-AGC-5-3 | 66.9550485 | -0.4992574 | 0.31691742 | -1.5753549 | 0.11517455 |
| tRNA-Ala-AGC-10-1 | 66.8530965 | -0.4957074 | 0.31619001 | -1.5677515 | 0.11693914 |
| tRNA-Ala-AGC-12-1 | 67.9931028 | -0.4816408 | 0.31074815 | -1.5499394 | 0.12115607 |
| tRNA-Arg-TCG-3-1 | 23.4064501 | -0.7167665 | 0.47010208 | -1.524704 | 0.12733295 |
| tRNA-Arg-TCG-3-2 | 23.4064501 | -0.7167665 | 0.47010208 | -1.524704 | 0.12733295 |
| tRNA-Ala-AGC-5-1 | 68.668781 | -0.4644458 | 0.31555737 | -1.4718267 | 0.14106767 |
| tRNA-Tyr-GTA-2-1 | 35.6118649 | 0.60742509 | 0.42479747 | 1.4299169 | 0.15274087 |
| tRNA-Glu-CTC-5-1 | 4.82959767 | 1.53203577 | 1.08903405 | 1.40678408 | 0.15949142 |
| tRNA-Lys-CTT-10-1 | 2.16766622 | -1.3988658 | 0.99885117 | -1.4004747 | 0.16137122 |
| tRNA-Phe-GAA-2-1 | 0.91742194 | -2.1077573 | 1.51956363 | -1.3870807 | 0.16541717 |
| tRNA-Gly-GCC-5-1 | 466.407171 | 0.47555303 | 0.35360795 | 1.34485954 | 0.1786706 |
| tRNA-Cys-GCA-13-1 | 21.9633135 | 0.45408824 | 0.34424971 | 1.31906643 | 0.18714691 |
| tRNA-Leu-TAA-3-1 | 0.76610446 | -2.240794 | 1.71161727 | -1.3091677 | 0.19047756 |
| tRNA-Glu-CTC-3-1 | 18.9331099 | 0.56058371 | 0.4291543 | 1.30625211 | 0.19146684 |
| tRNA-Trp-CCA-2-1 | 1815.9669 | -0.3710379 | 0.28552608 | -1.2994887 | 0.19377627 |
| tRNA-Trp-CCA-4-1 | 26.4858953 | 0.46757417 | 0.36469645 | 1.28209135 | 0.1998106 |
| tRNA-Trp-CCA-4-2 | 26.4858953 | 0.46757417 | 0.36469645 | 1.28209135 | 0.1998106 |
| tRNA-Cys-GCA-19-1 | 6.12119462 | 0.73301174 | 0.58550949 | 1.2519212 | 0.21059858 |
| tRNA-Lys-CTT-16-1 | 1.96092478 | -1.2336037 | 1.02052964 | -1.2087877 | 0.22674442 |
| tRNA-Ala-AGC-2-1 | 83.4787422 | 0.35058201 | 0.2904589 | 1.20699351 | 0.22743464 |
| tRNA-Lys-TTT-2-1 | 27.0666458 | -0.6487531 | 0.55004986 | -1.1794441 | 0.23822138 |
| tRNA-Lys-TTT-2-2 | 26.9172173 | -0.6489154 | 0.55134705 | -1.1769636 | 0.23921004 |
| tRNA-Ala-AGC-2-2 | 85.7093237 | 0.32964729 | 0.28349932 | 1.16277984 | 0.24491884 |
| tRNA-Asn-GTT-3-1 | 29.9718303 | -0.3918253 | 0.35073219 | -1.1171638 | 0.26392431 |
| tRNA-Arg-TCG-2-1 | 63.146195 | -0.4101635 | 0.3689914 | -1.1115802 | 0.2663187 |
| tRNA-Leu-CAG-4-1 | 27.9424695 | 0.49710882 | 0.45094569 | 1.1023696 | 0.27030102 |
| tRNA-Arg-ACG-1-3 | 19.0353373 | -0.4879683 | 0.44383158 | -1.0994447 | 0.27157413 |
| tRNA-Lys-CTT-18-1 | 7.24927805 | 0.80497318 | 0.74834457 | 1.07567184 | 0.28207405 |
| tRNA-Thr-TGT-3-1 | 51.8670138 | -0.3257339 | 0.30376058 | -1.0723377 | 0.28356837 |
| tRNA-Pro-AGG-3-1 | 7.72797027 | -0.6219248 | 0.59920068 | -1.0379241 | 0.2993054 |
| tRNA-Glu-CTC-2-1 | 367.676935 | 0.80741355 | 0.80830685 | 0.99889486 | 0.31784563 |
| tRNA-Cys-GCA-18-1 | 1.4036703 | 1.58974072 | 1.60787393 | 0.98872224 | 0.32279906 |
| tRNA-Tyr-GTA-3-1 | 9.85513622 | -0.5399466 | 0.54866246 | -0.9841144 | 0.32505927 |
| tRNA-Gln-CTG-7-1 | 2.46529023 | -0.9388857 | 0.95607557 | -0.9820204 | 0.32608979 |
| tRNA-Ala-CGC-5-1 | 9.54769646 | 0.88247214 | 0.92401955 | 0.95503622 | 0.33955938 |
| tRNA-Cys-GCA-11-1 | 1.7644337 | -0.905456 | 0.9842964 | -0.9199018 | 0.3576241 |
| tRNA-Ile-AAT-1-8 | 31.1021011 | -0.2919466 | 0.32125537 | -0.9087681 | 0.36347252 |
| tRNA-Arg-TCT-3-1 | 2.75609899 | 0.91458888 | 1.04604072 | 0.87433392 | 0.38193643 |
| tRNA-Thr-CGT-3-1 | 1.32481075 | 1.82742317 | 2.10835634 | 0.86675252 | 0.38607762 |
| tRNA-Ser-GCT-5-1 | 13.6209205 | -0.4615293 | 0.53534232 | -0.86212 | 0.38862148 |
| tRNA-Thr-AGT-2-1 | 50.0235375 | 0.38042775 | 0.47566074 | 0.79978799 | 0.42383365 |
| tRNA-Lys-CTT-13-1 | 7.44524461 | 0.58799814 | 0.75197778 | 0.78193553 | 0.43425247 |

| | | | | | |
|---|---|---|---|---|---|
| tRNA-Tyr-GTA-1-1 | 3.24003894 | -0.6602105 | 0.8561407 | -0.7711472 | 0.44061969 |
| tRNA-Cys-GCA-15-1 | 5.35612196 | 0.44378698 | 0.57657789 | 0.76969129 | 0.44148304 |
| tRNA-Gln-CTG-2-2 | 314.041103 | -0.2251393 | 0.30352772 | -0.7417422 | 0.45824353 |
| tRNA-Lys-TTT-1-1 | 614.218266 | -0.2962618 | 0.40336813 | -0.73447 | 0.46266231 |
| tRNA-Lys-TTT-1-5 | 604.229473 | -0.2957778 | 0.40370331 | -0.7326614 | 0.46376496 |
| tRNA-Ile-AAT-2-1 | 7.08639499 | -0.540188 | 0.74218162 | -0.727838 | 0.46671279 |
| tRNA-Gln-CTG-2-4 | 312.494498 | -0.220914 | 0.30401314 | -0.7266594 | 0.46743466 |
| tRNA-Gln-TTG-6-1 | 5.84371917 | -0.5767575 | 0.79513191 | -0.7253608 | 0.46823071 |
| tRNA-Asp-GTC-1-10 | 404.065018 | 0.25852589 | 0.36005184 | 0.71802409 | 0.47274243 |
| tRNA-Asp-GTC-1-5 | 404.36788 | 0.25689483 | 0.3614387 | 0.7107563 | 0.47723526 |
| tRNA-Asp-GTC-1-6 | 403.617022 | 0.2546531 | 0.36082997 | 0.70574265 | 0.48034819 |
| tRNA-Asp-GTC-1-1 | 404.200032 | 0.25447854 | 0.36110444 | 0.70472281 | 0.48098275 |
| tRNA-Asp-GTC-1-13 | 403.162405 | 0.25371405 | 0.36042074 | 0.70393856 | 0.48147104 |
| tRNA-Asp-GTC-1-3 | 402.920073 | 0.25371554 | 0.36052561 | 0.70373792 | 0.481596 |
| tRNA-Asp-GTC-1-4 | 402.920073 | 0.25371554 | 0.36052561 | 0.70373792 | 0.481596 |
| tRNA-Asp-GTC-1-9 | 402.920073 | 0.25371554 | 0.36052561 | 0.70373792 | 0.481596 |
| tRNA-Asp-GTC-1-7 | 403.159636 | 0.25371634 | 0.36058166 | 0.70363073 | 0.48166277 |
| tRNA-Asp-GTC-1-2 | 403.039162 | 0.25371651 | 0.36059412 | 0.70360692 | 0.4816776 |
| tRNA-Asp-GTC-1-12 | 403.134072 | 0.25266899 | 0.36052769 | 0.70083104 | 0.48340847 |
| tRNA-Thr-AGT-4-1 | 13.5343955 | 0.39299035 | 0.56148237 | 0.69991575 | 0.48397992 |
| tRNA-Tyr-GTA-1-2 | 3.21237119 | -0.5997956 | 0.85845674 | -0.6986905 | 0.48474546 |
| tRNA-Asp-GTC-1-8 | 403.237111 | 0.25191481 | 0.36057314 | 0.69865106 | 0.48477012 |
| tRNA-Tyr-GTA-1-5 | 3.28708544 | -0.5999316 | 0.85994701 | -0.6976379 | 0.48540369 |
| tRNA-Asp-GTC-1-11 | 404.495896 | 0.25047665 | 0.35954027 | 0.69665815 | 0.48601676 |
| tRNA-Gln-CTG-2-1 | 313.078969 | -0.2094547 | 0.30388465 | -0.6892573 | 0.49066139 |
| tRNA-Arg-ACG-1-1 | 13.1807709 | -0.2958484 | 0.43043158 | -0.6873296 | 0.49187504 |
| tRNA-Gln-CTG-2-3 | 313.626515 | -0.2083961 | 0.3055426 | -0.6820526 | 0.4952057 |
| tRNA-Lys-CTT-7-1 | 0.28965768 | 1.87187882 | 2.76036932 | 0.67812622 | 0.49769167 |
| tRNA-Arg-ACG-1-2 | 12.9566281 | -0.299165 | 0.44731176 | -0.6688064 | 0.503619 |
| tRNA-Lys-TTT-1-4 | 600.173947 | -0.2723541 | 0.41098867 | -0.6626803 | 0.50753536 |
| tRNA-Thr-TGT-1-1 | 4.06582087 | -0.5198714 | 0.78626959 | -0.6611872 | 0.50849225 |
| tRNA-Lys-TTT-1-3 | 636.401634 | -0.259314 | 0.39480275 | -0.6568192 | 0.51129718 |
| tRNA-Lys-CTT-15-1 | 0.79335361 | -1.0688857 | 1.63314769 | -0.6544942 | 0.51279346 |
| tRNA-Thr-TGT-2-1 | 29.0184782 | -0.1802535 | 0.2918619 | -0.6175985 | 0.53684005 |
| tRNA-Lys-TTT-1-2 | 574.090964 | -0.2605147 | 0.43116941 | -0.6042049 | 0.54570742 |
| tRNA-Leu-TAA-4-1 | 0.84691153 | 1.1890119 | 2.00613732 | 0.5926872 | 0.55339051 |
| tRNA-Phe-GAA-1-1 | 22.3229563 | -0.2455383 | 0.42296622 | -0.5805151 | 0.56156732 |
| tRNA-Trp-CCA-1-1 | 25.9166198 | 0.21574823 | 0.3811498 | 0.56604578 | 0.57136266 |
| tRNA-Glu-CTC-4-1 | 6.42515408 | 0.49105443 | 0.92775515 | 0.52929314 | 0.59660212 |
| tRNA-Cys-GCA-8-1 | 44.8066896 | -0.2165254 | 0.41466891 | -0.5221646 | 0.60155573 |
| tRNA-Lys-TTT-1-6 | 603.189306 | -0.2106882 | 0.40956741 | -0.5144164 | 0.60696087 |
| tRNA-Cys-GCA-25-1 | 0.77105215 | -0.8782054 | 1.77170933 | -0.4956825 | 0.6201184 |
| tRNA-Tyr-GTA-1-4 | 3.89448409 | -0.3859211 | 0.80387252 | -0.4800775 | 0.63117232 |
| tRNA-Gln-TTG-3-2 | 184.203703 | 0.11536004 | 0.24159991 | 0.47748378 | 0.63301767 |
| tRNA-Ile-TAT-2-3 | 2.17857793 | -0.5127638 | 1.07775706 | -0.4757693 | 0.63423872 |
| tRNA-Gln-TTG-3-1 | 184.544006 | 0.11363141 | 0.24099477 | 0.47150985 | 0.63727669 |

| | | | | | |
|---|---|---|---|---|---|
| tRNA-Sup-TTA-1-1 | 0.10509283 | 1.93415058 | 4.18419562 | 0.46225147 | 0.643901 |
| tRNA-Cys-GCA-17-1 | 5.85840989 | 0.24216108 | 0.56100334 | 0.43165711 | 0.66599065 |
| tRNA-Pro-GGG-1-1 | 0.09277906 | 1.80845071 | 4.19078188 | 0.43153062 | 0.66608259 |
| tRNA-Gly-TCC-2-1 | 243.066674 | -0.2944649 | 0.68363552 | -0.4307338 | 0.66666192 |
| tRNA-Arg-TCG-4-1 | 24.7520366 | 0.19698406 | 0.46182466 | 0.42653431 | 0.66971856 |
| tRNA-Lys-TTT-3-1 | 0.26740619 | 1.67688443 | 4.01013349 | 0.41816175 | 0.67582886 |
| tRNA-iMet-CAT-2-1 | 53.4682505 | -0.1011509 | 0.25447193 | -0.3974932 | 0.69100377 |
| tRNA-Gln-CTG-3-1 | 309.696451 | -0.1142887 | 0.28832036 | -0.3963947 | 0.69181385 |
| tRNA-Gln-CTG-3-2 | 309.662189 | -0.1136507 | 0.28828439 | -0.394231 | 0.69341047 |
| tRNA-Gln-CTG-3-3 | 309.634521 | -0.1131383 | 0.28842942 | -0.3922564 | 0.69486875 |
| tRNA-Gln-TTG-4-1 | 0.42758692 | -0.9859493 | 2.51833464 | -0.3915084 | 0.69542145 |
| tRNA-Tyr-GTA-1-3 | 3.5270186 | -0.3303537 | 0.84676216 | -0.3901376 | 0.69643482 |
| tRNA-Gln-TTG-1-1 | 168.678218 | 0.08878835 | 0.22803023 | 0.38937096 | 0.69700175 |
| tRNA-Gln-TTG-5-1 | 41.1598104 | -0.1545705 | 0.40508377 | -0.3815766 | 0.70277548 |
| tRNA-Arg-TCT-5-1 | 0.24748819 | 1.26608761 | 3.5592368 | 0.35571885 | 0.72205113 |
| tRNA-Arg-ACG-3-1 | 39.6139245 | -0.1542439 | 0.43510591 | -0.3544974 | 0.72296616 |
| tRNA-Arg-CCG-3-1 | 685.49171 | 0.13465083 | 0.39791467 | 0.33839122 | 0.73506839 |
| tRNA-Ile-TAT-1-1 | 7.23774635 | 0.26671601 | 0.80323193 | 0.33205355 | 0.73984882 |
| tRNA-Ile-GAT-1-1 | 0.15286675 | -1.3704124 | 4.16628127 | -0.3289294 | 0.74220905 |
| tRNA-Arg-ACG-3-2 | 30.7468997 | -0.1153069 | 0.35779674 | -0.3222693 | 0.74724866 |
| tRNA-Val-TAC-1-1 | 296.977326 | 0.08528372 | 0.27472592 | 0.31043202 | 0.75623244 |
| tRNA-Tyr-GTA-4-1 | 3.5628694 | -0.3288738 | 1.09294783 | -0.3009053 | 0.76348669 |
| tRNA-Pro-TGG-3-1 | 0.08630054 | -1.2433174 | 4.17138082 | -0.2980589 | 0.76565818 |
| tRNA-Gln-CTG-6-1 | 107.84331 | 0.08015064 | 0.27217922 | 0.29447744 | 0.7683931 |
| tRNA-Pro-TGG-5-1 | 0.08300326 | -1.2031731 | 4.17308171 | -0.2883177 | 0.77310359 |
| tRNA-Val-TAC-1-2 | 294.870895 | 0.07685943 | 0.27416415 | 0.28034092 | 0.77921596 |
| tRNA-Val-TAC-1-3 | 294.870895 | 0.07685943 | 0.27416415 | 0.28034092 | 0.77921596 |
| tRNA-Lys-CTT-4-1 | 0.11278317 | 1.17249337 | 4.20595667 | 0.27876972 | 0.78042155 |
| tRNA-Ile-TAT-3-1 | 0.11139863 | 1.17249337 | 4.20595667 | 0.27876972 | 0.78042155 |
| tRNA-Thr-AGT-1-2 | 14.1257781 | 0.12180445 | 0.46896766 | 0.25972889 | 0.7950729 |
| tRNA-Thr-AGT-1-3 | 14.1257781 | 0.12180445 | 0.46896766 | 0.25972889 | 0.7950729 |
| tRNA-Thr-AGT-1-1 | 14.1257781 | 0.12180445 | 0.46896766 | 0.25972889 | 0.7950729 |
| tRNA-Pro-AGG-2-1 | 1.9991773 | 0.24573143 | 0.95433592 | 0.25748945 | 0.79680096 |
| tRNA-Ser-GGA-1-1 | 0.54460155 | 0.63872101 | 2.63202331 | 0.24267301 | 0.80825872 |
| tRNA-Cys-GCA-23-1 | 1.97794458 | 0.22488116 | 0.9281479 | 0.24229022 | 0.80855529 |
| tRNA-Arg-TCT-2-1 | 20.4532262 | -0.1183666 | 0.48937754 | -0.2418718 | 0.80887949 |
| tRNA-Cys-GCA-26-1 | 2.44593532 | 0.20132183 | 0.85597817 | 0.23519506 | 0.81405734 |
| tRNA-Tyr-GTA-6-1 | 4.68458016 | 0.22980159 | 1.05230209 | 0.21837986 | 0.82713315 |
| tRNA-Ala-AGC-14-1 | 0.05863278 | -0.8389589 | 4.18869939 | -0.200291 | 0.84125298 |
| tRNA-Trp-CCA-6-1 | 19.3788719 | 0.07603156 | 0.38751403 | 0.19620338 | 0.84445098 |
| tRNA-Asp-GTC-3-1 | 1.85075566 | 0.18697587 | 1.23104202 | 0.15188423 | 0.87927824 |
| tRNA-Thr-AGT-3-1 | 14.0743672 | 0.06821715 | 0.49516784 | 0.13776572 | 0.89042558 |
| tRNA-Met-CAT-6-1 | 1.81608629 | -0.1388754 | 1.05555335 | -0.1315664 | 0.89532723 |
| tRNA-Arg-ACG-2-1 | 9.19852009 | 0.06157535 | 0.50298516 | 0.12241981 | 0.90256655 |
| tRNA-Lys-CTT-14-1 | 0.22859632 | 0.45119441 | 3.72718624 | 0.12105497 | 0.9036475 |
| tRNA-Asn-GTT-4-1 | 0.21611973 | 0.45120328 | 3.77234667 | 0.11960812 | 0.90479359 |

| | | | | | |
|---|---|---|---|---|---|
| tRNA-Tyr-GTA-5-1 | 0.29380037 | 0.4511042 | 4.20595667 | 0.10725365 | 0.91458775 |
| tRNA-Leu-TAA-5-1 | 2.15092461 | -0.0844288 | 0.91445749 | -0.0923266 | 0.92643855 |
| tRNA-Leu-TAA-5-2 | 2.15092461 | -0.0844288 | 0.91445749 | -0.0923266 | 0.92643855 |
| tRNA-Leu-TAA-5-3 | 2.15092461 | -0.0844288 | 0.91445749 | -0.0923266 | 0.92643855 |
| tRNA-Cys-GCA-20-1 | 4.07688079 | 0.04635321 | 0.70397696 | 0.06584478 | 0.94750141 |
| tRNA-Val-CAC-5-1 | 0.23619037 | -0.2701779 | 4.12226822 | -0.0655411 | 0.94774321 |
| tRNA-iMet-CAT-4-1 | 0.11553966 | -0.2701379 | 4.20595667 | -0.0642274 | 0.94878912 |
| tRNA-Thr-CGT-4-1 | 3.57577703 | 0.02183562 | 0.87671872 | 0.02490607 | 0.98012989 |
| tRNA-Ile-TAT-2-1 | 1.06638822 | 0.01786908 | 1.45802579 | 0.01225567 | 0.99022163 |
| tRNA-Ile-TAT-2-2 | 1.06638822 | 0.01786908 | 1.45802579 | 0.01225567 | 0.99022163 |
| tRNA-Thr-TGT-3-2 | 61.3470488 | 0.00206786 | 0.29088089 | 0.00710895 | 0.99432792 |

| padj |
|---|
| 4.06E-27 |
| 4.06E-27 |
| 4.06E-27 |
| 4.06E-27 |
| 4.06E-27 |
| 4.06E-27 |
| 4.06E-27 |
| 4.06E-27 |
| 1.50E-16 |
| 2.90E-12 |
| 6.70E-11 |
| 6.70E-11 |
| 6.70E-11 |
| 6.70E-11 |
| 6.70E-11 |
| 6.70E-11 |
| 6.70E-11 |
| 2.76E-10 |
| 2.76E-10 |
| 3.25E-10 |
| 7.22E-10 |
| 7.22E-10 |
| 7.22E-10 |
| 1.84E-09 |
| 4.26E-08 |
| 4.84E-08 |
| 1.82E-07 |
| 1.82E-07 |
| 2.09E-07 |
| 2.17E-07 |
| 3.76E-07 |
| 3.76E-07 |
| 3.95E-07 |
| 4.75E-07 |
| 5.25E-07 |
| 5.25E-07 |
| 7.11E-07 |
| 7.11E-07 |
| 7.11E-07 |
| 7.11E-07 |
| 8.26E-07 |
| 8.62E-07 |
| 1.13E-06 |
| 1.13E-06 |

| |
|---|
| 1.13E-06 |
| 1.13E-06 |
| 1.30E-06 |
| 3.83E-06 |
| 4.12E-06 |
| 7.03E-06 |
| 8.02E-06 |
| 8.84E-06 |
| 9.39E-06 |
| 1.32E-05 |
| 1.32E-05 |
| 1.32E-05 |
| 1.39E-05 |
| 1.39E-05 |
| 1.39E-05 |
| 1.60E-05 |
| 1.63E-05 |
| 2.40E-05 |
| 2.40E-05 |
| 2.45E-05 |
| 2.52E-05 |
| 2.52E-05 |
| 2.52E-05 |
| 2.57E-05 |
| 2.57E-05 |
| 2.57E-05 |
| 2.57E-05 |
| 2.57E-05 |
| 3.13E-05 |
| 3.46E-05 |
| 4.64E-05 |
| 5.03E-05 |
| 5.14E-05 |
| 5.44E-05 |
| 5.74E-05 |
| 0.00015232 |
| 0.00015823 |
| 0.00015871 |
| 0.00016981 |
| 0.00017505 |
| 0.00018379 |
| 0.00018379 |
| 0.00018379 |
| 0.00019963 |
| 0.00021802 |
| 0.00030628 |

| |
|---|
| 0.00035011 |
| 0.00035684 |
| 0.00035684 |
| 0.00035684 |
| 0.00035684 |
| 0.00035684 |
| 0.00035767 |
| 0.00036099 |
| 0.0006132 |
| 0.00061404 |
| 0.00061404 |
| 0.0006579 |
| 0.00065973 |
| 0.00065973 |
| 0.00065973 |
| 0.00065973 |
| 0.00066625 |
| 0.00070435 |
| 0.00076806 |
| 0.00080144 |
| 0.00080144 |
| 0.00080144 |
| 0.00088992 |
| 0.00088992 |
| 0.00088992 |
| 0.00092121 |
| 0.00093238 |
| 0.00093238 |
| 0.00096418 |
| 0.00125715 |
| 0.001291 |
| 0.00134655 |
| 0.00162708 |
| 0.00169888 |
| 0.00236118 |
| 0.00338902 |
| 0.00420302 |
| 0.00468694 |
| 0.00487858 |
| 0.00493337 |
| 0.00572098 |
| 0.00572098 |
| 0.00572098 |
| 0.00572098 |
| 0.00572098 |
| 0.00572098 |

| |
|---|
| 0.00572098 |
| 0.00572098 |
| 0.00572098 |
| 0.00572098 |
| 0.00572098 |
| 0.00572098 |
| 0.00572098 |
| 0.00615001 |
| 0.00647631 |
| 0.00874865 |
| 0.00933729 |
| 0.0108557 |
| 0.01107101 |
| 0.01153095 |
| 0.01206332 |
| 0.01206332 |
| 0.01210316 |
| 0.01214226 |
| 0.01273794 |
| 0.01449928 |
| 0.01493422 |
| 0.01602014 |
| 0.01602654 |
| 0.01867421 |
| 0.01939651 |
| 0.0197637 |
| 0.02055051 |
| 0.02218039 |
| 0.02518186 |
| 0.02518186 |
| 0.02555958 |
| 0.02671375 |
| 0.02808402 |
| 0.02826718 |
| 0.03230863 |
| 0.03230863 |
| 0.03230863 |
| 0.03230863 |
| 0.03230863 |
| 0.03230863 |
| 0.03230863 |
| 0.03237449 |
| 0.03237449 |
| 0.03255225 |
| 0.03366356 |
| 0.03366356 |

| |
|---|
| 0.03664772 |
| 0.03691475 |
| 0.0394224 |
| 0.04035521 |
| 0.04147555 |
| 0.0433433 |
| 0.04673307 |
| 0.04673307 |
| 0.05034252 |
| 0.05568191 |
| 0.07183788 |
| 0.07205172 |
| 0.0753358 |
| 0.0753358 |
| 0.0753358 |
| 0.07598238 |
| 0.08458137 |
| 0.08481933 |
| 0.08489605 |
| 0.08652068 |
| 0.0930662 |
| 0.0930662 |
| 0.0930662 |
| 0.0930662 |
| 0.0930662 |
| 0.0930662 |
| 0.0930662 |
| 0.0930662 |
| 0.0930662 |
| 0.0930662 |
| 0.0930662 |
| 0.0930662 |
| 0.0930662 |
| 0.0930662 |
| 0.0930662 |
| 0.09514512 |
| 0.09514512 |
| 0.09514512 |
| 0.10149422 |
| 0.10715357 |
| 0.10729877 |
| 0.10820601 |
| 0.10820601 |
| 0.10820601 |
| 0.10820601 |
| 0.10820601 |

| |
|---|
| 0.10820601 |
| 0.10820601 |
| 0.10820601 |
| 0.10820601 |
| 0.10820601 |
| 0.10820601 |
| 0.10820601 |
| 0.10820601 |
| 0.10820601 |
| 0.10820601 |
| 0.10820601 |
| 0.10820601 |
| 0.10820601 |
| 0.10820601 |
| 0.10820601 |
| 0.10820601 |
| 0.10820601 |
| 0.10820601 |
| 0.10820601 |
| 0.10820601 |
| 0.10820601 |
| 0.10820601 |
| 0.10820601 |
| 0.10820601 |
| 0.10820601 |
| 0.10820601 |
| 0.10820601 |
| 0.10820601 |
| 0.10820601 |
| 0.10820601 |
| 0.10820601 |
| 0.10827501 |
| 0.11005771 |
| 0.11379092 |
| 0.114305 |
| 0.11946025 |
| 0.11974584 |
| 0.12061783 |
| 0.12061783 |
| 0.12107393 |
| 0.12219296 |
| 0.12223509 |
| 0.12880629 |
| 0.13411828 |
| 0.13961674 |
| 0.14167742 |

| |
|---|
| 0.16295179 |
| 0.17105736 |
| 0.17240606 |
| 0.17461948 |
| 0.17461948 |
| 0.17666163 |
| 0.18238084 |
| 0.19032451 |
| 0.19032451 |
| 0.21011135 |
| 0.22669961 |
| 0.23589116 |
| 0.23783981 |
| 0.24295647 |
| 0.26151441 |
| 0.27297635 |
| 0.27687975 |
| 0.27736463 |
| 0.27975209 |
| 0.28650808 |
| 0.28650808 |
| 0.30095676 |
| 0.32283508 |
| 0.32283508 |
| 0.33701554 |
| 0.33728616 |
| 0.34418827 |
| 0.36966882 |
| 0.37179145 |
| 0.37610965 |
| 0.37664215 |
| 0.38992589 |
| 0.39071473 |
| 0.41105905 |
| 0.43510906 |
| 0.44046452 |
| 0.44210251 |
| 0.44210251 |
| 0.45889335 |
| 0.4817675 |
| 0.48809167 |
| 0.51126301 |
| 0.51517613 |
| 0.5169399 |
| 0.56201139 |
| 0.57402748 |

| |
|---|
| 0.57996063 |
| 0.57996063 |
| 0.59763107 |
| 0.59763107 |
| 0.59763107 |
| 0.59763107 |
| 0.59763107 |
| 0.59763107 |
| 0.59763107 |
| 0.59763107 |
| 0.59763107 |
| 0.59763107 |
| 0.59763107 |
| 0.59763107 |
| 0.59763107 |
| 0.59763107 |
| 0.59763107 |
| 0.59763107 |
| 0.59763107 |
| 0.59763107 |
| 0.59763107 |
| 0.59763107 |
| 0.59763107 |
| 0.59763107 |
| 0.60133856 |
| 0.60133856 |
| 0.60366574 |
| 0.6049528 |
| 0.61040354 |
| 0.61279836 |
| 0.61279836 |
| 0.61442815 |
| 0.61448055 |
| 0.64147837 |
| 0.65023729 |
| 0.65753985 |
| 0.66538649 |
| 0.67510169 |
| 0.70296015 |
| 0.70682798 |
| 0.71120345 |
| 0.72461349 |
| 0.73502186 |
| 0.73502186 |
| 0.73502186 |
| 0.73652469 |

| |
|---|
| 0.74215292 |
| 0.76215673 |
| 0.76215673 |
| 0.76215673 |
| 0.76358746 |
| 0.76848282 |
| 0.77792016 |
| 0.77792016 |
| 0.77792016 |
| 0.77792016 |
| 0.77792016 |
| 0.77792016 |
| 0.77792016 |
| 0.78230007 |
| 0.80056201 |
| 0.80056201 |
| 0.81183794 |
| 0.81498972 |
| 0.81546605 |
| 0.81887613 |
| 0.82657965 |
| 0.83235792 |
| 0.83257946 |
| 0.83341098 |
| 0.83574257 |
| 0.83574257 |
| 0.83574257 |
| 0.83574257 |
| 0.83574257 |
| 0.84472883 |
| 0.84472883 |
| 0.84472883 |
| 0.84472883 |
| 0.85113439 |
| 0.85113439 |
| 0.85113439 |
| 0.85445721 |
| 0.86603298 |
| 0.878642 |
| 0.87980977 |
| 0.91384447 |
| 0.92316181 |
| 0.92597413 |
| 0.9289507 |
| 0.9289507 |
| 0.9289507 |

0.93673274
0.94202766
0.94202766
0.94202766
0.95784677
0.95784677
0.95784677
0.98713082
0.99256813
0.99256813
0.99432792

| HS692_tRNA_pvalues_INTERACTION_ | | | | HS692_tRNA_pvalues_INT | |
|---|---|---|---|---|---|
| | baseMean | log2FoldChar | lfcSE | stat | pvalue |
| tRNA-His-GTG-2-7 | 28999.5299 | 1.90598032 | 0.3246549 | 5.87078863 | 4.34E-09 |
| tRNA-His-GTG-2-4 | 28998.4942 | 1.90641319 | 0.32474763 | 5.87044531 | 4.35E-09 |
| tRNA-His-GTG-2-5 | 28998.4942 | 1.90641319 | 0.32474763 | 5.87044531 | 4.35E-09 |
| tRNA-His-GTG-2-6 | 28998.4942 | 1.90641319 | 0.32474763 | 5.87044531 | 4.35E-09 |
| tRNA-His-GTG-2-1 | 29001.8773 | 1.90506895 | 0.32474776 | 5.8663035 | 4.46E-09 |
| tRNA-His-GTG-2-3 | 29001.8773 | 1.90506895 | 0.32474776 | 5.8663035 | 4.46E-09 |
| tRNA-His-GTG-2-2 | 29001.8918 | 1.90507028 | 0.32474836 | 5.86629686 | 4.46E-09 |
| tRNA-His-GTG-2-8 | 29001.817 | 1.90506341 | 0.32474841 | 5.86627475 | 4.46E-09 |
| tRNA-His-GTG-3-1 | 196.814482 | 1.63361493 | 0.37963662 | 4.30310157 | 1.68E-05 |
| tRNA-Pro-AGG-1-6 | 498.174598 | 1.24106994 | 0.353373 | 3.51206781 | 0.00044463 |
| tRNA-Pro-AGG-1-4 | 479.374246 | 1.256949 | 0.35908912 | 3.50038175 | 0.00046459 |
| tRNA-Pro-AGG-1-3 | 474.29805 | 1.24721022 | 0.35771443 | 3.48660862 | 0.00048919 |
| tRNA-Pro-AGG-1-2 | 472.481965 | 1.23715102 | 0.35634228 | 3.47180531 | 0.00051697 |
| tRNA-Pro-AGG-1-5 | 474.180973 | 1.2422944 | 0.3593391 | 3.45716456 | 0.00054589 |
| tRNA-Gly-GCC-6-1 | 530.397599 | 1.70430919 | 0.51879206 | 3.28514892 | 0.00101928 |
| tRNA-Pro-TGG-2-2 | 501.388162 | 1.11738386 | 0.34435577 | 3.24485301 | 0.00117511 |
| tRNA-Pro-TGG-2-3 | 499.10422 | 1.12135582 | 0.34593606 | 3.24151182 | 0.00118898 |
| tRNA-Pro-TGG-2-1 | 502.629255 | 1.12298301 | 0.34651668 | 3.24077623 | 0.00119205 |
| tRNA-Pro-TGG-4-1 | 196.596347 | 1.76520389 | 0.54595743 | 3.23322624 | 0.00122401 |
| tRNA-Pro-TGG-2-4 | 500.779936 | 1.11038375 | 0.3440964 | 3.22695542 | 0.00125115 |
| tRNA-Thr-TGT-1-1 | 4.06582087 | 4.96737464 | 1.58741652 | 3.12921944 | 0.00175271 |
| tRNA-Pro-AGG-1-1 | 498.548964 | 1.0895805 | 0.35481653 | 3.07082789 | 0.00213466 |
| tRNA-Glu-CTC-4-1 | 6.42515408 | 5.22163049 | 1.70912537 | 3.05514773 | 0.0022495 |
| tRNA-Pro-CGG-1-1 | 571.437924 | 1.08239559 | 0.35470372 | 3.05154843 | 0.00227664 |
| tRNA-Pro-CGG-1-3 | 563.110855 | 1.0803078 | 0.35481507 | 3.04470663 | 0.00232907 |
| tRNA-Pro-CGG-1-2 | 574.314626 | 1.06823468 | 0.35598459 | 3.00078912 | 0.00269281 |
| tRNA-Pro-TGG-1-1 | 358.920056 | 0.97021261 | 0.33934819 | 2.85904755 | 0.00424915 |
| tRNA-Trp-CCA-2-1 | 1815.9669 | -1.1817616 | 0.42075461 | -2.8086718 | 0.00497463 |
| tRNA-Thr-CGT-4-1 | 3.57577703 | 4.32298984 | 1.67494586 | 2.58097289 | 0.00985223 |
| tRNA-Lys-CTT-3-7 | 3829.01173 | 0.91853693 | 0.35609879 | 2.57944409 | 0.00989595 |
| tRNA-Lys-CTT-3-5 | 3828.89617 | 0.91776069 | 0.35615267 | 2.57687439 | 0.00996982 |
| tRNA-Lys-CTT-3-6 | 3828.89617 | 0.91776069 | 0.35615267 | 2.57687439 | 0.00996982 |
| tRNA-Lys-CTT-3-4 | 3854.03886 | 0.89811862 | 0.35604643 | 2.52247612 | 0.01165319 |
| tRNA-Lys-CTT-3-1 | 3852.60864 | 0.89847206 | 0.35625307 | 2.52200509 | 0.0116688 |
| tRNA-Lys-CTT-3-3 | 3858.00155 | 0.89838981 | 0.3564877 | 2.52011446 | 0.01173167 |
| tRNA-Lys-CTT-3-2 | 3876.20711 | 0.88502137 | 0.35149292 | 2.51789246 | 0.01180594 |
| tRNA-Arg-TCT-1-1 | 8.71325274 | 2.59410317 | 1.04305189 | 2.48703174 | 0.01288139 |
| tRNA-Leu-TAG-3-1 | 121.013674 | 0.93128737 | 0.38397946 | 2.42535723 | 0.01529333 |
| tRNA-Glu-CTC-2-1 | 367.676935 | 2.85099157 | 1.19412221 | 2.38752076 | 0.01696244 |
| tRNA-Gln-CTG-1-1 | 241.967211 | 0.83005567 | 0.35711085 | 2.32436416 | 0.02010598 |
| tRNA-Glu-CTC-3-1 | 18.9331099 | 1.65058827 | 0.73429795 | 2.24784538 | 0.02458605 |
| tRNA-Leu-CAA-2-1 | 362.290665 | -1.4955332 | 0.69436654 | -2.1538094 | 0.03125512 |
| tRNA-Ile-AAT-3-1 | 23.6073554 | -1.1999281 | 0.58094033 | -2.0654928 | 0.03887639 |
| tRNA-Thr-AGT-2-1 | 50.0235375 | 1.53004841 | 0.74479132 | 2.05433168 | 0.0399436 |

| | | | | | |
|---|---|---|---|---|---|
| tRNA-Glu-TTC-1-3 | 7516.66951 | 0.74635409 | 0.3677857 | 2.02931787 | 0.04242592 |
| tRNA-Glu-TTC-1-4 | 7540.22739 | 0.73769578 | 0.36624086 | 2.01423668 | 0.04398469 |
| tRNA-Ile-TAT-1-1 | 7.23774635 | 2.53020471 | 1.25972231 | 2.00854163 | 0.04458577 |
| tRNA-Glu-TTC-1-1 | 7533.07318 | 0.73546632 | 0.36640293 | 2.00726101 | 0.04472188 |
| tRNA-Glu-TTC-1-2 | 7533.07318 | 0.73546632 | 0.36640293 | 2.00726101 | 0.04472188 |
| tRNA-Ile-AAT-1-7 | 25.4197922 | -1.106405 | 0.5512907 | -2.0069357 | 0.04475651 |
| tRNA-Arg-ACG-3-1 | 39.6139245 | 1.30795076 | 0.6541441 | 1.99948413 | 0.045556 |
| tRNA-Arg-ACG-3-2 | 30.7468997 | 1.07548923 | 0.5544559 | 1.93972006 | 0.05241372 |
| tRNA-Thr-AGT-6-1 | 1.50954656 | -4.768021 | 2.47887907 | -1.9234585 | 0.05442249 |
| tRNA-Ala-TGC-1-1 | 24.514306 | 1.20524407 | 0.62801379 | 1.9191363 | 0.05496709 |
| tRNA-Ile-TAT-2-3 | 2.17857793 | 3.40861537 | 1.79376447 | 1.90025804 | 0.05739926 |
| tRNA-Gln-CTG-6-1 | 107.84331 | 0.78395357 | 0.41516411 | 1.88829804 | 0.05898595 |
| tRNA-Tyr-GTA-3-1 | 9.85513622 | -1.5126407 | 0.80702668 | -1.8743379 | 0.06088387 |
| tRNA-Gly-CCC-1-2 | 1464.64312 | 0.64838355 | 0.36243854 | 1.7889476 | 0.07362326 |
| tRNA-Gly-CCC-1-1 | 1464.64312 | 0.64838355 | 0.36243854 | 1.7889476 | 0.07362326 |
| tRNA-Arg-CCT-4-1 | 107.841334 | 0.79472362 | 0.44665121 | 1.77929358 | 0.07519164 |
| tRNA-Val-CAC-3-1 | 4033.67882 | 0.70959021 | 0.40094157 | 1.76980952 | 0.07675888 |
| tRNA-Val-AAC-3-1 | 3954.19939 | 0.70408036 | 0.40128971 | 1.75454374 | 0.07933738 |
| tRNA-Gly-CCC-4-1 | 1065.83419 | 0.8719639 | 0.49935959 | 1.74616432 | 0.0807824 |
| tRNA-Val-AAC-2-1 | 3983.63556 | 0.69291258 | 0.40001464 | 1.73221807 | 0.08323474 |
| tRNA-Val-AAC-2-2 | 3983.63556 | 0.69291258 | 0.40001464 | 1.73221807 | 0.08323474 |
| tRNA-Gly-CCC-3-1 | 1073.04047 | 0.85072249 | 0.49525292 | 1.71775362 | 0.08584156 |
| tRNA-Asn-GTT-3-4 | 24.0922481 | -1.0561671 | 0.62678923 | -1.6850435 | 0.09198019 |
| tRNA-Asn-GTT-3-5 | 24.0922481 | -1.0561671 | 0.62678923 | -1.6850435 | 0.09198019 |
| tRNA-Gly-TCC-1-2 | 390.66018 | 0.89191618 | 0.545771 | 1.63423155 | 0.10221024 |
| tRNA-Gly-TCC-1-3 | 390.66018 | 0.89191618 | 0.545771 | 1.63423155 | 0.10221024 |
| tRNA-Gly-TCC-1-4 | 390.66018 | 0.89191618 | 0.545771 | 1.63423155 | 0.10221024 |
| tRNA-Gly-TCC-1-7 | 393.953649 | 0.88379247 | 0.5427196 | 1.62845136 | 0.10342921 |
| tRNA-Gly-TCC-1-1 | 398.738556 | 0.86704416 | 0.53455045 | 1.62200622 | 0.10480202 |
| tRNA-Asn-GTT-3-9 | 24.368177 | -1.0093062 | 0.6260005 | -1.612309 | 0.10689472 |
| tRNA-Asn-GTT-3-3 | 23.967007 | -1.0215976 | 0.63382398 | -1.6118001 | 0.10700544 |
| tRNA-Cys-GCA-18-1 | 1.4036703 | -3.7163195 | 2.32259305 | -1.6000735 | 0.10958229 |
| tRNA-Gly-TCC-1-5 | 394.166849 | 0.86562101 | 0.54542571 | 1.58705574 | 0.11250002 |
| tRNA-Asn-GTT-1-1 | 15.3438029 | -1.1385974 | 0.71751948 | -1.5868522 | 0.11254612 |
| tRNA-Cys-GCA-7-1 | 272.909989 | -0.5018498 | 0.31763925 | -1.5799364 | 0.11412143 |
| tRNA-Arg-TCG-1-1 | 61.4602617 | -0.6054517 | 0.38972015 | -1.5535551 | 0.12029057 |
| tRNA-Tyr-GTA-2-1 | 35.6118649 | -0.973613 | 0.63281206 | -1.5385501 | 0.12391418 |
| tRNA-Trp-CCA-5-1 | 500.535416 | -0.7039413 | 0.4585431 | -1.5351693 | 0.12474224 |
| tRNA-Ala-CGC-4-1 | 1.76644092 | 2.96029908 | 1.92913118 | 1.53452451 | 0.12490067 |
| tRNA-Glu-CTC-1-4 | 12540.0237 | 0.58433004 | 0.38313862 | 1.52511391 | 0.12723069 |
| tRNA-Glu-CTC-1-7 | 12547.3672 | 0.58264652 | 0.38288499 | 1.52172725 | 0.12807744 |
| tRNA-Glu-CTC-1-8 | 12519.5887 | 0.58282115 | 0.38325598 | 1.52070985 | 0.12833267 |
| tRNA-Glu-CTC-1-5 | 12503.0878 | 0.58240419 | 0.38330875 | 1.51941275 | 0.12865864 |
| tRNA-Glu-CTC-1-6 | 12503.0878 | 0.58240419 | 0.38330875 | 1.51941275 | 0.12865864 |
| tRNA-Glu-CTC-1-9 | 12504.4328 | 0.58227117 | 0.38350901 | 1.51827246 | 0.12894573 |
| tRNA-Glu-CTC-1-1 | 12513.9485 | 0.58233487 | 0.3835958 | 1.518095 | 0.12899045 |

| | | | | | |
|---|---|---|---|---|---|
| tRNA-Glu-CTC-1-2 | 12513.9485 | 0.58233487 | 0.3835958 | 1.518095 | 0.12899045 |
| tRNA-Glu-CTC-1-3 | 12513.9485 | 0.58233487 | 0.3835958 | 1.518095 | 0.12899045 |
| tRNA-His-GTG-1-1 | 169.594431 | 0.66885439 | 0.44628402 | 1.4987191 | 0.13394652 |
| tRNA-Thr-TGT-2-1 | 29.0184782 | 0.7108 | 0.48001816 | 1.4807773 | 0.13866593 |
| tRNA-Ile-AAT-1-1 | 22.4318702 | -0.8198868 | 0.55410134 | -1.4796694 | 0.1389615 |
| tRNA-Leu-TAA-2-1 | 3119.15501 | -0.7407996 | 0.50161939 | -1.4768161 | 0.13972494 |
| tRNA-Asn-GTT-3-2 | 59.2855903 | -0.9548173 | 0.64680306 | -1.4762103 | 0.13988745 |
| tRNA-Trp-CCA-3-1 | 38.9630914 | -0.8162093 | 0.55482551 | -1.4711099 | 0.1412614 |
| tRNA-Cys-GCA-5-1 | 283.906768 | -0.4627113 | 0.31845484 | -1.4529886 | 0.14622691 |
| tRNA-Cys-GCA-4-13 | 269.242604 | -0.4742363 | 0.32646143 | -1.4526565 | 0.14631915 |
| tRNA-Lys-TTT-1-2 | 574.090964 | -0.920102 | 0.63560439 | -1.4476017 | 0.14772847 |
| tRNA-Cys-GCA-4-9 | 269.103436 | -0.4718502 | 0.32655393 | -1.4449379 | 0.14847533 |
| tRNA-Asp-GTC-4-1 | 215.689033 | 0.80748977 | 0.56285315 | 1.43463667 | 0.15139066 |
| tRNA-Cys-GCA-4-12 | 267.618534 | -0.4718185 | 0.32902777 | -1.4339777 | 0.15157862 |
| tRNA-SeC-TCA-1-1 | 10932.2345 | -0.5011883 | 0.35017289 | -1.4312595 | 0.15235585 |
| tRNA-Arg-ACG-2-1 | 9.19852009 | 1.17873511 | 0.82620052 | 1.42669374 | 0.15366819 |
| tRNA-Cys-GCA-9-1 | 267.201577 | -0.4710067 | 0.33091757 | -1.4233355 | 0.15463891 |
| tRNA-Cys-GCA-4-8 | 267.435139 | -0.4713517 | 0.33129408 | -1.4227593 | 0.15480594 |
| tRNA-Cys-GCA-24-1 | 267.526793 | -0.4711261 | 0.33128306 | -1.4221256 | 0.1549898 |
| tRNA-Cys-GCA-4-22 | 267.388749 | -0.4705626 | 0.33122441 | -1.4206761 | 0.15541095 |
| tRNA-Cys-GCA-4-23 | 267.388749 | -0.4705626 | 0.33122441 | -1.4206761 | 0.15541095 |
| tRNA-Cys-GCA-4-25 | 267.388749 | -0.4705626 | 0.33122441 | -1.4206761 | 0.15541095 |
| tRNA-Cys-GCA-4-28 | 267.388749 | -0.4705626 | 0.33122441 | -1.4206761 | 0.15541095 |
| tRNA-Cys-GCA-4-1 | 267.466104 | -0.4706352 | 0.3314108 | -1.420096 | 0.15557972 |
| tRNA-Cys-GCA-4-21 | 267.466104 | -0.4706352 | 0.3314108 | -1.420096 | 0.15557972 |
| tRNA-Cys-GCA-4-15 | 267.419715 | -0.469846 | 0.33134119 | -1.4180127 | 0.15618705 |
| tRNA-Cys-GCA-4-2 | 267.317117 | -0.4684989 | 0.33064976 | -1.4169038 | 0.15651104 |
| tRNA-Cys-GCA-4-4 | 267.317117 | -0.4684989 | 0.33064976 | -1.4169038 | 0.15651104 |
| tRNA-Cys-GCA-4-10 | 267.317117 | -0.4684989 | 0.33064976 | -1.4169038 | 0.15651104 |
| tRNA-Cys-GCA-4-11 | 267.317117 | -0.4684989 | 0.33064976 | -1.4169038 | 0.15651104 |
| tRNA-Cys-GCA-12-1 | 267.348082 | -0.4677823 | 0.33076688 | -1.4142356 | 0.15729275 |
| tRNA-Cys-GCA-4-16 | 267.348082 | -0.4677823 | 0.33076688 | -1.4142356 | 0.15729275 |
| tRNA-Cys-GCA-6-1 | 267.348082 | -0.4677823 | 0.33076688 | -1.4142356 | 0.15729275 |
| tRNA-Cys-GCA-4-17 | 267.348082 | -0.4677823 | 0.33076688 | -1.4142356 | 0.15729275 |
| tRNA-Cys-GCA-4-18 | 267.348082 | -0.4677823 | 0.33076688 | -1.4142356 | 0.15729275 |
| tRNA-Cys-GCA-4-19 | 267.348082 | -0.4677823 | 0.33076688 | -1.4142356 | 0.15729275 |
| tRNA-Cys-GCA-4-20 | 267.348082 | -0.4677823 | 0.33076688 | -1.4142356 | 0.15729275 |
| tRNA-Cys-GCA-4-24 | 267.348082 | -0.4677823 | 0.33076688 | -1.4142356 | 0.15729275 |
| tRNA-Cys-GCA-10-1 | 267.348082 | -0.4677823 | 0.33076688 | -1.4142356 | 0.15729275 |
| tRNA-Cys-GCA-4-26 | 267.348082 | -0.4677823 | 0.33076688 | -1.4142356 | 0.15729275 |
| tRNA-Cys-GCA-4-27 | 267.348082 | -0.4677823 | 0.33076688 | -1.4142356 | 0.15729275 |
| tRNA-Cys-GCA-4-29 | 267.348082 | -0.4677823 | 0.33076688 | -1.4142356 | 0.15729275 |
| tRNA-Ala-CGC-5-1 | 9.54769646 | 2.36832639 | 1.67543249 | 1.41356121 | 0.15749078 |
| tRNA-Cys-GCA-14-1 | 267.406934 | -0.4664962 | 0.33068932 | -1.4106781 | 0.15833956 |
| tRNA-Cys-GCA-4-3 | 267.406934 | -0.4664962 | 0.33068932 | -1.4106781 | 0.15833956 |
| tRNA-Cys-GCA-4-5 | 267.406934 | -0.4664962 | 0.33068932 | -1.4106781 | 0.15833956 |

| | | | | | |
|---|---|---|---|---|---|
| tRNA-Cys-GCA-4-6 | 267.406934 | -0.4664962 | 0.33068932 | -1.4106781 | 0.15833956 |
| tRNA-Cys-GCA-4-7 | 267.406934 | -0.4664962 | 0.33068932 | -1.4106781 | 0.15833956 |
| tRNA-Cys-GCA-4-14 | 267.406934 | -0.4664962 | 0.33068932 | -1.4106781 | 0.15833956 |
| tRNA-Lys-TTT-1-4 | 600.173947 | -0.8525759 | 0.60591197 | -1.4070953 | 0.15939912 |
| tRNA-Arg-CCT-2-1 | 19.3057441 | -0.8947287 | 0.63899624 | -1.4002096 | 0.16145056 |
| tRNA-Thr-AGT-4-1 | 13.5343955 | 1.30657994 | 0.93317808 | 1.40013998 | 0.16147141 |
| tRNA-Gln-CTG-2-4 | 312.494498 | 0.62428233 | 0.45024143 | 1.38655018 | 0.16557897 |
| tRNA-Gln-CTG-3-1 | 309.696451 | 0.59200373 | 0.42764444 | 1.38433633 | 0.16625549 |
| tRNA-Gln-CTG-2-3 | 313.626515 | 0.62611214 | 0.45251258 | 1.38363478 | 0.16647031 |
| tRNA-Gln-CTG-3-2 | 309.662189 | 0.59136471 | 0.42759105 | 1.38301469 | 0.16666036 |
| tRNA-Gln-CTG-3-3 | 309.634521 | 0.59085627 | 0.4278023 | 1.38114328 | 0.16723491 |
| tRNA-Gln-CTG-2-2 | 314.041103 | 0.61922577 | 0.44951391 | 1.37754528 | 0.16834373 |
| tRNA-Gly-CCC-2-1 | 3966.66658 | 0.73679209 | 0.53562935 | 1.37556333 | 0.16895686 |
| tRNA-Gly-CCC-2-2 | 3966.66658 | 0.73679209 | 0.53562935 | 1.37556333 | 0.16895686 |
| tRNA-Gln-CTG-2-1 | 313.078969 | 0.61777791 | 0.45009896 | 1.37253796 | 0.16989603 |
| tRNA-Gly-TCC-1-6 | 427.670255 | 0.75637274 | 0.55507968 | 1.36263812 | 0.17299659 |
| tRNA-Lys-CTT-2-2 | 3471.13671 | 0.41269246 | 0.30300452 | 1.36200102 | 0.17319756 |
| tRNA-Lys-CTT-2-1 | 3466.48099 | 0.40915009 | 0.30299291 | 1.35036193 | 0.17689992 |
| tRNA-Ala-AGC-6-1 | 18.0796148 | 0.97865578 | 0.72480893 | 1.35022589 | 0.17694354 |
| tRNA-Leu-CAG-2-3 | 715.525508 | 0.58838969 | 0.4382376 | 1.34262714 | 0.17939274 |
| tRNA-Lys-TTT-1-5 | 604.229473 | -0.7983208 | 0.59515841 | -1.3413585 | 0.17980408 |
| tRNA-Ile-AAT-1-3 | 22.9496487 | -0.7235456 | 0.54003538 | -1.3398114 | 0.18030667 |
| tRNA-Lys-TTT-1-3 | 636.401634 | -0.7788476 | 0.58200586 | -1.3382127 | 0.18082712 |
| tRNA-Phe-GAA-3-1 | 8.07769192 | 1.16990138 | 0.87451405 | 1.33777311 | 0.18097041 |
| tRNA-Asp-GTC-1-8 | 403.237111 | 0.70155001 | 0.53234709 | 1.31784324 | 0.18755613 |
| tRNA-Lys-TTT-1-6 | 603.189306 | -0.7953842 | 0.6037512 | -1.3174039 | 0.18770327 |
| tRNA-Leu-AAG-2-1 | 811.615521 | -0.5021359 | 0.38162594 | -1.3157804 | 0.18824776 |
| tRNA-Asp-GTC-1-13 | 403.162405 | 0.699907 | 0.53212135 | 1.31531463 | 0.18840418 |
| tRNA-Asp-GTC-1-6 | 403.617022 | 0.70055806 | 0.53272633 | 1.31504305 | 0.18849544 |
| tRNA-Asp-GTC-1-1 | 404.200032 | 0.70088866 | 0.53310878 | 1.31471978 | 0.1886041 |
| tRNA-Asp-GTC-1-7 | 403.159636 | 0.69988321 | 0.53235754 | 1.31468638 | 0.18861533 |
| tRNA-Asp-GTC-1-12 | 403.134072 | 0.69954401 | 0.53228192 | 1.31423591 | 0.18876683 |
| tRNA-Asp-GTC-1-2 | 403.039162 | 0.69887165 | 0.53237635 | 1.31273985 | 0.18927063 |
| tRNA-Asp-GTC-1-11 | 404.495896 | 0.69606681 | 0.53081771 | 1.31131044 | 0.1897529 |
| tRNA-Asp-GTC-1-3 | 402.920073 | 0.6978713 | 0.53227634 | 1.31110712 | 0.18982157 |
| tRNA-Asp-GTC-1-4 | 402.920073 | 0.6978713 | 0.53227634 | 1.31110712 | 0.18982157 |
| tRNA-Asp-GTC-1-9 | 402.920073 | 0.6978713 | 0.53227634 | 1.31110712 | 0.18982157 |
| tRNA-Lys-TTT-1-1 | 614.218266 | -0.7772818 | 0.59458889 | -1.3072592 | 0.1911247 |
| tRNA-Ala-AGC-7-1 | 17.8601853 | 0.93874057 | 0.72273337 | 1.29887536 | 0.19398671 |
| tRNA-Asp-GTC-1-5 | 404.36788 | 0.69083671 | 0.53361396 | 1.29463762 | 0.19544527 |
| tRNA-Asp-GTC-1-10 | 404.065018 | 0.68789043 | 0.5315598 | 1.29409792 | 0.1956316 |
| tRNA-Leu-AAG-3-1 | 798.841555 | -0.4985281 | 0.38596136 | -1.2916528 | 0.1964774 |
| tRNA-Gln-CTG-4-1 | 120.62929 | 0.6775224 | 0.52690547 | 1.28585191 | 0.19849475 |
| tRNA-Tyr-GTA-3-2 | 9.28585231 | -1.0506264 | 0.82225217 | -1.2777423 | 0.20134031 |
| tRNA-Ile-AAT-1-2 | 21.8607986 | -0.7124529 | 0.55769522 | -1.2774951 | 0.2014275 |
| tRNA-Ile-AAT-1-5 | 21.8607986 | -0.7124529 | 0.55769522 | -1.2774951 | 0.2014275 |

| | | | | | |
|---|---|---|---|---|---|
| tRNA-Trp-CCA-4-1 | 26.4858953 | -0.7343532 | 0.57731993 | -1.272004 | 0.20337171 |
| tRNA-Trp-CCA-4-2 | 26.4858953 | -0.7343532 | 0.57731993 | -1.272004 | 0.20337171 |
| tRNA-Glu-TTC-3-1 | 144.563688 | 0.62594804 | 0.49284881 | 1.27006099 | 0.20406291 |
| tRNA-Glu-TTC-3-2 | 144.563688 | 0.62594804 | 0.49284881 | 1.27006099 | 0.20406291 |
| tRNA-Arg-CCT-2-2 | 19.5824216 | -0.798801 | 0.64519829 | -1.2380705 | 0.21568993 |
| tRNA-Ser-GCT-1-1 | 1032.90289 | -0.6516482 | 0.53134354 | -1.2264159 | 0.2200422 |
| tRNA-Ile-AAT-1-6 | 22.6951187 | -0.6765566 | 0.55184187 | -1.2259973 | 0.2201997 |
| tRNA-Ala-AGC-2-2 | 85.7093237 | 0.53286823 | 0.43569281 | 1.22303654 | 0.22131589 |
| tRNA-Ser-GCT-3-1 | 1110.78244 | -0.6034343 | 0.50277619 | -1.2002046 | 0.23005988 |
| tRNA-Asn-GTT-3-6 | 24.7445466 | -0.6887736 | 0.57448156 | -1.1989481 | 0.23054814 |
| tRNA-Ser-GCT-4-2 | 1098.30062 | -0.59957 | 0.51221529 | -1.1705429 | 0.24178256 |
| tRNA-Ser-GCT-2-1 | 1106.56683 | -0.5956191 | 0.50915533 | -1.169818 | 0.24207423 |
| tRNA-Leu-CAG-2-1 | 695.833168 | 0.50070356 | 0.42841019 | 1.16874802 | 0.24250517 |
| tRNA-Ser-GCT-4-1 | 1098.54502 | -0.5969505 | 0.51214676 | -1.1655848 | 0.24378235 |
| tRNA-Arg-CCG-2-1 | 92.2280519 | -0.595353 | 0.51177533 | -1.1633092 | 0.24470407 |
| tRNA-Ser-GCT-4-3 | 1102.79447 | -0.5960426 | 0.51248409 | -1.163046 | 0.24481082 |
| tRNA-Met-CAT-5-1 | 8.28216092 | 1.10475793 | 0.95267043 | 1.15964335 | 0.24619404 |
| tRNA-Lys-CTT-13-1 | 7.44524461 | 1.41861886 | 1.22614063 | 1.15697892 | 0.24728097 |
| tRNA-Met-CAT-1-2 | 282.708567 | -0.6234454 | 0.5413577 | -1.1516331 | 0.24947188 |
| tRNA-Leu-TAG-2-1 | 796.037019 | -0.4337793 | 0.37688616 | -1.1509557 | 0.24975047 |
| tRNA-Asn-GTT-3-7 | 27.6734014 | -0.6245991 | 0.54580649 | -1.14436 | 0.25247435 |
| tRNA-Lys-TTT-2-2 | 26.9172173 | -0.9338322 | 0.81845268 | -1.1409728 | 0.25388126 |
| tRNA-iMet-CAT-3-1 | 1272.12042 | 0.77325295 | 0.68010414 | 1.13696257 | 0.25555393 |
| tRNA-Leu-CAG-1-4 | 673.662652 | 0.49077848 | 0.43189447 | 1.13633888 | 0.25581476 |
| tRNA-Arg-TCG-3-1 | 23.4064501 | -0.8169032 | 0.7189803 | -1.1361968 | 0.25587419 |
| tRNA-Arg-TCG-3-2 | 23.4064501 | -0.8169032 | 0.7189803 | -1.1361968 | 0.25587419 |
| tRNA-Ala-AGC-2-1 | 83.4787422 | 0.50364521 | 0.44616269 | 1.12883758 | 0.25896636 |
| tRNA-Ile-AAT-1-4 | 22.5194002 | -0.6253794 | 0.56095339 | -1.1148509 | 0.26491431 |
| tRNA-Arg-ACG-1-3 | 19.0353373 | 0.75767797 | 0.68523509 | 1.10571976 | 0.26884784 |
| tRNA-Leu-CAG-1-1 | 670.982084 | 0.47089388 | 0.42778606 | 1.10076956 | 0.27099696 |
| tRNA-Leu-CAG-1-2 | 670.982084 | 0.47089388 | 0.42778606 | 1.10076956 | 0.27099696 |
| tRNA-Leu-CAG-1-3 | 670.982084 | 0.47089388 | 0.42778606 | 1.10076956 | 0.27099696 |
| tRNA-Leu-CAG-2-2 | 682.200845 | 0.46165348 | 0.41965437 | 1.10008023 | 0.27129717 |
| tRNA-Leu-CAG-1-5 | 671.968702 | 0.4705054 | 0.42834487 | 1.09842661 | 0.27201825 |
| tRNA-Met-CAT-1-1 | 279.447803 | -0.6061397 | 0.5519147 | -1.0982488 | 0.27209586 |
| tRNA-Glu-TTC-2-1 | 5697.21165 | 0.3860517 | 0.3517137 | 1.09763055 | 0.27236585 |
| tRNA-Glu-TTC-2-2 | 5698.90117 | 0.38452707 | 0.35067045 | 1.09654828 | 0.27283891 |
| tRNA-Ala-AGC-1-1 | 59.737632 | 0.48566122 | 0.44429831 | 1.09309715 | 0.27435115 |
| tRNA-Tyr-GTA-1-4 | 3.89448409 | -1.2727121 | 1.16948318 | -1.0882688 | 0.27647644 |
| tRNA-Lys-TTT-2-1 | 27.0666458 | -0.8871515 | 0.81596145 | -1.0872468 | 0.27692775 |
| tRNA-Trp-CCA-6-1 | 19.3788719 | -0.6504069 | 0.60154426 | -1.0812286 | 0.27959543 |
| tRNA-Lys-CTT-1-1 | 19.2527981 | -0.969645 | 0.90322449 | -1.0735371 | 0.28303021 |
| tRNA-iMet-CAT-1-1 | 1324.81683 | 0.68583032 | 0.64693651 | 1.06011998 | 0.28909002 |
| tRNA-Gly-GCC-5-1 | 466.407171 | 0.55146539 | 0.52284781 | 1.05473404 | 0.29154698 |
| tRNA-Val-AAC-4-1 | 1646.3544 | 0.62820992 | 0.60467381 | 1.03892364 | 0.29884025 |
| tRNA-Ile-AAT-1-8 | 31.1021011 | -0.5119573 | 0.49905417 | -1.0258551 | 0.30495988 |

| | | | | | |
|---|---|---|---|---|---|
| tRNA-iMet-CAT-1-2 | 1316.71192 | 0.66564513 | 0.64890909 | 1.02579104 | 0.30499009 |
| tRNA-iMet-CAT-1-4 | 1316.71192 | 0.66564513 | 0.64890909 | 1.02579104 | 0.30499009 |
| tRNA-iMet-CAT-1-5 | 1316.68596 | 0.66529983 | 0.64874903 | 1.02551187 | 0.30512173 |
| tRNA-iMet-CAT-1-6 | 1323.30472 | 0.66516741 | 0.64884231 | 1.02516036 | 0.30528753 |
| tRNA-iMet-CAT-1-3 | 1325.69778 | 0.66364422 | 0.64853694 | 1.0232944 | 0.30616867 |
| tRNA-iMet-CAT-1-7 | 1318.84617 | 0.66282893 | 0.64784209 | 1.02313347 | 0.30624474 |
| tRNA-Asn-GTT-3-1 | 29.9718303 | -0.5384281 | 0.53012403 | -1.0156644 | 0.3097892 |
| tRNA-Asn-GTT-3-8 | 36.4757383 | -0.6294378 | 0.62389658 | -1.0088816 | 0.31303141 |
| tRNA-Gly-GCC-1-1 | 1698.42143 | 0.3619942 | 0.36090569 | 1.00301603 | 0.31585313 |
| tRNA-Gly-GCC-1-2 | 1698.42143 | 0.3619942 | 0.36090569 | 1.00301603 | 0.31585313 |
| tRNA-Gly-GCC-1-3 | 1698.42143 | 0.3619942 | 0.36090569 | 1.00301603 | 0.31585313 |
| tRNA-Lys-CTT-18-1 | 7.24927805 | 1.20575219 | 1.2160357 | 0.99154342 | 0.3214203 |
| tRNA-Cys-GCA-15-1 | 5.35612196 | -0.9232223 | 0.94635055 | -0.9755606 | 0.32928224 |
| tRNA-Asn-GTT-2-1 | 9.19200847 | -0.822016 | 0.86500359 | -0.9503036 | 0.34195801 |
| tRNA-Gly-TCC-2-1 | 243.066674 | 0.9527287 | 1.0060522 | 0.94699728 | 0.34364016 |
| tRNA-Met-CAT-4-1 | 9.82922744 | 1.02411574 | 1.08909413 | 0.94033721 | 0.34704462 |
| tRNA-Arg-TCG-4-1 | 24.7520366 | -0.6447692 | 0.7057988 | -0.9135312 | 0.36096324 |
| tRNA-Glu-CTC-5-1 | 4.82959767 | 1.53127802 | 1.72956421 | 0.88535482 | 0.37596529 |
| tRNA-Leu-TAA-5-1 | 2.15092461 | 1.36790919 | 1.58055367 | 0.86546203 | 0.38678525 |
| tRNA-Leu-TAA-5-2 | 2.15092461 | 1.36790919 | 1.58055367 | 0.86546203 | 0.38678525 |
| tRNA-Leu-TAA-5-3 | 2.15092461 | 1.36790919 | 1.58055367 | 0.86546203 | 0.38678525 |
| tRNA-Gln-TTG-6-1 | 5.84371917 | -0.9544518 | 1.13530103 | -0.8407037 | 0.40051393 |
| tRNA-Gln-TTG-3-2 | 184.203703 | -0.2900828 | 0.36053419 | -0.8045915 | 0.42105543 |
| tRNA-Val-AAC-5-1 | 33823.3676 | 0.3079381 | 0.3836031 | 0.80275184 | 0.42211818 |
| tRNA-Asp-GTC-2-1 | 1891.71821 | 0.23542611 | 0.29374003 | 0.80147778 | 0.4228551 |
| tRNA-Gln-TTG-3-1 | 184.544006 | -0.2865962 | 0.35967517 | -0.7968196 | 0.4255558 |
| tRNA-Arg-CCT-3-1 | 21.1317531 | 0.47667553 | 0.60183218 | 0.79204062 | 0.42833699 |
| tRNA-Leu-CAA-3-1 | 144.22626 | -0.4091213 | 0.51809456 | -0.7896653 | 0.42972329 |
| tRNA-Cys-GCA-25-1 | 0.77105215 | 2.05815536 | 2.61829607 | 0.7860667 | 0.43182841 |
| tRNA-Gln-TTG-4-1 | 0.42758692 | 2.86880636 | 3.65013463 | 0.78594536 | 0.4318995 |
| tRNA-Leu-CAA-1-1 | 79.2749203 | -0.3218113 | 0.42406426 | -0.7588738 | 0.44792805 |
| tRNA-Cys-GCA-11-1 | 1.7644337 | 1.41722077 | 1.86905456 | 0.75825543 | 0.44829808 |
| tRNA-Leu-TAA-3-1 | 0.76610446 | 1.98876389 | 2.74514461 | 0.72446598 | 0.4687797 |
| tRNA-Ser-CGA-3-1 | 53.5815648 | -0.4013647 | 0.58200542 | -0.6896236 | 0.49043094 |
| tRNA-Cys-GCA-19-1 | 6.12119462 | -0.6528799 | 0.9537264 | -0.6845568 | 0.49362361 |
| tRNA-Val-TAC-1-2 | 294.870895 | 0.27832054 | 0.40685177 | 0.6840834 | 0.4939225 |
| tRNA-Val-TAC-1-3 | 294.870895 | 0.27832054 | 0.40685177 | 0.6840834 | 0.4939225 |
| tRNA-Val-CAC-2-5 | 459699.745 | 0.24973082 | 0.37180121 | 0.67167833 | 0.5017885 |
| tRNA-Val-CAC-2-4 | 459696.983 | 0.24973033 | 0.37180463 | 0.67167084 | 0.50179327 |
| tRNA-Val-CAC-2-2 | 459699.312 | 0.24971482 | 0.37179899 | 0.67163933 | 0.50181333 |
| tRNA-Val-CAC-2-1 | 459699.535 | 0.24971599 | 0.3718011 | 0.67163864 | 0.50181378 |
| tRNA-Val-CAC-2-3 | 459702.447 | 0.24957893 | 0.3718038 | 0.67126513 | 0.50205164 |
| tRNA-Ile-TAT-2-1 | 1.06638822 | 1.51939374 | 2.27360788 | 0.66827431 | 0.50395851 |
| tRNA-Ile-TAT-2-2 | 1.06638822 | 1.51939374 | 2.27360788 | 0.66827431 | 0.50395851 |
| tRNA-Val-TAC-1-1 | 296.977326 | 0.27147065 | 0.40763492 | 0.66596514 | 0.50543338 |
| tRNA-Val-AAC-1-2 | 454075.646 | 0.24252643 | 0.3711823 | 0.65338899 | 0.51350553 |

| | | | | | |
|---|---|---|---|---|---|
| tRNA-Val-AAC-1-1 | 454079.297 | 0.24245266 | 0.37118364 | 0.65318789 | 0.51363515 |
| tRNA-Arg-TCT-3-1 | 2.75609899 | -1.0693049 | 1.66566511 | -0.6419687 | 0.5208935 |
| tRNA-Gln-CTG-7-1 | 2.46529023 | 1.0165209 | 1.5917736 | 0.63860897 | 0.52307735 |
| tRNA-Ser-CGA-1-1 | 54.8551022 | -0.3579479 | 0.56549959 | -0.6329765 | 0.526749 |
| tRNA-Lys-CTT-15-1 | 0.79335361 | -1.5480629 | 2.46750271 | -0.6273804 | 0.53040989 |
| tRNA-Trp-CCA-3-2 | 40.7546967 | -0.3121104 | 0.50496684 | -0.618081 | 0.53652192 |
| tRNA-Gln-TTG-5-1 | 41.1598104 | 0.37054327 | 0.6006808 | 0.61687217 | 0.53731904 |
| tRNA-Tyr-GTA-4-1 | 3.5628694 | -0.9961556 | 1.63609987 | -0.6088599 | 0.54261729 |
| tRNA-Arg-TCG-2-1 | 63.146195 | 0.3370217 | 0.55377664 | 0.60858779 | 0.5427977 |
| tRNA-Ala-AGC-4-1 | 82.8674516 | 0.26491398 | 0.43715756 | 0.60599199 | 0.54452007 |
| tRNA-Leu-CAA-4-1 | 81.1825193 | -0.2607647 | 0.43307927 | -0.6021177 | 0.54709581 |
| tRNA-Ala-AGC-4-2 | 85.6153283 | 0.27314736 | 0.45580906 | 0.59925829 | 0.54900066 |
| tRNA-Leu-TAA-4-1 | 0.84691153 | 1.68883897 | 2.84295623 | 0.59404325 | 0.55248318 |
| tRNA-Cys-GCA-17-1 | 5.85840989 | 0.56385373 | 0.95143348 | 0.592636 | 0.55342478 |
| tRNA-Phe-GAA-1-5 | 15.7564053 | 0.38220733 | 0.65506518 | 0.58346458 | 0.5595806 |
| tRNA-Lys-TTT-5-1 | 24.1426889 | 0.35777463 | 0.61987592 | 0.57717137 | 0.5638237 |
| tRNA-Lys-CTT-7-1 | 0.28965768 | -2.3370518 | 4.06938174 | -0.5743014 | 0.56576384 |
| tRNA-Pro-AGG-2-1 | 1.9991773 | 0.91687986 | 1.62064694 | 0.56574929 | 0.57156422 |
| tRNA-Met-CAT-3-1 | 30.246461 | 0.33793739 | 0.59840469 | 0.56473051 | 0.57225708 |
| tRNA-Tyr-GTA-5-1 | 0.29380037 | -3.4746178 | 6.15639213 | -0.5643919 | 0.57248745 |
| tRNA-Val-CAC-1-1 | 2394.51039 | 0.23853667 | 0.42873659 | 0.55637116 | 0.57795715 |
| tRNA-Ser-AGA-2-1 | 1069.48828 | 0.23952257 | 0.43744344 | 0.54755096 | 0.58400027 |
| tRNA-Ser-CGA-2-1 | 183.99309 | -0.2381903 | 0.43726352 | -0.5447294 | 0.58593964 |
| tRNA-Ser-AGA-2-5 | 1066.67852 | 0.23758278 | 0.43941827 | 0.54067569 | 0.58873114 |
| tRNA-Ala-CGC-3-1 | 259.731206 | 0.26669081 | 0.49424777 | 0.5395893 | 0.58948029 |
| tRNA-Ala-CGC-3-2 | 259.731206 | 0.26669081 | 0.49424777 | 0.5395893 | 0.58948029 |
| tRNA-Ala-CGC-3-3 | 259.731206 | 0.26669081 | 0.49424777 | 0.5395893 | 0.58948029 |
| tRNA-Arg-CCT-1-1 | 19.0254253 | -0.3630949 | 0.67346797 | -0.539142 | 0.58978888 |
| tRNA-Ser-TGA-2-1 | 1071.95997 | 0.23553472 | 0.43703858 | 0.53893348 | 0.58993275 |
| tRNA-Ala-TGC-4-1 | 365.042126 | 0.25847621 | 0.48000272 | 0.53848905 | 0.59023946 |
| tRNA-Ser-AGA-1-1 | 1069.59806 | 0.23553824 | 0.43815544 | 0.53756776 | 0.59087549 |
| tRNA-Ala-CGC-6-1 | 259.620535 | 0.26524435 | 0.49410876 | 0.5368137 | 0.59139631 |
| tRNA-Ser-AGA-2-2 | 1066.4647 | 0.2359274 | 0.44011429 | 0.53605939 | 0.59191751 |
| tRNA-Ser-TGA-2-2 | 1069.79911 | 0.2336298 | 0.43816244 | 0.53320362 | 0.59389263 |
| tRNA-Ser-AGA-2-3 | 1066.21343 | 0.23396832 | 0.44053476 | 0.53110071 | 0.595349 |
| tRNA-Ser-AGA-2-4 | 1066.21343 | 0.23396832 | 0.44053476 | 0.53110071 | 0.595349 |
| tRNA-Ser-AGA-2-6 | 1066.21343 | 0.23396832 | 0.44053476 | 0.53110071 | 0.595349 |
| tRNA-Ala-CGC-7-1 | 260.051064 | 0.26336243 | 0.4971622 | 0.5297314 | 0.59629818 |
| tRNA-Ala-AGC-5-1 | 68.668781 | 0.24772751 | 0.46979771 | 0.52730675 | 0.59798059 |
| tRNA-Asp-GTC-3-1 | 1.85075566 | 1.00112897 | 1.91327787 | 0.52325331 | 0.60079799 |
| tRNA-Cys-GCA-16-1 | 19.0894496 | 0.38213628 | 0.75520774 | 0.50600155 | 0.61285556 |
| tRNA-Phe-GAA-1-4 | 15.4291739 | -0.336778 | 0.70358159 | -0.4786623 | 0.6321789 |
| tRNA-Sup-TTA-1-1 | 0.10509283 | -2.9074342 | 6.17614301 | -0.4707524 | 0.63781756 |
| tRNA-Gly-GCC-2-3 | 82825.9032 | -0.2212773 | 0.47124899 | -0.4695551 | 0.63867294 |
| tRNA-Gly-GCC-4-1 | 82804.0993 | -0.2211098 | 0.47133434 | -0.4691146 | 0.63898774 |
| tRNA-Gly-GCC-2-1 | 82739.3231 | -0.2210737 | 0.47181544 | -0.4685596 | 0.63938445 |

| | | | | | |
|---|---|---|---|---|---|
| tRNA-Gly-GCC-2-2 | 82743.4953 | -0.2210413 | 0.47184635 | -0.4684604 | 0.63945538 |
| tRNA-Gly-GCC-2-4 | 82742.828 | -0.2208971 | 0.47181855 | -0.4681823 | 0.63965423 |
| tRNA-Gly-GCC-2-5 | 82743.8984 | -0.2208819 | 0.4718269 | -0.4681418 | 0.63968316 |
| tRNA-Gly-GCC-2-8 | 82771.4561 | -0.2207648 | 0.47178061 | -0.4679396 | 0.63982778 |
| tRNA-Gly-GCC-2-6 | 82751.08 | -0.2207546 | 0.47181404 | -0.4678847 | 0.63986702 |
| tRNA-Gly-GCC-2-7 | 82764.1107 | -0.2207017 | 0.47174068 | -0.4678453 | 0.63989522 |
| tRNA-Gly-GCC-3-1 | 82830.6429 | -0.2198168 | 0.47189641 | -0.4658159 | 0.64134729 |
| tRNA-Pro-GGG-1-1 | 0.09277906 | -2.7817369 | 6.18060659 | -0.4500751 | 0.65265632 |
| tRNA-Ala-TGC-3-1 | 408.032316 | 0.21448626 | 0.47933884 | 0.44746272 | 0.654541 |
| tRNA-Lys-TTT-3-1 | 0.26740619 | -2.6501864 | 5.93726026 | -0.4463652 | 0.65533346 |
| tRNA-Pro-AGG-3-1 | 7.72797027 | -0.4207864 | 0.94706479 | -0.4443058 | 0.65682149 |
| tRNA-Gly-CCC-5-1 | 81882.7096 | -0.2118592 | 0.47840281 | -0.4428468 | 0.65787655 |
| tRNA-Trp-CCA-1-1 | 25.9166198 | -0.2590102 | 0.58716694 | -0.4411184 | 0.65912725 |
| tRNA-Ala-TGC-7-1 | 341.857216 | 0.20999476 | 0.48420196 | 0.43369252 | 0.66451174 |
| tRNA-Ala-TGC-7-2 | 341.857216 | 0.20999476 | 0.48420196 | 0.43369252 | 0.66451174 |
| tRNA-Ala-TGC-2-1 | 532.573921 | 0.20486221 | 0.47430207 | 0.43192349 | 0.66579702 |
| tRNA-Ala-TGC-8-1 | 340.79098 | 0.20860434 | 0.48394112 | 0.43105314 | 0.66642973 |
| tRNA-Ala-AGC-3-1 | 359.507349 | 0.20748243 | 0.48258646 | 0.42993836 | 0.66724048 |
| tRNA-Tyr-GTA-1-2 | 3.21237119 | -0.5439657 | 1.26898471 | -0.4286621 | 0.66816914 |
| tRNA-Ala-TGC-6-1 | 593.053412 | -0.2168475 | 0.51037094 | -0.4248822 | 0.67092258 |
| tRNA-Arg-TCT-5-1 | 0.24748819 | -2.239413 | 5.30208862 | -0.4223643 | 0.67275913 |
| tRNA-Ala-AGC-12-1 | 67.9931028 | 0.19138635 | 0.46335379 | 0.41304582 | 0.67957305 |
| tRNA-Ala-AGC-8-1 | 345.193346 | 0.19720394 | 0.48146191 | 0.40959406 | 0.68210376 |
| tRNA-Leu-AAG-1-2 | 1011.53652 | -0.1426462 | 0.34960501 | -0.408021 | 0.68325826 |
| tRNA-Leu-AAG-1-3 | 1011.657 | -0.1414372 | 0.34977396 | -0.4043675 | 0.68594248 |
| tRNA-Ile-AAT-2-1 | 7.08639499 | -0.432993 | 1.07328296 | -0.4034286 | 0.68663295 |
| tRNA-Leu-AAG-1-1 | 1011.72894 | -0.140735 | 0.34906268 | -0.4031799 | 0.68681591 |
| tRNA-Arg-TCT-4-1 | 12.319731 | 0.42862264 | 1.06641581 | 0.40192825 | 0.68773683 |
| tRNA-Ala-TGC-5-3 | 2624.467 | 0.20398057 | 0.53463394 | 0.38153315 | 0.70280768 |
| tRNA-Tyr-GTA-1-1 | 3.24003894 | -0.4835935 | 1.26773523 | -0.3814625 | 0.70286008 |
| tRNA-Ala-TGC-5-1 | 2624.57532 | 0.20385744 | 0.5346475 | 0.38129318 | 0.70298572 |
| tRNA-Ala-TGC-5-2 | 2624.57532 | 0.20385744 | 0.5346475 | 0.38129318 | 0.70298572 |
| tRNA-Ala-CGC-2-1 | 443.307682 | 0.14856715 | 0.39294204 | 0.37808923 | 0.70536431 |
| tRNA-Ala-CGC-1-2 | 394.195456 | 0.17553264 | 0.46906979 | 0.37421433 | 0.70824486 |
| tRNA-Ala-AGC-5-2 | 66.9550485 | 0.17590791 | 0.47204553 | 0.3726503 | 0.70940873 |
| tRNA-Ala-AGC-5-3 | 66.9550485 | 0.17590791 | 0.47204553 | 0.3726503 | 0.70940873 |
| tRNA-Gln-TTG-2-1 | 92.281875 | 0.14941417 | 0.40697569 | 0.3671329 | 0.71351989 |
| tRNA-Arg-ACG-1-1 | 13.1807709 | 0.24821898 | 0.6814682 | 0.36424146 | 0.71567771 |
| tRNA-Thr-CGT-3-1 | 1.32481075 | -1.0479319 | 2.88556225 | -0.3631638 | 0.71648249 |
| tRNA-Ala-AGC-10-1 | 66.8530965 | 0.16777025 | 0.47099839 | 0.35620132 | 0.7216898 |
| tRNA-Arg-TCT-2-1 | 20.4532262 | 0.26495215 | 0.75603165 | 0.35045114 | 0.72600015 |
| tRNA-Arg-CCG-1-1 | 24.2194041 | 0.208932 | 0.61643886 | 0.33893385 | 0.73465956 |
| tRNA-Tyr-GTA-1-3 | 3.5270186 | -0.4202177 | 1.25106746 | -0.3358873 | 0.73695586 |
| tRNA-Ser-TGA-1-1 | 426.511732 | 0.16083818 | 0.47915023 | 0.3356738 | 0.73711687 |
| tRNA-Leu-CAG-4-1 | 27.9424695 | 0.22348645 | 0.68846387 | 0.32461609 | 0.74547165 |
| tRNA-Lys-CTT-6-1 | 1.4433289 | 0.72242517 | 2.23639149 | 0.32303162 | 0.7466713 |

| | | | | | |
|---|---|---|---|---|---|
| tRNA-Tyr-GTA-1-5 | 3.28708544 | -0.4000901 | 1.2635639 | -0.3166362 | 0.75151964 |
| tRNA-Ala-CGC-1-1 | 391.59381 | 0.15105592 | 0.48496244 | 0.31147962 | 0.75543603 |
| tRNA-Val-CAC-5-1 | 0.23619037 | -1.8254353 | 6.05552675 | -0.3014495 | 0.76307178 |
| tRNA-Tyr-GTA-6-1 | 4.68458016 | 0.4494987 | 1.52187869 | 0.29535777 | 0.76772059 |
| tRNA-Thr-AGT-3-1 | 14.0743672 | -0.2264148 | 0.78089996 | -0.2899409 | 0.77186147 |
| tRNA-Thr-AGT-1-2 | 14.1257781 | -0.2043446 | 0.74244103 | -0.2752335 | 0.78313685 |
| tRNA-Thr-AGT-1-3 | 14.1257781 | -0.2043446 | 0.74244103 | -0.2752335 | 0.78313685 |
| tRNA-Thr-AGT-1-1 | 14.1257781 | -0.2043446 | 0.74244103 | -0.2752335 | 0.78313685 |
| tRNA-Leu-TAG-1-1 | 1006.84179 | -0.0888032 | 0.34249525 | -0.2592829 | 0.79541698 |
| tRNA-Thr-CGT-1-1 | 20.4722709 | 0.20231309 | 0.82737196 | 0.24452495 | 0.80682428 |
| tRNA-Lys-CTT-10-1 | 2.16766622 | 0.35227603 | 1.4709345 | 0.23949131 | 0.81072463 |
| tRNA-Gln-TTG-1-1 | 168.678218 | -0.0802852 | 0.34168339 | -0.2349696 | 0.81423236 |
| tRNA-Leu-TAA-1-1 | 54.3850394 | -0.1453773 | 0.62005676 | -0.234458 | 0.81462945 |
| tRNA-Lys-CTT-4-1 | 0.11278317 | -1.42446 | 6.1909051 | -0.2300891 | 0.8180225 |
| tRNA-Ile-TAT-3-1 | 0.11139863 | -1.42446 | 6.1909051 | -0.2300891 | 0.8180225 |
| tRNA-Arg-ACG-1-2 | 12.9566281 | 0.16051769 | 0.70650959 | 0.22719818 | 0.82026964 |
| tRNA-Phe-GAA-1-3 | 14.6979462 | 0.1519703 | 0.70074529 | 0.21686953 | 0.82831003 |
| tRNA-Ser-GCT-5-1 | 13.6209205 | 0.17996174 | 0.83567891 | 0.21534795 | 0.82949605 |
| tRNA-Cys-GCA-3-1 | 3172.77916 | -0.0702096 | 0.34067025 | -0.2060926 | 0.83671856 |
| tRNA-Cys-GCA-2-1 | 3180.73817 | -0.0694484 | 0.34200071 | -0.203065 | 0.83908423 |
| tRNA-Leu-CAG-3-1 | 10.3740117 | 0.14892713 | 0.73855988 | 0.20164531 | 0.84019402 |
| tRNA-Gly-ACC-1-1 | 42.0915647 | -0.0945076 | 0.4686882 | -0.2016428 | 0.84019596 |
| tRNA-Cys-GCA-3-4 | 3170.81706 | -0.0672936 | 0.34042169 | -0.1976773 | 0.84329754 |
| tRNA-Lys-CTT-16-1 | 1.96092478 | -0.3010813 | 1.53610017 | -0.1960037 | 0.84460728 |
| tRNA-Thr-CGT-2-1 | 49.5938764 | 0.11699018 | 0.61315109 | 0.19080156 | 0.84868107 |
| tRNA-Cys-GCA-3-3 | 3185.20845 | -0.0640572 | 0.34078994 | -0.1879669 | 0.85090262 |
| tRNA-Thr-TGT-3-2 | 61.3470488 | 0.08316193 | 0.4443853 | 0.18713924 | 0.85155147 |
| tRNA-Cys-GCA-3-2 | 3185.66778 | -0.063307 | 0.34078789 | -0.1857664 | 0.85262793 |
| tRNA-Arg-CCG-3-1 | 685.49171 | -0.1066724 | 0.58743637 | -0.1815896 | 0.85590477 |
| tRNA-Cys-GCA-1-1 | 3162.87586 | -0.0619012 | 0.34197796 | -0.1810092 | 0.85636034 |
| tRNA-Cys-GCA-1-2 | 3179.62572 | -0.0597894 | 0.34297514 | -0.1743258 | 0.8616094 |
| tRNA-Thr-TGT-3-1 | 51.8670138 | 0.06805577 | 0.4645262 | 0.14650576 | 0.88352214 |
| tRNA-Cys-GCA-8-1 | 44.8066896 | -0.083611 | 0.62725359 | -0.133297 | 0.8939585 |
| tRNA-Cys-GCA-26-1 | 2.44593532 | 0.1901828 | 1.45773544 | 0.13046455 | 0.8961989 |
| tRNA-Lys-CTT-14-1 | 0.22859632 | -0.7031897 | 5.48620315 | -0.1281742 | 0.89801112 |
| tRNA-Cys-GCA-20-1 | 4.07688079 | 0.14457364 | 1.13331019 | 0.12756758 | 0.89849119 |
| tRNA-Ser-GGA-1-1 | 0.54460155 | 0.45275724 | 3.95945614 | 0.11434834 | 0.90896166 |
| tRNA-Met-CAT-2-2 | 238.585464 | -0.0701744 | 0.63896642 | -0.1098248 | 0.91254831 |
| tRNA-iMet-CAT-2-1 | 53.4682505 | 0.04285768 | 0.39286474 | 0.10909015 | 0.91313098 |
| tRNA-Cys-GCA-23-1 | 1.97794458 | -0.1644557 | 1.53541842 | -0.1071081 | 0.91470324 |
| tRNA-Phe-GAA-1-1 | 22.3229563 | 0.06523615 | 0.65735504 | 0.09924035 | 0.92094744 |
| tRNA-Val-CAC-4-1 | 473.594272 | -0.0323747 | 0.3668727 | -0.0882449 | 0.92968201 |
| tRNA-Met-CAT-2-1 | 238.96371 | -0.0510177 | 0.64250589 | -0.0794043 | 0.93671103 |
| tRNA-Ile-AAT-4-1 | 3.20242932 | -0.0846177 | 1.28504666 | -0.065848 | 0.94749888 |
| tRNA-Ile-GAT-1-1 | 0.15286675 | 0.39703912 | 6.16402281 | 0.06441234 | 0.94864191 |
| tRNA-Cys-GCA-13-1 | 21.9633135 | -0.0355974 | 0.55553924 | -0.0640772 | 0.94890876 |

| | | | | | |
|---|---|---|---|---|---|
| tRNA-Phe-GAA-2-1 | 0.91742194 | 0.11809002 | 2.40795473 | 0.04904163 | 0.96088612 |
| tRNA-Gln-CTG-5-1 | 85.5342562 | 0.03145722 | 0.64389298 | 0.04885474 | 0.96103506 |
| tRNA-Pro-TGG-3-1 | 0.08630054 | 0.2699482 | 6.16747031 | 0.04376968 | 0.965088 |
| tRNA-Pro-TGG-5-1 | 0.08300326 | 0.22980528 | 6.16862069 | 0.03725392 | 0.97028255 |
| tRNA-Met-CAT-6-1 | 1.81608629 | -0.0578995 | 1.66421301 | -0.0347909 | 0.97224647 |
| tRNA-Ser-GCT-6-1 | 12.6153113 | 0.02196172 | 0.77841062 | 0.02821354 | 0.97749184 |
| tRNA-Val-CAC-6-1 | 8198.89709 | -0.0107778 | 0.3869941 | -0.0278501 | 0.97778169 |
| tRNA-Phe-GAA-1-2 | 14.5147963 | 0.01578125 | 0.70228601 | 0.02247125 | 0.98207204 |
| tRNA-Ala-AGC-14-1 | 0.05863278 | -0.1343971 | 6.17919541 | -0.0217499 | 0.98264743 |
| tRNA-iMet-CAT-4-1 | 0.11553966 | 0.01813291 | 6.1909051 | 0.00292896 | 0.99766303 |
| tRNA-Asn-GTT-4-1 | 0.21611973 | -0.0157914 | 5.54895486 | -0.0028458 | 0.99772936 |

| ERACTION_ |
| --- |
| padj |
| 2.36E-07 |
| 2.36E-07 |
| 2.36E-07 |
| 2.36E-07 |
| 2.36E-07 |
| 2.36E-07 |
| 2.36E-07 |
| 2.36E-07 |
| 0.00079159 |
| 0.01649372 |
| 0.01649372 |
| 0.01649372 |
| 0.01649372 |
| 0.01649372 |
| 0.02646181 |
| 0.02646181 |
| 0.02646181 |
| 0.02646181 |
| 0.02646181 |
| 0.02646181 |
| 0.03530466 |
| 0.03940793 |
| 0.03940793 |
| 0.03940793 |
| 0.03940793 |
| 0.04380994 |
| 0.06657003 |
| 0.0751525 |
| 0.13178852 |
| 0.13178852 |
| 0.13178852 |
| 0.13178852 |
| 0.13871975 |
| 0.13871975 |
| 0.13871975 |
| 0.13871975 |
| 0.14726563 |
| 0.17023891 |
| 0.18397728 |
| 0.21262079 |
| 0.25365609 |
| 0.31478372 |
| 0.3778468 |
| 0.3778468 |

| |
|---|
| 0.3778468 |
| 0.3778468 |
| 0.3778468 |
| 0.3778468 |
| 0.3778468 |
| 0.3778468 |
| 0.3778468 |
| 0.42636545 |
| 0.43057551 |
| 0.43057551 |
| 0.44145252 |
| 0.44555459 |
| 0.45182237 |
| 0.46407855 |
| 0.46407855 |
| 0.46407855 |
| 0.46407855 |
| 0.46407855 |
| 0.46407855 |
| 0.46407855 |
| 0.46407855 |
| 0.46407855 |
| 0.46407855 |
| 0.46407855 |
| 0.46407855 |
| 0.46407855 |
| 0.46407855 |
| 0.46407855 |
| 0.46407855 |
| 0.46407855 |
| 0.46407855 |
| 0.46407855 |
| 0.46407855 |
| 0.46407855 |
| 0.46407855 |
| 0.46407855 |
| 0.46407855 |
| 0.46407855 |
| 0.46407855 |
| 0.46407855 |
| 0.46407855 |
| 0.46407855 |
| 0.46407855 |
| 0.46407855 |
| 0.46407855 |
| 0.46407855 |

0.46407855
0.46407855
0.46407855
0.46407855
0.46407855
0.46407855
0.46407855
0.46407855
0.46407855
0.46407855
0.46407855
0.46407855
0.46407855
0.46407855
0.46407855
0.46407855
0.46407855
0.46407855
0.46407855
0.46407855
0.46407855
0.46407855
0.46407855
0.46407855
0.46407855
0.46407855
0.46407855
0.46407855
0.46407855
0.46407855
0.46407855
0.46407855
0.46407855
0.46407855
0.46407855
0.46407855
0.46407855
0.46407855
0.46407855
0.46407855
0.46407855
0.46407855
0.46407855
0.46407855
0.46407855
0.46407855

0.46407855
0.46407855
0.46407855
0.46407855
0.46407855
0.46407855
0.46407855
0.46407855
0.46407855
0.46407855
0.46407855
0.46407855
0.46407855
0.46407855
0.46407855
0.46407855
0.46407855
0.46407855
0.46407855
0.46407855
0.46407855
0.46407855
0.46407855
0.46407855
0.46407855
0.46407855
0.46407855
0.46407855
0.46407855
0.46407855
0.46407855
0.46407855
0.46407855
0.46407855
0.46407855
0.46407855
0.46407855
0.46407855
0.46407855
0.46407855
0.46407855
0.46407855
0.46407855
0.46407855
0.46407855
0.46407855

0.46407855
0.46407855
0.46407855
0.46407855
0.48789755
0.49271907
0.49271907
0.49271907
0.50792638
0.50792638
0.52035953
0.52035953
0.52035953
0.52035953
0.52035953
0.52035953
0.52035953
0.52035953
0.52035953
0.52035953
0.52035953
0.52035953
0.52035953
0.52035953
0.52035953
0.52035953
0.52412808
0.52699022
0.52699022
0.52699022
0.52699022
0.52699022
0.52699022
0.52699022
0.52699022
0.52699022
0.52699022
0.52750243
0.52765964
0.52765964
0.53035366
0.53447223
0.54348923
0.54568307
0.55359627
0.55359627

| |
|---|
| 0.55359627 |
| 0.55359627 |
| 0.55359627 |
| 0.55359627 |
| 0.55359627 |
| 0.55359627 |
| 0.55762057 |
| 0.55902039 |
| 0.55902039 |
| 0.55902039 |
| 0.55902039 |
| 0.56650328 |
| 0.57795182 |
| 0.59772 |
| 0.59818843 |
| 0.60163883 |
| 0.62321409 |
| 0.6464769 |
| 0.65706892 |
| 0.65706892 |
| 0.65706892 |
| 0.67766956 |
| 0.70698699 |
| 0.70698699 |
| 0.70698699 |
| 0.7081143 |
| 0.7081143 |
| 0.7081143 |
| 0.7081143 |
| 0.7081143 |
| 0.7293465 |
| 0.7293465 |
| 0.75974642 |
| 0.78314403 |
| 0.78314403 |
| 0.78314403 |
| 0.78314403 |
| 0.78314403 |
| 0.78314403 |
| 0.78314403 |
| 0.78314403 |
| 0.78314403 |
| 0.78314403 |
| 0.78314403 |
| 0.78314403 |
| 0.79006425 |

| |
|---|
| 0.79006425 |
| 0.79832591 |
| 0.79877876 |
| 0.80149219 |
| 0.80416983 |
| 0.80813351 |
| 0.80813351 |
| 0.80813351 |
| 0.80813351 |
| 0.80813351 |
| 0.80813351 |
| 0.80813351 |
| 0.80813351 |
| 0.80813351 |
| 0.80813351 |
| 0.80813351 |
| 0.80813351 |
| 0.80813351 |
| 0.80813351 |
| 0.80813351 |
| 0.80813351 |
| 0.80813351 |
| 0.80813351 |
| 0.80813351 |
| 0.80813351 |
| 0.80813351 |
| 0.80813351 |
| 0.80813351 |
| 0.80813351 |
| 0.80813351 |
| 0.80813351 |
| 0.80813351 |
| 0.80813351 |
| 0.80813351 |
| 0.80813351 |
| 0.80813351 |
| 0.80813351 |
| 0.80813351 |
| 0.80813351 |
| 0.80935525 |
| 0.82297747 |
| 0.82963274 |
| 0.82963274 |
| 0.82963274 |
| 0.82963274 |
| 0.82963274 |

| |
|---|
| 0.82963274 |
| 0.82963274 |
| 0.82963274 |
| 0.82963274 |
| 0.82963274 |
| 0.82963274 |
| 0.82963274 |
| 0.83373318 |
| 0.83373318 |
| 0.83373318 |
| 0.83373318 |
| 0.83373318 |
| 0.83373318 |
| 0.83373318 |
| 0.83373318 |
| 0.83373318 |
| 0.83373318 |
| 0.83373318 |
| 0.83373318 |
| 0.83453698 |
| 0.83453698 |
| 0.83595598 |
| 0.83595598 |
| 0.83595598 |
| 0.83595598 |
| 0.83595598 |
| 0.83595598 |
| 0.83595598 |
| 0.84292105 |
| 0.84292105 |
| 0.84292105 |
| 0.84292105 |
| 0.84292105 |
| 0.84292105 |
| 0.84292105 |
| 0.84292105 |
| 0.84421196 |
| 0.84421196 |
| 0.84421196 |
| 0.84798552 |
| 0.85068716 |
| 0.8565946 |
| 0.8565946 |
| 0.8565946 |
| 0.86295617 |
| 0.86295617 |

0.86619294
0.86834087
0.87474082
0.87769137
0.88004691
0.88574034
0.88574034
0.88574034
0.89723035
0.90767731
0.90819821
0.90819821
0.90819821
0.90819821
0.90819821
0.90830906
0.91374174
0.91374174
0.91474854
0.91474854
0.91474854
0.91474854
0.91474854
0.91474854
0.91474854
0.91474854
0.91474854
0.91474854
0.91474854
0.91474854
0.91803722
0.93901976
0.94542729
0.94542729
0.94542729
0.94542729
0.95300362
0.95300362
0.95300362
0.95300362
0.95715176
0.96386149
0.96877449
0.97424371
0.97424371
0.97424371

| |
|---|
| 0.98192713 |
| 0.98192713 |
| 0.9836921 |
| 0.98623563 |
| 0.98623563 |
| 0.98711612 |
| 0.98711612 |
| 0.98731559 |
| 0.98731559 |
| 0.99772936 |
| 0.99772936 |

| HS692_miRNA_pvalues_TREATMENT_p | | | | HS692_miRNA_pvalues_TREATMENT_p | | |
|---|---|---|---|---|---|---|
| | baseMean | log2FoldChai | lfcSE | stat | pvalue | padj |
| miR-150-5p | 67.1664819 | 1.89159062 | 0.54958696 | 3.44184044 | 0.00057777 | 0.27064526 |
| miR-1247-3p | 10.5798277 | -4.5183069 | 1.35124982 | -3.3437984 | 0.0008264 | 0.27064526 |
| miR-410-3p | 4.64296563 | -4.4148953 | 1.5512113 | -2.8460954 | 0.00442589 | 0.96632008 |
| miR-148b-3p | 110.979738 | 1.44744749 | 0.53222739 | 2.71960355 | 0.00653602 | 0.99310104 |
| miR-147-5p | 13.8428067 | -1.824124 | 0.71943562 | -2.5354931 | 0.01122892 | 0.99310104 |
| miR-880-3p | 8.06722271 | -3.2446189 | 1.28581319 | -2.5233983 | 0.01162267 | 0.99310104 |
| miR-7230-3p | 46.0677911 | 1.08878636 | 0.44210186 | 2.46275004 | 0.0137876 | 0.99310104 |
| miR-878-3p | 5.15579097 | 2.241357 | 0.99111617 | 2.26144731 | 0.02373157 | 0.99310104 |
| miR-6538 | 16.2051488 | 1.48455689 | 0.68206797 | 2.17655272 | 0.02951396 | 0.99310104 |
| miR-130b-5p | 38.0105853 | -1.4830221 | 0.68579646 | -2.1624813 | 0.03058109 | 0.99310104 |
| miR-195a-5p | 150.334642 | 0.69154004 | 0.32269351 | 2.14302436 | 0.03211114 | 0.99310104 |
| miR-148a-5p | 40.6169889 | -1.0579062 | 0.50032798 | -2.1144254 | 0.03447895 | 0.99310104 |
| miR-345-3p | 5.44312551 | 2.88251789 | 1.4679985 | 1.96357005 | 0.04957997 | 0.99310104 |
| miR-434-3p | 23.72971 | -1.37189 | 0.7149573 | -1.9188419 | 0.05500435 | 0.99310104 |
| miR-541-5p | 85.6242488 | -0.7357164 | 0.38970126 | -1.8878983 | 0.0590396 | 0.99310104 |
| miR-125a-3p | 3.50064133 | 3.29843887 | 1.76773263 | 1.86591503 | 0.06205325 | 0.99310104 |
| miR-34c-3p | 262.034111 | 0.69549207 | 0.37520632 | 1.85362569 | 0.06379273 | 0.99310104 |
| miR-205-5p | 15.6252425 | -1.6401431 | 0.90135356 | -1.8196445 | 0.06881316 | 0.99310104 |
| miR-409-3p | 31.25176 | -0.8474123 | 0.46845631 | -1.8089462 | 0.07045936 | 0.99310104 |
| miR-10b-5p | 392080.628 | 0.60552586 | 0.33900204 | 1.78620125 | 0.07406668 | 0.99310104 |
| miR-16-2-3p | 12.7825411 | -1.3760094 | 0.77745822 | -1.7698821 | 0.07674679 | 0.99310104 |
| miR-574-3p | 9.99368537 | 1.7686421 | 1.00663357 | 1.75698699 | 0.07892003 | 0.99310104 |
| miR-30b-5p | 328.954044 | -0.602079 | 0.34378418 | -1.7513284 | 0.07988936 | 0.99310104 |
| miR-141-3p | 43.6745166 | 1.11601689 | 0.64689214 | 1.7251978 | 0.08449183 | 0.99310104 |
| miR-468-5p | 8.93206347 | 1.34475931 | 0.82803672 | 1.62403343 | 0.10436867 | 0.99310104 |
| miR-296-5p | 15.7564006 | -1.2714864 | 0.78673814 | -1.6161494 | 0.10606201 | 0.99310104 |
| miR-212-5p | 1.00092627 | 4.61961482 | 2.86103674 | 1.61466463 | 0.10638334 | 0.99310104 |
| miR-10a-5p | 171033.381 | 0.57593409 | 0.35795652 | 1.60894984 | 0.10762731 | 0.99310104 |
| miR-31-5p | 5.93726446 | -1.739364 | 1.08459237 | -1.603703 | 0.10877954 | 0.99310104 |
| miR-196b-5p | 26.3362485 | 1.09040005 | 0.68280059 | 1.59695241 | 0.11027632 | 0.99310104 |
| miR-496a-3p | 1.14879701 | -3.8644673 | 2.42251947 | -1.5952265 | 0.11066159 | 0.99310104 |
| miR-15b-5p | 189.084791 | 0.52829449 | 0.33190711 | 1.59169378 | 0.11145353 | 0.99310104 |
| miR-7217-3p | 97.335608 | -0.7709961 | 0.48919134 | -1.5760624 | 0.11501142 | 0.99310104 |
| miR-295-3p | 3.42954705 | -2.5230381 | 1.62637407 | -1.5513271 | 0.12082332 | 0.99310104 |
| miR-465a-5p | 68.8682584 | -0.7732824 | 0.50476411 | -1.5319678 | 0.12553037 | 0.99310104 |
| miR-425-5p | 443.12411 | -0.5146606 | 0.33841376 | -1.5208028 | 0.12830934 | 0.99310104 |
| miR-672-5p | 40.9597656 | 0.76053183 | 0.50654411 | 1.50141284 | 0.13324881 | 0.99310104 |
| miR-30c-2-3p | 53.1794842 | -0.6176285 | 0.41347202 | -1.4937612 | 0.13523803 | 0.99310104 |
| miR-125b-2-3p | 14.8456703 | -1.1272334 | 0.75891513 | -1.4853221 | 0.13745853 | 0.99310104 |
| miR-669c-5p | 27.7189097 | 0.93527454 | 0.63172563 | 1.48050751 | 0.13873786 | 0.99310104 |
| miR-212-3p | 2.43251707 | 2.68181862 | 1.81937372 | 1.47403394 | 0.14047245 | 0.99310104 |
| miR-23b-3p | 63.4340343 | 0.56948567 | 0.38713346 | 1.4710319 | 0.14128249 | 0.99310104 |
| miR-148a-3p | 4078.96285 | -0.4897932 | 0.33460025 | -1.463816 | 0.14324423 | 0.99310104 |
| miR-471-3p | 5.65147346 | 1.53163741 | 1.07078103 | 1.43039274 | 0.15260433 | 0.99310104 |

| | | | | | | |
|---|---|---|---|---|---|---|
| let-7g-5p | 1132.43314 | -0.4723854 | 0.33061755 | -1.428797 | 0.1530626 | 0.99310104 |
| miR-22-3p | 1217.90203 | -0.5556252 | 0.39274677 | -1.414716 | 0.15715177 | 0.99310104 |
| miR-878-5p | 70.3483372 | -0.6213546 | 0.43971546 | -1.4130834 | 0.15763121 | 0.99310104 |
| miR-429-3p | 97.3046123 | -0.8308946 | 0.58956934 | -1.4093247 | 0.15873918 | 0.99310104 |
| miR-25-3p | 1357.9244 | -0.4477694 | 0.31868904 | -1.4050355 | 0.16001073 | 0.99310104 |
| miR-133a-3p | 13.2516827 | 1.14571369 | 0.82865666 | 1.38261569 | 0.16678274 | 0.99310104 |
| miR-30d-5p | 1760.5325 | -0.474497 | 0.34539544 | -1.373779 | 0.1695103 | 0.99310104 |
| miR-335-3p | 3.1411732 | -2.1500141 | 1.56685903 | -1.372181 | 0.17000711 | 0.99310104 |
| miR-741-3p | 127.971851 | -0.6588198 | 0.48399567 | -1.3612101 | 0.1734473 | 0.99310104 |
| miR-676-3p | 309.355277 | -0.4919613 | 0.36160932 | -1.3604773 | 0.17367893 | 0.99310104 |
| miR-18a-3p | 2.41760672 | -2.3091962 | 1.71049759 | -1.3500143 | 0.17701139 | 0.99310104 |
| miR-3066-5p | 1.6247547 | -2.8264096 | 2.09558282 | -1.3487463 | 0.17741847 | 0.99310104 |
| miR-139-5p | 10.5989984 | -1.2935282 | 0.96071227 | -1.3464263 | 0.17816508 | 0.99310104 |
| miR-199a-3p | 14.4442038 | 0.91286967 | 0.68627085 | 1.33018862 | 0.18345613 | 0.99310104 |
| miR-3086-5p | 1.32850623 | -3.3645941 | 2.57202198 | -1.3081514 | 0.19082196 | 0.99310104 |
| miR-1198-5p | 3.55459749 | 1.5051274 | 1.15242384 | 1.30605368 | 0.1915343 | 0.99310104 |
| miR-132-5p | 0.97063471 | 4.41524814 | 3.41203686 | 1.29402123 | 0.19565809 | 0.99310104 |
| miR-1981-3p | 3.38152849 | 2.07954239 | 1.60834267 | 1.29297222 | 0.19602067 | 0.99310104 |
| miR-92b-3p | 72.4786355 | 0.63356812 | 0.51157733 | 1.23846013 | 0.2155455 | 0.99310104 |
| miR-28a-5p | 6.70795392 | 1.09127592 | 0.88116033 | 1.2384533 | 0.21554803 | 0.99310104 |
| miR-191-3p | 12.6768131 | -0.9472964 | 0.77972666 | -1.2149083 | 0.22440106 | 0.99310104 |
| miR-98-5p | 115.641042 | -0.6024355 | 0.49835326 | -1.2088523 | 0.2267196 | 0.99310104 |
| miR-3081-3p | 2.78968103 | -2.5982313 | 2.15378406 | -1.2063565 | 0.22768008 | 0.99310104 |
| miR-204-5p | 276.046951 | 0.4488429 | 0.37365652 | 1.2012179 | 0.22966669 | 0.99310104 |
| miR-203-3p | 21.1211209 | -0.8595581 | 0.72576094 | -1.1843543 | 0.23627283 | 0.99310104 |
| miR-1843a-3p | 3.48033372 | 1.57605104 | 1.33376083 | 1.18165941 | 0.23734087 | 0.99310104 |
| miR-298-5p | 7.23727778 | 0.92525087 | 0.78463958 | 1.17920494 | 0.23831658 | 0.99310104 |
| miR-467c-5p | 3.46959286 | 1.67855193 | 1.43970199 | 1.16590235 | 0.24365393 | 0.99310104 |
| miR-742-5p | 0.86779662 | -3.118034 | 2.67436749 | -1.1658959 | 0.24365655 | 0.99310104 |
| miR-485-5p | 4.73326633 | 1.50243681 | 1.28871978 | 1.16583669 | 0.24368048 | 0.99310104 |
| miR-7240-5p | 0.67367875 | 4.85051478 | 4.22907456 | 1.14694473 | 0.25140446 | 0.99310104 |
| let-7d-3p | 83.7901595 | 0.47183721 | 0.41424901 | 1.13901832 | 0.25469551 | 0.99310104 |
| miR-883b-5p | 3.33547215 | 1.51998221 | 1.33559839 | 1.13805334 | 0.25509821 | 0.99310104 |
| miR-7068-5p | 2.11119042 | -2.488329 | 2.19131282 | -1.1355426 | 0.25614806 | 0.99310104 |
| miR-211-5p | 2.84205704 | 2.11144212 | 1.87258393 | 1.1275554 | 0.25950773 | 0.99310104 |
| miR-187-3p | 1.50236409 | -2.7329178 | 2.42677735 | -1.126151 | 0.2601016 | 0.99310104 |
| miR-26a-1-3p | 0.46171449 | 4.96220531 | 4.429797 | 1.12018797 | 0.26263367 | 0.99310104 |
| miR-139-3p | 0.76411505 | -3.1509405 | 2.82298582 | -1.116173 | 0.2643481 | 0.99310104 |
| miR-669a-5p | 18.083587 | -0.6834295 | 0.61424585 | -1.1126318 | 0.26586658 | 0.99310104 |
| miR-93-5p | 224.45357 | -0.3972429 | 0.35911225 | -1.1061803 | 0.26864847 | 0.99310104 |
| miR-470-5p | 5636.05708 | 0.42454193 | 0.38417015 | 1.1050883 | 0.26912133 | 0.99310104 |
| miR-145a-5p | 61.9461242 | 0.83771851 | 0.75842662 | 1.10454787 | 0.26935555 | 0.99310104 |
| miR-1843b-3p | 32.6640078 | 0.57182242 | 0.52230015 | 1.09481573 | 0.27359737 | 0.99310104 |
| miR-206-3p | 1.83282319 | -2.3372302 | 2.14781553 | -1.0881894 | 0.27651149 | 0.99310104 |
| miR-214-3p | 3.82059379 | 1.3077995 | 1.20395843 | 1.08624971 | 0.27736853 | 0.99310104 |
| miR-7225-5p | 0.93754262 | 3.59318495 | 3.32663493 | 1.08012602 | 0.28008607 | 0.99310104 |

| | | | | | | |
|---|---|---|---|---|---|---|
| miR-145a-3p | 15.9145371 | 1.00404034 | 0.93486287 | 1.07399745 | 0.28282383 | 0.99310104 |
| miR-200a-3p | 6.9552802 | -1.0539664 | 0.98160796 | -1.0737141 | 0.28295082 | 0.99310104 |
| miR-300-3p | 2.81409544 | 1.95303228 | 1.82913116 | 1.06773769 | 0.28563885 | 0.99310104 |
| miR-324-5p | 0.83313136 | -3.2306036 | 3.03866656 | -1.0631649 | 0.28770719 | 0.99310104 |
| miR-182-5p | 3490.89211 | -0.3210466 | 0.30358943 | -1.0575027 | 0.29028223 | 0.99310104 |
| miR-144-5p | 1.6177354 | -2.3381996 | 2.22237103 | -1.0521194 | 0.29274478 | 0.99310104 |
| miR-421-3p | 1.79965379 | -2.3757615 | 2.26369231 | -1.0495073 | 0.29394472 | 0.99310104 |
| miR-883a-5p | 2.23124848 | -2.3927232 | 2.30278745 | -1.0390552 | 0.29877909 | 0.99310104 |
| miR-669o-5p | 4.38164049 | 1.43750483 | 1.39199378 | 1.03269487 | 0.30174672 | 0.99310104 |
| miR-23a-3p | 88.1362464 | 0.47129409 | 0.45699022 | 1.03130017 | 0.30240008 | 0.99310104 |
| miR-184-3p | 2339.78942 | 0.35660603 | 0.34663565 | 1.02876327 | 0.30359093 | 0.99310104 |
| miR-3070-5p | 0.64518778 | 4.43044742 | 4.31468205 | 1.02683057 | 0.30450025 | 0.99310104 |
| miR-27b-3p | 459.865937 | -0.3259598 | 0.319267 | -1.0209632 | 0.3072719 | 0.99310104 |
| miR-146b-5p | 843.96163 | 0.31542638 | 0.3107919 | 1.01491182 | 0.31014786 | 0.99310104 |
| miR-181b-5p | 32.2043317 | -0.7708578 | 0.761406 | -1.0124136 | 0.31134032 | 0.99310104 |
| miR-467a-3p | 5.38817566 | 1.17925574 | 1.16650554 | 1.01093025 | 0.31204981 | 0.99310104 |
| miR-881-3p | 131.778904 | -0.3721243 | 0.37298823 | -0.9976838 | 0.31843272 | 0.99310104 |
| miR-3064-5p | 1.90725672 | 4.43261098 | 4.44834915 | 0.99646202 | 0.31902571 | 0.99310104 |
| miR-7036b-5p | 0.32535072 | 4.43185938 | 4.44837887 | 0.9962864 | 0.31911101 | 0.99310104 |
| miR-7214-5p | 17.3857301 | 0.83656874 | 0.84394227 | 0.991263 | 0.32155718 | 0.99310104 |
| miR-7688-5p | 0.7748139 | -2.8102062 | 2.84492663 | -0.9877957 | 0.32325274 | 0.99310104 |
| miR-6905-3p | 0.30780966 | 4.3788677 | 4.45063185 | 0.98387551 | 0.32517672 | 0.99310104 |
| miR-7234-3p | 30.7430264 | 0.58530941 | 0.60233465 | 0.97173458 | 0.33118261 | 0.99310104 |
| miR-130b-3p | 37.360924 | 0.46663264 | 0.48076896 | 0.97059643 | 0.33174928 | 0.99310104 |
| miR-328-3p | 14.3504627 | 0.60198024 | 0.62048863 | 0.97017126 | 0.33196113 | 0.99310104 |
| miR-741-5p | 6.63699476 | -1.1619033 | 1.19767531 | -0.9701321 | 0.33198066 | 0.99310104 |
| miR-202-5p | 4.12620884 | 1.25919145 | 1.29857739 | 0.96966993 | 0.33221104 | 0.99310104 |
| miR-30c-1-3p | 4.35338289 | 1.17207973 | 1.20933571 | 0.96919302 | 0.33244889 | 0.99310104 |
| miR-449a-5p | 84.0485271 | -0.7211289 | 0.74748226 | -0.9647439 | 0.33467311 | 0.99310104 |
| miR-221-5p | 15.7134515 | 0.69769249 | 0.72552698 | 0.96163549 | 0.33623274 | 0.99310104 |
| miR-465c-5p | 181.957168 | -0.3216077 | 0.33764379 | -0.9525058 | 0.34084051 | 0.99310104 |
| miR-15a-3p | 0.82085359 | 2.53523283 | 2.68035053 | 0.94585869 | 0.34422067 | 0.99310104 |
| miR-222-5p | 1.11668659 | -2.3276644 | 2.46583735 | -0.9439651 | 0.34518748 | 0.99310104 |
| miR-200b-5p | 1.6344838 | -1.6953312 | 1.80643472 | -0.9384957 | 0.34798972 | 0.99310104 |
| miR-365-3p | 9.67637868 | -1.0168668 | 1.09295274 | -0.930385 | 0.35217181 | 0.99310104 |
| miR-3084-3p | 3.69419631 | 1.50633904 | 1.62094635 | 0.92929605 | 0.35273568 | 0.99310104 |
| miR-410-5p | 0.2535448 | 4.11443407 | 4.46317093 | 0.92186343 | 0.35659982 | 0.99310104 |
| miR-6942-5p | 0.2535448 | 4.11443407 | 4.46317093 | 0.92186343 | 0.35659982 | 0.99310104 |
| let-7b-5p | 480.909708 | 0.4121744 | 0.45019354 | 0.91554935 | 0.35990331 | 0.99310104 |
| miR-15a-5p | 15.6104589 | 0.78858231 | 0.86558365 | 0.91104113 | 0.3622737 | 0.99310104 |
| miR-192-5p | 525.686551 | -0.3662934 | 0.40309528 | -0.9087017 | 0.36350762 | 0.99310104 |
| miR-7214-3p | 14.3740518 | -0.7480009 | 0.82579714 | -0.9057926 | 0.36504566 | 0.99310104 |
| miR-210-3p | 1.31819247 | 2.23352611 | 2.50202097 | 0.89268881 | 0.37202385 | 0.99310104 |
| miR-3062-5p | 1.1157459 | -2.2129028 | 2.49588129 | -0.8866218 | 0.37528254 | 0.99310104 |
| miR-96-5p | 7.48396364 | 1.16654701 | 1.31686207 | 0.8858536 | 0.37569642 | 0.99310104 |
| miR-451a | 17.1113885 | 0.88317186 | 1.01498877 | 0.87012968 | 0.38422954 | 0.99310104 |

| | | | | | | |
|---|---|---|---|---|---|---|
| miR-7221-5p | 0.75652567 | -3.0165179 | 3.46788961 | -0.8698426 | 0.38438645 | 0.99310104 |
| miR-32-3p | 0.56166506 | 3.26269771 | 3.75403556 | 0.86911742 | 0.38478291 | 0.99310104 |
| miR-465b-3p | 162.106103 | -0.3939733 | 0.45418779 | -0.8674238 | 0.38570987 | 0.99310104 |
| miR-191-5p | 8236.5581 | -0.2742712 | 0.31761938 | -0.8635215 | 0.38785079 | 0.99310104 |
| miR-6915-3p | 0.21690048 | 3.83654941 | 4.47900797 | 0.85656231 | 0.39168682 | 0.99310104 |
| miR-323-3p | 0.21690048 | 3.83654941 | 4.47900797 | 0.85656231 | 0.39168682 | 0.99310104 |
| miR-7117-3p | 0.21690048 | 3.83654941 | 4.47900797 | 0.85656231 | 0.39168682 | 0.99310104 |
| miR-3085-5p | 0.21690048 | 3.83654941 | 4.47900797 | 0.85656231 | 0.39168682 | 0.99310104 |
| miR-199b-3p | 7.49644541 | 0.70163693 | 0.81915134 | 0.85654127 | 0.39169846 | 0.99310104 |
| miR-342-3p | 86.1737741 | 0.31447502 | 0.36714952 | 0.8565312 | 0.39170402 | 0.99310104 |
| miR-3535 | 2527.66528 | 0.45828595 | 0.53616275 | 0.85475157 | 0.3926887 | 0.99310104 |
| miR-465c-3p | 161.520091 | -0.3874134 | 0.45424971 | -0.8528643 | 0.39373454 | 0.99310104 |
| miR-224-5p | 4.18875103 | -0.9178156 | 1.08547881 | -0.8455399 | 0.39780946 | 0.99310104 |
| miR-375-3p | 1526.53454 | -0.3027582 | 0.35860308 | -0.8442711 | 0.3985179 | 0.99310104 |
| miR-143-3p | 18524.9756 | 0.43130092 | 0.51140418 | 0.84336605 | 0.39902376 | 0.99310104 |
| miR-671-5p | 1.29064859 | -2.3242151 | 2.76655486 | -0.8401117 | 0.40084577 | 0.99310104 |
| miR-1249-3p | 7.14171159 | 0.80149858 | 0.96065833 | 0.83432221 | 0.40409944 | 0.99310104 |
| miR-7234-5p | 0.84396123 | -2.735975 | 3.28415123 | -0.8330844 | 0.40479715 | 0.99310104 |
| miR-330-5p | 1.08453659 | -2.620625 | 3.15125384 | -0.8316134 | 0.40562719 | 0.99310104 |
| miR-193b-3p | 5.28269816 | 0.9788816 | 1.1810492 | 0.82882373 | 0.40720416 | 0.99310104 |
| miR-106b-3p | 71.543458 | -0.3333622 | 0.40327358 | -0.8266404 | 0.40844089 | 0.99310104 |
| miR-7a-2-3p | 0.43155262 | 3.70555632 | 4.4875688 | 0.82573805 | 0.4089527 | 0.99310104 |
| miR-8103 | 0.1901586 | 3.70555191 | 4.48756916 | 0.825737 | 0.4089533 | 0.99310104 |
| miR-539-5p | 0.1901586 | 3.70555191 | 4.48756916 | 0.825737 | 0.4089533 | 0.99310104 |
| miR-200c-5p | 0.1901586 | 3.70555191 | 4.48756916 | 0.825737 | 0.4089533 | 0.99310104 |
| miR-144-3p | 0.3307791 | 3.70554553 | 4.48756916 | 0.82573558 | 0.4089541 | 0.99310104 |
| miR-935 | 0.27163409 | 3.70553114 | 4.4875697 | 0.82573227 | 0.40895598 | 0.99310104 |
| miR-7242-3p | 19.1694289 | 0.7047585 | 0.85455005 | 0.82471296 | 0.40953457 | 0.99310104 |
| miR-99a-3p | 0.84195733 | -2.5412364 | 3.08699769 | -0.8232064 | 0.4103906 | 0.99310104 |
| miR-6516-3p | 1.25306265 | 2.30116073 | 2.80643952 | 0.81995736 | 0.41224042 | 0.99310104 |
| miR-7215-3p | 0.89167789 | 2.43049788 | 2.98019414 | 0.81555018 | 0.41475745 | 0.99310104 |
| miR-7233-3p | 86.2479996 | 0.31529783 | 0.38869652 | 0.81116711 | 0.41726971 | 0.99310104 |
| miR-183-5p | 637.348891 | 0.3319405 | 0.40941885 | 0.81076018 | 0.41750341 | 0.99310104 |
| miR-21a-3p | 18.0654067 | -0.6797343 | 0.84068405 | -0.8085491 | 0.41877457 | 0.99310104 |
| miR-1930-5p | 0.38185085 | 3.62433698 | 4.49326793 | 0.80661493 | 0.41988837 | 0.99310104 |
| miR-30d-3p | 4.34013111 | -1.061937 | 1.32805295 | -0.7996194 | 0.42393132 | 0.99310104 |
| miR-151-3p | 493.324752 | 0.22213459 | 0.27988267 | 0.7936704 | 0.42738734 | 0.99310104 |
| miR-466b-3p | 3.80569996 | -0.96313 | 1.21999892 | -0.7894515 | 0.42984817 | 0.99310104 |
| miR-465a-3p | 81.2636666 | -0.3759607 | 0.47668961 | -0.7886908 | 0.43029272 | 0.99310104 |
| miR-339-5p | 2.66677989 | -1.4098634 | 1.79349488 | -0.7860984 | 0.43180986 | 0.99310104 |
| miR-20a-3p | 0.81996891 | 2.87667664 | 3.6594603 | 0.78609314 | 0.43181292 | 0.99310104 |
| miR-423-3p | 47.2313136 | -0.3895881 | 0.49589764 | -0.785622 | 0.432089 | 0.99310104 |
| miR-128-3p | 148.185578 | -0.2923289 | 0.37328199 | -0.7831316 | 0.43354983 | 0.99310104 |
| miR-1306-5p | 11.0575603 | 0.72715271 | 0.93183 | 0.7803491 | 0.43518542 | 0.99310104 |
| miR-582-3p | 0.6706355 | 3.10623712 | 3.98971416 | 0.77856132 | 0.43623817 | 0.99310104 |
| miR-883b-3p | 3.94986695 | -0.8752717 | 1.12802379 | -0.7759337 | 0.43778813 | 0.99310104 |

| | | | | | | |
|---|---|---|---|---|---|---|
| miR-7242-5p | 6.32206472 | 0.95662838 | 1.23684656 | 0.77344143 | 0.43926118 | 0.99310104 |
| let-7i-3p | 1.70454211 | 1.89700724 | 2.45519 | 0.77265191 | 0.43972842 | 0.99310104 |
| miR-99b-5p | 638.306367 | 0.35074344 | 0.45467163 | 0.77142143 | 0.44045717 | 0.99310104 |
| miR-5099 | 84.7086433 | 0.57833117 | 0.75148586 | 0.76958357 | 0.44154695 | 0.99310104 |
| miR-465b-5p | 134.218272 | -0.4395903 | 0.57185016 | -0.7687159 | 0.44206198 | 0.99310104 |
| miR-664-5p | 2.60168405 | 1.08141354 | 1.41421138 | 0.76467603 | 0.44446449 | 0.99310104 |
| miR-466d-3p | 0.63912589 | 2.36842419 | 3.10274662 | 0.76333149 | 0.44526573 | 0.99310104 |
| miR-7060-5p | 1.17256607 | -1.9040677 | 2.50600008 | -0.7598035 | 0.44737202 | 0.99310104 |
| miR-142a-5p | 317.260067 | 0.23805776 | 0.31600323 | 0.75333963 | 0.45124585 | 0.99310104 |
| miR-134-5p | 1.98854484 | 1.4934974 | 1.98257277 | 0.75331278 | 0.45126198 | 0.99310104 |
| miR-471-5p | 12.6499637 | 0.76200922 | 1.01521082 | 0.7505921 | 0.45289818 | 0.99310104 |
| miR-195a-3p | 1.16652442 | -2.1651056 | 2.8958968 | -0.747646 | 0.45467372 | 0.99310104 |
| miR-381-3p | 1.4691769 | -1.3931608 | 1.86782542 | -0.7458731 | 0.45574407 | 0.99310104 |
| miR-344-3p | 2.01375743 | -3.3095253 | 4.43913289 | -0.7455342 | 0.45594886 | 0.99310104 |
| miR-17-5p | 2.20817842 | -1.1666505 | 1.57156909 | -0.7423476 | 0.45787677 | 0.99310104 |
| miR-20a-5p | 9.68228301 | 0.664869 | 0.89762441 | 0.74069843 | 0.45887631 | 0.99310104 |
| miR-124-5p | 0.55117314 | 2.50256438 | 3.45980803 | 0.72332463 | 0.46948046 | 0.99310104 |
| miR-1188-3p | 1.17284112 | -2.0924642 | 2.89899604 | -0.7217893 | 0.47042405 | 0.99310104 |
| miR-7a-5p | 95.5240462 | -0.3620567 | 0.50671568 | -0.7145164 | 0.4749079 | 0.99310104 |
| miR-3068-3p | 13.5648049 | -0.5452263 | 0.76324311 | -0.7143547 | 0.47500786 | 0.99310104 |
| miR-677-3p | 3.47500083 | -1.098831 | 1.54598836 | -0.7107628 | 0.47723123 | 0.99310104 |
| miR-7226-3p | 0.1267724 | 3.20416227 | 4.51909666 | 0.70902716 | 0.47830762 | 0.99310104 |
| miR-140-5p | 1.3537423 | 1.7356015 | 2.45314246 | 0.70750131 | 0.479255 | 0.99310104 |
| miR-132-3p | 1.31694205 | -1.6081228 | 2.28430189 | -0.7039888 | 0.48143978 | 0.99310104 |
| miR-34a-5p | 3.37585197 | 1.16400337 | 1.65819988 | 0.70196807 | 0.48269908 | 0.99310104 |
| miR-31-3p | 1.39114034 | 1.76382803 | 2.51517241 | 0.7012752 | 0.48313129 | 0.99310104 |
| miR-499-5p | 0.9508207 | -2.0789042 | 2.9665904 | -0.7007723 | 0.48344515 | 0.99310104 |
| miR-100-5p | 91.5998735 | 0.33116889 | 0.47278219 | 0.7004682 | 0.48363496 | 0.99310104 |
| miR-1191b-5p | 0.35921461 | 3.11510747 | 4.47899991 | 0.69549175 | 0.48674718 | 0.99310104 |
| miR-411-5p | 2.37450332 | -1.1940741 | 1.72740492 | -0.6912532 | 0.48940646 | 0.99310104 |
| miR-29c-3p | 0.92929929 | -2.3350135 | 3.38130697 | -0.6905654 | 0.48983871 | 0.99310104 |
| miR-29a-3p | 39.0011119 | -0.4905474 | 0.71065623 | -0.6902738 | 0.490022 | 0.99310104 |
| miR-122-5p | 0.85811582 | -2.5705312 | 3.73407236 | -0.6883989 | 0.49120163 | 0.99310104 |
| miR-874-3p | 1.12678457 | 1.46634132 | 2.15288694 | 0.68110466 | 0.49580527 | 0.99310104 |
| miR-497a-5p | 2.74225735 | -1.3642794 | 2.00382837 | -0.6808365 | 0.49597497 | 0.99310104 |
| miR-455-3p | 0.56760543 | -2.7908648 | 4.10430319 | -0.6799851 | 0.49651392 | 0.99310104 |
| miR-128-1-5p | 0.81916492 | -2.319629 | 3.41736469 | -0.6787771 | 0.49727908 | 0.99310104 |
| miR-369-3p | 0.54660629 | -2.5379722 | 3.78520919 | -0.6704972 | 0.50254088 | 0.99310104 |
| miR-468-3p | 6.10232842 | 0.70208616 | 1.05289148 | 0.66681721 | 0.5048889 | 0.99310104 |
| miR-7222-5p | 3.5716854 | 0.84602295 | 1.27366194 | 0.66424451 | 0.50653383 | 0.99310104 |
| miR-7230-5p | 12.6123363 | 0.39316701 | 0.59414043 | 0.66174087 | 0.5081373 | 0.99310104 |
| let-7a-5p | 12110.1372 | -0.2050731 | 0.31105616 | -0.6592801 | 0.50971592 | 0.99310104 |
| miR-125a-5p | 566.450555 | -0.329662 | 0.50066037 | -0.6584544 | 0.51024618 | 0.99310104 |
| miR-101a-3p | 23.1599465 | 0.42549098 | 0.65792391 | 0.64671761 | 0.51781472 | 0.99310104 |
| miR-8114 | 4.63926344 | -0.9588129 | 1.4871906 | -0.6447142 | 0.5191124 | 0.99310104 |
| miR-127-3p | 133.600536 | -0.31578 | 0.49872286 | -0.6331772 | 0.52661791 | 0.99310104 |

| | | | | | | |
|---|---|---|---|---|---|---|
| miR-19a-3p | 11.4926159 | -0.6417691 | 1.0162056 | -0.6315347 | 0.52769097 | 0.99310104 |
| miR-34b-3p | 1915.43036 | 0.24097526 | 0.38191002 | 0.6309739 | 0.52805759 | 0.99310104 |
| miR-93-3p | 0.72809689 | 2.19143348 | 3.48133005 | 0.62948168 | 0.52903376 | 0.99310104 |
| miR-871-5p | 716.00589 | -0.2352527 | 0.37505812 | -0.6272433 | 0.53049979 | 0.99310104 |
| miR-200a-5p | 1.2069913 | 1.82829944 | 2.95695705 | 0.61830436 | 0.53637473 | 0.99310104 |
| miR-187-5p | 2.16057964 | 1.58796072 | 2.58443558 | 0.61443231 | 0.53892969 | 0.99310104 |
| miR-294-3p | 0.92402984 | -1.8745335 | 3.05624125 | -0.6133461 | 0.53964755 | 0.99310104 |
| miR-672-3p | 0.69653624 | -2.403278 | 3.92256894 | -0.6126796 | 0.54008821 | 0.99310104 |
| miR-598-3p | 0.51310928 | -2.5840396 | 4.23344052 | -0.6103876 | 0.54160509 | 0.99310104 |
| miR-7229-5p | 78.3795612 | 0.26341224 | 0.43431644 | 0.60649843 | 0.54418383 | 0.99310104 |
| miR-181c-5p | 168.593623 | -0.2775202 | 0.46087711 | -0.6021566 | 0.54706987 | 0.99310104 |
| miR-500-3p | 0.52674753 | 2.54959364 | 4.25386799 | 0.5993589 | 0.54893358 | 0.99310104 |
| miR-99a-5p | 82.9188033 | 0.32163117 | 0.53741869 | 0.59847411 | 0.54952363 | 0.99310104 |
| miR-674-5p | 0.79662291 | -2.0175141 | 3.39418093 | -0.5944038 | 0.55224205 | 0.99310104 |
| miR-668-3p | 0.98864856 | 1.72141644 | 2.90410131 | 0.59275358 | 0.55334608 | 0.99310104 |
| miR-615-3p | 2.63498668 | -0.7775111 | 1.31597624 | -0.5908246 | 0.55463796 | 0.99310104 |
| miR-677-5p | 22.6023362 | -0.4810206 | 0.81443033 | -0.5906221 | 0.55477365 | 0.99310104 |
| miR-483-3p | 3.44937194 | -0.901721 | 1.53318848 | -0.5881345 | 0.55644205 | 0.99310104 |
| miR-181d-5p | 11.1097067 | -0.460101 | 0.78287684 | -0.5877054 | 0.55673002 | 0.99310104 |
| miR-1932 | 0.35891615 | -2.6073107 | 4.44670972 | -0.5863461 | 0.55764299 | 0.99310104 |
| miR-3572-5p | 0.35891615 | -2.6073107 | 4.44670972 | -0.5863461 | 0.55764299 | 0.99310104 |
| miR-135a-1-3p | 0.35891615 | -2.6073107 | 4.44670972 | -0.5863461 | 0.55764299 | 0.99310104 |
| miR-666-3p | 1.05680737 | -1.7566727 | 3.01763947 | -0.5821347 | 0.56047594 | 0.99310104 |
| miR-34c-5p | 1365.01758 | -0.3297719 | 0.57123165 | -0.5772997 | 0.56373701 | 0.99310104 |
| miR-532-5p | 35.9742044 | -0.2699123 | 0.47388138 | -0.5695778 | 0.56896412 | 0.99310104 |
| miR-301b-3p | 0.78326378 | -1.9424346 | 3.4155134 | -0.5687094 | 0.56955335 | 0.99310104 |
| miR-342-5p | 6.79951387 | -0.6737613 | 1.18666996 | -0.5677748 | 0.57018788 | 0.99310104 |
| miR-8111 | 0.53598151 | 2.5496707 | 4.54293138 | 0.5612391 | 0.57463455 | 0.99310104 |
| miR-6541 | 0.29020579 | 2.54964803 | 4.54293138 | 0.56123411 | 0.57463796 | 0.99310104 |
| miR-6971-3p | 0.21444515 | 2.54963244 | 4.54293138 | 0.56123068 | 0.57464029 | 0.99310104 |
| miR-293-3p | 0.25461773 | 2.54961118 | 4.54293138 | 0.56122599 | 0.57464349 | 0.99310104 |
| miR-182-3p | 0.24206941 | 2.54957736 | 4.54293138 | 0.56121855 | 0.57464856 | 0.99310104 |
| miR-320-3p | 35.3779332 | 0.33787803 | 0.60237204 | 0.56091255 | 0.57485716 | 0.99310104 |
| miR-7229-3p | 9.20503501 | 0.45786124 | 0.82205831 | 0.55696929 | 0.57754841 | 0.99310104 |
| miR-7685-5p | 0.32628741 | -2.4712391 | 4.44864582 | -0.5555037 | 0.5785502 | 0.99310104 |
| miR-379-5p | 4.0139086 | -0.7475825 | 1.34986333 | -0.5538209 | 0.57970145 | 0.99310104 |
| miR-200c-3p | 43.9662858 | -0.2641535 | 0.48003642 | -0.550278 | 0.58212872 | 0.99310104 |
| miR-669p-5p | 2.94739795 | -0.6310174 | 1.17755709 | -0.5358699 | 0.59204845 | 0.99310104 |
| let-7f-5p | 17411.4308 | -0.2219603 | 0.41755249 | -0.5315747 | 0.5950206 | 0.99310104 |
| miR-666-5p | 1.02168433 | -1.4977795 | 2.82164427 | -0.530818 | 0.59554491 | 0.99310104 |
| miR-449c-5p | 1.29789064 | -1.1421567 | 2.16241101 | -0.5281867 | 0.59736976 | 0.99310104 |
| miR-378d | 0.71288438 | -1.6257686 | 3.08367355 | -0.5272181 | 0.59804212 | 0.99310104 |
| miR-7658-3p | 0.65540536 | -2.3022868 | 4.41213121 | -0.5218083 | 0.60180379 | 0.99310104 |
| miR-700-5p | 0.74366835 | -1.6939815 | 3.24725092 | -0.5216663 | 0.60190267 | 0.99310104 |
| miR-532-3p | 4.07129501 | 0.70617948 | 1.35937608 | 0.51948794 | 0.60342052 | 0.99310104 |
| miR-3082-3p | 0.48479468 | -2.0376699 | 3.92572953 | -0.5190551 | 0.60372232 | 0.99310104 |

| | | | | | | |
|---|---|---|---|---|---|---|
| miR-1981-5p | 0.81363115 | 1.78178637 | 3.44467856 | 0.51725766 | 0.60497631 | 0.99310104 |
| miR-181a-1-3p | 1.32505335 | -1.3908587 | 2.70133531 | -0.5148782 | 0.60663812 | 0.99310104 |
| miR-190b-5p | 22.2117223 | 0.32971413 | 0.64058602 | 0.51470704 | 0.60675775 | 0.99310104 |
| miR-215-5p | 0.69339238 | -1.6932811 | 3.29486936 | -0.5139145 | 0.60731178 | 0.99310104 |
| miR-301a-3p | 10.7435234 | -0.5017122 | 0.98068416 | -0.5115941 | 0.60893514 | 0.99310104 |
| miR-322-3p | 0.64858961 | -1.6505379 | 3.22861148 | -0.5112222 | 0.60919547 | 0.99310104 |
| miR-682 | 0.8790814 | 1.58791661 | 3.1276634 | 0.50770061 | 0.61166332 | 0.99310104 |
| miR-184-5p | 0.73658487 | 1.82824391 | 3.60680442 | 0.50688745 | 0.61223379 | 0.99310104 |
| miR-345-5p | 2.83472097 | -0.7409182 | 1.46590251 | -0.5054348 | 0.61325345 | 0.99310104 |
| miR-3064-3p | 0.2773443 | -2.2395098 | 4.45238426 | -0.5029911 | 0.6149705 | 0.99310104 |
| miR-7648-3p | 1.1435915 | 1.5879059 | 3.15753651 | 0.50289391 | 0.61503886 | 0.99310104 |
| miR-27a-5p | 0.67508012 | 1.82832242 | 3.6405679 | 0.50220803 | 0.6155212 | 0.99310104 |
| miR-1291 | 0.31862991 | 2.20216327 | 4.42373015 | 0.49780687 | 0.61862017 | 0.99310104 |
| miR-98-3p | 0.65380206 | 1.77734564 | 3.59984927 | 0.49372779 | 0.62149843 | 0.99310104 |
| miR-150-3p | 0.64869789 | -1.9731949 | 4.03451466 | -0.4890786 | 0.62478604 | 0.99310104 |
| miR-196b-3p | 1.24694237 | 1.58793213 | 3.2708207 | 0.48548431 | 0.62733284 | 0.99310104 |
| miR-199b-5p | 0.26929608 | -2.1536892 | 4.45392478 | -0.4835486 | 0.62870625 | 0.99310104 |
| miR-7009-5p | 0.26102993 | -2.1531657 | 4.45393445 | -0.48343 | 0.62879043 | 0.99310104 |
| miR-466c-3p | 1.50123109 | -0.8190689 | 1.69428838 | -0.4834295 | 0.62879082 | 0.99310104 |
| miR-5121 | 6.97170829 | -0.5779126 | 1.20458173 | -0.479762 | 0.6313966 | 0.99310104 |
| miR-6960-5p | 0.67675021 | 1.69020067 | 3.52429553 | 0.4795854 | 0.63152223 | 0.99310104 |
| miR-125b-1-3p | 6.8504028 | -0.5586389 | 1.16844222 | -0.4781057 | 0.63257497 | 0.99310104 |
| miR-200b-3p | 43.7631029 | -0.2570832 | 0.54166427 | -0.4746173 | 0.6350598 | 0.99310104 |
| miR-21a-5p | 686.905345 | 0.20571179 | 0.43382151 | 0.47418533 | 0.63536775 | 0.99310104 |
| miR-872-5p | 34.3884674 | 0.31429387 | 0.66388755 | 0.47341432 | 0.63591761 | 0.99310104 |
| miR-1906 | 1.02708116 | 1.58795426 | 3.35818741 | 0.47286052 | 0.63631269 | 0.99310104 |
| miR-141-5p | 1.32835685 | -1.4758721 | 3.1218288 | -0.4727588 | 0.63638526 | 0.99310104 |
| miR-7a-1-3p | 1.40768575 | 0.92585189 | 1.96161769 | 0.47198386 | 0.63693831 | 0.99310104 |
| miR-32-5p | 2.22823869 | -0.8678184 | 1.862635 | -0.465909 | 0.64128063 | 0.99310104 |
| miR-362-5p | 0.24471556 | -2.0613328 | 4.45568566 | -0.4626298 | 0.64362976 | 0.99310104 |
| miR-127-5p | 0.54338521 | -1.9214539 | 4.16691637 | -0.4611213 | 0.64471159 | 0.99310104 |
| miR-615-5p | 4.90884478 | -0.7408308 | 1.6080469 | -0.4607022 | 0.64501226 | 0.99310104 |
| miR-362-3p | 0.23916252 | -2.0271973 | 4.45636482 | -0.4548993 | 0.64918168 | 0.99310104 |
| miR-703 | 0.59496855 | -1.650625 | 3.66351486 | -0.4505577 | 0.65230833 | 0.99310104 |
| miR-152-3p | 3.69076093 | 0.6974867 | 1.55222 | 0.44934784 | 0.65318075 | 0.99310104 |
| miR-149-5p | 2.78691366 | -0.851633 | 1.9020524 | -0.4477442 | 0.6543378 | 0.99310104 |
| miR-30b-3p | 2.45497289 | 0.61184247 | 1.40893898 | 0.43425761 | 0.66410138 | 0.99310104 |
| miR-540-5p | 0.46396931 | 1.92406023 | 4.43097161 | 0.43422987 | 0.66412152 | 0.99310104 |
| miR-329-5p | 0.23145904 | -1.9261704 | 4.45846904 | -0.4320251 | 0.6657232 | 0.99310104 |
| miR-671-3p | 18.2438899 | -0.2745651 | 0.63958227 | -0.4292882 | 0.66771349 | 0.99310104 |
| miR-143-5p | 1.38733169 | -0.9890296 | 2.31495371 | -0.4272351 | 0.66920807 | 0.99310104 |
| miR-467e-5p | 4.54618643 | 0.59448319 | 1.39614869 | 0.42580221 | 0.67025198 | 0.99310104 |
| miR-3077-3p | 0.2189207 | -1.8895721 | 4.4592674 | -0.4237405 | 0.67175509 | 0.99310104 |
| miR-7241-5p | 0.58243179 | -1.8895257 | 4.45926781 | -0.42373 | 0.6717627 | 0.99310104 |
| miR-5114 | 11.0690558 | -0.4653243 | 1.09961443 | -0.4231704 | 0.67217093 | 0.99310104 |
| miR-190a-5p | 0.225906 | -1.886958 | 4.45932518 | -0.4231488 | 0.67218671 | 0.99310104 |

| | | | | | | |
|---|---|---|---|---|---|---|
| miR-744-3p | 0.37696495 | -1.8869435 | 4.45932531 | -0.4231455 | 0.67218909 | 0.99310104 |
| miR-483-5p | 1.26550568 | -1.0953578 | 2.59698642 | -0.4217803 | 0.67318535 | 0.99310104 |
| miR-3057-5p | 3.32793103 | 0.60843386 | 1.44684298 | 0.42052515 | 0.67410186 | 0.99310104 |
| miR-30e-5p | 167.016417 | -0.4506432 | 1.07240616 | -0.4202169 | 0.67432698 | 0.99310104 |
| miR-5100 | 0.5482102 | -1.6003093 | 3.81822232 | -0.4191242 | 0.67512537 | 0.99310104 |
| miR-1306-3p | 0.35422355 | -1.8580069 | 4.45997233 | -0.4165961 | 0.67697389 | 0.99310104 |
| miR-409-5p | 0.29204608 | -1.8579532 | 4.45997297 | -0.416584 | 0.67698275 | 0.99310104 |
| miR-19b-3p | 42.3812045 | -0.234127 | 0.56478039 | -0.4145452 | 0.67847489 | 0.99310104 |
| miR-340-5p | 230.688377 | -0.2193437 | 0.5305617 | -0.4134179 | 0.67930043 | 0.99310104 |
| miR-3072-3p | 0.89318298 | -1.5204614 | 3.67812024 | -0.41338 | 0.67932821 | 0.99310104 |
| miR-505-5p | 0.83056638 | -1.3775547 | 3.35781821 | -0.4102529 | 0.68162042 | 0.99310104 |
| miR-669d-5p | 1.09656679 | -0.9449597 | 2.30772868 | -0.4094761 | 0.68219032 | 0.99310104 |
| miR-199a-5p | 7.36944671 | 0.49412878 | 1.20802302 | 0.40903921 | 0.68251089 | 0.99310104 |
| miR-8112 | 2.01044339 | -0.7735043 | 1.90802332 | -0.4053956 | 0.68518675 | 0.99310104 |
| miR-669m-3p | 0.20749561 | -1.801589 | 4.46126914 | -0.4038288 | 0.68633862 | 0.99310104 |
| miR-511-5p | 0.75405694 | -1.3946206 | 3.46168645 | -0.4028732 | 0.68704153 | 0.99310104 |
| miR-28c | 0.12476461 | 1.82836721 | 4.54292645 | 0.40246463 | 0.68734211 | 0.99310104 |
| miR-466c-5p | 0.16928888 | 1.8282266 | 4.54292366 | 0.40243392 | 0.68736471 | 0.99310104 |
| miR-3100-5p | 0.16928888 | 1.8282266 | 4.54292366 | 0.40243392 | 0.68736471 | 0.99310104 |
| miR-450b-5p | 0.13014528 | 1.8282265 | 4.54292366 | 0.4024339 | 0.68736472 | 0.99310104 |
| miR-743b-3p | 512.187236 | -0.1609056 | 0.40000366 | -0.4022602 | 0.68749252 | 0.99310104 |
| miR-1894-3p | 0.2031438 | -1.7854349 | 4.46164972 | -0.4001737 | 0.68902858 | 0.99310104 |
| miR-434-5p | 3.20697696 | 0.62843624 | 1.59128705 | 0.39492324 | 0.69289954 | 0.99310104 |
| miR-5107-3p | 0.17016248 | 1.75013105 | 4.49856529 | 0.38904205 | 0.69724504 | 0.99310104 |
| miR-1839-3p | 0.33311346 | 1.75008983 | 4.49856492 | 0.38903292 | 0.6972518 | 0.99310104 |
| miR-669m-5p | 0.3916325 | -1.7327246 | 4.4629209 | -0.388249 | 0.69783177 | 0.99310104 |
| miR-6988-5p | 0.19525535 | -1.7305913 | 4.46297309 | -0.3877665 | 0.69818886 | 0.99310104 |
| miR-124-3p | 2.25959209 | -0.6100834 | 1.58331603 | -0.3853201 | 0.7000003 | 0.99310104 |
| miR-673-5p | 0.41086014 | -1.584342 | 4.14975073 | -0.3817921 | 0.70261559 | 0.99310104 |
| miR-341-3p | 0.45586568 | 1.67269512 | 4.38372329 | 0.3815695 | 0.70278071 | 0.99310104 |
| miR-101b-3p | 13.5057654 | 0.28576349 | 0.75019958 | 0.38091662 | 0.70326512 | 0.99310104 |
| miR-872-3p | 3.0419223 | 0.57354266 | 1.51247281 | 0.37920858 | 0.70453298 | 0.99310104 |
| miR-743b-5p | 13.3933767 | -0.3050325 | 0.80453433 | -0.3791416 | 0.70458269 | 0.99310104 |
| miR-1964-3p | 0.68235136 | -1.494681 | 3.94271802 | -0.3790991 | 0.70461424 | 0.99310104 |
| miR-5113 | 0.69305464 | 1.58793355 | 4.22204595 | 0.37610523 | 0.70683867 | 0.99310104 |
| miR-1948-5p | 0.19465085 | -1.6784109 | 4.46427827 | -0.3759647 | 0.70694316 | 0.99310104 |
| miR-26b-5p | 124.394455 | -0.157069 | 0.42028707 | -0.3737184 | 0.70861387 | 0.99310104 |
| miR-1956 | 0.7075449 | 1.58783175 | 4.27822443 | 0.3711427 | 0.71053125 | 0.99310104 |
| miR-8118 | 0.68553564 | 1.58780589 | 4.29024021 | 0.3700972 | 0.71131007 | 0.99310104 |
| miR-664-3p | 13.0924885 | 0.27334956 | 0.74542166 | 0.36670461 | 0.71383937 | 0.99310104 |
| miR-204-3p | 0.66535337 | 1.58790841 | 4.34239765 | 0.36567549 | 0.71460723 | 0.99310104 |
| miR-7675-3p | 0.17945807 | -1.6210738 | 4.46576612 | -0.3630002 | 0.71660476 | 0.99310104 |
| miR-7224-3p | 0.1867624 | -1.6174545 | 4.46586198 | -0.3621819 | 0.71721609 | 0.99310104 |
| miR-223-3p | 2.14767617 | -0.6377846 | 1.76150956 | -0.3620671 | 0.71730192 | 0.99310104 |
| miR-760-3p | 0.23692155 | 1.60702032 | 4.46201153 | 0.36015602 | 0.71873046 | 0.99310104 |
| miR-1969 | 0.56552509 | -1.2448418 | 3.45705819 | -0.360087 | 0.71878206 | 0.99310104 |

| | | | | | | |
|---|---|---|---|---|---|---|
| miR-501-3p | 4.84182452 | -0.4364762 | 1.22604273 | -0.3560041 | 0.72183753 | 0.99310104 |
| miR-5617-3p | 0.60926336 | 1.58784643 | 4.49592061 | 0.35317493 | 0.7239573 | 0.99310104 |
| miR-1843b-5p | 13.0562397 | 0.27610598 | 0.78429801 | 0.35204217 | 0.72480664 | 0.99310104 |
| miR-1936 | 0.75529474 | 1.58792987 | 4.54293138 | 0.3495386 | 0.726685 | 0.99310104 |
| miR-7006-3p | 0.60423579 | 1.5879231 | 4.54293138 | 0.34953711 | 0.72668612 | 0.99310104 |
| miR-7017-3p | 0.54526664 | 1.58792082 | 4.54293138 | 0.34953661 | 0.72668649 | 0.99310104 |
| miR-5620-5p | 0.54526664 | 1.58792082 | 4.54293138 | 0.34953661 | 0.72668649 | 0.99310104 |
| miR-7652-3p | 0.48387344 | 1.5879167 | 4.54293138 | 0.3495357 | 0.72668717 | 0.99310104 |
| miR-433-3p | 0.45438887 | 1.58791542 | 4.54293138 | 0.34953542 | 0.72668739 | 0.99310104 |
| miR-210-5p | 0.43827431 | 1.5879144 | 4.54293138 | 0.34953519 | 0.72668755 | 0.99310104 |
| miR-7238-5p | 0.45317684 | 1.58791433 | 4.54293138 | 0.34953518 | 0.72668756 | 0.99310104 |
| miR-328-5p | 0.45317684 | 1.58791433 | 4.54293138 | 0.34953518 | 0.72668756 | 0.99310104 |
| miR-6990-5p | 0.42753127 | 1.58791371 | 4.54293138 | 0.34953504 | 0.72668767 | 0.99310104 |
| miR-183-3p | 0.42753127 | 1.58791371 | 4.54293138 | 0.34953504 | 0.72668767 | 0.99310104 |
| miR-3475-3p | 0.42753127 | 1.58791371 | 4.54293138 | 0.34953504 | 0.72668767 | 0.99310104 |
| miR-5125 | 0.39979633 | 1.58791183 | 4.54293138 | 0.34953463 | 0.72668798 | 0.99310104 |
| miR-7070-5p | 0.39979633 | 1.58791183 | 4.54293138 | 0.34953463 | 0.72668798 | 0.99310104 |
| miR-7073-3p | 0.39979633 | 1.58791183 | 4.54293138 | 0.34953463 | 0.72668798 | 0.99310104 |
| miR-6922-3p | 0.36351109 | 1.58790881 | 4.54293138 | 0.34953396 | 0.72668848 | 0.99310104 |
| miR-6340 | 0.36351109 | 1.58790881 | 4.54293138 | 0.34953396 | 0.72668848 | 0.99310104 |
| miR-3092-3p | 0.36351109 | 1.58790881 | 4.54293138 | 0.34953396 | 0.72668848 | 0.99310104 |
| miR-455-5p | 0.35813958 | 1.5879084 | 4.54293138 | 0.34953387 | 0.72668855 | 0.99310104 |
| miR-218-1-3p | 0.31983706 | 1.58790527 | 4.54293138 | 0.34953319 | 0.72668906 | 0.99310104 |
| miR-3103-5p | 0.27263332 | 1.5879008 | 4.54293138 | 0.3495322 | 0.7266898 | 0.99310104 |
| miR-504-5p | 0.27263332 | 1.5879008 | 4.54293138 | 0.3495322 | 0.7266898 | 0.99310104 |
| miR-7090-5p | 0.25651876 | 1.58789944 | 4.54293138 | 0.3495319 | 0.72669003 | 0.99310104 |
| miR-196a-1-3p | 0.3021179 | 1.5878992 | 4.54293138 | 0.34953185 | 0.72669007 | 0.99310104 |
| miR-673-3p | 0.3021179 | 1.5878992 | 4.54293138 | 0.34953185 | 0.72669007 | 0.99310104 |
| miR-719 | 0.3021179 | 1.5878992 | 4.54293138 | 0.34953185 | 0.72669007 | 0.99310104 |
| miR-546 | 0.2398778 | 1.58789795 | 4.54293138 | 0.34953157 | 0.72669027 | 0.99310104 |
| miR-134-3p | 0.2398778 | 1.58789795 | 4.54293138 | 0.34953157 | 0.72669027 | 0.99310104 |
| miR-412-5p | 0.2398778 | 1.58789795 | 4.54293138 | 0.34953157 | 0.72669027 | 0.99310104 |
| miR-6992-5p | 0.2398778 | 1.58789795 | 4.54293138 | 0.34953157 | 0.72669027 | 0.99310104 |
| miR-1960 | 0.2398778 | 1.58789795 | 4.54293138 | 0.34953157 | 0.72669027 | 0.99310104 |
| miR-7028-5p | 0.2398778 | 1.58789795 | 4.54293138 | 0.34953157 | 0.72669027 | 0.99310104 |
| miR-330-3p | 0.2398778 | 1.58789795 | 4.54293138 | 0.34953157 | 0.72669027 | 0.99310104 |
| miR-138-5p | 0.2398778 | 1.58789795 | 4.54293138 | 0.34953157 | 0.72669027 | 0.99310104 |
| miR-3109-3p | 0.2398778 | 1.58789795 | 4.54293138 | 0.34953157 | 0.72669027 | 0.99310104 |
| miR-7075-5p | 0.18175555 | 1.58789104 | 4.54293138 | 0.34953005 | 0.72669141 | 0.99310104 |
| miR-201-5p | 0.18175555 | 1.58789104 | 4.54293138 | 0.34953005 | 0.72669141 | 0.99310104 |
| miR-5119 | 0.17101251 | 1.58788975 | 4.54293138 | 0.34952977 | 0.72669163 | 0.99310104 |
| miR-7648-5p | 0.17101251 | 1.58788975 | 4.54293138 | 0.34952977 | 0.72669163 | 0.99310104 |
| miR-877-3p | 0.17101251 | 1.58788975 | 4.54293138 | 0.34952977 | 0.72669163 | 0.99310104 |
| miR-193a-5p | 0.15991853 | 1.58788834 | 4.54293138 | 0.34952946 | 0.72669186 | 0.99310104 |
| miR-665-3p | 0.15991853 | 1.58788834 | 4.54293138 | 0.34952946 | 0.72669186 | 0.99310104 |
| miR-874-5p | 0.15991853 | 1.58788834 | 4.54293138 | 0.34952946 | 0.72669186 | 0.99310104 |

| | | | | | | |
|---|---|---|---|---|---|---|
| miR-3090-3p | 0.15991853 | 1.58788834 | 4.54293138 | 0.34952946 | 0.72669186 | 0.99310104 |
| miR-7007-3p | 0.15991853 | 1.58788834 | 4.54293138 | 0.34952946 | 0.72669186 | 0.99310104 |
| miR-25-5p | 0.55995929 | 1.58788754 | 4.54293138 | 0.34952928 | 0.72669199 | 0.99310104 |
| miR-692 | 0.24139402 | 1.58786707 | 4.54293138 | 0.34952478 | 0.72669537 | 0.99310104 |
| miR-378c | 0.15231584 | 1.58785418 | 4.54293138 | 0.34952194 | 0.7266975 | 0.99310104 |
| miR-130a-5p | 0.23253444 | 1.58785418 | 4.54293138 | 0.34952194 | 0.7266975 | 0.99310104 |
| miR-7010-5p | 0.23253444 | 1.58785418 | 4.54293138 | 0.34952194 | 0.7266975 | 0.99310104 |
| miR-1933-3p | 0.38353075 | 1.58783135 | 4.54293138 | 0.34951691 | 0.72670128 | 0.99310104 |
| miR-324-3p | 0.12132248 | 1.58782301 | 4.54293138 | 0.34951508 | 0.72670265 | 0.99310104 |
| miR-3473e | 0.12132248 | 1.58782301 | 4.54293138 | 0.34951508 | 0.72670265 | 0.99310104 |
| miR-7026-3p | 0.12132248 | 1.58782301 | 4.54293138 | 0.34951508 | 0.72670265 | 0.99310104 |
| miR-1668 | 0.49396646 | 1.58781736 | 4.54293138 | 0.34951383 | 0.72670359 | 0.99310104 |
| miR-7662-3p | 0.16295098 | 1.58781446 | 4.54293138 | 0.34951319 | 0.72670407 | 0.99310104 |
| miR-129-5p | 0.16295098 | 1.58781446 | 4.54293138 | 0.34951319 | 0.72670407 | 0.99310104 |
| miR-7012-5p | 0.16295098 | 1.58781446 | 4.54293138 | 0.34951319 | 0.72670407 | 0.99310104 |
| miR-7042-5p | 0.57187003 | 1.58781217 | 4.54293138 | 0.34951269 | 0.72670444 | 0.99310104 |
| miR-7663-5p | 0.18198372 | 1.58780996 | 4.54293138 | 0.3495122 | 0.72670481 | 0.99310104 |
| miR-3960 | 0.18198372 | 1.58780996 | 4.54293138 | 0.3495122 | 0.72670481 | 0.99310104 |
| miR-7036b-3p | 0.18198372 | 1.58780996 | 4.54293138 | 0.3495122 | 0.72670481 | 0.99310104 |
| miR-376b-3p | 0.20042876 | 1.58780697 | 4.54293138 | 0.34951155 | 0.7267053 | 0.99310104 |
| miR-7663-3p | 0.24442647 | 1.58780073 | 4.54293138 | 0.34951017 | 0.72670634 | 0.99310104 |
| miR-7657-5p | 0.24442647 | 1.58780073 | 4.54293138 | 0.34951017 | 0.72670634 | 0.99310104 |
| miR-363-3p | 0.24442647 | 1.58780073 | 4.54293138 | 0.34951017 | 0.72670634 | 0.99310104 |
| miR-542-5p | 0.26109 | 1.58779799 | 4.54293138 | 0.34950957 | 0.72670679 | 0.99310104 |
| miR-7653-3p | 0.26723835 | 1.58779724 | 4.54293138 | 0.3495094 | 0.72670691 | 0.99310104 |
| miR-7670-3p | 0.26723835 | 1.58779724 | 4.54293138 | 0.3495094 | 0.72670691 | 0.99310104 |
| miR-7060-3p | 0.26723835 | 1.58779724 | 4.54293138 | 0.3495094 | 0.72670691 | 0.99310104 |
| miR-6947-3p | 0.39289638 | 1.58779374 | 4.54293138 | 0.34950863 | 0.72670749 | 0.99310104 |
| miR-365-1-5p | 0.39289638 | 1.58779374 | 4.54293138 | 0.34950863 | 0.72670749 | 0.99310104 |
| miR-3081-5p | 0.39289638 | 1.58779374 | 4.54293138 | 0.34950863 | 0.72670749 | 0.99310104 |
| miR-5132-3p | 0.39289638 | 1.58779374 | 4.54293138 | 0.34950863 | 0.72670749 | 0.99310104 |
| miR-6944-5p | 0.32590196 | 1.58779081 | 4.54293138 | 0.34950799 | 0.72670797 | 0.99310104 |
| miR-7661-3p | 0.32590196 | 1.58779081 | 4.54293138 | 0.34950799 | 0.72670797 | 0.99310104 |
| miR-331-5p | 0.40085752 | 1.58778299 | 4.54293138 | 0.34950627 | 0.72670927 | 0.99310104 |
| miR-1188-5p | 0.40085752 | 1.58778299 | 4.54293138 | 0.34950627 | 0.72670927 | 0.99310104 |
| miR-3086-3p | 0.58934457 | 1.58776927 | 4.54293138 | 0.34950325 | 0.72671153 | 0.99310104 |
| miR-344b-3p | 0.18251593 | -1.5541331 | 4.46757711 | -0.3478693 | 0.72793831 | 0.99310104 |
| miR-3088-3p | 0.35352844 | -1.5540877 | 4.4675779 | -0.3478591 | 0.72794599 | 0.99310104 |
| miR-449a-3p | 0.17115386 | -1.5516222 | 4.46764663 | -0.3473019 | 0.72836452 | 0.99310104 |
| miR-322-5p | 0.79284218 | 1.05503218 | 3.04761372 | 0.34618304 | 0.72920516 | 0.99310104 |
| miR-186-5p | 151.5697 | -0.1395183 | 0.40665548 | -0.3430873 | 0.73153282 | 0.99310104 |
| miR-30a-5p | 3490.53891 | -0.18372 | 0.53551102 | -0.3430742 | 0.73154267 | 0.99310104 |
| miR-7118-3p | 0.56316804 | -1.5296942 | 4.46825951 | -0.3423468 | 0.73208994 | 0.99310104 |
| miR-331-3p | 0.17027165 | -1.5296237 | 4.46826074 | -0.3423309 | 0.73210189 | 0.99310104 |
| miR-1948-3p | 0.17027165 | -1.5296237 | 4.46826074 | -0.3423309 | 0.73210189 | 0.99310104 |
| miR-1298-5p | 0.17027165 | -1.5296237 | 4.46826074 | -0.3423309 | 0.73210189 | 0.99310104 |

| | | | | | | |
|---|---|---|---|---|---|---|
| miR-1843a-5p | 6.54927118 | -0.331456 | 0.98292446 | -0.3372142 | 0.73595547 | 0.99310104 |
| let-7f-1-3p | 0.16931384 | -1.4978921 | 4.46916263 | -0.3351617 | 0.73750312 | 0.99310104 |
| miR-1247-5p | 11.0116551 | -0.3428703 | 1.02586745 | -0.3342247 | 0.73820999 | 0.99310104 |
| miR-743a-5p | 7.53008851 | 0.29975682 | 0.90789506 | 0.33016681 | 0.74127392 | 0.99310104 |
| miR-205-3p | 3.40748315 | 0.52515559 | 1.6030684 | 0.327594 | 0.74321866 | 0.99310104 |
| miR-652-3p | 4.68064068 | -0.4249988 | 1.29766119 | -0.3275113 | 0.74328116 | 0.99310104 |
| miR-92a-3p | 1248.67192 | -0.1121479 | 0.34651056 | -0.3236493 | 0.74620356 | 0.99310104 |
| let-7d-5p | 257.583474 | 0.10519721 | 0.32592841 | 0.32276169 | 0.74687573 | 0.99310104 |
| miR-125b-5p | 139.941512 | 0.12594794 | 0.39398798 | 0.31967457 | 0.74921504 | 0.99310104 |
| miR-669f-3p | 0.16506736 | -1.4273832 | 4.47123648 | -0.3192368 | 0.74954695 | 0.99310104 |
| miR-871-3p | 944.913414 | 0.09822457 | 0.31037814 | 0.31646743 | 0.75164774 | 0.99310104 |
| miR-15b-3p | 1.2431204 | 0.82100036 | 2.63080518 | 0.3120719 | 0.75498588 | 0.99310104 |
| miR-881-5p | 6.08016002 | 0.36224411 | 1.16253975 | 0.31159718 | 0.75534668 | 0.99310104 |
| miR-222-3p | 46.8754024 | -0.1366606 | 0.43912179 | -0.3112134 | 0.7556384 | 0.99310104 |
| miR-194-5p | 4.27779194 | 0.33362345 | 1.07487455 | 0.31038362 | 0.75626925 | 0.99310104 |
| miR-7025-5p | 0.15186527 | -1.3722202 | 4.47292843 | -0.3067834 | 0.75900824 | 0.99310104 |
| miR-3095-3p | 1.07401383 | -0.765262 | 2.51934571 | -0.3037543 | 0.76131512 | 0.99310104 |
| miR-5107-5p | 1.60470767 | -0.7019893 | 2.3182448 | -0.3028107 | 0.76203413 | 0.99310104 |
| miR-6981-5p | 0.14594713 | -1.3091191 | 4.4749417 | -0.2925444 | 0.76987042 | 0.99310104 |
| miR-221-3p | 75.1680938 | -0.1800004 | 0.6154338 | -0.2924773 | 0.76992174 | 0.99310104 |
| let-7b-3p | 1.47839757 | -0.7646602 | 2.62247917 | -0.2915791 | 0.77060843 | 0.99310104 |
| miR-674-3p | 2.69871643 | -0.4433588 | 1.52838133 | -0.2900839 | 0.77175204 | 0.99310104 |
| miR-351-5p | 70.8202101 | -0.1635766 | 0.56438833 | -0.2898299 | 0.77194636 | 0.99310104 |
| miR-6946-3p | 0.31863131 | -1.2935107 | 4.47545234 | -0.2890235 | 0.77256343 | 0.99310104 |
| miR-669a-3p | 3.87813555 | 0.32937308 | 1.1434592 | 0.2880497 | 0.7733087 | 0.99310104 |
| miR-877-5p | 0.73376402 | -0.9811523 | 3.44144424 | -0.285099 | 0.7755683 | 0.99310104 |
| miR-8094 | 4.54244484 | 0.47457324 | 1.67769402 | 0.28287234 | 0.7772747 | 0.99310104 |
| miR-202-3p | 0.34051606 | -1.2557137 | 4.45277107 | -0.2820072 | 0.77793796 | 0.99310104 |
| miR-6390 | 0.14068817 | -1.2587396 | 4.47661093 | -0.2811814 | 0.77857129 | 0.99310104 |
| miR-135a-5p | 0.43928871 | -1.1567833 | 4.138717 | -0.2795029 | 0.77985893 | 0.99310104 |
| miR-5122 | 1.07048901 | 0.70881741 | 2.55417999 | 0.27751271 | 0.78138645 | 0.99310104 |
| miR-7237-3p | 1.07297075 | 0.82300762 | 2.97918998 | 0.27625215 | 0.78235441 | 0.99310104 |
| miR-365-2-5p | 0.36655662 | -1.1748729 | 4.32065273 | -0.2719202 | 0.78568336 | 0.99310104 |
| miR-30a-3p | 101.129552 | 0.09292528 | 0.34249826 | 0.27131605 | 0.78614796 | 0.99310104 |
| miR-744-5p | 17.7411866 | 0.21711774 | 0.80155712 | 0.27086996 | 0.78649105 | 0.99310104 |
| miR-7210-5p | 320.93364 | -0.1069702 | 0.4012156 | -0.2666152 | 0.78976548 | 0.99310104 |
| miR-326-3p | 0.9228141 | 0.87984794 | 3.31467903 | 0.26543986 | 0.79067063 | 0.99310104 |
| miR-130a-3p | 344.123953 | -0.0907114 | 0.34618491 | -0.2620317 | 0.79329704 | 0.99310104 |
| miR-5128 | 0.13051496 | -1.1712873 | 4.4796446 | -0.2614688 | 0.79373101 | 0.99310104 |
| miR-467a-5p | 10.9428867 | 0.18146651 | 0.6958764 | 0.26077405 | 0.79426676 | 0.99310104 |
| miR-26a-2-3p | 0.31192578 | -1.1528184 | 4.48030748 | -0.2573079 | 0.79694111 | 0.99310104 |
| miR-431-3p | 0.40280355 | -1.152809 | 4.48030762 | -0.2573058 | 0.79694274 | 0.99310104 |
| miR-495-3p | 0.13381221 | -1.1414945 | 4.48071897 | -0.254757 | 0.79891079 | 0.99310104 |
| miR-181a-2-3p | 7.44225008 | -0.3027338 | 1.18923954 | -0.2545608 | 0.79906235 | 0.99310104 |
| miR-155-5p | 2.95756793 | -0.3599141 | 1.42835135 | -0.2519787 | 0.80105753 | 0.99310104 |
| miR-7043-3p | 0.70262989 | -0.9090505 | 3.6315517 | -0.2503201 | 0.80233978 | 0.99310104 |

| | | | | | | |
|---|---|---|---|---|---|---|
| miR-16-1-3p | 0.76760196 | 0.86653298 | 3.47747218 | 0.24918473 | 0.80321789 | 0.99310104 |
| miR-466e-3p | 0.3926779 | -0.7929247 | 3.19706699 | -0.2480163 | 0.80412181 | 0.99310104 |
| miR-466p-3p | 0.3926779 | -0.7929247 | 3.19706699 | -0.2480163 | 0.80412181 | 0.99310104 |
| miR-466a-3p | 0.3926779 | -0.7929247 | 3.19706699 | -0.2480163 | 0.80412181 | 0.99310104 |
| miR-540-3p | 0.80058379 | 0.62221192 | 2.52100391 | 0.24681117 | 0.80505437 | 0.99310104 |
| miR-30c-5p | 1036.86164 | -0.0655291 | 0.26881131 | -0.2437738 | 0.80740603 | 0.99310104 |
| miR-1927 | 0.67772998 | 0.86651817 | 3.63276081 | 0.23852883 | 0.81147096 | 0.99310104 |
| miR-16-5p | 3180.11971 | -0.0795013 | 0.33782341 | -0.2353339 | 0.81394959 | 0.99310104 |
| miR-653-5p | 0.48559004 | -1.0486682 | 4.48420687 | -0.2338581 | 0.81509513 | 0.99310104 |
| miR-106b-5p | 0.12162261 | -1.0485979 | 4.48420845 | -0.2338424 | 0.81510736 | 0.99310104 |
| miR-1955-5p | 0.39425593 | -1.0485582 | 4.48420909 | -0.2338335 | 0.81511427 | 0.99310104 |
| miR-329-3p | 0.11899313 | -1.021067 | 4.48509005 | -0.2276581 | 0.81991206 | 0.99310104 |
| miR-26a-5p | 3518.98588 | -0.0686103 | 0.30434149 | -0.2254387 | 0.82163803 | 0.99310104 |
| miR-879-5p | 0.17702488 | -0.995923 | 4.48563612 | -0.2220249 | 0.82429448 | 0.99310104 |
| miR-218-5p | 1.2749519 | -0.4816494 | 2.18258428 | -0.2206785 | 0.82534276 | 0.99310104 |
| miR-1947-5p | 8.86575276 | -0.1943141 | 0.88467806 | -0.2196439 | 0.8261485 | 0.99310104 |
| miR-18a-5p | 6.18124308 | -0.2479156 | 1.13679171 | -0.2180835 | 0.82736402 | 0.99310104 |
| miR-7233-5p | 3.96187147 | -0.252378 | 1.16473695 | -0.2166824 | 0.82845588 | 0.99310104 |
| miR-151-5p | 403.539251 | 0.07555157 | 0.35180447 | 0.21475443 | 0.82995878 | 0.99310104 |
| miR-29b-3p | 0.79210748 | -0.6545712 | 3.08267952 | -0.2123384 | 0.83184304 | 0.99310104 |
| miR-425-3p | 0.614012 | 0.86642596 | 4.10932575 | 0.21084383 | 0.83300914 | 0.99310104 |
| miR-146a-5p | 771.592558 | 0.07707097 | 0.36779953 | 0.20954615 | 0.83402192 | 0.99310104 |
| miR-223-5p | 5.10379837 | -0.2713878 | 1.29623858 | -0.2093656 | 0.83416281 | 0.99310104 |
| miR-1983 | 1.00427594 | -0.6673987 | 3.23362539 | -0.2063933 | 0.83648368 | 0.99310104 |
| miR-5101 | 0.10847519 | -0.9174559 | 4.48739999 | -0.2044516 | 0.83800064 | 0.99310104 |
| miR-7118-5p | 0.10847519 | -0.9174559 | 4.48739999 | -0.2044516 | 0.83800064 | 0.99310104 |
| miR-6986-5p | 0.10847519 | -0.9174559 | 4.48739999 | -0.2044516 | 0.83800064 | 0.99310104 |
| miR-7013-5p | 0.10847519 | -0.9174559 | 4.48739999 | -0.2044516 | 0.83800064 | 0.99310104 |
| miR-361-5p | 7.51965324 | -0.2118894 | 1.03676971 | -0.2043746 | 0.83806074 | 0.99310104 |
| miR-361-3p | 6.2232463 | 0.28148545 | 1.39494156 | 0.20179014 | 0.84008079 | 0.99310104 |
| miR-7241-3p | 57.5906637 | -0.0962064 | 0.48178792 | -0.1996862 | 0.84172599 | 0.99310104 |
| miR-467d-3p | 0.42010048 | 0.61896219 | 3.17741683 | 0.19480044 | 0.84554918 | 0.99310104 |
| miR-9769-3p | 0.6205535 | 0.86654379 | 4.47987567 | 0.19343032 | 0.84662197 | 0.99310104 |
| miR-350-3p | 6.88109379 | -0.2214846 | 1.15593254 | -0.1916069 | 0.84805014 | 0.99310104 |
| miR-92a-1-5p | 0.44965109 | 0.86641201 | 4.53958552 | 0.19085707 | 0.84863757 | 0.99310104 |
| miR-700-3p | 0.07697561 | 0.86658818 | 4.54292652 | 0.19075549 | 0.84871716 | 0.99310104 |
| miR-702-3p | 0.42434974 | 0.86652852 | 4.54292462 | 0.19074244 | 0.84872739 | 0.99310104 |
| miR-382-5p | 0.2782138 | 0.86651791 | 4.54292451 | 0.19074011 | 0.84872921 | 0.99310104 |
| miR-377-5p | 0.1016543 | 0.86649101 | 4.54292425 | 0.1907342 | 0.84873384 | 0.99310104 |
| miR-3084-5p | 0.10317053 | 0.86644944 | 4.54292376 | 0.19072507 | 0.848741 | 0.99310104 |
| miR-7236-3p | 0.34344628 | 0.86639273 | 4.54292302 | 0.19071262 | 0.84875075 | 0.99310104 |
| miR-3066-3p | 0.46849696 | 0.83387076 | 4.45523101 | 0.18716667 | 0.85152996 | 0.99310104 |
| miR-6948-3p | 0.9026544 | 0.57163016 | 3.06271449 | 0.18664167 | 0.85194159 | 0.99310104 |
| miR-669o-3p | 0.21123546 | 0.82926129 | 4.51120117 | 0.18382272 | 0.85415253 | 0.99310104 |
| miR-297c-3p | 0.50000017 | 0.50288945 | 2.75639862 | 0.18244438 | 0.855234 | 0.99310104 |
| miR-297b-3p | 0.50000017 | 0.50288945 | 2.75639862 | 0.18244438 | 0.855234 | 0.99310104 |

| | | | | | | |
|---|---|---|---|---|---|---|
| miR-669b-5p | 0.09759216 | -0.797603 | 4.49028027 | -0.1776288 | 0.85901452 | 0.99310104 |
| miR-7221-3p | 12.4474012 | 0.16813572 | 0.96140212 | 0.17488595 | 0.86116925 | 0.99310104 |
| miR-380-3p | 0.34172456 | -0.782487 | 4.49066084 | -0.1742476 | 0.86167086 | 0.99310104 |
| miR-7222-3p | 2.3458522 | 0.28452447 | 1.64300177 | 0.17317356 | 0.862515 | 0.99310104 |
| miR-196a-5p | 21.734875 | -0.1116143 | 0.65046225 | -0.1715923 | 0.86375804 | 0.99310104 |
| miR-7223-5p | 1.38907708 | 0.45184018 | 2.71306871 | 0.16654211 | 0.86773035 | 0.99310104 |
| miR-669l-5p | 0.75556817 | -0.5267374 | 3.2579694 | -0.1616766 | 0.87156053 | 0.99310104 |
| miR-8097 | 0.64813936 | -0.6601204 | 4.0872322 | -0.1615079 | 0.87169337 | 0.99310104 |
| miR-217-3p | 0.12167728 | -0.7174129 | 4.49234054 | -0.1596969 | 0.87311984 | 0.99310104 |
| miR-7021-5p | 0.12167728 | -0.7174129 | 4.49234054 | -0.1596969 | 0.87311984 | 0.99310104 |
| miR-7054-5p | 0.12167728 | -0.7174129 | 4.49234054 | -0.1596969 | 0.87311984 | 0.99310104 |
| miR-3473d | 0.12167728 | -0.7174129 | 4.49234054 | -0.1596969 | 0.87311984 | 0.99310104 |
| miR-5626-3p | 0.12167728 | -0.7174129 | 4.49234054 | -0.1596969 | 0.87311984 | 0.99310104 |
| miR-7646-5p | 0.12167728 | -0.7174129 | 4.49234054 | -0.1596969 | 0.87311984 | 0.99310104 |
| miR-370-3p | 0.21255506 | -0.7173818 | 4.49234114 | -0.15969 | 0.87312531 | 0.99310104 |
| miR-7082-5p | 0.29268979 | -0.7173527 | 4.49234169 | -0.1596835 | 0.87313042 | 0.99310104 |
| miR-7653-5p | 0.72175358 | -0.5631838 | 3.56530606 | -0.1579623 | 0.87448653 | 0.99310104 |
| miR-181c-3p | 0.7050816 | 0.51776266 | 3.30769752 | 0.15653265 | 0.87561319 | 0.99310104 |
| miR-17-3p | 0.59741414 | -0.5268503 | 3.47163816 | -0.1517584 | 0.87937747 | 0.99310104 |
| miR-152-5p | 3.40236558 | -0.2345034 | 1.55647424 | -0.1506632 | 0.88024138 | 0.99310104 |
| miR-7015-3p | 0.20810263 | -0.6565975 | 4.49397872 | -0.1461061 | 0.88383767 | 0.99310104 |
| miR-8104 | 0.08678015 | -0.6565639 | 4.49397923 | -0.1460986 | 0.88384357 | 0.99310104 |
| miR-203-5p | 0.08678015 | -0.6565639 | 4.49397923 | -0.1460986 | 0.88384357 | 0.99310104 |
| miR-6940-3p | 0.08678015 | -0.6565639 | 4.49397923 | -0.1460986 | 0.88384357 | 0.99310104 |
| miR-3547-5p | 0.08678015 | -0.6565639 | 4.49397923 | -0.1460986 | 0.88384357 | 0.99310104 |
| miR-7216-3p | 0.08678015 | -0.6565639 | 4.49397923 | -0.1460986 | 0.88384357 | 0.99310104 |
| miR-7668-5p | 0.08678015 | -0.6565639 | 4.49397923 | -0.1460986 | 0.88384357 | 0.99310104 |
| miR-6236 | 0.08678015 | -0.6565639 | 4.49397923 | -0.1460986 | 0.88384357 | 0.99310104 |
| let-7c-1-3p | 0.2378391 | -0.6565502 | 4.49397937 | -0.1460955 | 0.88384599 | 0.99310104 |
| miR-1938 | 0.48657648 | -0.6565167 | 4.49397972 | -0.146088 | 0.88385188 | 0.99310104 |
| miR-7232-5p | 0.65927838 | -0.4711887 | 3.32395895 | -0.1417553 | 0.88727331 | 0.99310104 |
| miR-344d-3p | 0.12847132 | 0.63734012 | 4.50419258 | 0.1414993 | 0.88747551 | 0.99310104 |
| miR-676-5p | 2.23226047 | -0.2531361 | 1.80415179 | -0.1403076 | 0.88841699 | 0.99310104 |
| miR-7210-3p | 14.8382315 | 0.09503151 | 0.67848125 | 0.14006505 | 0.88860859 | 0.99310104 |
| let-7c-5p | 5728.75798 | 0.05593471 | 0.40921069 | 0.13668928 | 0.8912764 | 0.99310104 |
| miR-34b-5p | 24.8292741 | -0.1990048 | 1.49755628 | -0.1328864 | 0.89428325 | 0.99310104 |
| miR-467e-3p | 0.1311008 | 0.5954044 | 4.50277562 | 0.13223053 | 0.89480195 | 0.99310104 |
| miR-297a-3p | 1.07183877 | 0.2543851 | 1.952631 | 0.13027812 | 0.89634639 | 0.99310104 |
| let-7i-5p | 646.879343 | -0.0516076 | 0.41921411 | -0.1231057 | 0.90202343 | 0.99310104 |
| miR-293-5p | 6.37104336 | -0.1471604 | 1.20787357 | -0.1218342 | 0.9030303 | 0.99310104 |
| miR-743a-3p | 10.6800043 | 0.08168642 | 0.69206064 | 0.11803361 | 0.90604103 | 0.99310104 |
| miR-7215-5p | 0.27346293 | -0.5245019 | 4.46949761 | -0.1173514 | 0.90658158 | 0.99310104 |
| miR-431-5p | 1.10369239 | 0.28198783 | 2.4084257 | 0.11708388 | 0.90679358 | 0.99310104 |
| miR-27a-3p | 18.3808121 | 0.08410767 | 0.74450966 | 0.11297056 | 0.9100539 | 0.99310104 |
| miR-28a-3p | 11.530027 | 0.09618288 | 0.8538254 | 0.11264936 | 0.91030855 | 0.99310104 |
| miR-27b-5p | 1.90329209 | 0.25197946 | 2.29512657 | 0.10978892 | 0.91257678 | 0.99310104 |

| | | | | | | |
|---|---|---|---|---|---|---|
| miR-7032-3p | 0.07297357 | -0.4466206 | 4.50017122 | -0.0992452 | 0.92094355 | 0.99310104 |
| miR-7059-5p | 0.07297357 | -0.4466206 | 4.50017122 | -0.0992452 | 0.92094355 | 0.99310104 |
| miR-3102-5p.2- | 0.16385134 | -0.4466071 | 4.50017138 | -0.0992422 | 0.92094593 | 0.99310104 |
| miR-485-3p | 1.01910843 | 0.27404078 | 2.79419256 | 0.09807512 | 0.92187264 | 0.99310104 |
| miR-22-5p | 0.53459881 | -0.3158976 | 3.27125457 | -0.0965677 | 0.92306968 | 0.99310104 |
| miR-708-3p | 0.62483669 | 0.36159424 | 3.7450339 | 0.096553 | 0.92308139 | 0.99310104 |
| miR-467b-5p | 0.9405871 | 0.18032959 | 1.90966216 | 0.0944301 | 0.92476751 | 0.99310104 |
| miR-6240 | 22.4600502 | -0.0621407 | 0.68706009 | -0.0904443 | 0.92793413 | 0.99310104 |
| miR-10b-3p | 5.87406405 | 0.10153068 | 1.13344996 | 0.08957668 | 0.92862362 | 0.99310104 |
| miR-185-5p | 2.9484287 | -0.1549249 | 1.76478566 | -0.0877868 | 0.93004611 | 0.99310104 |
| miR-467d-5p | 6.87334942 | -0.0922435 | 1.05418352 | -0.0875024 | 0.93027221 | 0.99310104 |
| miR-133b-3p | 0.118135 | 0.39360602 | 4.53352384 | 0.08682121 | 0.93081363 | 0.99310104 |
| miR-6970-5p | 0.20933333 | -0.3473608 | 4.47494119 | -0.0776236 | 0.93812751 | 0.99310104 |
| miR-181a-5p | 616.065004 | 0.03448733 | 0.46744661 | 0.07377811 | 0.94118694 | 0.99310104 |
| miR-450a-5p | 0.24706883 | -0.3244657 | 4.50419219 | -0.0720364 | 0.94257296 | 0.99310104 |
| miR-1931 | 0.06508512 | -0.3244192 | 4.50419309 | -0.072026 | 0.94258119 | 0.99310104 |
| miR-6972-3p | 0.06508512 | -0.3244192 | 4.50419309 | -0.072026 | 0.94258119 | 0.99310104 |
| miR-9-3p | 0.27670588 | 0.30517725 | 4.49397856 | 0.06790803 | 0.94585884 | 0.99310104 |
| miR-7217-5p | 23.1520426 | 0.06245233 | 0.95996626 | 0.0650568 | 0.94812878 | 0.99310104 |
| miR-26b-3p | 0.95444117 | -0.1580515 | 2.80795386 | -0.0562871 | 0.95511312 | 0.99310104 |
| miR-1943-5p | 0.23012752 | 0.24430387 | 4.49233903 | 0.05438233 | 0.95663055 | 0.99310104 |
| miR-7218-3p | 9.6558705 | -0.0560062 | 1.05648156 | -0.053012 | 0.95772237 | 0.99310104 |
| miR-1982-5p | 1.15280594 | 0.13190276 | 2.57423576 | 0.05123958 | 0.95913461 | 0.99310104 |
| miR-340-3p | 3.66941818 | -0.0716651 | 1.47547908 | -0.0485707 | 0.96126141 | 0.99310104 |
| miR-6911-3p | 1.14584515 | 0.13941584 | 2.8939567 | 0.04817482 | 0.96157692 | 0.99310104 |
| miR-7232-3p | 49.1727142 | 0.02043754 | 0.44484298 | 0.04594327 | 0.96335547 | 0.99310104 |
| miR-5126 | 2.20662156 | -0.0702777 | 1.56009288 | -0.0450471 | 0.96406974 | 0.99310104 |
| miR-484 | 5.68165104 | -0.0485038 | 1.22607894 | -0.0395601 | 0.96844386 | 0.99310104 |
| miR-1a-3p | 8.02350501 | 0.03747471 | 1.02443086 | 0.036581 | 0.97081909 | 0.99310104 |
| miR-423-5p | 599.714497 | 0.01450993 | 0.40019752 | 0.03625691 | 0.97107751 | 0.99310104 |
| miR-3963 | 0.39910525 | 0.13506082 | 3.868487 | 0.03491309 | 0.97214905 | 0.99310104 |
| miR-883a-3p | 19.3266994 | 0.0201185 | 0.61146594 | 0.03290208 | 0.97375267 | 0.99310104 |
| miR-140-3p | 43.1775835 | -0.0170678 | 0.52028309 | -0.0328049 | 0.97383016 | 0.99310104 |
| let-7e-3p | 0.13426785 | 0.13384955 | 4.52251126 | 0.02959629 | 0.97638903 | 0.99310104 |
| miR-6916-5p | 0.04339008 | 0.13383503 | 4.52251098 | 0.02959308 | 0.97639158 | 0.99310104 |
| miR-217-5p | 0.04339008 | 0.13383503 | 4.52251098 | 0.02959308 | 0.97639158 | 0.99310104 |
| miR-301a-5p | 0.04339008 | 0.13383503 | 4.52251098 | 0.02959308 | 0.97639158 | 0.99310104 |
| miR-6931-3p | 0.04339008 | 0.13383503 | 4.52251098 | 0.02959308 | 0.97639158 | 0.99310104 |
| miR-377-3p | 0.04339008 | 0.13383503 | 4.52251098 | 0.02959308 | 0.97639158 | 0.99310104 |
| miR-369-5p | 0.04339008 | 0.13383503 | 4.52251098 | 0.02959308 | 0.97639158 | 0.99310104 |
| miR-7218-5p | 0.04339008 | 0.13383503 | 4.52251098 | 0.02959308 | 0.97639158 | 0.99310104 |
| miR-6985-3p | 0.04339008 | 0.13383503 | 4.52251098 | 0.02959308 | 0.97639158 | 0.99310104 |
| miR-208b-3p | 0.16471256 | 0.13379887 | 4.52251 | 0.02958509 | 0.97639796 | 0.99310104 |
| miR-3079-5p | 0.28781655 | 0.13377737 | 4.52250942 | 0.02958034 | 0.97640175 | 0.99310104 |
| miR-7b-5p | 1.36611929 | 0.06617371 | 2.24368791 | 0.02949328 | 0.97647118 | 0.99310104 |
| miR-10a-3p | 2.983913 | 0.0532576 | 1.90195794 | 0.02800146 | 0.97766098 | 0.99310104 |

| | | | | | | |
|---|---|---|---|---|---|---|
| miR-1839-5p | 125.16915 | -0.0141869 | 0.51052012 | -0.027789 | 0.97783043 | 0.99310104 |
| miR-335-5p | 5.31225984 | -0.0295236 | 1.08187995 | -0.0272891 | 0.97822913 | 0.99310104 |
| miR-7068-3p | 12.7803931 | -0.0256004 | 0.95887128 | -0.0266984 | 0.97870026 | 0.99310104 |
| miR-296-3p | 0.32202775 | -0.1143043 | 4.43870484 | -0.0257517 | 0.97945537 | 0.99310104 |
| let-7e-5p | 645.74885 | -0.0061294 | 0.40238546 | -0.0152326 | 0.98784661 | 0.99681925 |
| miR-9-5p | 6.40917121 | 0.0145229 | 1.09079213 | 0.01331409 | 0.98937721 | 0.99681925 |
| miR-6516-5p | 0.21692543 | 0.04430475 | 4.48739968 | 0.00987315 | 0.9921225 | 0.99681925 |
| miR-148b-5p | 1.41662624 | 0.01445312 | 1.94792302 | 0.00741976 | 0.99407994 | 0.99681925 |
| miR-7073-5p | 0.78357044 | 0.02331175 | 3.85781938 | 0.00604273 | 0.99517863 | 0.99681925 |
| let-7a-1-3p | 1.18944819 | -0.0105996 | 2.2475695 | -0.004716 | 0.99623717 | 0.99681925 |
| let-7c-2-3p | 1.18944819 | -0.0105996 | 2.2475695 | -0.004716 | 0.99623717 | 0.99681925 |
| miR-350-5p | 0.29106583 | 0.01801746 | 4.51735061 | 0.0039885 | 0.99681765 | 0.99681925 |
| miR-300-5p | 0.04864904 | 0.01800835 | 4.5173506 | 0.00398648 | 0.99681925 | 0.99681925 |
| miR-103-3p | 203.394758 | -0.0804817 | 0.60473546 | -0.1330858 | NA | NA |
| miR-378a-3p | 17.4797114 | 0.95346947 | 0.93672059 | 1.01788034 | NA | NA |
| miR-107-3p | 66.4487281 | 0.05943307 | 0.75436027 | 0.07878606 | NA | NA |
| miR-463-5p | 236.689323 | -0.0744636 | 0.61245687 | -0.1215819 | NA | NA |

| HS692_miRNA_pvalues_TIME_padj05 | | | | HS692_miRNA_pvalues_TIME_padj05 | | |
|---|---|---|---|---|---|---|
| | baseMean | log2FoldChai | lfcSE | stat | pvalue | padj |
| miR-7232-3p | 49.1727142 | -1.4225362 | 0.43426568 | -3.2757279 | 0.0010539 | 0.47899824 |
| miR-7217-3p | 97.335608 | -1.4616605 | 0.45935196 | -3.1820056 | 0.00146259 | 0.47899824 |
| miR-142a-5p | 317.260067 | 0.87240827 | 0.29136079 | 2.99425421 | 0.00275117 | 0.48745876 |
| miR-196a-5p | 21.734875 | -1.9305344 | 0.64998563 | -2.9701187 | 0.00297685 | 0.48745876 |
| miR-5114 | 11.0690558 | 2.50805166 | 0.87566448 | 2.86416969 | 0.00418104 | 0.5477162 |
| miR-6538 | 16.2051488 | 1.74947178 | 0.63041627 | 2.77510569 | 0.00551838 | 0.60242284 |
| miR-128-3p | 148.185578 | 0.92126846 | 0.34004887 | 2.70922374 | 0.00674408 | 0.63105361 |
| miR-150-5p | 67.1664819 | 1.34445656 | 0.51426252 | 2.61433899 | 0.00894003 | 0.73196486 |
| let-7b-5p | 480.909708 | -0.9884304 | 0.41906456 | -2.3586589 | 0.0183411 | 0.98715323 |
| miR-5099 | 84.7086433 | 1.62705137 | 0.69166792 | 2.35235917 | 0.01865475 | 0.98715323 |
| miR-145a-5p | 61.9461242 | -1.7099318 | 0.72965783 | -2.3434708 | 0.01910526 | 0.98715323 |
| miR-26b-5p | 124.394455 | -0.9001833 | 0.39394415 | -2.2850532 | 0.02230971 | 0.98715323 |
| miR-335-3p | 3.1411732 | -3.6692983 | 1.61155656 | -2.276866 | 0.02279423 | 0.98715323 |
| miR-379-5p | 4.0139086 | -3.3286168 | 1.49398437 | -2.2280131 | 0.02587964 | 0.98715323 |
| miR-127-3p | 133.600536 | -1.0205949 | 0.46630399 | -2.1886901 | 0.02861937 | 0.98715323 |
| miR-21a-3p | 18.0654067 | 1.44045552 | 0.66209292 | 2.1756093 | 0.02958449 | 0.98715323 |
| miR-7233-3p | 86.2479996 | -0.7985034 | 0.36990902 | -2.158648 | 0.03087749 | 0.98715323 |
| miR-7241-3p | 57.5906637 | -0.9829321 | 0.45604221 | -2.1553533 | 0.0311342 | 0.98715323 |
| miR-871-5p | 716.00589 | 0.73048512 | 0.34641869 | 2.10867697 | 0.03497247 | 0.98715323 |
| miR-100-5p | 91.5998735 | -0.9311264 | 0.44774101 | -2.0796094 | 0.03756138 | 0.98715323 |
| miR-99b-5p | 638.306367 | -0.8697018 | 0.42268408 | -2.0575693 | 0.03963149 | 0.98715323 |
| miR-677-3p | 3.47500083 | -3.3810713 | 1.65229309 | -2.0462903 | 0.04072782 | 0.98715323 |
| miR-7229-5p | 78.3795612 | -0.8427886 | 0.41263627 | -2.0424492 | 0.041107 | 0.98715323 |
| miR-3068-3p | 13.5648049 | 1.28295617 | 0.64602301 | 1.98592953 | 0.04704115 | 0.98715323 |
| miR-192-5p | 525.686551 | 0.73542269 | 0.37194229 | 1.97724944 | 0.04801344 | 0.98715323 |
| miR-139-5p | 10.5989984 | 1.47073942 | 0.75392041 | 1.95078872 | 0.05108218 | 0.98715323 |
| miR-18a-3p | 2.41760672 | -3.1413159 | 1.6132565 | -1.9471893 | 0.05151204 | 0.98715323 |
| miR-182-5p | 3490.89211 | 0.53342842 | 0.28085893 | 1.89927527 | 0.05752829 | 0.98715323 |
| miR-434-3p | 23.72971 | -1.2423248 | 0.66001076 | -1.8822795 | 0.05979807 | 0.98715323 |
| miR-26a-5p | 3518.98588 | -0.5297742 | 0.28193205 | -1.8790847 | 0.06023293 | 0.98715323 |
| miR-151-3p | 493.324752 | -0.4800109 | 0.26098893 | -1.8392004 | 0.06588572 | 0.98715323 |
| miR-429-3p | 97.3046123 | 0.95475056 | 0.53231477 | 1.79358271 | 0.0728798 | 0.98715323 |
| miR-6240 | 22.4600502 | 1.09393962 | 0.61239249 | 1.78633743 | 0.07404464 | 0.98715323 |
| miR-471-5p | 12.6499637 | 1.63365084 | 0.92955963 | 1.75744597 | 0.07884182 | 0.98715323 |
| miR-541-5p | 85.6242488 | -0.633533 | 0.36065387 | -1.7566232 | 0.07898206 | 0.98715323 |
| miR-532-5p | 35.9742044 | 0.73754877 | 0.42345487 | 1.74174114 | 0.08155375 | 0.98715323 |
| miR-328-3p | 14.3504627 | -1.0765637 | 0.62524712 | -1.7218211 | 0.08510193 | 0.98715323 |
| miR-7648-3p | 1.1435915 | 4.72443899 | 2.80993186 | 1.68133579 | 0.09269771 | 0.98715323 |
| miR-218-5p | 1.2749519 | -3.6420419 | 2.19156954 | -1.6618418 | 0.0965445 | 0.98715323 |
| miR-881-3p | 131.778904 | 0.56587737 | 0.34112074 | 1.65887706 | 0.09714057 | 0.98715323 |
| miR-1198-5p | 3.55459749 | -2.272394 | 1.40054679 | -1.6225049 | 0.10469529 | 0.98715323 |
| miR-212-5p | 1.00092627 | 4.34742254 | 2.6950097 | 1.61313799 | 0.10671453 | 0.98715323 |
| miR-99a-5p | 82.9188033 | -0.8154868 | 0.50660635 | -1.6097051 | 0.10746226 | 0.98715323 |
| miR-677-5p | 22.6023362 | 1.16886825 | 0.72902482 | 1.60333121 | 0.10886155 | 0.98715323 |

| | | | | | | |
|---|---|---|---|---|---|---|
| miR-132-5p | 0.97063471 | 5.05357624 | 3.15865153 | 1.59991572 | 0.10961728 | 0.98715323 |
| miR-1247-3p | 10.5798277 | 1.06963318 | 0.67082616 | 1.5945013 | 0.1108238 | 0.98715323 |
| miR-200a-5p | 1.2069913 | 4.15693589 | 2.60849759 | 1.59361309 | 0.11102272 | 0.98715323 |
| miR-205-3p | 3.40748315 | 2.20511236 | 1.40766796 | 1.56650036 | 0.11723152 | 0.98715323 |
| miR-6911-3p | 1.14584515 | 3.91154919 | 2.52271696 | 1.55053035 | 0.12101428 | 0.98715323 |
| miR-465c-3p | 161.520091 | 0.64375021 | 0.41691956 | 1.54406335 | 0.12257299 | 0.98715323 |
| miR-485-5p | 4.73326633 | -2.2770757 | 1.47524283 | -1.543526 | 0.12270319 | 0.98715323 |
| miR-465b-3p | 162.106103 | 0.64119106 | 0.41684879 | 1.53818619 | 0.1240031 | 0.98715323 |
| miR-30b-5p | 328.954044 | 0.48272262 | 0.31573594 | 1.52888082 | 0.12629399 | 0.98715323 |
| miR-147-5p | 13.8428067 | 0.82905734 | 0.54324584 | 1.52611815 | 0.12698044 | 0.98715323 |
| miR-615-5p | 4.90884478 | 2.04025575 | 1.34492204 | 1.51700671 | 0.12926499 | 0.98715323 |
| miR-7230-3p | 46.0677911 | -0.6530259 | 0.43322031 | -1.5073761 | 0.13171428 | 0.98715323 |
| miR-222-3p | 46.8754024 | 0.59436978 | 0.39524589 | 1.50379749 | 0.13263352 | 0.98715323 |
| miR-7234-3p | 30.7430264 | -0.8713303 | 0.5816328 | -1.4980762 | 0.13411346 | 0.98715323 |
| miR-468-3p | 6.10232842 | -1.6257391 | 1.08664426 | -1.4961098 | 0.13462506 | 0.98715323 |
| miR-465a-3p | 81.2636666 | 0.64866024 | 0.43445074 | 1.49305819 | 0.13542195 | 0.98715323 |
| miR-7214-5p | 17.3857301 | -1.2195785 | 0.82485359 | -1.4785394 | 0.13926346 | 0.98715323 |
| miR-7217-5p | 23.1520426 | -1.3172985 | 0.91053401 | -1.4467318 | 0.14797205 | 0.98715323 |
| miR-7210-5p | 320.93364 | -0.5389926 | 0.3727395 | -1.4460303 | 0.14816869 | 0.98715323 |
| miR-672-5p | 40.9597656 | 0.66683106 | 0.4720508 | 1.41262565 | 0.15776582 | 0.98715323 |
| miR-423-3p | 47.2313136 | -0.6524299 | 0.46448907 | -1.4046183 | 0.16013481 | 0.98715323 |
| miR-682 | 0.8790814 | 3.95039267 | 2.81851251 | 1.40158777 | 0.16103838 | 0.98715323 |
| miR-1249-3p | 7.14171159 | -1.3632746 | 0.9814061 | -1.3891034 | 0.1648013 | 0.98715323 |
| miR-125a-5p | 566.450555 | -0.6362469 | 0.46409389 | -1.3709444 | 0.1703923 | 0.98715323 |
| miR-1843a-3p | 3.48033372 | 1.69050146 | 1.23969869 | 1.36363898 | 0.17268122 | 0.98715323 |
| miR-155-5p | 2.95756793 | -1.9729132 | 1.45765719 | -1.3534823 | 0.17590161 | 0.98715323 |
| miR-125b-2-3p | 14.8456703 | 0.84444571 | 0.62780247 | 1.34508187 | 0.1785988 | 0.98715323 |
| miR-5121 | 6.97170829 | 1.34619903 | 1.01472446 | 1.3266646 | 0.18461966 | 0.98715323 |
| let-7g-5p | 1132.43314 | 0.40513151 | 0.30552022 | 1.32603828 | 0.18482702 | 0.98715323 |
| miR-194-5p | 4.27779194 | -1.4362243 | 1.11241471 | -1.2910871 | 0.19667349 | 0.98715323 |
| miR-741-3p | 127.971851 | -0.5785553 | 0.44827994 | -1.2906117 | 0.19683834 | 0.98715323 |
| miR-434-5p | 3.20697696 | -2.1467415 | 1.66511395 | -1.289246 | 0.19731258 | 0.98715323 |
| let-7d-3p | 83.7901595 | -0.504731 | 0.39205991 | -1.2873823 | 0.19796107 | 0.98715323 |
| miR-342-5p | 6.79951387 | 1.28975126 | 1.00843977 | 1.27895716 | 0.20091214 | 0.98715323 |
| miR-125b-1-3p | 6.8504028 | -1.4242013 | 1.12806545 | -1.2625165 | 0.20676298 | 0.98715323 |
| miR-7215-3p | 0.89167789 | 3.41844223 | 2.72650697 | 1.25378085 | 0.20992167 | 0.98715323 |
| miR-296-5p | 15.7564006 | -0.8943101 | 0.71681622 | -1.2476142 | 0.21217237 | 0.98715323 |
| miR-467d-5p | 6.87334942 | 1.16874172 | 0.93875305 | 1.24499379 | 0.21313403 | 0.98715323 |
| miR-212-3p | 2.43251707 | -2.4291768 | 1.95358683 | -1.2434445 | 0.21370409 | 0.98715323 |
| miR-671-5p | 1.29064859 | -3.1358989 | 2.53522324 | -1.236932 | 0.21611231 | 0.98715323 |
| miR-125a-3p | 3.50064133 | -2.3661815 | 1.91994109 | -1.232424 | 0.21779074 | 0.98715323 |
| miR-8118 | 0.68553564 | 4.78024378 | 3.88818959 | 1.22942662 | 0.2189119 | 0.98715323 |
| miR-1839-5p | 125.16915 | 0.57275715 | 0.47066312 | 1.2169153 | 0.22363644 | 0.98715323 |
| miR-381-3p | 1.4691769 | -2.1880157 | 1.79865455 | -1.2164736 | 0.22380458 | 0.98715323 |
| miR-181d-5p | 11.1097067 | -0.89926 | 0.74465986 | -1.2076118 | 0.22719662 | 0.98715323 |
| miR-31-5p | 5.93726446 | -1.1598407 | 0.96263191 | -1.2048642 | 0.22825575 | 0.98715323 |

| | | | | | | |
|---|---|---|---|---|---|---|
| miR-27a-3p | 18.3808121 | -0.858146 | 0.71341653 | -1.2028682 | 0.22902734 | 0.98715323 |
| miR-214-3p | 3.82059379 | -1.5406635 | 1.29315789 | -1.1913963 | 0.23349804 | 0.98715323 |
| miR-7242-3p | 19.1694289 | -0.9829225 | 0.82613871 | -1.189779 | 0.23413327 | 0.98715323 |
| miR-878-5p | 70.3483372 | 0.47325685 | 0.39834445 | 1.18805936 | 0.23481002 | 0.98715323 |
| miR-1956 | 0.7075449 | 4.59099543 | 3.88360152 | 1.18214894 | 0.23714661 | 0.98715323 |
| miR-501-3p | 4.84182452 | -1.4163778 | 1.19948501 | -1.1808216 | 0.2376736 | 0.98715323 |
| miR-340-5p | 230.688377 | 0.57454557 | 0.48992235 | 1.17272784 | 0.24090496 | 0.98715323 |
| miR-3086-3p | 0.58934457 | 4.83439839 | 4.12529104 | 1.17189268 | 0.24124015 | 0.98715323 |
| miR-221-5p | 15.7134515 | -0.8423171 | 0.72229444 | -1.1661686 | 0.24354628 | 0.98715323 |
| miR-669a-5p | 18.083587 | 0.62195949 | 0.53668373 | 1.15889387 | 0.24649945 | 0.98715323 |
| miR-7230-5p | 12.6123363 | -0.668179 | 0.58366887 | -1.1447913 | 0.25229563 | 0.98715323 |
| miR-152-5p | 3.40236558 | 1.53746599 | 1.35603266 | 1.13379717 | 0.25687964 | 0.98715323 |
| miR-184-5p | 0.73658487 | 3.66642401 | 3.25646525 | 1.12589072 | 0.26021178 | 0.98715323 |
| miR-1947-5p | 8.86575276 | 0.88306396 | 0.7844568 | 1.1257012 | 0.26029201 | 0.98715323 |
| let-7a-5p | 12110.1372 | 0.32329803 | 0.28795561 | 1.12273564 | 0.26154979 | 0.98715323 |
| miR-196b-3p | 1.24694237 | 3.28810945 | 2.99110306 | 1.09929661 | 0.27163871 | 0.98715323 |
| miR-223-3p | 2.14767617 | -1.8739141 | 1.7254283 | -1.0860574 | 0.27745362 | 0.98715323 |
| miR-1983 | 1.00427594 | 3.08622894 | 2.84869822 | 1.0833822 | 0.27863881 | 0.98715323 |
| miR-574-3p | 9.99368537 | -1.1056776 | 1.02612821 | -1.0775239 | 0.28124629 | 0.98715323 |
| miR-181c-5p | 168.593623 | 0.45714632 | 0.42457942 | 1.07670391 | 0.28161257 | 0.98715323 |
| let-7e-5p | 645.74885 | -0.397387 | 0.37314746 | -1.0649598 | 0.28689413 | 0.98715323 |
| miR-184-3p | 2339.78942 | -0.3410459 | 0.32121003 | -1.0617537 | 0.28834751 | 0.98715323 |
| miR-3086-5p | 1.32850623 | -2.4931397 | 2.35294125 | -1.0595844 | 0.28933373 | 0.98715323 |
| miR-7042-5p | 0.57187003 | 4.36901706 | 4.14197439 | 1.05481508 | 0.2915099 | 0.98715323 |
| miR-331-5p | 0.40085752 | 4.36899428 | 4.14197482 | 1.05480947 | 0.29151247 | 0.98715323 |
| miR-1188-5p | 0.40085752 | 4.36899428 | 4.14197482 | 1.05480947 | 0.29151247 | 0.98715323 |
| miR-881-5p | 6.08016002 | -1.2011695 | 1.15385794 | -1.041003 | 0.29787417 | 0.98715323 |
| miR-425-3p | 0.614012 | 3.87205562 | 3.72379414 | 1.03981463 | 0.29842603 | 0.98715323 |
| miR-365-3p | 9.67637868 | -1.0505756 | 1.01557385 | -1.034465 | 0.30091883 | 0.98715323 |
| miR-187-5p | 2.16057964 | 2.45942421 | 2.39272599 | 1.02787541 | 0.30400844 | 0.98715323 |
| miR-532-3p | 4.07129501 | 1.27009913 | 1.24519722 | 1.01999837 | 0.30772924 | 0.98715323 |
| miR-743b-3p | 512.187236 | 0.37517415 | 0.36986007 | 1.0143678 | 0.31040728 | 0.98715323 |
| miR-32-5p | 2.22823869 | 1.52351482 | 1.51390911 | 1.00634497 | 0.31424965 | 0.98715323 |
| miR-500-3p | 0.52674753 | 3.88809203 | 3.88499887 | 1.00079618 | 0.31692536 | 0.98715323 |
| miR-1668 | 0.49396646 | 4.14522298 | 4.14865037 | 0.99917385 | 0.31771048 | 0.98715323 |
| miR-148b-3p | 110.979738 | -0.4983259 | 0.50126568 | -0.9941352 | 0.32015703 | 0.98715323 |
| miR-6944-5p | 0.32590196 | 4.10368008 | 4.1497803 | 0.98889092 | 0.32271651 | 0.98715323 |
| miR-7661-3p | 0.32590196 | 4.10368008 | 4.1497803 | 0.98889092 | 0.32271651 | 0.98715323 |
| miR-15a-3p | 0.82085359 | 2.47184003 | 2.50203185 | 0.98793308 | 0.32318542 | 0.98715323 |
| miR-140-5p | 1.3537423 | 2.19544013 | 2.26060261 | 0.97117473 | 0.33146128 | 0.98715323 |
| miR-496a-3p | 1.14879701 | -2.0910754 | 2.15325231 | -0.9711242 | 0.33148644 | 0.98715323 |
| miR-125b-5p | 139.941512 | -0.3556908 | 0.36813448 | -0.9661979 | 0.33394515 | 0.98715323 |
| miR-6947-3p | 0.39289638 | 4.00430235 | 4.15261483 | 0.96428456 | 0.33490328 | 0.98715323 |
| miR-365-1-5p | 0.39289638 | 4.00430235 | 4.15261483 | 0.96428456 | 0.33490328 | 0.98715323 |
| miR-3081-5p | 0.39289638 | 4.00430235 | 4.15261483 | 0.96428456 | 0.33490328 | 0.98715323 |
| miR-5132-3p | 0.39289638 | 4.00430235 | 4.15261483 | 0.96428456 | 0.33490328 | 0.98715323 |

| | | | | | | |
|---|---|---|---|---|---|---|
| let-7c-5p | 5728.75798 | -0.3629109 | 0.37892746 | -0.9577319 | 0.33819795 | 0.98715323 |
| miR-351-5p | 70.8202101 | -0.5025643 | 0.52671815 | -0.9541427 | 0.34001144 | 0.98715323 |
| miR-5617-3p | 0.60926336 | 3.88612446 | 4.11217204 | 0.94502964 | 0.34464375 | 0.98715323 |
| miR-669l-5p | 0.75556817 | -2.9183994 | 3.09688128 | -0.9423672 | 0.34600466 | 0.98715323 |
| miR-669d-5p | 1.09656679 | -2.0651853 | 2.19172114 | -0.9422665 | 0.34605623 | 0.98715323 |
| miR-7653-3p | 0.26723835 | 3.8857357 | 4.15625181 | 0.93491344 | 0.34983291 | 0.98715323 |
| miR-7670-3p | 0.26723835 | 3.8857357 | 4.15625181 | 0.93491344 | 0.34983291 | 0.98715323 |
| miR-7060-3p | 0.26723835 | 3.8857357 | 4.15625181 | 0.93491344 | 0.34983291 | 0.98715323 |
| miR-883b-3p | 3.94986695 | -0.9784937 | 1.05312448 | -0.929134 | 0.35281967 | 0.98715323 |
| miR-542-5p | 0.26109 | 3.86028655 | 4.15707055 | 0.92860742 | 0.35309258 | 0.98715323 |
| let-7d-5p | 257.583474 | -0.2816649 | 0.30353014 | -0.9279634 | 0.35342655 | 0.98715323 |
| miR-7a-5p | 95.5240462 | 0.43155118 | 0.46553113 | 0.92700822 | 0.35392227 | 0.98715323 |
| miR-149-5p | 2.78691366 | 1.44320246 | 1.55940885 | 0.92548048 | 0.35471603 | 0.98715323 |
| miR-106b-3p | 71.543458 | 0.33682863 | 0.36774691 | 0.91592513 | 0.35970616 | 0.98715323 |
| miR-10b-3p | 5.87406405 | -1.035489 | 1.13486845 | -0.9124309 | 0.36154193 | 0.98715323 |
| miR-199a-3p | 14.4442038 | -0.6194082 | 0.68249549 | -0.9075639 | 0.3641087 | 0.98715323 |
| miR-7663-3p | 0.24442647 | 3.767261 | 4.16018382 | 0.90555157 | 0.36517325 | 0.98715323 |
| miR-7657-5p | 0.24442647 | 3.767261 | 4.16018382 | 0.90555157 | 0.36517325 | 0.98715323 |
| miR-363-3p | 0.24442647 | 3.767261 | 4.16018382 | 0.90555157 | 0.36517325 | 0.98715323 |
| miR-34c-5p | 1365.01758 | 0.47715076 | 0.52864505 | 0.90259195 | 0.3667425 | 0.98715323 |
| miR-361-5p | 7.51965324 | -0.8913609 | 0.99078397 | -0.8996521 | 0.36830543 | 0.98715323 |
| miR-743b-5p | 13.3933767 | 0.64689199 | 0.72519436 | 0.89202569 | 0.37237916 | 0.98715323 |
| miR-130a-3p | 344.123953 | 0.28503219 | 0.3199798 | 0.89078181 | 0.37304624 | 0.98715323 |
| miR-133a-3p | 13.2516827 | 0.69146506 | 0.77919488 | 0.88740965 | 0.37485839 | 0.98715323 |
| miR-449c-5p | 1.29789064 | -1.8130073 | 2.05056696 | -0.8841493 | 0.37661564 | 0.98715323 |
| miR-1933-3p | 0.38353075 | 3.67061919 | 4.16362823 | 0.88159148 | 0.37799776 | 0.98715323 |
| miR-139-3p | 0.76411505 | -2.2794859 | 2.58828913 | -0.8806921 | 0.37848448 | 0.98715323 |
| miR-471-3p | 5.65147346 | 0.89288204 | 1.01443634 | 0.88017552 | 0.37876423 | 0.98715323 |
| miR-582-3p | 0.6706355 | 3.21478278 | 3.70189739 | 0.86841488 | 0.38516725 | 0.98715323 |
| miR-7233-5p | 3.96187147 | -0.974381 | 1.12563725 | -0.8656261 | 0.38669525 | 0.98715323 |
| miR-204-5p | 276.046951 | -0.300792 | 0.34843744 | -0.8632597 | 0.38799472 | 0.98715323 |
| miR-30c-2-3p | 53.1794842 | 0.31715263 | 0.37020372 | 0.85669758 | 0.39161204 | 0.98715323 |
| miR-7234-5p | 0.84396123 | -2.58587 | 3.01977423 | -0.8563124 | 0.39182503 | 0.98715323 |
| miR-7068-5p | 2.11119042 | 1.52137424 | 1.77895707 | 0.85520571 | 0.39243728 | 0.98715323 |
| miR-376b-3p | 0.20042876 | 3.55577074 | 4.16801678 | 0.85310855 | 0.39359911 | 0.98715323 |
| miR-324-5p | 0.83313136 | -2.35915 | 2.78962626 | -0.8456868 | 0.39772751 | 0.98715323 |
| miR-7068-3p | 12.7803931 | 0.72377047 | 0.85775597 | 0.84379531 | 0.3987838 | 0.98715323 |
| miR-878-3p | 5.15579097 | 0.82291809 | 0.97532948 | 0.84373344 | 0.39881838 | 0.98715323 |
| miR-1a-3p | 8.02350501 | -0.8234218 | 0.97625022 | -0.8434536 | 0.39897481 | 0.98715323 |
| miR-132-3p | 1.31694205 | -1.7753067 | 2.1112566 | -0.8408768 | 0.40041695 | 0.98715323 |
| miR-484 | 5.68165104 | -0.9942474 | 1.18421073 | -0.8395865 | 0.40114027 | 0.98715323 |
| miR-181b-5p | 32.2043317 | -0.5905328 | 0.70370334 | -0.8391786 | 0.40136907 | 0.98715323 |
| miR-29b-3p | 0.79210748 | -2.4230399 | 2.88834837 | -0.8389015 | 0.40152456 | 0.98715323 |
| miR-10a-3p | 2.983913 | 1.40279386 | 1.69234705 | 0.82890437 | 0.40715852 | 0.98715323 |
| miR-7663-5p | 0.18198372 | 3.45426533 | 4.1721817 | 0.82792783 | 0.40771137 | 0.98715323 |
| miR-3960 | 0.18198372 | 3.45426533 | 4.1721817 | 0.82792783 | 0.40771137 | 0.98715323 |

| | | | | | | |
|---|---|---|---|---|---|---|
| miR-7036b-3p | 0.18198372 | 3.45426533 | 4.1721817 | 0.82792783 | 0.40771137 | 0.98715323 |
| miR-18a-5p | 6.18124308 | -0.8936063 | 1.08191939 | -0.8259454 | 0.40883509 | 0.98715323 |
| miR-30d-5p | 1760.5325 | 0.26304594 | 0.31946935 | 0.82338396 | 0.41028968 | 0.98715323 |
| miR-7236-3p | 0.34344628 | 3.38201336 | 4.1497781 | 0.81498656 | 0.41508001 | 0.98715323 |
| miR-186-5p | 151.5697 | 0.3035758 | 0.37507755 | 0.80936808 | 0.41830346 | 0.98715323 |
| miR-223-5p | 5.10379837 | 0.91793227 | 1.13491684 | 0.80881016 | 0.41862435 | 0.98715323 |
| miR-19a-3p | 11.4926159 | 0.7330737 | 0.91206524 | 0.80375139 | 0.42154057 | 0.98715323 |
| miR-92a-1-5p | 0.44965109 | 3.31682197 | 4.14848496 | 0.79952609 | 0.42398542 | 0.98715323 |
| miR-361-3p | 6.2232463 | 1.01689805 | 1.27316377 | 0.7987174 | 0.4244543 | 0.98715323 |
| miR-3084-3p | 3.69419631 | 1.20588845 | 1.52234214 | 0.79212709 | 0.42828658 | 0.98715323 |
| miR-7662-3p | 0.16295098 | 3.30166149 | 4.17898794 | 0.79006246 | 0.42949129 | 0.98715323 |
| miR-129-5p | 0.16295098 | 3.30166149 | 4.17898794 | 0.79006246 | 0.42949129 | 0.98715323 |
| miR-7012-5p | 0.16295098 | 3.30166149 | 4.17898794 | 0.79006246 | 0.42949129 | 0.98715323 |
| miR-669a-3p | 3.87813555 | -0.8963873 | 1.14419686 | -0.7834205 | 0.43338021 | 0.98715323 |
| miR-145a-3p | 15.9145371 | -0.6975896 | 0.89790509 | -0.7769079 | 0.4372131 | 0.98715323 |
| miR-200b-5p | 1.6344838 | -1.2320446 | 1.58600779 | -0.7768213 | 0.43726422 | 0.98715323 |
| miR-130b-3p | 37.360924 | 0.34787304 | 0.44850889 | 0.77562127 | 0.43797264 | 0.98715323 |
| miR-1927 | 0.67772998 | 2.57226253 | 3.32974839 | 0.77250958 | 0.43981267 | 0.98715323 |
| miR-344-3p | 2.01375743 | -3.1594398 | 4.09362001 | -0.7717961 | 0.44023522 | 0.98715323 |
| miR-221-3p | 75.1680938 | 0.43490652 | 0.56405434 | 0.77103657 | 0.44068525 | 0.98715323 |
| miR-34a-5p | 3.37585197 | 1.18821824 | 1.54405052 | 0.76954622 | 0.44156912 | 0.98715323 |
| miR-880-3p | 8.06722271 | -0.7594224 | 0.99818112 | -0.7608062 | 0.44677283 | 0.98715323 |
| miR-141-3p | 43.6745166 | 0.46186599 | 0.60790387 | 0.75976813 | 0.4473932 | 0.98715323 |
| miR-431-5p | 1.10369239 | -1.740489 | 2.30785236 | -0.7541596 | 0.4507534 | 0.98715323 |
| miR-134-5p | 1.98854484 | 1.38835665 | 1.85071349 | 0.75017373 | 0.45315008 | 0.98715323 |
| miR-25-3p | 1357.9244 | 0.22096331 | 0.29469104 | 0.74981347 | 0.45336705 | 0.98715323 |
| miR-465c-5p | 181.957168 | 0.2325004 | 0.31075507 | 0.74817895 | 0.45435223 | 0.98715323 |
| miR-182-3p | 0.24206941 | 3.10134686 | 4.18901102 | 0.740353 | 0.45908583 | 0.98715323 |
| miR-181a-2-3p | 7.44225008 | 0.76538265 | 1.06496141 | 0.7186952 | 0.47232874 | 0.98715323 |
| miR-324-3p | 0.12132248 | 3.01180373 | 4.19330715 | 0.71824067 | 0.47260891 | 0.98715323 |
| miR-3473e | 0.12132248 | 3.01180373 | 4.19330715 | 0.71824067 | 0.47260891 | 0.98715323 |
| miR-7026-3p | 0.12132248 | 3.01180373 | 4.19330715 | 0.71824067 | 0.47260891 | 0.98715323 |
| miR-16-1-3p | 0.76760196 | 2.29903423 | 3.20242384 | 0.71790442 | 0.47281622 | 0.98715323 |
| miR-127-5p | 0.54338521 | -2.7331352 | 3.84121271 | -0.7115292 | 0.47675634 | 0.98715323 |
| miR-34c-3p | 262.034111 | -0.2480935 | 0.35019176 | -0.7084503 | 0.47866563 | 0.98715323 |
| miR-199b-3p | 7.49644541 | -0.5791598 | 0.81949241 | -0.7067299 | 0.47973433 | 0.98715323 |
| miR-96-5p | 7.48396364 | 0.86863473 | 1.23078397 | 0.70575727 | 0.4803391 | 0.98715323 |
| miR-122-5p | 0.85811582 | -2.4204298 | 3.43925069 | -0.7037666 | 0.48157814 | 0.98715323 |
| miR-7210-3p | 14.8382315 | -0.4535158 | 0.64509115 | -0.703026 | 0.48203956 | 0.98715323 |
| miR-3963 | 0.39910525 | 2.47167602 | 3.51744975 | 0.70268979 | 0.48224909 | 0.98715323 |
| miR-7242-5p | 6.32206472 | -0.8620185 | 1.22748942 | -0.7022615 | 0.48251611 | 0.98715323 |
| miR-191-3p | 12.6768131 | 0.46953319 | 0.66989462 | 0.70090605 | 0.48336165 | 0.98715323 |
| miR-425-5p | 443.12411 | -0.2191162 | 0.31298749 | -0.7000798 | 0.48387748 | 0.98715323 |
| miR-455-3p | 0.56760543 | -2.6407645 | 3.78316778 | -0.69803 | 0.48515846 | 0.98715323 |
| miR-6516-3p | 1.25306265 | -1.923906 | 2.76909632 | -0.6947776 | 0.48719472 | 0.98715323 |
| miR-741-5p | 6.63699476 | 0.71675066 | 1.04139145 | 0.68826248 | 0.4912875 | 0.98715323 |

| | | | | | | |
|---|---|---|---|---|---|---|
| miR-7222-5p | 3.5716854 | -0.8654791 | 1.26688375 | -0.6831559 | 0.49450836 | 0.98715323 |
| miR-7223-5p | 1.38907708 | -1.8024979 | 2.64495592 | -0.681485 | 0.49556463 | 0.98715323 |
| miR-92a-3p | 1248.67192 | -0.217724 | 0.32098385 | -0.6783022 | 0.4975801 | 0.98715323 |
| miR-7073-5p | 0.78357044 | 2.34229212 | 3.50712666 | 0.66786642 | 0.50421887 | 0.98715323 |
| miR-666-3p | 1.05680737 | -1.8470102 | 2.77122062 | -0.666497 | 0.50509349 | 0.98715323 |
| miR-339-5p | 2.66677989 | -1.0577004 | 1.6025108 | -0.660027 | 0.50923652 | 0.98715323 |
| miR-3095-3p | 1.07401383 | -1.5457554 | 2.3504698 | -0.6576368 | 0.51077155 | 0.98715323 |
| miR-31-3p | 1.39114034 | 1.54045135 | 2.35062901 | 0.6553358 | 0.51225157 | 0.98715323 |
| miR-148a-3p | 4078.96285 | 0.20182901 | 0.3096516 | 0.65179385 | 0.51453417 | 0.98715323 |
| miR-5126 | 2.20662156 | 0.89397296 | 1.37692366 | 0.64925383 | 0.51617432 | 0.98715323 |
| miR-21a-5p | 686.905345 | 0.26026886 | 0.40173676 | 0.64785921 | 0.51707601 | 0.98715323 |
| miR-30c-5p | 1036.86164 | 0.16070447 | 0.24879874 | 0.64592156 | 0.51833016 | 0.98715323 |
| miR-30a-5p | 3490.53891 | 0.3201759 | 0.49573426 | 0.64586196 | 0.51836876 | 0.98715323 |
| miR-101b-3p | 13.5057654 | 0.44788186 | 0.69363913 | 0.64569866 | 0.51847453 | 0.98715323 |
| miR-222-5p | 1.11668659 | -1.4562072 | 2.25672055 | -0.6452758 | 0.51874847 | 0.98715323 |
| miR-708-3p | 0.62483669 | 2.17301229 | 3.38002851 | 0.64289762 | 0.52029053 | 0.98715323 |
| miR-511-5p | 0.75405694 | 1.94850418 | 3.07912886 | 0.63281021 | 0.52685759 | 0.98715323 |
| miR-301a-3p | 10.7435234 | 0.55419626 | 0.88435883 | 0.62666447 | 0.5308792 | 0.98715323 |
| miR-673-5p | 0.41086014 | -2.3960215 | 3.82595746 | -0.6262541 | 0.5311483 | 0.98715323 |
| miR-598-3p | 0.51310928 | -2.4339392 | 3.90367239 | -0.6234999 | 0.53295609 | 0.98715323 |
| miR-124-5p | 0.55117314 | 2.00813084 | 3.24041799 | 0.61971352 | 0.53544641 | 0.98715323 |
| miR-30d-3p | 4.34013111 | 0.69942457 | 1.13337802 | 0.61711499 | 0.53715888 | 0.98715323 |
| miR-20a-5p | 9.68228301 | 0.51408848 | 0.83800644 | 0.61346602 | 0.53956824 | 0.98715323 |
| miR-204-3p | 0.66535337 | 2.45937019 | 4.02029605 | 0.61173858 | 0.54071073 | 0.98715323 |
| miR-674-3p | 2.69871643 | -0.8936267 | 1.46090212 | -0.6116951 | 0.5407395 | 0.98715323 |
| miR-3072-3p | 0.89318298 | 1.99951487 | 3.28070937 | 0.60947638 | 0.54220872 | 0.98715323 |
| miR-16-2-3p | 12.7825411 | 0.39710676 | 0.65387578 | 0.60731223 | 0.54364373 | 0.98715323 |
| miR-181a-5p | 616.065004 | -0.2600537 | 0.43323206 | -0.6002643 | 0.54833011 | 0.98715323 |
| miR-1932 | 0.35891615 | -2.4572084 | 4.10183531 | -0.599051 | 0.5491389 | 0.98715323 |
| miR-3572-5p | 0.35891615 | -2.4572084 | 4.10183531 | -0.599051 | 0.5491389 | 0.98715323 |
| miR-135a-1-3p | 0.35891615 | -2.4572084 | 4.10183531 | -0.599051 | 0.5491389 | 0.98715323 |
| miR-181a-1-3p | 1.32505335 | -1.4811945 | 2.47699266 | -0.597981 | 0.54985262 | 0.98715323 |
| miR-674-5p | 0.79662291 | -1.867408 | 3.12463929 | -0.5976395 | 0.55008048 | 0.98715323 |
| miR-7060-5p | 1.17256607 | -1.3422686 | 2.25933582 | -0.5940988 | 0.55244606 | 0.98715323 |
| miR-152-3p | 3.69076093 | -0.8938655 | 1.51396002 | -0.5904155 | 0.55491211 | 0.98715323 |
| miR-99a-3p | 0.84195733 | -1.6697826 | 2.83647121 | -0.5886831 | 0.55607389 | 0.98715323 |
| miR-190b-5p | 22.2117223 | 0.35005506 | 0.59467898 | 0.58864542 | 0.55609916 | 0.98715323 |
| miR-872-3p | 3.0419223 | 0.82141159 | 1.39600534 | 0.58840147 | 0.55626285 | 0.98715323 |
| miR-499-5p | 0.9508207 | 1.51775679 | 2.58369302 | 0.58743697 | 0.55691027 | 0.98715323 |
| miR-8094 | 4.54244484 | 0.89615505 | 1.52741383 | 0.58671398 | 0.55739582 | 0.98715323 |
| miR-25-5p | 0.55995929 | 2.45935097 | 4.20595667 | 0.58473046 | 0.55872897 | 0.98715323 |
| miR-692 | 0.24139402 | 2.45933342 | 4.20595667 | 0.58472628 | 0.55873178 | 0.98715323 |
| miR-378c | 0.15231584 | 2.45932237 | 4.20595667 | 0.58472366 | 0.55873354 | 0.98715323 |
| miR-130a-5p | 0.23253444 | 2.45932237 | 4.20595667 | 0.58472366 | 0.55873354 | 0.98715323 |
| miR-7010-5p | 0.23253444 | 2.45932237 | 4.20595667 | 0.58472366 | 0.55873354 | 0.98715323 |
| miR-293-3p | 0.25461773 | 2.45929685 | 4.20595667 | 0.58471759 | 0.55873762 | 0.98715323 |

| | | | | | | |
|---|---|---|---|---|---|---|
| miR-7a-2-3p | 0.43155262 | 2.45927722 | 4.20595667 | 0.58471292 | 0.55874076 | 0.98715323 |
| miR-144-3p | 0.3307791 | 2.45926618 | 4.20595667 | 0.5847103 | 0.55874253 | 0.98715323 |
| miR-1930-5p | 0.38185085 | 2.45925557 | 4.20595667 | 0.58470778 | 0.55874422 | 0.98715323 |
| miR-935 | 0.27163409 | 2.45925342 | 4.20595667 | 0.58470726 | 0.55874457 | 0.98715323 |
| miR-1981-5p | 0.81363115 | 1.8633922 | 3.1902584 | 0.58408817 | 0.55916099 | 0.98715323 |
| miR-7a-1-3p | 1.40768575 | 1.05634819 | 1.8178317 | 0.5811034 | 0.56117077 | 0.98715323 |
| miR-465b-5p | 134.218272 | 0.30376728 | 0.52721123 | 0.57617756 | 0.56449517 | 0.98715323 |
| miR-34b-3p | 1915.43036 | -0.2032075 | 0.35380064 | -0.574356 | 0.56572691 | 0.98715323 |
| miR-3535 | 2527.66528 | 0.2845869 | 0.49647271 | 0.57321762 | 0.56649736 | 0.98715323 |
| miR-470-5p | 5636.05708 | 0.20249742 | 0.35571929 | 0.56926183 | 0.56917846 | 0.98715323 |
| miR-485-3p | 1.01910843 | -1.529224 | 2.70022319 | -0.5663324 | 0.57116782 | 0.98715323 |
| miR-7685-5p | 0.32628741 | -2.3211359 | 4.10393416 | -0.565588 | 0.57167388 | 0.98715323 |
| miR-1306-5p | 11.0575603 | -0.5055647 | 0.89400338 | -0.5655065 | 0.57172931 | 0.98715323 |
| miR-3079-5p | 0.28781655 | 2.31322511 | 4.13787194 | 0.55903739 | 0.57613621 | 0.98715323 |
| miR-1906 | 1.02708116 | 1.73807167 | 3.10908528 | 0.55902991 | 0.57614131 | 0.98715323 |
| miR-449a-5p | 84.0485271 | 0.37802396 | 0.6874218 | 0.54991559 | 0.58237727 | 0.98715323 |
| miR-30a-3p | 101.129552 | -0.1751964 | 0.32003792 | -0.5474237 | 0.58408765 | 0.98715323 |
| miR-7221-3p | 12.4474012 | -0.4941915 | 0.90673107 | -0.5450254 | 0.58573603 | 0.98715323 |
| miR-669p-5p | 2.94739795 | 0.54277224 | 0.99757668 | 0.54409074 | 0.58637903 | 0.98715323 |
| miR-143-5p | 1.38733169 | -1.1649092 | 2.14182957 | -0.5438851 | 0.58652053 | 0.98715323 |
| miR-206-3p | 1.83282319 | -1.0031603 | 1.84495012 | -0.543733 | 0.58662521 | 0.98715323 |
| miR-143-3p | 18524.9756 | -0.2561943 | 0.47349295 | -0.541073 | 0.58845727 | 0.98715323 |
| miR-7688-5p | 0.7748139 | -1.3702079 | 2.58792837 | -0.5294613 | 0.59648549 | 0.98715323 |
| miR-7658-3p | 0.65540536 | -2.1521892 | 4.07042124 | -0.5287387 | 0.59698673 | 0.98715323 |
| miR-467e-5p | 4.54618643 | -0.7065937 | 1.34558247 | -0.5251211 | 0.59949902 | 0.98715323 |
| miR-8112 | 2.01044339 | -0.9233113 | 1.76410356 | -0.5233884 | 0.60070401 | 0.98715323 |
| miR-1982-5p | 1.15280594 | -1.2991443 | 2.48597545 | -0.5225894 | 0.60126004 | 0.98715323 |
| miR-3082-3p | 0.48479468 | -1.8875655 | 3.61913458 | -0.5215516 | 0.60198256 | 0.98715323 |
| miR-23a-3p | 88.1362464 | -0.2233965 | 0.42882933 | -0.520945 | 0.60240506 | 0.98715323 |
| miR-483-3p | 3.44937194 | -0.7320894 | 1.41083399 | -0.5189054 | 0.6038267 | 0.98715323 |
| miR-22-3p | 1217.90203 | 0.18839096 | 0.36321835 | 0.51867139 | 0.60398992 | 0.98715323 |
| miR-410-3p | 4.64296563 | 0.51357112 | 0.99489099 | 0.51620844 | 0.60570884 | 0.98715323 |
| miR-743a-5p | 7.53008851 | 0.42987691 | 0.84005361 | 0.51172557 | 0.60884309 | 0.98715323 |
| miR-497a-5p | 2.74225735 | -0.9337931 | 1.82582173 | -0.5114372 | 0.60904498 | 0.98715323 |
| miR-676-5p | 2.23226047 | -0.8797593 | 1.72501175 | -0.5100019 | 0.61005012 | 0.98715323 |
| miR-3064-3p | 0.2773443 | -2.0894053 | 4.10798642 | -0.5086203 | 0.61101839 | 0.98715323 |
| miR-335-5p | 5.31225984 | -0.5209425 | 1.02492479 | -0.5082738 | 0.61126131 | 0.98715323 |
| miR-93-5p | 224.45357 | 0.16806773 | 0.33101371 | 0.50773646 | 0.61163817 | 0.98715323 |
| miR-202-3p | 0.34051606 | -2.0673916 | 4.10840566 | -0.5032102 | 0.61481651 | 0.98715323 |
| miR-365-2-5p | 0.36655662 | -1.9865503 | 3.98588604 | -0.4983962 | 0.61820484 | 0.98715323 |
| miR-28a-5p | 6.70795392 | -0.4413549 | 0.89062055 | -0.4955588 | 0.62020572 | 0.98715323 |
| miR-668-3p | 0.98864856 | -1.3930382 | 2.82441754 | -0.4932126 | 0.62186241 | 0.98715323 |
| miR-150-3p | 0.64869789 | -1.8230926 | 3.72049613 | -0.4900133 | 0.62412449 | 0.98715323 |
| miR-7b-5p | 1.36611929 | 0.98896613 | 2.02367152 | 0.48869894 | 0.62505485 | 0.98715323 |
| miR-199b-5p | 0.26929608 | -2.0035842 | 4.10965607 | -0.4875309 | 0.62588217 | 0.98715323 |
| miR-7009-5p | 0.26102993 | -2.0030607 | 4.10966656 | -0.4874022 | 0.62597331 | 0.98715323 |

| | | | | | | |
|---|---|---|---|---|---|---|
| miR-16-5p | 3180.11971 | 0.1505079 | 0.3127471 | 0.48124474 | 0.63034256 | 0.98715323 |
| miR-181c-3p | 0.7050816 | 1.43729043 | 3.00510056 | 0.47828364 | 0.63244834 | 0.98715323 |
| miR-369-3p | 0.54660629 | -1.6665207 | 3.48692117 | -0.4779347 | 0.63269667 | 0.98715323 |
| miR-195a-3p | 1.16652442 | -1.2380418 | 2.60024429 | -0.4761252 | 0.6339852 | 0.98715323 |
| miR-200b-3p | 43.7631029 | 0.23463143 | 0.49530065 | 0.47371517 | 0.63570304 | 0.98715323 |
| miR-28a-3p | 11.530027 | 0.36726336 | 0.78699048 | 0.46666812 | 0.64073734 | 0.98715323 |
| miR-362-5p | 0.24471556 | -1.9112273 | 4.11156444 | -0.4648419 | 0.6420447 | 0.98715323 |
| miR-93-3p | 0.72809689 | 1.51499408 | 3.26668645 | 0.46377089 | 0.6428119 | 0.98715323 |
| miR-151-5p | 403.539251 | 0.1509749 | 0.32585738 | 0.46331588 | 0.64313796 | 0.98715323 |
| miR-23b-3p | 63.4340343 | -0.1695595 | 0.36754768 | -0.4613264 | 0.64456443 | 0.98715323 |
| miR-362-3p | 0.23916252 | -1.8770916 | 4.11230046 | -0.4564578 | 0.64806082 | 0.98715323 |
| miR-15b-3p | 1.2431204 | 1.10377239 | 2.42728362 | 0.45473565 | 0.64929943 | 0.98715323 |
| let-7i-5p | 646.879343 | 0.17584701 | 0.38804866 | 0.45315711 | 0.65043561 | 0.98715323 |
| miR-5113 | 0.69305464 | 1.7380513 | 3.90887065 | 0.44464283 | 0.65657789 | 0.98715323 |
| miR-1839-3p | 0.33311346 | 1.84758686 | 4.15677631 | 0.44447589 | 0.65669855 | 0.98715323 |
| miR-703 | 0.59496855 | -1.5005185 | 3.37689789 | -0.4443482 | 0.65679085 | 0.98715323 |
| miR-664-3p | 13.0924885 | 0.30539546 | 0.69207195 | 0.44127703 | 0.65901245 | 0.98715323 |
| let-7a-1-3p | 1.18944819 | -0.9463446 | 2.15023174 | -0.4401128 | 0.65985539 | 0.98715323 |
| let-7c-2-3p | 1.18944819 | -0.9463446 | 2.15023174 | -0.4401128 | 0.65985539 | 0.98715323 |
| miR-98-5p | 115.641042 | 0.20103243 | 0.45766296 | 0.43925868 | 0.66047411 | 0.98715323 |
| miR-141-5p | 1.32835685 | 1.21669258 | 2.78225942 | 0.43730379 | 0.66189105 | 0.98715323 |
| miR-5122 | 1.07048901 | -1.0817728 | 2.48880374 | -0.4346557 | 0.66381233 | 0.98715323 |
| miR-7240-5p | 0.67367875 | 1.73794666 | 4.02178366 | 0.4321333 | 0.66564453 | 0.98715323 |
| miR-329-5p | 0.23145904 | -1.776064 | 4.11458069 | -0.4316513 | 0.66599488 | 0.98715323 |
| miR-29a-3p | 39.0011119 | 0.28011726 | 0.65053076 | 0.43059803 | 0.66676067 | 0.98715323 |
| miR-3070-5p | 0.64518778 | 1.73795853 | 4.08478342 | 0.4254714 | 0.67049307 | 0.98715323 |
| miR-301b-3p | 0.78326378 | 1.28602676 | 3.03623082 | 0.42356028 | 0.67188653 | 0.98715323 |
| miR-3077-3p | 0.2189207 | -1.7394656 | 4.11544578 | -0.4226676 | 0.6725378 | 0.98715323 |
| miR-7241-5p | 0.58243179 | -1.7394261 | 4.11544623 | -0.422658 | 0.67254484 | 0.98715323 |
| miR-190a-5p | 0.225906 | -1.7368515 | 4.11550839 | -0.422026 | 0.67300605 | 0.98715323 |
| miR-744-3p | 0.37696495 | -1.7368391 | 4.11550853 | -0.422023 | 0.67300826 | 0.98715323 |
| miR-19b-3p | 42.3812045 | 0.21775063 | 0.519303 | 0.41931326 | 0.67498721 | 0.98715323 |
| miR-345-5p | 2.83472097 | -0.5638283 | 1.34524893 | -0.4191256 | 0.67512432 | 0.98715323 |
| miR-409-5p | 0.29204608 | -1.7078486 | 4.11621031 | -0.414908 | 0.67820926 | 0.98715323 |
| miR-27b-5p | 1.90329209 | 0.86787334 | 2.09440404 | 0.41437723 | 0.67859786 | 0.98715323 |
| miR-340-3p | 3.66941818 | 0.55556056 | 1.34134051 | 0.41418309 | 0.67874002 | 0.98715323 |
| miR-7218-3p | 9.6558705 | -0.409936 | 0.99096567 | -0.4136732 | 0.67911344 | 0.98715323 |
| miR-700-3p | 0.07697561 | 1.73805425 | 4.20595142 | 0.41323688 | 0.67943307 | 0.98715323 |
| miR-1936 | 0.75529474 | 1.73804702 | 4.20595667 | 0.41323465 | 0.67943471 | 0.98715323 |
| miR-7006-3p | 0.60423579 | 1.73804122 | 4.20595667 | 0.41323327 | 0.67943572 | 0.98715323 |
| miR-7017-3p | 0.54526664 | 1.73803926 | 4.20595667 | 0.4132328 | 0.67943606 | 0.98715323 |
| miR-5620-5p | 0.54526664 | 1.73803926 | 4.20595667 | 0.4132328 | 0.67943606 | 0.98715323 |
| miR-7652-3p | 0.48387344 | 1.73803573 | 4.20595667 | 0.41323196 | 0.67943667 | 0.98715323 |
| miR-433-3p | 0.45438887 | 1.73803463 | 4.20595667 | 0.4132317 | 0.67943686 | 0.98715323 |
| miR-210-5p | 0.43827431 | 1.73803376 | 4.20595667 | 0.41323149 | 0.67943702 | 0.98715323 |
| miR-7238-5p | 0.45317684 | 1.7380337 | 4.20595667 | 0.41323148 | 0.67943703 | 0.98715323 |

| | | | | | | |
|---|---|---|---|---|---|---|
| miR-328-5p | 0.45317684 | 1.7380337 | 4.20595667 | 0.41323148 | 0.67943703 | 0.98715323 |
| miR-6990-5p | 0.42753127 | 1.73803317 | 4.20595667 | 0.41323135 | 0.67943712 | 0.98715323 |
| miR-183-3p | 0.42753127 | 1.73803317 | 4.20595667 | 0.41323135 | 0.67943712 | 0.98715323 |
| miR-3475-3p | 0.42753127 | 1.73803317 | 4.20595667 | 0.41323135 | 0.67943712 | 0.98715323 |
| miR-5125 | 0.39979633 | 1.73803156 | 4.20595667 | 0.41323097 | 0.6794374 | 0.98715323 |
| miR-7070-5p | 0.39979633 | 1.73803156 | 4.20595667 | 0.41323097 | 0.6794374 | 0.98715323 |
| miR-7073-3p | 0.39979633 | 1.73803156 | 4.20595667 | 0.41323097 | 0.6794374 | 0.98715323 |
| miR-6922-3p | 0.36351109 | 1.73802897 | 4.20595667 | 0.41323036 | 0.67943785 | 0.98715323 |
| miR-6340 | 0.36351109 | 1.73802897 | 4.20595667 | 0.41323036 | 0.67943785 | 0.98715323 |
| miR-3092-3p | 0.36351109 | 1.73802897 | 4.20595667 | 0.41323036 | 0.67943785 | 0.98715323 |
| miR-455-5p | 0.35813958 | 1.73802862 | 4.20595667 | 0.41323027 | 0.67943791 | 0.98715323 |
| miR-218-1-3p | 0.31983706 | 1.73802594 | 4.20595667 | 0.41322963 | 0.67943838 | 0.98715323 |
| miR-3103-5p | 0.27263332 | 1.7380221 | 4.20595667 | 0.41322872 | 0.67943905 | 0.98715323 |
| miR-504-5p | 0.27263332 | 1.7380221 | 4.20595667 | 0.41322872 | 0.67943905 | 0.98715323 |
| miR-7090-5p | 0.25651876 | 1.73802093 | 4.20595667 | 0.41322844 | 0.67943925 | 0.98715323 |
| miR-196a-1-3p | 0.3021179 | 1.73802073 | 4.20595667 | 0.4132284 | 0.67943929 | 0.98715323 |
| miR-673-3p | 0.3021179 | 1.73802073 | 4.20595667 | 0.4132284 | 0.67943929 | 0.98715323 |
| miR-719 | 0.3021179 | 1.73802073 | 4.20595667 | 0.4132284 | 0.67943929 | 0.98715323 |
| miR-546 | 0.2398778 | 1.73801966 | 4.20595667 | 0.41322814 | 0.67943947 | 0.98715323 |
| miR-134-3p | 0.2398778 | 1.73801966 | 4.20595667 | 0.41322814 | 0.67943947 | 0.98715323 |
| miR-412-5p | 0.2398778 | 1.73801966 | 4.20595667 | 0.41322814 | 0.67943947 | 0.98715323 |
| miR-6992-5p | 0.2398778 | 1.73801966 | 4.20595667 | 0.41322814 | 0.67943947 | 0.98715323 |
| miR-1960 | 0.2398778 | 1.73801966 | 4.20595667 | 0.41322814 | 0.67943947 | 0.98715323 |
| miR-7028-5p | 0.2398778 | 1.73801966 | 4.20595667 | 0.41322814 | 0.67943947 | 0.98715323 |
| miR-330-3p | 0.2398778 | 1.73801966 | 4.20595667 | 0.41322814 | 0.67943947 | 0.98715323 |
| miR-138-5p | 0.2398778 | 1.73801966 | 4.20595667 | 0.41322814 | 0.67943947 | 0.98715323 |
| miR-3109-3p | 0.2398778 | 1.73801966 | 4.20595667 | 0.41322814 | 0.67943947 | 0.98715323 |
| miR-7075-5p | 0.18175555 | 1.73801373 | 4.20595667 | 0.41322673 | 0.6794405 | 0.98715323 |
| miR-201-5p | 0.18175555 | 1.73801373 | 4.20595667 | 0.41322673 | 0.6794405 | 0.98715323 |
| miR-5119 | 0.17101251 | 1.73801263 | 4.20595667 | 0.41322647 | 0.6794407 | 0.98715323 |
| miR-7648-5p | 0.17101251 | 1.73801263 | 4.20595667 | 0.41322647 | 0.6794407 | 0.98715323 |
| miR-877-3p | 0.17101251 | 1.73801263 | 4.20595667 | 0.41322647 | 0.6794407 | 0.98715323 |
| miR-193a-5p | 0.15991853 | 1.73801142 | 4.20595667 | 0.41322618 | 0.67944091 | 0.98715323 |
| miR-665-3p | 0.15991853 | 1.73801142 | 4.20595667 | 0.41322618 | 0.67944091 | 0.98715323 |
| miR-874-5p | 0.15991853 | 1.73801142 | 4.20595667 | 0.41322618 | 0.67944091 | 0.98715323 |
| miR-3090-3p | 0.15991853 | 1.73801142 | 4.20595667 | 0.41322618 | 0.67944091 | 0.98715323 |
| miR-7007-3p | 0.15991853 | 1.73801142 | 4.20595667 | 0.41322618 | 0.67944091 | 0.98715323 |
| miR-8111 | 0.53598151 | 1.73800764 | 4.20595667 | 0.41322529 | 0.67944156 | 0.98715323 |
| miR-3064-5p | 1.90725672 | 1.73799103 | 4.20595667 | 0.41322134 | 0.67944446 | 0.98715323 |
| miR-6541 | 0.29020579 | 1.73798821 | 4.20595667 | 0.41322066 | 0.67944495 | 0.98715323 |
| miR-6971-3p | 0.21444515 | 1.73797485 | 4.20595667 | 0.41321749 | 0.67944728 | 0.98715323 |
| miR-7226-3p | 0.1267724 | 1.73794472 | 4.20595667 | 0.41321033 | 0.67945252 | 0.98715323 |
| miR-8103 | 0.1901586 | 1.73793142 | 4.20595667 | 0.41320716 | 0.67945484 | 0.98715323 |
| miR-539-5p | 0.1901586 | 1.73793142 | 4.20595667 | 0.41320716 | 0.67945484 | 0.98715323 |
| miR-200c-5p | 0.1901586 | 1.73793142 | 4.20595667 | 0.41320716 | 0.67945484 | 0.98715323 |
| miR-6915-3p | 0.21690048 | 1.73792794 | 4.20595667 | 0.41320634 | 0.67945545 | 0.98715323 |

| | | | | | | |
|---|---|---|---|---|---|---|
| miR-323-3p | 0.21690048 | 1.73792794 | 4.20595667 | 0.41320634 | 0.67945545 | 0.98715323 |
| miR-7117-3p | 0.21690048 | 1.73792794 | 4.20595667 | 0.41320634 | 0.67945545 | 0.98715323 |
| miR-3085-5p | 0.21690048 | 1.73792794 | 4.20595667 | 0.41320634 | 0.67945545 | 0.98715323 |
| miR-410-5p | 0.2535448 | 1.73792057 | 4.20595667 | 0.41320458 | 0.67945673 | 0.98715323 |
| miR-6942-5p | 0.2535448 | 1.73792057 | 4.20595667 | 0.41320458 | 0.67945673 | 0.98715323 |
| miR-3084-5p | 0.10317053 | 1.7379155 | 4.20594845 | 0.41320419 | 0.67945702 | 0.98715323 |
| miR-6905-3p | 0.30780966 | 1.73791355 | 4.20595667 | 0.41320291 | 0.67945795 | 0.98715323 |
| miR-7036b-5p | 0.32535072 | 1.73791214 | 4.20595667 | 0.41320258 | 0.6794582 | 0.98715323 |
| miR-26a-1-3p | 0.46171449 | 1.73789807 | 4.20595667 | 0.41319923 | 0.67946065 | 0.98715323 |
| miR-1191b-5p | 0.35921461 | 1.73784467 | 4.20594735 | 0.41318745 | 0.67946928 | 0.98715323 |
| miR-700-5p | 0.74366835 | 1.18232568 | 2.89185888 | 0.40884626 | 0.68265249 | 0.98715323 |
| miR-148a-5p | 40.6169889 | -0.1829897 | 0.44859963 | -0.4079131 | 0.68333746 | 0.98715323 |
| miR-224-5p | 4.18875103 | -0.3918504 | 0.96796623 | -0.4048182 | 0.68561114 | 0.98715323 |
| miR-146b-5p | 843.96163 | -0.1156989 | 0.28834833 | -0.4012471 | 0.68823821 | 0.98715323 |
| miR-669m-3p | 0.20749561 | -1.6514819 | 4.11761472 | -0.4010773 | 0.68836319 | 0.98715323 |
| miR-652-3p | 4.68064068 | 0.4569867 | 1.14094275 | 0.4005343 | 0.68876303 | 0.98715323 |
| miR-1894-3p | 0.2031438 | -1.6353278 | 4.11802708 | -0.3971144 | 0.69128312 | 0.98715323 |
| miR-505-5p | 0.83056638 | -1.2274465 | 3.0940509 | -0.3967118 | 0.69157999 | 0.98715323 |
| miR-653-5p | 0.48559004 | 1.61407381 | 4.08148592 | 0.3954623 | 0.69250174 | 0.98715323 |
| miR-30c-1-3p | 4.35338289 | 0.44565043 | 1.15760499 | 0.38497626 | 0.70025501 | 0.98715323 |
| miR-883a-3p | 19.3266994 | -0.2202241 | 0.57323977 | -0.3841745 | 0.7008491 | 0.98715323 |
| miR-296-3p | 0.32202775 | -1.5805237 | 4.11946018 | -0.3836725 | 0.70122117 | 0.98715323 |
| miR-6988-5p | 0.19525535 | -1.5804839 | 4.11946086 | -0.3836628 | 0.70122839 | 0.98715323 |
| miR-322-5p | 0.79284218 | -1.102859 | 2.9386864 | -0.3752898 | 0.70744497 | 0.98715323 |
| miR-26b-3p | 0.95444117 | -0.9697304 | 2.58398964 | -0.3752841 | 0.70744915 | 0.98715323 |
| miR-540-5p | 0.46396931 | 1.54197106 | 4.12190082 | 0.37409223 | 0.7083357 | 0.98715323 |
| miR-208b-3p | 0.16471256 | 1.55776572 | 4.17117316 | 0.37345985 | 0.70880622 | 0.98715323 |
| miR-450a-5p | 0.24706883 | 1.5419744 | 4.12996395 | 0.37336268 | 0.70887853 | 0.98715323 |
| miR-7221-5p | 0.75652567 | -1.1748579 | 3.14677191 | -0.3733534 | 0.70888545 | 0.98715323 |
| miR-1948-5p | 0.19465085 | -1.5283031 | 4.12087487 | -0.3708686 | 0.7107354 | 0.98715323 |
| miR-744-5p | 17.7411866 | -0.2796814 | 0.75548367 | -0.3702018 | 0.71123213 | 0.98715323 |
| miR-1964-3p | 0.68235136 | -1.3445765 | 3.63721052 | -0.3696724 | 0.71162658 | 0.98715323 |
| miR-7214-3p | 14.3740518 | -0.2782066 | 0.75302379 | -0.3694526 | 0.71179038 | 0.98715323 |
| miR-195a-5p | 150.334642 | -0.1119118 | 0.3033038 | -0.368976 | 0.71214562 | 0.98715323 |
| miR-666-5p | 1.02168433 | -0.9388168 | 2.56028225 | -0.3666849 | 0.71385408 | 0.98715323 |
| miR-467c-5p | 3.46959286 | 0.50140463 | 1.38567339 | 0.36184907 | 0.71746482 | 0.98715323 |
| miR-669o-5p | 4.38164049 | -0.491001 | 1.36683435 | -0.359225 | 0.71942679 | 0.98715323 |
| miR-7675-3p | 0.17945807 | -1.4709657 | 4.12248671 | -0.3568152 | 0.72123019 | 0.98715323 |
| miR-7224-3p | 0.1867624 | -1.4673463 | 4.12259055 | -0.3559282 | 0.72189432 | 0.98715323 |
| miR-211-5p | 2.84205704 | 0.63478952 | 1.78689618 | 0.35524701 | 0.72240455 | 0.98715323 |
| miR-5107-5p | 1.60470767 | 0.70692421 | 2.04623297 | 0.34547592 | 0.72973661 | 0.98715323 |
| miR-3066-3p | 0.46849696 | 1.39687425 | 4.09248178 | 0.34132693 | 0.73285748 | 0.98715323 |
| miR-92b-3p | 72.4786355 | -0.1636321 | 0.4805465 | -0.3405125 | 0.7334706 | 0.98715323 |
| miR-380-3p | 0.34172456 | 1.39695294 | 4.10303739 | 0.340468 | 0.73350412 | 0.98715323 |
| miR-344b-3p | 0.18251593 | -1.4040246 | 4.12444847 | -0.3404151 | 0.73354394 | 0.98715323 |
| miR-3088-3p | 0.35352844 | -1.4039834 | 4.12444933 | -0.340405 | 0.73355152 | 0.98715323 |

| | | | | | | |
|---|---|---|---|---|---|---|
| miR-326-3p | 0.9228141 | -1.0765369 | 3.16740733 | -0.3398795 | 0.73394726 | 0.98715323 |
| miR-449a-3p | 0.17115386 | -1.4015137 | 4.12452378 | -0.3398001 | 0.73400706 | 0.98715323 |
| let-7b-3p | 1.47839757 | 0.80372774 | 2.37814892 | 0.33796359 | 0.73539063 | 0.98715323 |
| miR-743a-3p | 10.6800043 | 0.21529953 | 0.63886337 | 0.33700402 | 0.73611387 | 0.98715323 |
| miR-300-3p | 2.81409544 | -0.6130724 | 1.82456484 | -0.3360102 | 0.73686316 | 0.98715323 |
| miR-466d-3p | 0.63912589 | -1.0101278 | 3.01206701 | -0.3353603 | 0.73735328 | 0.98715323 |
| miR-7229-3p | 9.20503501 | -0.2634965 | 0.787528 | -0.3345868 | 0.73793679 | 0.98715323 |
| miR-331-3p | 0.17027165 | -1.379515 | 4.12518898 | -0.3344126 | 0.73806826 | 0.98715323 |
| miR-1948-3p | 0.17027165 | -1.379515 | 4.12518898 | -0.3344126 | 0.73806826 | 0.98715323 |
| miR-1298-5p | 0.17027165 | -1.379515 | 4.12518898 | -0.3344126 | 0.73806826 | 0.98715323 |
| miR-3066-5p | 1.6247547 | -0.5675749 | 1.71544372 | -0.3308618 | 0.74074884 | 0.98715323 |
| let-7f-1-3p | 0.16931384 | -1.3477832 | 4.12616589 | -0.326643 | 0.74393791 | 0.98715323 |
| miR-7215-5p | 0.27346293 | -1.3361755 | 4.12652871 | -0.3238013 | 0.74608845 | 0.98715323 |
| miR-200c-3p | 43.9662858 | -0.1426394 | 0.44389204 | -0.3213381 | 0.74795418 | 0.98715323 |
| miR-140-3p | 43.1775835 | 0.15389612 | 0.48097115 | 0.31996955 | 0.74899141 | 0.98715323 |
| miR-330-5p | 1.08453659 | 0.87697078 | 2.76655259 | 0.31699046 | 0.75125084 | 0.98715323 |
| miR-375-3p | 1526.53454 | -0.104327 | 0.33197626 | -0.3142606 | 0.75332315 | 0.98715323 |
| miR-669f-3p | 0.16506736 | -1.277274 | 4.1284121 | -0.3093863 | 0.75702773 | 0.98715323 |
| miR-5100 | 0.5482102 | 1.0632664 | 3.44851899 | 0.30832552 | 0.75783465 | 0.98715323 |
| miR-10b-5p | 392080.628 | -0.09612 | 0.31385685 | -0.3062541 | 0.75941115 | 0.98715323 |
| miR-27a-5p | 0.67508012 | 1.01665087 | 3.37051971 | 0.3016303 | 0.76293391 | 0.98715323 |
| miR-1247-5p | 11.0116551 | 0.27773846 | 0.92554831 | 0.30007992 | 0.7641162 | 0.98715323 |
| miR-421-3p | 1.79965379 | 0.57222568 | 1.91384248 | 0.29899309 | 0.76494532 | 0.98715323 |
| miR-7025-5p | 0.15186527 | -1.2221106 | 4.13024452 | -0.295893 | 0.76731176 | 0.98715323 |
| miR-32-3p | 0.56166506 | 1.0165978 | 3.55558618 | 0.28591567 | 0.77494271 | 0.98715323 |
| miR-193b-3p | 5.28269816 | 0.3145675 | 1.11880714 | 0.2811633 | 0.77858515 | 0.98715323 |
| miR-8097 | 0.64813936 | 1.05085595 | 3.74565416 | 0.28055339 | 0.77905297 | 0.98715323 |
| miR-6970-5p | 0.20933333 | -1.1590334 | 4.13242425 | -0.280473 | 0.77911465 | 0.98715323 |
| miR-6981-5p | 0.14594713 | -1.1590092 | 4.1324248 | -0.2804671 | 0.77911917 | 0.98715323 |
| miR-8114 | 4.63926344 | -0.3725438 | 1.33566221 | -0.2789207 | 0.78030568 | 0.98715323 |
| miR-6946-3p | 0.31863131 | -1.1434048 | 4.13297777 | -0.276654 | 0.7820458 | 0.98715323 |
| miR-874-3p | 1.12678457 | -0.5923081 | 2.15459837 | -0.2749042 | 0.78338987 | 0.98715323 |
| miR-466b-3p | 3.80569996 | -0.2978428 | 1.08806349 | -0.2737366 | 0.78428708 | 0.98715323 |
| miR-6960-5p | 0.67675021 | -0.9065582 | 3.35662855 | -0.27008 | 0.78709871 | 0.98715323 |
| miR-6390 | 0.14068817 | -1.1086293 | 4.13423237 | -0.2681584 | 0.78857736 | 0.98715323 |
| miR-877-5p | 0.73376402 | -0.8310429 | 3.17445461 | -0.2617908 | 0.79348276 | 0.98715323 |
| miR-883b-5p | 3.33547215 | -0.3453283 | 1.32625655 | -0.2603782 | 0.79457205 | 0.98715323 |
| miR-615-3p | 2.63498668 | -0.3079906 | 1.18392589 | -0.2601435 | 0.79475311 | 0.98715323 |
| miR-7232-5p | 0.65927838 | 0.77715398 | 2.99675935 | 0.25933146 | 0.79537951 | 0.98715323 |
| let-7i-3p | 1.70454211 | 0.59959006 | 2.36200567 | 0.25384785 | 0.7996131 | 0.98715323 |
| miR-483-5p | 1.26550568 | 0.56958606 | 2.27169496 | 0.25073175 | 0.80202151 | 0.98715323 |
| miR-322-3p | 0.64858961 | -0.7308911 | 2.91882158 | -0.2504062 | 0.80227322 | 0.98715323 |
| miR-17-5p | 2.20817842 | 0.33359413 | 1.3393311 | 0.24907518 | 0.80330263 | 0.98715323 |
| miR-15a-5p | 15.6104589 | 0.20224775 | 0.81305025 | 0.24875185 | 0.80355274 | 0.98715323 |
| miR-341-3p | 0.45586568 | -1.0212476 | 4.13751539 | -0.2468263 | 0.80504266 | 0.98715323 |
| miR-5128 | 0.13051496 | -1.0211765 | 4.13751716 | -0.246809 | 0.80505604 | 0.98715323 |

| | | | | | | |
|---|---|---|---|---|---|---|
| miR-342-3p | 86.1737741 | 0.08418592 | 0.34274442 | 0.24562304 | 0.80597406 | 0.98715323 |
| miR-135a-5p | 0.43928871 | 0.92635358 | 3.77434585 | 0.24543421 | 0.80612025 | 0.98715323 |
| miR-9769-3p | 0.6205535 | 1.01665799 | 4.14757659 | 0.24512097 | 0.80636277 | 0.98715323 |
| miR-411-5p | 2.37450332 | 0.3499566 | 1.44057114 | 0.24292906 | 0.80806035 | 0.98715323 |
| miR-26a-2-3p | 0.31192578 | -1.0027119 | 4.13823487 | -0.2423042 | 0.80854442 | 0.98715323 |
| miR-431-3p | 0.40280355 | -1.0027038 | 4.13823502 | -0.2423023 | 0.80854594 | 0.98715323 |
| miR-28c | 0.12476461 | 1.01670749 | 4.20595134 | 0.24173068 | 0.80898885 | 0.98715323 |
| miR-702-3p | 0.42434974 | 1.01664442 | 4.20594937 | 0.2417158 | 0.80900038 | 0.98715323 |
| miR-382-5p | 0.2782138 | 1.01663515 | 4.20594926 | 0.2417136 | 0.80900209 | 0.98715323 |
| miR-377-5p | 0.1016543 | 1.01661168 | 4.20594897 | 0.24170804 | 0.8090064 | 0.98715323 |
| miR-466c-5p | 0.16928888 | 1.01656688 | 4.20594833 | 0.24169742 | 0.80901463 | 0.98715323 |
| miR-3100-5p | 0.16928888 | 1.01656688 | 4.20594833 | 0.24169742 | 0.80901463 | 0.98715323 |
| miR-450b-5p | 0.13014528 | 1.01656678 | 4.20594833 | 0.2416974 | 0.80901464 | 0.98715323 |
| miR-294-3p | 0.92402984 | -0.6598321 | 2.73688597 | -0.2410886 | 0.80948642 | 0.98715323 |
| miR-345-3p | 5.44312551 | 0.35494339 | 1.48069384 | 0.23971424 | 0.8105518 | 0.98715323 |
| miR-495-3p | 0.13381221 | -0.9913836 | 4.13868038 | -0.239541 | 0.81068612 | 0.98715323 |
| miR-320-3p | 35.3779332 | 0.13268034 | 0.56328556 | 0.23554721 | 0.81378404 | 0.98715323 |
| miR-423-5p | 599.714497 | 0.08705019 | 0.37062053 | 0.23487686 | 0.81430431 | 0.98715323 |
| miR-3081-3p | 2.78968103 | 0.41931451 | 1.79460416 | 0.23365293 | 0.81525444 | 0.98715323 |
| miR-664-5p | 2.60168405 | 0.3174085 | 1.35929894 | 0.23350897 | 0.8153662 | 0.98715323 |
| miR-466c-3p | 1.50123109 | -0.3534901 | 1.51763134 | -0.2329222 | 0.81582178 | 0.98715323 |
| miR-7222-3p | 2.3458522 | 0.33944251 | 1.52410585 | 0.22271584 | 0.82375667 | 0.98715323 |
| let-7f-5p | 17411.4308 | 0.08581787 | 0.38657272 | 0.22199671 | 0.82431644 | 0.98715323 |
| miR-7118-3p | 0.56316804 | 0.88692724 | 4.07078169 | 0.2178764 | 0.82752542 | 0.98715323 |
| miR-10a-5p | 171033.381 | 0.07192002 | 0.33140677 | 0.21701433 | 0.82819718 | 0.98715323 |
| miR-106b-5p | 0.12162261 | -0.8984864 | 4.14245807 | -0.2168969 | 0.82828868 | 0.98715323 |
| miR-1955-5p | 0.39425593 | -0.8984524 | 4.14245877 | -0.2168887 | 0.8282951 | 0.98715323 |
| miR-350-5p | 0.29106583 | 0.88947416 | 4.17831271 | 0.21287879 | 0.8314215 | 0.98715323 |
| miR-742-5p | 0.86779662 | -0.4946545 | 2.34327433 | -0.2110954 | 0.83281281 | 0.98715323 |
| miR-329-3p | 0.11899313 | -0.8709554 | 4.14341242 | -0.2102024 | 0.83350968 | 0.98715323 |
| miR-669m-5p | 0.3916325 | -0.8612739 | 4.11940432 | -0.2090773 | 0.83438791 | 0.98715323 |
| miR-297c-3p | 0.50000017 | -0.5374332 | 2.59570497 | -0.2070471 | 0.83597307 | 0.98715323 |
| miR-297b-3p | 0.50000017 | -0.5374332 | 2.59570497 | -0.2070471 | 0.83597307 | 0.98715323 |
| miR-883a-5p | 2.23124848 | 0.38764521 | 1.90405723 | 0.20358905 | 0.83867465 | 0.98715323 |
| miR-124-3p | 2.25959209 | -0.2895976 | 1.4403759 | -0.201057 | 0.84065403 | 0.98715323 |
| miR-27b-3p | 459.865937 | -0.0562771 | 0.29532298 | -0.1905612 | 0.84886939 | 0.98715323 |
| miR-203-3p | 21.1211209 | -0.123893 | 0.65651558 | -0.188713 | 0.85031776 | 0.98715323 |
| miR-671-3p | 18.2438899 | -0.1103888 | 0.58897978 | -0.1874237 | 0.85132844 | 0.98715323 |
| miR-7015-3p | 0.20810263 | 0.76736183 | 4.14022027 | 0.18534324 | 0.85295983 | 0.98715323 |
| miR-6516-5p | 0.21692543 | -0.7673655 | 4.14591245 | -0.1850897 | 0.85315872 | 0.98715323 |
| miR-5101 | 0.10847519 | -0.7673437 | 4.14591279 | -0.1850844 | 0.85316286 | 0.98715323 |
| miR-7118-5p | 0.10847519 | -0.7673437 | 4.14591279 | -0.1850844 | 0.85316286 | 0.98715323 |
| miR-6986-5p | 0.10847519 | -0.7673437 | 4.14591279 | -0.1850844 | 0.85316286 | 0.98715323 |
| miR-7013-5p | 0.10847519 | -0.7673437 | 4.14591279 | -0.1850844 | 0.85316286 | 0.98715323 |
| miR-409-3p | 31.25176 | 0.07599093 | 0.41442262 | 0.18336578 | 0.85451102 | 0.98715323 |
| miR-467d-3p | 0.42010048 | 0.50393322 | 2.93613085 | 0.17163173 | 0.86372706 | 0.98715323 |

| | | | | | | |
|---|---|---|---|---|---|---|
| miR-669c-5p | 27.7189097 | -0.0960586 | 0.60008324 | -0.1600755 | 0.87282162 | 0.98715323 |
| miR-7225-5p | 0.93754262 | -0.5140675 | 3.23073481 | -0.1591178 | 0.87357605 | 0.98715323 |
| miR-144-5p | 1.6177354 | -0.2944611 | 1.86386926 | -0.1579838 | 0.87446958 | 0.98715323 |
| miR-669b-5p | 0.09759216 | -0.64749 | 4.14903021 | -0.1560582 | 0.87598718 | 0.98715323 |
| miR-210-3p | 1.31819247 | -0.3823803 | 2.47709074 | -0.1543667 | 0.87732061 | 0.98715323 |
| miR-30e-5p | 167.016417 | 0.15151875 | 0.99214965 | 0.15271763 | 0.87862095 | 0.98715323 |
| miR-295-3p | 3.42954705 | -0.1847508 | 1.28768728 | -0.1434749 | 0.88591516 | 0.98715323 |
| miR-34b-5p | 24.8292741 | 0.19765157 | 1.38450723 | 0.14275951 | 0.88648012 | 0.98715323 |
| miR-467a-3p | 5.38817566 | 0.15859494 | 1.11720557 | 0.14195681 | 0.88711412 | 0.98715323 |
| miR-350-3p | 6.88109379 | -0.1469007 | 1.07098254 | -0.1371645 | 0.8909008 | 0.98715323 |
| miR-1943-5p | 0.23012752 | -0.5673652 | 4.15125826 | -0.1366731 | 0.89128922 | 0.98715323 |
| miR-217-3p | 0.12167728 | -0.5672994 | 4.1512599 | -0.1366572 | 0.89130179 | 0.98715323 |
| miR-7021-5p | 0.12167728 | -0.5672994 | 4.1512599 | -0.1366572 | 0.89130179 | 0.98715323 |
| miR-7054-5p | 0.12167728 | -0.5672994 | 4.1512599 | -0.1366572 | 0.89130179 | 0.98715323 |
| miR-3473d | 0.12167728 | -0.5672994 | 4.1512599 | -0.1366572 | 0.89130179 | 0.98715323 |
| miR-5626-3p | 0.12167728 | -0.5672994 | 4.1512599 | -0.1366572 | 0.89130179 | 0.98715323 |
| miR-7646-5p | 0.12167728 | -0.5672994 | 4.1512599 | -0.1366572 | 0.89130179 | 0.98715323 |
| miR-370-3p | 0.21255506 | -0.5672705 | 4.15126054 | -0.1366502 | 0.89130732 | 0.98715323 |
| miR-7082-5p | 0.29268979 | -0.5672434 | 4.15126114 | -0.1366436 | 0.89131248 | 0.98715323 |
| miR-7043-3p | 0.70262989 | 0.43518568 | 3.2826327 | 0.13257215 | 0.89453176 | 0.98715323 |
| miR-133b-3p | 0.118135 | 0.5437239 | 4.19579342 | 0.12958786 | 0.89689251 | 0.98715323 |
| miR-297a-3p | 1.07183877 | -0.2383134 | 1.84930299 | -0.1288666 | 0.89746319 | 0.98715323 |
| miR-467a-5p | 10.9428867 | 0.0823284 | 0.65017238 | 0.1266255 | 0.89923681 | 0.98715323 |
| miR-7653-5p | 0.72175358 | -0.4130733 | 3.29167924 | -0.1254902 | 0.90013552 | 0.98715323 |
| miR-8104 | 0.08678015 | -0.5064501 | 4.15303321 | -0.121947 | 0.90294096 | 0.98715323 |
| miR-203-5p | 0.08678015 | -0.5064501 | 4.15303321 | -0.121947 | 0.90294096 | 0.98715323 |
| miR-6940-3p | 0.08678015 | -0.5064501 | 4.15303321 | -0.121947 | 0.90294096 | 0.98715323 |
| miR-3547-5p | 0.08678015 | -0.5064501 | 4.15303321 | -0.121947 | 0.90294096 | 0.98715323 |
| miR-7216-3p | 0.08678015 | -0.5064501 | 4.15303321 | -0.121947 | 0.90294096 | 0.98715323 |
| miR-7668-5p | 0.08678015 | -0.5064501 | 4.15303321 | -0.121947 | 0.90294096 | 0.98715323 |
| miR-6236 | 0.08678015 | -0.5064501 | 4.15303321 | -0.121947 | 0.90294096 | 0.98715323 |
| let-7c-1-3p | 0.2378391 | -0.5064385 | 4.15303337 | -0.1219442 | 0.90294317 | 0.98715323 |
| miR-1938 | 0.48657648 | -0.5064102 | 4.15303374 | -0.1219374 | 0.90294858 | 0.98715323 |
| miR-196b-5p | 26.3362485 | -0.0787922 | 0.6510596 | -0.1210215 | 0.903674 | 0.98715323 |
| miR-30b-3p | 2.45497289 | -0.1616185 | 1.35537468 | -0.1192427 | 0.90508308 | 0.98715323 |
| miR-128-1-5p | 0.81916492 | 0.36254477 | 3.06265219 | 0.11837608 | 0.90576968 | 0.98715323 |
| miR-1969 | 0.56552509 | -0.3733856 | 3.18721663 | -0.117151 | 0.90674041 | 0.98715323 |
| miR-298-5p | 7.23727778 | -0.0883141 | 0.76815758 | -0.1149687 | 0.90846992 | 0.98715323 |
| miR-187-3p | 1.50236409 | 0.23553239 | 2.07613685 | 0.11344743 | 0.90967584 | 0.98715323 |
| miR-7237-3p | 1.07297075 | 0.30776426 | 2.78745558 | 0.11041046 | 0.91208386 | 0.98715323 |
| miR-1843b-3p | 32.6640078 | 0.05430941 | 0.49308113 | 0.11014296 | 0.912296 | 0.98715323 |
| miR-98-3p | 0.65380206 | 0.37343752 | 3.41297118 | 0.10941713 | 0.91287165 | 0.98715323 |
| miR-17-3p | 0.59741414 | 0.34460793 | 3.20491668 | 0.10752477 | 0.91437267 | 0.98715323 |
| miR-185-5p | 2.9484287 | 0.15609754 | 1.62385058 | 0.09612802 | 0.9234189 | 0.98715323 |
| miR-466e-3p | 0.3926779 | -0.2793868 | 2.92068836 | -0.0956579 | 0.92379232 | 0.98715323 |
| miR-466p-3p | 0.3926779 | -0.2793868 | 2.92068836 | -0.0956579 | 0.92379232 | 0.98715323 |

| | | | | | | |
|---|---|---|---|---|---|---|
| miR-466a-3p | 0.3926779 | -0.2793868 | 2.92068836 | -0.0956579 | 0.92379232 | 0.98715323 |
| miR-672-3p | 0.69653624 | 0.32119344 | 3.54283902 | 0.0906599 | 0.92776283 | 0.98715323 |
| miR-872-5p | 34.3884674 | -0.0547474 | 0.61959464 | -0.08836 | 0.92959053 | 0.98715323 |
| miR-130b-5p | 38.0105853 | 0.05255694 | 0.61252925 | 0.08580316 | 0.9316229 | 0.98715323 |
| miR-199a-5p | 7.36944671 | 0.09644345 | 1.13652197 | 0.08485841 | 0.93237395 | 0.98715323 |
| miR-467b-5p | 0.9405871 | -0.1474856 | 1.79096817 | -0.0823496 | 0.93436869 | 0.98715323 |
| miR-205-5p | 15.6252425 | -0.0615721 | 0.76801504 | -0.0801705 | 0.93610167 | 0.98715323 |
| miR-1843a-5p | 6.54927118 | -0.0660302 | 0.90198107 | -0.0732057 | 0.94164242 | 0.98715323 |
| miR-468-5p | 8.93206347 | 0.05788135 | 0.80660215 | 0.07175948 | 0.94279332 | 0.98715323 |
| miR-7032-3p | 0.07297357 | -0.2965056 | 4.15973292 | -0.07128 | 0.94317495 | 0.98715323 |
| miR-7059-5p | 0.07297357 | -0.2965056 | 4.15973292 | -0.07128 | 0.94317495 | 0.98715323 |
| miR-3102-5p.2- | 0.16385134 | -0.2964942 | 4.15973309 | -0.0712772 | 0.94317713 | 0.98715323 |
| miR-1306-3p | 0.35422355 | -0.2846699 | 4.09569036 | -0.0695048 | 0.94458785 | 0.98715323 |
| miR-202-5p | 4.12620884 | -0.0875593 | 1.26347514 | -0.0693004 | 0.94475054 | 0.98715323 |
| let-7e-3p | 0.13426785 | 0.2839659 | 4.1838917 | 0.06787123 | 0.94588814 | 0.98715323 |
| miR-6916-5p | 0.04339008 | 0.2839535 | 4.1838914 | 0.06786828 | 0.94589049 | 0.98715323 |
| miR-217-5p | 0.04339008 | 0.2839535 | 4.1838914 | 0.06786828 | 0.94589049 | 0.98715323 |
| miR-301a-5p | 0.04339008 | 0.2839535 | 4.1838914 | 0.06786828 | 0.94589049 | 0.98715323 |
| miR-6931-3p | 0.04339008 | 0.2839535 | 4.1838914 | 0.06786828 | 0.94589049 | 0.98715323 |
| miR-377-3p | 0.04339008 | 0.2839535 | 4.1838914 | 0.06786828 | 0.94589049 | 0.98715323 |
| miR-369-5p | 0.04339008 | 0.2839535 | 4.1838914 | 0.06786828 | 0.94589049 | 0.98715323 |
| miR-7218-5p | 0.04339008 | 0.2839535 | 4.1838914 | 0.06786828 | 0.94589049 | 0.98715323 |
| miR-6985-3p | 0.04339008 | 0.2839535 | 4.1838914 | 0.06786828 | 0.94589049 | 0.98715323 |
| miR-5107-3p | 0.17016248 | 0.28391236 | 4.18389005 | 0.06785847 | 0.9458983 | 0.98715323 |
| miR-6948-3p | 0.9026544 | -0.1933172 | 2.85629152 | -0.0676812 | 0.94603942 | 0.98715323 |
| miR-22-5p | 0.53459881 | 0.19703096 | 2.993175 | 0.06582674 | 0.94751577 | 0.98715323 |
| miR-215-5p | 0.69339238 | 0.18732346 | 2.98531124 | 0.06274839 | 0.94996687 | 0.98715323 |
| miR-465a-5p | 68.8682584 | -0.028121 | 0.46183766 | -0.0608895 | 0.95144724 | 0.98715323 |
| miR-1843b-5p | 13.0562397 | 0.04305572 | 0.73628396 | 0.05847706 | 0.95336864 | 0.98715323 |
| miR-101a-3p | 23.1599465 | -0.0343955 | 0.61856752 | -0.055605 | 0.95565644 | 0.98715323 |
| miR-467e-3p | 0.1311008 | -0.2162626 | 4.16255039 | -0.0519543 | 0.95856507 | 0.98715323 |
| miR-9-3p | 0.27670588 | 0.21485182 | 4.15303249 | 0.05173372 | 0.95874087 | 0.98715323 |
| miR-9-5p | 6.40917121 | 0.05221925 | 1.01199311 | 0.0516004 | 0.9588471 | 0.98715323 |
| miR-20a-3p | 0.81996891 | 0.17432423 | 3.48964628 | 0.0499547 | 0.96015849 | 0.98715323 |
| miR-1981-3p | 3.38152849 | 0.07160778 | 1.5875652 | 0.04510541 | 0.96402329 | 0.98715323 |
| miR-146a-5p | 771.592558 | -0.0151773 | 0.34077692 | -0.0445373 | 0.96447609 | 0.98715323 |
| miR-183-5p | 637.348891 | 0.01681388 | 0.37960953 | 0.04429257 | 0.96467119 | 0.98715323 |
| miR-3062-5p | 1.1157459 | 0.09625351 | 2.18867212 | 0.04397804 | 0.96492191 | 0.98715323 |
| miR-1291 | 0.31862991 | -0.1743644 | 4.16408227 | -0.0418734 | 0.9665996 | 0.98715323 |
| miR-760-3p | 0.23692155 | -0.17435 | 4.16408261 | -0.04187 | 0.96660235 | 0.98715323 |
| miR-344d-3p | 0.12847132 | -0.1743266 | 4.16408316 | -0.0418643 | 0.96660684 | 0.98715323 |
| miR-1931 | 0.06508512 | -0.1743034 | 4.1640837 | -0.0418588 | 0.96661129 | 0.98715323 |
| miR-6972-3p | 0.06508512 | -0.1743034 | 4.1640837 | -0.0418588 | 0.96661129 | 0.98715323 |
| miR-29c-3p | 0.92929929 | -0.1267522 | 3.04727211 | -0.0415953 | 0.96682132 | 0.98715323 |
| miR-191-5p | 8236.5581 | 0.01221123 | 0.29404331 | 0.04152868 | 0.96687443 | 0.98715323 |
| miR-300-5p | 0.04864904 | 0.16812614 | 4.1783127 | 0.04023781 | 0.96790354 | 0.98715323 |

| | | | | | | |
|---|---|---|---|---|---|---|
| miR-148b-5p | 1.41662624 | -0.0704595 | 1.81703193 | -0.0387772 | 0.96906798 | 0.98715323 |
| miR-200a-3p | 6.9552802 | 0.0296476 | 0.83211005 | 0.03562942 | 0.97157785 | 0.98796816 |
| miR-871-3p | 944.913414 | -0.0093486 | 0.28761014 | -0.0325044 | 0.97406984 | 0.98796816 |
| miR-1188-3p | 1.17284112 | 0.08309063 | 2.58856589 | 0.0320991 | 0.97439302 | 0.98796816 |
| miR-879-5p | 0.17702488 | -0.1244679 | 4.14400352 | -0.0300357 | 0.9760386 | 0.98810709 |
| miR-378d | 0.71288438 | 0.06691302 | 2.79818517 | 0.02391301 | 0.980922 | 0.99151168 |
| miR-15b-5p | 189.084791 | -0.0068272 | 0.30999286 | -0.0220236 | 0.98242913 | 0.99151168 |
| miR-3057-5p | 3.32793103 | -0.0163714 | 1.38353617 | -0.011833 | 0.99055884 | 0.99744768 |
| miR-293-5p | 6.37104336 | -0.0112474 | 1.11740343 | -0.0100657 | 0.9919689 | 0.99744768 |
| miR-451a | 17.1113885 | 0.00850562 | 0.95304341 | 0.00892469 | 0.99287922 | 0.99744768 |
| miR-669o-3p | 0.21123546 | 0.01759355 | 4.17166336 | 0.0042174 | 0.99663502 | 0.99747269 |
| miR-540-3p | 0.80058379 | 0.00982845 | 2.3760192 | 0.00413652 | 0.99669955 | 0.99747269 |
| miR-676-3p | 309.355277 | 0.00105704 | 0.33371284 | 0.00316752 | 0.99747269 | 0.99747269 |
| miR-103-3p | 203.394758 | 0.0253609 | 0.56002225 | 0.04528553 | NA | NA |
| miR-378a-3p | 17.4797114 | 0.90418124 | 0.8706665 | 1.0384932 | NA | NA |
| miR-107-3p | 66.4487281 | 0.60163361 | 0.6937356 | 0.86723763 | NA | NA |
| miR-463-5p | 236.689323 | 0.83798409 | 0.56559811 | 1.48158926 | NA | NA |

| HS692_miRNA_pvalues_INTERACTION | | | | HS692_miRNA_pvalues_INTERACTION | | |
|---|---|---|---|---|---|---|
| | baseMean | log2FoldChar | lfcSE | stat | pvalue | padj |
| miR-1247-3p | 10.5798277 | 5.67703005 | 1.50485821 | 3.77246841 | 0.00016164 | 0.10587449 |
| miR-148b-3p | 110.979738 | -2.4235976 | 0.74419325 | -3.2566777 | 0.00112724 | 0.36917224 |
| miR-195a-5p | 150.334642 | -1.1402083 | 0.44986051 | -2.5345819 | 0.01125816 | 0.9999244 |
| miR-221-3p | 75.1680938 | 2.04538128 | 0.833531 | 2.45387547 | 0.01413259 | 0.9999244 |
| miR-18a-3p | 2.41760672 | 5.71579155 | 2.3757661 | 2.40587302 | 0.01613387 | 0.9999244 |
| miR-7217-3p | 97.335608 | 1.61032594 | 0.67592372 | 2.3824078 | 0.01719984 | 0.9999244 |
| miR-190b-5p | 22.2117223 | -2.1614975 | 0.91976068 | -2.3500651 | 0.01877013 | 0.9999244 |
| miR-205-5p | 15.6252425 | 2.77416874 | 1.1885156 | 2.33414584 | 0.01958809 | 0.9999244 |
| miR-21a-3p | 18.0654067 | 2.44155009 | 1.05716704 | 2.30952158 | 0.02091466 | 0.9999244 |
| miR-410-3p | 4.64296563 | 4.22238375 | 1.86508047 | 2.26391505 | 0.02357934 | 0.9999244 |
| miR-677-3p | 3.47500083 | 5.11487461 | 2.27682845 | 2.246491 | 0.02467258 | 0.9999244 |
| miR-423-3p | 47.2313136 | 1.5125183 | 0.68438121 | 2.21005235 | 0.02710153 | 0.9999244 |
| miR-471-3p | 5.65147346 | -3.5084807 | 1.59091581 | -2.2053214 | 0.02743155 | 0.9999244 |
| miR-434-3p | 23.72971 | 2.09744594 | 0.9851743 | 2.12900999 | 0.03325343 | 0.9999244 |
| miR-471-5p | 12.6499637 | -2.9450439 | 1.41803798 | -2.0768441 | 0.03781595 | 0.9999244 |
| miR-878-3p | 5.15579097 | -3.0005808 | 1.4586394 | -2.0571094 | 0.03967571 | 0.9999244 |
| miR-881-3p | 131.778904 | -1.0560752 | 0.51398597 | -2.0546772 | 0.0399102 | 0.9999244 |
| miR-880-3p | 8.06722271 | 3.36162611 | 1.65070089 | 2.03648409 | 0.04170177 | 0.9999244 |
| miR-127-3p | 133.600536 | 1.39087951 | 0.68515226 | 2.03002982 | 0.04235351 | 0.9999244 |
| miR-211-5p | 2.84205704 | -5.5264572 | 2.75947995 | -2.0027169 | 0.04520768 | 0.9999244 |
| miR-296-5p | 15.7564006 | 2.10676424 | 1.07722542 | 1.95573202 | 0.05049673 | 0.9999244 |
| miR-125a-5p | 566.450555 | 1.22739529 | 0.6831748 | 1.79660505 | 0.07239834 | 0.9999244 |
| miR-340-5p | 230.688377 | -1.2822998 | 0.72585946 | -1.7665951 | 0.07729606 | 0.9999244 |
| miR-351-5p | 70.8202101 | 1.36451158 | 0.77441162 | 1.76199781 | 0.07806967 | 0.9999244 |
| miR-150-5p | 67.1664819 | -1.2992715 | 0.75191327 | -1.727954 | 0.08399647 | 0.9999244 |
| miR-99b-5p | 638.306367 | 1.05718548 | 0.62120453 | 1.70183156 | 0.08878695 | 0.9999244 |
| miR-148a-5p | 40.6169889 | 1.15391575 | 0.68129693 | 1.69370462 | 0.09032143 | 0.9999244 |
| miR-429-3p | 97.3046123 | 1.33528565 | 0.79485294 | 1.67991534 | 0.09297379 | 0.9999244 |
| miR-200a-3p | 6.9552802 | 2.15620463 | 1.29551324 | 1.66436325 | 0.09603986 | 0.9999244 |
| miR-200b-3p | 43.7631029 | 1.22221327 | 0.73625152 | 1.66004855 | 0.09690469 | 0.9999244 |
| miR-872-5p | 34.3884674 | -1.5409524 | 0.93129287 | -1.6546379 | 0.09799798 | 0.9999244 |
| miR-743a-5p | 7.53008851 | -2.1933652 | 1.33213412 | -1.6465048 | 0.09965988 | 0.9999244 |
| miR-132-5p | 0.97063471 | -7.5975565 | 4.64960022 | -1.6340236 | 0.10225389 | 0.9999244 |
| miR-3081-3p | 2.78968103 | 4.55188044 | 2.79298843 | 1.6297527 | 0.10315377 | 0.9999244 |
| miR-365-3p | 9.67637868 | 2.42858177 | 1.50288188 | 1.61594986 | 0.10610515 | 0.9999244 |
| miR-532-3p | 4.07129501 | -3.1775508 | 1.9765807 | -1.6075998 | 0.10792285 | 0.9999244 |
| miR-3086-5p | 1.32850623 | 5.63290022 | 3.52815545 | 1.5965567 | 0.11036456 | 0.9999244 |
| miR-379-5p | 4.0139086 | 3.25005048 | 2.05457999 | 1.58185638 | 0.11368236 | 0.9999244 |
| miR-145a-5p | 61.9461242 | 1.66042068 | 1.05439876 | 1.57475591 | 0.1153128 | 0.9999244 |
| miR-883b-5p | 3.33547215 | -3.2727948 | 2.10426497 | -1.555315 | 0.11987108 | 0.9999244 |
| miR-142a-5p | 317.260067 | -0.6698344 | 0.43152853 | -1.5522367 | 0.1206056 | 0.9999244 |
| miR-222-3p | 46.8754024 | 0.91198698 | 0.59281349 | 1.53840457 | 0.12394973 | 0.9999244 |
| miR-7230-5p | 12.6123363 | -1.3894573 | 0.91064796 | -1.5257897 | 0.12706224 | 0.9999244 |
| miR-7068-3p | 12.7803931 | 1.94946014 | 1.27835847 | 1.52497142 | 0.12726623 | 0.9999244 |

| | | | | | | |
|---|---|---|---|---|---|---|
| miR-743b-3p | 512.187236 | -0.8326224 | 0.54648171 | -1.5236053 | 0.12760733 | 0.9999244 |
| miR-676-3p | 309.355277 | 0.74909895 | 0.49344208 | 1.51810918 | 0.12898688 | 0.9999244 |
| miR-10b-3p | 5.87406405 | 2.424824 | 1.60743411 | 1.50850599 | 0.13142507 | 0.9999244 |
| miR-31-5p | 5.93726446 | 2.23953237 | 1.48609471 | 1.50699168 | 0.13181279 | 0.9999244 |
| miR-1247-5p | 11.0116551 | 2.05323985 | 1.37967585 | 1.48820453 | 0.13669696 | 0.9999244 |
| miR-15b-5p | 189.084791 | -0.6807752 | 0.45812062 | -1.4860174 | 0.13727451 | 0.9999244 |
| miR-339-5p | 2.66677989 | 3.58830468 | 2.41562836 | 1.48545394 | 0.13742362 | 0.9999244 |
| miR-34c-3p | 262.034111 | -0.764409 | 0.51608137 | -1.4811793 | 0.13855881 | 0.9999244 |
| miR-671-3p | 18.2438899 | 1.29679774 | 0.87630757 | 1.47984313 | 0.13891512 | 0.9999244 |
| miR-222-5p | 1.11668659 | 4.85181579 | 3.36151415 | 1.44334237 | 0.14892405 | 0.9999244 |
| miR-671-5p | 1.29064859 | 5.43270445 | 3.786915 | 1.43459899 | 0.1514014 | 0.9999244 |
| miR-468-5p | 8.93206347 | -1.7114515 | 1.19947587 | -1.4268328 | 0.15362809 | 0.9999244 |
| miR-193b-3p | 5.28269816 | -2.4525165 | 1.7221249 | -1.4241223 | 0.15441107 | 0.9999244 |
| miR-200c-3p | 43.9662858 | 0.92300984 | 0.65961678 | 1.39931224 | 0.16171937 | 0.9999244 |
| miR-20a-5p | 9.68228301 | -1.7695444 | 1.27268865 | -1.3903985 | 0.16440791 | 0.9999244 |
| miR-335-3p | 3.1411732 | 3.27375555 | 2.36052596 | 1.38687547 | 0.16547974 | 0.9999244 |
| miR-320-3p | 35.3779332 | 1.13620276 | 0.8253931 | 1.37655955 | 0.16864846 | 0.9999244 |
| miR-34c-5p | 1365.01758 | -1.067552 | 0.77887029 | -1.3706416 | 0.17048671 | 0.9999244 |
| miR-16-5p | 3180.11971 | -0.6312418 | 0.46066103 | -1.3702956 | 0.17059465 | 0.9999244 |
| miR-669c-5p | 27.7189097 | -1.216098 | 0.88891837 | -1.3680649 | 0.17129176 | 0.9999244 |
| miR-133a-3p | 13.2516827 | -1.578311 | 1.15767281 | -1.3633481 | 0.17277283 | 0.9999244 |
| miR-7215-3p | 0.89167789 | -5.5304971 | 4.06744642 | -1.3596976 | 0.17392564 | 0.9999244 |
| miR-212-5p | 1.00092627 | -5.1976139 | 3.89289955 | -1.3351523 | 0.1818265 | 0.9999244 |
| miR-467d-5p | 6.87334942 | -1.9776681 | 1.48468724 | -1.3320436 | 0.18284588 | 0.9999244 |
| miR-181d-5p | 11.1097067 | 1.45636526 | 1.09814076 | 1.32621 | 0.18477015 | 0.9999244 |
| miR-741-5p | 6.63699476 | -2.3539751 | 1.77982384 | -1.3225888 | 0.18597217 | 0.9999244 |
| miR-8114 | 4.63926344 | 2.60672057 | 2.00487997 | 1.30018785 | 0.19353659 | 0.9999244 |
| miR-130b-5p | 38.0105853 | 1.19638252 | 0.92772144 | 1.28959241 | 0.19719221 | 0.9999244 |
| miR-21a-5p | 686.905345 | -0.7634598 | 0.59220036 | -1.2891917 | 0.19733145 | 0.9999244 |
| miR-344-3p | 2.01375743 | 7.74111181 | 6.04177735 | 1.281264 | 0.20010095 | 0.9999244 |
| miR-7222-5p | 3.5716854 | -2.5421173 | 2.02553175 | -1.255037 | 0.20946533 | 0.9999244 |
| miR-470-5p | 5636.05708 | -0.6562331 | 0.52367412 | -1.2531325 | 0.21015749 | 0.9999244 |
| miR-15a-5p | 15.6104589 | -1.5098309 | 1.21343364 | -1.2442633 | 0.21340268 | 0.9999244 |
| miR-741-3p | 127.971851 | 0.8093953 | 0.66345824 | 1.21996419 | 0.22247845 | 0.9999244 |
| miR-101b-3p | 13.5057654 | -1.2787183 | 1.05558285 | -1.211386 | 0.2257475 | 0.9999244 |
| miR-7234-5p | 0.84396123 | 5.43821426 | 4.49820219 | 1.20897506 | 0.22667243 | 0.9999244 |
| miR-125b-1-3p | 6.8504028 | 1.96586899 | 1.64342389 | 1.19620325 | 0.23161725 | 0.9999244 |
| miR-181b-5p | 32.2043317 | 1.23931302 | 1.04429482 | 1.18674631 | 0.2353277 | 0.9999244 |
| miR-871-5p | 716.00589 | -0.6056564 | 0.51151212 | -1.1840509 | 0.23639292 | 0.9999244 |
| miR-28a-3p | 11.530027 | -1.4161763 | 1.20596659 | -1.1743081 | 0.24027166 | 0.9999244 |
| miR-467a-3p | 5.38817566 | -1.9715785 | 1.67946548 | -1.1739321 | 0.24042222 | 0.9999244 |
| miR-9-5p | 6.40917121 | -1.8660442 | 1.59269089 | -1.1716298 | 0.24134571 | 0.9999244 |
| miR-421-3p | 1.79965379 | 3.47210506 | 2.96664426 | 1.17038133 | 0.24184754 | 0.9999244 |
| miR-467c-5p | 3.46959286 | -2.3886097 | 2.06978535 | -1.1540374 | 0.24848483 | 0.9999244 |
| miR-31-3p | 1.39114034 | -4.0732058 | 3.5479694 | -1.1480386 | 0.25095263 | 0.9999244 |
| miR-8112 | 2.01044339 | 2.9820572 | 2.60840533 | 1.14324916 | 0.25293516 | 0.9999244 |

| | | | | | | |
|---|---|---|---|---|---|---|
| miR-878-5p | 70.3483372 | -0.6878893 | 0.60594701 | -1.1352301 | 0.2562789 | 0.9999244 |
| miR-7232-3p | 49.1727142 | 0.71520332 | 0.63382663 | 1.12838951 | 0.25915546 | 0.9999244 |
| miR-196a-5p | 21.734875 | 1.06000468 | 0.94067131 | 1.1268598 | 0.25980176 | 0.9999244 |
| miR-203-3p | 21.1211209 | 1.10527688 | 0.98939152 | 1.1171279 | 0.26393964 | 0.9999244 |
| miR-181a-1-3p | 1.32505335 | 4.08556014 | 3.67765449 | 1.11091462 | 0.2666051 | 0.9999244 |
| miR-298-5p | 7.23727778 | -1.2790368 | 1.15300617 | -1.1093062 | 0.26729813 | 0.9999244 |
| miR-451a | 17.1113885 | -1.5695704 | 1.41539051 | -1.108931 | 0.26745996 | 0.9999244 |
| miR-872-3p | 3.0419223 | -2.4086582 | 2.17493116 | -1.1074641 | 0.26809333 | 0.9999244 |
| miR-151-3p | 493.324752 | 0.42155042 | 0.38391249 | 1.09803778 | 0.27218799 | 0.9999244 |
| miR-674-5p | 0.79662291 | 5.07598309 | 4.62658764 | 1.09713324 | 0.27258315 | 0.9999244 |
| miR-125b-2-3p | 14.8456703 | 1.09105642 | 0.99451025 | 1.09707911 | 0.27260681 | 0.9999244 |
| miR-674-3p | 2.69871643 | 2.32823713 | 2.12980475 | 1.09316929 | 0.27431948 | 0.9999244 |
| miR-6538 | 16.2051488 | -0.9972665 | 0.92303661 | -1.0804193 | 0.27995551 | 0.9999244 |
| miR-134-5p | 1.98854484 | -3.0290476 | 2.81103827 | -1.0775547 | 0.28123251 | 0.9999244 |
| miR-122-5p | 0.85811582 | 5.46718308 | 5.10102248 | 1.0717818 | 0.28381805 | 0.9999244 |
| miR-669o-5p | 4.38164049 | -2.1887331 | 2.05470656 | -1.065229 | 0.28677231 | 0.9999244 |
| miR-541-5p | 85.6242488 | 0.57027993 | 0.53974872 | 1.05656559 | 0.29070989 | 0.9999244 |
| miR-19a-3p | 11.4926159 | -1.4986887 | 1.41948488 | -1.0557976 | 0.29106072 | 0.9999244 |
| miR-7060-5p | 1.17256607 | 3.57398424 | 3.40948608 | 1.0482472 | 0.29452473 | 0.9999244 |
| miR-148a-3p | 4078.96285 | 0.47717147 | 0.45602154 | 1.04637922 | 0.29538598 | 0.9999244 |
| miR-221-5p | 15.7134515 | 1.07681728 | 1.03079197 | 1.04465044 | 0.29618456 | 0.9999244 |
| miR-7b-5p | 1.36611929 | -3.3016036 | 3.16051447 | -1.0446412 | 0.29618883 | 0.9999244 |
| miR-505-5p | 0.83056638 | 4.7428633 | 4.55575736 | 1.04107022 | 0.29784296 | 0.9999244 |
| let-7b-3p | 1.47839757 | 3.58281935 | 3.48514733 | 1.02802522 | 0.30393797 | 0.9999244 |
| miR-7a-5p | 95.5240462 | -0.7106337 | 0.69527452 | -1.0220909 | 0.3067379 | 0.9999244 |
| miR-1843a-3p | 3.48033372 | -1.8543881 | 1.83347385 | -1.0114069 | 0.31182172 | 0.9999244 |
| miR-1a-3p | 8.02350501 | -1.5289945 | 1.51848469 | -1.0069213 | 0.31397261 | 0.9999244 |
| miR-30c-2-3p | 53.1794842 | 0.56589312 | 0.56234157 | 1.00631564 | 0.31426376 | 0.9999244 |
| let-7a-5p | 12110.1372 | -0.4237176 | 0.42395162 | -0.9994481 | 0.31757767 | 0.9999244 |
| miR-501-3p | 4.84182452 | 1.73462554 | 1.74303966 | 0.99517274 | 0.31965226 | 0.9999244 |
| miR-139-3p | 0.76411505 | 3.92126675 | 3.9403408 | 0.99515929 | 0.3196588 | 0.9999244 |
| miR-192-5p | 525.686551 | -0.5468178 | 0.54973145 | -0.9946999 | 0.31988226 | 0.9999244 |
| miR-465c-5p | 181.957168 | -0.4573554 | 0.46389364 | -0.9859056 | 0.32417942 | 0.9999244 |
| miR-187-3p | 1.50236409 | 3.17066234 | 3.22081932 | 0.98442726 | 0.32490548 | 0.9999244 |
| miR-409-3p | 31.25176 | 0.62791359 | 0.63966011 | 0.98163631 | 0.32627905 | 0.9999244 |
| miR-652-3p | 4.68064068 | 1.68384788 | 1.72882991 | 0.97398123 | 0.33006586 | 0.9999244 |
| miR-3062-5p | 1.1157459 | 3.23160893 | 3.33848476 | 0.96798673 | 0.333051 | 0.9999244 |
| miR-743b-5p | 13.3933767 | -1.0772392 | 1.11925789 | -0.9624584 | 0.33581937 | 0.9999244 |
| miR-200b-5p | 1.6344838 | 2.36024267 | 2.47447331 | 0.95383638 | 0.34016647 | 0.9999244 |
| miR-7214-3p | 14.3740518 | 1.07910261 | 1.13176756 | 0.95346664 | 0.34035369 | 0.9999244 |
| miR-666-3p | 1.05680737 | 3.93176345 | 4.12937389 | 0.95214518 | 0.34102336 | 0.9999244 |
| miR-582-3p | 0.6706355 | -5.1711867 | 5.44255162 | -0.9501401 | 0.34204106 | 0.9999244 |
| miR-191-3p | 12.6768131 | 0.98623915 | 1.04138489 | 0.94704576 | 0.34361546 | 0.9999244 |
| miR-26b-3p | 0.95444117 | 3.58398172 | 3.79287933 | 0.94492374 | 0.34469781 | 0.9999244 |
| miR-125b-5p | 139.941512 | 0.50422303 | 0.54199685 | 0.9303062 | 0.35221257 | 0.9999244 |
| miR-7a-1-3p | 1.40768575 | -2.5764946 | 2.81172706 | -0.9163388 | 0.35948923 | 0.9999244 |

| | | | | | | |
|---|---|---|---|---|---|---|
| miR-350-3p | 6.88109379 | -1.5210292 | 1.66482094 | -0.9136293 | 0.36091165 | 0.9999244 |
| miR-431-5p | 1.10369239 | 3.04929583 | 3.34109442 | 0.91266377 | 0.3614194 | 0.9999244 |
| miR-342-3p | 86.1737741 | -0.4634581 | 0.50932852 | -0.9099395 | 0.36285444 | 0.9999244 |
| miR-664-5p | 2.60168405 | -1.8755748 | 2.06396005 | -0.9087263 | 0.36349461 | 0.9999244 |
| miR-877-5p | 0.73376402 | 4.22548525 | 4.66241008 | 0.90628777 | 0.36478355 | 0.9999244 |
| miR-152-5p | 3.40236558 | -1.9586847 | 2.16296991 | -0.9055534 | 0.36517229 | 0.9999244 |
| miR-128-3p | 148.185578 | -0.4603088 | 0.50876231 | -0.904762 | 0.36559147 | 0.9999244 |
| miR-7230-3p | 46.0677911 | -0.5662556 | 0.63086914 | -0.8975802 | 0.3694094 | 0.9999244 |
| miR-124-5p | 0.55117314 | -4.2644284 | 4.76709757 | -0.8945545 | 0.3710253 | 0.9999244 |
| miR-1188-3p | 1.17284112 | 3.46021429 | 3.89380945 | 0.888645 | 0.3741939 | 0.9999244 |
| miR-7688-5p | 0.7748139 | 3.4868069 | 3.92560019 | 0.88822262 | 0.37442101 | 0.9999244 |
| miR-30b-3p | 2.45497289 | -1.8428934 | 2.11204287 | -0.8725644 | 0.38290055 | 0.9999244 |
| miR-8094 | 4.54244484 | 1.94389583 | 2.24190444 | 0.86707345 | 0.38590176 | 0.9999244 |
| miR-99a-3p | 0.84195733 | 3.69397932 | 4.26611995 | 0.86588735 | 0.38655194 | 0.9999244 |
| miR-155-5p | 2.95756793 | 1.80519611 | 2.08880459 | 0.8642245 | 0.38746457 | 0.9999244 |
| miR-703 | 0.59496855 | 4.31290551 | 4.99917337 | 0.86272373 | 0.38828938 | 0.9999244 |
| miR-17-5p | 2.20817842 | -1.9599862 | 2.28671053 | -0.8571204 | 0.39137836 | 0.9999244 |
| miR-669d-5p | 1.09656679 | 2.77380376 | 3.24505127 | 0.85477964 | 0.39267315 | 0.9999244 |
| miR-150-3p | 0.64869789 | 4.61289799 | 5.51000785 | 0.83718537 | 0.40248838 | 0.9999244 |
| miR-1964-3p | 0.68235136 | 4.46708301 | 5.36710671 | 0.83230747 | 0.40523541 | 0.9999244 |
| miR-99a-5p | 82.9188033 | 0.60841871 | 0.7423824 | 0.81954894 | 0.41247329 | 0.9999244 |
| miR-19b-3p | 42.3812045 | -0.6339 | 0.78124155 | -0.8114008 | 0.41713556 | 0.9999244 |
| miR-293-5p | 6.37104336 | -1.3921217 | 1.72136763 | -0.8087301 | 0.41867044 | 0.9999244 |
| miR-3095-3p | 1.07401383 | 2.79297506 | 3.48517396 | 0.80138756 | 0.42290731 | 0.9999244 |
| miR-423-5p | 599.714497 | 0.43684885 | 0.54598522 | 0.80011113 | 0.42364641 | 0.9999244 |
| miR-485-5p | 4.73326633 | 1.57505581 | 1.9748702 | 0.79754903 | 0.42513224 | 0.9999244 |
| miR-27a-3p | 18.3808121 | 0.83070205 | 1.04358152 | 0.7960107 | 0.42602581 | 0.9999244 |
| miR-324-5p | 0.83313136 | 3.3638113 | 4.23584643 | 0.79412966 | 0.42711995 | 0.9999244 |
| miR-7658-3p | 0.65540536 | 4.77340732 | 6.0327172 | 0.79125329 | 0.4287962 | 0.9999244 |
| miR-26a-1-3p | 0.46171449 | -4.8288572 | 6.10837571 | -0.7905305 | 0.42921803 | 0.9999244 |
| miR-182-5p | 3490.89211 | 0.32577809 | 0.4137074 | 0.78746013 | 0.43101256 | 0.9999244 |
| miR-214-3p | 3.82059379 | -1.5091746 | 1.9223305 | -0.7850755 | 0.43240929 | 0.9999244 |
| miR-7653-5p | 0.72175358 | 3.74355368 | 4.82499185 | 0.77586736 | 0.43782731 | 0.9999244 |
| miR-7068-5p | 2.11119042 | 2.19314942 | 2.83719378 | 0.77299951 | 0.43952267 | 0.9999244 |
| miR-100-5p | 91.5998735 | 0.50479897 | 0.65573036 | 0.76982704 | 0.4414025 | 0.9999244 |
| miR-187-5p | 2.16057964 | 2.57308734 | 3.3870447 | 0.75968509 | 0.44744285 | 0.9999244 |
| miR-34b-3p | 1915.43036 | -0.3956474 | 0.52097299 | -0.7594394 | 0.44758976 | 0.9999244 |
| miR-483-3p | 3.44937194 | 1.60216965 | 2.11423464 | 0.75780125 | 0.44856998 | 0.9999244 |
| miR-455-3p | 0.56760543 | 4.27126085 | 5.65562746 | 0.75522316 | 0.45011509 | 0.9999244 |
| miR-93-5p | 224.45357 | -0.3686277 | 0.49227899 | -0.7488186 | 0.45396655 | 0.9999244 |
| miR-3086-3p | 0.58934457 | -4.550919 | 6.13639276 | -0.7416277 | 0.45831292 | 0.9999244 |
| miR-3082-3p | 0.48479468 | 3.98661473 | 5.38795611 | 0.73991225 | 0.45935324 | 0.9999244 |
| miR-7225-5p | 0.93754262 | -3.4599028 | 4.68542534 | -0.7384394 | 0.46024747 | 0.9999244 |
| miR-195a-3p | 1.16652442 | 2.90979814 | 3.9487453 | 0.73689183 | 0.46118813 | 0.9999244 |
| miR-883a-5p | 2.23124848 | 2.20888134 | 3.02055278 | 0.73128381 | 0.46460582 | 0.9999244 |
| miR-300-3p | 2.81409544 | -1.9552446 | 2.67513457 | -0.7308958 | 0.46484281 | 0.9999244 |

| | | | | | | |
|---|---|---|---|---|---|---|
| miR-30d-5p | 1760.5325 | 0.34352102 | 0.4707869 | 0.72967412 | 0.46558941 | 0.9999244 |
| miR-151-5p | 403.539251 | -0.3501776 | 0.48116688 | -0.7277674 | 0.46675598 | 0.9999244 |
| miR-666-5p | 1.02168433 | 2.79881046 | 3.85003486 | 0.72695717 | 0.46725219 | 0.9999244 |
| miR-1843b-5p | 13.0562397 | 0.78288022 | 1.08212251 | 0.72346727 | 0.46939286 | 0.9999244 |
| miR-140-5p | 1.3537423 | -2.4187351 | 3.35118274 | -0.7217557 | 0.47044471 | 0.9999244 |
| miR-15a-3p | 0.82085359 | -2.6234981 | 3.64891666 | -0.71898 | 0.47215324 | 0.9999244 |
| miR-497a-5p | 2.74225735 | 1.97081627 | 2.74603351 | 0.71769564 | 0.47294497 | 0.9999244 |
| miR-7241-5p | 0.58243179 | 4.36048018 | 6.09176252 | 0.71579944 | 0.4741152 | 0.9999244 |
| miR-7036b-5p | 0.32535072 | -4.2985207 | 6.12186371 | -0.7021588 | 0.48258012 | 0.9999244 |
| miR-496a-3p | 1.14879701 | 2.37441075 | 3.38712163 | 0.70101136 | 0.48329592 | 0.9999244 |
| miR-295-3p | 3.42954705 | 1.49917197 | 2.14343597 | 0.69942466 | 0.48428668 | 0.9999244 |
| miR-935 | 0.27163409 | -4.2935337 | 6.15039817 | -0.6980904 | 0.48512069 | 0.9999244 |
| miR-93-3p | 0.72809689 | -3.3648122 | 4.8206788 | -0.6979955 | 0.48518 | 0.9999244 |
| miR-184-3p | 2339.78942 | -0.33001 | 0.47290053 | -0.6978423 | 0.48527581 | 0.9999244 |
| miR-598-3p | 0.51310928 | 4.0638163 | 5.82800737 | 0.69729086 | 0.48562078 | 0.9999244 |
| miR-202-5p | 4.12620884 | -1.2979594 | 1.86186883 | -0.6971272 | 0.4857232 | 0.9999244 |
| miR-6905-3p | 0.30780966 | -4.2455299 | 6.12350089 | -0.6933174 | 0.48811036 | 0.9999244 |
| miR-96-5p | 7.48396364 | -1.2475081 | 1.81656725 | -0.6867393 | 0.49224705 | 0.9999244 |
| miR-205-3p | 3.40748315 | -1.472957 | 2.14991064 | -0.6851248 | 0.4932652 | 0.9999244 |
| miR-301a-3p | 10.7435234 | -0.9314938 | 1.35966027 | -0.6850931 | 0.49328518 | 0.9999244 |
| miR-1930-5p | 0.38185085 | -4.2123409 | 6.15455676 | -0.6844264 | 0.49370597 | 0.9999244 |
| miR-1306-5p | 11.0575603 | -0.9045463 | 1.32437459 | -0.6829988 | 0.4946076 | 0.9999244 |
| miR-224-5p | 4.18875103 | 1.0050994 | 1.50037044 | 0.66990083 | 0.50292101 | 0.9999244 |
| miR-500-3p | 0.52674753 | -3.8450786 | 5.76089207 | -0.667445 | 0.50448794 | 0.9999244 |
| miR-331-5p | 0.40085752 | -4.0855245 | 6.14762019 | -0.6645701 | 0.5063255 | 0.9999244 |
| miR-1188-5p | 0.40085752 | -4.0855245 | 6.14762019 | -0.6645701 | 0.5063255 | 0.9999244 |
| miR-22-3p | 1217.90203 | 0.35408572 | 0.53537367 | 0.66138052 | 0.50836831 | 0.9999244 |
| miR-361-5p | 7.51965324 | 0.96295321 | 1.4581474 | 0.66039497 | 0.5090004 | 0.9999244 |
| miR-484 | 5.68165104 | 1.13936744 | 1.72723761 | 0.65964719 | 0.50948026 | 0.9999244 |
| miR-8118 | 0.68553564 | -3.7754434 | 5.78994297 | -0.6520692 | 0.51435655 | 0.9999244 |
| miR-744-5p | 17.7411866 | 0.72206928 | 1.1091575 | 0.65100699 | 0.51504198 | 0.9999244 |
| miR-410-5p | 0.2535448 | -3.981101 | 6.13261983 | -0.6491681 | 0.51622975 | 0.9999244 |
| miR-6942-5p | 0.2535448 | -3.981101 | 6.13261983 | -0.6491681 | 0.51622975 | 0.9999244 |
| miR-29b-3p | 0.79210748 | 2.74344054 | 4.28892444 | 0.639657 | 0.52239562 | 0.9999244 |
| miR-144-5p | 1.6177354 | 1.88019344 | 2.98366916 | 0.6301615 | 0.52858892 | 0.9999244 |
| miR-1981-3p | 3.38152849 | -1.4360119 | 2.28924489 | -0.6272863 | 0.53047159 | 0.9999244 |
| miR-139-5p | 10.5989984 | 0.77234041 | 1.23204608 | 0.62687624 | 0.53074036 | 0.9999244 |
| miR-5121 | 6.97170829 | 0.98723707 | 1.57524066 | 0.62672142 | 0.53084186 | 0.9999244 |
| miR-6944-5p | 0.32590196 | -3.8202158 | 6.15288142 | -0.6208824 | 0.534677 | 0.9999244 |
| miR-7661-3p | 0.32590196 | -3.8202158 | 6.15288142 | -0.6208824 | 0.534677 | 0.9999244 |
| miR-411-5p | 2.37450332 | 1.40698325 | 2.27964343 | 0.61719444 | 0.53710648 | 0.9999244 |
| miR-184-5p | 0.73658487 | -2.9935288 | 4.85326179 | -0.6168076 | 0.53736163 | 0.9999244 |
| miR-881-5p | 6.08016002 | 1.02086966 | 1.65927281 | 0.61525124 | 0.53838881 | 0.9999244 |
| miR-145a-3p | 15.9145371 | -0.8077353 | 1.31782558 | -0.6129304 | 0.53992238 | 0.9999244 |
| miR-182-3p | 0.24206941 | -3.7796688 | 6.17940608 | -0.6116557 | 0.5407656 | 0.9999244 |
| miR-29c-3p | 0.92929929 | 2.79220272 | 4.58966647 | 0.60836724 | 0.54294393 | 0.9999244 |

| | | | | | | |
|---|---|---|---|---|---|---|
| miR-130b-3p | 37.360924 | -0.4035603 | 0.6663852 | -0.6055961 | 0.54478299 | 0.9999244 |
| miR-101a-3p | 23.1599465 | -0.5563623 | 0.91895633 | -0.6054284 | 0.54489438 | 0.9999244 |
| miR-6947-3p | 0.39289638 | -3.7208401 | 6.15479335 | -0.6045435 | 0.54548238 | 0.9999244 |
| miR-365-1-5p | 0.39289638 | -3.7208401 | 6.15479335 | -0.6045435 | 0.54548238 | 0.9999244 |
| miR-3081-5p | 0.39289638 | -3.7208401 | 6.15479335 | -0.6045435 | 0.54548238 | 0.9999244 |
| miR-5132-3p | 0.39289638 | -3.7208401 | 6.15479335 | -0.6045435 | 0.54548238 | 0.9999244 |
| miR-6915-3p | 0.21690048 | -3.7032212 | 6.14415443 | -0.6027227 | 0.54669319 | 0.9999244 |
| miR-323-3p | 0.21690048 | -3.7032212 | 6.14415443 | -0.6027227 | 0.54669319 | 0.9999244 |
| miR-7117-3p | 0.21690048 | -3.7032212 | 6.14415443 | -0.6027227 | 0.54669319 | 0.9999244 |
| miR-3085-5p | 0.21690048 | -3.7032212 | 6.14415443 | -0.6027227 | 0.54669319 | 0.9999244 |
| miR-1191b-5p | 0.35921461 | -3.7031268 | 6.14414856 | -0.6027079 | 0.54670303 | 0.9999244 |
| miR-92a-3p | 1248.67192 | 0.28462643 | 0.47279576 | 0.60200717 | 0.54716937 | 0.9999244 |
| miR-141-5p | 1.32835685 | 2.49360036 | 4.16918086 | 0.59810319 | 0.54977108 | 0.9999244 |
| miR-345-5p | 2.83472097 | 1.21071448 | 2.03325228 | 0.59545709 | 0.55153798 | 0.9999244 |
| let-7i-5p | 646.879343 | -0.3387052 | 0.57215886 | -0.5919776 | 0.55386559 | 0.9999244 |
| miR-146b-5p | 843.96163 | -0.2492366 | 0.4247814 | -0.5867409 | 0.55737772 | 0.9999244 |
| miR-7653-3p | 0.26723835 | -3.6022759 | 6.15724759 | -0.5850465 | 0.55851648 | 0.9999244 |
| miR-7670-3p | 0.26723835 | -3.6022759 | 6.15724759 | -0.5850465 | 0.55851648 | 0.9999244 |
| miR-7060-3p | 0.26723835 | -3.6022759 | 6.15724759 | -0.5850465 | 0.55851648 | 0.9999244 |
| miR-542-5p | 0.26109 | -3.5768273 | 6.15780024 | -0.5808612 | 0.561334 | 0.9999244 |
| miR-8103 | 0.1901586 | -3.572226 | 6.15039778 | -0.5808122 | 0.56136703 | 0.9999244 |
| miR-539-5p | 0.1901586 | -3.572226 | 6.15039778 | -0.5808122 | 0.56136703 | 0.9999244 |
| miR-200c-5p | 0.1901586 | -3.572226 | 6.15039778 | -0.5808122 | 0.56136703 | 0.9999244 |
| miR-144-3p | 0.3307791 | -3.5722133 | 6.15039778 | -0.5808101 | 0.56136843 | 0.9999244 |
| miR-540-5p | 0.46396931 | -3.5071405 | 6.08056051 | -0.5767791 | 0.56408866 | 0.9999244 |
| miR-98-5p | 115.641042 | 0.38805355 | 0.68008213 | 0.57059808 | 0.56827212 | 0.9999244 |
| miR-7663-3p | 0.24442647 | -3.4838037 | 6.15990223 | -0.5655615 | 0.57169189 | 0.9999244 |
| miR-7657-5p | 0.24442647 | -3.4838037 | 6.15990223 | -0.5655615 | 0.57169189 | 0.9999244 |
| miR-363-3p | 0.24442647 | -3.4838037 | 6.15990223 | -0.5655615 | 0.57169189 | 0.9999244 |
| miR-183-5p | 637.348891 | 0.3156843 | 0.55876956 | 0.56496331 | 0.57209872 | 0.9999244 |
| miR-672-5p | 40.9597656 | -0.391036 | 0.69652833 | -0.5614072 | 0.57451996 | 0.9999244 |
| miR-708-3p | 0.62483669 | -2.8513981 | 5.08732785 | -0.5604903 | 0.57514503 | 0.9999244 |
| miR-16-2-3p | 12.7825411 | 0.58227901 | 1.04306376 | 0.55823914 | 0.5766811 | 0.9999244 |
| miR-30c-1-3p | 4.35338289 | -0.9468 | 1.7010296 | -0.5566041 | 0.57779798 | 0.9999244 |
| miR-434-5p | 3.20697696 | 1.29770538 | 2.34179101 | 0.55415081 | 0.57947564 | 0.9999244 |
| miR-328-3p | 14.3504627 | -0.5096855 | 0.92231077 | -0.5526179 | 0.58052507 | 0.9999244 |
| miR-335-5p | 5.31225984 | -0.8647453 | 1.57237439 | -0.5499615 | 0.58234579 | 0.9999244 |
| miR-143-5p | 1.38733169 | 1.76659254 | 3.2159176 | 0.54932767 | 0.5827806 | 0.9999244 |
| miR-467e-5p | 4.54618643 | -1.0957029 | 2.01191259 | -0.5446076 | 0.58602344 | 0.9999244 |
| miR-196b-5p | 26.3362485 | -0.513012 | 0.95198994 | -0.5388839 | 0.58996698 | 0.9999244 |
| miR-1938 | 0.48657648 | 3.27369793 | 6.1126975 | 0.535557 | 0.59226475 | 0.9999244 |
| miR-381-3p | 1.4691769 | 1.47555569 | 2.75759866 | 0.53508718 | 0.59258957 | 0.9999244 |
| miR-376b-3p | 0.20042876 | -3.2723178 | 6.16519461 | -0.5307728 | 0.59557621 | 0.9999244 |
| miR-1906 | 1.02708116 | 2.37461868 | 4.48419707 | 0.5295527 | 0.59642209 | 0.9999244 |
| miR-677-5p | 22.6023362 | -0.5849552 | 1.10600176 | -0.5288917 | 0.59688055 | 0.9999244 |
| miR-431-3p | 0.40280355 | 3.23532806 | 6.12006123 | 0.52864309 | 0.59705306 | 0.9999244 |

| | | | | | | |
|---|---|---|---|---|---|---|
| miR-676-5p | 2.23226047 | 1.33734513 | 2.53039494 | 0.52851241 | 0.59714374 | 0.9999244 |
| miR-210-3p | 1.31819247 | -1.8984818 | 3.60749138 | -0.5262609 | 0.59870692 | 0.9999244 |
| let-7a-1-3p | 1.18944819 | -1.690092 | 3.25229654 | -0.5196611 | 0.6032998 | 0.9999244 |
| let-7c-2-3p | 1.18944819 | -1.690092 | 3.25229654 | -0.5196611 | 0.6032998 | 0.9999244 |
| miR-1839-3p | 0.33311346 | -3.1804982 | 6.14003945 | -0.5179931 | 0.60446307 | 0.9999244 |
| miR-425-3p | 0.614012 | -2.8672846 | 5.54550723 | -0.5170464 | 0.60512376 | 0.9999244 |
| miR-7663-5p | 0.18198372 | -3.1708144 | 6.16801086 | -0.5140741 | 0.60720021 | 0.9999244 |
| miR-3960 | 0.18198372 | -3.1708144 | 6.16801086 | -0.5140741 | 0.60720021 | 0.9999244 |
| miR-7036b-3p | 0.18198372 | -3.1708144 | 6.16801086 | -0.5140741 | 0.60720021 | 0.9999244 |
| miR-200a-5p | 1.2069913 | -1.9821259 | 3.86111258 | -0.5133561 | 0.60770222 | 0.9999244 |
| miR-1955-5p | 0.39425593 | 3.13107983 | 6.12291763 | 0.51137056 | 0.6090916 | 0.9999244 |
| miR-466d-3p | 0.63912589 | -2.2351602 | 4.39029435 | -0.509114 | 0.61067234 | 0.9999244 |
| let-7i-3p | 1.70454211 | 1.68939955 | 3.32748461 | 0.5077107 | 0.61165624 | 0.9999244 |
| miR-3088-3p | 0.35352844 | 3.10127353 | 6.12850798 | 0.50604055 | 0.61282819 | 0.9999244 |
| miR-7236-3p | 0.34344628 | -3.0985725 | 6.15287993 | -0.5035971 | 0.61454453 | 0.9999244 |
| miR-7240-5p | 0.67367875 | -2.9044101 | 5.81377358 | -0.499574 | 0.61737507 | 0.9999244 |
| miR-7226-3p | 0.1267724 | -3.0708453 | 6.17343748 | -0.4974287 | 0.61888674 | 0.9999244 |
| miR-378d | 0.71288438 | 2.08849164 | 4.21389827 | 0.49561985 | 0.62016264 | 0.9999244 |
| miR-215-5p | 0.69339238 | 2.2201089 | 4.48408948 | 0.49510807 | 0.62052383 | 0.9999244 |
| miR-7223-5p | 1.38907708 | 1.88937277 | 3.81724566 | 0.49495708 | 0.62063042 | 0.9999244 |
| miR-1198-5p | 3.55459749 | 0.91432263 | 1.85592766 | 0.49264993 | 0.62225996 | 0.9999244 |
| miR-181c-5p | 168.593623 | -0.30987 | 0.62972471 | -0.4920722 | 0.62266832 | 0.9999244 |
| miR-669a-5p | 18.083587 | -0.4141126 | 0.84353119 | -0.4909274 | 0.6234778 | 0.9999244 |
| miR-7662-3p | 0.16295098 | -3.0182138 | 6.17261641 | -0.4889683 | 0.62486414 | 0.9999244 |
| miR-129-5p | 0.16295098 | -3.0182138 | 6.17261641 | -0.4889683 | 0.62486414 | 0.9999244 |
| miR-7012-5p | 0.16295098 | -3.0182138 | 6.17261641 | -0.4889683 | 0.62486414 | 0.9999244 |
| miR-191-5p | 8236.5581 | 0.2114678 | 0.43291031 | 0.48847947 | 0.62521026 | 0.9999244 |
| miR-223-5p | 5.10379837 | 0.84258658 | 1.72768439 | 0.48769705 | 0.62576444 | 0.9999244 |
| miR-664-3p | 13.0924885 | 0.49620912 | 1.0244437 | 0.48436934 | 0.62812377 | 0.9999244 |
| miR-669a-3p | 3.87813555 | 0.79697765 | 1.65002498 | 0.48300944 | 0.62908903 | 0.9999244 |
| miR-7a-2-3p | 0.43155262 | -2.9479087 | 6.1458323 | -0.4796598 | 0.63146932 | 0.9999244 |
| miR-6240 | 22.4600502 | 0.44103587 | 0.92255749 | 0.47805787 | 0.63260902 | 0.9999244 |
| miR-1983 | 1.00427594 | -2.0814962 | 4.36160366 | -0.4772318 | 0.63319703 | 0.9999244 |
| miR-10a-5p | 171033.381 | 0.22710058 | 0.48781635 | 0.46554525 | 0.64154104 | 0.9999244 |
| miR-6946-3p | 0.31863131 | 2.84070316 | 6.13424999 | 0.46308891 | 0.64330064 | 0.9999244 |
| miR-883b-3p | 3.94986695 | 0.74835931 | 1.61719832 | 0.46275049 | 0.64354322 | 0.9999244 |
| miR-3066-5p | 1.6247547 | 1.30958669 | 2.8642334 | 0.45722066 | 0.64751245 | 0.9999244 |
| miR-3535 | 2527.66528 | 0.33229975 | 0.73077552 | 0.45472206 | 0.64930921 | 0.9999244 |
| miR-425-5p | 443.12411 | 0.20965834 | 0.46276407 | 0.45305664 | 0.65050796 | 0.9999244 |
| miR-6948-3p | 0.9026544 | 1.88393712 | 4.17094436 | 0.45168119 | 0.65149867 | 0.9999244 |
| miR-26a-2-3p | 0.31192578 | 2.76250296 | 6.13538821 | 0.45025724 | 0.65252497 | 0.9999244 |
| miR-3066-3p | 0.46849696 | -2.7298942 | 6.0884205 | -0.4483748 | 0.65388273 | 0.9999244 |
| miR-1932 | 0.35891615 | 2.74048778 | 6.12065353 | 0.44774431 | 0.65433774 | 0.9999244 |
| miR-3572-5p | 0.35891615 | 2.74048778 | 6.12065353 | 0.44774431 | 0.65433774 | 0.9999244 |
| miR-135a-1-3p | 0.35891615 | 2.74048778 | 6.12065353 | 0.44774431 | 0.65433774 | 0.9999244 |
| miR-744-3p | 0.37696495 | 2.74147449 | 6.12982403 | 0.44723543 | 0.65470508 | 0.9999244 |

| | | | | | | |
|---|---|---|---|---|---|---|
| miR-1969 | 0.56552509 | 2.11686019 | 4.75358403 | 0.44531877 | 0.65608941 | 0.9999244 |
| miR-146a-5p | 771.592558 | 0.22290714 | 0.50197089 | 0.44406387 | 0.65699641 | 0.9999244 |
| miR-467d-3p | 0.42010048 | -1.9473595 | 4.39042331 | -0.4435471 | 0.65737006 | 0.9999244 |
| miR-7229-5p | 78.3795612 | 0.26826241 | 0.60589625 | 0.44275304 | 0.65794437 | 0.9999244 |
| miR-409-5p | 0.29204608 | 2.71248488 | 6.13029515 | 0.44247215 | 0.65814758 | 0.9999244 |
| miR-324-3p | 0.12132248 | -2.728362 | 6.18231899 | -0.4413169 | 0.65898358 | 0.9999244 |
| miR-3473e | 0.12132248 | -2.728362 | 6.18231899 | -0.4413169 | 0.65898358 | 0.9999244 |
| miR-7026-3p | 0.12132248 | -2.728362 | 6.18231899 | -0.4413169 | 0.65898358 | 0.9999244 |
| miR-98-3p | 0.65380206 | -2.2018238 | 5.01439204 | -0.4391008 | 0.66058847 | 0.9999244 |
| miR-743a-3p | 10.6800043 | -0.4263435 | 0.97255841 | -0.4383732 | 0.66111579 | 0.9999244 |
| miR-6911-3p | 1.14584515 | -1.6692148 | 3.82230868 | -0.4367033 | 0.66232654 | 0.9999244 |
| miR-147-5p | 13.8428067 | 0.40830142 | 0.93734167 | 0.43559508 | 0.66313054 | 0.9999244 |
| miR-7042-5p | 0.57187003 | -2.6715767 | 6.14031058 | -0.4350882 | 0.66349841 | 0.9999244 |
| miR-1956 | 0.7075449 | -2.4942176 | 5.75547115 | -0.4333646 | 0.6647499 | 0.9999244 |
| let-7b-5p | 480.909708 | 0.26619655 | 0.61616013 | 0.43202495 | 0.66572328 | 0.9999244 |
| miR-7242-5p | 6.32206472 | 0.75549466 | 1.75414145 | 0.43069198 | 0.66669235 | 0.9999244 |
| miR-5114 | 11.0690558 | 0.59901218 | 1.39403397 | 0.42969698 | 0.66741608 | 0.9999244 |
| miR-7242-3p | 19.1694289 | 0.51365118 | 1.19832373 | 0.42864141 | 0.6681842 | 0.9999244 |
| miR-7685-5p | 0.32628741 | 2.60441974 | 6.12206012 | 0.42541558 | 0.67053375 | 0.9999244 |
| miR-143-3p | 18524.9756 | -0.2954524 | 0.69696735 | -0.4239114 | 0.67163043 | 0.9999244 |
| miR-29a-3p | 39.0011119 | 0.40483625 | 0.96969036 | 0.41749023 | 0.67631987 | 0.9999244 |
| miR-1668 | 0.49396646 | -2.5161114 | 6.14755568 | -0.4092865 | 0.68232943 | 0.9999244 |
| miR-574-3p | 9.99368537 | -0.5973388 | 1.46132056 | -0.4087664 | 0.68271107 | 0.9999244 |
| miR-124-3p | 2.25959209 | 0.89315822 | 2.19043194 | 0.40775438 | 0.68345401 | 0.9999244 |
| miR-1981-5p | 0.81363115 | -1.916266 | 4.70580151 | -0.4072135 | 0.68385117 | 0.9999244 |
| miR-7217-5p | 23.1520426 | 0.52986967 | 1.3359558 | 0.39662216 | 0.69164611 | 0.9999244 |
| miR-27b-3p | 459.865937 | 0.17254988 | 0.43645145 | 0.39534726 | 0.69258663 | 0.9999244 |
| miR-148b-5p | 1.41662624 | -1.1099076 | 2.80863757 | -0.3951765 | 0.69271263 | 0.9999244 |
| miR-293-3p | 0.25461773 | -2.4162994 | 6.1909051 | -0.3902982 | 0.69631602 | 0.9999244 |
| miR-3064-3p | 0.2773443 | 2.37269666 | 6.12477694 | 0.38739315 | 0.69846517 | 0.9999244 |
| miR-5099 | 84.7086433 | -0.391182 | 1.02117034 | -0.3830723 | 0.70166619 | 0.9999244 |
| miR-322-5p | 0.79284218 | -1.6431221 | 4.32389775 | -0.3800095 | 0.70393838 | 0.9999244 |
| miR-92a-1-5p | 0.44965109 | -2.3120499 | 6.1495437 | -0.375971 | 0.70693848 | 0.9999244 |
| miR-199b-5p | 0.26929608 | 2.28687829 | 6.12589678 | 0.37331323 | 0.70891533 | 0.9999244 |
| miR-7009-5p | 0.26102993 | 2.28635478 | 6.12590382 | 0.37322734 | 0.70897925 | 0.9999244 |
| miR-369-3p | 0.54660629 | 1.94983575 | 5.22905415 | 0.37288498 | 0.70923405 | 0.9999244 |
| miR-9769-3p | 0.6205535 | 2.23139904 | 6.05010729 | 0.36881975 | 0.71226209 | 0.9999244 |
| miR-7082-5p | 0.29268979 | 2.26455978 | 6.14658157 | 0.36842589 | 0.7125557 | 0.9999244 |
| miR-181a-5p | 616.065004 | 0.2314311 | 0.6380101 | 0.36273893 | 0.71679992 | 0.9999244 |
| miR-17-3p | 0.59741414 | 1.71840383 | 4.74225371 | 0.36236016 | 0.7170829 | 0.9999244 |
| miR-3070-5p | 0.64518778 | -2.1293806 | 5.90828417 | -0.3604059 | 0.7185436 | 0.9999244 |
| miR-1843b-3p | 32.6640078 | -0.261746 | 0.72759605 | -0.3597408 | 0.71904095 | 0.9999244 |
| miR-362-5p | 0.24471556 | 2.1945244 | 6.12717704 | 0.3581624 | 0.72022179 | 0.9999244 |
| miR-6516-3p | 1.25306265 | -1.4465618 | 4.04160698 | -0.3579175 | 0.72040506 | 0.9999244 |
| miR-127-5p | 0.54338521 | 2.05464761 | 5.74568952 | 0.35759809 | 0.72064411 | 0.9999244 |
| miR-485-3p | 1.01910843 | -1.428511 | 4.00041149 | -0.357091 | 0.72102367 | 0.9999244 |

| | | | | | | |
|---|---|---|---|---|---|---|
| miR-532-5p | 35.9742044 | -0.2308613 | 0.64670552 | -0.3569805 | 0.72110638 | 0.9999244 |
| miR-511-5p | 0.75405694 | -1.6651265 | 4.70168497 | -0.3541553 | 0.72322252 | 0.9999244 |
| let-7e-5p | 645.74885 | 0.19452505 | 0.54963237 | 0.35391846 | 0.72339999 | 0.9999244 |
| miR-362-3p | 0.23916252 | 2.16038977 | 6.12767089 | 0.35256296 | 0.72441611 | 0.9999244 |
| miR-3084-3p | 3.69419631 | -0.7804063 | 2.22021041 | -0.3515011 | 0.72521248 | 0.9999244 |
| miR-149-5p | 2.78691366 | 0.85254807 | 2.46750256 | 0.34551051 | 0.72971061 | 0.9999244 |
| miR-3072-3p | 0.89318298 | -1.7161361 | 4.9964515 | -0.343471 | 0.73124416 | 0.9999244 |
| miR-483-5p | 1.26550568 | -1.2135547 | 3.57602581 | -0.3393585 | 0.73433969 | 0.9999244 |
| miR-1291 | 0.31862991 | -2.0688809 | 6.10397878 | -0.3389397 | 0.73465513 | 0.9999244 |
| miR-7648-3p | 1.1435915 | -1.4016336 | 4.14466382 | -0.3381779 | 0.73522916 | 0.9999244 |
| miR-329-5p | 0.23145904 | 2.05936551 | 6.1292012 | 0.33599248 | 0.73687654 | 0.9999244 |
| miR-1933-3p | 0.38353075 | -2.0415168 | 6.15767265 | -0.3315403 | 0.74023639 | 0.9999244 |
| miR-3079-5p | 0.28781655 | -2.0298149 | 6.14485755 | -0.3303274 | 0.74115258 | 0.9999244 |
| miR-3077-3p | 0.2189207 | 2.02276823 | 6.1297819 | 0.32999024 | 0.74140733 | 0.9999244 |
| miR-190a-5p | 0.225906 | 2.02015422 | 6.12982393 | 0.32956154 | 0.74173129 | 0.9999244 |
| miR-194-5p | 4.27779194 | 0.52453392 | 1.60056278 | 0.32771843 | 0.74312457 | 0.9999244 |
| miR-615-3p | 2.63498668 | -0.6336135 | 1.93361116 | -0.327684 | 0.74315058 | 0.9999244 |
| miR-130a-3p | 344.123953 | -0.1531851 | 0.47313214 | -0.3237682 | 0.74611353 | 0.9999244 |
| miR-7229-3p | 9.20503501 | -0.377614 | 1.16894736 | -0.3230376 | 0.74666675 | 0.9999244 |
| miR-199a-5p | 7.36944671 | 0.53643575 | 1.66594618 | 0.32200065 | 0.7474522 | 0.9999244 |
| let-7d-5p | 257.583474 | 0.14335188 | 0.44775701 | 0.32015554 | 0.74885043 | 0.9999244 |
| miR-669m-3p | 0.20749561 | 1.93478742 | 6.13123812 | 0.31556227 | 0.75233478 | 0.9999244 |
| miR-465b-5p | 134.218272 | -0.2457058 | 0.78138232 | -0.3144502 | 0.75317912 | 0.9999244 |
| miR-5113 | 0.69305464 | 1.78840992 | 5.68961309 | 0.31432892 | 0.75327124 | 0.9999244 |
| miR-1894-3p | 0.2031438 | 1.91863381 | 6.13151502 | 0.3129135 | 0.75434638 | 0.9999244 |
| miR-1936 | 0.75529474 | 1.90337826 | 6.12955386 | 0.31052476 | 0.75616193 | 0.9999244 |
| miR-1947-5p | 8.86575276 | -0.3706866 | 1.20560918 | -0.3074683 | 0.75848694 | 0.9999244 |
| miR-23b-3p | 63.4340343 | -0.1661357 | 0.54087737 | -0.3071597 | 0.75872179 | 0.9999244 |
| let-7f-5p | 17411.4308 | -0.1747197 | 0.56905937 | -0.3070324 | 0.75881868 | 0.9999244 |
| miR-152-3p | 3.69076093 | -0.6852099 | 2.23355997 | -0.3067793 | 0.75901137 | 0.9999244 |
| miR-499-5p | 0.9508207 | -1.2343896 | 4.03332947 | -0.3060473 | 0.75956862 | 0.9999244 |
| miR-7221-5p | 0.75652567 | 1.45818348 | 4.77878771 | 0.30513669 | 0.76026205 | 0.9999244 |
| miR-15b-3p | 1.2431204 | -1.100068 | 3.61870268 | -0.3039951 | 0.7611316 | 0.9999244 |
| miR-6988-5p | 0.19525535 | 1.86379167 | 6.13247794 | 0.30392146 | 0.76118772 | 0.9999244 |
| miR-673-5p | 0.41086014 | 1.71754408 | 5.71973849 | 0.30028367 | 0.76396079 | 0.9999244 |
| miR-1948-5p | 0.19465085 | 1.81161262 | 6.13342776 | 0.29536708 | 0.76771348 | 0.9999244 |
| miR-7675-3p | 0.17945807 | 1.75427709 | 6.13451068 | 0.28596854 | 0.77490221 | 0.9999244 |
| miR-223-3p | 2.14767617 | -0.7600048 | 2.65816976 | -0.2859128 | 0.7749449 | 0.9999244 |
| miR-7222-3p | 2.3458522 | -0.6563333 | 2.2980156 | -0.2856087 | 0.77517783 | 0.9999244 |
| miR-7224-3p | 0.1867624 | 1.75065781 | 6.13458046 | 0.28537531 | 0.77535662 | 0.9999244 |
| miR-8097 | 0.64813936 | 1.57235563 | 5.54197689 | 0.28371747 | 0.77662691 | 0.9999244 |
| miR-3963 | 0.39910525 | -1.4669384 | 5.24621671 | -0.2796183 | 0.77977034 | 0.9999244 |
| miR-466b-3p | 3.80569996 | 0.46735179 | 1.69822037 | 0.27520091 | 0.78316189 | 0.9999244 |
| miR-344b-3p | 0.18251593 | 1.68733812 | 6.13582902 | 0.27499758 | 0.7833181 | 0.9999244 |
| miR-449a-3p | 0.17115386 | 1.6848273 | 6.13587963 | 0.27458611 | 0.78363424 | 0.9999244 |
| miR-28c | 0.12476461 | -1.6950809 | 6.19090148 | -0.273802 | 0.78423682 | 0.9999244 |

| | | | | | | |
|---|---|---|---|---|---|---|
| miR-6971-3p | 0.21444515 | -1.6949923 | 6.1909051 | -0.2737875 | 0.78424795 | 0.9999244 |
| miR-466c-5p | 0.16928888 | -1.6949403 | 6.19089944 | -0.2737793 | 0.78425421 | 0.9999244 |
| miR-3100-5p | 0.16928888 | -1.6949403 | 6.19089944 | -0.2737793 | 0.78425421 | 0.9999244 |
| miR-450b-5p | 0.13014528 | -1.6949402 | 6.19089944 | -0.2737793 | 0.78425422 | 0.9999244 |
| let-7d-3p | 83.7901595 | -0.15727 | 0.57697394 | -0.2725773 | 0.78517813 | 0.9999244 |
| miR-331-3p | 0.17027165 | 1.66282935 | 6.13632674 | 0.27098123 | 0.78640547 | 0.9999244 |
| miR-1948-3p | 0.17027165 | 1.66282935 | 6.13632674 | 0.27098123 | 0.78640547 | 0.9999244 |
| miR-1298-5p | 0.17027165 | 1.66282935 | 6.13632674 | 0.27098123 | 0.78640547 | 0.9999244 |
| miR-345-3p | 5.44312551 | -0.5600339 | 2.0675416 | -0.2708695 | 0.78649143 | 0.9999244 |
| let-7f-1-3p | 0.16931384 | 1.6310986 | 6.13698343 | 0.26578182 | 0.79040725 | 0.9999244 |
| let-7c-5p | 5728.75798 | 0.14685642 | 0.55780233 | 0.26327681 | 0.79233723 | 0.9999244 |
| miR-1249-3p | 7.14171159 | -0.3753945 | 1.42641982 | -0.2631725 | 0.79241762 | 0.9999244 |
| miR-5107-3p | 0.17016248 | -1.6168526 | 6.15842553 | -0.2625432 | 0.7929027 | 0.9999244 |
| miR-7218-3p | 9.6558705 | 0.38399403 | 1.46777255 | 0.26161685 | 0.79361685 | 0.9999244 |
| miR-702-3p | 0.42434974 | 1.60449763 | 6.15325748 | 0.26075581 | 0.79428083 | 0.9999244 |
| miR-7233-3p | 86.2479996 | -0.1404076 | 0.54535805 | -0.2574596 | 0.79682402 | 0.9999244 |
| miR-7006-3p | 0.60423579 | 1.57549028 | 6.1354568 | 0.25678451 | 0.79734513 | 0.9999244 |
| miR-32-3p | 0.56166506 | -1.3163137 | 5.14534226 | -0.2558263 | 0.79808499 | 0.9999244 |
| miR-370-3p | 0.21255506 | 1.57194378 | 6.15388153 | 0.2554394 | 0.79838375 | 0.9999244 |
| miR-181a-2-3p | 7.44225008 | 0.40941856 | 1.60767831 | 0.25466448 | 0.79898225 | 0.9999244 |
| miR-669f-3p | 0.16506736 | 1.56059161 | 6.13849369 | 0.25423038 | 0.79931758 | 0.9999244 |
| miR-140-3p | 43.1775835 | 0.18183984 | 0.71554068 | 0.25412928 | 0.79939568 | 0.9999244 |
| miR-341-3p | 0.45586568 | -1.5394296 | 6.07504988 | -0.253402 | 0.7999576 | 0.9999244 |
| miR-27a-5p | 0.67508012 | 1.23489176 | 4.89438456 | 0.25230787 | 0.8008031 | 0.9999244 |
| miR-361-3p | 6.2232463 | -0.476342 | 1.90216137 | -0.2504214 | 0.80226146 | 0.9999244 |
| miR-7221-3p | 12.4474012 | -0.3368688 | 1.34702624 | -0.2500833 | 0.80252294 | 0.9999244 |
| miR-669p-5p | 2.94739795 | -0.4031028 | 1.63160635 | -0.2470588 | 0.80486269 | 0.9999244 |
| let-7c-1-3p | 0.2378391 | 1.51111383 | 6.15507742 | 0.24550688 | 0.80606399 | 0.9999244 |
| miR-7025-5p | 0.15186527 | 1.50543006 | 6.13972607 | 0.24519499 | 0.80630546 | 0.9999244 |
| miR-883a-3p | 19.3266994 | -0.207404 | 0.86000142 | -0.241167 | 0.80942565 | 0.9999244 |
| miR-760-3p | 0.23692155 | -1.4737485 | 6.13177718 | -0.2403461 | 0.81006199 | 0.9999244 |
| miR-5126 | 2.20662156 | -0.5126092 | 2.13471273 | -0.2401303 | 0.81022926 | 0.9999244 |
| miR-7017-3p | 0.54526664 | 1.4649439 | 6.13776557 | 0.23867707 | 0.811356 | 0.9999244 |
| miR-5620-5p | 0.54526664 | 1.4649439 | 6.13776557 | 0.23867707 | 0.811356 | 0.9999244 |
| miR-26b-5p | 124.394455 | 0.13893102 | 0.58240045 | 0.23854896 | 0.81145534 | 0.9999244 |
| miR-700-3p | 0.07697561 | -1.4546473 | 6.19090154 | -0.2349653 | 0.81423563 | 0.9999244 |
| miR-378c | 0.15231584 | -1.4545594 | 6.1909051 | -0.234951 | 0.81424676 | 0.9999244 |
| miR-130a-5p | 0.23253444 | -1.4545594 | 6.1909051 | -0.234951 | 0.81424676 | 0.9999244 |
| miR-7010-5p | 0.23253444 | -1.4545594 | 6.1909051 | -0.234951 | 0.81424676 | 0.9999244 |
| miR-3084-5p | 0.10317053 | -1.4545086 | 6.19089952 | -0.234943 | 0.81425297 | 0.9999244 |
| miR-6981-5p | 0.14594713 | 1.44233067 | 6.14119278 | 0.23486165 | 0.81431612 | 0.9999244 |
| miR-340-3p | 3.66941818 | -0.4766404 | 2.03455547 | -0.2342725 | 0.81477343 | 0.9999244 |
| miR-874-3p | 1.12678457 | -0.7268667 | 3.1251261 | -0.2325879 | 0.81608139 | 0.9999244 |
| miR-16-1-3p | 0.76760196 | 1.07337882 | 4.65289094 | 0.23069073 | 0.81755507 | 0.9999244 |
| miR-30e-5p | 167.016417 | -0.3368312 | 1.46275607 | -0.2302716 | 0.81788072 | 0.9999244 |
| miR-342-5p | 6.79951387 | 0.35965871 | 1.56971579 | 0.22912346 | 0.81877296 | 0.9999244 |

| | | | | | | |
|---|---|---|---|---|---|---|
| miR-202-3p | 0.34051606 | 1.38891809 | 6.1250581 | 0.22675999 | 0.82061038 | 0.9999244 |
| miR-6390 | 0.14068817 | 1.39195245 | 6.14240909 | 0.22661344 | 0.82072434 | 0.9999244 |
| let-7g-5p | 1132.43314 | -0.102069 | 0.45080786 | -0.2264136 | 0.82087978 | 0.9999244 |
| miR-1982-5p | 1.15280594 | 0.82138286 | 3.64898064 | 0.22509926 | 0.82190205 | 0.9999244 |
| miR-5617-3p | 0.60926336 | -1.3463458 | 6.05729769 | -0.2222684 | 0.82410496 | 0.9999244 |
| miR-7233-5p | 3.96187147 | 0.37323163 | 1.68366787 | 0.2216777 | 0.82456479 | 0.9999244 |
| miR-7210-5p | 320.93364 | 0.12130725 | 0.54973847 | 0.22066356 | 0.8253544 | 0.9999244 |
| miR-365-2-5p | 0.36655662 | 1.30808176 | 5.94581096 | 0.22000056 | 0.82587072 | 0.9999244 |
| miR-653-5p | 0.48559004 | -1.330692 | 6.10703463 | -0.217895 | 0.82751096 | 0.9999244 |
| miR-7214-5p | 17.3857301 | -0.2605476 | 1.20166552 | -0.2168221 | 0.828347 | 0.9999244 |
| miR-301b-3p | 0.78326378 | -1.0026614 | 4.65294724 | -0.2154895 | 0.82938568 | 0.9999244 |
| miR-5128 | 0.13051496 | 1.30450245 | 6.14462015 | 0.21229993 | 0.83187305 | 0.9999244 |
| miR-3057-5p | 3.32793103 | 0.42714786 | 2.01777529 | 0.21169248 | 0.83234695 | 0.9999244 |
| miR-742-5p | 0.86779662 | 0.77800053 | 3.68260152 | 0.21126384 | 0.8326814 | 0.9999244 |
| miR-3102-5p.2 | 0.16385134 | 1.30117628 | 6.15959934 | 0.21124366 | 0.83269715 | 0.9999244 |
| miR-26a-5p | 3518.98588 | 0.08698929 | 0.41511407 | 0.20955514 | 0.83401489 | 0.9999244 |
| miR-495-3p | 0.13381221 | 1.2747105 | 6.14540337 | 0.20742503 | 0.83567793 | 0.9999244 |
| miR-208b-3p | 0.16471256 | -1.2743711 | 6.16732947 | -0.2066326 | 0.83629682 | 0.9999244 |
| miR-7652-3p | 0.48387344 | 1.26537133 | 6.14240319 | 0.20600591 | 0.8367863 | 0.9999244 |
| miR-450a-5p | 0.24706883 | -1.2585855 | 6.13953627 | -0.2049968 | 0.83757459 | 0.9999244 |
| miR-196b-3p | 1.24694237 | 0.88607296 | 4.33596755 | 0.20435415 | 0.83807676 | 0.9999244 |
| miR-185-5p | 2.9484287 | -0.4919609 | 2.45392979 | -0.2004788 | 0.84110614 | 0.9999244 |
| miR-30a-3p | 101.129552 | -0.094108 | 0.47523908 | -0.1980225 | 0.84302746 | 0.9999244 |
| miR-433-3p | 0.45438887 | 1.20320273 | 6.14398209 | 0.19583435 | 0.84473982 | 0.9999244 |
| miR-106b-5p | 0.12162261 | 1.18181631 | 6.1479478 | 0.1922294 | 0.84756252 | 0.9999244 |
| miR-668-3p | 0.98864856 | 0.77965762 | 4.07598092 | 0.19128098 | 0.84830546 | 0.9999244 |
| miR-218-5p | 1.2749519 | 0.6148827 | 3.24507021 | 0.18948209 | 0.84971499 | 0.9999244 |
| miR-132-3p | 1.31694205 | 0.61513374 | 3.26163392 | 0.18859681 | 0.85040884 | 0.9999244 |
| miR-210-5p | 0.43827431 | 1.15397629 | 6.14528078 | 0.18778252 | 0.85104714 | 0.9999244 |
| miR-329-3p | 0.11899313 | 1.15428621 | 6.14859079 | 0.18773183 | 0.85108688 | 0.9999244 |
| miR-7238-5p | 0.45317684 | 1.15067632 | 6.14536941 | 0.18724282 | 0.85147025 | 0.9999244 |
| miR-328-5p | 0.45317684 | 1.15067632 | 6.14536941 | 0.18724282 | 0.85147025 | 0.9999244 |
| miR-669m-5p | 0.3916325 | 1.14459651 | 6.13243997 | 0.18664618 | 0.85193805 | 0.9999244 |
| miR-382-5p | 0.2782138 | 1.15015454 | 6.16789013 | 0.18647455 | 0.85207263 | 0.9999244 |
| miR-199a-3p | 14.4442038 | 0.18256057 | 0.98187385 | 0.18593078 | 0.85249904 | 0.9999244 |
| miR-682 | 0.8790814 | -0.7651245 | 4.13204354 | -0.1851685 | 0.85309684 | 0.9999244 |
| miR-6990-5p | 0.42753127 | 1.12029795 | 6.14619479 | 0.18227505 | 0.85536688 | 0.9999244 |
| miR-183-3p | 0.42753127 | 1.12029795 | 6.14619479 | 0.18227505 | 0.85536688 | 0.9999244 |
| miR-3475-3p | 0.42753127 | 1.12029795 | 6.14619479 | 0.18227505 | 0.85536688 | 0.9999244 |
| miR-380-3p | 0.34172456 | -1.1135725 | 6.1214576 | -0.181913 | 0.85565103 | 0.9999244 |
| miR-199b-3p | 7.49644541 | 0.20482197 | 1.18359485 | 0.17305075 | 0.86261154 | 0.9999244 |
| miR-3068-3p | 13.5648049 | 0.17441183 | 1.01047918 | 0.17260309 | 0.86296342 | 0.9999244 |
| miR-5101 | 0.10847519 | 1.05067785 | 6.1502758 | 0.17083427 | 0.86435408 | 0.9999244 |
| miR-7118-5p | 0.10847519 | 1.05067785 | 6.1502758 | 0.17083427 | 0.86435408 | 0.9999244 |
| miR-6986-5p | 0.10847519 | 1.05067785 | 6.1502758 | 0.17083427 | 0.86435408 | 0.9999244 |
| miR-7013-5p | 0.10847519 | 1.05067785 | 6.1502758 | 0.17083427 | 0.86435408 | 0.9999244 |

| | | | | | | |
|---|---|---|---|---|---|---|
| miR-5107-5p | 1.60470767 | -0.5408185 | 3.17906661 | -0.1701186 | 0.86491684 | 0.9999244 |
| miR-5125 | 0.39979633 | 1.02932767 | 6.1487714 | 0.1674038 | 0.86705234 | 0.9999244 |
| miR-7070-5p | 0.39979633 | 1.02932767 | 6.1487714 | 0.1674038 | 0.86705234 | 0.9999244 |
| miR-7073-3p | 0.39979633 | 1.02932767 | 6.1487714 | 0.1674038 | 0.86705234 | 0.9999244 |
| miR-294-3p | 0.92402984 | 0.70273148 | 4.20984927 | 0.16692557 | 0.86742861 | 0.9999244 |
| miR-25-3p | 1357.9244 | 0.07172278 | 0.43458291 | 0.16503819 | 0.86891392 | 0.9999244 |
| miR-615-5p | 4.90884478 | -0.3452951 | 2.11315447 | -0.1634027 | 0.87020138 | 0.9999244 |
| miR-7210-3p | 14.8382315 | 0.15486982 | 0.95766868 | 0.16171545 | 0.87152994 | 0.9999244 |
| miR-7232-5p | 0.65927838 | -0.734231 | 4.54774151 | -0.1614496 | 0.87173931 | 0.9999244 |
| miR-6541 | 0.29020579 | -0.9398778 | 6.18126086 | -0.1520528 | 0.87914532 | 0.9999244 |
| miR-669b-5p | 0.09759216 | 0.9308281 | 6.15237741 | 0.15129568 | 0.87974248 | 0.9999244 |
| miR-204-3p | 0.66535337 | 0.88180806 | 5.8590111 | 0.15050459 | 0.88036653 | 0.9999244 |
| miR-5100 | 0.5482102 | -0.7799018 | 5.2117918 | -0.1496418 | 0.88104725 | 0.9999244 |
| miR-9-3p | 0.27670588 | -0.8932598 | 6.15507682 | -0.1451257 | 0.88461164 | 0.9999244 |
| miR-7237-3p | 1.07297075 | 0.59148766 | 4.08139429 | 0.14492294 | 0.88477171 | 0.9999244 |
| miR-449a-5p | 84.0485271 | -0.1467236 | 1.02023189 | -0.143814 | 0.88564738 | 0.9999244 |
| miR-669l-5p | 0.75556817 | 0.65996247 | 4.59545214 | 0.14361209 | 0.8858068 | 0.9999244 |
| miR-6922-3p | 0.36351109 | 0.88313618 | 6.15326228 | 0.14352325 | 0.88587695 | 0.9999244 |
| miR-6340 | 0.36351109 | 0.88313618 | 6.15326228 | 0.14352325 | 0.88587695 | 0.9999244 |
| miR-3092-3p | 0.36351109 | 0.88313618 | 6.15326228 | 0.14352325 | 0.88587695 | 0.9999244 |
| miR-350-5p | 0.29106583 | 0.87036771 | 6.16248715 | 0.14123643 | 0.88768316 | 0.9999244 |
| miR-455-5p | 0.35813958 | 0.86307135 | 6.15391434 | 0.14024754 | 0.88846441 | 0.9999244 |
| miR-92b-3p | 72.4786355 | -0.0978772 | 0.70608177 | -0.1386201 | 0.88975033 | 0.9999244 |
| miR-7241-3p | 57.5906637 | -0.093757 | 0.67732551 | -0.1384223 | 0.88990664 | 0.9999244 |
| miR-217-3p | 0.12167728 | 0.85064008 | 6.1538811 | 0.13822823 | 0.89006005 | 0.9999244 |
| miR-7021-5p | 0.12167728 | 0.85064008 | 6.1538811 | 0.13822823 | 0.89006005 | 0.9999244 |
| miR-7054-5p | 0.12167728 | 0.85064008 | 6.1538811 | 0.13822823 | 0.89006005 | 0.9999244 |
| miR-3473d | 0.12167728 | 0.85064008 | 6.1538811 | 0.13822823 | 0.89006005 | 0.9999244 |
| miR-5626-3p | 0.12167728 | 0.85064008 | 6.1538811 | 0.13822823 | 0.89006005 | 0.9999244 |
| miR-7646-5p | 0.12167728 | 0.85064008 | 6.1538811 | 0.13822823 | 0.89006005 | 0.9999244 |
| miR-330-5p | 1.08453659 | -0.5936138 | 4.29558753 | -0.1381915 | 0.89008904 | 0.9999244 |
| miR-466c-3p | 1.50123109 | 0.32963247 | 2.39363734 | 0.13771195 | 0.89046807 | 0.9999244 |
| miR-128-1-5p | 0.81916492 | 0.64214484 | 4.67824905 | 0.13726179 | 0.89082387 | 0.9999244 |
| miR-692 | 0.24139402 | -0.8302548 | 6.18637047 | -0.1342071 | 0.89323881 | 0.9999244 |
| miR-10b-5p | 392080.628 | 0.06046567 | 0.46198419 | 0.13088255 | 0.89586822 | 0.9999244 |
| miR-8104 | 0.08678015 | 0.78979278 | 6.15507731 | 0.12831566 | 0.89789918 | 0.9999244 |
| miR-203-5p | 0.08678015 | 0.78979278 | 6.15507731 | 0.12831566 | 0.89789918 | 0.9999244 |
| miR-6940-3p | 0.08678015 | 0.78979278 | 6.15507731 | 0.12831566 | 0.89789918 | 0.9999244 |
| miR-3547-5p | 0.08678015 | 0.78979278 | 6.15507731 | 0.12831566 | 0.89789918 | 0.9999244 |
| miR-7216-3p | 0.08678015 | 0.78979278 | 6.15507731 | 0.12831566 | 0.89789918 | 0.9999244 |
| miR-7668-5p | 0.08678015 | 0.78979278 | 6.15507731 | 0.12831566 | 0.89789918 | 0.9999244 |
| miR-6236 | 0.08678015 | 0.78979278 | 6.15507731 | 0.12831566 | 0.89789918 | 0.9999244 |
| miR-5122 | 1.07048901 | -0.4503915 | 3.67193172 | -0.1226579 | 0.902378 | 0.9999244 |
| miR-204-5p | 276.046951 | 0.06259533 | 0.51258525 | 0.12211692 | 0.90280642 | 0.9999244 |
| miR-141-3p | 43.6745166 | 0.10377186 | 0.88796939 | 0.11686423 | 0.90696764 | 0.9999244 |
| let-7e-3p | 0.13426785 | 0.72073503 | 6.17593784 | 0.1167005 | 0.90709739 | 0.9999244 |

| | | | | | | |
|---|---|---|---|---|---|---|
| miR-218-1-3p | 0.31983706 | 0.71168699 | 6.1591321 | 0.11554988 | 0.90800928 | 0.9999244 |
| miR-30c-5p | 1036.86164 | -0.0416752 | 0.3669918 | -0.1135588 | 0.90958753 | 0.9999244 |
| miR-206-3p | 1.83282319 | 0.33904326 | 2.99857522 | 0.11306812 | 0.90997655 | 0.9999244 |
| miR-106b-3p | 71.543458 | 0.06140056 | 0.55257435 | 0.11111729 | 0.91152334 | 0.9999244 |
| miR-468-3p | 6.10232842 | -0.1742385 | 1.57304894 | -0.1107648 | 0.91180284 | 0.9999244 |
| miR-7215-5p | 0.27346293 | 0.65772575 | 6.13722735 | 0.10716985 | 0.91465423 | 0.9999244 |
| miR-326-3p | 0.9228141 | 0.48576261 | 4.63410983 | 0.10482328 | 0.91651603 | 0.9999244 |
| miR-7118-3p | 0.56316804 | -0.6035661 | 6.09988706 | -0.0989471 | 0.92118027 | 0.9999244 |
| miR-32-5p | 2.22823869 | -0.2386746 | 2.44871314 | -0.0974694 | 0.92235363 | 0.9999244 |
| miR-1927 | 0.67772998 | 0.47406708 | 4.86534874 | 0.09743743 | 0.92237902 | 0.9999244 |
| miR-297c-3p | 0.50000017 | -0.3696414 | 3.87124436 | -0.0954839 | 0.9239305 | 0.9999244 |
| miR-297b-3p | 0.50000017 | -0.3696414 | 3.87124436 | -0.0954839 | 0.9239305 | 0.9999244 |
| miR-7032-3p | 0.07297357 | 0.57985502 | 6.15959922 | 0.09413843 | 0.92499919 | 0.9999244 |
| miR-7059-5p | 0.07297357 | 0.57985502 | 6.15959922 | 0.09413843 | 0.92499919 | 0.9999244 |
| miR-1306-3p | 0.35422355 | 0.56800301 | 6.11653699 | 0.0928635 | 0.92601201 | 0.9999244 |
| miR-30d-3p | 4.34013111 | -0.1669403 | 1.80175418 | -0.0926543 | 0.9261782 | 0.9999244 |
| miR-466e-3p | 0.3926779 | -0.3990438 | 4.41371015 | -0.0904101 | 0.92796136 | 0.9999244 |
| miR-466p-3p | 0.3926779 | -0.3990438 | 4.41371015 | -0.0904101 | 0.92796136 | 0.9999244 |
| miR-466a-3p | 0.3926779 | -0.3990438 | 4.41371015 | -0.0904101 | 0.92796136 | 0.9999244 |
| miR-30a-5p | 3490.53891 | -0.0641831 | 0.72984953 | -0.0879401 | 0.92992426 | 0.9999244 |
| miR-27b-5p | 1.90329209 | -0.2596878 | 3.1317871 | -0.08292 | 0.93391515 | 0.9999244 |
| miR-465a-5p | 68.8682584 | 0.05721712 | 0.69332421 | 0.08252578 | 0.93422862 | 0.9999244 |
| miR-344d-3p | 0.12847132 | -0.5040854 | 6.16253751 | -0.0817984 | 0.93480706 | 0.9999244 |
| miR-3103-5p | 0.27263332 | 0.49468678 | 6.16604837 | 0.08022752 | 0.9360563 | 0.9999244 |
| miR-504-5p | 0.27263332 | 0.49468678 | 6.16604837 | 0.08022752 | 0.9360563 | 0.9999244 |
| miR-34a-5p | 3.37585197 | 0.18026624 | 2.24793561 | 0.0801919 | 0.93608463 | 0.9999244 |
| miR-7015-3p | 0.20810263 | -0.4839929 | 6.14643999 | -0.0787436 | 0.93723656 | 0.9999244 |
| miR-6970-5p | 0.20933333 | 0.48058937 | 6.14119241 | 0.07825669 | 0.93762387 | 0.9999244 |
| miR-449c-5p | 1.29789064 | 0.23746002 | 3.13704059 | 0.07569555 | 0.93966131 | 0.9999244 |
| miR-467e-3p | 0.1311008 | -0.4621508 | 6.16150204 | -0.0750062 | 0.94020978 | 0.9999244 |
| miR-133b-3p | 0.118135 | 0.46098545 | 6.18400582 | 0.0745448 | 0.9405769 | 0.9999244 |
| miR-1931 | 0.06508512 | 0.45765684 | 6.16253787 | 0.07426435 | 0.94080005 | 0.9999244 |
| miR-6972-3p | 0.06508512 | 0.45765684 | 6.16253787 | 0.07426435 | 0.94080005 | 0.9999244 |
| miR-10a-3p | 2.983913 | -0.1822793 | 2.557103 | -0.0712835 | 0.94317213 | 0.9999244 |
| miR-7090-5p | 0.25651876 | 0.42879094 | 6.16789504 | 0.06951982 | 0.94457586 | 0.9999244 |
| miR-6960-5p | 0.67675021 | 0.33214606 | 4.89258903 | 0.06788759 | 0.94587512 | 0.9999244 |
| miR-196a-1-3[illegible] | 0.3021179 | 0.41717006 | 6.16822915 | 0.06763206 | 0.94607853 | 0.9999244 |
| miR-673-3p | 0.3021179 | 0.41717006 | 6.16822915 | 0.06763206 | 0.94607853 | 0.9999244 |
| miR-719 | 0.3021179 | 0.41717006 | 6.16822915 | 0.06763206 | 0.94607853 | 0.9999244 |
| miR-34b-5p | 24.8292741 | 0.13590504 | 2.0444405 | 0.06647542 | 0.94699933 | 0.9999244 |
| miR-879-5p | 0.17702488 | 0.40781446 | 6.14898908 | 0.0663222 | 0.94712131 | 0.9999244 |
| miR-30b-5p | 328.954044 | -0.0300482 | 0.46887262 | -0.064086 | 0.94890172 | 0.9999244 |
| miR-465c-3p | 161.520091 | -0.0379266 | 0.61910454 | -0.0612604 | 0.95115184 | 0.9999244 |
| miR-18a-5p | 6.18124308 | 0.09597242 | 1.62009807 | 0.05923865 | 0.95276203 | 0.9999244 |
| miR-546 | 0.2398778 | 0.35664103 | 6.17001152 | 0.05780233 | 0.95390608 | 0.9999244 |
| miR-134-3p | 0.2398778 | 0.35664103 | 6.17001152 | 0.05780233 | 0.95390608 | 0.9999244 |

| | | | | | | |
|---|---|---|---|---|---|---|
| miR-412-5p | 0.2398778 | 0.35664103 | 6.17001152 | 0.05780233 | 0.95390608 | 0.9999244 |
| miR-6992-5p | 0.2398778 | 0.35664103 | 6.17001152 | 0.05780233 | 0.95390608 | 0.9999244 |
| miR-1960 | 0.2398778 | 0.35664103 | 6.17001152 | 0.05780233 | 0.95390608 | 0.9999244 |
| miR-7028-5p | 0.2398778 | 0.35664103 | 6.17001152 | 0.05780233 | 0.95390608 | 0.9999244 |
| miR-330-3p | 0.2398778 | 0.35664103 | 6.17001152 | 0.05780233 | 0.95390608 | 0.9999244 |
| miR-138-5p | 0.2398778 | 0.35664103 | 6.17001152 | 0.05780233 | 0.95390608 | 0.9999244 |
| miR-3109-3p | 0.2398778 | 0.35664103 | 6.17001152 | 0.05780233 | 0.95390608 | 0.9999244 |
| miR-465a-3p | 81.2636666 | -0.032482 | 0.64966322 | -0.0499983 | 0.96012378 | 0.9999244 |
| miR-212-3p | 2.43251707 | -0.1349775 | 2.73689095 | -0.0493178 | 0.960666 | 0.9999244 |
| miR-7043-3p | 0.70262989 | 0.23662906 | 4.96550076 | 0.04765462 | 0.9619915 | 0.9999244 |
| miR-297a-3p | 1.07183877 | -0.1229085 | 2.7847938 | -0.0441356 | 0.96479634 | 0.9999244 |
| miR-465b-3p | 162.106103 | -0.0270539 | 0.61899534 | -0.0437062 | 0.96513863 | 0.9999244 |
| miR-296-3p | 0.32202775 | 0.2475332 | 6.11484038 | 0.04048073 | 0.96770987 | 0.9999244 |
| miR-700-5p | 0.74366835 | -0.1776204 | 4.42982349 | -0.0400965 | 0.9680162 | 0.9999244 |
| miR-7073-5p | 0.78357044 | 0.20315518 | 5.19403924 | 0.03911314 | 0.96880019 | 0.9999244 |
| miR-22-5p | 0.53459881 | -0.1541137 | 4.50228367 | -0.0342301 | 0.97269365 | 0.9999244 |
| miR-181c-3p | 0.7050816 | 0.15211275 | 4.46355832 | 0.03407881 | 0.97281431 | 0.9999244 |
| miR-125a-3p | 3.50064133 | -0.076427 | 2.63984456 | -0.0289513 | 0.97690341 | 0.9999244 |
| miR-1839-5p | 125.16915 | 0.01999353 | 0.69648359 | 0.0287064 | 0.97709875 | 0.9999244 |
| miR-3064-5p | 1.90725672 | 0.16103808 | 6.04724678 | 0.02662998 | 0.97875486 | 0.9999244 |
| miR-25-5p | 0.55995929 | 0.1618287 | 6.15326208 | 0.02629966 | 0.97901833 | 0.9999244 |
| miR-8111 | 0.53598151 | 0.15855737 | 6.14619457 | 0.02579765 | 0.97941874 | 0.9999244 |
| miR-186-5p | 151.5697 | 0.01347011 | 0.55618413 | 0.02421879 | 0.98067809 | 0.9999244 |
| miR-467a-5p | 10.9428867 | -0.0234734 | 0.9740294 | -0.0240993 | 0.98077343 | 0.9999244 |
| miR-28a-5p | 6.70795392 | 0.02855373 | 1.27278648 | 0.02243403 | 0.98210174 | 0.9999244 |
| miR-1843a-5p | 6.54927118 | -0.0264578 | 1.37342685 | -0.0192641 | 0.98463042 | 0.9999244 |
| miR-300-5p | 0.04864904 | 0.1152384 | 6.1721602 | 0.01867068 | 0.98510382 | 0.9999244 |
| miR-1943-5p | 0.23012752 | -0.1110596 | 6.15387999 | -0.0180471 | 0.98560129 | 0.9999244 |
| miR-193a-5p | 0.15991853 | -0.108947 | 6.1863707 | -0.0176108 | 0.98594933 | 0.9999244 |
| miR-665-3p | 0.15991853 | -0.108947 | 6.1863707 | -0.0176108 | 0.98594933 | 0.9999244 |
| miR-874-5p | 0.15991853 | -0.108947 | 6.1863707 | -0.0176108 | 0.98594933 | 0.9999244 |
| miR-3090-3p | 0.15991853 | -0.108947 | 6.1863707 | -0.0176108 | 0.98594933 | 0.9999244 |
| miR-7007-3p | 0.15991853 | -0.108947 | 6.1863707 | -0.0176108 | 0.98594933 | 0.9999244 |
| miR-20a-3p | 0.81996891 | -0.0818594 | 5.0218258 | -0.0163007 | 0.98699448 | 0.9999244 |
| miR-7234-3p | 30.7430264 | 0.01239298 | 0.85010766 | 0.01457813 | 0.98836875 | 0.9999244 |
| miR-6516-5p | 0.21692543 | 0.08893418 | 6.15027557 | 0.01446019 | 0.98846284 | 0.9999244 |
| miR-23a-3p | 88.1362464 | 0.00895488 | 0.63084832 | 0.01419497 | 0.98867443 | 0.9999244 |
| miR-135a-5p | 0.43928871 | 0.07834756 | 5.65171779 | 0.01386261 | 0.98893959 | 0.9999244 |
| miR-322-3p | 0.64858961 | 0.05244985 | 4.45085924 | 0.01178421 | 0.99059778 | 0.9999244 |
| miR-672-3p | 0.69653624 | -0.0378484 | 5.36440668 | -0.0070555 | 0.9943706 | 0.9999244 |
| miR-5119 | 0.17101251 | -0.0406238 | 6.18364776 | -0.0065696 | 0.99475829 | 0.9999244 |
| miR-7648-5p | 0.17101251 | -0.0406238 | 6.18364776 | -0.0065696 | 0.99475829 | 0.9999244 |
| miR-877-3p | 0.17101251 | -0.0406238 | 6.18364776 | -0.0065696 | 0.99475829 | 0.9999244 |
| miR-375-3p | 1526.53454 | 0.00296839 | 0.48915308 | 0.00606843 | 0.99515812 | 0.9999244 |
| miR-467b-5p | 0.9405871 | 0.01454775 | 2.71253663 | 0.00536315 | 0.99572084 | 0.9999244 |
| miR-669o-3p | 0.21123546 | 0.02533323 | 6.16766138 | 0.00410743 | 0.99672275 | 0.9999244 |

| | | | | | | |
|---|---|---|---|---|---|---|
| miR-540-3p | 0.80058379 | 0.01421235 | 3.51960795 | 0.00403805 | 0.99677811 | 0.9999244 |
| miR-7075-5p | 0.18175555 | 0.02186323 | 6.18126114 | 0.00353702 | 0.99717787 | 0.9999244 |
| miR-201-5p | 0.18175555 | 0.02186323 | 6.18126114 | 0.00353702 | 0.99717787 | 0.9999244 |
| miR-377-5p | 0.1016543 | -0.011887 | 6.19089987 | -0.0019201 | 0.998468 | 0.9999244 |
| miR-871-3p | 944.913414 | 0.00029536 | 0.42379892 | 0.00069694 | 0.99944393 | 0.9999244 |
| miR-6916-5p | 0.04339008 | -0.0005852 | 6.17593764 | -9.48E-05 | 0.9999244 | 0.9999244 |
| miR-217-5p | 0.04339008 | -0.0005852 | 6.17593764 | -9.48E-05 | 0.9999244 | 0.9999244 |
| miR-301a-5p | 0.04339008 | -0.0005852 | 6.17593764 | -9.48E-05 | 0.9999244 | 0.9999244 |
| miR-6931-3p | 0.04339008 | -0.0005852 | 6.17593764 | -9.48E-05 | 0.9999244 | 0.9999244 |
| miR-377-3p | 0.04339008 | -0.0005852 | 6.17593764 | -9.48E-05 | 0.9999244 | 0.9999244 |
| miR-369-5p | 0.04339008 | -0.0005852 | 6.17593764 | -9.48E-05 | 0.9999244 | 0.9999244 |
| miR-7218-5p | 0.04339008 | -0.0005852 | 6.17593764 | -9.48E-05 | 0.9999244 | 0.9999244 |
| miR-6985-3p | 0.04339008 | -0.0005852 | 6.17593764 | -9.48E-05 | 0.9999244 | 0.9999244 |
| miR-103-3p | 203.394758 | 1.55962481 | 0.82439317 | 1.89184587 | NA | NA |
| miR-378a-3p | 17.4797114 | -1.0398491 | 1.28656646 | -0.8082358 | NA | NA |
| miR-107-3p | 66.4487281 | 2.06760163 | 1.02257577 | 2.02195445 | NA | NA |
| miR-463-5p | 236.689323 | -0.4625632 | 0.83496126 | -0.5539936 | NA | NA |

| | | | | | | |
|---|---|---|---|---|---|---|
| q-PCR ARG-CCT-2 | | q-PCR ARG-CCT-2 | | q-PCR ARG-CCT-2 | | |
| File Name | 20201022_130950_CT027050__SYBR_MIRCUR.pcrd | | | | | |
| Created By User | | | | | | |
| Notes | | | | | | |
| ID | | | | | | |
| Run Started | 10/22/2020 21:11:00 UTC | | | | | |
| Run Ended | 10/22/2020 23:14:53 UTC | | | | | |
| Sample Vol | 8 | | | | | |
| Lid Temp | 95 | | | | | |
| Protocol File | Unknown.prcl | | | | | |
| Plate Setup F | DefaultPlate.pltd | | | | | |
| Base Serial N | CT027050 | | | | | |
| Optical Head | 786BR04591 | | | | | |
| CFX Maestro | 4.1.2433.1219. | | | | | |
| Well group | All Wells | | | | | |
| Amplificatio | 3 | | | | | |
| Melt step | 5 | | | | | |
| Well | Fluor | Sample | Target | Cq | location | treatment |
| A19 | SYBR | 19 | ArCCT2 | 28.2098782 | cauda | dex |
| A20 | SYBR | 19 | ArCCT2 | 28.2867647 | cauda | dex |
| A21 | SYBR | 19 | ArCCT2 | 28.2478036 | cauda | dex |
| B19 | SYBR | 20 | ArCCT2 | 29.2999789 | cauda | dex |
| B20 | SYBR | 20 | ArCCT2 | 29.2975449 | cauda | dex |
| B21 | SYBR | 20 | ArCCT2 | 29.2816311 | cauda | dex |
| C19 | SYBR | 21 | ArCCT2 | 30.3740823 | cauda | dex |
| C20 | SYBR | 21 | ArCCT2 | 30.271482 | cauda | dex |
| C21 | SYBR | 21 | ArCCT2 | 30.2967472 | cauda | dex |
| D19 | SYBR | 22 | ArCCT2 | 30.6366676 | cauda | dex |
| D20 | SYBR | 22 | ArCCT2 | 30.6454238 | cauda | dex |
| D21 | SYBR | 22 | ArCCT2 | 30.6923239 | cauda | dex |
| F19 | SYBR | 24 | ArCCT2 | 28.7498383 | cauda | con |
| F20 | SYBR | 24 | ArCCT2 | 28.7032726 | cauda | con |
| F21 | SYBR | 24 | ArCCT2 | 28.7943957 | cauda | con |
| G19 | SYBR | 25 | ArCCT2 | 37.1792466 | cauda | con |
| G20 | SYBR | 25 | ArCCT2 | 37.0074551 | cauda | con |
| G21 | SYBR | 25 | ArCCT2 | 37.0596246 | cauda | con |
| H19 | SYBR | 26 | ArCCT2 | 37.5324818 | cauda | con |
| H20 | SYBR | 26 | ArCCT2 | 37.2848173 | cauda | con |
| H21 | SYBR | 26 | ArCCT2 | 37.703623 | cauda | con |
| I19 | SYBR | 27 | ArCCT2 | 38.5575068 | cauda | con |
| I20 | SYBR | 27 | ArCCT2 | 38.4399242 | cauda | con |
| I21 | SYBR | 27 | ArCCT2 | 38.6528506 | cauda | con |
| J19 | SYBR | 28 | ArCCT2 | 40.1996679 | cauda | con |
| J20 | SYBR | 28 | ArCCT2 | 40.024747 | cauda | con |
| J21 | SYBR | 28 | ArCCT2 | 40.2294765 | cauda | con |

| | | | | | | |
|---|---|---|---|---|---|---|
| K19 | SYBR | 29 | ArCCT2 | 40.369128 | caput | dex |
| K20 | SYBR | 29 | ArCCT2 | 40.1978648 | caput | dex |
| K21 | SYBR | 29 | ArCCT2 | 39.9124594 | caput | dex |
| L19 | SYBR | 30 | ArCCT2 | 37.2877977 | caput | dex |
| L20 | SYBR | 30 | ArCCT2 | 37.0486182 | caput | dex |
| L21 | SYBR | 30 | ArCCT2 | 37.0535183 | caput | dex |
| N19 | SYBR | 32 | ArCCT2 | 32.1826726 | caput | dex |
| N20 | SYBR | 32 | ArCCT2 | 31.6480699 | caput | dex |
| N21 | SYBR | 32 | ArCCT2 | 32.07918 | caput | dex |
| O19 | SYBR | 33 | ArCCT2 | 35.1956435 | caput | dex |
| O20 | SYBR | 33 | ArCCT2 | 35.5422798 | caput | dex |
| O21 | SYBR | 33 | ArCCT2 | 35.0558139 | caput | dex |
| P19 | SYBR | 34 | ArCCT2 | 36.3702654 | caput | con |
| P20 | SYBR | 34 | ArCCT2 | 36.2133965 | caput | con |
| P21 | SYBR | 34 | ArCCT2 | 36.1373413 | caput | con |
| A22 | SYBR | 35 | ArCCT2 | 35.9419236 | caput | con |
| A23 | SYBR | 35 | ArCCT2 | 35.952297 | caput | con |
| A24 | SYBR | 35 | ArCCT2 | 35.9395674 | caput | con |
| B22 | SYBR | 36 | ArCCT2 | 33.4862946 | caput | con |
| B23 | SYBR | 36 | ArCCT2 | 33.5109333 | caput | con |
| B24 | SYBR | 36 | ArCCT2 | 33.6265019 | caput | con |
| C22 | SYBR | 37 | ArCCT2 | 38.5600538 | caput | con |
| C23 | SYBR | 37 | ArCCT2 | 39.2110809 | caput | con |
| C24 | SYBR | 37 | ArCCT2 | 38.7246656 | caput | con |
| D22 | SYBR | 38 | ArCCT2 | 36.7884502 | caput | con |
| D23 | SYBR | 38 | ArCCT2 | 36.9899836 | caput | con |
| D24 | SYBR | 38 | ArCCT2 | 36.5163583 | caput | con |
| A01 | SYBR | 19 | rnu | 21.7134895 | cauda | dex |
| A02 | SYBR | 19 | rnu | 21.6767274 | cauda | dex |
| A03 | SYBR | 19 | rnu | 21.8072906 | cauda | dex |
| B01 | SYBR | 20 | rnu | 23.0751087 | cauda | dex |
| B02 | SYBR | 20 | rnu | 22.9133109 | cauda | dex |
| B03 | SYBR | 20 | rnu | 23.0405274 | cauda | dex |
| C01 | SYBR | 21 | rnu | 22.193542 | cauda | dex |
| C02 | SYBR | 21 | rnu | 22.0426328 | cauda | dex |
| C03 | SYBR | 21 | rnu | 21.9780556 | cauda | dex |
| D01 | SYBR | 22 | rnu | 22.1163447 | cauda | dex |
| D02 | SYBR | 22 | rnu | 22.2360917 | cauda | dex |
| D03 | SYBR | 22 | rnu | 22.1988217 | cauda | dex |
| F01 | SYBR | 24 | Rnu | 21.6307178 | cauda | con |
| F02 | SYBR | 24 | Rnu | 21.8135557 | cauda | con |
| F03 | SYBR | 24 | Rnu | 21.8517386 | cauda | con |
| G01 | SYBR | 25 | Rnu | 28.1461182 | cauda | con |
| G02 | SYBR | 25 | Rnu | 28.2219395 | cauda | con |
| G03 | SYBR | 25 | Rnu | 28.1431924 | cauda | con |
| H01 | SYBR | 26 | Rnu | 28.6733126 | cauda | con |
| H02 | SYBR | 26 | Rnu | 28.5871014 | cauda | con |
| H03 | SYBR | 26 | Rnu | 28.6423902 | cauda | con |

| | | | | | | |
|---|---|---|---|---|---|---|
| I01 | SYBR | 27 | Rnu | 28.5066562 | cauda | con |
| I02 | SYBR | 27 | Rnu | 28.7160174 | cauda | con |
| I03 | SYBR | 27 | Rnu | 28.7184343 | cauda | con |
| J01 | SYBR | 28 | Rnu | 28.4197119 | cauda | con |
| J02 | SYBR | 28 | Rnu | 28.4034915 | cauda | con |
| J03 | SYBR | 28 | Rnu | 28.2528356 | cauda | con |
| K01 | SYBR | 29 | Rnu | 24.8325175 | caput | dex |
| K02 | SYBR | 29 | Rnu | 24.5825323 | caput | dex |
| K03 | SYBR | 29 | Rnu | 24.7648725 | caput | dex |
| L01 | SYBR | 30 | Rnu | 22.58569 | caput | dex |
| L02 | SYBR | 30 | Rnu | 22.642813 | caput | dex |
| L03 | SYBR | 30 | Rnu | 22.5845381 | caput | dex |
| N01 | SYBR | 32 | Rnu | 19.7493271 | caput | dex |
| N02 | SYBR | 32 | Rnu | 19.5208021 | caput | dex |
| N03 | SYBR | 32 | Rnu | 19.7751695 | caput | dex |
| O01 | SYBR | 33 | Rnu | 22.5797257 | caput | dex |
| O02 | SYBR | 33 | Rnu | 22.7297704 | caput | dex |
| O03 | SYBR | 33 | Rnu | 22.6687455 | caput | dex |
| P01 | SYBR | 34 | Rnu | 24.1646463 | caput | con |
| P02 | SYBR | 34 | Rnu | 24.3068973 | caput | con |
| P03 | SYBR | 34 | Rnu | 24.2134487 | caput | con |
| A04 | SYBR | 35 | rnu | 24.8087321 | caput | con |
| A05 | SYBR | 35 | rnu | 24.8499021 | caput | con |
| A06 | SYBR | 35 | rnu | 24.7083429 | caput | con |
| B04 | SYBR | 36 | rnu | 21.0468771 | caput | con |
| B05 | SYBR | 36 | rnu | 21.1630316 | caput | con |
| B06 | SYBR | 36 | rnu | 21.1637841 | caput | con |
| C04 | SYBR | 37 | rnu | 20.7726061 | caput | con |
| C05 | SYBR | 37 | rnu | 20.599424 | caput | con |
| C06 | SYBR | 37 | rnu | 20.6869102 | caput | con |
| D04 | SYBR | 38 | rnu | 24.1717524 | caput | con |
| D05 | SYBR | 38 | rnu | 23.7070299 | caput | con |
| D06 | SYBR | 38 | rnu | 23.4634021 | caput | con |

| | Control | Control | Control | Control | Control | Treated |
|---|---|---|---|---|---|---|
| Cauda sperm | **293.994297** | 77.2671768 | 79.3836466 | 38.8654931 | 10.4893868 | 406.702353 |
| Caput sperm | 9.00965721 | 16.312118 | 6.80595149 | 0.12832337 | 4.59247933 | 0.84107082 |

| Treated | Treated | Treated | Treated |
| --- | --- | --- | --- |
| 477.737146 | 122.852395 | 104.625459 | |
| 1.57772397 | | 7.439821 | 5.97239925 |
