## Supplementary Table 3 for "Single paternal Dexamethasone challenge programs offspring metabolism and reveals circRNAs as novel candidates in RNA-mediated inheritance"

| BMI | BMI | BMI | BMI | BMI | BMI |
|---|---|---|---|---|---|
| | | Females Vehicle | Females Vehicle | Females Vehicle | Females Vehicle |
| Birth | 23.May.18 | | | | |
| 3 weeks | 21.Jun.18 | 0.283355226 | 0.313581315 | 0.345993757 | 0.319979188 |
| 5 weeks | 05.Jul.18 | 0.323121475 | 0.309339679 | 0.319750149 | 0.328125 |
| 8 weeks | 27.Jul.18 | 0.286197319 | 0.286197319 | 0.272913899 | 0.280246914 |
| 12 weeks | 24.Aug.18 | 0.310341163 | 0.335708248 | 0.287098299 | 0.289196676 |
| | | males Vehicle | males Vehicle | males Vehicle | males Vehicle |
| Birth | 23.May.18 | | | | |
| 3 weeks | 21.Jun.18 | 0.287641341 | 0.310859063 | 0.286705625 | 0.365636147 |
| 5 weeks | 05.Jul.18 | 0.362629758 | 0.3515625 | 0.353604633 | 0.339671941 |
| 8 weeks | 27.Jul.18 | 0.319010417 | 0.303304663 | 0.32345679 | 0.376984127 |
| 12 weeks | 24.Aug.18 | 0.366753472 | 0.384789498 | 0.327977839 | 0.395408163 |

| BMI Females Vehicle | BMI Females Vehicle | BMI Females Vehicle | BMI Females Vehicle | BMI Females Vehicle |
|---|---|---|---|---|
| 0.307418133 | 0.295918367 | 0.304783951 | 0.325239474 | 0.318367347 |
| 0.318262939 | 0.296124256 | 0.34129146 | 0.308171745 | 0.33021542 |
| 0.300878099 | 0.265831758 | 0.280267643 | 0.264721074 | 0.34691358 |
| 0.317816635 | 0.287894554 | 0.330864198 | 0.30551698 | 0.349173315 |

| males Vehicle | males Vehicle | males Vehicle | males Vehicle | males Vehicle |
|---|---|---|---|---|
| 0.321893491 | 0.289792388 | 0.286960514 | 0.233747261 | 0.251020408 |
| 0.390311419 | 0.33293698 | 0.353955979 | 0.346519929 | 0.326169791 |
| 0.361328125 | 0.351851852 | 0.341975309 | 0.374483471 | 0.358542604 |
| 0.415581597 | 0.417823934 | 0.342915346 | 0.435937689 | 0.340166205 |

| BMI Females Vehicle | BMI Females Vehicle | BMI Females Vehicle | BMI Females Vehicle | BMI Females Vehicle |
|---|---|---|---|---|
| 0.303511208 | 0.283558793 | 0.24994749 | 0.266750683 | 0.265820274 |
| 0.30625 | 0.297441999 | 0.3109375 | 0.314052235 | 0.30625 |
| 0.260838063 | 0.294809689 | 0.297265161 | 0.253592561 | 0.257543239 |
| 0.318404016 | 0.289461248 | 0.281366985 | 0.278951817 | 0.304311074 |
| males Vehicle | males Vehicle | males Vehicle | males Vehicle | males Vehicle |
| 0.253917873 | 0.281141869 | 0.266750683 | 0.275482094 | 0.249950407 |
| 0.37037037 | 0.368530684 | 0.307643006 | 0.344387755 | 0.346929888 |
| 0.341783612 | 0.327160494 | 0.308650519 | 0.347365702 | 0.339330999 |
| 0.405160706 | 0.388839793 | 0.365432099 | 0.386314859 | 0.397295013 |

| | | | | |
|---|---|---|---|---|
| BMI | BMI | BMI | BMI | BMI |
| Females Vehicle | Females Vehicle | Females Vehicle | | |
| 0.261333333 | 0.264589979 | 0.285714286 | | |
| 0.31462585 | 0.337596669 | 0.336838897 | | |
| 0.297530864 | 0.281190926 | 0.24898144 | | |
| 0.334357278 | 0.333175803 | 0.314622001 | | |
| males Vehicle | males Vehicle | males Vehicle | males Vehicle | males Vehicle |
| 0.269387755 | 0.266466504 | 0.259869074 | 0.291836735 | 0.268132716 |
| 0.368530684 | 0.334948097 | 0.312314099 | 0.346020761 | 0.351433479 |
| 0.337078652 | 0.378357438 | 0.348639456 | 0.337611285 | 0.341783612 |
| 0.368313006 | 0.401234568 | 0.40879017 | 0.38252603 | 0.402897239 |

| BMI | BMI | BMI | BMI | BMI | BMI |
|---|---|---|---|---|---|
| | | Females Dex | Females Dex | Females Dex | Females Dex |
| | | 0.3686088 | 0.35277778 | 0.36483769 | 0.39833532 |
| | | 0.328473 | 0.34972299 | 0.33859303 | 0.35493827 |
| | | 0.32951081 | 0.32370446 | 0.32387543 | |
| | | 0.31481482 | 0.31708284 | 0.30864198 | |

| males Vehicle | males Vehicle | males Dex | males Dex | males Dex | males Dex |
|---|---|---|---|---|---|
| 0.273755207 | 0.268132716 | 0.37627551 | 0.346681 | 0.350474 | 0.36111111 |
| 0.3625 | 0.323704456 | 0.2984375 | 0.36982249 | 0.3609375 | 0.3703125 |
| 0.344161333 | 0.297731569 | 0.30172958 | 0.36422903 | 0.383391 | 0.37858302 |
| 0.472165197 | 0.405245747 | 0.35925926 | 0.40740741 | 0.35916824 | 0.34349031 |

| BMI Females Dex | BMI Females Dex | BMI Females Dex | BMI Females Dex | BMI Females Dex | BMI Females Dex |
|---|---|---|---|---|---|
| 0.32249937 | 0.34528848 | 0.331360947 | 0.40725327 | 0.304557866 | 0.33203125 |
| 0.32375346 | 0.31509695 | 0.35891684 | 0.490306041 | 0.340625 | 0.325518637 |
| 0.30773697 | 0.31975015 | 0.356949703 | 0.317460317 | 0.364152893 | 0.296193772 |
| 0.31234568 | 0.34072472 | 0.358987603 | 0.322424828 | 0.379234594 | 0.311798124 |

| males Dex | males Dex | males Dex | males Dex | males Dex | males Dex |
|---|---|---|---|---|---|
| 0.448 | 0.32214506 | 0.300925926 | 0.320784139 | 0.342365049 | 0.337234676 |
| 0.36289737 | 0.35572562 | 0.37799116 | 0.368828079 | 0.401568415 | 0.392390012 |
| 0.34499055 | 0.37747759 | 0.370470525 | 0.389747655 | 0.42962963 | 0.406427221 |
| 0.3634349 | 0.44076802 | 0.367672563 | 0.393431002 | 0.426638918 | 0.412326389 |

| BMI | BMI | BMI | BMI | BMI | BMI |
| --- | --- | --- | --- | --- | --- |
| Females Dex | Females Dex | Females Dex | Females Dex | Females Dex | |
| 0.294117647 | 0.302768166 | 0.275738941 | 0.271771474 | 0.298256669 | |
| 0.345167653 | 0.362666667 | 0.362624389 | 0.304076266 | 0.328891887 | |
| 0.321712018 | 0.310034602 | 0.348639456 | 0.294753921 | 0.304705215 | |
| 0.318518519 | 0.349382716 | 0.347784084 | 0.298273155 | 0.314440481 | |
| males Dex | males Dex | males Dex | males Dex | males Dex | males Dex |
| 0.298438935 | 0.302768166 | 0.377526754 | 0.300357068 | 0.281453476 | 0.295918367 |
| 0.370315289 | 0.364366449 | 0.3421875 | 0.35430839 | 0.390184423 | 0.352736246 |
| 0.38200692 | | 0.325851812 | 0.375308642 | 0.396160087 | 0.377854671 |
| 0.36389414 | | 0.341746166 | 0.38252603 | | 0.39310903 |

| BMI | BMI | BMI | BMI | BMI | BMI |
|---|---|---|---|---|---|
| males Dex | males Dex | males Dex | males Dex | males Dex | males Dex |
| 0.297959184 | 0.285714286 | 0.298442907 | 0.289792388 | 0.276816609 | 0.271771474 |
| 0.365397924 | 0.342935528 | 0.368480726 | 0.359994194 | 0.347222222 | 0.345804989 |
| 0.421849648 | 0.382653061 | 0.399018595 | 0.364197531 | 0.365444215 | 0.396353547 |
| 0.420857903 | 0.402726928 | 0.41942344 | 0.366681756 | 0.384011593 | 0.406427221 |

| GTT | GTT | GTT | GTT | GTT | GTT |
|---|---|---|---|---|---|
| Minutes | | females vehicle | females vehicle | females vehicle | females vehicle |
| 0 | | 6.4 | 6.7 | 6.3 | 8 |
| 15 | | 16.8 | 19.2 | 18.8 | 13.2 |
| 30 | | 12.1 | 14.2 | 17.7 | 11.8 |
| 60 | | 8.3 | 10.4 | 9.1 | 9.1 |
| 120 | | 6.4 | 9.8 | 8.2 | 10.7 |

| Minutes | | males vehicle | males vehicle | males vehicle | males vehicle |
|---|---|---|---|---|---|
| 0 | | 9.4 | 9 | 9.3 | 8.2 |
| 15 | | 16.7 | 19.9 | 26.5 | 22.1 |
| 30 | | 17.5 | 25.5 | 21.5 | 21.4 |
| 60 | | 22.7 | 12.9 | 18.1 | 16.1 |
| 120 | | 13.8 | 9.4 | 12.4 | 9.8 |

| GTT females vehicle | GTT females vehicle | GTT females vehicle | GTT females vehicle | GTT females vehicle | GTT females vehicle |
|---|---|---|---|---|---|
| 7 | 6.8 | 7 | 8.7 | 6.4 | 6.7 |
| 12.5 | 17.4 | 14.5 | 13.6 | 17.6 | 20.5 |
| 12.5 | 13.8 | 10.5 | 13.6 | 12.4 | 10.1 |
| 13.2 | 10.3 | 8.7 | 11 | 11.2 | 10.3 |
| 10.2 | 6.6 | 6.3 | 10.7 | 7.7 | 7.5 |

| males vehicle | males vehicle | males vehicle | males vehicle | males vehicle | males vehicle |
|---|---|---|---|---|---|
| 11.2 | 7.7 | 10.4 | 10.2 | 10 | 9.4 |
| 24.6 | 20.8 | 25.1 | 20.4 | 19.3 | 21.9 |
| 19.2 | 16.4 | 16.9 | 19.3 | 23.5 | 21.8 |
| 16.1 | 13.7 | 13.8 | 17.1 | 26.2 | 15.2 |
| 9.9 | 9.3 | 12.1 | 13 | 22.5 | 11.2 |

| GTT | GTT | GTT | GTT | GTT | GTT | GTT |
|---|---|---|---|---|---|---|
| females vehicle | females vehicle | Females Dex | Females Dex | Females Dex | Females Dex | Females Dex |
| 7.3 | 7.4 | 7.6 | 7.1 | 8.3 | 6.6 | 7.4 |
| 13.4 | 10.2 | 20 | 18.6 | 16 | 19.1 | 18.6 |
| 10.7 | 8.2 | 12.8 | 13.5 | 19.9 | 13.5 | 19 |
| 8.7 | 7.8 | 13.6 | 10.2 | 12.6 | 9.5 | 9.5 |
| 6.7 | 8.3 | 8.6 | 7.4 | 11.8 | 8.4 | 8.5 |
| males vehicle | males vehicle | males Dex | males Dex | males Dex | males Dex | males Dex |
| 9.7 | 9.5 | 8.3 | 7.8 | 11.8 | 9.5 | 8.7 |
| 21.4 | 20.6 | 23.9 | 19.7 | 25.8 | 18.6 | 19.9 |
| 22.1 | 23.8 | 18.4 | 17.8 | 25.1 | 19.7 | 15.8 |
| 21.4 | 15.6 | 12.3 | 14.8 | 24.6 | 20.8 | 14.1 |
| 12.5 | 14.4 | 9.8 | 13 | 18.4 | 14.7 | 10.4 |

| GTT Females Dex | GTT Females Dex | GTT Females Dex | GTT Females Dex | GTT Females Dex | GTT Females Dex | GTT Females Dex |
|---|---|---|---|---|---|---|
| 7.3 | 7.4 | 6.5 | 7.5 | 8.6 | 7 | 7.1 |
| 16.6 | 18.3 | 16 | 20.6 | 20.6 | 21.6 | 20.4 |
| 14.2 | 12.7 | 14.2 | 14.4 | 23 | 14.1 | 13.3 |
| 10.7 | 9.9 | 11.8 | 12.2 | 18.9 | 11.8 | 11.2 |
| 7.9 | 10.9 | 8.3 | 9.9 | 12.1 | 7.3 | 8.7 |

| males Dex | males Dex | males Dex | males Dex | males Dex | males Dex | males Dex |
|---|---|---|---|---|---|---|
| 7.9 | 9.7 | 8.3 | 6.2 | 7.6 | 7.9 | 7.5 |
| 19.1 | 15.3 | 19.8 | 23.5 | 16.5 | 16.9 | 19 |
| 16.8 | 19 | 15.4 | 17.1 | 16.9 | 19.3 | 15.8 |
| 11.8 | 13.5 | 12.5 | 10.5 | 15.4 | 11.3 | 11.5 |
| 8.1 | 11.2 | 8 | 6.4 | 10.4 | 7.9 | 8 |

| ITT | ITT | ITT | ITT | ITT | ITT |
|---|---|---|---|---|---|
| Minutes | | females vehicle | females vehicle | females vehicle | females vehicle |
| 0 | | 8.3 | 8.3 | 7.1 | 7.4 |
| 15 | | 6.1 | 5.8 | 5.3 | 5.5 |
| 30 | | 5.7 | 5.4 | 4.9 | 6.2 |
| 60 | | 5.7 | 5.3 | 4.9 | 5.6 |
| 120 | | 7.4 | 5 | 5.2 | 4.6 |

| Minutes | | males vehicle | males vehicle | males vehicle | males vehicle |
|---|---|---|---|---|---|
| 0 | | 9.9 | 8 | 8.3 | 8.4 |
| 15 | | 10.3 | 7.9 | 8.5 | 10.2 |
| 30 | | 10.3 | 6.2 | 6.6 | 7.2 |
| 60 | | 9.7 | 5.4 | 6.3 | 8.6 |
| 120 | | 10.4 | 6.2 | 11.6 | 9.2 |

| ITT females vehicle | ITT females vehicle | ITT females vehicle | ITT females vehicle | ITT females vehicle | ITT females vehicle |
|---|---|---|---|---|---|
| 7.1 | 6.2 | 7.4 | 6.5 | 7.5 | 7.3 |
| 8.1 | 5.3 | 4.6 | 6 | 5.4 | 5.5 |
| 7.1 | 4.5 | 4.4 | 5.5 | 4.7 | 4.4 |
| 6.3 | 3.7 | 4.7 | 4.7 | 4.6 | 4.7 |
| 6.1 | 3.7 | 4.7 | 5.2 | 6.7 | 5.4 |

| males vehicle | males vehicle | males vehicle | males vehicle | males vehicle |
|---|---|---|---|---|
| 11.1 | 11.2 | 12.8 | 9.2 | 10.3 |
| 11.2 | 10.2 | 10.8 | 8.6 | 10.2 |
| 8.9 | 9.2 | 9.5 | 5.9 | 10.5 |
| 9.2 | 9.8 | 10.2 | 6.1 | 8.8 |
| 10.9 | 12 | 9.8 | 9.2 | 11.2 |

| ITT females vehicle | ITT females vehicle | ITT females Dex | ITT females Dex | ITT females Dex | ITT females Dex | ITT females Dex |
|---|---|---|---|---|---|---|
| 7.4 | 7.1 | 6.9 | 6.5 | 9.5 | 6.3 | 8.2 |
| 5.3 | 6.3 | 5.3 | 5.4 | 6.1 | 5.3 | 6.6 |
| 5.5 | 4.9 | 6 | 4.3 | 5.3 | 4.5 | 5.8 |
| 3.9 | 3.3 | 4.4 | 4.2 | 4.7 | 4.6 | 3.8 |
| 5 | 4.7 | 5.3 | 4.9 | 7.5 | 5 | 4.6 |

| males Dex | males Dex | males Dex | males Dex | males Dex |
|---|---|---|---|---|
| 10.2 | 8.9 | 9.2 | 10.1 | 8.9 |
| 11.2 | 9.4 | 8.7 | 10.8 | 9.7 |
| 10.5 | 8.9 | 8.4 | 9.6 | 9.5 |
| 11.3 | 7.9 | 8.6 | 9.8 | 9.2 |
| 10.5 | 8.9 | 8 | 10.2 | 9.1 |

| ITT females Dex | ITT females Dex | ITT females Dex | ITT females Dex | ITT females Dex | ITT females Dex | ITT females Dex |
|---|---|---|---|---|---|---|
| 7.8 | 7.3 | 7.5 | 6.5 | 8 | 7.5 | 7.4 |
| 4.9 | 4.6 | 8.4 | 6.3 | 6 | 5.4 | 6.3 |
| 6.4 | 4.8 | 7 | 5.4 | 6.3 | 5.2 | 5.9 |
| 3.9 | 4.2 | 8.5 | 4.3 | 4.9 | 5 | 5.2 |
| 3.8 | 4.9 | 6.9 | 5 | 5.2 | 7 | 6.3 |

| males Dex | males Dex | males Dex |
|---|---|---|
| 8.7 | 9.9 | 9.5 |
| 12.4 | 11.3 | 12.3 |
| 13.1 | 9.8 | 12.5 |
| 10.2 | 9.8 | 12 |
| 11.7 | 9 | 9.2 |

| Necropsy | Necropsy Females vehicle offspring | Necropsy Females vehicle offspring | Necropsy Females vehicle offspring | Necropsy Females vehicle offspring | Necropsy Females vehicle offspring | Necropsy Females vehicle offspring |
|---|---|---|---|---|---|---|
| Liver (g) | 1.557 | 1.331 | 1.264 | 1.388 | 1.3 | 1.227 |
| Kidney (g) | 0.351 | 0.389 | 0.393 | 0.39 | 0.339 | 0.378 |
| Inguinal WA | 0.665 | 0.545 | 0.204 | 0.334 | 0.27 | 0.374 |
| Gonadal WA | 1.994 | 1.509 | 0.619 | 1.182 | 0.963 | 1.221 |
| Interscapular | 0.161 | 0.155 | 0.11 | 0.132 | 0.09 | 0.126 |

| | Males vehicle offspring | Males vehicle offspring | Males vehicle offspring | Males vehicle offspring | Males vehicle offspring | Males vehicle offspring |
|---|---|---|---|---|---|---|
| Liver (g) | 1.756 | 1.918 | 2.075 | 1.88 | 1.59 | 1.941 |
| Kidney (g) | 0.478 | 0.48 | 0.487 | 0.423 | 0.415 | 0.51 |
| Inguinal WA | 0.624 | 0.193 | 0.76 | 0.754 | 0.543 | 0.43 |
| Gonadal WA | 1.904 | 0.483 | 2.099 | 2.284 | 1.58 | 1.334 |
| Interscapular | 0.227 | 0.175 | 0.262 | 0.31 | 0.298 | 0.172 |

| Necropsy Females vehicle offspring | Necropsy Females vehicle offspring | Necropsy Females vehicle offspring | Necropsy Females Dex offspring | Necropsy Females Dex offspring | Necropsy Females Dex offspring | Necropsy Females Dex offspring |
|---|---|---|---|---|---|---|
| 1.376 | 1.333 | 1.615 | 1.382 | 1.389 | 1.569 | 1.433 |
| 0.38 | 0.397 | 0.388 | 0.379 | 0.388 | 0.407 | 0.39 |
| 0.414 | 0.468 | 0.804 | 0.312 | 0.249 | 0.662 | 0.695 |
| 1.292 | 1.51 | 2.42 | 0.961 | 0.823 | 1.762 | 1.798 |
| 0.227 | 0.133 | 0.236 | 0.123 | 0.123 | 0.241 | 0.167 |

| Males vehicle offspring | Males vehicle offspring | Males vehicle offspring | Males Dex offspring | Males Dex offspring | Males Dex offspring | Males Dex offspring |
|---|---|---|---|---|---|---|
| 1.254 | 1.918 | 1.753 | 1.695 | 1.651 | 2.103 | 1.307 |
| 0.457 | 0.428 | 0.459 | 0.42 | 0.463 | 0.484 | 0.404 |
| 0.564 | 0.857 | 0.738 | 0.448 | 0.615 | 0.856 | 0.443 |
| 1.74 | 2.303 | 1.985 | 1.725 | 1.873 | 2.156 | 1.495 |
| 0.205 | 0.255 | 0.208 | 0.194 | 0.264 | 0.22 | 0.147 |

| Necropsy Females Dex offspring | Necropsy Females Dex offspring |
|---|---|
| 1.519 | 1.376 |
| 0.379 | 0.403 |
| 0.624 | 0.355 |
| 1.972 | 1.09 |
| 0.144 | 0.154 |

| Males Dex offspring | Males Dex offspring | Males Dex offspring | Males Dex offspring |
|---|---|---|---|
| 1.597 | 1.858 | 1.866 | 1.671 |
| 0.473 | 0.449 | 0.502 | 0.448 |
| 0.638 | 0.749 | 0.65 | 0.569 |
| 1.9 | 1.886 | 2.107 | 2.222 |
| 0.204 | 0.176 | 0.232 | 0.289 |
