## Supplementary Table 4 for "Single paternal Dexamethasone challenge programs offspring metabolism and reveals circRNAs as novel candidates in RNA-mediated inheritance"

| gene | Total | Cluster1 | Cluster2 |
|---|---|---|---|
| Pemt | TRUE | FALSE | FALSE |
| Nhp2 | TRUE | FALSE | FALSE |
| Bcap31 | TRUE | TRUE | FALSE |
| Rps11 | TRUE | FALSE | FALSE |
| Hspb1 | TRUE | FALSE | FALSE |
| Pgrmc1 | TRUE | TRUE | FALSE |
| Cdc20 | TRUE | FALSE | FALSE |
| D8Ertd738e | TRUE | FALSE | FALSE |
| Cops6 | TRUE | FALSE | FALSE |
| Gyg | TRUE | FALSE | FALSE |
| Supt4a | TRUE | FALSE | FALSE |
| Bmp15 | TRUE | FALSE | TRUE |
| Ccndbp1 | TRUE | FALSE | FALSE |
| Hsp90ab1 | TRUE | FALSE | FALSE |
| Guca1a | TRUE | FALSE | FALSE |
| Dpy30 | TRUE | FALSE | FALSE |
| Fth1 | TRUE | FALSE | FALSE |
| Sat1 | TRUE | TRUE | FALSE |
| Hprt | TRUE | TRUE | FALSE |
| Alppl2 | TRUE | FALSE | TRUE |
| Med27 | TRUE | FALSE | FALSE |
| Dut | TRUE | FALSE | FALSE |
| Knstrn | TRUE | FALSE | FALSE |
| Rbm38 | TRUE | TRUE | TRUE |
| Cks1b | TRUE | FALSE | FALSE |
| Rps3a1 | TRUE | FALSE | FALSE |
| Tesc | TRUE | FALSE | TRUE |
| Cnbp | TRUE | FALSE | FALSE |
| Pqbp1 | TRUE | FALSE | FALSE |
| Rbbp7 | TRUE | TRUE | FALSE |
| Haus7 | TRUE | FALSE | FALSE |
| Prps1 | TRUE | TRUE | FALSE |
| Mphosph6 | TRUE | TRUE | TRUE |
| Bcl2l10 | TRUE | FALSE | TRUE |
| Cdo1 | TRUE | FALSE | TRUE |
| Chmp2a | TRUE | FALSE | FALSE |
| Rfpl4 | TRUE | FALSE | TRUE |
| Oaz1 | TRUE | FALSE | TRUE |
| Lmo1 | TRUE | FALSE | FALSE |
| Rps27l | TRUE | FALSE | TRUE |
| Wee2 | TRUE | TRUE | FALSE |
| Omt2b | TRUE | FALSE | TRUE |
| Nubp2 | TRUE | FALSE | FALSE |
| Polr2j | TRUE | FALSE | TRUE |
| Tcl1 | TRUE | TRUE | TRUE |
| Selenow | TRUE | TRUE | FALSE |
| Oosp1 | TRUE | FALSE | FALSE |

| | | | |
|---|---|---|---|
| Immp1l | TRUE | FALSE | FALSE |
| Crb3 | TRUE | FALSE | FALSE |
| Tubg2 | TRUE | TRUE | FALSE |
| Dppa3 | TRUE | FALSE | FALSE |
| Cox6b2 | TRUE | FALSE | TRUE |
| Grwd1 | TRUE | FALSE | FALSE |
| 1700013H16I | TRUE | FALSE | FALSE |
| Oosp3 | TRUE | FALSE | FALSE |
| Gm6054 | TRUE | FALSE | FALSE |
| Gm839 | TRUE | FALSE | FALSE |
| Dppa5a | TRUE | FALSE | FALSE |
| Tcl1b2 | TRUE | FALSE | TRUE |
| Snrpa | TRUE | FALSE | FALSE |
| Cks2 | TRUE | FALSE | TRUE |
| mt-Rnr1 | TRUE | TRUE | FALSE |
| mt-Nd1 | TRUE | FALSE | TRUE |
| mt-Nd2 | TRUE | TRUE | FALSE |
| mt-Co1 | TRUE | FALSE | TRUE |
| mt-Nd6 | TRUE | TRUE | FALSE |
| mt-Cytb | TRUE | TRUE | FALSE |
| Gm13023 | TRUE | FALSE | FALSE |
| Utp3 | TRUE | FALSE | FALSE |
| Uqcc3 | TRUE | FALSE | FALSE |
| Fthl17f | TRUE | TRUE | FALSE |
| Gm9 | TRUE | FALSE | TRUE |
| AU022751 | TRUE | TRUE | TRUE |
| Gm6091 | TRUE | TRUE | FALSE |
| 7420426K07I | TRUE | FALSE | TRUE |
| Obox5 | TRUE | FALSE | TRUE |
| Accsl | TRUE | FALSE | FALSE |
| Gm23935 | TRUE | TRUE | TRUE |
| Mir743 | TRUE | FALSE | FALSE |
| Ube2d3 | TRUE | FALSE | FALSE |
| Eif5a | TRUE | TRUE | FALSE |
| Rab7 | TRUE | FALSE | FALSE |
| Gm11448 | TRUE | FALSE | FALSE |
| Gm7212 | TRUE | FALSE | FALSE |
| Gm11517 | TRUE | FALSE | FALSE |
| Gm11839 | TRUE | FALSE | FALSE |
| Gm15361 | TRUE | FALSE | FALSE |
| Gm12617 | TRUE | TRUE | FALSE |
| Gm15452 | TRUE | FALSE | FALSE |
| Gm26479 | TRUE | FALSE | FALSE |
| Khdc1b | TRUE | FALSE | FALSE |
| Gm11381 | TRUE | FALSE | FALSE |
| Gm26049 | TRUE | FALSE | FALSE |
| Gm1965 | TRUE | TRUE | FALSE |
| Gm4202 | TRUE | FALSE | FALSE |

| | | | |
|---|---|---|---|
| Rpl41 | TRUE | FALSE | FALSE |
| Gm12625 | TRUE | FALSE | FALSE |
| Rfpl4b | TRUE | TRUE | FALSE |
| Gm12622 | TRUE | TRUE | TRUE |
| Gm29290 | TRUE | FALSE | FALSE |
| Gm36976 | TRUE | FALSE | TRUE |
| | TRUE | TRUE | TRUE |
| Gm5 | TRUE | FALSE | FALSE |
| AA545190 | TRUE | FALSE | FALSE |
| C86187 | TRUE | TRUE | FALSE |
