## Supplementary Table 5 for "Single paternal Dexamethasone challenge programs offspring metabolism and reveals circRNAs as novel candidates in RNA-mediated inheritance"

| HS548_counts | | HS548_counts | | HS548_counts | | HS548_cou |
|---|---|---|---|---|---|---|
| Ens_ID | Gene | Chrom | Strand | Start | End | RNA_type |
| ENSMUSG00000000085 | Scmh1 | chr4 | + | 120405280 | 120530186 | regular |
| ENSMUSG00000000125 | Wnt3 | chr11 | + | 103774149 | 103817957 | regular |
| ENSMUSG00000000127 | Fer | chr17 | + | 63896017 | 64139494 | regular |
| ENSMUSG00000000131 | Xpo6 | chr7 | - | 126101714 | 126200501 | regular |
| ENSMUSG00000000167 | Pih1d2 | chr9 | + | 50617320 | 50625000 | regular |
| ENSMUSG00000000215 | Ins2 | chr7 | - | 142678655 | 142743381 | regular |
| ENSMUSG00000000275 | Trim25 | chr11 | + | 88999375 | 89020293 | regular |
| ENSMUSG00000000346 | Dazap2 | chr15 | + | 100615627 | 100620731 | regular |
| ENSMUSG00000000365 | Rnf17 | chr14 | + | 56402580 | 56525032 | regular |
| ENSMUSG00000000411 | Tssk3 | chr4 | - | 129489013 | 129491022 | regular |
| ENSMUSG00000000439 | Mkrn2 | chr6 | + | 115601901 | 115622624 | regular |
| ENSMUSG00000000552 | Zfp385a | chr15 | - | 103313894 | 103340086 | regular |
| ENSMUSG00000000568 | Hnrnpd | chr5 | - | 99955934 | 99978938 | regular |
| ENSMUSG00000000682 | Cd52 | chr4 | - | 134082447 | 134095082 | regular |
| ENSMUSG00000000740 | Rpl13 | chr8 | + | 123102349 | 123105244 | regular |
| ENSMUSG00000000751 | Rpa1 | chr11 | - | 75298165 | 75348324 | regular |
| ENSMUSG00000000787 | Ddx3x | chrX | + | 13280969 | 13294052 | regular |
| ENSMUSG00000000804 | Usp32 | chr11 | - | 84984441 | 85140161 | regular |
| ENSMUSG00000000811 | Txnrd3 | chr6 | + | 89643987 | 89675529 | regular |
| ENSMUSG00000000823 | Zfp512b | chr2 | - | 181582102 | 181592803 | regular |
| ENSMUSG00000000915 | Hip1r | chr5 | + | 123973627 | 124005558 | regular |
| ENSMUSG00000001052 | Sec24b | chr3 | - | 129982758 | 130061553 | regular |
| ENSMUSG00000001089 | Luzp1 | chr4 | + | 136469760 | 136554780 | regular |
| ENSMUSG00000001100 | Poldip2 | chr11 | + | 78512192 | 78522736 | regular |
| ENSMUSG00000001120 | Pcbp3 | chr10 | - | 76761856 | 76961887 | regular |
| ENSMUSG00000001151 | Pcnt | chr10 | - | 76351262 | 76442786 | regular |
| ENSMUSG00000001156 | Mxd1 | chr6 | - | 86647041 | 86669161 | regular |
| ENSMUSG00000001173 | Ocrl | chrX | + | 47912386 | 47965868 | regular |
| ENSMUSG00000001175 | Calm1 | chr12 | + | 100199434 | 100209814 | regular |
| ENSMUSG00000001211 | Agpat3 | chr10 | - | 78269177 | 78352489 | regular |
| ENSMUSG00000001280 | Sp1 | chr15 | + | 102406142 | 102436404 | regular |
| ENSMUSG00000001289 | Pfdn5 | chr15 | + | 102326115 | 102331873 | regular |
| ENSMUSG00000001415 | Smg5 | chr3 | + | 88336259 | 88362338 | regular |
| ENSMUSG00000001416 | Cct3 | chr3 | + | 88297115 | 88321767 | regular |
| ENSMUSG00000001419 | Mef2d | chr3 | + | 88142371 | 88172086 | regular |
| ENSMUSG00000001424 | Snd1 | chr6 | + | 28475138 | 28935162 | regular |
| ENSMUSG00000001440 | Kpnb1 | chr11 | - | 97159713 | 97187881 | regular |
| ENSMUSG00000001441 | Npepps | chr11 | - | 97205841 | 97280638 | regular |
| ENSMUSG00000001472 | Tcf25 | chr8 | + | 123373752 | 123403835 | regular |
| ENSMUSG00000001517 | Foxm1 | chr6 | + | 128362966 | 128376146 | regular |
| ENSMUSG00000001552 | Jup | chr11 | - | 100368957 | 100397763 | regular |
| ENSMUSG00000001558 | Klhl10 | chr11 | + | 100441916 | 100457022 | regular |

| | | | | | | |
|---|---|---|---|---|---|---|
| ENSMUSG00000001576 | Ergic1 | chr17 | + | 26561488 | 26656931 | regular |
| ENSMUSG00000001687 | Ubl3 | chr5 | - | 148504634 | 148552789 | regular |
| ENSMUSG00000001774 | Chordc1 | chr9 | + | 18292124 | 18317442 | regular |
| ENSMUSG00000001794 | Capns1 | chr7 | - | 30186935 | 30198811 | regular |
| ENSMUSG00000001829 | Clpb | chr7 | + | 101663632 | 101795506 | regular |
| ENSMUSG00000001855 | Nup214 | chr2 | + | 31974435 | 32053975 | regular |
| ENSMUSG00000001909 | Trmt1 | chr8 | + | 84686306 | 84699808 | regular |
| ENSMUSG00000001924 | Uba1 | chrX | + | 20658325 | 20683179 | regular |
| ENSMUSG00000001948 | Spa17 | chr9 | - | 37603289 | 37613722 | regular |
| ENSMUSG00000002028 | Kmt2a | chr9 | - | 44803354 | 44881296 | regular |
| ENSMUSG00000002043 | Trappc6a | chr7 | + | 19508394 | 19516145 | regular |
| ENSMUSG00000002052 | Supt6 | chr11 | - | 78206745 | 78245987 | regular |
| ENSMUSG00000002055 | Spag5 | chr11 | + | 78301528 | 78322457 | regular |
| ENSMUSG00000002102 | Psmc3 | chr2 | + | 91054008 | 91066369 | regular |
| ENSMUSG00000002107 | Celf2 | chr2 | - | 6539693 | 7509563 | regular |
| ENSMUSG00000002190 | Clgn | chr8 | + | 83389866 | 83428552 | regular |
| ENSMUSG00000002210 | Smg9 | chr7 | + | 24399618 | 24422778 | regular |
| ENSMUSG00000002324 | Rec8 | chr14 | + | 55618036 | 55625395 | regular |
| ENSMUSG00000002372 | Ranbp3 | chr17 | + | 56673224 | 56711769 | regular |
| ENSMUSG00000002395 | Use1 | chr8 | + | 71366847 | 71369732 | regular |
| ENSMUSG00000002409 | Dyrk1b | chr7 | + | 28179468 | 28187294 | regular |
| ENSMUSG00000002413 | Braf | chr6 | - | 39603236 | 39725463 | regular |
| ENSMUSG00000002455 | Prpf6 | chr2 | + | 181591867 | 181655660 | regular |
| ENSMUSG00000002546 | Golga2 | chr2 | + | 32287383 | 32307921 | regular |
| ENSMUSG00000002625 | Akap8l | chr17 | - | 32321424 | 32350581 | regular |
| ENSMUSG00000002658 | Gtf2f1 | chr17 | - | 57003404 | 57011288 | regular |
| ENSMUSG00000002718 | Cse1l | chr2 | + | 166906039 | 166946389 | regular |
| ENSMUSG00000002733 | Plekha3 | chr2 | + | 76675280 | 76696828 | regular |
| ENSMUSG00000002748 | Baz1b | chr5 | + | 135187263 | 135246129 | regular |
| ENSMUSG00000002763 | Pex6 | chr17 | + | 46711462 | 46725541 | regular |
| ENSMUSG00000002768 | Mea1 | chr17 | + | 46681136 | 46683105 | regular |
| ENSMUSG00000002833 | Hdgfl2 | chr17 | + | 56079633 | 56100607 | regular |
| ENSMUSG00000002835 | Chaf1a | chr17 | + | 56040415 | 56068026 | regular |
| ENSMUSG00000002885 | Adgre5 | chr8 | - | 83723250 | 83741326 | regular |
| ENSMUSG00000002968 | Med25 | chr7 | - | 44876764 | 44892712 | regular |
| ENSMUSG00000002985 | Apoe | chr7 | - | 19696108 | 19699188 | regular |
| ENSMUSG00000002996 | Hbp1 | chr12 | - | 31926253 | 31950535 | regular |
| ENSMUSG00000003033 | Ap1m1 | chr8 | + | 72240017 | 72257385 | regular |
| ENSMUSG00000003068 | Stk11 | chr10 | + | 80115802 | 80130682 | regular |
| ENSMUSG00000003119 | Cdk12 | chr11 | + | 98203058 | 98278504 | regular |
| ENSMUSG00000003123 | Lipe | chr7 | - | 25379526 | 25398710 | regular |
| ENSMUSG00000003131 | Pafah1b2 | chr9 | - | 45962858 | 46012690 | regular |
| ENSMUSG00000003153 | Slc2a3 | chr6 | - | 122727808 | 122801640 | regular |
| ENSMUSG00000003226 | Ranbp2 | chr10 | + | 58446919 | 58494356 | regular |
| ENSMUSG00000003269 | Cyth2 | chr7 | - | 45806636 | 45814581 | regular |
| ENSMUSG00000003341 | Atp8b3 | chr10 | - | 80519583 | 80539124 | regular |

| | | | | | | |
|---|---|---|---|---|---|---|
| ENSMUSG00000003345 | Csnk1g2 | chr10 | + | 80622837 | 80640749 | regular |
| ENSMUSG00000003382 | Etv3 | chr3 | + | 87525406 | 87540156 | regular |
| ENSMUSG00000003429 | Rps11 | chr7 | - | 45122378 | 45124439 | regular |
| ENSMUSG00000003435 | Supt5 | chr7 | - | 28314890 | 28338746 | regular |
| ENSMUSG00000003531 | Dgcr6 | chr16 | + | 18052859 | 18071632 | regular |
| ENSMUSG00000003604 | Aven | chr2 | + | 112492963 | 112634573 | regular |
| ENSMUSG00000003660 | Snrnp200 | chr2 | + | 127208385 | 127240451 | regular |
| ENSMUSG00000003731 | Kpna6 | chr4 | - | 129643979 | 129672767 | regular |
| ENSMUSG00000003746 | Man1a | chr10 | - | 53904784 | 54076609 | regular |
| ENSMUSG00000003808 | Farsa | chr8 | + | 84856988 | 84869257 | regular |
| ENSMUSG00000003810 | Mast2 | chr4 | - | 116306761 | 116464183 | regular |
| ENSMUSG00000003814 | Calr | chr8 | - | 84841849 | 84846934 | regular |
| ENSMUSG00000003824 | Syce2 | chr8 | + | 84872110 | 84888221 | regular |
| ENSMUSG00000003847 | Nfat5 | chr8 | + | 107293469 | 107379517 | regular |
| ENSMUSG00000003970 | Rpl8 | chr15 | + | 76904070 | 76906318 | regular |
| ENSMUSG00000004018 | Fancl | chr11 | + | 26386134 | 26471876 | regular |
| ENSMUSG00000004032 | Gstm5 | chr3 | + | 107895820 | 107898686 | regular |
| ENSMUSG00000004035 | Gstm7 | chr3 | - | 107926333 | 107931817 | regular |
| ENSMUSG00000004040 | Stat3 | chr11 | - | 100885097 | 100939540 | regular |
| ENSMUSG00000004070 | Hmox2 | chr16 | + | 4726360 | 4766742 | regular |
| ENSMUSG00000004110 | Cacna1e | chr1 | - | 154390730 | 154884501 | regular |
| ENSMUSG00000004127 | Trmt10a | chr3 | + | 138143447 | 138159821 | regular |
| ENSMUSG00000004207 | Psap | chr10 | + | 60277626 | 60302597 | regular |
| ENSMUSG00000004347 | Pde1c | chr6 | - | 56069803 | 56569103 | regular |
| ENSMUSG00000004356 | Utp20 | chr10 | - | 88746606 | 88826804 | regular |
| ENSMUSG00000004364 | Cul3 | chr1 | - | 80264922 | 80340480 | regular |
| ENSMUSG00000004455 | Ppp1cc | chr5 | + | 122158277 | 122175273 | regular |
| ENSMUSG00000004535 | Tax1bp1 | chr6 | + | 52713728 | 52766780 | regular |
| ENSMUSG00000004677 | Myo9b | chr8 | + | 71272713 | 71360713 | regular |
| ENSMUSG00000004789 | Dlst | chr12 | + | 85110832 | 85134845 | regular |
| ENSMUSG00000004798 | Ulk2 | chr11 | - | 61775648 | 61855073 | regular |
| ENSMUSG00000004865 | Srpk1 | chr17 | - | 28587647 | 28622521 | regular |
| ENSMUSG00000004897 | Hdgf | chr3 | + | 87906320 | 87916132 | regular |
| ENSMUSG00000004929 | Thop1 | chr10 | + | 81070034 | 81082559 | regular |
| ENSMUSG00000004934 | Pias4 | chr10 | - | 81153265 | 81167923 | regular |
| ENSMUSG00000004937 | Sgta | chr10 | - | 81044074 | 81060181 | regular |
| ENSMUSG00000004980 | Hnrnpa2b | chr6 | - | 51460931 | 51469894 | regular |
| ENSMUSG00000005045 | Chd5 | chr4 | + | 152338650 | 152390194 | regular |
| ENSMUSG00000005087 | Cd44 | chr2 | - | 102811140 | 102901665 | regular |
| ENSMUSG00000005103 | Wdr1 | chr5 | - | 38526812 | 38563221 | regular |
| ENSMUSG00000005198 | Polr2a | chr11 | - | 69733996 | 69758637 | regular |
| ENSMUSG00000005225 | Plekha8 | chr6 | + | 54595110 | 54645839 | regular |
| ENSMUSG00000005237 | Dnah2 | chr11 | - | 69420808 | 69549110 | regular |
| ENSMUSG00000005312 | Ubqln1 | chr13 | - | 58176155 | 58215653 | regular |
| ENSMUSG00000005417 | Mprip | chr11 | + | 59661304 | 59780860 | regular |
| ENSMUSG00000005442 | Cic | chr7 | + | 25267703 | 25294159 | regular |

| | | | | | | |
|---|---|---|---|---|---|---|
| ENSMUSG00000005483 | Dnajb1 | chr8 | + | 83608192 | 83612653 | regular |
| ENSMUSG00000005510 | Ndufs3 | chr2 | - | 90894633 | 90904827 | regular |
| ENSMUSG00000005566 | Trim28 | chr7 | + | 12999113 | 13031035 | regular |
| ENSMUSG00000005575 | Ube2m | chr7 | - | 13035119 | 13038275 | regular |
| ENSMUSG00000005609 | Ctr9 | chr7 | + | 111028950 | 111056377 | regular |
| ENSMUSG00000005610 | Eif4g2 | chr7 | - | 111067749 | 111083030 | regular |
| ENSMUSG00000005615 | Pcyt1a | chr16 | + | 32430920 | 32475070 | regular |
| ENSMUSG00000005621 | Zfp592 | chr7 | + | 80993680 | 81045164 | regular |
| ENSMUSG00000005683 | Cs | chr10 | + | 128337733 | 128362479 | regular |
| ENSMUSG00000005698 | Ctcf | chr8 | + | 105636567 | 105682922 | regular |
| ENSMUSG00000005802 | Slc30a4 | chr2 | - | 122681232 | 122702663 | regular |
| ENSMUSG00000005886 | Ncoa2 | chr1 | - | 13139104 | 13374083 | regular |
| ENSMUSG00000005893 | Nr2c2 | chr6 | + | 92091389 | 92174294 | regular |
| ENSMUSG00000005973 | Rcn1 | chr2 | - | 105386290 | 105399319 | regular |
| ENSMUSG00000005981 | Trap1 | chr16 | - | 4039970 | 4077827 | regular |
| ENSMUSG00000006005 | Tpr | chr1 | + | 150392837 | 150449935 | regular |
| ENSMUSG00000006014 | Prg4 | chr1 | - | 150449411 | 150466165 | regular |
| ENSMUSG00000006024 | Napa | chr7 | + | 16098457 | 16117975 | regular |
| ENSMUSG00000006127 | Inpp5k | chr11 | + | 75630987 | 75648871 | regular |
| ENSMUSG00000006169 | Clint1 | chr11 | + | 45852050 | 45910625 | regular |
| ENSMUSG00000006241 | Ccdc159 | chr9 | + | 21927470 | 21935872 | regular |
| ENSMUSG00000006299 | Aamp | chr1 | - | 74279839 | 74284738 | regular |
| ENSMUSG00000006304 | Arpc2 | chr1 | + | 74236083 | 74268209 | regular |
| ENSMUSG00000006307 | Kmt2b | chr7 | - | 30568857 | 30588726 | regular |
| ENSMUSG00000006333 | Rps9 | chr7 | + | 3703992 | 3706897 | regular |
| ENSMUSG00000006356 | Crip2 | chr12 | + | 113140235 | 113145506 | regular |
| ENSMUSG00000006418 | Rnf114 | chr2 | + | 167492644 | 167516173 | regular |
| ENSMUSG00000006435 | Neurl1a | chr19 | + | 47178819 | 47259440 | regular |
| ENSMUSG00000006498 | Ptbp1 | chr10 | + | 79854426 | 79864771 | regular |
| ENSMUSG00000006527 | Sfmbt1 | chr14 | + | 30714848 | 30822721 | regular |
| ENSMUSG00000006575 | Rundc3a | chr11 | + | 102393402 | 102402555 | regular |
| ENSMUSG00000006599 | Gtf2h1 | chr7 | + | 46796102 | 46823800 | regular |
| ENSMUSG00000006673 | Qrich1 | chr9 | + | 108516805 | 108560163 | regular |
| ENSMUSG00000006676 | Usp19 | chr9 | + | 108490601 | 108502337 | regular |
| ENSMUSG00000006699 | Cdc42 | chr4 | - | 137319695 | 137357720 | regular |
| ENSMUSG00000006705 | Pknox1 | chr17 | + | 31564748 | 31607684 | regular |
| ENSMUSG00000006818 | Sod2 | chr17 | + | 13007838 | 13018119 | regular |
| ENSMUSG00000006932 | Ctnnb1 | chr9 | + | 120929215 | 120960507 | regular |
| ENSMUSG00000006998 | Psmd2 | chr16 | + | 20651651 | 20663414 | regular |
| ENSMUSG00000007029 | Vars | chr17 | + | 35000986 | 35016322 | regular |
| ENSMUSG00000007033 | Hspa1l | chr17 | + | 34972702 | 34979285 | regular |
| ENSMUSG00000007107 | Atp1a4 | chr1 | - | 172223512 | 172258414 | regular |
| ENSMUSG00000007411 | Mark3 | chr12 | + | 111574522 | 111656221 | regular |
| ENSMUSG00000007617 | Homer1 | chr13 | + | 93299634 | 93405129 | regular |
| ENSMUSG00000007739 | Cct4 | chr11 | + | 22990518 | 23003780 | regular |
| ENSMUSG00000007815 | Rhoa | chr9 | + | 108306128 | 108337934 | regular |

| | | | | | | |
|---|---|---|---|---|---|---|
| ENSMUSG00000007817 | Zmiz1 | chr14 | + | 25459184 | 25666743 | regular |
| ENSMUSG00000007850 | Hnrnph1 | chr11 | + | 50376989 | 50386528 | regular |
| ENSMUSG00000007880 | Arid1a | chr4 | - | 133679007 | 133756769 | regular |
| ENSMUSG00000007892 | Rplp1 | chr9 | - | 61913283 | 61914542 | regular |
| ENSMUSG00000007907 | Cabs1 | chr5 | + | 87979438 | 87981544 | regular |
| ENSMUSG00000007950 | Abhd8 | chr8 | - | 71456704 | 71463655 | regular |
| ENSMUSG00000008200 | Fnbp4 | chr2 | + | 90745369 | 90781021 | regular |
| ENSMUSG00000008348 | Ubc | chr5 | - | 125385964 | 125390202 | regular |
| ENSMUSG00000008393 | Carhsp1 | chr16 | - | 8658586 | 8672153 | regular |
| ENSMUSG00000008429 | Herpud2 | chr9 | - | 25108131 | 25151820 | regular |
| ENSMUSG00000008668 | Rps18 | chr17 | - | 33951998 | 33956001 | regular |
| ENSMUSG00000008682 | Rpl10 | chrX | + | 74270811 | 74273135 | regular |
| ENSMUSG00000008683 | Rps15a | chr7 | - | 118104371 | 118116188 | regular |
| ENSMUSG00000008730 | Hipk1 | chr3 | - | 103739814 | 103791563 | regular |
| ENSMUSG00000008763 | Man1a2 | chr3 | - | 100562207 | 100685503 | regular |
| ENSMUSG00000008813 | Tppp2 | chr14 | + | 51918424 | 51920700 | regular |
| ENSMUSG00000009013 | Dynll1 | chr5 | - | 115297109 | 115300999 | regular |
| ENSMUSG00000009073 | Nf2 | chr11 | - | 4765844 | 4849536 | regular |
| ENSMUSG00000009079 | Ewsr1 | chr11 | - | 5069688 | 5099266 | regular |
| ENSMUSG00000009112 | Bcl2l13 | chr6 | + | 120836211 | 120892842 | regular |
| ENSMUSG00000009207 | Lnpk | chr2 | - | 74520290 | 74579435 | regular |
| ENSMUSG00000009394 | Syn2 | chr6 | + | 115134901 | 115282006 | regular |
| ENSMUSG00000009470 | Tnpo1 | chr13 | - | 98839018 | 98926384 | regular |
| ENSMUSG00000009575 | Cbx5 | chr15 | - | 103191543 | 103239816 | regular |
| ENSMUSG00000009621 | Vav2 | chr2 | - | 27262103 | 27427033 | regular |
| ENSMUSG00000009741 | Ubp1 | chr9 | + | 113930933 | 113977202 | regular |
| ENSMUSG00000009927 | Rps25 | chr9 | + | 44407138 | 44410427 | regular |
| ENSMUSG00000010097 | Nxf1 | chr19 | + | 8757072 | 8772475 | regular |
| ENSMUSG00000010277 | 2610507B | chr11 | + | 78261751 | 78290623 | regular |
| ENSMUSG00000010342 | Tex14 | chr11 | + | 87405064 | 87555823 | regular |
| ENSMUSG00000010453 | Kansl3 | chr1 | - | 36335729 | 36369181 | regular |
| ENSMUSG00000010461 | Eya4 | chr10 | - | 23102962 | 23350786 | regular |
| ENSMUSG00000010517 | Faf1 | chr4 | + | 109676587 | 109963960 | regular |
| ENSMUSG00000010538 | Tsacc | chr3 | - | 88282756 | 88296999 | regular |
| ENSMUSG00000010608 | Rbm25 | chr12 | + | 83631235 | 83683123 | regular |
| ENSMUSG00000010721 | Lmbr1 | chr5 | - | 29229801 | 29378390 | regular |
| ENSMUSG00000010755 | Cars | chr7 | - | 143557229 | 143600090 | regular |
| ENSMUSG00000011158 | Brf1 | chr12 | - | 112960084 | 113000704 | regular |
| ENSMUSG00000011179 | Odc1 | chr12 | + | 17544793 | 17551505 | regular |
| ENSMUSG00000011350 | Gm5893 | chr7 | + | 24818794 | 24840275 | regular |
| ENSMUSG00000011831 | Evi5 | chr5 | - | 107744794 | 107875107 | regular |
| ENSMUSG00000011832 | Evi5l | chr8 | + | 4166566 | 4211257 | regular |
| ENSMUSG00000011877 | Git1 | chr11 | + | 77493561 | 77507786 | regular |
| ENSMUSG00000012114 | Med15 | chr16 | - | 17651220 | 17722930 | regular |
| ENSMUSG00000012126 | Ubxn11 | chr4 | + | 134102569 | 134127593 | regular |
| ENSMUSG00000012535 | Tnpo3 | chr6 | - | 29540826 | 29609887 | regular |

| | | | | | | |
|---|---|---|---|---|---|---|
| ENSMUSG00000012609 | Ttll5 | chr12 | + | 85824658 | 86061893 | regular |
| ENSMUSG00000012819 | Cdh23 | chr10 | - | 60302747 | 60696490 | regular |
| ENSMUSG00000012848 | Rps5 | chr7 | + | 12922289 | 12926686 | regular |
| ENSMUSG00000013629 | Cad | chr5 | + | 31054779 | 31078479 | regular |
| ENSMUSG00000013663 | Pten | chr19 | + | 32757496 | 32826160 | regular |
| ENSMUSG00000013787 | Ehmt2 | chr17 | + | 34898468 | 34914052 | regular |
| ENSMUSG00000013822 | Elof1 | chr9 | - | 22112988 | 22117172 | regular |
| ENSMUSG00000014030 | Pax5 | chr4 | - | 44524756 | 44710487 | regular |
| ENSMUSG00000014077 | Chp1 | chr2 | + | 119547696 | 119587027 | regular |
| ENSMUSG00000014195 | Dnajc7 | chr11 | - | 100582817 | 100620168 | regular |
| ENSMUSG00000014294 | Ndufa2 | chr18 | - | 36742331 | 36744557 | regular |
| ENSMUSG00000014496 | Ankrd28 | chr14 | - | 31698767 | 31830651 | regular |
| ENSMUSG00000014602 | Kif1a | chr1 | - | 93015463 | 93101951 | regular |
| ENSMUSG00000014769 | Psmb1 | chr17 | - | 15475720 | 15498276 | regular |
| ENSMUSG00000014850 | Msh3 | chr13 | - | 92211871 | 92355003 | regular |
| ENSMUSG00000014867 | Surf4 | chr2 | - | 26920039 | 26933928 | regular |
| ENSMUSG00000015090 | Ptgds | chr2 | - | 25466708 | 25470046 | regular |
| ENSMUSG00000015095 | Fbxw5 | chr2 | + | 25500749 | 25505471 | regular |
| ENSMUSG00000015120 | Ube2i | chr17 | - | 25261915 | 25274622 | regular |
| ENSMUSG00000015149 | Sirt2 | chr7 | + | 28766734 | 28788661 | regular |
| ENSMUSG00000015165 | Hnrnpl | chr7 | + | 28808540 | 28822266 | regular |
| ENSMUSG00000015243 | Abca1 | chr4 | - | 53030786 | 53159895 | regular |
| ENSMUSG00000015305 | Sash1 | chr10 | - | 8722218 | 8886070 | regular |
| ENSMUSG00000015340 | Cybb | chrX | - | 9435251 | 9487771 | regular |
| ENSMUSG00000015357 | Clpx | chr9 | + | 65294259 | 65330658 | regular |
| ENSMUSG00000015365 | Mov10l1 | chr15 | + | 88982908 | 89055152 | regular |
| ENSMUSG00000015501 | Hivep2 | chr10 | + | 13966074 | 14151374 | regular |
| ENSMUSG00000015597 | Zfp318 | chr17 | + | 46383730 | 46420920 | regular |
| ENSMUSG00000015599 | Ttbk1 | chr17 | - | 46442447 | 46487675 | regular |
| ENSMUSG00000015656 | Hspa8 | chr9 | + | 40800983 | 40810087 | regular |
| ENSMUSG00000015733 | Capza2 | chr6 | + | 17636233 | 17666972 | regular |
| ENSMUSG00000015747 | Vps45 | chr3 | - | 95999831 | 96058466 | regular |
| ENSMUSG00000015755 | Tab2 | chr10 | - | 7905652 | 7956230 | regular |
| ENSMUSG00000015837 | Sqstm1 | chr11 | - | 50199365 | 50210827 | regular |
| ENSMUSG00000015932 | Dstn | chr2 | + | 143915319 | 143943324 | regular |
| ENSMUSG00000015961 | Adss | chr1 | - | 177762961 | 177796511 | regular |
| ENSMUSG00000015980 | Lrrc27 | chr7 | + | 139212987 | 139242979 | regular |
| ENSMUSG00000016253 | Nelfcd | chr2 | + | 174415803 | 174427502 | regular |
| ENSMUSG00000016541 | Atxn10 | chr15 | + | 85336380 | 85463836 | regular |
| ENSMUSG00000016559 | H3f3b | chr11 | - | 116021911 | 116027962 | regular |
| ENSMUSG00000016619 | Nup50 | chr15 | + | 84923427 | 84942955 | regular |
| ENSMUSG00000016831 | Tox4 | chr14 | + | 52279145 | 52296401 | regular |
| ENSMUSG00000016921 | Srsf6 | chr2 | + | 162931527 | 162937121 | regular |
| ENSMUSG00000017119 | Nbr1 | chr11 | + | 101552148 | 101581951 | regular |
| ENSMUSG00000017132 | Cyth1 | chr11 | - | 118132018 | 118248592 | regular |
| ENSMUSG00000017146 | Brca1 | chr11 | - | 101488763 | 101551955 | regular |

| | | | | | | |
|---|---|---|---|---|---|---|
| ENSMUSG00000017291 | Taok1 | chr11 | - | 77529161 | 77607815 | regular |
| ENSMUSG00000017404 | Rpl19 | chr11 | + | 98026709 | 98030492 | regular |
| ENSMUSG00000017421 | Zfp207 | chr11 | + | 80383278 | 80405733 | regular |
| ENSMUSG00000017707 | Serinc3 | chr2 | - | 163623271 | 163645131 | regular |
| ENSMUSG00000017774 | Myo1c | chr11 | + | 75650503 | 75673910 | regular |
| ENSMUSG00000017832 | Hspb9 | chr11 | + | 100713849 | 100714575 | regular |
| ENSMUSG00000017837 | Nkiras2 | chr11 | + | 100619243 | 100627607 | regular |
| ENSMUSG00000017843 | Ppp2r5c | chr12 | + | 110447119 | 110583062 | regular |
| ENSMUSG00000018076 | Med13l | chr5 | + | 118560678 | 118765438 | regular |
| ENSMUSG00000018196 | Glrx2 | chr1 | + | 143716337 | 143749676 | regular |
| ENSMUSG00000018209 | Stk4 | chr2 | + | 164070321 | 164155524 | regular |
| ENSMUSG00000018211 | Wfdc15b | chr2 | - | 164214453 | 164221660 | regular |
| ENSMUSG00000018339 | Gpx3 | chr11 | + | 54902452 | 54910377 | regular |
| ENSMUSG00000018372 | Cep95 | chr11 | + | 106789251 | 106819930 | regular |
| ENSMUSG00000018379 | Srsf1 | chr11 | + | 88047372 | 88053755 | regular |
| ENSMUSG00000018395 | Kif3a | chr11 | + | 53567378 | 53601967 | regular |
| ENSMUSG00000018401 | Mtmr4 | chr11 | + | 87592161 | 87616302 | regular |
| ENSMUSG00000018412 | Kansl1 | chr11 | - | 104333228 | 104468861 | regular |
| ENSMUSG00000018428 | Akap1 | chr11 | - | 88830791 | 88864586 | regular |
| ENSMUSG00000018474 | Chd3 | chr11 | - | 69343272 | 69369406 | regular |
| ENSMUSG00000018476 | Kdm6b | chr11 | - | 69398507 | 69413675 | regular |
| ENSMUSG00000018501 | Ncor1 | chr11 | - | 62316425 | 62458541 | regular |
| ENSMUSG00000018509 | Cenpv | chr11 | - | 62524945 | 62539261 | regular |
| ENSMUSG00000018548 | Trim37 | chr11 | + | 87127076 | 87220683 | regular |
| ENSMUSG00000018554 | Ybx2 | chr11 | + | 69935795 | 69941605 | regular |
| ENSMUSG00000018567 | Gabarap | chr11 | + | 69991142 | 69994951 | regular |
| ENSMUSG00000018570 | 2810408A | chr11 | - | 69897351 | 69900987 | regular |
| ENSMUSG00000018659 | Pnpo | chr11 | - | 96937824 | 96943986 | regular |
| ENSMUSG00000018669 | Cdk5rap3 | chr11 | - | 96907425 | 96916496 | regular |
| ENSMUSG00000018677 | Slc25a39 | chr11 | - | 102402984 | 102407946 | regular |
| ENSMUSG00000018697 | Aatf | chr11 | - | 84422854 | 84513522 | regular |
| ENSMUSG00000018707 | Dync1h1 | chr12 | + | 110601451 | 110666945 | regular |
| ENSMUSG00000018736 | Ndel1 | chr11 | - | 68821433 | 68871858 | regular |
| ENSMUSG00000018796 | Acsl1 | chr8 | + | 46471036 | 46536051 | regular |
| ENSMUSG00000018924 | Alox15 | chr11 | - | 70344151 | 70352031 | regular |
| ENSMUSG00000018995 | Nars2 | chr7 | + | 96951504 | 97064758 | regular |
| ENSMUSG00000019027 | Dnah1 | chr14 | - | 31260374 | 31323896 | regular |
| ENSMUSG00000019132 | BC005537 | chr13 | + | 24801656 | 24816197 | regular |
| ENSMUSG00000019139 | Isyna1 | chr8 | + | 70594372 | 70597290 | regular |
| ENSMUSG00000019188 | H13 | chr2 | + | 152669460 | 152708670 | regular |
| ENSMUSG00000019210 | Atp6v1e1 | chr6 | - | 120794304 | 120822793 | regular |
| ENSMUSG00000019338 | Zfp687 | chr3 | - | 95006589 | 95015448 | regular |
| ENSMUSG00000019370 | Calm3 | chr7 | - | 16915378 | 16924114 | regular |
| ENSMUSG00000019428 | Fkbp8 | chr8 | + | 70527723 | 70535328 | regular |
| ENSMUSG00000019505 | Ubb | chr11 | + | 62551170 | 62553213 | regular |
| ENSMUSG00000019578 | Ubxn6 | chr17 | - | 56067044 | 56075028 | regular |

| | | | | | | |
|---|---|---|---|---|---|---|
| ENSMUSG00000019590 | Cyb561 | chr11 | - | 105933701 | 105953336 | regular |
| ENSMUSG00000019699 | Akt3 | chr1 | - | 177020072 | 177258203 | regular |
| ENSMUSG00000019726 | Lyst | chr13 | + | 13590396 | 13778803 | regular |
| ENSMUSG00000019763 | Rmnd1 | chr10 | - | 4401914 | 4432388 | regular |
| ENSMUSG00000019795 | Pcmt1 | chr10 | - | 7629372 | 7681136 | regular |
| ENSMUSG00000019802 | Sec63 | chr10 | + | 42761495 | 42832514 | regular |
| ENSMUSG00000019804 | Snx3 | chr10 | + | 42502029 | 42535381 | regular |
| ENSMUSG00000019818 | Cd164 | chr10 | + | 41519413 | 41531048 | regular |
| ENSMUSG00000019820 | Utrn | chr10 | - | 12382187 | 12869365 | regular |
| ENSMUSG00000019834 | Slc22a16 | chr10 | + | 40570335 | 40604132 | regular |
| ENSMUSG00000019841 | Rev3l | chr10 | + | 39732117 | 39875211 | regular |
| ENSMUSG00000019850 | Tnfaip3 | chr10 | - | 19000909 | 19015657 | regular |
| ENSMUSG00000019856 | Fam184a | chr10 | - | 53633144 | 53751123 | regular |
| ENSMUSG00000019876 | Pkib | chr10 | + | 57631980 | 57741112 | regular |
| ENSMUSG00000019892 | Lrriq1 | chr10 | - | 103046030 | 103236322 | regular |
| ENSMUSG00000019906 | Lin7a | chr10 | + | 107271685 | 107421474 | regular |
| ENSMUSG00000019907 | Ppp1r12a | chr10 | + | 108162192 | 108284475 | regular |
| ENSMUSG00000019917 | Sep.10 | chr10 | - | 59141626 | 59221847 | regular |
| ENSMUSG00000019943 | Atp2b1 | chr10 | + | 98914405 | 99026143 | regular |
| ENSMUSG00000019947 | Arid5b | chr10 | - | 68092519 | 68278740 | regular |
| ENSMUSG00000019951 | Uhrf1bp1l | chr10 | + | 89744990 | 89819871 | regular |
| ENSMUSG00000019952 | Poc1b | chr10 | + | 99107035 | 99198074 | regular |
| ENSMUSG00000019969 | Psen1 | chr12 | + | 83688151 | 83735199 | regular |
| ENSMUSG00000019971 | Cep290 | chr10 | + | 100487557 | 100574840 | regular |
| ENSMUSG00000019977 | Hbs1l | chr10 | + | 21295978 | 21368898 | regular |
| ENSMUSG00000019978 | Epb41l2 | chr10 | + | 25359797 | 25523519 | regular |
| ENSMUSG00000019986 | Ahi1 | chr10 | + | 20952546 | 21080429 | regular |
| ENSMUSG00000019996 | Map7 | chr10 | + | 20148470 | 20281590 | regular |
| ENSMUSG00000020014 | Cfap54 | chr10 | - | 92775618 | 93081618 | regular |
| ENSMUSG00000020015 | Cdk17 | chr10 | + | 93160874 | 93267071 | regular |
| ENSMUSG00000020020 | Usp44 | chr10 | + | 93831554 | 93858088 | regular |
| ENSMUSG00000020021 | Fgd6 | chr10 | + | 94036000 | 94145339 | regular |
| ENSMUSG00000020024 | Cep83 | chr10 | + | 94688653 | 94790853 | regular |
| ENSMUSG00000020029 | Nudt4 | chr10 | - | 95547006 | 95564146 | regular |
| ENSMUSG00000020048 | Hsp90b1 | chr10 | - | 86690208 | 86705509 | regular |
| ENSMUSG00000020056 | Washc3 | chr10 | + | 88201092 | 88246158 | regular |
| ENSMUSG00000020064 | Herc4 | chr10 | + | 63243809 | 63317878 | regular |
| ENSMUSG00000020074 | Ccar1 | chr10 | - | 62743927 | 62792286 | regular |
| ENSMUSG00000020075 | Ddx21 | chr10 | - | 62580250 | 62602281 | regular |
| ENSMUSG00000020076 | Ddx50 | chr10 | - | 62615894 | 62651218 | regular |
| ENSMUSG00000020091 | Eif4ebp2 | chr10 | - | 61428490 | 61452661 | regular |
| ENSMUSG00000020102 | Slc16a7 | chr10 | - | 125219269 | 125389465 | regular |
| ENSMUSG00000020114 | Cand1 | chr10 | - | 119199254 | 119240055 | regular |
| ENSMUSG00000020120 | Plek | chr11 | - | 16971205 | 17052381 | regular |
| ENSMUSG00000020124 | Usp15 | chr10 | - | 123105005 | 123196995 | regular |
| ENSMUSG00000020131 | Pcsk4 | chr10 | - | 80321282 | 80329498 | regular |

| | | | | | | |
|---|---|---|---|---|---|---|
| ENSMUSG00000020143 | Dock2 | chr11 | - | 34226814 | 34783892 | regular |
| ENSMUSG00000020152 | Actr2 | chr11 | - | 20062303 | 20112913 | regular |
| ENSMUSG00000020156 | Mum1 | chr10 | + | 80226433 | 80243903 | regular |
| ENSMUSG00000020166 | Cnot2 | chr10 | - | 116485160 | 116581511 | regular |
| ENSMUSG00000020184 | Mdm2 | chr10 | - | 117688874 | 117710758 | regular |
| ENSMUSG00000020189 | Osbpl8 | chr10 | + | 111164751 | 111297249 | regular |
| ENSMUSG00000020196 | Cabin1 | chr10 | - | 75646111 | 75764341 | regular |
| ENSMUSG00000020198 | Ap3d1 | chr10 | - | 80706955 | 80742264 | regular |
| ENSMUSG00000020211 | Sf3a2 | chr10 | + | 80798197 | 80804924 | regular |
| ENSMUSG00000020212 | Mdm1 | chr10 | + | 118141810 | 118168997 | regular |
| ENSMUSG00000020220 | Vps13d | chr4 | - | 144972621 | 145195005 | regular |
| ENSMUSG00000020235 | Fzr1 | chr10 | - | 81366221 | 81378516 | regular |
| ENSMUSG00000020246 | Hcfc2 | chr10 | + | 82696159 | 82742428 | regular |
| ENSMUSG00000020250 | Txnrd1 | chr10 | + | 82833950 | 82897712 | regular |
| ENSMUSG00000020255 | D10Wsu1( | chr10 | + | 83360220 | 83488505 | regular |
| ENSMUSG00000020275 | Rel | chr11 | - | 23736846 | 23770970 | regular |
| ENSMUSG00000020300 | Cpeb4 | chr11 | + | 31872210 | 31935634 | regular |
| ENSMUSG00000020307 | Cdc34 | chr10 | + | 79682194 | 79688398 | regular |
| ENSMUSG00000020315 | Sptbn1 | chr11 | - | 30099394 | 30268175 | regular |
| ENSMUSG00000020317 | Theg | chr10 | - | 79576378 | 79587331 | regular |
| ENSMUSG00000020319 | Wdpcp | chr11 | + | 21572234 | 21898989 | regular |
| ENSMUSG00000020326 | Ccng1 | chr11 | - | 40748551 | 40755311 | regular |
| ENSMUSG00000020358 | Hnrnpab | chr11 | - | 51600099 | 51606847 | regular |
| ENSMUSG00000020361 | Hspa4 | chr11 | - | 53259813 | 53300457 | regular |
| ENSMUSG00000020362 | Cnot6 | chr11 | - | 49671502 | 49712723 | regular |
| ENSMUSG00000020368 | Canx | chr11 | - | 50293960 | 50325673 | regular |
| ENSMUSG00000020372 | Rack1 | chr11 | + | 48800331 | 48806434 | regular |
| ENSMUSG00000020385 | Clk4 | chr11 | + | 51261729 | 51281766 | regular |
| ENSMUSG00000020389 | Cdkl3 | chr11 | + | 52004220 | 52089784 | regular |
| ENSMUSG00000020393 | Kremen1 | chr11 | - | 5191551 | 5261558 | regular |
| ENSMUSG00000020412 | Ascc2 | chr11 | + | 4637746 | 4685699 | regular |
| ENSMUSG00000020419 | Hormad2 | chr11 | - | 4345813 | 4441105 | regular |
| ENSMUSG00000020435 | Osbp2 | chr11 | - | 3703730 | 3863903 | regular |
| ENSMUSG00000020437 | Myo1g | chr11 | - | 6506547 | 6520965 | regular |
| ENSMUSG00000020456 | Ogdh | chr11 | + | 6291632 | 6356642 | regular |
| ENSMUSG00000020458 | Rtn4 | chr11 | + | 29692946 | 29744331 | regular |
| ENSMUSG00000020460 | Rps27a | chr11 | - | 29545845 | 29548109 | regular |
| ENSMUSG00000020462 | Cfap36 | chr11 | - | 29221531 | 29247409 | regular |
| ENSMUSG00000020463 | Ppp4r3b | chr11 | + | 29172889 | 29220797 | regular |
| ENSMUSG00000020472 | Zkscan17 | chr11 | - | 59485519 | 59526751 | regular |
| ENSMUSG00000020475 | Pgam2 | chr11 | - | 5801639 | 5803733 | regular |
| ENSMUSG00000020481 | Ankrd36 | chr11 | + | 5569683 | 5689337 | regular |
| ENSMUSG00000020525 | Ppm1d | chr11 | + | 85311243 | 85347066 | regular |
| ENSMUSG00000020530 | Ggnbp2 | chr11 | - | 84832360 | 84870817 | regular |
| ENSMUSG00000020532 | Acaca | chr11 | + | 84129671 | 84401651 | regular |
| ENSMUSG00000020576 | Nbas | chr12 | + | 13269132 | 13583811 | regular |

| | | | | | | |
|---|---|---|---|---|---|---|
| ENSMUSG00000020580 | Rock2 | chr12 | + | 16894894 | 16987823 | regular |
| ENSMUSG00000020585 | Laptm4a | chr12 | + | 8921663 | 8938742 | regular |
| ENSMUSG00000020590 | Snx13 | chr12 | + | 35047185 | 35147479 | regular |
| ENSMUSG00000020593 | Lpin1 | chr12 | - | 16535668 | 16610966 | regular |
| ENSMUSG00000020594 | Pum2 | chr12 | + | 8674133 | 8752581 | regular |
| ENSMUSG00000020608 | Smc6 | chr12 | + | 11265885 | 11319785 | regular |
| ENSMUSG00000020610 | Amz2 | chr11 | + | 109425945 | 109452813 | regular |
| ENSMUSG00000020612 | Prkar1a | chr11 | + | 109649404 | 109669656 | regular |
| ENSMUSG00000020622 | Nt5c1b | chr12 | + | 10369972 | 10390175 | regular |
| ENSMUSG00000020636 | Allc | chr12 | - | 28553754 | 28582523 | regular |
| ENSMUSG00000020640 | Itsn2 | chr12 | + | 4592637 | 4713962 | regular |
| ENSMUSG00000020647 | Ncoa1 | chr12 | - | 4247361 | 4477182 | regular |
| ENSMUSG00000020650 | Bcap29 | chr12 | - | 31590966 | 31634658 | regular |
| ENSMUSG00000020657 | Dnajc27 | chr12 | + | 4082582 | 4110606 | regular |
| ENSMUSG00000020681 | Ace | chr11 | + | 105967944 | 105989964 | regular |
| ENSMUSG00000020694 | Tlk2 | chr11 | + | 105178806 | 105283959 | regular |
| ENSMUSG00000020697 | Lig3 | chr11 | + | 82781107 | 82804274 | regular |
| ENSMUSG00000020698 | Cct6b | chr11 | - | 82719249 | 82764321 | regular |
| ENSMUSG00000020716 | Nf1 | chr11 | + | 79339692 | 79581612 | regular |
| ENSMUSG00000020719 | Ddx5 | chr11 | - | 106780354 | 106789185 | regular |
| ENSMUSG00000020728 | Cep112 | chr11 | + | 108425191 | 108860615 | regular |
| ENSMUSG00000020733 | Slc9a3r1 | chr11 | + | 115163340 | 115181181 | regular |
| ENSMUSG00000020737 | Jpt1 | chr11 | - | 115497352 | 115514387 | regular |
| ENSMUSG00000020743 | Mif4gd | chr11 | - | 115607917 | 115612969 | regular |
| ENSMUSG00000020745 | Pafah1b1 | chr11 | - | 74673948 | 74724670 | regular |
| ENSMUSG00000020799 | Tekt1 | chr11 | - | 72344721 | 72362442 | regular |
| ENSMUSG00000020802 | Ube2o | chr11 | - | 116537739 | 116581447 | regular |
| ENSMUSG00000020817 | Rabep1 | chr11 | + | 70844777 | 70943105 | regular |
| ENSMUSG00000020823 | Sec14l1 | chr11 | + | 117115167 | 117159268 | regular |
| ENSMUSG00000020841 | Cpd | chr11 | - | 76778423 | 76847018 | regular |
| ENSMUSG00000020844 | Nxn | chr11 | - | 76257197 | 76399140 | regular |
| ENSMUSG00000020849 | Ywhae | chr11 | + | 75732868 | 75765845 | regular |
| ENSMUSG00000020850 | Prpf8 | chr11 | + | 75486815 | 75509449 | regular |
| ENSMUSG00000020859 | Spag9 | chr11 | + | 93996090 | 94126085 | regular |
| ENSMUSG00000020863 | Luc7l3 | chr11 | - | 94287889 | 94321988 | regular |
| ENSMUSG00000020867 | Spata20 | chr11 | - | 94478903 | 94486179 | regular |
| ENSMUSG00000020878 | Lrrc46 | chr11 | - | 97034601 | 97041407 | regular |
| ENSMUSG00000020882 | Cacnb1 | chr11 | - | 98001507 | 98023034 | regular |
| ENSMUSG00000020893 | Per1 | chr11 | + | 69095216 | 69109960 | regular |
| ENSMUSG00000020900 | Myh10 | chr11 | + | 68691558 | 68816632 | regular |
| ENSMUSG00000020917 | Acly | chr11 | - | 100476352 | 100528000 | regular |
| ENSMUSG00000020929 | Eftud2 | chr11 | - | 102838472 | 102880985 | regular |
| ENSMUSG00000020936 | Nmt1 | chr11 | + | 103028189 | 103068912 | regular |
| ENSMUSG00000020946 | Gosr2 | chr11 | - | 103676848 | 103697898 | regular |
| ENSMUSG00000020954 | Strn3 | chr12 | - | 51609631 | 51691897 | regular |
| ENSMUSG00000020982 | Nemf | chr12 | - | 69310521 | 69357165 | regular |

| | | | | | | |
|---|---|---|---|---|---|---|
| ENSMUSG00000020986 | Sec23a | chr12 | - | 58958382 | 59012017 | regular |
| ENSMUSG00000021000 | Ctage5 | chr12 | + | 59095798 | 59191583 | regular |
| ENSMUSG00000021003 | Galc | chr12 | - | 98202293 | 98259459 | regular |
| ENSMUSG00000021007 | Spata7 | chr12 | + | 98628156 | 98669815 | regular |
| ENSMUSG00000021012 | Zc3h14 | chr12 | + | 98746963 | 98787753 | regular |
| ENSMUSG00000021024 | Psma6 | chr12 | + | 55384221 | 55418454 | regular |
| ENSMUSG00000021027 | Ralgapa1 | chr12 | - | 55602895 | 55821167 | regular |
| ENSMUSG00000021065 | Fut8 | chr12 | + | 77238124 | 77476338 | regular |
| ENSMUSG00000021081 | Serpina1f | chr12 | - | 103688043 | 103695529 | regular |
| ENSMUSG00000021090 | Lrrc9 | chr12 | + | 72441865 | 72530750 | regular |
| ENSMUSG00000021096 | Ppm1a | chr12 | + | 72757454 | 72799819 | regular |
| ENSMUSG00000021097 | Clmn | chr12 | - | 104763116 | 104865076 | regular |
| ENSMUSG00000021111 | Papola | chr12 | + | 105784693 | 105838944 | regular |
| ENSMUSG00000021127 | Zfp36l1 | chr12 | - | 80107753 | 80113013 | regular |
| ENSMUSG00000021134 | Srsf5 | chr12 | + | 80945503 | 80950507 | regular |
| ENSMUSG00000021140 | Pcnx | chr12 | + | 81860022 | 82000924 | regular |
| ENSMUSG00000021143 | Pacs2 | chr12 | + | 113014507 | 113074401 | regular |
| ENSMUSG00000021147 | Wdr37 | chr13 | - | 8802967 | 8871909 | regular |
| ENSMUSG00000021156 | Zmynd11 | chr13 | - | 9684832 | 9765330 | regular |
| ENSMUSG00000021171 | Esyt2 | chr12 | + | 116281195 | 116391050 | regular |
| ENSMUSG00000021179 | Nrde2 | chr12 | - | 100125451 | 100159653 | regular |
| ENSMUSG00000021182 | Ccdc88c | chr12 | - | 100911522 | 101029056 | regular |
| ENSMUSG00000021188 | Trip11 | chr12 | - | 101834042 | 101913267 | regular |
| ENSMUSG00000021189 | Atxn3 | chr12 | - | 101918900 | 101958246 | regular |
| ENSMUSG00000021192 | Golga5 | chr12 | + | 102469134 | 102497907 | regular |
| ENSMUSG00000021196 | Pfkp | chr13 | - | 6579767 | 6648777 | regular |
| ENSMUSG00000021203 | Otub2 | chr12 | + | 103388681 | 103406350 | regular |
| ENSMUSG00000021218 | Gdi2 | chr13 | + | 3538062 | 3567871 | regular |
| ENSMUSG00000021242 | Npc2 | chr12 | - | 84754561 | 84773152 | regular |
| ENSMUSG00000021244 | Ylpm1 | chr12 | + | 84996320 | 85070515 | regular |
| ENSMUSG00000021254 | Gpatch2l | chr12 | + | 86241857 | 86291784 | regular |
| ENSMUSG00000021257 | Angel1 | chr12 | - | 86700501 | 86726460 | regular |
| ENSMUSG00000021258 | Ccnk | chr12 | + | 108179578 | 108203359 | regular |
| ENSMUSG00000021266 | Wars | chr12 | - | 108860029 | 108894174 | regular |
| ENSMUSG00000021270 | Hsp90aa1 | chr12 | - | 110690604 | 110702728 | regular |
| ENSMUSG00000021273 | Fdft1 | chr14 | - | 63145149 | 63179578 | regular |
| ENSMUSG00000021282 | Eif5 | chr12 | + | 111538015 | 111546752 | regular |
| ENSMUSG00000021285 | Ppp1r13b | chr12 | - | 111828456 | 111908110 | regular |
| ENSMUSG00000021326 | Trim27 | chr13 | + | 21179444 | 21194724 | regular |
| ENSMUSG00000021339 | Mrs2 | chr13 | - | 24987482 | 25020379 | regular |
| ENSMUSG00000021356 | Irf4 | chr13 | + | 30749225 | 30766976 | regular |
| ENSMUSG00000021357 | Exoc2 | chr13 | - | 30813918 | 30974093 | regular |
| ENSMUSG00000021366 | Hivep1 | chr13 | + | 42052020 | 42192537 | regular |
| ENSMUSG00000021374 | Nup153 | chr13 | - | 46679904 | 46727940 | regular |
| ENSMUSG00000021391 | Cenpp | chr13 | - | 49464022 | 49652785 | regular |
| ENSMUSG00000021392 | Nol8 | chr13 | + | 49653077 | 49679016 | regular |

| | | | | | | |
|---|---|---|---|---|---|---|
| ENSMUSG00000021414 | Fam217a | chr13 | - | 34909959 | 34924310 | regular |
| ENSMUSG00000021420 | Fars2 | chr13 | + | 36117411 | 36726280 | regular |
| ENSMUSG00000021457 | Syk | chr13 | + | 52583172 | 52648792 | regular |
| ENSMUSG00000021458 | 2010111I( | chr13 | + | 62964892 | 63326096 | regular |
| ENSMUSG00000021460 | Auh | chr13 | - | 52835118 | 52929681 | regular |
| ENSMUSG00000021461 | Fancc | chr13 | - | 63285042 | 63497278 | regular |
| ENSMUSG00000021476 | Habp4 | chr13 | + | 64161823 | 64186537 | regular |
| ENSMUSG00000021488 | Nsd1 | chr13 | + | 55209781 | 55318325 | regular |
| ENSMUSG00000021500 | Ddx46 | chr13 | + | 55635026 | 55681256 | regular |
| ENSMUSG00000021546 | Hnrnpk | chr13 | - | 58391141 | 58403343 | regular |
| ENSMUSG00000021552 | Gkap1 | chr13 | - | 58233345 | 58275699 | regular |
| ENSMUSG00000021557 | Agtpbp1 | chr13 | - | 59445741 | 59585227 | regular |
| ENSMUSG00000021572 | Cep72 | chr13 | - | 74036499 | 74062299 | regular |
| ENSMUSG00000021577 | Sdha | chr13 | - | 74322253 | 74350280 | regular |
| ENSMUSG00000021585 | Cast | chr13 | - | 74692367 | 74808810 | regular |
| ENSMUSG00000021615 | Xrcc4 | chr13 | - | 89774026 | 90089608 | regular |
| ENSMUSG00000021660 | Btf3 | chr13 | - | 98309895 | 98317006 | regular |
| ENSMUSG00000021669 | Col4a3bp | chr13 | + | 96542617 | 96640167 | regular |
| ENSMUSG00000021670 | Hmgcr | chr13 | - | 96648966 | 96670936 | regular |
| ENSMUSG00000021686 | Ap3b1 | chr13 | + | 94358959 | 94566317 | regular |
| ENSMUSG00000021687 | Scamp1 | chr13 | - | 94201309 | 94285857 | regular |
| ENSMUSG00000021690 | Jmy | chr13 | - | 93430100 | 93499808 | regular |
| ENSMUSG00000021693 | Kif2a | chr13 | - | 106958995 | 107022126 | regular |
| ENSMUSG00000021709 | Erbin | chr13 | - | 103818786 | 103920514 | regular |
| ENSMUSG00000021733 | Slc4a7 | chr14 | + | 14702278 | 14799940 | regular |
| ENSMUSG00000021738 | Atxn7 | chr14 | + | 13961439 | 14107302 | regular |
| ENSMUSG00000021754 | Map3k1 | chr13 | - | 111746427 | 111808993 | regular |
| ENSMUSG00000021758 | Ddx4 | chr13 | - | 112598332 | 112652475 | regular |
| ENSMUSG00000021767 | Kat6b | chr14 | + | 21481433 | 21672478 | regular |
| ENSMUSG00000021770 | Samd8 | chr14 | + | 21750530 | 21798726 | regular |
| ENSMUSG00000021771 | Vdac2 | chr14 | + | 21825237 | 21845879 | regular |
| ENSMUSG00000021785 | Ngly1 | chr14 | + | 16249279 | 16311926 | regular |
| ENSMUSG00000021807 | Rtraf | chr14 | - | 19811350 | 19823824 | regular |
| ENSMUSG00000021819 | Zswim8 | chr14 | + | 20707551 | 20723619 | regular |
| ENSMUSG00000021830 | Txndc16 | chr14 | - | 45133464 | 45220328 | regular |
| ENSMUSG00000021838 | Samd4 | chr14 | + | 46882853 | 47105815 | regular |
| ENSMUSG00000021846 | Peli2 | chr14 | + | 48120822 | 48281575 | regular |
| ENSMUSG00000021870 | Slmap | chr14 | - | 26413167 | 26534931 | regular |
| ENSMUSG00000021877 | Arf4 | chr14 | + | 26638073 | 26665084 | regular |
| ENSMUSG00000021879 | Dnah12 | chr14 | + | 26693273 | 26891703 | regular |
| ENSMUSG00000021891 | Mettl6 | chr14 | - | 31473577 | 31495040 | regular |
| ENSMUSG00000021902 | Phf7 | chr14 | - | 31237688 | 31251317 | regular |
| ENSMUSG00000021910 | Nisch | chr14 | - | 31170929 | 31216946 | regular |
| ENSMUSG00000021936 | Mapk8 | chr14 | - | 33377897 | 33447158 | regular |
| ENSMUSG00000021938 | Pspc1 | chr14 | - | 56720709 | 56778316 | regular |
| ENSMUSG00000021940 | Ptpn20 | chr14 | + | 33589206 | 33640754 | regular |

| | | | | | | |
|---|---|---|---|---|---|---|
| ENSMUSG00000021945 | Zmym2 | chr14 | + | 56886652 | 56962701 | regular |
| ENSMUSG00000021948 | Prkcd | chr14 | - | 30595353 | 30626210 | regular |
| ENSMUSG00000021969 | Zdhhc20 | chr14 | - | 57832702 | 57890276 | regular |
| ENSMUSG00000021981 | Cab39l | chr14 | + | 59440971 | 59585764 | regular |
| ENSMUSG00000021983 | Atp8a2 | chr14 | - | 59638539 | 60197179 | regular |
| ENSMUSG00000021987 | Mtmr6 | chr14 | + | 60265227 | 60302370 | regular |
| ENSMUSG00000021993 | Mipep | chr14 | + | 60784572 | 60905478 | regular |
| ENSMUSG00000021997 | Lrrc63 | chr14 | - | 75084302 | 75130881 | regular |
| ENSMUSG00000021998 | Lcp1 | chr14 | + | 75131100 | 75230842 | regular |
| ENSMUSG00000022010 | Tsc22d1 | chr14 | + | 76414960 | 76507765 | regular |
| ENSMUSG00000022013 | Dnajc15 | chr14 | - | 77820680 | 77875125 | regular |
| ENSMUSG00000022019 | Tdrd3 | chr14 | + | 87416638 | 87545504 | regular |
| ENSMUSG00000022021 | Diaph3 | chr14 | - | 86655366 | 87141235 | regular |
| ENSMUSG00000022031 | Elp3 | chr14 | - | 65530448 | 65593075 | regular |
| ENSMUSG00000022039 | Adam2 | chr14 | - | 66027328 | 66077733 | regular |
| ENSMUSG00000022056 | Adam7 | chr14 | - | 68497335 | 68533741 | regular |
| ENSMUSG00000022064 | Pibf1 | chr14 | + | 99099423 | 99254493 | regular |
| ENSMUSG00000022066 | Entpd4b | chr14 | + | 69555423 | 69585520 | regular |
| ENSMUSG00000022091 | Sorbs3 | chr14 | - | 70180467 | 70211989 | regular |
| ENSMUSG00000022100 | Xpo7 | chr14 | - | 70654245 | 70766628 | regular |
| ENSMUSG00000022108 | Itm2b | chr14 | - | 73362225 | 73385289 | regular |
| ENSMUSG00000022110 | Sucla2 | chr14 | + | 73525318 | 73596142 | regular |
| ENSMUSG00000022139 | Mbnl2 | chr14 | + | 120275668 | 120431697 | regular |
| ENSMUSG00000022141 | Nipbl | chr15 | - | 8291224 | 8444463 | regular |
| ENSMUSG00000022142 | Nup155 | chr15 | + | 8109312 | 8159857 | regular |
| ENSMUSG00000022155 | Mroh2b | chr15 | + | 4898736 | 4962205 | regular |
| ENSMUSG00000022185 | Acin1 | chr14 | - | 54642160 | 54686931 | regular |
| ENSMUSG00000022191 | Drosha | chr15 | + | 12824814 | 12935291 | regular |
| ENSMUSG00000022200 | Golph3 | chr15 | + | 12321449 | 12351591 | regular |
| ENSMUSG00000022201 | Zfr | chr15 | + | 12117830 | 12185683 | regular |
| ENSMUSG00000022214 | Dcaf11 | chr14 | + | 55560005 | 55570065 | regular |
| ENSMUSG00000022234 | Cct5 | chr15 | - | 31590799 | 31601804 | regular |
| ENSMUSG00000022236 | Ropn1l | chr15 | - | 31441210 | 31453737 | regular |
| ENSMUSG00000022246 | Rai14 | chr15 | - | 10568968 | 10714624 | regular |
| ENSMUSG00000022253 | Nadk2 | chr15 | + | 9071259 | 9110891 | regular |
| ENSMUSG00000022255 | Mtdh | chr15 | + | 34082693 | 34145624 | regular |
| ENSMUSG00000022263 | Trio | chr15 | - | 27730650 | 28025848 | regular |
| ENSMUSG00000022269 | Mar.11 | chr15 | + | 26309047 | 26409576 | regular |
| ENSMUSG00000022280 | Rnf19a | chr15 | - | 36239932 | 36283147 | regular |
| ENSMUSG00000022283 | Pabpc1 | chr15 | - | 36595660 | 36609668 | regular |
| ENSMUSG00000022285 | Ywhaz | chr15 | - | 36770769 | 36796929 | regular |
| ENSMUSG00000022307 | Oxr1 | chr15 | + | 41447481 | 41861048 | regular |
| ENSMUSG00000022312 | Eif3h | chr15 | - | 51786557 | 51865523 | regular |
| ENSMUSG00000022332 | Khdrbs3 | chr15 | + | 68928419 | 69093512 | regular |
| ENSMUSG00000022336 | Eif3e | chr15 | - | 43250057 | 43282719 | regular |
| ENSMUSG00000022369 | Mtbp | chr15 | + | 55557407 | 55626423 | regular |

| | | | | | | |
|---|---|---|---|---|---|---|
| ENSMUSG00000022377 | Asap1 | chr15 | - | 64086856 | 64382919 | regular |
| ENSMUSG00000022388 | Ttll8 | chr15 | - | 88890632 | 88954418 | regular |
| ENSMUSG00000022389 | Tef | chr15 | + | 81802420 | 81826861 | regular |
| ENSMUSG00000022391 | Rangap1 | chr15 | - | 81704247 | 81729919 | regular |
| ENSMUSG00000022401 | Xpnpep3 | chr15 | + | 81400137 | 81456807 | regular |
| ENSMUSG00000022403 | St13 | chr15 | - | 81363668 | 81400077 | regular |
| ENSMUSG00000022427 | Tomm22 | chr15 | + | 79670860 | 79672862 | regular |
| ENSMUSG00000022432 | Smc1b | chr15 | - | 85064688 | 85131964 | regular |
| ENSMUSG00000022441 | Efcab6 | chr15 | - | 83866711 | 84065379 | regular |
| ENSMUSG00000022443 | Myh9 | chr15 | - | 77760586 | 77842175 | regular |
| ENSMUSG00000022471 | Xrcc6 | chr15 | + | 81987834 | 82040085 | regular |
| ENSMUSG00000022472 | Desi1 | chr15 | - | 81992522 | 82033862 | regular |
| ENSMUSG00000022477 | Aco2 | chr15 | + | 81872308 | 81915133 | regular |
| ENSMUSG00000022487 | Gtsf1 | chr15 | - | 103402404 | 103430470 | regular |
| ENSMUSG00000022488 | Nckap1l | chr15 | + | 103453824 | 103498800 | regular |
| ENSMUSG00000022498 | Txndc11 | chr16 | - | 11074910 | 11134650 | regular |
| ENSMUSG00000022501 | Prm1 | chr16 | - | 10796325 | 10796886 | regular |
| ENSMUSG00000022517 | Mgrn1 | chr16 | + | 4886099 | 4938296 | regular |
| ENSMUSG00000022521 | Crebbp | chr16 | - | 4081327 | 4213997 | regular |
| ENSMUSG00000022533 | Atp13a3 | chr16 | - | 30312422 | 30388837 | regular |
| ENSMUSG00000022536 | Glyr1 | chr16 | - | 5013905 | 5049910 | regular |
| ENSMUSG00000022540 | Rogdi | chr16 | - | 5008729 | 5013553 | regular |
| ENSMUSG00000022542 | Sep.12 | chr16 | - | 4986857 | 4997852 | regular |
| ENSMUSG00000022558 | Mroh1 | chr15 | + | 76380260 | 76453038 | regular |
| ENSMUSG00000022562 | Oplah | chr15 | - | 76296596 | 76328015 | regular |
| ENSMUSG00000022564 | Grina | chr15 | + | 76246806 | 76249904 | regular |
| ENSMUSG00000022565 | Plec | chr15 | - | 76170973 | 76232574 | regular |
| ENSMUSG00000022568 | Scrib | chr15 | - | 76047157 | 76069784 | regular |
| ENSMUSG00000022587 | Ly6e | chr15 | + | 74955050 | 74959905 | regular |
| ENSMUSG00000022601 | Zbtb11 | chr16 | + | 55973882 | 56008913 | regular |
| ENSMUSG00000022603 | Mroh4 | chr15 | - | 74606028 | 74636353 | regular |
| ENSMUSG00000022607 | Ptk2 | chr15 | - | 73205332 | 73423280 | regular |
| ENSMUSG00000022614 | Lmf2 | chr15 | - | 89351003 | 89355659 | regular |
| ENSMUSG00000022637 | Cblb | chr16 | + | 52031224 | 52208048 | regular |
| ENSMUSG00000022641 | Bbx | chr16 | - | 50191843 | 50432390 | regular |
| ENSMUSG00000022656 | Nectin3 | chr16 | - | 46387705 | 46498525 | regular |
| ENSMUSG00000022664 | Slc35a5 | chr16 | - | 45139572 | 45158706 | regular |
| ENSMUSG00000022680 | Pdxdc1 | chr16 | - | 13833147 | 13903131 | regular |
| ENSMUSG00000022684 | Bfar | chr16 | + | 13671857 | 13703612 | regular |
| ENSMUSG00000022685 | Parn | chr16 | - | 13537963 | 13668170 | regular |
| ENSMUSG00000022707 | Gbe1 | chr16 | + | 70313948 | 70569716 | regular |
| ENSMUSG00000022708 | Zbtb20 | chr16 | + | 42875880 | 43642602 | regular |
| ENSMUSG00000022710 | Usp7 | chr16 | - | 8689594 | 8792308 | regular |
| ENSMUSG00000022770 | Dlg1 | chr16 | + | 31663442 | 31873356 | regular |
| ENSMUSG00000022773 | Ypel1 | chr16 | + | 17069695 | 17087045 | regular |
| ENSMUSG00000022781 | Pak2 | chr16 | - | 32016289 | 32079342 | regular |

| | | | | | | |
|---|---|---|---|---|---|---|
| ENSMUSG00000022783 | Spag6l | chr16 | - | 16753015 | 16829456 | regular |
| ENSMUSG00000022800 | Fyttd1 | chr16 | + | 32877499 | 32908963 | regular |
| ENSMUSG00000022801 | Lrch3 | chr16 | + | 32914099 | 33015647 | regular |
| ENSMUSG00000022805 | Maats1 | chr16 | - | 38297753 | 38342143 | regular |
| ENSMUSG00000022811 | Zfp148 | chr16 | + | 33380774 | 33503903 | regular |
| ENSMUSG00000022855 | Senp2 | chr16 | + | 22009483 | 22049269 | regular |
| ENSMUSG00000022858 | Tra2b | chr16 | - | 22244548 | 22266005 | regular |
| ENSMUSG00000022867 | Usp25 | chr16 | + | 77014068 | 77116779 | regular |
| ENSMUSG00000022884 | Eif4a2 | chr16 | + | 23107443 | 23114136 | regular |
| ENSMUSG00000022897 | Dyrk1a | chr16 | + | 94570009 | 94695517 | regular |
| ENSMUSG00000022901 | Cd86 | chr16 | - | 36603868 | 36666081 | regular |
| ENSMUSG00000022905 | Kpna1 | chr16 | + | 35978749 | 36037131 | regular |
| ENSMUSG00000022914 | Brwd1 | chr16 | - | 95992091 | 96082526 | regular |
| ENSMUSG00000022946 | Dopey2 | chr16 | + | 93711903 | 93810590 | regular |
| ENSMUSG00000022957 | Itsn1 | chr16 | + | 91729280 | 91920597 | regular |
| ENSMUSG00000022961 | Son | chr16 | + | 91647505 | 91679221 | regular |
| ENSMUSG00000022982 | Sod1 | chr16 | + | 90220741 | 90226322 | regular |
| ENSMUSG00000022995 | Enah | chr1 | - | 181896383 | 182019990 | regular |
| ENSMUSG00000022999 | Lmbr1l | chr15 | - | 98903920 | 98918231 | regular |
| ENSMUSG00000023010 | Tmbim6 | chr15 | + | 99392881 | 99410049 | regular |
| ENSMUSG00000023026 | Dip2b | chr15 | + | 100038663 | 100219473 | regular |
| ENSMUSG00000023027 | Atf1 | chr15 | + | 100227818 | 100261244 | regular |
| ENSMUSG00000023072 | Cep89 | chr7 | + | 35397034 | 35438689 | regular |
| ENSMUSG00000023084 | Lrrc71 | chr3 | - | 87736922 | 87748625 | regular |
| ENSMUSG00000023104 | Rfc2 | chr5 | + | 134581365 | 134601805 | regular |
| ENSMUSG00000023118 | Sympk | chr7 | + | 19024376 | 19054618 | regular |
| ENSMUSG00000023150 | Ivns1abp | chr1 | + | 151344476 | 151364422 | regular |
| ENSMUSG00000023151 | Lrrc69 | chr4 | - | 14623619 | 14796060 | regular |
| ENSMUSG00000023403 | Stk31 | chr6 | + | 49395603 | 49469501 | regular |
| ENSMUSG00000023456 | Tpi1 | chr6 | - | 124810585 | 124814296 | regular |
| ENSMUSG00000023467 | Tulp2 | chr7 | + | 45481876 | 45523632 | regular |
| ENSMUSG00000023495 | Pcbp4 | chr9 | + | 106453290 | 106464012 | regular |
| ENSMUSG00000023577 | Iqcf3 | chr9 | - | 106503224 | 106561631 | regular |
| ENSMUSG00000023806 | Rsph3b | chr17 | - | 6904715 | 6948356 | regular |
| ENSMUSG00000023852 | Chd1 | chr17 | + | 15704966 | 17394898 | regular |
| ENSMUSG00000023873 | 1700010I1 | chr17 | + | 8988332 | 9008319 | regular |
| ENSMUSG00000023923 | Tbc1d5 | chr17 | - | 50733123 | 51179352 | regular |
| ENSMUSG00000023927 | Satb1 | chr17 | - | 51736186 | 51833290 | regular |
| ENSMUSG00000023930 | Crisp2 | chr17 | - | 40764733 | 40807003 | regular |
| ENSMUSG00000023944 | Hsp90ab1 | chr17 | - | 45567774 | 45573271 | regular |
| ENSMUSG00000023949 | Tcte1 | chr17 | + | 45523433 | 45542679 | regular |
| ENSMUSG00000023952 | Gtpbp2 | chr17 | + | 46161031 | 46169370 | regular |
| ENSMUSG00000023953 | Polh | chr17 | - | 46172003 | 46202625 | regular |
| ENSMUSG00000024006 | Stk38 | chr17 | - | 28970884 | 29007945 | regular |
| ENSMUSG00000024008 | Cpne5 | chr17 | - | 29156549 | 29237797 | regular |
| ENSMUSG00000024014 | Pim1 | chr17 | + | 29490811 | 29496111 | regular |

| | | | | | | |
|---|---|---|---|---|---|---|
| ENSMUSG00000024033 | Rsph1 | chr17 | - | 31255018 | 31277356 | regular |
| ENSMUSG00000024042 | Sik1 | chr17 | - | 31844247 | 31855792 | regular |
| ENSMUSG00000024053 | Emilin2 | chr17 | - | 71252175 | 71310965 | regular |
| ENSMUSG00000024059 | Clip4 | chr17 | + | 71781946 | 71858351 | regular |
| ENSMUSG00000024073 | Birc6 | chr17 | + | 74528294 | 74703356 | regular |
| ENSMUSG00000024085 | Man2a1 | chr17 | + | 64600735 | 64755110 | regular |
| ENSMUSG00000024112 | Cacna1h | chr17 | - | 25374284 | 25433783 | regular |
| ENSMUSG00000024122 | Pdpk1 | chr17 | - | 24073679 | 24150924 | regular |
| ENSMUSG00000024135 | Srbd1 | chr17 | - | 85984664 | 86145175 | regular |
| ENSMUSG00000024154 | Gtf2a1l | chr17 | + | 88668659 | 88715152 | regular |
| ENSMUSG00000024160 | Spsb3 | chr17 | + | 24886642 | 24892152 | regular |
| ENSMUSG00000024163 | Mapk8ip3 | chr17 | - | 24892152 | 24936977 | regular |
| ENSMUSG00000024169 | Ift140 | chr17 | + | 25016090 | 25099495 | regular |
| ENSMUSG00000024180 | Tmem8 | chr17 | + | 26113315 | 26123254 | regular |
| ENSMUSG00000024188 | Luc7l | chr17 | + | 26252909 | 26285504 | regular |
| ENSMUSG00000024206 | Rfx2 | chr17 | - | 56775896 | 56831008 | regular |
| ENSMUSG00000024207 | Acsbg2 | chr17 | - | 56843102 | 56874447 | regular |
| ENSMUSG00000024208 | Uqcc2 | chr17 | - | 27122662 | 27133916 | regular |
| ENSMUSG00000024222 | Fkbp5 | chr17 | - | 28399094 | 28517524 | regular |
| ENSMUSG00000024223 | Armc12 | chr17 | + | 28530859 | 28538954 | regular |
| ENSMUSG00000024238 | Zeb1 | chr18 | + | 5591859 | 5775467 | regular |
| ENSMUSG00000024240 | Epc1 | chr18 | - | 6435950 | 6516108 | regular |
| ENSMUSG00000024242 | Map4k3 | chr17 | - | 80580511 | 80728093 | regular |
| ENSMUSG00000024248 | Cox7a2l | chr17 | - | 83501918 | 83514333 | regular |
| ENSMUSG00000024251 | Thada | chr17 | - | 84190055 | 84466196 | regular |
| ENSMUSG00000024269 | Tpgs2 | chr18 | - | 25127222 | 25169007 | regular |
| ENSMUSG00000024283 | Wac | chr18 | + | 7868831 | 7973547 | regular |
| ENSMUSG00000024286 | Ccny | chr18 | - | 9314043 | 9450150 | regular |
| ENSMUSG00000024290 | Rock1 | chr18 | - | 10064400 | 10181792 | regular |
| ENSMUSG00000024293 | Esco1 | chr18 | - | 10566601 | 10610352 | regular |
| ENSMUSG00000024294 | Mib1 | chr18 | + | 10725547 | 10818704 | regular |
| ENSMUSG00000024306 | Ccdc178 | chr18 | - | 21810896 | 22171396 | regular |
| ENSMUSG00000024317 | Rnf138 | chr18 | + | 21001340 | 21028223 | regular |
| ENSMUSG00000024335 | Brd2 | chr17 | - | 34112022 | 34122634 | regular |
| ENSMUSG00000024352 | Spata24 | chr18 | - | 35656689 | 35662186 | regular |
| ENSMUSG00000024357 | Sil1 | chr18 | - | 35266399 | 35498925 | regular |
| ENSMUSG00000024359 | Hspa9 | chr18 | - | 34937413 | 34954357 | regular |
| ENSMUSG00000024369 | Nelfe | chr17 | + | 34850390 | 34856372 | regular |
| ENSMUSG00000024387 | Csnk2b | chr17 | - | 35116195 | 35122053 | regular |
| ENSMUSG00000024392 | Bag6 | chr17 | + | 35135177 | 35147322 | regular |
| ENSMUSG00000024393 | Prrc2a | chr17 | - | 35149075 | 35164897 | regular |
| ENSMUSG00000024397 | Aif1 | chr17 | - | 35170990 | 35176068 | regular |
| ENSMUSG00000024400 | Wdr33 | chr18 | + | 31804056 | 31908987 | regular |
| ENSMUSG00000024404 | Riok3 | chr18 | + | 12128849 | 12157367 | regular |
| ENSMUSG00000024429 | Gnl1 | chr17 | + | 35979850 | 35989462 | regular |
| ENSMUSG00000024430 | Cabyr | chr18 | + | 12741323 | 12755146 | regular |

| | | | | | | |
|---|---|---|---|---|---|---|
| ENSMUSG00000024456 | Diaph1 | chr18 | - | 37843600 | 37935476 | regular |
| ENSMUSG00000024474 | Ik | chr18 | + | 36744655 | 36757639 | regular |
| ENSMUSG00000024483 | Ankhd1 | chr18 | + | 36559986 | 36665917 | regular |
| ENSMUSG00000024491 | Rbm27 | chr18 | + | 42275352 | 42341542 | regular |
| ENSMUSG00000024500 | Ppp2r2b | chr18 | - | 42637431 | 43059471 | regular |
| ENSMUSG00000024513 | Mbd2 | chr18 | + | 70568188 | 70626131 | regular |
| ENSMUSG00000024515 | Smad4 | chr18 | - | 73639008 | 73703780 | regular |
| ENSMUSG00000024542 | Cep192 | chr18 | + | 67800106 | 67885170 | regular |
| ENSMUSG00000024566 | Atp9b | chr18 | - | 80734140 | 80934058 | regular |
| ENSMUSG00000024576 | Csnk1a1 | chr18 | + | 61555273 | 61590061 | regular |
| ENSMUSG00000024608 | Rps14 | chr18 | + | 60774509 | 60778546 | regular |
| ENSMUSG00000024610 | Cd74 | chr18 | + | 60803847 | 60812646 | regular |
| ENSMUSG00000024622 | Hmgxb3 | chr18 | - | 61131278 | 61177050 | regular |
| ENSMUSG00000024654 | Asrgl1 | chr19 | - | 9109867 | 9135636 | regular |
| ENSMUSG00000024661 | Fth1 | chr19 | + | 9982702 | 9985092 | regular |
| ENSMUSG00000024663 | Rab3il1 | chr19 | + | 10001668 | 10038380 | regular |
| ENSMUSG00000024687 | Osbp | chr19 | + | 11965843 | 11994105 | regular |
| ENSMUSG00000024695 | Zfp91 | chr19 | - | 12763659 | 12796126 | regular |
| ENSMUSG00000024696 | Lpxn | chr19 | + | 12798605 | 12833807 | regular |
| ENSMUSG00000024740 | Ddb1 | chr19 | + | 10605624 | 10629813 | regular |
| ENSMUSG00000024749 | Tmc1 | chr19 | - | 20783457 | 20954202 | regular |
| ENSMUSG00000024758 | Rtn3 | chr19 | - | 7425900 | 7483289 | regular |
| ENSMUSG00000024759 | Atl3 | chr19 | + | 7494039 | 7538608 | regular |
| ENSMUSG00000024767 | Otub1 | chr19 | - | 7198201 | 7206316 | regular |
| ENSMUSG00000024772 | Ehd1 | chr19 | + | 6276724 | 6300096 | regular |
| ENSMUSG00000024780 | Cdc37l1 | chr19 | + | 28990351 | 29026681 | regular |
| ENSMUSG00000024789 | Jak2 | chr19 | + | 29251827 | 29313080 | regular |
| ENSMUSG00000024853 | Sf3b2 | chr19 | - | 5273922 | 5295455 | regular |
| ENSMUSG00000024855 | Pacs1 | chr19 | - | 5133687 | 5273119 | regular |
| ENSMUSG00000024908 | Ppp6r3 | chr19 | - | 3454927 | 3575749 | regular |
| ENSMUSG00000024913 | Lrp5 | chr19 | - | 3584827 | 3686564 | regular |
| ENSMUSG00000024914 | Drap1 | chr19 | - | 5422804 | 5424979 | regular |
| ENSMUSG00000024921 | Smarca2 | chr19 | + | 26605049 | 26778322 | regular |
| ENSMUSG00000024937 | Ehbp1l1 | chr19 | - | 5707375 | 5726317 | regular |
| ENSMUSG00000024943 | Smc5 | chr19 | - | 23206450 | 23273897 | regular |
| ENSMUSG00000024948 | Map4k2 | chr19 | + | 6341134 | 6355615 | regular |
| ENSMUSG00000024949 | Sf1 | chr19 | + | 6363689 | 6378030 | regular |
| ENSMUSG00000024953 | Prdx5 | chr19 | - | 6906696 | 6910106 | regular |
| ENSMUSG00000024973 | Hrasls5 | chr19 | + | 7612540 | 7639642 | regular |
| ENSMUSG00000024974 | Smc3 | chr19 | + | 53600397 | 53645833 | regular |
| ENSMUSG00000024976 | Shoc2 | chr19 | + | 53944305 | 54033268 | regular |
| ENSMUSG00000024991 | Eif3a | chr19 | - | 60761115 | 60790693 | regular |
| ENSMUSG00000025019 | Lcor | chr19 | + | 41482644 | 41562246 | regular |
| ENSMUSG00000025025 | Mxi1 | chr19 | + | 53310505 | 53375810 | regular |
| ENSMUSG00000025026 | Add3 | chr19 | + | 53140444 | 53247399 | regular |
| ENSMUSG00000025041 | Nt5c2 | chr19 | - | 46886830 | 47015153 | regular |

| | | | | | | |
|---|---|---|---|---|---|---|
| ENSMUSG00000025047 | Pdcd11 | chr19 | + | 47090765 | 47131143 | regular |
| ENSMUSG00000025081 | Tdrd1 | chr19 | + | 56826208 | 56870012 | regular |
| ENSMUSG00000025103 | Btbd1 | chr7 | - | 81792073 | 81829431 | regular |
| ENSMUSG00000025104 | Hdgfl3 | chr7 | - | 81881250 | 81934473 | regular |
| ENSMUSG00000025130 | P4hb | chr11 | - | 120560297 | 120573253 | regular |
| ENSMUSG00000025132 | Arhgdia | chr11 | - | 120578103 | 120581624 | regular |
| ENSMUSG00000025139 | Tollip | chr7 | - | 141874812 | 141918507 | regular |
| ENSMUSG00000025142 | Aspscr1 | chr11 | + | 120672972 | 120709447 | regular |
| ENSMUSG00000025153 | Fasn | chr11 | - | 120805845 | 120824547 | regular |
| ENSMUSG00000025156 | Gps1 | chr11 | + | 120784271 | 120789102 | regular |
| ENSMUSG00000025159 | Mms19 | chr19 | - | 41941085 | 41981157 | regular |
| ENSMUSG00000025162 | Csnk1d | chr11 | - | 120961748 | 120991330 | regular |
| ENSMUSG00000025171 | Ubtd1 | chr19 | + | 41981762 | 42034641 | regular |
| ENSMUSG00000025178 | Pi4k2a | chr19 | + | 42090434 | 42122218 | regular |
| ENSMUSG00000025217 | Btrc | chr19 | + | 45363733 | 45530013 | regular |
| ENSMUSG00000025220 | Mgea5 | chr19 | - | 45750260 | 45783520 | regular |
| ENSMUSG00000025224 | Gbf1 | chr19 | + | 46152508 | 46286510 | regular |
| ENSMUSG00000025228 | Actr1a | chr19 | - | 46376813 | 46395735 | regular |
| ENSMUSG00000025234 | Arih1 | chr9 | - | 59388257 | 59486618 | regular |
| ENSMUSG00000025239 | Limd1 | chr9 | + | 123478705 | 123521552 | regular |
| ENSMUSG00000025241 | Fyco1 | chr9 | - | 123789499 | 123851899 | regular |
| ENSMUSG00000025245 | Lztfl1 | chr9 | - | 123694415 | 123717697 | regular |
| ENSMUSG00000025261 | Huwe1 | chrX | + | 151800806 | 151935417 | regular |
| ENSMUSG00000025278 | Flnb | chr14 | + | 7817956 | 7951588 | regular |
| ENSMUSG00000025283 | Sat1 | chrX | - | 155213131 | 155216449 | regular |
| ENSMUSG00000025290 | Rps24 | chr14 | + | 24487124 | 24496959 | regular |
| ENSMUSG00000025314 | Ptprj | chr2 | - | 90429753 | 90580647 | regular |
| ENSMUSG00000025326 | Ube3a | chr7 | + | 59228749 | 59311536 | regular |
| ENSMUSG00000025340 | Rabgef1 | chr5 | + | 130171797 | 130214342 | regular |
| ENSMUSG00000025357 | Dgka | chr10 | - | 128720133 | 128744855 | regular |
| ENSMUSG00000025362 | Rps26 | chr10 | - | 128624533 | 128626747 | regular |
| ENSMUSG00000025369 | Smarcc2 | chr10 | + | 128459247 | 128490482 | regular |
| ENSMUSG00000025372 | Baiap2 | chr11 | + | 119942762 | 120006782 | regular |
| ENSMUSG00000025384 | Faap100 | chr11 | - | 120369054 | 120378764 | regular |
| ENSMUSG00000025393 | Atp5b | chr10 | + | 128083272 | 128090391 | regular |
| ENSMUSG00000025404 | R3hdm2 | chr10 | + | 127380326 | 127499384 | regular |
| ENSMUSG00000025410 | Dctn2 | chr10 | + | 127266367 | 127281950 | regular |
| ENSMUSG00000025423 | Pias2 | chr18 | + | 77065207 | 77155708 | regular |
| ENSMUSG00000025428 | Atp5a1 | chr18 | + | 77773728 | 77782869 | regular |
| ENSMUSG00000025431 | Crisp1 | chr17 | - | 40293757 | 40319207 | regular |
| ENSMUSG00000025453 | Nnt | chr13 | - | 119335447 | 119408997 | regular |
| ENSMUSG00000025477 | Inpp5a | chr7 | + | 139389108 | 139579652 | regular |
| ENSMUSG00000025480 | Syce1 | chr7 | - | 140777228 | 140787852 | regular |
| ENSMUSG00000025495 | Ptdss2 | chr7 | + | 141122381 | 141157606 | regular |
| ENSMUSG00000025500 | Lmntd2 | chr7 | - | 141209991 | 141214080 | regular |
| ENSMUSG00000025508 | Rplp2 | chr7 | + | 141447369 | 141451585 | regular |

| | | | | | | |
|---|---|---|---|---|---|---|
| ENSMUSG00000025532 | Crcp | chr5 | + | 130029289 | 130060789 | regular |
| ENSMUSG00000025558 | Dock9 | chr14 | - | 121542045 | 121797837 | regular |
| ENSMUSG00000025571 | Tnrc6c | chr11 | + | 117654288 | 117763439 | regular |
| ENSMUSG00000025583 | Rptor | chr11 | + | 119602904 | 119899576 | regular |
| ENSMUSG00000025584 | Pde8a | chr7 | + | 81213595 | 81334533 | regular |
| ENSMUSG00000025648 | Pfkfb4 | chr9 | + | 108991777 | 109032228 | regular |
| ENSMUSG00000025651 | Uqcrc1 | chr9 | + | 108936632 | 108949623 | regular |
| ENSMUSG00000025742 | Prps2 | chrX | - | 167346321 | 167382749 | regular |
| ENSMUSG00000025745 | Hadha | chr5 | - | 30118303 | 30155162 | regular |
| ENSMUSG00000025757 | Hspa4l | chr3 | + | 40744494 | 40796103 | regular |
| ENSMUSG00000025762 | Larp1b | chr3 | + | 40950353 | 41040234 | regular |
| ENSMUSG00000025781 | Atp5c1 | chr2 | - | 10056015 | 10080510 | regular |
| ENSMUSG00000025793 | Hgs | chr11 | + | 120467634 | 120483984 | regular |
| ENSMUSG00000025794 | Rpl14 | chr9 | + | 120571443 | 120574654 | regular |
| ENSMUSG00000025808 | Ccdc7a | chr8 | - | 128734234 | 129065517 | regular |
| ENSMUSG00000025862 | Stag2 | chrX | + | 42149316 | 42277185 | regular |
| ENSMUSG00000025871 | 4833439L | chr13 | - | 54551217 | 54565435 | regular |
| ENSMUSG00000025885 | Myo5b | chr18 | + | 74440935 | 74771493 | regular |
| ENSMUSG00000025900 | Rp1 | chr1 | - | 3999556 | 4409241 | regular |
| ENSMUSG00000025903 | Lypla1 | chr1 | + | 4807787 | 4848410 | regular |
| ENSMUSG00000025907 | Rb1cc1 | chr1 | + | 6206196 | 6276648 | regular |
| ENSMUSG00000025909 | Sntg1 | chr1 | - | 8361474 | 9299878 | regular |
| ENSMUSG00000025920 | Stau2 | chr1 | - | 16228673 | 16520112 | regular |
| ENSMUSG00000025939 | Ube2w | chr1 | - | 16540789 | 16619489 | regular |
| ENSMUSG00000025949 | Pikfyve | chr1 | + | 65186682 | 65278695 | regular |
| ENSMUSG00000025958 | Creb1 | chr1 | + | 64532644 | 64604548 | regular |
| ENSMUSG00000025967 | Eef1b2 | chr1 | + | 63176824 | 63180486 | regular |
| ENSMUSG00000025978 | Rftn2 | chr1 | - | 55170158 | 55226782 | regular |
| ENSMUSG00000025982 | Sf3b1 | chr1 | - | 54985168 | 55027481 | regular |
| ENSMUSG00000025983 | Ccdc150 | chr1 | + | 54250682 | 54368727 | regular |
| ENSMUSG00000025986 | Slc39a10 | chr1 | - | 46807543 | 46892852 | regular |
| ENSMUSG00000026000 | Lancl1 | chr1 | - | 67000516 | 67038872 | regular |
| ENSMUSG00000026004 | Kansl1l | chr1 | - | 66719247 | 66817562 | regular |
| ENSMUSG00000026017 | Carf | chr1 | + | 60098246 | 60153953 | regular |
| ENSMUSG00000026018 | Ica1l | chr1 | - | 59982489 | 60043184 | regular |
| ENSMUSG00000026034 | Clk1 | chr1 | - | 58410188 | 58424066 | regular |
| ENSMUSG00000026082 | Rev1 | chr1 | - | 38052785 | 38129801 | regular |
| ENSMUSG00000026083 | Eif5b | chr1 | + | 37998009 | 38055579 | regular |
| ENSMUSG00000026094 | Stk17b | chr1 | - | 53755505 | 53785224 | regular |
| ENSMUSG00000026107 | Nabp1 | chr1 | - | 51465861 | 51478425 | regular |
| ENSMUSG00000026113 | Inpp4a | chr1 | + | 37299864 | 37410736 | regular |
| ENSMUSG00000026125 | Prss39 | chr1 | + | 34498409 | 34503063 | regular |
| ENSMUSG00000026131 | Dst | chr1 | + | 33908224 | 34308661 | regular |
| ENSMUSG00000026134 | Prim2 | chr1 | - | 33453809 | 33669795 | regular |
| ENSMUSG00000026150 | Mff | chr1 | + | 82724889 | 82752394 | regular |
| ENSMUSG00000026155 | Smap1 | chr1 | - | 23844845 | 23922317 | regular |

| | | | | | | |
|---|---|---|---|---|---|---|
| ENSMUSG00000026159 | Agfg1 | chr1 | + | 82839482 | 82901182 | regular |
| ENSMUSG00000026162 | Nhej1 | chr1 | - | 74967138 | 75125200 | regular |
| ENSMUSG00000026163 | Sphkap | chr1 | - | 83254138 | 83408200 | regular |
| ENSMUSG00000026173 | Plcd4 | chr1 | + | 74542887 | 74567794 | regular |
| ENSMUSG00000026182 | Tnp1 | chr1 | - | 73015073 | 73015909 | regular |
| ENSMUSG00000026193 | Fn1 | chr1 | - | 71585519 | 71653200 | regular |
| ENSMUSG00000026200 | Glb1l | chr1 | - | 75198235 | 75210813 | regular |
| ENSMUSG00000026203 | Dnajb2 | chr1 | + | 75236405 | 75245692 | regular |
| ENSMUSG00000026219 | Trip12 | chr1 | - | 84721188 | 84840516 | regular |
| ENSMUSG00000026226 | Spata3 | chr1 | + | 86016815 | 86029958 | regular |
| ENSMUSG00000026234 | Ncl | chr1 | - | 86344718 | 86359400 | regular |
| ENSMUSG00000026254 | Eif4e2 | chr1 | + | 87213913 | 87240488 | regular |
| ENSMUSG00000026269 | Rnpepl1 | chr1 | + | 92910782 | 92924384 | regular |
| ENSMUSG00000026276 | Sep.02 | chr1 | + | 93478963 | 93510260 | regular |
| ENSMUSG00000026277 | Stk25 | chr1 | - | 93619750 | 93658659 | regular |
| ENSMUSG00000026288 | Inpp5d | chr1 | + | 87620311 | 87720507 | regular |
| ENSMUSG00000026305 | Lrrfip1 | chr1 | + | 90998736 | 91128944 | regular |
| ENSMUSG00000026341 | Actr3 | chr1 | - | 125392904 | 125435727 | regular |
| ENSMUSG00000026353 | Ubxn4 | chr1 | + | 128243963 | 128279378 | regular |
| ENSMUSG00000026361 | Cdc73 | chr1 | - | 143598799 | 143702893 | regular |
| ENSMUSG00000026380 | Tfcp2l1 | chr1 | + | 118627944 | 118685168 | regular |
| ENSMUSG00000026383 | Epb41l5 | chr1 | - | 119545036 | 119649000 | regular |
| ENSMUSG00000026384 | Ptpn4 | chr1 | - | 119652466 | 119837613 | regular |
| ENSMUSG00000026395 | Ptprc | chr1 | - | 138062860 | 138175708 | regular |
| ENSMUSG00000026399 | Cd55 | chr1 | - | 130439026 | 130462744 | regular |
| ENSMUSG00000026434 | Nucks1 | chr1 | + | 131910533 | 131936321 | regular |
| ENSMUSG00000026457 | Adipor1 | chr1 | + | 134415377 | 134433351 | regular |
| ENSMUSG00000026463 | Atp2b4 | chr1 | - | 133699456 | 133801041 | regular |
| ENSMUSG00000026466 | Tor1aip1 | chr1 | - | 156004598 | 156036480 | regular |
| ENSMUSG00000026473 | Glul | chr1 | + | 153899943 | 153909723 | regular |
| ENSMUSG00000026482 | Rgl1 | chr1 | - | 152516759 | 152766351 | regular |
| ENSMUSG00000026491 | Ahctf1 | chr1 | - | 179744893 | 179803680 | regular |
| ENSMUSG00000026499 | Acbd3 | chr1 | + | 180726042 | 180754204 | regular |
| ENSMUSG00000026504 | Sdccag8 | chr1 | + | 176814659 | 177020437 | regular |
| ENSMUSG00000026516 | Nvl | chr1 | - | 181087137 | 181144204 | regular |
| ENSMUSG00000026523 | Wdr64 | chr1 | + | 175698592 | 175815734 | regular |
| ENSMUSG00000026553 | Copa | chr1 | + | 172082528 | 172122330 | regular |
| ENSMUSG00000026554 | Dcaf8 | chr1 | + | 172148083 | 172197005 | regular |
| ENSMUSG00000026565 | Pou2f1 | chr1 | - | 165865153 | 166002678 | regular |
| ENSMUSG00000026567 | Adcy10 | chr1 | + | 165485182 | 165576774 | regular |
| ENSMUSG00000026568 | Mpc2 | chr1 | + | 165460636 | 165481214 | regular |
| ENSMUSG00000026571 | Dcaf6 | chr1 | - | 165328697 | 165460475 | regular |
| ENSMUSG00000026575 | Nme7 | chr1 | + | 164304120 | 164437725 | regular |
| ENSMUSG00000026578 | Ccdc181 | chr1 | + | 164275584 | 164287847 | regular |
| ENSMUSG00000026585 | Kifap3 | chr1 | + | 163779582 | 163917109 | regular |
| ENSMUSG00000026594 | Ralgps2 | chr1 | - | 156804165 | 156939626 | regular |

| | | | | | | |
|---|---|---|---|---|---|---|
| ENSMUSG00000026601 | Axdnd1 | chr1 | - | 156323508 | 156421159 | regular |
| ENSMUSG00000026605 | Cenpf | chr1 | - | 189640605 | 189688086 | regular |
| ENSMUSG00000026609 | Ush2a | chr1 | + | 188262022 | 188965041 | regular |
| ENSMUSG00000026611 | Spata17 | chr1 | - | 187044647 | 187215465 | regular |
| ENSMUSG00000026615 | Eprs | chr1 | + | 185363043 | 185428360 | regular |
| ENSMUSG00000026650 | Meig1 | chr2 | - | 3409042 | 3422648 | regular |
| ENSMUSG00000026655 | Fam107b | chr2 | + | 3570487 | 3782142 | regular |
| ENSMUSG00000026663 | Atf6 | chr1 | - | 170704673 | 170867771 | regular |
| ENSMUSG00000026670 | Uap1 | chr1 | - | 170141937 | 170174957 | regular |
| ENSMUSG00000026672 | Optn | chr2 | - | 5020641 | 5064051 | regular |
| ENSMUSG00000026701 | Prdx6 | chr1 | - | 161240111 | 161251219 | regular |
| ENSMUSG00000026721 | Rabgap1l | chr1 | - | 160219173 | 160793211 | regular |
| ENSMUSG00000026739 | Bmi1 | chr2 | + | 18677017 | 18686629 | regular |
| ENSMUSG00000026743 | Mllt10 | chr2 | + | 18055236 | 18212388 | regular |
| ENSMUSG00000026766 | Mmadhc | chr2 | - | 50279880 | 50296801 | regular |
| ENSMUSG00000026773 | Pfkfb3 | chr2 | - | 11471432 | 11554077 | regular |
| ENSMUSG00000026775 | Yme1l1 | chr2 | + | 23156368 | 23199260 | regular |
| ENSMUSG00000026778 | Prkcq | chr2 | + | 11172107 | 11301222 | regular |
| ENSMUSG00000026781 | Acbd5 | chr2 | + | 23068166 | 23115558 | regular |
| ENSMUSG00000026790 | Odf2 | chr2 | + | 29889220 | 29931746 | regular |
| ENSMUSG00000026807 | Ak8 | chr2 | + | 28700163 | 28813165 | regular |
| ENSMUSG00000026809 | Spaca9 | chr2 | - | 28692079 | 28699727 | regular |
| ENSMUSG00000026817 | Ak1 | chr2 | + | 32621757 | 32635058 | regular |
| ENSMUSG00000026821 | Ralgds | chr2 | + | 28513124 | 28553081 | regular |
| ENSMUSG00000026827 | Gpd2 | chr2 | + | 57237634 | 57370719 | regular |
| ENSMUSG00000026832 | Cytip | chr2 | - | 58129136 | 58195532 | regular |
| ENSMUSG00000026842 | Abl1 | chr2 | + | 31688375 | 31804227 | regular |
| ENSMUSG00000026853 | Crat | chr2 | - | 30400470 | 30415813 | regular |
| ENSMUSG00000026864 | Hspa5 | chr2 | + | 34771969 | 34777547 | regular |
| ENSMUSG00000026867 | Gapvd1 | chr2 | - | 34674593 | 34755232 | regular |
| ENSMUSG00000026872 | Zeb2 | chr2 | - | 44983631 | 45117395 | regular |
| ENSMUSG00000026878 | Rab14 | chr2 | - | 35180204 | 35201120 | regular |
| ENSMUSG00000026915 | Strbp | chr2 | - | 37483227 | 37703859 | regular |
| ENSMUSG00000026922 | Agpat2 | chr2 | - | 26593056 | 26604417 | regular |
| ENSMUSG00000026924 | Sec16a | chr2 | - | 26409430 | 26445216 | regular |
| ENSMUSG00000026927 | Sdccag3 | chr2 | - | 26382801 | 26389316 | regular |
| ENSMUSG00000026931 | 1700019N | chr19 | - | 58785802 | 58794461 | regular |
| ENSMUSG00000026933 | Camsap1 | chr2 | - | 25926837 | 25983282 | regular |
| ENSMUSG00000026940 | Ccdc183 | chr2 | - | 25608634 | 25617678 | regular |
| ENSMUSG00000026969 | Fam166a | chr2 | + | 25218744 | 25222280 | regular |
| ENSMUSG00000026977 | Mar.07 | chr2 | + | 60209886 | 60250676 | regular |
| ENSMUSG00000026987 | Baz2b | chr2 | - | 59899362 | 60209839 | regular |
| ENSMUSG00000027006 | Dnajc10 | chr2 | + | 80315465 | 80354043 | regular |
| ENSMUSG00000027007 | Ssfa2 | chr2 | + | 79635351 | 79672966 | regular |
| ENSMUSG00000027009 | Itga4 | chr2 | + | 79255425 | 79333123 | regular |
| ENSMUSG00000027012 | Dync1i2 | chr2 | + | 71211705 | 71263303 | regular |

| | | | | | | |
|---|---|---|---|---|---|---|
| ENSMUSG00000027030 | Stk39 | chr2 | - | 68210444 | 68472268 | regular |
| ENSMUSG00000027067 | Ssrp1 | chr2 | + | 85037233 | 85047109 | regular |
| ENSMUSG00000027104 | Atf2 | chr2 | - | 73816508 | 73892639 | regular |
| ENSMUSG00000027157 | 4930430A | chr2 | - | 111162060 | 111229602 | regular |
| ENSMUSG00000027177 | Hipk3 | chr2 | - | 104426480 | 104494446 | regular |
| ENSMUSG00000027184 | Caprin1 | chr2 | - | 103762940 | 103797649 | regular |
| ENSMUSG00000027227 | Sord | chr2 | + | 122234748 | 122265340 | regular |
| ENSMUSG00000027248 | Pdia3 | chr2 | + | 121413774 | 121438687 | regular |
| ENSMUSG00000027282 | Mtch2 | chr2 | + | 90847154 | 90866810 | regular |
| ENSMUSG00000027286 | Lrrc57 | chr2 | - | 120604237 | 120609520 | regular |
| ENSMUSG00000027288 | Zfp106 | chr2 | - | 120506819 | 120563843 | regular |
| ENSMUSG00000027304 | Rtf1 | chr2 | + | 119675067 | 119735407 | regular |
| ENSMUSG00000027309 | 4930402H | chr2 | - | 130706199 | 130906406 | regular |
| ENSMUSG00000027326 | Knl1 | chr2 | + | 119047118 | 119105501 | regular |
| ENSMUSG00000027331 | Knstrn | chr2 | + | 118814002 | 118853957 | regular |
| ENSMUSG00000027344 | Fsip1 | chr2 | - | 118204887 | 118256966 | regular |
| ENSMUSG00000027346 | Gpcpd1 | chr2 | - | 132529081 | 132587729 | regular |
| ENSMUSG00000027355 | Tmco5 | chr2 | + | 116878690 | 116892494 | regular |
| ENSMUSG00000027363 | Usp8 | chr2 | + | 126707327 | 126759297 | regular |
| ENSMUSG00000027365 | Trpm7 | chr2 | - | 126791564 | 126876230 | regular |
| ENSMUSG00000027378 | Nphp1 | chr2 | - | 127740731 | 127788897 | regular |
| ENSMUSG00000027422 | Rrbp1 | chr2 | - | 143947394 | 144011263 | regular |
| ENSMUSG00000027423 | Snx5 | chr2 | - | 144250122 | 144270906 | regular |
| ENSMUSG00000027433 | Xrn2 | chr2 | + | 147012995 | 147078000 | regular |
| ENSMUSG00000027439 | Gzf1 | chr2 | + | 148681022 | 148692949 | regular |
| ENSMUSG00000027465 | Tbc1d20 | chr2 | + | 152293827 | 152313996 | regular |
| ENSMUSG00000027475 | Kif3b | chr2 | + | 153291412 | 153333390 | regular |
| ENSMUSG00000027479 | Mapre1 | chr2 | + | 153741273 | 153773310 | regular |
| ENSMUSG00000027506 | Tpd52 | chr3 | - | 8925592 | 9004723 | regular |
| ENSMUSG00000027517 | Ankrd60 | chr2 | - | 173568665 | 173578365 | regular |
| ENSMUSG00000027519 | Rab22a | chr2 | + | 173659759 | 173707343 | regular |
| ENSMUSG00000027520 | Zdbf2 | chr1 | + | 63273264 | 63314576 | regular |
| ENSMUSG00000027523 | Gnas | chr2 | + | 174284319 | 174346744 | regular |
| ENSMUSG00000027528 | Fabp9 | chr3 | - | 10193620 | 10197283 | regular |
| ENSMUSG00000027546 | Atp9a | chr2 | - | 168634437 | 168742409 | regular |
| ENSMUSG00000027550 | Lrrcc1 | chr3 | + | 14533787 | 14572658 | regular |
| ENSMUSG00000027562 | Car2 | chr3 | + | 14886272 | 14900770 | regular |
| ENSMUSG00000027564 | Cypt12 | chr3 | + | 17948409 | 17948959 | regular |
| ENSMUSG00000027602 | Map1lc3a | chr2 | + | 155276296 | 155278073 | regular |
| ENSMUSG00000027620 | Rbm39 | chr2 | - | 156147238 | 156180238 | regular |
| ENSMUSG00000027630 | Tbl1xr1 | chr3 | + | 22076651 | 22216594 | regular |
| ENSMUSG00000027665 | Pik3ca | chr3 | + | 32397670 | 32468486 | regular |
| ENSMUSG00000027676 | Ccdc39 | chr3 | - | 33812361 | 33844310 | regular |
| ENSMUSG00000027678 | Ncoa3 | chr2 | + | 165992635 | 166073242 | regular |
| ENSMUSG00000027680 | Fxr1 | chr3 | + | 34019942 | 34070322 | regular |
| ENSMUSG00000027708 | Dcun1d1 | chr3 | - | 35892104 | 35937445 | regular |

| | | | | | | |
|---|---|---|---|---|---|---|
| ENSMUSG00000027722 | Spata5 | chr3 | + | 37419895 | 37579096 | regular |
| ENSMUSG00000027751 | Supt20 | chr3 | + | 54692806 | 54728766 | regular |
| ENSMUSG00000027763 | Mbnl1 | chr3 | + | 60472829 | 60629750 | regular |
| ENSMUSG00000027778 | Ift80 | chr3 | - | 68892498 | 69004570 | regular |
| ENSMUSG00000027782 | Kpna4 | chr3 | - | 69067148 | 69127113 | regular |
| ENSMUSG00000027799 | Nbea | chr3 | - | 55625194 | 56183701 | regular |
| ENSMUSG00000027808 | Serp1 | chr3 | - | 58519816 | 58525892 | regular |
| ENSMUSG00000027823 | Gmps | chr3 | + | 63976105 | 64022579 | regular |
| ENSMUSG00000027829 | Ccnl1 | chr3 | - | 65946150 | 65958249 | regular |
| ENSMUSG00000027867 | Spag17 | chr3 | + | 99885405 | 100143322 | regular |
| ENSMUSG00000027878 | Notch2 | chr3 | + | 98013526 | 98150361 | regular |
| ENSMUSG00000027905 | Ddx20 | chr3 | - | 105678269 | 105687574 | regular |
| ENSMUSG00000027927 | Lelp1 | chr3 | - | 92134996 | 92142744 | regular |
| ENSMUSG00000027939 | Nup210l | chr3 | + | 90104131 | 90212048 | regular |
| ENSMUSG00000027940 | Tpm3 | chr3 | + | 90072648 | 90100902 | regular |
| ENSMUSG00000027942 | 4933434E | chr3 | + | 90051635 | 90068347 | regular |
| ENSMUSG00000027968 | Larp7 | chr3 | - | 127536713 | 127553349 | regular |
| ENSMUSG00000028013 | Ppa2 | chr3 | + | 133310109 | 133378235 | regular |
| ENSMUSG00000028018 | Gstcd | chr3 | - | 132981751 | 133092033 | regular |
| ENSMUSG00000028034 | Fubp1 | chr3 | + | 152210421 | 152236826 | regular |
| ENSMUSG00000028035 | Dnajb4 | chr3 | - | 152178510 | 152210302 | regular |
| ENSMUSG00000028053 | Ash1l | chr3 | + | 88950621 | 89079375 | regular |
| ENSMUSG00000028059 | Arhgef2 | chr3 | + | 88605965 | 88648052 | regular |
| ENSMUSG00000028063 | Lmna | chr3 | - | 88480146 | 88509956 | regular |
| ENSMUSG00000028080 | Lrba | chr3 | + | 86224679 | 86782692 | regular |
| ENSMUSG00000028081 | Rps3a1 | chr3 | - | 86137939 | 86142702 | regular |
| ENSMUSG00000028086 | Fbxw7 | chr3 | + | 84815267 | 84979198 | regular |
| ENSMUSG00000028106 | Rprd2 | chr3 | - | 95760340 | 95818863 | regular |
| ENSMUSG00000028121 | Bcar3 | chr3 | + | 122294135 | 122530191 | regular |
| ENSMUSG00000028132 | Tmem56 | chr3 | - | 121201760 | 121283098 | regular |
| ENSMUSG00000028136 | Snx27 | chr3 | - | 94497543 | 94582716 | regular |
| ENSMUSG00000028137 | Celf3 | chr3 | + | 94478294 | 94492198 | regular |
| ENSMUSG00000028138 | Adh5 | chr3 | + | 138443092 | 138455499 | regular |
| ENSMUSG00000028141 | Oaz3 | chr3 | - | 94432413 | 94443915 | regular |
| ENSMUSG00000028156 | Eif4e | chr3 | + | 138526178 | 138559696 | regular |
| ENSMUSG00000028158 | Mttp | chr3 | - | 138089853 | 138144968 | regular |
| ENSMUSG00000028161 | Ppp3ca | chr3 | + | 136670123 | 136937727 | regular |
| ENSMUSG00000028163 | Nfkb1 | chr3 | - | 135584654 | 135691547 | regular |
| ENSMUSG00000028188 | Spata1 | chr3 | - | 146457195 | 146499753 | regular |
| ENSMUSG00000028207 | Asph | chr4 | - | 9448068 | 9669344 | regular |
| ENSMUSG00000028234 | Rps20 | chr4 | - | 3831333 | 3835665 | regular |
| ENSMUSG00000028248 | Pnisr | chr4 | + | 21847582 | 21876475 | regular |
| ENSMUSG00000028256 | Odf2l | chr3 | + | 145118587 | 145153915 | regular |
| ENSMUSG00000028273 | Pdlim5 | chr3 | - | 142239589 | 142395696 | regular |
| ENSMUSG00000028277 | Ube2j1 | chr4 | + | 33031415 | 33052363 | regular |
| ENSMUSG00000028287 | 1700009N | chr4 | + | 39450294 | 39451776 | regular |

| | | | | | | |
|---|---|---|---|---|---|---|
| ENSMUSG00000028294 | Cfap206 | chr4 | - | 34688558 | 34730206 | regular |
| ENSMUSG00000028310 | Ppp3r2 | chr4 | - | 49678746 | 49682024 | regular |
| ENSMUSG00000028330 | Ncbp1 | chr4 | + | 46138612 | 46172403 | regular |
| ENSMUSG00000028337 | Coro2a | chr4 | - | 46536936 | 46602202 | regular |
| ENSMUSG00000028347 | Tmeff1 | chr4 | + | 48585173 | 48663131 | regular |
| ENSMUSG00000028354 | Fmn2 | chr1 | + | 174501824 | 174822729 | regular |
| ENSMUSG00000028382 | Ptbp3 | chr4 | - | 59471867 | 59549364 | regular |
| ENSMUSG00000028383 | Hsdl2 | chr4 | + | 59581562 | 59618689 | regular |
| ENSMUSG00000028392 | Bspry | chr4 | + | 62480052 | 62497298 | regular |
| ENSMUSG00000028410 | Dnaja1 | chr4 | + | 40722149 | 40737149 | regular |
| ENSMUSG00000028411 | Aptx | chr4 | - | 40682381 | 40703194 | regular |
| ENSMUSG00000028414 | Fktn | chr4 | + | 53713997 | 53777890 | regular |
| ENSMUSG00000028416 | Bag1 | chr4 | - | 40936397 | 40948294 | regular |
| ENSMUSG00000028423 | Nfx1 | chr4 | + | 40970905 | 41025993 | regular |
| ENSMUSG00000028426 | Rad23b | chr4 | + | 55350042 | 55392237 | regular |
| ENSMUSG00000028433 | Ubap2 | chr4 | - | 41194312 | 41275144 | regular |
| ENSMUSG00000028441 | 1110017D | chr4 | - | 41505008 | 41517333 | regular |
| ENSMUSG00000028452 | Vcp | chr4 | - | 42979962 | 43000507 | regular |
| ENSMUSG00000028456 | Unc13b | chr4 | + | 43058952 | 43264871 | regular |
| ENSMUSG00000028457 | Atp8b5 | chr4 | + | 43267158 | 43373833 | regular |
| ENSMUSG00000028458 | Tesk1 | chr4 | + | 43441938 | 43448064 | regular |
| ENSMUSG00000028465 | Tln1 | chr4 | - | 43531518 | 43562691 | regular |
| ENSMUSG00000028484 | Psip1 | chr4 | - | 83455679 | 83486459 | regular |
| ENSMUSG00000028487 | Bnc2 | chr4 | - | 84275094 | 84675275 | regular |
| ENSMUSG00000028495 | Rps6 | chr4 | - | 86854659 | 86857412 | regular |
| ENSMUSG00000028514 | Usp24 | chr4 | + | 106316212 | 106441322 | regular |
| ENSMUSG00000028520 | 4921539E | chr4 | - | 103230444 | 103290863 | regular |
| ENSMUSG00000028525 | Pde4b | chr4 | + | 102087542 | 102607259 | regular |
| ENSMUSG00000028530 | Jak1 | chr4 | - | 101152366 | 101265282 | regular |
| ENSMUSG00000028541 | B4galt2 | chr4 | - | 117869259 | 117883487 | regular |
| ENSMUSG00000028551 | Cdkn2c | chr4 | - | 109660875 | 109667189 | regular |
| ENSMUSG00000028552 | Eps15 | chr4 | + | 109280267 | 109387817 | regular |
| ENSMUSG00000028555 | Ttc39a | chr4 | + | 109406622 | 109444745 | regular |
| ENSMUSG00000028557 | Rnf11 | chr4 | - | 109451097 | 109484728 | regular |
| ENSMUSG00000028559 | Osbpl9 | chr4 | - | 109061144 | 109202272 | regular |
| ENSMUSG00000028560 | Usp1 | chr4 | + | 98923809 | 98935543 | regular |
| ENSMUSG00000028572 | Hook1 | chr4 | + | 95967239 | 96025413 | regular |
| ENSMUSG00000028576 | Ift74 | chr4 | + | 94614490 | 94693229 | regular |
| ENSMUSG00000028580 | Pum1 | chr4 | + | 130663320 | 130781564 | regular |
| ENSMUSG00000028582 | Cc2d1b | chr4 | + | 108619939 | 108634123 | regular |
| ENSMUSG00000028610 | Dmrtb1 | chr4 | - | 107676289 | 107684230 | regular |
| ENSMUSG00000028637 | Ccdc30 | chr4 | - | 119322892 | 119415521 | regular |
| ENSMUSG00000028639 | Ybx1 | chr4 | - | 119277980 | 119294604 | regular |
| ENSMUSG00000028649 | Macf1 | chr4 | - | 123349632 | 123684360 | regular |
| ENSMUSG00000028673 | Fuca1 | chr4 | + | 135920734 | 135940311 | regular |
| ENSMUSG00000028677 | Rnf220 | chr4 | - | 117271462 | 117497052 | regular |

| | | | | | | |
|---|---|---|---|---|---|---|
| ENSMUSG00000028678 | Kif2c | chr4 | - | 117159638 | 117182639 | regular |
| ENSMUSG00000028691 | Prdx1 | chr4 | + | 116685543 | 116700822 | regular |
| ENSMUSG00000028692 | Akr1a1 | chr4 | - | 116636509 | 116651680 | regular |
| ENSMUSG00000028693 | Nasp | chr4 | - | 116601051 | 116627941 | regular |
| ENSMUSG00000028706 | Nsun4 | chr4 | - | 116032841 | 116053876 | regular |
| ENSMUSG00000028730 | Cfap57 | chr4 | - | 118554550 | 118620777 | regular |
| ENSMUSG00000028745 | Capzb | chr4 | + | 139192898 | 139291818 | regular |
| ENSMUSG00000028750 | Pla2g2c | chr4 | + | 138724791 | 138746132 | regular |
| ENSMUSG00000028756 | Pink1 | chr4 | - | 138313408 | 138326307 | regular |
| ENSMUSG00000028758 | Kif17 | chr4 | + | 138250434 | 138301967 | regular |
| ENSMUSG00000028759 | Hp1bp3 | chr4 | + | 138216295 | 138244683 | regular |
| ENSMUSG00000028760 | Eif4g3 | chr4 | + | 137993021 | 138208508 | regular |
| ENSMUSG00000028772 | Zcchc17 | chr4 | - | 130315382 | 130359943 | regular |
| ENSMUSG00000028788 | Ptp4a2 | chr4 | + | 129811218 | 129850003 | regular |
| ENSMUSG00000028789 | Azin2 | chr4 | - | 128930232 | 128962442 | regular |
| ENSMUSG00000028790 | Khdrbs1 | chr4 | - | 129703163 | 129742303 | regular |
| ENSMUSG00000028793 | Rnf19b | chr4 | + | 129058270 | 129085886 | regular |
| ENSMUSG00000028800 | Hdac1 | chr4 | - | 129516103 | 129542713 | regular |
| ENSMUSG00000028809 | Srrm1 | chr4 | - | 135320483 | 135353321 | regular |
| ENSMUSG00000028813 | CK137956 | chr4 | - | 127927591 | 127970951 | regular |
| ENSMUSG00000028820 | Sfpq | chr4 | + | 127021323 | 127037013 | regular |
| ENSMUSG00000028843 | Sh3bgrl3 | chr4 | - | 134127405 | 134128789 | regular |
| ENSMUSG00000028845 | Tekt2 | chr4 | - | 126322120 | 126325688 | regular |
| ENSMUSG00000028868 | Wasf2 | chr4 | + | 133130504 | 133199756 | regular |
| ENSMUSG00000028878 | Fam76a | chr4 | - | 132899212 | 132922558 | regular |
| ENSMUSG00000028886 | Eya3 | chr4 | + | 132638986 | 132724765 | regular |
| ENSMUSG00000028896 | Rcc1 | chr4 | - | 132331918 | 132353605 | regular |
| ENSMUSG00000028926 | Cdk14 | chr5 | - | 4803390 | 5420312 | regular |
| ENSMUSG00000028932 | Psmc2 | chr5 | + | 21785282 | 21803787 | regular |
| ENSMUSG00000028936 | Rpl22 | chr4 | + | 152325741 | 152334071 | regular |
| ENSMUSG00000028937 | Acot7 | chr4 | + | 152178133 | 152271855 | regular |
| ENSMUSG00000028938 | Galntl5 | chr5 | + | 25181459 | 25220297 | regular |
| ENSMUSG00000028944 | Prkag2 | chr5 | - | 24862743 | 25100642 | regular |
| ENSMUSG00000028949 | Smarcd3 | chr5 | - | 24590817 | 24624411 | regular |
| ENSMUSG00000028952 | Zbtb48 | chr4 | - | 152019773 | 152027671 | regular |
| ENSMUSG00000028953 | Abcf2 | chr5 | - | 24565344 | 24577467 | regular |
| ENSMUSG00000028976 | Slc2a5 | chr4 | + | 150119282 | 150144169 | regular |
| ENSMUSG00000028991 | Mtor | chr4 | + | 148448610 | 148557683 | regular |
| ENSMUSG00000028995 | Fam126a | chr5 | - | 23915275 | 24030690 | regular |
| ENSMUSG00000029001 | Fbxo44 | chr4 | - | 148152799 | 148160482 | regular |
| ENSMUSG00000029004 | Kmt2e | chr5 | + | 23434440 | 23504235 | regular |
| ENSMUSG00000029014 | Dnajc2 | chr5 | - | 21757266 | 21785251 | regular |
| ENSMUSG00000029020 | Mfn2 | chr4 | - | 147873598 | 147904704 | regular |
| ENSMUSG00000029030 | Tprgl | chr4 | - | 154157484 | 154160666 | regular |
| ENSMUSG00000029064 | Gnb1 | chr4 | + | 155491360 | 155559269 | regular |
| ENSMUSG00000029068 | Ccnl2 | chr4 | + | 155812488 | 155824543 | regular |

| | | | | | | |
|---|---|---|---|---|---|---|
| ENSMUSG00000029073 | Cptp | chr4 | - | 155864722 | 155869440 | regular |
| ENSMUSG00000029074 | Ttll10 | chr4 | - | 156034839 | 156059414 | regular |
| ENSMUSG00000029076 | Sdf4 | chr4 | + | 155992871 | 156013610 | regular |
| ENSMUSG00000029086 | Prom1 | chr5 | - | 43993619 | 44102032 | regular |
| ENSMUSG00000029098 | Acox3 | chr5 | + | 35583039 | 35615352 | regular |
| ENSMUSG00000029104 | Htt | chr5 | + | 34761739 | 34912534 | regular |
| ENSMUSG00000029111 | Nelfa | chr5 | - | 33897915 | 33936413 | regular |
| ENSMUSG00000029128 | Rab28 | chr5 | - | 41624975 | 41708157 | regular |
| ENSMUSG00000029130 | Rnf32 | chr5 | + | 29195991 | 29228457 | regular |
| ENSMUSG00000029131 | Dnajb6 | chr5 | + | 29735687 | 29818134 | regular |
| ENSMUSG00000029134 | Plb1 | chr5 | + | 32232707 | 32366520 | regular |
| ENSMUSG00000029135 | Fosl2 | chr5 | + | 32135800 | 32157842 | regular |
| ENSMUSG00000029147 | Ppm1g | chr5 | - | 31202663 | 31220687 | regular |
| ENSMUSG00000029148 | Nrbp1 | chr5 | + | 31240863 | 31251566 | regular |
| ENSMUSG00000029155 | Spata18 | chr5 | + | 73651378 | 73679512 | regular |
| ENSMUSG00000029165 | Agbl5 | chr5 | + | 30888693 | 30906965 | regular |
| ENSMUSG00000029174 | Tbc1d1 | chr5 | + | 64156304 | 64351486 | regular |
| ENSMUSG00000029191 | Rfc1 | chr5 | - | 65261849 | 65335670 | regular |
| ENSMUSG00000029192 | Tbc1d14 | chr5 | - | 36490603 | 36593276 | regular |
| ENSMUSG00000029202 | Pds5a | chr5 | - | 65605720 | 65698273 | regular |
| ENSMUSG00000029203 | Ube2k | chr5 | + | 65537232 | 65598988 | regular |
| ENSMUSG00000029204 | Rhoh | chr5 | + | 65861212 | 65896700 | regular |
| ENSMUSG00000029206 | Nsun7 | chr5 | + | 66259896 | 66298026 | regular |
| ENSMUSG00000029235 | Pdcl2 | chr5 | - | 76312114 | 76331156 | regular |
| ENSMUSG00000029279 | Brdt | chr5 | + | 107331158 | 107387058 | regular |
| ENSMUSG00000029291 | Rufy3 | chr5 | + | 88565039 | 88651392 | regular |
| ENSMUSG00000029313 | Aff1 | chr5 | + | 103692373 | 103855322 | regular |
| ENSMUSG00000029320 | 1700016H | chr5 | - | 103648038 | 103656010 | regular |
| ENSMUSG00000029330 | Cds1 | chr5 | + | 101765129 | 101823858 | regular |
| ENSMUSG00000029338 | Antxr2 | chr5 | - | 97882782 | 98031043 | regular |
| ENSMUSG00000029381 | Shroom3 | chr5 | + | 92683434 | 92965318 | regular |
| ENSMUSG00000029397 | Rchy1 | chr5 | - | 91948903 | 91963068 | regular |
| ENSMUSG00000029403 | Cdkl2 | chr5 | - | 92006073 | 92043883 | regular |
| ENSMUSG00000029405 | G3bp2 | chr5 | - | 92052145 | 92083719 | regular |
| ENSMUSG00000029406 | Pitpnm2 | chr5 | - | 124118689 | 124249760 | regular |
| ENSMUSG00000029422 | Rsrc2 | chr5 | - | 123728425 | 123749414 | regular |
| ENSMUSG00000029423 | Piwil1 | chr5 | + | 128702523 | 128755474 | regular |
| ENSMUSG00000029427 | Zcchc8 | chr5 | - | 123698293 | 123721100 | regular |
| ENSMUSG00000029428 | Stx2 | chr5 | - | 128984556 | 129008574 | regular |
| ENSMUSG00000029433 | Diablo | chr5 | - | 123509764 | 123525429 | regular |
| ENSMUSG00000029439 | Sfswap | chr5 | + | 129501220 | 129571384 | regular |
| ENSMUSG00000029442 | Wdr66 | chr5 | + | 123252101 | 123327484 | regular |
| ENSMUSG00000029458 | Brap | chr5 | + | 121660562 | 121687256 | regular |
| ENSMUSG00000029463 | Fam216a | chr5 | - | 122364579 | 122372364 | regular |
| ENSMUSG00000029467 | Atp2a2 | chr5 | - | 122453512 | 122502225 | regular |
| ENSMUSG00000029469 | Ift81 | chr5 | - | 122550203 | 122614518 | regular |

| | | | | | | |
|---|---|---|---|---|---|---|
| ENSMUSG00000029472 | Anapc5 | chr5 | - | 122787458 | 122821339 | regular |
| ENSMUSG00000029475 | Kdm2b | chr5 | - | 122870664 | 122989823 | regular |
| ENSMUSG00000029477 | Morn3 | chr5 | - | 123035768 | 123047016 | regular |
| ENSMUSG00000029478 | Ncor2 | chr5 | - | 125017152 | 125179219 | regular |
| ENSMUSG00000029501 | Ankle2 | chr5 | + | 110231003 | 110256651 | regular |
| ENSMUSG00000029505 | Ep400 | chr5 | - | 110664372 | 110770717 | regular |
| ENSMUSG00000029516 | Cit | chr5 | + | 115845277 | 116008947 | regular |
| ENSMUSG00000029534 | St7 | chr6 | + | 17692932 | 17943025 | regular |
| ENSMUSG00000029577 | Ube3b | chr5 | + | 114380606 | 114421169 | regular |
| ENSMUSG00000029580 | Actb | chr5 | - | 142903114 | 142906754 | regular |
| ENSMUSG00000029614 | Rpl6 | chr5 | + | 121204480 | 121209241 | regular |
| ENSMUSG00000029621 | Arpc1a | chr5 | + | 145083829 | 145108761 | regular |
| ENSMUSG00000029629 | Phf14 | chr6 | + | 11907808 | 12081205 | regular |
| ENSMUSG00000029640 | Usp12 | chr5 | - | 146734808 | 146795006 | regular |
| ENSMUSG00000029647 | Pan3 | chr5 | + | 147430160 | 147548502 | regular |
| ENSMUSG00000029686 | Cul1 | chr6 | + | 47453397 | 47526139 | regular |
| ENSMUSG00000029703 | Lrwd1 | chr5 | - | 136122771 | 136136074 | regular |
| ENSMUSG00000029705 | Cux1 | chr5 | - | 136248134 | 136567490 | regular |
| ENSMUSG00000029707 | Fscn3 | chr6 | + | 28427788 | 28438622 | regular |
| ENSMUSG00000029726 | Mepce | chr5 | - | 137781905 | 137787655 | regular |
| ENSMUSG00000029766 | 1700012A | chr6 | + | 32050245 | 32058921 | regular |
| ENSMUSG00000029769 | Ccdc136 | chr6 | + | 29396308 | 29426994 | regular |
| ENSMUSG00000029772 | Ahcyl2 | chr6 | + | 29768010 | 29912310 | regular |
| ENSMUSG00000029788 | Cpa5 | chr6 | + | 30611009 | 30631745 | regular |
| ENSMUSG00000029814 | Igf2bp3 | chr6 | - | 49085222 | 49214957 | regular |
| ENSMUSG00000029817 | Tra2a | chr6 | - | 49243923 | 49264033 | regular |
| ENSMUSG00000029823 | Luc7l2 | chr6 | + | 38551333 | 38609470 | regular |
| ENSMUSG00000029826 | Zc3hav1 | chr6 | - | 38305285 | 38354603 | regular |
| ENSMUSG00000029828 | 4921507P | chr6 | - | 50573301 | 50596632 | regular |
| ENSMUSG00000029833 | Trim24 | chr6 | + | 37870810 | 37966296 | regular |
| ENSMUSG00000029867 | Llcfc1 | chr6 | + | 41684430 | 41685717 | regular |
| ENSMUSG00000029922 | Mkrn1 | chr6 | - | 39397803 | 39420462 | regular |
| ENSMUSG00000029992 | Gfpt1 | chr6 | + | 87042845 | 87092197 | regular |
| ENSMUSG00000030007 | Cct7 | chr6 | + | 85451513 | 85468475 | regular |
| ENSMUSG00000030016 | Zfp638 | chr6 | + | 83867108 | 83989550 | regular |
| ENSMUSG00000030030 | 1700003E | chr6 | + | 83156403 | 83162945 | regular |
| ENSMUSG00000030041 | M1ap | chr6 | + | 82946901 | 83030309 | regular |
| ENSMUSG00000030057 | Cnbp | chr6 | - | 87842614 | 87851106 | regular |
| ENSMUSG00000030059 | Tmf1 | chr6 | - | 97152996 | 97179122 | regular |
| ENSMUSG00000030067 | Foxp1 | chr6 | - | 98925337 | 99522721 | regular |
| ENSMUSG00000030068 | Gm20696 | chr6 | - | 99257483 | 99666797 | regular |
| ENSMUSG00000030077 | Chl1 | chr6 | + | 103510585 | 103750211 | regular |
| ENSMUSG00000030082 | Sec61a1 | chr6 | - | 88503578 | 88518905 | regular |
| ENSMUSG00000030086 | Chchd6 | chr6 | - | 89383145 | 89595652 | regular |
| ENSMUSG00000030096 | Slc6a6 | chr6 | + | 91684052 | 91759066 | regular |
| ENSMUSG00000030102 | Itpr1 | chr6 | + | 108213095 | 108551109 | regular |

| | | | | | | |
|---|---|---|---|---|---|---|
| ENSMUSG00000030126 | Tmcc1 | chr6 | - | 116018610 | 116193486 | regular |
| ENSMUSG00000030138 | Bms1 | chr6 | - | 118383380 | 118419474 | regular |
| ENSMUSG00000030161 | Gabarapl1 | chr6 | + | 129533159 | 129542346 | regular |
| ENSMUSG00000030172 | Erc1 | chr6 | - | 119570795 | 119848167 | regular |
| ENSMUSG00000030180 | Kdm5a | chr6 | + | 120364123 | 120444574 | regular |
| ENSMUSG00000030189 | Ybx3 | chr6 | - | 131364854 | 131388476 | regular |
| ENSMUSG00000030199 | Etv6 | chr6 | + | 134035699 | 134270158 | regular |
| ENSMUSG00000030200 | Bcl2l14 | chr6 | + | 134396317 | 134438736 | regular |
| ENSMUSG00000030203 | Dusp16 | chr6 | - | 134715467 | 134792625 | regular |
| ENSMUSG00000030206 | Gsg1 | chr6 | - | 135237325 | 135254351 | regular |
| ENSMUSG00000030213 | Atf7ip | chr6 | + | 136506166 | 136610862 | regular |
| ENSMUSG00000030216 | Wbp11 | chr6 | - | 136813653 | 136828233 | regular |
| ENSMUSG00000030230 | Plcz1 | chr6 | - | 139989672 | 140041457 | regular |
| ENSMUSG00000030231 | Plekha5 | chr6 | + | 140424053 | 140597110 | regular |
| ENSMUSG00000030232 | Aebp2 | chr6 | + | 140622662 | 140678472 | regular |
| ENSMUSG00000030254 | Rad18 | chr6 | - | 112619849 | 112696686 | regular |
| ENSMUSG00000030265 | Kras | chr6 | - | 145216698 | 145250239 | regular |
| ENSMUSG00000030275 | Etnk1 | chr6 | + | 143167220 | 143232307 | regular |
| ENSMUSG00000030276 | Ttll3 | chr6 | + | 113389259 | 113414587 | regular |
| ENSMUSG00000030287 | Itpr2 | chr6 | - | 146108298 | 146502223 | regular |
| ENSMUSG00000030292 | Smco2 | chr6 | + | 146850103 | 146871406 | regular |
| ENSMUSG00000030301 | Ccdc91 | chr6 | + | 147475827 | 147632612 | regular |
| ENSMUSG00000030313 | Dennd5b | chr6 | - | 148988070 | 149101680 | regular |
| ENSMUSG00000030323 | Ift122 | chr6 | + | 115853469 | 115926699 | regular |
| ENSMUSG00000030344 | Akap3 | chr6 | + | 126853097 | 126874308 | regular |
| ENSMUSG00000030357 | Fkbp4 | chr6 | - | 128429734 | 128438677 | regular |
| ENSMUSG00000030374 | Strn4 | chr7 | + | 16815888 | 16840931 | regular |
| ENSMUSG00000030403 | Vasp | chr7 | - | 19256928 | 19271817 | regular |
| ENSMUSG00000030432 | Rpl28 | chr7 | + | 4792873 | 4795066 | regular |
| ENSMUSG00000030451 | Herc2 | chr7 | + | 56050195 | 56231800 | regular |
| ENSMUSG00000030465 | Psd3 | chr8 | - | 67689081 | 68212027 | regular |
| ENSMUSG00000030498 | Gas2 | chr7 | + | 51862014 | 51994975 | regular |
| ENSMUSG00000030533 | Unc45a | chr7 | - | 80325291 | 80341005 | regular |
| ENSMUSG00000030536 | Iqgap1 | chr7 | - | 80711582 | 80825974 | regular |
| ENSMUSG00000030538 | Cib1 | chr7 | - | 80227146 | 80232813 | regular |
| ENSMUSG00000030555 | Ttc23 | chr7 | + | 67647409 | 67762912 | regular |
| ENSMUSG00000030556 | Lrrc28 | chr7 | - | 67513409 | 67645268 | regular |
| ENSMUSG00000030557 | Mef2a | chr7 | - | 67231162 | 67372858 | regular |
| ENSMUSG00000030583 | Sipa1l3 | chr7 | - | 29320371 | 29518641 | regular |
| ENSMUSG00000030588 | Yif1b | chr7 | + | 29238328 | 29247597 | regular |
| ENSMUSG00000030591 | Psmd8 | chr7 | - | 29174187 | 29180701 | regular |
| ENSMUSG00000030595 | Nfkbib | chr7 | - | 28758250 | 28767512 | regular |
| ENSMUSG00000030638 | Sh3gl3 | chr7 | + | 82173839 | 82307419 | regular |
| ENSMUSG00000030655 | Smg1 | chr7 | - | 118131307 | 118243670 | regular |
| ENSMUSG00000030662 | Ipo5 | chr14 | + | 120911223 | 120947999 | regular |
| ENSMUSG00000030671 | Pde3b | chr7 | + | 114415280 | 114539251 | regular |

| | | | | | | |
|---|---|---|---|---|---|---|
| ENSMUSG00000030688 | Stard10 | chr7 | + | 101317085 | 101346626 | regular |
| ENSMUSG00000030695 | Aldoa | chr7 | - | 126795233 | 126800751 | regular |
| ENSMUSG00000030726 | Pold3 | chr7 | - | 100082110 | 100121565 | regular |
| ENSMUSG00000030738 | Eif3c | chr7 | - | 126546454 | 126566411 | regular |
| ENSMUSG00000030744 | Rps3 | chr7 | - | 99477895 | 99483738 | regular |
| ENSMUSG00000030748 | Il4ra | chr7 | + | 125552119 | 125579474 | regular |
| ENSMUSG00000030754 | Copb1 | chr7 | - | 114215558 | 114254711 | regular |
| ENSMUSG00000030761 | Myo7a | chr7 | - | 98051059 | 98119524 | regular |
| ENSMUSG00000030763 | Lcmt1 | chr7 | + | 123369783 | 123430358 | regular |
| ENSMUSG00000030779 | Rbbp6 | chr7 | + | 122965685 | 123002557 | regular |
| ENSMUSG00000030792 | Dkkl1 | chr7 | - | 45207518 | 45212147 | regular |
| ENSMUSG00000030795 | Fus | chr7 | + | 127967456 | 127985701 | regular |
| ENSMUSG00000030798 | Cd37 | chr7 | - | 45233631 | 45239115 | regular |
| ENSMUSG00000030815 | Phkg2 | chr7 | + | 127573339 | 127583307 | regular |
| ENSMUSG00000030816 | Rnf40 | chr7 | + | 127588766 | 127604799 | regular |
| ENSMUSG00000030846 | Tial1 | chr7 | - | 128439776 | 128461717 | regular |
| ENSMUSG00000030850 | Ate1 | chr7 | - | 130391492 | 130520369 | regular |
| ENSMUSG00000030851 | Ldhc | chr7 | + | 46861202 | 46878142 | regular |
| ENSMUSG00000030870 | Ubfd1 | chr7 | + | 122067174 | 122082194 | regular |
| ENSMUSG00000030872 | Gga2 | chr7 | - | 121986721 | 122021222 | regular |
| ENSMUSG00000030882 | Dnhd1 | chr7 | + | 105650826 | 105721799 | regular |
| ENSMUSG00000030889 | Vwa3a | chr7 | + | 120739317 | 120805742 | regular |
| ENSMUSG00000030967 | Zranb1 | chr7 | + | 132931141 | 132986391 | regular |
| ENSMUSG00000030980 | Knop1 | chr7 | - | 118842221 | 118856254 | regular |
| ENSMUSG00000030987 | Stim1 | chr7 | + | 102267805 | 102437319 | regular |
| ENSMUSG00000030990 | Pgap2 | chr7 | + | 102210207 | 102238567 | regular |
| ENSMUSG00000030994 | D7Ertd443 | chr7 | - | 134265778 | 134385661 | regular |
| ENSMUSG00000031010 | Usp9x | chrX | + | 13071497 | 13173328 | regular |
| ENSMUSG00000031015 | Swap70 | chr7 | + | 110221710 | 110283506 | regular |
| ENSMUSG00000031027 | Stk33 | chr7 | - | 109279222 | 109439081 | regular |
| ENSMUSG00000031029 | Eif3f | chr7 | + | 108934416 | 108942951 | regular |
| ENSMUSG00000031065 | Cdk16 | chrX | + | 20687953 | 20699880 | regular |
| ENSMUSG00000031068 | Glrx3 | chr7 | + | 137437613 | 137468594 | regular |
| ENSMUSG00000031085 | Gm498 | chr7 | + | 143866837 | 143900046 | regular |
| ENSMUSG00000031153 | Gripap1 | chrX | + | 7789764 | 7820567 | regular |
| ENSMUSG00000031207 | Msn | chrX | + | 96096041 | 96168552 | regular |
| ENSMUSG00000031233 | Pgk2 | chr17 | - | 40207017 | 40208609 | regular |
| ENSMUSG00000031278 | Acsl4 | chrX | - | 142317992 | 142390535 | regular |
| ENSMUSG00000031320 | Rps4x | chrX | - | 102184940 | 102189394 | regular |
| ENSMUSG00000031328 | Flna | chrX | - | 74223460 | 74249820 | regular |
| ENSMUSG00000031392 | Irak1 | chrX | - | 74013913 | 74023918 | regular |
| ENSMUSG00000031393 | Mecp2 | chrX | - | 74026591 | 74135363 | regular |
| ENSMUSG00000031422 | Morf4l2 | chrX | - | 136732941 | 136743690 | regular |
| ENSMUSG00000031430 | Vsig1 | chrX | + | 140907607 | 140939472 | regular |
| ENSMUSG00000031447 | Lamp1 | chr8 | + | 13159160 | 13175338 | regular |
| ENSMUSG00000031452 | 1700029H | chr8 | - | 13550732 | 13562461 | regular |

| | | | | | | |
|---|---|---|---|---|---|---|
| ENSMUSG00000031453 | Rasa3 | chr8 | - | 13566947 | 13677603 | regular |
| ENSMUSG00000031485 | Plpbp | chr8 | + | 27042554 | 27056132 | regular |
| ENSMUSG00000031493 | Ggn | chr7 | + | 29170209 | 29173976 | regular |
| ENSMUSG00000031518 | Spata4 | chr8 | + | 54600780 | 54610098 | regular |
| ENSMUSG00000031529 | Tnks | chr8 | - | 34826459 | 34965690 | regular |
| ENSMUSG00000031536 | Polb | chr8 | - | 22628125 | 22653435 | regular |
| ENSMUSG00000031537 | Ikbkb | chr8 | - | 22659211 | 22706589 | regular |
| ENSMUSG00000031540 | Kat6a | chr8 | + | 22859534 | 22943259 | regular |
| ENSMUSG00000031545 | Gpat4 | chr8 | - | 23171264 | 23208346 | regular |
| ENSMUSG00000031552 | Adam18 | chr8 | - | 24602245 | 24674755 | regular |
| ENSMUSG00000031553 | Adam3 | chr8 | - | 24677224 | 24725852 | regular |
| ENSMUSG00000031554 | Adam5 | chr8 | - | 24727092 | 24824369 | regular |
| ENSMUSG00000031563 | Wwc2 | chr8 | - | 47823958 | 47990924 | regular |
| ENSMUSG00000031576 | Kcnu1 | chr8 | + | 25849622 | 25937939 | regular |
| ENSMUSG00000031583 | Wrn | chr8 | - | 33234383 | 33385527 | regular |
| ENSMUSG00000031585 | Gtf2e2 | chr8 | + | 33731832 | 33777173 | regular |
| ENSMUSG00000031592 | Pcm1 | chr8 | + | 41239751 | 41332344 | regular |
| ENSMUSG00000031617 | Tmem184 | chr8 | - | 77595981 | 77610698 | regular |
| ENSMUSG00000031620 | Iqcm | chr8 | + | 75448693 | 75984503 | regular |
| ENSMUSG00000031622 | Sin3b | chr8 | + | 72723284 | 72758201 | regular |
| ENSMUSG00000031631 | Cfap97 | chr8 | + | 46151770 | 46195590 | regular |
| ENSMUSG00000031644 | Nek1 | chr8 | + | 60993194 | 61131346 | regular |
| ENSMUSG00000031647 | Mfap3l | chr8 | + | 60632826 | 60676729 | regular |
| ENSMUSG00000031667 | Aktip | chr8 | - | 91111783 | 91199976 | regular |
| ENSMUSG00000031672 | Got2 | chr8 | - | 95864133 | 95888547 | regular |
| ENSMUSG00000031684 | Slc10a7 | chr8 | + | 78509345 | 78734003 | regular |
| ENSMUSG00000031701 | Dnaja2 | chr8 | - | 85537632 | 85555344 | regular |
| ENSMUSG00000031706 | Rfx1 | chr8 | + | 84066833 | 84096992 | regular |
| ENSMUSG00000031727 | Pmfbp1 | chr8 | + | 109494026 | 109542640 | regular |
| ENSMUSG00000031728 | Zfp821 | chr8 | + | 109705545 | 109724932 | regular |
| ENSMUSG00000031731 | Ap1g1 | chr8 | + | 109778553 | 109864204 | regular |
| ENSMUSG00000031751 | Amfr | chr8 | - | 93971587 | 94012842 | regular |
| ENSMUSG00000031765 | Mt1 | chr8 | + | 94179081 | 94180327 | regular |
| ENSMUSG00000031770 | Herpud1 | chr8 | + | 94386437 | 94395377 | regular |
| ENSMUSG00000031774 | Fam192a | chr8 | - | 94574137 | 94601997 | regular |
| ENSMUSG00000031776 | Arl2bp | chr8 | + | 94666599 | 94674425 | regular |
| ENSMUSG00000031786 | Drc7 | chr8 | + | 95055102 | 95078141 | regular |
| ENSMUSG00000031787 | Katnb1 | chr8 | + | 95081185 | 95103149 | regular |
| ENSMUSG00000031788 | Kifc3 | chr8 | - | 95099827 | 95202812 | regular |
| ENSMUSG00000031818 | Cox4i1 | chr8 | + | 120668221 | 120674207 | regular |
| ENSMUSG00000031822 | Gse1 | chr8 | + | 120228455 | 120581390 | regular |
| ENSMUSG00000031831 | Dnaaf1 | chr8 | + | 119575234 | 119598454 | regular |
| ENSMUSG00000031839 | Hsbp1 | chr8 | + | 119344487 | 119348922 | regular |
| ENSMUSG00000031847 | 1700030J: | chr8 | - | 116969593 | 116978959 | regular |
| ENSMUSG00000031849 | Comp | chr8 | + | 70373557 | 70382066 | regular |
| ENSMUSG00000031858 | Mau2 | chr8 | - | 70016122 | 70042734 | regular |

| | | | | | | |
|---|---|---|---|---|---|---|
| ENSMUSG00000031865 | Dctn1 | chr6 | + | 83165919 | 83200117 | regular |
| ENSMUSG00000031885 | Cbfb | chr8 | + | 105170673 | 105217989 | regular |
| ENSMUSG00000031893 | Tsnaxip1 | chr8 | + | 105827730 | 105844680 | regular |
| ENSMUSG00000031902 | Nfatc3 | chr8 | + | 106058839 | 106130537 | regular |
| ENSMUSG00000031924 | Cyb5b | chr8 | + | 107150639 | 107187471 | regular |
| ENSMUSG00000031930 | Wwp2 | chr8 | + | 107436364 | 107558595 | regular |
| ENSMUSG00000031949 | Adat1 | chr8 | - | 111966907 | 111992302 | regular |
| ENSMUSG00000031958 | Ldhd | chr8 | - | 111623784 | 111630374 | regular |
| ENSMUSG00000031960 | Aars | chr8 | + | 111033143 | 111057664 | regular |
| ENSMUSG00000031985 | Gnpat | chr8 | + | 124863032 | 124890057 | regular |
| ENSMUSG00000031986 | Sprtn | chr8 | + | 124897885 | 124906161 | regular |
| ENSMUSG00000032009 | Sesn3 | chr9 | + | 14275066 | 14333101 | regular |
| ENSMUSG00000032010 | Usp2 | chr9 | + | 44067020 | 44095627 | regular |
| ENSMUSG00000032050 | Rdx | chr9 | + | 52047172 | 52088735 | regular |
| ENSMUSG00000032053 | Pou2af1 | chr9 | + | 51213707 | 51240080 | regular |
| ENSMUSG00000032076 | Cadm1 | chr9 | + | 47530172 | 47857637 | regular |
| ENSMUSG00000032096 | Arcn1 | chr9 | - | 44741563 | 44767845 | regular |
| ENSMUSG00000032097 | Ddx6 | chr9 | + | 44604891 | 44640731 | regular |
| ENSMUSG00000032101 | Ddx25 | chr9 | - | 35536646 | 35558528 | regular |
| ENSMUSG00000032115 | Hyou1 | chr9 | + | 44379489 | 44392369 | regular |
| ENSMUSG00000032178 | Ilf3 | chr9 | + | 21367870 | 21405361 | regular |
| ENSMUSG00000032185 | Carm1 | chr9 | + | 21546893 | 21592623 | regular |
| ENSMUSG00000032187 | Smarca4 | chr9 | + | 21616168 | 21704230 | regular |
| ENSMUSG00000032212 | Sltm | chr9 | + | 70542753 | 70592234 | regular |
| ENSMUSG00000032216 | Nedd4 | chr9 | + | 72662345 | 72749852 | regular |
| ENSMUSG00000032217 | Rnf111 | chr9 | - | 70425423 | 70503725 | regular |
| ENSMUSG00000032220 | Myo1e | chr9 | + | 70207349 | 70399766 | regular |
| ENSMUSG00000032221 | Mns1 | chr9 | + | 72438010 | 72462025 | regular |
| ENSMUSG00000032228 | Tcf12 | chr9 | - | 71842687 | 72111871 | regular |
| ENSMUSG00000032232 | Cgnl1 | chr9 | - | 71626508 | 71771602 | regular |
| ENSMUSG00000032244 | Fem1b | chr9 | - | 62791820 | 62811934 | regular |
| ENSMUSG00000032253 | Phip | chr9 | - | 82866158 | 82975516 | regular |
| ENSMUSG00000032258 | Lca5 | chr9 | - | 83390292 | 83441127 | regular |
| ENSMUSG00000032280 | Tle3 | chr9 | + | 61372365 | 61418497 | regular |
| ENSMUSG00000032285 | Dnaja4 | chr9 | + | 54698872 | 54716315 | regular |
| ENSMUSG00000032290 | Ptpn9 | chr9 | + | 56994922 | 57062807 | regular |
| ENSMUSG00000032294 | Pkm | chr9 | + | 59656367 | 59679375 | regular |
| ENSMUSG00000032300 | 1700017B | chr9 | - | 57253116 | 57262612 | regular |
| ENSMUSG00000032316 | Clk3 | chr9 | - | 57750623 | 57765860 | regular |
| ENSMUSG00000032336 | Nptn | chr9 | + | 58582239 | 58657955 | regular |
| ENSMUSG00000032349 | Elovl5 | chr9 | + | 77917363 | 77984519 | regular |
| ENSMUSG00000032352 | Lrrc1 | chr9 | - | 77430822 | 77544870 | regular |
| ENSMUSG00000032355 | Mlip | chr9 | - | 77102080 | 77352969 | regular |
| ENSMUSG00000032382 | Snx1 | chr9 | - | 66088132 | 66126587 | regular |
| ENSMUSG00000032384 | Csnk1g1 | chr9 | + | 65908923 | 66045015 | regular |
| ENSMUSG00000032393 | Dpp8 | chr9 | + | 65032413 | 65082651 | regular |

| | | | | | | |
|---|---|---|---|---|---|---|
| ENSMUSG00000032396 | Dis3l | chr9 | - | 64306755 | 64341288 | regular |
| ENSMUSG00000032399 | Rpl4 | chr9 | + | 64173374 | 64178666 | regular |
| ENSMUSG00000032400 | Zwilch | chr9 | - | 64137143 | 64173104 | regular |
| ENSMUSG00000032409 | Atr | chr9 | + | 95857596 | 95951781 | regular |
| ENSMUSG00000032410 | Xrn1 | chr9 | + | 95954759 | 96057803 | regular |
| ENSMUSG00000032412 | Atp1b3 | chr9 | - | 96332654 | 96364442 | regular |
| ENSMUSG00000032418 | Me1 | chr9 | - | 86581370 | 86695953 | regular |
| ENSMUSG00000032423 | Syncrip | chr9 | - | 88447008 | 88482574 | regular |
| ENSMUSG00000032437 | Stt3b | chr9 | - | 115242580 | 115310421 | regular |
| ENSMUSG00000032479 | Map4 | chr9 | + | 109931459 | 110083955 | regular |
| ENSMUSG00000032481 | Smarcc1 | chr9 | + | 110117707 | 110240178 | regular |
| ENSMUSG00000032489 | Kif9 | chr9 | + | 110476957 | 110525179 | regular |
| ENSMUSG00000032496 | Ltf | chr9 | + | 111019270 | 111042767 | regular |
| ENSMUSG00000032497 | Lrrfip2 | chr9 | + | 111117591 | 111225668 | regular |
| ENSMUSG00000032504 | Pdcd6ip | chr9 | - | 113651743 | 113708259 | regular |
| ENSMUSG00000032512 | Wdr48 | chr9 | + | 119894877 | 119926587 | regular |
| ENSMUSG00000032514 | Ttc21a | chr9 | + | 119937605 | 119967793 | regular |
| ENSMUSG00000032518 | Rpsa | chr9 | + | 120127688 | 120132369 | regular |
| ENSMUSG00000032525 | Nktr | chr9 | + | 121719168 | 121756843 | regular |
| ENSMUSG00000032534 | Cep63 | chr9 | - | 102584587 | 102626534 | regular |
| ENSMUSG00000032536 | Trak1 | chr9 | + | 121297501 | 121474918 | regular |
| ENSMUSG00000032560 | Dnajc13 | chr9 | - | 104151281 | 104262930 | regular |
| ENSMUSG00000032562 | Gnai2 | chr9 | - | 107614124 | 107635367 | regular |
| ENSMUSG00000032566 | 1700080E | chr9 | - | 105143341 | 105145092 | regular |
| ENSMUSG00000032570 | Atp2c1 | chr9 | - | 105403538 | 105527319 | regular |
| ENSMUSG00000032580 | Rbm5 | chr9 | - | 107740370 | 107770996 | regular |
| ENSMUSG00000032582 | Rbm6 | chr9 | - | 107773558 | 107873237 | regular |
| ENSMUSG00000032594 | Ip6k1 | chr9 | + | 108002500 | 108048782 | regular |
| ENSMUSG00000032601 | Prkar2a | chr9 | + | 108689313 | 108750436 | regular |
| ENSMUSG00000032612 | Usp4 | chr9 | + | 108347852 | 108392545 | regular |
| ENSMUSG00000032621 | Srek1 | chr13 | - | 103739347 | 103774608 | regular |
| ENSMUSG00000032624 | Eml4 | chr17 | + | 83350930 | 83480361 | regular |
| ENSMUSG00000032637 | Atxn2l | chr7 | - | 126491707 | 126503437 | regular |
| ENSMUSG00000032688 | Malt1 | chr18 | + | 65430961 | 65478823 | regular |
| ENSMUSG00000032712 | 2810474C | chr6 | + | 149309413 | 149335663 | regular |
| ENSMUSG00000032740 | Ccdc88a | chr11 | + | 29373657 | 29510808 | regular |
| ENSMUSG00000032743 | D430042C | chr7 | + | 125707887 | 125874793 | regular |
| ENSMUSG00000032745 | Gpbp1 | chr13 | - | 111425679 | 111490111 | regular |
| ENSMUSG00000032777 | Gtf3c1 | chr7 | - | 125640953 | 125707780 | regular |
| ENSMUSG00000032802 | Srxn1 | chr2 | + | 152105515 | 152111376 | regular |
| ENSMUSG00000032803 | Cdv3 | chr9 | - | 103353093 | 103365840 | regular |
| ENSMUSG00000032812 | Arap1 | chr7 | + | 101348066 | 101412586 | regular |
| ENSMUSG00000032826 | Ank2 | chr3 | - | 126921611 | 127499350 | regular |
| ENSMUSG00000032846 | Zswim6 | chr13 | - | 107724616 | 107890064 | regular |
| ENSMUSG00000032869 | Psmf1 | chr2 | - | 151715811 | 151744186 | regular |
| ENSMUSG00000032872 | Cyb5r4 | chr9 | + | 87022013 | 87077774 | regular |

| | | | | | | |
|---|---|---|---|---|---|---|
| ENSMUSG00000032959 | Pebp1 | chr5 | - | 117282653 | 117287625 | regular |
| ENSMUSG00000032966 | Fkbp1a | chr2 | + | 151542482 | 151561692 | regular |
| ENSMUSG00000033004 | Mycbp2 | chr14 | - | 103113410 | 103346814 | regular |
| ENSMUSG00000033016 | Nfatc1 | chr18 | - | 80606204 | 80713071 | regular |
| ENSMUSG00000033022 | Cdo1 | chr18 | - | 46713204 | 46728342 | regular |
| ENSMUSG00000033039 | Micall1 | chr15 | + | 79108971 | 79136897 | regular |
| ENSMUSG00000033055 | Ankrd54 | chr15 | - | 79053093 | 79062893 | regular |
| ENSMUSG00000033075 | Senp1 | chr15 | - | 98038743 | 98093744 | regular |
| ENSMUSG00000033102 | Cdc14b | chr13 | - | 64189267 | 64275290 | regular |
| ENSMUSG00000033124 | Atg9a | chr1 | - | 75180859 | 75192196 | regular |
| ENSMUSG00000033216 | Eefsec | chr6 | - | 88257333 | 88446539 | regular |
| ENSMUSG00000033228 | Scaf11 | chr15 | - | 96411698 | 96460843 | regular |
| ENSMUSG00000033237 | Arid2 | chr15 | + | 96287517 | 96404992 | regular |
| ENSMUSG00000033257 | Ttll4 | chr1 | + | 74661744 | 74703730 | regular |
| ENSMUSG00000033306 | Lpp | chr16 | + | 24393349 | 24992576 | regular |
| ENSMUSG00000033319 | Fem1c | chr18 | - | 46501745 | 46525971 | regular |
| ENSMUSG00000033335 | Dnm2 | chr9 | + | 21424907 | 21507759 | regular |
| ENSMUSG00000033364 | Usp37 | chr1 | - | 74435510 | 74544284 | regular |
| ENSMUSG00000033392 | Clasp2 | chr9 | + | 113741472 | 113919682 | regular |
| ENSMUSG00000033411 | Ctdspl2 | chr2 | + | 121956000 | 122013642 | regular |
| ENSMUSG00000033423 | Eri3 | chr4 | + | 117550364 | 117674297 | regular |
| ENSMUSG00000033444 | Specc1l | chr10 | + | 75212072 | 75312743 | regular |
| ENSMUSG00000033478 | Fam160b: | chr19 | + | 57361008 | 57389594 | regular |
| ENSMUSG00000033486 | Catsper2 | chr2 | - | 121392630 | 121413792 | regular |
| ENSMUSG00000033487 | Fndc3a | chr14 | - | 72537945 | 72710003 | regular |
| ENSMUSG00000033499 | Larp4b | chr13 | + | 9093880 | 9174451 | regular |
| ENSMUSG00000033502 | Cdc14a | chr3 | - | 116272552 | 116424032 | regular |
| ENSMUSG00000033526 | Ppip5k1 | chr2 | - | 121310560 | 121355396 | regular |
| ENSMUSG00000033557 | Fam20b | chr1 | - | 156678531 | 156719086 | regular |
| ENSMUSG00000033632 | AW55491 | chr18 | + | 25168998 | 25467321 | regular |
| ENSMUSG00000033658 | Ddx19b | chr8 | - | 111003187 | 111031751 | regular |
| ENSMUSG00000033671 | Cep350 | chr1 | - | 155844963 | 155973255 | regular |
| ENSMUSG00000033701 | Acbd6 | chr1 | + | 155558119 | 155691330 | regular |
| ENSMUSG00000033705 | Stard9 | chr2 | + | 120629125 | 120731895 | regular |
| ENSMUSG00000033732 | Sf3b3 | chr8 | - | 110810238 | 110846787 | regular |
| ENSMUSG00000033739 | Fkbpl | chr17 | + | 34644763 | 34646324 | regular |
| ENSMUSG00000033760 | Rbm4b | chr19 | + | 4756542 | 4767065 | regular |
| ENSMUSG00000033767 | Tmem131 | chr3 | - | 83897654 | 84040175 | regular |
| ENSMUSG00000033769 | Exoc6b | chr6 | - | 84618486 | 85069513 | regular |
| ENSMUSG00000033793 | Atp6v1h | chr1 | + | 5070017 | 5162529 | regular |
| ENSMUSG00000033799 | Fam208b | chr13 | - | 3566034 | 3611108 | regular |
| ENSMUSG00000033813 | Tcea1 | chr1 | + | 4857813 | 4897909 | regular |
| ENSMUSG00000033819 | Ppp1r16a | chr15 | + | 76671614 | 76694919 | regular |
| ENSMUSG00000033826 | Dnah8 | chr17 | + | 30624353 | 30875264 | regular |
| ENSMUSG00000033882 | Rbm46 | chr3 | - | 82837227 | 82876483 | regular |
| ENSMUSG00000033904 | Ccp110 | chr7 | + | 118712551 | 118737024 | regular |

| | | | | | | |
|---|---|---|---|---|---|---|
| ENSMUSG00000033916 | Chmp2a | chr7 | - | 13032009 | 13034807 | regular |
| ENSMUSG00000033917 | Gde1 | chr7 | - | 118688544 | 118705778 | regular |
| ENSMUSG00000033943 | Mga | chr2 | + | 119897227 | 119969581 | regular |
| ENSMUSG00000033949 | Trim36 | chr18 | - | 46165301 | 46212607 | regular |
| ENSMUSG00000033983 | Coil | chr11 | + | 88970251 | 88991613 | regular |
| ENSMUSG00000033987 | Dnah17 | chr11 | - | 118021722 | 118130634 | regular |
| ENSMUSG00000034007 | Scaper | chr9 | - | 55549878 | 55938119 | regular |
| ENSMUSG00000034024 | Cct2 | chr10 | - | 117051000 | 117063814 | regular |
| ENSMUSG00000034042 | Gpbp1l1 | chr4 | + | 116557657 | 116593882 | regular |
| ENSMUSG00000034075 | Zdhhc5 | chr2 | - | 84687969 | 84715180 | regular |
| ENSMUSG00000034088 | Hdlbp | chr1 | - | 93405939 | 93478815 | regular |
| ENSMUSG00000034120 | Srsf2 | chr11 | - | 116849900 | 116853094 | regular |
| ENSMUSG00000034151 | Zbbx | chr3 | - | 75037906 | 75165034 | regular |
| ENSMUSG00000034154 | Ino80 | chr2 | - | 119373041 | 119477687 | regular |
| ENSMUSG00000034160 | Ogt | chrX | + | 101640059 | 101684351 | regular |
| ENSMUSG00000034163 | Zfc3h1 | chr10 | + | 115384958 | 115432772 | regular |
| ENSMUSG00000034187 | Nsf | chr11 | - | 103821781 | 103954056 | regular |
| ENSMUSG00000034201 | Gas2l1 | chr11 | - | 5054131 | 5065327 | regular |
| ENSMUSG00000034212 | Ankmy1 | chr1 | - | 92859802 | 92902906 | regular |
| ENSMUSG00000034235 | Usp54 | chr14 | - | 20548911 | 20641063 | regular |
| ENSMUSG00000034239 | Gm884 | chr11 | - | 103534576 | 103621140 | regular |
| ENSMUSG00000034243 | Golgb1 | chr16 | + | 36875139 | 36933085 | regular |
| ENSMUSG00000034245 | Hdac11 | chr6 | + | 91156664 | 91174692 | regular |
| ENSMUSG00000034252 | Senp6 | chr9 | + | 80066902 | 80144953 | regular |
| ENSMUSG00000034254 | Agpat1 | chr17 | + | 34604261 | 34613449 | regular |
| ENSMUSG00000034258 | Flvcr2 | chr12 | + | 85746538 | 85813585 | regular |
| ENSMUSG00000034269 | Setd5 | chr6 | + | 113077364 | 113153435 | regular |
| ENSMUSG00000034297 | Med13 | chr11 | - | 86267032 | 86357602 | regular |
| ENSMUSG00000034300 | Fam53c | chr18 | + | 34758905 | 34773760 | regular |
| ENSMUSG00000034303 | Ccdc15 | chr9 | - | 37275834 | 37348432 | regular |
| ENSMUSG00000034312 | Iqsec1 | chr6 | - | 90656087 | 90988685 | regular |
| ENSMUSG00000034334 | Fam151b | chr13 | - | 92449624 | 92484015 | regular |
| ENSMUSG00000034342 | Cbl | chr9 | - | 44142975 | 44234049 | regular |
| ENSMUSG00000034343 | Ube2f | chr1 | + | 91250303 | 91290337 | regular |
| ENSMUSG00000034349 | Smc4 | chr3 | + | 69004737 | 69034623 | regular |
| ENSMUSG00000034354 | Mtmr3 | chr11 | - | 4480867 | 4594863 | regular |
| ENSMUSG00000034377 | Tulp4 | chr17 | + | 6106436 | 6251128 | regular |
| ENSMUSG00000034390 | Cmip | chr8 | + | 117257063 | 117459430 | regular |
| ENSMUSG00000034401 | Spata6 | chr4 | + | 111719983 | 111829184 | regular |
| ENSMUSG00000034485 | Uaca | chr9 | + | 60794541 | 60880370 | regular |
| ENSMUSG00000034543 | Morc2a | chr11 | + | 3649493 | 3690477 | regular |
| ENSMUSG00000034544 | Rsrc1 | chr3 | + | 66981389 | 67358396 | regular |
| ENSMUSG00000034574 | Daam1 | chr12 | + | 71831077 | 71992333 | regular |
| ENSMUSG00000034601 | 2700049A | chr12 | + | 71136847 | 71243303 | regular |
| ENSMUSG00000034610 | Zcchc11 | chr4 | + | 108459425 | 108559421 | regular |
| ENSMUSG00000034647 | Ankrd12 | chr17 | - | 65967500 | 66077089 | regular |

| | | | | | | |
|---|---|---|---|---|---|---|
| ENSMUSG00000034667 | Xpot | chr10 | - | 121587379 | 121626332 | regular |
| ENSMUSG00000034707 | Gns | chr10 | + | 121365089 | 121397249 | regular |
| ENSMUSG00000034724 | Cnot6l | chr5 | - | 96070332 | 96164171 | regular |
| ENSMUSG00000034731 | Dgkh | chr14 | - | 78558749 | 78732776 | regular |
| ENSMUSG00000034751 | Mast4 | chr13 | - | 102732485 | 103334497 | regular |
| ENSMUSG00000034793 | G6pc3 | chr11 | + | 102189619 | 102194081 | regular |
| ENSMUSG00000034813 | Grip1 | chr10 | + | 119453829 | 120087261 | regular |
| ENSMUSG00000034842 | Art3 | chr5 | + | 92331826 | 92414628 | regular |
| ENSMUSG00000034892 | Rps29 | chr12 | - | 69157721 | 69159186 | regular |
| ENSMUSG00000034908 | Sidt2 | chr9 | - | 45937856 | 45955258 | regular |
| ENSMUSG00000034913 | Spert | chr14 | - | 75582876 | 75593116 | regular |
| ENSMUSG00000034928 | Rnf44 | chr13 | - | 54679398 | 54693907 | regular |
| ENSMUSG00000034940 | Synrg | chr11 | + | 83964427 | 84044578 | regular |
| ENSMUSG00000034994 | Eef2 | chr10 | + | 81176630 | 81182498 | regular |
| ENSMUSG00000035021 | Baz1a | chr12 | - | 54892988 | 55014348 | regular |
| ENSMUSG00000035027 | Map2k2 | chr10 | + | 81105914 | 81133975 | regular |
| ENSMUSG00000035032 | Nek11 | chr9 | - | 105162155 | 105395524 | regular |
| ENSMUSG00000035086 | Becn1 | chr11 | - | 101285951 | 101302286 | regular |
| ENSMUSG00000035126 | Wdr78 | chr4 | - | 103038064 | 103114555 | regular |
| ENSMUSG00000035131 | Brinp3 | chr1 | + | 146494759 | 146902472 | regular |
| ENSMUSG00000035133 | Arhgap5 | chr12 | + | 52503971 | 52571975 | regular |
| ENSMUSG00000035152 | Ap2b1 | chr11 | + | 83299023 | 83405035 | regular |
| ENSMUSG00000035161 | Ints6 | chr14 | - | 62676329 | 62761169 | regular |
| ENSMUSG00000035202 | Lars2 | chr9 | + | 123366926 | 123462666 | regular |
| ENSMUSG00000035203 | Epn1 | chr7 | + | 5080234 | 5098178 | regular |
| ENSMUSG00000035211 | Xrra1 | chr7 | + | 99859115 | 99917822 | regular |
| ENSMUSG00000035242 | Oaz1 | chr10 | + | 80826655 | 80829290 | regular |
| ENSMUSG00000035247 | Hectd1 | chr12 | - | 51743721 | 51829536 | regular |
| ENSMUSG00000035297 | Cops4 | chr5 | + | 100518308 | 100547803 | regular |
| ENSMUSG00000035325 | Sec31a | chr5 | - | 100361648 | 100416234 | regular |
| ENSMUSG00000035356 | Nfkbiz | chr16 | - | 55811374 | 55838899 | regular |
| ENSMUSG00000035378 | Shq1 | chr6 | - | 100568255 | 100671157 | regular |
| ENSMUSG00000035392 | Dennd1a | chr2 | - | 37798990 | 38287390 | regular |
| ENSMUSG00000035399 | Oser1 | chr2 | - | 163405821 | 163424697 | regular |
| ENSMUSG00000035435 | Abca17 | chr17 | - | 24264258 | 24351029 | regular |
| ENSMUSG00000035437 | Rabgap1 | chr2 | + | 37443278 | 37566454 | regular |
| ENSMUSG00000035473 | Galm | chr17 | + | 80127470 | 80185101 | regular |
| ENSMUSG00000035478 | Mbd3 | chr10 | - | 80392538 | 80399550 | regular |
| ENSMUSG00000035504 | Reep6 | chr10 | + | 80329952 | 80336441 | regular |
| ENSMUSG00000035517 | Tdrd7 | chr4 | + | 45965333 | 46034761 | regular |
| ENSMUSG00000035530 | Eif1 | chr11 | + | 100319884 | 100322096 | regular |
| ENSMUSG00000035539 | Ccdc180 | chr4 | + | 45890302 | 45950774 | regular |
| ENSMUSG00000035545 | Leng8 | chr7 | + | 4137038 | 4148177 | regular |
| ENSMUSG00000035569 | Ankrd11 | chr8 | - | 122883821 | 123042277 | regular |
| ENSMUSG00000035621 | Midn | chr10 | + | 80148271 | 80158368 | regular |
| ENSMUSG00000035623 | Rsf1 | chr7 | + | 97579888 | 97692778 | regular |

| | | | | | | |
|---|---|---|---|---|---|---|
| ENSMUSG00000035629 | Rubcn | chr16 | - | 32821702 | 32877766 | regular |
| ENSMUSG00000035649 | Zcchc7 | chr4 | + | 44755876 | 44932215 | regular |
| ENSMUSG00000035696 | Rnf38 | chr4 | - | 44126209 | 44233789 | regular |
| ENSMUSG00000035697 | Arhgap45 | chr10 | + | 80016652 | 80031472 | regular |
| ENSMUSG00000035765 | Dym | chr18 | + | 75018771 | 75286966 | regular |
| ENSMUSG00000035770 | Dync1li2 | chr8 | - | 104417679 | 104443047 | regular |
| ENSMUSG00000035845 | Alg12 | chr15 | - | 88805243 | 88819318 | regular |
| ENSMUSG00000035851 | Ythdc1 | chr5 | + | 86804220 | 86836659 | regular |
| ENSMUSG00000035891 | Cerk | chr15 | - | 86139127 | 86186141 | regular |
| ENSMUSG00000035901 | Dennd5a | chr7 | - | 109893779 | 109960470 | regular |
| ENSMUSG00000035919 | Bbs9 | chr9 | + | 22475714 | 22888280 | regular |
| ENSMUSG00000035992 | Fnip1 | chr11 | + | 54438198 | 54518235 | regular |
| ENSMUSG00000036002 | Fam214b | chr4 | - | 43032413 | 43046220 | regular |
| ENSMUSG00000036006 | Ripor2 | chr13 | + | 24582188 | 24733816 | regular |
| ENSMUSG00000036019 | Tmtc2 | chr10 | - | 105187662 | 105574451 | regular |
| ENSMUSG00000036046 | 5031439G | chr15 | - | 84943935 | 84988551 | regular |
| ENSMUSG00000036054 | Sugp2 | chr8 | + | 70234225 | 70279915 | regular |
| ENSMUSG00000036087 | Slain2 | chr5 | + | 72914303 | 72978829 | regular |
| ENSMUSG00000036097 | Slf2 | chr19 | + | 44931118 | 44983787 | regular |
| ENSMUSG00000036168 | Ccdc38 | chr10 | + | 93540631 | 93584327 | regular |
| ENSMUSG00000036180 | Gatad2a | chr8 | - | 69907075 | 69996384 | regular |
| ENSMUSG00000036196 | Slc26a8 | chr17 | - | 28637782 | 28689987 | regular |
| ENSMUSG00000036199 | Ndufa13 | chr8 | - | 69894179 | 69902558 | regular |
| ENSMUSG00000036202 | Rif1 | chr2 | + | 52072831 | 52122383 | regular |
| ENSMUSG00000036241 | Ube2r2 | chr4 | + | 41135742 | 41193380 | regular |
| ENSMUSG00000036244 | Tbc1d21 | chr9 | - | 58359703 | 58370455 | regular |
| ENSMUSG00000036257 | Pnpla8 | chr12 | + | 44221369 | 44322532 | regular |
| ENSMUSG00000036270 | Edc4 | chr8 | + | 105880880 | 105894908 | regular |
| ENSMUSG00000036273 | Lrrk2 | chr15 | + | 91673174 | 91816120 | regular |
| ENSMUSG00000036292 | Gramd1c | chr16 | - | 43980349 | 44063345 | regular |
| ENSMUSG00000036309 | Skp1a | chr11 | + | 52231994 | 52246858 | regular |
| ENSMUSG00000036371 | Serbp1 | chr6 | + | 67238175 | 67297736 | regular |
| ENSMUSG00000036377 | C530008M | chr5 | + | 76656511 | 76873554 | regular |
| ENSMUSG00000036398 | Ppp1r11 | chr17 | - | 36948355 | 36951741 | regular |
| ENSMUSG00000036403 | Cep135 | chr5 | + | 76588697 | 76646466 | regular |
| ENSMUSG00000036411 | 9530077C | chr9 | + | 22411512 | 22444681 | regular |
| ENSMUSG00000036427 | Gpi1 | chr7 | - | 34201329 | 34230336 | regular |
| ENSMUSG00000036438 | Calm2 | chr17 | - | 87433411 | 87446935 | regular |
| ENSMUSG00000036461 | Elf1 | chr14 | + | 79481193 | 79582494 | regular |
| ENSMUSG00000036478 | Btg1 | chr10 | + | 96617005 | 96622809 | regular |
| ENSMUSG00000036501 | Fam13b | chr18 | - | 34442351 | 34506823 | regular |
| ENSMUSG00000036529 | Sbf1 | chr15 | - | 89288235 | 89315311 | regular |
| ENSMUSG00000036550 | Cnot1 | chr8 | - | 95719450 | 95807464 | regular |
| ENSMUSG00000036552 | Ermard | chr17 | + | 14978863 | 15064232 | regular |
| ENSMUSG00000036555 | Iqce | chr5 | - | 140661826 | 140702378 | regular |
| ENSMUSG00000036557 | Stpg4 | chr17 | - | 87389570 | 87427741 | regular |

| | | | | | | |
|---|---|---|---|---|---|---|
| ENSMUSG00000036574 | Tex44 | chr1 | + | 86426328 | 86428052 | regular |
| ENSMUSG00000036594 | H2-Aa | chr17 | - | 34282743 | 34287827 | regular |
| ENSMUSG00000036606 | Plxnb2 | chr15 | - | 89155548 | 89171120 | regular |
| ENSMUSG00000036613 | Tssc1 | chr12 | + | 28751802 | 28869865 | regular |
| ENSMUSG00000036641 | Ccdc148 | chr2 | - | 58821069 | 59160683 | regular |
| ENSMUSG00000036646 | Man1b1 | chr2 | + | 25332337 | 25352212 | regular |
| ENSMUSG00000036698 | Ago2 | chr15 | - | 73095843 | 73184840 | regular |
| ENSMUSG00000036707 | Cab39 | chr1 | + | 85793440 | 85851576 | regular |
| ENSMUSG00000036712 | Cyld | chr8 | + | 88697027 | 88751945 | regular |
| ENSMUSG00000036752 | Tubb4b | chr2 | - | 25222159 | 25224702 | regular |
| ENSMUSG00000036768 | Kif15 | chr9 | + | 122951045 | 123018733 | regular |
| ENSMUSG00000036779 | Papd5 | chr8 | + | 88199212 | 88259722 | regular |
| ENSMUSG00000036792 | Mbd5 | chr2 | + | 48949507 | 49325405 | regular |
| ENSMUSG00000036825 | Ssx2ip | chr3 | + | 146404641 | 146440144 | regular |
| ENSMUSG00000036833 | Pnpla7 | chr2 | + | 24976032 | 25054057 | regular |
| ENSMUSG00000036872 | Abcc12 | chr8 | - | 86482259 | 86580686 | regular |
| ENSMUSG00000036879 | Phkb | chr8 | + | 85840958 | 86061376 | regular |
| ENSMUSG00000036882 | Arhgap33 | chr7 | - | 30522225 | 30535060 | regular |
| ENSMUSG00000036893 | Ehmt1 | chr2 | - | 24789927 | 24919614 | regular |
| ENSMUSG00000036918 | Ttc7 | chr17 | + | 87282885 | 87381769 | regular |
| ENSMUSG00000036928 | Stag3 | chr5 | + | 138280239 | 138312393 | regular |
| ENSMUSG00000036964 | Trim17 | chr11 | + | 58954684 | 58973098 | regular |
| ENSMUSG00000036990 | Otud4 | chr8 | + | 79639617 | 79677724 | regular |
| ENSMUSG00000036995 | Asap3 | chr4 | + | 136206364 | 136245216 | regular |
| ENSMUSG00000037012 | Hk1 | chr10 | - | 62268854 | 62379908 | regular |
| ENSMUSG00000037013 | Ss18 | chr18 | - | 14624197 | 14682914 | regular |
| ENSMUSG00000037020 | Wdr62 | chr7 | - | 30240137 | 30280419 | regular |
| ENSMUSG00000037058 | Paip2 | chr18 | + | 35598616 | 35617186 | regular |
| ENSMUSG00000037062 | Sh3glb1 | chr3 | - | 144683677 | 144720335 | regular |
| ENSMUSG00000037071 | Scd1 | chr19 | - | 44394450 | 44407709 | regular |
| ENSMUSG00000037075 | Rnf139 | chr15 | + | 58889228 | 58907057 | regular |
| ENSMUSG00000037101 | Ttc29 | chr8 | + | 78213296 | 78394326 | regular |
| ENSMUSG00000037110 | Ralgapa2 | chr2 | - | 146239878 | 146512344 | regular |
| ENSMUSG00000037112 | Sik2 | chr9 | - | 50892800 | 51009073 | regular |
| ENSMUSG00000037119 | Fam91a1 | chr15 | + | 58415467 | 58457740 | regular |
| ENSMUSG00000037143 | Cfap61 | chr2 | + | 145934783 | 146215039 | regular |
| ENSMUSG00000037148 | Arhgap10 | chr8 | - | 77250365 | 77517953 | regular |
| ENSMUSG00000037149 | Ddx1 | chr12 | - | 13216972 | 13249213 | regular |
| ENSMUSG00000037174 | Elf2 | chr3 | - | 51252719 | 51340663 | regular |
| ENSMUSG00000037196 | Pacrg | chr17 | - | 10403011 | 10840311 | regular |
| ENSMUSG00000037210 | Fam193a | chr5 | + | 34369932 | 34486456 | regular |
| ENSMUSG00000037236 | Matr3 | chr18 | + | 35562145 | 35593835 | regular |
| ENSMUSG00000037266 | Rsrp1 | chr4 | + | 134923591 | 134927671 | regular |
| ENSMUSG00000037270 | 4932438A | chr3 | + | 36863103 | 37053033 | regular |
| ENSMUSG00000037278 | Tmem97 | chr11 | - | 78541816 | 78550777 | regular |
| ENSMUSG00000037286 | Stag1 | chr9 | + | 100597797 | 100959375 | regular |

| | | | | | | |
|---|---|---|---|---|---|---|
| ENSMUSG00000037287 | Tbcel | chr9 | - | 42412315 | 42507809 | regular |
| ENSMUSG00000037331 | Larp1 | chr11 | + | 58009063 | 58062034 | regular |
| ENSMUSG00000037343 | Taf2 | chr15 | - | 55015130 | 55072152 | regular |
| ENSMUSG00000037363 | Letm2 | chr8 | - | 25578489 | 25597582 | regular |
| ENSMUSG00000037364 | Srrt | chr5 | - | 137295703 | 137307674 | regular |
| ENSMUSG00000037386 | Rims2 | chr15 | + | 39198260 | 39684372 | regular |
| ENSMUSG00000037395 | Rcor3 | chr1 | - | 192098545 | 192138062 | regular |
| ENSMUSG00000037416 | Dmxl1 | chr18 | + | 49832669 | 49965473 | regular |
| ENSMUSG00000037432 | Fer1l5 | chr1 | + | 36372290 | 36422110 | regular |
| ENSMUSG00000037437 | Adam32 | chr8 | - | 24836139 | 24948804 | regular |
| ENSMUSG00000037443 | Cep85 | chr4 | - | 134129857 | 134187112 | regular |
| ENSMUSG00000037458 | Azin1 | chr15 | - | 38487426 | 38519266 | regular |
| ENSMUSG00000037486 | Asxl2 | chr12 | + | 3426856 | 3506852 | regular |
| ENSMUSG00000037487 | Ubr5 | chr15 | - | 37967327 | 38078854 | regular |
| ENSMUSG00000037503 | Fam168b | chr1 | - | 34813218 | 34843065 | regular |
| ENSMUSG00000037533 | Rapgef6 | chr11 | + | 54522846 | 54699285 | regular |
| ENSMUSG00000037536 | Fbxo34 | chr14 | + | 47450420 | 47531962 | regular |
| ENSMUSG00000037568 | Vash2 | chr1 | - | 190947645 | 190979296 | regular |
| ENSMUSG00000037608 | Bclaf1 | chr10 | + | 20312468 | 20344613 | regular |
| ENSMUSG00000037622 | Wdtc1 | chr4 | - | 133292458 | 133353481 | regular |
| ENSMUSG00000037627 | Rgs22 | chr15 | - | 36009478 | 36140400 | regular |
| ENSMUSG00000037646 | Vps13b | chr15 | + | 35371159 | 35931229 | regular |
| ENSMUSG00000037652 | Phc3 | chr3 | - | 30899370 | 30969415 | regular |
| ENSMUSG00000037674 | Rfx7 | chr9 | + | 72532239 | 72622937 | regular |
| ENSMUSG00000037683 | Armc3 | chr2 | + | 19199301 | 19310241 | regular |
| ENSMUSG00000037697 | Ddhd1 | chr14 | - | 45588466 | 45658143 | regular |
| ENSMUSG00000037716 | Ccdc33 | chr9 | - | 58028676 | 58118823 | regular |
| ENSMUSG00000037742 | Eef1a1 | chr9 | - | 78478448 | 78489151 | regular |
| ENSMUSG00000037754 | Ppp1r16b | chr2 | + | 158665397 | 158766334 | regular |
| ENSMUSG00000037795 | N4bp2 | chr5 | + | 65763520 | 65830108 | regular |
| ENSMUSG00000037801 | Iqch | chr9 | - | 63421454 | 63602493 | regular |
| ENSMUSG00000037805 | Rpl10a | chr17 | + | 28328470 | 28331032 | regular |
| ENSMUSG00000037822 | Smim14 | chr5 | - | 65446843 | 65537184 | regular |
| ENSMUSG00000037857 | Nufip2 | chr11 | + | 77686154 | 77741921 | regular |
| ENSMUSG00000037876 | Jmjd1c | chr10 | + | 67096124 | 67256326 | regular |
| ENSMUSG00000037885 | Stk35 | chr2 | + | 129800516 | 129832287 | regular |
| ENSMUSG00000037922 | Bank1 | chr3 | - | 136053362 | 136326066 | regular |
| ENSMUSG00000037926 | Ssh2 | chr11 | + | 77216286 | 77460220 | regular |
| ENSMUSG00000037933 | Bicd2 | chr13 | + | 49341584 | 49387026 | regular |
| ENSMUSG00000037940 | Inpp4b | chr8 | + | 81342555 | 82127914 | regular |
| ENSMUSG00000037965 | Zc3h7a | chr16 | - | 11136591 | 11176393 | regular |
| ENSMUSG00000037971 | 1110032A | chr9 | - | 50762827 | 50775520 | regular |
| ENSMUSG00000037972 | Snn | chr16 | + | 11066297 | 11074983 | regular |
| ENSMUSG00000037979 | Ccdc92 | chr5 | - | 124834417 | 124862424 | regular |
| ENSMUSG00000037999 | Arap2 | chr5 | - | 62602444 | 62766159 | regular |
| ENSMUSG00000038010 | Ccdc138 | chr10 | + | 58497947 | 58576244 | regular |

| | | | | | | |
|---|---|---|---|---|---|---|
| ENSMUSG00000038011 | Dnah10 | chr5 | + | 124725084 | 124834308 | regular |
| ENSMUSG00000038014 | Fam120a | chr13 | - | 48879218 | 48968017 | regular |
| ENSMUSG00000038015 | Prm2 | chr16 | - | 10791379 | 10796134 | regular |
| ENSMUSG00000038039 | Gcc2 | chr10 | + | 58255496 | 58305599 | regular |
| ENSMUSG00000038042 | Ptpdc1 | chr13 | - | 48577871 | 48625664 | regular |
| ENSMUSG00000038056 | Kmt2c | chr5 | - | 25271797 | 25498783 | regular |
| ENSMUSG00000038057 | Dbil5 | chr11 | + | 76217607 | 76218666 | regular |
| ENSMUSG00000038060 | Dlec1 | chr9 | + | 119102477 | 119148246 | regular |
| ENSMUSG00000038068 | Rnf144b | chr13 | + | 47122655 | 47247991 | regular |
| ENSMUSG00000038070 | Cntln | chr4 | + | 84884308 | 85131921 | regular |
| ENSMUSG00000038095 | Sbno1 | chr5 | - | 124368701 | 124426001 | regular |
| ENSMUSG00000038102 | Trappc11 | chr8 | - | 47490114 | 47533470 | regular |
| ENSMUSG00000038116 | Phf20 | chr2 | + | 156196465 | 156309952 | regular |
| ENSMUSG00000038122 | Tbc1d32 | chr10 | - | 56014292 | 56228689 | regular |
| ENSMUSG00000038126 | Mphosph9 | chr5 | - | 124250958 | 124327972 | regular |
| ENSMUSG00000038127 | Ccdc50 | chr16 | + | 27388868 | 27452218 | regular |
| ENSMUSG00000038128 | Camk4 | chr18 | + | 32939040 | 33195767 | regular |
| ENSMUSG00000038165 | 1700022A | chr5 | + | 24645452 | 24648860 | regular |
| ENSMUSG00000038170 | Pde4dip | chr3 | - | 97689823 | 97888707 | regular |
| ENSMUSG00000038180 | Spag4 | chr2 | + | 156065174 | 156070617 | regular |
| ENSMUSG00000038252 | Ncapd2 | chr6 | - | 125168006 | 125191701 | regular |
| ENSMUSG00000038260 | Trpm4 | chr7 | - | 45302631 | 45333780 | regular |
| ENSMUSG00000038274 | Fau | chr19 | + | 6057887 | 6059524 | regular |
| ENSMUSG00000038290 | Smg6 | chr11 | + | 74925822 | 75164448 | regular |
| ENSMUSG00000038298 | Pdzk1 | chr3 | + | 96829283 | 96870926 | regular |
| ENSMUSG00000038324 | Trpc4ap | chr2 | - | 155634270 | 155692384 | regular |
| ENSMUSG00000038347 | Tcte2 | chr17 | - | 13482552 | 13761825 | regular |
| ENSMUSG00000038368 | Focad | chr4 | + | 88094628 | 88411011 | regular |
| ENSMUSG00000038369 | Ncoa6 | chr2 | - | 155390655 | 155473894 | regular |
| ENSMUSG00000038371 | Sbf2 | chr7 | - | 110308012 | 110614922 | regular |
| ENSMUSG00000038388 | Mpp6 | chr6 | + | 50110240 | 50198939 | regular |
| ENSMUSG00000038393 | Txnip | chr3 | + | 96557956 | 96561883 | regular |
| ENSMUSG00000038429 | Usp5 | chr6 | - | 124815018 | 124829484 | regular |
| ENSMUSG00000038467 | Chmp4b | chr2 | + | 154651704 | 154694785 | regular |
| ENSMUSG00000038481 | Cdk19 | chr10 | + | 40339563 | 40483818 | regular |
| ENSMUSG00000038485 | Socs7 | chr11 | + | 97362550 | 97398542 | regular |
| ENSMUSG00000038495 | Otud7b | chr3 | + | 96104526 | 96161129 | regular |
| ENSMUSG00000038518 | Jarid2 | chr13 | + | 44729473 | 44921643 | regular |
| ENSMUSG00000038523 | 1700003F | chr2 | + | 154547084 | 154550003 | regular |
| ENSMUSG00000038538 | Ubn2 | chr6 | + | 38433949 | 38524825 | regular |
| ENSMUSG00000038546 | Ranbp9 | chr13 | - | 43402674 | 43481282 | regular |
| ENSMUSG00000038564 | Ift172 | chr5 | - | 31253276 | 31291116 | regular |
| ENSMUSG00000038587 | Akap12 | chr10 | + | 4266379 | 4359470 | regular |
| ENSMUSG00000038594 | Cep85l | chr10 | - | 53273442 | 53379947 | regular |
| ENSMUSG00000038608 | Dock10 | chr1 | - | 80501072 | 80758527 | regular |
| ENSMUSG00000038612 | Mcl1 | chr3 | + | 95658787 | 95663176 | regular |

| | | | | | | |
|---|---|---|---|---|---|---|
| ENSMUSG00000038637 | Lrrc56 | chr7 | + | 141194156 | 141210055 | regular |
| ENSMUSG00000038658 | Ric1 | chr19 | + | 29522281 | 29606829 | regular |
| ENSMUSG00000038664 | Herc1 | chr9 | + | 66350449 | 66508775 | regular |
| ENSMUSG00000038679 | Trps1 | chr15 | - | 50654751 | 50890463 | regular |
| ENSMUSG00000038685 | Rtel1 | chr2 | + | 181319738 | 181356616 | regular |
| ENSMUSG00000038708 | Golga4 | chr9 | + | 118506266 | 118582519 | regular |
| ENSMUSG00000038733 | Wdr26 | chr1 | - | 181173227 | 181211552 | regular |
| ENSMUSG00000038756 | Ttll6 | chr11 | + | 96133785 | 96165451 | regular |
| ENSMUSG00000038759 | Nup205 | chr6 | + | 35177420 | 35247596 | regular |
| ENSMUSG00000038762 | Abcf1 | chr17 | - | 35956818 | 35969761 | regular |
| ENSMUSG00000038766 | Gabpb2 | chr3 | - | 95181765 | 95217916 | regular |
| ENSMUSG00000038773 | Kdm3b | chr18 | + | 34777046 | 34838660 | regular |
| ENSMUSG00000038774 | Ascc3 | chr10 | + | 50592668 | 50851485 | regular |
| ENSMUSG00000038828 | Tmem214 | chr5 | + | 30868011 | 30879180 | regular |
| ENSMUSG00000038831 | Ralgps1 | chr2 | - | 33133416 | 33371486 | regular |
| ENSMUSG00000038844 | Kif16b | chr2 | - | 142617473 | 142901531 | regular |
| ENSMUSG00000038855 | Itpkb | chr1 | + | 180330484 | 180424802 | regular |
| ENSMUSG00000038860 | Garnl3 | chr2 | - | 32986223 | 33131654 | regular |
| ENSMUSG00000038866 | Zcchc2 | chr1 | + | 105990405 | 106034074 | regular |
| ENSMUSG00000038900 | Rpl12 | chr2 | + | 32961558 | 32965345 | regular |
| ENSMUSG00000038909 | Kat7 | chr11 | - | 95274258 | 95310246 | regular |
| ENSMUSG00000038914 | Dido1 | chr2 | - | 180657963 | 180709999 | regular |
| ENSMUSG00000038932 | Tcfl5 | chr2 | - | 180621955 | 180642708 | regular |
| ENSMUSG00000038936 | Sccpdh | chr1 | + | 179668209 | 179687189 | regular |
| ENSMUSG00000038954 | Supt3 | chr17 | + | 44777151 | 45119290 | regular |
| ENSMUSG00000038967 | Pdk2 | chr11 | - | 95026257 | 95041354 | regular |
| ENSMUSG00000038970 | Lmtk2 | chr5 | + | 144100435 | 144188204 | regular |
| ENSMUSG00000038994 | Hils1 | chr11 | + | 94967592 | 94968456 | regular |
| ENSMUSG00000039000 | Ube3c | chr5 | + | 29569241 | 29676092 | regular |
| ENSMUSG00000039001 | Rps21 | chr2 | + | 180257376 | 180258445 | regular |
| ENSMUSG00000039007 | Cpq | chr15 | + | 33083128 | 33594552 | regular |
| ENSMUSG00000039021 | Ttc16 | chr2 | - | 32757025 | 32775633 | regular |
| ENSMUSG00000039033 | Tasp1 | chr2 | - | 139833479 | 140066805 | regular |
| ENSMUSG00000039041 | Adrm1 | chr2 | + | 180171484 | 180176286 | regular |
| ENSMUSG00000039046 | Usp6nl | chr2 | + | 6322666 | 6446390 | regular |
| ENSMUSG00000039050 | Osbpl2 | chr2 | + | 180119305 | 180162680 | regular |
| ENSMUSG00000039068 | Zzz3 | chr3 | + | 152395472 | 152462826 | regular |
| ENSMUSG00000039100 | Mar.06 | chr15 | - | 31455890 | 31531053 | regular |
| ENSMUSG00000039105 | Atp6v1g1 | chr4 | + | 63544771 | 63550750 | regular |
| ENSMUSG00000039110 | Mycbpap | chr11 | - | 94501346 | 94521742 | regular |
| ENSMUSG00000039158 | Akna | chr4 | - | 63367124 | 63403354 | regular |
| ENSMUSG00000039168 | Dap | chr15 | + | 31224313 | 31274341 | regular |
| ENSMUSG00000039191 | Rbpj | chr5 | + | 53466151 | 53657362 | regular |
| ENSMUSG00000039202 | Abhd2 | chr7 | + | 79273198 | 79365508 | regular |
| ENSMUSG00000039210 | Gpatch2 | chr1 | + | 187215507 | 187351704 | regular |
| ENSMUSG00000039218 | Srrm2 | chr17 | + | 23790661 | 23824741 | regular |

| | | | | | | |
|---|---|---|---|---|---|---|
| ENSMUSG00000039219 | Arid4b | chr13 | + | 14063231 | 14199603 | regular |
| ENSMUSG00000039220 | Ppp1r10 | chr17 | + | 35916433 | 35932283 | regular |
| ENSMUSG00000039224 | D1Pas1 | chr1 | + | 186967415 | 186970627 | regular |
| ENSMUSG00000039236 | Isg20 | chr7 | + | 78913423 | 78920396 | regular |
| ENSMUSG00000039254 | Pomt1 | chr2 | + | 32236589 | 32255005 | regular |
| ENSMUSG00000039262 | Prrc2b | chr2 | + | 32151081 | 32234537 | regular |
| ENSMUSG00000039275 | Foxk2 | chr11 | + | 121259989 | 121309896 | regular |
| ENSMUSG00000039285 | Azi2 | chr9 | + | 118040498 | 118069794 | regular |
| ENSMUSG00000039298 | Cdk5rap2 | chr4 | - | 70216855 | 70410443 | regular |
| ENSMUSG00000039318 | Rab3gap2 | chr1 | + | 185204116 | 185286759 | regular |
| ENSMUSG00000039335 | Spata16 | chr3 | + | 26637619 | 26983212 | regular |
| ENSMUSG00000039342 | Ankar | chr1 | - | 72642979 | 72700579 | regular |
| ENSMUSG00000039361 | Picalm | chr7 | + | 90130212 | 90213465 | regular |
| ENSMUSG00000039391 | Ccdc81 | chr7 | - | 89866147 | 89903629 | regular |
| ENSMUSG00000039456 | Morc3 | chr16 | + | 93832120 | 93876072 | regular |
| ENSMUSG00000039458 | Mtmr12 | chr15 | + | 12205027 | 12274496 | regular |
| ENSMUSG00000039473 | Ubn1 | chr16 | + | 5050067 | 5086285 | regular |
| ENSMUSG00000039477 | Tnrc18 | chr5 | - | 142724660 | 142817662 | regular |
| ENSMUSG00000039501 | Znfx1 | chr2 | - | 167035792 | 167063015 | regular |
| ENSMUSG00000039523 | Cep104 | chr4 | + | 153975193 | 154008732 | regular |
| ENSMUSG00000039530 | Tusc3 | chr8 | + | 39005844 | 39165114 | regular |
| ENSMUSG00000039543 | Cfap70 | chr14 | - | 20394192 | 20452226 | regular |
| ENSMUSG00000039568 | Ubald1 | chr16 | - | 4874777 | 4880315 | regular |
| ENSMUSG00000039585 | Myo9a | chr9 | + | 59750895 | 59928866 | regular |
| ENSMUSG00000039621 | Prex1 | chr2 | - | 166566341 | 166713832 | regular |
| ENSMUSG00000039630 | Hnrnpu | chr1 | - | 178321107 | 178337797 | regular |
| ENSMUSG00000039632 | Ccdc151 | chr9 | - | 21989870 | 22002634 | regular |
| ENSMUSG00000039652 | Cpeb3 | chr19 | - | 37021290 | 37208601 | regular |
| ENSMUSG00000039703 | Nploc4 | chr11 | - | 120380369 | 120437708 | regular |
| ENSMUSG00000039716 | Dock3 | chr9 | - | 106892824 | 107231909 | regular |
| ENSMUSG00000039717 | Ralyl | chr3 | + | 13471654 | 14182287 | regular |
| ENSMUSG00000039735 | Fnbp1l | chr3 | - | 122538718 | 122619715 | regular |
| ENSMUSG00000039738 | Slx4 | chr16 | - | 3979104 | 4003770 | regular |
| ENSMUSG00000039782 | Cpeb2 | chr5 | + | 43233169 | 43289724 | regular |
| ENSMUSG00000039801 | 2410089E | chr15 | + | 8169105 | 8271158 | regular |
| ENSMUSG00000039828 | Wdr70 | chr15 | - | 7873054 | 8099209 | regular |
| ENSMUSG00000039831 | Arhgap29 | chr3 | + | 121952540 | 122016753 | regular |
| ENSMUSG00000039842 | Mcph1 | chr8 | + | 18595130 | 18803189 | regular |
| ENSMUSG00000039844 | Rapgef1 | chr2 | + | 29619719 | 29740978 | regular |
| ENSMUSG00000039852 | Rere | chr4 | + | 150281645 | 150621966 | regular |
| ENSMUSG00000039886 | Tmem12C | chr5 | - | 135735484 | 135744448 | regular |
| ENSMUSG00000039901 | 9130011E | chr19 | - | 45818143 | 45998488 | regular |
| ENSMUSG00000039936 | Pik3cd | chr4 | - | 149649167 | 149702571 | regular |
| ENSMUSG00000039963 | Ccdc40 | chr11 | + | 119228571 | 119265236 | regular |
| ENSMUSG00000039977 | Deup1 | chr9 | - | 15559863 | 15627933 | regular |
| ENSMUSG00000039990 | Edrf1 | chr7 | + | 133637542 | 133672971 | regular |

| | | | | | | |
|---|---|---|---|---|---|---|
| ENSMUSG00000040003 | Magi2 | chr5 | + | 19227035 | 20704792 | regular |
| ENSMUSG00000040021 | Lats1 | chr10 | + | 7681213 | 7716460 | regular |
| ENSMUSG00000040022 | Rab11fip2 | chr19 | - | 59902883 | 59943654 | regular |
| ENSMUSG00000040026 | Saa3 | chr7 | - | 46711997 | 46715700 | regular |
| ENSMUSG00000040054 | Baz2a | chr10 | + | 128091576 | 128129303 | regular |
| ENSMUSG00000040084 | Bub1b | chr2 | + | 118598210 | 118641591 | regular |
| ENSMUSG00000040097 | Flywch1 | chr17 | - | 23752792 | 23771602 | regular |
| ENSMUSG00000040111 | Gramd1b | chr9 | - | 40293232 | 40531383 | regular |
| ENSMUSG00000040123 | Zmym5 | chr14 | - | 56790584 | 56811716 | regular |
| ENSMUSG00000040140 | Tdrd6 | chr17 | - | 43615334 | 43630299 | regular |
| ENSMUSG00000040152 | Thbs1 | chr2 | + | 118111875 | 118127133 | regular |
| ENSMUSG00000040163 | 1700034J( | chr6 | - | 146950328 | 146954422 | regular |
| ENSMUSG00000040225 | Prrc2c | chr1 | - | 162670724 | 162740556 | regular |
| ENSMUSG00000040242 | Fgfr1op2 | chr6 | + | 146577202 | 146599198 | regular |
| ENSMUSG00000040249 | Lrp1 | chr10 | - | 127538160 | 127621148 | regular |
| ENSMUSG00000040297 | Suco | chr1 | - | 161816113 | 161876682 | regular |
| ENSMUSG00000040312 | Cchcr1 | chr17 | + | 35517099 | 35531015 | regular |
| ENSMUSG00000040325 | Dcaf1 | chr9 | + | 106821873 | 106880992 | regular |
| ENSMUSG00000040351 | Ankib1 | chr5 | - | 3689999 | 3802925 | regular |
| ENSMUSG00000040356 | Skiv2l | chr17 | - | 34839227 | 34850210 | regular |
| ENSMUSG00000040365 | Trim41 | chr11 | - | 48806403 | 48817353 | regular |
| ENSMUSG00000040390 | Map3k10 | chr7 | - | 27656374 | 27674598 | regular |
| ENSMUSG00000040407 | Akap9 | chr5 | + | 3928053 | 4081310 | regular |
| ENSMUSG00000040412 | 5330417C | chr3 | - | 108455693 | 108536536 | regular |
| ENSMUSG00000040423 | Rc3h1 | chr1 | + | 160906417 | 160974978 | regular |
| ENSMUSG00000040451 | Sgms1 | chr19 | - | 32122726 | 32389714 | regular |
| ENSMUSG00000040473 | Cfap69 | chr5 | - | 5579277 | 5664239 | regular |
| ENSMUSG00000040481 | Bptf | chr11 | - | 107033080 | 107132127 | regular |
| ENSMUSG00000040506 | Ambra1 | chr2 | + | 91730133 | 91918849 | regular |
| ENSMUSG00000040524 | Zfp609 | chr9 | - | 65692390 | 65827564 | regular |
| ENSMUSG00000040548 | Tex2 | chr11 | - | 106502146 | 106613423 | regular |
| ENSMUSG00000040549 | Ckap5 | chr2 | + | 91526761 | 91620664 | regular |
| ENSMUSG00000040560 | Wdr7 | chr18 | + | 63708694 | 63989760 | regular |
| ENSMUSG00000040562 | Gstm2 | chr3 | - | 107981701 | 107986453 | regular |
| ENSMUSG00000040565 | Btaf1 | chr19 | + | 36926078 | 37012752 | regular |
| ENSMUSG00000040576 | Tmem27( | chr5 | - | 134901592 | 134906733 | regular |
| ENSMUSG00000040594 | Ranbp17 | chr11 | - | 33211794 | 33513746 | regular |
| ENSMUSG00000040629 | Mael | chr1 | - | 166201200 | 166238747 | regular |
| ENSMUSG00000040649 | Rimklb | chr6 | - | 122447295 | 122498761 | regular |
| ENSMUSG00000040651 | Fam208a | chr14 | + | 27428833 | 27483555 | regular |
| ENSMUSG00000040652 | Oaz2 | chr9 | + | 65668000 | 65690300 | regular |
| ENSMUSG00000040669 | Phc1 | chr6 | - | 122317730 | 122340561 | regular |
| ENSMUSG00000040681 | Hmgn1 | chr16 | - | 96120617 | 96127729 | regular |
| ENSMUSG00000040687 | Madd | chr2 | - | 91137359 | 91183837 | regular |
| ENSMUSG00000040690 | Col16a1 | chr4 | + | 130047839 | 130099283 | regular |
| ENSMUSG00000040699 | Limd2 | chr11 | - | 106156255 | 106160860 | regular |

| | | | | | | |
|---|---|---|---|---|---|---|
| ENSMUSG00000040725 | Hnrnpul1 | chr7 | - | 25721164 | 25754757 | regular |
| ENSMUSG00000040729 | Cep126 | chr9 | - | 8076460 | 8134294 | regular |
| ENSMUSG00000040731 | Eif4h | chr5 | - | 134619720 | 134639490 | regular |
| ENSMUSG00000040761 | Spen | chr4 | - | 141467889 | 141538597 | regular |
| ENSMUSG00000040782 | Rfwd2 | chr1 | + | 159232319 | 159347640 | regular |
| ENSMUSG00000040785 | Ttc3 | chr16 | + | 94370617 | 94522117 | regular |
| ENSMUSG00000040794 | C1qtnf4 | chr2 | + | 90885859 | 90890525 | regular |
| ENSMUSG00000040812 | Agbl2 | chr2 | + | 90782726 | 90834437 | regular |
| ENSMUSG00000040828 | Catsperd | chr17 | + | 56628142 | 56664456 | regular |
| ENSMUSG00000040829 | Zmynd15 | chr11 | + | 70459432 | 70466202 | regular |
| ENSMUSG00000040838 | Gm11639 | chr11 | + | 104685706 | 105117394 | regular |
| ENSMUSG00000040850 | Psme4 | chr11 | + | 30771725 | 30880361 | regular |
| ENSMUSG00000040852 | Plekhh2 | chr17 | + | 84511894 | 84622142 | regular |
| ENSMUSG00000040859 | Bsdc1 | chr4 | + | 129461580 | 129488498 | regular |
| ENSMUSG00000040865 | Ino80d | chr1 | - | 62958417 | 63114667 | regular |
| ENSMUSG00000040875 | Osbpl10 | chr9 | + | 114978568 | 115232225 | regular |
| ENSMUSG00000040913 | Fbxw4 | chr19 | - | 45578253 | 45660312 | regular |
| ENSMUSG00000040928 | S100pbp | chr4 | - | 129148004 | 129189727 | regular |
| ENSMUSG00000040929 | Rfx3 | chr19 | - | 27761720 | 28011166 | regular |
| ENSMUSG00000040936 | Ulk4 | chr9 | - | 120955350 | 121277197 | regular |
| ENSMUSG00000040940 | Arhgef1 | chr7 | + | 24902911 | 24926594 | regular |
| ENSMUSG00000040943 | Tet2 | chr3 | - | 133463678 | 133545139 | regular |
| ENSMUSG00000040952 | Rps19 | chr7 | + | 24884370 | 24889806 | regular |
| ENSMUSG00000041009 | Iqcf4 | chr9 | - | 106568318 | 106570996 | regular |
| ENSMUSG00000041028 | Ghitm | chr14 | - | 37120091 | 37135272 | regular |
| ENSMUSG00000041040 | Fam117b | chr1 | + | 59913005 | 59985346 | regular |
| ENSMUSG00000041112 | Elmo1 | chr13 | + | 20090595 | 20608353 | regular |
| ENSMUSG00000041144 | Dnah7b | chr1 | + | 46066314 | 46373546 | regular |
| ENSMUSG00000041161 | Otud3 | chr4 | - | 138895378 | 138913945 | regular |
| ENSMUSG00000041164 | Zmiz2 | chr11 | + | 6389073 | 6406158 | regular |
| ENSMUSG00000041165 | Spem1 | chr11 | - | 69820875 | 69822180 | regular |
| ENSMUSG00000041215 | Yeats2 | chr16 | + | 20141062 | 20232573 | regular |
| ENSMUSG00000041225 | Arhgap12 | chr18 | - | 6024426 | 6136098 | regular |
| ENSMUSG00000041229 | Phf8 | chrX | + | 151520671 | 151633859 | regular |
| ENSMUSG00000041235 | Chd7 | chr4 | + | 8690405 | 8867659 | regular |
| ENSMUSG00000041258 | Zfp236 | chr18 | - | 82593592 | 82692883 | regular |
| ENSMUSG00000041264 | Uspl1 | chr5 | + | 149184349 | 149215434 | regular |
| ENSMUSG00000041268 | Dmxl2 | chr9 | - | 54365157 | 54501626 | regular |
| ENSMUSG00000041297 | Cdk13 | chr13 | - | 17710314 | 17805097 | regular |
| ENSMUSG00000041301 | Cftr | chr6 | + | 18170686 | 18322768 | regular |
| ENSMUSG00000041323 | Ak7 | chr12 | + | 105705975 | 105782447 | regular |
| ENSMUSG00000041328 | Pcf11 | chr7 | - | 92643542 | 92669934 | regular |
| ENSMUSG00000041341 | Atg2b | chr12 | - | 105616135 | 105685211 | regular |
| ENSMUSG00000041343 | Ankrd42 | chr7 | - | 92581722 | 92637142 | regular |
| ENSMUSG00000041399 | 1700013G | chr4 | + | 137453283 | 137455461 | regular |
| ENSMUSG00000041408 | Wapl | chr14 | + | 34673927 | 34747983 | regular |

| | | | | | | |
|---|---|---|---|---|---|---|
| ENSMUSG00000041444 | Arhgap32 | chr9 | + | 32116135 | 32268446 | regular |
| ENSMUSG00000041453 | Rpl21 | chr5 | + | 146832889 | 146837032 | regular |
| ENSMUSG00000041560 | Nop53 | chr7 | - | 15936182 | 15946074 | regular |
| ENSMUSG00000041570 | Camsap2 | chr1 | - | 136268122 | 136346104 | regular |
| ENSMUSG00000041596 | Nlrp5-ps | chr7 | - | 14530651 | 14622479 | regular |
| ENSMUSG00000041629 | Fam104a | chr11 | - | 113661318 | 113684155 | regular |
| ENSMUSG00000041638 | Gcn1l1 | chr5 | + | 115565253 | 115622654 | regular |
| ENSMUSG00000041642 | Kif21b | chr1 | + | 136131388 | 136177998 | regular |
| ENSMUSG00000041645 | Ddx24 | chr12 | - | 103407981 | 103425830 | regular |
| ENSMUSG00000041650 | Pcca | chr14 | + | 122534323 | 122891100 | regular |
| ENSMUSG00000041673 | Lrrc18 | chr14 | + | 32991381 | 33015291 | regular |
| ENSMUSG00000041702 | Btbd7 | chr12 | - | 102780796 | 102878471 | regular |
| ENSMUSG00000041720 | Pi4ka | chr16 | - | 17280350 | 17406314 | regular |
| ENSMUSG00000041740 | Rnf10 | chr5 | - | 115241411 | 115272898 | regular |
| ENSMUSG00000041757 | Plekha6 | chr1 | + | 133164209 | 133303435 | regular |
| ENSMUSG00000041763 | Tpp2 | chr1 | + | 43933646 | 44003000 | regular |
| ENSMUSG00000041841 | Rpl37 | chr15 | + | 5116644 | 5119140 | regular |
| ENSMUSG00000041846 | Ppp4r3a | chr12 | - | 101039408 | 101083702 | regular |
| ENSMUSG00000041852 | Tcf20 | chr15 | - | 82808628 | 82912134 | regular |
| ENSMUSG00000041870 | Ankrd13a | chr5 | + | 114774676 | 114806200 | regular |
| ENSMUSG00000041890 | Git2 | chr5 | - | 114727406 | 114775517 | regular |
| ENSMUSG00000041895 | Wipi1 | chr11 | - | 109573330 | 109611967 | regular |
| ENSMUSG00000041915 | Ammecr1 | chr18 | + | 31759823 | 31784071 | regular |
| ENSMUSG00000041923 | Nol4 | chr18 | - | 22693180 | 23041653 | regular |
| ENSMUSG00000041945 | Mfsd9 | chr1 | - | 40772037 | 40790687 | regular |
| ENSMUSG00000041974 | Spidr | chr16 | - | 15889226 | 16146851 | regular |
| ENSMUSG00000042050 | Wdr60 | chr12 | - | 116206261 | 116263022 | regular |
| ENSMUSG00000042079 | Hnrnpf | chr6 | + | 117900323 | 117925622 | regular |
| ENSMUSG00000042156 | Dzip1 | chr14 | - | 118875519 | 118925460 | regular |
| ENSMUSG00000042197 | Zfp451 | chr1 | - | 33761544 | 33814595 | regular |
| ENSMUSG00000042207 | Kdm5b | chr1 | + | 134560170 | 134635285 | regular |
| ENSMUSG00000042213 | Zfand4 | chr6 | + | 116264221 | 116330302 | regular |
| ENSMUSG00000042228 | Lyn | chr4 | + | 3678114 | 3813122 | regular |
| ENSMUSG00000042246 | Tmc7 | chr7 | - | 118535840 | 118584736 | regular |
| ENSMUSG00000042293 | Gm5617 | chr9 | + | 48495441 | 48495972 | regular |
| ENSMUSG00000042298 | Ttc19 | chr11 | + | 62281472 | 62328451 | regular |
| ENSMUSG00000042302 | Ehbp1 | chr11 | - | 22005827 | 22342292 | regular |
| ENSMUSG00000042323 | Pbrm1 | chr14 | + | 31019137 | 31121592 | regular |
| ENSMUSG00000042331 | Specc1 | chr11 | + | 61956762 | 62223013 | regular |
| ENSMUSG00000042350 | Arel1 | chr12 | - | 84918147 | 84970900 | regular |
| ENSMUSG00000042354 | Gnl3 | chr14 | - | 31012432 | 31019152 | regular |
| ENSMUSG00000042390 | Gatad2b | chr3 | + | 90293177 | 90363407 | regular |
| ENSMUSG00000042404 | Dennd4b | chr3 | + | 90265184 | 90280669 | regular |
| ENSMUSG00000042408 | Zmym6 | chr4 | + | 127077382 | 127124372 | regular |
| ENSMUSG00000042446 | Zmym4 | chr4 | - | 126861938 | 126968135 | regular |
| ENSMUSG00000042507 | Elmsan1 | chr12 | - | 84149175 | 84218881 | regular |

| | | | | | | |
|---|---|---|---|---|---|---|
| ENSMUSG00000042520 | Ubap2l | chr3 | - | 90000139 | 90052628 | regular |
| ENSMUSG00000042541 | Sem1 | chr6 | - | 6557293 | 6578663 | regular |
| ENSMUSG00000042548 | Asxl1 | chr2 | + | 153345828 | 153404007 | regular |
| ENSMUSG00000042557 | Sin3a | chr9 | + | 57072039 | 57128366 | regular |
| ENSMUSG00000042567 | Nek10 | chr14 | + | 14803414 | 15012059 | regular |
| ENSMUSG00000042590 | Ipo11 | chr13 | - | 106794438 | 106936958 | regular |
| ENSMUSG00000042594 | Sh2b3 | chr5 | - | 121815487 | 121837646 | regular |
| ENSMUSG00000042599 | Kdm7a | chr6 | - | 39136622 | 39206789 | regular |
| ENSMUSG00000042605 | Atxn2 | chr5 | + | 121711336 | 121816493 | regular |
| ENSMUSG00000042606 | Hirip3 | chr7 | + | 126861971 | 126865377 | regular |
| ENSMUSG00000042608 | Stk40 | chr4 | + | 126103956 | 126141029 | regular |
| ENSMUSG00000042616 | Oscp1 | chr4 | + | 126058564 | 126092195 | regular |
| ENSMUSG00000042625 | Safb2 | chr17 | - | 56560964 | 56584585 | regular |
| ENSMUSG00000042628 | Zfyve1 | chr12 | - | 83546557 | 83597222 | regular |
| ENSMUSG00000042632 | Pla2g6 | chr15 | - | 79286227 | 79328390 | regular |
| ENSMUSG00000042644 | Itpr3 | chr17 | + | 27057303 | 27122223 | regular |
| ENSMUSG00000042662 | Dusp15 | chr2 | - | 152940996 | 152951698 | regular |
| ENSMUSG00000042678 | Myo15 | chr11 | + | 60469338 | 60528369 | regular |
| ENSMUSG00000042688 | Mapk6 | chr9 | - | 75369061 | 75410005 | regular |
| ENSMUSG00000042699 | Dhx9 | chr1 | - | 153455757 | 153487660 | regular |
| ENSMUSG00000042700 | Sipa1l1 | chr12 | + | 82169319 | 82451786 | regular |
| ENSMUSG00000042707 | Dnali1 | chr4 | - | 125055337 | 125065703 | regular |
| ENSMUSG00000042708 | Shcbp1l | chr1 | + | 153425161 | 153452574 | regular |
| ENSMUSG00000042726 | Trafd1 | chr5 | - | 121371724 | 121385632 | regular |
| ENSMUSG00000042744 | Hectd4 | chr5 | + | 121220218 | 121368577 | regular |
| ENSMUSG00000042772 | Smg7 | chr1 | - | 152836994 | 152902646 | regular |
| ENSMUSG00000042790 | Rnf214 | chr9 | - | 45863424 | 45906911 | regular |
| ENSMUSG00000042797 | Aqp11 | chr7 | - | 97724005 | 97738289 | regular |
| ENSMUSG00000042888 | Prr30 | chr14 | - | 101197689 | 101200330 | regular |
| ENSMUSG00000043003 | Rasef | chr4 | - | 73714578 | 73790994 | regular |
| ENSMUSG00000043036 | Ccdc63 | chr5 | - | 122108039 | 122140823 | regular |
| ENSMUSG00000043050 | Tnp2 | chr16 | - | 10787935 | 10788655 | regular |
| ENSMUSG00000043059 | Zfp513 | chr5 | - | 31198980 | 31202303 | regular |
| ENSMUSG00000043060 | Fscb | chr12 | - | 64471329 | 64474690 | regular |
| ENSMUSG00000043131 | Mob1a | chr6 | + | 83326015 | 83343776 | regular |
| ENSMUSG00000043154 | Ppp2r3a | chr9 | - | 101105083 | 101251795 | regular |
| ENSMUSG00000043241 | Upf2 | chr2 | + | 5951468 | 6056703 | regular |
| ENSMUSG00000043252 | Tmem64 | chr4 | + | 15265830 | 15286753 | regular |
| ENSMUSG00000043411 | Usp48 | chr4 | + | 137593754 | 137658537 | regular |
| ENSMUSG00000043445 | Pgp | chr17 | + | 24470472 | 24471591 | regular |
| ENSMUSG00000043535 | Setx | chr2 | + | 29124180 | 29182471 | regular |
| ENSMUSG00000043621 | Ubxn10 | chr4 | - | 138709836 | 138737167 | regular |
| ENSMUSG00000043629 | 1700019D | chr1 | - | 52922323 | 53020179 | regular |
| ENSMUSG00000043633 | Fam221b | chr4 | - | 43659621 | 43669145 | regular |
| ENSMUSG00000043716 | Rpl7 | chr1 | - | 16101294 | 16104662 | regular |
| ENSMUSG00000043866 | Taf10 | chr7 | - | 105739392 | 105744361 | regular |

| | | | | | | |
|---|---|---|---|---|---|---|
| ENSMUSG00000043909 | Trp53bp1 | chr2 | - | 121193280 | 121271407 | regular |
| ENSMUSG00000043913 | Ccdc60 | chr5 | - | 116124640 | 116288985 | regular |
| ENSMUSG00000043940 | Wdfy3 | chr5 | - | 101832955 | 102069921 | regular |
| ENSMUSG00000043962 | Thrap3 | chr4 | - | 126164081 | 126202760 | regular |
| ENSMUSG00000043987 | Cep164 | chr9 | - | 45766945 | 45828691 | regular |
| ENSMUSG00000044042 | Fmn1 | chr2 | + | 113327735 | 113716767 | regular |
| ENSMUSG00000044072 | Eml6 | chr11 | - | 29743047 | 30026033 | regular |
| ENSMUSG00000044083 | Efcab8 | chr2 | + | 153780878 | 153844752 | regular |
| ENSMUSG00000044084 | Spem2 | chr11 | - | 69816565 | 69818465 | regular |
| ENSMUSG00000044122 | Proca1 | chr11 | + | 78193391 | 78205763 | regular |
| ENSMUSG00000044176 | Spink10 | chr18 | + | 62548910 | 62661366 | regular |
| ENSMUSG00000044224 | Dnajc21 | chr15 | - | 10446755 | 10470516 | regular |
| ENSMUSG00000044252 | Osbpl1a | chr18 | - | 12755313 | 12941841 | regular |
| ENSMUSG00000044285 | Gm1821 | chr14 | + | 46084027 | 46085000 | regular |
| ENSMUSG00000044320 | 1700001C | chr2 | - | 30794768 | 30803661 | regular |
| ENSMUSG00000044340 | Phlpp1 | chr1 | + | 106171751 | 106394250 | regular |
| ENSMUSG00000044468 | Fam46c | chr3 | - | 100451627 | 100489324 | regular |
| ENSMUSG00000044471 | Lncpint | chr6 | - | 31093558 | 31218474 | regular |
| ENSMUSG00000044475 | Ascc1 | chr10 | + | 60002804 | 60099988 | regular |
| ENSMUSG00000044477 | Zfand3 | chr17 | + | 30005050 | 30210828 | regular |
| ENSMUSG00000044526 | Znrf4 | chr17 | - | 56511247 | 56512483 | regular |
| ENSMUSG00000044533 | Rps2 | chr17 | + | 24718115 | 24721929 | regular |
| ENSMUSG00000044566 | Cage1 | chr13 | - | 38006051 | 38037069 | regular |
| ENSMUSG00000044581 | 4932415D | chr10 | - | 82282115 | 82316582 | regular |
| ENSMUSG00000044627 | Swi5 | chr2 | - | 32278815 | 32288075 | regular |
| ENSMUSG00000044786 | Zfp36 | chr7 | - | 28376783 | 28380253 | regular |
| ENSMUSG00000044791 | Setd2 | chr9 | + | 110532596 | 110618633 | regular |
| ENSMUSG00000044792 | Isca1 | chr13 | - | 59755408 | 59769810 | regular |
| ENSMUSG00000044937 | Ttc41 | chr10 | + | 86705810 | 86776844 | regular |
| ENSMUSG00000044948 | Cfap43 | chr19 | - | 47737560 | 47919299 | regular |
| ENSMUSG00000044949 | Ubtd2 | chr11 | + | 32455369 | 32516687 | regular |
| ENSMUSG00000044950 | Pwwp2a | chr11 | + | 43681997 | 43721491 | regular |
| ENSMUSG00000045004 | Spata21 | chr4 | + | 141088332 | 141112760 | regular |
| ENSMUSG00000045036 | Tmem232 | chr17 | - | 65256004 | 65540782 | regular |
| ENSMUSG00000045078 | Rnf216 | chr5 | - | 142990892 | 143112994 | regular |
| ENSMUSG00000045107 | Saysd1 | chr14 | - | 20075635 | 20138534 | regular |
| ENSMUSG00000045128 | Rpl18a | chr8 | - | 70894721 | 70897412 | regular |
| ENSMUSG00000045210 | Vcpip1 | chr1 | - | 9718621 | 9748382 | regular |
| ENSMUSG00000045252 | Zfp574 | chr7 | + | 25072566 | 25083492 | regular |
| ENSMUSG00000045257 | Morn2 | chr17 | + | 80290202 | 80297476 | regular |
| ENSMUSG00000045275 | Lca5l | chr16 | - | 96158406 | 96192257 | regular |
| ENSMUSG00000045350 | Fam186a | chr15 | - | 99918347 | 99967075 | regular |
| ENSMUSG00000045467 | Ttll13 | chr7 | + | 80246375 | 80260821 | regular |
| ENSMUSG00000045482 | Trrap | chr5 | + | 144767731 | 144859778 | regular |
| ENSMUSG00000045521 | Tssk2 | chr16 | + | 17898636 | 17900023 | regular |
| ENSMUSG00000045709 | Smkr-ps | chr6 | + | 29985328 | 29993776 | regular |

| | | | | | | |
|---|---|---|---|---|---|---|
| ENSMUSG00000045962 | Wnk1 | chr6 | - | 119923968 | 120038672 | regular |
| ENSMUSG00000045983 | Eif4g1 | chr16 | + | 20668312 | 20692884 | regular |
| ENSMUSG00000046111 | Cep295 | chr9 | - | 15316914 | 15357788 | regular |
| ENSMUSG00000046173 | Pabpc6 | chr17 | - | 9666496 | 9669704 | regular |
| ENSMUSG00000046192 | Iqub | chr6 | - | 24444864 | 24515067 | regular |
| ENSMUSG00000046196 | Ttc39d | chr17 | + | 80207459 | 80217936 | regular |
| ENSMUSG00000046230 | Vps13a | chr19 | - | 16615365 | 16780933 | regular |
| ENSMUSG00000046330 | Rpl37a | chr1 | + | 72711289 | 72713813 | regular |
| ENSMUSG00000046337 | Fam178b | chr1 | - | 36562691 | 36683183 | regular |
| ENSMUSG00000046364 | Rpl27a | chr7 | + | 109519146 | 109522367 | regular |
| ENSMUSG00000046434 | Hnrnpa1 | chr15 | + | 103240431 | 103244955 | regular |
| ENSMUSG00000046585 | Cfap58 | chr19 | + | 47937711 | 48035379 | regular |
| ENSMUSG00000046727 | Cystm1 | chr18 | + | 36345618 | 36402980 | regular |
| ENSMUSG00000046755 | Kif2b | chr11 | - | 91575314 | 91577558 | regular |
| ENSMUSG00000046782 | Ttc6 | chr12 | + | 57564112 | 57737928 | regular |
| ENSMUSG00000046957 | Spz1 | chr13 | - | 92574635 | 92576172 | regular |
| ENSMUSG00000046991 | Wdr27 | chr17 | - | 14818671 | 14943158 | regular |
| ENSMUSG00000047021 | Cfap65 | chr1 | - | 74902070 | 74935599 | regular |
| ENSMUSG00000047126 | Cltc | chr11 | - | 86694350 | 86757565 | regular |
| ENSMUSG00000047139 | Cd24a | chr10 | + | 43578283 | 43584265 | regular |
| ENSMUSG00000047141 | Zfp654 | chr16 | - | 64780346 | 64851652 | regular |
| ENSMUSG00000047150 | 1700001C | chr17 | - | 47412733 | 47437376 | regular |
| ENSMUSG00000047153 | Khnyn | chr14 | + | 55884946 | 55898775 | regular |
| ENSMUSG00000047193 | Dync2h1 | chr9 | - | 6928502 | 7184446 | regular |
| ENSMUSG00000047215 | Rpl9 | chr5 | - | 65388363 | 65391444 | regular |
| ENSMUSG00000047248 | C2cd3 | chr7 | + | 100372232 | 100470152 | regular |
| ENSMUSG00000047284 | Neurl4 | chr11 | + | 69901071 | 69913820 | regular |
| ENSMUSG00000047361 | Gm973 | chr1 | + | 59516263 | 59636417 | regular |
| ENSMUSG00000047369 | Dnah14 | chr1 | + | 181576558 | 181815774 | regular |
| ENSMUSG00000047388 | Atmin | chr8 | + | 116943392 | 116960445 | regular |
| ENSMUSG00000047412 | Zbtb44 | chr9 | + | 31030643 | 31075885 | regular |
| ENSMUSG00000047417 | Rexo1 | chr10 | - | 80540921 | 80561560 | regular |
| ENSMUSG00000047441 | Antxrl | chr14 | + | 34052706 | 34076405 | regular |
| ENSMUSG00000047446 | Arl4a | chr12 | - | 40005446 | 40038025 | regular |
| ENSMUSG00000047454 | Gphn | chr12 | + | 78226378 | 78684772 | regular |
| ENSMUSG00000047502 | Mroh7 | chr4 | - | 106680416 | 106730925 | regular |
| ENSMUSG00000047518 | Slfnl1 | chr4 | + | 120532230 | 120536661 | regular |
| ENSMUSG00000047528 | Als2cr12 | chr1 | - | 58646902 | 58695989 | regular |
| ENSMUSG00000047648 | Fbxo30 | chr10 | + | 11281329 | 11298052 | regular |
| ENSMUSG00000047654 | Tssk6 | chr8 | + | 69902183 | 69903518 | regular |
| ENSMUSG00000047675 | Rps8 | chr4 | - | 117153826 | 117156243 | regular |
| ENSMUSG00000047720 | 4922502D | chr6 | - | 129322163 | 129331796 | regular |
| ENSMUSG00000047866 | Lonp2 | chr8 | + | 86624042 | 86723873 | regular |
| ENSMUSG00000047867 | Gimap6 | chr6 | - | 48701581 | 48708225 | regular |
| ENSMUSG00000047888 | Tnrc6b | chr15 | + | 80711312 | 80941085 | regular |
| ENSMUSG00000047921 | Trappc9 | chr15 | - | 72589620 | 73061204 | regular |

| | | | | | | |
|---|---|---|---|---|---|---|
| ENSMUSG00000047940 | Stpg2 | chr3 | + | 139205693 | 139710299 | regular |
| ENSMUSG00000048000 | Gigyf2 | chr1 | + | 87326997 | 87450796 | regular |
| ENSMUSG00000048039 | Isg20l2 | chr3 | + | 87930313 | 87940686 | regular |
| ENSMUSG00000048076 | Arf1 | chr11 | - | 59211411 | 59228270 | regular |
| ENSMUSG00000048077 | H1fnt | chr15 | - | 98255981 | 98257307 | regular |
| ENSMUSG00000048154 | Kmt2d | chr15 | - | 98831668 | 98871204 | regular |
| ENSMUSG00000048206 | Dnajb8 | chr6 | + | 88222267 | 88254415 | regular |
| ENSMUSG00000048249 | Crebrf | chr17 | + | 26715649 | 26776635 | regular |
| ENSMUSG00000048264 | Dip2c | chr13 | + | 9276527 | 9668928 | regular |
| ENSMUSG00000048271 | Rbm33 | chr5 | + | 28317120 | 28419239 | regular |
| ENSMUSG00000048410 | Zfp407 | chr18 | - | 84128026 | 84589725 | regular |
| ENSMUSG00000048416 | Mlf1 | chr3 | + | 67374096 | 67400003 | regular |
| ENSMUSG00000048478 | Spata33 | chr8 | + | 123212831 | 123222046 | regular |
| ENSMUSG00000048520 | Fbxl13 | chr5 | - | 21483846 | 21645634 | regular |
| ENSMUSG00000048578 | Mlec | chr5 | - | 115142980 | 115158179 | regular |
| ENSMUSG00000048602 | Morc2b | chr17 | - | 33135587 | 33139683 | regular |
| ENSMUSG00000048668 | Rhno1 | chr6 | - | 128356999 | 128362911 | regular |
| ENSMUSG00000048686 | Hmgb4 | chr4 | - | 128260211 | 128260957 | regular |
| ENSMUSG00000048696 | Mex3d | chr10 | - | 80380354 | 80387659 | regular |
| ENSMUSG00000048701 | Ccdc6 | chr10 | + | 70097120 | 70193200 | regular |
| ENSMUSG00000048720 | Tbc1d12 | chr19 | + | 38836578 | 38919923 | regular |
| ENSMUSG00000048731 | Ggnbp1 | chr17 | + | 26973216 | 27036378 | regular |
| ENSMUSG00000048758 | Rpl29 | chr9 | + | 106429453 | 106431568 | regular |
| ENSMUSG00000048772 | Tmem53 | chr4 | + | 117251950 | 117268582 | regular |
| ENSMUSG00000048799 | Cep120 | chr18 | - | 53681723 | 53744547 | regular |
| ENSMUSG00000048827 | Pkd1l3 | chr8 | + | 109614516 | 109674386 | regular |
| ENSMUSG00000048874 | Phf3 | chr1 | - | 30802338 | 30873921 | regular |
| ENSMUSG00000049008 | BB01443 | chr8 | - | 15041443 | 15046078 | regular |
| ENSMUSG00000049106 | Dcaf5 | chr12 | - | 80335847 | 80436601 | regular |
| ENSMUSG00000049123 | Catsperg2 | chr7 | - | 29697218 | 29727032 | regular |
| ENSMUSG00000049252 | Lrp1b | chr2 | - | 40595245 | 42653624 | regular |
| ENSMUSG00000049295 | Zfp219 | chr14 | - | 52006076 | 52020733 | regular |
| ENSMUSG00000049339 | Retreg2 | chr1 | + | 75142777 | 75147913 | regular |
| ENSMUSG00000049353 | Rd3 | chr1 | + | 191977369 | 191988283 | regular |
| ENSMUSG00000049354 | Dcaf7 | chr11 | + | 106036871 | 106059324 | regular |
| ENSMUSG00000049470 | Aff4 | chr11 | + | 53350832 | 53421830 | regular |
| ENSMUSG00000049482 | Ctu2 | chr8 | + | 122476142 | 122484138 | regular |
| ENSMUSG00000049506 | Sppl2c | chr11 | + | 104186326 | 104191163 | regular |
| ENSMUSG00000049517 | Rps23 | chr13 | + | 90922957 | 90924950 | regular |
| ENSMUSG00000049550 | Clip1 | chr5 | - | 123577794 | 123684618 | regular |
| ENSMUSG00000049571 | Cfap46 | chr7 | - | 139600950 | 139683817 | regular |
| ENSMUSG00000049606 | Zfp644 | chr5 | - | 106616738 | 106697287 | regular |
| ENSMUSG00000049653 | Spatc1 | chr15 | + | 76268088 | 76292572 | regular |
| ENSMUSG00000049659 | Aftph | chr11 | - | 20685083 | 20741589 | regular |
| ENSMUSG00000049670 | Morn4 | chr19 | - | 42074938 | 42086370 | regular |
| ENSMUSG00000049676 | Catsperg1 | chr7 | - | 29181320 | 29214035 | regular |

| | | | | | | |
|---|---|---|---|---|---|---|
| ENSMUSG00000049739 | Zfp646 | chr7 | + | 127876220 | 127885996 | regular |
| ENSMUSG00000049744 | Arhgap15 | chr2 | + | 43748823 | 44395953 | regular |
| ENSMUSG00000049760 | 2410015N | chr17 | - | 56607451 | 56609771 | regular |
| ENSMUSG00000049775 | Tmsb4x | chrX | - | 167207092 | 167209315 | regular |
| ENSMUSG00000049792 | Bag5 | chr12 | - | 111709487 | 111713257 | regular |
| ENSMUSG00000049811 | Fam161a | chr11 | + | 23007530 | 23030788 | regular |
| ENSMUSG00000049866 | Arl4c | chr1 | - | 88673124 | 88702221 | regular |
| ENSMUSG00000049916 | 2610318N | chr16 | - | 17113398 | 17125140 | regular |
| ENSMUSG00000049971 | Glt1d1 | chr5 | + | 127632261 | 127709374 | regular |
| ENSMUSG00000050017 | Pitpnb | chr5 | + | 111330762 | 111388359 | regular |
| ENSMUSG00000050035 | Fhl4 | chr10 | - | 85096286 | 85102495 | regular |
| ENSMUSG00000050050 | Ccdc158 | chr5 | - | 92607953 | 92675271 | regular |
| ENSMUSG00000050058 | Prm3 | chr16 | - | 10790504 | 10790914 | regular |
| ENSMUSG00000050074 | Spink8 | chr9 | + | 109812279 | 109826633 | regular |
| ENSMUSG00000050089 | Akap4 | chrX | + | 7067518 | 7078609 | regular |
| ENSMUSG00000050107 | Haspin | chr11 | - | 73135484 | 73138294 | regular |
| ENSMUSG00000050122 | Vwa3b | chr1 | + | 37026595 | 37187613 | regular |
| ENSMUSG00000050299 | Gm9843 | chr16 | - | 76403208 | 76403712 | regular |
| ENSMUSG00000050471 | Fam118b | chr9 | - | 35216964 | 35267805 | regular |
| ENSMUSG00000050553 | Gk2 | chr5 | - | 97455141 | 97457021 | regular |
| ENSMUSG00000050565 | Tor1aip2 | chr1 | + | 156035402 | 156068861 | regular |
| ENSMUSG00000050612 | Txndc2 | chr17 | - | 65637504 | 65642204 | regular |
| ENSMUSG00000050623 | Catsperz | chr19 | - | 6922425 | 6925380 | regular |
| ENSMUSG00000050677 | Ccdc96 | chr5 | + | 36484587 | 36488172 | regular |
| ENSMUSG00000050702 | 4930563N | chr9 | - | 55962588 | 56010550 | regular |
| ENSMUSG00000050708 | Ftl1 | chr7 | - | 45457943 | 45459884 | regular |
| ENSMUSG00000050721 | Plekho2 | chr9 | - | 65552697 | 65580040 | regular |
| ENSMUSG00000050821 | Fam131a | chr16 | + | 20693281 | 20703048 | regular |
| ENSMUSG00000050860 | Phospho1 | chr11 | + | 95824498 | 95832140 | regular |
| ENSMUSG00000050944 | Efcab5 | chr11 | - | 77089914 | 77188968 | regular |
| ENSMUSG00000050994 | Adgb | chr10 | - | 10335702 | 10472326 | regular |
| ENSMUSG00000050996 | Cetn1 | chr18 | - | 9615523 | 9619478 | regular |
| ENSMUSG00000051113 | Fam71e1 | chr7 | + | 44496580 | 44501486 | regular |
| ENSMUSG00000051223 | Bzw1 | chr1 | + | 58392897 | 58407353 | regular |
| ENSMUSG00000051236 | Msrb3 | chr10 | - | 120781095 | 120899101 | regular |
| ENSMUSG00000051276 | Actrt2 | chr4 | - | 154666432 | 154667867 | regular |
| ENSMUSG00000051306 | Usp42 | chr5 | - | 143710324 | 143732280 | regular |
| ENSMUSG00000051329 | Nup160 | chr2 | + | 90677214 | 90736328 | regular |
| ENSMUSG00000051343 | Rab11fip5 | chr6 | - | 85334961 | 85374634 | regular |
| ENSMUSG00000051427 | Ccdc157 | chr11 | - | 4141122 | 4160293 | regular |
| ENSMUSG00000051435 | Fhad1 | chr4 | - | 141890437 | 142015082 | regular |
| ENSMUSG00000051437 | Ubqlnl | chr7 | - | 104148258 | 104150556 | regular |
| ENSMUSG00000051495 | Irf2bp2 | chr8 | - | 126588295 | 126593986 | regular |
| ENSMUSG00000051506 | Wdfy4 | chr14 | - | 32959546 | 33185508 | regular |
| ENSMUSG00000051586 | Mical3 | chr6 | - | 120931706 | 121130999 | regular |
| ENSMUSG00000051618 | Ubqln3 | chr7 | - | 104140622 | 104143279 | regular |

| | | | | | | |
|---|---|---|---|---|---|---|
| ENSMUSG00000051648 | Kctd19 | chr8 | - | 105382806 | 105413502 | regular |
| ENSMUSG00000051674 | Dcun1d4 | chr5 | + | 73480999 | 73560794 | regular |
| ENSMUSG00000051728 | 4930563D | chr16 | - | 92318762 | 92321408 | regular |
| ENSMUSG00000051732 | Pabpc2 | chr18 | + | 39773496 | 39776082 | regular |
| ENSMUSG00000051736 | Fam229b | chr10 | - | 39118803 | 39133914 | regular |
| ENSMUSG00000051747 | Ttn | chr2 | - | 76703979 | 76982547 | regular |
| ENSMUSG00000051900 | Abca16 | chr7 | + | 120409646 | 120544813 | regular |
| ENSMUSG00000051910 | Sox6 | chr7 | - | 115470871 | 116038796 | regular |
| ENSMUSG00000051934 | Spats2 | chr15 | + | 99126577 | 99213206 | regular |
| ENSMUSG00000052056 | Zfp217 | chr2 | - | 170108642 | 170148103 | regular |
| ENSMUSG00000052075 | 1700029F | chr13 | - | 97021863 | 97034362 | regular |
| ENSMUSG00000052139 | Babam2 | chr5 | + | 31697683 | 32084962 | regular |
| ENSMUSG00000052146 | Rps10 | chr17 | - | 27630421 | 27636629 | regular |
| ENSMUSG00000052221 | Ppp1r36 | chr12 | + | 76417537 | 76439492 | regular |
| ENSMUSG00000052273 | Dnah3 | chr7 | - | 119922670 | 120095280 | regular |
| ENSMUSG00000052296 | Ppp6r1 | chr7 | - | 4631494 | 4659019 | regular |
| ENSMUSG00000052299 | Ltn1 | chr16 | - | 87376650 | 87432606 | regular |
| ENSMUSG00000052305 | Hbb-bs | chr7 | - | 103826533 | 103828096 | regular |
| ENSMUSG00000052337 | Immt | chr6 | + | 71831330 | 71877388 | regular |
| ENSMUSG00000052384 | Nrros | chr16 | - | 32142784 | 32165594 | regular |
| ENSMUSG00000052397 | Ezr | chr17 | - | 6738040 | 6782784 | regular |
| ENSMUSG00000052407 | Ccdc171 | chr4 | + | 83525544 | 83864670 | regular |
| ENSMUSG00000052459 | Atp6v1a | chr16 | - | 44085401 | 44139705 | regular |
| ENSMUSG00000052469 | Tcp10c | chr17 | + | 13354571 | 13377223 | regular |
| ENSMUSG00000052512 | Nav2 | chr7 | + | 48908715 | 49610090 | regular |
| ENSMUSG00000052533 | Nup188 | chr2 | + | 30286396 | 30344266 | regular |
| ENSMUSG00000052681 | Rap1b | chr10 | - | 117813870 | 117846035 | regular |
| ENSMUSG00000052698 | Tln2 | chr9 | - | 67217086 | 67559703 | regular |
| ENSMUSG00000052707 | Tnrc6a | chr7 | + | 123123884 | 123195296 | regular |
| ENSMUSG00000052783 | Grk4 | chr5 | + | 34660378 | 34755305 | regular |
| ENSMUSG00000052837 | Junb | chr8 | - | 84974483 | 84978718 | regular |
| ENSMUSG00000052861 | Dnah6 | chr6 | - | 73017605 | 73221651 | regular |
| ENSMUSG00000052889 | Prkcb | chr7 | + | 122288750 | 122634402 | regular |
| ENSMUSG00000052915 | Msl1 | chr11 | + | 98795515 | 98807859 | regular |
| ENSMUSG00000052926 | Rnaseh2a | chr8 | - | 84956609 | 84969767 | regular |
| ENSMUSG00000052955 | Cpvl | chr6 | - | 53873278 | 53978671 | regular |
| ENSMUSG00000052997 | Uba2 | chr7 | - | 34140687 | 34169599 | regular |
| ENSMUSG00000053111 | Fank1 | chr7 | + | 133776860 | 133881532 | regular |
| ENSMUSG00000053153 | Spag16 | chr1 | + | 69826969 | 70725132 | regular |
| ENSMUSG00000053192 | Mllt11 | chr3 | - | 95219134 | 95232171 | regular |
| ENSMUSG00000053253 | Ndfip2 | chr14 | + | 105258572 | 105309298 | regular |
| ENSMUSG00000053279 | Aldh1a1 | chr19 | + | 20492714 | 20643462 | regular |
| ENSMUSG00000053293 | Pom121 | chr5 | - | 135376140 | 135394546 | regular |
| ENSMUSG00000053332 | Gas5 | chr1 | + | 161034421 | 161038539 | regular |
| ENSMUSG00000053333 | Dis3l2 | chr1 | + | 86703807 | 87050095 | regular |
| ENSMUSG00000053470 | Kdm3a | chr6 | - | 71588971 | 71632990 | regular |

| | | | | | | |
|---|---|---|---|---|---|---|
| ENSMUSG00000053510 | Nrd1 | chr4 | + | 109000654 | 109061777 | regular |
| ENSMUSG00000053565 | Eif3k | chr7 | - | 28971371 | 28981864 | regular |
| ENSMUSG00000053624 | Gykl1 | chr18 | + | 52693678 | 52695668 | regular |
| ENSMUSG00000053641 | Dennd4a | chr9 | + | 64811339 | 64919667 | regular |
| ENSMUSG00000053754 | Chd8 | chr14 | - | 52198150 | 52257780 | regular |
| ENSMUSG00000053774 | Ubxn7 | chr16 | + | 32332251 | 32393747 | regular |
| ENSMUSG00000053819 | Camk2d | chr3 | + | 126596301 | 126846326 | regular |
| ENSMUSG00000053838 | Nudcd3 | chr11 | - | 6105690 | 6200415 | regular |
| ENSMUSG00000053896 | 4933409G | chr2 | + | 68582412 | 68616387 | regular |
| ENSMUSG00000053907 | Mat2a | chr6 | - | 72432798 | 72439558 | regular |
| ENSMUSG00000053929 | Cyhr1 | chr15 | - | 76643394 | 76660117 | regular |
| ENSMUSG00000054003 | Tdrd9 | chr12 | + | 111971558 | 112068854 | regular |
| ENSMUSG00000054199 | Gon4l | chr3 | + | 88835230 | 88910103 | regular |
| ENSMUSG00000054277 | Arfgap3 | chr15 | - | 83299738 | 83350247 | regular |
| ENSMUSG00000054280 | Prr14l | chr5 | - | 32789819 | 32854256 | regular |
| ENSMUSG00000054387 | Mdm4 | chr1 | - | 132959483 | 133030561 | regular |
| ENSMUSG00000054452 | Aes | chr10 | + | 81559487 | 81566362 | regular |
| ENSMUSG00000054611 | Kdm2a | chr19 | - | 4314418 | 4398285 | regular |
| ENSMUSG00000054823 | Nsd3 | chr8 | + | 25601600 | 25719667 | regular |
| ENSMUSG00000054901 | Arhgef33 | chr17 | + | 80307406 | 80400081 | regular |
| ENSMUSG00000054909 | Wbscr25 | chr5 | - | 134987432 | 135001350 | regular |
| ENSMUSG00000055013 | Agap1 | chr1 | + | 89454805 | 89897617 | regular |
| ENSMUSG00000055024 | Ep300 | chr15 | + | 81585350 | 81652077 | regular |
| ENSMUSG00000055065 | Ddx17 | chr15 | - | 79527706 | 79546741 | regular |
| ENSMUSG00000055137 | Sugct | chr13 | - | 16857471 | 17695553 | regular |
| ENSMUSG00000055204 | Ankrd17 | chr5 | - | 90227165 | 90366577 | regular |
| ENSMUSG00000055239 | Kcmf1 | chr6 | - | 72841113 | 72899979 | regular |
| ENSMUSG00000055302 | Mrfap1 | chr5 | - | 36794866 | 36796772 | regular |
| ENSMUSG00000055319 | Sec23ip | chr7 | + | 128744942 | 128784836 | regular |
| ENSMUSG00000055322 | Tns1 | chr1 | - | 73910230 | 74124449 | regular |
| ENSMUSG00000055436 | Srsf11 | chr3 | - | 158010472 | 158036639 | regular |
| ENSMUSG00000055531 | Cpsf6 | chr10 | - | 117344672 | 117380015 | regular |
| ENSMUSG00000055602 | Tcp10b | chr17 | + | 13061103 | 13082481 | regular |
| ENSMUSG00000055670 | Zzef1 | chr11 | + | 72796225 | 72927120 | regular |
| ENSMUSG00000055692 | Tmem191 | chr16 | + | 17276308 | 17283229 | regular |
| ENSMUSG00000055720 | Ubl7 | chr9 | + | 57910978 | 57929968 | regular |
| ENSMUSG00000055762 | Eef1d | chr15 | - | 75894795 | 75909556 | regular |
| ENSMUSG00000055839 | Elob | chr17 | - | 23824739 | 23829109 | regular |
| ENSMUSG00000055850 | Rnf181 | chr6 | - | 72359713 | 72366956 | regular |
| ENSMUSG00000055891 | Ubl4b | chr3 | - | 107553686 | 107555073 | regular |
| ENSMUSG00000055932 | Fto | chr8 | + | 91313524 | 91668439 | regular |
| ENSMUSG00000056018 | Ccdc7b | chr8 | + | 129067133 | 129183732 | regular |
| ENSMUSG00000056050 | Mia3 | chr1 | - | 183326724 | 183369553 | regular |
| ENSMUSG00000056076 | Eif3b | chr5 | + | 140419327 | 140443360 | regular |
| ENSMUSG00000056201 | Cfl1 | chr19 | + | 5490454 | 5495201 | regular |
| ENSMUSG00000056211 | R3hdm1 | chr1 | + | 128103300 | 128237736 | regular |

| | | | | | | |
|---|---|---|---|---|---|---|
| ENSMUSG00000056234 | Ncoa4 | chr14 | + | 32159864 | 32179855 | regular |
| ENSMUSG00000056267 | Cep70 | chr9 | + | 99243366 | 99300404 | regular |
| ENSMUSG00000056268 | Dennd1b | chr1 | + | 138963434 | 139178960 | regular |
| ENSMUSG00000056342 | Usp34 | chr11 | + | 23306894 | 23490560 | regular |
| ENSMUSG00000056429 | Tgoln1 | chr6 | - | 72608431 | 72617000 | regular |
| ENSMUSG00000056458 | Mok | chr12 | - | 110807797 | 110840939 | regular |
| ENSMUSG00000056508 | 1700001K | chr12 | - | 110667687 | 110682619 | regular |
| ENSMUSG00000056509 | Gm9999 | chr7 | + | 46983579 | 46987803 | regular |
| ENSMUSG00000056536 | Pign | chr1 | - | 105518421 | 105663677 | regular |
| ENSMUSG00000056537 | Rlim | chrX | - | 103957162 | 103981284 | regular |
| ENSMUSG00000056579 | Tug1 | chr11 | - | 3639784 | 3649673 | regular |
| ENSMUSG00000056602 | Fry | chr5 | + | 150118644 | 150497753 | regular |
| ENSMUSG00000056608 | Chd9 | chr8 | + | 90828351 | 91054516 | regular |
| ENSMUSG00000056692 | D17Wsu9 | chr17 | - | 27751231 | 27820558 | regular |
| ENSMUSG00000056763 | Cspp1 | chr1 | + | 10037986 | 10136768 | regular |
| ENSMUSG00000056770 | Setd3 | chr12 | - | 108106430 | 108179314 | regular |
| ENSMUSG00000056851 | Pcbp2 | chr15 | + | 102470631 | 102500059 | regular |
| ENSMUSG00000056899 | Immp2l | chr12 | + | 41024089 | 41955588 | regular |
| ENSMUSG00000056962 | Jmjd6 | chr11 | - | 116837431 | 116843449 | regular |
| ENSMUSG00000056987 | Fam71d | chr12 | + | 78691534 | 78734516 | regular |
| ENSMUSG00000056999 | Ide | chr19 | - | 37268742 | 37337852 | regular |
| ENSMUSG00000057069 | Ero1lb | chr13 | + | 12565873 | 12611396 | regular |
| ENSMUSG00000057098 | Ebf1 | chr11 | + | 44617316 | 45008091 | regular |
| ENSMUSG00000057110 | Cntrl | chr2 | + | 35109491 | 35178822 | regular |
| ENSMUSG00000057113 | Npm1 | chr11 | - | 33152286 | 33163206 | regular |
| ENSMUSG00000057132 | Rpgrip1 | chr14 | + | 52110703 | 52163546 | regular |
| ENSMUSG00000057176 | Ccdc189 | chr7 | - | 127582382 | 127588595 | regular |
| ENSMUSG00000057230 | Aak1 | chr6 | + | 86849516 | 87003223 | regular |
| ENSMUSG00000057236 | Rbbp4 | chr4 | - | 129307099 | 129335370 | regular |
| ENSMUSG00000057240 | Ms4a13 | chr19 | - | 11169417 | 11196737 | regular |
| ENSMUSG00000057246 | BC051142 | chr17 | + | 34398819 | 34460734 | regular |
| ENSMUSG00000057265 | Bbof1 | chr12 | + | 84409070 | 84442887 | regular |
| ENSMUSG00000057286 | St6galnac | chr11 | - | 116675302 | 116694868 | regular |
| ENSMUSG00000057322 | Rpl38 | chr11 | + | 114668523 | 114672331 | regular |
| ENSMUSG00000057335 | Cep170 | chr1 | - | 176733652 | 176814067 | regular |
| ENSMUSG00000057406 | Nsd2 | chr5 | + | 33820724 | 33897975 | regular |
| ENSMUSG00000057440 | Mpp7 | chr18 | - | 7347961 | 7626863 | regular |
| ENSMUSG00000057531 | Dtnbp1 | chr13 | - | 44922074 | 45002147 | regular |
| ENSMUSG00000057551 | Zfp317 | chr9 | + | 19622101 | 19649731 | regular |
| ENSMUSG00000057637 | Prdm2 | chr4 | - | 143107390 | 143212995 | regular |
| ENSMUSG00000057672 | Pkn1 | chr8 | - | 83666535 | 83699179 | regular |
| ENSMUSG00000057719 | Sh3rf2 | chr18 | + | 42053666 | 42158960 | regular |
| ENSMUSG00000057738 | Sptan1 | chr2 | + | 29965559 | 30031451 | regular |
| ENSMUSG00000057841 | Rpl32 | chr6 | - | 115805504 | 115808747 | regular |
| ENSMUSG00000057863 | Rpl36 | chr17 | + | 56613394 | 56614243 | regular |
| ENSMUSG00000058006 | Mdn1 | chr4 | + | 32657118 | 32775217 | regular |

| | | | | | | |
|---|---|---|---|---|---|---|
| ENSMUSG00000058013 | Sep.11 | chr5 | + | 93093436 | 93176447 | regular |
| ENSMUSG00000058019 | Ces5a | chr8 | - | 93499063 | 93535830 | regular |
| ENSMUSG00000058135 | Gstm1 | chr3 | - | 108012254 | 108017973 | regular |
| ENSMUSG00000058183 | Mmel1 | chr4 | + | 154869584 | 154895528 | regular |
| ENSMUSG00000058230 | Arhgap35 | chr7 | - | 16493718 | 16614993 | regular |
| ENSMUSG00000058290 | Espl1 | chr15 | + | 102296292 | 102324356 | regular |
| ENSMUSG00000058318 | Phf21a | chr2 | + | 92093116 | 92364666 | regular |
| ENSMUSG00000058443 | Rpl10-ps3 | chr9 | - | 50344200 | 50344968 | regular |
| ENSMUSG00000058546 | Rpl23a | chr11 | - | 78180931 | 78183584 | regular |
| ENSMUSG00000058558 | Rpl5 | chr5 | + | 107900501 | 107909005 | regular |
| ENSMUSG00000058587 | Tmod3 | chr9 | - | 75497706 | 75559657 | regular |
| ENSMUSG00000058589 | Anks1b | chr10 | + | 89873508 | 90973300 | regular |
| ENSMUSG00000058655 | Eif4b | chr15 | + | 102073772 | 102097172 | regular |
| ENSMUSG00000058690 | Ccser2 | chr14 | - | 36874935 | 36968777 | regular |
| ENSMUSG00000058704 | Memo1 | chr17 | - | 74200705 | 74294883 | regular |
| ENSMUSG00000058709 | Egln2 | chr7 | - | 27158657 | 27166802 | regular |
| ENSMUSG00000058761 | Rnf169 | chr7 | - | 99920253 | 99980448 | regular |
| ENSMUSG00000058799 | Nap1l1 | chr10 | + | 111473222 | 111498150 | regular |
| ENSMUSG00000058881 | Zfp516 | chr18 | + | 82910878 | 83005314 | regular |
| ENSMUSG00000058997 | Vwa8 | chr14 | + | 78849051 | 79202310 | regular |
| ENSMUSG00000059040 | Eno1b | chr18 | + | 48045334 | 48048378 | regular |
| ENSMUSG00000059070 | Rpl18 | chr7 | + | 45715456 | 45720836 | regular |
| ENSMUSG00000059114 | Lrrc74a | chr12 | + | 86734368 | 86763797 | regular |
| ENSMUSG00000059119 | Nap1l4 | chr7 | - | 143513578 | 143549106 | regular |
| ENSMUSG00000059208 | Hnrnpm | chr17 | - | 33646232 | 33686860 | regular |
| ENSMUSG00000059263 | Usp47 | chr7 | + | 112023503 | 112111661 | regular |
| ENSMUSG00000059273 | Zc3h4 | chr7 | + | 16400909 | 16437696 | regular |
| ENSMUSG00000059288 | Cdyl | chr13 | + | 35659832 | 35874063 | regular |
| ENSMUSG00000059291 | Rpl11 | chr4 | - | 136028264 | 136053428 | regular |
| ENSMUSG00000059343 | Aldoart1 | chr4 | - | 72850582 | 72852634 | regular |
| ENSMUSG00000059430 | Actg2 | chr6 | - | 83512904 | 83536265 | regular |
| ENSMUSG00000059439 | Bcas3 | chr11 | + | 85353166 | 85826058 | regular |
| ENSMUSG00000059456 | Ptk2b | chr14 | - | 66153256 | 66281052 | regular |
| ENSMUSG00000059474 | Mbtd1 | chr11 | + | 93885851 | 93946985 | regular |
| ENSMUSG00000059495 | Arhgef12 | chr9 | - | 42963841 | 43107239 | regular |
| ENSMUSG00000059540 | Tcea2 | chr2 | + | 181680309 | 181688071 | regular |
| ENSMUSG00000059562 | Ccdc154 | chr17 | + | 25162460 | 25171913 | regular |
| ENSMUSG00000059796 | Eif4a1 | chr11 | - | 69666935 | 69672423 | regular |
| ENSMUSG00000059834 | Sclt1 | chr3 | - | 41626719 | 41742514 | regular |
| ENSMUSG00000059854 | Hydin | chr8 | + | 110266976 | 110610253 | regular |
| ENSMUSG00000059891 | Tsks | chr7 | + | 44943203 | 44958035 | regular |
| ENSMUSG00000059895 | Ptp4a3 | chr15 | + | 73723144 | 73758766 | regular |
| ENSMUSG00000059920 | 4930453N | chr16 | - | 64751750 | 64771161 | regular |
| ENSMUSG00000059970 | Hspa2 | chr12 | + | 76404175 | 76406938 | regular |
| ENSMUSG00000060002 | Chpt1 | chr10 | - | 88452744 | 88504073 | regular |
| ENSMUSG00000060036 | Rpl3 | chr15 | - | 80077793 | 80091868 | regular |

| | | | | | | |
|---|---|---|---|---|---|---|
| ENSMUSG00000060098 | Prmt7 | chr8 | + | 106210935 | 106252794 | regular |
| ENSMUSG00000060126 | Tpt1 | chr14 | + | 75845092 | 75848525 | regular |
| ENSMUSG00000060143 | Gm10076 | chr14 | + | 105681827 | 105682211 | regular |
| ENSMUSG00000060176 | Kif27 | chr13 | - | 58287501 | 58359122 | regular |
| ENSMUSG00000060260 | Pwwp2b | chr7 | + | 139248481 | 139269903 | regular |
| ENSMUSG00000060261 | Gtf2i | chr5 | - | 134237833 | 134314760 | regular |
| ENSMUSG00000060268 | Gm1661 | chr4 | - | 117213332 | 117252125 | regular |
| ENSMUSG00000060279 | Ap2a1 | chr7 | - | 44900372 | 44929496 | regular |
| ENSMUSG00000060743 | H3f3a | chr1 | - | 180800831 | 180813943 | regular |
| ENSMUSG00000060771 | Tsga10 | chr1 | - | 37754775 | 37866429 | regular |
| ENSMUSG00000060802 | B2m | chr2 | + | 122147685 | 122153083 | regular |
| ENSMUSG00000060843 | Ctnna3 | chr10 | + | 63430097 | 65003667 | regular |
| ENSMUSG00000060938 | Rpl26 | chr11 | + | 68901582 | 68906989 | regular |
| ENSMUSG00000061028 | Clasrp | chr7 | - | 19581034 | 19604486 | regular |
| ENSMUSG00000061032 | Rrp1 | chr10 | - | 78400383 | 78413043 | regular |
| ENSMUSG00000061099 | Gapdhs | chr7 | - | 30729774 | 30743681 | regular |
| ENSMUSG00000061130 | Ppm1b | chr17 | + | 84956740 | 85023991 | regular |
| ENSMUSG00000061136 | Prpf40a | chr2 | - | 53134703 | 53191284 | regular |
| ENSMUSG00000061186 | Sfmbt2 | chr2 | + | 10370509 | 10595253 | regular |
| ENSMUSG00000061232 | H2-K1 | chr17 | - | 33996016 | 34000333 | regular |
| ENSMUSG00000061288 | Taok3 | chr5 | + | 117120128 | 117275219 | regular |
| ENSMUSG00000061298 | Agbl4 | chr4 | + | 110397660 | 111664324 | regular |
| ENSMUSG00000061315 | Naca | chr10 | + | 128035574 | 128048637 | regular |
| ENSMUSG00000061322 | Dnaic1 | chr4 | + | 41569774 | 41638158 | regular |
| ENSMUSG00000061411 | Nol4l | chr2 | - | 153407461 | 153529971 | regular |
| ENSMUSG00000061436 | Hipk2 | chr6 | - | 38694389 | 38876165 | regular |
| ENSMUSG00000061455 | Stx17 | chr4 | + | 48124914 | 48186507 | regular |
| ENSMUSG00000061477 | Rps7 | chr12 | - | 28630853 | 28635953 | regular |
| ENSMUSG00000061525 | 4921509C | chr2 | - | 151470541 | 151476153 | regular |
| ENSMUSG00000061533 | Cep128 | chr12 | - | 90998491 | 91384409 | regular |
| ENSMUSG00000061589 | Dot1l | chr10 | + | 80755205 | 80795461 | regular |
| ENSMUSG00000061607 | Mdc1 | chr17 | + | 35841514 | 35859670 | regular |
| ENSMUSG00000061665 | Cd2ap | chr17 | - | 42792950 | 42876424 | regular |
| ENSMUSG00000061666 | Gdpd1 | chr11 | - | 87033866 | 87074062 | regular |
| ENSMUSG00000061689 | Dlgap4 | chr2 | + | 156613704 | 156764363 | regular |
| ENSMUSG00000061755 | Bod1l | chr5 | - | 41787537 | 41844315 | regular |
| ENSMUSG00000061787 | Rps17 | chr7 | - | 81342731 | 81345254 | regular |
| ENSMUSG00000061802 | Armc4 | chr18 | - | 7088232 | 7297901 | regular |
| ENSMUSG00000061838 | Suclg2 | chr6 | - | 95473008 | 95718800 | regular |
| ENSMUSG00000061859 | Patj | chr4 | + | 98395784 | 98719603 | regular |
| ENSMUSG00000061904 | Slc25a3 | chr10 | - | 91116573 | 91124059 | regular |
| ENSMUSG00000061923 | Odf1 | chr15 | + | 38219202 | 38226735 | regular |
| ENSMUSG00000061950 | Ppp4r1 | chr17 | + | 65782572 | 65841926 | regular |
| ENSMUSG00000061983 | Rps12 | chr10 | - | 23785182 | 23787275 | regular |
| ENSMUSG00000062014 | Gmfb | chr14 | - | 46808148 | 46822242 | regular |
| ENSMUSG00000062017 | Abca14 | chr7 | + | 120203960 | 120325352 | regular |

| | | | | | | |
|---|---|---|---|---|---|---|
| ENSMUSG00000062028 | Irgc1 | chr7 | - | 24431921 | 24452297 | regular |
| ENSMUSG00000062078 | Qk | chr17 | - | 10206470 | 10319361 | regular |
| ENSMUSG00000062154 | Tex33 | chr15 | - | 78378399 | 78395912 | regular |
| ENSMUSG00000062168 | Ppef1 | chrX | - | 160623093 | 160735765 | regular |
| ENSMUSG00000062202 | Btbd9 | chr17 | - | 30215523 | 30576287 | regular |
| ENSMUSG00000062210 | Tnfaip8 | chr18 | + | 49979426 | 50107173 | regular |
| ENSMUSG00000062270 | Morf4l1 | chr9 | - | 90091664 | 90114774 | regular |
| ENSMUSG00000062328 | Rpl17 | chr18 | + | 75000476 | 75003381 | regular |
| ENSMUSG00000062345 | Serpinb2 | chr1 | + | 107511422 | 107535478 | regular |
| ENSMUSG00000062382 | Ftl1-ps1 | chr13 | + | 74399693 | 74407346 | regular |
| ENSMUSG00000062397 | Zfp706 | chr15 | - | 36997026 | 37007773 | regular |
| ENSMUSG00000062604 | Srpk2 | chr5 | - | 23503263 | 23684617 | regular |
| ENSMUSG00000062647 | Rpl7a | chr2 | + | 26910763 | 26913318 | regular |
| ENSMUSG00000062651 | H2al2a | chr2 | - | 17996416 | 17996958 | regular |
| ENSMUSG00000062732 | Lypd4 | chr7 | - | 24864619 | 24869941 | regular |
| ENSMUSG00000062825 | Actg1 | chr11 | - | 120345689 | 120348542 | regular |
| ENSMUSG00000062859 | Tcp11 | chr17 | - | 28066746 | 28080639 | regular |
| ENSMUSG00000062901 | Klhl24 | chr16 | + | 20097553 | 20127744 | regular |
| ENSMUSG00000062939 | Stat4 | chr1 | + | 51987147 | 52107189 | regular |
| ENSMUSG00000062995 | Ica1 | chr6 | - | 8630526 | 8778488 | regular |
| ENSMUSG00000062997 | Rpl35 | chr2 | - | 39001579 | 39005624 | regular |
| ENSMUSG00000063015 | Ccni | chr5 | - | 93181932 | 93206495 | regular |
| ENSMUSG00000063065 | Mapk3 | chr7 | + | 126759600 | 126765819 | regular |
| ENSMUSG00000063077 | Kif1b | chr4 | - | 149176318 | 149307693 | regular |
| ENSMUSG00000063129 | Aldoart2 | chr12 | + | 55565238 | 55566896 | regular |
| ENSMUSG00000063229 | Ldha | chr7 | + | 46841474 | 46855627 | regular |
| ENSMUSG00000063273 | Naa15 | chr3 | + | 51415147 | 51476507 | regular |
| ENSMUSG00000063286 | Gm8995 | chr7 | - | 106504238 | 106511671 | regular |
| ENSMUSG00000063358 | Mapk1 | chr16 | + | 16983381 | 17047453 | regular |
| ENSMUSG00000063409 | Lrrc43 | chr5 | + | 123489304 | 123508205 | regular |
| ENSMUSG00000063450 | Syne2 | chr12 | + | 75818133 | 76110926 | regular |
| ENSMUSG00000063457 | Rps15 | chr10 | + | 80292452 | 80294114 | regular |
| ENSMUSG00000063511 | Snrnp70 | chr7 | - | 45376452 | 45395742 | regular |
| ENSMUSG00000063524 | Eno1 | chr4 | + | 150236720 | 150248879 | regular |
| ENSMUSG00000063550 | Nup98 | chr7 | - | 102119397 | 102210176 | regular |
| ENSMUSG00000063576 | Klhdc3 | chr17 | - | 46674549 | 46680930 | regular |
| ENSMUSG00000063810 | Alms1 | chr6 | + | 85587530 | 85702753 | regular |
| ENSMUSG00000063820 | Arl9 | chr5 | + | 77004054 | 77010606 | regular |
| ENSMUSG00000063870 | Chd4 | chr6 | + | 125095980 | 125130591 | regular |
| ENSMUSG00000063882 | Uqcrh | chr4 | - | 116066964 | 116075071 | regular |
| ENSMUSG00000063889 | Crem | chr18 | - | 3266047 | 3337748 | regular |
| ENSMUSG00000063972 | Nr6a1 | chr2 | - | 38723369 | 38927688 | regular |
| ENSMUSG00000064061 | Dzip3 | chr16 | - | 48924231 | 48994165 | regular |
| ENSMUSG00000064068 | Mtx1 | chr3 | - | 89209080 | 89227088 | regular |
| ENSMUSG00000064138 | Fam172a | chr13 | + | 77702032 | 78166235 | regular |
| ENSMUSG00000064145 | Arih2 | chr9 | - | 108602941 | 108649386 | regular |

| | | | | | | |
|---|---|---|---|---|---|---|
| ENSMUSG00000064181 | Rab3ip | chr10 | - | 116905783 | 116950769 | regular |
| ENSMUSG00000064202 | 4430402I1 | chr19 | - | 28893041 | 28967800 | regular |
| ENSMUSG00000064225 | Paqr9 | chr9 | + | 95559656 | 95568023 | regular |
| ENSMUSG00000064289 | Tank | chr2 | + | 61578584 | 61654171 | regular |
| ENSMUSG00000064299 | 4921528I0 | chr9 | - | 114237696 | 114317028 | regular |
| ENSMUSG00000064302 | Clasp1 | chr1 | + | 118389057 | 118612678 | regular |
| ENSMUSG00000064307 | Lrrc51 | chr7 | - | 101912983 | 101933895 | regular |
| ENSMUSG00000064330 | Pde6h | chr6 | + | 136923831 | 136968865 | regular |
| ENSMUSG00000064337 | mt-Rnr1 | chrM | + | 69 | 1024 | regular |
| ENSMUSG00000064339 | mt-Rnr2 | chrM | + | 1093 | 2675 | regular |
| ENSMUSG00000064341 | mt-Nd1 | chrM | + | 2750 | 3707 | regular |
| ENSMUSG00000064345 | mt-Nd2 | chrM | + | 3913 | 4951 | regular |
| ENSMUSG00000064351 | mt-Co1 | chrM | + | 5327 | 6872 | regular |
| ENSMUSG00000064354 | mt-Co2 | chrM | + | 7012 | 7696 | regular |
| ENSMUSG00000064357 | mt-Atp6 | chrM | + | 7926 | 8607 | regular |
| ENSMUSG00000064358 | mt-Co3 | chrM | + | 8606 | 9390 | regular |
| ENSMUSG00000064360 | mt-Nd3 | chrM | + | 9458 | 9806 | regular |
| ENSMUSG00000064363 | mt-Nd4 | chrM | + | 10166 | 11544 | regular |
| ENSMUSG00000064367 | mt-Nd5 | chrM | + | 11741 | 13565 | regular |
| ENSMUSG00000064370 | mt-Cytb | chrM | + | 14144 | 15288 | regular |
| ENSMUSG00000064702 | Gm24950 | chr11 | + | 101641747 | 101641938 | regular |
| ENSMUSG00000064923 | Gm22042 | chr4 | - | 51812482 | 51812647 | regular |
| ENSMUSG00000064941 | Gm23238 | chr19 | + | 18732740 | 18732904 | regular |
| ENSMUSG00000064999 | Gm26035 | chrX | - | 152923452 | 152923760 | regular |
| ENSMUSG00000065037 | Rn7sk | chr9 | - | 78175302 | 78175633 | regular |
| ENSMUSG00000065176 | Rnu12 | chr15 | + | 83149643 | 83149794 | regular |
| ENSMUSG00000065251 | Gm23971 | chr11 | - | 101645739 | 101645930 | regular |
| ENSMUSG00000065701 | Rny1 | chr6 | - | 47788068 | 47788180 | regular |
| ENSMUSG00000065767 | Gm23849 | chr11 | + | 101627365 | 101627556 | regular |
| ENSMUSG00000065773 | Rnu1b6 | chr3 | - | 96460023 | 96460189 | regular |
| ENSMUSG00000065820 | Gm26316 | chr11 | + | 101604849 | 101605040 | regular |
| ENSMUSG00000065824 | Gm26315 | chr2 | - | 7076093 | 7076414 | regular |
| ENSMUSG00000065911 | Gm24447 | chr15 | + | 93234362 | 93234700 | regular |
| ENSMUSG00000065944 | Rnu2-10 | chr11 | + | 101658237 | 101658428 | regular |
| ENSMUSG00000065954 | Tacc1 | chr8 | - | 25154551 | 25256588 | regular |
| ENSMUSG00000066036 | Ubr4 | chr4 | + | 139352608 | 139489588 | regular |
| ENSMUSG00000066037 | Hnrnpr | chr4 | + | 136310941 | 136359447 | regular |
| ENSMUSG00000066278 | Vps37b | chr5 | - | 124004640 | 124032270 | regular |
| ENSMUSG00000066306 | Numa1 | chr7 | + | 101934110 | 102014964 | regular |
| ENSMUSG00000066368 | Actl11 | chr9 | + | 107928468 | 107932461 | regular |
| ENSMUSG00000066406 | Akap13 | chr7 | + | 75455533 | 75754609 | regular |
| ENSMUSG00000066643 | Wdr35 | chr12 | + | 8973891 | 9028847 | regular |
| ENSMUSG00000066900 | Suds3 | chr5 | - | 117091679 | 117116113 | regular |
| ENSMUSG00000067028 | Cntnap5b | chr1 | + | 99772764 | 100485942 | regular |
| ENSMUSG00000067274 | Rplp0 | chr5 | + | 115559466 | 115563727 | regular |
| ENSMUSG00000067288 | Rps28 | chr17 | - | 33819026 | 33824562 | regular |

| | | | | | | |
|---|---|---|---|---|---|---|
| ENSMUSG00000067338 | Tuba3b | chr6 | + | 145615756 | 145621479 | regular |
| ENSMUSG00000067367 | Lyar | chr5 | + | 38220469 | 38234306 | regular |
| ENSMUSG00000067377 | Tspan6 | chrX | - | 133891067 | 133898429 | regular |
| ENSMUSG00000067629 | Syngap1 | chr17 | + | 26941252 | 26972434 | regular |
| ENSMUSG00000067702 | Tuba3a | chr6 | - | 125278273 | 125286042 | regular |
| ENSMUSG00000067851 | Arfgef1 | chr1 | - | 10137570 | 10232670 | regular |
| ENSMUSG00000068036 | Afdn | chr17 | + | 13760538 | 13906150 | regular |
| ENSMUSG00000068039 | Tcp1 | chr17 | + | 12915700 | 12925067 | regular |
| ENSMUSG00000068167 | Csnka2ip | chr16 | - | 64477805 | 64602408 | regular |
| ENSMUSG00000068240 | Gm11808 | chr4 | - | 3973091 | 3973595 | regular |
| ENSMUSG00000068663 | Clec16a | chr16 | + | 10545338 | 10744878 | regular |
| ENSMUSG00000068735 | Trp53i11 | chr2 | + | 93187547 | 93201759 | regular |
| ENSMUSG00000068747 | Sort1 | chr3 | + | 108284081 | 108361511 | regular |
| ENSMUSG00000068758 | Il3ra | chr14 | + | 14346274 | 14356011 | regular |
| ENSMUSG00000068823 | Csde1 | chr3 | + | 103020425 | 103058186 | regular |
| ENSMUSG00000068917 | Clk2 | chr3 | + | 89164794 | 89176921 | regular |
| ENSMUSG00000068921 | Dap3 | chr3 | - | 88920802 | 88951181 | regular |
| ENSMUSG00000068922 | Msto1 | chr3 | - | 88905106 | 88913999 | regular |
| ENSMUSG00000069094 | Pde7a | chr3 | - | 19223107 | 19311322 | regular |
| ENSMUSG00000069135 | Fgfr1op | chr17 | + | 8165500 | 8196804 | regular |
| ENSMUSG00000069170 | Adgrv1 | chr13 | - | 81095067 | 81633154 | regular |
| ENSMUSG00000069495 | Epc2 | chr2 | + | 49451485 | 49551948 | regular |
| ENSMUSG00000069516 | Lyz2 | chr10 | - | 117277330 | 117282321 | regular |
| ENSMUSG00000069565 | Dazap1 | chr10 | + | 80261485 | 80288408 | regular |
| ENSMUSG00000069769 | Msi2 | chr11 | - | 88339381 | 88718513 | regular |
| ENSMUSG00000069833 | Ahnak | chr19 | + | 8989283 | 9076919 | regular |
| ENSMUSG00000069895 | Atxn1l | chr8 | - | 109726450 | 109737739 | regular |
| ENSMUSG00000069917 | Hba-a2 | chr11 | + | 32296488 | 32297298 | regular |
| ENSMUSG00000069919 | Hba-a1 | chr11 | + | 32283510 | 32284465 | regular |
| ENSMUSG00000069971 | 4933402J( | chr8 | + | 87563852 | 87589424 | regular |
| ENSMUSG00000070031 | Sp140 | chr1 | + | 85600377 | 85645037 | regular |
| ENSMUSG00000070047 | Fat1 | chr8 | + | 44935446 | 45052257 | regular |
| ENSMUSG00000070283 | Ndufaf3 | chr9 | - | 108565536 | 108567434 | regular |
| ENSMUSG00000070319 | Eif3g | chr9 | - | 20894348 | 20898623 | regular |
| ENSMUSG00000070327 | Rnf213 | chr11 | + | 119393099 | 119487418 | regular |
| ENSMUSG00000070331 | Qrich2 | chr11 | - | 116441324 | 116466241 | regular |
| ENSMUSG00000070332 | Trim80 | chr11 | + | 115440544 | 115448270 | regular |
| ENSMUSG00000070345 | Hsf5 | chr11 | + | 87617163 | 87659542 | regular |
| ENSMUSG00000070372 | Capza1 | chr3 | - | 104822778 | 104864505 | regular |
| ENSMUSG00000070529 | Wfdc10 | chr2 | + | 164656045 | 164657368 | regular |
| ENSMUSG00000070530 | Wfdc16 | chr2 | - | 164634706 | 164638796 | regular |
| ENSMUSG00000070531 | Wfdc6b | chr2 | + | 164613521 | 164618212 | regular |
| ENSMUSG00000070544 | Top1 | chr2 | + | 160645887 | 160722764 | regular |
| ENSMUSG00000070565 | Rasal2 | chr1 | - | 157135181 | 157412595 | regular |
| ENSMUSG00000070568 | Slc6a21 | chr7 | + | 45277512 | 45288998 | regular |
| ENSMUSG00000070639 | Lrrc8b | chr5 | + | 105415774 | 105490074 | regular |

| | | | | | | |
|---|---|---|---|---|---|---|
| ENSMUSG00000070733 | Fryl | chr5 | - | 73019986 | 73256619 | regular |
| ENSMUSG00000070738 | Dgkd | chr1 | + | 87853286 | 87945180 | regular |
| ENSMUSG00000070780 | Rbm47 | chr5 | - | 66016548 | 66173118 | regular |
| ENSMUSG00000070871 | Ccnyl1 | chr1 | + | 64690454 | 64725644 | regular |
| ENSMUSG00000070979 | Actl7a | chr4 | + | 56743412 | 56744925 | regular |
| ENSMUSG00000070980 | Actl7b | chr4 | - | 56740004 | 56741443 | regular |
| ENSMUSG00000071054 | Safb | chr17 | + | 56584824 | 56606294 | regular |
| ENSMUSG00000071064 | Zfp827 | chr8 | + | 79028436 | 79193766 | regular |
| ENSMUSG00000071076 | Jund | chr8 | + | 70698948 | 70700616 | regular |
| ENSMUSG00000071179 | Serpina16 | chr12 | - | 103668807 | 103676139 | regular |
| ENSMUSG00000071226 | Cecr2 | chr6 | + | 120666368 | 120771190 | regular |
| ENSMUSG00000071252 | 2210408I2 | chr13 | + | 77135539 | 77613784 | regular |
| ENSMUSG00000071322 | Tcp10a | chr17 | + | 7324645 | 7345974 | regular |
| ENSMUSG00000071324 | Armc2 | chr10 | - | 41914989 | 42018442 | regular |
| ENSMUSG00000071359 | Tbpl1 | chr10 | - | 22703878 | 22731938 | regular |
| ENSMUSG00000071415 | Rpl23 | chr11 | - | 97777526 | 97782437 | regular |
| ENSMUSG00000071454 | Dtnb | chr12 | + | 3572380 | 3781796 | regular |
| ENSMUSG00000071550 | Cfap44 | chr16 | + | 44394795 | 44482428 | regular |
| ENSMUSG00000071636 | Rimbp3 | chr16 | + | 17208134 | 17213921 | regular |
| ENSMUSG00000071644 | Eef1g | chr19 | + | 8967040 | 8978479 | regular |
| ENSMUSG00000071655 | Ubxn1 | chr19 | + | 8871558 | 8875663 | regular |
| ENSMUSG00000071657 | Bscl2 | chr19 | + | 8837466 | 8848683 | regular |
| ENSMUSG00000071659 | Hnrnpul2 | chr19 | + | 8819400 | 8834142 | regular |
| ENSMUSG00000071669 | Snx29 | chr16 | + | 11322907 | 11755472 | regular |
| ENSMUSG00000071713 | Csf2rb | chr15 | + | 78325989 | 78351000 | regular |
| ENSMUSG00000071724 | Smpd5 | chr15 | + | 76294433 | 76296896 | regular |
| ENSMUSG00000071866 | Ppia | chr11 | + | 6415442 | 6419817 | regular |
| ENSMUSG00000072214 | Sep.05 | chr16 | - | 18621810 | 18629938 | regular |
| ENSMUSG00000072295 | C2cd6 | chr1 | - | 59050505 | 59094900 | regular |
| ENSMUSG00000072487 | Mroh5 | chr15 | - | 73761409 | 73839699 | regular |
| ENSMUSG00000072663 | Spef2 | chr15 | - | 9578192 | 9748868 | regular |
| ENSMUSG00000072723 | Gm10044 | chr14 | - | 7744062 | 7791651 | regular |
| ENSMUSG00000072791 | Abcb5 | chr12 | - | 118867823 | 118966421 | regular |
| ENSMUSG00000072825 | Cep170b | chr12 | + | 112720454 | 112746592 | regular |
| ENSMUSG00000072878 | 1700123L | chr6 | - | 96164496 | 96166243 | regular |
| ENSMUSG00000073411 | H2-D1 | chr17 | + | 35262729 | 35267499 | regular |
| ENSMUSG00000073418 | C4b | chr17 | - | 34728379 | 34743882 | regular |
| ENSMUSG00000073471 | Rsph3a | chr17 | + | 7945652 | 7979824 | regular |
| ENSMUSG00000073643 | Wdfy1 | chr1 | - | 79702261 | 79776143 | regular |
| ENSMUSG00000073702 | Rpl31 | chr1 | + | 39367841 | 39371911 | regular |
| ENSMUSG00000073725 | Lmbrd1 | chr1 | + | 24678629 | 24766301 | regular |
| ENSMUSG00000073730 | 4933415F | chr1 | - | 23100472 | 23102253 | regular |
| ENSMUSG00000073731 | AI507597 | chr4 | + | 141614025 | 141615604 | regular |
| ENSMUSG00000073758 | Sh3d21 | chr4 | - | 126150601 | 126163491 | regular |
| ENSMUSG00000073856 | Iqck | chr7 | + | 118855751 | 118972652 | regular |
| ENSMUSG00000073940 | Hbb-bt | chr7 | - | 103812523 | 103813996 | regular |

| | | | | | | |
|---|---|---|---|---|---|---|
| ENSMUSG00000074063 | Osgin1 | chr8 | + | 119434123 | 119446256 | regular |
| ENSMUSG00000074146 | 4930579C | chr9 | - | 89113110 | 89177887 | regular |
| ENSMUSG00000074149 | Gm10634 | chr9 | + | 88719628 | 88806264 | regular |
| ENSMUSG00000074151 | Nlrc5 | chr8 | + | 94434355 | 94527272 | regular |
| ENSMUSG00000074212 | Dnajb14 | chr3 | + | 137867674 | 137916557 | regular |
| ENSMUSG00000074247 | Dda1 | chr8 | + | 71469198 | 71476098 | regular |
| ENSMUSG00000074435 | Smcp | chr3 | - | 92583866 | 92588997 | regular |
| ENSMUSG00000074476 | Spc24 | chr9 | - | 21755441 | 21760303 | regular |
| ENSMUSG00000074579 | Lekr1 | chr3 | + | 65666227 | 65831159 | regular |
| ENSMUSG00000074582 | Arfgef2 | chr2 | + | 166805587 | 166898052 | regular |
| ENSMUSG00000074619 | 1700034I2 | chr3 | - | 40900195 | 40902543 | regular |
| ENSMUSG00000074627 | Mroh8 | chr2 | - | 157208549 | 157279549 | regular |
| ENSMUSG00000074634 | Tmem267 | chr13 | + | 119488038 | 119611059 | regular |
| ENSMUSG00000074679 | Defb28 | chr2 | + | 152518254 | 152521447 | regular |
| ENSMUSG00000074746 | Pdzd8 | chr19 | - | 59296083 | 59345780 | regular |
| ENSMUSG00000074748 | Atxn7l3b | chr10 | - | 112925429 | 112929001 | regular |
| ENSMUSG00000074749 | Kiz | chr2 | + | 146855863 | 146970097 | regular |
| ENSMUSG00000074771 | Ankef1 | chr2 | + | 136501909 | 136562091 | regular |
| ENSMUSG00000074781 | Ube2n | chr10 | + | 95515144 | 95545657 | regular |
| ENSMUSG00000074817 | Papolb | chr5 | - | 142525837 | 142530076 | regular |
| ENSMUSG00000074884 | Serf2 | chr2 | + | 121449194 | 121458313 | regular |
| ENSMUSG00000075014 | Gm10800 | chr2 | - | 98666546 | 98667301 | regular |
| ENSMUSG00000075015 | Gm10801 | chr2 | + | 98662236 | 98664083 | regular |
| ENSMUSG00000075033 | Nxpe3 | chr16 | - | 55839952 | 55895279 | regular |
| ENSMUSG00000075227 | Znhit2 | chr19 | + | 6061191 | 6062472 | regular |
| ENSMUSG00000075249 | Fsip2 | chr2 | + | 82943633 | 83008937 | regular |
| ENSMUSG00000075302 | Erich2 | chr2 | + | 70508818 | 70540884 | regular |
| ENSMUSG00000075376 | Rc3h2 | chr2 | - | 37370068 | 37422903 | regular |
| ENSMUSG00000075415 | Fnbp1 | chr2 | - | 31026205 | 31142008 | regular |
| ENSMUSG00000075511 | 1700001L | chr15 | - | 83357525 | 83367282 | regular |
| ENSMUSG00000075520 | Malrd1 | chr2 | + | 15526478 | 16255555 | regular |
| ENSMUSG00000075524 | 4930407I1 | chr15 | + | 82059150 | 82066538 | regular |
| ENSMUSG00000075569 | Rsph10b | chr5 | + | 143933034 | 143985719 | regular |
| ENSMUSG00000075706 | Gpx4 | chr10 | + | 80047165 | 80056439 | regular |
| ENSMUSG00000076258 | Gm23935 | chr16 | + | 11144124 | 11144181 | regular |
| ENSMUSG00000076281 | Gm24270 | chr9 | - | 56223714 | 56223771 | regular |
| ENSMUSG00000076432 | Ywhaq | chr12 | - | 21390070 | 21417637 | regular |
| ENSMUSG00000076433 | Cep295nl | chr11 | - | 118332359 | 118342500 | regular |
| ENSMUSG00000076436 | Oxct2a | chr4 | - | 123321874 | 123323634 | regular |
| ENSMUSG00000076438 | Oxct2b | chr4 | + | 123116265 | 123118000 | regular |
| ENSMUSG00000076617 | Ighm | chr12 | - | 113418557 | 113422730 | regular |
| ENSMUSG00000077239 | Snord66 | chr16 | + | 20684238 | 20684313 | regular |
| ENSMUSG00000077714 | Snord17 | chr2 | - | 144265978 | 144266216 | regular |
| ENSMUSG00000078161 | Erich3 | chr3 | + | 154663858 | 154767790 | regular |
| ENSMUSG00000078193 | Gm2000 | chr1 | + | 156366039 | 156366484 | regular |
| ENSMUSG00000078429 | Ctdsp2 | chr10 | + | 126978716 | 126999975 | regular |

| | | | | | | |
|---|---|---|---|---|---|---|
| ENSMUSG00000078439 | Smim24 | chr10 | + | 81393050 | 81395392 | regular |
| ENSMUSG00000078490 | Cfap74 | chr4 | + | 155409189 | 155466823 | regular |
| ENSMUSG00000078578 | Ube2d3 | chr3 | + | 135438148 | 135468198 | regular |
| ENSMUSG00000078611 | Gm5901 | chr7 | + | 105375037 | 105378377 | regular |
| ENSMUSG00000078627 | Mar.10 | chr11 | - | 105360797 | 105456735 | regular |
| ENSMUSG00000078632 | Lrrc37a | chr11 | - | 103451954 | 103504597 | regular |
| ENSMUSG00000078671 | Chd2 | chr7 | - | 73426637 | 73541830 | regular |
| ENSMUSG00000078796 | Zfp541 | chr7 | + | 16061813 | 16096334 | regular |
| ENSMUSG00000078812 | Eif5a | chr11 | - | 69916713 | 69921958 | regular |
| ENSMUSG00000078907 | Fam186b | chr15 | - | 99271017 | 99287180 | regular |
| ENSMUSG00000078908 | Mon1b | chr8 | + | 113635586 | 113645179 | regular |
| ENSMUSG00000079005 | Gm14147 | chr2 | + | 151098244 | 151103818 | regular |
| ENSMUSG00000079006 | Gm14151 | chr2 | - | 151086785 | 151092350 | regular |
| ENSMUSG00000079022 | Col22a1 | chr15 | - | 71795794 | 72034227 | regular |
| ENSMUSG00000079173 | Zan | chr5 | - | 137378636 | 137477064 | regular |
| ENSMUSG00000079235 | Ccdc13 | chr9 | - | 121797626 | 121839461 | regular |
| ENSMUSG00000079324 | 4932414N | chr2 | + | 68656485 | 68748467 | regular |
| ENSMUSG00000079356 | Mettl4-ps | chr17 | + | 71526895 | 71556543 | regular |
| ENSMUSG00000079429 | Mroh2a | chr1 | + | 88226985 | 88262289 | regular |
| ENSMUSG00000079435 | Rpl36a | chrX | + | 134585653 | 134588062 | regular |
| ENSMUSG00000079523 | Tmsb10 | chr6 | - | 72957346 | 72958748 | regular |
| ENSMUSG00000079557 | Mar.02 | chr17 | - | 33685691 | 33718670 | regular |
| ENSMUSG00000079606 | Gm595 | chrX | - | 48841465 | 48877713 | regular |
| ENSMUSG00000079608 | Stard6 | chr18 | + | 70472453 | 70501066 | regular |
| ENSMUSG00000079641 | Rpl39 | chrX | - | 37082519 | 37085402 | regular |
| ENSMUSG00000079652 | Fam71f2 | chr6 | + | 29279592 | 29290680 | regular |
| ENSMUSG00000079710 | Gm3448 | chr17 | - | 14964188 | 15041537 | regular |
| ENSMUSG00000080316 | Spaca6 | chr17 | + | 17827157 | 17843009 | regular |
| ENSMUSG00000081984 | Dnajb3 | chr1 | - | 88204731 | 88205782 | regular |
| ENSMUSG00000083012 | Fam220a | chr5 | + | 143548705 | 143564531 | regular |
| ENSMUSG00000083899 | Gm12346 | chr11 | + | 77293443 | 77295898 | regular |
| ENSMUSG00000084786 | Ubl5 | chr9 | + | 20642877 | 20647140 | regular |
| ENSMUSG00000085143 | Gm11520 | chr11 | + | 95310428 | 95324496 | regular |
| ENSMUSG00000085438 | 1700020I1 | chr2 | + | 119594296 | 119607502 | regular |
| ENSMUSG00000085576 | Dpy19l2 | chr9 | - | 24557046 | 24696293 | regular |
| ENSMUSG00000085720 | Gm7854 | chr5 | - | 43151685 | 43235355 | regular |
| ENSMUSG00000085793 | Lin52 | chr12 | + | 84451421 | 84546145 | regular |
| ENSMUSG00000086211 | Gm12462 | chr4 | + | 44664414 | 44685347 | regular |
| ENSMUSG00000086324 | Gm15564 | chr16 | - | 35966751 | 35983230 | regular |
| ENSMUSG00000086451 | 4933431K | chr8 | + | 77610859 | 77769682 | regular |
| ENSMUSG00000086459 | 1700030C | chr11 | + | 23497752 | 23499661 | regular |
| ENSMUSG00000086583 | Gm15500 | chr7 | - | 114705216 | 114706235 | regular |
| ENSMUSG00000086808 | Gm11981 | chr11 | - | 6716000 | 6720037 | regular |
| ENSMUSG00000086856 | Gm16080 | chr4 | - | 131694606 | 131710508 | regular |
| ENSMUSG00000086898 | Itpr3os | chr17 | - | 27099657 | 27101440 | regular |
| ENSMUSG00000087279 | 4930544D | chr11 | + | 70615847 | 70616890 | regular |

| | | | | | | |
|---|---|---|---|---|---|---|
| ENSMUSG00000087574 | C030037D | chr11 | + | 88718644 | 88728893 | regular |
| ENSMUSG00000089872 | Rps6kc1 | chr1 | - | 190700201 | 190911770 | regular |
| ENSMUSG00000089992 | G6pd2 | chr5 | + | 61808815 | 61811163 | regular |
| ENSMUSG00000090173 | Fbxw10 | chr11 | + | 62847068 | 62877465 | regular |
| ENSMUSG00000090202 | 4930503B | chr3 | - | 146645192 | 146651317 | regular |
| ENSMUSG00000090213 | Tmem189 | chr2 | - | 167642607 | 167689822 | regular |
| ENSMUSG00000090369 | 4933411K | chr19 | + | 42052247 | 42053627 | regular |
| ENSMUSG00000090386 | Mir99ahg | chr16 | + | 77236316 | 77691403 | regular |
| ENSMUSG00000090394 | 4930523C | chr1 | + | 160044381 | 160080208 | regular |
| ENSMUSG00000090457 | 4930571K | chr7 | + | 125368860 | 125371013 | regular |
| ENSMUSG00000090558 | Gm17202 | chr5 | - | 107404822 | 107431910 | regular |
| ENSMUSG00000090733 | Rps27 | chr3 | - | 90212521 | 90213651 | regular |
| ENSMUSG00000090841 | Myl6 | chr10 | - | 128490859 | 128494145 | regular |
| ENSMUSG00000090862 | Rps13 | chr7 | - | 116331504 | 116334195 | regular |
| ENSMUSG00000090877 | Hspa1b | chr17 | - | 34956435 | 34959238 | regular |
| ENSMUSG00000091028 | Gm10722 | chr9 | + | 3000921 | 3002330 | regular |
| ENSMUSG00000091415 | Ak9 | chr10 | + | 41303979 | 41434534 | regular |
| ENSMUSG00000091476 | Gm16432 | chr1 | + | 177983422 | 178172704 | regular |
| ENSMUSG00000091896 | Ube2d2a | chr18 | + | 35771558 | 35807172 | regular |
| ENSMUSG00000091971 | Hspa1a | chr17 | - | 34969189 | 34972156 | regular |
| ENSMUSG00000092274 | Neat1 | chr19 | - | 5824707 | 5845478 | regular |
| ENSMUSG00000092341 | Malat1 | chr19 | - | 5795689 | 5802672 | regular |
| ENSMUSG00000092518 | Fam71e2 | chr7 | - | 4753225 | 4771302 | regular |
| ENSMUSG00000092519 | Actl9 | chr17 | + | 33432895 | 33434268 | regular |
| ENSMUSG00000092702 | Gm24514 | chr18 | - | 55622635 | 55622920 | regular |
| ENSMUSG00000092746 | Rn7s6 | chr6 | + | 69516338 | 69516629 | regular |
| ENSMUSG00000092805 | Gm26461 | chr9 | + | 88553235 | 88553525 | regular |
| ENSMUSG00000093674 | Rpl41 | chr10 | - | 128548113 | 128549305 | regular |
| ENSMUSG00000093815 | Gm26444 | chr12 | + | 54729601 | 54729765 | regular |
| ENSMUSG00000093834 | Rnu1b2 | chr3 | + | 96374020 | 96374185 | regular |
| ENSMUSG00000094306 | Gm24924 | chr18 | - | 14647766 | 14647930 | regular |
| ENSMUSG00000094377 | Gm24407 | chr5 | + | 115490253 | 115490394 | regular |
| ENSMUSG00000094445 | 1700015G | chr7 | - | 52011682 | 52015728 | regular |
| ENSMUSG00000094483 | Purb | chr11 | - | 6467598 | 6475917 | regular |
| ENSMUSG00000094812 | Gm22614 | chr3 | + | 96488728 | 96488893 | regular |
| ENSMUSG00000094826 | Gm23804 | chr12 | + | 54714217 | 54714381 | regular |
| ENSMUSG00000095186 | Gm10718 | chr9 | + | 3023546 | 3025218 | regular |
| ENSMUSG00000095260 | Gm25890 | chr3 | - | 96486554 | 96486719 | regular |
| ENSMUSG00000095280 | Gm21738 | chr14 | - | 19415856 | 19418930 | regular |
| ENSMUSG00000095463 | Entpd4 | chr14 | + | 69337173 | 69367270 | regular |
| ENSMUSG00000095547 | Gm10719 | chr9 | + | 3017407 | 3021593 | regular |
| ENSMUSG00000095580 | Rnu1b1 | chr3 | - | 96359952 | 96360117 | regular |
| ENSMUSG00000095590 | Gm24305 | chr12 | - | 54734554 | 54734718 | regular |
| ENSMUSG00000095616 | Gm26244 | chr18 | - | 10150976 | 10151140 | regular |
| ENSMUSG00000095701 | Gm24830 | chr3 | + | 96328424 | 96328590 | regular |
| ENSMUSG00000095738 | Gm25313 | chr9 | - | 65196571 | 65196687 | regular |

| | | | | | | |
|---|---|---|---|---|---|---|
| ENSMUSG00000095789 | Nupr1l | chr5 | + | 129908539 | 129913789 | regular |
| ENSMUSG00000095891 | Gm10717 | chr9 | + | 3025416 | 3033289 | regular |
| ENSMUSG00000095892 | Rnu5g | chr9 | - | 65201725 | 65201841 | regular |
| ENSMUSG00000095969 | Rnu1a1 | chr11 | + | 87422866 | 87423030 | regular |
| ENSMUSG00000096054 | Syne1 | chr10 | - | 5020916 | 5551482 | regular |
| ENSMUSG00000096141 | Dnah7a | chr1 | - | 53397005 | 53706784 | regular |
| ENSMUSG00000096205 | Gm22068 | chr11 | + | 87426715 | 87426879 | regular |
| ENSMUSG00000096206 | Gm22317 | chr12 | - | 54738458 | 54738622 | regular |
| ENSMUSG00000096214 | Gm22634 | chr12 | + | 54710307 | 54710471 | regular |
| ENSMUSG00000096243 | Gm24265 | chr5 | + | 115489458 | 115489599 | regular |
| ENSMUSG00000096349 | Gm22513 | chr12 | + | 54696781 | 54696945 | regular |
| ENSMUSG00000096385 | Gm11168 | chr9 | + | 3003345 | 3006948 | regular |
| ENSMUSG00000096573 | 1700009J( | chr10 | + | 77893416 | 77896114 | regular |
| ENSMUSG00000096659 | Gm25679 | chr12 | - | 54719162 | 54719326 | regular |
| ENSMUSG00000096838 | Gm26232 | chr3 | + | 96450445 | 96450611 | regular |
| ENSMUSG00000096887 | Gm20594 | chr6 | + | 79817915 | 79818364 | regular |
| ENSMUSG00000097039 | Pvt1 | chr15 | + | 62037985 | 62378096 | regular |
| ENSMUSG00000097055 | Gm4419 | chr12 | + | 21417322 | 21421064 | regular |
| ENSMUSG00000097075 | D830044I: | chr7 | - | 126971721 | 126976098 | regular |
| ENSMUSG00000097312 | Gm26870 | chr9 | - | 3000281 | 3038313 | regular |
| ENSMUSG00000097585 | E230029C | chr7 | + | 89980722 | 90049069 | regular |
| ENSMUSG00000097589 | Dleu2 | chr14 | - | 61602838 | 61682373 | regular |
| ENSMUSG00000097971 | Gm26917 | chr17 | + | 39843012 | 39846341 | regular |
| ENSMUSG00000098178 | Gm42418 | chr17 | + | 39846957 | 39848788 | regular |
| ENSMUSG00000098557 | Kctd12 | chr14 | - | 102976580 | 102982637 | regular |
| ENSMUSG00000098973 | Mir6236 | chr9 | + | 110281286 | 110281409 | regular |
| ENSMUSG00000099021 | Rn7s1 | chr12 | + | 69159294 | 69159594 | regular |
| ENSMUSG00000099250 | Rn7s2 | chr12 | - | 69361181 | 69361481 | regular |
| ENSMUSG00000099353 | 1700031M | chr15 | + | 98398418 | 98416268 | regular |
| ENSMUSG00000099398 | Ms4a14 | chr19 | - | 11301248 | 11314454 | regular |
| ENSMUSG00000099930 | Gm2396 | chr9 | - | 88878993 | 88937342 | regular |
| ENSMUSG00000100153 | Gm5601 | chr7 | + | 119739925 | 119742226 | regular |
| ENSMUSG00000100169 | Gm29087 | chr7 | + | 38500431 | 38537077 | regular |
| ENSMUSG00000101012 | 1700008K | chr17 | - | 49112262 | 49113535 | regular |
| ENSMUSG00000101225 | 1700008J( | chr7 | - | 127510437 | 127512869 | regular |
| ENSMUSG00000101337 | Dnah7c | chr1 | + | 46425591 | 46807476 | regular |
| ENSMUSG00000101609 | Kcnq1ot1 | chr7 | - | 143212154 | 143296549 | regular |
| ENSMUSG00000101855 | 1700010N | chr7 | + | 38791660 | 38860532 | regular |
| ENSMUSG00000101959 | Ldhal6b | chr17 | - | 5417322 | 5418767 | regular |
| ENSMUSG00000101963 | 1700001J: | chr9 | + | 40050364 | 40053028 | regular |
| ENSMUSG00000101968 | 1700027A | chr1 | + | 72984843 | 73025507 | regular |
| ENSMUSG00000104301 | Wdr49 | chr3 | - | 75274987 | 75482156 | regular |
| ENSMUSG00000104388 | Gm37033 | chr1 | + | 180357648 | 180364998 | regular |
| ENSMUSG00000104529 | Rbakdn | chr5 | - | 143164778 | 143165751 | regular |
| ENSMUSG00000105703 | Gm43305 | chr14 | - | 54292442 | 54297456 | regular |
| ENSMUSG00000106106 | CT010467 | chr17 | - | 39846352 | 39848827 | regular |

| | | | | | | |
|---|---|---|---|---|---|---|
| ENSMUSG00000107008 | Gm2762 | chr13 | + | 53393656 | 53455630 | regular |
| ENSMUSG00000108448 | Gm21269 | chr7 | + | 73816545 | 73838898 | regular |
| ENSMUSG00000108738 | Gm44777 | chr7 | + | 116334254 | 116338633 | regular |
| ENSMUSG00000109170 | Gm45086 | chr7 | - | 48724379 | 48726822 | regular |
| ENSMUSG00000109179 | Gm35339 | chr15 | + | 76354471 | 76363795 | regular |
| ENSMUSG00000109278 | Gm45144 | chr7 | - | 88853042 | 88864892 | regular |
| ENSMUSG00000109336 | Samd4b | chr7 | - | 28399521 | 28598144 | regular |
| ENSMUSG00000109561 | Ankrd31 | chr13 | + | 96748271 | 96910039 | regular |
| ENSMUSG00000109564 | Muc16 | chr9 | - | 18495454 | 18674530 | regular |
| ENSMUSG00000109737 | Gm39566 | chr11 | - | 69810964 | 69815298 | regular |
| ENSMUSG00000109864 | Eid3 | chr10 | + | 82866625 | 82867930 | regular |
| ENSMUSG00000110020 | Gm45441 | chr7 | + | 45830894 | 45835249 | regular |
| ENSMUSG00000110170 | St6galnac | chr11 | - | 116677482 | 116681290 | regular |
| ENSMUSG00000110333 | Gm45861 | chr8 | + | 27447099 | 27620917 | regular |
| ENSMUSG00000110622 | Gm16486 | chr8 | + | 70703456 | 70717650 | regular |
| ENSMUSG00000110841 | Gpx4-ps2 | chr10 | + | 45875290 | 45876086 | regular |
| ENSMUSG00000111028 | Gm5922 | chr9 | - | 120133462 | 120136329 | regular |
| ENSMUSG00000111375 | A830010M | chr5 | + | 107437996 | 107512556 | regular |
| ENSMUSG00000111734 | Gm29825 | chr9 | + | 114118087 | 114192640 | regular |
| ENSMUSG00000112023 | Lilr4b | chr10 | + | 51480631 | 51486703 | regular |
| ENSMUSG00000112148 | Lilrb4a | chr10 | + | 51490955 | 51496611 | regular |
| ENSMUSG00000112323 | Gm32828 | chr12 | - | 7524001 | 7859766 | regular |
| ENSMUSG00000112627 | 4933412E | chr10 | + | 116949756 | 116963279 | regular |
| ENSMUSG00000112657 | BC106175 | chr12 | - | 8478873 | 8482157 | regular |
| ENSMUSG00000112785 | AC163296 | chr12 | + | 58813882 | 58815197 | regular |
| ENSMUSG00000113255 | AC123954 | chr13 | + | 49493317 | 49496589 | regular |
| ENSMUST00000037248 | Hirip3 | chr7 | + | 126864319 | 126864661 | circular |
| ENSMUST00000039798 | Bbs9 | chr9 | + | 22643744 | 22659145 | circular |
| ENSMUST00000039798 | Bbs9 | chr9 | + | 22643744 | 22679064 | circular |
| ENSMUST00000042373 | Shcbp1l | chr1 | + | 153435917 | 153437630 | circular |
| ENSMUST00000046656 | Tasp1 | chr2 | - | 139977606 | 140057499 | circular |
| ENSMUST00000046656 | Tasp1 | chr2 | - | 140008698 | 140057499 | circular |
| ENSMUST00000046656 | Tasp1 | chr2 | - | 140042093 | 140057499 | circular |
| ENSMUST00000063071 | Kctd19 | chr8 | - | 105395270 | 105396480 | circular |
| ENSMUST00000066715 | Zbtb48 | chr4 | - | 152020189 | 152020708 | circular |
| ENSMUST00000092143 | Msrb3 | chr10 | - | 120825107 | 120852092 | circular |
| ENSMUST00000106587 | Agbl4 | chr4 | + | 110478464 | 110580175 | circular |
| ENSMUST00000108075 | Usp32 | chr11 | - | 85077264 | 85104026 | circular |
| ENSMUST00000162036 | Rsrc1 | chr3 | + | 66994516 | 66996590 | circular |
| ENSMUST00000163257 | Fam172a | chr13 | + | 77951984 | 78006472 | circular |
| ENSMUST00000165566 | Rfx3 | chr19 | - | 27849710 | 27900903 | circular |
| ENSMUST00000166726 | Mast4 | chr13 | - | 103140162 | 103171556 | circular |
| ENSMUST00000168527 | Dennd1b | chr1 | + | 139133670 | 139144000 | circular |
| ENSMUST00000201614 | Med13l | chr5 | + | 118593332 | 118593570 | circular |
| ENSMUST00000209052 | Gm45086 | chr7 | - | 48724379 | 48726822 | circular |
| ENSMUST00000214747 | Scaper | chr9 | - | 55807012 | 55816809 | circular |

| nts | HS548_counts | | HS548_counts | | HS548_counts | |
|---|---|---|---|---|---|---|
| Sperm | Sperm | Sperm | Sperm | Sperm | Sperm | Sperm |
| Vehicle | Vehicle | Vehicle | Vehicle | Dexamethas | Dexamethas | Dexamethas |
| KG_7_10 | KG_7_11 | KG_7_12 | KG_7_13 | KG_7_14 | KG_7_15 | KG_7_16 |
| 32.5215755 | 36.2200255 | 9.71350712 | 21.2016871 | 23.2170925 | 21.583257 | 25.7868495 |
| 2.95650687 | 9.80478397 | 9.71350712 | 18.1728746 | 10.4851386 | 2.27192179 | 12.1086076 |
| 79.8256854 | 51.9076799 | 38.8540285 | 58.5570405 | 71.1491546 | 70.4295755 | 72.8758789 |
| 17.7390412 | 44.4099039 | 9.71350712 | 26.2497078 | 16.4766463 | 14.7674916 | 30.4957524 |
| 14.7825343 | 5.07541759 | 29.1405214 | 7.06722902 | 4.49363081 | 10.2236481 | 6.95123768 |
| 2.95650687 | 8.30522878 | 9.71350712 | 5.04802073 | 9.7362001 | 2.27192179 | 6.95123768 |
| 2.95650687 | 4.49866559 | 0 | 5.04802073 | 9.7362001 | 3.40788269 | 5.83007031 |
| 0 | 4.26796479 | 0 | 9.08643731 | 8.23832316 | 9.08768716 | 3.13926863 |
| 5.91301374 | 8.42057918 | 9.71350712 | 5.04802073 | 4.49363081 | 9.08768716 | 9.41780589 |
| 41.3910961 | 15.9183552 | 0 | 34.326541 | 42.6894927 | 34.0788269 | 36.9985231 |
| 11.8260275 | 11.3043392 | 19.4270142 | 12.1152497 | 8.98726163 | 9.08768716 | 14.1267088 |
| 8.8695206 | 14.0727488 | 0 | 15.1440622 | 11.234077 | 10.2236481 | 16.3690436 |
| 11.8260275 | 3.22981119 | 9.71350712 | 1.00960415 | 3.74469235 | 9.08768716 | 5.60583684 |
| 32.5215755 | 46.7169119 | 58.2810427 | 69.6626861 | 61.4129545 | 78.3813018 | 20.853713 |
| 18.8920789 | 9.21649694 | 9.71350712 | 20.5656364 | 15.750176 | 24.7525879 | 11.5143889 |
| 2.95650687 | 7.38242558 | 9.71350712 | 7.06722902 | 9.7362001 | 10.2236481 | 11.6601406 |
| 14.7825343 | 3.34516159 | 0 | 6.05762487 | 8.98726163 | 12.4955698 | 8.07240505 |
| 133.042809 | 38.0656319 | 48.5675356 | 82.7875399 | 89.1236778 | 64.749771 | 84.5360195 |
| 5.91301374 | 13.7266976 | 0 | 20.1920829 | 7.48938469 | 10.2236481 | 10.9874402 |
| 2.95650687 | 14.3034496 | 0 | 11.1056456 | 8.23832316 | 2.27192179 | 4.260436 |
| 2.95650687 | 8.30522878 | 0 | 5.04802073 | 10.4851386 | 4.54384358 | 13.9024754 |
| 8.8695206 | 7.61312638 | 0 | 7.06722902 | 3.74469235 | 12.4955698 | 5.83007031 |
| 44.347603 | 54.5607391 | 67.9945498 | 65.6242695 | 62.9108314 | 63.6138101 | 67.0458086 |
| 5.91301374 | 8.99733118 | 19.4270142 | 13.1248539 | 12.731954 | 12.4955698 | 10.9874402 |
| 14.7825343 | 14.1880992 | 0 | 7.06722902 | 3.74469235 | 12.4955698 | 7.84817157 |
| 17.7390412 | 50.1774239 | 19.4270142 | 20.1920829 | 20.2213387 | 18.1753743 | 21.9748804 |
| 5.91301374 | 2.76840959 | 0 | 9.08643731 | 8.23832316 | 4.54384358 | 7.39970463 |
| 2.95650687 | 7.84382718 | 9.71350712 | 5.04802073 | 7.48938469 | 11.359609 | 2.69080168 |
| 5.91301374 | 17.0718592 | 9.71350712 | 18.1728746 | 19.4724002 | 20.4472961 | 17.4902109 |
| 38.4345893 | 38.6423839 | 9.71350712 | 24.2304995 | 21.7192156 | 32.942866 | 34.9804219 |
| 47.3041099 | 41.9875455 | 38.8540285 | 43.4129783 | 41.1916158 | 37.4867095 | 47.0890294 |
| 14.7825343 | 17.7639616 | 29.1405214 | 25.2401036 | 14.9787694 | 22.7192179 | 21.0779465 |
| 8.8695206 | 12.688544 | 0 | 9.08643731 | 14.9787694 | 9.08768716 | 17.041744 |
| 0 | 11.8810912 | 19.4270142 | 15.1440622 | 10.1106693 | 12.4955698 | 9.7541561 |
| 2.95650687 | 10.0354848 | 0 | 7.06722902 | 7.48938469 | 9.08768716 | 14.3509423 |
| 14.7825343 | 8.76663038 | 9.71350712 | 18.1728746 | 17.9745233 | 13.6315307 | 12.781308 |
| 7.39126717 | 8.76663038 | 0 | 12.1152497 | 11.9830155 | 14.7674916 | 10.3147398 |
| 26.6085618 | 26.6459423 | 0 | 18.1728746 | 23.966031 | 14.7674916 | 24.4414486 |
| 8.8695206 | 8.07452798 | 9.71350712 | 6.05762487 | 8.23832316 | 6.81576537 | 11.8843741 |
| 2.95650687 | 8.30522878 | 9.71350712 | 8.07683317 | 2.24681541 | 9.08768716 | 5.38160336 |
| 35.4780824 | 28.6068991 | 9.71350712 | 12.1152497 | 14.2298309 | 21.583257 | 18.1629114 |
| 292.69418 | 215.243846 | 184.556635 | 238.266578 | 187.234617 | 193.113352 | 237.911715 |

| | | | | | | |
|---|---|---|---|---|---|---|
| 0 | 8.07452798 | 9.71350712 | 9.08643731 | 2.24681541 | 9.08768716 | 4.03620252 |
| 67.999658 | 47.5243647 | 19.4270142 | 39.3745617 | 41.9405543 | 43.166514 | 46.1920955 |
| 11.8260275 | 16.0337056 | 9.71350712 | 12.1152497 | 12.731954 | 9.08768716 | 13.0055415 |
| 8.8695206 | 14.8802016 | 9.71350712 | 34.326541 | 17.9745233 | 7.95172627 | 23.5445147 |
| 371.041612 | 354.183402 | 233.124171 | 322.568525 | 287.217903 | 306.709442 | 391.735878 |
| 11.8260275 | 8.53592958 | 0 | 11.6104477 | 11.9830155 | 6.24778492 | 8.52087199 |
| 2.95650687 | 7.03637438 | 9.71350712 | 3.02881244 | 7.48938469 | 2.27192179 | 6.50277073 |
| 14.7825343 | 23.7621823 | 9.71350712 | 17.1632705 | 11.9830155 | 12.4955698 | 13.6782419 |
| 56.1736305 | 12.9192448 | 97.1350712 | 71.6818943 | 41.1916158 | 42.0305531 | 41.2589591 |
| 20.6955481 | 7.03637438 | 9.71350712 | 11.1056456 | 20.2213387 | 9.08768716 | 17.4902109 |
| 76.8691786 | 54.2146879 | 29.1405214 | 67.6434778 | 80.8853547 | 57.9340057 | 73.1001124 |
| 56.1736305 | 79.7071262 | 29.1405214 | 46.4417907 | 29.9575388 | 36.3507487 | 39.4650913 |
| 11.8260275 | 20.4170207 | 0 | 8.07683317 | 8.23832316 | 15.9034525 | 13.2297749 |
| 17.7390412 | 19.0328159 | 9.71350712 | 25.2401036 | 16.4766463 | 20.4472961 | 19.5083122 |
| 8.8695206 | 10.4968864 | 0 | 9.08643731 | 16.626434 | 17.0394134 | 19.5083122 |
| 2.95650687 | 11.7657408 | 9.71350712 | 19.1824788 | 11.234077 | 5.67980448 | 11.6601406 |
| 14.7825343 | 14.7648512 | 9.71350712 | 16.1536663 | 11.9830155 | 11.359609 | 20.1810126 |
| 14.7825343 | 28.8375999 | 0 | 16.1536663 | 12.731954 | 17.0394134 | 19.2840787 |
| 2.95650687 | 7.49777598 | 0 | 9.08643731 | 7.48938469 | 3.40788269 | 6.05430379 |
| 14.7825343 | 8.76663038 | 0 | 12.1152497 | 9.7362001 | 24.9911397 | 9.86627284 |
| 224.694522 | 263.691014 | 135.9891 | 240.285787 | 208.204894 | 177.2099 | 288.812714 |
| 11.8260275 | 12.688544 | 9.71350712 | 15.1440622 | 14.9787694 | 19.3113352 | 16.8175105 |
| 2.95650687 | 7.03637438 | 9.71350712 | 5.04802073 | 8.98726163 | 5.67980448 | 6.95123768 |
| 2.95650687 | 13.0345952 | 0 | 1.00960415 | 2.24681541 | 10.2236481 | 5.94218705 |
| 5.91301374 | 5.53681919 | 9.71350712 | 11.1056456 | 10.4851386 | 3.40788269 | 11.2116737 |
| 26.6085618 | 21.2244735 | 38.8540285 | 16.1536663 | 14.9787694 | 12.4955698 | 10.3147398 |
| 8.8695206 | 15.4569536 | 0 | 10.0960415 | 7.48938469 | 13.6315307 | 14.1267088 |
| 8.8695206 | 11.7657408 | 0 | 12.1152497 | 13.4808924 | 18.1753743 | 9.86627284 |
| 11.8260275 | 6.22892158 | 9.71350712 | 8.07683317 | 11.234077 | 14.7674916 | 10.5389733 |
| 100.521233 | 1308.18888 | 48.5675356 | 78.7491234 | 58.4172006 | 51.1182403 | 67.2700421 |
| 57.6518839 | 49.7160223 | 38.8540285 | 64.1098633 | 77.1406623 | 51.6862207 | 77.1363149 |
| 260.172604 | 1609.59948 | 97.1350712 | 180.719142 | 173.004786 | 153.354721 | 178.489845 |
| 12.8016747 | 7.26707518 | 0 | 9.08643731 | 8.23832316 | 10.2236481 | 7.39970463 |
| 8.8695206 | 13.842048 | 0 | 9.08643731 | 9.7362001 | 9.08768716 | 10.0905063 |
| 5.91301374 | 23.5314815 | 19.4270142 | 12.1152497 | 5.99150775 | 9.08768716 | 11.4359072 |
| 26.6085618 | 7.26707518 | 0 | 14.134458 | 17.2255848 | 18.1753743 | 19.2840787 |
| 32.5215755 | 12.2271424 | 19.4270142 | 29.2785202 | 23.2170925 | 27.2630615 | 30.4957524 |
| 2.95650687 | 10.2661856 | 0 | 11.1056456 | 12.731954 | 12.4955698 | 11.2116737 |
| 0 | 6.34427198 | 0 | 22.2112912 | 5.99150775 | 4.54384358 | 11.8843741 |
| 29.5650687 | 40.9493919 | 19.4270142 | 35.3361451 | 25.4639079 | 30.6709442 | 38.0075738 |
| 100.521233 | 98.7399421 | 58.2810427 | 80.7683317 | 63.6597699 | 59.0699666 | 103.595865 |
| 17.7390412 | 24.8003359 | 67.9945498 | 35.3361451 | 32.9532926 | 29.5349833 | 34.5319549 |
| 17.7390412 | 27.6264207 | 9.71350712 | 16.325299 | 23.2170925 | 15.9034525 | 19.2840787 |
| 20.6955481 | 23.6468319 | 19.4270142 | 16.1536663 | 18.7234617 | 31.8069051 | 17.041744 |
| 5.91301374 | 6.80567358 | 9.71350712 | 6.05762487 | 3.74469235 | 12.4955698 | 4.48466947 |
| 130.086302 | 266.690124 | 126.275593 | 190.815184 | 128.817417 | 121.547816 | 173.332475 |

| | | | | | | |
|---|---|---|---|---|---|---|
| 505.562674 | 368.775228 | 417.680806 | 482.590782 | 414.911912 | 399.858235 | 527.621363 |
| 5.91301374 | 2.65305919 | 0 | 5.04802073 | 7.48938469 | 5.67980448 | 5.60583684 |
| 7.89387334 | 14.9759424 | 0 | 13.2964866 | 7.05500038 | 17.0394134 | 10.3528595 |
| 8.8695206 | 5.42146879 | 19.4270142 | 6.05762487 | 11.234077 | 5.67980448 | 8.96933894 |
| 0 | 5.65216958 | 9.71350712 | 10.0960415 | 7.48938469 | 1.1359609 | 4.70890294 |
| 2.95650687 | 8.65127998 | 9.71350712 | 12.1152497 | 9.7362001 | 6.81576537 | 8.52087199 |
| 5.91301374 | 11.9964416 | 9.71350712 | 15.1440622 | 8.23832316 | 6.81576537 | 8.52087199 |
| 8.8695206 | 16.8411584 | 29.1405214 | 11.1056456 | 8.23832316 | 6.81576537 | 11.4359072 |
| 8.8695206 | 7.84382718 | 9.71350712 | 3.02881244 | 12.731954 | 4.54384358 | 8.07240505 |
| 8.8695206 | 24.5696351 | 0 | 16.1536663 | 14.2298309 | 19.3113352 | 14.5751758 |
| 35.4780824 | 46.8322623 | 19.4270142 | 35.3361451 | 31.4554157 | 27.2630615 | 41.9316596 |
| 8.8695206 | 17.7639616 | 0 | 9.08643731 | 13.4808924 | 9.08768716 | 11.4359072 |
| 115.303768 | 68.8641886 | 58.2810427 | 86.8259565 | 94.3662471 | 87.4689889 | 91.0387903 |
| 14.7825343 | 5.53681919 | 0 | 10.0960415 | 17.2255848 | 12.4955698 | 14.5751758 |
| 14.7825343 | 18.8021151 | 9.71350712 | 16.1536663 | 22.4681541 | 14.7674916 | 10.7632067 |
| 14.7825343 | 27.2226943 | 9.71350712 | 17.1632705 | 14.2298309 | 4.54384358 | 11.8843741 |
| 526.258222 | 567.639317 | 301.118721 | 473.504344 | 446.367328 | 453.248397 | 454.745484 |
| 11.8260275 | 14.1880992 | 9.71350712 | 15.1440622 | 16.4766463 | 14.7674916 | 5.38160336 |
| 8.8695206 | 10.381536 | 9.71350712 | 13.1248539 | 14.2298309 | 13.6315307 | 13.0055415 |
| 5.91301374 | 16.0337056 | 9.71350712 | 24.2304995 | 13.4808924 | 13.6315307 | 16.1448101 |
| 11.8260275 | 18.1100128 | 9.71350712 | 12.1152497 | 23.2170925 | 14.7674916 | 21.9748804 |
| 31.0433221 | 4.07532962 | 19.4270142 | 7.06722902 | 9.7362001 | 9.08768716 | 10.3147398 |
| 20.6955481 | 14.5918256 | 9.71350712 | 5.5528228 | 6.74044622 | 10.2236481 | 8.52087199 |
| 76.8691786 | 49.0239199 | 67.9945498 | 91.8739773 | 71.1491546 | 82.9251454 | 88.7964555 |
| 0 | 6.11357118 | 0 | 12.1152497 | 2.99575388 | 3.40788269 | 5.15736989 |
| 47.3041099 | 32.2981119 | 38.8540285 | 62.595457 | 53.1746313 | 47.7103576 | 45.2951617 |
| 5.17388702 | 12.3228832 | 24.2837678 | 13.4580233 | 11.9830155 | 13.6315307 | 9.11957537 |
| 100.521233 | 44.7559551 | 77.708057 | 98.9412063 | 89.1236778 | 99.9645588 | 104.717032 |
| 8.8695206 | 6.45962238 | 9.71350712 | 3.02881244 | 5.99150775 | 11.359609 | 7.84817157 |
| 5.91301374 | 5.42146879 | 0 | 6.05762487 | 5.24256928 | 11.359609 | 7.6239381 |
| 82.7821923 | 68.6334878 | 38.8540285 | 83.7971441 | 48.6810005 | 53.3901621 | 71.978945 |
| 2.95650687 | 16.0337056 | 9.71350712 | 5.04802073 | 11.234077 | 9.08768716 | 10.7632067 |
| 26.6085618 | 10.8429376 | 19.4270142 | 21.2016871 | 19.4724002 | 38.6226704 | 20.4052461 |
| 14.7825343 | 24.3389343 | 19.4270142 | 31.2977285 | 17.2255848 | 15.9034525 | 18.8356118 |
| 8.8695206 | 7.61312638 | 9.71350712 | 4.03841658 | 8.98726163 | 5.67980448 | 5.83007031 |
| 11.8260275 | 10.0354848 | 0 | 10.0960415 | 8.98726163 | 9.08768716 | 14.3509423 |
| 11.8260275 | 4.03726399 | 0 | 9.08643731 | 20.2213387 | 25.5591201 | 15.6963431 |
| 23.6520549 | 61.5971134 | 9.71350712 | 37.3553534 | 25.4639079 | 56.7980448 | 37.2227566 |
| 2.95650687 | 5.53681919 | 0 | 2.01920829 | 2.99575388 | 9.08768716 | 8.29663852 |
| 23.6520549 | 4.15261439 | 9.71350712 | 17.1632705 | 14.2298309 | 6.81576537 | 13.0055415 |
| 29.5650687 | 30.1064543 | 9.71350712 | 9.08643731 | 13.4808924 | 15.9034525 | 18.8356118 |
| 2.95650687 | 3.92191359 | 0 | 6.05762487 | 7.48938469 | 14.7674916 | 5.15736989 |
| 32.5215755 | 40.3726399 | 9.71350712 | 33.3169368 | 23.2170925 | 31.8069051 | 33.635021 |
| 17.7390412 | 6.34427198 | 0 | 10.0960415 | 10.4851386 | 10.2236481 | 10.0905063 |
| 79.8256854 | 53.0035087 | 9.71350712 | 32.3073327 | 35.9490465 | 36.9187291 | 41.9316596 |
| 8.8695206 | 9.92013437 | 0 | 8.07683317 | 9.7362001 | 7.95172627 | 14.3509423 |

| | | | | | | |
|---|---|---|---|---|---|---|
| 11.8260275 | 6.92102398 | 0 | 14.134458 | 8.23832316 | 9.08768716 | 12.332841 |
| 2.95650687 | 2.99911039 | 0 | 3.53361451 | 9.36173086 | 2.83990224 | 4.59678621 |
| 8.8695206 | 24.5696351 | 0 | 10.0960415 | 18.7234617 | 11.359609 | 9.19357241 |
| 11.8260275 | 7.72847678 | 9.71350712 | 7.06722902 | 11.234077 | 12.4955698 | 10.3147398 |
| 5.91301374 | 13.1499456 | 0 | 2.01920829 | 6.74044622 | 10.2236481 | 4.48466947 |
| 62.0866442 | 52.3690815 | 38.8540285 | 82.7875399 | 79.3874777 | 102.236481 | 60.5430379 |
| 2.95650687 | 13.7266976 | 0 | 10.0960415 | 8.98726163 | 7.95172627 | 5.38160336 |
| 5.91301374 | 14.6495008 | 0 | 20.1920829 | 12.731954 | 13.6315307 | 13.2297749 |
| 5.91301374 | 8.30522878 | 9.71350712 | 6.05762487 | 5.99150775 | 11.359609 | 4.93313642 |
| 14.7825343 | 5.53681919 | 0 | 5.04802073 | 8.23832316 | 13.6315307 | 9.19357241 |
| 11.8260275 | 15.3416032 | 0 | 8.07683317 | 9.7362001 | 9.08768716 | 11.2116737 |
| 59.1301374 | 24.8003359 | 19.4270142 | 32.3073327 | 40.4426773 | 40.8945922 | 38.1196905 |
| 26.6085618 | 13.6113472 | 0 | 15.1440622 | 23.2170925 | 14.7674916 | 23.2081645 |
| 5.91301374 | 1.6149056 | 0 | 7.06722902 | 8.98726163 | 17.0394134 | 3.13926863 |
| 5.91301374 | 30.4525055 | 9.71350712 | 6.05762487 | 11.234077 | 7.95172627 | 9.19357241 |
| 0 | 9.92013437 | 0 | 9.08643731 | 14.2298309 | 11.359609 | 13.2297749 |
| 8.8695206 | 8.07452798 | 9.71350712 | 7.06722902 | 10.4851386 | 20.4472961 | 7.17547115 |
| 2.95650687 | 18.8021151 | 9.71350712 | 10.0960415 | 12.731954 | 23.8551788 | 12.332841 |
| 5.91301374 | 10.2661856 | 0 | 6.05762487 | 8.23832316 | 4.54384358 | 7.39970463 |
| 2.95650687 | 5.76751998 | 0 | 13.1248539 | 14.9787694 | 11.359609 | 8.74510547 |
| 23.6520549 | 36.3353759 | 19.4270142 | 26.2497078 | 24.7149695 | 17.0394134 | 27.1322503 |
| 17.7390412 | 9.80478397 | 0 | 11.1056456 | 11.234077 | 6.81576537 | 11.4359072 |
| 5.91301374 | 4.72936639 | 0 | 8.07683317 | 3.74469235 | 10.2236481 | 6.72700421 |
| 8.8695206 | 11.4196896 | 9.71350712 | 7.06722902 | 8.98726163 | 6.81576537 | 8.74510547 |
| 13.3042809 | 17.8216368 | 9.71350712 | 39.3745617 | 42.3150235 | 53.9581425 | 21.8627637 |
| 8.8695206 | 10.7275872 | 9.71350712 | 9.08643731 | 9.7362001 | 3.40788269 | 9.64203936 |
| 5.91301374 | 9.80478397 | 0 | 10.0960415 | 7.48938469 | 14.7674916 | 8.29663852 |
| 23.6520549 | 16.0337056 | 9.71350712 | 15.1440622 | 15.7277078 | 6.81576537 | 21.30218 |
| 8.8695206 | 10.09316 | 0 | 5.04802073 | 8.98726163 | 6.81576537 | 8.74510547 |
| 14.7825343 | 15.2262528 | 19.4270142 | 15.1440622 | 14.2298309 | 12.4955698 | 14.1267088 |
| 26.6085618 | 29.5297023 | 9.71350712 | 25.2401036 | 28.4596618 | 23.8551788 | 23.7687482 |
| 8.8695206 | 5.30611839 | 9.71350712 | 5.04802073 | 2.99575388 | 10.2236481 | 4.48466947 |
| 11.8260275 | 37.1428287 | 19.4270142 | 26.2497078 | 22.4681541 | 23.8551788 | 26.4595499 |
| 14.7825343 | 14.6495008 | 9.71350712 | 6.05762487 | 4.49363081 | 9.08768716 | 11.8843741 |
| 5.91301374 | 15.9183552 | 0 | 10.0960415 | 15.7277078 | 6.81576537 | 8.74510547 |
| 2.95650687 | 11.9964416 | 0 | 7.06722902 | 11.9830155 | 9.08768716 | 6.05430379 |
| 4.4347603 | 5.19076799 | 19.4270142 | 2.52401036 | 10.4851386 | 10.2236481 | 5.26948663 |
| 0 | 8.88198078 | 9.71350712 | 13.1248539 | 11.9830155 | 9.08768716 | 10.5389733 |
| 17.7390412 | 11.7080656 | 19.4270142 | 9.59123938 | 11.9830155 | 16.471433 | 10.202623 |
| 292.69418 | 1517.60753 | 58.2810427 | 243.314599 | 251.643326 | 211.288727 | 364.267278 |
| 10.8503802 | 19.9175535 | 0 | 25.2401036 | 17.0982652 | 16.2783196 | 18.2369084 |
| 35.4780824 | 32.5288127 | 9.71350712 | 39.3745617 | 28.0851926 | 26.1271006 | 34.8683051 |
| 8.8695206 | 7.61312638 | 0 | 19.1824788 | 12.731954 | 6.81576537 | 10.5389733 |
| 5.91301374 | 4.03726399 | 19.4270142 | 11.1056456 | 9.7362001 | 7.95172627 | 5.38160336 |
| 5.91301374 | 10.2661856 | 9.71350712 | 8.07683317 | 14.2298309 | 11.359609 | 11.4359072 |
| 7.39126717 | 8.18987838 | 9.71350712 | 8.58163524 | 10.1106693 | 9.65566761 | 9.30568915 |

| | | | | | | |
|---|---|---|---|---|---|---|
| 8.8695206 | 6.92102398 | 0 | 6.05762487 | 7.48938469 | 7.95172627 | 4.93313642 |
| 8.8695206 | 5.53681919 | 9.71350712 | 14.134458 | 20.9702771 | 18.1753743 | 19.9567791 |
| 26.6085618 | 11.53504 | 9.71350712 | 16.1536663 | 13.4808924 | 12.4955698 | 13.2297749 |
| 42.3667434 | 31.7017503 | 9.71350712 | 55.528228 | 55.4214467 | 71.9404035 | 34.1395463 |
| 141.91233 | 94.4719773 | 106.848578 | 127.210122 | 127.31954 | 120.411855 | 106.735133 |
| 26.6085618 | 18.9174655 | 9.71350712 | 13.1248539 | 11.234077 | 23.8551788 | 17.2659775 |
| 14.7825343 | 5.30611839 | 0 | 5.04802073 | 8.98726163 | 11.359609 | 7.17547115 |
| 118.260275 | 97.3522768 | 19.4270142 | 73.7011026 | 80.1364162 | 76.10938 | 96.4159089 |
| 11.8260275 | 13.265296 | 9.71350712 | 11.1056456 | 7.48938469 | 5.67980448 | 14.2388256 |
| 8.8695206 | 19.8402687 | 0 | 16.1536663 | 7.48938469 | 17.0394134 | 17.041744 |
| 20.9320686 | 16.1248324 | 16.9986375 | 24.7151095 | 20.4984459 | 25.7863123 | 17.0125936 |
| 26.7268221 | 36.8648342 | 0 | 44.1197012 | 42.7194503 | 50.3685061 | 13.2858333 |
| 13.3042809 | 15.9471928 | 19.4270142 | 18.4252757 | 10.1705844 | 23.6620655 | 8.35269689 |
| 32.5215755 | 53.4072351 | 29.1405214 | 61.5858529 | 47.1831235 | 43.166514 | 63.90654 |
| 8.8695206 | 7.84382718 | 0 | 21.2016871 | 25.4639079 | 12.4955698 | 21.7506469 |
| 118.260275 | 37.0274783 | 48.5675356 | 77.7395192 | 85.3789855 | 65.8857319 | 83.1906187 |
| 26.6085618 | 12.3424928 | 9.71350712 | 7.57203109 | 11.234077 | 9.65566761 | 11.0995569 |
| 14.7825343 | 8.07452798 | 9.71350712 | 15.1440622 | 12.731954 | 23.8551788 | 13.2297749 |
| 17.7390412 | 9.20842241 | 19.4270142 | 15.6488643 | 9.7362001 | 18.1753743 | 17.041744 |
| 0 | 17.9946624 | 19.4270142 | 9.08643731 | 8.23832316 | 11.359609 | 12.5570745 |
| 8.8695206 | 3.46051199 | 0 | 11.1056456 | 7.48938469 | 9.08768716 | 11.6601406 |
| 11.8260275 | 8.07452798 | 0 | 7.06722902 | 9.7362001 | 7.95172627 | 6.27853726 |
| 16.2607878 | 14.4188 | 38.8540285 | 16.6584684 | 13.1064232 | 15.3354721 | 12.332841 |
| 2.95650687 | 8.99733118 | 9.71350712 | 10.0960415 | 11.9830155 | 12.4955698 | 4.70890294 |
| 2.95650687 | 20.8784223 | 19.4270142 | 5.04802073 | 3.74469235 | 9.08768716 | 7.17547115 |
| 8.8695206 | 8.76663038 | 9.71350712 | 10.0960415 | 14.2298309 | 13.6315307 | 10.3147398 |
| 15.2851405 | 15.5919135 | 0 | 20.8382296 | 15.7277078 | 9.65566761 | 8.82807185 |
| 29.5650687 | 20.1863199 | 19.4270142 | 15.1440622 | 20.5958079 | 26.1271006 | 21.0779465 |
| 23.6520549 | 38.2963327 | 19.4270142 | 30.2881244 | 23.2170925 | 31.8069051 | 26.9080168 |
| 2.95650687 | 7.15172478 | 9.71350712 | 13.1248539 | 6.74044622 | 15.9034525 | 10.5389733 |
| 11.8260275 | 9.34338237 | 29.1405214 | 9.08643731 | 12.731954 | 9.08768716 | 9.64203936 |
| 44.347603 | 27.2226943 | 38.8540285 | 34.326541 | 31.4554157 | 39.7586313 | 30.0472855 |
| 14.7825343 | 16.0337056 | 0 | 15.1440622 | 11.9830155 | 13.6315307 | 12.5570745 |
| 5.91301374 | 10.0354848 | 19.4270142 | 9.08643731 | 12.731954 | 5.67980448 | 5.15736989 |
| 11.8260275 | 14.4188 | 19.4270142 | 20.1920829 | 18.7833768 | 12.4955698 | 19.5083122 |
| 5.91301374 | 11.7657408 | 9.71350712 | 8.07683317 | 8.23832316 | 11.359609 | 13.0055415 |
| 20.6955481 | 38.1809823 | 0 | 23.2208954 | 20.9702771 | 15.9034525 | 23.5445147 |
| 14.7825343 | 10.1508352 | 0 | 5.04802073 | 10.4851386 | 11.359609 | 8.07240505 |
| 79.0865587 | 69.3625023 | 38.8540285 | 81.7779358 | 43.9626881 | 75.7345129 | 58.4016082 |
| 8.8695206 | 10.5545616 | 19.4270142 | 25.7449057 | 19.8468694 | 19.3113352 | 18.9477285 |
| 17.7390412 | 6.22892158 | 19.4270142 | 13.1248539 | 22.4681541 | 9.08768716 | 14.3509423 |
| 20.6955481 | 14.1880992 | 9.71350712 | 11.1056456 | 8.23832316 | 11.359609 | 18.8356118 |
| 177.390412 | 241.543737 | 116.562085 | 159.517455 | 155.779202 | 155.626643 | 205.397862 |
| 5.91301374 | 17.6486112 | 9.71350712 | 9.08643731 | 6.74044622 | 9.08768716 | 4.260436 |
| 59.1301374 | 35.0665215 | 48.5675356 | 49.4706031 | 39.6937389 | 52.2542012 | 68.166976 |
| 2.95650687 | 5.19076799 | 0 | 3.02881244 | 8.23832316 | 11.359609 | 6.72700421 |

| | | | | | | |
|---|---|---|---|---|---|---|
| 70.9561648 | 73.593555 | 77.708057 | 78.7491234 | 54.6725082 | 57.9340057 | 63.458073 |
| 8.8695206 | 6.34427198 | 0 | 11.1056456 | 4.49363081 | 5.67980448 | 9.64203936 |
| 17.7390412 | 28.2608479 | 9.71350712 | 26.2497078 | 22.4681541 | 28.3990224 | 16.8175105 |
| 5.91301374 | 13.842048 | 0 | 17.1632705 | 7.48938469 | 6.81576537 | 3.81196905 |
| 4.4347603 | 7.30514081 | 0 | 7.7436638 | 13.3535729 | 9.46255426 | 13.7522389 |
| 41.3910961 | 40.4879903 | 29.1405214 | 37.3553534 | 38.1958619 | 37.4867095 | 40.1377918 |
| 32.5215755 | 46.9476127 | 29.1405214 | 40.3841658 | 40.4426773 | 38.6226704 | 44.8466947 |
| 5.91301374 | 8.18987838 | 0 | 1.00960415 | 7.48938469 | 3.40788269 | 8.29663852 |
| 0 | 7.20939998 | 0 | 8.07683317 | 4.49363081 | 10.2236481 | 4.260436 |
| 11.8260275 | 44.1792031 | 9.71350712 | 20.1920829 | 7.48938469 | 18.1753743 | 18.6113783 |
| 5.91301374 | 6.57497278 | 0 | 3.02881244 | 7.48938469 | 5.67980448 | 7.84817157 |
| 8.8695206 | 23.5314815 | 9.71350712 | 11.1056456 | 8.98726163 | 13.6315307 | 8.96933894 |
| 17.7390412 | 47.1783135 | 29.1405214 | 12.1152497 | 20.9702771 | 18.1753743 | 23.5445147 |
| 5.91301374 | 5.30611839 | 0 | 5.04802073 | 8.23832316 | 7.95172627 | 6.390654 |
| 8.8695206 | 6.34427198 | 9.71350712 | 3.02881244 | 8.23832316 | 7.95172627 | 6.05430379 |
| 5.91301374 | 7.15172478 | 0 | 5.04802073 | 4.49363081 | 9.08768716 | 6.50277073 |
| 14.7825343 | 12.111792 | 0 | 7.06722902 | 11.234077 | 21.583257 | 6.05430379 |
| 35.4780824 | 50.9848767 | 38.8540285 | 30.2881244 | 38.9448004 | 32.942866 | 36.7742897 |
| 2.95650687 | 12.5731936 | 0 | 13.9628253 | 9.98334979 | 9.08768716 | 8.03428536 |
| 8.8695206 | 14.6495008 | 0 | 10.0960415 | 9.7362001 | 9.08768716 | 8.74510547 |
| 5.91301374 | 12.3424928 | 9.71350712 | 9.08643731 | 9.7362001 | 9.08768716 | 11.2116737 |
| 35.4780824 | 14.8802016 | 0 | 12.1152497 | 20.2213387 | 32.942866 | 16.1448101 |
| 5.91301374 | 2.30700799 | 0 | 2.01920829 | 3.74469235 | 12.4955698 | 5.15736989 |
| 8.8695206 | 4.61401599 | 0 | 4.03841658 | 3.74469235 | 18.1753743 | 7.6239381 |
| 8.8695206 | 3.92191359 | 9.71350712 | 21.2016871 | 12.731954 | 10.2236481 | 13.4540084 |
| 5.91301374 | 10.958288 | 9.71350712 | 7.06722902 | 9.7362001 | 9.08768716 | 9.19357241 |
| 8.8695206 | 3.22981119 | 0 | 6.05762487 | 9.7362001 | 2.27192179 | 10.3147398 |
| 11.8260275 | 7.49777598 | 9.71350712 | 7.06722902 | 11.9830155 | 12.4955698 | 9.86627284 |
| 112.347261 | 79.7071262 | 29.1405214 | 92.8835814 | 65.1576468 | 84.0611063 | 104.492799 |
| 24.6277022 | 28.0301471 | 19.4270142 | 16.4363555 | 18.3115456 | 26.1271006 | 22.7036392 |
| 5.91301374 | 2.42235839 | 0 | 2.01920829 | 3.74469235 | 11.359609 | 4.03620252 |
| 14.7825343 | 2.76840959 | 9.71350712 | 9.08643731 | 7.48938469 | 6.81576537 | 8.29663852 |
| 8.8695206 | 4.15261439 | 0 | 6.05762487 | 8.23832316 | 4.54384358 | 4.48466947 |
| 11.8260275 | 11.0736384 | 0 | 10.0960415 | 11.9830155 | 18.1753743 | 11.8843741 |
| 2.95650687 | 1.8456064 | 4.85675356 | 3.02881244 | 5.61703852 | 17.6073939 | 5.26948663 |
| 8.8695206 | 7.72847678 | 0 | 3.02881244 | 6.74044622 | 9.08768716 | 5.83007031 |
| 23.6520549 | 20.3016703 | 9.71350712 | 18.1728746 | 14.9787694 | 17.0394134 | 12.5570745 |
| 35.4780824 | 29.8757535 | 29.1405214 | 16.1536663 | 20.2213387 | 19.3113352 | 30.0472855 |
| 20.6955481 | 21.1091231 | 19.4270142 | 19.1824788 | 14.9787694 | 19.3113352 | 15.6963431 |
| 10.347774 | 20.1286447 | 9.71350712 | 15.6488643 | 13.4808924 | 19.1182219 | 14.0885891 |
| 17.7390412 | 5.42146879 | 9.71350712 | 14.134458 | 7.48938469 | 7.95172627 | 7.39970463 |
| 2.95650687 | 4.84471679 | 0 | 3.02881244 | 2.99575388 | 11.359609 | 6.27853726 |
| 8.8695206 | 5.88287038 | 0 | 10.0960415 | 9.7362001 | 10.2236481 | 6.05430379 |
| 189.21644 | 231.046851 | 77.708057 | 230.189745 | 181.243109 | 189.70547 | 236.117848 |
| 14.7825343 | 13.265296 | 0 | 7.06722902 | 12.731954 | 9.08768716 | 15.0236427 |
| 11.8260275 | 15.572304 | 0 | 5.04802073 | 5.99150775 | 10.2236481 | 9.41780589 |

| | | | | | | |
|---|---|---|---|---|---|---|
| 11.8260275 | 7.84382718 | 0 | 13.1248539 | 9.7362001 | 10.2236481 | 10.9874402 |
| 15.5216611 | 9.8693802 | 16.2215569 | 20.5252523 | 25.0519918 | 29.5349833 | 16.5641267 |
| 8.8695206 | 11.3043392 | 9.71350712 | 14.134458 | 9.7362001 | 13.6315307 | 11.8843741 |
| 20.6955481 | 19.1481663 | 0 | 16.1536663 | 30.7064772 | 21.583257 | 23.3202812 |
| 8.8695206 | 8.76663038 | 0 | 2.01920829 | 8.98726163 | 4.54384358 | 9.19357241 |
| 144.868837 | 113.043392 | 126.275593 | 154.469434 | 142.298309 | 136.315307 | 152.927229 |
| 2.95650687 | 11.0736384 | 0 | 12.1152497 | 2.24681541 | 7.95172627 | 11.6601406 |
| 5.91301374 | 13.9573984 | 9.71350712 | 21.2016871 | 17.9745233 | 31.8069051 | 15.6963431 |
| 38.4345893 | 50.5234751 | 29.1405214 | 18.1728746 | 26.2128464 | 20.4472961 | 30.7199859 |
| 14.7825343 | 6.92102398 | 9.71350712 | 15.1440622 | 8.98726163 | 9.08768716 | 9.64203936 |
| 8.8695206 | 16.3797568 | 9.71350712 | 22.2112912 | 15.7277078 | 13.6315307 | 13.2297749 |
| 14.7825343 | 17.30256 | 19.4270142 | 19.1824788 | 14.2298309 | 29.5349833 | 8.07240505 |
| 76.8691786 | 179.600572 | 87.4215641 | 176.680725 | 128.068478 | 197.657196 | 52.9190998 |
| 23.6520549 | 42.9103487 | 9.71350712 | 32.3073327 | 26.2128464 | 21.583257 | 27.3564838 |
| 5.91301374 | 5.42146879 | 0 | 1.00960415 | 8.23832316 | 5.67980448 | 10.0905063 |
| 2.95650687 | 3.34516159 | 29.1405214 | 5.04802073 | 11.234077 | 12.4955698 | 6.27853726 |
| 0 | 14.0727488 | 0 | 14.134458 | 13.4808924 | 20.4472961 | 19.0598453 |
| 62.0866442 | 32.0674111 | 48.5675356 | 29.2785202 | 50.1788774 | 45.4384358 | 40.3620252 |
| 103.47774 | 82.5908862 | 67.9945498 | 103.989227 | 83.1321701 | 69.2936146 | 108.080534 |
| 2.95650687 | 5.07541759 | 19.4270142 | 8.07683317 | 9.13704932 | 15.9034525 | 10.5389733 |
| 0 | 10.958288 | 0 | 4.03841658 | 9.7362001 | 7.95172627 | 10.0905063 |
| 20.6955481 | 28.8375999 | 48.5675356 | 41.39377 | 26.9617849 | 32.942866 | 31.6169198 |
| 0 | 7.49777598 | 19.4270142 | 5.04802073 | 8.23832316 | 6.81576537 | 6.27853726 |
| 23.6520549 | 27.6840959 | 9.71350712 | 47.4513948 | 47.1831235 | 45.4384358 | 39.9135583 |
| 28.0868152 | 29.3566767 | 38.8540285 | 37.3553534 | 35.5745773 | 34.0788269 | 39.9135583 |
| 5.91301374 | 17.879312 | 0 | 14.134458 | 8.98726163 | 21.583257 | 8.29663852 |
| 135.999316 | 68.8641886 | 87.4215641 | 99.9508104 | 91.3704932 | 98.8285979 | 117.946807 |
| 26.6085618 | 54.5607391 | 0 | 20.1920829 | 27.7107234 | 17.0394134 | 27.8049507 |
| 5.91301374 | 19.0328159 | 0 | 8.07683317 | 11.234077 | 4.54384358 | 12.5570745 |
| 50.2606167 | 43.1410495 | 19.4270142 | 45.4321866 | 35.200108 | 23.8551788 | 42.155893 |
| 5.91301374 | 22.2626271 | 19.4270142 | 19.1824788 | 10.4851386 | 11.359609 | 12.1086076 |
| 56.1736305 | 72.0939998 | 19.4270142 | 53.5090197 | 49.429939 | 53.3901621 | 58.3007031 |
| 8.8695206 | 12.3424928 | 0 | 11.1056456 | 5.24256928 | 7.95172627 | 9.41780589 |
| 478.954113 | 420.682908 | 378.826778 | 484.60999 | 455.354589 | 400.994196 | 527.621363 |
| 0 | 13.3806464 | 0 | 6.05762487 | 11.9830155 | 6.81576537 | 5.83007031 |
| 5.91301374 | 7.72847678 | 0 | 8.07683317 | 8.98726163 | 7.95172627 | 8.96933894 |
| 17.7390412 | 32.5288127 | 0 | 16.1536663 | 16.4766463 | 18.1753743 | 14.5751758 |
| 8.8695206 | 22.7240287 | 9.71350712 | 23.2208954 | 19.4724002 | 18.1753743 | 21.30218 |
| 5.91301374 | 30.6832063 | 9.71350712 | 15.1440622 | 18.7234617 | 23.8551788 | 21.30218 |
| 17.7390412 | 10.4968864 | 0 | 6.05762487 | 4.49363081 | 9.08768716 | 6.50277073 |
| 2.95650687 | 3.22981119 | 9.71350712 | 7.06722902 | 7.48938469 | 4.54384358 | 4.70890294 |
| 5.91301374 | 16.9565088 | 0 | 6.05762487 | 8.23832316 | 11.359609 | 8.96933894 |
| 14.7825343 | 16.2644064 | 9.71350712 | 13.1248539 | 13.4808924 | 11.359609 | 9.19357241 |
| 20.6955481 | 15.4569536 | 0 | 10.0960415 | 14.2298309 | 19.3113352 | 15.0236427 |
| 75.6865758 | 80.5065045 | 72.8513034 | 71.853527 | 48.5836385 | 53.6968715 | 64.3258566 |
| 56.1736305 | 72.7861022 | 29.1405214 | 56.5378322 | 38.1958619 | 54.526123 | 79.401073 |

| | | | | | | |
|---|---|---|---|---|---|---|
| 11.8260275 | 9.57408317 | 9.71350712 | 9.08643731 | 8.23832316 | 14.7674916 | 3.13926863 |
| 50.2606167 | 12.8038944 | 19.4270142 | 46.4417907 | 45.6852466 | 31.8069051 | 43.0528269 |
| 29.5650687 | 34.1437183 | 38.8540285 | 25.2401036 | 26.2128464 | 24.9911397 | 28.0291842 |
| 4.4347603 | 7.55545118 | 0 | 12.6200518 | 5.99150775 | 4.54384358 | 5.60583684 |
| 5.91301374 | 7.49777598 | 0 | 6.05762487 | 8.23832316 | 9.08768716 | 11.6601406 |
| 5.91301374 | 4.26796479 | 9.71350712 | 3.02881244 | 7.48938469 | 4.54384358 | 6.27853726 |
| 8.8695206 | 6.92102398 | 0 | 6.05762487 | 6.74044622 | 10.2236481 | 6.72700421 |
| 8.8695206 | 5.76751998 | 0 | 7.06722902 | 8.98726163 | 7.95172627 | 4.48466947 |
| 0 | 11.0736384 | 0 | 4.03841658 | 8.23832316 | 10.2236481 | 6.27853726 |
| 5.91301374 | 14.0727488 | 9.71350712 | 11.1056456 | 7.48938469 | 10.2236481 | 9.64203936 |
| 8.8695206 | 12.2271424 | 0 | 24.2304995 | 13.4808924 | 22.7192179 | 28.9261181 |
| 2.95650687 | 3.22981119 | 0 | 8.07683317 | 9.7362001 | 12.4955698 | 12.332841 |
| 14.7825343 | 1.3842048 | 0 | 9.08643731 | 8.23832316 | 11.359609 | 8.74510547 |
| 2.95650687 | 10.0354848 | 9.71350712 | 8.07683317 | 11.234077 | 6.81576537 | 4.260436 |
| 20.6955481 | 12.8038944 | 9.71350712 | 16.1536663 | 7.48938469 | 18.1753743 | 12.781308 |
| 17.7390412 | 18.2253632 | 0 | 12.1152497 | 14.9787694 | 12.4955698 | 10.3147398 |
| 5.91301374 | 4.15261439 | 9.71350712 | 5.04802073 | 8.98726163 | 7.95172627 | 6.72700421 |
| 5.91301374 | 1.8456064 | 0 | 10.0960415 | 11.234077 | 4.54384358 | 6.95123768 |
| 20.6955481 | 6.11357118 | 9.71350712 | 7.57203109 | 8.98726163 | 14.7674916 | 12.6691913 |
| 11.8260275 | 7.03637438 | 9.71350712 | 6.05762487 | 10.4851386 | 7.95172627 | 8.07240505 |
| 11.8260275 | 38.1809823 | 19.4270142 | 14.134458 | 20.2213387 | 26.1271006 | 19.2840787 |
| 11.8260275 | 28.9529503 | 19.4270142 | 14.134458 | 12.731954 | 14.7674916 | 22.4233474 |
| 5.91301374 | 6.45962238 | 0 | 8.07683317 | 8.23832316 | 6.81576537 | 7.84817157 |
| 2.95650687 | 2.99911039 | 0 | 7.06722902 | 8.23832316 | 9.08768716 | 5.15736989 |
| 23.6520549 | 29.0879103 | 29.1405214 | 29.6116896 | 27.3362541 | 37.1118425 | 29.0001151 |
| 35.4780824 | 13.3806464 | 9.71350712 | 33.3169368 | 23.966031 | 23.8551788 | 26.6837834 |
| 23.6520549 | 19.4942175 | 9.71350712 | 30.2881244 | 23.2170925 | 13.6315307 | 21.30218 |
| 8.8695206 | 5.65216958 | 0 | 14.134458 | 7.48938469 | 6.81576537 | 8.96933894 |
| 32.5215755 | 19.6095679 | 19.4270142 | 25.2401036 | 26.9617849 | 21.583257 | 34.5319549 |
| 8.8695206 | 10.7275872 | 9.71350712 | 9.08643731 | 10.4851386 | 12.4955698 | 6.72700421 |
| 17.7390412 | 13.6113472 | 0 | 11.1056456 | 10.4851386 | 9.08768716 | 13.2297749 |
| 5.91301374 | 14.995552 | 9.71350712 | 5.04802073 | 2.99575388 | 9.08768716 | 10.7632067 |
| 5.91301374 | 5.19076799 | 38.8540285 | 16.1536663 | 10.4851386 | 13.6315307 | 14.7994093 |
| 17.7390412 | 19.4942175 | 9.71350712 | 30.2881244 | 20.2213387 | 29.5349833 | 14.0145921 |
| 5.91301374 | 3.80656319 | 9.71350712 | 26.2497078 | 13.4808924 | 18.1753743 | 11.6601406 |
| 11.8260275 | 15.4569536 | 0 | 9.08643731 | 14.9787694 | 15.9034525 | 11.2116737 |
| 17.7390412 | 15.3416032 | 19.4270142 | 8.07683317 | 14.9787694 | 12.4955698 | 13.6782419 |
| 14.7825343 | 11.8810912 | 19.4270142 | 13.1248539 | 14.2298309 | 7.95172627 | 14.3509423 |
| 8.8695206 | 6.34427198 | 0 | 14.134458 | 6.74044622 | 5.67980448 | 8.29663852 |
| 11.8260275 | 4.26796479 | 0 | 5.04802073 | 9.7362001 | 10.2236481 | 9.64203936 |
| 26.9042125 | 6.11357118 | 19.4270142 | 21.191591 | 10.1106693 | 12.9726734 | 5.94442938 |
| 5.91301374 | 5.99822078 | 0 | 11.1056456 | 6.74044622 | 6.81576537 | 4.93313642 |
| 20.6955481 | 22.2626271 | 19.4270142 | 12.1152497 | 9.7362001 | 20.4472961 | 11.2116737 |
| 29.5650687 | 16.2644064 | 0 | 8.07683317 | 17.2255848 | 30.6709442 | 18.3871448 |
| 23.6520549 | 17.6486112 | 9.71350712 | 21.2016871 | 11.234077 | 24.9911397 | 18.8356118 |
| 11.8260275 | 17.879312 | 9.71350712 | 9.08643731 | 8.23832316 | 6.81576537 | 11.6601406 |

| | | | | | | |
|---|---|---|---|---|---|---|
| 8.8695206 | 24.2235839 | 9.71350712 | 15.1440622 | 26.9617849 | 13.6315307 | 21.9748804 |
| 5.91301374 | 4.72936639 | 0 | 3.53361451 | 8.23832316 | 10.2236481 | 7.28758789 |
| 0 | 17.0718592 | 9.71350712 | 12.1152497 | 17.2255848 | 14.7674916 | 13.4540084 |
| 11.8260275 | 15.1109024 | 29.1405214 | 12.1152497 | 8.23832316 | 11.359609 | 12.1086076 |
| 8.8695206 | 6.69032318 | 9.71350712 | 11.1056456 | 12.731954 | 11.359609 | 11.6601406 |
| 14.7825343 | 3.79156764 | 0 | 8.07683317 | 11.234077 | 4.54384358 | 11.6601406 |
| 11.8260275 | 11.6503904 | 0 | 14.134458 | 8.23832316 | 6.81576537 | 12.781308 |
| 17.7390412 | 24.4542847 | 0 | 15.1440622 | 11.9830155 | 14.7674916 | 19.2840787 |
| 5.91301374 | 65.9804286 | 9.71350712 | 16.1536663 | 14.2298309 | 17.0394134 | 19.7325457 |
| 0 | 1.8456064 | 0 | 10.0960415 | 3.74469235 | 9.08768716 | 6.05430379 |
| 11.8260275 | 13.9573984 | 9.71350712 | 13.1248539 | 14.2298309 | 18.1753743 | 12.1086076 |
| 0 | 6.92102398 | 9.71350712 | 6.05762487 | 8.23832316 | 6.81576537 | 7.39970463 |
| 2.95650687 | 9.45873277 | 0 | 9.08643731 | 4.49363081 | 9.08768716 | 8.96933894 |
| 2.95650687 | 5.19076799 | 0 | 5.04802073 | 2.24681541 | 9.08768716 | 5.15736989 |
| 11.8260275 | 9.45181175 | 0 | 3.02881244 | 8.05108854 | 3.47604034 | 11.240824 |
| 14.7825343 | 14.6495008 | 19.4270142 | 8.07683317 | 7.48938469 | 13.6315307 | 16.1448101 |
| 2.95650687 | 8.42057918 | 0 | 2.01920829 | 8.23832316 | 5.67980448 | 5.60583684 |
| 2.95650687 | 8.93965598 | 0 | 8.07683317 | 8.98726163 | 9.08768716 | 7.39970463 |
| 23.6520549 | 51.5616287 | 19.4270142 | 26.2497078 | 28.4596618 | 34.0788269 | 33.4107876 |
| 17.7390412 | 12.4578432 | 9.71350712 | 12.1152497 | 22.4681541 | 22.7192179 | 21.30218 |
| 32.5215755 | 41.2954431 | 29.1405214 | 37.3553534 | 33.7022311 | 18.1753743 | 31.1684528 |
| 5.91301374 | 4.03726399 | 9.71350712 | 10.0960415 | 4.49363081 | 13.6315307 | 3.13926863 |
| 14.7825343 | 10.1508352 | 29.1405214 | 16.1536663 | 8.23832316 | 14.7674916 | 8.52087199 |
| 8.8695206 | 12.2271424 | 0 | 11.1056456 | 13.4808924 | 18.1753743 | 8.07240505 |
| 8.8695206 | 7.03637438 | 0 | 6.05762487 | 8.98726163 | 3.40788269 | 5.38160336 |
| 20.6955481 | 6.22892158 | 9.71350712 | 13.1248539 | 14.2298309 | 23.8551788 | 6.72700421 |
| 14.7825343 | 43.8331519 | 19.4270142 | 37.3553534 | 32.2043542 | 48.8463185 | 19.7325457 |
| 2.95650687 | 4.03726399 | 19.4270142 | 13.1248539 | 8.98726163 | 7.95172627 | 7.39970463 |
| 5.91301374 | 8.07452798 | 19.4270142 | 14.134458 | 5.99150775 | 11.359609 | 10.0905063 |
| 41.3910961 | 20.8784223 | 9.71350712 | 29.2785202 | 26.2128464 | 17.0394134 | 28.0291842 |
| 2.95650687 | 5.65216958 | 0 | 7.06722902 | 7.48938469 | 6.81576537 | 5.60583684 |
| 14.7825343 | 6.11357118 | 0 | 4.03841658 | 10.4851386 | 9.08768716 | 10.0905063 |
| 742.083224 | 477.435304 | 505.10237 | 608.7913 | 554.214467 | 590.699666 | 749.388269 |
| 8.8695206 | 8.30522878 | 9.71350712 | 1.00960415 | 11.234077 | 2.27192179 | 9.64203936 |
| 26.6085618 | 60.5589598 | 29.1405214 | 14.134458 | 25.4639079 | 32.942866 | 25.562616 |
| 0 | 5.76751998 | 9.71350712 | 8.07683317 | 10.4851386 | 4.54384358 | 8.74510547 |
| 5.91301374 | 5.54835423 | 9.71350712 | 9.8638325 | 10.0432649 | 14.4835014 | 7.25395287 |
| 5.91301374 | 2.99911039 | 0 | 10.0960415 | 8.98726163 | 17.0394134 | 5.60583684 |
| 2.95650687 | 5.07541759 | 0 | 4.03841658 | 10.4851386 | 7.95172627 | 4.03620252 |
| 14.7825343 | 12.2271424 | 19.4270142 | 14.134458 | 11.9830155 | 12.4955698 | 13.6782419 |
| 44.347603 | 32.9902143 | 19.4270142 | 25.2401036 | 21.7192156 | 31.8069051 | 26.6837834 |
| 82.7821923 | 37.7195807 | 106.848578 | 77.7395192 | 53.1746313 | 46.5743967 | 65.7004077 |
| 8.8695206 | 8.30522878 | 0 | 11.1056456 | 5.99150775 | 9.08768716 | 9.86627284 |
| 8.8695206 | 7.26707518 | 9.71350712 | 20.1920829 | 14.2298309 | 13.6315307 | 16.8175105 |
| 23.6520549 | 33.3939407 | 0 | 22.7160933 | 20.9702771 | 22.5261046 | 18.8356118 |
| 38.4345893 | 75.5545118 | 38.8540285 | 61.5858529 | 43.4384312 | 38.6226704 | 65.4761743 |

| | | | | | | |
|---|---|---|---|---|---|---|
| 0 | 2.65305919 | 0 | 3.02881244 | 9.7362001 | 5.67980448 | 7.17547115 |
| 5.91301374 | 32.0674111 | 0 | 9.08643731 | 10.4851386 | 4.54384358 | 7.17547115 |
| 8.8695206 | 3.57586239 | 29.1405214 | 3.02881244 | 8.98726163 | 3.40788269 | 8.74510547 |
| 322.259249 | 201.171097 | 213.697157 | 270.573911 | 211.949587 | 216.968531 | 310.339127 |
| 20.6955481 | 40.9493919 | 19.4270142 | 21.2016871 | 20.9702771 | 21.583257 | 19.2840787 |
| 11.8260275 | 2.65305919 | 0 | 16.1536663 | 10.4851386 | 12.4955698 | 13.4540084 |
| 59.1301374 | 86.7435006 | 82.5648105 | 70.6722902 | 70.0257469 | 71.5655364 | 78.8180659 |
| 0 | 6.57497278 | 0 | 6.05762487 | 8.98726163 | 11.359609 | 6.95123768 |
| 38.4345893 | 50.8695263 | 19.4270142 | 45.4321866 | 32.2043542 | 40.8945922 | 44.1739943 |
| 8.8695206 | 15.4569536 | 9.71350712 | 17.1632705 | 5.24256928 | 4.54384358 | 9.19357241 |
| 2.95650687 | 8.99733118 | 0 | 2.01920829 | 4.49363081 | 9.08768716 | 10.5389733 |
| 29.5650687 | 20.3016703 | 48.5675356 | 13.1248539 | 18.7234617 | 17.0394134 | 19.0598453 |
| 5.91301374 | 5.99822078 | 19.4270142 | 9.08643731 | 7.48938469 | 14.7674916 | 8.74510547 |
| 0 | 11.1889888 | 0 | 7.06722902 | 8.23832316 | 6.81576537 | 8.07240505 |
| 50.2606167 | 80.8606302 | 67.9945498 | 63.6050612 | 47.1831235 | 47.7103576 | 63.2338395 |
| 48.2797571 | 17.7835711 | 9.71350712 | 49.2989704 | 34.9529583 | 37.7706998 | 50.2350251 |
| 26.6085618 | 16.149056 | 9.71350712 | 20.1920829 | 15.7277078 | 10.2236481 | 15.2478762 |
| 50.2606167 | 15.572304 | 9.71350712 | 19.1824788 | 20.9702771 | 20.4472961 | 25.562616 |
| 41.3910961 | 51.5616287 | 19.4270142 | 24.2304995 | 31.4554157 | 30.6709442 | 33.4107876 |
| 45.8258564 | 26.4729167 | 29.1405214 | 44.4225824 | 54.298039 | 48.2783381 | 46.864796 |
| 56.1736305 | 13.265296 | 29.1405214 | 39.3745617 | 35.9490465 | 42.0305531 | 44.9588114 |
| 7.39126717 | 7.49777598 | 9.71350712 | 7.06722902 | 17.2255848 | 31.2389246 | 5.94218705 |
| 51.236264 | 56.9830974 | 24.2837678 | 35.6693145 | 37.6940731 | 33.8857135 | 57.067419 |
| 0 | 11.0736384 | 0 | 11.1056456 | 4.49363081 | 2.27192179 | 6.27853726 |
| 17.7390412 | 18.7444399 | 19.4270142 | 25.7449057 | 21.7192156 | 22.1512375 | 23.6566315 |
| 14.7825343 | 12.8038944 | 0 | 16.1536663 | 6.74044622 | 5.67980448 | 13.0055415 |
| 5.91301374 | 6.80567358 | 0 | 4.03841658 | 6.74044622 | 10.2236481 | 5.83007031 |
| 8.8695206 | 7.03637438 | 0 | 10.0960415 | 8.23832316 | 4.54384358 | 6.72700421 |
| 17.7390412 | 31.6060095 | 38.8540285 | 30.2881244 | 23.2170925 | 29.5349833 | 36.7742897 |
| 2.95650687 | 2.07630719 | 9.71350712 | 4.03841658 | 4.49363081 | 9.08768716 | 4.70890294 |
| 8.8695206 | 12.2271424 | 0 | 5.04802073 | 5.99150775 | 10.2236481 | 7.84817157 |
| 11.8260275 | 15.9183552 | 0 | 13.1248539 | 10.4851386 | 18.1753743 | 9.41780589 |
| 8.8695206 | 9.57408317 | 9.71350712 | 8.07683317 | 12.731954 | 12.4955698 | 10.0905063 |
| 29.5650687 | 34.1437183 | 38.8540285 | 39.3745617 | 41.9405543 | 36.3507487 | 40.8104922 |
| 0 | 3.46051199 | 9.71350712 | 1.00960415 | 11.234077 | 6.81576537 | 8.29663852 |
| 70.9561648 | 95.7408317 | 9.71350712 | 84.8067482 | 68.9023391 | 55.6620839 | 92.1599576 |
| 29.5650687 | 25.2617375 | 0 | 29.2785202 | 31.4554157 | 23.8551788 | 26.9080168 |
| 11.8260275 | 22.8393791 | 9.71350712 | 9.08643731 | 7.48938469 | 12.4955698 | 7.39970463 |
| 20.6955481 | 45.3327071 | 0 | 18.1728746 | 18.7234617 | 14.7674916 | 21.7506469 |
| 23.6520549 | 19.1481663 | 9.71350712 | 18.1728746 | 14.9787694 | 24.9911397 | 16.0326934 |
| 8.8695206 | 15.2262528 | 9.71350712 | 13.1248539 | 14.2298309 | 20.4472961 | 9.86627284 |
| 2.95650687 | 8.53592958 | 0 | 11.1056456 | 8.23832316 | 5.67980448 | 4.70890294 |
| 23.6520549 | 10.381536 | 0 | 16.1536663 | 18.7234617 | 13.6315307 | 16.8175105 |
| 14.7825343 | 25.8384895 | 29.1405214 | 18.1728746 | 21.3447464 | 17.6073939 | 14.1267088 |
| 0 | 5.30611839 | 0 | 5.04802073 | 4.49363081 | 9.08768716 | 4.48466947 |
| 5.91301374 | 1.6149056 | 0 | 10.0960415 | 5.24256928 | 11.359609 | 10.3147398 |

| | | | | | | |
|---|---|---|---|---|---|---|
| 14.7825343 | 23.4161311 | 0 | 32.3073327 | 18.7234617 | 11.359609 | 16.593277 |
| 14.7825343 | 17.5332608 | 0 | 5.04802073 | 6.74044622 | 11.359609 | 7.17547115 |
| 2.95650687 | 3.69121279 | 0 | 5.04802073 | 5.24256928 | 10.2236481 | 4.260436 |
| 26.6085618 | 29.2990015 | 38.8540285 | 34.326541 | 33.7022311 | 21.583257 | 32.5138537 |
| 14.7825343 | 16.3220816 | 0 | 13.1248539 | 7.48938469 | 12.4955698 | 13.4540084 |
| 14.7825343 | 13.3806464 | 0 | 14.134458 | 10.4851386 | 17.0394134 | 10.0905063 |
| 0 | 10.381536 | 0 | 10.0960415 | 11.234077 | 4.54384358 | 10.3147398 |
| 29.5650687 | 24.7230512 | 19.4270142 | 13.1248539 | 15.7277078 | 10.2236481 | 17.4902109 |
| 5.91301374 | 11.53504 | 0 | 7.06722902 | 8.23832316 | 12.4955698 | 5.60583684 |
| 14.7825343 | 34.1437183 | 19.4270142 | 18.1728746 | 13.4808924 | 12.4955698 | 17.9386779 |
| 168.520891 | 114.658297 | 97.1350712 | 141.34458 | 131.064232 | 126.091659 | 141.042855 |
| 133.042809 | 119.733715 | 48.5675356 | 119.133289 | 100.357755 | 88.6049498 | 128.261547 |
| 8.8695206 | 9.22803198 | 0 | 11.1056456 | 14.2298309 | 15.9034525 | 11.2116737 |
| 11.8260275 | 7.15172478 | 9.71350712 | 4.03841658 | 13.1064232 | 14.1995112 | 13.1176582 |
| 4.4347603 | 8.24755358 | 14.5702607 | 13.1248539 | 15.7277078 | 13.0635503 | 10.8753235 |
| 8.8695206 | 16.6104576 | 0 | 9.08643731 | 7.48938469 | 13.6315307 | 8.96933894 |
| 5.91301374 | 7.03637438 | 0 | 11.1056456 | 5.99150775 | 7.95172627 | 10.9874402 |
| 2.95650687 | 13.842048 | 9.71350712 | 7.06722902 | 11.234077 | 12.4955698 | 13.6782419 |
| 5.91301374 | 5.99822078 | 0 | 6.05762487 | 8.23832316 | 5.67980448 | 8.29663852 |
| 2.95650687 | 7.38242558 | 9.71350712 | 8.07683317 | 4.49363081 | 9.08768716 | 4.70890294 |
| 5.91301374 | 4.26796479 | 0 | 6.05762487 | 4.49363081 | 9.08768716 | 5.15736989 |
| 14.7825343 | 13.4959968 | 19.4270142 | 14.134458 | 11.9830155 | 14.7674916 | 17.7144444 |
| 5.91301374 | 4.72936639 | 9.71350712 | 8.07683317 | 13.4808924 | 17.0394134 | 12.332841 |
| 2.95650687 | 10.4968864 | 9.71350712 | 4.03841658 | 8.23832316 | 11.359609 | 7.17547115 |
| 8.8695206 | 8.53592958 | 0 | 9.08643731 | 6.74044622 | 14.7674916 | 13.4540084 |
| 5.91301374 | 5.53681919 | 0 | 11.1056456 | 3.74469235 | 5.67980448 | 7.17547115 |
| 5.91301374 | 21.3398239 | 0 | 30.2881244 | 18.7234617 | 11.359609 | 20.4052461 |
| 11.8260275 | 10.958288 | 19.4270142 | 24.2304995 | 27.3362541 | 23.2871984 | 18.9477285 |
| 2.95650687 | 6.45962238 | 0 | 4.03841658 | 4.49363081 | 9.08768716 | 4.70890294 |
| 29.5650687 | 27.7994463 | 29.1405214 | 29.2785202 | 36.697985 | 34.0788269 | 34.9804219 |
| 26.6085618 | 10.0354848 | 0 | 15.1440622 | 14.2298309 | 7.95172627 | 16.593277 |
| 2.95650687 | 33.3362655 | 0 | 6.05762487 | 10.4851386 | 5.67980448 | 8.74510547 |
| 38.4345893 | 38.7577343 | 58.2810427 | 44.4225824 | 32.2043542 | 27.2630615 | 36.7742897 |
| 5.91301374 | 10.8429376 | 19.4270142 | 18.1728746 | 15.7277078 | 24.9911397 | 7.84817157 |
| 16.7633939 | 27.6506443 | 32.3459787 | 31.9741633 | 28.8565992 | 33.3177331 | 32.475734 |
| 5.91301374 | 8.93965598 | 0 | 11.1056456 | 4.86810005 | 7.38374582 | 7.28758789 |
| 38.4345893 | 27.7994463 | 38.8540285 | 51.3181787 | 45.9323963 | 62.4778492 | 54.8250843 |
| 14.7825343 | 5.53681919 | 0 | 3.02881244 | 10.4851386 | 6.81576537 | 7.39970463 |
| 2.95650687 | 9.45873277 | 9.71350712 | 5.04802073 | 7.48938469 | 11.359609 | 8.96933894 |
| 11.8260275 | 14.3034496 | 19.4270142 | 8.07683317 | 7.48938469 | 9.08768716 | 6.05430379 |
| 11.8260275 | 3.46051199 | 9.71350712 | 3.02881244 | 5.99150775 | 10.2236481 | 6.50277073 |
| 20.6955481 | 26.9919935 | 19.4270142 | 26.2497078 | 22.4681541 | 21.583257 | 30.7199859 |
| 14.7825343 | 23.5314815 | 9.71350712 | 27.2593119 | 18.7234617 | 17.0394134 | 15.2478762 |
| 17.7390412 | 43.1410495 | 48.5675356 | 38.3649575 | 20.9702771 | 27.2630615 | 42.7164767 |
| 2.95650687 | 8.07452798 | 9.71350712 | 3.02881244 | 8.23832316 | 2.27192179 | 5.83007031 |
| 5.91301374 | 4.38331519 | 0 | 5.04802073 | 11.9830155 | 6.81576537 | 7.84817157 |

| | | | | | | |
|---|---|---|---|---|---|---|
| 8.8695206 | 8.42057918 | 9.71350712 | 11.1056456 | 12.731954 | 11.359609 | 14.3509423 |
| 8.8695206 | 13.0345952 | 19.4270142 | 12.1152497 | 16.4766463 | 10.2236481 | 20.1810126 |
| 20.6955481 | 14.7648512 | 0 | 5.04802073 | 22.4681541 | 13.6315307 | 18.6113783 |
| 20.6955481 | 47.0629631 | 29.1405214 | 36.3457492 | 23.966031 | 26.1271006 | 34.7561884 |
| 0 | 1.8456064 | 9.71350712 | 7.06722902 | 5.24256928 | 9.08768716 | 2.91503516 |
| 8.8695206 | 6.69032318 | 0 | 3.02881244 | 8.23832316 | 5.67980448 | 7.17547115 |
| 17.7390412 | 10.4968864 | 0 | 18.1728746 | 10.8596078 | 11.359609 | 12.5570745 |
| 8.8695206 | 16.9565088 | 19.4270142 | 17.1632705 | 34.4511696 | 28.3990224 | 17.7144444 |
| 2.95650687 | 11.1889888 | 0 | 7.06722902 | 9.7362001 | 6.81576537 | 8.07240505 |
| 13.3042809 | 10.09316 | 0 | 13.4580233 | 7.73653438 | 19.3113352 | 7.51182136 |
| 7.39126717 | 9.45873277 | 29.1405214 | 19.1824788 | 21.7192156 | 21.0152766 | 17.9386779 |
| 47.3041099 | 14.3034496 | 29.1405214 | 32.3073327 | 33.7022311 | 38.6226704 | 34.9804219 |
| 0 | 13.0345952 | 0 | 8.07683317 | 10.4851386 | 4.54384358 | 8.96933894 |
| 0 | 4.49866559 | 9.71350712 | 13.1248539 | 3.74469235 | 6.81576537 | 4.93313642 |
| 8.8695206 | 8.88198078 | 0 | 10.0960415 | 10.8596078 | 12.4955698 | 11.5480239 |
| 11.8260275 | 9.45873277 | 9.71350712 | 4.03841658 | 14.9787694 | 12.4955698 | 12.5570745 |
| 2.95650687 | 2.88375999 | 0 | 8.58163524 | 7.73653438 | 6.81576537 | 6.390654 |
| 14.7825343 | 10.0354848 | 9.71350712 | 17.1632705 | 18.7234617 | 13.6315307 | 10.9874402 |
| 8.8695206 | 9.22803198 | 19.4270142 | 8.07683317 | 6.74044622 | 9.08768716 | 6.27853726 |
| 20.6955481 | 7.38242558 | 29.1405214 | 17.1632705 | 12.731954 | 15.9034525 | 25.114149 |
| 29.5650687 | 13.7266976 | 19.4270142 | 23.2208954 | 17.9745233 | 14.7674916 | 20.1810126 |
| 5.91301374 | 7.38242558 | 9.71350712 | 15.1440622 | 8.98726163 | 5.67980448 | 9.86627284 |
| 2.95650687 | 4.05687356 | 0 | 11.1056456 | 8.98726163 | 6.24778492 | 8.52087199 |
| 5.91301374 | 6.57497278 | 9.71350712 | 13.1248539 | 16.4766463 | 14.7674916 | 14.7994093 |
| 8.8695206 | 9.92013437 | 0 | 15.1440622 | 11.9830155 | 9.08768716 | 14.3509423 |
| 14.7825343 | 7.03637438 | 38.8540285 | 16.1536663 | 12.3574847 | 12.4955698 | 15.9205766 |
| 11.8260275 | 8.18987838 | 0 | 3.02881244 | 8.98726163 | 9.08768716 | 8.74510547 |
| 8.8695206 | 7.61312638 | 0 | 12.1152497 | 13.4808924 | 15.9034525 | 8.07240505 |
| 11.8260275 | 8.18987838 | 0 | 14.134458 | 16.4766463 | 23.8551788 | 20.6294796 |
| 5.91301374 | 15.3416032 | 0 | 13.1248539 | 5.24256928 | 9.08768716 | 8.29663852 |
| 11.8260275 | 11.9964416 | 9.71350712 | 17.6680725 | 13.4808924 | 14.1995112 | 9.52992263 |
| 8.8695206 | 10.6122368 | 0 | 23.2208954 | 10.4851386 | 18.1753743 | 15.2478762 |
| 1.47825343 | 4.44099039 | 0 | 9.08643731 | 14.9787694 | 12.4955698 | 2.91503516 |
| 5.91301374 | 2.88375999 | 0 | 4.03841658 | 5.99150775 | 10.2236481 | 6.50277073 |
| 8.8695206 | 7.95917758 | 0 | 6.05762487 | 9.7362001 | 4.54384358 | 7.69793515 |
| 20.6955481 | 26.0691903 | 0 | 17.1632705 | 10.4851386 | 7.95172627 | 11.6601406 |
| 23.6520549 | 14.4188 | 9.71350712 | 6.05762487 | 7.48938469 | 7.95172627 | 13.0055415 |
| 8.8695206 | 10.0354848 | 0 | 15.1440622 | 5.24256928 | 6.81576537 | 7.84817157 |
| 11.8260275 | 7.61312638 | 0 | 4.03841658 | 5.24256928 | 12.4955698 | 3.3635021 |
| 8.8695206 | 7.95917758 | 19.4270142 | 7.06722902 | 11.234077 | 12.4955698 | 12.1086076 |
| 11.8260275 | 22.3779775 | 0 | 13.1248539 | 14.2298309 | 9.08768716 | 13.0055415 |
| 23.6520549 | 38.8730847 | 9.71350712 | 35.3361451 | 27.7107234 | 29.5349833 | 26.4595499 |
| 11.8260275 | 1.6149056 | 9.71350712 | 2.01920829 | 8.98726163 | 11.359609 | 5.83007031 |
| 5.91301374 | 3.34516159 | 9.71350712 | 5.04802073 | 7.48938469 | 1.1359609 | 4.03620252 |
| 5.91301374 | 16.725808 | 19.4270142 | 15.1440622 | 8.23832316 | 13.6315307 | 9.64203936 |
| 11.8260275 | 7.72847678 | 9.71350712 | 10.0960415 | 7.48938469 | 13.6315307 | 10.5389733 |

| | | | | | | |
|---|---|---|---|---|---|---|
| 13.3042809 | 4.72936639 | 0 | 5.5528228 | 8.98726163 | 6.81576537 | 7.6239381 |
| 50.2606167 | 44.1792031 | 0 | 40.3841658 | 27.7107234 | 32.942866 | 37.6712236 |
| 17.7390412 | 28.3761983 | 9.71350712 | 18.1728746 | 23.966031 | 10.2236481 | 17.2659775 |
| 14.7825343 | 8.76663038 | 9.71350712 | 6.05762487 | 11.234077 | 9.08768716 | 7.6239381 |
| 5.91301374 | 18.3407136 | 9.71350712 | 7.06722902 | 8.98726163 | 11.359609 | 12.1086076 |
| 0 | 5.76751998 | 9.71350712 | 3.02881244 | 8.98726163 | 5.67980448 | 6.95123768 |
| 23.6520549 | 42.5642975 | 19.4270142 | 23.2208954 | 19.4724002 | 12.4955698 | 23.0960478 |
| 8.8695206 | 9.45873277 | 19.4270142 | 14.134458 | 7.48938469 | 14.7674916 | 13.4741894 |
| 2.95650687 | 8.88198078 | 9.71350712 | 11.1056456 | 7.48938469 | 4.54384358 | 11.3977875 |
| 2.95650687 | 6.34427198 | 19.4270142 | 7.06722902 | 8.98726163 | 9.08768716 | 13.4540084 |
| 8.8695206 | 10.8429376 | 0 | 4.03841658 | 8.98726163 | 4.54384358 | 4.260436 |
| 11.8260275 | 7.49777598 | 19.4270142 | 13.1248539 | 11.9830155 | 12.4955698 | 12.781308 |
| 29.5650687 | 26.9919935 | 9.71350712 | 19.6872808 | 22.4681541 | 20.4472961 | 21.30218 |
| 0 | 8.42057918 | 0 | 5.04802073 | 4.49363081 | 9.08768716 | 6.95123768 |
| 8.8695206 | 8.07452798 | 9.71350712 | 17.1632705 | 12.731954 | 21.583257 | 13.2297749 |
| 8.8695206 | 10.2661856 | 9.71350712 | 13.1248539 | 8.98726163 | 17.0394134 | 5.38160336 |
| 14.7825343 | 4.44099039 | 19.4270142 | 9.08643731 | 10.4851386 | 15.9034525 | 11.8843741 |
| 13.3042809 | 14.995552 | 4.85675356 | 11.6104477 | 8.23832316 | 17.0394134 | 13.5661251 |
| 2.95650687 | 17.0718592 | 0 | 9.08643731 | 8.98726163 | 3.40788269 | 6.95123768 |
| 11.8260275 | 10.4968864 | 9.71350712 | 9.08643731 | 7.48938469 | 4.54384358 | 6.50277073 |
| 8.8695206 | 22.2626271 | 0 | 19.1824788 | 23.966031 | 42.0305531 | 11.8843741 |
| 11.8260275 | 10.0354848 | 0 | 20.1920829 | 12.731954 | 7.95172627 | 15.0236427 |
| 2.95650687 | 7.49777598 | 9.71350712 | 6.05762487 | 8.98726163 | 5.67980448 | 8.29663852 |
| 20.6955481 | 14.5341504 | 9.71350712 | 25.2401036 | 29.9575388 | 23.8551788 | 22.1991139 |
| 38.4345893 | 110.505683 | 19.4270142 | 25.2401036 | 35.9490465 | 22.7192179 | 33.8592545 |
| 47.3041099 | 69.7869918 | 19.4270142 | 41.39377 | 51.6767544 | 43.166514 | 50.0040646 |
| 41.3910961 | 23.6468319 | 38.8540285 | 42.4033741 | 38.1958619 | 27.2630615 | 35.8773558 |
| 2.95650687 | 19.4942175 | 9.71350712 | 13.1248539 | 13.4808924 | 2.27192179 | 9.64203936 |
| 0 | 3.92191359 | 0 | 7.06722902 | 8.98726163 | 11.359609 | 6.05430379 |
| 5.91301374 | 2.99911039 | 19.4270142 | 13.1248539 | 13.4808924 | 18.1753743 | 14.5751758 |
| 0 | 16.6104576 | 0 | 5.04802073 | 5.99150775 | 9.08768716 | 7.17547115 |
| 5.91301374 | 11.6503904 | 0 | 19.1824788 | 9.7362001 | 14.7674916 | 11.6601406 |
| 5.91301374 | 12.688544 | 9.71350712 | 14.134458 | 6.74044622 | 13.6315307 | 12.332841 |
| 23.6520549 | 17.879312 | 9.71350712 | 20.1920829 | 18.7234617 | 14.7674916 | 17.2659775 |
| 11.8260275 | 6.69032318 | 0 | 14.134458 | 10.4851386 | 10.2236481 | 9.19357241 |
| 14.7825343 | 8.18987838 | 0 | 3.02881244 | 17.9745233 | 10.2236481 | 15.4721097 |
| 56.1736305 | 101.162301 | 19.4270142 | 37.3553534 | 40.4426773 | 31.8069051 | 41.2589591 |
| 0 | 11.6503904 | 9.71350712 | 4.03841658 | 9.7362001 | 7.95172627 | 5.60583684 |
| 62.0866442 | 49.8313727 | 77.708057 | 53.5090197 | 62.9108314 | 45.4384358 | 69.4002601 |
| 67.999658 | 45.0247215 | 38.8540285 | 66.6338736 | 48.6810005 | 80.4601102 | 50.8628788 |
| 8.8695206 | 12.4382336 | 9.71350712 | 18.6776767 | 13.608212 | 27.8310419 | 12.781308 |
| 94.6082198 | 32.5288127 | 77.708057 | 61.5858529 | 59.1661391 | 71.5655364 | 74.4455132 |
| 4.4347603 | 6.05589598 | 0 | 15.1440622 | 11.234077 | 12.4955698 | 12.332841 |
| 29.5650687 | 9.68943357 | 9.71350712 | 17.1632705 | 23.966031 | 21.583257 | 26.0110829 |
| 7.39126717 | 6.17124638 | 9.71350712 | 10.6008435 | 10.1106693 | 6.62265202 | 3.81196905 |
| 11.8260275 | 13.1499456 | 0 | 10.0960415 | 19.4724002 | 13.6315307 | 15.9205766 |

| | | | | | | |
|---|---|---|---|---|---|---|
| 44.347603 | 27.9147967 | 9.71350712 | 21.2016871 | 24.7149695 | 27.2630615 | 23.9929817 |
| 14.7825343 | 11.4196896 | 0 | 15.1440622 | 11.234077 | 14.7674916 | 13.0055415 |
| 17.7390412 | 149.840169 | 9.71350712 | 24.2304995 | 19.4724002 | 17.0394134 | 24.2172151 |
| 5.91301374 | 20.1863199 | 0 | 20.1920829 | 14.9787694 | 13.6315307 | 22.1991139 |
| 11.8260275 | 16.6104576 | 9.71350712 | 10.0960415 | 10.4851386 | 18.1753743 | 11.4359072 |
| 5.91301374 | 9.34338237 | 29.1405214 | 10.0960415 | 14.9787694 | 9.08768716 | 9.19357241 |
| 5.91301374 | 11.4196896 | 0 | 7.06722902 | 6.74044622 | 10.2236481 | 4.48466947 |
| 11.8260275 | 1.6149056 | 19.4270142 | 11.1056456 | 9.7362001 | 12.4955698 | 5.38160336 |
| 8.8695206 | 18.3407136 | 0 | 21.2016871 | 14.9787694 | 10.2236481 | 15.2478762 |
| 8.8695206 | 6.80567358 | 0 | 9.08643731 | 14.9787694 | 15.9034525 | 28.1413009 |
| 2.95650687 | 9.45873277 | 19.4270142 | 5.04802073 | 10.4851386 | 1.1359609 | 5.60583684 |
| 11.8260275 | 12.5731936 | 9.71350712 | 13.1248539 | 8.23832316 | 4.54384358 | 7.6239381 |
| 0 | 2.76840959 | 9.71350712 | 4.54321866 | 7.48938469 | 3.97586313 | 3.81196905 |
| 50.2606167 | 34.1437183 | 48.5675356 | 46.4417907 | 50.9278159 | 42.0305531 | 56.2826019 |
| 17.7390412 | 18.2253632 | 29.1405214 | 18.1728746 | 17.9745233 | 23.8551788 | 21.9748804 |
| 8.8695206 | 10.2661856 | 0 | 4.03841658 | 7.48938469 | 3.40788269 | 6.05430379 |
| 233.564043 | 181.330828 | 262.264692 | 276.631536 | 199.966571 | 164.71433 | 189.477285 |
| 5.91301374 | 10.1508352 | 0 | 3.02881244 | 5.24256928 | 12.4955698 | 4.48466947 |
| 20.6955481 | 17.311788 | 9.71350712 | 14.134458 | 15.7277078 | 21.583257 | 25.3383825 |
| 0 | 1.730256 | 0 | 4.03841658 | 4.49363081 | 11.359609 | 4.48466947 |
| 5.91301374 | 7.84382718 | 0 | 4.03841658 | 7.48938469 | 4.54384358 | 7.6239381 |
| 8.8695206 | 8.42057918 | 9.71350712 | 7.06722902 | 7.48938469 | 6.81576537 | 8.74510547 |
| 14.7825343 | 4.38331519 | 9.71350712 | 10.0960415 | 5.99150775 | 11.359609 | 10.3147398 |
| 26.6085618 | 22.9547295 | 9.71350712 | 13.1248539 | 15.7277078 | 17.0394134 | 20.1810126 |
| 2.95650687 | 15.1109024 | 0 | 14.134458 | 8.23832316 | 4.54384358 | 13.0055415 |
| 14.7825343 | 47.6397151 | 19.4270142 | 19.1824788 | 21.7192156 | 15.9034525 | 21.5264135 |
| 79.8256854 | 95.7408317 | 48.5675356 | 61.5858529 | 39.6937389 | 61.3418884 | 81.3967509 |
| 8.8695206 | 39.6805375 | 0 | 15.1440622 | 13.4808924 | 17.0394134 | 17.041744 |
| 17.7390412 | 20.5323711 | 9.71350712 | 17.1632705 | 17.2255848 | 22.7192179 | 6.50277073 |
| 2.95650687 | 5.65216958 | 9.71350712 | 2.01920829 | 4.49363081 | 9.08768716 | 4.48466947 |
| 8.8695206 | 31.1446079 | 19.4270142 | 20.1920829 | 16.4766463 | 17.0394134 | 18.8356118 |
| 26.6085618 | 23.3007807 | 9.71350712 | 15.1440622 | 10.4851386 | 17.0394134 | 15.0236427 |
| 5.91301374 | 28.8375999 | 0 | 3.02881244 | 10.4851386 | 1.1359609 | 6.95123768 |
| 5.91301374 | 10.958288 | 0 | 2.01920829 | 15.7277078 | 9.08768716 | 15.6963431 |
| 8.8695206 | 25.6654639 | 9.71350712 | 19.6872808 | 11.9830155 | 13.6315307 | 15.2478762 |
| 2.95650687 | 12.3424928 | 0 | 11.1056456 | 7.48938469 | 9.08768716 | 7.84817157 |
| 0 | 6.11357118 | 9.71350712 | 7.06722902 | 7.48938469 | 11.359609 | 4.93313642 |
| 70.9561648 | 60.5589598 | 48.5675356 | 57.5474363 | 67.4044622 | 53.3901621 | 60.7672713 |
| 8.8695206 | 19.0328159 | 19.4270142 | 8.07683317 | 10.4851386 | 5.67980448 | 10.5389733 |
| 14.7825343 | 32.4134623 | 0 | 7.06722902 | 13.4808924 | 2.27192179 | 10.0905063 |
| 2.95650687 | 9.92013437 | 19.4270142 | 16.1536663 | 15.7277078 | 20.4472961 | 15.6963431 |
| 17.7390412 | 23.1854303 | 0 | 10.0960415 | 18.7234617 | 15.9034525 | 17.4902109 |
| 11.8260275 | 27.2226943 | 19.4270142 | 25.2401036 | 17.9745233 | 17.0394134 | 17.7144444 |
| 11.8260275 | 10.0354848 | 9.71350712 | 7.06722902 | 8.23832316 | 14.7674916 | 10.7632067 |
| 2.95650687 | 14.4188 | 9.71350712 | 10.0960415 | 8.23832316 | 7.95172627 | 8.74510547 |
| 11.8260275 | 6.92102398 | 0 | 7.06722902 | 6.74044622 | 11.359609 | 11.5480239 |

| | | | | | | |
|---|---|---|---|---|---|---|
| 14.7825343 | 10.7275872 | 19.4270142 | 10.0960415 | 8.23832316 | 9.08768716 | 11.6601406 |
| 5.91301374 | 9.22803198 | 19.4270142 | 2.01920829 | 8.23832316 | 15.9034525 | 5.83007031 |
| 8.8695206 | 6.11357118 | 19.4270142 | 9.08643731 | 16.4766463 | 13.6315307 | 10.5389733 |
| 5.91301374 | 2.76840959 | 0 | 8.07683317 | 2.24681541 | 9.08768716 | 4.48466947 |
| 38.4345893 | 8.76663038 | 9.71350712 | 25.2401036 | 37.4469235 | 34.0788269 | 25.7868495 |
| 17.7390412 | 35.5279231 | 9.71350712 | 14.134458 | 20.9702771 | 13.6315307 | 19.3961955 |
| 5.91301374 | 4.49866559 | 0 | 4.87638802 | 12.3574847 | 11.1664956 | 5.45560041 |
| 23.6520549 | 11.7657408 | 19.4270142 | 11.1056456 | 17.9745233 | 18.1753743 | 11.8843741 |
| 8.8695206 | 4.61401599 | 0 | 6.05762487 | 9.7362001 | 10.2236481 | 10.202623 |
| 5.91301374 | 7.38242558 | 0 | 5.04802073 | 4.49363081 | 9.08768716 | 6.05430379 |
| 5.91301374 | 6.92102398 | 9.71350712 | 4.03841658 | 13.4808924 | 4.54384358 | 9.64203936 |
| 8.8695206 | 4.15261439 | 0 | 5.04802073 | 8.23832316 | 4.54384358 | 5.15736989 |
| 32.5215755 | 64.365523 | 38.8540285 | 41.39377 | 39.6937389 | 28.3990224 | 37.895457 |
| 8.8695206 | 22.3779775 | 0 | 8.07683317 | 11.234077 | 4.54384358 | 8.07240505 |
| 5.91301374 | 16.149056 | 0 | 9.08643731 | 11.234077 | 14.7674916 | 10.7632067 |
| 29.5650687 | 42.6796479 | 38.8540285 | 24.2304995 | 25.4639079 | 32.942866 | 28.0291842 |
| 8.8695206 | 6.61303841 | 9.71350712 | 7.06722902 | 16.4766463 | 19.8793157 | 8.29663852 |
| 32.5215755 | 46.7169119 | 48.5675356 | 35.3361451 | 69.0521268 | 43.166514 | 57.4037692 |
| 41.3910961 | 63.5580702 | 29.1405214 | 50.4802073 | 47.932062 | 51.1182403 | 61.6642052 |
| 8.8695206 | 22.5510031 | 0 | 9.08643731 | 10.1106693 | 6.81576537 | 8.18452178 |
| 2.95650687 | 8.65127998 | 0 | 10.0960415 | 4.49363081 | 11.359609 | 10.9874402 |
| 8.8695206 | 4.96006719 | 0 | 11.6104477 | 7.48938469 | 5.11182403 | 5.83007031 |
| 14.7825343 | 5.76751998 | 9.71350712 | 8.07683317 | 8.98726163 | 5.67980448 | 10.0905063 |
| 5.91301374 | 20.3016703 | 0 | 9.08643731 | 8.23832316 | 4.54384358 | 11.2116737 |
| 8.8695206 | 15.1109024 | 19.4270142 | 13.1248539 | 8.98726163 | 3.40788269 | 9.64203936 |
| 14.7825343 | 24.4669733 | 0 | 12.1152497 | 7.61670423 | 9.08768716 | 14.5931145 |
| 2.95650687 | 13.0345952 | 9.71350712 | 7.06722902 | 12.731954 | 14.7674916 | 12.781308 |
| 8.8695206 | 13.3806464 | 19.4270142 | 9.08643731 | 14.2298309 | 15.9034525 | 10.5389733 |
| 8.8695206 | 4.03726399 | 0 | 2.01920829 | 9.7362001 | 9.08768716 | 8.29663852 |
| 121.216782 | 64.500483 | 63.1377963 | 82.2827379 | 98.8598779 | 96.9315432 | 105.501849 |
| 17.7390412 | 22.1472767 | 9.71350712 | 25.2401036 | 17.9745233 | 14.7674916 | 20.4052461 |
| 0 | 18.5714143 | 19.4270142 | 18.1728746 | 12.731954 | 10.2236481 | 20.4052461 |
| 62.0866442 | 29.7604031 | 77.708057 | 93.8931855 | 52.4256928 | 64.749771 | 73.5485793 |
| 10.347774 | 13.0922704 | 0 | 10.6008435 | 10.4851386 | 12.4955698 | 10.7632067 |
| 2.95650687 | 4.65208162 | 9.71350712 | 6.05762487 | 7.48938469 | 7.95172627 | 9.41780589 |
| 5.91301374 | 9.11268158 | 9.71350712 | 11.1056456 | 16.4766463 | 10.2236481 | 8.52087199 |
| 85.7386992 | 64.8269246 | 67.9945498 | 72.6914985 | 60.664016 | 59.0699666 | 68.6154429 |
| 14.7825343 | 11.4196896 | 0 | 4.03841658 | 11.234077 | 17.0394134 | 17.041744 |
| 260.172604 | 151.570425 | 271.978199 | 377.59195 | 327.286111 | 251.047358 | 291.055049 |
| 14.7825343 | 30.4525055 | 9.71350712 | 26.2497078 | 28.4596618 | 49.9822794 | 17.7144444 |
| 32.5215755 | 17.3602352 | 19.4270142 | 30.2881244 | 29.9575388 | 22.7192179 | 29.5988185 |
| 17.7390412 | 18.5714143 | 29.1405214 | 11.1056456 | 12.731954 | 15.9034525 | 15.2478762 |
| 8.8695206 | 10.958288 | 0 | 3.02881244 | 9.7362001 | 4.54384358 | 8.96933894 |
| 2.95650687 | 4.72936639 | 0 | 6.05762487 | 8.23832316 | 17.0394134 | 8.29663852 |
| 5.91301374 | 30.5678559 | 9.71350712 | 14.134458 | 9.7362001 | 6.81576537 | 6.390654 |
| 159.651371 | 142.169368 | 38.8540285 | 117.114081 | 125.447194 | 103.940422 | 145.303291 |

| | | | | | | |
|---|---|---|---|---|---|---|
| 17.7390412 | 4.49866559 | 9.71350712 | 18.1728746 | 11.9830155 | 23.8551788 | 15.4721097 |
| 5.91301374 | 9.11268158 | 0 | 9.08643731 | 8.98726163 | 6.81576537 | 7.39970463 |
| 2.95650687 | 8.76663038 | 9.71350712 | 2.01920829 | 5.24256928 | 9.08768716 | 4.70890294 |
| 11.8260275 | 9.22803198 | 9.71350712 | 10.0960415 | 8.23832316 | 9.08768716 | 7.39970463 |
| 14.7825343 | 18.6867647 | 9.71350712 | 19.1824788 | 31.4554157 | 21.583257 | 23.7687482 |
| 5.91301374 | 12.3424928 | 19.4270142 | 13.1248539 | 14.9787694 | 22.7192179 | 16.8175105 |
| 20.6955481 | 11.8810912 | 9.71350712 | 9.08643731 | 3.74469235 | 9.08768716 | 10.9874402 |
| 17.7390412 | 13.265296 | 9.71350712 | 17.1632705 | 14.9787694 | 9.08768716 | 11.8843741 |
| 11.8260275 | 7.72847678 | 19.4270142 | 6.05762487 | 8.23832316 | 6.81576537 | 9.64203936 |
| 11.8260275 | 27.1073439 | 9.71350712 | 11.1056456 | 17.2255848 | 6.81576537 | 15.2478762 |
| 38.4345893 | 42.7373231 | 19.4270142 | 44.4225824 | 27.3362541 | 34.6468073 | 41.2589591 |
| 17.7390412 | 18.9174655 | 19.4270142 | 15.1440622 | 17.9745233 | 19.3113352 | 20.6294796 |
| 11.8260275 | 22.9547295 | 9.71350712 | 20.1920829 | 14.9787694 | 34.0788269 | 23.9929817 |
| 29.5650687 | 28.3761983 | 9.71350712 | 28.2689161 | 14.2298309 | 18.1753743 | 23.5445147 |
| 5.91301374 | 2.88375999 | 0 | 3.02881244 | 8.23832316 | 6.81576537 | 4.03620252 |
| 23.6520549 | 39.2191359 | 19.4270142 | 31.2977285 | 29.9575388 | 23.8551788 | 35.4288888 |
| 88.695206 | 71.6325982 | 67.9945498 | 71.6818943 | 64.4087083 | 54.526123 | 68.8396764 |
| 5.91301374 | 15.8030048 | 0 | 8.07683317 | 9.7362001 | 7.95172627 | 14.5751758 |
| 11.8260275 | 5.76751998 | 0 | 11.1056456 | 9.7362001 | 14.7674916 | 7.39970463 |
| 8.8695206 | 11.9964416 | 0 | 9.08643731 | 3.74469235 | 11.359609 | 10.9874402 |
| 5.91301374 | 13.265296 | 0 | 14.134458 | 8.98726163 | 11.359609 | 13.0055415 |
| 8.8695206 | 8.18987838 | 0 | 5.04802073 | 5.99150775 | 11.359609 | 8.96933894 |
| 20.6955481 | 47.8704159 | 29.1405214 | 27.2593119 | 25.4639079 | 14.7674916 | 21.9748804 |
| 0 | 14.707176 | 19.4270142 | 13.629656 | 16.8511156 | 14.7674916 | 6.83912094 |
| 35.4780824 | 43.0256991 | 0 | 14.134458 | 24.7149695 | 20.4472961 | 35.2046553 |
| 93.1299663 | 75.035435 | 87.4215641 | 85.8163524 | 77.1406623 | 67.5896733 | 83.9754358 |
| 0 | 2.42235839 | 0 | 8.07683317 | 8.61279239 | 9.08768716 | 5.4937201 |
| 2.95650687 | 5.19076799 | 9.71350712 | 8.07683317 | 7.48938469 | 4.54384358 | 5.38160336 |
| 2.95650687 | 2.65305919 | 9.71350712 | 6.05762487 | 5.24256928 | 9.08768716 | 7.39970463 |
| 13.3042809 | 9.22803198 | 0 | 19.1824788 | 11.234077 | 11.359609 | 12.781308 |
| 8.8695206 | 20.3016703 | 0 | 29.2785202 | 28.4596618 | 29.5349833 | 26.2353164 |
| 2.95650687 | 1.6149056 | 9.71350712 | 8.07683317 | 10.4851386 | 9.08768716 | 8.96933894 |
| 56.1736305 | 55.1374911 | 58.2810427 | 110.046852 | 88.3747393 | 77.2453409 | 104.044332 |
| 14.7825343 | 14.0727488 | 9.71350712 | 8.07683317 | 17.2255848 | 21.583257 | 20.853713 |
| 11.8260275 | 13.265296 | 0 | 5.04802073 | 8.98726163 | 6.81576537 | 12.1086076 |
| 32.5215755 | 23.4161311 | 19.4270142 | 18.1728746 | 26.2128464 | 17.0394134 | 23.9929817 |
| 8.8695206 | 7.61312638 | 0 | 16.1536663 | 6.74044622 | 7.95172627 | 8.96933894 |
| 14.7825343 | 12.688544 | 19.4270142 | 14.134458 | 14.2298309 | 23.8551788 | 17.2659775 |
| 11.8260275 | 28.0301471 | 9.71350712 | 13.1248539 | 8.98726163 | 12.4955698 | 9.86627284 |
| 29.5650687 | 45.2173567 | 0 | 35.3361451 | 29.2086003 | 22.7192179 | 25.114149 |
| 26.6085618 | 33.4516159 | 38.8540285 | 28.2689161 | 24.7149695 | 24.9911397 | 31.6169198 |
| 2.95650687 | 9.22803198 | 0 | 6.05762487 | 7.48938469 | 6.81576537 | 6.72700421 |
| 8.8695206 | 7.15172478 | 9.71350712 | 6.05762487 | 9.7362001 | 4.54384358 | 6.05430379 |
| 11.8260275 | 13.265296 | 9.71350712 | 21.2016871 | 11.234077 | 15.9034525 | 13.2297749 |
| 11.8260275 | 12.8038944 | 9.71350712 | 12.1152497 | 8.23832316 | 7.95172627 | 11.2116737 |
| 35.4780824 | 22.8393791 | 19.4270142 | 24.2304995 | 14.2298309 | 18.1753743 | 18.1629114 |

| | | | | | | |
|---|---|---|---|---|---|---|
| 5.91301374 | 17.0718592 | 9.71350712 | 12.1152497 | 11.234077 | 11.359609 | 11.4359072 |
| 5.91301374 | 4.49866559 | 0 | 3.02881244 | 6.74044622 | 9.08768716 | 6.05430379 |
| 11.8260275 | 6.45962238 | 10.2963175 | 10.0960415 | 17.2255848 | 13.6315307 | 12.1086076 |
| 2.95650687 | 3.46051199 | 9.71350712 | 11.1056456 | 8.23832316 | 9.08768716 | 10.9874402 |
| 17.7390412 | 25.2617375 | 38.8540285 | 24.2304995 | 21.7192156 | 31.8069051 | 30.7199859 |
| 2.95650687 | 2.99911039 | 0 | 12.1152497 | 10.4851386 | 14.7674916 | 9.64203936 |
| 20.6955481 | 2.30700799 | 0 | 6.05762487 | 10.4851386 | 6.81576537 | 5.15736989 |
| 41.3910961 | 42.7949983 | 29.1405214 | 43.4129783 | 41.1916158 | 29.5349833 | 41.7074261 |
| 29.5650687 | 58.598003 | 29.1405214 | 37.3553534 | 38.1958619 | 32.942866 | 44.3982278 |
| 11.8260275 | 11.3043392 | 9.71350712 | 9.08643731 | 15.7277078 | 11.359609 | 11.8843741 |
| 6.888661 | 27.7994463 | 0 | 25.2401036 | 12.9791037 | 30.4778308 | 13.7163616 |
| 32.5215755 | 27.3380447 | 0 | 7.06722902 | 17.9745233 | 17.0394134 | 27.3564838 |
| 5.91301374 | 5.88287038 | 0 | 19.1824788 | 14.9787694 | 5.67980448 | 17.7144444 |
| 32.5215755 | 33.7976671 | 29.1405214 | 36.3457492 | 37.4469235 | 51.1182403 | 35.4288888 |
| 44.347603 | 27.5687455 | 9.71350712 | 40.8889679 | 38.5703312 | 44.3024749 | 24.6656821 |
| 11.8260275 | 23.9928831 | 19.4270142 | 20.1920829 | 22.4681541 | 27.2630615 | 24.2172151 |
| 14.7825343 | 7.38242558 | 0 | 5.04802073 | 4.49363081 | 9.08768716 | 6.27853726 |
| 17.7390412 | 12.5731936 | 0 | 16.1536663 | 14.2298309 | 13.0635503 | 13.4540084 |
| 29.5650687 | 39.6805375 | 29.1405214 | 21.2016871 | 26.2128464 | 14.7674916 | 25.3383825 |
| 11.8260275 | 28.4915487 | 9.71350712 | 13.1248539 | 10.4851386 | 11.359609 | 10.9874402 |
| 5.91301374 | 2.88375999 | 9.71350712 | 11.1056456 | 11.234077 | 10.2236481 | 7.84817157 |
| 5.91301374 | 4.26796479 | 9.71350712 | 3.02881244 | 7.48938469 | 13.6315307 | 4.48466947 |
| 168.520891 | 175.90936 | 58.2810427 | 171.632705 | 136.306801 | 135.179347 | 138.576287 |
| 14.7825343 | 8.53592958 | 9.71350712 | 7.06722902 | 7.48938469 | 10.2236481 | 7.6239381 |
| 26.6085618 | 26.2998911 | 29.1405214 | 29.2785202 | 20.9702771 | 39.7586313 | 26.4595499 |
| 5.91301374 | 6.57497278 | 0 | 9.08643731 | 5.24256928 | 10.2236481 | 6.27853726 |
| 0 | 4.96006719 | 0 | 10.0960415 | 6.74044622 | 10.2236481 | 7.17547115 |
| 5.91301374 | 7.49777598 | 0 | 12.1152497 | 8.98726163 | 11.359609 | 7.84817157 |
| 8.8695206 | 11.4196896 | 0 | 8.07683317 | 8.23832316 | 5.67980448 | 7.17547115 |
| 8.8695206 | 14.0727488 | 9.71350712 | 5.04802073 | 7.48938469 | 10.2236481 | 11.8843741 |
| 5.91301374 | 7.84382718 | 0 | 12.1152497 | 4.49363081 | 4.54384358 | 8.74510547 |
| 5.91301374 | 3.11446079 | 0 | 8.07683317 | 8.23832316 | 6.81576537 | 5.83007031 |
| 11.8260275 | 10.7275872 | 19.4270142 | 12.1152497 | 10.4851386 | 29.5349833 | 21.0779465 |
| 11.8260275 | 8.76663038 | 0 | 9.08643731 | 7.48938469 | 17.0394134 | 13.2297749 |
| 5.91301374 | 2.88375999 | 19.4270142 | 12.1152497 | 11.234077 | 9.08768716 | 9.64203936 |
| 5.91301374 | 9.34338237 | 0 | 9.08643731 | 7.48938469 | 6.81576537 | 6.27853726 |
| 5.91301374 | 18.9174655 | 29.1405214 | 19.1824788 | 20.9702771 | 19.3113352 | 22.1991139 |
| 5.91301374 | 4.72936639 | 0 | 5.04802073 | 7.48938469 | 9.08768716 | 5.60583684 |
| 67.999658 | 46.4862111 | 67.9945498 | 74.7107068 | 57.6682621 | 40.8945922 | 65.7004077 |
| 2.95650687 | 5.30611839 | 9.71350712 | 12.1152497 | 16.4766463 | 13.6315307 | 13.2297749 |
| 8.8695206 | 6.22892158 | 9.71350712 | 9.08643731 | 11.9830155 | 9.08768716 | 9.64203936 |
| 14.7825343 | 13.9573984 | 19.4270142 | 15.6488643 | 10.4851386 | 15.9034525 | 13.3418917 |
| 0 | 9.57408317 | 9.71350712 | 6.05762487 | 8.98726163 | 6.81576537 | 7.17547115 |
| 11.8260275 | 10.8429376 | 0 | 4.03841658 | 14.9787694 | 7.95172627 | 12.5570745 |
| 17.7390412 | 5.53681919 | 0 | 4.03841658 | 8.23832316 | 6.81576537 | 5.60583684 |
| 8.8695206 | 17.0718592 | 9.71350712 | 6.05762487 | 8.98726163 | 5.67980448 | 8.74510547 |

| | | | | | | |
|---|---|---|---|---|---|---|
| 5.91301374 | 15.1109024 | 0 | 8.07683317 | 11.234077 | 7.95172627 | 9.41780589 |
| 2.95650687 | 13.0345952 | 29.1405214 | 8.07683317 | 8.23832316 | 7.95172627 | 11.2116737 |
| 4.4347603 | 10.1508352 | 29.1405214 | 9.08643731 | 5.24256928 | 11.359609 | 9.08145568 |
| 20.6955481 | 2.53770879 | 19.4270142 | 16.1536663 | 13.4808924 | 12.4955698 | 11.4359072 |
| 2.95650687 | 7.26707518 | 9.71350712 | 6.05762487 | 5.24256928 | 13.6315307 | 6.72700421 |
| 29.5650687 | 28.7222495 | 9.71350712 | 31.2977285 | 33.7022311 | 23.8551788 | 24.8899156 |
| 2.95650687 | 13.842048 | 0 | 7.06722902 | 10.4851386 | 9.08768716 | 7.84817157 |
| 8.8695206 | 11.4196896 | 0 | 7.06722902 | 7.48938469 | 11.359609 | 7.84817157 |
| 17.7390412 | 20.1863199 | 0 | 12.1152497 | 7.48938469 | 15.9034525 | 8.96933894 |
| 14.7825343 | 13.0922704 | 0 | 5.5528228 | 8.23832316 | 10.2236481 | 13.3418917 |
| 35.4780824 | 91.5882174 | 29.1405214 | 14.134458 | 23.2170925 | 24.9911397 | 27.5807172 |
| 23.6520549 | 21.6858751 | 48.5675356 | 22.2112912 | 17.2255848 | 24.9911397 | 32.0653867 |
| 32.5215755 | 32.5288127 | 19.4270142 | 32.3073327 | 41.9405543 | 27.2630615 | 40.3620252 |
| 0 | 19.1481663 | 9.71350712 | 7.06722902 | 11.9830155 | 5.67980448 | 9.64203936 |
| 5.91301374 | 5.42146879 | 0 | 7.06722902 | 7.48938469 | 2.27192179 | 5.38160336 |
| 11.8260275 | 15.9183552 | 0 | 22.2112912 | 20.2213387 | 21.583257 | 22.8718143 |
| 8.8695206 | 25.3770879 | 9.71350712 | 14.134458 | 9.7362001 | 18.1753743 | 19.7325457 |
| 14.7825343 | 11.1889888 | 19.4270142 | 13.1248539 | 11.9830155 | 12.4955698 | 8.96933894 |
| 17.7390412 | 52.8304831 | 9.71350712 | 28.2689161 | 30.7064772 | 30.6709442 | 25.562616 |
| 0 | 5.88287038 | 0 | 3.02881244 | 7.48938469 | 6.81576537 | 5.38160336 |
| 14.7825343 | 14.0727488 | 0 | 13.4580233 | 2.99575388 | 13.6315307 | 10.7632067 |
| 5.91301374 | 3.92191359 | 0 | 7.06722902 | 6.36597699 | 10.2236481 | 5.00713346 |
| 32.5215755 | 40.1419391 | 19.4270142 | 15.1440622 | 26.9617849 | 23.8551788 | 28.2534177 |
| 62.0866442 | 123.078876 | 48.5675356 | 51.4898114 | 39.6937389 | 84.0611063 | 61.2157383 |
| 11.8260275 | 13.6690224 | 19.4270142 | 22.2112912 | 24.3405002 | 25.3660068 | 6.72700421 |
| 26.6085618 | 34.7493079 | 29.1405214 | 34.7000945 | 31.208266 | 30.1029637 | 24.9056119 |
| 33.4972228 | 12.6112592 | 0 | 7.06722902 | 7.99117346 | 8.71282007 | 11.0614372 |
| 17.7390412 | 32.2981119 | 9.71350712 | 14.134458 | 20.2213387 | 13.6315307 | 15.9205766 |
| 2.95650687 | 2.30700799 | 29.1405214 | 6.05762487 | 7.48938469 | 9.08768716 | 7.6239381 |
| 8.8695206 | 11.53504 | 0 | 6.05762487 | 10.4851386 | 4.54384358 | 8.96933894 |
| 4.93736647 | 11.9191568 | 9.71350712 | 29.6116896 | 21.2174268 | 26.8881944 | 13.323953 |
| 65.0431511 | 54.6760895 | 38.8540285 | 47.4513948 | 47.932062 | 31.8069051 | 47.0890294 |
| 14.7825343 | 11.7657408 | 19.4270142 | 13.1248539 | 8.98726163 | 14.7674916 | 12.781308 |
| 11.8260275 | 10.0354848 | 0 | 7.06722902 | 5.99150775 | 9.08768716 | 11.6601406 |
| 29.5650687 | 18.8021151 | 0 | 17.1632705 | 10.4851386 | 14.7674916 | 11.5480239 |
| 2.95650687 | 19.0328159 | 0 | 9.08643731 | 6.36597699 | 10.2236481 | 6.05430379 |
| 11.8260275 | 15.9183552 | 0 | 12.1152497 | 14.2298309 | 10.2236481 | 14.1267088 |
| 17.7390412 | 15.1109024 | 9.71350712 | 16.1536663 | 9.7362001 | 11.359609 | 10.0905063 |
| 14.7825343 | 10.7275872 | 0 | 8.07683317 | 8.98726163 | 13.6315307 | 7.84817157 |
| 70.9561648 | 40.6033407 | 38.8540285 | 84.8067482 | 74.8938469 | 127.22762 | 25.562616 |
| 11.8260275 | 18.3407136 | 0 | 8.07683317 | 14.9787694 | 10.2236481 | 14.1267088 |
| 17.7390412 | 21.3398239 | 19.4270142 | 20.1920829 | 14.2298309 | 20.4472961 | 21.9748804 |
| 8.8695206 | 10.6122368 | 9.71350712 | 14.134458 | 9.7362001 | 10.2236481 | 11.4359072 |
| 8.8695206 | 14.1880992 | 0 | 11.1056456 | 7.48938469 | 13.6315307 | 8.07240505 |
| 0 | 15.1109024 | 0 | 7.06722902 | 7.48938469 | 5.67980448 | 8.07240505 |
| 2.95650687 | 4.44099039 | 0 | 10.0960415 | 12.3574847 | 6.81576537 | 6.27853726 |

| | | | | | | |
|---|---|---|---|---|---|---|
| 2.95650687 | 6.34427198 | 0 | 8.07683317 | 7.48938469 | 3.40788269 | 4.260436 |
| 11.8260275 | 24.5696351 | 0 | 9.08643731 | 5.99150775 | 10.2236481 | 8.74510547 |
| 5.91301374 | 8.88198078 | 19.4270142 | 7.06722902 | 8.98726163 | 6.81576537 | 8.52087199 |
| 5.91301374 | 33.6823167 | 19.4270142 | 13.1248539 | 17.9745233 | 13.6315307 | 13.0055415 |
| 26.6085618 | 19.0328159 | 0 | 15.1440622 | 11.234077 | 14.7674916 | 17.7144444 |
| 20.6955481 | 11.9964416 | 0 | 8.07683317 | 12.731954 | 9.08768716 | 9.19357241 |
| 5.91301374 | 15.1109024 | 9.71350712 | 11.1056456 | 10.4851386 | 12.4955698 | 15.2478762 |
| 2.95650687 | 2.42235839 | 0 | 4.03841658 | 5.99150775 | 14.1995112 | 2.80291842 |
| 11.8260275 | 7.15172478 | 0 | 10.0960415 | 5.99150775 | 9.08768716 | 6.50277073 |
| 88.695206 | 65.8650782 | 38.8540285 | 102.979623 | 75.6427854 | 84.0611063 | 82.2936848 |
| 22.1738015 | 10.8429376 | 58.2810427 | 15.6488643 | 17.2255848 | 14.7674916 | 17.7144444 |
| 5.91301374 | 7.61312638 | 9.71350712 | 2.52401036 | 5.24256928 | 9.08768716 | 5.38160336 |
| 17.7390412 | 44.4099039 | 9.71350712 | 17.1632705 | 13.4808924 | 14.7674916 | 12.1086076 |
| 4.4347603 | 12.4578432 | 4.85675356 | 23.7256974 | 20.2213387 | 31.8069051 | 9.30568915 |
| 50.2606167 | 21.4551743 | 38.8540285 | 30.2881244 | 43.4384312 | 31.8069051 | 34.7561884 |
| 2.95650687 | 1.96095679 | 0 | 7.06722902 | 3.74469235 | 13.6315307 | 7.84817157 |
| 2.95650687 | 4.78704159 | 0 | 6.05762487 | 8.23832316 | 6.24778492 | 4.93313642 |
| 2.95650687 | 6.11357118 | 9.71350712 | 10.0960415 | 11.234077 | 13.6315307 | 8.96933894 |
| 44.347603 | 23.0700799 | 9.71350712 | 23.2208954 | 14.2298309 | 29.5349833 | 25.114149 |
| 8.8695206 | 10.1508352 | 9.71350712 | 10.0960415 | 14.2298309 | 14.7674916 | 12.1086076 |
| 29.5650687 | 15.283928 | 0 | 28.7737182 | 25.0894387 | 18.1753743 | 21.7506469 |
| 26.6085618 | 11.53504 | 9.71350712 | 10.0960415 | 13.4808924 | 13.6315307 | 15.4721097 |
| 11.8260275 | 13.842048 | 0 | 7.06722902 | 10.4851386 | 3.40788269 | 6.50277073 |
| 8.8695206 | 4.84471679 | 9.71350712 | 3.02881244 | 4.49363081 | 10.2236481 | 6.05430379 |
| 0 | 13.265296 | 19.4270142 | 5.04802073 | 8.98726163 | 4.54384358 | 4.93313642 |
| 29.5650687 | 17.30256 | 19.4270142 | 28.7737182 | 26.2128464 | 13.6315307 | 24.1050984 |
| 16.2607878 | 10.0354848 | 9.71350712 | 13.1248539 | 16.4766463 | 31.2389246 | 11.4359072 |
| 14.7825343 | 0.8074528 | 19.4270142 | 5.04802073 | 12.731954 | 9.08768716 | 6.05430379 |
| 20.6955481 | 18.1100128 | 19.4270142 | 17.1632705 | 32.9532926 | 17.0394134 | 21.5264135 |
| 2.95650687 | 1.730256 | 19.4270142 | 8.07683317 | 13.4808924 | 12.4955698 | 14.7994093 |
| 8.8695206 | 12.3424928 | 9.71350712 | 7.06722902 | 11.9830155 | 15.3354721 | 8.74510547 |
| 8.8695206 | 10.1508352 | 0 | 6.05762487 | 9.7362001 | 7.95172627 | 6.05430379 |
| 8.8695206 | 16.4951072 | 9.71350712 | 9.08643731 | 14.2298309 | 5.67980448 | 12.781308 |
| 5.91301374 | 11.0736384 | 0 | 12.1152497 | 8.23832316 | 6.81576537 | 8.07240505 |
| 17.7390412 | 17.4179104 | 0 | 13.1248539 | 14.2298309 | 10.2236481 | 13.6782419 |
| 8.8695206 | 5.19076799 | 9.71350712 | 8.07683317 | 11.234077 | 9.08768716 | 15.4721097 |
| 35.4780824 | 33.1055647 | 19.4270142 | 21.2016871 | 14.9787694 | 15.9034525 | 20.1810126 |
| 0 | 0.8074528 | 19.4270142 | 12.1152497 | 8.23832316 | 11.359609 | 7.17547115 |
| 5.91301374 | 6.69032318 | 0 | 5.04802073 | 14.2298309 | 6.81576537 | 14.7994093 |
| 11.8260275 | 6.69032318 | 9.71350712 | 6.05762487 | 11.234077 | 11.359609 | 10.5389733 |
| 29.5650687 | 8.76663038 | 0 | 13.1248539 | 9.7362001 | 12.4955698 | 13.6782419 |
| 14.7825343 | 15.8030048 | 9.71350712 | 12.1152497 | 8.98726163 | 5.67980448 | 8.29663852 |
| 26.6085618 | 21.0514479 | 38.8540285 | 44.4225824 | 34.0767003 | 44.3024749 | 42.2680098 |
| 26.6085618 | 49.2546207 | 0 | 17.1632705 | 22.4681541 | 22.7192179 | 28.0291842 |
| 20.6955481 | 12.111792 | 9.71350712 | 14.134458 | 13.4808924 | 15.9034525 | 17.4902109 |
| 38.4345893 | 33.9130175 | 19.4270142 | 45.4321866 | 31.4554157 | 22.7192179 | 37.895457 |

| | | | | | | |
|---|---|---|---|---|---|---|
| 20.6955481 | 10.958288 | 29.1405214 | 10.0960415 | 12.731954 | 11.359609 | 14.3509423 |
| 5.91301374 | 17.9946624 | 0 | 7.06722902 | 7.48938469 | 5.67980448 | 9.64203936 |
| 2.95650687 | 8.99733118 | 9.71350712 | 1.00960415 | 4.49363081 | 9.08768716 | 4.70890294 |
| 5.91301374 | 5.88287038 | 0 | 6.05762487 | 11.234077 | 3.40788269 | 10.1645034 |
| 38.4345893 | 24.5696351 | 38.8540285 | 55.528228 | 34.4511696 | 18.1753743 | 28.9261181 |
| 115.303768 | 163.797568 | 19.4270142 | 94.9027897 | 139.302555 | 243.095632 | 83.4148522 |
| 5.91301374 | 10.6122368 | 0 | 7.06722902 | 7.48938469 | 2.27192179 | 6.50277073 |
| 14.7825343 | 30.7985567 | 9.71350712 | 13.1248539 | 15.7277078 | 13.6315307 | 10.7632067 |
| 109.390754 | 118.926262 | 58.2810427 | 104.998831 | 74.8938469 | 86.3330281 | 86.890471 |
| 11.8260275 | 16.6104576 | 9.71350712 | 27.2593119 | 20.9702771 | 14.7674916 | 21.7506469 |
| 2.95650687 | 8.53592958 | 29.1405214 | 10.0960415 | 11.234077 | 15.9034525 | 10.5389733 |
| 23.6520549 | 32.8748639 | 29.1405214 | 31.2977285 | 34.4511696 | 19.3113352 | 28.2534177 |
| 23.6520549 | 34.2590687 | 19.4270142 | 20.1920829 | 21.7192156 | 15.9034525 | 23.5445147 |
| 23.6520549 | 8.24755358 | 0 | 15.1440622 | 10.4851386 | 12.4955698 | 11.6982603 |
| 0 | 7.61312638 | 0 | 1.00960415 | 10.4851386 | 7.95172627 | 5.83007031 |
| 11.8260275 | 16.725808 | 0 | 8.07683317 | 17.9745233 | 15.9034525 | 15.9205766 |
| 5.91301374 | 2.19165759 | 0 | 3.02881244 | 8.98726163 | 5.67980448 | 5.60583684 |
| 2.4539007 | 3.40283679 | 0 | 9.41960668 | 5.86418821 | 9.28080052 | 6.05430379 |
| 14.7825343 | 10.1508352 | 9.71350712 | 3.02881244 | 5.99150775 | 9.08768716 | 6.27853726 |
| 11.8260275 | 22.7240287 | 19.4270142 | 14.134458 | 11.9830155 | 14.8924473 | 15.4721097 |
| 2.95650687 | 3.92191359 | 9.71350712 | 14.134458 | 5.99150775 | 12.4955698 | 3.13926863 |
| 20.6955481 | 15.8030048 | 38.8540285 | 10.0960415 | 15.7277078 | 7.95172627 | 14.5751758 |
| 17.7390412 | 31.4906591 | 29.1405214 | 21.2016871 | 19.4724002 | 14.7674916 | 30.9442193 |
| 26.6085618 | 7.38242558 | 19.4270142 | 7.06722902 | 19.4724002 | 21.583257 | 25.7868495 |
| 11.8260275 | 6.22892158 | 0 | 3.53361451 | 9.7362001 | 9.08768716 | 7.28758789 |
| 0 | 2.50656419 | 9.71350712 | 7.06722902 | 8.98726163 | 10.2236481 | 4.03620252 |
| 26.6085618 | 40.1419391 | 19.4270142 | 35.3361451 | 24.7149695 | 35.2147878 | 38.1196905 |
| 5.91301374 | 9.57408317 | 9.71350712 | 3.11967681 | 9.7362001 | 9.08768716 | 6.72700421 |
| 2.95650687 | 19.9556191 | 0 | 8.07683317 | 10.4851386 | 5.67980448 | 7.6239381 |
| 53.2171236 | 41.7949103 | 19.4270142 | 36.6789186 | 31.4554157 | 42.5985336 | 39.7274445 |
| 5.91301374 | 11.0736384 | 9.71350712 | 4.03841658 | 9.7362001 | 6.81576537 | 5.83007031 |
| 41.3910961 | 49.2546207 | 0 | 33.3169368 | 28.4596618 | 38.6226704 | 26.9080168 |
| 2.95650687 | 5.99822078 | 0 | 5.04802073 | 2.99575388 | 11.359609 | 7.39970463 |
| 26.6085618 | 8.07452798 | 0 | 31.8025306 | 25.4639079 | 18.7433548 | 21.9748804 |
| 8.8695206 | 7.15172478 | 0 | 3.02881244 | 9.7362001 | 5.67980448 | 5.38160336 |
| 35.4780824 | 37.6042303 | 48.5675356 | 29.2785202 | 20.9702771 | 18.1753743 | 23.0960478 |
| 0 | 13.4959968 | 9.71350712 | 8.07683317 | 8.23832316 | 6.81576537 | 9.64203936 |
| 5.91301374 | 22.1472767 | 9.71350712 | 21.2016871 | 23.966031 | 18.1753743 | 16.593277 |
| 11.8260275 | 14.7648512 | 29.1405214 | 16.1536663 | 12.731954 | 11.359609 | 19.0598453 |
| 44.347603 | 54.5607391 | 48.5675356 | 35.3361451 | 22.4681541 | 24.9911397 | 37.4469901 |
| 14.7825343 | 18.3407136 | 19.4270142 | 7.06722902 | 8.98726163 | 7.95172627 | 8.74510547 |
| 79.8256854 | 70.9404958 | 87.4215641 | 66.6338736 | 58.4172006 | 62.4778492 | 68.6154429 |
| 26.6085618 | 6.11357118 | 0 | 12.1152497 | 6.74044622 | 6.81576537 | 8.96933894 |
| 11.8260275 | 11.1889888 | 9.71350712 | 10.0960415 | 13.4808924 | 5.67980448 | 8.74510547 |
| 20.6955481 | 23.9928831 | 9.71350712 | 26.2497078 | 25.4639079 | 22.7192179 | 26.9080168 |
| 35.4780824 | 30.2218047 | 29.1405214 | 33.3169368 | 44.1873697 | 39.7586313 | 42.155893 |

| | | | | | | |
|---|---|---|---|---|---|---|
| 23.6520549 | 10.4968864 | 9.71350712 | 21.2016871 | 14.9787694 | 13.6315307 | 17.7144444 |
| 5.91301374 | 5.88287038 | 19.4270142 | 10.0960415 | 9.7362001 | 3.40788269 | 5.83007031 |
| 8.8695206 | 11.8810912 | 0 | 12.1152497 | 12.731954 | 6.81576537 | 9.86627284 |
| 38.4345893 | 34.0283679 | 0 | 40.3841658 | 30.7064772 | 31.8069051 | 45.0709282 |
| 5.91301374 | 4.84471679 | 29.1405214 | 11.1056456 | 10.4851386 | 18.1753743 | 11.4359072 |
| 17.7390412 | 14.7648512 | 19.4270142 | 14.134458 | 11.234077 | 12.4955698 | 14.7994093 |
| 5.91301374 | 4.49866559 | 9.71350712 | 7.06722902 | 7.48938469 | 5.67980448 | 8.74510547 |
| 5.91301374 | 12.3424928 | 9.71350712 | 9.08643731 | 9.7362001 | 9.08768716 | 7.84817157 |
| 0 | 2.99911039 | 9.71350712 | 7.06722902 | 2.24681541 | 9.08768716 | 2.91503516 |
| 11.8260275 | 5.76751998 | 0 | 7.06722902 | 7.48938469 | 15.9034525 | 11.6601406 |
| 0 | 8.53592958 | 0 | 4.03841658 | 2.99575388 | 9.08768716 | 4.03620252 |
| 44.347603 | 21.2244735 | 19.4270142 | 34.326541 | 44.1873697 | 31.8069051 | 40.1377918 |
| 11.8260275 | 5.76751998 | 9.71350712 | 8.07683317 | 11.234077 | 12.4955698 | 8.63298873 |
| 56.1736305 | 59.2901054 | 29.1405214 | 52.4994156 | 31.4554157 | 39.7586313 | 41.0347257 |
| 2.95650687 | 2.19165759 | 0 | 12.1152497 | 5.99150775 | 5.67980448 | 14.5751758 |
| 5.91301374 | 1.96095679 | 0 | 7.06722902 | 7.48938469 | 5.67980448 | 5.15736989 |
| 5.91301374 | 9.45873277 | 9.71350712 | 7.06722902 | 18.7234617 | 17.0394134 | 13.2297749 |
| 94.6082198 | 56.4063454 | 67.9945498 | 62.595457 | 50.9278159 | 64.749771 | 67.0458086 |
| 2.95650687 | 5.47914399 | 0 | 4.03841658 | 7.48938469 | 11.359609 | 4.48466947 |
| 591.301374 | 273.726498 | 534.242892 | 619.896945 | 581.92519 | 530.493738 | 643.101602 |
| 11.8260275 | 17.30256 | 9.71350712 | 9.08643731 | 11.9830155 | 4.54384358 | 12.332841 |
| 11.8260275 | 19.9556191 | 0 | 12.1152497 | 6.74044622 | 15.9034525 | 13.2297749 |
| 29.5650687 | 26.4152415 | 9.71350712 | 29.2785202 | 22.4681541 | 15.9034525 | 30.2715189 |
| 14.7825343 | 11.8810912 | 0 | 3.02881244 | 3.74469235 | 12.4955698 | 4.70890294 |
| 29.5650687 | 32.0674111 | 29.1405214 | 20.1920829 | 23.966031 | 29.5349833 | 25.7868495 |
| 2.95650687 | 3.11446079 | 0 | 5.04802073 | 9.7362001 | 7.0997556 | 8.96933894 |
| 23.6520549 | 23.0700799 | 19.4270142 | 17.6680725 | 19.4724002 | 22.7192179 | 26.4595499 |
| 79.8256854 | 66.7878814 | 77.708057 | 65.6242695 | 50.1788774 | 56.7980448 | 70.8577776 |
| 26.6085618 | 9.80478397 | 19.4270142 | 12.1152497 | 13.4808924 | 14.7674916 | 13.2297749 |
| 5.91301374 | 17.5332608 | 0 | 11.1056456 | 10.4851386 | 5.67980448 | 11.6601406 |
| 14.7825343 | 13.0345952 | 9.71350712 | 9.08643731 | 19.4724002 | 18.1753743 | 15.6963431 |
| 0 | 7.95917758 | 0 | 8.07683317 | 7.48938469 | 5.67980448 | 7.6239381 |
| 38.4345893 | 22.4933279 | 67.9945498 | 56.5378322 | 39.6937389 | 37.4867095 | 39.9135583 |
| 5.91301374 | 14.1880992 | 9.71350712 | 16.1536663 | 9.7362001 | 6.81576537 | 15.0236427 |
| 35.4780824 | 48.2164671 | 29.1405214 | 15.1440622 | 20.9702771 | 9.08768716 | 19.9567791 |
| 29.5650687 | 27.2226943 | 19.4270142 | 24.2304995 | 31.0809465 | 24.4231593 | 27.0201336 |
| 8.8695206 | 10.381536 | 9.71350712 | 8.07683317 | 9.7362001 | 7.95172627 | 7.84817157 |
| 70.9561648 | 44.6982799 | 14.5702607 | 42.9081762 | 58.7916698 | 48.8463185 | 67.6063923 |
| 2.95650687 | 7.15172478 | 9.71350712 | 18.1728746 | 6.74044622 | 14.7674916 | 11.8843741 |
| 8.8695206 | 10.6122368 | 9.71350712 | 10.0960415 | 12.731954 | 11.359609 | 15.4721097 |
| 2.95650687 | 0.2307008 | 9.71350712 | 6.05762487 | 7.48938469 | 10.2236481 | 8.07240505 |
| 8.8695206 | 7.61312638 | 9.71350712 | 15.1440622 | 14.9787694 | 9.08768716 | 19.9567791 |
| 0 | 7.15172478 | 9.71350712 | 3.02881244 | 6.74044622 | 11.359609 | 5.60583684 |
| 5.91301374 | 5.76751998 | 0 | 2.01920829 | 8.23832316 | 5.67980448 | 4.260436 |
| 11.8260275 | 1.2688544 | 0 | 9.08643731 | 11.234077 | 10.2236481 | 6.95123768 |
| 0 | 3.11446079 | 0 | 2.01920829 | 3.74469235 | 9.08768716 | 7.6239381 |

| | | | | | | |
|---|---|---|---|---|---|---|
| 47.3041099 | 27.1073439 | 19.4270142 | 40.3841658 | 44.1873697 | 35.2147878 | 37.4469901 |
| 5.91301374 | 8.07452798 | 0 | 5.04802073 | 6.74044622 | 11.359609 | 4.03620252 |
| 8.8695206 | 27.1073439 | 9.71350712 | 18.1728746 | 17.2255848 | 17.0394134 | 16.8175105 |
| 8.8695206 | 0.4614016 | 0 | 9.08643731 | 11.9830155 | 13.6315307 | 8.96933894 |
| 2.95650687 | 22.1472767 | 29.1405214 | 12.1152497 | 12.731954 | 15.9034525 | 11.4359072 |
| 8.8695206 | 17.879312 | 9.71350712 | 28.7737182 | 26.5873156 | 26.1271006 | 28.0291842 |
| 482.886267 | 450.09726 | 315.688981 | 511.3645 | 420.154481 | 414.057746 | 480.532334 |
| 11.8260275 | 10.4968864 | 9.71350712 | 8.07683317 | 8.98726163 | 5.67980448 | 10.5389733 |
| 8.8695206 | 18.2253632 | 9.71350712 | 10.0960415 | 11.9830155 | 12.4955698 | 8.8572222 |
| 82.7821923 | 66.3264798 | 19.4270142 | 64.6146653 | 56.1703852 | 48.8463185 | 64.3550069 |
| 26.6085618 | 11.1889888 | 0 | 15.1440622 | 23.966031 | 13.0635503 | 22.8718143 |
| 20.6955481 | 14.0727488 | 29.1405214 | 8.07683317 | 11.234077 | 12.4955698 | 14.5751758 |
| 35.4780824 | 89.9733118 | 38.8540285 | 19.1824788 | 21.7192156 | 20.4472961 | 29.1503516 |
| 5.91301374 | 8.18987838 | 9.71350712 | 9.08643731 | 11.234077 | 3.40788269 | 7.17547115 |
| 26.6085618 | 16.725808 | 29.1405214 | 28.2689161 | 17.9745233 | 24.9911397 | 25.562616 |
| 2.95650687 | 3.92191359 | 0 | 5.04802073 | 13.4808924 | 10.2236481 | 9.86627284 |
| 5.91301374 | 6.34427198 | 19.4270142 | 7.06722902 | 8.23832316 | 7.95172627 | 9.19357241 |
| 14.7825343 | 0.3460512 | 9.71350712 | 11.1056456 | 7.48938469 | 7.95172627 | 5.15736989 |
| 14.7825343 | 9.57408317 | 0 | 10.0960415 | 11.234077 | 19.3113352 | 11.4359072 |
| 5.91301374 | 4.72936639 | 0 | 3.02881244 | 5.24256928 | 11.359609 | 7.6239381 |
| 20.6955481 | 26.7612927 | 0 | 23.2208954 | 20.9702771 | 23.8551788 | 19.5083122 |
| 5.91301374 | 5.76751998 | 0 | 20.1920829 | 14.9787694 | 23.8551788 | 20.1810126 |
| 17.7390412 | 17.4179104 | 0 | 7.06722902 | 13.4808924 | 14.7674916 | 14.3509423 |
| 26.6085618 | 6.22892158 | 0 | 17.1632705 | 17.9745233 | 10.2236481 | 15.2478762 |
| 17.7390412 | 17.30256 | 9.71350712 | 15.1440622 | 10.4851386 | 6.81576537 | 9.64203936 |
| 138.955823 | 191.597014 | 87.4215641 | 162.546267 | 144.545125 | 115.868011 | 200.016258 |
| 11.8260275 | 15.9183552 | 29.1405214 | 10.0960415 | 8.98726163 | 22.7192179 | 17.4902109 |
| 0 | 7.38242558 | 0 | 1.00960415 | 4.86810005 | 9.65566761 | 3.47561884 |
| 2.95650687 | 5.53681919 | 0 | 6.56242695 | 6.74044622 | 17.6073939 | 5.94218705 |
| 47.3041099 | 21.2244735 | 67.9945498 | 43.4129783 | 19.4724002 | 20.4472961 | 31.1684528 |
| 0 | 8.88198078 | 0 | 6.05762487 | 7.48938469 | 7.95172627 | 4.93313642 |
| 11.8260275 | 19.8402687 | 0 | 6.05762487 | 7.48938469 | 3.40788269 | 9.64203936 |
| 8.8695206 | 19.2635167 | 0 | 37.3553534 | 23.966031 | 52.2542012 | 13.0055415 |
| 14.7825343 | 6.11357118 | 0 | 14.134458 | 11.234077 | 15.9034525 | 7.6239381 |
| 29.5650687 | 39.3344863 | 19.4270142 | 21.2016871 | 14.2298309 | 19.3113352 | 22.8718143 |
| 2.95650687 | 3.46051199 | 0 | 11.1056456 | 8.98726163 | 3.40788269 | 4.48466947 |
| 73.9126717 | 73.7089054 | 48.5675356 | 82.7875399 | 63.6597699 | 49.9822794 | 76.9120814 |
| 8.8695206 | 5.07541759 | 0 | 4.03841658 | 8.98726163 | 4.54384358 | 6.95123768 |
| 35.4780824 | 59.9822078 | 38.8540285 | 38.3649575 | 34.4511696 | 26.1271006 | 40.1377918 |
| 8.8695206 | 10.8429376 | 9.71350712 | 6.05762487 | 15.7277078 | 21.583257 | 16.1448101 |
| 2.95650687 | 11.7657408 | 0 | 6.05762487 | 9.7362001 | 5.67980448 | 8.74510547 |
| 11.8260275 | 9.57408317 | 19.4270142 | 7.06722902 | 8.23832316 | 6.81576537 | 4.03620252 |
| 11.8260275 | 2.99911039 | 0 | 11.1056456 | 8.23832316 | 13.6315307 | 8.74510547 |
| 8.8695206 | 9.22803198 | 9.71350712 | 7.06722902 | 8.23832316 | 13.6315307 | 8.29663852 |
| 100.521233 | 38.2963327 | 87.4215641 | 114.085268 | 79.3874777 | 74.9734191 | 84.9844865 |
| 36.9563358 | 29.5873775 | 72.8513034 | 54.5186239 | 50.1788774 | 59.0699666 | 56.3947186 |

| | | | | | | |
|---|---|---|---|---|---|---|
| 5.91301374 | 2.53770879 | 0 | 3.02881244 | 7.48938469 | 9.08768716 | 5.83007031 |
| 2.95650687 | 14.0727488 | 0 | 20.1920829 | 17.2255848 | 14.7674916 | 15.9205766 |
| 2.95650687 | 8.07452798 | 9.71350712 | 5.04802073 | 20.2213387 | 5.67980448 | 10.9874402 |
| 29.5650687 | 22.2626271 | 29.1405214 | 32.3073327 | 15.7277078 | 23.8551788 | 24.6656821 |
| 11.8260275 | 6.34427198 | 9.71350712 | 15.1440622 | 10.4851386 | 9.08768716 | 11.4359072 |
| 56.1736305 | 36.6814271 | 29.1405214 | 42.4033741 | 33.7022311 | 36.3507487 | 37.4469901 |
| 11.8260275 | 6.80567358 | 9.71350712 | 18.1728746 | 23.2170925 | 23.8551788 | 8.52087199 |
| 2.95650687 | 6.92102398 | 0 | 11.1056456 | 5.99150775 | 6.81576537 | 6.27853726 |
| 0 | 6.69032318 | 0 | 8.07683317 | 11.234077 | 13.6315307 | 10.3147398 |
| 32.5215755 | 41.0647423 | 58.2810427 | 42.4033741 | 35.200108 | 36.3507487 | 44.6224612 |
| 5.91301374 | 12.5731936 | 0 | 6.05762487 | 7.48938469 | 17.0394134 | 13.2297749 |
| 5.91301374 | 5.76751998 | 9.71350712 | 8.07683317 | 7.48938469 | 11.359609 | 6.72700421 |
| 65.0431511 | 74.170307 | 116.562085 | 58.5570405 | 40.4426773 | 37.4867095 | 61.6642052 |
| 20.6955481 | 19.7249183 | 48.5675356 | 28.2689161 | 15.7277078 | 18.1753743 | 21.5264135 |
| 2.95650687 | 4.84471679 | 0 | 5.5528228 | 7.11491546 | 9.08768716 | 7.6239381 |
| 14.7825343 | 21.3594335 | 0 | 8.07683317 | 10.4851386 | 9.08768716 | 9.82815314 |
| 5.91301374 | 3.63353759 | 0 | 5.04802073 | 6.36597699 | 9.08768716 | 7.17547115 |
| 11.8260275 | 4.61401599 | 0 | 8.07683317 | 7.48938469 | 10.2236481 | 8.07240505 |
| 11.8260275 | 21.3398239 | 0 | 20.1920829 | 23.2170925 | 15.9034525 | 20.853713 |
| 26.6085618 | 9.22803198 | 19.4270142 | 24.2304995 | 20.9702771 | 24.9911397 | 23.5445147 |
| 32.5215755 | 19.2635167 | 38.8540285 | 20.1920829 | 22.4681541 | 30.6709442 | 23.0960478 |
| 35.4780824 | 28.0301471 | 19.4270142 | 24.2304995 | 23.2170925 | 23.8551788 | 26.4595499 |
| 41.3910961 | 46.8322623 | 9.71350712 | 25.2401036 | 29.2086003 | 42.0305531 | 34.5319549 |
| 5.91301374 | 7.38242558 | 19.4270142 | 2.01920829 | 10.4851386 | 14.7674916 | 7.84817157 |
| 50.2606167 | 38.2963327 | 19.4270142 | 50.4802073 | 41.9405543 | 47.7103576 | 41.4831926 |
| 15.7581816 | 21.2786882 | 2.42837678 | 22.6959012 | 25.7260364 | 32.1249741 | 13.5280055 |
| 11.8260275 | 3.11446079 | 9.71350712 | 5.04802073 | 5.24256928 | 9.08768716 | 8.52087199 |
| 7.89387334 | 3.34516159 | 0 | 10.8431485 | 10.8596078 | 14.1086343 | 6.31665695 |
| 0 | 2.30700799 | 9.71350712 | 5.04802073 | 7.48938469 | 5.67980448 | 6.95123768 |
| 53.2171236 | 79.361075 | 48.5675356 | 61.5858529 | 70.4002161 | 53.3901621 | 69.2881433 |
| 26.6085618 | 24.8003359 | 29.1405214 | 28.2689161 | 21.7192156 | 17.0394134 | 26.6837834 |
| 91.6517129 | 1096.7516 | 48.5675356 | 98.9412063 | 69.6512776 | 70.4295755 | 104.044332 |
| 5.91301374 | 4.03726399 | 9.71350712 | 9.08643731 | 11.9830155 | 11.359609 | 3.13926863 |
| 712.518155 | 961.676282 | 475.961849 | 640.089028 | 536.988882 | 513.454325 | 593.994471 |
| 5.91301374 | 13.265296 | 29.1405214 | 11.6104477 | 9.7362001 | 15.9034525 | 8.18452178 |
| 2.95650687 | 6.22892158 | 0 | 8.07683317 | 10.4851386 | 5.67980448 | 4.70890294 |
| 8.8695206 | 7.26707518 | 9.71350712 | 6.05762487 | 6.74044622 | 9.08768716 | 8.07240505 |
| 17.7390412 | 24.6849855 | 0 | 12.6200518 | 9.7362001 | 21.583257 | 20.1810126 |
| 2.95650687 | 2.53770879 | 0 | 11.1056456 | 10.4851386 | 9.08768716 | 6.72700421 |
| 11.8260275 | 3.46051199 | 9.71350712 | 4.03841658 | 11.234077 | 10.2236481 | 7.84817157 |
| 36.4537297 | 29.7407936 | 9.71350712 | 40.3841658 | 44.9363081 | 37.4867095 | 30.6818662 |
| 20.6955481 | 7.72847678 | 9.71350712 | 11.1056456 | 20.9702771 | 14.7674916 | 16.1448101 |
| 0 | 2.99911039 | 0 | 5.04802073 | 5.24256928 | 9.08768716 | 4.260436 |
| 16.2607878 | 21.9742511 | 19.4270142 | 23.7256974 | 23.2170925 | 19.8793157 | 19.5083122 |
| 20.6955481 | 17.1872096 | 0 | 29.2785202 | 17.9745233 | 10.2236481 | 20.4052461 |
| 127.129795 | 159.414252 | 126.275593 | 144.373393 | 134.059986 | 130.635503 | 167.950872 |

| | | | | | | |
|---|---|---|---|---|---|---|
| 17.7390412 | 17.879312 | 29.1405214 | 23.2208954 | 14.9787694 | 17.0394134 | 13.4540084 |
| 11.8260275 | 10.2661856 | 9.71350712 | 12.1152497 | 9.7362001 | 13.6315307 | 7.17547115 |
| 8.8695206 | 10.2661856 | 0 | 11.1056456 | 7.48938469 | 12.4955698 | 5.83007031 |
| 14.7825343 | 12.4578432 | 19.4270142 | 18.1728746 | 13.4808924 | 21.583257 | 9.86627284 |
| 17.7390412 | 4.61401599 | 0 | 10.0960415 | 14.9787694 | 3.40788269 | 11.6601406 |
| 8.8695206 | 17.0718592 | 19.4270142 | 17.1632705 | 27.7107234 | 20.4472961 | 25.114149 |
| 20.6955481 | 11.7657408 | 9.71350712 | 15.1440622 | 14.9787694 | 18.1753743 | 14.5751758 |
| 11.8260275 | 9.45873277 | 29.1405214 | 17.1632705 | 14.9787694 | 9.08768716 | 15.4721097 |
| 65.0431511 | 25.1463871 | 48.5675356 | 47.4513948 | 44.9363081 | 47.7103576 | 44.3982278 |
| 9.84516787 | 9.05500638 | 3.20545735 | 8.66240357 | 9.36173086 | 8.32659336 | 14.6693538 |
| 2.95650687 | 4.49866559 | 0 | 4.03841658 | 1.49787694 | 9.08768716 | 3.3635021 |
| 14.7825343 | 11.0736384 | 0 | 6.05762487 | 8.23832316 | 2.27192179 | 8.74510547 |
| 97.5647266 | 63.4427198 | 58.2810427 | 110.046852 | 103.353509 | 102.236481 | 125.794979 |
| 5.91301374 | 8.30522878 | 0 | 7.06722902 | 13.4808924 | 5.67980448 | 9.86627284 |
| 33.999829 | 26.8766431 | 77.708057 | 42.9081762 | 46.4341851 | 43.7344945 | 43.1649437 |
| 26.6085618 | 26.1845407 | 38.8540285 | 24.2304995 | 25.4639079 | 23.8551788 | 29.823052 |
| 8.8695206 | 12.4578432 | 0 | 16.1536663 | 14.9787694 | 15.9034525 | 14.1267088 |
| 10.347774 | 8.18987838 | 0 | 9.08643731 | 7.11491546 | 13.6315307 | 9.7541561 |
| 17.7390412 | 13.1499456 | 9.71350712 | 18.1728746 | 11.9830155 | 7.95172627 | 11.2116737 |
| 5.91301374 | 22.1472767 | 0 | 11.1056456 | 14.2298309 | 17.0394134 | 14.3509423 |
| 159.651371 | 130.807353 | 135.9891 | 132.258143 | 137.804678 | 122.683777 | 147.097159 |
| 23.6520549 | 12.8038944 | 9.71350712 | 13.1248539 | 17.2255848 | 22.7192179 | 12.5570745 |
| 11.8260275 | 1.7879312 | 9.71350712 | 8.07683317 | 9.7362001 | 7.95172627 | 6.27853726 |
| 38.4345893 | 52.2537311 | 38.8540285 | 29.2785202 | 26.9617849 | 18.1753743 | 32.2896202 |
| 23.6520549 | 12.5812681 | 14.0845853 | 25.8862503 | 23.479221 | 29.7962543 | 11.8664354 |
| 5.91301374 | 10.0354848 | 0 | 12.1152497 | 6.74044622 | 11.359609 | 11.4359072 |
| 5.91301374 | 1.96095679 | 9.71350712 | 10.0960415 | 8.23832316 | 13.6315307 | 11.2116737 |
| 8.8695206 | 3.92191359 | 9.71350712 | 7.06722902 | 7.48938469 | 5.67980448 | 6.72700421 |
| 2.95650687 | 12.688544 | 0 | 7.06722902 | 9.7362001 | 14.7674916 | 10.7632067 |
| 26.6085618 | 32.4134623 | 53.4242892 | 31.2977285 | 34.4511696 | 28.3990224 | 36.1015892 |
| 20.6955481 | 6.69032318 | 0 | 4.03841658 | 10.4851386 | 14.7674916 | 10.3147398 |
| 8.8695206 | 12.2271424 | 0 | 11.1056456 | 5.99150775 | 10.2236481 | 8.07240505 |
| 2.95650687 | 5.53681919 | 0 | 5.04802073 | 6.74044622 | 9.08768716 | 4.93313642 |
| 2.95650687 | 10.4968864 | 9.71350712 | 7.06722902 | 7.48938469 | 6.81576537 | 4.03620252 |
| 38.4345893 | 27.2803695 | 33.9972749 | 18.1728746 | 17.2255848 | 17.0394134 | 19.0598453 |
| 11.8260275 | 3.69121279 | 0 | 12.6200518 | 9.36173086 | 6.24778492 | 7.17547115 |
| 35.4780824 | 18.3407136 | 58.2810427 | 36.3457492 | 38.9448004 | 39.7586313 | 29.5988185 |
| 5.91301374 | 1.0381536 | 9.71350712 | 13.1248539 | 8.98726163 | 9.08768716 | 8.74510547 |
| 14.7825343 | 17.30256 | 9.71350712 | 14.134458 | 16.4766463 | 12.4955698 | 21.7506469 |
| 5.91301374 | 57.7905502 | 29.1405214 | 9.08643731 | 7.48938469 | 10.2236481 | 7.84817157 |
| 2.95650687 | 15.2262528 | 29.1405214 | 9.08643731 | 5.99150775 | 10.2236481 | 9.86627284 |
| 50.2606167 | 58.021251 | 77.708057 | 51.4898114 | 61.4129545 | 38.6226704 | 46.7145595 |
| 17.7390412 | 12.5616585 | 12.9189645 | 23.7256974 | 18.5961422 | 24.7980263 | 16.2569268 |
| 44.347603 | 60.7896606 | 38.8540285 | 61.5858529 | 95.864124 | 73.8374582 | 87.6752882 |
| 20.6955481 | 15.572304 | 9.71350712 | 12.1152497 | 17.2255848 | 17.0394134 | 13.0055415 |
| 2.95650687 | 8.76663038 | 0 | 5.04802073 | 7.48938469 | 9.08768716 | 8.07240505 |

| | | | | | | |
|---|---|---|---|---|---|---|
| 75.3909251 | 69.4986158 | 19.4270142 | 70.6722902 | 58.4172006 | 46.0064163 | 73.5485793 |
| 0 | 6.53690715 | 0 | 6.05762487 | 9.36173086 | 4.91871068 | 3.13926863 |
| 11.8260275 | 12.4578432 | 9.71350712 | 13.1248539 | 6.74044622 | 17.0394134 | 4.48466947 |
| 26.6085618 | 19.2635167 | 9.71350712 | 23.2208954 | 26.9617849 | 22.7192179 | 23.3202812 |
| 50.2606167 | 72.9014526 | 38.8540285 | 72.6914985 | 65.9065853 | 54.526123 | 61.4399717 |
| 8.8695206 | 7.61312638 | 9.71350712 | 24.2304995 | 18.7234617 | 10.2236481 | 15.0236427 |
| 26.6085618 | 37.2581791 | 29.1405214 | 29.2785202 | 28.4596618 | 17.0394134 | 29.1503516 |
| 23.6520549 | 26.5305919 | 9.71350712 | 19.1824788 | 14.9787694 | 15.9034525 | 16.8175105 |
| 20.6955481 | 19.0328159 | 19.4270142 | 17.1632705 | 6.74044622 | 15.9034525 | 15.0236427 |
| 73.9126717 | 91.0114654 | 67.9945498 | 81.7779358 | 69.6512776 | 71.5655364 | 107.856301 |
| 2.95650687 | 4.84471679 | 9.71350712 | 8.07683317 | 3.74469235 | 12.4955698 | 5.38160336 |
| 38.4345893 | 35.7586239 | 29.1405214 | 47.4513948 | 35.200108 | 51.1182403 | 52.0221659 |
| 5.91301374 | 3.92191359 | 19.4270142 | 11.1056456 | 6.74044622 | 11.359609 | 5.60583684 |
| 5.91301374 | 15.1109024 | 9.71350712 | 16.2546267 | 8.98726163 | 18.1753743 | 10.9874402 |
| 130.086302 | 145.802905 | 67.9945498 | 112.06606 | 127.31954 | 103.372441 | 164.58737 |
| 5.91301374 | 8.07452798 | 0 | 5.04802073 | 7.48938469 | 9.08768716 | 7.6239381 |
| 38.4345893 | 26.6459423 | 9.71350712 | 32.3073327 | 33.7022311 | 30.6709442 | 51.125232 |
| 2.95650687 | 12.5731936 | 9.71350712 | 3.02881244 | 8.98726163 | 0 | 6.91311799 |
| 8.8695206 | 3.46051199 | 19.4270142 | 6.05762487 | 11.9830155 | 10.2236481 | 7.17547115 |
| 11.8260275 | 10.4968864 | 29.1405214 | 6.05762487 | 14.2298309 | 9.08768716 | 9.86627284 |
| 2.95650687 | 4.84471679 | 0 | 14.134458 | 16.4766463 | 13.6315307 | 12.332841 |
| 8.8695206 | 10.2661856 | 9.71350712 | 19.1824788 | 25.4639079 | 17.0394134 | 3.81196905 |
| 2.95650687 | 12.8038944 | 0 | 11.1056456 | 3.74469235 | 5.67980448 | 8.07240505 |
| 14.7825343 | 11.8810912 | 19.4270142 | 14.134458 | 8.98726163 | 7.95172627 | 7.17547115 |
| 22.1738015 | 14.707176 | 29.1405214 | 20.1920829 | 26.2128464 | 25.5591201 | 19.171962 |
| 2.95650687 | 9.45873277 | 9.71350712 | 7.06722902 | 11.9830155 | 5.67980448 | 6.05430379 |
| 53.2171236 | 37.8349311 | 19.4270142 | 37.3553534 | 26.9617849 | 40.8945922 | 50.0040646 |
| 8.8695206 | 8.07452798 | 0 | 7.06722902 | 6.74044622 | 10.2236481 | 7.17547115 |
| 14.7825343 | 14.4188 | 0 | 9.08643731 | 7.48938469 | 6.81576537 | 7.84817157 |
| 2.95650687 | 6.05589598 | 9.71350712 | 10.6008435 | 9.36173086 | 15.9034525 | 3.69985231 |
| 17.7390412 | 26.4152415 | 9.71350712 | 14.134458 | 14.9787694 | 11.359609 | 16.1448101 |
| 2.95650687 | 7.95917758 | 0 | 13.1248539 | 8.23832316 | 5.67980448 | 7.6239381 |
| 17.7390412 | 17.5332608 | 9.71350712 | 15.1440622 | 11.234077 | 9.08768716 | 13.0055415 |
| 2.95650687 | 5.53681919 | 0 | 16.1536663 | 7.48938469 | 23.8551788 | 4.93313642 |
| 38.4345893 | 53.7532863 | 29.1405214 | 27.2593119 | 28.4596618 | 13.6315307 | 28.0291842 |
| 0 | 8.53592958 | 9.71350712 | 6.05762487 | 5.99150775 | 16.471433 | 6.390654 |
| 8.8695206 | 14.4188 | 0 | 7.06722902 | 8.98726163 | 13.6315307 | 8.07240505 |
| 2.95650687 | 12.5731936 | 0 | 7.06722902 | 8.98726163 | 13.6315307 | 8.74510547 |
| 11.8260275 | 16.6104576 | 0 | 11.1056456 | 7.48938469 | 3.40788269 | 6.95123768 |
| 32.5215755 | 33.4516159 | 9.71350712 | 29.2785202 | 27.7107234 | 19.3113352 | 28.2534177 |
| 22.6764077 | 17.6486112 | 0 | 22.2112912 | 22.4681541 | 22.7192179 | 18.8356118 |
| 2.95650687 | 3.11446079 | 0 | 2.01920829 | 6.74044622 | 9.08768716 | 5.15736989 |
| 20.6955481 | 19.6095679 | 0 | 12.1152497 | 10.4851386 | 15.9034525 | 18.8356118 |
| 26.6085618 | 23.3007807 | 19.4270142 | 18.1728746 | 25.4639079 | 19.3113352 | 17.041744 |
| 23.6520549 | 16.4951072 | 29.1405214 | 16.1536663 | 16.4766463 | 107.916285 | 11.4359072 |
| 8.8695206 | 9.80478397 | 29.1405214 | 11.1056456 | 17.2255848 | 14.7674916 | 11.8843741 |

| | | | | | | |
|---|---|---|---|---|---|---|
| 17.7390412 | 24.6849855 | 0 | 30.2881244 | 17.9745233 | 17.0394134 | 26.9080168 |
| 20.6955481 | 26.8766431 | 0 | 14.134458 | 14.9787694 | 20.4472961 | 23.3202812 |
| 0 | 1.3842048 | 0 | 4.03841658 | 8.98726163 | 14.7674916 | 5.60583684 |
| 50.2606167 | 72.5554014 | 48.5675356 | 80.7683317 | 72.6470315 | 68.1576537 | 67.0458086 |
| 11.8260275 | 8.99733118 | 0 | 11.1056456 | 4.49363081 | 7.95172627 | 8.29663852 |
| 14.7825343 | 21.6858751 | 0 | 9.59123938 | 17.9745233 | 11.359609 | 15.4721097 |
| 5.91301374 | 5.99822078 | 0 | 6.05762487 | 4.49363081 | 12.4955698 | 6.72700421 |
| 17.7390412 | 5.65216958 | 0 | 14.134458 | 7.48938469 | 9.08768716 | 9.19357241 |
| 5.91301374 | 6.80567358 | 0 | 7.06722902 | 7.48938469 | 5.67980448 | 8.29663852 |
| 28.2937707 | 17.3394721 | 24.575173 | 16.0527059 | 23.8761584 | 22.4352277 | 23.0332624 |
| 97.5647266 | 295.181673 | 106.848578 | 61.5858529 | 83.8811085 | 45.4384358 | 90.3660898 |
| 17.7390412 | 8.99733118 | 0 | 5.04802073 | 12.731954 | 5.67980448 | 13.4540084 |
| 14.7825343 | 15.6876544 | 9.71350712 | 15.1440622 | 20.2213387 | 19.3113352 | 18.8356118 |
| 8.8695206 | 7.03637438 | 0 | 8.07683317 | 7.48938469 | 3.40788269 | 7.84817157 |
| 369.563358 | 387.115941 | 281.691706 | 336.198181 | 257.634833 | 252.183319 | 362.80976 |
| 41.3910961 | 50.2927743 | 19.4270142 | 31.2977285 | 26.2128464 | 35.2147878 | 31.3926863 |
| 14.7825343 | 19.6095679 | 19.4270142 | 12.1152497 | 18.7234617 | 21.583257 | 20.853713 |
| 2.95650687 | 10.8429376 | 19.4270142 | 23.2208954 | 8.98726163 | 7.95172627 | 11.4359072 |
| 23.6520549 | 18.456064 | 19.4270142 | 27.2593119 | 18.7234617 | 21.583257 | 27.3564838 |
| 2.95650687 | 4.26796479 | 9.71350712 | 6.05762487 | 13.4808924 | 6.81576537 | 7.51182136 |
| 59.1301374 | 27.5687455 | 67.9945498 | 68.6530819 | 45.6852466 | 52.2542012 | 53.329447 |
| 0 | 2.53770879 | 0 | 3.02881244 | 9.7362001 | 5.67980448 | 6.50277073 |
| 5.91301374 | 4.96006719 | 9.71350712 | 12.1152497 | 6.74044622 | 9.08768716 | 6.50277073 |
| 5.91301374 | 2.65305919 | 9.71350712 | 7.06722902 | 5.99150775 | 11.359609 | 6.95123768 |
| 8.8695206 | 5.07541759 | 19.4270142 | 11.1056456 | 6.74044622 | 6.81576537 | 7.39970463 |
| 14.7825343 | 5.19076799 | 0 | 13.1248539 | 9.7362001 | 7.95172627 | 9.19357241 |
| 5.91301374 | 8.07452798 | 9.71350712 | 9.08643731 | 10.4851386 | 10.2236481 | 10.7632067 |
| 88.695206 | 83.7443902 | 19.4270142 | 86.8259565 | 77.8896008 | 69.2936146 | 89.6933894 |
| 0 | 13.7266976 | 9.71350712 | 7.06722902 | 6.74044622 | 9.08768716 | 10.3147398 |
| 11.8260275 | 27.7994463 | 9.71350712 | 21.2016871 | 29.2086003 | 19.3113352 | 28.0291842 |
| 0 | 7.38242558 | 9.71350712 | 4.03841658 | 7.48938469 | 3.40788269 | 6.05430379 |
| 11.8260275 | 13.9573984 | 0 | 9.08643731 | 8.23832316 | 5.67980448 | 10.3147398 |
| 8.8695206 | 16.149056 | 9.71350712 | 15.1440622 | 16.4766463 | 22.7192179 | 20.853713 |
| 14.7825343 | 4.49866559 | 0 | 7.06722902 | 8.23832316 | 5.67980448 | 9.86627284 |
| 8.8695206 | 7.84382718 | 9.71350712 | 11.1056456 | 10.4851386 | 12.4955698 | 7.39970463 |
| 5.91301374 | 9.92013437 | 29.1405214 | 7.06722902 | 5.99150775 | 9.08768716 | 9.64203936 |
| 14.7825343 | 7.84382718 | 0 | 7.06722902 | 8.23832316 | 9.08768716 | 10.3147398 |
| 5.91301374 | 2.42235839 | 9.71350712 | 7.06722902 | 5.99150775 | 9.08768716 | 5.83007031 |
| 2.95650687 | 11.9964416 | 9.71350712 | 7.06722902 | 8.98726163 | 9.08768716 | 9.86627284 |
| 8.8695206 | 10.7275872 | 0 | 8.07683317 | 10.4851386 | 6.81576537 | 10.7632067 |
| 2.95650687 | 1.6149056 | 0 | 5.04802073 | 7.48938469 | 6.81576537 | 8.29663852 |
| 8.8695206 | 9.68943357 | 19.4270142 | 10.0960415 | 8.98726163 | 7.95172627 | 8.74510547 |
| 17.7390412 | 13.7266976 | 19.4270142 | 20.1920829 | 10.4851386 | 15.9034525 | 13.4540084 |
| 14.7825343 | 7.67080158 | 19.4270142 | 17.1632705 | 23.5915618 | 11.359609 | 12.2207243 |
| 2.95650687 | 10.7275872 | 9.71350712 | 7.06722902 | 13.4808924 | 4.54384358 | 6.16642052 |
| 14.7825343 | 8.76663038 | 38.8540285 | 17.1632705 | 15.7277078 | 14.7674916 | 17.041744 |

| | | | | | | |
|---|---|---|---|---|---|---|
| 14.7825343 | 10.958288 | 0.77708057 | 14.134458 | 11.234077 | 9.08768716 | 13.9024754 |
| 11.8260275 | 4.26796479 | 0 | 12.6200518 | 7.11491546 | 8.51970672 | 9.7541561 |
| 32.5215755 | 14.0727488 | 19.4270142 | 17.1632705 | 20.2213387 | 15.9034525 | 18.6113783 |
| 23.6520549 | 37.6042303 | 0 | 22.2112912 | 16.4766463 | 14.7674916 | 21.30218 |
| 5.91301374 | 6.11357118 | 0 | 3.02881244 | 5.99150775 | 10.2236481 | 9.19357241 |
| 100.521233 | 108.314025 | 77.708057 | 85.8163524 | 90.6215547 | 69.2936146 | 113.462138 |
| 11.8260275 | 12.111792 | 0 | 9.08643731 | 15.7277078 | 12.4955698 | 9.19357241 |
| 8.8695206 | 10.4968864 | 19.4270142 | 12.1152497 | 8.98726163 | 6.81576537 | 6.27853726 |
| 94.6082198 | 98.8552925 | 38.8540285 | 64.6146653 | 59.1661391 | 60.2059275 | 80.0513501 |
| 47.3041099 | 112.370899 | 38.8540285 | 56.5378322 | 65.4047965 | 70.4295755 | 55.4484533 |
| 22.9129282 | 15.6046021 | 9.71350712 | 17.3651913 | 20.460999 | 38.4295571 | 13.8643557 |
| 5.91301374 | 11.4196896 | 0 | 7.06722902 | 7.48938469 | 13.6315307 | 3.58773558 |
| 8.8695206 | 1.8456064 | 0 | 8.07683317 | 9.7362001 | 6.81576537 | 8.40875526 |
| 14.7825343 | 11.8810912 | 9.71350712 | 10.0960415 | 9.7362001 | 7.95172627 | 9.52992263 |
| 29.5650687 | 44.6406047 | 29.1405214 | 22.2112912 | 26.2128464 | 22.7192179 | 27.1322503 |
| 0 | 13.842048 | 0 | 2.01920829 | 7.48938469 | 9.08768716 | 8.52087199 |
| 2.95650687 | 9.11268158 | 0 | 5.04802073 | 7.48938469 | 7.95172627 | 6.72700421 |
| 79.8256854 | 70.4790942 | 67.9945498 | 80.7683317 | 74.1449084 | 56.7980448 | 84.5360195 |
| 20.6955481 | 19.3788671 | 0 | 20.1920829 | 8.98726163 | 19.3113352 | 18.6113783 |
| 8.8695206 | 9.11268158 | 0 | 4.03841658 | 8.98726163 | 2.27192179 | 6.72700421 |
| 133.042809 | 133.460412 | 67.9945498 | 142.354185 | 138.553617 | 127.22762 | 140.594388 |
| 650.431511 | 391.499257 | 349.686256 | 590.618425 | 528.001621 | 540.717386 | 667.094584 |
| 44.347603 | 20.7630719 | 9.71350712 | 24.2304995 | 21.7192156 | 18.1753743 | 25.562616 |
| 8.8695206 | 21.6858751 | 0 | 8.07683317 | 15.7277078 | 13.6315307 | 10.0905063 |
| 11.8260275 | 11.0736384 | 0 | 9.08643731 | 11.9830155 | 7.95172627 | 11.2116737 |
| 0 | 1.6149056 | 9.71350712 | 11.1056456 | 5.99150775 | 6.81576537 | 7.39970463 |
| 22.1738015 | 21.7435503 | 19.4270142 | 12.1152497 | 34.8256388 | 32.3748855 | 32.2896202 |
| 8.8695206 | 7.38242558 | 0 | 2.01920829 | 8.98726163 | 7.95172627 | 9.41780589 |
| 2.95650687 | 13.9573984 | 0 | 18.1728746 | 8.98726163 | 6.81576537 | 10.7632067 |
| 26.6085618 | 20.4170207 | 0 | 14.134458 | 26.2128464 | 23.8551788 | 19.2840787 |
| 20.6955481 | 5.88287038 | 19.4270142 | 18.1728746 | 20.2213387 | 15.9034525 | 18.3871448 |
| 65.0431511 | 63.7310958 | 29.1405214 | 47.4513948 | 66.2810545 | 69.8615951 | 66.8215751 |
| 5.91301374 | 9.22803198 | 0 | 10.0960415 | 14.2298309 | 12.4955698 | 11.4359072 |
| 8.8695206 | 8.88198078 | 9.71350712 | 16.1536663 | 15.3532386 | 14.7674916 | 10.5389733 |
| 5.91301374 | 2.30700799 | 0 | 6.05762487 | 10.4851386 | 6.81576537 | 6.72700421 |
| 73.9126717 | 64.8269246 | 58.2810427 | 90.8643731 | 77.1406623 | 55.6620839 | 90.1418564 |
| 5.91301374 | 23.9928831 | 0 | 18.1728746 | 14.2298309 | 20.4472961 | 21.30218 |
| 8.8695206 | 7.84382718 | 0 | 8.07683317 | 7.48938469 | 9.08768716 | 6.95123768 |
| 17.7390412 | 12.4578432 | 9.71350712 | 15.1440622 | 17.9745233 | 14.7674916 | 12.5570745 |
| 20.6955481 | 14.4188 | 9.71350712 | 14.134458 | 20.9702771 | 17.0394134 | 23.5445147 |
| 8.8695206 | 9.22803198 | 0 | 1.00960415 | 8.23832316 | 4.54384358 | 6.27853726 |
| 118.260275 | 62.8659678 | 87.4215641 | 133.267747 | 119.830155 | 101.10052 | 104.044332 |
| 14.7825343 | 16.0337056 | 29.1405214 | 27.2593119 | 17.9745233 | 24.9911397 | 21.9748804 |
| 0 | 8.99733118 | 9.71350712 | 12.1152497 | 4.49363081 | 7.95172627 | 6.72700421 |
| 2.95650687 | 4.15261439 | 19.4270142 | 8.07683317 | 11.234077 | 9.08768716 | 9.41780589 |
| 0 | 8.30522878 | 0 | 6.05762487 | 11.9830155 | 7.95172627 | 9.64203936 |

| | | | | | | |
|---|---|---|---|---|---|---|
| 56.1736305 | 33.6823167 | 48.5675356 | 47.4513948 | 44.9363081 | 51.1182403 | 56.8050658 |
| 8.8695206 | 8.30522878 | 9.71350712 | 8.07683317 | 5.24256928 | 11.359609 | 6.05430379 |
| 5.91301374 | 5.30611839 | 19.4270142 | 5.04802073 | 5.99150775 | 9.08768716 | 6.27853726 |
| 23.6520549 | 18.9174655 | 67.9945498 | 29.2785202 | 34.4511696 | 32.942866 | 31.6169198 |
| 8.8695206 | 11.8810912 | 0 | 9.08643731 | 10.4851386 | 6.81576537 | 11.4359072 |
| 26.6085618 | 22.8393791 | 19.4270142 | 24.2304995 | 27.7107234 | 28.3990224 | 25.3383825 |
| 11.8260275 | 4.03726399 | 9.71350712 | 4.03841658 | 8.23832316 | 2.27192179 | 4.70890294 |
| 269.042125 | 182.253632 | 184.556635 | 228.170537 | 220.18791 | 215.83257 | 234.772447 |
| 8.8695206 | 2.42235839 | 0 | 9.08643731 | 11.9830155 | 4.54384358 | 6.27853726 |
| 251.303084 | 353.548975 | 106.848578 | 279.660348 | 202.213387 | 219.240453 | 298.006286 |
| 23.6520549 | 12.9192448 | 0 | 11.1056456 | 20.9702771 | 7.95172627 | 22.1991139 |
| 11.8260275 | 10.4968864 | 9.71350712 | 15.1440622 | 14.9787694 | 15.9034525 | 15.9205766 |
| 8.8695206 | 10.958288 | 9.71350712 | 28.2689161 | 20.2213387 | 12.4955698 | 15.0236427 |
| 20.6955481 | 8.88198078 | 0 | 15.1440622 | 15.7277078 | 18.1753743 | 15.9205766 |
| 17.7390412 | 14.8802016 | 0 | 14.134458 | 13.4808924 | 11.359609 | 14.6872925 |
| 2.95650687 | 6.22892158 | 9.71350712 | 4.03841658 | 12.731954 | 9.08768716 | 6.72700421 |
| 5.91301374 | 3.11446079 | 0 | 2.01920829 | 5.24256928 | 9.08768716 | 5.60583684 |
| 8.8695206 | 17.6486112 | 0 | 16.1536663 | 16.4766463 | 9.08768716 | 12.781308 |
| 14.7825343 | 22.2626271 | 9.71350712 | 10.0960415 | 11.234077 | 14.7674916 | 22.4233474 |
| 8.8695206 | 2.53770879 | 29.1405214 | 7.06722902 | 9.7362001 | 9.08768716 | 8.52087199 |
| 11.8260275 | 24.6849855 | 0 | 16.1536663 | 11.234077 | 11.359609 | 11.2116737 |
| 159.651371 | 50.8695263 | 87.4215641 | 117.114081 | 104.851386 | 123.819738 | 120.189142 |
| 5.91301374 | 10.7275872 | 0 | 3.02881244 | 8.23832316 | 7.95172627 | 7.17547115 |
| 67.999658 | 79.361075 | 29.1405214 | 30.2881244 | 37.4469235 | 34.0788269 | 59.4218705 |
| 14.7825343 | 17.9946624 | 0 | 18.1728746 | 14.9787694 | 15.9034525 | 11.4359072 |
| 11.8260275 | 8.88198078 | 19.4270142 | 9.08643731 | 14.2298309 | 15.9034525 | 13.4540084 |
| 11.8260275 | 5.19076799 | 9.71350712 | 6.05762487 | 8.23832316 | 7.95172627 | 7.6239381 |
| 32.5215755 | 22.1472767 | 19.4270142 | 30.2881244 | 32.9532926 | 29.5349833 | 42.8285934 |
| 34.2363495 | 11.3816239 | 38.8540285 | 36.9313196 | 19.8468694 | 22.628341 | 16.5551574 |
| 20.6955481 | 29.4143519 | 0 | 20.1920829 | 24.7149695 | 34.0788269 | 21.30218 |
| 17.7390412 | 15.1109024 | 38.8540285 | 31.2977285 | 24.7149695 | 23.8551788 | 23.7687482 |
| 47.3041099 | 29.2990015 | 9.71350712 | 32.3073327 | 40.4426773 | 36.3507487 | 40.8104922 |
| 62.0866442 | 190.155134 | 9.71350712 | 40.3841658 | 26.9617849 | 26.1271006 | 28.2534177 |
| 35.4780824 | 29.2990015 | 9.71350712 | 28.2689161 | 41.9405543 | 37.4867095 | 36.7742897 |
| 85.7386992 | 110.505683 | 29.1405214 | 95.9123938 | 88.3747393 | 70.4295755 | 113.013671 |
| 8.8695206 | 11.4196896 | 0 | 11.1056456 | 14.2298309 | 7.95172627 | 8.52087199 |
| 35.4780824 | 58.8287038 | 19.4270142 | 34.326541 | 23.2170925 | 19.3113352 | 29.7109352 |
| 11.8260275 | 16.9565088 | 19.4270142 | 13.1248539 | 20.9702771 | 15.9034525 | 19.3961955 |
| 2.95650687 | 6.11357118 | 0 | 2.01920829 | 6.74044622 | 9.08768716 | 7.39970463 |
| 5.91301374 | 17.30256 | 9.71350712 | 11.1056456 | 11.234077 | 3.40788269 | 12.781308 |
| 8.8695206 | 14.995552 | 19.4270142 | 16.1536663 | 12.731954 | 11.359609 | 10.9874402 |
| 20.6955481 | 8.30522878 | 19.4270142 | 6.05762487 | 17.2255848 | 12.4955698 | 14.5751758 |
| 38.4345893 | 18.8597903 | 9.71350712 | 18.1728746 | 21.7192156 | 20.4472961 | 21.0779465 |
| 85.7386992 | 160.394731 | 97.1350712 | 74.7107068 | 77.1406623 | 85.1970672 | 91.3751405 |
| 41.3910961 | 146.552683 | 33.9972749 | 72.1866964 | 76.7661931 | 44.8704554 | 65.0277073 |
| 5.91301374 | 7.15172478 | 29.1405214 | 10.0960415 | 20.9702771 | 15.9034525 | 12.781308 |

| | | | | | | |
|---|---|---|---|---|---|---|
| 47.3041099 | 45.3327071 | 9.71350712 | 65.6242695 | 49.429939 | 55.6620839 | 67.9427425 |
| 56.1736305 | 57.2714734 | 19.4270142 | 34.831343 | 42.6894927 | 35.7827682 | 45.6315119 |
| 11.8260275 | 17.879312 | 9.71350712 | 10.0960415 | 10.8596078 | 7.95172627 | 11.0995569 |
| 23.6520549 | 24.5696351 | 9.71350712 | 23.2208954 | 23.2170925 | 24.9911397 | 27.1322503 |
| 23.6520549 | 20.2439951 | 29.1405214 | 33.8217389 | 20.2213387 | 46.5743967 | 14.2388256 |
| 5.91301374 | 9.57408317 | 0 | 6.05762487 | 7.48938469 | 5.67980448 | 5.60583684 |
| 8.8695206 | 6.34427198 | 0 | 3.02881244 | 2.99575388 | 10.2236481 | 6.05430379 |
| 35.4780824 | 21.9165759 | 9.71350712 | 20.1920829 | 12.731954 | 21.583257 | 23.7687482 |
| 8.8695206 | 9.57408317 | 38.8540285 | 14.134458 | 11.9830155 | 17.0394134 | 18.1629114 |
| 14.7825343 | 4.03726399 | 0 | 20.1920829 | 7.48938469 | 13.6315307 | 21.7775549 |
| 17.7390412 | 31.0292575 | 9.71350712 | 20.1920829 | 23.966031 | 9.08768716 | 19.2840787 |
| 20.6955481 | 16.0337056 | 19.4270142 | 15.6488643 | 18.3489925 | 9.65566761 | 15.5842264 |
| 29.5650687 | 114.658297 | 0 | 16.1536663 | 16.4766463 | 18.1753743 | 21.0779465 |
| 212.868494 | 329.440742 | 87.4215641 | 215.045683 | 192.477187 | 197.657196 | 255.850393 |
| 2.95650687 | 12.8038944 | 19.4270142 | 16.1536663 | 12.731954 | 20.4472961 | 16.8175105 |
| 0 | 2.07630719 | 0 | 5.04802073 | 11.234077 | 1.1359609 | 2.91503516 |
| 2.95650687 | 11.53504 | 0 | 10.0960415 | 7.48938469 | 3.40788269 | 6.72700421 |
| 50.2606167 | 92.9724221 | 29.1405214 | 70.6722902 | 56.1703852 | 64.749771 | 63.6823065 |
| 2.95650687 | 6.11357118 | 0 | 3.02881244 | 7.48938469 | 5.67980448 | 3.13926863 |
| 0 | 4.15261439 | 0 | 1.00960415 | 7.48938469 | 3.40788269 | 3.3635021 |
| 8.8695206 | 17.5332608 | 9.71350712 | 12.1152497 | 10.4851386 | 13.6315307 | 13.4540084 |
| 17.7390412 | 19.8402687 | 9.71350712 | 6.05762487 | 7.48938469 | 15.9034525 | 6.27853726 |
| 11.8260275 | 12.9192448 | 9.71350712 | 14.134458 | 14.2298309 | 12.4955698 | 11.2116737 |
| 2.95650687 | 4.03726399 | 0 | 5.04802073 | 6.74044622 | 11.359609 | 5.15736989 |
| 14.7825343 | 10.6122368 | 0 | 7.06722902 | 5.99150775 | 11.359609 | 10.3147398 |
| 8.8695206 | 17.4179104 | 9.71350712 | 17.1632705 | 6.74044622 | 15.9034525 | 10.0905063 |
| 8.8695206 | 7.61312638 | 9.71350712 | 5.04802073 | 1.49787694 | 10.2236481 | 6.27853726 |
| 1.9808596 | 10.7471967 | 29.1405214 | 16.9916378 | 16.7912005 | 18.8342316 | 12.6512526 |
| 5.91301374 | 5.42146879 | 19.4270142 | 2.01920829 | 4.49363081 | 10.2236481 | 8.29663852 |
| 11.8260275 | 9.68943357 | 19.4270142 | 12.1152497 | 8.98726163 | 13.6315307 | 10.9874402 |
| 11.8260275 | 17.5332608 | 0 | 15.1440622 | 9.7362001 | 20.4472961 | 7.17547115 |
| 41.3910961 | 40.7186911 | 0 | 34.326541 | 30.7064772 | 36.3507487 | 33.4107876 |
| 8.8695206 | 6.51729758 | 4.85675356 | 10.6008435 | 13.4808924 | 10.2236481 | 11.9964908 |
| 38.4345893 | 49.4853215 | 19.4270142 | 63.6050612 | 35.200108 | 45.4384358 | 47.9859633 |
| 5.91301374 | 19.7249183 | 0 | 4.03841658 | 6.74044622 | 10.2236481 | 4.93313642 |
| 8.8695206 | 4.26796479 | 19.4270142 | 2.01920829 | 6.74044622 | 11.359609 | 10.3147398 |
| 195.129453 | 146.725708 | 116.562085 | 178.699934 | 176.749479 | 152.21876 | 194.634655 |
| 0 | 4.15261439 | 0 | 6.05762487 | 3.74469235 | 10.2236481 | 2.91503516 |
| 19.2172946 | 14.8029168 | 19.4270142 | 43.3322099 | 31.3955006 | 54.844192 | 20.6474182 |
| 26.6085618 | 47.9857663 | 0 | 25.2401036 | 32.9532926 | 32.942866 | 32.0653867 |
| 5.91301374 | 6.80567358 | 0 | 12.1152497 | 16.4766463 | 20.4472961 | 4.03620252 |
| 4.4347603 | 2.65305919 | 9.71350712 | 4.03841658 | 5.99150775 | 15.9034525 | 4.83895836 |
| 4.4347603 | 8.18987838 | 0 | 5.5528228 | 4.86810005 | 9.65566761 | 2.91503516 |
| 5.91301374 | 7.95917758 | 9.71350712 | 9.08643731 | 11.234077 | 5.67980448 | 6.05430379 |
| 14.7825343 | 14.5341504 | 0 | 19.1824788 | 11.9830155 | 11.359609 | 10.5389733 |
| 124.173288 | 59.0594046 | 67.9945498 | 111.056456 | 87.6258009 | 70.4295755 | 100.456596 |

| | | | | | | |
|---|---|---|---|---|---|---|
| 2.95650687 | 10.958288 | 9.71350712 | 7.06722902 | 14.9787694 | 10.2236481 | 12.1086076 |
| 2.95650687 | 4.15261439 | 0 | 2.01920829 | 4.49363081 | 10.2236481 | 3.58773558 |
| 32.5215755 | 20.1863199 | 19.4270142 | 27.2593119 | 26.2128464 | 23.8551788 | 32.2896202 |
| 8.8695206 | 18.2253632 | 19.4270142 | 18.1728746 | 17.2255848 | 7.95172627 | 13.9024754 |
| 41.3910961 | 104.161411 | 19.4270142 | 51.4898114 | 39.6937389 | 52.2542012 | 54.2645006 |
| 2.95650687 | 8.99733118 | 19.4270142 | 15.1440622 | 8.98726163 | 9.08768716 | 12.332841 |
| 0 | 8.99733118 | 0 | 7.06722902 | 7.48938469 | 11.359609 | 10.3147398 |
| 5.91301374 | 6.22892158 | 0 | 9.08643731 | 10.4851386 | 7.95172627 | 3.81196905 |
| 1114.60309 | 898.348913 | 679.945498 | 920.758981 | 899.475101 | 858.786437 | 1002.99633 |
| 2.95650687 | 18.3407136 | 9.71350712 | 8.07683317 | 8.23832316 | 9.08768716 | 7.39970463 |
| 26.6085618 | 41.6414943 | 19.4270142 | 40.3841658 | 26.9617849 | 17.0394134 | 28.9261181 |
| 26.6085618 | 33.6823167 | 9.71350712 | 42.4033741 | 39.6937389 | 37.4867095 | 34.9804219 |
| 8.8695206 | 10.4968864 | 9.71350712 | 12.1152497 | 7.48938469 | 6.81576537 | 9.64203936 |
| 17.7390412 | 26.9919935 | 9.71350712 | 9.08643731 | 7.48938469 | 9.08768716 | 14.3509423 |
| 5.91301374 | 11.8810912 | 9.71350712 | 12.1152497 | 14.2298309 | 13.6315307 | 12.1086076 |
| 5.91301374 | 11.8810912 | 0 | 5.04802073 | 7.48938469 | 6.81576537 | 4.93313642 |
| 11.8260275 | 8.30522878 | 9.71350712 | 11.1056456 | 14.9787694 | 18.1753743 | 15.9205766 |
| 5.91301374 | 13.7266976 | 0 | 8.07683317 | 14.2298309 | 3.40788269 | 8.29663852 |
| 8.8695206 | 10.7275872 | 0 | 22.2112912 | 20.2213387 | 10.2236481 | 11.6601406 |
| 5.91301374 | 11.53504 | 9.71350712 | 13.1248539 | 11.9830155 | 17.0394134 | 10.0905063 |
| 1.47825343 | 7.26707518 | 9.71350712 | 15.1440622 | 11.9830155 | 18.1753743 | 8.18452178 |
| 23.6520549 | 14.995552 | 9.71350712 | 19.1824788 | 17.9745233 | 18.1753743 | 21.5264135 |
| 162.607878 | 122.848176 | 165.129621 | 160.527059 | 157.277078 | 135.179347 | 162.569268 |
| 2.95650687 | 2.53770879 | 9.71350712 | 6.05762487 | 11.9830155 | 6.81576537 | 4.48466947 |
| 2.95650687 | 5.30611839 | 0 | 7.57203109 | 8.98726163 | 14.7674916 | 2.35445147 |
| 35.4780824 | 39.6805375 | 19.4270142 | 36.3457492 | 29.9575388 | 30.6709442 | 35.4288888 |
| 5.91301374 | 2.99911039 | 29.1405214 | 5.04802073 | 6.74044622 | 9.08768716 | 6.27853726 |
| 11.8260275 | 14.0727488 | 9.71350712 | 10.0960415 | 5.24256928 | 12.4955698 | 10.0905063 |
| 41.3910961 | 22.3779775 | 29.1405214 | 38.3649575 | 35.200108 | 48.8463185 | 37.6712236 |
| 11.8260275 | 20.9937727 | 9.71350712 | 18.1728746 | 10.4851386 | 9.08768716 | 11.8843741 |
| 5.91301374 | 14.4188 | 0 | 4.03841658 | 9.7362001 | 9.08768716 | 5.15736989 |
| 14.7825343 | 30.9139071 | 0 | 6.05762487 | 5.99150775 | 9.08768716 | 5.38160336 |
| 0 | 8.88198078 | 9.71350712 | 16.1536663 | 3.74469235 | 4.54384358 | 3.58773558 |
| 53.2171236 | 56.6370462 | 19.4270142 | 39.3745617 | 36.697985 | 28.3990224 | 47.7617299 |
| 8.8695206 | 3.57586239 | 0 | 2.01920829 | 11.9830155 | 3.40788269 | 5.60583684 |
| 5.91301374 | 14.0727488 | 29.1405214 | 10.0960415 | 8.23832316 | 13.6315307 | 10.9874402 |
| 5.91301374 | 3.92191359 | 0 | 7.06722902 | 7.48938469 | 10.2236481 | 9.86627284 |
| 5.91301374 | 33.1055647 | 0 | 5.04802073 | 7.48938469 | 12.4955698 | 8.29663852 |
| 8.8695206 | 4.61401599 | 0 | 8.07683317 | 10.4851386 | 5.67980448 | 6.27853726 |
| 14.7825343 | 15.8030048 | 0 | 21.2016871 | 26.2128464 | 32.942866 | 13.6782419 |
| 0 | 3.46051199 | 9.71350712 | 1.00960415 | 7.48938469 | 3.40788269 | 4.03620252 |
| 23.6520549 | 28.2608479 | 48.5675356 | 28.2689161 | 23.2170925 | 27.2630615 | 27.1322503 |
| 5.91301374 | 14.3034496 | 9.71350712 | 14.134458 | 14.2298309 | 11.359609 | 5.38160336 |
| 0 | 11.1889888 | 0 | 6.05762487 | 2.99575388 | 12.4955698 | 10.9874402 |
| 29.5650687 | 22.9547295 | 29.1405214 | 28.2689161 | 22.4681541 | 21.583257 | 32.2896202 |
| 8.8695206 | 20.8784223 | 19.4270142 | 6.05762487 | 13.4808924 | 7.95172627 | 9.41780589 |

| | | | | | | |
|---|---|---|---|---|---|---|
| 2.95650687 | 6.11357118 | 0 | 6.05762487 | 11.234077 | 4.54384358 | 6.05430379 |
| 14.7825343 | 2.65305919 | 0 | 9.08643731 | 8.23832316 | 6.81576537 | 8.52087199 |
| 11.8260275 | 11.8810912 | 9.71350712 | 7.06722902 | 4.49363081 | 9.08768716 | 9.41780589 |
| 8.8695206 | 6.80567358 | 0 | 2.01920829 | 8.98726163 | 13.6315307 | 8.96933894 |
| 2.95650687 | 25.0310367 | 9.71350712 | 33.3169368 | 25.4639079 | 45.4384358 | 9.86627284 |
| 53.2171236 | 52.4844319 | 19.4270142 | 25.2401036 | 31.4554157 | 20.4472961 | 33.1865541 |
| 17.7390412 | 8.42057918 | 1.359891 | 9.08643731 | 7.48938469 | 5.67980448 | 6.05430379 |
| 5.91301374 | 12.688544 | 9.71350712 | 7.06722902 | 11.9830155 | 13.6315307 | 3.13926863 |
| 17.7390412 | 13.1499456 | 9.71350712 | 14.134458 | 7.48938469 | 6.81576537 | 15.0236427 |
| 41.3910961 | 91.8189182 | 38.8540285 | 44.4225824 | 47.932062 | 45.4384358 | 53.3675667 |
| 0 | 11.1889888 | 0 | 2.01920829 | 6.74044622 | 9.08768716 | 6.05430379 |
| 5.91301374 | 10.7275872 | 0 | 13.1248539 | 3.74469235 | 12.4955698 | 8.74510547 |
| 65.0431511 | 69.7869918 | 48.5675356 | 52.4994156 | 30.7064772 | 31.8069051 | 56.0583684 |
| 8.8695206 | 6.45962238 | 19.4270142 | 5.04802073 | 7.48938469 | 10.2236481 | 5.15736989 |
| 11.8260275 | 4.72936639 | 0 | 4.03841658 | 5.99150775 | 9.08768716 | 3.81196905 |
| 8.8695206 | 5.88287038 | 38.8540285 | 14.134458 | 4.49363081 | 2.27192179 | 9.86627284 |
| 5.91301374 | 3.11446079 | 9.71350712 | 12.1152497 | 11.9830155 | 10.2236481 | 5.67983388 |
| 2.95650687 | 14.4764752 | 0 | 9.59123938 | 14.9787694 | 24.9911397 | 13.5661251 |
| 0 | 16.4951072 | 9.71350712 | 3.02881244 | 5.24256928 | 9.08768716 | 9.19357241 |
| 8.8695206 | 13.3806464 | 0 | 7.06722902 | 9.7362001 | 10.2236481 | 6.50277073 |
| 11.8260275 | 10.4968864 | 19.4270142 | 9.08643731 | 14.9787694 | 17.0394134 | 13.0055415 |
| 14.7825343 | 40.9493919 | 29.1405214 | 14.134458 | 9.7362001 | 7.95172627 | 16.3690436 |
| 8.8695206 | 13.1499456 | 9.71350712 | 17.1632705 | 9.7362001 | 14.7674916 | 13.4540084 |
| 47.3041099 | 15.4569536 | 29.1405214 | 32.3073327 | 34.4511696 | 21.583257 | 29.5988185 |
| 2.95650687 | 17.7639616 | 0 | 14.134458 | 8.98726163 | 5.67980448 | 6.27853726 |
| 2.95650687 | 5.99822078 | 9.71350712 | 9.08643731 | 11.6085463 | 7.38374582 | 10.5389733 |
| 2.95650687 | 15.3416032 | 19.4270142 | 6.05762487 | 11.9830155 | 10.2236481 | 12.5570745 |
| 14.7825343 | 3.22981119 | 19.4270142 | 13.1248539 | 12.731954 | 13.6315307 | 9.86627284 |
| 0 | 8.18987838 | 0 | 5.04802073 | 6.74044622 | 10.2236481 | 6.72700421 |
| 5.91301374 | 7.38242558 | 0 | 5.04802073 | 8.23832316 | 7.95172627 | 5.60583684 |
| 14.7825343 | 8.30522878 | 0 | 13.1248539 | 8.98726163 | 14.7674916 | 7.84817157 |
| 8.8695206 | 17.30256 | 0 | 8.07683317 | 11.234077 | 6.81576537 | 8.96933894 |
| 5.91301374 | 9.34338237 | 0 | 6.05762487 | 7.48938469 | 5.67980448 | 6.50277073 |
| 44.347603 | 36.5660767 | 38.8540285 | 33.3169368 | 31.4554157 | 34.0788269 | 40.1377918 |
| 5.91301374 | 2.42235839 | 0 | 11.1056456 | 10.4851386 | 11.359609 | 6.05430379 |
| 17.7390412 | 19.4942175 | 9.71350712 | 16.1536663 | 10.4851386 | 10.7916285 | 11.2116737 |
| 86.7143464 | 62.0965807 | 67.9945498 | 77.7395192 | 63.1879386 | 54.526123 | 73.5104596 |
| 5.91301374 | 9.80478397 | 9.71350712 | 7.06722902 | 9.7362001 | 5.67980448 | 11.8843741 |
| 8.8695206 | 17.5332608 | 0 | 20.1920829 | 9.7362001 | 21.583257 | 17.2659775 |
| 2.95650687 | 3.11446079 | 9.71350712 | 5.04802073 | 2.99575388 | 9.08768716 | 3.81196905 |
| 2.95650687 | 17.1872096 | 9.71350712 | 5.04802073 | 11.9830155 | 6.81576537 | 9.41780589 |
| 44.347603 | 47.8704159 | 58.2810427 | 39.3745617 | 38.9448004 | 23.8551788 | 45.9678621 |
| 5.91301374 | 11.0736384 | 0 | 7.06722902 | 7.48938469 | 4.54384358 | 5.38160336 |
| 0 | 8.42057918 | 19.4270142 | 5.04802073 | 8.23832316 | 9.08768716 | 6.72700421 |
| 14.7825343 | 10.381536 | 9.71350712 | 19.1824788 | 15.7277078 | 17.0394134 | 16.593277 |
| 14.7825343 | 20.8784223 | 0 | 23.7256974 | 11.9830155 | 13.6315307 | 18.1629114 |

| | | | | | | |
|---|---|---|---|---|---|---|
| 11.8260275 | 16.9565088 | 0 | 12.1152497 | 10.4851386 | 13.6315307 | 10.7632067 |
| 35.4780824 | 20.8207471 | 38.8540285 | 52.4994156 | 50.5533467 | 77.8133213 | 25.562616 |
| 44.347603 | 79.1303742 | 19.4270142 | 27.2593119 | 18.7234617 | 20.4472961 | 28.4776511 |
| 5.91301374 | 11.53504 | 0 | 14.134458 | 11.9830155 | 14.7674916 | 10.5389733 |
| 2.95650687 | 6.57497278 | 0 | 4.03841658 | 14.9787694 | 5.67980448 | 7.6239381 |
| 115.303768 | 99.2013437 | 48.5675356 | 108.027644 | 127.31954 | 99.3965784 | 129.719064 |
| 14.7825343 | 11.5927152 | 8.06221091 | 9.08643731 | 14.4769806 | 15.3354721 | 17.1538607 |
| 2.95650687 | 1.153504 | 0 | 3.02881244 | 14.2298309 | 9.08768716 | 7.96028831 |
| 20.6955481 | 24.6849855 | 9.71350712 | 21.2016871 | 18.7234617 | 12.4955698 | 16.1448101 |
| 56.1736305 | 92.9147469 | 48.5675356 | 47.4513948 | 44.9363081 | 38.6226704 | 52.2463993 |
| 5.91301374 | 7.67080158 | 0 | 6.05762487 | 12.731954 | 9.08768716 | 9.19357241 |
| 38.4345893 | 35.0665215 | 48.5675356 | 30.2881244 | 33.7022311 | 34.0788269 | 35.4288888 |
| 14.7825343 | 6.22892158 | 0 | 8.07683317 | 11.9830155 | 18.1753743 | 3.58773558 |
| 17.7390412 | 10.0354848 | 0 | 15.1440622 | 21.7192156 | 17.0394134 | 23.7687482 |
| 11.8260275 | 1.96095679 | 0 | 3.02881244 | 7.48938469 | 5.67980448 | 5.60583684 |
| 14.7825343 | 16.4951072 | 29.1405214 | 22.2112912 | 13.4808924 | 15.9034525 | 21.30218 |
| 14.7825343 | 14.3034496 | 9.71350712 | 14.134458 | 11.9830155 | 11.359609 | 15.9205766 |
| 35.4780824 | 15.6876544 | 9.71350712 | 27.996323 | 19.6596348 | 30.0802445 | 21.9233067 |
| 38.4345893 | 16.3797568 | 29.1405214 | 21.2016871 | 31.4554157 | 32.942866 | 31.1684528 |
| 23.6520549 | 14.1880992 | 19.4270142 | 29.2785202 | 23.2170925 | 22.7192179 | 33.8592545 |
| 14.7825343 | 8.88198078 | 0 | 4.03841658 | 8.23832316 | 10.2236481 | 8.29663852 |
| 2.95650687 | 6.11357118 | 0 | 6.05762487 | 11.234077 | 6.81576537 | 4.70890294 |
| 14.7825343 | 7.38242558 | 9.71350712 | 5.04802073 | 11.234077 | 11.359609 | 4.03620252 |
| 17.7390412 | 20.9937727 | 19.4270142 | 14.134458 | 20.9702771 | 18.1753743 | 25.3383825 |
| 2.95650687 | 9.57408317 | 0 | 11.1056456 | 8.98726163 | 5.67980448 | 8.52087199 |
| 20.6955481 | 19.3788671 | 19.4270142 | 10.0960415 | 14.9787694 | 11.359609 | 11.8843741 |
| 29.5650687 | 23.0573914 | 9.71350712 | 16.6584684 | 22.8426233 | 23.8551788 | 26.1231997 |
| 0 | 8.65127998 | 0 | 5.04802073 | 7.48938469 | 6.81576537 | 6.95123768 |
| 32.5215755 | 16.149056 | 19.4270142 | 21.2016871 | 17.9745233 | 17.0394134 | 22.8718143 |
| 5.91301374 | 14.7648512 | 0 | 6.05762487 | 8.23832316 | 9.08768716 | 7.84817157 |
| 17.7390412 | 1.8456064 | 0 | 6.05762487 | 10.4851386 | 9.08768716 | 8.07240505 |
| 11.8260275 | 7.95917758 | 19.4270142 | 6.05762487 | 5.99150775 | 10.2236481 | 6.72700421 |
| 59.1301374 | 87.0895518 | 29.1405214 | 42.4033741 | 38.9448004 | 44.3024749 | 53.8160336 |
| 2.95650687 | 7.26707518 | 0 | 11.1056456 | 12.731954 | 3.40788269 | 10.0905063 |
| 0 | 1.96095679 | 0 | 6.05762487 | 13.4808924 | 5.67980448 | 3.58773558 |
| 2.95650687 | 7.84382718 | 9.71350712 | 11.1056456 | 12.731954 | 6.81576537 | 8.52087199 |
| 41.3910961 | 37.3735295 | 67.9945498 | 45.4321866 | 29.9575388 | 35.2147878 | 44.3982278 |
| 8.8695206 | 6.22892158 | 0 | 7.06722902 | 11.234077 | 10.2236481 | 8.18452178 |
| 14.7825343 | 14.0727488 | 9.71350712 | 15.1440622 | 10.4851386 | 7.95172627 | 10.3147398 |
| 0 | 8.65127998 | 9.71350712 | 5.04802073 | 2.24681541 | 12.4955698 | 2.91503516 |
| 28.825942 | 26.0634228 | 19.4270142 | 30.0357233 | 30.8937118 | 25.5591201 | 28.6727343 |
| 17.7390412 | 5.76751998 | 9.71350712 | 7.06722902 | 11.234077 | 22.7192179 | 12.332841 |
| 11.8260275 | 11.8810912 | 9.71350712 | 5.04802073 | 5.99150775 | 11.359609 | 8.07240505 |
| 18.7146885 | 11.3043392 | 19.4270142 | 21.2016871 | 8.23832316 | 11.359609 | 12.5570745 |
| 11.8260275 | 32.4134623 | 9.71350712 | 16.1536663 | 7.48938469 | 9.08768716 | 11.4359072 |
| 11.8260275 | 11.9964416 | 0 | 16.1536663 | 10.4851386 | 8.51970672 | 8.40875526 |

| | | | | | | |
|---|---|---|---|---|---|---|
| 235.042296 | 226.778886 | 108.499875 | 203.940037 | 246.153607 | 238.744901 | 260.895646 |
| 5.91301374 | 10.6122368 | 0 | 6.05762487 | 5.99150775 | 9.08768716 | 8.07240505 |
| 13.8068871 | 15.72572 | 32.3459787 | 22.0396585 | 22.4681541 | 19.6862023 | 20.5173628 |
| 14.7825343 | 7.95917758 | 9.71350712 | 8.07683317 | 20.2213387 | 12.4955698 | 13.6782419 |
| 2.95650687 | 4.15261439 | 0 | 11.1056456 | 7.48938469 | 10.2236481 | 3.3635021 |
| 26.6085618 | 24.3389343 | 19.4270142 | 11.1056456 | 14.9787694 | 12.4955698 | 10.7632067 |
| 11.8260275 | 9.11268158 | 19.4270142 | 9.08643731 | 5.24256928 | 9.08768716 | 9.64203936 |
| 14.7825343 | 22.1472767 | 9.71350712 | 12.1152497 | 10.4851386 | 7.95172627 | 13.2297749 |
| 5.91301374 | 10.1508352 | 0 | 12.1152497 | 8.98726163 | 13.6315307 | 7.39970463 |
| 59.1301374 | 95.2794301 | 19.4270142 | 62.595457 | 49.429939 | 53.3901621 | 74.4455132 |
| 50.2606167 | 189.982108 | 9.71350712 | 59.5666446 | 52.4256928 | 53.3901621 | 63.0096061 |
| 0 | 2.07630719 | 0 | 4.03841658 | 8.23832316 | 3.40788269 | 4.70890294 |
| 12.8312398 | 19.2808193 | 32.2488436 | 17.7589369 | 7.86385392 | 16.0284082 | 17.1112564 |
| 5.91301374 | 11.0736384 | 0 | 11.1056456 | 8.98726163 | 10.2236481 | 8.52087199 |
| 44.347603 | 47.2936639 | 38.8540285 | 48.460999 | 50.1788774 | 39.7586313 | 54.7129675 |
| 79.8256854 | 82.821587 | 38.8540285 | 64.6146653 | 69.6512776 | 80.6532236 | 78.0332488 |
| 23.6520549 | 12.4578432 | 0 | 13.1248539 | 2.99575388 | 6.81576537 | 14.5751758 |
| 11.8260275 | 5.65216958 | 9.71350712 | 4.03841658 | 5.24256928 | 9.08768716 | 7.84817157 |
| 23.6520549 | 6.80567358 | 19.4270142 | 4.03841658 | 14.2298309 | 6.81576537 | 8.29663852 |
| 13.3042809 | 4.90239199 | 0 | 15.6488643 | 4.86810005 | 3.97586313 | 10.8753235 |
| 14.7825343 | 22.8393791 | 19.4270142 | 15.1440622 | 15.7277078 | 21.583257 | 18.6113783 |
| 5.91301374 | 12.4578432 | 0 | 20.1920829 | 23.2170925 | 19.3113352 | 18.3871448 |
| 47.3041099 | 44.5252543 | 9.71350712 | 33.3169368 | 35.9490465 | 24.9911397 | 36.3258227 |
| 20.6955481 | 29.2990015 | 38.8540285 | 30.2881244 | 28.4596618 | 18.1753743 | 26.0110829 |
| 14.7825343 | 23.1854303 | 9.71350712 | 17.1632705 | 14.2298309 | 14.7674916 | 14.7994093 |
| 17.7390412 | 9.45873277 | 0 | 7.06722902 | 8.23832316 | 7.95172627 | 9.19357241 |
| 11.8260275 | 15.6876544 | 19.4270142 | 10.0960415 | 18.7234617 | 22.7192179 | 13.6782419 |
| 11.8260275 | 23.7621823 | 0 | 14.134458 | 10.4851386 | 10.2236481 | 15.2613302 |
| 8.8695206 | 3.92191359 | 0 | 4.03841658 | 8.98726163 | 5.67980448 | 5.83007031 |
| 35.4780824 | 25.2617375 | 38.8540285 | 25.2401036 | 27.7781278 | 29.5349833 | 33.1865541 |
| 14.7825343 | 21.9165759 | 9.71350712 | 12.1152497 | 16.3493268 | 10.7916285 | 16.9677469 |
| 35.4780824 | 10.381536 | 19.4270142 | 36.3457492 | 18.7234617 | 30.6709442 | 22.6475808 |
| 2.95650687 | 6.45962238 | 0 | 0 | 8.23832316 | 4.54384358 | 4.48466947 |
| 20.6955481 | 20.7630719 | 0 | 14.134458 | 10.4851386 | 13.6315307 | 14.1267088 |
| 5.91301374 | 12.111792 | 9.71350712 | 6.05762487 | 8.98726163 | 4.54384358 | 9.64203936 |
| 2.95650687 | 6.22892158 | 0 | 4.03841658 | 7.48938469 | 7.95172627 | 5.15736989 |
| 72.4344183 | 34.3167439 | 9.71350712 | 49.4706031 | 56.1703852 | 48.8463185 | 58.3007031 |
| 5.91301374 | 17.30256 | 0 | 10.0960415 | 15.7277078 | 10.2236481 | 12.5570745 |
| 20.6955481 | 14.6495008 | 0 | 9.08643731 | 8.98726163 | 11.359609 | 7.39970463 |
| 17.7390412 | 6.69032318 | 0 | 6.05762487 | 6.74044622 | 12.4955698 | 8.29663852 |
| 20.6955481 | 23.4242057 | 9.71350712 | 16.1536663 | 19.4724002 | 22.7192179 | 17.7144444 |
| 0 | 5.72945435 | 21.0783104 | 8.41000253 | 8.98726163 | 9.46255426 | 5.86819 |
| 8.8695206 | 18.8021151 | 0 | 14.134458 | 8.98726163 | 11.359609 | 18.3871448 |
| 223.216269 | 227.990065 | 126.275593 | 191.319986 | 200.34104 | 182.889704 | 199.343558 |
| 5.91301374 | 6.45962238 | 0 | 13.1248539 | 5.99150775 | 6.81576537 | 6.05430379 |
| 2.95650687 | 7.49777598 | 0 | 6.05762487 | 5.99150775 | 10.2236481 | 6.27853726 |

| | | | | | | |
|---|---|---|---|---|---|---|
| 5.91301374 | 1.8456064 | 9.71350712 | 1.00960415 | 2.24681541 | 9.08768716 | 3.02715189 |
| 17.7390412 | 30.4525055 | 0 | 25.2401036 | 16.4766463 | 21.583257 | 23.7687482 |
| 5.91301374 | 15.9183552 | 9.71350712 | 16.1536663 | 11.234077 | 14.7674916 | 11.6601406 |
| 14.7825343 | 22.6086783 | 0 | 11.1056456 | 8.98726163 | 5.67980448 | 10.7632067 |
| 14.7825343 | 21.3398239 | 0 | 8.07683317 | 11.234077 | 10.7916285 | 13.6782419 |
| 70.9561648 | 61.7124638 | 19.4270142 | 68.6530819 | 47.932062 | 73.8374582 | 57.4037692 |
| 407.997948 | 124.463081 | 388.540285 | 425.043345 | 364.733034 | 289.670028 | 380.075738 |
| 14.7825343 | 15.9183552 | 9.71350712 | 15.1440622 | 8.23832316 | 21.583257 | 11.4359072 |
| 4.4347603 | 3.17213599 | 0 | 6.39079424 | 14.2298309 | 7.95172627 | 7.92216862 |
| 5.91301374 | 10.6122368 | 19.4270142 | 14.134458 | 7.48938469 | 13.6315307 | 10.7632067 |
| 29.5650687 | 33.7976671 | 48.5675356 | 44.9273845 | 33.7022311 | 44.3024749 | 27.5807172 |
| 8.8695206 | 10.958288 | 0 | 8.07683317 | 9.7362001 | 3.40788269 | 8.52087199 |
| 44.347603 | 26.5305919 | 38.8540285 | 50.4802073 | 37.4469235 | 40.8945922 | 43.7255273 |
| 14.7825343 | 10.4968864 | 0 | 8.07683317 | 7.48938469 | 7.95172627 | 8.74510547 |
| 5.91301374 | 7.84382718 | 0 | 13.1248539 | 8.98726163 | 10.2236481 | 7.6239381 |
| 11.8260275 | 10.2661856 | 9.71350712 | 5.04802073 | 3.74469235 | 10.2236481 | 5.60583684 |
| 8.8695206 | 5.30611839 | 0 | 7.06722902 | 8.23832316 | 5.67980448 | 10.0905063 |
| 0 | 4.61401599 | 9.71350712 | 4.03841658 | 9.7362001 | 12.4955698 | 5.60583684 |
| 5.91301374 | 8.53592958 | 0 | 0 | 4.49363081 | 10.2236481 | 6.72700421 |
| 5.91301374 | 7.49777598 | 0 | 4.03841658 | 12.731954 | 12.4955698 | 10.3147398 |
| 10.347774 | 13.3229712 | 0 | 11.6104477 | 11.9830155 | 15.3354721 | 11.0995569 |
| 26.6085618 | 10.6122368 | 0 | 18.1728746 | 20.2213387 | 29.5349833 | 23.0960478 |
| 5.91301374 | 16.2644064 | 9.71350712 | 3.02881244 | 5.99150775 | 9.08768716 | 5.60583684 |
| 5.91301374 | 5.53681919 | 0 | 13.1248539 | 9.7362001 | 9.08768716 | 10.5389733 |
| 38.4345893 | 34.2590687 | 9.71350712 | 35.3361451 | 21.7192156 | 22.7192179 | 29.823052 |
| 17.7390412 | 20.3016703 | 9.71350712 | 18.1728746 | 11.9830155 | 10.2236481 | 22.1991139 |
| 26.6085618 | 8.65127998 | 19.4270142 | 11.1056456 | 18.7234617 | 21.583257 | 19.5083122 |
| 14.7825343 | 11.4196896 | 9.71350712 | 13.1248539 | 14.9787694 | 19.3113352 | 19.0598453 |
| 8.8695206 | 5.42146879 | 9.71350712 | 3.02881244 | 5.99150775 | 12.4955698 | 11.4359072 |
| 17.7390412 | 4.40292476 | 9.71350712 | 5.04802073 | 5.99150775 | 9.08768716 | 5.83007031 |
| 23.6520549 | 11.0736384 | 19.4270142 | 7.06722902 | 14.9787694 | 19.3113352 | 20.1810126 |
| 11.8260275 | 6.45962238 | 9.71350712 | 11.1056456 | 9.7362001 | 7.95172627 | 11.6601406 |
| 26.6085618 | 21.5128495 | 0 | 11.1056456 | 21.7192156 | 12.4955698 | 18.7974921 |
| 8.8695206 | 5.65216958 | 9.71350712 | 14.134458 | 13.1064232 | 21.0152766 | 6.95123768 |
| 2.95650687 | 5.88287038 | 0 | 4.03841658 | 6.74044622 | 11.359609 | 8.52087199 |
| 11.8260275 | 27.4533951 | 0 | 27.764114 | 23.966031 | 32.942866 | 27.244367 |
| 49.7580106 | 49.3319054 | 100.340529 | 66.4622409 | 47.8047425 | 45.4384358 | 62.1866692 |
| 0 | 5.76751998 | 0 | 4.03841658 | 8.23832316 | 4.54384358 | 8.07240505 |
| 41.3910961 | 79.361075 | 38.8540285 | 40.3841658 | 41.9405543 | 44.3024749 | 41.0347257 |
| 8.8695206 | 21.1091231 | 9.71350712 | 6.05762487 | 11.234077 | 9.08768716 | 8.52087199 |
| 11.8260275 | 16.149056 | 0 | 7.06722902 | 10.4851386 | 4.54384358 | 12.332841 |
| 26.6085618 | 11.1889888 | 29.1405214 | 24.2304995 | 26.2128464 | 21.583257 | 28.2534177 |
| 79.8256854 | 26.7612927 | 38.8540285 | 46.4417907 | 53.1746313 | 51.1182403 | 58.7491701 |
| 5.91301374 | 4.61401599 | 0 | 8.07683317 | 11.234077 | 5.67980448 | 4.93313642 |
| 26.6085618 | 14.995552 | 19.4270142 | 18.1728746 | 14.2298309 | 27.2630615 | 21.5264135 |
| 8.8695206 | 4.03726399 | 14.5702607 | 11.1056456 | 14.9787694 | 15.3354721 | 11.9964908 |

| | | | | | | |
|---|---|---|---|---|---|---|
| 2.95650687 | 7.26707518 | 9.71350712 | 1.00960415 | 7.86385392 | 5.67980448 | 3.92408579 |
| 14.7825343 | 10.381536 | 9.71350712 | 3.02881244 | 7.48938469 | 11.359609 | 8.07240505 |
| 20.6955481 | 9.80478397 | 0 | 17.1632705 | 17.9745233 | 20.4472961 | 19.7325457 |
| 8.8695206 | 23.8775327 | 9.71350712 | 9.08643731 | 13.4808924 | 15.9034525 | 12.332841 |
| 73.9126717 | 88.9351582 | 9.71350712 | 81.7779358 | 67.4044622 | 52.2542012 | 81.3967509 |
| 5.91301374 | 12.5731936 | 0 | 16.1536663 | 2.99575388 | 10.2236481 | 11.4359072 |
| 8.8695206 | 11.0736384 | 19.4270142 | 5.04802073 | 7.48938469 | 5.67980448 | 6.95123768 |
| 2.95650687 | 7.49777598 | 9.71350712 | 7.06722902 | 6.74044622 | 9.08768716 | 4.93313642 |
| 14.7825343 | 25.3770879 | 9.71350712 | 13.1248539 | 9.7362001 | 11.359609 | 7.77417453 |
| 11.8260275 | 16.2644064 | 0 | 6.05762487 | 8.23832316 | 6.81576537 | 10.7632067 |
| 94.6082198 | 75.5545118 | 135.9891 | 102.979623 | 72.6470315 | 71.5655364 | 87.4510547 |
| 50.2606167 | 65.6343774 | 67.9945498 | 49.4706031 | 35.200108 | 23.8551788 | 43.2770604 |
| 2.95650687 | 10.7275872 | 0 | 13.1248539 | 9.7362001 | 13.6315307 | 13.9024754 |
| 84.2604457 | 102.277739 | 97.1350712 | 145.382997 | 122.32412 | 191.216298 | 88.0116384 |
| 26.6085618 | 29.0106255 | 67.9945498 | 24.7353016 | 33.3277619 | 24.9911397 | 28.5897679 |
| 2.95650687 | 6.45962238 | 0 | 12.1152497 | 12.731954 | 17.0394134 | 12.781308 |
| 8.8695206 | 5.76751998 | 0 | 11.1056456 | 8.98726163 | 10.2236481 | 7.6239381 |
| 0 | 3.92191359 | 0 | 8.07683317 | 10.4851386 | 4.54384358 | 3.81196905 |
| 0 | 5.76751998 | 0 | 5.04802073 | 7.48938469 | 4.54384358 | 6.50277073 |
| 118.260275 | 176.947513 | 67.9945498 | 91.8739773 | 110.842893 | 87.4689889 | 101.801997 |
| 2.95650687 | 0.6921024 | 0 | 9.08643731 | 13.4808924 | 17.0394134 | 6.50277073 |
| 35.4780824 | 65.519027 | 29.1405214 | 37.3553534 | 43.4384312 | 34.0788269 | 47.0890294 |
| 2.95650687 | 19.3788671 | 0 | 8.07683317 | 11.234077 | 15.9034525 | 15.4721097 |
| 1262.63539 | 13.130336 | 8679.60142 | 372.190568 | 190.941863 | 268.166289 | 50.7485197 |
| 56.1736305 | 33.4516159 | 9.71350712 | 24.2304995 | 29.9575388 | 42.0305531 | 39.0166244 |
| 59.1301374 | 58.9440542 | 9.71350712 | 41.39377 | 39.6937389 | 35.2147878 | 49.1071307 |
| 5.91301374 | 5.07541759 | 0 | 1.51440622 | 7.11491546 | 9.08768716 | 5.26948663 |
| 26.6085618 | 20.5323711 | 9.71350712 | 13.1248539 | 19.4724002 | 30.1029637 | 18.1629114 |
| 14.7825343 | 7.15172478 | 0 | 9.08643731 | 3.74469235 | 11.359609 | 5.60583684 |
| 11.8260275 | 4.61401599 | 9.71350712 | 6.05762487 | 12.731954 | 9.08768716 | 5.60583684 |
| 5.91301374 | 8.88198078 | 0 | 3.02881244 | 7.48938469 | 6.81576537 | 6.95123768 |
| 14.7825343 | 12.688544 | 9.71350712 | 8.07683317 | 12.731954 | 13.6315307 | 12.5570745 |
| 67.999658 | 89.742611 | 29.1405214 | 42.4033741 | 50.9278159 | 22.7192179 | 50.4525315 |
| 28.0868152 | 22.0319263 | 24.2837678 | 32.3073327 | 31.3280962 | 28.3990224 | 34.4198382 |
| 5.91301374 | 9.68943357 | 19.4270142 | 10.0960415 | 7.48938469 | 11.9275894 | 12.781308 |
| 8.8695206 | 3.11446079 | 19.4270142 | 15.1440622 | 9.7362001 | 6.81576537 | 10.3147398 |
| 29.5650687 | 7.15172478 | 0 | 10.0960415 | 9.7362001 | 6.81576537 | 12.781308 |
| 8.8695206 | 13.1499456 | 9.71350712 | 10.0960415 | 7.48938469 | 7.95172627 | 11.6601406 |
| 357.737331 | 304.525055 | 456.534835 | 360.42868 | 292.086003 | 256.727162 | 397.117482 |
| 8.8695206 | 15.8030048 | 9.71350712 | 15.1440622 | 17.9745233 | 15.9034525 | 16.1448101 |
| 6.14953428 | 17.0730127 | 8.45075119 | 15.9214574 | 14.5593638 | 19.1295815 | 15.0236427 |
| 5.91301374 | 6.69032318 | 0 | 10.0960415 | 9.7362001 | 12.4955698 | 9.41780589 |
| 14.7825343 | 18.2253632 | 19.4270142 | 16.1536663 | 19.4724002 | 12.4955698 | 19.9567791 |
| 17.7390412 | 18.456064 | 19.4270142 | 22.2112912 | 19.4724002 | 26.1271006 | 28.0673039 |
| 35.4780824 | 106.122368 | 19.4270142 | 27.2593119 | 17.2255848 | 23.8551788 | 26.6837834 |
| 11.8260275 | 3.11446079 | 9.71350712 | 15.1440622 | 14.2298309 | 12.4955698 | 11.6601406 |

| | | | | | | |
|---|---|---|---|---|---|---|
| 11.8260275 | 8.76663038 | 0 | 9.08643731 | 5.24256928 | 9.08768716 | 8.29663852 |
| 5.91301374 | 11.4196896 | 0 | 8.07683317 | 8.98726163 | 12.4955698 | 6.27853726 |
| 85.7386992 | 90.896115 | 97.1350712 | 66.6338736 | 86.8768624 | 85.1970672 | 97.541561 |
| 5.91301374 | 9.34338237 | 9.71350712 | 3.02881244 | 9.7362001 | 6.81576537 | 15.6963431 |
| 17.7390412 | 24.5696351 | 29.1405214 | 28.2689161 | 25.4639079 | 17.0394134 | 38.1196905 |
| 2.95650687 | 5.42146879 | 0 | 9.08643731 | 9.7362001 | 3.40788269 | 3.58773558 |
| 29.5650687 | 64.4808734 | 19.4270142 | 26.2497078 | 16.4766463 | 17.0394134 | 30.0472855 |
| 2.95650687 | 3.22981119 | 9.71350712 | 12.1152497 | 7.48938469 | 5.67980448 | 16.1448101 |
| 2.95650687 | 8.53592958 | 0 | 4.03841658 | 7.48938469 | 9.08768716 | 9.19357241 |
| 5.91301374 | 10.7275872 | 0 | 4.03841658 | 7.48938469 | 4.54384358 | 9.86627284 |
| 298.607194 | 312.368882 | 485.675356 | 334.178972 | 282.349803 | 237.415827 | 311.460295 |
| 35.4780824 | 22.0319263 | 48.5675356 | 27.2593119 | 25.4639079 | 20.4472961 | 24.2172151 |
| 29.5650687 | 23.7621823 | 9.71350712 | 21.2016871 | 19.4724002 | 18.1753743 | 26.4595499 |
| 2.95650687 | 7.84382718 | 9.71350712 | 3.02881244 | 8.23832316 | 2.27192179 | 4.03620252 |
| 2.95650687 | 7.72847678 | 0 | 4.03841658 | 7.48938469 | 4.54384358 | 4.70890294 |
| 487.823633 | 271.18879 | 233.124171 | 359.419076 | 315.303095 | 298.757715 | 386.354275 |
| 47.3041099 | 34.9511711 | 9.71350712 | 26.2497078 | 24.7149695 | 22.7192179 | 35.6531223 |
| 5.91301374 | 13.6113472 | 0 | 13.1248539 | 9.7362001 | 14.7674916 | 14.5751758 |
| 2.95650687 | 2.30700799 | 0 | 19.1824788 | 9.7362001 | 11.359609 | 7.39970463 |
| 5.91301374 | 4.84471679 | 0 | 23.2208954 | 9.7362001 | 18.1753743 | 13.2297749 |
| 8.8695206 | 13.6113472 | 9.71350712 | 17.1632705 | 14.9787694 | 14.7674916 | 16.8175105 |
| 0 | 14.3034496 | 0 | 13.1248539 | 7.48938469 | 11.359609 | 8.52087199 |
| 11.8260275 | 7.95917758 | 0 | 8.07683317 | 10.4851386 | 3.40788269 | 8.07240505 |
| 5.91301374 | 4.72936639 | 9.71350712 | 10.0960415 | 13.4808924 | 13.6315307 | 7.6239381 |
| 47.3041099 | 29.1836511 | 19.4270142 | 42.4033741 | 29.2086003 | 27.2630615 | 28.9261181 |
| 17.7390412 | 12.4578432 | 0 | 10.0960415 | 7.48938469 | 1.1359609 | 11.4359072 |
| 13.3042809 | 11.6503904 | 9.71350712 | 21.2016871 | 20.5958079 | 20.4472961 | 13.2297749 |
| 11.8260275 | 29.0683007 | 0 | 20.1920829 | 17.2255848 | 6.81576537 | 17.041744 |
| 8.8695206 | 5.19076799 | 9.71350712 | 2.01920829 | 8.23832316 | 9.08768716 | 5.15736989 |
| 0 | 3.92191359 | 0 | 10.0960415 | 8.23832316 | 9.08768716 | 5.60583684 |
| 2.95650687 | 7.15172478 | 19.4270142 | 6.05762487 | 8.23832316 | 6.81576537 | 7.39970463 |
| 5.91301374 | 3.11446079 | 0 | 16.1536663 | 11.234077 | 12.4955698 | 13.0055415 |
| 23.6520549 | 17.6486112 | 48.5675356 | 29.2785202 | 18.7234617 | 18.1753743 | 19.2840787 |
| 23.6520549 | 19.3788671 | 9.71350712 | 15.1440622 | 25.4639079 | 19.3113352 | 24.8899156 |
| 5.91301374 | 5.42146879 | 19.4270142 | 7.06722902 | 7.48938469 | 9.08768716 | 9.64203936 |
| 8.8695206 | 8.88198078 | 0 | 4.03841658 | 5.99150775 | 12.4955698 | 6.72700421 |
| 104.955994 | 98.4515661 | 53.4242892 | 90.8643731 | 95.4896548 | 81.221204 | 95.5234597 |
| 8.8695206 | 9.22803198 | 9.71350712 | 12.1152497 | 7.36206515 | 12.4955698 | 6.83912094 |
| 0 | 3.22981119 | 0 | 5.04802073 | 10.4851386 | 4.54384358 | 5.60583684 |
| 8.8695206 | 7.61312638 | 9.71350712 | 9.59123938 | 11.6085463 | 20.4472961 | 13.3418917 |
| 20.6955481 | 22.7240287 | 9.71350712 | 13.1248539 | 24.7149695 | 24.9911397 | 24.2172151 |
| 8.8695206 | 20.7630719 | 0 | 15.1440622 | 11.9830155 | 11.359609 | 12.1086076 |
| 25.1303084 | 38.5847087 | 19.4270142 | 16.1536663 | 20.2213387 | 31.8069051 | 23.6566315 |
| 24.6277022 | 13.6494128 | 19.4270142 | 12.7916845 | 9.4890504 | 6.44089828 | 8.74510547 |
| 29.5650687 | 40.6033407 | 29.1405214 | 45.4321866 | 39.6937389 | 36.3507487 | 47.0890294 |
| 2.95650687 | 4.96006719 | 0 | 9.08643731 | 8.23832316 | 5.67980448 | 5.15736989 |

| | | | | | | |
|---|---|---|---|---|---|---|
| 14.7825343 | 19.9556191 | 19.4270142 | 20.1920829 | 14.2298309 | 15.9034525 | 18.3871448 |
| 11.8260275 | 16.6104576 | 0 | 10.0960415 | 16.4766463 | 17.0394134 | 12.332841 |
| 38.4345893 | 47.5243647 | 38.8540285 | 26.2497078 | 29.9575388 | 20.4472961 | 26.2353164 |
| 26.6085618 | 10.4968864 | 19.4270142 | 17.1632705 | 7.48938469 | 13.6315307 | 13.6782419 |
| 26.6085618 | 32.1827615 | 77.708057 | 42.4033741 | 45.6852466 | 40.8945922 | 43.9497608 |
| 2.95650687 | 6.22892158 | 9.71350712 | 1.00960415 | 8.23832316 | 5.67980448 | 7.39970463 |
| 17.7390412 | 15.6876544 | 29.1405214 | 9.59123938 | 13.4808924 | 15.9034525 | 19.171962 |
| 5.91301374 | 6.57497278 | 0 | 16.1536663 | 5.99150775 | 14.7674916 | 12.1086076 |
| 8.8695206 | 5.07541759 | 0 | 1.00960415 | 4.49363081 | 9.08768716 | 4.70890294 |
| 40.88849 | 41.7568447 | 9.71350712 | 26.5828772 | 30.4593275 | 32.942866 | 30.2333992 |
| 5.91301374 | 19.9556191 | 19.4270142 | 6.05762487 | 11.9830155 | 6.81576537 | 12.781308 |
| 11.8260275 | 3.69121279 | 0 | 6.05762487 | 8.98726163 | 12.4955698 | 7.6239381 |
| 8.8695206 | 8.53592958 | 0 | 9.08643731 | 13.4808924 | 15.9034525 | 12.781308 |
| 17.7390412 | 10.4968864 | 19.4270142 | 24.2304995 | 16.4766463 | 21.583257 | 16.593277 |
| 0 | 9.92013437 | 19.4270142 | 11.1056456 | 5.99150775 | 5.67980448 | 8.74510547 |
| 8.8695206 | 19.0328159 | 0 | 11.1056456 | 15.7277078 | 21.583257 | 14.1267088 |
| 14.7825343 | 13.7266976 | 19.4270142 | 8.07683317 | 11.234077 | 12.4955698 | 13.4540084 |
| 0 | 10.958288 | 0 | 4.03841658 | 7.48938469 | 3.40788269 | 5.38160336 |
| 0 | 2.76840959 | 0 | 3.02881244 | 7.48938469 | 4.54384358 | 5.15736989 |
| 56.1736305 | 84.551843 | 48.5675356 | 76.7299151 | 50.9278159 | 61.3418884 | 86.5541208 |
| 5.91301374 | 19.0328159 | 9.71350712 | 13.1248539 | 14.9787694 | 7.95172627 | 13.2297749 |
| 20.6955481 | 11.0736384 | 9.71350712 | 11.1056456 | 18.7234617 | 3.40788269 | 16.8175105 |
| 23.6520549 | 16.8411584 | 19.4270142 | 12.1152497 | 11.234077 | 22.7192179 | 10.5389733 |
| 5.91301374 | 11.4196896 | 0 | 11.1056456 | 7.48938469 | 5.67980448 | 11.6601406 |
| 957.908225 | 528.304831 | 573.09692 | 755.183901 | 688.274453 | 615.690805 | 860.832305 |
| 2.95650687 | 9.51640797 | 0 | 14.134458 | 7.48938469 | 7.95172627 | 8.29663852 |
| 23.6520549 | 33.9130175 | 9.71350712 | 20.1920829 | 21.7192156 | 26.1271006 | 21.0779465 |
| 20.6955481 | 14.0727488 | 9.71350712 | 12.1152497 | 11.9830155 | 9.08768716 | 9.86627284 |
| 23.6520549 | 24.8003359 | 48.5675356 | 36.3457492 | 23.2170925 | 23.8551788 | 29.5988185 |
| 7.39126717 | 8.76663038 | 0 | 11.1056456 | 14.8514498 | 9.65566761 | 16.1066904 |
| 50.2606167 | 56.7523966 | 48.5675356 | 87.8355607 | 90.6215547 | 81.7891845 | 85.6571869 |
| 50.2606167 | 22.7240287 | 19.4270142 | 28.2689161 | 26.9617849 | 12.4955698 | 25.7868495 |
| 17.7390412 | 15.9183552 | 9.71350712 | 7.06722902 | 11.234077 | 18.1753743 | 14.5751758 |
| 17.7390412 | 16.3797568 | 19.4270142 | 7.06722902 | 8.23832316 | 11.359609 | 11.6601406 |
| 2.95650687 | 7.03637438 | 9.71350712 | 2.01920829 | 9.7362001 | 11.359609 | 4.93313642 |
| 32.5215755 | 30.4525055 | 9.71350712 | 21.2016871 | 25.4639079 | 19.3113352 | 27.1322503 |
| 82.7821923 | 153.06998 | 38.8540285 | 54.5186239 | 55.4214467 | 65.8857319 | 62.7853726 |
| 0 | 3.92191359 | 19.4270142 | 8.07683317 | 7.48938469 | 2.27192179 | 5.83007031 |
| 65.0431511 | 34.1437183 | 48.5675356 | 55.528228 | 68.9023391 | 52.2542012 | 66.5973416 |
| 14.7825343 | 18.6867647 | 29.1405214 | 13.1248539 | 9.7362001 | 13.6315307 | 15.0236427 |
| 62.0866442 | 36.5660767 | 67.9945498 | 44.4225824 | 41.1916158 | 42.0305531 | 52.9190998 |
| 5.91301374 | 4.03726399 | 9.71350712 | 17.1632705 | 14.2298309 | 12.4955698 | 13.6782419 |
| 11.8260275 | 7.15172478 | 9.71350712 | 5.04802073 | 6.74044622 | 9.08768716 | 10.5389733 |
| 26.6085618 | 42.3335967 | 9.71350712 | 24.2304995 | 40.4426773 | 31.8069051 | 37.895457 |
| 19.2172946 | 26.0691903 | 0 | 10.6008435 | 7.11491546 | 11.359609 | 9.7541561 |
| 20.6955481 | 12.4578432 | 38.8540285 | 12.1152497 | 22.4681541 | 10.2236481 | 15.6963431 |

| | | | | | | |
|---|---|---|---|---|---|---|
| 14.7825343 | 16.725808 | 9.71350712 | 9.08643731 | 11.234077 | 4.54384358 | 9.64203936 |
| 16.2607878 | 25.3770879 | 9.71350712 | 35.8409472 | 19.4724002 | 28.3990224 | 25.3383825 |
| 2.95650687 | 2.99911039 | 0 | 3.02881244 | 8.23832316 | 4.54384358 | 4.48466947 |
| 35.4780824 | 11.3043392 | 38.8540285 | 28.2689161 | 21.7192156 | 26.1271006 | 24.6656821 |
| 5.91301374 | 5.65216958 | 9.71350712 | 9.08643731 | 11.234077 | 10.2236481 | 9.19357241 |
| 56.1736305 | 30.3371551 | 29.1405214 | 64.6146653 | 59.1661391 | 43.166514 | 57.4037692 |
| 8.8695206 | 3.92191359 | 9.71350712 | 9.08643731 | 5.99150775 | 10.2236481 | 8.96933894 |
| 17.7390412 | 9.80478397 | 0 | 9.08643731 | 11.234077 | 15.9034525 | 13.0055415 |
| 20.6955481 | 36.1046751 | 0 | 30.2881244 | 14.9787694 | 26.1271006 | 27.1322503 |
| 35.4780824 | 14.995552 | 19.4270142 | 26.2497078 | 21.7192156 | 13.6315307 | 21.7506469 |
| 2.95650687 | 13.3806464 | 0 | 10.0960415 | 6.74044622 | 18.1753743 | 9.4312599 |
| 2.95650687 | 5.76751998 | 0 | 8.07683317 | 12.731954 | 11.359609 | 10.0905063 |
| 38.4345893 | 14.4188 | 29.1405214 | 21.2016871 | 22.4681541 | 19.3113352 | 29.2243486 |
| 65.0431511 | 85.8206974 | 38.8540285 | 38.3649575 | 65.9065853 | 42.0305531 | 49.7798311 |
| 2.95650687 | 7.49777598 | 9.71350712 | 2.01920829 | 8.23832316 | 4.54384358 | 6.95123768 |
| 8.8695206 | 7.72847678 | 19.4270142 | 5.04802073 | 11.234077 | 13.6315307 | 12.5570745 |
| 7.39126717 | 10.8429376 | 9.71350712 | 8.07683317 | 14.2298309 | 7.95172627 | 9.64203936 |
| 14.7825343 | 35.6432735 | 0 | 17.1632705 | 14.9787694 | 13.6315307 | 15.0236427 |
| 5.91301374 | 10.7275872 | 9.71350712 | 15.1440622 | 11.1067575 | 13.6315307 | 5.60583684 |
| 44.347603 | 58.2519518 | 29.1405214 | 30.2881244 | 30.7064772 | 28.3990224 | 52.0221659 |
| 62.0866442 | 43.8331519 | 38.8540285 | 58.5570405 | 65.9065853 | 54.526123 | 59.4218705 |
| 65.0431511 | 66.2111294 | 29.1405214 | 76.7299151 | 76.3917238 | 65.8857319 | 69.7366103 |
| 8.8695206 | 5.07541759 | 19.4270142 | 5.04802073 | 12.731954 | 9.08768716 | 8.29663852 |
| 23.6520549 | 37.3735295 | 0 | 12.1152497 | 5.24256928 | 7.95172627 | 10.5389733 |
| 8.8695206 | 20.6477215 | 0 | 5.04802073 | 9.7362001 | 5.67980448 | 11.8843741 |
| 26.6085618 | 22.9547295 | 9.71350712 | 13.1248539 | 14.9787694 | 13.6315307 | 16.8175105 |
| 2.95650687 | 10.0354848 | 9.71350712 | 12.1152497 | 12.731954 | 10.2236481 | 11.6601406 |
| 212.484149 | 290.61149 | 176.494424 | 446.214744 | 359.19089 | 579.851239 | 193.782568 |
| 2.95650687 | 4.84471679 | 9.71350712 | 5.04802073 | 11.9830155 | 7.95172627 | 6.00721476 |
| 0 | 9.92013437 | 38.8540285 | 9.08643731 | 6.74044622 | 10.2236481 | 10.7632067 |
| 32.5215755 | 12.8038944 | 9.71350712 | 16.1536663 | 20.9702771 | 11.359609 | 18.1629114 |
| 7.39126717 | 12.97692 | 0 | 18.6776767 | 11.234077 | 19.6862023 | 9.24963078 |
| 5.91301374 | 7.38242558 | 9.71350712 | 7.06722902 | 7.48938469 | 9.08768716 | 7.6239381 |
| 0 | 5.88287038 | 9.71350712 | 3.02881244 | 5.24256928 | 11.359609 | 5.15736989 |
| 8.8695206 | 24.6849855 | 29.1405214 | 16.1536663 | 24.7149695 | 26.1271006 | 22.6475808 |
| 62.0866442 | 47.4090143 | 29.1405214 | 46.4417907 | 47.1831235 | 27.2630615 | 49.5555977 |
| 2.95650687 | 8.76663038 | 0 | 3.02881244 | 14.9787694 | 10.2236481 | 9.41780589 |
| 11.8260275 | 27.3380447 | 0 | 20.1920829 | 31.4554157 | 27.2630615 | 23.5445147 |
| 8.8695206 | 10.0354848 | 0 | 4.03841658 | 6.74044622 | 15.9034525 | 8.96933894 |
| 5.91301374 | 11.6503904 | 9.71350712 | 8.07683317 | 8.98726163 | 9.08768716 | 9.86627284 |
| 17.7390412 | 8.07452798 | 0 | 6.05762487 | 11.234077 | 10.2236481 | 8.52087199 |
| 2.95650687 | 14.5341504 | 9.71350712 | 6.05762487 | 6.74044622 | 10.2236481 | 7.6239381 |
| 2.95650687 | 6.57497278 | 9.71350712 | 10.0960415 | 7.48938469 | 9.08768716 | 4.03620252 |
| 26.6085618 | 24.5696351 | 19.4270142 | 18.1728746 | 24.7149695 | 28.3990224 | 32.9623206 |
| 17.7390412 | 5.88287038 | 0 | 11.1056456 | 11.234077 | 13.6315307 | 12.8934247 |
| 5.91301374 | 2.74533951 | 9.71350712 | 8.07683317 | 8.23832316 | 5.67980448 | 7.6239381 |

| | | | | | | |
|---|---|---|---|---|---|---|
| 14.7825343 | 35.9893247 | 0 | 30.2881244 | 21.7192156 | 34.0788269 | 28.2534177 |
| 11.8260275 | 25.0310367 | 9.71350712 | 11.1056456 | 14.9787694 | 14.7674916 | 10.7632067 |
| 1019.99487 | 250.771769 | 1291.89645 | 1016.67137 | 972.871071 | 741.782465 | 1050.30959 |
| 8.8695206 | 8.07452798 | 29.1405214 | 10.0960415 | 6.74044622 | 13.6315307 | 11.6601406 |
| 11.8260275 | 18.456064 | 9.71350712 | 19.1824788 | 15.7277078 | 18.7433548 | 15.8084599 |
| 11.8260275 | 11.1889888 | 0 | 16.1536663 | 14.9787694 | 21.583257 | 16.593277 |
| 242.433563 | 106.468419 | 184.556635 | 191.824788 | 207.455956 | 176.073939 | 186.113783 |
| 8.8695206 | 8.30522878 | 9.71350712 | 6.05762487 | 5.24256928 | 9.08768716 | 5.60583684 |
| 2.95650687 | 7.84382718 | 0 | 9.08643731 | 7.48938469 | 2.27192179 | 6.72700421 |
| 32.5215755 | 5.88287038 | 19.4270142 | 15.1440622 | 18.7234617 | 15.9034525 | 24.2172151 |
| 2.95650687 | 24.5696351 | 0 | 25.2401036 | 21.7192156 | 11.359609 | 22.8718143 |
| 11.8260275 | 14.1880992 | 0 | 8.07683317 | 10.4851386 | 5.67980448 | 8.07240505 |
| 5.91301374 | 3.34516159 | 19.4270142 | 10.0960415 | 8.98726163 | 9.08768716 | 9.41780589 |
| 5.91301374 | 6.80567358 | 0 | 6.05762487 | 11.234077 | 7.95172627 | 6.27853726 |
| 29.5650687 | 30.5678559 | 9.71350712 | 27.2593119 | 26.2128464 | 28.3990224 | 25.3383825 |
| 41.3910961 | 14.6495008 | 19.4270142 | 19.6872808 | 30.332008 | 34.6468073 | 22.5354641 |
| 14.7825343 | 10.958288 | 0 | 9.08643731 | 13.4808924 | 7.95172627 | 10.7632067 |
| 2.95650687 | 5.42146879 | 0 | 7.06722902 | 8.98726163 | 3.40788269 | 5.83007031 |
| 5.91301374 | 8.07452798 | 0 | 12.1152497 | 17.2255848 | 21.583257 | 6.95123768 |
| 2.95650687 | 29.2990015 | 0 | 14.134458 | 10.4851386 | 11.359609 | 16.593277 |
| 11.8260275 | 10.1508352 | 0 | 3.02881244 | 9.7362001 | 7.95172627 | 7.39970463 |
| 20.6955481 | 18.9174655 | 9.71350712 | 18.1728746 | 16.4766463 | 21.583257 | 16.8175105 |
| 36.9563358 | 18.6867647 | 24.2837678 | 28.2689161 | 32.5788234 | 52.8221816 | 32.7380871 |
| 14.7825343 | 29.4143519 | 0 | 27.2593119 | 20.2213387 | 26.1271006 | 23.0960478 |
| 147.825343 | 66.672531 | 145.702607 | 133.267747 | 132.562109 | 113.59609 | 129.831181 |
| 8.8695206 | 6.22892158 | 0 | 4.03841658 | 5.24256928 | 12.4955698 | 8.52087199 |
| 17.7390412 | 26.9343183 | 9.71350712 | 13.629656 | 11.234077 | 13.6315307 | 13.4540084 |
| 20.6955481 | 27.2226943 | 19.4270142 | 29.2785202 | 23.966031 | 17.0394134 | 21.5264135 |
| 8.8695206 | 10.8429376 | 9.71350712 | 23.2208954 | 9.7362001 | 15.9034525 | 9.19357241 |
| 17.7390412 | 30.1064543 | 9.71350712 | 16.1536663 | 17.9745233 | 18.1753743 | 16.593277 |
| 29.5650687 | 13.3806464 | 29.1405214 | 27.2593119 | 30.7064772 | 30.6709442 | 27.5807172 |
| 11.8260275 | 12.8038944 | 0 | 5.04802073 | 11.9830155 | 7.95172627 | 15.6963431 |
| 14.7825343 | 9.92013437 | 9.71350712 | 3.02881244 | 9.7362001 | 9.08768716 | 8.07240505 |
| 11.8260275 | 3.57586239 | 0 | 2.01920829 | 8.23832316 | 14.7674916 | 4.93313642 |
| 2.95650687 | 10.7275872 | 0 | 7.06722902 | 14.2298309 | 6.81576537 | 9.41780589 |
| 79.8256854 | 84.4364926 | 38.8540285 | 75.7203109 | 63.6597699 | 60.2059275 | 74.6697467 |
| 26.6085618 | 19.3788671 | 9.71350712 | 13.1248539 | 10.4851386 | 24.9911397 | 23.7687482 |
| 2.95650687 | 7.61312638 | 9.71350712 | 7.06722902 | 11.3089709 | 9.08768716 | 13.0055415 |
| 8.8695206 | 17.0718592 | 9.71350712 | 26.2497078 | 25.4639079 | 14.7674916 | 24.6656821 |
| 85.7386992 | 30.3371551 | 58.2810427 | 82.7875399 | 68.1534007 | 67.0216928 | 75.6787973 |
| 29.5650687 | 23.8775327 | 24.2837678 | 25.2401036 | 15.7277078 | 20.4472961 | 26.4595499 |
| 17.7390412 | 40.2572895 | 19.4270142 | 14.134458 | 16.4766463 | 17.0394134 | 30.9442193 |
| 1806.4257 | 568.56212 | 1447.31256 | 1529.04548 | 1345.84243 | 1342.1378 | 1841.29317 |
| 5.91301374 | 15.9183552 | 19.4270142 | 17.1632705 | 14.9787694 | 13.6315307 | 19.7325457 |
| 8.8695206 | 4.03726399 | 0 | 6.05762487 | 9.7362001 | 2.27192179 | 5.38160336 |
| 14.7825343 | 10.8429376 | 9.71350712 | 10.0960415 | 17.9745233 | 5.67980448 | 17.041744 |

| | | | | | | |
|---|---|---|---|---|---|---|
| 8.8695206 | 15.2262528 | 19.4270142 | 12.1152497 | 10.4851386 | 10.2236481 | 12.781308 |
| 8.8695206 | 2.76840959 | 0 | 3.02881244 | 7.48938469 | 0 | 6.05430379 |
| 103.47774 | 147.225176 | 67.9945498 | 57.2142669 | 84.1282582 | 58.3088728 | 68.4652065 |
| 11.8260275 | 7.03637438 | 0 | 5.04802073 | 5.99150775 | 11.359609 | 7.84817157 |
| 2.95650687 | 9.68943357 | 0 | 2.01920829 | 8.98726163 | 9.08768716 | 4.03620252 |
| 20.6955481 | 12.5731936 | 9.71350712 | 17.1632705 | 26.2128464 | 24.9911397 | 21.7506469 |
| 11.8260275 | 11.4196896 | 0 | 16.1536663 | 21.7192156 | 20.4472961 | 15.6963431 |
| 5.91301374 | 11.9964416 | 0 | 14.134458 | 7.48938469 | 7.95172627 | 12.332841 |
| 20.6955481 | 32.0674111 | 38.8540285 | 22.2112912 | 25.4639079 | 24.9911397 | 26.0110829 |
| 17.7390412 | 30.7985567 | 0 | 20.1920829 | 20.9702771 | 24.9911397 | 30.0472855 |
| 5.91301374 | 4.26796479 | 9.71350712 | 2.01920829 | 8.23832316 | 4.54384358 | 7.17547115 |
| 11.8260275 | 11.6503904 | 9.71350712 | 6.05762487 | 11.9830155 | 3.40788269 | 12.332841 |
| 26.6085618 | 29.1836511 | 9.71350712 | 21.2016871 | 21.3447464 | 20.4472961 | 26.0110829 |
| 5.91301374 | 11.9964416 | 19.4270142 | 8.07683317 | 7.48938469 | 9.08768716 | 6.05430379 |
| 23.6520549 | 23.4161311 | 19.4270142 | 18.1728746 | 19.4724002 | 20.4472961 | 12.781308 |
| 5.91301374 | 7.03637438 | 9.71350712 | 13.1248539 | 15.7277078 | 10.2236481 | 14.5751758 |
| 2.95650687 | 9.57408317 | 9.71350712 | 4.03841658 | 20.9702771 | 12.4955698 | 14.1401628 |
| 14.7825343 | 29.1836511 | 0 | 12.1152497 | 9.7362001 | 7.95172627 | 11.4359072 |
| 11.8260275 | 2.88375999 | 9.71350712 | 9.08643731 | 10.4851386 | 9.08768716 | 11.4359072 |
| 20.6955481 | 12.1060244 | 0 | 22.7160933 | 21.6593005 | 26.4792485 | 11.8395274 |
| 8.8695206 | 11.7657408 | 0 | 9.08643731 | 14.2298309 | 4.54384358 | 10.3147398 |
| 11.8260275 | 20.4170207 | 0 | 19.1824788 | 14.9787694 | 15.9034525 | 15.9205766 |
| 23.6520549 | 13.265296 | 19.4270142 | 21.2016871 | 27.7107234 | 17.0394134 | 23.9929817 |
| 5.91301374 | 12.8038944 | 9.71350712 | 17.1632705 | 10.4851386 | 11.359609 | 8.07240505 |
| 20.6955481 | 10.7275872 | 9.71350712 | 13.1248539 | 11.9830155 | 6.81576537 | 10.0905063 |
| 2.95650687 | 11.3043392 | 0 | 8.07683317 | 5.99150775 | 9.08768716 | 7.39970463 |
| 5.91301374 | 10.8429376 | 0 | 8.41000253 | 7.48938469 | 15.9034525 | 9.41780589 |
| 23.6520549 | 16.2644064 | 0 | 14.134458 | 10.4851386 | 13.6315307 | 13.9024754 |
| 14.7825343 | 19.6095679 | 0 | 19.1824788 | 14.2298309 | 17.0394134 | 15.2478762 |
| 2.95650687 | 2.59538399 | 0 | 4.03841658 | 2.99575388 | 10.2236481 | 3.13926863 |
| 2.95650687 | 6.11357118 | 9.71350712 | 6.05762487 | 8.98726163 | 4.54384358 | 6.72700421 |
| 29.5650687 | 40.8340415 | 0 | 15.1440622 | 20.9702771 | 10.2236481 | 25.114149 |
| 546.953771 | 314.67589 | 446.821327 | 450.283449 | 527.252682 | 410.081883 | 514.167355 |
| 0 | 14.5341504 | 9.71350712 | 3.53361451 | 7.86385392 | 8.51970672 | 6.50277073 |
| 20.6955481 | 23.1854303 | 19.4270142 | 14.134458 | 11.9830155 | 10.2236481 | 18.8356118 |
| 5.91301374 | 8.99733118 | 9.71350712 | 13.1248539 | 12.731954 | 15.9034525 | 13.6782419 |
| 8.8695206 | 4.72936639 | 9.71350712 | 13.1248539 | 16.4766463 | 9.08768716 | 13.2297749 |
| 5.91301374 | 7.95917758 | 0 | 4.03841658 | 9.7362001 | 2.27192179 | 10.0905063 |
| 10.347774 | 4.07532962 | 0 | 6.56242695 | 4.11916158 | 8.89457381 | 3.3635021 |
| 8.8695206 | 8.53592958 | 0 | 5.04802073 | 6.74044622 | 9.08768716 | 11.6601406 |
| 5.91301374 | 20.6477215 | 0 | 4.03841658 | 9.7362001 | 6.81576537 | 14.3509423 |
| 8.8695206 | 7.84382718 | 0 | 10.0960415 | 7.48938469 | 21.583257 | 4.93313642 |
| 0 | 6.17124638 | 0 | 5.04802073 | 7.48938469 | 10.2236481 | 3.95772081 |
| 8.8695206 | 23.7621823 | 9.71350712 | 14.134458 | 14.9787694 | 17.0394134 | 9.52992263 |
| 8.8695206 | 10.2661856 | 19.4270142 | 8.07683317 | 8.98726163 | 5.67980448 | 11.4359072 |
| 50.2606167 | 49.3699711 | 38.8540285 | 43.4129783 | 50.1788774 | 56.7980448 | 49.5555977 |

| | | | | | | |
|---|---|---|---|---|---|---|
| 14.7825343 | 19.2635167 | 0 | 17.1632705 | 14.9787694 | 22.7192179 | 17.2659775 |
| 14.7825343 | 21.5901343 | 0 | 8.07683317 | 8.23832316 | 11.9275894 | 11.2116737 |
| 44.347603 | 19.1481663 | 48.5675356 | 44.4225824 | 35.9490465 | 36.3507487 | 34.083488 |
| 0 | 2.53770879 | 0 | 16.1536663 | 3.74469235 | 13.6315307 | 4.48466947 |
| 11.8260275 | 11.1889888 | 29.1405214 | 5.04802073 | 8.98726163 | 15.9034525 | 11.8843741 |
| 8.8695206 | 9.11268158 | 9.71350712 | 4.03841658 | 8.23832316 | 12.4955698 | 8.52087199 |
| 8.8695206 | 8.76663038 | 29.1405214 | 6.05762487 | 9.7362001 | 9.08768716 | 12.1086076 |
| 11.8260275 | 10.0354848 | 9.71350712 | 10.0960415 | 11.9830155 | 12.4955698 | 11.2116737 |
| 26.6085618 | 25.9538399 | 29.1405214 | 26.2497078 | 33.7022311 | 23.8551788 | 30.0472855 |
| 5.91301374 | 7.15172478 | 0 | 1.00960415 | 7.48938469 | 11.359609 | 6.72700421 |
| 14.7825343 | 24.5696351 | 0 | 20.1920829 | 26.2128464 | 29.5349833 | 28.2534177 |
| 20.6955481 | 11.1889888 | 9.71350712 | 18.1728746 | 18.7234617 | 15.9034525 | 15.2478762 |
| 47.3041099 | 30.3371551 | 29.1405214 | 39.3745617 | 26.9617849 | 32.942866 | 28.0291842 |
| 8.8695206 | 4.84471679 | 0 | 6.05762487 | 8.23832316 | 7.95172627 | 5.60583684 |
| 17.7390412 | 1.96095679 | 19.4270142 | 4.03841658 | 8.98726163 | 6.81576537 | 7.6239381 |
| 20.6955481 | 15.6876544 | 0 | 5.04802073 | 5.24256928 | 9.08768716 | 8.52087199 |
| 0 | 8.07452798 | 0 | 3.02881244 | 8.98726163 | 7.95172627 | 4.03620252 |
| 14.7825343 | 13.9573984 | 0 | 5.04802073 | 8.98726163 | 10.2236481 | 8.29663852 |
| 26.6085618 | 27.2226943 | 9.71350712 | 25.2401036 | 30.7064772 | 21.583257 | 28.7018846 |
| 20.6955481 | 15.3416032 | 19.4270142 | 17.1632705 | 7.48938469 | 10.2236481 | 21.0779465 |
| 35.4780824 | 39.1037855 | 19.4270142 | 35.3361451 | 26.9617849 | 21.583257 | 28.9261181 |
| 20.6955481 | 24.5696351 | 0 | 14.134458 | 14.9787694 | 9.08768716 | 13.4540084 |
| 0 | 3.22981119 | 9.71350712 | 6.05762487 | 8.23832316 | 5.67980448 | 4.70890294 |
| 32.5215755 | 17.5332608 | 38.8540285 | 14.134458 | 22.4681541 | 22.7192179 | 20.1810126 |
| 11.8260275 | 9.34338237 | 0 | 10.0960415 | 7.48938469 | 11.359609 | 7.6239381 |
| 44.347603 | 35.9316495 | 0 | 42.4033741 | 41.1916158 | 30.6709442 | 41.2589591 |
| 0 | 2.53770879 | 9.71350712 | 5.04802073 | 4.49363081 | 9.08768716 | 6.50277073 |
| 59.1301374 | 72.5554014 | 0 | 49.4706031 | 42.6894927 | 52.2542012 | 62.7853726 |
| 14.7825343 | 11.3043392 | 0 | 7.06722902 | 8.23832316 | 15.9034525 | 8.29663852 |
| 38.4345893 | 28.6068991 | 48.5675356 | 30.2881244 | 45.6852466 | 40.8945922 | 47.9859633 |
| 5.91301374 | 9.22803198 | 9.71350712 | 10.0960415 | 10.4851386 | 3.40788269 | 12.5570745 |
| 2.95650687 | 5.99822078 | 9.71350712 | 8.07683317 | 8.23832316 | 3.40788269 | 6.50277073 |
| 5.91301374 | 27.6840959 | 9.71350712 | 16.1536663 | 14.2298309 | 11.359609 | 12.781308 |
| 17.7390412 | 11.8810912 | 0 | 9.08643731 | 15.7277078 | 13.6315307 | 13.6782419 |
| 13.3042809 | 5.53681919 | 29.1405214 | 23.7256974 | 19.4724002 | 21.0152766 | 16.0326934 |
| 23.6520549 | 17.0718592 | 9.71350712 | 10.0960415 | 20.9702771 | 9.08768716 | 12.4449578 |
| 8.8695206 | 19.3788671 | 29.1405214 | 16.1536663 | 16.4766463 | 14.7674916 | 15.0236427 |
| 32.5215755 | 40.6033407 | 9.71350712 | 17.1632705 | 25.4639079 | 17.0394134 | 23.9929817 |
| 23.6520549 | 27.2226943 | 19.4270142 | 20.1920829 | 15.7277078 | 14.7674916 | 22.6475808 |
| 41.3910961 | 21.3398239 | 19.4270142 | 39.3745617 | 29.2086003 | 24.9911397 | 36.9985231 |
| 29.5650687 | 11.0736384 | 9.71350712 | 8.07683317 | 10.4851386 | 12.4955698 | 8.07240505 |
| 2.95650687 | 5.07541759 | 0 | 5.04802073 | 1.49787694 | 9.08768716 | 4.70890294 |
| 5.91301374 | 6.69032318 | 0 | 2.01920829 | 5.24256928 | 11.359609 | 8.74510547 |
| 17.7390412 | 26.6459423 | 0 | 9.08643731 | 11.9830155 | 17.0394134 | 17.7144444 |
| 53.2171236 | 33.5669663 | 9.71350712 | 56.5378322 | 63.6597699 | 47.7103576 | 54.2645006 |
| 14.7825343 | 35.7586239 | 19.4270142 | 7.06722902 | 10.4851386 | 6.81576537 | 13.2297749 |

| | | | | | | |
|---|---|---|---|---|---|---|
| 20.6955481 | 8.88198078 | 9.71350712 | 16.1536663 | 15.7277078 | 13.6315307 | 13.6782419 |
| 11.8260275 | 5.30611839 | 9.71350712 | 1.00960415 | 8.98726163 | 3.40788269 | 4.260436 |
| 26.6085618 | 11.3043392 | 38.8540285 | 16.1536663 | 22.4681541 | 19.3113352 | 20.6294796 |
| 2.95650687 | 7.95917758 | 0 | 3.02881244 | 8.98726163 | 6.81576537 | 4.03620252 |
| 44.347603 | 53.2918847 | 38.8540285 | 33.3169368 | 33.7022311 | 35.2147878 | 30.7199859 |
| 44.347603 | 64.1348222 | 29.1405214 | 29.2785202 | 24.7149695 | 27.2630615 | 33.4107876 |
| 17.7390412 | 11.7657408 | 0 | 16.1536663 | 8.98726163 | 20.4472961 | 12.1086076 |
| 5.91301374 | 19.8402687 | 0 | 10.0960415 | 11.234077 | 10.2236481 | 7.84817157 |
| 1.47825343 | 2.59538399 | 0 | 2.52401036 | 5.24256928 | 12.4955698 | 3.02715189 |
| 17.7390412 | 38.8730847 | 0 | 28.2689161 | 21.7192156 | 29.5349833 | 23.0960478 |
| 79.8256854 | 49.7160223 | 19.4270142 | 57.5474363 | 44.9363081 | 169.258173 | 46.1920955 |
| 5.91301374 | 12.3424928 | 9.71350712 | 11.1056456 | 14.9787694 | 10.2236481 | 9.41780589 |
| 13.3042809 | 14.4188 | 9.71350712 | 17.1632705 | 13.4808924 | 23.8551788 | 24.1050984 |
| 5.91301374 | 3.69121279 | 0 | 11.1056456 | 5.61703852 | 10.7916285 | 7.17547115 |
| 17.7390412 | 17.1872096 | 9.71350712 | 9.08643731 | 8.23832316 | 13.6315307 | 11.6601406 |
| 11.8260275 | 13.7266976 | 9.71350712 | 15.1440622 | 13.4808924 | 11.359609 | 12.332841 |
| 29.5650687 | 19.7249183 | 0 | 23.2208954 | 25.4639079 | 30.6709442 | 31.8411532 |
| 8.8695206 | 22.3779775 | 38.8540285 | 21.2016871 | 11.234077 | 22.7192179 | 18.3871448 |
| 17.7390412 | 9.34338237 | 19.4270142 | 19.1824788 | 11.234077 | 20.4472961 | 15.0236427 |
| 11.8260275 | 49.0239199 | 0 | 15.1440622 | 17.2255848 | 17.0394134 | 20.4052461 |
| 14.7825343 | 10.1508352 | 0 | 6.05762487 | 8.23832316 | 3.40788269 | 8.07240505 |
| 11.8260275 | 2.88375999 | 0 | 3.02881244 | 7.48938469 | 2.27192179 | 9.19357241 |
| 35.4780824 | 19.2635167 | 0 | 20.1920829 | 30.7064772 | 40.8945922 | 39.9135583 |
| 17.7390412 | 10.381536 | 0 | 5.04802073 | 10.4851386 | 6.81576537 | 11.4359072 |
| 5.91301374 | 16.725808 | 0 | 12.1152497 | 5.99150775 | 10.2236481 | 13.4540084 |
| 141.91233 | 93.7798749 | 67.9945498 | 98.9412063 | 96.6130625 | 79.5172627 | 106.959367 |
| 11.8260275 | 11.9964416 | 0 | 15.1440622 | 14.2298309 | 11.359609 | 10.9874402 |
| 87.9560793 | 10.4680488 | 44.4878626 | 30.2780283 | 32.2268223 | 40.1334984 | 24.3741786 |
| 8.8695206 | 25.6077887 | 0 | 6.05762487 | 12.731954 | 12.4955698 | 14.1267088 |
| 2.95650687 | 3.69121279 | 0 | 10.0960415 | 9.7362001 | 17.0394134 | 11.4359072 |
| 29.5650687 | 21.2244735 | 38.8540285 | 24.2304995 | 28.4596618 | 29.5349833 | 34.7561884 |
| 17.7390412 | 5.76751998 | 0 | 12.1152497 | 5.24256928 | 3.40788269 | 10.0905063 |
| 35.4780824 | 35.5279231 | 29.1405214 | 34.326541 | 42.6894927 | 34.0788269 | 40.1377918 |
| 5.91301374 | 11.5154304 | 38.8540285 | 15.6488643 | 13.4808924 | 13.6315307 | 3.02715189 |
| 2.95650687 | 8.53592958 | 19.4270142 | 15.1440622 | 9.7362001 | 17.0394134 | 9.64203936 |
| 26.6085618 | 27.7994463 | 9.71350712 | 18.1728746 | 17.2255848 | 15.9034525 | 25.7868495 |
| 8.8695206 | 36.2200255 | 0 | 26.2497078 | 29.2086003 | 21.583257 | 23.9929817 |
| 0 | 22.3779775 | 0 | 12.1152497 | 5.99150775 | 15.9034525 | 13.2297749 |
| 8.8695206 | 5.99822078 | 0 | 6.05762487 | 8.23832316 | 10.2236481 | 11.8843741 |
| 5.91301374 | 1.153504 | 0 | 12.1152497 | 8.23832316 | 6.81576537 | 10.0905063 |
| 11.8260275 | 36.2580911 | 0 | 16.1536663 | 14.2298309 | 11.9275894 | 15.43399 |
| 59.1301374 | 63.212019 | 58.2810427 | 68.6530819 | 46.4341851 | 64.749771 | 65.4761743 |
| 2.95650687 | 1.8456064 | 0 | 5.04802073 | 12.3574847 | 8.51970672 | 3.69985231 |
| 5.91301374 | 7.84382718 | 0 | 5.04802073 | 7.48938469 | 1.1359609 | 8.07240505 |
| 5.91301374 | 9.45873277 | 0 | 6.05762487 | 6.74044622 | 13.6315307 | 6.05430379 |
| 29.5650687 | 35.5279231 | 9.71350712 | 23.2208954 | 25.4639079 | 17.0394134 | 33.4107876 |

| | | | | | | |
|---|---|---|---|---|---|---|
| 45.8258564 | 20.0709695 | 48.5675356 | 38.3649575 | 32.9532926 | 24.9911397 | 26.9080168 |
| 5.91301374 | 12.688544 | 9.71350712 | 13.1248539 | 2.99575388 | 9.08768716 | 8.52087199 |
| 20.6955481 | 9.92013437 | 9.71350712 | 28.2689161 | 23.2170925 | 21.583257 | 18.6113783 |
| 17.7390412 | 12.2271424 | 0 | 19.1824788 | 8.98726163 | 19.8793157 | 16.1448101 |
| 6.888661 | 10.5741711 | 0 | 7.65279942 | 9.98334979 | 4.54384358 | 8.73613613 |
| 5.91301374 | 4.49866559 | 19.4270142 | 23.2208954 | 8.98726163 | 12.4955698 | 12.5570745 |
| 32.5215755 | 27.7994463 | 0 | 24.2304995 | 16.4766463 | 17.0394134 | 25.114149 |
| 5.91301374 | 81.3220318 | 0 | 17.1632705 | 15.7277078 | 10.2236481 | 15.4721097 |
| 2.95650687 | 22.2626271 | 0 | 9.08643731 | 9.7362001 | 3.40788269 | 9.19357241 |
| 38.4345893 | 32.5288127 | 29.1405214 | 36.3457492 | 32.2043542 | 23.8551788 | 29.1503516 |
| 17.7390412 | 10.7275872 | 9.71350712 | 35.3361451 | 25.4639079 | 21.583257 | 24.4414486 |
| 11.8260275 | 33.6823167 | 9.71350712 | 30.2881244 | 30.7064772 | 22.7192179 | 26.6837834 |
| 53.2171236 | 33.5669663 | 29.1405214 | 38.3649575 | 38.9448004 | 43.166514 | 45.7436286 |
| 2.95650687 | 4.03726399 | 0 | 6.05762487 | 4.49363081 | 10.2236481 | 8.52087199 |
| 8.8695206 | 8.53592958 | 0 | 14.134458 | 3.74469235 | 5.67980448 | 9.19357241 |
| 14.7825343 | 7.61312638 | 0 | 10.237386 | 9.7362001 | 5.67980448 | 8.29663852 |
| 11.8260275 | 15.4569536 | 0 | 14.134458 | 8.98726163 | 11.359609 | 10.5389733 |
| 14.7825343 | 8.30522878 | 9.71350712 | 8.07683317 | 4.49363081 | 11.359609 | 8.74510547 |
| 124.173288 | 96.2022333 | 116.562085 | 154.469434 | 164.766463 | 88.6049498 | 155.618031 |
| 41.3910961 | 56.0602942 | 29.1405214 | 39.3745617 | 30.7064772 | 31.8069051 | 45.7436286 |
| 20.6955481 | 7.38242558 | 0 | 4.03841658 | 10.4851386 | 12.4955698 | 8.29663852 |
| 2.95650687 | 3.57586239 | 0 | 4.03841658 | 7.48938469 | 3.40788269 | 4.93313642 |
| 42.3667434 | 33.3028139 | 1.94270142 | 43.110097 | 50.5233891 | 47.1423772 | 43.214275 |
| 56.1736305 | 23.6468319 | 29.1405214 | 55.528228 | 34.4511696 | 30.6709442 | 48.4344303 |
| 5.91301374 | 4.96006719 | 0 | 6.05762487 | 8.23832316 | 5.67980448 | 4.260436 |
| 17.7390412 | 19.8402687 | 29.1405214 | 26.2497078 | 24.7149695 | 21.583257 | 26.4595499 |
| 20.6955481 | 13.3806464 | 9.71350712 | 17.1632705 | 11.234077 | 9.08768716 | 15.4721097 |
| 56.1736305 | 49.9663326 | 9.71350712 | 60.5762487 | 74.272228 | 62.6709626 | 68.1288563 |
| 53.2171236 | 10.4968864 | 29.1405214 | 26.2497078 | 21.7192156 | 21.583257 | 29.5988185 |
| 29.5650687 | 46.2555103 | 48.5675356 | 27.2593119 | 41.1916158 | 23.8551788 | 50.0040646 |
| 91.6517129 | 95.9715325 | 67.9945498 | 135.286956 | 88.3747393 | 81.7891845 | 132.297749 |
| 14.7825343 | 21.6858751 | 29.1405214 | 18.1728746 | 8.98726163 | 13.6315307 | 12.1086076 |
| 2.95650687 | 10.1508352 | 0 | 10.0960415 | 8.98726163 | 10.2236481 | 6.50277073 |
| 8.8695206 | 14.6495008 | 9.71350712 | 7.06722902 | 8.98726163 | 10.2236481 | 6.05430379 |
| 14.7825343 | 8.65127998 | 9.71350712 | 6.05762487 | 11.9830155 | 7.95172627 | 16.593277 |
| 26.6085618 | 24.9156863 | 19.4270142 | 27.2593119 | 14.2298309 | 18.1753743 | 22.4233474 |
| 8.8695206 | 8.07452798 | 9.71350712 | 4.03841658 | 7.48938469 | 4.54384358 | 6.95123768 |
| 2.95650687 | 7.72847678 | 9.71350712 | 4.03841658 | 10.4851386 | 3.40788269 | 6.95123768 |
| 20.6955481 | 15.9183552 | 0 | 15.2652147 | 18.7234617 | 13.6315307 | 20.0688959 |
| 14.7825343 | 15.3416032 | 0 | 14.134458 | 13.4808924 | 9.08768716 | 10.7632067 |
| 14.7825343 | 8.53592958 | 9.71350712 | 21.2016871 | 21.7192156 | 19.3113352 | 19.7325457 |
| 5.91301374 | 13.4959968 | 19.4270142 | 12.1152497 | 9.7362001 | 12.4955698 | 10.9874402 |
| 14.7825343 | 52.7151327 | 9.71350712 | 18.1728746 | 21.7192156 | 11.359609 | 22.5354641 |
| 23.6520549 | 22.1472767 | 19.4270142 | 25.2401036 | 14.9787694 | 30.6709442 | 17.2659775 |
| 14.7825343 | 14.4188 | 0 | 14.134458 | 10.4851386 | 5.67980448 | 9.64203936 |
| 14.7825343 | 9.11268158 | 9.71350712 | 21.2016871 | 15.7277078 | 20.4472961 | 11.6601406 |

| | | | | | | |
|---|---|---|---|---|---|---|
| 14.7825343 | 16.6104576 | 0 | 9.08643731 | 14.2298309 | 4.54384358 | 8.96933894 |
| 77.5491751 | 18.8101897 | 60.5151494 | 92.8936774 | 87.550907 | 93.8530892 | 76.6228202 |
| 11.8260275 | 5.30611839 | 0 | 8.07683317 | 7.11491546 | 11.9275894 | 6.83912094 |
| 23.6520549 | 10.8429376 | 19.4270142 | 8.07683317 | 6.74044622 | 9.08768716 | 11.4359072 |
| 82.7821923 | 48.9085695 | 29.1405214 | 49.4706031 | 61.4129545 | 49.414299 | 59.8703374 |
| 19.2172946 | 23.2431055 | 9.71350712 | 11.6104477 | 14.2298309 | 19.8793157 | 15.0236427 |
| 14.7825343 | 12.5155184 | 0 | 9.59123938 | 4.11916158 | 12.4955698 | 8.40875526 |
| 41.3910961 | 82.7062366 | 29.1405214 | 47.4513948 | 44.9363081 | 39.7586313 | 55.8341349 |
| 5.91301374 | 5.30611839 | 0 | 3.02881244 | 8.98726163 | 9.08768716 | 9.86627284 |
| 41.3910961 | 55.5988926 | 48.5675356 | 42.4033741 | 40.4426773 | 34.0788269 | 36.5500562 |
| 35.4780824 | 37.8349311 | 48.5675356 | 42.4033741 | 30.7064772 | 40.8945922 | 38.1196905 |
| 41.3910961 | 29.6450527 | 38.8540285 | 25.2401036 | 19.4724002 | 34.0788269 | 27.5067202 |
| 53.4240791 | 135.421369 | 20.0098247 | 47.4513948 | 51.8040739 | 48.9371954 | 52.4706328 |
| 2.95650687 | 13.6113472 | 9.71350712 | 10.0960415 | 16.4766463 | 19.3113352 | 12.1086076 |
| 2.95650687 | 11.7657408 | 0 | 7.06722902 | 8.23832316 | 13.6315307 | 9.41780589 |
| 11.8260275 | 13.1499456 | 19.4270142 | 7.06722902 | 12.731954 | 9.08768716 | 10.9874402 |
| 8.8695206 | 7.55545118 | 9.71350712 | 16.1536663 | 16.1021771 | 13.6315307 | 8.63298873 |
| 2.95650687 | 5.19076799 | 0 | 6.05762487 | 10.4851386 | 4.54384358 | 7.6239381 |
| 85.7386992 | 63.212019 | 116.562085 | 95.9123938 | 70.4002161 | 74.9734191 | 102.474697 |
| 2.95650687 | 17.5332608 | 9.71350712 | 17.1632705 | 9.7362001 | 13.6315307 | 9.19357241 |
| 5.91301374 | 5.42146879 | 0 | 6.05762487 | 6.74044622 | 9.08768716 | 8.74510547 |
| 17.7390412 | 20.9937727 | 9.71350712 | 13.1248539 | 5.99150775 | 7.95172627 | 10.7632067 |
| 8.8695206 | 14.8802016 | 9.71350712 | 10.0960415 | 8.98726163 | 14.7674916 | 8.52087199 |
| 62.0866442 | 28.7222495 | 48.5675356 | 39.3745617 | 40.4426773 | 29.5349833 | 52.6948663 |
| 0 | 6.92102398 | 29.1405214 | 7.06722902 | 7.48938469 | 2.27192179 | 6.27853726 |
| 2.95650687 | 7.03637438 | 0 | 4.03841658 | 11.9830155 | 7.95172627 | 6.95123768 |
| 5.91301374 | 8.88198078 | 0 | 10.0960415 | 8.23832316 | 5.67980448 | 8.74510547 |
| 0.73912672 | 5.44799938 | 7.57653555 | 5.75474363 | 10.3203721 | 7.41782465 | 5.41972306 |
| 70.9561648 | 50.0620735 | 38.8540285 | 78.7491234 | 55.4214467 | 62.4778492 | 77.1363149 |
| 23.6520549 | 21.5705247 | 0 | 15.1440622 | 13.4808924 | 26.1271006 | 16.3690436 |
| 47.3041099 | 38.7577343 | 19.4270142 | 36.3457492 | 32.9532926 | 24.9911397 | 32.5138537 |
| 32.5215755 | 14.8802016 | 0 | 25.2401036 | 20.9702771 | 18.1753743 | 25.114149 |
| 7.39126717 | 4.90239199 | 9.71350712 | 4.54321866 | 7.48938469 | 12.4955698 | 9.97838957 |
| 0 | 11.8810912 | 19.4270142 | 13.1248539 | 10.4851386 | 7.95172627 | 9.64203936 |
| 124.173288 | 98.2785405 | 9.71350712 | 72.6914985 | 56.1703852 | 74.9734191 | 110.098636 |
| 0 | 5.07541759 | 0 | 11.1056456 | 6.74044622 | 4.54384358 | 8.96933894 |
| 32.5215755 | 13.6113472 | 29.1405214 | 31.2977285 | 32.9532926 | 35.2147878 | 30.4957524 |
| 23.6520549 | 14.8802016 | 9.71350712 | 12.1152497 | 16.4766463 | 12.4955698 | 10.3147398 |
| 11.8260275 | 9.92013437 | 0 | 11.1056456 | 12.731954 | 11.359609 | 13.9024754 |
| 32.5215755 | 46.1401599 | 29.1405214 | 43.4129783 | 52.4256928 | 44.3024749 | 46.1920955 |
| 8.8695206 | 7.72847678 | 0 | 5.04802073 | 9.7362001 | 10.2236481 | 7.17547115 |
| 0 | 3.92191359 | 19.4270142 | 3.02881244 | 4.49363081 | 9.08768716 | 6.50277073 |
| 26.6085618 | 13.0345952 | 9.71350712 | 5.04802073 | 7.48938469 | 10.2236481 | 9.64203936 |
| 0 | 10.6122368 | 0 | 6.05762487 | 8.98726163 | 10.2236481 | 7.84817157 |
| 14.7825343 | 17.4179104 | 19.4270142 | 21.2016871 | 31.4554157 | 26.1271006 | 26.2353164 |
| 11.8260275 | 5.07541759 | 0 | 5.04802073 | 8.98726163 | 3.40788269 | 6.50277073 |

| | | | | | | |
|---|---|---|---|---|---|---|
| 32.5215755 | 15.9183552 | 29.1405214 | 22.2112912 | 21.7192156 | 19.3113352 | 18.1629114 |
| 2.95650687 | 1.96095679 | 0 | 2.01920829 | 7.48938469 | 2.27192179 | 3.13926863 |
| 17.7390412 | 25.2617375 | 0 | 10.0960415 | 15.3532386 | 11.359609 | 19.7325457 |
| 14.7825343 | 8.99733118 | 0 | 12.1152497 | 14.2298309 | 20.4472961 | 13.9024754 |
| 0 | 17.0718592 | 9.71350712 | 13.1248539 | 17.2255848 | 10.2236481 | 16.593277 |
| 2.95650687 | 14.995552 | 0 | 7.06722902 | 9.7362001 | 4.54384358 | 10.3147398 |
| 8.8695206 | 14.8802016 | 0 | 4.03841658 | 8.98726163 | 4.54384358 | 10.0905063 |
| 8.8695206 | 9.57408317 | 29.1405214 | 8.07683317 | 13.4808924 | 7.95172627 | 9.86627284 |
| 14.7825343 | 10.8429376 | 0 | 7.06722902 | 13.4808924 | 9.08768716 | 12.1086076 |
| 23.6520549 | 89.9733118 | 9.71350712 | 20.1920829 | 26.2128464 | 24.9911397 | 39.4650913 |
| 2.95650687 | 4.15261439 | 0 | 10.0960415 | 3.74469235 | 9.08768716 | 5.83007031 |
| 2.95650687 | 11.6503904 | 9.71350712 | 8.07683317 | 3.74469235 | 9.08768716 | 8.07240505 |
| 14.7825343 | 12.8038944 | 19.4270142 | 13.1248539 | 13.8553617 | 24.4231593 | 17.2659775 |
| 17.7390412 | 14.6495008 | 0 | 15.1440622 | 11.9830155 | 5.67980448 | 15.6963431 |
| 8.8695206 | 12.3424928 | 9.71350712 | 8.07683317 | 11.234077 | 19.3113352 | 14.1267088 |
| 17.7390412 | 49.0239199 | 9.71350712 | 22.2112912 | 14.9787694 | 21.583257 | 21.7506469 |
| 26.6085618 | 22.8393791 | 0 | 20.1920829 | 12.731954 | 15.9034525 | 20.6294796 |
| 38.4345893 | 47.5243647 | 29.1405214 | 15.1440622 | 28.4596618 | 15.9034525 | 24.8899156 |
| 368.085105 | 314.618215 | 267.121446 | 365.476701 | 353.498957 | 302.733579 | 383.327123 |
| 20.6955481 | 3.11446079 | 9.71350712 | 10.0960415 | 8.23832316 | 9.08768716 | 8.07240505 |
| 44.347603 | 26.5305919 | 9.71350712 | 22.2112912 | 20.9702771 | 23.8551788 | 30.7199859 |
| 5.91301374 | 12.4578432 | 0 | 11.1056456 | 5.24256928 | 5.67980448 | 7.6239381 |
| 91.6517129 | 60.7896606 | 58.2810427 | 75.7203109 | 100.357755 | 78.3813018 | 90.1418564 |
| 14.7825343 | 15.6876544 | 9.71350712 | 19.1824788 | 13.4808924 | 18.1753743 | 13.6782419 |
| 50.2606167 | 74.747059 | 48.5675356 | 40.3841658 | 32.9532926 | 43.166514 | 52.4706328 |
| 47.3041099 | 45.2173567 | 48.5675356 | 29.2785202 | 36.697985 | 30.6709442 | 41.0347257 |
| 23.6520549 | 27.3380447 | 29.1405214 | 16.1536663 | 11.234077 | 12.4955698 | 11.4359072 |
| 41.3910961 | 87.6663038 | 38.8540285 | 58.5570405 | 57.6682621 | 59.0699666 | 62.7853726 |
| 195.129453 | 304.986457 | 77.708057 | 159.517455 | 158.774955 | 168.122213 | 221.542672 |
| 5.91301374 | 3.11446079 | 0 | 4.03841658 | 11.234077 | 9.08768716 | 7.17547115 |
| 2.95650687 | 6.11357118 | 19.4270142 | 7.06722902 | 5.99150775 | 9.08768716 | 8.52087199 |
| 230.607536 | 248.580111 | 242.837678 | 226.151329 | 174.502663 | 134.043386 | 233.202812 |
| 26.6085618 | 54.3300383 | 19.4270142 | 24.2304995 | 14.2298309 | 14.7674916 | 29.5988185 |
| 38.4345893 | 64.5962238 | 0 | 62.595457 | 47.932062 | 43.166514 | 54.2645006 |
| 2.95650687 | 7.15172478 | 0 | 0 | 9.7362001 | 12.4955698 | 4.03620252 |
| 8.8695206 | 9.22803198 | 0 | 6.05762487 | 5.99150775 | 11.359609 | 6.27853726 |
| 29.5650687 | 11.9964416 | 19.4270142 | 24.2304995 | 31.4554157 | 18.1753743 | 21.7506469 |
| 14.7825343 | 4.84471679 | 9.71350712 | 9.08643731 | 8.98726163 | 15.9034525 | 4.48466947 |
| 5.91301374 | 9.45873277 | 9.71350712 | 5.04802073 | 10.4851386 | 9.08768716 | 10.7632067 |
| 17.7390412 | 35.0665215 | 9.71350712 | 24.2304995 | 20.2213387 | 26.1271006 | 36.7742897 |
| 32.5215755 | 22.1472767 | 29.1405214 | 35.3361451 | 39.6937389 | 49.9822794 | 36.3258227 |
| 41.3910961 | 36.3353759 | 19.4270142 | 37.3553534 | 24.7149695 | 29.5349833 | 32.5138537 |
| 14.7825343 | 16.6104576 | 0 | 11.1056456 | 8.23832316 | 7.38374582 | 7.96028831 |
| 11.8260275 | 3.57586239 | 0 | 3.02881244 | 5.99150775 | 10.2236481 | 7.6239381 |
| 12.5651542 | 15.8087723 | 19.4270142 | 26.6131653 | 32.7885262 | 49.0394319 | 13.846417 |
| 0 | 4.49866559 | 9.71350712 | 11.1056456 | 2.24681541 | 12.4955698 | 8.07240505 |

| | | | | | | |
|---|---|---|---|---|---|---|
| 29.5650687 | 29.7604031 | 19.4270142 | 42.4033741 | 31.4554157 | 22.7192179 | 29.1503516 |
| 11.8260275 | 29.9911039 | 9.71350712 | 19.1824788 | 11.234077 | 13.6315307 | 18.6113783 |
| 11.8260275 | 15.6876544 | 9.71350712 | 17.1632705 | 11.234077 | 13.6315307 | 15.0236427 |
| 13.3042809 | 3.74888799 | 9.71350712 | 6.05762487 | 13.1064232 | 12.4955698 | 11.2497934 |
| 8.8695206 | 6.34427198 | 9.71350712 | 4.03841658 | 7.48938469 | 11.359609 | 9.86627284 |
| 47.3041099 | 24.9156863 | 29.1405214 | 36.3457492 | 37.4469235 | 19.3113352 | 28.7018846 |
| 17.7390412 | 12.111792 | 38.8540285 | 4.03841658 | 8.98726163 | 14.7674916 | 9.64203936 |
| 5.91301374 | 4.61401599 | 0 | 7.06722902 | 8.23832316 | 6.81576537 | 6.27853726 |
| 203.998974 | 176.601462 | 155.416114 | 183.747955 | 153.532386 | 137.451268 | 217.282236 |
| 0 | 2.07630719 | 19.4270142 | 7.06722902 | 2.99575388 | 9.08768716 | 5.15736989 |
| 5.91301374 | 11.6503904 | 0 | 8.07683317 | 11.9830155 | 19.3113352 | 4.93313642 |
| 8.8695206 | 3.11446079 | 9.71350712 | 8.07683317 | 9.7362001 | 3.40788269 | 10.3147398 |
| 20.6955481 | 6.92102398 | 9.71350712 | 8.07683317 | 10.4851386 | 4.54384358 | 7.39970463 |
| 33.7928735 | 23.6237619 | 14.5702607 | 25.9569226 | 20.947809 | 29.2055546 | 24.7217405 |
| 88.695206 | 119.733715 | 19.4270142 | 97.9316021 | 69.6512776 | 65.8857319 | 110.547102 |
| 85.7386992 | 72.440051 | 126.275593 | 83.7971441 | 82.3832316 | 60.2059275 | 93.5053585 |
| 319.302742 | 361.046751 | 233.124171 | 301.87164 | 283.098741 | 245.367553 | 329.623206 |
| 70.9561648 | 42.3335967 | 67.9945498 | 54.5186239 | 57.6682621 | 47.7103576 | 72.6516454 |
| 26.6085618 | 11.3043392 | 29.1405214 | 16.1536663 | 14.2298309 | 12.4955698 | 15.2478762 |
| 17.7390412 | 20.9937727 | 38.8540285 | 24.2304995 | 15.7277078 | 20.4472961 | 15.4721097 |
| 85.7386992 | 73.3628542 | 48.5675356 | 56.5378322 | 48.6810005 | 38.6226704 | 73.5485793 |
| 15.5807912 | 9.90513882 | 0.48567536 | 31.0453275 | 23.0298579 | 25.331928 | 7.7853862 |
| 14.7825343 | 17.6486112 | 29.1405214 | 21.2016871 | 21.7192156 | 27.2630615 | 22.4233474 |
| 56.1736305 | 87.6086286 | 29.1405214 | 50.4802073 | 35.200108 | 35.2147878 | 36.9985231 |
| 0 | 4.38331519 | 9.71350712 | 12.1152497 | 14.9787694 | 15.9034525 | 5.60583684 |
| 2.95650687 | 6.34427198 | 0 | 4.03841658 | 11.9830155 | 6.81576537 | 8.96933894 |
| 35.4780824 | 19.7249183 | 0 | 23.2208954 | 26.2128464 | 32.942866 | 30.4957524 |
| 8.8695206 | 12.5155184 | 0 | 10.6008435 | 9.7362001 | 18.1753743 | 9.97838957 |
| 8.8695206 | 19.7249183 | 0 | 20.1920829 | 6.74044622 | 20.4472961 | 15.2478762 |
| 41.3910961 | 42.7949983 | 19.4270142 | 38.3649575 | 46.4341851 | 36.3507487 | 32.9623206 |
| 5.91301374 | 2.30700799 | 0 | 3.02881244 | 7.48938469 | 6.81576537 | 4.260436 |
| 5.91301374 | 2.19165759 | 19.4270142 | 16.1536663 | 9.7362001 | 13.6315307 | 17.7144444 |
| 50.2606167 | 48.7932191 | 19.4270142 | 58.5570405 | 53.9235698 | 38.6226704 | 56.0583684 |
| 20.6955481 | 18.3407136 | 0 | 22.2112912 | 35.9490465 | 23.8551788 | 22.4233474 |
| 26.6085618 | 9.22803198 | 9.71350712 | 15.1440622 | 13.4808924 | 9.08768716 | 14.3509423 |
| 8.8695206 | 11.3043392 | 9.71350712 | 15.1440622 | 16.4766463 | 6.81576537 | 13.2297749 |
| 24.6277022 | 19.6960807 | 33.9972749 | 31.1260958 | 27.3362541 | 38.7135473 | 35.2607137 |
| 0 | 5.07541759 | 0 | 4.03841658 | 8.23832316 | 11.359609 | 4.70890294 |
| 26.6085618 | 34.6051199 | 29.1405214 | 19.1824788 | 24.7149695 | 22.7192179 | 32.0653867 |
| 5.91301374 | 5.88287038 | 0 | 9.08643731 | 3.74469235 | 10.2236481 | 8.74510547 |
| 5.91301374 | 5.30611839 | 0 | 5.04802073 | 7.48938469 | 4.54384358 | 6.27853726 |
| 0 | 8.53592958 | 9.71350712 | 19.1824788 | 8.98726163 | 4.54384358 | 10.3147398 |
| 14.7825343 | 20.7630719 | 29.1405214 | 19.1824788 | 15.7277078 | 22.7192179 | 21.5264135 |
| 8.8695206 | 21.3398239 | 0 | 12.1152497 | 5.99150775 | 10.2236481 | 10.9874402 |
| 59.1301374 | 73.1898286 | 33.9972749 | 84.3019462 | 48.3065313 | 46.5743967 | 68.3912094 |
| 17.7390412 | 7.49777598 | 19.4270142 | 28.2689161 | 27.7107234 | 14.7674916 | 16.8175105 |

| | | | | | | |
|---|---|---|---|---|---|---|
| 8.8695206 | 31.3753087 | 0 | 28.2689161 | 30.7064772 | 26.1271006 | 26.2353164 |
| 79.8256854 | 136.574873 | 58.2810427 | 52.4994156 | 66.6555237 | 57.9340057 | 66.8215751 |
| 14.7825343 | 14.7648512 | 9.71350712 | 10.0960415 | 16.4766463 | 14.7674916 | 10.9874402 |
| 8.8695206 | 8.42057918 | 0 | 10.0960415 | 11.234077 | 7.95172627 | 6.27853726 |
| 11.8260275 | 8.18987838 | 19.4270142 | 6.05762487 | 8.98726163 | 7.95172627 | 6.27853726 |
| 47.3041099 | 39.3344863 | 29.1405214 | 34.326541 | 29.9575388 | 31.8069051 | 25.562616 |
| 26.6085618 | 14.6495008 | 0 | 38.3649575 | 20.2213387 | 29.5349833 | 22.1991139 |
| 2.4539007 | 17.1445299 | 9.71350712 | 11.3580466 | 15.0386845 | 23.5711886 | 13.2477136 |
| 14.7825343 | 16.0337056 | 0 | 12.1152497 | 12.731954 | 11.359609 | 13.2297749 |
| 14.9894898 | 10.528031 | 8.06221091 | 20.2728512 | 22.2210044 | 27.4902537 | 10.4089178 |
| 1.47825343 | 2.55731836 | 0 | 2.6047787 | 7.00257469 | 10.8825054 | 3.4150758 |
| 2.95650687 | 3.34516159 | 0 | 15.1440622 | 8.23832316 | 20.4472961 | 11.2116737 |
| 8.8695206 | 6.57497278 | 0 | 12.1152497 | 8.23832316 | 3.40788269 | 7.84817157 |
| 0 | 3.92191359 | 0 | 9.08643731 | 9.7362001 | 13.6315307 | 8.07240505 |
| 8.8695206 | 7.61312638 | 0 | 7.06722902 | 15.7277078 | 10.2236481 | 7.84817157 |
| 212.868494 | 76.477315 | 155.416114 | 179.709538 | 182.740986 | 186.297587 | 213.470267 |
| 20.6955481 | 14.5341504 | 0 | 6.05762487 | 8.98726163 | 7.95172627 | 12.1086076 |
| 20.6955481 | 37.3735295 | 9.71350712 | 31.2977285 | 37.4469235 | 38.6226704 | 37.4469901 |
| 23.6520549 | 36.7967775 | 9.71350712 | 22.2112912 | 18.7234617 | 38.6226704 | 16.8175105 |
| 17.7390412 | 7.67080158 | 9.71350712 | 24.2304995 | 22.8426233 | 23.8551788 | 7.39970463 |
| 41.3910961 | 20.5323711 | 67.9945498 | 60.5762487 | 67.4044622 | 68.1576537 | 58.7491701 |
| 8.8695206 | 15.3416032 | 19.4270142 | 21.2016871 | 15.7277078 | 13.6315307 | 17.4902109 |
| 2.95650687 | 4.72936639 | 9.71350712 | 4.03841658 | 8.98726163 | 4.54384358 | 9.41780589 |
| 53.2171236 | 58.3673022 | 38.8540285 | 50.4802073 | 46.4341851 | 49.9822794 | 49.7798311 |
| 35.4780824 | 20.4181742 | 9.71350712 | 21.4742802 | 18.4388651 | 14.0859151 | 12.8597897 |
| 5.91301374 | 18.1100128 | 29.1405214 | 13.1248539 | 14.9787694 | 13.6315307 | 16.593277 |
| 5.91301374 | 23.1854303 | 0 | 6.05762487 | 4.49363081 | 13.6315307 | 9.41780589 |
| 5.91301374 | 9.34338237 | 0 | 4.03841658 | 6.74044622 | 11.359609 | 5.60583684 |
| 29.5650687 | 22.3779775 | 38.8540285 | 35.3361451 | 32.2043542 | 21.583257 | 31.8411532 |
| 20.6955481 | 12.8038944 | 19.4270142 | 15.1440622 | 10.4851386 | 18.1753743 | 18.6113783 |
| 5.91301374 | 9.57408317 | 9.71350712 | 12.1152497 | 5.99150775 | 4.54384358 | 8.96933894 |
| 23.6520549 | 27.2226943 | 19.4270142 | 16.1536663 | 10.4851386 | 9.08768716 | 18.8356118 |
| 14.7825343 | 16.9565088 | 9.71350712 | 14.134458 | 9.7362001 | 4.54384358 | 9.41780589 |
| 20.6955481 | 11.9964416 | 38.8540285 | 12.1152497 | 14.9787694 | 17.0394134 | 16.1896568 |
| 5.91301374 | 16.3797568 | 0 | 12.1152497 | 9.7362001 | 10.2236481 | 15.2478762 |
| 59.1301374 | 50.4081247 | 58.2810427 | 62.595457 | 38.9448004 | 52.2542012 | 68.6154429 |
| 56.1736305 | 35.2972223 | 48.5675356 | 40.3841658 | 40.4426773 | 35.2147878 | 48.8828972 |
| 11.8260275 | 26.2998911 | 9.71350712 | 18.6776767 | 19.4724002 | 13.6315307 | 20.7415963 |
| 26.6085618 | 24.2235839 | 9.71350712 | 38.3649575 | 30.7064772 | 19.3113352 | 29.5988185 |
| 322.259249 | 287.453196 | 87.4215641 | 237.256974 | 256.885895 | 224.920257 | 321.102334 |
| 14.5460138 | 9.22457146 | 9.71350712 | 8.31913816 | 24.0559036 | 25.5364009 | 11.7677727 |
| 20.6955481 | 28.7222495 | 38.8540285 | 17.1632705 | 17.2255848 | 17.0394134 | 15.4721097 |
| 5.91301374 | 6.80567358 | 29.1405214 | 13.1248539 | 10.4851386 | 4.54384358 | 10.7632067 |
| 5.91301374 | 3.92191359 | 0 | 3.02881244 | 9.7362001 | 5.67980448 | 4.260436 |
| 23.6520549 | 12.9192448 | 19.4270142 | 15.1440622 | 18.7234617 | 24.9911397 | 16.1448101 |
| 17.7390412 | 14.0727488 | 9.71350712 | 10.0960415 | 8.23832316 | 9.08768716 | 13.2297749 |

| | | | | | | |
|---|---|---|---|---|---|---|
| 2.95650687 | 7.61312638 | 9.71350712 | 4.03841658 | 10.4851386 | 4.54384358 | 5.15736989 |
| 26.6085618 | 10.1508352 | 0 | 20.1920829 | 16.4766463 | 23.8551788 | 18.8356118 |
| 17.7390412 | 49.2546207 | 29.1405214 | 49.4706031 | 38.5703312 | 37.4867095 | 57.2916525 |
| 0 | 4.49866559 | 0 | 7.06722902 | 13.8553617 | 20.4472961 | 6.27853726 |
| 35.4780824 | 19.6095679 | 0 | 31.2977285 | 38.1958619 | 32.942866 | 31.6169198 |
| 20.6955481 | 17.30256 | 9.71350712 | 13.1248539 | 14.9787694 | 28.3990224 | 22.6475808 |
| 47.3041099 | 45.5634079 | 9.71350712 | 66.6338736 | 49.429939 | 54.526123 | 68.166976 |
| 76.8691786 | 33.2209151 | 87.4215641 | 72.6914985 | 74.1449084 | 65.8857319 | 77.3605484 |
| 20.6955481 | 10.8429376 | 9.71350712 | 10.0960415 | 5.99150775 | 13.6315307 | 13.9024754 |
| 20.6955481 | 16.6104576 | 48.5675356 | 29.2785202 | 26.9617849 | 24.9911397 | 29.5988185 |
| 17.7390412 | 18.8021151 | 29.1405214 | 31.2977285 | 26.9617849 | 23.8551788 | 23.3202812 |
| 20.6955481 | 34.7204703 | 9.71350712 | 22.2112912 | 28.4596618 | 17.0394134 | 20.853713 |
| 32.5215755 | 9.11268158 | 9.71350712 | 20.1920829 | 17.9745233 | 11.359609 | 12.781308 |
| 17.7390412 | 18.2253632 | 0 | 9.08643731 | 8.23832316 | 6.81576537 | 12.5570745 |
| 0 | 5.30611839 | 0 | 5.04802073 | 6.74044622 | 18.1753743 | 7.39970463 |
| 0 | 4.84471679 | 9.71350712 | 7.06722902 | 6.74044622 | 10.2236481 | 9.41780589 |
| 26.6085618 | 12.3424928 | 9.71350712 | 29.2785202 | 24.7149695 | 14.7674916 | 29.1503516 |
| 0 | 10.0354848 | 29.1405214 | 7.06722902 | 7.48938469 | 5.67980448 | 5.38160336 |
| 94.6082198 | 90.2040126 | 49.150346 | 48.460999 | 62.1618929 | 53.3901621 | 82.7421517 |
| 14.7825343 | 3.22981119 | 19.4270142 | 7.06722902 | 5.99150775 | 15.9034525 | 9.41780589 |
| 26.6085618 | 31.2599583 | 29.1405214 | 15.1440622 | 14.2298309 | 6.81576537 | 20.4052461 |
| 14.7825343 | 6.69032318 | 9.71350712 | 18.1728746 | 13.4808924 | 6.81576537 | 14.7994093 |
| 13.8364521 | 18.3730117 | 24.2837678 | 25.5530809 | 20.5508716 | 19.2431776 | 17.102287 |
| 23.6520549 | 38.4116831 | 19.4270142 | 28.2689161 | 26.2128464 | 21.583257 | 27.8049507 |
| 5.91301374 | 5.99822078 | 9.71350712 | 4.03841658 | 8.23832316 | 5.67980448 | 5.38160336 |
| 115.303768 | 96.4329341 | 48.5675356 | 81.7779358 | 71.898093 | 55.6620839 | 82.7421517 |
| 2.95650687 | 6.22892158 | 0 | 9.08643731 | 8.98726163 | 5.67980448 | 9.7541561 |
| 115.303768 | 119.617211 | 48.5675356 | 94.9027897 | 86.1279239 | 86.3330281 | 106.735133 |
| 47.3041099 | 37.7195807 | 9.71350712 | 34.326541 | 36.697985 | 23.8551788 | 45.2951617 |
| 29.5650687 | 33.0478895 | 4.85675356 | 31.8025306 | 35.5745773 | 28.9670028 | 34.8683051 |
| 8.8695206 | 6.45962238 | 9.71350712 | 4.03841658 | 8.23832316 | 2.27192179 | 6.50277073 |
| 14.7825343 | 23.6468319 | 19.4270142 | 4.03841658 | 11.234077 | 2.27192179 | 4.260436 |
| 11.8260275 | 15.572304 | 0 | 3.02881244 | 7.48938469 | 6.81576537 | 10.3147398 |
| 0 | 6.69032318 | 0 | 6.05762487 | 0.74893847 | 9.08768716 | 4.48466947 |
| 2.95650687 | 2.76840959 | 0 | 5.04802073 | 8.23832316 | 6.81576537 | 6.95123768 |
| 14.7825343 | 17.1872096 | 77.708057 | 25.2401036 | 18.7234617 | 37.4867095 | 28.7018846 |
| 17.7390412 | 171.756745 | 9.71350712 | 23.2208954 | 25.4639079 | 17.0394134 | 39.4650913 |
| 8.8695206 | 22.1472767 | 29.1405214 | 15.1440622 | 5.24256928 | 9.08768716 | 10.7632067 |
| 7.00692128 | 15.5030937 | 0 | 16.7897169 | 14.1025114 | 19.8565965 | 7.60824176 |
| 20.6955481 | 7.26707518 | 0 | 18.1728746 | 11.9830155 | 30.6709442 | 15.7411898 |
| 50.2606167 | 49.3699711 | 19.4270142 | 17.1632705 | 25.4639079 | 17.0394134 | 27.5807172 |
| 11.8260275 | 18.5714143 | 19.4270142 | 11.1056456 | 16.4766463 | 19.3113352 | 11.4359072 |
| 44.347603 | 38.5270335 | 0 | 44.4225824 | 38.9448004 | 31.8069051 | 51.3494654 |
| 5.91301374 | 16.3797568 | 0 | 25.2401036 | 14.9787694 | 15.9034525 | 15.2478762 |
| 8.8695206 | 7.38242558 | 0 | 15.1440622 | 7.48938469 | 7.95172627 | 8.52087199 |
| 50.2606167 | 58.4249774 | 43.710782 | 55.0234259 | 40.8171466 | 38.05469 | 60.2066876 |

| | | | | | | |
|---|---|---|---|---|---|---|
| 2.95650687 | 12.4578432 | 9.71350712 | 13.1248539 | 14.2298309 | 10.2236481 | 16.1448101 |
| 2.95650687 | 7.61312638 | 9.71350712 | 3.02881244 | 8.23832316 | 6.81576537 | 9.19357241 |
| 11.8260275 | 6.80567358 | 19.4270142 | 19.1824788 | 16.4766463 | 9.08768716 | 14.1267088 |
| 17.7390412 | 8.76663038 | 0 | 15.1440622 | 26.9617849 | 20.4472961 | 11.6601406 |
| 65.0431511 | 33.4516159 | 19.4270142 | 52.4994156 | 55.4214467 | 63.6138101 | 69.5123768 |
| 11.8260275 | 14.0727488 | 9.71350712 | 19.1824788 | 18.7234617 | 15.9034525 | 22.4233474 |
| 8.8695206 | 5.42146879 | 0 | 10.0960415 | 8.23832316 | 12.4955698 | 2.91503516 |
| 56.1736305 | 57.8482254 | 24.2837678 | 35.6693145 | 37.4469235 | 34.6468073 | 48.2101968 |
| 47.3041099 | 25.2617375 | 19.4270142 | 26.2497078 | 17.2255848 | 24.9911397 | 32.0653867 |
| 5.91301374 | 7.38242558 | 0 | 11.1056456 | 5.24256928 | 3.40788269 | 8.29663852 |
| 192.172946 | 254.116931 | 174.843128 | 239.276183 | 194.724002 | 206.744883 | 211.003699 |
| 20.6955481 | 7.61312638 | 0 | 17.1632705 | 20.2213387 | 21.583257 | 14.5751758 |
| 23.6520549 | 4.26796479 | 29.1405214 | 16.1536663 | 19.4724002 | 13.6315307 | 16.3690436 |
| 17.7390412 | 11.9964416 | 0 | 15.1440622 | 17.2255848 | 19.3113352 | 8.29663852 |
| 147.825343 | 184.906691 | 165.129621 | 241.295391 | 158.026017 | 173.802017 | 184.992616 |
| 8.8695206 | 29.4143519 | 9.71350712 | 13.1248539 | 9.7362001 | 15.9034525 | 15.4721097 |
| 29.5650687 | 62.7506174 | 38.8540285 | 49.4706031 | 47.932062 | 39.7586313 | 66.1488747 |
| 13.3042809 | 9.68943357 | 4.85675356 | 20.1920829 | 12.3574847 | 21.0152766 | 14.5751758 |
| 8.8695206 | 6.57497278 | 0 | 8.07683317 | 5.99150775 | 10.2236481 | 6.95123768 |
| 2.95650687 | 13.3806464 | 0 | 11.1056456 | 9.7362001 | 7.95172627 | 7.73605484 |
| 0 | 8.53592958 | 9.71350712 | 6.05762487 | 9.7362001 | 11.359609 | 7.39970463 |
| 14.7825343 | 4.38331519 | 48.5675356 | 29.2785202 | 16.4766463 | 9.08768716 | 17.041744 |
| 5.91301374 | 5.88287038 | 0 | 7.06722902 | 7.48938469 | 11.359609 | 5.38160336 |
| 8.8695206 | 4.96006719 | 9.71350712 | 6.05762487 | 8.98726163 | 4.54384358 | 10.7632067 |
| 29.5650687 | 30.2218047 | 19.4270142 | 11.1056456 | 14.2298309 | 9.08768716 | 16.3690436 |
| 19.7790309 | 14.9413373 | 4.3710782 | 15.3055988 | 27.2688497 | 39.361045 | 17.5529963 |
| 20.6955481 | 29.9911039 | 19.4270142 | 22.2112912 | 26.2128464 | 28.3990224 | 34.7561884 |
| 11.8260275 | 7.72847678 | 9.71350712 | 8.07683317 | 7.48938469 | 9.08768716 | 10.3147398 |
| 20.6955481 | 5.30611839 | 9.71350712 | 16.1536663 | 15.7277078 | 6.81576537 | 14.3509423 |
| 26.6085618 | 13.4959968 | 9.71350712 | 16.1536663 | 16.4766463 | 17.0394134 | 17.4902109 |
| 5.91301374 | 5.30611839 | 0 | 17.1632705 | 5.99150775 | 14.7674916 | 14.3509423 |
| 5.91301374 | 4.84471679 | 0 | 13.1248539 | 8.23832316 | 6.81576537 | 9.19357241 |
| 26.6085618 | 28.9529503 | 9.71350712 | 16.1536663 | 16.4766463 | 10.2236481 | 26.4595499 |
| 19.2172946 | 8.68011758 | 0 | 20.1920829 | 17.9745233 | 14.1995112 | 13.9024754 |
| 141.91233 | 283.877334 | 97.1350712 | 137.306164 | 116.834401 | 86.3330281 | 134.540084 |
| 5.91301374 | 15.2262528 | 9.71350712 | 9.08643731 | 14.2298309 | 10.2236481 | 10.0905063 |
| 14.7825343 | 22.1472767 | 29.1405214 | 12.1152497 | 22.4681541 | 10.2236481 | 13.0055415 |
| 23.6520549 | 22.7240287 | 9.71350712 | 9.08643731 | 8.23832316 | 10.2236481 | 10.0905063 |
| 26.6085618 | 25.1463871 | 0 | 18.1728746 | 20.2213387 | 24.9911397 | 22.8718143 |
| 5.91301374 | 5.19076799 | 0 | 3.02881244 | 8.23832316 | 4.54384358 | 5.38160336 |
| 26.6085618 | 14.7648512 | 19.4270142 | 46.4417907 | 47.1831235 | 31.8069051 | 41.4831926 |
| 35.4780824 | 49.2546207 | 9.71350712 | 48.460999 | 30.7064772 | 22.7192179 | 31.8411532 |
| 2.95650687 | 4.49866559 | 0 | 2.01920829 | 8.98726163 | 7.95172627 | 6.95123768 |
| 11.8260275 | 15.6876544 | 9.71350712 | 8.07683317 | 11.9830155 | 11.359609 | 15.0236427 |
| 29.5650687 | 25.7231391 | 0 | 21.2016871 | 16.4766463 | 30.6709442 | 21.30218 |
| 47.3041099 | 63.4427198 | 58.2810427 | 42.4033741 | 38.9448004 | 30.6709442 | 47.7617299 |

| | | | | | | |
|---|---|---|---|---|---|---|
| 65.0431511 | 86.9742014 | 87.4215641 | 80.7683317 | 69.6512776 | 65.8857319 | 97.7657945 |
| 17.7390412 | 6.92102398 | 9.71350712 | 35.3361451 | 31.4554157 | 24.9911397 | 29.1503516 |
| 133.042809 | 76.3619646 | 67.9945498 | 111.056456 | 150.536632 | 118.139933 | 132.521983 |
| 11.8260275 | 12.4578432 | 9.71350712 | 17.1632705 | 11.9830155 | 12.4955698 | 10.5389733 |
| 5.91301374 | 51.6769791 | 0 | 18.1728746 | 18.7234617 | 17.0394134 | 19.5083122 |
| 14.7825343 | 18.6867647 | 9.71350712 | 16.1536663 | 8.98726163 | 15.9034525 | 8.74510547 |
| 2.95650687 | 19.9556191 | 0 | 3.02881244 | 5.99150775 | 13.6315307 | 6.50277073 |
| 8.8695206 | 13.1499456 | 9.71350712 | 16.1536663 | 11.234077 | 14.7674916 | 18.8356118 |
| 17.7390412 | 18.3407136 | 9.71350712 | 16.1536663 | 14.9787694 | 15.9034525 | 15.2478762 |
| 11.8260275 | 22.3779775 | 19.4270142 | 21.2016871 | 16.4766463 | 21.583257 | 19.2840787 |
| 23.6520549 | 38.0079567 | 0 | 16.1536663 | 15.7277078 | 24.9911397 | 21.0779465 |
| 5.91301374 | 12.2271424 | 0 | 8.07683317 | 9.7362001 | 4.54384358 | 7.17547115 |
| 2.4539007 | 6.55536321 | 0 | 10.2676742 | 10.9869273 | 14.0063978 | 5.71795358 |
| 56.1736305 | 48.1011167 | 58.2810427 | 60.5762487 | 60.664016 | 55.6620839 | 72.6516454 |
| 41.3910961 | 29.0683007 | 29.1405214 | 17.1632705 | 20.2213387 | 17.0394134 | 23.3202812 |
| 32.5215755 | 45.7941087 | 19.4270142 | 14.6392601 | 11.234077 | 20.4472961 | 28.0291842 |
| 14.7825343 | 27.5687455 | 9.71350712 | 24.2304995 | 16.4766463 | 19.3113352 | 17.7144444 |
| 72.4344183 | 21.3974991 | 14.5702607 | 9.08643731 | 30.332008 | 13.6315307 | 30.1594022 |
| 11.8260275 | 14.3034496 | 38.8540285 | 16.1536663 | 12.731954 | 21.583257 | 19.0598453 |
| 17.7390412 | 5.99822078 | 0 | 5.04802073 | 7.48938469 | 4.54384358 | 6.27853726 |
| 14.7825343 | 7.49777598 | 0 | 15.1440622 | 20.9702771 | 19.3113352 | 11.6601406 |
| 38.4345893 | 23.9928831 | 19.4270142 | 34.326541 | 29.2086003 | 31.8069051 | 32.9623206 |
| 5.91301374 | 9.11268158 | 9.71350712 | 1.00960415 | 11.234077 | 6.81576537 | 6.05430379 |
| 28.5894214 | 20.7630719 | 12.9189645 | 20.3637156 | 22.5954736 | 21.2083899 | 21.7887666 |
| 17.7390412 | 17.1872096 | 0 | 9.08643731 | 5.24256928 | 11.359609 | 6.05430379 |
| 17.7390412 | 27.9147967 | 9.71350712 | 9.08643731 | 8.98726163 | 12.4955698 | 12.781308 |
| 2.95650687 | 4.15261439 | 9.71350712 | 7.06722902 | 5.24256928 | 10.2236481 | 7.6239381 |
| 32.5215755 | 48.9085695 | 19.4270142 | 40.3841658 | 46.4341851 | 38.6226704 | 52.9190998 |
| 8.8695206 | 10.0354848 | 0 | 7.06722902 | 8.23832316 | 10.2236481 | 4.93313642 |
| 17.7390412 | 10.381536 | 9.71350712 | 7.06722902 | 5.24256928 | 9.08768716 | 7.6239381 |
| 8.8695206 | 8.18987838 | 0 | 10.0960415 | 12.731954 | 4.54384358 | 13.4540084 |
| 29.5650687 | 23.6468319 | 9.71350712 | 22.2112912 | 18.7234617 | 22.7192179 | 23.3202812 |
| 8.8695206 | 26.2998911 | 29.1405214 | 14.134458 | 13.4808924 | 18.1753743 | 13.9024754 |
| 11.8260275 | 11.0736384 | 19.4270142 | 10.0960415 | 11.9830155 | 21.583257 | 17.9386779 |
| 76.8691786 | 75.1507854 | 19.4270142 | 82.7875399 | 75.6427854 | 71.5655364 | 95.7476932 |
| 23.6520549 | 15.572304 | 29.1405214 | 28.2689161 | 19.4724002 | 11.359609 | 17.4902109 |
| 2.95650687 | 1.7879312 | 0 | 4.03841658 | 2.24681541 | 10.2236481 | 4.93313642 |
| 41.3910961 | 45.4480575 | 19.4270142 | 53.5090197 | 43.4384312 | 40.8945922 | 53.1433332 |
| 118.260275 | 80.3992286 | 116.562085 | 123.171706 | 125.072724 | 82.9251454 | 113.910605 |
| 11.8260275 | 13.842048 | 9.71350712 | 12.1152497 | 18.7234617 | 7.95172627 | 15.2478762 |
| 11.8260275 | 5.65216958 | 9.71350712 | 3.02881244 | 5.24256928 | 11.359609 | 8.29663852 |
| 5.91301374 | 21.4551743 | 38.8540285 | 33.3169368 | 23.2170925 | 35.7827682 | 5.38160336 |
| 17.7390412 | 21.3398239 | 0 | 14.6392601 | 11.9830155 | 12.4955698 | 14.0145921 |
| 5.91301374 | 2.07630719 | 0 | 1.00960415 | 7.48938469 | 4.54384358 | 4.70890294 |
| 35.4780824 | 21.9165759 | 48.5675356 | 40.3841658 | 25.4639079 | 31.8069051 | 29.374585 |
| 11.8260275 | 16.8411584 | 9.71350712 | 20.1920829 | 12.731954 | 32.942866 | 16.1448101 |

| | | | | | | |
|---|---|---|---|---|---|---|
| 144.868837 | 102.315805 | 135.9891 | 127.210122 | 124.323786 | 112.460129 | 132.97045 |
| 2.95650687 | 11.53504 | 9.71350712 | 7.06722902 | 10.4851386 | 11.359609 | 6.72700421 |
| 5.91301374 | 12.2271424 | 0 | 9.08643731 | 7.48938469 | 6.81576537 | 6.95123768 |
| 38.4345893 | 46.0248095 | 29.1405214 | 29.2785202 | 70.4002161 | 36.3507487 | 50.2282981 |
| 14.7825343 | 9.80478397 | 0 | 5.04802073 | 9.7362001 | 15.9034525 | 7.84817157 |
| 2.95650687 | 1.153504 | 0 | 4.03841658 | 7.48938469 | 9.08768716 | 6.27853726 |
| 8.8695206 | 5.88287038 | 9.71350712 | 4.03841658 | 12.731954 | 13.6315307 | 12.781308 |
| 5.91301374 | 4.15261439 | 0 | 8.07683317 | 9.7362001 | 4.54384358 | 8.96933894 |
| 106.434247 | 53.0611839 | 97.1350712 | 82.7875399 | 84.630047 | 67.0216928 | 94.6265258 |
| 8.8695206 | 10.09316 | 9.71350712 | 5.04802073 | 2.99575388 | 9.08768716 | 9.19357241 |
| 29.5650687 | 21.1091231 | 19.4270142 | 41.898572 | 31.0809465 | 25.5591201 | 32.6259704 |
| 11.8260275 | 22.9547295 | 9.71350712 | 7.06722902 | 10.4851386 | 11.359609 | 14.5751758 |
| 16.2607878 | 10.2085104 | 14.5702607 | 9.59123938 | 13.7280421 | 13.2566636 | 15.2097565 |
| 11.8260275 | 22.6086783 | 19.4270142 | 12.1152497 | 8.98726163 | 10.7916285 | 10.7632067 |
| 13.3042809 | 20.7053967 | 67.9945498 | 24.7353016 | 26.2128464 | 26.1271006 | 25.7868495 |
| 0 | 8.42057918 | 0 | 9.08643731 | 15.7277078 | 15.9034525 | 14.7994093 |
| 20.6955481 | 13.9573984 | 0 | 28.2689161 | 18.7234617 | 30.6709442 | 13.2297749 |
| 35.4780824 | 19.2635167 | 19.4270142 | 22.2112912 | 17.9745233 | 17.0394134 | 23.3202812 |
| 8.8695206 | 11.3043392 | 9.71350712 | 11.1056456 | 17.2255848 | 17.0394134 | 15.9205766 |
| 11.8260275 | 4.72936639 | 9.71350712 | 7.06722902 | 13.4808924 | 9.08768716 | 15.0236427 |
| 830.77843 | 619.200946 | 1223.9019 | 885.422836 | 698.010653 | 572.524291 | 832.915237 |
| 14.7825343 | 10.1508352 | 0 | 13.1248539 | 9.7362001 | 6.81576537 | 10.3147398 |
| 29.5650687 | 40.8340415 | 29.1405214 | 31.2977285 | 43.4384312 | 32.942866 | 46.416329 |
| 8.8695206 | 12.3424928 | 0 | 8.07683317 | 19.4724002 | 11.359609 | 15.2478762 |
| 14.7825343 | 51.4462783 | 9.71350712 | 17.1632705 | 27.7107234 | 21.583257 | 22.4233474 |
| 11.8260275 | 5.99822078 | 0 | 16.1536663 | 15.7277078 | 15.9034525 | 13.6782419 |
| 8.8695206 | 18.8021151 | 38.8540285 | 35.3361451 | 23.2170925 | 18.1753743 | 22.8718143 |
| 4.4347603 | 7.09404958 | 9.71350712 | 9.59123938 | 7.86385392 | 17.6073939 | 4.93313642 |
| 5.91301374 | 9.34338237 | 9.71350712 | 17.1632705 | 16.4766463 | 11.359609 | 13.4921281 |
| 14.7825343 | 24.1082335 | 9.71350712 | 24.2304995 | 22.4681541 | 15.9034525 | 27.5807172 |
| 17.7390412 | 7.61312638 | 0 | 6.05762487 | 18.7234617 | 15.9034525 | 8.74510547 |
| 14.7825343 | 2.99911039 | 9.71350712 | 11.1056456 | 11.9830155 | 9.08768716 | 10.3147398 |
| 25.1303084 | 22.0123168 | 22.6324716 | 22.2920595 | 28.2724272 | 12.6886832 | 20.4792431 |
| 2.95650687 | 12.2271424 | 0 | 8.07683317 | 9.7362001 | 7.95172627 | 9.19357241 |
| 11.8260275 | 9.80478397 | 0 | 3.02881244 | 8.98726163 | 9.08768716 | 8.07240505 |
| 8.8695206 | 11.0736384 | 9.71350712 | 12.1152497 | 7.48938469 | 7.95172627 | 8.96933894 |
| 25.1303084 | 15.4569536 | 0 | 34.326541 | 20.9702771 | 19.3113352 | 29.4867018 |
| 29.5650687 | 33.2578272 | 0 | 26.3304761 | 27.2089346 | 30.1029637 | 36.9043451 |
| 67.999658 | 73.1321534 | 9.71350712 | 61.5858529 | 50.1788774 | 60.2059275 | 79.3786496 |
| 112.347261 | 101.739053 | 67.9945498 | 98.9412063 | 77.8896008 | 84.0611063 | 83.4148522 |
| 26.6085618 | 22.9547295 | 29.1405214 | 21.2016871 | 23.2170925 | 14.7674916 | 21.9748804 |
| 2.95650687 | 3.22981119 | 9.71350712 | 10.0960415 | 9.7362001 | 4.54384358 | 5.38160336 |
| 5.91301374 | 3.80656319 | 0 | 11.1056456 | 8.98726163 | 10.2236481 | 7.84817157 |
| 59.1301374 | 46.3708607 | 9.71350712 | 36.3457492 | 37.4469235 | 39.7586313 | 48.6586638 |
| 2.95650687 | 12.7266096 | 0 | 7.57203109 | 18.3489925 | 14.1995112 | 7.51182136 |
| 11.8260275 | 11.3043392 | 0 | 20.1920829 | 14.2298309 | 11.359609 | 21.5264135 |

| | | | | | | |
|---|---|---|---|---|---|---|
| 4.4347603 | 9.92013437 | 9.71350712 | 4.03841658 | 6.74044622 | 10.2236481 | 6.72700421 |
| 5.91301374 | 11.0736384 | 38.8540285 | 11.1056456 | 14.9787694 | 3.40788269 | 8.07240505 |
| 124.173288 | 92.6644366 | 135.9891 | 108.027644 | 118.332278 | 98.8285979 | 102.02623 |
| 14.7825343 | 11.9964416 | 19.4270142 | 18.1728746 | 14.2298309 | 20.4472961 | 20.1810126 |
| 5.91301374 | 4.49866559 | 9.71350712 | 5.04802073 | 8.98726163 | 6.81576537 | 3.13926863 |
| 5.91301374 | 4.03726399 | 0 | 7.06722902 | 7.48938469 | 3.40788269 | 5.38160336 |
| 20.6955481 | 20.7630719 | 0 | 27.2593119 | 21.7192156 | 22.7192179 | 26.0110829 |
| 29.5650687 | 47.6397151 | 19.4270142 | 26.2497078 | 19.4724002 | 14.7674916 | 24.8899156 |
| 17.7390412 | 13.3806464 | 9.71350712 | 21.2016871 | 17.2255848 | 13.6315307 | 23.7687482 |
| 5.91301374 | 1.6149056 | 9.71350712 | 7.06722902 | 8.98726163 | 13.6315307 | 7.39970463 |
| 11.8260275 | 5.82519518 | 19.4270142 | 4.54321866 | 8.23832316 | 7.95172627 | 10.5389733 |
| 8.8695206 | 15.6876544 | 0 | 4.03841658 | 9.7362001 | 7.95172627 | 6.50277073 |
| 10.0521233 | 12.4013215 | 0 | 12.2969785 | 12.499783 | 13.0635503 | 8.39530125 |
| 17.7390412 | 7.84382718 | 0 | 21.2016871 | 17.9745233 | 17.0394134 | 16.1448101 |
| 20.6955481 | 7.38242558 | 9.71350712 | 8.07683317 | 19.4724002 | 12.4955698 | 14.7994093 |
| 36.4537297 | 28.70264 | 58.2810427 | 63.6050612 | 46.8086543 | 71.7586498 | 56.9171826 |
| 14.7825343 | 46.1401599 | 9.71350712 | 22.2112912 | 20.9702771 | 29.5349833 | 11.7722574 |
| 11.8260275 | 11.0736384 | 0 | 19.1824788 | 18.7234617 | 14.7674916 | 23.5445147 |
| 2.95650687 | 4.15261439 | 0 | 7.06722902 | 3.74469235 | 9.08768716 | 7.84817157 |
| 14.7825343 | 13.1499456 | 0 | 8.07683317 | 15.7277078 | 9.08768716 | 10.0905063 |
| 8.8695206 | 13.3806464 | 9.71350712 | 11.1056456 | 7.48938469 | 11.359609 | 10.0905063 |
| 8.8695206 | 17.590936 | 19.4270142 | 10.6008435 | 13.8553617 | 14.1995112 | 15.6963431 |
| 2.95650687 | 2.99911039 | 0 | 2.01920829 | 10.4851386 | 5.67980448 | 16.1448101 |
| 5.91301374 | 3.11446079 | 0 | 8.07683317 | 8.23832316 | 6.81576537 | 8.96933894 |
| 10.347774 | 1.96095679 | 4.85675356 | 11.6104477 | 12.3574847 | 10.7916285 | 6.61488747 |
| 50.2606167 | 44.0638527 | 48.5675356 | 50.4802073 | 35.200108 | 51.1182403 | 47.7617299 |
| 8.8695206 | 11.9964416 | 0 | 14.134458 | 11.234077 | 6.81576537 | 12.1086076 |
| 11.8260275 | 8.65127998 | 0 | 14.134458 | 5.99150775 | 12.4955698 | 13.4540084 |
| 11.8260275 | 20.6477215 | 9.71350712 | 7.57203109 | 7.48938469 | 10.2236481 | 12.332841 |
| 17.7390412 | 4.15261439 | 0 | 8.07683317 | 11.9830155 | 9.08768716 | 12.5570745 |
| 35.4780824 | 75.2084606 | 29.1405214 | 54.5186239 | 37.4469235 | 43.166514 | 40.3620252 |
| 2.95650687 | 6.34427198 | 9.71350712 | 18.1728746 | 14.2298309 | 15.9034525 | 13.0055415 |
| 322.259249 | 196.441731 | 194.270142 | 260.47787 | 285.345557 | 226.056218 | 272.44367 |
| 6.888661 | 12.8615696 | 0 | 13.4580233 | 8.48547285 | 3.78274978 | 8.63971574 |
| 20.6955481 | 23.0700799 | 0 | 8.07683317 | 15.7277078 | 20.4472961 | 17.4902109 |
| 23.6520549 | 26.6459423 | 19.4270142 | 29.2785202 | 26.2128464 | 19.3113352 | 24.6656821 |
| 35.4780824 | 17.7639616 | 19.4270142 | 25.2401036 | 35.9490465 | 28.3990224 | 29.1503516 |
| 76.8691786 | 36.1046751 | 58.2810427 | 42.4033741 | 51.6767544 | 52.2542012 | 47.3132629 |
| 14.7825343 | 14.995552 | 0 | 8.07683317 | 11.9830155 | 12.4955698 | 10.0905063 |
| 17.7390412 | 30.9139071 | 0 | 16.1536663 | 22.4681541 | 22.7192179 | 17.7144444 |
| 8.8695206 | 5.07541759 | 0 | 5.04802073 | 7.48938469 | 6.81576537 | 8.74510547 |
| 2.95650687 | 6.22892158 | 0 | 6.05762487 | 7.48938469 | 1.1359609 | 5.38160336 |
| 11.8260275 | 31.4906591 | 19.4270142 | 29.2785202 | 26.2128464 | 27.2630615 | 31.1684528 |
| 11.8260275 | 30.3371551 | 19.4270142 | 23.7256974 | 25.8608453 | 23.6620655 | 14.0885891 |
| 35.4780824 | 22.362982 | 12.9189645 | 45.9369886 | 42.8767274 | 42.5985336 | 41.4405882 |
| 2.95650687 | 29.4143519 | 0 | 13.1248539 | 18.7234617 | 15.9034525 | 18.6113783 |

| | | | | | | |
|---|---|---|---|---|---|---|
| 2.95650687 | 6.11357118 | 0 | 3.02881244 | 9.7362001 | 6.81576537 | 4.93313642 |
| 2.95650687 | 20.9937727 | 9.71350712 | 14.134458 | 9.7362001 | 23.8551788 | 7.39970463 |
| 0 | 10.7275872 | 0 | 4.03841658 | 7.48938469 | 17.0394134 | 3.47561884 |
| 14.7825343 | 12.5731936 | 9.71350712 | 9.08643731 | 10.4851386 | 18.1753743 | 13.9024754 |
| 23.6520549 | 22.2626271 | 29.1405214 | 21.2016871 | 19.4724002 | 12.4955698 | 15.9205766 |
| 26.6085618 | 20.1863199 | 19.4270142 | 27.2593119 | 20.2213387 | 9.08768716 | 24.4414486 |
| 14.7825343 | 10.6122368 | 19.4270142 | 19.1824788 | 14.9787694 | 23.8551788 | 15.9205766 |
| 3.93215413 | 7.56583272 | 0 | 8.70278774 | 7.04751099 | 11.5754415 | 3.12805696 |
| 10.9982055 | 8.8300731 | 19.4270142 | 9.02586106 | 8.74760132 | 17.3461229 | 6.68664218 |
| 14.2799282 | 9.37221997 | 0 | 20.1113146 | 14.6642152 | 32.9087871 | 12.4785928 |
| 5.91301374 | 3.80656319 | 0 | 6.05762487 | 15.7277078 | 6.81576537 | 10.3147398 |
| 8.8695206 | 2.53770879 | 0 | 23.2208954 | 23.2170925 | 14.7674916 | 17.2659775 |
| 2.95650687 | 10.09316 | 19.4270142 | 7.06722902 | 12.731954 | 11.9275894 | 9.19357241 |
| 17.7390412 | 15.4569536 | 9.71350712 | 19.1824788 | 13.4808924 | 17.0394134 | 13.6782419 |
| 14.7825343 | 11.1889888 | 0 | 5.04802073 | 9.7362001 | 7.95172627 | 11.2116737 |
| 177.390412 | 88.012355 | 106.848578 | 114.085268 | 116.085463 | 114.73205 | 155.618031 |
| 8.8695206 | 7.15172478 | 29.1405214 | 16.1536663 | 9.7362001 | 13.6315307 | 9.19357241 |
| 14.7825343 | 4.38331519 | 9.71350712 | 7.57203109 | 8.98726163 | 6.81576537 | 8.52087199 |
| 8.8695206 | 8.42057918 | 9.71350712 | 9.08643731 | 9.7362001 | 9.08768716 | 10.9874402 |
| 65.0431511 | 101.508352 | 58.2810427 | 56.5378322 | 71.1491546 | 60.2059275 | 68.6154429 |
| 21.937281 | 14.6298912 | 22.6324716 | 13.3570628 | 16.723796 | 19.6862023 | 20.5061512 |
| 5.91301374 | 13.418712 | 9.71350712 | 19.8589135 | 12.5447194 | 18.1753743 | 7.81005188 |
| 29.5650687 | 28.6068991 | 9.71350712 | 29.2785202 | 29.2086003 | 37.4867095 | 37.6712236 |
| 5.91301374 | 9.57408317 | 0 | 8.07683317 | 11.234077 | 14.7674916 | 8.74510547 |
| 2.95650687 | 4.26796479 | 19.4270142 | 5.04802073 | 5.24256928 | 9.08768716 | 6.05430379 |
| 23.6520549 | 23.1854303 | 29.1405214 | 27.5924813 | 25.4639079 | 27.2630615 | 31.3926863 |
| 11.8260275 | 8.99733118 | 0 | 1.00960415 | 10.4851386 | 4.54384358 | 6.50277073 |
| 2.95650687 | 5.99822078 | 9.71350712 | 5.04802073 | 8.98726163 | 3.40788269 | 6.05430379 |
| 5.91301374 | 3.64276562 | 9.71350712 | 27.5420011 | 24.8048421 | 18.1753743 | 12.3709607 |
| 14.7825343 | 17.4179104 | 0 | 16.1536663 | 18.7234617 | 18.7433548 | 17.041744 |
| 11.8260275 | 12.8038944 | 19.4270142 | 11.1056456 | 6.74044622 | 4.54384358 | 8.74510547 |
| 8.8695206 | 20.9937727 | 9.71350712 | 27.2593119 | 14.2298309 | 18.1753743 | 26.4595499 |
| 8.8695206 | 32.5288127 | 0 | 9.08643731 | 8.23832316 | 11.359609 | 13.4540084 |
| 10.347774 | 8.93965598 | 19.4270142 | 11.6104477 | 14.2298309 | 21.583257 | 14.911526 |
| 15.2851405 | 23.7852524 | 29.1405214 | 7.06722902 | 15.1060889 | 18.1753743 | 13.5661251 |
| 26.6085618 | 40.8340415 | 19.4270142 | 25.2401036 | 23.2170925 | 19.3113352 | 29.5988185 |
| 11.8260275 | 11.4196896 | 29.1405214 | 6.05762487 | 16.4766463 | 15.9034525 | 20.6294796 |
| 10.347774 | 9.80478397 | 0 | 8.07683317 | 6.74044622 | 16.471433 | 7.77417453 |
| 44.347603 | 15.572304 | 38.8540285 | 47.4513948 | 47.932062 | 36.3507487 | 41.4831926 |
| 73.9126717 | 113.620144 | 29.1405214 | 83.7971441 | 79.3874777 | 71.5655364 | 97.7657945 |
| 109.390754 | 47.7550655 | 29.1405214 | 61.5858529 | 58.4172006 | 68.1576537 | 73.5485793 |
| 8.8695206 | 17.6486112 | 0 | 11.1056456 | 12.731954 | 11.359609 | 19.2840787 |
| 94.6082198 | 70.7294046 | 72.8513034 | 114.085268 | 96.6130625 | 69.2936146 | 84.4239028 |
| 38.4345893 | 22.9547295 | 0 | 31.2977285 | 26.9617849 | 31.8069051 | 27.8049507 |
| 26.6085618 | 13.6113472 | 0 | 13.1248539 | 16.4766463 | 28.9670028 | 15.4721097 |
| 18.3303426 | 23.6906651 | 12.1418839 | 25.4117363 | 36.2636007 | 61.3759672 | 16.6941821 |

| | | | | | | |
|---|---|---|---|---|---|---|
| 8.8695206 | 3.11446079 | 9.71350712 | 4.03841658 | 2.99575388 | 9.08768716 | 5.83007031 |
| 43.3719557 | 19.2542887 | 4.85675356 | 54.0138218 | 47.932062 | 55.6961627 | 26.2061661 |
| 17.5025207 | 9.04577834 | 9.71350712 | 26.3304761 | 24.6101181 | 23.7983808 | 20.8492284 |
| 32.5215755 | 17.879312 | 29.1405214 | 14.134458 | 34.4511696 | 20.4472961 | 28.4776511 |
| 17.7390412 | 21.2244735 | 19.4270142 | 31.2977285 | 17.2255848 | 20.4472961 | 26.4595499 |
| 17.7390412 | 13.265296 | 0 | 9.08643731 | 8.23832316 | 6.81576537 | 8.74510547 |
| 14.7825343 | 24.8003359 | 9.71350712 | 11.1056456 | 16.4766463 | 11.359609 | 16.1448101 |
| 20.6955481 | 17.879312 | 0 | 13.1248539 | 17.9745233 | 13.6315307 | 17.4902109 |
| 2.12868494 | 7.19209742 | 2.42837678 | 11.2873743 | 7.50436346 | 7.37238621 | 5.35693768 |
| 47.3041099 | 13.3806464 | 38.8540285 | 25.2401036 | 32.2043542 | 22.7192179 | 36.7742897 |
| 17.7390412 | 10.2661856 | 9.71350712 | 22.2112912 | 17.9745233 | 22.7192179 | 10.9874402 |
| 8.8695206 | 25.1463871 | 19.4270142 | 23.2208954 | 18.7234617 | 15.9034525 | 22.1991139 |
| 63.3283771 | 54.823738 | 40.5053247 | 70.2482565 | 54.1707195 | 56.2300643 | 47.1607842 |
| 8.8695206 | 22.3779775 | 9.71350712 | 13.1248539 | 5.24256928 | 7.95172627 | 13.2297749 |
| 41.3910961 | 25.2617375 | 38.8540285 | 27.2593119 | 20.2213387 | 21.583257 | 32.9623206 |
| 239.477056 | 305.447858 | 223.410664 | 217.064891 | 180.494171 | 143.131073 | 234.099746 |
| 130.086302 | 73.3628542 | 72.8513034 | 109.54205 | 111.719151 | 92.0128325 | 119.628558 |
| 8.8695206 | 8.07452798 | 19.4270142 | 4.03841658 | 8.98726163 | 15.9034525 | 6.95123768 |
| 76.8691786 | 58.021251 | 58.2810427 | 51.4898114 | 47.1831235 | 46.5743967 | 58.7491701 |
| 11.8260275 | 4.16530293 | 9.71350712 | 8.07683317 | 9.7362001 | 11.359609 | 11.9494018 |
| 8.8695206 | 6.11357118 | 0 | 5.04802073 | 8.23832316 | 12.4955698 | 9.86627284 |
| 47.3041099 | 57.9059006 | 19.4270142 | 35.3361451 | 21.7192156 | 20.4472961 | 32.0653867 |
| 7.39126717 | 10.0354848 | 19.4270142 | 21.2016871 | 15.2259191 | 11.359609 | 10.2586814 |
| 29.5650687 | 62.8659678 | 19.4270142 | 18.1728746 | 17.2255848 | 10.2236481 | 24.8899156 |
| 59.1301374 | 78.8996734 | 19.4270142 | 62.595457 | 52.4256928 | 52.2542012 | 63.2338395 |
| 5.91301374 | 3.80656319 | 0 | 10.0960415 | 7.48938469 | 4.54384358 | 5.83007031 |
| 11.8260275 | 14.5341504 | 0 | 9.08643731 | 10.4851386 | 9.08768716 | 15.0236427 |
| 5.41040757 | 11.0621033 | 1.65129621 | 13.1248539 | 22.5205798 | 18.6865567 | 9.36623219 |
| 12.8016747 | 55.5066123 | 9.71350712 | 15.1440622 | 17.4727345 | 10.2236481 | 16.499099 |
| 103.47774 | 125.501235 | 77.708057 | 111.056456 | 126.570601 | 97.692637 | 120.413375 |
| 11.8260275 | 27.6840959 | 29.1405214 | 11.1056456 | 22.4681541 | 11.359609 | 27.1322503 |
| 23.6520549 | 8.99733118 | 0 | 6.05762487 | 12.731954 | 10.2236481 | 10.9874402 |
| 1.47825343 | 7.55545118 | 0 | 9.59123938 | 10.4851386 | 11.359609 | 3.92408579 |
| 20.6955481 | 31.3753087 | 0 | 11.1056456 | 13.4808924 | 6.81576537 | 14.3509423 |
| 10.347774 | 8.59360478 | 0 | 7.06722902 | 9.7362001 | 17.0394134 | 10.65109 |
| 5.91301374 | 4.15261439 | 0 | 2.01920829 | 7.48938469 | 2.27192179 | 5.15736989 |
| 4.4347603 | 3.36477116 | 9.71350712 | 12.9532212 | 15.8550274 | 18.9364681 | 5.26948663 |
| 14.7825343 | 6.34427198 | 9.71350712 | 11.1056456 | 15.7277078 | 14.7674916 | 11.6601406 |
| 4.4347603 | 2.76840959 | 0 | 8.07683317 | 7.86385392 | 8.51970672 | 4.93313642 |
| 0 | 14.7648512 | 0 | 3.02881244 | 4.49363081 | 9.08768716 | 8.29663852 |
| 5.91301374 | 10.0354848 | 9.71350712 | 12.6200518 | 11.9830155 | 11.9275894 | 10.9874402 |
| 807.126375 | 586.441432 | 689.659005 | 718.838152 | 673.295684 | 589.563705 | 699.608437 |
| 20.6955481 | 79.0150238 | 19.4270142 | 20.1920829 | 20.9702771 | 14.7674916 | 26.3474331 |
| 107.79424 | 40.7948224 | 50.510237 | 84.9783809 | 64.5734748 | 62.3301743 | 55.6637175 |
| 2.95650687 | 6.69032318 | 0 | 8.07683317 | 7.48938469 | 5.67980448 | 4.93313642 |
| 0 | 10.1508352 | 0 | 8.07683317 | 12.731954 | 7.95172627 | 9.41780589 |

| | | | | | | |
|---|---|---|---|---|---|---|
| 67.999658 | 94.5873277 | 29.1405214 | 64.6146653 | 59.9150775 | 39.7586313 | 67.4942755 |
| 2.95650687 | 5.42146879 | 19.4270142 | 13.1248539 | 12.731954 | 15.9034525 | 10.65109 |
| 5.91301374 | 12.688544 | 0 | 11.1056456 | 5.24256928 | 4.54384358 | 9.64203936 |
| 2.95650687 | 5.88287038 | 0 | 6.05762487 | 5.24256928 | 10.2236481 | 6.72700421 |
| 5.91301374 | 24.9156863 | 0 | 12.1152497 | 12.731954 | 7.95172627 | 13.0055415 |
| 0 | 4.61401599 | 0 | 2.01920829 | 7.48938469 | 2.27192179 | 3.81196905 |
| 9.72690759 | 15.2343273 | 9.71350712 | 14.4272432 | 22.2284938 | 17.8573053 | 19.9657485 |
| 30.15637 | 20.4470118 | 20.7869052 | 38.728415 | 34.4137227 | 36.0894776 | 17.6695977 |
| 2.95650687 | 4.26796479 | 0 | 5.04802073 | 2.24681541 | 11.359609 | 5.60583684 |
| 4.99649661 | 4.7905021 | 4.3710782 | 6.56242695 | 7.66912992 | 9.45119465 | 3.35229043 |
| 7.39126717 | 8.82430558 | 19.4270142 | 7.57203109 | 5.61703852 | 12.4955698 | 5.26948663 |
| 47.3041099 | 60.0975582 | 48.5675356 | 57.5474363 | 49.429939 | 62.4778492 | 61.6911132 |
| 22.6468426 | 10.2777206 | 0 | 18.9401738 | 10.7547564 | 24.6844303 | 8.77425582 |
| 20.6955481 | 12.1636996 | 0 | 14.6392601 | 10.4851386 | 6.81576537 | 8.96933894 |
| 2.95650687 | 17.0718592 | 9.71350712 | 12.1152497 | 5.24256928 | 4.54384358 | 5.83007031 |
| 10.347774 | 23.5868497 | 1.94270142 | 23.7256974 | 19.1129097 | 25.0820166 | 13.8441747 |
| 555.823291 | 511.579023 | 281.691706 | 517.926927 | 410.418281 | 439.616867 | 624.938691 |
| 14.7825343 | 9.80478397 | 9.71350712 | 14.134458 | 8.98726163 | 9.08768716 | 10.0905063 |
| 8.8695206 | 14.4188 | 9.71350712 | 8.07683317 | 7.48938469 | 12.4955698 | 7.6239381 |
| 17.7390412 | 3.69121279 | 0 | 18.1728746 | 14.9787694 | 11.359609 | 14.3509423 |
| 23.1494488 | 10.1831333 | 8.06221091 | 20.949286 | 13.6681271 | 10.2236481 | 11.0883453 |
| 19.2172946 | 14.1684896 | 0 | 31.2977285 | 18.9706114 | 32.7497526 | 10.5008536 |
| 11.8260275 | 20.4170207 | 0 | 3.02881244 | 8.23832316 | 5.67980448 | 5.15736989 |
| 8.8695206 | 9.34338237 | 0 | 2.01920829 | 8.98726163 | 10.2236481 | 10.7632067 |
| 14.7825343 | 11.7080656 | 0 | 19.1824788 | 11.234077 | 7.95172627 | 17.041744 |
| 14.7825343 | 24.2235839 | 9.71350712 | 16.1536663 | 18.7234617 | 10.7916285 | 14.9496457 |
| 5.91301374 | 3.22981119 | 0 | 9.08643731 | 5.24256928 | 10.2236481 | 7.06335442 |
| 5.91301374 | 6.34427198 | 0 | 6.05762487 | 10.2379889 | 12.4955698 | 10.8372038 |
| 11.8260275 | 7.26707518 | 0 | 6.05762487 | 3.74469235 | 10.2236481 | 7.17547115 |
| 2.95650687 | 1.96095679 | 0 | 3.02881244 | 12.731954 | 3.40788269 | 9.64203936 |
| 11.8260275 | 14.7648512 | 9.71350712 | 22.2112912 | 15.7277078 | 13.6315307 | 13.0055415 |
| 67.999658 | 52.4648223 | 48.5675356 | 54.0138218 | 61.0384852 | 72.9854875 | 66.7655167 |
| 20.6955481 | 3.22981119 | 0 | 6.05762487 | 11.9830155 | 18.1753743 | 12.1086076 |
| 81.0674183 | 70.286459 | 51.7729929 | 46.8355363 | 54.298039 | 64.5566577 | 66.5480103 |
| 162.607878 | 214.782444 | 126.275593 | 182.73835 | 129.566355 | 132.907425 | 186.338017 |
| 26.6085618 | 40.7186911 | 19.4270142 | 14.134458 | 22.4681541 | 20.4472961 | 21.9748804 |
| 72.9370244 | 76.6503406 | 116.562085 | 74.3775374 | 78.5112197 | 72.1335169 | 84.2938474 |
| 5.91301374 | 10.381536 | 0 | 7.06722902 | 7.48938469 | 6.81576537 | 7.6239381 |
| 59.1301374 | 67.5953342 | 38.8540285 | 32.3073327 | 41.1916158 | 36.3507487 | 45.9678621 |
| 14.7825343 | 4.49866559 | 0 | 11.1056456 | 10.4851386 | 11.359609 | 5.38160336 |
| 56.1736305 | 49.2546207 | 38.8540285 | 38.3649575 | 40.689827 | 35.2147878 | 42.155893 |
| 38.4345893 | 14.4188 | 38.8540285 | 16.1536663 | 20.9702771 | 21.583257 | 27.3564838 |
| 11.8260275 | 7.26707518 | 0 | 19.1824788 | 13.4808924 | 13.6315307 | 15.2478762 |
| 29.5650687 | 21.5705247 | 19.4270142 | 39.3745617 | 29.9575388 | 23.8551788 | 40.5862587 |
| 41.3910961 | 50.8695263 | 29.1405214 | 38.3649575 | 55.4214467 | 39.7586313 | 58.5249366 |
| 11.3234213 | 16.3797568 | 24.2837678 | 7.06722902 | 13.1064232 | 10.7916285 | 10.202623 |

| | | | | | | |
|---|---|---|---|---|---|---|
| 17.7390412 | 12.3424928 | 19.4270142 | 10.0960415 | 11.9830155 | 9.08768716 | 11.4359072 |
| 5.91301374 | 12.9192448 | 9.71350712 | 9.08643731 | 8.98726163 | 5.67980448 | 8.07240505 |
| 8.8695206 | 15.1109024 | 19.4270142 | 10.0960415 | 18.7234617 | 13.6315307 | 17.041744 |
| 8.8695206 | 2.30700799 | 0 | 6.05762487 | 5.99150775 | 11.359609 | 6.95123768 |
| 56.1736305 | 73.7665806 | 33.9972749 | 35.8409472 | 44.1873697 | 38.6226704 | 41.9316596 |
| 38.4345893 | 59.4054558 | 9.71350712 | 32.3073327 | 32.9532926 | 29.5349833 | 29.5988185 |
| 53.2171236 | 60.905011 | 19.4270142 | 50.4802073 | 31.4554157 | 40.8945922 | 42.8285934 |
| 14.7825343 | 15.2262528 | 0 | 16.1536663 | 11.9830155 | 13.6315307 | 15.0236427 |
| 8919.78122 | 1083.71701 | 33705.8697 | 10166.7137 | 16879.5752 | 7768.83656 | 12818.9792 |
| 30076.0418 | 3011.22218 | 95653.7614 | 35492.1289 | 53579.807 | 28650.6377 | 46746.6249 |
| 144.868837 | 34.6051199 | 135.9891 | 142.354185 | 105.600324 | 79.5172627 | 66.8215751 |
| 42.8693496 | 0 | 48.5675356 | 36.3457492 | 23.966031 | 28.9670028 | 16.1448101 |
| 916.014523 | 258.442571 | 723.65628 | 870.783576 | 739.449419 | 587.09867 | 651.772711 |
| 155.216611 | 27.4337856 | 106.848578 | 147.907007 | 127.07239 | 108.859133 | 118.058924 |
| 79.8256854 | 4.21028959 | 82.5648105 | 86.3211545 | 65.532116 | 40.7014789 | 50.9009985 |
| 184.781679 | 44.2368783 | 121.418839 | 162.546267 | 164.391994 | 113.59609 | 123.104177 |
| 17.7390412 | 0 | 0 | 13.1248539 | 15.7277078 | 10.2236481 | 8.74510547 |
| 78.347432 | 16.0337056 | 87.4215641 | 80.7683317 | 61.4129545 | 57.9340057 | 44.8466947 |
| 94.6082198 | 26.0115151 | 116.562085 | 87.8355607 | 70.4002161 | 61.3418884 | 55.6838985 |
| 186.259933 | 28.8375999 | 194.270142 | 125.190914 | 85.3789855 | 72.7014973 | 61.8884387 |
| 12.4173288 | 146.614972 | 1.94270142 | 8.22827379 | 10.5825006 | 11.6890376 | 9.12854471 |
| 14.5460138 | 79.5617847 | 3.20545735 | 14.5988759 | 7.96870531 | 11.7685549 | 9.46937959 |
| 22.5581474 | 249.867422 | 25.3522536 | 25.8559622 | 17.0608183 | 25.9907853 | 20.799897 |
| 10.0521233 | 41.6091962 | 0 | 8.63211545 | 6.36597699 | 15.5058662 | 13.5212785 |
| 489.508842 | 1465.57181 | 414.475349 | 483.347985 | 358.928761 | 503.310194 | 442.609969 |
| 31.0433221 | 197.133833 | 48.5675356 | 26.7545099 | 14.6043001 | 35.2147878 | 11.6601406 |
| 12.4173288 | 146.614972 | 1.94270142 | 8.22827379 | 10.5825006 | 11.6890376 | 9.12854471 |
| 5.91301374 | 146.148956 | 9.71350712 | 29.2785202 | 9.7362001 | 4.54384358 | 10.0905063 |
| 12.4173288 | 146.614972 | 1.94270142 | 8.22827379 | 10.5825006 | 11.6890376 | 9.01642797 |
| 14.5460138 | 85.1620466 | 3.20545735 | 15.5378078 | 7.96870531 | 11.7799145 | 10.0030553 |
| 11.2938562 | 102.323879 | 1.94270142 | 7.21866964 | 9.27185825 | 9.41711582 | 7.42212797 |
| 85.9456546 | 474.673816 | 25.935064 | 84.5543472 | 84.8172816 | 134.690883 | 127.411702 |
| 99.2499355 | 497.701216 | 25.935064 | 92.6311804 | 93.0556048 | 144.346551 | 138.997846 |
| 12.4173288 | 147.191724 | 1.94270142 | 8.22827379 | 10.5825006 | 11.6890376 | 9.01642797 |
| 5.91301374 | 6.11357118 | 0 | 12.1152497 | 9.7362001 | 12.4955698 | 9.41780589 |
| 70.9561648 | 74.5163582 | 58.2810427 | 50.4802073 | 44.9363081 | 67.0216928 | 59.4218705 |
| 5.91301374 | 4.84471679 | 9.71350712 | 5.04802073 | 8.98726163 | 6.81576537 | 10.202623 |
| 5.91301374 | 12.111792 | 9.71350712 | 5.04802073 | 9.7362001 | 5.67980448 | 9.19357241 |
| 0 | 8.07452798 | 9.71350712 | 9.08643731 | 10.4851386 | 15.9034525 | 7.39970463 |
| 56.1736305 | 105.891667 | 77.708057 | 78.7491234 | 47.932062 | 64.749771 | 93.9538254 |
| 14.7825343 | 13.4959968 | 0 | 14.134458 | 14.9787694 | 32.942866 | 25.2262658 |
| 0 | 12.8038944 | 0 | 3.02881244 | 4.49363081 | 12.4955698 | 6.05430379 |
| 2.95650687 | 5.19076799 | 9.71350712 | 5.04802073 | 7.48938469 | 7.95172627 | 5.83007031 |
| 8.8695206 | 8.18987838 | 0 | 9.08643731 | 7.48938469 | 11.359609 | 3.13926863 |
| 45.5893359 | 37.2178065 | 19.4270142 | 68.5723136 | 50.8004964 | 78.6993708 | 35.9334141 |
| 22.1738015 | 18.0788681 | 9.71350712 | 17.1632705 | 14.6642152 | 8.80369694 | 15.996816 |

| | | | | | | |
|---|---|---|---|---|---|---|
| 1.47825343 | 24.1082335 | 24.2837678 | 18.1728746 | 11.234077 | 19.3113352 | 20.2931294 |
| 8.8695206 | 15.4569536 | 38.8540285 | 18.6776767 | 14.9787694 | 19.3113352 | 17.4902109 |
| 35.4780824 | 13.7266976 | 19.4270142 | 28.2689161 | 14.2298309 | 14.7674916 | 13.2297749 |
| 20.6955481 | 9.57408317 | 19.4270142 | 13.1248539 | 9.7362001 | 14.7674916 | 13.2297749 |
| 4.4347603 | 21.2244735 | 24.2837678 | 16.1536663 | 13.4808924 | 15.9034525 | 16.0326934 |
| 14.7825343 | 8.30522878 | 0 | 9.08643731 | 9.7362001 | 13.6315307 | 10.9874402 |
| 17.7390412 | 7.61312638 | 9.71350712 | 15.1440622 | 14.2298309 | 27.2630615 | 18.1629114 |
| 47.3041099 | 42.3335967 | 0 | 41.39377 | 37.8213927 | 42.5985336 | 33.1865541 |
| 14.7825343 | 5.76751998 | 48.5675356 | 31.2977285 | 17.2255848 | 15.9034525 | 13.2297749 |
| 20.0451166 | 18.2288237 | 16.2215569 | 31.7823385 | 21.8690033 | 25.5591201 | 25.7240641 |
| 5.91301374 | 4.96006719 | 19.4270142 | 7.06722902 | 7.48938469 | 6.81576537 | 7.39970463 |
| 8.8695206 | 3.34516159 | 0 | 8.07683317 | 7.48938469 | 1.1359609 | 7.84817157 |
| 20.6955481 | 22.6086783 | 19.4270142 | 7.06722902 | 14.2298309 | 11.359609 | 9.64203936 |
| 14.7825343 | 5.19076799 | 0 | 6.05762487 | 4.49363081 | 9.08768716 | 7.17547115 |
| 23.6520549 | 11.6919165 | 19.4270142 | 46.9465928 | 39.3192696 | 34.0788269 | 21.7506469 |
| 5.91301374 | 10.381536 | 0 | 5.04802073 | 13.4808924 | 3.40788269 | 10.0905063 |
| 2.95650687 | 16.0337056 | 0 | 6.05762487 | 7.48938469 | 7.95172627 | 6.72700421 |
| 23.6520549 | 61.8693404 | 67.9945498 | 37.3553534 | 33.7022311 | 27.2630615 | 48.2101968 |
| 2.95650687 | 3.57586239 | 0 | 2.01920829 | 7.48938469 | 4.54384358 | 5.83007031 |
| 11.8260275 | 9.68943357 | 0 | 7.06722902 | 10.4851386 | 14.7674916 | 13.0055415 |
| 14.7825343 | 26.1845407 | 0 | 34.326541 | 23.966031 | 14.7674916 | 24.4414486 |
| 2.95650687 | 6.45962238 | 0 | 7.06722902 | 8.98726163 | 6.81576537 | 5.60583684 |
| 29.5650687 | 52.8304831 | 0 | 18.6776767 | 20.5958079 | 43.166514 | 22.983931 |
| 8.8695206 | 5.42146879 | 9.71350712 | 6.05762487 | 2.24681541 | 11.359609 | 4.93313642 |
| 16.2607878 | 10.958288 | 29.1405214 | 18.1728746 | 19.4724002 | 18.1753743 | 12.1086076 |
| 118.260275 | 99.4320445 | 106.848578 | 94.9027897 | 135.557863 | 186.297587 | 161.223867 |
| 5.91301374 | 14.0727488 | 19.4270142 | 12.1152497 | 8.23832316 | 10.2236481 | 18.6113783 |
| 51.7388702 | 24.1082335 | 33.9972749 | 8.07683317 | 28.4596618 | 7.95172627 | 24.5535654 |
| 48.7823633 | 25.4924383 | 53.4242892 | 8.07683317 | 29.2086003 | 6.81576537 | 24.3293319 |
| 32.5215755 | 13.6113472 | 29.1405214 | 22.2112912 | 27.7107234 | 13.6315307 | 25.3383825 |
| 2.21738015 | 7.49085496 | 9.71350712 | 5.9667605 | 13.6606377 | 7.39510543 | 9.51422628 |
| 2.95650687 | 5.30611839 | 9.71350712 | 9.08643731 | 2.99575388 | 12.4955698 | 7.17547115 |
| 8.8695206 | 9.80478397 | 0 | 6.05762487 | 7.48938469 | 9.08768716 | 5.15736989 |
| 2.95650687 | 9.57408317 | 0 | 6.05762487 | 7.48938469 | 6.81576537 | 4.93313642 |
| 17.7390412 | 34.2590687 | 0 | 8.07683317 | 12.731954 | 26.1271006 | 17.7144444 |
| 23.6520549 | 45.7941087 | 9.71350712 | 49.4706031 | 32.9532926 | 40.8945922 | 48.2101968 |
| 14.7825343 | 12.2271424 | 0 | 4.03841658 | 7.48938469 | 4.54384358 | 6.95123768 |
| 8.8695206 | 12.688544 | 0 | 10.0960415 | 8.98726163 | 2.27192179 | 8.96933894 |
| 10.347774 | 9.82439354 | 9.71350712 | 8.75326794 | 12.3574847 | 7.38374582 | 8.70698578 |
| 0 | 7.84382718 | 0 | 6.05762487 | 8.23832316 | 9.08768716 | 2.91503516 |
| 5.91301374 | 13.9573984 | 9.71350712 | 17.1632705 | 13.4808924 | 19.3113352 | 5.60583684 |
| 2.95650687 | 5.42146879 | 0 | 11.6104477 | 15.3532386 | 14.1995112 | 2.46656821 |
| 2.95650687 | 2.88375999 | 9.71350712 | 6.05762487 | 8.98726163 | 9.08768716 | 5.15736989 |
| 23.6520549 | 23.9928831 | 0 | 14.134458 | 24.7149695 | 24.9911397 | 26.2353164 |
| 11.8260275 | 36.6814271 | 0 | 18.1728746 | 14.9787694 | 15.9034525 | 18.1629114 |
| 399.128427 | 458.97924 | 213.697157 | 394.755221 | 332.52868 | 331.700581 | 434.788705 |

| | | | | | | |
|---|---|---|---|---|---|---|
| 20.6955481 | 21.6858751 | 9.71350712 | 3.02881244 | 6.74044622 | 9.08768716 | 13.5661251 |
| 11.8260275 | 35.6432735 | 9.71350712 | 20.1920829 | 26.2128464 | 27.2630615 | 27.3564838 |
| 0 | 4.49866559 | 0 | 9.08643731 | 9.7362001 | 12.4955698 | 3.3635021 |
| 4.4347603 | 8.13220318 | 4.85675356 | 11.6104477 | 6.74044622 | 5.67980448 | 8.74510547 |
| 11.8260275 | 26.8766431 | 19.4270142 | 33.3169368 | 16.4766463 | 19.3113352 | 19.5083122 |
| 5.91301374 | 19.6095679 | 0 | 10.0960415 | 7.48938469 | 14.7674916 | 8.96933894 |
| 5.91301374 | 6.57497278 | 9.71350712 | 11.1056456 | 10.1106693 | 11.9275894 | 10.9874402 |
| 2.95650687 | 4.72936639 | 19.4270142 | 11.1056456 | 5.99150775 | 9.08768716 | 8.52087199 |
| 17.7390412 | 8.53592958 | 0 | 6.05762487 | 8.23832316 | 14.1995112 | 12.332841 |
| 8.8695206 | 13.265296 | 0 | 9.08643731 | 12.731954 | 14.7674916 | 4.48466947 |
| 14.7825343 | 18.2253632 | 9.71350712 | 13.1248539 | 21.7192156 | 14.7674916 | 19.7325457 |
| 17.7390412 | 19.8402687 | 0 | 20.1920829 | 20.9702771 | 21.583257 | 22.1991139 |
| 23.1494488 | 19.3592576 | 12.9189645 | 28.016515 | 24.7748846 | 19.1182219 | 25.3765022 |
| 5.91301374 | 11.9964416 | 0 | 18.1728746 | 9.7362001 | 7.95172627 | 10.0905063 |
| 0 | 5.76751998 | 0 | 10.0960415 | 7.48938469 | 2.27192179 | 4.93313642 |
| 32.5215755 | 7.15172478 | 19.4270142 | 27.5924813 | 21.3447464 | 31.0458113 | 12.1086076 |
| 138.955823 | 158.8375 | 67.9945498 | 113.075664 | 119.081217 | 98.8285979 | 133.64315 |
| 5.91301374 | 2.76840959 | 0 | 4.03841658 | 13.4808924 | 11.359609 | 11.4359072 |
| 17.7390412 | 22.0319263 | 0 | 19.1824788 | 14.9787694 | 9.08768716 | 16.1448101 |
| 53.7197298 | 53.9932151 | 29.9176019 | 61.9190223 | 68.4679548 | 68.7256342 | 57.7961778 |
| 5.91301374 | 2.53770879 | 0 | 5.04802073 | 7.48938469 | 7.95172627 | 2.91503516 |
| 2.95650687 | 15.2262528 | 0 | 9.08643731 | 5.99150775 | 11.359609 | 10.0905063 |
| 8.8695206 | 5.65216958 | 0 | 7.06722902 | 8.98726163 | 9.08768716 | 6.50277073 |
| 14.7825343 | 18.456064 | 0 | 8.07683317 | 15.7277078 | 18.1753743 | 29.1503516 |
| 0 | 7.09404958 | 0 | 2.01920829 | 3.74469235 | 9.65566761 | 6.83912094 |
| 2.95650687 | 14.7648512 | 0 | 5.04802073 | 8.98726163 | 3.40788269 | 4.48466947 |
| 7.98256854 | 13.5836631 | 0 | 13.5589837 | 16.1246452 | 13.8928018 | 10.2945588 |
| 17.7390412 | 19.2635167 | 9.71350712 | 13.1248539 | 20.9702771 | 13.6315307 | 18.3871448 |
| 5.91301374 | 11.8810912 | 0 | 11.1056456 | 4.49363081 | 14.7674916 | 7.17547115 |
| 17.7390412 | 24.1082335 | 38.8540285 | 18.1728746 | 15.7277078 | 14.7674916 | 17.7144444 |
| 0 | 4.26796479 | 9.71350712 | 7.06722902 | 1.49787694 | 11.359609 | 6.95123768 |
| 38.4345893 | 10.958288 | 19.4270142 | 16.1536663 | 29.9575388 | 14.7674916 | 18.8356118 |
| 0 | 3.11446079 | 0 | 2.01920829 | 9.7362001 | 15.9034525 | 0.89693389 |
| 17.7390412 | 6.34427198 | 19.4270142 | 11.1056456 | 8.98726163 | 9.08768716 | 6.27853726 |
| 17.7390412 | 20.4170207 | 9.71350712 | 17.1632705 | 10.4851386 | 11.359609 | 14.1267088 |
| 5.91301374 | 11.2847296 | 0 | 7.31963006 | 6.80785068 | 12.4955698 | 16.2838348 |
| 10.347774 | 21.3398239 | 0 | 5.04802073 | 5.99150775 | 13.6315307 | 9.97838957 |
| 13.3042809 | 12.400168 | 0 | 13.629656 | 9.7362001 | 10.2236481 | 10.3147398 |
| 839.64795 | 969.168291 | 369.113271 | 452.302657 | 412.665096 | 381.682861 | 526.049487 |
| 8.3077843 | 15.4823306 | 1.94270142 | 13.3166787 | 9.11458117 | 9.58750996 | 11.7879537 |
| 2.95650687 | 4.26796479 | 0 | 8.07683317 | 13.4808924 | 4.54384358 | 8.52087199 |
| 141.91233 | 122.732825 | 135.9891 | 145.382997 | 112.962389 | 106.780324 | 130.728115 |
| 8.8695206 | 8.30522878 | 9.71350712 | 15.1440622 | 9.7362001 | 15.9034525 | 14.7994093 |
| 2.95650687 | 17.5332608 | 19.4270142 | 13.1248539 | 14.2298309 | 11.359609 | 12.5570745 |
| 11.8260275 | 13.3806464 | 0 | 7.06722902 | 8.23832316 | 3.40788269 | 6.50277073 |
| 39.9128427 | 16.6681328 | 14.5702607 | 3.02881244 | 16.8511156 | 5.67980448 | 14.463059 |

| | | | | | | |
|---|---|---|---|---|---|---|
| 144.868837 | 149.494118 | 135.9891 | 106.008435 | 126.570601 | 86.3330281 | 134.315851 |
| 66.5214045 | 58.1458295 | 35.6485711 | 34.6597103 | 31.9422257 | 40.5878828 | 36.9200414 |
| 29.5650687 | 33.0478895 | 38.8540285 | 33.7409705 | 28.1376183 | 22.9804889 | 23.5400301 |
| 2.95650687 | 15.9183552 | 0 | 7.06722902 | 8.23832316 | 5.67980448 | 8.74510547 |
| 94.6082198 | 88.3584062 | 67.9945498 | 104.998831 | 77.1406623 | 86.3330281 | 86.5541208 |
| 8.8695206 | 7.61312638 | 0 | 4.03841658 | 7.48938469 | 7.95172627 | 4.48466947 |
| 836.691444 | 389.076898 | 1214.18839 | 584.5608 | 595.406083 | 473.695693 | 447.794247 |
| 14.7825343 | 4.61401599 | 0 | 13.1248539 | 15.7277078 | 13.6315307 | 13.6782419 |
| 38.4345893 | 9.34338237 | 29.1405214 | 26.2497078 | 29.2086003 | 22.7192179 | 27.3564838 |
| 8.8695206 | 35.6432735 | 19.4270142 | 18.1728746 | 32.2043542 | 19.3113352 | 19.5083122 |
| 78.347432 | 49.0815951 | 33.9972749 | 71.1770923 | 68.5278699 | 66.4537124 | 55.4977847 |
| 5.91301374 | 11.3043392 | 0 | 7.06722902 | 8.23832316 | 3.40788269 | 6.27853726 |
| 91.6517129 | 59.1170798 | 29.1405214 | 67.6434778 | 46.0597158 | 48.8463185 | 47.3132629 |
| 17.7390412 | 18.2253632 | 0 | 22.2112912 | 23.966031 | 23.8551788 | 8.07240505 |
| 141.91233 | 100.124147 | 106.848578 | 147.402205 | 136.306801 | 101.10052 | 148.218326 |
| 8.8695206 | 9.68943357 | 0 | 9.08643731 | 17.2255848 | 7.95172627 | 8.52087199 |
| 17.7390412 | 13.4959968 | 9.71350712 | 30.2881244 | 23.966031 | 22.7192179 | 26.2353164 |
| 73.9126717 | 69.9023422 | 67.9945498 | 85.8163524 | 91.3704932 | 84.0611063 | 79.8271166 |
| 65.0431511 | 56.3717403 | 48.5675356 | 90.6927404 | 66.5282042 | 60.9670213 | 71.5820518 |
| 5.91301374 | 9.80478397 | 9.71350712 | 11.1056456 | 6.74044622 | 15.9034525 | 9.86627284 |
| 4.4347603 | 10.4392112 | 14.5702607 | 12.6200518 | 7.48938469 | 17.6073939 | 7.6239381 |
| 179.519097 | 4081.12598 | 562.509197 | 380.721723 | 59.4207781 | 167.168005 | 210.292878 |
| 11.5599419 | 1826.39017 | 106.945713 | 72.1362162 | 7.39202269 | 25.331928 | 111.36107 |
| 11.8260275 | 23.9928831 | 29.1405214 | 9.08643731 | 9.7362001 | 7.95172627 | 9.64203936 |
| 20.6955481 | 12.5731936 | 19.4270142 | 14.134458 | 11.234077 | 14.7674916 | 14.7994093 |
| 17.7390412 | 9.80478397 | 29.1405214 | 35.3361451 | 33.7022311 | 26.1271006 | 30.0472855 |
| 17.7390412 | 9.92013437 | 0 | 7.06722902 | 8.23832316 | 9.08768716 | 5.83007031 |
| 5.91301374 | 3.11446079 | 0 | 11.1056456 | 11.9830155 | 5.67980448 | 10.9874402 |
| 2.95650687 | 5.07541759 | 0 | 3.02881244 | 8.23832316 | 11.359609 | 8.29663852 |
| 70.9561648 | 60.7896606 | 19.4270142 | 58.0522384 | 45.6852466 | 62.4778492 | 56.9553023 |
| 5.91301374 | 3.92191359 | 9.71350712 | 13.1248539 | 8.23832316 | 4.54384358 | 10.0905063 |
| 8.8695206 | 10.8429376 | 9.71350712 | 12.1152497 | 9.7362001 | 14.7674916 | 9.86627284 |
| 2.95650687 | 5.07541759 | 0 | 6.05762487 | 5.99150775 | 9.08768716 | 6.27853726 |
| 32.5215755 | 79.361075 | 53.4242892 | 62.090655 | 52.9274816 | 51.1182403 | 58.2267061 |
| 692.650429 | 297.947775 | 72.0742228 | 740.524449 | 1554.60903 | 726.640106 | 2847.63282 |
| 23.0016234 | 4.59325292 | 28.3634408 | 13.1046618 | 44.0001351 | 11.5527223 | 85.4127724 |
| 7.15474662 | 3.99689135 | 3.20545735 | 7.52155089 | 8.29823824 | 3.78274978 | 5.51165878 |
| 5.91301374 | 9.45873277 | 9.71350712 | 11.1056456 | 6.74044622 | 15.9034525 | 12.5570745 |
| 28.0868152 | 19.5518927 | 9.71350712 | 16.1536663 | 17.2255848 | 17.6073939 | 20.6294796 |
| 51.7388702 | 46.0824847 | 19.4270142 | 40.3841658 | 20.9702771 | 35.7827682 | 43.5012939 |
| 5.91301374 | 23.1854303 | 0 | 6.05762487 | 15.7277078 | 15.9034525 | 20.4052461 |
| 11.8260275 | 106.81447 | 0 | 20.1920829 | 28.4596618 | 14.7674916 | 25.114149 |
| 23.6520549 | 380.771669 | 0 | 14.134458 | 12.731954 | 15.9034525 | 36.7742897 |
| 5.91301374 | 3.69121279 | 19.4270142 | 11.1056456 | 11.234077 | 10.2236481 | 8.52087199 |
| 7.89387334 | 4.3579381 | 12.9189645 | 12.8724529 | 9.54896548 | 4.54384358 | 6.15520885 |
| 32.5215755 | 30.9715823 | 4.85675356 | 19.6872808 | 22.0936848 | 18.7433548 | 25.562616 |

| | | | | | | |
|---|---|---|---|---|---|---|
| 70.9561648 | 82.4755358 | 38.8540285 | 74.7107068 | 77.1406623 | 67.0216928 | 75.1182136 |
| 11.8260275 | 15.3416032 | 0 | 8.07683317 | 11.9830155 | 19.3113352 | 14.3509423 |
| 14.7825343 | 7.20939998 | 14.5702607 | 18.1728746 | 24.3405002 | 23.8551788 | 24.4033289 |
| 2.95650687 | 4.49866559 | 0 | 15.1440622 | 2.99575388 | 2.27192179 | 7.17547115 |
| 32.5215755 | 12.5731936 | 19.4270142 | 29.2785202 | 30.7064772 | 29.5349833 | 29.823052 |
| 28.0868152 | 21.5705247 | 9.71350712 | 21.7064891 | 22.4681541 | 27.8310419 | 23.8808649 |
| 20.6955481 | 13.7266976 | 0 | 15.1440622 | 24.7149695 | 15.9034525 | 25.7868495 |
| 8.8695206 | 6.57497278 | 0 | 8.07683317 | 11.9830155 | 14.7674916 | 8.96933894 |
| 8.8695206 | 15.1881871 | 9.71350712 | 10.7724762 | 9.36173086 | 18.1753743 | 10.5008536 |
| 29.5650687 | 13.1499456 | 9.71350712 | 14.134458 | 22.4681541 | 11.359609 | 15.9205766 |
| 17.7390412 | 2.99911039 | 29.1405214 | 29.2785202 | 34.4511696 | 23.8551788 | 26.2353164 |
| 26.1059556 | 39.6459324 | 14.5702607 | 16.1536663 | 14.1025114 | 17.4142805 | 13.7724199 |
| 23.1494488 | 41.83759 | 14.5702607 | 16.1536663 | 15.6003883 | 17.4142805 | 14.4451204 |
| 8.8695206 | 4.96006719 | 0 | 20.1920829 | 8.23832316 | 5.67980448 | 8.96933894 |
| 20.6955481 | 41.1800927 | 9.71350712 | 18.1728746 | 19.4724002 | 28.3990224 | 26.4595499 |
| 8.8695206 | 5.19076799 | 0 | 6.05762487 | 10.4851386 | 20.4472961 | 11.6601406 |
| 168.520891 | 81.8987838 | 203.98365 | 132.258143 | 140.800432 | 123.819738 | 124.449578 |
| 17.7390412 | 5.76751998 | 19.4270142 | 20.1920829 | 25.4639079 | 10.7916285 | 16.8175105 |
| 23.6520549 | 40.7186911 | 19.4270142 | 19.1824788 | 26.9617849 | 24.9911397 | 31.3926863 |
| 2.95650687 | 2.21126716 | 3.20545735 | 11.1056456 | 8.23832316 | 9.56479074 | 4.74702263 |
| 15.2851405 | 13.1499456 | 14.5702607 | 9.59123938 | 10.5076067 | 11.359609 | 10.3528595 |
| 8.8695206 | 15.9183552 | 19.4270142 | 5.04802073 | 9.7362001 | 9.08768716 | 13.4540084 |
| 5.91301374 | 3.34516159 | 0 | 8.07683317 | 7.48938469 | 6.81576537 | 8.29663852 |
| 33.999829 | 31.3753087 | 33.9972749 | 58.5570405 | 38.1958619 | 45.4384358 | 47.6496131 |
| 6.65214045 | 14.9967055 | 2.42837678 | 12.9532212 | 9.26436886 | 10.3145249 | 3.90614711 |
| 47.3041099 | 84.6671934 | 106.848578 | 69.6626861 | 53.1746313 | 38.6226704 | 71.0820111 |
| 17.7390412 | 9.68943357 | 6.50804977 | 17.3349032 | 8.98726163 | 11.1664956 | 13.0055415 |
| 5.91301374 | 9.22803198 | 0 | 2.01920829 | 8.98726163 | 1.1359609 | 5.84352432 |
| 20.6955481 | 17.1872096 | 19.4270142 | 17.1632705 | 19.4724002 | 10.2236481 | 14.3509423 |
| 50.2606167 | 36.3930511 | 58.2810427 | 32.8121347 | 36.697985 | 36.3507487 | 50.1722397 |
| 1.9808596 | 5.26805275 | 8.06221091 | 8.75326794 | 9.60888056 | 7.38374582 | 5.64395653 |
| 11.8260275 | 25.9538399 | 19.4270142 | 16.1536663 | 17.2255848 | 11.359609 | 15.4721097 |
| 5.91301374 | 12.688544 | 9.71350712 | 13.1248539 | 8.23832316 | 3.40788269 | 9.86627284 |
| 5.91301374 | 7.95917758 | 0 | 12.1152497 | 9.7362001 | 11.359609 | 6.95123768 |
| 5.91301374 | 3.69121279 | 0 | 11.1056456 | 7.48938469 | 5.67980448 | 8.07240505 |
| 8.8695206 | 25.0310367 | 19.4270142 | 8.07683317 | 16.4766463 | 11.359609 | 9.64203936 |
| 28.0868152 | 109.409854 | 0 | 30.2881244 | 23.966031 | 16.471433 | 25.0020323 |
| 5.91301374 | 8.99733118 | 0 | 7.06722902 | 8.23832316 | 5.67980448 | 8.29663852 |
| 506.893102 | 1.3842048 | 2524.73477 | 110.117524 | 78.848242 | 87.8324964 | 14.5639641 |
| 8.8695206 | 20.0709695 | 9.71350712 | 21.2016871 | 13.4808924 | 15.9034525 | 16.3690436 |
| 11.8260275 | 9.34338237 | 19.4270142 | 15.1440622 | 17.9745233 | 12.4955698 | 13.0055415 |
| 5.41040757 | 2.02786003 | 0 | 8.3292342 | 11.7957809 | 11.8935106 | 5.65741054 |
| 17.7390412 | 28.2608479 | 19.4270142 | 12.1152497 | 13.4808924 | 9.08768716 | 19.9567791 |
| 2.95650687 | 3.46051199 | 0 | 8.07683317 | 7.48938469 | 6.81576537 | 4.93313642 |
| 59.1301374 | 49.7160223 | 48.5675356 | 36.3457492 | 39.6937389 | 37.4867095 | 49.5555977 |
| 2.95650687 | 16.8411584 | 0 | 16.1536663 | 15.7277078 | 17.0394134 | 18.1629114 |

| | | | | | | |
|---|---|---|---|---|---|---|
| 20.6955481 | 13.9573984 | 9.71350712 | 18.1728746 | 21.7192156 | 20.4472961 | 17.9386779 |
| 17.7390412 | 9.34338237 | 0 | 13.1248539 | 12.731954 | 13.6315307 | 16.3690436 |
| 8.8695206 | 5.76751998 | 0 | 11.1056456 | 10.4851386 | 5.67980448 | 11.3237904 |
| 5.91301374 | 5.65216958 | 0 | 6.05762487 | 8.23832316 | 4.54384358 | 6.27853726 |
| 8.8695206 | 6.92102398 | 9.71350712 | 6.05762487 | 7.48938469 | 0 | 4.93313642 |
| 26.6085618 | 14.8802016 | 9.71350712 | 16.1536663 | 24.7149695 | 21.583257 | 20.853713 |
| 449.389044 | 313.637737 | 359.399763 | 497.734844 | 326.537172 | 307.845403 | 421.783164 |
| 14.7825343 | 7.95917758 | 0 | 9.08643731 | 12.731954 | 22.7192179 | 11.4359072 |
| 2.95650687 | 6.80567358 | 0 | 5.04802073 | 8.98726163 | 9.08768716 | 8.07240505 |
| 505.562674 | 407.763663 | 301.118721 | 500.763656 | 388.699065 | 377.139017 | 450.036581 |
| 26.6085618 | 60.6743102 | 9.71350712 | 13.1248539 | 12.731954 | 17.0394134 | 15.2478762 |
| 18.7146885 | 54.4373141 | 0 | 24.2304995 | 20.3711264 | 25.649997 | 15.1536981 |
| 2.95650687 | 4.49866559 | 0 | 4.03841658 | 8.61279239 | 9.93965784 | 3.47561884 |
| 2.95650687 | 10.2892557 | 1.65129621 | 7.26914985 | 12.5671875 | 12.3478949 | 12.0480645 |
| 7.89387334 | 7.11365915 | 0 | 7.06722902 | 6.23865745 | 13.4384174 | 21.6004105 |
| 15.6399213 | 150.638394 | 33.9972749 | 12.1354418 | 3.68477727 | 6.52041554 | 10.9022315 |
| 2.95650687 | 8.42057918 | 9.71350712 | 9.08643731 | 7.48938469 | 11.359609 | 12.332841 |
| 20.6955481 | 7.44010078 | 0 | 15.1440622 | 9.98334979 | 10.7916285 | 10.2766201 |
| 2.95650687 | 4.21028959 | 4.85675356 | 4.03841658 | 8.23832316 | 6.24778492 | 5.15736989 |
| 4.93736647 | 3.53779676 | 0 | 5.04802073 | 6.61312668 | 8.89457381 | 18.2369084 |
| 76.8691786 | 86.5127998 | 48.5675356 | 65.6242695 | 101.855632 | 171.530095 | 155.169564 |
| 280.868152 | 156.848859 | 213.697157 | 269.564307 | 453.107774 | 485.055302 | 425.846274 |
| 11.8260275 | 24.1082335 | 9.71350712 | 22.2112912 | 11.9830155 | 20.4472961 | 17.041744 |
| 11.8260275 | 36.5660767 | 0 | 29.2785202 | 20.9702771 | 20.4472961 | 17.4902109 |
| 13.0677604 | 22.9904882 | 3.88540285 | 10.3585385 | 13.7505103 | 11.5300031 | 16.7771485 |
| 183.953857 | 391.800321 | 29.8204669 | 171.249055 | 206.31008 | 149.32206 | 280.509348 |
| 250.475262 | 763.73961 | 126.081322 | 205.666461 | 316.419014 | 275.163808 | 367.368427 |
| 56.9127572 | 64.7404118 | 58.2810427 | 93.1359824 | 71.8606461 | 89.502359 | 79.6544568 |
| 22.5581474 | 257.662802 | 25.3522536 | 26.794894 | 17.2705211 | 26.4792485 | 21.5084748 |
| 14.5460138 | 86.6777508 | 3.20545735 | 15.9416495 | 8.59032424 | 13.2907425 | 9.4312599 |
| 22.5581474 | 256.540442 | 25.3522536 | 26.794894 | 17.2705211 | 26.3429332 | 21.7327083 |
| 180.346919 | 1897.86013 | 276.834953 | 265.52589 | 91.6176429 | 128.079591 | 169.146036 |
| 11.8260275 | 15.572304 | 0 | 10.0960415 | 10.4851386 | 6.81576537 | 7.17547115 |
| 2.95650687 | 3.11446079 | 0 | 10.0960415 | 7.48938469 | 7.95172627 | 5.83007031 |
| 14.5460138 | 81.1789973 | 3.20545735 | 15.2046384 | 7.96870531 | 12.0525451 | 9.64203936 |
| 22.5581474 | 257.662802 | 25.3522536 | 26.794894 | 17.2705211 | 26.4792485 | 21.5084748 |
| 10.6138597 | 92.4072052 | 5.24529384 | 16.5171238 | 3.75218173 | 5.55484878 | 7.98943866 |
| 14.5460138 | 81.1789973 | 3.20545735 | 15.2046384 | 7.96870531 | 12.0525451 | 9.64203936 |
| 8.54430485 | 608.14807 | 64.4005522 | 31.0655196 | 6.83780822 | 21.2424687 | 24.6634398 |
| 13.3042809 | 14.995552 | 4.85675356 | 11.6104477 | 8.23832316 | 17.0394134 | 13.5661251 |
| 2.74955139 | 76.6295775 | 9.61637205 | 9.22778189 | 2.98077511 | 8.89457381 | 8.37512024 |
| 14.5460138 | 86.6777508 | 3.20545735 | 15.9416495 | 8.59032424 | 13.2907425 | 9.4312599 |
| 22.2624967 | 248.036811 | 25.3522536 | 25.5530809 | 16.8361368 | 26.0135045 | 20.7505656 |
| 22.5581474 | 256.472386 | 25.3522536 | 26.794894 | 17.2705211 | 26.3429332 | 21.5084748 |
| 14.5460138 | 85.1620466 | 3.20545735 | 15.5378078 | 7.96870531 | 11.7799145 | 10.0030553 |
| 13.3042809 | 1.90328159 | 11.6562085 | 3.02881244 | 3.99184204 | 10.2236481 | 2.95315485 |

| | | | | | | |
|---|---|---|---|---|---|---|
| 79.8256854 | 50.9848767 | 38.8540285 | 51.4898114 | 65.9065853 | 52.2542012 | 59.646104 |
| 16.349483 | 142.547717 | 20.4955 | 21.5651446 | 7.3845333 | 9.86014057 | 14.7029889 |
| 13.3042809 | 1.90328159 | 11.6562085 | 3.02881244 | 3.99184204 | 11.359609 | 2.95315485 |
| 22.5581474 | 257.662802 | 25.3522536 | 26.794894 | 17.2705211 | 26.4792485 | 21.5084748 |
| 8.8695206 | 12.5731936 | 0 | 16.1536663 | 8.23832316 | 17.0394134 | 15.6963431 |
| 152.260104 | 115.081633 | 106.848578 | 126.200518 | 126.697921 | 112.653242 | 121.94489 |
| 22.5581474 | 257.662802 | 25.3522536 | 27.8044982 | 17.2705211 | 26.4792485 | 21.7327083 |
| 22.5581474 | 257.676644 | 25.3522536 | 26.794894 | 17.2705211 | 26.4224504 | 21.6205915 |
| 22.5581474 | 257.662802 | 25.3522536 | 26.794894 | 17.2705211 | 26.4792485 | 21.6205915 |
| 192.172946 | 1991.8707 | 373.970024 | 270.573911 | 92.3665814 | 118.991904 | 167.576402 |
| 136.501922 | 912.144821 | 39.631109 | 127.714924 | 57.353708 | 108.620581 | 82.5650073 |
| 11.5303768 | 98.3004571 | 12.1418839 | 11.0248773 | 3.08562649 | 9.62158878 | 11.6197786 |
| 103.47774 | 108.890777 | 126.275593 | 126.200518 | 85.3789855 | 82.9251454 | 91.2630237 |
| 22.5581474 | 257.676644 | 25.3522536 | 26.794894 | 17.2705211 | 26.4224504 | 21.5084748 |
| 14.5460138 | 85.1620466 | 3.20545735 | 15.5378078 | 7.96870531 | 11.7799145 | 10.0030553 |
| 4.4347603 | 0.8074528 | 53.4242892 | 7.57203109 | 21.3447464 | 11.359609 | 13.0055415 |
| 11.8260275 | 20.1863199 | 0 | 15.1440622 | 5.99150775 | 6.81576537 | 11.4359072 |
| 14.7825343 | 24.4542847 | 0 | 21.2016871 | 23.966031 | 17.0394134 | 20.1810126 |
| 50.2606167 | 47.4090143 | 48.5675356 | 66.6338736 | 62.1618929 | 57.9340057 | 75.3424471 |
| 214.524138 | 599.616755 | 339.487074 | 195.681476 | 67.614165 | 134.838558 | 115.995976 |
| 17.7390412 | 24.4542847 | 9.71350712 | 10.0960415 | 13.4808924 | 7.95172627 | 11.6601406 |
| 5.91301374 | 1.96095679 | 0 | 2.01920829 | 5.24256928 | 10.2236481 | 5.62601785 |
| 1188.51576 | 1227.2129 | 417.680806 | 845.03867 | 1645.41782 | 1306.35503 | 2483.83419 |
| 15307.3143 | 643.116544 | 235426.272 | 5590.00652 | 3405.42322 | 3955.22272 | 738.737179 |
| 8.8695206 | 11.53504 | 0 | 0 | 2.24681541 | 9.08768716 | 6.95123768 |
| 82.7821923 | 7.95917758 | 305.975474 | 14.6392601 | 21.3447464 | 21.0152766 | 21.30218 |
| 1206.13654 | 3941.17711 | 273.435225 | 1042.45666 | 1477.88028 | 1164.23496 | 2048.94906 |
| 1206.13654 | 3941.26362 | 273.435225 | 1042.45666 | 1477.73049 | 1163.8601 | 2048.893 |
| 20.6955481 | 19.3788671 | 9.71350712 | 26.2497078 | 23.966031 | 20.4472961 | 22.8718143 |
| 35.4780824 | 22.4933279 | 19.4270142 | 30.2881244 | 17.2255848 | 24.9911397 | 23.0960478 |
| 28.0868152 | 32.9233111 | 22.6324716 | 24.5636689 | 25.8009303 | 29.5690621 | 27.5874442 |
| 2.21738015 | 8.68934561 | 4.85675356 | 5.3811901 | 5.99150775 | 9.08768716 | 5.08337284 |
| 8.54430485 | 31.3833832 | 9.71350712 | 3.73553534 | 16.236986 | 13.9495998 | 14.5415408 |
| 8.8695206 | 27.7994463 | 0 | 4.03841658 | 8.23832316 | 9.08768716 | 8.29663852 |
| 17.7390412 | 9.92013437 | 38.8540285 | 25.2401036 | 18.7234617 | 14.7674916 | 15.2478762 |
| 42.8693496 | 70.3833534 | 48.5675356 | 75.2155089 | 61.9147432 | 71.1906693 | 73.1741094 |
| 8.8695206 | 13.842048 | 19.4270142 | 19.1824788 | 20.9702771 | 18.1753743 | 28.7018846 |
| 40.8589249 | 72.2001222 | 17.775718 | 21.6560089 | 18.416397 | 10.8143477 | 20.4725161 |
| 5.91301374 | 19.6095679 | 38.8540285 | 10.0960415 | 10.4851386 | 7.95172627 | 11.4359072 |
| 2.95650687 | 5.28650882 | 9.71350712 | 13.1248539 | 14.2298309 | 11.9275894 | 10.725087 |
| 56.1736305 | 59.0594046 | 38.8540285 | 51.4898114 | 38.1958619 | 28.3990224 | 45.7436286 |
| 8.8695206 | 5.65216958 | 0 | 4.03841658 | 10.4851386 | 13.6315307 | 4.70890294 |
| 8.8695206 | 6.92102398 | 9.71350712 | 6.05762487 | 11.234077 | 9.08768716 | 11.2116737 |
| 17.7390412 | 14.1880992 | 0 | 12.1152497 | 17.9745233 | 10.2236481 | 15.0236427 |
| 12.5651542 | 6.85181374 | 19.6212844 | 17.8800894 | 14.4170655 | 21.8558876 | 7.60824176 |
| 3040132.2 | 25513.9316 | 19001320.6 | 978033.016 | 659634.357 | 804522.005 | 133822.163 |

| | | | | | | |
|---|---|---|---|---|---|---|
| 5.91301374 | 16.4951072 | 9.71350712 | 16.1536663 | 8.98726163 | 15.9034525 | 9.64203936 |
| 17.7390412 | 16.8411584 | 0 | 5.04802073 | 8.23832316 | 4.54384358 | 6.95123768 |
| 8.8695206 | 28.0301471 | 0 | 4.03841658 | 9.7362001 | 7.95172627 | 10.9874402 |
| 121.216782 | 234.161311 | 116.562085 | 80.7683317 | 70.4002161 | 64.749771 | 103.371631 |
| 14.7825343 | 23.7621823 | 0 | 11.1056456 | 20.2213387 | 17.0394134 | 20.853713 |
| 5.91301374 | 11.3043392 | 0 | 11.1056456 | 5.24256928 | 6.81576537 | 6.72700421 |
| 8.8695206 | 11.3043392 | 0 | 13.1248539 | 15.7277078 | 11.359609 | 11.8843741 |
| 17.7390412 | 3.80656319 | 29.1405214 | 21.2016871 | 24.7149695 | 21.583257 | 24.2172151 |
| 2.95650687 | 11.7657408 | 0 | 15.1440622 | 8.23832316 | 5.67980448 | 11.8843741 |
| 0 | 10.8429376 | 0 | 10.0960415 | 9.7362001 | 9.08768716 | 7.84817157 |
| 5.91301374 | 9.22803198 | 9.71350712 | 7.06722902 | 5.24256928 | 9.08768716 | 6.05430379 |
| 5.91301374 | 15.6876544 | 19.4270142 | 10.0960415 | 12.731954 | 4.54384358 | 7.39970463 |
| 29.5650687 | 8.07452798 | 97.1350712 | 21.2016871 | 14.2298309 | 10.2236481 | 19.0598453 |
| 32.5215755 | 13.6113472 | 9.71350712 | 31.2977285 | 26.9617849 | 24.9911397 | 27.8049507 |
| 56.1736305 | 58.021251 | 67.9945498 | 52.4994156 | 43.4384312 | 36.3507487 | 61.8884387 |
| 11.8260275 | 20.9937727 | 14.5702607 | 20.1920829 | 15.8550274 | 18.7433548 | 19.6585486 |
| 8.8695206 | 10.381536 | 58.2810427 | 10.0960415 | 14.9787694 | 10.2236481 | 9.64203936 |
| 17.7390412 | 7.61312638 | 19.4270142 | 10.0960415 | 10.4851386 | 11.359609 | 13.9024754 |
| 44.347603 | 87.3779278 | 43.710782 | 47.4513948 | 41.9405543 | 52.2542012 | 43.7255273 |
| 19.2172946 | 7.49777598 | 0 | 6.56242695 | 11.6085463 | 17.0394134 | 13.0055415 |
| 0 | 6.34427198 | 0 | 5.5528228 | 5.99150775 | 11.359609 | 5.60583684 |
| 20.6955481 | 28.0301471 | 38.8540285 | 37.3553534 | 27.7107234 | 30.6709442 | 32.2896202 |
| 8.8695206 | 2.65305919 | 9.71350712 | 4.03841658 | 5.24256928 | 9.08768716 | 6.72700421 |
| 452.345551 | 791.995844 | 456.534835 | 465.427511 | 487.558943 | 355.55576 | 492.192474 |
| 5.91301374 | 5.30611839 | 0 | 7.06722902 | 8.98726163 | 11.359609 | 4.03620252 |
| 36.9563358 | 14.707176 | 72.8513034 | 39.3745617 | 34.4511696 | 31.8069051 | 30.1594022 |
| 17.7390412 | 26.2998911 | 9.71350712 | 12.1152497 | 14.9787694 | 9.08768716 | 18.3871448 |
| 8.8695206 | 26.8766431 | 0 | 16.1536663 | 19.4724002 | 4.54384358 | 18.8356118 |
| 5.91301374 | 7.84382718 | 0 | 5.04802073 | 11.9830155 | 2.27192179 | 9.86627284 |
| 14.7825343 | 4.96006719 | 38.8540285 | 11.1056456 | 14.2298309 | 13.6315307 | 15.4721097 |
| 8.8695206 | 2.88375999 | 0 | 11.1056456 | 8.23832316 | 4.54384358 | 11.4359072 |
| 0 | 5.07541759 | 0 | 8.07683317 | 24.7149695 | 13.6315307 | 15.0236427 |
| 35.4780824 | 7.49777598 | 9.71350712 | 8.07683317 | 22.4681541 | 14.7674916 | 23.7687482 |
| 23.6520549 | 13.842048 | 29.1405214 | 8.07683317 | 15.7277078 | 9.08768716 | 18.8356118 |
| 29.5650687 | 19.8402687 | 19.4270142 | 10.0960415 | 11.9830155 | 3.40788269 | 10.0905063 |
| 8.8695206 | 33.9130175 | 0 | 11.1056456 | 14.2298309 | 6.81576537 | 14.3509423 |
| 23.6520549 | 14.6495008 | 0 | 7.06722902 | 8.23832316 | 4.54384358 | 6.27853726 |
| 17.7390412 | 1.6149056 | 0 | 3.02881244 | 11.9830155 | 4.54384358 | 7.39970463 |
| 11.8260275 | 1.6149056 | 19.4270142 | 8.07683317 | 10.4851386 | 10.2236481 | 8.74510547 |
| 5.91301374 | 17.879312 | 0 | 5.04802073 | 11.234077 | 5.67980448 | 9.19357241 |
| 11.8260275 | 8.53592958 | 19.4270142 | 15.1440622 | 24.7149695 | 9.08768716 | 15.4721097 |
| 14.7825343 | 17.4179104 | 9.71350712 | 15.1440622 | 5.99150775 | 3.40788269 | 20.853713 |
| 8.8695206 | 2.42235839 | 0 | 6.05762487 | 9.7362001 | 12.4955698 | 9.19357241 |
| 11.8260275 | 12.8038944 | 0 | 4.03841658 | 9.7362001 | 2.27192179 | 6.27853726 |
| 26.6085618 | 56.5216958 | 9.71350712 | 16.1536663 | 17.9745233 | 13.6315307 | 22.8718143 |
| 5.91301374 | 11.3043392 | 19.4270142 | 9.08643731 | 11.9830155 | 21.583257 | 11.2116737 |

Sperm
Dexamethasone
KG_7_17
21.2158687
7.71486134
67.0228579
19.8513026
8.67921901
4.82178834
6.75050367
6.26832484
11.0901132
19.2871533
11.572292
8.19704017
7.71486134
34.716876
11.6301535
6.75050367
20.4926004
85.8278324
17.358438
4.3396095
7.71486134
10.6079343
63.1654272
5.786146
7.71486134
19.2871533
7.2326825
4.82178834
18.8049745
20.7336898
51.5931352
23.8678523
11.0901132
13.0188285
15.4297227
9.64357667
11.572292
17.8406168
9.64357667
4.82178834
8.19704017
235.78545

| |
|---|
| 5.30396717 |
| 43.9747096 |
| 19.2871533 |
| 21.6980475 |
| 233.133466 |
| 5.786146 |
| 7.71486134 |
| 12.5366497 |
| 68.9515732 |
| 16.8762592 |
| 54.0040294 |
| 36.1634125 |
| 11.572292 |
| 22.1802263 |
| 30.8594454 |
| 18.3227957 |
| 17.358438 |
| 12.5366497 |
| 4.3396095 |
| 8.19704017 |
| 163.458625 |
| 18.3227957 |
| 9.16139784 |
| 6.02723542 |
| 5.30396717 |
| 12.5366497 |
| 11.0901132 |
| 14.9475438 |
| 15.9119015 |
| 32.7881607 |
| 57.86146 |
| 71.3624674 |
| 7.2326825 |
| 11.572292 |
| 6.75050367 |
| 11.572292 |
| 24.1089417 |
| 7.2326825 |
| 7.71486134 |
| 44.8426315 |
| 63.1654272 |
| 35.6812337 |
| 10.6079343 |
| 21.6980475 |
| 5.30396717 |
| 154.779406 |

| |
|---|
| 386.707425 |
| 5.30396717 |
| 17.6814978 |
| 11.0901132 |
| 7.2326825 |
| 6.75050367 |
| 13.0188285 |
| 9.64357667 |
| 14.465365 |
| 9.64357667 |
| 34.716876 |
| 13.0188285 |
| 56.8971024 |
| 18.3227957 |
| 13.9831862 |
| 10.1257555 |
| 466.749111 |
| 8.19704017 |
| 15.4297227 |
| 19.7693322 |
| 27.4841935 |
| 15.4297227 |
| 7.2326825 |
| 86.3100112 |
| 7.2326825 |
| 54.4862082 |
| 16.7171402 |
| 113.312026 |
| 6.26832484 |
| 5.30396717 |
| 58.3436389 |
| 21.2158687 |
| 14.9475438 |
| 14.465365 |
| 5.786146 |
| 12.5366497 |
| 21.2158687 |
| 34.716876 |
| 11.572292 |
| 10.6079343 |
| 12.0544708 |
| 6.99159309 |
| 25.5554782 |
| 13.0188285 |
| 33.9936078 |
| 6.26832484 |

| |
|---|
| 11.572292 |
| 4.3396095 |
| 13.9831862 |
| 3.37525184 |
| 5.786146 |
| 79.0773287 |
| 8.67921901 |
| 11.572292 |
| 4.3396095 |
| 8.67921901 |
| 8.67921901 |
| 40.0208432 |
| 22.6624052 |
| 7.71486134 |
| 9.64357667 |
| 15.9119015 |
| 10.6079343 |
| 17.8406168 |
| 9.16139784 |
| 11.0901132 |
| 22.6624052 |
| 9.16139784 |
| 14.465365 |
| 14.465365 |
| 14.465365 |
| 9.64357667 |
| 13.9831862 |
| 7.2326825 |
| 8.67921901 |
| 8.67921901 |
| 15.4297227 |
| 6.26832484 |
| 25.5554782 |
| 5.786146 |
| 12.5366497 |
| 9.64357667 |
| 2.65198358 |
| 11.572292 |
| 6.99159309 |
| 112.829847 |
| 23.8678523 |
| 33.2703395 |
| 14.9475438 |
| 8.19704017 |
| 9.64357667 |
| 8.92030842 |

| |
|---|
| 7.2326825 |
| 15.4297227 |
| 14.465365 |
| 35.1990549 |
| 144.171471 |
| 11.572292 |
| 9.16139784 |
| 75.2150763 |
| 5.06287775 |
| 15.4297227 |
| 17.5754185 |
| 20.0055998 |
| 9.68215098 |
| 45.8069892 |
| 30.3772665 |
| 82.4525806 |
| 7.2326825 |
| 13.0188285 |
| 19.0460639 |
| 7.71486134 |
| 11.0901132 |
| 7.2326825 |
| 22.1802263 |
| 5.786146 |
| 9.16139784 |
| 19.2871533 |
| 8.67921901 |
| 21.6980475 |
| 28.4485512 |
| 9.16139784 |
| 8.67921901 |
| 34.2346972 |
| 17.8406168 |
| 5.30396717 |
| 28.9693043 |
| 5.786146 |
| 10.6079343 |
| 8.19704017 |
| 56.3667057 |
| 24.1089417 |
| 20.7336898 |
| 13.5010073 |
| 132.599179 |
| 6.26832484 |
| 33.7525184 |
| 8.19704017 |

| |
|---|
| 61.7188907 |
| 6.26832484 |
| 17.358438 |
| 5.786146 |
| 16.7171402 |
| 21.6980475 |
| 35.6812337 |
| 18.8049745 |
| 5.786146 |
| 11.572292 |
| 4.82178834 |
| 11.572292 |
| 20.7336898 |
| 7.2326825 |
| 13.9831862 |
| 6.26832484 |
| 8.19704017 |
| 23.144584 |
| 7.95595075 |
| 8.67921901 |
| 12.5366497 |
| 26.5198358 |
| 4.3396095 |
| 17.358438 |
| 8.19704017 |
| 5.30396717 |
| 13.9831862 |
| 10.6079343 |
| 54.968387 |
| 36.0235807 |
| 8.19704017 |
| 6.26832484 |
| 10.1257555 |
| 9.64357667 |
| 6.26832484 |
| 5.30396717 |
| 16.8762592 |
| 12.5366497 |
| 11.0901132 |
| 13.7420968 |
| 9.64357667 |
| 10.6079343 |
| 6.75050367 |
| 178.888347 |
| 15.4297227 |
| 7.71486134 |

| |
|---|
| 18.3227957 |
| 13.8626415 |
| 18.8049745 |
| 27.9663724 |
| 10.1257555 |
| 99.3288397 |
| 8.19704017 |
| 21.2158687 |
| 24.5911205 |
| 11.0901132 |
| 16.8762592 |
| 15.4297227 |
| 111.865489 |
| 24.5911205 |
| 9.64357667 |
| 4.3396095 |
| 17.8406168 |
| 45.3248104 |
| 67.0228579 |
| 11.0901132 |
| 7.71486134 |
| 48.7000622 |
| 6.75050367 |
| 35.1990549 |
| 39.5386644 |
| 9.64357667 |
| 97.8823032 |
| 18.3227957 |
| 5.786146 |
| 21.6980475 |
| 13.5010073 |
| 64.1297849 |
| 7.2326825 |
| 448.426315 |
| 4.82178834 |
| 10.1257555 |
| 13.0188285 |
| 20.7336898 |
| 13.5010073 |
| 8.19704017 |
| 6.75050367 |
| 5.30396717 |
| 11.572292 |
| 8.19704017 |
| 62.2348221 |
| 49.182241 |

| |
|---|
| 2.41089417 |
| 52.075314 |
| 28.2556797 |
| 7.47377192 |
| 6.75050367 |
| 5.30396717 |
| 8.19704017 |
| 8.67921901 |
| 9.16139784 |
| 6.26832484 |
| 26.5198358 |
| 13.9831862 |
| 10.6079343 |
| 7.2326825 |
| 13.9831862 |
| 7.71486134 |
| 9.64357667 |
| 5.786146 |
| 12.0544708 |
| 6.26832484 |
| 16.3940803 |
| 17.358438 |
| 10.6079343 |
| 6.26832484 |
| 20.41063 |
| 27.9663724 |
| 20.251511 |
| 8.67921901 |
| 33.2703395 |
| 10.1257555 |
| 7.2326825 |
| 12.0544708 |
| 14.465365 |
| 16.8762592 |
| 14.9475438 |
| 16.8762592 |
| 11.0901132 |
| 19.2871533 |
| 12.5366497 |
| 9.40248726 |
| 3.57294516 |
| 4.3396095 |
| 13.9831862 |
| 41.4673797 |
| 20.7336898 |
| 6.75050367 |

| |
|---|
| 27.4841935 |
| 5.786146 |
| 8.67921901 |
| 8.67921901 |
| 12.2955603 |
| 11.572292 |
| 13.0188285 |
| 11.572292 |
| 8.19704017 |
| 5.786146 |
| 14.9475438 |
| 6.75050367 |
| 8.19704017 |
| 9.64357667 |
| 11.5144305 |
| 26.5198358 |
| 6.26832484 |
| 7.2326825 |
| 42.4317374 |
| 15.4297227 |
| 30.3772665 |
| 6.26832484 |
| 10.1257555 |
| 15.4297227 |
| 6.26832484 |
| 13.5010073 |
| 21.2158687 |
| 9.16139784 |
| 8.67921901 |
| 23.6267628 |
| 8.19704017 |
| 12.5366497 |
| 510.145206 |
| 8.19704017 |
| 27.0020147 |
| 11.0901132 |
| 8.36098098 |
| 13.9831862 |
| 7.2326825 |
| 6.26832484 |
| 21.2158687 |
| 67.5050367 |
| 10.1257555 |
| 23.144584 |
| 14.9475438 |
| 43.8782739 |

| |
|---|
| 7.2326825 |
| 6.26832484 |
| 6.75050367 |
| 240.607238 |
| 23.6267628 |
| 19.7693322 |
| 59.549086 |
| 6.75050367 |
| 40.9852009 |
| 13.5010073 |
| 11.0901132 |
| 21.2158687 |
| 4.82178834 |
| 7.71486134 |
| 44.3604527 |
| 46.3711384 |
| 14.9475438 |
| 21.6980475 |
| 31.823803 |
| 65.335232 |
| 45.3248104 |
| 4.3396095 |
| 17.4404084 |
| 7.2326825 |
| 22.9034946 |
| 8.67921901 |
| 9.16139784 |
| 7.71486134 |
| 25.0732993 |
| 4.82178834 |
| 6.75050367 |
| 15.9119015 |
| 10.1257555 |
| 54.4862082 |
| 7.2326825 |
| 68.4693944 |
| 27.4841935 |
| 9.64357667 |
| 9.16139784 |
| 18.8049745 |
| 9.64357667 |
| 10.1257555 |
| 13.9831862 |
| 18.3227957 |
| 7.2326825 |
| 13.5010073 |

| |
|---|
| 20.251511 |
| 7.71486134 |
| 7.2326825 |
| 50.6287775 |
| 13.0188285 |
| 6.75050367 |
| 8.67921901 |
| 17.8406168 |
| 11.0901132 |
| 21.2158687 |
| 144.171471 |
| 109.936774 |
| 13.5010073 |
| 8.19704017 |
| 14.9475438 |
| 8.19704017 |
| 13.0188285 |
| 16.3940803 |
| 7.71486134 |
| 7.71486134 |
| 4.3396095 |
| 22.6624052 |
| 13.9831862 |
| 13.5010073 |
| 16.3940803 |
| 15.9119015 |
| 14.465365 |
| 26.7609253 |
| 6.75050367 |
| 43.8782739 |
| 26.5198358 |
| 4.3396095 |
| 36.6455914 |
| 14.465365 |
| 36.563621 |
| 8.43812959 |
| 50.6287775 |
| 8.19704017 |
| 6.75050367 |
| 4.82178834 |
| 15.9119015 |
| 24.1089417 |
| 19.7693322 |
| 30.8594454 |
| 4.82178834 |
| 8.19704017 |

| |
|---|
| 19.7693322 |
| 9.64357667 |
| 31.3416242 |
| 26.037657 |
| 6.75050367 |
| 5.30396717 |
| 9.16139784 |
| 16.8762592 |
| 10.1257555 |
| 14.2242756 |
| 25.0732993 |
| 38.0921279 |
| 6.75050367 |
| 7.2326825 |
| 13.0188285 |
| 13.9831862 |
| 6.18635444 |
| 12.5366497 |
| 5.30396717 |
| 18.8049745 |
| 13.9831862 |
| 13.5010073 |
| 10.1257555 |
| 15.4297227 |
| 9.64357667 |
| 13.9831862 |
| 15.4297227 |
| 12.5366497 |
| 20.251511 |
| 9.16139784 |
| 7.71486134 |
| 10.6079343 |
| 8.19704017 |
| 5.30396717 |
| 8.73708047 |
| 12.5366497 |
| 7.71486134 |
| 10.6079343 |
| 4.82178834 |
| 12.5366497 |
| 12.5366497 |
| 33.7525184 |
| 7.2326825 |
| 4.82178834 |
| 11.0901132 |
| 11.572292 |

| |
|---|
| 9.16139784 |
| 33.7525184 |
| 25.0732993 |
| 10.1257555 |
| 8.67921901 |
| 10.1257555 |
| 20.251511 |
| 10.1257555 |
| 14.9475438 |
| 8.67921901 |
| 12.0544708 |
| 11.0901132 |
| 27.4841935 |
| 6.26832484 |
| 13.0188285 |
| 8.19704017 |
| 10.6079343 |
| 11.8133814 |
| 4.3396095 |
| 12.0544708 |
| 24.5911205 |
| 15.9119015 |
| 13.0188285 |
| 33.7525184 |
| 26.037657 |
| 42.9139162 |
| 36.1634125 |
| 7.71486134 |
| 7.2326825 |
| 23.144584 |
| 7.2326825 |
| 14.9475438 |
| 10.1257555 |
| 29.4129089 |
| 3.85743067 |
| 22.6624052 |
| 26.037657 |
| 7.2326825 |
| 59.7901754 |
| 89.9263525 |
| 17.1173486 |
| 67.0228579 |
| 13.7420968 |
| 24.1089417 |
| 8.43812959 |
| 12.0544708 |

| |
|---|
| 27.4841935 |
| 15.9119015 |
| 17.8406168 |
| 14.9475438 |
| 4.3396095 |
| 19.7693322 |
| 4.82178834 |
| 8.19704017 |
| 16.3940803 |
| 29.4129089 |
| 8.19704017 |
| 9.64357667 |
| 4.82178834 |
| 27.4841935 |
| 13.5010073 |
| 5.30396717 |
| 237.714165 |
| 6.75050367 |
| 25.5554782 |
| 6.75050367 |
| 6.26832484 |
| 6.26832484 |
| 11.0901132 |
| 12.5366497 |
| 4.82178834 |
| 10.6079343 |
| 48.2178834 |
| 11.0901132 |
| 11.572292 |
| 8.19704017 |
| 14.465365 |
| 11.572292 |
| 6.75050367 |
| 19.2871533 |
| 20.7336898 |
| 14.9475438 |
| 6.26832484 |
| 49.182241 |
| 5.30396717 |
| 9.64357667 |
| 13.5010073 |
| 24.1089417 |
| 23.6267628 |
| 12.5366497 |
| 8.19704017 |
| 9.64357667 |

| |
|---|
| 12.0544708 |
| 10.6079343 |
| 6.26832484 |
| 9.64357667 |
| 38.0921279 |
| 19.2871533 |
| 6.90962269 |
| 22.1802263 |
| 8.19704017 |
| 12.0544708 |
| 18.8049745 |
| 4.3396095 |
| 41.2262903 |
| 9.16139784 |
| 10.6079343 |
| 42.9139162 |
| 11.3312026 |
| 47.7357045 |
| 36.6455914 |
| 9.64357667 |
| 13.0188285 |
| 8.52009999 |
| 10.6079343 |
| 7.71486134 |
| 10.1257555 |
| 8.67921901 |
| 10.6079343 |
| 6.75050367 |
| 6.75050367 |
| 74.7377192 |
| 15.4297227 |
| 7.2326825 |
| 68.9515732 |
| 17.1173486 |
| 12.5366497 |
| 13.5010073 |
| 72.8090039 |
| 27.4841935 |
| 268.57361 |
| 43.1550056 |
| 32.7881607 |
| 5.786146 |
| 4.82178834 |
| 10.1257555 |
| 4.3396095 |
| 92.3372466 |

| |
|---|
| 12.5366497 |
| 5.786146 |
| 6.75050367 |
| 11.0901132 |
| 27.4841935 |
| 16.3940803 |
| 10.6079343 |
| 12.0544708 |
| 8.67921901 |
| 21.6980475 |
| 18.3227957 |
| 20.251511 |
| 26.5198358 |
| 15.4297227 |
| 4.3396095 |
| 20.251511 |
| 61.7188907 |
| 9.64357667 |
| 5.786146 |
| 6.75050367 |
| 18.8049745 |
| 5.30396717 |
| 26.5198358 |
| 16.1529909 |
| 18.3227957 |
| 56.8971024 |
| 10.8490238 |
| 5.786146 |
| 7.71486134 |
| 10.6079343 |
| 29.4129089 |
| 11.572292 |
| 61.7188907 |
| 20.251511 |
| 6.26832484 |
| 16.3940803 |
| 11.0901132 |
| 9.16139784 |
| 19.2871533 |
| 29.4129089 |
| 36.1634125 |
| 4.82178834 |
| 9.16139784 |
| 11.0901132 |
| 4.82178834 |
| 20.251511 |

| |
|---|
| 18.8049745 |
| 10.8490238 |
| 16.3940803 |
| 17.358438 |
| 18.8049745 |
| 11.0901132 |
| 6.75050367 |
| 50.1465987 |
| 28.4485512 |
| 14.9475438 |
| 4.2576391 |
| 39.0564855 |
| 8.19704017 |
| 42.4317374 |
| 39.0564855 |
| 14.465365 |
| 9.16139784 |
| 24.1089417 |
| 19.2871533 |
| 7.71486134 |
| 8.67921901 |
| 5.786146 |
| 89.2030842 |
| 11.0901132 |
| 22.1802263 |
| 9.64357667 |
| 9.64357667 |
| 13.5010073 |
| 11.0901132 |
| 11.572292 |
| 9.64357667 |
| 4.3396095 |
| 27.0020147 |
| 8.19704017 |
| 16.3940803 |
| 9.16139784 |
| 22.1802263 |
| 7.71486134 |
| 53.5218505 |
| 20.7336898 |
| 7.71486134 |
| 20.7336898 |
| 9.16139784 |
| 8.19704017 |
| 9.64357667 |
| 5.30396717 |

| |
|---|
| 10.1257555 |
| 9.16139784 |
| 10.3668449 |
| 18.3227957 |
| 10.1257555 |
| 18.8049745 |
| 8.19704017 |
| 1.4465365 |
| 7.2326825 |
| 4.09852009 |
| 13.9831862 |
| 19.2871533 |
| 22.1802263 |
| 7.2326825 |
| 6.26832484 |
| 23.6267628 |
| 15.9119015 |
| 7.2326825 |
| 28.4485512 |
| 5.786146 |
| 15.6708121 |
| 7.47377192 |
| 39.0564855 |
| 57.3792812 |
| 13.1007989 |
| 34.875995 |
| 15.7527825 |
| 16.3940803 |
| 1.92871533 |
| 11.0901132 |
| 26.2401721 |
| 45.3248104 |
| 15.9119015 |
| 6.75050367 |
| 12.5366497 |
| 7.71486134 |
| 9.16139784 |
| 17.358438 |
| 12.0544708 |
| 54.7272976 |
| 6.75050367 |
| 26.5198358 |
| 17.8406168 |
| 9.64357667 |
| 4.3396095 |
| 6.99159309 |

| |
|---|
| 5.786146 |
| 5.30396717 |
| 9.16139784 |
| 17.8406168 |
| 19.2871533 |
| 3.85743067 |
| 9.16139784 |
| 4.3396095 |
| 14.465365 |
| 68.4693944 |
| 17.358438 |
| 5.786146 |
| 13.5010073 |
| 24.8322099 |
| 34.716876 |
| 8.67921901 |
| 6.02723542 |
| 8.19704017 |
| 24.5911205 |
| 14.465365 |
| 29.6539983 |
| 7.2326825 |
| 12.0544708 |
| 5.30396717 |
| 8.19704017 |
| 30.6183559 |
| 14.9475438 |
| 5.786146 |
| 29.4129089 |
| 13.9831862 |
| 12.0544708 |
| 7.2326825 |
| 7.2326825 |
| 10.1257555 |
| 13.0188285 |
| 18.3227957 |
| 20.251511 |
| 16.3940803 |
| 18.8049745 |
| 13.5010073 |
| 9.16139784 |
| 12.5366497 |
| 42.6728268 |
| 23.144584 |
| 15.4297227 |
| 31.3416242 |

| |
|---|
| 15.4297227 |
| 4.3396095 |
| 5.786146 |
| 6.26832484 |
| 27.0020147 |
| 210.229971 |
| 4.82178834 |
| 18.8049745 |
| 107.52588 |
| 16.8762592 |
| 15.9119015 |
| 24.5911205 |
| 12.5366497 |
| 13.7420968 |
| 6.26832484 |
| 21.2158687 |
| 10.6079343 |
| 7.2326825 |
| 12.0544708 |
| 15.9119015 |
| 3.37525184 |
| 13.5010073 |
| 24.5911205 |
| 35.6812337 |
| 12.5366497 |
| 7.71486134 |
| 20.7336898 |
| 2.893073 |
| 10.1257555 |
| 38.3332173 |
| 5.786146 |
| 28.4485512 |
| 8.67921901 |
| 29.4129089 |
| 5.786146 |
| 25.0732993 |
| 7.2326825 |
| 20.251511 |
| 22.6624052 |
| 36.6455914 |
| 6.75050367 |
| 74.7377192 |
| 11.572292 |
| 13.0188285 |
| 27.4841935 |
| 40.503022 |

| |
|---|
| 20.251511 |
| 8.67921901 |
| 9.64357667 |
| 22.1802263 |
| 13.0188285 |
| 28.93073 |
| 10.1257555 |
| 11.572292 |
| 4.3396095 |
| 14.465365 |
| 6.26832484 |
| 38.5743067 |
| 11.3312026 |
| 37.1277702 |
| 4.3396095 |
| 12.0544708 |
| 10.1257555 |
| 59.3079965 |
| 3.85743067 |
| 726.643502 |
| 5.30396717 |
| 10.1257555 |
| 27.0020147 |
| 5.786146 |
| 30.8594454 |
| 9.64357667 |
| 12.5366497 |
| 66.0585002 |
| 10.1257555 |
| 12.0544708 |
| 31.3416242 |
| 6.75050367 |
| 38.5743067 |
| 12.0544708 |
| 19.2871533 |
| 32.3059819 |
| 7.71486134 |
| 44.6015421 |
| 6.75050367 |
| 14.9475438 |
| 9.64357667 |
| 27.4841935 |
| 4.82178834 |
| 4.82178834 |
| 8.67921901 |
| 8.67921901 |

| |
|---|
| 35.1990549 |
| 5.786146 |
| 29.4129089 |
| 5.30396717 |
| 7.2326825 |
| 29.1718194 |
| 359.223231 |
| 15.9119015 |
| 9.64357667 |
| 38.5743067 |
| 31.3416242 |
| 11.0901132 |
| 17.358438 |
| 9.64357667 |
| 21.2158687 |
| 10.1257555 |
| 13.5010073 |
| 10.6079343 |
| 25.5554782 |
| 8.19704017 |
| 25.0732993 |
| 18.8049745 |
| 15.9794065 |
| 19.7693322 |
| 9.64357667 |
| 91.1317996 |
| 20.251511 |
| 6.99159309 |
| 4.58069892 |
| 34.2346972 |
| 5.30396717 |
| 13.0188285 |
| 27.0020147 |
| 11.572292 |
| 17.8406168 |
| 8.67921901 |
| 50.1465987 |
| 4.82178834 |
| 27.0020147 |
| 28.4485512 |
| 12.5366497 |
| 8.67921901 |
| 20.251511 |
| 7.71486134 |
| 84.1402065 |
| 57.3792812 |

| |
|---|
| 7.71486134 |
| 21.6980475 |
| 29.8950877 |
| 18.3227957 |
| 12.5366497 |
| 37.609949 |
| 14.465365 |
| 8.83833802 |
| 9.16139784 |
| 40.0208432 |
| 13.5010073 |
| 10.6079343 |
| 47.2535257 |
| 14.465365 |
| 10.6079343 |
| 8.31758488 |
| 15.1886333 |
| 6.75050367 |
| 13.9831862 |
| 29.4129089 |
| 27.4841935 |
| 37.1277702 |
| 22.1802263 |
| 2.893073 |
| 38.0921279 |
| 22.8118806 |
| 12.5366497 |
| 2.57001318 |
| 7.2326825 |
| 69.9159309 |
| 18.3227957 |
| 19.2871533 |
| 5.786146 |
| 469.160005 |
| 13.2599179 |
| 4.3396095 |
| 13.5010073 |
| 24.1089417 |
| 9.16139784 |
| 14.465365 |
| 36.8866808 |
| 17.8406168 |
| 4.3396095 |
| 26.7609253 |
| 25.0732993 |
| 101.257555 |

| |
|---|
| 26.037657 |
| 9.64357667 |
| 11.0901132 |
| 9.64357667 |
| 6.26832484 |
| 13.5010073 |
| 26.5198358 |
| 13.9831862 |
| 30.8594454 |
| 11.2492322 |
| 4.82178834 |
| 7.71486134 |
| 118.615993 |
| 8.19704017 |
| 48.7000622 |
| 30.8594454 |
| 14.9475438 |
| 12.5366497 |
| 11.572292 |
| 13.9831862 |
| 88.7209054 |
| 20.7336898 |
| 11.572292 |
| 20.7336898 |
| 27.1755991 |
| 11.0901132 |
| 9.64357667 |
| 8.19704017 |
| 10.1257555 |
| 28.4485512 |
| 5.30396717 |
| 9.64357667 |
| 6.26832484 |
| 6.75050367 |
| 23.3856734 |
| 8.67921901 |
| 42.4317374 |
| 11.572292 |
| 21.2158687 |
| 4.82178834 |
| 11.0901132 |
| 63.1654272 |
| 21.3749877 |
| 109.936774 |
| 10.6079343 |
| 7.2326825 |

| |
|---|
| 64.3708743 |
| 5.786146 |
| 7.71486134 |
| 43.396095 |
| 54.4862082 |
| 24.5911205 |
| 23.6267628 |
| 10.1257555 |
| 9.64357667 |
| 69.433752 |
| 8.19704017 |
| 53.0396717 |
| 8.19704017 |
| 15.6708121 |
| 88.2387266 |
| 10.1257555 |
| 30.3772665 |
| 5.46308618 |
| 13.5010073 |
| 9.64357667 |
| 19.7693322 |
| 7.2326825 |
| 11.572292 |
| 13.0188285 |
| 27.2431041 |
| 9.64357667 |
| 21.6980475 |
| 4.3396095 |
| 7.2326825 |
| 5.06287775 |
| 11.0901132 |
| 8.67921901 |
| 10.6079343 |
| 9.64357667 |
| 15.9119015 |
| 10.1257555 |
| 12.5366497 |
| 11.0901132 |
| 11.0901132 |
| 22.1802263 |
| 33.2703395 |
| 7.95595075 |
| 8.67921901 |
| 15.9119015 |
| 12.0544708 |
| 15.4297227 |

| |
|---|
| 15.4297227 |
| 15.9119015 |
| 7.71486134 |
| 58.8258177 |
| 6.75050367 |
| 15.4297227 |
| 7.2326825 |
| 4.3396095 |
| 9.16139784 |
| 22.9372471 |
| 76.1842557 |
| 12.5366497 |
| 13.9831862 |
| 4.3396095 |
| 297.022162 |
| 25.5554782 |
| 20.7336898 |
| 9.64357667 |
| 20.7336898 |
| 16.6351698 |
| 64.1297849 |
| 9.64357667 |
| 13.0188285 |
| 10.6079343 |
| 10.1257555 |
| 8.19704017 |
| 12.0544708 |
| 52.075314 |
| 9.64357667 |
| 19.2871533 |
| 4.82178834 |
| 9.16139784 |
| 15.4297227 |
| 9.16139784 |
| 5.786146 |
| 14.465365 |
| 7.71486134 |
| 7.2326825 |
| 9.64357667 |
| 10.1257555 |
| 6.26832484 |
| 11.0901132 |
| 18.8049745 |
| 15.9119015 |
| 11.8133814 |
| 29.8950877 |

| |
|---|
| 5.786146 |
| 6.50941425 |
| 13.5010073 |
| 10.1257555 |
| 6.26832484 |
| 79.0773287 |
| 12.5366497 |
| 9.16139784 |
| 49.182241 |
| 69.1926626 |
| 14.6823455 |
| 5.786146 |
| 14.9475438 |
| 13.2599179 |
| 29.4129089 |
| 8.67921901 |
| 7.2326825 |
| 77.1486134 |
| 10.1257555 |
| 7.71486134 |
| 81.4882229 |
| 567.524487 |
| 17.8406168 |
| 12.5366497 |
| 9.64357667 |
| 10.1257555 |
| 46.289168 |
| 13.5010073 |
| 10.6079343 |
| 18.8049745 |
| 17.358438 |
| 48.4589728 |
| 13.9831862 |
| 13.9831862 |
| 10.6079343 |
| 36.6455914 |
| 16.3940803 |
| 7.71486134 |
| 16.8762592 |
| 37.609949 |
| 9.16139784 |
| 99.8110186 |
| 18.3227957 |
| 9.64357667 |
| 9.64357667 |
| 14.9475438 |

| |
|---|
| 50.1465987 |
| 9.16139784 |
| 6.75050367 |
| 35.1990549 |
| 11.0901132 |
| 38.5743067 |
| 6.75050367 |
| 187.567566 |
| 4.3396095 |
| 192.389355 |
| 26.5198358 |
| 22.6624052 |
| 21.6980475 |
| 19.7693322 |
| 13.7420968 |
| 5.786146 |
| 13.0188285 |
| 13.0188285 |
| 16.3940803 |
| 13.9831862 |
| 13.0188285 |
| 126.330854 |
| 10.1257555 |
| 35.1990549 |
| 13.9831862 |
| 13.0188285 |
| 5.786146 |
| 20.251511 |
| 28.93073 |
| 25.5554782 |
| 30.8594454 |
| 30.8594454 |
| 17.8406168 |
| 45.3248104 |
| 59.7901754 |
| 7.2326825 |
| 23.6267628 |
| 16.8762592 |
| 4.82178834 |
| 8.67921901 |
| 14.9475438 |
| 13.5010073 |
| 19.7693322 |
| 68.4693944 |
| 75.4609875 |
| 20.251511 |

| |
|---|
| 43.396095 |
| 38.0921279 |
| 7.2326825 |
| 36.6455914 |
| 20.7336898 |
| 11.572292 |
| 5.786146 |
| 13.5010073 |
| 17.8406168 |
| 28.93073 |
| 23.144584 |
| 20.7336898 |
| 7.71486134 |
| 186.12103 |
| 10.1257555 |
| 5.786146 |
| 5.30396717 |
| 54.968387 |
| 7.71486134 |
| 4.82178834 |
| 10.6079343 |
| 4.82178834 |
| 13.5010073 |
| 8.19704017 |
| 9.64357667 |
| 7.71486134 |
| 6.26832484 |
| 13.1007989 |
| 12.5366497 |
| 18.8049745 |
| 6.75050367 |
| 27.0020147 |
| 14.465365 |
| 45.3248104 |
| 8.19704017 |
| 18.3227957 |
| 100.775376 |
| 4.82178834 |
| 28.7282149 |
| 29.8950877 |
| 8.19704017 |
| 8.92030842 |
| 6.50941425 |
| 8.67921901 |
| 18.8049745 |
| 93.5426937 |

16.3940803
5.786146
21.2158687
22.1802263
39.0564855
9.64357667
8.67921901
12.5366497
825.972342
8.19704017
35.1990549
56.4149235
13.9831862
7.71486134
14.465365
5.30396717
22.6624052
9.64357667
13.0188285
4.82178834
16.1529909
17.8406168
138.385325
5.786146
8.19704017
40.9852009
9.16139784
6.26832484
41.4673797
8.67921901
8.19704017
7.2326825
8.67921901
33.2703395
4.3396095
7.2326825
12.0544708
7.2326825
6.75050367
10.1257555
4.3396095
29.4129089
16.8762592
11.572292
13.0188285
8.19704017

| |
|---|
| 6.50941425 |
| 6.26832484 |
| 14.465365 |
| 14.9475438 |
| 17.8406168 |
| 12.5366497 |
| 6.26832484 |
| 3.85743067 |
| 13.5010073 |
| 34.716876 |
| 7.2326825 |
| 8.67921901 |
| 44.3604527 |
| 8.67921901 |
| 6.75050367 |
| 8.67921901 |
| 7.71486134 |
| 26.037657 |
| 13.9831862 |
| 4.3396095 |
| 19.7693322 |
| 9.16139784 |
| 13.0188285 |
| 32.3059819 |
| 8.19704017 |
| 15.4297227 |
| 24.1089417 |
| 23.6267628 |
| 14.9475438 |
| 6.26832484 |
| 12.5366497 |
| 15.4297227 |
| 10.1257555 |
| 28.4485512 |
| 4.82178834 |
| 10.6079343 |
| 54.4476339 |
| 7.2326825 |
| 22.3393454 |
| 5.30396717 |
| 13.0188285 |
| 37.1277702 |
| 4.82178834 |
| 9.16139784 |
| 24.5911205 |
| 27.9663724 |

| |
|---|
| 10.6079343 |
| 60.9956225 |
| 11.572292 |
| 12.5366497 |
| 7.2326825 |
| 96.6768561 |
| 13.0188285 |
| 10.6079343 |
| 27.4841935 |
| 43.396095 |
| 6.75050367 |
| 24.5911205 |
| 3.85743067 |
| 25.5554782 |
| 6.26832484 |
| 19.7693322 |
| 7.2326825 |
| 18.8339052 |
| 24.5911205 |
| 32.7881607 |
| 7.2326825 |
| 6.26832484 |
| 3.85743067 |
| 9.16139784 |
| 5.786146 |
| 18.3227957 |
| 19.2871533 |
| 6.26832484 |
| 27.4841935 |
| 12.0544708 |
| 9.64357667 |
| 8.19704017 |
| 41.4673797 |
| 28.4485512 |
| 6.75050367 |
| 9.64357667 |
| 43.396095 |
| 13.9831862 |
| 8.19704017 |
| 6.26832484 |
| 31.5827136 |
| 13.5010073 |
| 9.16139784 |
| 9.64357667 |
| 13.0188285 |
| 10.1257555 |

| |
|---|
| 102.622121 |
| 4.3396095 |
| 24.7502395 |
| 20.251511 |
| 4.82178834 |
| 8.67921901 |
| 6.26832484 |
| 9.64357667 |
| 12.5366497 |
| 48.7000622 |
| 55.4505659 |
| 9.16139784 |
| 18.6554991 |
| 6.75050367 |
| 42.9139162 |
| 67.0228579 |
| 12.0544708 |
| 6.26832484 |
| 9.16139784 |
| 12.0544708 |
| 13.5010073 |
| 13.0188285 |
| 14.9475438 |
| 29.8950877 |
| 24.1089417 |
| 11.572292 |
| 13.0188285 |
| 12.0544708 |
| 7.2326825 |
| 35.1990549 |
| 12.7777391 |
| 32.7881607 |
| 4.3396095 |
| 9.64357667 |
| 7.2326825 |
| 5.30396717 |
| 45.5658998 |
| 10.6079343 |
| 7.71486134 |
| 9.16139784 |
| 21.6980475 |
| 8.52009999 |
| 7.2326825 |
| 216.739386 |
| 9.64357667 |
| 5.786146 |

| |
|---|
| 6.26832484 |
| 13.0188285 |
| 13.0188285 |
| 13.9831862 |
| 15.9119015 |
| 41.4673797 |
| 479.767939 |
| 15.9119015 |
| 11.3312026 |
| 14.465365 |
| 37.609949 |
| 6.75050367 |
| 47.7357045 |
| 4.3396095 |
| 10.6079343 |
| 12.5366497 |
| 14.9475438 |
| 4.3396095 |
| 5.30396717 |
| 9.16139784 |
| 11.3312026 |
| 30.3772665 |
| 5.786146 |
| 15.9119015 |
| 16.3940803 |
| 13.9831862 |
| 21.6980475 |
| 18.3227957 |
| 4.3396095 |
| 5.30396717 |
| 19.2871533 |
| 9.64357667 |
| 14.9475438 |
| 10.1257555 |
| 10.6079343 |
| 23.144584 |
| 53.8449104 |
| 8.19704017 |
| 35.1990549 |
| 10.6079343 |
| 9.64357667 |
| 42.4317374 |
| 67.5050367 |
| 10.1257555 |
| 32.7881607 |
| 16.3940803 |

| |
|---|
| 5.54505659 |
| 13.9831862 |
| 29.4129089 |
| 13.9831862 |
| 42.9139162 |
| 6.26832484 |
| 6.26832484 |
| 7.2326825 |
| 4.09852009 |
| 5.786146 |
| 79.0773287 |
| 24.1089417 |
| 12.5366497 |
| 105.838254 |
| 35.1990549 |
| 8.19704017 |
| 8.19704017 |
| 8.19704017 |
| 7.71486134 |
| 84.8634747 |
| 13.9831862 |
| 37.8510384 |
| 13.0188285 |
| 30.9414158 |
| 30.8594454 |
| 45.8069892 |
| 4.58069892 |
| 21.6980475 |
| 7.71486134 |
| 13.0188285 |
| 5.786146 |
| 11.0901132 |
| 40.503022 |
| 22.9034946 |
| 12.0544708 |
| 9.16139784 |
| 7.2326825 |
| 4.3396095 |
| 308.594454 |
| 17.8406168 |
| 10.3909539 |
| 8.19704017 |
| 14.465365 |
| 32.5470713 |
| 11.572292 |
| 12.5366497 |

| |
|---|
| 11.572292 |
| 13.5010073 |
| 90.6496207 |
| 8.19704017 |
| 27.9663724 |
| 4.3396095 |
| 16.3940803 |
| 11.572292 |
| 17.358438 |
| 12.0544708 |
| 376.166995 |
| 22.1802263 |
| 18.3227957 |
| 9.88466609 |
| 5.30396717 |
| 200.586395 |
| 26.5198358 |
| 14.465365 |
| 8.19704017 |
| 21.6980475 |
| 15.4297227 |
| 9.16139784 |
| 5.786146 |
| 12.5366497 |
| 24.1089417 |
| 5.786146 |
| 15.9119015 |
| 8.67921901 |
| 5.786146 |
| 6.75050367 |
| 9.64357667 |
| 21.6980475 |
| 22.6624052 |
| 32.7881607 |
| 7.2326825 |
| 5.30396717 |
| 92.5783361 |
| 8.67921901 |
| 10.6079343 |
| 24.5911205 |
| 17.358438 |
| 12.0544708 |
| 25.0732993 |
| 7.07356349 |
| 22.1802263 |
| 10.1257555 |

| |
|---|
| 15.4297227 |
| 16.8762592 |
| 23.144584 |
| 13.0188285 |
| 40.503022 |
| 6.26832484 |
| 19.0460639 |
| 16.8762592 |
| 9.16139784 |
| 33.0292501 |
| 7.2326825 |
| 12.0544708 |
| 17.8406168 |
| 29.4129089 |
| 10.1257555 |
| 21.2158687 |
| 7.71486134 |
| 5.30396717 |
| 7.71486134 |
| 30.8594454 |
| 18.3227957 |
| 15.9119015 |
| 13.0188285 |
| 4.82178834 |
| 696.266236 |
| 5.786146 |
| 18.3227957 |
| 11.572292 |
| 33.2703395 |
| 27.4841935 |
| 91.1317996 |
| 23.6267628 |
| 12.5366497 |
| 11.0901132 |
| 5.30396717 |
| 23.144584 |
| 56.4149235 |
| 11.0901132 |
| 50.1465987 |
| 9.16139784 |
| 53.5218505 |
| 20.251511 |
| 6.75050367 |
| 41.9495585 |
| 11.8133814 |
| 17.358438 |

| |
|---|
| 10.6079343 |
| 21.2158687 |
| 5.786146 |
| 24.1089417 |
| 7.71486134 |
| 63.1654272 |
| 11.572292 |
| 16.8762592 |
| 27.4841935 |
| 23.144584 |
| 10.6079343 |
| 6.26832484 |
| 28.93073 |
| 61.7188907 |
| 6.75050367 |
| 10.6079343 |
| 7.2326825 |
| 11.572292 |
| 14.7064544 |
| 36.1634125 |
| 64.6119637 |
| 73.7733615 |
| 10.6079343 |
| 7.2326825 |
| 9.64357667 |
| 13.0188285 |
| 11.0901132 |
| 298.849619 |
| 9.16139784 |
| 11.572292 |
| 14.9475438 |
| 13.6601264 |
| 12.0544708 |
| 8.19704017 |
| 28.93073 |
| 40.0208432 |
| 19.2871533 |
| 34.716876 |
| 9.64357667 |
| 9.16139784 |
| 8.19704017 |
| 11.0901132 |
| 8.67921901 |
| 25.0732993 |
| 18.8049745 |
| 13.5010073 |

| |
|---|
| 26.037657 |
| 12.2955603 |
| 963.39331 |
| 16.3940803 |
| 15.9119015 |
| 21.6980475 |
| 154.779406 |
| 4.3396095 |
| 4.82178834 |
| 24.5911205 |
| 25.5554782 |
| 10.1257555 |
| 7.47377192 |
| 4.82178834 |
| 31.3416242 |
| 29.8950877 |
| 10.6079343 |
| 4.3396095 |
| 8.19704017 |
| 10.6079343 |
| 5.786146 |
| 5.786146 |
| 32.0648924 |
| 21.2158687 |
| 123.437781 |
| 4.3396095 |
| 10.6079343 |
| 14.465365 |
| 12.5366497 |
| 20.251511 |
| 25.5554782 |
| 18.8049745 |
| 4.82178834 |
| 5.786146 |
| 7.2326825 |
| 51.1109564 |
| 12.5366497 |
| 10.6079343 |
| 11.572292 |
| 84.1402065 |
| 13.5010073 |
| 19.7693322 |
| 1587.33272 |
| 15.4297227 |
| 12.5366497 |
| 17.358438 |

| |
|---|
| 12.0544708 |
| 9.16139784 |
| 62.5241294 |
| 9.64357667 |
| 5.786146 |
| 30.8594454 |
| 19.2871533 |
| 13.0188285 |
| 19.2871533 |
| 14.465365 |
| 8.19704017 |
| 8.67921901 |
| 22.6624052 |
| 7.2326825 |
| 9.16139784 |
| 12.5366497 |
| 21.2158687 |
| 6.26832484 |
| 9.16139784 |
| 7.67628703 |
| 9.64357667 |
| 18.8049745 |
| 30.8594454 |
| 12.0544708 |
| 7.71486134 |
| 6.26832484 |
| 9.16139784 |
| 3.85743067 |
| 14.9475438 |
| 4.3396095 |
| 7.71486134 |
| 10.1257555 |
| 491.82241 |
| 2.65198358 |
| 14.9475438 |
| 15.4297227 |
| 18.3227957 |
| 13.0188285 |
| 5.06287775 |
| 7.71486134 |
| 11.572292 |
| 8.19704017 |
| 9.16139784 |
| 10.1257555 |
| 14.9475438 |
| 77.6307922 |

| |
|---|
| 21.6980475 |
| 9.88466609 |
| 26.037657 |
| 8.19704017 |
| 13.0188285 |
| 9.64357667 |
| 10.1257555 |
| 5.786146 |
| 22.1802263 |
| 5.30396717 |
| 26.5198358 |
| 8.19704017 |
| 32.7881607 |
| 7.2326825 |
| 10.6079343 |
| 10.6079343 |
| 6.75050367 |
| 8.19704017 |
| 26.037657 |
| 18.3227957 |
| 18.8049745 |
| 11.0901132 |
| 4.3396095 |
| 27.0020147 |
| 6.75050367 |
| 59.3079965 |
| 5.786146 |
| 27.0020147 |
| 11.572292 |
| 37.1277702 |
| 6.26832484 |
| 5.30396717 |
| 17.8406168 |
| 8.19704017 |
| 25.7965676 |
| 16.8762592 |
| 15.4297227 |
| 28.4485512 |
| 20.251511 |
| 38.0921279 |
| 7.2326825 |
| 4.3396095 |
| 6.75050367 |
| 16.8762592 |
| 44.3604527 |
| 11.0901132 |

| |
|---|
| 13.0188285 |
| 4.82178834 |
| 25.5554782 |
| 6.26832484 |
| 32.0648924 |
| 16.8762592 |
| 10.1257555 |
| 13.9831862 |
| 4.82178834 |
| 26.5198358 |
| 73.7733615 |
| 11.0901132 |
| 33.7525184 |
| 9.64357667 |
| 9.16139784 |
| 18.3227957 |
| 17.8406168 |
| 21.2158687 |
| 15.4297227 |
| 9.64357667 |
| 4.82178834 |
| 4.82178834 |
| 44.8426315 |
| 5.786146 |
| 9.16139784 |
| 101.257555 |
| 14.9475438 |
| 19.4076981 |
| 12.0544708 |
| 11.0901132 |
| 33.2703395 |
| 7.71486134 |
| 24.1089417 |
| 13.2599179 |
| 14.9475438 |
| 11.0901132 |
| 14.465365 |
| 19.2871533 |
| 7.2326825 |
| 10.1257555 |
| 6.99159309 |
| 59.3079965 |
| 3.61634125 |
| 5.786146 |
| 4.82178834 |
| 18.8049745 |

| |
|---|
| 38.0921279 |
| 8.19704017 |
| 18.8049745 |
| 18.3227957 |
| 4.55176819 |
| 22.6624052 |
| 7.71486134 |
| 10.1257555 |
| 6.75050367 |
| 19.7693322 |
| 27.4841935 |
| 29.4129089 |
| 41.4673797 |
| 8.19704017 |
| 9.64357667 |
| 8.19704017 |
| 2.41089417 |
| 9.64357667 |
| 172.620022 |
| 29.8950877 |
| 7.71486134 |
| 4.82178834 |
| 18.2504689 |
| 29.8950877 |
| 4.3396095 |
| 30.3772665 |
| 13.5010073 |
| 60.3543246 |
| 31.823803 |
| 29.8950877 |
| 118.133814 |
| 15.9119015 |
| 14.465365 |
| 11.0901132 |
| 14.9475438 |
| 23.6267628 |
| 4.98090735 |
| 5.786146 |
| 23.144584 |
| 7.71486134 |
| 19.7693322 |
| 14.9475438 |
| 12.0544708 |
| 23.144584 |
| 10.1257555 |
| 20.251511 |

| |
|---|
| 8.67921901 |
| 70.6054466 |
| 5.06287775 |
| 10.6079343 |
| 59.549086 |
| 12.7777391 |
| 9.40248726 |
| 46.289168 |
| 8.67921901 |
| 31.3416242 |
| 33.7525184 |
| 25.5554782 |
| 45.8069892 |
| 8.67921901 |
| 5.786146 |
| 21.2158687 |
| 16.1529909 |
| 12.0544708 |
| 82.4525806 |
| 14.9475438 |
| 17.8406168 |
| 9.64357667 |
| 5.30396717 |
| 36.6455914 |
| 5.30396717 |
| 9.16139784 |
| 9.64357667 |
| 8.05238652 |
| 71.8446462 |
| 23.144584 |
| 27.9663724 |
| 27.4841935 |
| 19.2871533 |
| 5.30396717 |
| 67.0228579 |
| 9.64357667 |
| 38.5743067 |
| 13.5010073 |
| 9.16139784 |
| 38.0921279 |
| 4.82178834 |
| 13.9831862 |
| 9.64357667 |
| 9.64357667 |
| 28.93073 |
| 4.82178834 |

| |
|---|
| 24.1089417 |
| 8.67921901 |
| 14.9475438 |
| 16.8762592 |
| 12.5366497 |
| 5.786146 |
| 7.2326825 |
| 16.3940803 |
| 13.5010073 |
| 19.2871533 |
| 8.19704017 |
| 7.2326825 |
| 21.6980475 |
| 14.465365 |
| 6.26832484 |
| 13.9831862 |
| 11.0901132 |
| 17.358438 |
| 283.038975 |
| 15.4297227 |
| 25.0732993 |
| 8.19704017 |
| 54.0040294 |
| 15.9119015 |
| 28.4485512 |
| 41.4673797 |
| 18.3227957 |
| 44.8426315 |
| 151.886333 |
| 6.75050367 |
| 5.786146 |
| 165.869519 |
| 8.67921901 |
| 31.3416242 |
| 8.19704017 |
| 5.30396717 |
| 21.6980475 |
| 1.92871533 |
| 8.67921901 |
| 16.8762592 |
| 40.503022 |
| 23.6267628 |
| 7.47377192 |
| 13.0188285 |
| 27.6963522 |
| 3.37525184 |

| |
|---|
| 30.8594454 |
| 20.7336898 |
| 7.2326825 |
| 17.358438 |
| 11.572292 |
| 34.716876 |
| 10.1257555 |
| 5.30396717 |
| 169.244771 |
| 6.26832484 |
| 11.0901132 |
| 11.0901132 |
| 8.19704017 |
| 25.4638642 |
| 46.7713469 |
| 90.6496207 |
| 216.257207 |
| 54.0040294 |
| 12.5366497 |
| 23.144584 |
| 25.5554782 |
| 19.0798164 |
| 35.1990549 |
| 47.2535257 |
| 12.7777391 |
| 8.19704017 |
| 35.6812337 |
| 9.07942744 |
| 12.5366497 |
| 37.609949 |
| 4.3396095 |
| 20.251511 |
| 42.9139162 |
| 15.9119015 |
| 15.4297227 |
| 8.19704017 |
| 44.1772247 |
| 6.26832484 |
| 19.2871533 |
| 9.64357667 |
| 4.3396095 |
| 9.64357667 |
| 20.251511 |
| 7.71486134 |
| 35.4401443 |
| 13.0188285 |

| |
|---|
| 39.0564855 |
| 62.6832484 |
| 16.8762592 |
| 12.0544708 |
| 9.64357667 |
| 24.5911205 |
| 31.823803 |
| 6.75050367 |
| 8.19704017 |
| 12.2714513 |
| 8.269367 |
| 11.572292 |
| 5.30396717 |
| 14.465365 |
| 10.6079343 |
| 165.869519 |
| 8.19704017 |
| 34.716876 |
| 27.0020147 |
| 18.0817063 |
| 59.3079965 |
| 16.3940803 |
| 6.75050367 |
| 32.7881607 |
| 9.97145828 |
| 15.4297227 |
| 8.67921901 |
| 8.19704017 |
| 36.6455914 |
| 13.0188285 |
| 8.19704017 |
| 15.9119015 |
| 6.26832484 |
| 11.8133814 |
| 14.465365 |
| 42.4317374 |
| 32.7881607 |
| 18.3227957 |
| 20.7336898 |
| 113.312026 |
| 17.9900923 |
| 30.3772665 |
| 9.64357667 |
| 8.19704017 |
| 20.251511 |
| 10.6079343 |

| |
|---|
| 4.82178834 |
| 23.144584 |
| 47.4946151 |
| 5.54505659 |
| 27.0020147 |
| 12.5366497 |
| 57.86146 |
| 93.0605149 |
| 10.1257555 |
| 30.8594454 |
| 36.6455914 |
| 18.3227957 |
| 21.2158687 |
| 13.5010073 |
| 9.64357667 |
| 8.67921901 |
| 22.1802263 |
| 7.2326825 |
| 33.2703395 |
| 12.0544708 |
| 13.0188285 |
| 10.1257555 |
| 17.2812894 |
| 19.7693322 |
| 5.786146 |
| 56.4149235 |
| 9.40248726 |
| 66.0585002 |
| 36.1634125 |
| 27.0020147 |
| 4.3396095 |
| 4.82178834 |
| 10.6079343 |
| 4.3396095 |
| 5.786146 |
| 33.2703395 |
| 7.71486134 |
| 9.16139784 |
| 13.2068783 |
| 18.3227957 |
| 20.7336898 |
| 18.3227957 |
| 29.4129089 |
| 16.3940803 |
| 5.786146 |
| 33.7525184 |

| |
|---|
| 14.465365 |
| 9.16139784 |
| 8.67921901 |
| 14.465365 |
| 49.6644199 |
| 10.6079343 |
| 5.30396717 |
| 45.3248104 |
| 26.5198358 |
| 9.16139784 |
| 155.261584 |
| 22.6624052 |
| 13.0188285 |
| 7.71486134 |
| 176.959632 |
| 20.251511 |
| 39.5386644 |
| 16.1529909 |
| 5.786146 |
| 13.0188285 |
| 6.26832484 |
| 15.9119015 |
| 11.0901132 |
| 9.16139784 |
| 15.4297227 |
| 9.16139784 |
| 28.4485512 |
| 5.786146 |
| 5.786146 |
| 22.1802263 |
| 18.8049745 |
| 13.9831862 |
| 5.786146 |
| 15.4297227 |
| 89.2030842 |
| 5.30396717 |
| 15.9119015 |
| 8.19704017 |
| 20.251511 |
| 6.75050367 |
| 48.2178834 |
| 39.0564855 |
| 5.30396717 |
| 14.9475438 |
| 27.0020147 |
| 31.3416242 |

| |
|---|
| 76.1842557 |
| 27.0020147 |
| 107.043701 |
| 22.6624052 |
| 18.3227957 |
| 9.64357667 |
| 4.3396095 |
| 21.2158687 |
| 18.8049745 |
| 13.9831862 |
| 20.7336898 |
| 9.16139784 |
| 7.39180152 |
| 67.5050367 |
| 30.3772665 |
| 18.8049745 |
| 19.2871533 |
| 47.9767939 |
| 16.8762592 |
| 7.2326825 |
| 21.2158687 |
| 28.4485512 |
| 6.26832484 |
| 28.6896406 |
| 4.3396095 |
| 12.0544708 |
| 6.26832484 |
| 31.823803 |
| 5.786146 |
| 4.3396095 |
| 6.26832484 |
| 16.3940803 |
| 10.6079343 |
| 12.2955603 |
| 63.647606 |
| 23.6267628 |
| 4.82178834 |
| 41.9495585 |
| 90.1674419 |
| 6.75050367 |
| 6.75050367 |
| 15.9119015 |
| 11.0901132 |
| 6.75050367 |
| 27.9663724 |
| 8.19704017 |

| |
|---|
| 129.706106 |
| 9.16139784 |
| 9.16139784 |
| 65.5763214 |
| 12.0544708 |
| 5.786146 |
| 27.9663724 |
| 5.30396717 |
| 91.1317996 |
| 6.26832484 |
| 18.3227957 |
| 7.2326825 |
| 16.5531994 |
| 8.19704017 |
| 22.9034946 |
| 19.0460639 |
| 11.572292 |
| 30.6183559 |
| 17.358438 |
| 11.572292 |
| 998.592364 |
| 10.6079343 |
| 30.8594454 |
| 16.8762592 |
| 17.358438 |
| 16.8762592 |
| 27.9663724 |
| 5.06287775 |
| 13.9831862 |
| 20.251511 |
| 14.9475438 |
| 14.465365 |
| 25.5554782 |
| 7.71486134 |
| 3.85743067 |
| 9.16139784 |
| 22.6624052 |
| 17.517557 |
| 61.2367119 |
| 59.7901754 |
| 13.5010073 |
| 5.30396717 |
| 9.16139784 |
| 41.4673797 |
| 8.92030842 |
| 30.8594454 |

| |
|---|
| 5.54505659 |
| 15.4297227 |
| 147.546723 |
| 18.3227957 |
| 4.82178834 |
| 4.3396095 |
| 16.8762592 |
| 21.6980475 |
| 19.2871533 |
| 17.8406168 |
| 13.2599179 |
| 5.30396717 |
| 13.0815118 |
| 14.465365 |
| 13.0188285 |
| 54.3270892 |
| 14.465365 |
| 10.1257555 |
| 9.64357667 |
| 11.0901132 |
| 13.5010073 |
| 24.1089417 |
| 23.6267628 |
| 12.0544708 |
| 21.1338983 |
| 43.396095 |
| 5.30396717 |
| 11.0901132 |
| 10.6079343 |
| 8.19704017 |
| 54.968387 |
| 14.465365 |
| 189.014103 |
| 3.77546027 |
| 25.0732993 |
| 27.4841935 |
| 27.9663724 |
| 40.0208432 |
| 7.2326825 |
| 18.8049745 |
| 6.26832484 |
| 4.3396095 |
| 32.3059819 |
| 31.4235946 |
| 27.4359756 |
| 13.9831862 |

| |
|---|
| 8.67921901 |
| 10.6079343 |
| 4.82178834 |
| 10.1257555 |
| 14.465365 |
| 12.0544708 |
| 17.8406168 |
| 5.96937396 |
| 12.7680955 |
| 14.238741 |
| 14.9475438 |
| 18.3227957 |
| 17.517557 |
| 15.9119015 |
| 9.88466609 |
| 97.8823032 |
| 13.0188285 |
| 16.3940803 |
| 9.16139784 |
| 58.3436389 |
| 14.2242756 |
| 6.02723542 |
| 28.93073 |
| 8.19704017 |
| 13.5010073 |
| 30.3772665 |
| 6.02723542 |
| 5.30396717 |
| 18.7230041 |
| 11.0901132 |
| 8.19704017 |
| 28.93073 |
| 16.3940803 |
| 12.7777391 |
| 12.0544708 |
| 12.5366497 |
| 15.4297227 |
| 9.40248726 |
| 51.1109564 |
| 84.3812959 |
| 47.7357045 |
| 13.0188285 |
| 84.7043557 |
| 40.503022 |
| 11.572292 |
| 32.918349 |

| |
|---|
| 4.3396095 |
| 50.6673518 |
| 16.0661987 |
| 26.5198358 |
| 13.0188285 |
| 10.1257555 |
| 14.465365 |
| 10.1257555 |
| 7.80647532 |
| 46.289168 |
| 28.93073 |
| 20.251511 |
| 55.4891402 |
| 7.71486134 |
| 35.6812337 |
| 156.225942 |
| 87.7565477 |
| 8.19704017 |
| 46.289168 |
| 11.8133814 |
| 9.16139784 |
| 18.3227957 |
| 19.6102132 |
| 17.358438 |
| 40.0208432 |
| 6.75050367 |
| 7.71486134 |
| 10.8972416 |
| 10.7670534 |
| 115.240741 |
| 16.3940803 |
| 13.0188285 |
| 4.82178834 |
| 9.16139784 |
| 11.0901132 |
| 8.67921901 |
| 11.9725004 |
| 9.64357667 |
| 7.47377192 |
| 7.2326825 |
| 8.67921901 |
| 540.522472 |
| 26.5198358 |
| 73.7685398 |
| 7.2326825 |
| 6.26832484 |

| |
|---|
| 27.9663724 |
| 14.9475438 |
| 7.71486134 |
| 6.26832484 |
| 7.2326825 |
| 5.786146 |
| 26.8043214 |
| 32.2143679 |
| 17.358438 |
| 3.85743067 |
| 9.40248726 |
| 54.4862082 |
| 14.7257416 |
| 2.893073 |
| 9.64357667 |
| 20.936205 |
| 326.917249 |
| 13.5010073 |
| 14.465365 |
| 14.9475438 |
| 15.2706037 |
| 13.2599179 |
| 4.82178834 |
| 8.67921901 |
| 12.0544708 |
| 17.9997359 |
| 6.75050367 |
| 13.5010073 |
| 5.786146 |
| 6.75050367 |
| 12.0544708 |
| 40.4210516 |
| 11.0901132 |
| 30.4592369 |
| 142.242756 |
| 13.5010073 |
| 60.4314732 |
| 7.2326825 |
| 30.8594454 |
| 5.786146 |
| 41.4673797 |
| 21.2158687 |
| 14.465365 |
| 14.9475438 |
| 31.823803 |
| 17.1173486 |

| |
|---|
| 11.0901132 |
| 11.0901132 |
| 10.6079343 |
| 11.572292 |
| 46.289168 |
| 33.7525184 |
| 31.823803 |
| 10.1257555 |
| 12210.6968 |
| 45348.9193 |
| 164.422982 |
| 41.2262903 |
| 1016.19189 |
| 171.737635 |
| 111.383311 |
| 136.45661 |
| 11.3312026 |
| 94.7481408 |
| 118.615993 |
| 129.706106 |
| 2.87860764 |
| 3.43793508 |
| 5.82472031 |
| 10.2559438 |
| 606.228982 |
| 21.6980475 |
| 2.87860764 |
| 12.5366497 |
| 2.87860764 |
| 3.91047034 |
| 1.79370526 |
| 55.8218436 |
| 69.5639403 |
| 2.87860764 |
| 14.465365 |
| 58.8258177 |
| 8.67921901 |
| 8.19704017 |
| 13.5010073 |
| 59.7901754 |
| 33.2703395 |
| 10.1257555 |
| 6.75050367 |
| 8.19704017 |
| 35.719808 |
| 15.8492183 |

18.0817063
30.3772665
9.64357667
7.71486134
17.1173486
17.8406168
14.9475438
34.9579654
25.0732993
17.8020425
6.26832484
6.75050367
7.71486134
4.3396095
26.5198358
10.1257555
4.82178834
36.6455914
5.30396717
9.40248726
17.358438
10.6079343
31.1005348
4.82178834
15.9119015
143.689292
5.786146
27.2431041
28.6896406
24.5911205
10.2463002
5.786146
4.3396095
7.71486134
13.0188285
46.289168
4.3396095
11.0901132
7.71486134
2.41089417
11.0901132
7.71486134
8.35615919
21.6980475
16.3940803
305.219202

| |
|---|
| 18.3227957 |
| 30.3772665 |
| 7.2326825 |
| 9.16139784 |
| 17.358438 |
| 6.26832484 |
| 12.0544708 |
| 8.67921901 |
| 8.67921901 |
| 11.572292 |
| 22.6624052 |
| 20.7336898 |
| 26.037657 |
| 6.26832484 |
| 7.71486134 |
| 30.4978112 |
| 96.4357667 |
| 8.67921901 |
| 22.1802263 |
| 68.3102754 |
| 5.786146 |
| 9.64357667 |
| 6.75050367 |
| 11.0901132 |
| 7.2326825 |
| 7.2326825 |
| 6.93855342 |
| 16.3940803 |
| 10.6079343 |
| 17.358438 |
| 5.30396717 |
| 19.2871533 |
| 4.3396095 |
| 4.3396095 |
| 27.0020147 |
| 12.3630653 |
| 10.1257555 |
| 15.6708121 |
| 658.651465 |
| 17.5464878 |
| 13.9831862 |
| 153.815048 |
| 16.8762592 |
| 24.1089417 |
| 8.67921901 |
| 20.4926004 |

| |
|---|
| 64.6119637 |
| 50.8071837 |
| 33.5114289 |
| 5.786146 |
| 87.2743689 |
| 4.3396095 |
| 720.375177 |
| 18.3227957 |
| 39.5386644 |
| 27.0020147 |
| 77.1486134 |
| 5.30396717 |
| 44.3604527 |
| 7.2326825 |
| 120.06253 |
| 4.82178834 |
| 34.2346972 |
| 83.8991171 |
| 54.4042378 |
| 14.465365 |
| 13.5010073 |
| 506.109369 |
| 92.7953165 |
| 11.572292 |
| 6.26832484 |
| 39.0564855 |
| 7.71486134 |
| 9.64357667 |
| 14.9475438 |
| 40.0208432 |
| 8.67921901 |
| 13.0188285 |
| 12.0544708 |
| 31.3416242 |
| 15.0680886 |
| 5.6656013 |
| 6.02723542 |
| 12.5366497 |
| 10.6079343 |
| 17.8406168 |
| 34.2346972 |
| 6.75050367 |
| 14.9475438 |
| 15.9119015 |
| 8.76118941 |
| 14.2242756 |

| |
|---|
| 74.2555404 |
| 17.8406168 |
| 31.5007432 |
| 7.2326825 |
| 48.2178834 |
| 37.8510384 |
| 34.716876 |
| 11.572292 |
| 8.92030842 |
| 15.4297227 |
| 18.8049745 |
| 13.9012158 |
| 13.9012158 |
| 3.85743067 |
| 15.4297227 |
| 10.1257555 |
| 146.100187 |
| 15.9119015 |
| 24.1089417 |
| 5.14484815 |
| 18.0961716 |
| 7.2326825 |
| 14.9475438 |
| 41.2262903 |
| 4.59516428 |
| 55.9327447 |
| 20.1695406 |
| 4.82178834 |
| 15.4297227 |
| 32.4265266 |
| 9.07942744 |
| 10.6079343 |
| 7.2326825 |
| 12.0544708 |
| 9.16139784 |
| 5.30396717 |
| 15.4297227 |
| 6.75050367 |
| 10.728479 |
| 19.7693322 |
| 12.5366497 |
| 7.32429648 |
| 8.67921901 |
| 5.786146 |
| 40.9852009 |
| 13.5010073 |

| |
|---|
| 11.572292 |
| 11.572292 |
| 10.8490238 |
| 8.19704017 |
| 11.0901132 |
| 23.144584 |
| 337.043005 |
| 6.75050367 |
| 10.1257555 |
| 351.990549 |
| 15.9119015 |
| 15.2030986 |
| 3.61634125 |
| 8.72743689 |
| 14.2242756 |
| 13.5154727 |
| 21.6980475 |
| 10.7670534 |
| 7.2326825 |
| 13.0188285 |
| 103.668449 |
| 577.650243 |
| 21.2158687 |
| 9.64357667 |
| 2.73877577 |
| 24.4223579 |
| 74.5737784 |
| 63.0014864 |
| 6.2827902 |
| 3.6790245 |
| 6.2442159 |
| 258.770915 |
| 7.71486134 |
| 9.64357667 |
| 3.46204403 |
| 6.2827902 |
| 19.5427081 |
| 3.46204403 |
| 68.9322861 |
| 11.8133814 |
| 12.5028972 |
| 3.6790245 |
| 5.76203706 |
| 6.2442159 |
| 3.91047034 |
| 20.0104216 |

| |
|---|
| 27.9663724 |
| 24.7020216 |
| 20.0104216 |
| 6.76496904 |
| 12.5366497 |
| 118.697963 |
| 6.2827902 |
| 6.2827902 |
| 6.2827902 |
| 285.29075 |
| 18.9833807 |
| 17.4018341 |
| 97.8823032 |
| 6.2827902 |
| 3.91047034 |
| 8.92030842 |
| 10.6079343 |
| 10.1257555 |
| 74.7377192 |
| 409.528949 |
| 14.465365 |
| 7.2326825 |
| 871.779331 |
| 461.686233 |
| 6.75050367 |
| 22.0982559 |
| 211.54632 |
| 211.54632 |
| 29.8950877 |
| 30.3772665 |
| 29.0271658 |
| 5.62702699 |
| 7.90773287 |
| 5.786146 |
| 24.1089417 |
| 64.6939341 |
| 41.4673797 |
| 11.6976585 |
| 13.0188285 |
| 12.5366497 |
| 40.503022 |
| 6.26832484 |
| 20.251511 |
| 11.572292 |
| 4.90375874 |
| 314000.329 |

| |
|---|
| 12.5366497 |
| 8.19704017 |
| 4.82178834 |
| 44.8426315 |
| 11.572292 |
| 5.30396717 |
| 13.0188285 |
| 24.1089417 |
| 13.5010073 |
| 9.64357667 |
| 5.30396717 |
| 8.19704017 |
| 16.3940803 |
| 40.0208432 |
| 49.6644199 |
| 7.71486134 |
| 13.5010073 |
| 5.30396717 |
| 50.1465987 |
| 18.3227957 |
| 5.30396717 |
| 36.1634125 |
| 6.26832484 |
| 380.921279 |
| 5.30396717 |
| 42.9139162 |
| 6.26832484 |
| 6.26832484 |
| 4.3396095 |
| 9.64357667 |
| 1.92871533 |
| 11.572292 |
| 19.7693322 |
| 13.0188285 |
| 8.19704017 |
| 7.71486134 |
| 10.6079343 |
| 4.82178834 |
| 28.93073 |
| 4.82178834 |
| 12.5366497 |
| 8.67921901 |
| 19.7693322 |
| 10.6079343 |
| 14.9475438 |
| 5.786146 |

| HS548_p_values | | HS548_p_values | | HS548_p_values | | HS548_p_v |
|---|---|---|---|---|---|---|
| Ens_ID | Gene | Chrom | Strand | Start | End | RNA_type |
| ENSMUSG00000032423 | Syncrip | chr9 | - | 88447008 | 88482574 | regular |
| ENSMUSG00000057098 | Ebf1 | chr11 | + | 44617316 | 45008091 | regular |
| ENSMUSG00000004980 | Hnrnpa2b1 | chr6 | - | 51460931 | 51469894 | regular |
| ENSMUSG00000071550 | Cfap44 | chr16 | + | 44394795 | 44482428 | regular |
| ENSMUSG00000029076 | Sdf4 | chr4 | + | 155992871 | 156013610 | regular |
| ENSMUSG00000022443 | Myh9 | chr15 | - | 77760586 | 77842175 | regular |
| ENSMUST00000046656 | Tasp1 | chr2 | - | 140008698 | 140057499 | circular |
| ENSMUSG00000037674 | Rfx7 | chr9 | + | 72532239 | 72622937 | regular |
| ENSMUSG00000053774 | Ubxn7 | chr16 | + | 32332251 | 32393747 | regular |
| ENSMUSG00000063409 | Lrrc43 | chr5 | + | 123489304 | 123508205 | regular |
| ENSMUSG00000025983 | Ccdc150 | chr1 | + | 54250682 | 54368727 | regular |
| ENSMUSG00000024008 | Cpne5 | chr17 | - | 29156549 | 29237797 | regular |
| ENSMUSG00000019850 | Tnfaip3 | chr10 | - | 19000909 | 19015657 | regular |
| ENSMUSG00000029204 | Rhoh | chr5 | + | 65861212 | 65896700 | regular |
| ENSMUSG00000027344 | Fsip1 | chr2 | - | 118204887 | 118256966 | regular |
| ENSMUSG00000091971 | Hspa1a | chr17 | - | 34969189 | 34972156 | regular |
| ENSMUSG00000101855 | 1700010N08 | chr7 | + | 38791660 | 38860532 | regular |
| ENSMUSG00000106106 | CT010467.1 | chr17 | - | 39846352 | 39848827 | regular |
| ENSMUST00000168527 | Dennd1b | chr1 | + | 139133670 | 139144000 | circular |
| ENSMUSG00000053819 | Camk2d | chr3 | + | 126596301 | 126846326 | regular |
| ENSMUSG00000020580 | Rock2 | chr12 | + | 16894894 | 16987823 | regular |
| ENSMUSG00000058587 | Tmod3 | chr9 | - | 75497706 | 75559657 | regular |
| ENSMUSG00000029439 | Sfswap | chr5 | + | 129501220 | 129571384 | regular |
| ENSMUSG00000026832 | Cytip | chr2 | - | 58129136 | 58195532 | regular |
| ENSMUSG00000020863 | Luc7l3 | chr11 | - | 94287889 | 94321988 | regular |
| ENSMUSG00000021188 | Trip11 | chr12 | - | 101834042 | 101913267 | regular |
| ENSMUSG00000034892 | Rps29 | chr12 | - | 69157721 | 69159186 | regular |
| ENSMUSG00000097589 | Dleu2 | chr14 | - | 61602838 | 61682373 | regular |
| ENSMUSG00000026305 | Lrrfip1 | chr1 | + | 90998736 | 91128944 | regular |
| ENSMUSG00000043962 | Thrap3 | chr4 | - | 126164081 | 126202760 | regular |
| ENSMUSG00000026094 | Stk17b | chr1 | - | 53755505 | 53785224 | regular |
| ENSMUSG00000054387 | Mdm4 | chr1 | - | 132959483 | 133030561 | regular |
| ENSMUSG00000002107 | Celf2 | chr2 | - | 6539693 | 7509563 | regular |
| ENSMUSG00000015932 | Dstn | chr2 | + | 143915319 | 143943324 | regular |
| ENSMUSG00000038855 | Itpkb | chr1 | + | 180330484 | 180424802 | regular |
| ENSMUSG00000013629 | Cad | chr5 | + | 31054779 | 31078479 | regular |
| ENSMUSG00000034042 | Gpbp1l1 | chr4 | + | 116557657 | 116593882 | regular |
| ENSMUSG00000027012 | Dync1i2 | chr2 | + | 71211705 | 71263303 | regular |
| ENSMUSG00000032400 | Zwilch | chr9 | - | 64137143 | 64173104 | regular |
| ENSMUSG00000032185 | Carm1 | chr9 | + | 21546893 | 21592623 | regular |
| ENSMUSG00000022255 | Mtdh | chr15 | + | 34082693 | 34145624 | regular |
| ENSMUSG00000024974 | Smc3 | chr19 | + | 53600397 | 53645833 | regular |
| ENSMUSG00000028582 | Cc2d1b | chr4 | + | 108619939 | 108634123 | regular |
| ENSMUSG00000031751 | Amfr | chr8 | - | 93971587 | 94012842 | regular |

| | | | | | | |
|---|---|---|---|---|---|---|
| ENSMUSG00000022533 | Atp13a3 | chr16 | - | 30312422 | 30388837 | regular |
| ENSMUSG00000026395 | Ptprc | chr1 | - | 138062860 | 138175708 | regular |
| ENSMUSG00000027620 | Rbm39 | chr2 | - | 156147238 | 156180238 | regular |
| ENSMUSG00000075415 | Fnbp1 | chr2 | - | 31026205 | 31142008 | regular |
| ENSMUSG00000027722 | Spata5 | chr3 | + | 37419895 | 37579096 | regular |
| ENSMUSG00000007850 | Hnrnph1 | chr11 | + | 50376989 | 50386528 | regular |
| ENSMUSG00000056537 | Rlim | chrX | - | 103957162 | 103981284 | regular |
| ENSMUSG00000049506 | Sppl2c | chr11 | + | 104186326 | 104191163 | regular |
| ENSMUSG00000024306 | Ccdc178 | chr18 | - | 21810896 | 22171396 | regular |
| ENSMUSG00000052512 | Nav2 | chr7 | + | 48908715 | 49610090 | regular |
| ENSMUSG00000040524 | Zfp609 | chr9 | - | 65692390 | 65827564 | regular |
| ENSMUSG00000053332 | Gas5 | chr1 | + | 161034421 | 161038539 | regular |
| ENSMUSG00000098178 | Gm42418 | chr17 | + | 39846957 | 39848788 | regular |
| ENSMUSG00000032869 | Psmf1 | chr2 | - | 151715811 | 151744186 | regular |
| ENSMUSG00000079235 | Ccdc13 | chr9 | - | 121797626 | 121839461 | regular |
| ENSMUSG00000021767 | Kat6b | chr14 | + | 21481433 | 21672478 | regular |
| ENSMUSG00000049295 | Zfp219 | chr14 | - | 52006076 | 52020733 | regular |
| ENSMUSG00000026034 | Clk1 | chr1 | - | 58410188 | 58424066 | regular |
| ENSMUSG00000033228 | Scaf11 | chr15 | - | 96411698 | 96460843 | regular |
| ENSMUSG00000091028 | Gm10722 | chr9 | + | 3000921 | 3002330 | regular |
| ENSMUSG00000078578 | Ube2d3 | chr3 | + | 135438148 | 135468198 | regular |
| ENSMUSG00000031207 | Msn | chrX | + | 96096041 | 96168552 | regular |
| ENSMUSG00000043003 | Rasef | chr4 | - | 73714578 | 73790994 | regular |
| ENSMUSG00000028121 | Bcar3 | chr3 | + | 122294135 | 122530191 | regular |
| ENSMUSG00000035621 | Midn | chr10 | + | 80148271 | 80158368 | regular |
| ENSMUSG00000030016 | Zfp638 | chr6 | + | 83867108 | 83989550 | regular |
| ENSMUSG00000021910 | Nisch | chr14 | - | 31170929 | 31216946 | regular |
| ENSMUSG00000046755 | Kif2b | chr11 | - | 91575314 | 91577558 | regular |
| ENSMUSG00000042228 | Lyn | chr4 | + | 3678114 | 3813122 | regular |
| ENSMUSG00000003847 | Nfat5 | chr8 | + | 107293469 | 107379517 | regular |
| ENSMUSG00000038860 | Garnl3 | chr2 | - | 32986223 | 33131654 | regular |
| ENSMUSG00000027763 | Mbnl1 | chr3 | + | 60472829 | 60629750 | regular |
| ENSMUSG00000092341 | Malat1 | chr19 | - | 5795689 | 5802672 | regular |
| ENSMUSG00000000823 | Zfp512b | chr2 | - | 181582102 | 181592803 | regular |
| ENSMUSG00000025793 | Hgs | chr11 | + | 120467634 | 120483984 | regular |
| ENSMUST00000066715 | Zbtb48 | chr4 | - | 152020189 | 152020708 | circular |
| ENSMUSG00000028820 | Sfpq | chr4 | + | 127021323 | 127037013 | regular |
| ENSMUSG00000021838 | Samd4 | chr14 | + | 46882853 | 47105815 | regular |
| ENSMUSG00000030265 | Kras | chr6 | - | 145216698 | 145250239 | regular |
| ENSMUSG00000047441 | Antxrl | chr14 | + | 34052706 | 34076405 | regular |
| ENSMUSG00000041895 | Wipi1 | chr11 | - | 109573330 | 109611967 | regular |
| ENSMUSG00000028207 | Asph | chr4 | - | 9448068 | 9669344 | regular |
| ENSMUSG00000022201 | Zfr | chr15 | + | 12117830 | 12185683 | regular |
| ENSMUSG00000004018 | Fancl | chr11 | + | 26386134 | 26471876 | regular |
| ENSMUSG00000046434 | Hnrnpa1 | chr15 | + | 103240431 | 103244955 | regular |
| ENSMUSG00000029014 | Dnajc2 | chr5 | - | 21757266 | 21785251 | regular |

| | | | | | | |
|---|---|---|---|---|---|---|
| ENSMUSG00000065773 | Rnu1b6 | chr3 | - | 96460023 | 96460189 | regular |
| ENSMUSG00000095701 | Gm24830 | chr3 | + | 96328424 | 96328590 | regular |
| ENSMUSG00000096838 | Gm26232 | chr3 | + | 96450445 | 96450611 | regular |
| ENSMUSG00000013663 | Pten | chr19 | + | 32757496 | 32826160 | regular |
| ENSMUSG00000025558 | Dock9 | chr14 | - | 121542045 | 121797837 | regular |
| ENSMUSG00000064923 | Gm22042 | chr4 | - | 51812482 | 51812647 | regular |
| ENSMUSG00000054901 | Arhgef33 | chr17 | + | 80307406 | 80400081 | regular |
| ENSMUSG00000094812 | Gm22614 | chr3 | + | 96488728 | 96488893 | regular |
| ENSMUSG00000095260 | Gm25890 | chr3 | - | 96486554 | 96486719 | regular |
| ENSMUSG00000031786 | Drc7 | chr8 | + | 95055102 | 95078141 | regular |
| ENSMUSG00000066406 | Akap13 | chr7 | + | 75455533 | 75754609 | regular |
| ENSMUSG00000029202 | Pds5a | chr5 | - | 65605720 | 65698273 | regular |
| ENSMUSG00000012114 | Med15 | chr16 | - | 17651220 | 17722930 | regular |
| ENSMUSG00000036893 | Ehmt1 | chr2 | - | 24789927 | 24919614 | regular |
| ENSMUSG00000036478 | Btg1 | chr10 | + | 96617005 | 96622809 | regular |
| ENSMUSG00000051329 | Nup160 | chr2 | + | 90677214 | 90736328 | regular |
| ENSMUSG00000035202 | Lars2 | chr9 | + | 123366926 | 123462666 | regular |
| ENSMUSG00000024740 | Ddb1 | chr19 | + | 10605624 | 10629813 | regular |
| ENSMUSG00000029823 | Luc7l2 | chr6 | + | 38551333 | 38609470 | regular |
| ENSMUSG00000020585 | Laptm4a | chr12 | + | 8921663 | 8938742 | regular |
| ENSMUSG00000035133 | Arhgap5 | chr12 | + | 52503971 | 52571975 | regular |
| ENSMUSG00000024006 | Stk38 | chr17 | - | 28970884 | 29007945 | regular |
| ENSMUSG00000002763 | Pex6 | chr17 | + | 46711462 | 46725541 | regular |
| ENSMUSG00000027309 | 4930402H24I | chr2 | - | 130706199 | 130906406 | regular |
| ENSMUSG00000034647 | Ankrd12 | chr17 | - | 65967500 | 66077089 | regular |
| ENSMUSG00000030671 | Pde3b | chr7 | + | 114415280 | 114539251 | regular |
| ENSMUSG00000030533 | Unc45a | chr7 | - | 80325291 | 80341005 | regular |
| ENSMUSG00000061298 | Agbl4 | chr4 | + | 110397660 | 111664324 | regular |
| ENSMUSG00000090558 | Gm17202 | chr5 | - | 107404822 | 107431910 | regular |
| ENSMUSG00000036461 | Elf1 | chr14 | + | 79481193 | 79582494 | regular |
| ENSMUSG00000093834 | Rnu1b2 | chr3 | + | 96374020 | 96374185 | regular |
| ENSMUSG00000095580 | Rnu1b1 | chr3 | - | 96359952 | 96360117 | regular |
| ENSMUSG00000063286 | Gm8995 | chr7 | - | 106504238 | 106511671 | regular |
| ENSMUSG00000027968 | Larp7 | chr3 | - | 127536713 | 127553349 | regular |
| ENSMUSG00000031592 | Pcm1 | chr8 | + | 41239751 | 41332344 | regular |
| ENSMUSG00000023403 | Stk31 | chr6 | + | 49395603 | 49469501 | regular |
| ENSMUSG00000079005 | Gm14147 | chr2 | + | 151098244 | 151103818 | regular |
| ENSMUSG00000062732 | Lypd4 | chr7 | - | 24864619 | 24869941 | regular |
| ENSMUSG00000025862 | Stag2 | chrX | + | 42149316 | 42277185 | regular |
| ENSMUSG00000036552 | Ermard | chr17 | + | 14978863 | 15064232 | regular |
| ENSMUSG00000049354 | Dcaf7 | chr11 | + | 106036871 | 106059324 | regular |
| ENSMUSG00000037922 | Bank1 | chr3 | - | 136053362 | 136326066 | regular |
| ENSMUSG00000065176 | Rnu12 | chr15 | + | 83149643 | 83149794 | regular |
| ENSMUSG00000109170 | Gm45086 | chr7 | - | 48724379 | 48726822 | regular |
| ENSMUSG00000032497 | Lrrfip2 | chr9 | + | 111117591 | 111225668 | regular |
| ENSMUSG00000021339 | Mrs2 | chr13 | - | 24987482 | 25020379 | regular |

| | | | | | | |
|---|---|---|---|---|---|---|
| ENSMUSG00000002968 | Med25 | chr7 | - | 44876764 | 44892712 | regular |
| ENSMUSG00000065701 | Rny1 | chr6 | - | 47788068 | 47788180 | regular |
| ENSMUSG00000096243 | Gm24265 | chr5 | + | 115489458 | 115489599 | regular |
| ENSMUSG00000061322 | Dnaic1 | chr4 | + | 41569774 | 41638158 | regular |
| ENSMUSG00000043633 | Fam221b | chr4 | - | 43659621 | 43669145 | regular |
| ENSMUSG00000040407 | Akap9 | chr5 | + | 3928053 | 4081310 | regular |
| ENSMUSG00000037071 | Scd1 | chr19 | - | 44394450 | 44407709 | regular |
| ENSMUSG00000090877 | Hspa1b | chr17 | - | 34956435 | 34959238 | regular |
| ENSMUSG00000031728 | Zfp821 | chr8 | + | 109705545 | 109724932 | regular |
| ENSMUSG00000063511 | Snrnp70 | chr7 | - | 45376452 | 45395742 | regular |
| ENSMUSG00000022637 | Cblb | chr16 | + | 52031224 | 52208048 | regular |
| ENSMUSG00000047675 | Rps8 | chr4 | - | 117153826 | 117156243 | regular |
| ENSMUSG00000033016 | Nfatc1 | chr18 | - | 80606204 | 80713071 | regular |
| ENSMUSG00000064289 | Tank | chr2 | + | 61578584 | 61654171 | regular |
| ENSMUSG00000024283 | Wac | chr18 | + | 7868831 | 7973547 | regular |
| ENSMUSG00000000787 | Ddx3x | chrX | + | 13280969 | 13294052 | regular |
| ENSMUSG00000092274 | Neat1 | chr19 | - | 5824707 | 5845478 | regular |
| ENSMUSG00000054277 | Arfgap3 | chr15 | - | 83299738 | 83350247 | regular |
| ENSMUSG00000035845 | Alg12 | chr15 | - | 88805243 | 88819318 | regular |
| ENSMUSG00000043629 | 1700019D03 | chr1 | - | 52922323 | 53020179 | regular |
| ENSMUSG00000020076 | Ddx50 | chr10 | - | 62615894 | 62651218 | regular |
| ENSMUSG00000034928 | Rnf44 | chr13 | - | 54679398 | 54693907 | regular |
| ENSMUSG00000042541 | Sem1 | chr6 | - | 6557293 | 6578663 | regular |
| ENSMUSG00000027439 | Gzf1 | chr2 | + | 148681022 | 148692949 | regular |
| ENSMUSG00000021709 | Erbin | chr13 | - | 103818786 | 103920514 | regular |
| ENSMUSG00000023927 | Satb1 | chr17 | - | 51736186 | 51833290 | regular |
| ENSMUSG00000041645 | Ddx24 | chr12 | - | 103407981 | 103425830 | regular |
| ENSMUSG00000031393 | Mecp2 | chrX | - | 74026591 | 74135363 | regular |
| ENSMUSG00000086324 | Gm15564 | chr16 | - | 35966751 | 35983230 | regular |
| ENSMUSG00000032216 | Nedd4 | chr9 | + | 72662345 | 72749852 | regular |
| ENSMUSG00000008763 | Man1a2 | chr3 | - | 100562207 | 100685503 | regular |
| ENSMUSG00000029629 | Phf14 | chr6 | + | 11907808 | 12081205 | regular |
| ENSMUSG00000002658 | Gtf2f1 | chr17 | - | 57003404 | 57011288 | regular |
| ENSMUSG00000094377 | Gm24407 | chr5 | + | 115490253 | 115490394 | regular |
| ENSMUSG00000079006 | Gm14151 | chr2 | - | 151086785 | 151092350 | regular |
| ENSMUSG00000027422 | Rrbp1 | chr2 | - | 143947394 | 144011263 | regular |
| ENSMUSG00000015340 | Cybb | chrX | - | 9435251 | 9487771 | regular |
| ENSMUSG00000015733 | Capza2 | chr6 | + | 17636233 | 17666972 | regular |
| ENSMUSG00000037278 | Tmem97 | chr11 | - | 78541816 | 78550777 | regular |
| ENSMUSG00000031902 | Nfatc3 | chr8 | + | 106058839 | 106130537 | regular |
| ENSMUSG00000079641 | Rpl39 | chrX | - | 37082519 | 37085402 | regular |
| ENSMUSG00000043059 | Zfp513 | chr5 | - | 31198980 | 31202303 | regular |
| ENSMUSG00000048578 | Mlec | chr5 | - | 115142980 | 115158179 | regular |
| ENSMUSG00000057110 | Cntrl | chr2 | + | 35109491 | 35178822 | regular |
| ENSMUSG00000073725 | Lmbrd1 | chr1 | + | 24678629 | 24766301 | regular |
| ENSMUSG00000026977 | Mar.07 | chr2 | + | 60209886 | 60250676 | regular |

| | | | | | | |
|---|---|---|---|---|---|---|
| ENSMUSG00000096205 | Gm22068 | chr11 | + | 87426715 | 87426879 | regular |
| ENSMUSG00000095969 | Rnu1a1 | chr11 | + | 87422866 | 87423030 | regular |
| ENSMUSG00000096659 | Gm25679 | chr12 | - | 54719162 | 54719326 | regular |
| ENSMUSG00000095616 | Gm26244 | chr18 | - | 10150976 | 10151140 | regular |
| ENSMUSG00000064941 | Gm23238 | chr19 | + | 18732740 | 18732904 | regular |
| ENSMUSG00000093815 | Gm26444 | chr12 | + | 54729601 | 54729765 | regular |
| ENSMUSG00000094826 | Gm23804 | chr12 | + | 54714217 | 54714381 | regular |
| ENSMUSG00000096206 | Gm22317 | chr12 | - | 54738458 | 54738622 | regular |
| ENSMUSG00000096214 | Gm22634 | chr12 | + | 54710307 | 54710471 | regular |
| ENSMUSG00000094306 | Gm24924 | chr18 | - | 14647766 | 14647930 | regular |
| ENSMUSG00000095590 | Gm24305 | chr12 | - | 54734554 | 54734718 | regular |
| ENSMUSG00000032712 | 2810474O19 | chr6 | + | 149309413 | 149335663 | regular |
| ENSMUSG00000048076 | Arf1 | chr11 | - | 59211411 | 59228270 | regular |
| ENSMUSG00000023852 | Chd1 | chr17 | + | 15704966 | 17394898 | regular |
| ENSMUSG00000036779 | Papd5 | chr8 | + | 88199212 | 88259722 | regular |
| ENSMUSG00000033039 | Micall1 | chr15 | + | 79108971 | 79136897 | regular |
| ENSMUSG00000039990 | Edrf1 | chr7 | + | 133637542 | 133672971 | regular |
| ENSMUSG00000061525 | 4921509C19I | chr2 | - | 151470541 | 151476153 | regular |
| ENSMUSG00000024513 | Mbd2 | chr18 | + | 70568188 | 70626131 | regular |
| ENSMUSG00000018379 | Srsf1 | chr11 | + | 88047372 | 88053755 | regular |
| ENSMUSG00000018474 | Chd3 | chr11 | - | 69343272 | 69369406 | regular |
| ENSMUSG00000026672 | Optn | chr2 | - | 5020641 | 5064051 | regular |
| ENSMUSG00000026924 | Sec16a | chr2 | - | 26409430 | 26445216 | regular |
| ENSMUSG00000101609 | Kcnq1ot1 | chr7 | - | 143212154 | 143296549 | regular |
| ENSMUSG00000024483 | Ankhd1 | chr18 | + | 36559986 | 36665917 | regular |
| ENSMUSG00000015305 | Sash1 | chr10 | - | 8722218 | 8886070 | regular |
| ENSMUSG00000075033 | Nxpe3 | chr16 | - | 55839952 | 55895279 | regular |
| ENSMUSG00000036792 | Mbd5 | chr2 | + | 48949507 | 49325405 | regular |
| ENSMUSG00000042790 | Rnf214 | chr9 | - | 45863424 | 45906911 | regular |
| ENSMUSG00000015501 | Hivep2 | chr10 | + | 13966074 | 14151374 | regular |
| ENSMUSG00000028952 | Zbtb48 | chr4 | - | 152019773 | 152027671 | regular |
| ENSMUSG00000022884 | Eif4a2 | chr16 | + | 23107443 | 23114136 | regular |
| ENSMUSG00000061232 | H2-K1 | chr17 | - | 33996016 | 34000333 | regular |
| ENSMUSG00000095891 | Gm10717 | chr9 | + | 3025416 | 3033289 | regular |
| ENSMUSG00000024921 | Smarca2 | chr19 | + | 26605049 | 26778322 | regular |
| ENSMUSG00000024290 | Rock1 | chr18 | - | 10064400 | 10181792 | regular |
| ENSMUSG00000042390 | Gatad2b | chr3 | + | 90293177 | 90363407 | regular |
| ENSMUSG00000097039 | Pvt1 | chr15 | + | 62037985 | 62378096 | regular |
| ENSMUSG00000078671 | Chd2 | chr7 | - | 73426637 | 73541830 | regular |
| ENSMUSG00000074476 | Spc24 | chr9 | - | 21755441 | 21760303 | regular |
| ENSMUSG00000104388 | Gm37033 | chr1 | + | 180357648 | 180364998 | regular |
| ENSMUSG00000039632 | Ccdc151 | chr9 | - | 21989870 | 22002634 | regular |
| ENSMUSG00000014195 | Dnajc7 | chr11 | - | 100582817 | 100620168 | regular |
| ENSMUSG00000026568 | Mpc2 | chr1 | + | 165460636 | 165481214 | regular |
| ENSMUSG00000030556 | Lrrc28 | chr7 | - | 67513409 | 67645268 | regular |
| ENSMUSG00000021819 | Zswim8 | chr14 | + | 20707551 | 20723619 | regular |

| | | | | | | |
|---|---|---|---|---|---|---|
| ENSMUSG00000021392 | Nol8 | chr13 | + | 49653077 | 49679016 | regular |
| ENSMUSG00000020882 | Cacnb1 | chr11 | - | 98001507 | 98023034 | regular |
| ENSMUSG00000075014 | Gm10800 | chr2 | - | 98666546 | 98667301 | regular |
| ENSMUSG00000028649 | Macf1 | chr4 | - | 123349632 | 123684360 | regular |
| ENSMUSG00000032504 | Pdcd6ip | chr9 | - | 113651743 | 113708259 | regular |
| ENSMUSG00000024491 | Rbm27 | chr18 | + | 42275352 | 42341542 | regular |
| ENSMUSG00000032812 | Arap1 | chr7 | + | 101348066 | 101412586 | regular |
| ENSMUSG00000031576 | Kcnu1 | chr8 | + | 25849622 | 25937939 | regular |
| ENSMUSG00000032290 | Ptpn9 | chr9 | + | 56994922 | 57062807 | regular |
| ENSMUSG00000061666 | Gdpd1 | chr11 | - | 87033866 | 87074062 | regular |
| ENSMUSG00000061028 | Clasrp | chr7 | - | 19581034 | 19604486 | regular |
| ENSMUSG00000112023 | Lilr4b | chr10 | + | 51480631 | 51486703 | regular |
| ENSMUSG00000028809 | Srrm1 | chr4 | - | 135320483 | 135353321 | regular |
| ENSMUSG00000063065 | Mapk3 | chr7 | + | 126759600 | 126765819 | regular |
| ENSMUSG00000079606 | Gm595 | chrX | - | 48841465 | 48877713 | regular |
| ENSMUSG00000096349 | Gm22513 | chr12 | + | 54696781 | 54696945 | regular |
| ENSMUSG00000040084 | Bub1b | chr2 | + | 118598210 | 118641591 | regular |
| ENSMUSG00000041890 | Git2 | chr5 | - | 114727406 | 114775517 | regular |
| ENSMUSG00000038708 | Golga4 | chr9 | + | 118506266 | 118582519 | regular |
| ENSMUSG00000032097 | Ddx6 | chr9 | + | 44604891 | 44640731 | regular |
| ENSMUSG00000026987 | Baz2b | chr2 | - | 59899362 | 60209839 | regular |
| ENSMUSG00000028137 | Celf3 | chr3 | + | 94478294 | 94492198 | regular |
| ENSMUSG00000025153 | Fasn | chr11 | - | 120805845 | 120824547 | regular |
| ENSMUSG00000028995 | Fam126a | chr5 | - | 23915275 | 24030690 | regular |
| ENSMUSG00000052997 | Uba2 | chr7 | - | 34140687 | 34169599 | regular |
| ENSMUST00000209052 | Gm45086 | chr7 | - | 48724379 | 48726822 | circular |
| ENSMUSG00000026872 | Zeb2 | chr2 | - | 44983631 | 45117395 | regular |
| ENSMUSG00000028520 | 4921539E11I | chr4 | - | 103230444 | 103290863 | regular |
| ENSMUSG00000039068 | Zzz3 | chr3 | + | 152395472 | 152462826 | regular |
| ENSMUSG00000048720 | Tbc1d12 | chr19 | + | 38836578 | 38919923 | regular |
| ENSMUSG00000040681 | Hmgn1 | chr16 | - | 96120617 | 96127729 | regular |
| ENSMUSG00000030096 | Slc6a6 | chr6 | + | 91684052 | 91759066 | regular |
| ENSMUSG00000022901 | Cd86 | chr16 | - | 36603868 | 36666081 | regular |
| ENSMUSG00000051747 | Ttn | chr2 | - | 76703979 | 76982547 | regular |
| ENSMUSG00000086583 | Gm15500 | chr7 | - | 114705216 | 114706235 | regular |
| ENSMUSG00000021356 | Irf4 | chr13 | + | 30749225 | 30766976 | regular |
| ENSMUSG00000022091 | Sorbs3 | chr14 | - | 70180467 | 70211989 | regular |
| ENSMUSG00000038347 | Tcte2 | chr17 | - | 13482552 | 13761825 | regular |
| ENSMUSG00000021457 | Syk | chr13 | + | 52583172 | 52648792 | regular |
| ENSMUSG00000062651 | H2al2a | chr2 | - | 17996416 | 17996958 | regular |
| ENSMUSG00000040652 | Oaz2 | chr9 | + | 65668000 | 65690300 | regular |
| ENSMUSG00000002833 | Hdgfl2 | chr17 | + | 56079633 | 56100607 | regular |
| ENSMUSG00000002748 | Baz1b | chr5 | + | 135187263 | 135246129 | regular |
| ENSMUSG00000067377 | Tspan6 | chrX | - | 133891067 | 133898429 | regular |
| ENSMUSG00000020982 | Nemf | chr12 | - | 69310521 | 69357165 | regular |
| ENSMUSG00000022982 | Sod1 | chr16 | + | 90220741 | 90226322 | regular |

| | | | | | | |
|---|---|---|---|---|---|---|
| ENSMUSG00000002052 | Supt6 | chr11 | - | 78206745 | 78245987 | regular |
| ENSMUSG00000045482 | Trrap | chr5 | + | 144767731 | 144859778 | regular |
| ENSMUSG00000056509 | Gm9999 | chr7 | + | 46983579 | 46987803 | regular |
| ENSMUSG00000027940 | Tpm3 | chr3 | + | 90072648 | 90100902 | regular |
| ENSMUSG00000034243 | Golgb1 | chr16 | + | 36875139 | 36933085 | regular |
| ENSMUSG00000040123 | Zmym5 | chr14 | - | 56790584 | 56811716 | regular |
| ENSMUSG00000029235 | Pdcl2 | chr5 | - | 76312114 | 76331156 | regular |
| ENSMUSG00000047215 | Rpl9 | chr5 | - | 65388363 | 65391444 | regular |
| ENSMUSG00000066037 | Hnrnpr | chr4 | + | 136310941 | 136359447 | regular |
| ENSMUSG00000034349 | Smc4 | chr3 | + | 69004737 | 69034623 | regular |
| ENSMUSG00000024474 | Ik | chr18 | + | 36744655 | 36757639 | regular |
| ENSMUSG00000070544 | Top1 | chr2 | + | 160645887 | 160722764 | regular |
| ENSMUSG00000012535 | Tnpo3 | chr6 | - | 29540826 | 29609887 | regular |
| ENSMUSG00000098973 | Mir6236 | chr9 | + | 110281286 | 110281409 | regular |
| ENSMUSG00000032217 | Rnf111 | chr9 | - | 70425423 | 70503725 | regular |
| ENSMUSG00000027829 | Ccnl1 | chr3 | - | 65946150 | 65958249 | regular |
| ENSMUSG00000028248 | Pnisr | chr4 | + | 21847582 | 21876475 | regular |
| ENSMUSG00000028932 | Psmc2 | chr5 | + | 21785282 | 21803787 | regular |
| ENSMUSG00000022263 | Trio | chr15 | - | 27730650 | 28025848 | regular |
| ENSMUSG00000037568 | Vash2 | chr1 | - | 190947645 | 190979296 | regular |
| ENSMUSG00000036371 | Serbp1 | chr6 | + | 67238175 | 67297736 | regular |
| ENSMUSG00000065944 | Rnu2-10 | chr11 | + | 101658237 | 101658428 | regular |
| ENSMUSG00000065767 | Gm23849 | chr11 | + | 101627365 | 101627556 | regular |
| ENSMUSG00000064702 | Gm24950 | chr11 | + | 101641747 | 101641938 | regular |
| ENSMUSG00000065251 | Gm23971 | chr11 | - | 101645739 | 101645930 | regular |
| ENSMUSG00000065820 | Gm26316 | chr11 | + | 101604849 | 101605040 | regular |
| ENSMUSG00000075015 | Gm10801 | chr2 | + | 98662236 | 98664083 | regular |
| ENSMUSG00000059430 | Actg2 | chr6 | - | 83512904 | 83536265 | regular |
| ENSMUSG00000026523 | Wdr64 | chr1 | + | 175698592 | 175815734 | regular |
| ENSMUSG00000051495 | Irf2bp2 | chr8 | - | 126588295 | 126593986 | regular |
| ENSMUSG00000022200 | Golph3 | chr15 | + | 12321449 | 12351591 | regular |
| ENSMUSG00000040859 | Bsdc1 | chr4 | + | 129461580 | 129488498 | regular |
| ENSMUSG00000030889 | Vwa3a | chr7 | + | 120739317 | 120805742 | regular |
| ENSMUSG00000020462 | Cfap36 | chr11 | - | 29221531 | 29247409 | regular |
| ENSMUSG00000029478 | Ncor2 | chr5 | - | 125017152 | 125179219 | regular |
| ENSMUSG00000034235 | Usp54 | chr14 | - | 20548911 | 20641063 | regular |
| ENSMUSG00000057322 | Rpl38 | chr11 | + | 114668523 | 114672331 | regular |
| ENSMUSG00000095186 | Gm10718 | chr9 | + | 3023546 | 3025218 | regular |
| ENSMUSG00000044949 | Ubtd2 | chr11 | + | 32455369 | 32516687 | regular |
| ENSMUSG00000006005 | Tpr | chr1 | + | 150392837 | 150449935 | regular |
| ENSMUSG00000021127 | Zfp36l1 | chr12 | - | 80107753 | 80113013 | regular |
| ENSMUSG00000026773 | Pfkfb3 | chr2 | - | 11471432 | 11554077 | regular |
| ENSMUSG00000020091 | Eif4ebp2 | chr10 | - | 61428490 | 61452661 | regular |
| ENSMUSG00000024335 | Brd2 | chr17 | - | 34112022 | 34122634 | regular |
| ENSMUSG00000034724 | Cnot6l | chr5 | - | 96070332 | 96164171 | regular |
| ENSMUSG00000026775 | Yme1l1 | chr2 | + | 23156368 | 23199260 | regular |

| | | | | | | |
|---|---|---|---|---|---|---|
| ENSMUSG00000052397 | Ezr | chr17 | - | 6738040 | 6782784 | regular |
| ENSMUSG00000028487 | Bnc2 | chr4 | - | 84275094 | 84675275 | regular |
| ENSMUSG00000109278 | Gm45144 | chr7 | - | 88853042 | 88864892 | regular |
| ENSMUSG00000027365 | Trpm7 | chr2 | - | 126791564 | 126876230 | regular |
| ENSMUSG00000023010 | Tmbim6 | chr15 | + | 99392881 | 99410049 | regular |
| ENSMUSG00000020419 | Hormad2 | chr11 | - | 4345813 | 4441105 | regular |
| ENSMUSG00000108448 | Gm21269 | chr7 | + | 73816545 | 73838898 | regular |
| ENSMUSG00000020841 | Cpd | chr11 | - | 76778423 | 76847018 | regular |
| ENSMUSG00000025159 | Mms19 | chr19 | - | 41941085 | 41981157 | regular |
| ENSMUSG00000030292 | Smco2 | chr6 | + | 146850103 | 146871406 | regular |
| ENSMUSG00000028086 | Fbxw7 | chr3 | + | 84815267 | 84979198 | regular |
| ENSMUSG00000048602 | Morc2b | chr17 | - | 33135587 | 33139683 | regular |
| ENSMUSG00000044581 | 4932415D10 | chr10 | - | 82282115 | 82316582 | regular |
| ENSMUSG00000030067 | Foxp1 | chr6 | - | 98925337 | 99522721 | regular |
| ENSMUSG00000031068 | Glrx3 | chr7 | + | 137437613 | 137468594 | regular |
| ENSMUSG00000019907 | Ppp1r12a | chr10 | + | 108162192 | 108284475 | regular |
| ENSMUSG00000095280 | Gm21738 | chr14 | - | 19415856 | 19418930 | regular |
| ENSMUSG00000062345 | Serpinb2 | chr1 | + | 107511422 | 107535478 | regular |
| ENSMUSG00000041215 | Yeats2 | chr16 | + | 20141062 | 20232573 | regular |
| ENSMUSG00000042644 | Itpr3 | chr17 | + | 27057303 | 27122223 | regular |
| ENSMUSG00000020636 | Allc | chr12 | - | 28553754 | 28582523 | regular |
| ENSMUSG00000085793 | Lin52 | chr12 | + | 84451421 | 84546145 | regular |
| ENSMUSG00000023118 | Sympk | chr7 | + | 19024376 | 19054618 | regular |
| ENSMUSG00000033075 | Senp1 | chr15 | - | 98038743 | 98093744 | regular |
| ENSMUSG00000072791 | Abcb5 | chr12 | - | 118867823 | 118966421 | regular |
| ENSMUSG00000020120 | Plek | chr11 | - | 16971205 | 17052381 | regular |
| ENSMUSG00000096385 | Gm11168 | chr9 | + | 3003345 | 3006948 | regular |
| ENSMUSG00000037343 | Taf2 | chr15 | - | 55015130 | 55072152 | regular |
| ENSMUSG00000026107 | Nabp1 | chr1 | - | 51465861 | 51478425 | regular |
| ENSMUSG00000030990 | Pgap2 | chr7 | + | 102210207 | 102238567 | regular |
| ENSMUSG00000026082 | Rev1 | chr1 | - | 38052785 | 38129801 | regular |
| ENSMUST00000106587 | Agbl4 | chr4 | + | 110478464 | 110580175 | circular |
| ENSMUSG00000029134 | Plb1 | chr5 | + | 32232707 | 32366520 | regular |
| ENSMUSG00000022684 | Bfar | chr16 | + | 13671857 | 13703612 | regular |
| ENSMUSG00000037652 | Phc3 | chr3 | - | 30899370 | 30969415 | regular |
| ENSMUSG00000059562 | Ccdc154 | chr17 | + | 25162460 | 25171913 | regular |
| ENSMUSG00000094445 | 1700015G11 | chr7 | - | 52011682 | 52015728 | regular |
| ENSMUSG00000050702 | 4930563M21 | chr9 | - | 55962588 | 56010550 | regular |
| ENSMUSG00000038766 | Gabpb2 | chr3 | - | 95181765 | 95217916 | regular |
| ENSMUSG00000112627 | 4933412E12 | chr10 | + | 116949756 | 116963279 | regular |
| ENSMUSG00000036202 | Rif1 | chr2 | + | 52072831 | 52122383 | regular |
| ENSMUSG00000023104 | Rfc2 | chr5 | + | 134581365 | 134601805 | regular |
| ENSMUSG00000069094 | Pde7a | chr3 | - | 19223107 | 19311322 | regular |
| ENSMUSG00000031765 | Mt1 | chr8 | + | 94179081 | 94180327 | regular |
| ENSMUSG00000022472 | Desi1 | chr15 | - | 81992522 | 82033862 | regular |
| ENSMUSG00000025393 | Atp5b | chr10 | + | 128083272 | 128090391 | regular |

| | | | | | | |
|---|---|---|---|---|---|---|
| ENSMUSG00000027546 | Atp9a | chr2 | - | 168634437 | 168742409 | regular |
| ENSMUSG00000041846 | Ppp4r3a | chr12 | - | 101039408 | 101083702 | regular |
| ENSMUSG00000018697 | Aatf | chr11 | - | 84422854 | 84513522 | regular |
| ENSMUSG00000005087 | Cd44 | chr2 | - | 102811140 | 102901665 | regular |
| ENSMUSG00000062078 | Qk | chr17 | - | 10206470 | 10319361 | regular |
| ENSMUSG00000055436 | Srsf11 | chr3 | - | 158010472 | 158036639 | regular |
| ENSMUSG00000021414 | Fam217a | chr13 | - | 34909959 | 34924310 | regular |
| ENSMUSG00000027676 | Ccdc39 | chr3 | - | 33812361 | 33844310 | regular |
| ENSMUSG00000001924 | Uba1 | chrX | + | 20658325 | 20683179 | regular |
| ENSMUSG00000052533 | Nup188 | chr2 | + | 30286396 | 30344266 | regular |
| ENSMUSG00000028750 | Pla2g2c | chr4 | + | 138724791 | 138746132 | regular |
| ENSMUSG00000038010 | Ccdc138 | chr10 | + | 58497947 | 58576244 | regular |
| ENSMUSG00000037236 | Matr3 | chr18 | + | 35562145 | 35593835 | regular |
| ENSMUSG00000030102 | Itpr1 | chr6 | + | 108213095 | 108551109 | regular |
| ENSMUSG00000025404 | R3hdm2 | chr10 | + | 127380326 | 127499384 | regular |
| ENSMUSG00000101963 | 1700001J11F | chr9 | + | 40050364 | 40053028 | regular |
| ENSMUSG00000026434 | Nucks1 | chr1 | + | 131910533 | 131936321 | regular |
| ENSMUSG00000022946 | Dopey2 | chr16 | + | 93711903 | 93810590 | regular |
| ENSMUSG00000034269 | Setd5 | chr6 | + | 113077364 | 113153435 | regular |
| ENSMUSG00000021552 | Gkap1 | chr13 | - | 58233345 | 58275699 | regular |
| ENSMUSG00000068747 | Sort1 | chr3 | + | 108284081 | 108361511 | regular |
| ENSMUSG00000035027 | Map2k2 | chr10 | + | 81105914 | 81133975 | regular |
| ENSMUSG00000090733 | Rps27 | chr3 | - | 90212521 | 90213651 | regular |
| ENSMUSG00000005973 | Rcn1 | chr2 | - | 105386290 | 105399319 | regular |
| ENSMUSG00000039543 | Cfap70 | chr14 | - | 20394192 | 20452226 | regular |
| ENSMUSG00000037999 | Arap2 | chr5 | - | 62602444 | 62766159 | regular |
| ENSMUSG00000034187 | Nsf | chr11 | - | 103821781 | 103954056 | regular |
| ENSMUSG00000062270 | Morf4l1 | chr9 | - | 90091664 | 90114774 | regular |
| ENSMUSG00000027288 | Zfp106 | chr2 | - | 120506819 | 120563843 | regular |
| ENSMUSG00000061288 | Taok3 | chr5 | + | 117120128 | 117275219 | regular |
| ENSMUSG00000040828 | Catsperd | chr17 | + | 56628142 | 56664456 | regular |
| ENSMUSG00000055204 | Ankrd17 | chr5 | - | 90227165 | 90366577 | regular |
| ENSMUSG00000021770 | Samd8 | chr14 | + | 21750530 | 21798726 | regular |
| ENSMUSG00000033916 | Chmp2a | chr7 | - | 13032009 | 13034807 | regular |
| ENSMUSG00000021615 | Xrcc4 | chr13 | - | 89774026 | 90089608 | regular |
| ENSMUSG00000027927 | Lelp1 | chr3 | - | 92134996 | 92142744 | regular |
| ENSMUSG00000021993 | Mipep | chr14 | + | 60784572 | 60905478 | regular |
| ENSMUSG00000040225 | Prrc2c | chr1 | - | 162670724 | 162740556 | regular |
| ENSMUSG00000020317 | Theg | chr10 | - | 79576378 | 79587331 | regular |
| ENSMUSG00000056763 | Cspp1 | chr1 | + | 10037986 | 10136768 | regular |
| ENSMUSG00000054823 | Nsd3 | chr8 | + | 25601600 | 25719667 | regular |
| ENSMUSG00000030980 | Knop1 | chr7 | - | 118842221 | 118856254 | regular |
| ENSMUSG00000046585 | Cfap58 | chr19 | + | 47937711 | 48035379 | regular |
| ENSMUSG00000044786 | Zfp36 | chr7 | - | 28376783 | 28380253 | regular |
| ENSMUSG00000022389 | Tef | chr15 | + | 81802420 | 81826861 | regular |
| ENSMUSG00000078796 | Zfp541 | chr7 | + | 16061813 | 16096334 | regular |

| | | | | | | |
|---|---|---|---|---|---|---|
| ENSMUSG00000075376 | Rc3h2 | chr2 | - | 37370068 | 37422903 | regular |
| ENSMUSG00000047417 | Rexo1 | chr10 | - | 80540921 | 80561560 | regular |
| ENSMUSG00000042557 | Sin3a | chr9 | + | 57072039 | 57128366 | regular |
| ENSMUSG00000022858 | Tra2b | chr16 | - | 22244548 | 22266005 | regular |
| ENSMUSG00000028756 | Pink1 | chr4 | - | 138313408 | 138326307 | regular |
| ENSMUSG00000038518 | Jarid2 | chr13 | + | 44729473 | 44921643 | regular |
| ENSMUSG00000038467 | Chmp4b | chr2 | + | 154651704 | 154694785 | regular |
| ENSMUSG00000005225 | Plekha8 | chr6 | + | 54595110 | 54645839 | regular |
| ENSMUSG00000032745 | Gpbp1 | chr13 | - | 111425679 | 111490111 | regular |
| ENSMUSG00000003382 | Etv3 | chr3 | + | 87525406 | 87540156 | regular |
| ENSMUSG00000043987 | Cep164 | chr9 | - | 45766945 | 45828691 | regular |
| ENSMUSG00000047153 | Khnyn | chr14 | + | 55884946 | 55898775 | regular |
| ENSMUSG00000027363 | Usp8 | chr2 | + | 126707327 | 126759297 | regular |
| ENSMUSG00000051236 | Msrb3 | chr10 | - | 120781095 | 120899101 | regular |
| ENSMUSG00000022564 | Grina | chr15 | + | 76246806 | 76249904 | regular |
| ENSMUSG00000026615 | Eprs | chr1 | + | 185363043 | 185428360 | regular |
| ENSMUSG00000022142 | Nup155 | chr15 | + | 8109312 | 8159857 | regular |
| ENSMUSG00000026125 | Prss39 | chr1 | + | 34498409 | 34503063 | regular |
| ENSMUSG00000027942 | 4933434E20I | chr3 | + | 90051635 | 90068347 | regular |
| ENSMUSG00000091415 | Ak9 | chr10 | + | 41303979 | 41434534 | regular |
| ENSMUSG00000022685 | Parn | chr16 | - | 13537963 | 13668170 | regular |
| ENSMUSG00000025742 | Prps2 | chrX | - | 167346321 | 167382749 | regular |
| ENSMUSG00000037754 | Ppp1r16b | chr2 | + | 158665397 | 158766334 | regular |
| ENSMUSG00000061950 | Ppp4r1 | chr17 | + | 65782572 | 65841926 | regular |
| ENSMUSG00000031930 | Wwp2 | chr8 | + | 107436364 | 107558595 | regular |
| ENSMUSG00000037458 | Azin1 | chr15 | - | 38487426 | 38519266 | regular |
| ENSMUSG00000016831 | Tox4 | chr14 | + | 52279145 | 52296401 | regular |
| ENSMUSG00000044627 | Swi5 | chr2 | - | 32278815 | 32288075 | regular |
| ENSMUSG00000036270 | Edc4 | chr8 | + | 105880880 | 105894908 | regular |
| ENSMUSG00000022811 | Zfp148 | chr16 | + | 33380774 | 33503903 | regular |
| ENSMUSG00000033949 | Trim36 | chr18 | - | 46165301 | 46212607 | regular |
| ENSMUSG00000042446 | Zmym4 | chr4 | - | 126861938 | 126968135 | regular |
| ENSMUSG00000037533 | Rapgef6 | chr11 | + | 54522846 | 54699285 | regular |
| ENSMUSG00000014496 | Ankrd28 | chr14 | - | 31698767 | 31830651 | regular |
| ENSMUSG00000029826 | Zc3hav1 | chr6 | - | 38305285 | 38354603 | regular |
| ENSMUSG00000101012 | 1700008K24I | chr17 | - | 49112262 | 49113535 | regular |
| ENSMUSG00000030287 | Itpr2 | chr6 | - | 146108298 | 146502223 | regular |
| ENSMUSG00000050050 | Ccdc158 | chr5 | - | 92607953 | 92675271 | regular |
| ENSMUSG00000038733 | Wdr26 | chr1 | - | 181173227 | 181211552 | regular |
| ENSMUSG00000025041 | Nt5c2 | chr19 | - | 46886830 | 47015153 | regular |
| ENSMUSG00000005981 | Trap1 | chr16 | - | 4039970 | 4077827 | regular |
| ENSMUSG00000020152 | Actr2 | chr11 | - | 20062303 | 20112913 | regular |
| ENSMUSG00000022607 | Ptk2 | chr15 | - | 73205332 | 73423280 | regular |
| ENSMUSG00000033769 | Exoc6b | chr6 | - | 84618486 | 85069513 | regular |
| ENSMUSG00000002028 | Kmt2a | chr9 | - | 44803354 | 44881296 | regular |
| ENSMUSG00000009927 | Rps25 | chr9 | + | 44407138 | 44410427 | regular |

| | | | | | | |
|---|---|---|---|---|---|---|
| ENSMUSG00000022332 | Khdrbs3 | chr15 | + | 68928419 | 69093512 | regular |
| ENSMUSG00000009621 | Vav2 | chr2 | - | 27262103 | 27427033 | regular |
| ENSMUSG00000038844 | Kif16b | chr2 | - | 142617473 | 142901531 | regular |
| ENSMUSG00000003731 | Kpna6 | chr4 | - | 129643979 | 129672767 | regular |
| ENSMUSG00000020893 | Per1 | chr11 | + | 69095216 | 69109960 | regular |
| ENSMUSG00000020131 | Pcsk4 | chr10 | - | 80321282 | 80329498 | regular |
| ENSMUSG00000041341 | Atg2b | chr12 | - | 105616135 | 105685211 | regular |
| ENSMUSG00000038866 | Zcchc2 | chr1 | + | 105990405 | 106034074 | regular |
| ENSMUSG00000068036 | Afdn | chr17 | + | 13760538 | 13906150 | regular |
| ENSMUSG00000032777 | Gtf3c1 | chr7 | - | 125640953 | 125707780 | regular |
| ENSMUSG00000035891 | Cerk | chr15 | - | 86139127 | 86186141 | regular |
| ENSMUSG00000036606 | Plxnb2 | chr15 | - | 89155548 | 89171120 | regular |
| ENSMUSG00000040729 | Cep126 | chr9 | - | 8076460 | 8134294 | regular |
| ENSMUSG00000077714 | Snord17 | chr2 | - | 144265978 | 144266216 | regular |
| ENSMUSG00000032688 | Malt1 | chr18 | + | 65430961 | 65478823 | regular |
| ENSMUST00000046656 | Tasp1 | chr2 | - | 140042093 | 140057499 | circular |
| ENSMUSG00000055065 | Ddx17 | chr15 | - | 79527706 | 79546741 | regular |
| ENSMUSG00000019971 | Cep290 | chr10 | + | 100487557 | 100574840 | regular |
| ENSMUSG00000031667 | Aktip | chr8 | - | 91111783 | 91199976 | regular |
| ENSMUST00000092143 | Msrb3 | chr10 | - | 120825107 | 120852092 | circular |
| ENSMUSG00000095547 | Gm10719 | chr9 | + | 3017407 | 3021593 | regular |
| ENSMUSG00000032355 | Mlip | chr9 | - | 77102080 | 77352969 | regular |
| ENSMUSG00000034312 | Iqsec1 | chr6 | - | 90656087 | 90988685 | regular |
| ENSMUSG00000019841 | Rev3l | chr10 | + | 39732117 | 39875211 | regular |
| ENSMUSG00000038056 | Kmt2c | chr5 | - | 25271797 | 25498783 | regular |
| ENSMUSG00000028188 | Spata1 | chr3 | - | 146457195 | 146499753 | regular |
| ENSMUSG00000049571 | Cfap46 | chr7 | - | 139600950 | 139683817 | regular |
| ENSMUSG00000069833 | Ahnak | chr19 | + | 8989283 | 9076919 | regular |
| ENSMUSG00000031015 | Swap70 | chr7 | + | 110221710 | 110283506 | regular |
| ENSMUSG00000022191 | Drosha | chr15 | + | 12824814 | 12935291 | regular |
| ENSMUSG00000021987 | Mtmr6 | chr14 | + | 60265227 | 60302370 | regular |
| ENSMUSG00000032621 | Srek1 | chr13 | - | 103739347 | 103774608 | regular |
| ENSMUSG00000030850 | Ate1 | chr7 | - | 130391492 | 130520369 | regular |
| ENSMUSG00000040913 | Fbxw4 | chr19 | - | 45578253 | 45660312 | regular |
| ENSMUSG00000020189 | Osbpl8 | chr10 | + | 111164751 | 111297249 | regular |
| ENSMUSG00000030798 | Cd37 | chr7 | - | 45233631 | 45239115 | regular |
| ENSMUSG00000078161 | Erich3 | chr3 | + | 154663858 | 154767790 | regular |
| ENSMUSG00000035851 | Ythdc1 | chr5 | + | 86804220 | 86836659 | regular |
| ENSMUSG00000001151 | Pcnt | chr10 | - | 76351262 | 76442786 | regular |
| ENSMUSG00000041235 | Chd7 | chr4 | + | 8690405 | 8867659 | regular |
| ENSMUSG00000009207 | Lnpk | chr2 | - | 74520290 | 74579435 | regular |
| ENSMUSG00000026162 | Nhej1 | chr1 | - | 74967138 | 75125200 | regular |
| ENSMUSG00000018401 | Mtmr4 | chr11 | + | 87592161 | 87616302 | regular |
| ENSMUSG00000047867 | Gimap6 | chr6 | - | 48701581 | 48708225 | regular |
| ENSMUSG00000020743 | Mif4gd | chr11 | - | 115607917 | 115612969 | regular |
| ENSMUSG00000042707 | Dnali1 | chr4 | - | 125055337 | 125065703 | regular |

| | | | | | | |
|---|---|---|---|---|---|---|
| ENSMUSG00000022010 | Tsc22d1 | chr14 | + | 76414960 | 76507765 | regular |
| ENSMUSG00000110020 | Gm45441 | chr7 | + | 45830894 | 45835249 | regular |
| ENSMUSG00000052783 | Grk4 | chr5 | + | 34660378 | 34755305 | regular |
| ENSMUSG00000004207 | Psap | chr10 | + | 60277626 | 60302597 | regular |
| ENSMUSG00000030846 | Tial1 | chr7 | - | 128439776 | 128461717 | regular |
| ENSMUSG00000060036 | Rpl3 | chr15 | - | 80077793 | 80091868 | regular |
| ENSMUSG00000024180 | Tmem8 | chr17 | + | 26113315 | 26123254 | regular |
| ENSMUSG00000030726 | Pold3 | chr7 | - | 100082110 | 100121565 | regular |
| ENSMUSG00000009741 | Ubp1 | chr9 | + | 113930933 | 113977202 | regular |
| ENSMUSG00000044224 | Dnajc21 | chr15 | - | 10446755 | 10470516 | regular |
| ENSMUSG00000024242 | Map4k3 | chr17 | - | 80580511 | 80728093 | regular |
| ENSMUSG00000056579 | Tug1 | chr11 | - | 3639784 | 3649673 | regular |
| ENSMUSG00000034297 | Med13 | chr11 | - | 86267032 | 86357602 | regular |
| ENSMUSG00000056962 | Jmjd6 | chr11 | - | 116837431 | 116843449 | regular |
| ENSMUSG00000097971 | Gm26917 | chr17 | + | 39843012 | 39846341 | regular |
| ENSMUSG00000032178 | Ilf3 | chr9 | + | 21367870 | 21405361 | regular |
| ENSMUSG00000059474 | Mbtd1 | chr11 | + | 93885851 | 93946985 | regular |
| ENSMUSG00000024943 | Smc5 | chr19 | - | 23206450 | 23273897 | regular |
| ENSMUSG00000017707 | Serinc3 | chr2 | - | 163623271 | 163645131 | regular |
| ENSMUSG00000059208 | Hnrnpm | chr17 | - | 33646232 | 33686860 | regular |
| ENSMUSG00000092519 | Actl9 | chr17 | + | 33432895 | 33434268 | regular |
| ENSMUSG00000075569 | Rsph10b | chr5 | + | 143933034 | 143985719 | regular |
| ENSMUSG00000031552 | Adam18 | chr8 | - | 24602245 | 24674755 | regular |
| ENSMUSG00000038658 | Ric1 | chr19 | + | 29522281 | 29606829 | regular |
| ENSMUSG00000041399 | 1700013G24 | chr4 | + | 137453283 | 137455461 | regular |
| ENSMUSG00000023084 | Lrrc71 | chr3 | - | 87736922 | 87748625 | regular |
| ENSMUSG00000028944 | Prkag2 | chr5 | - | 24862743 | 25100642 | regular |
| ENSMUSG00000034201 | Gas2l1 | chr11 | - | 5054131 | 5065327 | regular |
| ENSMUSG00000021192 | Golga5 | chr12 | + | 102469134 | 102497907 | regular |
| ENSMUSG00000078627 | Mar.10 | chr11 | - | 105360797 | 105456735 | regular |
| ENSMUSG00000022312 | Eif3h | chr15 | - | 51786557 | 51865523 | regular |
| ENSMUSG00000022064 | Pibf1 | chr14 | + | 99099423 | 99254493 | regular |
| ENSMUSG00000040649 | Rimklb | chr6 | - | 122447295 | 122498761 | regular |
| ENSMUSG00000024188 | Luc7l | chr17 | + | 26252909 | 26285504 | regular |
| ENSMUSG00000032010 | Usp2 | chr9 | + | 44067020 | 44095627 | regular |
| ENSMUSG00000026567 | Adcy10 | chr1 | + | 165485182 | 165576774 | regular |
| ENSMUSG00000061787 | Rps17 | chr7 | - | 81342731 | 81345254 | regular |
| ENSMUSG00000028937 | Acot7 | chr4 | + | 152178133 | 152271855 | regular |
| ENSMUSG00000020719 | Ddx5 | chr11 | - | 106780354 | 106789185 | regular |
| ENSMUSG00000110170 | St6galnac2 | chr11 | - | 116677482 | 116681290 | regular |
| ENSMUSG00000033705 | Stard9 | chr2 | + | 120629125 | 120731895 | regular |
| ENSMUSG00000073856 | Iqck | chr7 | + | 118855751 | 118972652 | regular |
| ENSMUSG00000026399 | Cd55 | chr1 | - | 130439026 | 130462744 | regular |
| ENSMUSG00000030595 | Nfkbib | chr7 | - | 28758250 | 28767512 | regular |
| ENSMUSG00000053641 | Dennd4a | chr9 | + | 64811339 | 64919667 | regular |
| ENSMUSG00000021218 | Gdi2 | chr13 | + | 3538062 | 3567871 | regular |

| | | | | | | |
|---|---|---|---|---|---|---|
| ENSMUSG00000076617 | Ighm | chr12 | - | 113418557 | 113422730 | regular |
| ENSMUSG00000024073 | Birc6 | chr17 | + | 74528294 | 74703356 | regular |
| ENSMUSG00000086808 | Gm11981 | chr11 | - | 6716000 | 6720037 | regular |
| ENSMUSG00000011831 | Evi5 | chr5 | - | 107744794 | 107875107 | regular |
| ENSMUSG00000042678 | Myo15 | chr11 | + | 60469338 | 60528369 | regular |
| ENSMUSG00000070332 | Trim80 | chr11 | + | 115440544 | 115448270 | regular |
| ENSMUSG00000005698 | Ctcf | chr8 | + | 105636567 | 105682922 | regular |
| ENSMUSG00000036244 | Tbc1d21 | chr9 | - | 58359703 | 58370455 | regular |
| ENSMUSG00000004929 | Thop1 | chr10 | + | 81070034 | 81082559 | regular |
| ENSMUSG00000041323 | Ak7 | chr12 | + | 105705975 | 105782447 | regular |
| ENSMUSG00000039530 | Tusc3 | chr8 | + | 39005844 | 39165114 | regular |
| ENSMUSG00000057240 | Ms4a13 | chr19 | - | 11169417 | 11196737 | regular |
| ENSMUSG00000021000 | Ctage5 | chr12 | + | 59095798 | 59191583 | regular |
| ENSMUSG00000021879 | Dnah12 | chr14 | + | 26693273 | 26891703 | regular |
| ENSMUSG00000004110 | Cacna1e | chr1 | - | 154390730 | 154884501 | regular |
| ENSMUSG00000039050 | Osbpl2 | chr2 | + | 180119305 | 180162680 | regular |
| ENSMUSG00000031701 | Dnaja2 | chr8 | - | 85537632 | 85555344 | regular |
| ENSMUSG00000097585 | E230029C05I | chr7 | + | 89980722 | 90049069 | regular |
| ENSMUSG00000109561 | Ankrd31 | chr13 | + | 96748271 | 96910039 | regular |
| ENSMUSG00000057113 | Npm1 | chr11 | - | 33152286 | 33163206 | regular |
| ENSMUSG00000085143 | Gm11520 | chr11 | + | 95310428 | 95324496 | regular |
| ENSMUSG00000034390 | Cmip | chr8 | + | 117257063 | 117459430 | regular |
| ENSMUSG00000020184 | Mdm2 | chr10 | - | 117688874 | 117710758 | regular |
| ENSMUSG00000051435 | Fhad1 | chr4 | - | 141890437 | 142015082 | regular |
| ENSMUSG00000030216 | Wbp11 | chr6 | - | 136813653 | 136828233 | regular |
| ENSMUSG00000000568 | Hnrnpd | chr5 | - | 99955934 | 99978938 | regular |
| ENSMUSG00000031453 | Rasa3 | chr8 | - | 13566947 | 13677603 | regular |
| ENSMUSG00000033557 | Fam20b | chr1 | - | 156678531 | 156719086 | regular |
| ENSMUSG00000026655 | Fam107b | chr2 | + | 3570487 | 3782142 | regular |
| ENSMUSG00000006169 | Clint1 | chr11 | + | 45852050 | 45910625 | regular |
| ENSMUSG00000084786 | Ubl5 | chr9 | + | 20642877 | 20647140 | regular |
| ENSMUSG00000035392 | Dennd1a | chr2 | - | 37798990 | 38287390 | regular |
| ENSMUSG00000070031 | Sp140 | chr1 | + | 85600377 | 85645037 | regular |
| ENSMUSG00000032410 | Xrn1 | chr9 | + | 95954759 | 96057803 | regular |
| ENSMUSG00000065954 | Tacc1 | chr8 | - | 25154551 | 25256588 | regular |
| ENSMUSG00000040390 | Map3k10 | chr7 | - | 27656374 | 27674598 | regular |
| ENSMUSG00000034252 | Senp6 | chr9 | + | 80066902 | 80144953 | regular |
| ENSMUSG00000058230 | Arhgap35 | chr7 | - | 16493718 | 16614993 | regular |
| ENSMUSG00000029992 | Gfpt1 | chr6 | + | 87042845 | 87092197 | regular |
| ENSMUSG00000024387 | Csnk2b | chr17 | - | 35116195 | 35122053 | regular |
| ENSMUSG00000085720 | Gm7854 | chr5 | - | 43151685 | 43235355 | regular |
| ENSMUSG00000043131 | Mob1a | chr6 | + | 83326015 | 83343776 | regular |
| ENSMUSG00000030231 | Plekha5 | chr6 | + | 140424053 | 140597110 | regular |
| ENSMUSG00000005510 | Ndufs3 | chr2 | - | 90894633 | 90904827 | regular |
| ENSMUSG00000037287 | Tbcel | chr9 | - | 42412315 | 42507809 | regular |
| ENSMUSG00000032479 | Map4 | chr9 | + | 109931459 | 110083955 | regular |

| | | | | | | |
|---|---|---|---|---|---|---|
| ENSMUSG00000050677 | Ccdc96 | chr5 | + | 36484587 | 36488172 | regular |
| ENSMUSG00000062202 | Btbd9 | chr17 | - | 30215523 | 30576287 | regular |
| ENSMUSG00000043716 | Rpl7 | chr1 | - | 16101294 | 16104662 | regular |
| ENSMUSG00000026743 | Mllt10 | chr2 | + | 18055236 | 18212388 | regular |
| ENSMUSG00000029427 | Zcchc8 | chr5 | - | 123698293 | 123721100 | regular |
| ENSMUSG00000034245 | Hdac11 | chr6 | + | 91156664 | 91174692 | regular |
| ENSMUSG00000034544 | Rsrc1 | chr3 | + | 66981389 | 67358396 | regular |
| ENSMUSG00000049775 | Tmsb4x | chrX | - | 167207092 | 167209315 | regular |
| ENSMUSG00000019027 | Dnah1 | chr14 | - | 31260374 | 31323896 | regular |
| ENSMUSG00000014030 | Pax5 | chr4 | - | 44524756 | 44710487 | regular |
| ENSMUSG00000030357 | Fkbp4 | chr6 | - | 128429734 | 128438677 | regular |
| ENSMUSG00000044950 | Pwwp2a | chr11 | + | 43681997 | 43721491 | regular |
| ENSMUSG00000090394 | 4930523C07I | chr1 | + | 160044381 | 160080208 | regular |
| ENSMUSG00000022141 | Nipbl | chr15 | - | 8291224 | 8444463 | regular |
| ENSMUSG00000020608 | Smc6 | chr12 | + | 11265885 | 11319785 | regular |
| ENSMUSG00000037148 | Arhgap10 | chr8 | - | 77250365 | 77517953 | regular |
| ENSMUSG00000061838 | Suclg2 | chr6 | - | 95473008 | 95718800 | regular |
| ENSMUSG00000030213 | Atf7ip | chr6 | + | 136506166 | 136610862 | regular |
| ENSMUSG00000031774 | Fam192a | chr8 | - | 94574137 | 94601997 | regular |
| ENSMUSG00000021660 | Btf3 | chr13 | - | 98309895 | 98317006 | regular |
| ENSMUSG00000021938 | Pspc1 | chr14 | - | 56720709 | 56778316 | regular |
| ENSMUSG00000056851 | Pcbp2 | chr15 | + | 102470631 | 102500059 | regular |
| ENSMUSG00000024085 | Man2a1 | chr17 | + | 64600735 | 64755110 | regular |
| ENSMUSG00000052296 | Ppp6r1 | chr7 | - | 4631494 | 4659019 | regular |
| ENSMUSG00000001552 | Jup | chr11 | - | 100368957 | 100397763 | regular |
| ENSMUSG00000038828 | Tmem214 | chr5 | + | 30868011 | 30879180 | regular |
| ENSMUSG00000021257 | Angel1 | chr12 | - | 86700501 | 86726460 | regular |
| ENSMUSG00000034254 | Agpat1 | chr17 | + | 34604261 | 34613449 | regular |
| ENSMUSG00000021090 | Lrrc9 | chr12 | + | 72441865 | 72530750 | regular |
| ENSMUSG00000038831 | Ralgps1 | chr2 | - | 33133416 | 33371486 | regular |
| ENSMUST00000162036 | Rsrc1 | chr3 | + | 66994516 | 66996590 | circular |
| ENSMUSG00000026739 | Bmi1 | chr2 | + | 18677017 | 18686629 | regular |
| ENSMUSG00000035325 | Sec31a | chr5 | - | 100361648 | 100416234 | regular |
| ENSMUSG00000064999 | Gm26035 | chrX | - | 152923452 | 152923760 | regular |
| ENSMUSG00000045210 | Vcpip1 | chr1 | - | 9718621 | 9748382 | regular |
| ENSMUSG00000026200 | Glb1l | chr1 | - | 75198235 | 75210813 | regular |
| ENSMUSG00000075249 | Fsip2 | chr2 | + | 82943633 | 83008937 | regular |
| ENSMUSG00000091896 | Ube2d2a | chr18 | + | 35771558 | 35807172 | regular |
| ENSMUSG00000018395 | Kif3a | chr11 | + | 53567378 | 53601967 | regular |
| ENSMUSG00000028256 | Odf2l | chr3 | + | 145118587 | 145153915 | regular |
| ENSMUSG00000021488 | Nsd1 | chr13 | + | 55209781 | 55318325 | regular |
| ENSMUSG00000053293 | Pom121 | chr5 | - | 135376140 | 135394546 | regular |
| ENSMUST00000037248 | Hirip3 | chr7 | + | 126864319 | 126864661 | circular |
| ENSMUSG00000057265 | Bbof1 | chr12 | + | 84409070 | 84442887 | regular |
| ENSMUSG00000071054 | Safb | chr17 | + | 56584824 | 56606294 | regular |
| ENSMUSG00000039220 | Ppp1r10 | chr17 | + | 35916433 | 35932283 | regular |

| | | | | | | |
|---|---|---|---|---|---|---|
| ENSMUSG00000068921 | Dap3 | chr3 | - | 88920802 | 88951181 | regular |
| ENSMUSG00000052889 | Prkcb | chr7 | + | 122288750 | 122634402 | regular |
| ENSMUSG00000020458 | Rtn4 | chr11 | + | 29692946 | 29744331 | regular |
| ENSMUSG00000020530 | Ggnbp2 | chr11 | - | 84832360 | 84870817 | regular |
| ENSMUSG00000028692 | Akr1a1 | chr4 | - | 116636509 | 116651680 | regular |
| ENSMUSG00000031706 | Rfx1 | chr8 | + | 84066833 | 84096992 | regular |
| ENSMUSG00000001419 | Mef2d | chr3 | + | 88142371 | 88172086 | regular |
| ENSMUSG00000074634 | Tmem267 | chr13 | + | 119488038 | 119611059 | regular |
| ENSMUSG00000022568 | Scrib | chr15 | - | 76047157 | 76069784 | regular |
| ENSMUSG00000010755 | Cars | chr7 | - | 143557229 | 143600090 | regular |
| ENSMUSG00000070980 | Actl7b | chr4 | - | 56740004 | 56741443 | regular |
| ENSMUSG00000056211 | R3hdm1 | chr1 | + | 128103300 | 128237736 | regular |
| ENSMUSG00000074627 | Mroh8 | chr2 | - | 157208549 | 157279549 | regular |
| ENSMUSG00000020211 | Sf3a2 | chr10 | + | 80798197 | 80804924 | regular |
| ENSMUSG00000031787 | Katnb1 | chr8 | + | 95081185 | 95103149 | regular |
| ENSMUSG00000032846 | Zswim6 | chr13 | - | 107724616 | 107890064 | regular |
| ENSMUSG00000024429 | Gnl1 | chr17 | + | 35979850 | 35989462 | regular |
| ENSMUSG00000027184 | Caprin1 | chr2 | - | 103762940 | 103797649 | regular |
| ENSMUSG00000061755 | Bod1l | chr5 | - | 41787537 | 41844315 | regular |
| ENSMUSG00000047141 | Zfp654 | chr16 | - | 64780346 | 64851652 | regular |
| ENSMUSG00000038994 | Hils1 | chr11 | + | 94967592 | 94968456 | regular |
| ENSMUSG00000040812 | Agbl2 | chr2 | + | 90782726 | 90834437 | regular |
| ENSMUSG00000021693 | Kif2a | chr13 | - | 106958995 | 107022126 | regular |
| ENSMUSG00000040794 | C1qtnf4 | chr2 | + | 90885859 | 90890525 | regular |
| ENSMUSG00000043535 | Setx | chr2 | + | 29124180 | 29182471 | regular |
| ENSMUSG00000024749 | Tmc1 | chr19 | - | 20783457 | 20954202 | regular |
| ENSMUSG00000030536 | Iqgap1 | chr7 | - | 80711582 | 80825974 | regular |
| ENSMUSG00000003435 | Supt5 | chr7 | - | 28314890 | 28338746 | regular |
| ENSMUSG00000024430 | Cabyr | chr18 | + | 12741323 | 12755146 | regular |
| ENSMUSG00000045128 | Rpl18a | chr8 | - | 70894721 | 70897412 | regular |
| ENSMUSG00000041974 | Spidr | chr16 | - | 15889226 | 16146851 | regular |
| ENSMUSG00000024294 | Mib1 | chr18 | + | 10725547 | 10818704 | regular |
| ENSMUSG00000021003 | Galc | chr12 | - | 98202293 | 98259459 | regular |
| ENSMUSG00000048701 | Ccdc6 | chr10 | + | 70097120 | 70193200 | regular |
| ENSMUSG00000051618 | Ubqln3 | chr7 | - | 104140622 | 104143279 | regular |
| ENSMUSG00000026807 | Ak8 | chr2 | + | 28700163 | 28813165 | regular |
| ENSMUSG00000050994 | Adgb | chr10 | - | 10335702 | 10472326 | regular |
| ENSMUSG00000055531 | Cpsf6 | chr10 | - | 117344672 | 117380015 | regular |
| ENSMUSG00000019834 | Slc22a16 | chr10 | + | 40570335 | 40604132 | regular |
| ENSMUSG00000027602 | Map1lc3a | chr2 | + | 155276296 | 155278073 | regular |
| ENSMUSG00000001120 | Pcbp3 | chr10 | - | 76761856 | 76961887 | regular |
| ENSMUSG00000058589 | Anks1b | chr10 | + | 89873508 | 90973300 | regular |
| ENSMUSG00000027009 | Itga4 | chr2 | + | 79255425 | 79333123 | regular |
| ENSMUSG00000022707 | Gbe1 | chr16 | + | 70313948 | 70569716 | regular |
| ENSMUSG00000064307 | Lrrc51 | chr7 | - | 101912983 | 101933895 | regular |
| ENSMUSG00000101968 | 1700027A15 | chr1 | + | 72984843 | 73025507 | regular |

| | | | | | | |
|---|---|---|---|---|---|---|
| ENSMUSG00000021754 | Map3k1 | chr13 | - | 111746427 | 111808993 | regular |
| ENSMUSG00000037395 | Rcor3 | chr1 | - | 192098545 | 192138062 | regular |
| ENSMUSG00000020166 | Cnot2 | chr10 | - | 116485160 | 116581511 | regular |
| ENSMUSG00000021111 | Papola | chr12 | + | 105784693 | 105838944 | regular |
| ENSMUSG00000022427 | Tomm22 | chr15 | + | 79670860 | 79672862 | regular |
| ENSMUSG00000036398 | Ppp1r11 | chr17 | - | 36948355 | 36951741 | regular |
| ENSMUSG00000038664 | Herc1 | chr9 | + | 66350449 | 66508775 | regular |
| ENSMUSG00000037196 | Pacrg | chr17 | - | 10403011 | 10840311 | regular |
| ENSMUSG00000056050 | Mia3 | chr1 | - | 183326724 | 183369553 | regular |
| ENSMUSG00000029534 | St7 | chr6 | + | 17692932 | 17943025 | regular |
| ENSMUSG00000070979 | Actl7a | chr4 | + | 56743412 | 56744925 | regular |
| ENSMUSG00000067851 | Arfgef1 | chr1 | - | 10137570 | 10232670 | regular |
| ENSMUSG00000059540 | Tcea2 | chr2 | + | 181680309 | 181688071 | regular |
| ENSMUSG00000049482 | Ctu2 | chr8 | + | 122476142 | 122484138 | regular |
| ENSMUSG00000021134 | Srsf5 | chr12 | + | 80945503 | 80950507 | regular |
| ENSMUSG00000039202 | Abhd2 | chr7 | + | 79273198 | 79365508 | regular |
| ENSMUSG00000024696 | Lpxn | chr19 | + | 12798605 | 12833807 | regular |
| ENSMUSG00000031029 | Eif3f | chr7 | + | 108934416 | 108942951 | regular |
| ENSMUSG00000001576 | Ergic1 | chr17 | + | 26561488 | 26656931 | regular |
| ENSMUSG00000044526 | Znrf4 | chr17 | - | 56511247 | 56512483 | regular |
| ENSMUSG00000101959 | Ldhal6b | chr17 | - | 5417322 | 5418767 | regular |
| ENSMUSG00000039021 | Ttc16 | chr2 | - | 32757025 | 32775633 | regular |
| ENSMUSG00000038274 | Fau | chr19 | + | 6057887 | 6059524 | regular |
| ENSMUSG00000049550 | Clip1 | chr5 | - | 123577794 | 123684618 | regular |
| ENSMUSG00000031949 | Adat1 | chr8 | - | 111966907 | 111992302 | regular |
| ENSMUSG00000033526 | Ppip5k1 | chr2 | - | 121310560 | 121355396 | regular |
| ENSMUSG00000046196 | Ttc39d | chr17 | + | 80207459 | 80217936 | regular |
| ENSMUSG00000051674 | Dcun1d4 | chr5 | + | 73480999 | 73560794 | regular |
| ENSMUSG00000094483 | Purb | chr11 | - | 6467598 | 6475917 | regular |
| ENSMUSG00000026269 | Rnpepl1 | chr1 | + | 92910782 | 92924384 | regular |
| ENSMUSG00000031858 | Mau2 | chr8 | - | 70016122 | 70042734 | regular |
| ENSMUSG00000024758 | Rtn3 | chr19 | - | 7425900 | 7483289 | regular |
| ENSMUSG00000021807 | Rtraf | chr14 | - | 19811350 | 19823824 | regular |
| ENSMUSG00000062154 | Tex33 | chr15 | - | 78378399 | 78395912 | regular |
| ENSMUSG00000032228 | Tcf12 | chr9 | - | 71842687 | 72111871 | regular |
| ENSMUSG00000026193 | Fn1 | chr1 | - | 71585519 | 71653200 | regular |
| ENSMUSG00000020733 | Slc9a3r1 | chr11 | + | 115163340 | 115181181 | regular |
| ENSMUSG00000066306 | Numa1 | chr7 | + | 101934110 | 102014964 | regular |
| ENSMUSG00000042625 | Safb2 | chr17 | - | 56560964 | 56584585 | regular |
| ENSMUSG00000018736 | Ndel1 | chr11 | - | 68821433 | 68871858 | regular |
| ENSMUSG00000044791 | Setd2 | chr9 | + | 110532596 | 110618633 | regular |
| ENSMUSG00000061689 | Dlgap4 | chr2 | + | 156613704 | 156764363 | regular |
| ENSMUSG00000037926 | Ssh2 | chr11 | + | 77216286 | 77460220 | regular |
| ENSMUSG00000020114 | Cand1 | chr10 | - | 119199254 | 119240055 | regular |
| ENSMUSG00000042744 | Hectd4 | chr5 | + | 121220218 | 121368577 | regular |
| ENSMUSG00000025967 | Eef1b2 | chr1 | + | 63176824 | 63180486 | regular |

| | | | | | | |
|---|---|---|---|---|---|---|
| ENSMUSG00000062028 | Irgc1 | chr7 | - | 24431921 | 24452297 | regular |
| ENSMUSG00000029501 | Ankle2 | chr5 | + | 110231003 | 110256651 | regular |
| ENSMUSG00000031822 | Gse1 | chr8 | + | 120228455 | 120581390 | regular |
| ENSMUSG00000020799 | Tekt1 | chr11 | - | 72344721 | 72362442 | regular |
| ENSMUSG00000034300 | Fam53c | chr18 | + | 34758905 | 34773760 | regular |
| ENSMUSG00000026234 | Ncl | chr1 | - | 86344718 | 86359400 | regular |
| ENSMUSG00000031153 | Gripap1 | chrX | + | 7789764 | 7820567 | regular |
| ENSMUSG00000021945 | Zmym2 | chr14 | + | 56886652 | 56962701 | regular |
| ENSMUSG00000108738 | Gm44777 | chr7 | + | 116334254 | 116338633 | regular |
| ENSMUSG00000058799 | Nap1l1 | chr10 | + | 111473222 | 111498150 | regular |
| ENSMUSG00000028868 | Wasf2 | chr4 | + | 133130504 | 133199756 | regular |
| ENSMUSG00000026203 | Dnajb2 | chr1 | + | 75236405 | 75245692 | regular |
| ENSMUSG00000025885 | Myo5b | chr18 | + | 74440935 | 74771493 | regular |
| ENSMUSG00000076438 | Oxct2b | chr4 | + | 123116265 | 123118000 | regular |
| ENSMUSG00000005893 | Nr2c2 | chr6 | + | 92091389 | 92174294 | regular |
| ENSMUSG00000097312 | Gm26870 | chr9 | - | 3000281 | 3038313 | regular |
| ENSMUSG00000039458 | Mtmr12 | chr15 | + | 12205027 | 12274496 | regular |
| ENSMUSG00000028484 | Psip1 | chr4 | - | 83455679 | 83486459 | regular |
| ENSMUSG00000031585 | Gtf2e2 | chr8 | + | 33731832 | 33777173 | regular |
| ENSMUSG00000019943 | Atp2b1 | chr10 | + | 98914405 | 99026143 | regular |
| ENSMUSG00000017843 | Ppp2r5c | chr12 | + | 110447119 | 110583062 | regular |
| ENSMUSG00000061802 | Armc4 | chr18 | - | 7088232 | 7297901 | regular |
| ENSMUSG00000019906 | Lin7a | chr10 | + | 107271685 | 107421474 | regular |
| ENSMUSG00000032221 | Mns1 | chr9 | + | 72438010 | 72462025 | regular |
| ENSMUSG00000071324 | Armc2 | chr10 | - | 41914989 | 42018442 | regular |
| ENSMUSG00000037075 | Rnf139 | chr15 | + | 58889228 | 58907057 | regular |
| ENSMUSG00000005198 | Polr2a | chr11 | - | 69733996 | 69758637 | regular |
| ENSMUSG00000028559 | Osbpl9 | chr4 | - | 109061144 | 109202272 | regular |
| ENSMUSG00000021282 | Eif5 | chr12 | + | 111538015 | 111546752 | regular |
| ENSMUSG00000038932 | Tcfl5 | chr2 | - | 180621955 | 180642708 | regular |
| ENSMUSG00000041301 | Cftr | chr6 | + | 18170686 | 18322768 | regular |
| ENSMUSG00000070568 | Slc6a21 | chr7 | + | 45277512 | 45288998 | regular |
| ENSMUSG00000034793 | G6pc3 | chr11 | + | 102189619 | 102194081 | regular |
| ENSMUSG00000027177 | Hipk3 | chr2 | - | 104426480 | 104494446 | regular |
| ENSMUSG00000046782 | Ttc6 | chr12 | + | 57564112 | 57737928 | regular |
| ENSMUSG00000060261 | Gtf2i | chr5 | - | 134237833 | 134314760 | regular |
| ENSMUSG00000041444 | Arhgap32 | chr9 | + | 32116135 | 32268446 | regular |
| ENSMUSG00000027517 | Ankrd60 | chr2 | - | 173568665 | 173578365 | regular |
| ENSMUSG00000041720 | Pi4ka | chr16 | - | 17280350 | 17406314 | regular |
| ENSMUSG00000001517 | Foxm1 | chr6 | + | 128362966 | 128376146 | regular |
| ENSMUSG00000021460 | Auh | chr13 | - | 52835118 | 52929681 | regular |
| ENSMUSG00000001156 | Mxd1 | chr6 | - | 86647041 | 86669161 | regular |
| ENSMUSG00000032399 | Rpl4 | chr9 | + | 64173374 | 64178666 | regular |
| ENSMUSG00000036529 | Sbf1 | chr15 | - | 89288235 | 89315311 | regular |
| ENSMUSG00000037416 | Dmxl1 | chr18 | + | 49832669 | 49965473 | regular |
| ENSMUSG00000029817 | Tra2a | chr6 | - | 49243923 | 49264033 | regular |

| | | | | | | |
|---|---|---|---|---|---|---|
| ENSMUSG00000024953 | Prdx5 | chr19 | - | 6906696 | 6910106 | regular |
| ENSMUSG00000027346 | Gpcpd1 | chr2 | - | 132529081 | 132587729 | regular |
| ENSMUSG00000042302 | Ehbp1 | chr11 | - | 22005827 | 22342292 | regular |
| ENSMUSG00000006304 | Arpc2 | chr1 | + | 74236083 | 74268209 | regular |
| ENSMUSG00000071724 | Smpd5 | chr15 | + | 76294433 | 76296896 | regular |
| ENSMUSG00000044566 | Cage1 | chr13 | - | 38006051 | 38037069 | regular |
| ENSMUSG00000052915 | Msl1 | chr11 | + | 98795515 | 98807859 | regular |
| ENSMUSG00000008200 | Fnbp4 | chr2 | + | 90745369 | 90781021 | regular |
| ENSMUSG00000019856 | Fam184a | chr10 | - | 53633144 | 53751123 | regular |
| ENSMUSG00000024789 | Jak2 | chr19 | + | 29251827 | 29313080 | regular |
| ENSMUSG00000000915 | Hip1r | chr5 | + | 123973627 | 124005558 | regular |
| ENSMUST00000108075 | Usp32 | chr11 | - | 85077264 | 85104026 | circular |
| ENSMUSG00000040356 | Skiv2l | chr17 | - | 34839227 | 34850210 | regular |
| ENSMUSG00000034813 | Grip1 | chr10 | + | 119453829 | 120087261 | regular |
| ENSMUSG00000071655 | Ubxn1 | chr19 | + | 8871558 | 8875663 | regular |
| ENSMUSG00000057236 | Rbbp4 | chr4 | - | 129307099 | 129335370 | regular |
| ENSMUSG00000021670 | Hmgcr | chr13 | - | 96648966 | 96670936 | regular |
| ENSMUSG00000039335 | Spata16 | chr3 | + | 26637619 | 26983212 | regular |
| ENSMUSG00000027939 | Nup210l | chr3 | + | 90104131 | 90212048 | regular |
| ENSMUSG00000000131 | Xpo6 | chr7 | - | 126101714 | 126200501 | regular |
| ENSMUSG00000031631 | Cfap97 | chr8 | + | 46151770 | 46195590 | regular |
| ENSMUSG00000020590 | Snx13 | chr12 | + | 35047185 | 35147479 | regular |
| ENSMUSG00000016541 | Atxn10 | chr15 | + | 85336380 | 85463836 | regular |
| ENSMUSG00000058655 | Eif4b | chr15 | + | 102073772 | 102097172 | regular |
| ENSMUSG00000052681 | Rap1b | chr10 | - | 117813870 | 117846035 | regular |
| ENSMUSG00000073731 | AI507597 | chr4 | + | 141614025 | 141615604 | regular |
| ENSMUSG00000054199 | Gon4l | chr3 | + | 88835230 | 88910103 | regular |
| ENSMUSG00000074749 | Kiz | chr2 | + | 146855863 | 146970097 | regular |
| ENSMUSG00000063576 | Klhdc3 | chr17 | - | 46674549 | 46680930 | regular |
| ENSMUSG00000022773 | Ypel1 | chr16 | + | 17069695 | 17087045 | regular |
| ENSMUSG00000037364 | Srrt | chr5 | - | 137295703 | 137307674 | regular |
| ENSMUSG00000026969 | Fam166a | chr2 | + | 25218744 | 25222280 | regular |
| ENSMUSG00000040651 | Fam208a | chr14 | + | 27428833 | 27483555 | regular |
| ENSMUSG00000003746 | Man1a | chr10 | - | 53904784 | 54076609 | regular |
| ENSMUSG00000028557 | Rnf11 | chr4 | - | 109451097 | 109484728 | regular |
| ENSMUSG00000014850 | Msh3 | chr13 | - | 92211871 | 92355003 | regular |
| ENSMUSG00000015597 | Zfp318 | chr17 | + | 46383730 | 46420920 | regular |
| ENSMUSG00000022477 | Aco2 | chr15 | + | 81872308 | 81915133 | regular |
| ENSMUSG00000026466 | Tor1aip1 | chr1 | - | 156004598 | 156036480 | regular |
| ENSMUSG00000035623 | Rsf1 | chr7 | + | 97579888 | 97692778 | regular |
| ENSMUSG00000026288 | Inpp5d | chr1 | + | 87620311 | 87720507 | regular |
| ENSMUSG00000025903 | Lypla1 | chr1 | + | 4807787 | 4848410 | regular |
| ENSMUSG00000041915 | Ammecr1l | chr18 | + | 31759823 | 31784071 | regular |
| ENSMUSG00000034485 | Uaca | chr9 | + | 60794541 | 60880370 | regular |
| ENSMUSG00000061099 | Gapdhs | chr7 | - | 30729774 | 30743681 | regular |
| ENSMUSG00000005615 | Pcyt1a | chr16 | + | 32430920 | 32475070 | regular |

| | | | | | | |
|---|---|---|---|---|---|---|
| ENSMUSG00000028456 | Unc13b | chr4 | + | 43058952 | 43264871 | regular |
| ENSMUSG00000027157 | 4930430A15 | chr2 | - | 111162060 | 111229602 | regular |
| ENSMUSG00000020463 | Ppp4r3b | chr11 | + | 29172889 | 29220797 | regular |
| ENSMUSG00000025220 | Mgea5 | chr19 | - | 45750260 | 45783520 | regular |
| ENSMUSG00000038014 | Fam120a | chr13 | - | 48879218 | 48968017 | regular |
| ENSMUSG00000026182 | Tnp1 | chr1 | - | 73015073 | 73015909 | regular |
| ENSMUSG00000040249 | Lrp1 | chr10 | - | 127538160 | 127621148 | regular |
| ENSMUSG00000038393 | Txnip | chr3 | + | 96557956 | 96561883 | regular |
| ENSMUSG00000033444 | Specc1l | chr10 | + | 75212072 | 75312743 | regular |
| ENSMUSG00000007029 | Vars | chr17 | + | 35000986 | 35016322 | regular |
| ENSMUSG00000019590 | Cyb561 | chr11 | - | 105933701 | 105953336 | regular |
| ENSMUSG00000051113 | Fam71e1 | chr7 | + | 44496580 | 44501486 | regular |
| ENSMUSG00000065037 | Rn7sk | chr9 | - | 78175302 | 78175633 | regular |
| ENSMUSG00000074151 | Nlrc5 | chr8 | + | 94434355 | 94527272 | regular |
| ENSMUSG00000050708 | Ftl1 | chr7 | - | 45457943 | 45459884 | regular |
| ENSMUSG00000022603 | Mroh4 | chr15 | - | 74606028 | 74636353 | regular |
| ENSMUSG00000017404 | Rpl19 | chr11 | + | 98026709 | 98030492 | regular |
| ENSMUSG00000025794 | Rpl14 | chr9 | + | 120571443 | 120574654 | regular |
| ENSMUSG00000042662 | Dusp15 | chr2 | - | 152940996 | 152951698 | regular |
| ENSMUSG00000025982 | Sf3b1 | chr1 | - | 54985168 | 55027481 | regular |
| ENSMUSG00000041650 | Pcca | chr14 | + | 122534323 | 122891100 | regular |
| ENSMUSG00000006676 | Usp19 | chr9 | + | 108490601 | 108502337 | regular |
| ENSMUSG00000039936 | Pik3cd | chr4 | - | 149649167 | 149702571 | regular |
| ENSMUSG00000035569 | Ankrd11 | chr8 | - | 122883821 | 123042277 | regular |
| ENSMUSG00000021065 | Fut8 | chr12 | + | 77238124 | 77476338 | regular |
| ENSMUSG00000000275 | Trim25 | chr11 | + | 88999375 | 89020293 | regular |
| ENSMUSG00000045521 | Tssk2 | chr16 | + | 17898636 | 17900023 | regular |
| ENSMUSG00000018677 | Slc25a39 | chr11 | - | 102402984 | 102407946 | regular |
| ENSMUSG00000076281 | Gm24270 | chr9 | - | 56223714 | 56223771 | regular |
| ENSMUSG00000027665 | Pik3ca | chr3 | + | 32397670 | 32468486 | regular |
| ENSMUSG00000027808 | Serp1 | chr3 | - | 58519816 | 58525892 | regular |
| ENSMUSG00000057672 | Pkn1 | chr8 | - | 83666535 | 83699179 | regular |
| ENSMUSG00000034908 | Sidt2 | chr9 | - | 45937856 | 45955258 | regular |
| ENSMUSG00000035697 | Arhgap45 | chr10 | + | 80016652 | 80031472 | regular |
| ENSMUSG00000030254 | Rad18 | chr6 | - | 112619849 | 112696686 | regular |
| ENSMUSG00000027520 | Zdbf2 | chr1 | + | 63273264 | 63314576 | regular |
| ENSMUSG00000032418 | Me1 | chr9 | - | 86581370 | 86695953 | regular |
| ENSMUSG00000022801 | Lrch3 | chr16 | + | 32914099 | 33015647 | regular |
| ENSMUSG00000028693 | Nasp | chr4 | - | 116601051 | 116627941 | regular |
| ENSMUSG00000018076 | Med13l | chr5 | + | 118560678 | 118765438 | regular |
| ENSMUSG00000027282 | Mtch2 | chr2 | + | 90847154 | 90866810 | regular |
| ENSMUSG00000020594 | Pum2 | chr12 | + | 8674133 | 8752581 | regular |
| ENSMUSG00000030779 | Rbbp6 | chr7 | + | 122965685 | 123002557 | regular |
| ENSMUSG00000032115 | Hyou1 | chr9 | + | 44379489 | 44392369 | regular |
| ENSMUSG00000022995 | Enah | chr1 | - | 181896383 | 182019990 | regular |
| ENSMUSG00000060176 | Kif27 | chr13 | - | 58287501 | 58359122 | regular |

| | | | | | | |
|---|---|---|---|---|---|---|
| ENSMUSG00000032076 | Cadm1 | chr9 | + | 47530172 | 47857637 | regular |
| ENSMUSG00000022521 | Crebbp | chr16 | - | 4081327 | 4213997 | regular |
| ENSMUSG00000038608 | Dock10 | chr1 | - | 80501072 | 80758527 | regular |
| ENSMUSG00000038070 | Cntln | chr4 | + | 84884308 | 85131921 | regular |
| ENSMUSG00000032384 | Csnk1g1 | chr9 | + | 65908923 | 66045015 | regular |
| ENSMUSG00000021285 | Ppp1r13b | chr12 | - | 111828456 | 111908110 | regular |
| ENSMUSG00000019370 | Calm3 | chr7 | - | 16915378 | 16924114 | regular |
| ENSMUSG00000072487 | Mroh5 | chr15 | - | 73761409 | 73839699 | regular |
| ENSMUSG00000025500 | Lmntd2 | chr7 | - | 141209991 | 141214080 | regular |
| ENSMUSG00000056602 | Fry | chr5 | + | 150118644 | 150497753 | regular |
| ENSMUSG00000040054 | Baz2a | chr10 | + | 128091576 | 128129303 | regular |
| ENSMUSG00000038368 | Focad | chr4 | + | 88094628 | 88411011 | regular |
| ENSMUSG00000020532 | Acaca | chr11 | + | 84129671 | 84401651 | regular |
| ENSMUSG00000022562 | Oplah | chr15 | - | 76296596 | 76328015 | regular |
| ENSMUSG00000015243 | Abca1 | chr4 | - | 53030786 | 53159895 | regular |
| ENSMUSG00000079652 | Fam71f2 | chr6 | + | 29279592 | 29290680 | regular |
| ENSMUSG00000039046 | Usp6nl | chr2 | + | 6322666 | 6446390 | regular |
| ENSMUSG00000021891 | Mettl6 | chr14 | - | 31473577 | 31495040 | regular |
| ENSMUSG00000002985 | Apoe | chr7 | - | 19696108 | 19699188 | regular |
| ENSMUSG00000030763 | Lcmt1 | chr7 | + | 123369783 | 123430358 | regular |
| ENSMUSG00000029405 | G3bp2 | chr5 | - | 92052145 | 92083719 | regular |
| ENSMUSG00000031485 | Plpbp | chr8 | + | 27042554 | 27056132 | regular |
| ENSMUSG00000026341 | Actr3 | chr1 | - | 125392904 | 125435727 | regular |
| ENSMUSG00000023944 | Hsp90ab1 | chr17 | - | 45567774 | 45573271 | regular |
| ENSMUSG00000015149 | Sirt2 | chr7 | + | 28766734 | 28788661 | regular |
| ENSMUSG00000076258 | Gm23935 | chr16 | + | 11144124 | 11144181 | regular |
| ENSMUSG00000022108 | Itm2b | chr14 | - | 73362225 | 73385289 | regular |
| ENSMUSG00000019505 | Ubb | chr11 | + | 62551170 | 62553213 | regular |
| ENSMUSG00000020640 | Itsn2 | chr12 | + | 4592637 | 4713962 | regular |
| ENSMUSG00000026277 | Stk25 | chr1 | - | 93619750 | 93658659 | regular |
| ENSMUSG00000045983 | Eif4g1 | chr16 | + | 20668312 | 20692884 | regular |
| ENSMUSG00000029165 | Agbl5 | chr5 | + | 30888693 | 30906965 | regular |
| ENSMUSG00000048827 | Pkd1l3 | chr8 | + | 109614516 | 109674386 | regular |
| ENSMUSG00000058013 | Sep.11 | chr5 | + | 93093436 | 93176447 | regular |
| ENSMUSG00000032872 | Cyb5r4 | chr9 | + | 87022013 | 87077774 | regular |
| ENSMUSG00000020212 | Mdm1 | chr10 | + | 118141810 | 118168997 | regular |
| ENSMUSG00000039100 | Mar.06 | chr15 | - | 31455890 | 31531053 | regular |
| ENSMUSG00000090213 | Tmem189 | chr2 | - | 167642607 | 167689822 | regular |
| ENSMUSG00000063273 | Naa15 | chr3 | + | 51415147 | 51476507 | regular |
| ENSMUSG00000029867 | Llcfc1 | chr6 | + | 41684430 | 41685717 | regular |
| ENSMUSG00000028034 | Fubp1 | chr3 | + | 152210421 | 152236826 | regular |
| ENSMUSG00000022781 | Pak2 | chr16 | - | 32016289 | 32079342 | regular |
| ENSMUSG00000021585 | Cast | chr13 | - | 74692367 | 74808810 | regular |
| ENSMUSG00000037697 | Ddhd1 | chr14 | - | 45588466 | 45658143 | regular |
| ENSMUSG00000019188 | H13 | chr2 | + | 152669460 | 152708670 | regular |
| ENSMUSG00000041144 | Dnah7b | chr1 | + | 46066314 | 46373546 | regular |

| | | | | | | |
|---|---|---|---|---|---|---|
| ENSMUSG00000040152 | Thbs1 | chr2 | + | 118111875 | 118127133 | regular |
| ENSMUSG00000035478 | Mbd3 | chr10 | - | 80392538 | 80399550 | regular |
| ENSMUSG00000042079 | Hnrnpf | chr6 | + | 117900323 | 117925622 | regular |
| ENSMUSG00000000751 | Rpa1 | chr11 | - | 75298165 | 75348324 | regular |
| ENSMUSG00000047284 | Neurl4 | chr11 | + | 69901071 | 69913820 | regular |
| ENSMUSG00000034401 | Spata6 | chr4 | + | 111719983 | 111829184 | regular |
| ENSMUSG00000038612 | Mcl1 | chr3 | + | 95658787 | 95663176 | regular |
| ENSMUSG00000090386 | Mir99ahg | chr16 | + | 77236316 | 77691403 | regular |
| ENSMUSG00000008683 | Rps15a | chr7 | - | 118104371 | 118116188 | regular |
| ENSMUSG00000029313 | Aff1 | chr5 | + | 103692373 | 103855322 | regular |
| ENSMUSG00000037058 | Paip2 | chr18 | + | 35598616 | 35617186 | regular |
| ENSMUSG00000039738 | Slx4 | chr16 | - | 3979104 | 4003770 | regular |
| ENSMUSG00000021140 | Pcnx | chr12 | + | 81860022 | 82000924 | regular |
| ENSMUSG00000036712 | Cyld | chr8 | + | 88697027 | 88751945 | regular |
| ENSMUSG00000020986 | Sec23a | chr12 | - | 58958382 | 59012017 | regular |
| ENSMUSG00000029191 | Rfc1 | chr5 | - | 65261849 | 65335670 | regular |
| ENSMUSG00000028416 | Bag1 | chr4 | - | 40936397 | 40948294 | regular |
| ENSMUSG00000025326 | Ube3a | chr7 | + | 59228749 | 59311536 | regular |
| ENSMUSG00000039001 | Rps21 | chr2 | + | 180257376 | 180258445 | regular |
| ENSMUSG00000089992 | G6pd2 | chr5 | + | 61808815 | 61811163 | regular |
| ENSMUSG00000020681 | Ace | chr11 | + | 105967944 | 105989964 | regular |
| ENSMUSG00000038170 | Pde4dip | chr3 | - | 97689823 | 97888707 | regular |
| ENSMUSG00000020612 | Prkar1a | chr11 | + | 109649404 | 109669656 | regular |
| ENSMUSG00000032534 | Cep63 | chr9 | - | 102584587 | 102626534 | regular |
| ENSMUSG00000049792 | Bag5 | chr12 | - | 111709487 | 111713257 | regular |
| ENSMUSG00000028354 | Fmn2 | chr1 | + | 174501824 | 174822729 | regular |
| ENSMUSG00000038116 | Phf20 | chr2 | + | 156196465 | 156309952 | regular |
| ENSMUSG00000032637 | Atxn2l | chr7 | - | 126491707 | 126503437 | regular |
| ENSMUSG00000040365 | Trim41 | chr11 | - | 48806403 | 48817353 | regular |
| ENSMUSG00000029068 | Ccnl2 | chr4 | + | 155812488 | 155824543 | regular |
| ENSMUSG00000048874 | Phf3 | chr1 | - | 30802338 | 30873921 | regular |
| ENSMUSG00000039886 | Tmem120a | chr5 | - | 135735484 | 135744448 | regular |
| ENSMUSG00000040943 | Tet2 | chr3 | - | 133463678 | 133545139 | regular |
| ENSMUSG00000059456 | Ptk2b | chr14 | - | 66153256 | 66281052 | regular |
| ENSMUSG00000022614 | Lmf2 | chr15 | - | 89351003 | 89355659 | regular |
| ENSMUSG00000036168 | Ccdc38 | chr10 | + | 93540631 | 93584327 | regular |
| ENSMUSG00000051437 | Ubqlnl | chr7 | - | 104148258 | 104150556 | regular |
| ENSMUSG00000058290 | Espl1 | chr15 | + | 102296292 | 102324356 | regular |
| ENSMUST00000166726 | Mast4 | chr13 | - | 103140162 | 103171556 | circular |
| ENSMUSG00000014769 | Psmb1 | chr17 | - | 15475720 | 15498276 | regular |
| ENSMUSG00000026173 | Plcd4 | chr1 | + | 74542887 | 74567794 | regular |
| ENSMUSG00000053838 | Nudcd3 | chr11 | - | 6105690 | 6200415 | regular |
| ENSMUSG00000019917 | Sep.10 | chr10 | - | 59141626 | 59221847 | regular |
| ENSMUSG00000061477 | Rps7 | chr12 | - | 28630853 | 28635953 | regular |
| ENSMUSG00000002413 | Braf | chr6 | - | 39603236 | 39725463 | regular |
| ENSMUSG00000023952 | Gtpbp2 | chr17 | + | 46161031 | 46169370 | regular |

| | | | | | | |
|---|---|---|---|---|---|---|
| ENSMUSG00000018659 | Pnpo | chr11 | - | 96937824 | 96943986 | regular |
| ENSMUSG00000021870 | Slmap | chr14 | - | 26413167 | 26534931 | regular |
| ENSMUSG00000026821 | Ralgds | chr2 | + | 28513124 | 28553081 | regular |
| ENSMUSG00000030588 | Yif1b | chr7 | + | 29238328 | 29247597 | regular |
| ENSMUSG00000038180 | Spag4 | chr2 | + | 156065174 | 156070617 | regular |
| ENSMUSG00000034154 | Ino80 | chr2 | - | 119373041 | 119477687 | regular |
| ENSMUSG00000030583 | Sipa1l3 | chr7 | - | 29320371 | 29518641 | regular |
| ENSMUSG00000063870 | Chd4 | chr6 | + | 125095980 | 125130591 | regular |
| ENSMUSG00000037857 | Nufip2 | chr11 | + | 77686154 | 77741921 | regular |
| ENSMUSG00000054003 | Tdrd9 | chr12 | + | 111971558 | 112068854 | regular |
| ENSMUSG00000039801 | 2410089E03I | chr15 | + | 8169105 | 8271158 | regular |
| ENSMUSG00000048686 | Hmgb4 | chr4 | - | 128260211 | 128260957 | regular |
| ENSMUSG00000044122 | Proca1 | chr11 | + | 78193391 | 78205763 | regular |
| ENSMUSG00000030655 | Smg1 | chr7 | - | 118131307 | 118243670 | regular |
| ENSMUSG00000029828 | 4921507P07I | chr6 | - | 50573301 | 50596632 | regular |
| ENSMUSG00000036646 | Man1b1 | chr2 | + | 25332337 | 25352212 | regular |
| ENSMUSG00000029469 | Ift81 | chr5 | - | 122550203 | 122614518 | regular |
| ENSMUSG00000025245 | Lztfl1 | chr9 | - | 123694415 | 123717697 | regular |
| ENSMUSG00000009073 | Nf2 | chr11 | - | 4765844 | 4849536 | regular |
| ENSMUSG00000049744 | Arhgap15 | chr2 | + | 43748823 | 44395953 | regular |
| ENSMUSG00000071669 | Snx29 | chr16 | + | 11322907 | 11755472 | regular |
| ENSMUSG00000041638 | Gcn1l1 | chr5 | + | 115565253 | 115622654 | regular |
| ENSMUSG00000043050 | Tnp2 | chr16 | - | 10787935 | 10788655 | regular |
| ENSMUSG00000018548 | Trim37 | chr11 | + | 87127076 | 87220683 | regular |
| ENSMUSG00000039568 | Ubald1 | chr16 | - | 4874777 | 4880315 | regular |
| ENSMUSG00000038954 | Supt3 | chr17 | + | 44777151 | 45119290 | regular |
| ENSMUSG00000006014 | Prg4 | chr1 | - | 150449411 | 150466165 | regular |
| ENSMUSG00000029397 | Rchy1 | chr5 | - | 91948903 | 91963068 | regular |
| ENSMUSG00000021846 | Peli2 | chr14 | + | 48120822 | 48281575 | regular |
| ENSMUSG00000104301 | Wdr49 | chr3 | - | 75274987 | 75482156 | regular |
| ENSMUSG00000109864 | Eid3 | chr10 | + | 82866625 | 82867930 | regular |
| ENSMUSG00000070283 | Ndufaf3 | chr9 | - | 108565536 | 108567434 | regular |
| ENSMUSG00000021758 | Ddx4 | chr13 | - | 112598332 | 112652475 | regular |
| ENSMUSG00000022498 | Txndc11 | chr16 | - | 11074910 | 11134650 | regular |
| ENSMUSG00000038060 | Dlec1 | chr9 | + | 119102477 | 119148246 | regular |
| ENSMUSG00000010538 | Tsacc | chr3 | - | 88282756 | 88296999 | regular |
| ENSMUSG00000072825 | Cep170b | chr12 | + | 112720454 | 112746592 | regular |
| ENSMUSG00000031529 | Tnks | chr8 | - | 34826459 | 34965690 | regular |
| ENSMUSG00000022039 | Adam2 | chr14 | - | 66027328 | 66077733 | regular |
| ENSMUSG00000048000 | Gigyf2 | chr1 | + | 87326997 | 87450796 | regular |
| ENSMUSG00000040829 | Zmynd15 | chr11 | + | 70459432 | 70466202 | regular |
| ENSMUSG00000036292 | Gramd1c | chr16 | - | 43980349 | 44063345 | regular |
| ENSMUSG00000043913 | Ccdc60 | chr5 | - | 116124640 | 116288985 | regular |
| ENSMUSG00000073411 | H2-D1 | chr17 | + | 35262729 | 35267499 | regular |
| ENSMUSG00000036241 | Ube2r2 | chr4 | + | 41135742 | 41193380 | regular |
| ENSMUSG00000036707 | Cab39 | chr1 | + | 85793440 | 85851576 | regular |

| | | | | | | |
|---|---|---|---|---|---|---|
| ENSMUSG00000015980 | Lrrc27 | chr7 | + | 139212987 | 139242979 | regular |
| ENSMUSG00000039716 | Dock3 | chr9 | - | 106892824 | 107231909 | regular |
| ENSMUSG00000008393 | Carhsp1 | chr16 | - | 8658586 | 8672153 | regular |
| ENSMUSG00000021244 | Ylpm1 | chr12 | + | 84996320 | 85070515 | regular |
| ENSMUSG00000026809 | Spaca9 | chr2 | - | 28692079 | 28699727 | regular |
| ENSMUSG00000030872 | Gga2 | chr7 | - | 121986721 | 122021222 | regular |
| ENSMUSG00000043621 | Ubxn10 | chr4 | - | 138709836 | 138737167 | regular |
| ENSMUSG00000033478 | Fam160b1 | chr19 | + | 57361008 | 57389594 | regular |
| ENSMUSG00000053624 | Gykl1 | chr18 | + | 52693678 | 52695668 | regular |
| ENSMUSG00000019802 | Sec63 | chr10 | + | 42761495 | 42832514 | regular |
| ENSMUSG00000039456 | Morc3 | chr16 | + | 93832120 | 93876072 | regular |
| ENSMUSG00000026594 | Ralgps2 | chr1 | - | 156804165 | 156939626 | regular |
| ENSMUSG00000048520 | Fbxl13 | chr5 | - | 21483846 | 21645634 | regular |
| ENSMUSG00000027506 | Tpd52 | chr3 | - | 8925592 | 9004723 | regular |
| ENSMUST00000165566 | Rfx3 | chr19 | - | 27849710 | 27900903 | circular |
| ENSMUSG00000090862 | Rps13 | chr7 | - | 116331504 | 116334195 | regular |
| ENSMUSG00000028552 | Eps15 | chr4 | + | 109280267 | 109387817 | regular |
| ENSMUSG00000064302 | Clasp1 | chr1 | + | 118389057 | 118612678 | regular |
| ENSMUSG00000018372 | Cep95 | chr11 | + | 106789251 | 106819930 | regular |
| ENSMUSG00000035086 | Becn1 | chr11 | - | 101285951 | 101302286 | regular |
| ENSMUSG00000050996 | Cetn1 | chr18 | - | 9615523 | 9619478 | regular |
| ENSMUSG00000038127 | Ccdc50 | chr16 | + | 27388868 | 27452218 | regular |
| ENSMUSG00000035539 | Ccdc180 | chr4 | + | 45890302 | 45950774 | regular |
| ENSMUSG00000025583 | Rptor | chr11 | + | 119602904 | 119899576 | regular |
| ENSMUSG00000025508 | Rplp2 | chr7 | + | 141447369 | 141451585 | regular |
| ENSMUSG00000055891 | Ubl4b | chr3 | - | 107553686 | 107555073 | regular |
| ENSMUSG00000022369 | Mtbp | chr15 | + | 55557407 | 55626423 | regular |
| ENSMUSG00000096573 | 1700009J07F | chr10 | + | 77893416 | 77896114 | regular |
| ENSMUSG00000011350 | Gm5893 | chr7 | + | 24818794 | 24840275 | regular |
| ENSMUSG00000029135 | Fosl2 | chr5 | + | 32135800 | 32157842 | regular |
| ENSMUSG00000110333 | Gm45861 | chr8 | + | 27447099 | 27620917 | regular |
| ENSMUSG00000022139 | Mbnl2 | chr14 | + | 120275668 | 120431697 | regular |
| ENSMUSG00000038102 | Trappc11 | chr8 | - | 47490114 | 47533470 | regular |
| ENSMUSG00000020460 | Rps27a | chr11 | - | 29545845 | 29548109 | regular |
| ENSMUSG00000034601 | 2700049A03I | chr12 | + | 71136847 | 71243303 | regular |
| ENSMUSG00000002324 | Rec8 | chr14 | + | 55618036 | 55625395 | regular |
| ENSMUSG00000019795 | Pcmt1 | chr10 | - | 7629372 | 7681136 | regular |
| ENSMUSG00000036403 | Cep135 | chr5 | + | 76588697 | 76646466 | regular |
| ENSMUSG00000029098 | Acox3 | chr5 | + | 35583039 | 35615352 | regular |
| ENSMUSG00000015120 | Ube2i | chr17 | - | 25261915 | 25274622 | regular |
| ENSMUSG00000020358 | Hnrnpab | chr11 | - | 51600099 | 51606847 | regular |
| ENSMUSG00000028382 | Ptbp3 | chr4 | - | 59471867 | 59549364 | regular |
| ENSMUSG00000024112 | Cacna1h | chr17 | - | 25374284 | 25433783 | regular |
| ENSMUSG00000022587 | Ly6e | chr15 | + | 74955050 | 74959905 | regular |
| ENSMUSG00000027778 | Ift80 | chr3 | - | 68892498 | 69004570 | regular |
| ENSMUSG00000026499 | Acbd3 | chr1 | + | 180726042 | 180754204 | regular |

| | | | | | | |
|---|---|---|---|---|---|---|
| ENSMUSG00000035242 | Oaz1 | chr10 | + | 80826655 | 80829290 | regular |
| ENSMUSG00000069135 | Fgfr1op | chr17 | + | 8165500 | 8196804 | regular |
| ENSMUSG00000050623 | Catsperz | chr19 | - | 6922425 | 6925380 | regular |
| ENSMUSG00000024392 | Bag6 | chr17 | + | 35135177 | 35147322 | regular |
| ENSMUSG00000040242 | Fgfr1op2 | chr6 | + | 146577202 | 146599198 | regular |
| ENSMUSG00000070780 | Rbm47 | chr5 | - | 66016548 | 66173118 | regular |
| ENSMUSG00000031553 | Adam3 | chr8 | - | 24677224 | 24725852 | regular |
| ENSMUSG00000058558 | Rpl5 | chr5 | + | 107900501 | 107909005 | regular |
| ENSMUSG00000021458 | 2010111I01R | chr13 | + | 62964892 | 63326096 | regular |
| ENSMUSG00000037112 | Sik2 | chr9 | - | 50892800 | 51009073 | regular |
| ENSMUSG00000014602 | Kif1a | chr1 | - | 93015463 | 93101951 | regular |
| ENSMUSG00000028878 | Fam76a | chr4 | - | 132899212 | 132922558 | regular |
| ENSMUSG00000026131 | Dst | chr1 | + | 33908224 | 34308661 | regular |
| ENSMUSG00000006333 | Rps9 | chr7 | + | 3703992 | 3706897 | regular |
| ENSMUSG00000033335 | Dnm2 | chr9 | + | 21424907 | 21507759 | regular |
| ENSMUSG00000042404 | Dennd4b | chr3 | + | 90265184 | 90280669 | regular |
| ENSMUST00000163257 | Fam172a | chr13 | + | 77951984 | 78006472 | circular |
| ENSMUSG00000063077 | Kif1b | chr4 | - | 149176318 | 149307693 | regular |
| ENSMUSG00000028610 | Dmrtb1 | chr4 | - | 107676289 | 107684230 | regular |
| ENSMUSG00000027878 | Notch2 | chr3 | + | 98013526 | 98150361 | regular |
| ENSMUSG00000032481 | Smarcc1 | chr9 | + | 110117707 | 110240178 | regular |
| ENSMUSG00000026721 | Rabgap1l | chr1 | - | 160219173 | 160793211 | regular |
| ENSMUSG00000038546 | Ranbp9 | chr13 | - | 43402674 | 43481282 | regular |
| ENSMUSG00000048077 | H1fnt | chr15 | - | 98255981 | 98257307 | regular |
| ENSMUSG00000059070 | Rpl18 | chr7 | + | 45715456 | 45720836 | regular |
| ENSMUSG00000025949 | Pikfyve | chr1 | + | 65186682 | 65278695 | regular |
| ENSMUSG00000019763 | Rmnd1 | chr10 | - | 4401914 | 4432388 | regular |
| ENSMUSG00000004040 | Stat3 | chr11 | - | 100885097 | 100939540 | regular |
| ENSMUSG00000018669 | Cdk5rap3 | chr11 | - | 96907425 | 96916496 | regular |
| ENSMUSG00000032285 | Dnaja4 | chr9 | + | 54698872 | 54716315 | regular |
| ENSMUSG00000020319 | Wdpcp | chr11 | + | 21572234 | 21898989 | regular |
| ENSMUSG00000017146 | Brca1 | chr11 | - | 101488763 | 101551955 | regular |
| ENSMUSG00000025495 | Ptdss2 | chr7 | + | 141122381 | 141157606 | regular |
| ENSMUSG00000024160 | Spsb3 | chr17 | + | 24886642 | 24892152 | regular |
| ENSMUSG00000031985 | Gnpat | chr8 | + | 124863032 | 124890057 | regular |
| ENSMUSG00000030323 | Ift122 | chr6 | + | 115853469 | 115926699 | regular |
| ENSMUSG00000022565 | Plec | chr15 | - | 76170973 | 76232574 | regular |
| ENSMUSG00000039191 | Rbpj | chr5 | + | 53466151 | 53657362 | regular |
| ENSMUSG00000029403 | Cdkl2 | chr5 | - | 92006073 | 92043883 | regular |
| ENSMUSG00000024033 | Rsph1 | chr17 | - | 31255018 | 31277356 | regular |
| ENSMUSG00000052299 | Ltn1 | chr16 | - | 87376650 | 87432606 | regular |
| ENSMUSG00000042616 | Oscp1 | chr4 | + | 126058564 | 126092195 | regular |
| ENSMUSG00000020697 | Lig3 | chr11 | + | 82781107 | 82804274 | regular |
| ENSMUSG00000060143 | Gm10076 | chr14 | + | 105681827 | 105682211 | regular |
| ENSMUSG00000005312 | Ubqln1 | chr13 | - | 58176155 | 58215653 | regular |
| ENSMUSG00000032525 | Nktr | chr9 | + | 121719168 | 121756843 | regular |

| | | | | | | |
|---|---|---|---|---|---|---|
| ENSMUSG00000055720 | Ubl7 | chr9 | + | 57910978 | 57929968 | regular |
| ENSMUSG00000017291 | Taok1 | chr11 | - | 77529161 | 77607815 | regular |
| ENSMUSG00000005610 | Eif4g2 | chr7 | - | 111067749 | 111083030 | regular |
| ENSMUSG00000032220 | Myo1e | chr9 | + | 70207349 | 70399766 | regular |
| ENSMUSG00000040936 | Ulk4 | chr9 | - | 120955350 | 121277197 | regular |
| ENSMUSG00000111734 | Gm29825 | chr9 | + | 114118087 | 114192640 | regular |
| ENSMUSG00000019338 | Zfp687 | chr3 | - | 95006589 | 95015448 | regular |
| ENSMUSG00000057440 | Mpp7 | chr18 | - | 7347961 | 7626863 | regular |
| ENSMUSG00000074063 | Osgin1 | chr8 | + | 119434123 | 119446256 | regular |
| ENSMUSG00000056899 | Immp2l | chr12 | + | 41024089 | 41955588 | regular |
| ENSMUSG00000022914 | Brwd1 | chr16 | - | 95992091 | 96082526 | regular |
| ENSMUSG00000031554 | Adam5 | chr8 | - | 24727092 | 24824369 | regular |
| ENSMUSG00000107008 | Gm2762 | chr13 | + | 53393656 | 53455630 | regular |
| ENSMUSG00000028410 | Dnaja1 | chr4 | + | 40722149 | 40737149 | regular |
| ENSMUSG00000021686 | Ap3b1 | chr13 | + | 94358959 | 94566317 | regular |
| ENSMUSG00000027905 | Ddx20 | chr3 | - | 105678269 | 105687574 | regular |
| ENSMUSG00000012609 | Ttll5 | chr12 | + | 85824658 | 86061893 | regular |
| ENSMUSG00000051728 | 4930563D23I | chr16 | - | 92318762 | 92321408 | regular |
| ENSMUSG00000048772 | Tmem53 | chr4 | + | 117251950 | 117268582 | regular |
| ENSMUSG00000042606 | Hirip3 | chr7 | + | 126861971 | 126865377 | regular |
| ENSMUSG00000028525 | Pde4b | chr4 | + | 102087542 | 102607259 | regular |
| ENSMUSG00000075302 | Erich2 | chr2 | + | 70508818 | 70540884 | regular |
| ENSMUSG00000029707 | Fscn3 | chr6 | + | 28427788 | 28438622 | regular |
| ENSMUSG00000021156 | Zmynd11 | chr13 | - | 9684832 | 9765330 | regular |
| ENSMUSG00000020198 | Ap3d1 | chr10 | - | 80706955 | 80742264 | regular |
| ENSMUSG00000085576 | Dpy19l2 | chr9 | - | 24557046 | 24696293 | regular |
| ENSMUSG00000077239 | Snord66 | chr16 | + | 20684238 | 20684313 | regular |
| ENSMUSG00000027751 | Supt20 | chr3 | + | 54692806 | 54728766 | regular |
| ENSMUSG00000064061 | Dzip3 | chr16 | - | 48924231 | 48994165 | regular |
| ENSMUSG00000038388 | Mpp6 | chr6 | + | 50110240 | 50198939 | regular |
| ENSMUSG00000036768 | Kif15 | chr9 | + | 122951045 | 123018733 | regular |
| ENSMUSG00000027708 | Dcun1d1 | chr3 | - | 35892104 | 35937445 | regular |
| ENSMUSG00000029004 | Kmt2e | chr5 | + | 23434440 | 23504235 | regular |
| ENSMUSG00000032396 | Dis3l | chr9 | - | 64306755 | 64341288 | regular |
| ENSMUSG00000042207 | Kdm5b | chr1 | + | 134560170 | 134635285 | regular |
| ENSMUSG00000028141 | Oaz3 | chr3 | - | 94432413 | 94443915 | regular |
| ENSMUSG00000028745 | Capzb | chr4 | + | 139192898 | 139291818 | regular |
| ENSMUSG00000025362 | Rps26 | chr10 | - | 128624533 | 128626747 | regular |
| ENSMUSG00000026766 | Mmadhc | chr2 | - | 50279880 | 50296801 | regular |
| ENSMUSG00000059291 | Rpl11 | chr4 | - | 136028264 | 136053428 | regular |
| ENSMUSG00000025648 | Pfkfb4 | chr9 | + | 108991777 | 109032228 | regular |
| ENSMUSG00000034610 | Zcchc11 | chr4 | + | 108459425 | 108559421 | regular |
| ENSMUSG00000024976 | Shoc2 | chr19 | + | 53944305 | 54033268 | regular |
| ENSMUSG00000004535 | Tax1bp1 | chr6 | + | 52713728 | 52766780 | regular |
| ENSMUSG00000019699 | Akt3 | chr1 | - | 177020072 | 177258203 | regular |
| ENSMUSG00000035126 | Wdr78 | chr4 | - | 103038064 | 103114555 | regular |

| | | | | | | |
|---|---|---|---|---|---|---|
| ENSMUSG00000028845 | Tekt2 | chr4 | - | 126322120 | 126325688 | regular |
| ENSMUSG00000037876 | Jmjd1c | chr10 | + | 67096124 | 67256326 | regular |
| ENSMUSG00000040022 | Rab11fip2 | chr19 | - | 59902883 | 59943654 | regular |
| ENSMUSG00000047412 | Zbtb44 | chr9 | + | 31030643 | 31075885 | regular |
| ENSMUSG00000004035 | Gstm7 | chr3 | - | 107926333 | 107931817 | regular |
| ENSMUSG00000021366 | Hivep1 | chr13 | + | 42052020 | 42192537 | regular |
| ENSMUSG00000041258 | Zfp236 | chr18 | - | 82593592 | 82692883 | regular |
| ENSMUSG00000056770 | Setd3 | chr12 | - | 108106430 | 108179314 | regular |
| ENSMUSG00000051910 | Sox6 | chr7 | - | 115470871 | 116038796 | regular |
| ENSMUSG00000031392 | Irak1 | chrX | - | 74013913 | 74023918 | regular |
| ENSMUSG00000074579 | Lekr1 | chr3 | + | 65666227 | 65831159 | regular |
| ENSMUSG00000045962 | Wnk1 | chr6 | - | 119923968 | 120038672 | regular |
| ENSMUSG00000021147 | Wdr37 | chr13 | - | 8802967 | 8871909 | regular |
| ENSMUSG00000037101 | Ttc29 | chr8 | + | 78213296 | 78394326 | regular |
| ENSMUSG00000041229 | Phf8 | chrX | + | 151520671 | 151633859 | regular |
| ENSMUSG00000086856 | Gm16080 | chr4 | - | 131694606 | 131710508 | regular |
| ENSMUSG00000022403 | St13 | chr15 | - | 81363668 | 81400077 | regular |
| ENSMUSG00000042246 | Tmc7 | chr7 | - | 118535840 | 118584736 | regular |
| ENSMUSG00000028730 | Cfap57 | chr4 | - | 118554550 | 118620777 | regular |
| ENSMUSG00000073418 | C4b | chr17 | - | 34728379 | 34743882 | regular |
| ENSMUSG00000001440 | Kpnb1 | chr11 | - | 97159713 | 97187881 | regular |
| ENSMUSG00000033917 | Gde1 | chr7 | - | 118688544 | 118705778 | regular |
| ENSMUSG00000022641 | Bbx | chr16 | - | 50191843 | 50432390 | regular |
| ENSMUSG00000095892 | Rnu5g | chr9 | - | 65201725 | 65201841 | regular |
| ENSMUSG00000040548 | Tex2 | chr11 | - | 106502146 | 106613423 | regular |
| ENSMUSG00000097075 | D830044I16R | chr7 | - | 126971721 | 126976098 | regular |
| ENSMUSG00000045036 | Tmem232 | chr17 | - | 65256004 | 65540782 | regular |
| ENSMUSG00000043940 | Wdfy3 | chr5 | - | 101832955 | 102069921 | regular |
| ENSMUSG00000040576 | Tmem270 | chr5 | - | 134901592 | 134906733 | regular |
| ENSMUSG00000029577 | Ube3b | chr5 | + | 114380606 | 114421169 | regular |
| ENSMUSG00000039318 | Rab3gap2 | chr1 | + | 185204116 | 185286759 | regular |
| ENSMUSG00000078490 | Cfap74 | chr4 | + | 155409189 | 155466823 | regular |
| ENSMUSG00000025228 | Actr1a | chr19 | - | 46376813 | 46395735 | regular |
| ENSMUSG00000028013 | Ppa2 | chr3 | + | 133310109 | 133378235 | regular |
| ENSMUSG00000039844 | Rapgef1 | chr2 | + | 29619719 | 29740978 | regular |
| ENSMUST00000042373 | Shcbp1l | chr1 | + | 153435917 | 153437630 | circular |
| ENSMUSG00000028414 | Fktn | chr4 | + | 53713997 | 53777890 | regular |
| ENSMUSG00000021258 | Ccnk | chr12 | + | 108179578 | 108203359 | regular |
| ENSMUSG00000033216 | Eefsec | chr6 | - | 88257333 | 88446539 | regular |
| ENSMUSG00000073643 | Wdfy1 | chr1 | - | 79702261 | 79776143 | regular |
| ENSMUSG00000079022 | Col22a1 | chr15 | - | 71795794 | 72034227 | regular |
| ENSMUSG00000060126 | Tpt1 | chr14 | + | 75845092 | 75848525 | regular |
| ENSMUSG00000064370 | mt-Cytb | chrM | + | 14144 | 15288 | regular |
| ENSMUSG00000062995 | Ica1 | chr6 | - | 8630526 | 8778488 | regular |
| ENSMUSG00000053253 | Ndfip2 | chr14 | + | 105258572 | 105309298 | regular |
| ENSMUSG00000078632 | Lrrc37a | chr11 | - | 103451954 | 103504597 | regular |

| | | | | | | |
|---|---|---|---|---|---|---|
| ENSMUSG00000031727 | Pmfbp1 | chr8 | + | 109494026 | 109542640 | regular |
| ENSMUSG00000042298 | Ttc19 | chr11 | + | 62281472 | 62328451 | regular |
| ENSMUSG00000071713 | Csf2rb | chr15 | + | 78325989 | 78351000 | regular |
| ENSMUSG00000005417 | Mprip | chr11 | + | 59661304 | 59780860 | regular |
| ENSMUSG00000009013 | Dynll1 | chr5 | - | 115297109 | 115300999 | regular |
| ENSMUSG00000071226 | Cecr2 | chr6 | + | 120666368 | 120771190 | regular |
| ENSMUSG00000056342 | Usp34 | chr11 | + | 23306894 | 23490560 | regular |
| ENSMUSG00000039218 | Srrm2 | chr17 | + | 23790661 | 23824741 | regular |
| ENSMUSG00000025178 | Pi4k2a | chr19 | + | 42090434 | 42122218 | regular |
| ENSMUSG00000041870 | Ankrd13a | chr5 | + | 114774676 | 114806200 | regular |
| ENSMUSG00000025423 | Pias2 | chr18 | + | 77065207 | 77155708 | regular |
| ENSMUSG00000026083 | Eif5b | chr1 | + | 37998009 | 38055579 | regular |
| ENSMUSG00000029064 | Gnb1 | chr4 | + | 155491360 | 155559269 | regular |
| ENSMUSG00000031776 | Arl2bp | chr8 | + | 94666599 | 94674425 | regular |
| ENSMUSG00000019210 | Atp6v1e1 | chr6 | - | 120794304 | 120822793 | regular |
| ENSMUSG00000024914 | Drap1 | chr19 | - | 5422804 | 5424979 | regular |
| ENSMUSG00000074435 | Smcp | chr3 | - | 92583866 | 92588997 | regular |
| ENSMUSG00000028576 | Ift74 | chr4 | + | 94614490 | 94693229 | regular |
| ENSMUSG00000030189 | Ybx3 | chr6 | - | 131364854 | 131388476 | regular |
| ENSMUSG00000006241 | Ccdc159 | chr9 | + | 21927470 | 21935872 | regular |
| ENSMUSG00000003808 | Farsa | chr8 | + | 84856988 | 84869257 | regular |
| ENSMUSG00000034343 | Ube2f | chr1 | + | 91250303 | 91290337 | regular |
| ENSMUSG00000036196 | Slc26a8 | chr17 | - | 28637782 | 28689987 | regular |
| ENSMUSG00000059834 | Sclt1 | chr3 | - | 41626719 | 41742514 | regular |
| ENSMUSG00000060771 | Tsga10 | chr1 | - | 37754775 | 37866429 | regular |
| ENSMUSG00000038936 | Sccpdh | chr1 | + | 179668209 | 179687189 | regular |
| ENSMUSG00000021557 | Agtpbp1 | chr13 | - | 59445741 | 59585227 | regular |
| ENSMUSG00000095738 | Gm25313 | chr9 | - | 65196571 | 65196687 | regular |
| ENSMUSG00000066900 | Suds3 | chr5 | - | 117091679 | 117116113 | regular |
| ENSMUSG00000041453 | Rpl21 | chr5 | + | 146832889 | 146837032 | regular |
| ENSMUSG00000052707 | Tnrc6a | chr7 | + | 123123884 | 123195296 | regular |
| ENSMUSG00000015755 | Tab2 | chr10 | - | 7905652 | 7956230 | regular |
| ENSMUSG00000092702 | Gm24514 | chr18 | - | 55622635 | 55622920 | regular |
| ENSMUSG00000022100 | Xpo7 | chr14 | - | 70654245 | 70766628 | regular |
| ENSMUSG00000003660 | Snrnp200 | chr2 | + | 127208385 | 127240451 | regular |
| ENSMUSG00000020475 | Pgam2 | chr11 | - | 5801639 | 5803733 | regular |
| ENSMUSG00000010608 | Rbm25 | chr12 | + | 83631235 | 83683123 | regular |
| ENSMUSG00000026361 | Cdc73 | chr1 | - | 143598799 | 143702893 | regular |
| ENSMUSG00000055932 | Fto | chr8 | + | 91313524 | 91668439 | regular |
| ENSMUSG00000031770 | Herpud1 | chr8 | + | 94386437 | 94395377 | regular |
| ENSMUSG00000067288 | Rps28 | chr17 | - | 33819026 | 33824562 | regular |
| ENSMUSG00000037013 | Ss18 | chr18 | - | 14624197 | 14682914 | regular |
| ENSMUSG00000031672 | Got2 | chr8 | - | 95864133 | 95888547 | regular |
| ENSMUSG00000034913 | Spert | chr14 | - | 75582876 | 75593116 | regular |
| ENSMUSG00000000411 | Tssk3 | chr4 | - | 129489013 | 129491022 | regular |
| ENSMUSG00000070345 | Hsf5 | chr11 | + | 87617163 | 87659542 | regular |

| | | | | | | |
|---|---|---|---|---|---|---|
| ENSMUSG00000019969 | Psen1 | chr12 | + | 83688151 | 83735199 | regular |
| ENSMUSG00000051736 | Fam229b | chr10 | - | 39118803 | 39133914 | regular |
| ENSMUSG00000003341 | Atp8b3 | chr10 | - | 80519583 | 80539124 | regular |
| ENSMUSG00000024169 | Ift140 | chr17 | + | 25016090 | 25099495 | regular |
| ENSMUSG00000031320 | Rps4x | chrX | - | 102184940 | 102189394 | regular |
| ENSMUSG00000028793 | Rnf19b | chr4 | + | 129058270 | 129085886 | regular |
| ENSMUSG00000064299 | 4921528I07R | chr9 | - | 114237696 | 114317028 | regular |
| ENSMUSG00000047888 | Tnrc6b | chr15 | + | 80711312 | 80941085 | regular |
| ENSMUSG00000037683 | Armc3 | chr2 | + | 19199301 | 19310241 | regular |
| ENSMUSG00000028161 | Ppp3ca | chr3 | + | 136670123 | 136937727 | regular |
| ENSMUSG00000031537 | Ikbkb | chr8 | - | 22659211 | 22706589 | regular |
| ENSMUSG00000040423 | Rc3h1 | chr1 | + | 160906417 | 160974978 | regular |
| ENSMUSG00000055137 | Sugct | chr13 | - | 16857471 | 17695553 | regular |
| ENSMUSG00000020143 | Dock2 | chr11 | - | 34226814 | 34783892 | regular |
| ENSMUSG00000079356 | Mettl4-ps1 | chr17 | + | 71526895 | 71556543 | regular |
| ENSMUSG00000074146 | 4930579C12I | chr9 | - | 89113110 | 89177887 | regular |
| ENSMUSG00000025920 | Stau2 | chr1 | - | 16228673 | 16520112 | regular |
| ENSMUSG00000020728 | Cep112 | chr11 | + | 108425191 | 108860615 | regular |
| ENSMUSG00000051427 | Ccdc157 | chr11 | - | 4141122 | 4160293 | regular |
| ENSMUSG00000065824 | Gm26315 | chr2 | - | 7076093 | 7076414 | regular |
| ENSMUSG00000030761 | Myo7a | chr7 | - | 98051059 | 98119524 | regular |
| ENSMUSG00000025239 | Limd1 | chr9 | + | 123478705 | 123521552 | regular |
| ENSMUSG00000020694 | Tlk2 | chr11 | + | 105178806 | 105283959 | regular |
| ENSMUSG00000070565 | Rasal2 | chr1 | - | 157135181 | 157412595 | regular |
| ENSMUSG00000004798 | Ulk2 | chr11 | - | 61775648 | 61855073 | regular |
| ENSMUSG00000089872 | Rps6kc1 | chr1 | - | 190700201 | 190911770 | regular |
| ENSMUSG00000060802 | B2m | chr2 | + | 122147685 | 122153083 | regular |
| ENSMUSG00000024576 | Csnk1a1 | chr18 | + | 61555273 | 61590061 | regular |
| ENSMUSG00000078429 | Ctdsp2 | chr10 | + | 126978716 | 126999975 | regular |
| ENSMUSG00000040111 | Gramd1b | chr9 | - | 40293232 | 40531383 | regular |
| ENSMUSG00000027523 | Gnas | chr2 | + | 174284319 | 174346744 | regular |
| ENSMUSG00000023873 | 1700010I14R | chr17 | + | 8988332 | 9008319 | regular |
| ENSMUSG00000021143 | Pacs2 | chr12 | + | 113014507 | 113074401 | regular |
| ENSMUSG00000020946 | Gosr2 | chr11 | - | 103676848 | 103697898 | regular |
| ENSMUSG00000055319 | Sec23ip | chr7 | + | 128744942 | 128784836 | regular |
| ENSMUSG00000026491 | Ahctf1 | chr1 | - | 179744893 | 179803680 | regular |
| ENSMUSG00000029647 | Pan3 | chr5 | + | 147430160 | 147548502 | regular |
| ENSMUSG00000022855 | Senp2 | chr16 | + | 22009483 | 22049269 | regular |
| ENSMUSG00000001415 | Smg5 | chr3 | + | 88336259 | 88362338 | regular |
| ENSMUSG00000038967 | Pdk2 | chr11 | - | 95026257 | 95041354 | regular |
| ENSMUSG00000022542 | Sep.12 | chr16 | - | 4986857 | 4997852 | regular |
| ENSMUSG00000068922 | Msto1 | chr3 | - | 88905106 | 88913999 | regular |
| ENSMUSG00000041841 | Rpl37 | chr15 | + | 5116644 | 5119140 | regular |
| ENSMUSG00000049739 | Zfp646 | chr7 | + | 127876220 | 127885996 | regular |
| ENSMUSG00000029463 | Fam216a | chr5 | - | 122364579 | 122372364 | regular |
| ENSMUSG00000002372 | Ranbp3 | chr17 | + | 56673224 | 56711769 | regular |

| | | | | | | |
|---|---|---|---|---|---|---|
| ENSMUSG00000016253 | Nelfcd | chr2 | + | 174415803 | 174427502 | regular |
| ENSMUSG00000056692 | D17Wsu92e | chr17 | - | 27751231 | 27820558 | regular |
| ENSMUSG00000020014 | Cfap54 | chr10 | - | 92775618 | 93081618 | regular |
| ENSMUST00000039798 | Bbs9 | chr9 | + | 22643744 | 22659145 | circular |
| ENSMUSG00000039391 | Ccdc81 | chr7 | - | 89866147 | 89903629 | regular |
| ENSMUSG00000025142 | Aspscr1 | chr11 | + | 120672972 | 120709447 | regular |
| ENSMUSG00000009079 | Ewsr1 | chr11 | - | 5069688 | 5099266 | regular |
| ENSMUSG00000028452 | Vcp | chr4 | - | 42979962 | 43000507 | regular |
| ENSMUSG00000025278 | Flnb | chr14 | + | 7817956 | 7951588 | regular |
| ENSMUSG00000039041 | Adrm1 | chr2 | + | 180171484 | 180176286 | regular |
| ENSMUSG00000058318 | Phf21a | chr2 | + | 92093116 | 92364666 | regular |
| ENSMUSG00000023467 | Tulp2 | chr7 | + | 45481876 | 45523632 | regular |
| ENSMUSG00000062210 | Tnfaip8 | chr18 | + | 49979426 | 50107173 | regular |
| ENSMUSG00000020024 | Cep83 | chr10 | + | 94688653 | 94790853 | regular |
| ENSMUSG00000083899 | Gm12346 | chr11 | + | 77293443 | 77295898 | regular |
| ENSMUSG00000041297 | Cdk13 | chr13 | - | 17710314 | 17805097 | regular |
| ENSMUSG00000004897 | Hdgf | chr3 | + | 87906320 | 87916132 | regular |
| ENSMUSG00000000085 | Scmh1 | chr4 | + | 120405280 | 120530186 | regular |
| ENSMUSG00000020859 | Spag9 | chr11 | + | 93996090 | 94126085 | regular |
| ENSMUSG00000036273 | Lrrk2 | chr15 | + | 91673174 | 91816120 | regular |
| ENSMUSG00000034940 | Synrg | chr11 | + | 83964427 | 84044578 | regular |
| ENSMUSG00000064202 | 4430402I18R | chr19 | - | 28893041 | 28967800 | regular |
| ENSMUSG00000034574 | Daam1 | chr12 | + | 71831077 | 71992333 | regular |
| ENSMUSG00000021771 | Vdac2 | chr14 | + | 21825237 | 21845879 | regular |
| ENSMUSG00000004356 | Utp20 | chr10 | - | 88746606 | 88826804 | regular |
| ENSMUSG00000020456 | Ogdh | chr11 | + | 6291632 | 6356642 | regular |
| ENSMUSG00000039977 | Deup1 | chr9 | - | 15559863 | 15627933 | regular |
| ENSMUSG00000021024 | Psma6 | chr12 | + | 55384221 | 55418454 | regular |
| ENSMUSG00000071644 | Eef1g | chr19 | + | 8967040 | 8978479 | regular |
| ENSMUSG00000022214 | Dcaf11 | chr14 | + | 55560005 | 55570065 | regular |
| ENSMUSG00000022805 | Maats1 | chr16 | - | 38297753 | 38342143 | regular |
| ENSMUSG00000090841 | Myl6 | chr10 | - | 128490859 | 128494145 | regular |
| ENSMUSG00000026504 | Sdccag8 | chr1 | + | 176814659 | 177020437 | regular |
| ENSMUSG00000035131 | Brinp3 | chr1 | + | 146494759 | 146902472 | regular |
| ENSMUSG00000024937 | Ehbp1l1 | chr19 | - | 5707375 | 5726317 | regular |
| ENSMUSG00000026867 | Gapvd1 | chr2 | - | 34674593 | 34755232 | regular |
| ENSMUSG00000037270 | 4932438A13I | chr3 | + | 36863103 | 37053033 | regular |
| ENSMUSG00000028551 | Cdkn2c | chr4 | - | 109660875 | 109667189 | regular |
| ENSMUSG00000004789 | Dlst | chr12 | + | 85110832 | 85134845 | regular |
| ENSMUSG00000028136 | Snx27 | chr3 | - | 94497543 | 94582716 | regular |
| ENSMUSG00000046991 | Wdr27 | chr17 | - | 14818671 | 14943158 | regular |
| ENSMUSG00000049676 | Catsperg1 | chr7 | - | 29181320 | 29214035 | regular |
| ENSMUSG00000047021 | Cfap65 | chr1 | - | 74902070 | 74935599 | regular |
| ENSMUSG00000034120 | Srsf2 | chr11 | - | 116849900 | 116853094 | regular |
| ENSMUSG00000065911 | Gm24447 | chr15 | + | 93234362 | 93234700 | regular |
| ENSMUSG00000057246 | BC051142 | chr17 | + | 34398819 | 34460734 | regular |

| | | | | | | |
|---|---|---|---|---|---|---|
| ENSMUSG00000006527 | Sfmbt1 | chr14 | + | 30714848 | 30822721 | regular |
| ENSMUSG00000036825 | Ssx2ip | chr3 | + | 146404641 | 146440144 | regular |
| ENSMUSG00000048696 | Mex3d | chr10 | - | 80380354 | 80387659 | regular |
| ENSMUSG00000007739 | Cct4 | chr11 | + | 22990518 | 23003780 | regular |
| ENSMUSG00000040312 | Cchcr1 | chr17 | + | 35517099 | 35531015 | regular |
| ENSMUSG00000022867 | Usp25 | chr16 | + | 77014068 | 77116779 | regular |
| ENSMUSG00000038039 | Gcc2 | chr10 | + | 58255496 | 58305599 | regular |
| ENSMUSG00000037149 | Ddx1 | chr12 | - | 13216972 | 13249213 | regular |
| ENSMUSG00000027433 | Xrn2 | chr2 | + | 147012995 | 147078000 | regular |
| ENSMUSG00000078908 | Mon1b | chr8 | + | 113635586 | 113645179 | regular |
| ENSMUSG00000075706 | Gpx4 | chr10 | + | 80047165 | 80056439 | regular |
| ENSMUSG00000037979 | Ccdc92 | chr5 | - | 124834417 | 124862424 | regular |
| ENSMUSG00000030344 | Akap3 | chr6 | + | 126853097 | 126874308 | regular |
| ENSMUSG00000036872 | Abcc12 | chr8 | - | 86482259 | 86580686 | regular |
| ENSMUSG00000069495 | Epc2 | chr2 | + | 49451485 | 49551948 | regular |
| ENSMUSG00000028423 | Nfx1 | chr4 | + | 40970905 | 41025993 | regular |
| ENSMUSG00000020802 | Ube2o | chr11 | - | 116537739 | 116581447 | regular |
| ENSMUSG00000028800 | Hdac1 | chr4 | - | 129516103 | 129542713 | regular |
| ENSMUSG00000033055 | Ankrd54 | chr15 | - | 79053093 | 79062893 | regular |
| ENSMUSG00000112657 | BC106175 | chr12 | - | 8478873 | 8482157 | regular |
| ENSMUSG00000040838 | Gm11639 | chr11 | + | 104685706 | 105117394 | regular |
| ENSMUSG00000039477 | Tnrc18 | chr5 | - | 142724660 | 142817662 | regular |
| ENSMUSG00000028953 | Abcf2 | chr5 | - | 24565344 | 24577467 | regular |
| ENSMUSG00000036613 | Tssc1 | chr12 | + | 28751802 | 28869865 | regular |
| ENSMUSG00000032352 | Lrrc1 | chr9 | - | 77430822 | 77544870 | regular |
| ENSMUSG00000020250 | Txnrd1 | chr10 | + | 82833950 | 82897712 | regular |
| ENSMUSG00000060098 | Prmt7 | chr8 | + | 106210935 | 106252794 | regular |
| ENSMUSG00000052139 | Babam2 | chr5 | + | 31697683 | 32084962 | regular |
| ENSMUSG00000025369 | Smarcc2 | chr10 | + | 128459247 | 128490482 | regular |
| ENSMUSG00000048249 | Crebrf | chr17 | + | 26715649 | 26776635 | regular |
| ENSMUSG00000038068 | Rnf144b | chr13 | + | 47122655 | 47247991 | regular |
| ENSMUSG00000011179 | Odc1 | chr12 | + | 17544793 | 17551505 | regular |
| ENSMUSG00000022401 | Xpnpep3 | chr15 | + | 81400137 | 81456807 | regular |
| ENSMUSG00000025025 | Mxi1 | chr19 | + | 53310505 | 53375810 | regular |
| ENSMUSG00000048416 | Mlf1 | chr3 | + | 67374096 | 67400003 | regular |
| ENSMUSG00000063229 | Ldha | chr7 | + | 46841474 | 46855627 | regular |
| ENSMUSG00000076433 | Cep295nl | chr11 | - | 118332359 | 118342500 | regular |
| ENSMUSG00000022285 | Ywhaz | chr15 | - | 36770769 | 36796929 | regular |
| ENSMUSG00000001472 | Tcf25 | chr8 | + | 123373752 | 123403835 | regular |
| ENSMUSG00000032096 | Arcn1 | chr9 | - | 44741563 | 44767845 | regular |
| ENSMUSG00000030738 | Eif3c | chr7 | - | 126546454 | 126566411 | regular |
| ENSMUSG00000044937 | Ttc41 | chr10 | + | 86705810 | 86776844 | regular |
| ENSMUSG00000024610 | Cd74 | chr18 | + | 60803847 | 60812646 | regular |
| ENSMUSG00000005802 | Slc30a4 | chr2 | - | 122681232 | 122702663 | regular |
| ENSMUSG00000028760 | Eif4g3 | chr4 | + | 137993021 | 138208508 | regular |
| ENSMUSG00000042605 | Atxn2 | chr5 | + | 121711336 | 121816493 | regular |

| | | | | | | |
|---|---|---|---|---|---|---|
| ENSMUSG00000040562 | Gstm2 | chr3 | - | 107981701 | 107986453 | regular |
| ENSMUSG00000020102 | Slc16a7 | chr10 | - | 125219269 | 125389465 | regular |
| ENSMUSG00000040865 | Ino80d | chr1 | - | 62958417 | 63114667 | regular |
| ENSMUSG00000022601 | Zbtb11 | chr16 | + | 55973882 | 56008913 | regular |
| ENSMUSG00000020020 | Usp44 | chr10 | + | 93831554 | 93858088 | regular |
| ENSMUSG00000057176 | Ccdc189 | chr7 | - | 127582382 | 127588595 | regular |
| ENSMUSG00000028465 | Tln1 | chr4 | - | 43531518 | 43562691 | regular |
| ENSMUSG00000038371 | Sbf2 | chr7 | - | 110308012 | 110614922 | regular |
| ENSMUSG00000047921 | Trappc9 | chr15 | - | 72589620 | 73061204 | regular |
| ENSMUSG00000070531 | Wfdc6b | chr2 | + | 164613521 | 164618212 | regular |
| ENSMUSG00000029174 | Tbc1d1 | chr5 | + | 64156304 | 64351486 | regular |
| ENSMUSG00000071076 | Jund | chr8 | + | 70698948 | 70700616 | regular |
| ENSMUSG00000036377 | C530008M17 | chr5 | + | 76656511 | 76873554 | regular |
| ENSMUSG00000030172 | Erc1 | chr6 | - | 119570795 | 119848167 | regular |
| ENSMUSG00000002768 | Mea1 | chr17 | + | 46681136 | 46683105 | regular |
| ENSMUSG00000040481 | Bptf | chr11 | - | 107033080 | 107132127 | regular |
| ENSMUSG00000009112 | Bcl2l13 | chr6 | + | 120836211 | 120892842 | regular |
| ENSMUSG00000074149 | Gm10634 | chr9 | + | 88719628 | 88806264 | regular |
| ENSMUSG00000027286 | Lrrc57 | chr2 | - | 120604237 | 120609520 | regular |
| ENSMUSG00000001175 | Calm1 | chr12 | + | 100199434 | 100209814 | regular |
| ENSMUSG00000046337 | Fam178b | chr1 | - | 36562691 | 36683183 | regular |
| ENSMUSG00000023026 | Dip2b | chr15 | + | 100038663 | 100219473 | regular |
| ENSMUSG00000050107 | Haspin | chr11 | - | 73135484 | 73138294 | regular |
| ENSMUSG00000025130 | P4hb | chr11 | - | 120560297 | 120573253 | regular |
| ENSMUSG00000007892 | Rplp1 | chr9 | - | 61913283 | 61914542 | regular |
| ENSMUSG00000022710 | Usp7 | chr16 | - | 8689594 | 8792308 | regular |
| ENSMUSG00000036882 | Arhgap33 | chr7 | - | 30522225 | 30535060 | regular |
| ENSMUSG00000079435 | Rpl36a | chrX | + | 134585653 | 134588062 | regular |
| ENSMUSG00000061186 | Sfmbt2 | chr2 | + | 10370509 | 10595253 | regular |
| ENSMUSG00000037119 | Fam91a1 | chr15 | + | 58415467 | 58457740 | regular |
| ENSMUSG00000030203 | Dusp16 | chr6 | - | 134715467 | 134792625 | regular |
| ENSMUSG00000029001 | Fbxo44 | chr4 | - | 148152799 | 148160482 | regular |
| ENSMUSG00000036995 | Asap3 | chr4 | + | 136206364 | 136245216 | regular |
| ENSMUSG00000024654 | Asrgl1 | chr19 | - | 9109867 | 9135636 | regular |
| ENSMUSG00000042590 | Ipo11 | chr13 | - | 106794438 | 106936958 | regular |
| ENSMUSG00000023150 | Ivns1abp | chr1 | + | 151344476 | 151364422 | regular |
| ENSMUSG00000060268 | Gm1661 | chr4 | - | 117213332 | 117252125 | regular |
| ENSMUSG00000029621 | Arpc1a | chr5 | + | 145083829 | 145108761 | regular |
| ENSMUSG00000044477 | Zfand3 | chr17 | + | 30005050 | 30210828 | regular |
| ENSMUSG00000044083 | Efcab8 | chr2 | + | 153780878 | 153844752 | regular |
| ENSMUSG00000066368 | Actl11 | chr9 | + | 107928468 | 107932461 | regular |
| ENSMUSG00000023072 | Cep89 | chr7 | + | 35397034 | 35438689 | regular |
| ENSMUSG00000036557 | Stpg4 | chr17 | - | 87389570 | 87427741 | regular |
| ENSMUSG00000052146 | Rps10 | chr17 | - | 27630421 | 27636629 | regular |
| ENSMUSG00000026017 | Carf | chr1 | + | 60098246 | 60153953 | regular |
| ENSMUSG00000024122 | Pdpk1 | chr17 | - | 24073679 | 24150924 | regular |

| | | | | | | |
|---|---|---|---|---|---|---|
| ENSMUSG00000037174 | Elf2 | chr3 | - | 51252719 | 51340663 | regular |
| ENSMUSG00000021012 | Zc3h14 | chr12 | + | 98746963 | 98787753 | regular |
| ENSMUSG00000018995 | Nars2 | chr7 | + | 96951504 | 97064758 | regular |
| ENSMUSG00000027562 | Car2 | chr3 | + | 14886272 | 14900770 | regular |
| ENSMUSG00000075227 | Znhit2 | chr19 | + | 6061191 | 6062472 | regular |
| ENSMUSG00000021273 | Fdft1 | chr14 | - | 63145149 | 63179578 | regular |
| ENSMUSG00000027564 | Cypt12 | chr3 | + | 17948409 | 17948959 | regular |
| ENSMUSG00000024207 | Acsbg2 | chr17 | - | 56843102 | 56874447 | regular |
| ENSMUSG00000038260 | Trpm4 | chr7 | - | 45302631 | 45333780 | regular |
| ENSMUSG00000015095 | Fbxw5 | chr2 | + | 25500749 | 25505471 | regular |
| ENSMUSG00000028063 | Lmna | chr3 | - | 88480146 | 88509956 | regular |
| ENSMUSG00000017774 | Myo1c | chr11 | + | 75650503 | 75673910 | regular |
| ENSMUSG00000059114 | Lrrc74a | chr12 | + | 86734368 | 86763797 | regular |
| ENSMUSG00000036990 | Otud4 | chr8 | + | 79639617 | 79677724 | regular |
| ENSMUSG00000039361 | Picalm | chr7 | + | 90130212 | 90213465 | regular |
| ENSMUSG00000052075 | 1700029F12I | chr13 | - | 97021863 | 97034362 | regular |
| ENSMUSG00000024949 | Sf1 | chr19 | + | 6363689 | 6378030 | regular |
| ENSMUSG00000037822 | Smim14 | chr5 | - | 65446843 | 65537184 | regular |
| ENSMUSG00000026004 | Kansl1l | chr1 | - | 66719247 | 66817562 | regular |
| ENSMUSG00000003153 | Slc2a3 | chr6 | - | 122727808 | 122801640 | regular |
| ENSMUSG00000033237 | Arid2 | chr15 | + | 96287517 | 96404992 | regular |
| ENSMUSG00000004677 | Myo9b | chr8 | + | 71272713 | 71360713 | regular |
| ENSMUSG00000031958 | Ldhd | chr8 | - | 111623784 | 111630374 | regular |
| ENSMUSG00000020275 | Rel | chr11 | - | 23736846 | 23770970 | regular |
| ENSMUSG00000022019 | Tdrd3 | chr14 | + | 87416638 | 87545504 | regular |
| ENSMUSG00000022800 | Fyttd1 | chr16 | + | 32877499 | 32908963 | regular |
| ENSMUSG00000000127 | Fer | chr17 | + | 63896017 | 64139494 | regular |
| ENSMUSG00000032009 | Sesn3 | chr9 | + | 14275066 | 14333101 | regular |
| ENSMUSG00000021182 | Ccdc88c | chr12 | - | 100911522 | 101029056 | regular |
| ENSMUSG00000025871 | 4833439L19F | chr13 | - | 54551217 | 54565435 | regular |
| ENSMUSG00000041629 | Fam104a | chr11 | - | 113661318 | 113684155 | regular |
| ENSMUSG00000061607 | Mdc1 | chr17 | + | 35841514 | 35859670 | regular |
| ENSMUSG00000059495 | Arhgef12 | chr9 | - | 42963841 | 43107239 | regular |
| ENSMUSG00000035992 | Fnip1 | chr11 | + | 54438198 | 54518235 | regular |
| ENSMUSG00000002733 | Plekha3 | chr2 | + | 76675280 | 76696828 | regular |
| ENSMUSG00000038679 | Trps1 | chr15 | - | 50654751 | 50890463 | regular |
| ENSMUSG00000045257 | Morn2 | chr17 | + | 80290202 | 80297476 | regular |
| ENSMUSG00000024853 | Sf3b2 | chr19 | - | 5273922 | 5295455 | regular |
| ENSMUSG00000031788 | Kifc3 | chr8 | - | 95099827 | 95202812 | regular |
| ENSMUSG00000030557 | Mef2a | chr7 | - | 67231162 | 67372858 | regular |
| ENSMUSG00000041570 | Camsap2 | chr1 | - | 136268122 | 136346104 | regular |
| ENSMUSG00000067367 | Lyar | chr5 | + | 38220469 | 38234306 | regular |
| ENSMUSG00000057863 | Rpl36 | chr17 | + | 56613394 | 56614243 | regular |
| ENSMUSG00000021983 | Atp8a2 | chr14 | - | 59638539 | 60197179 | regular |
| ENSMUSG00000000125 | Wnt3 | chr11 | + | 103774149 | 103817957 | regular |
| ENSMUSG00000031885 | Cbfb | chr8 | + | 105170673 | 105217989 | regular |

| | | | | | | |
|---|---|---|---|---|---|---|
| ENSMUSG00000004865 | Srpk1 | chr17 | - | 28587647 | 28622521 | regular |
| ENSMUSG00000029131 | Dnajb6 | chr5 | + | 29735687 | 29818134 | regular |
| ENSMUSG00000032336 | Nptn | chr9 | + | 58582239 | 58657955 | regular |
| ENSMUSG00000005683 | Cs | chr10 | + | 128337733 | 128362479 | regular |
| ENSMUSG00000030007 | Cct7 | chr6 | + | 85451513 | 85468475 | regular |
| ENSMUSG00000063450 | Syne2 | chr12 | + | 75818133 | 76110926 | regular |
| ENSMUSG00000057551 | Zfp317 | chr9 | + | 19622101 | 19649731 | regular |
| ENSMUSG00000020844 | Nxn | chr11 | - | 76257197 | 76399140 | regular |
| ENSMUSG00000003123 | Lipe | chr7 | - | 25379526 | 25398710 | regular |
| ENSMUSG00000043411 | Usp48 | chr4 | + | 137593754 | 137658537 | regular |
| ENSMUSG00000036918 | Ttc7 | chr17 | + | 87282885 | 87381769 | regular |
| ENSMUSG00000038324 | Trpc4ap | chr2 | - | 155634270 | 155692384 | regular |
| ENSMUSG00000020576 | Nbas | chr12 | + | 13269132 | 13583811 | regular |
| ENSMUSG00000051900 | Abca16 | chr7 | + | 120409646 | 120544813 | regular |
| ENSMUSG00000020954 | Strn3 | chr12 | - | 51609631 | 51691897 | regular |
| ENSMUSG00000037386 | Rims2 | chr15 | + | 39198260 | 39684372 | regular |
| ENSMUSG00000030792 | Dkkl1 | chr7 | - | 45207518 | 45212147 | regular |
| ENSMUSG00000036501 | Fam13b | chr18 | - | 34442351 | 34506823 | regular |
| ENSMUSG00000021270 | Hsp90aa1 | chr12 | - | 110690604 | 110702728 | regular |
| ENSMUSG00000031583 | Wrn | chr8 | - | 33234383 | 33385527 | regular |
| ENSMUSG00000037020 | Wdr62 | chr7 | - | 30240137 | 30280419 | regular |
| ENSMUSG00000030882 | Dnhd1 | chr7 | + | 105650826 | 105721799 | regular |
| ENSMUSG00000031010 | Usp9x | chrX | + | 13071497 | 13173328 | regular |
| ENSMUSG00000021254 | Gpatch2l | chr12 | + | 86241857 | 86291784 | regular |
| ENSMUSG00000038485 | Socs7 | chr11 | + | 97362550 | 97398542 | regular |
| ENSMUSG00000021948 | Prkcd | chr14 | - | 30595353 | 30626210 | regular |
| ENSMUSG00000038369 | Ncoa6 | chr2 | - | 155390655 | 155473894 | regular |
| ENSMUSG00000029111 | Nelfa | chr5 | - | 33897915 | 33936413 | regular |
| ENSMUSG00000021189 | Atxn3 | chr12 | - | 101918900 | 101958246 | regular |
| ENSMUSG00000006818 | Sod2 | chr17 | + | 13007838 | 13018119 | regular |
| ENSMUSG00000024759 | Atl3 | chr19 | + | 7494039 | 7538608 | regular |
| ENSMUSG00000048206 | Dnajb8 | chr6 | + | 88222267 | 88254415 | regular |
| ENSMUSG00000058019 | Ces5a | chr8 | - | 93499063 | 93535830 | regular |
| ENSMUSG00000018567 | Gabarap | chr11 | + | 69991142 | 69994951 | regular |
| ENSMUSG00000020850 | Prpf8 | chr11 | + | 75486815 | 75509449 | regular |
| ENSMUSG00000026482 | Rgl1 | chr1 | - | 152516759 | 152766351 | regular |
| ENSMUSG00000026134 | Prim2 | chr1 | - | 33453809 | 33669795 | regular |
| ENSMUSG00000034543 | Morc2a | chr11 | + | 3649493 | 3690477 | regular |
| ENSMUSG00000021940 | Ptpn20 | chr14 | + | 33589206 | 33640754 | regular |
| ENSMUSG00000050299 | Gm9843 | chr16 | - | 76403208 | 76403712 | regular |
| ENSMUSG00000039630 | Hnrnpu | chr1 | - | 178321107 | 178337797 | regular |
| ENSMUSG00000033004 | Mycbp2 | chr14 | - | 103113410 | 103346814 | regular |
| ENSMUSG00000110841 | Gpx4-ps2 | chr10 | + | 45875290 | 45876086 | regular |
| ENSMUSG00000047720 | 4922502D21 | chr6 | - | 129322163 | 129331796 | regular |
| ENSMUSG00000034334 | Fam151b | chr13 | - | 92449624 | 92484015 | regular |
| ENSMUSG00000025340 | Rabgef1 | chr5 | + | 130171797 | 130214342 | regular |

| | | | | | | |
|---|---|---|---|---|---|---|
| ENSMUSG00000010342 | Tex14 | chr11 | + | 87405064 | 87555823 | regular |
| ENSMUSG00000026553 | Copa | chr1 | + | 172082528 | 172122330 | regular |
| ENSMUSG00000040565 | Btaf1 | chr19 | + | 36926078 | 37012752 | regular |
| ENSMUSG00000028949 | Smarcd3 | chr5 | - | 24590817 | 24624411 | regular |
| ENSMUSG00000020737 | Jpt1 | chr11 | - | 115497352 | 115514387 | regular |
| ENSMUSG00000017132 | Cyth1 | chr11 | - | 118132018 | 118248592 | regular |
| ENSMUSG00000028790 | Khdrbs1 | chr4 | - | 129703163 | 129742303 | regular |
| ENSMUSG00000042599 | Kdm7a | chr6 | - | 39136622 | 39206789 | regular |
| ENSMUSG00000041408 | Wapl | chr14 | + | 34673927 | 34747983 | regular |
| ENSMUSG00000034258 | Flvcr2 | chr12 | + | 85746538 | 85813585 | regular |
| ENSMUSG00000032294 | Pkm | chr9 | + | 59656367 | 59679375 | regular |
| ENSMUSG00000037933 | Bicd2 | chr13 | + | 49341584 | 49387026 | regular |
| ENSMUSG00000049670 | Morn4 | chr19 | - | 42074938 | 42086370 | regular |
| ENSMUSG00000062168 | Ppef1 | chrX | - | 160623093 | 160735765 | regular |
| ENSMUSG00000002455 | Prpf6 | chr2 | + | 181591867 | 181655660 | regular |
| ENSMUSG00000030662 | Ipo5 | chr14 | + | 120911223 | 120947999 | regular |
| ENSMUSG00000026790 | Odf2 | chr2 | + | 29889220 | 29931746 | regular |
| ENSMUSG00000037742 | Eef1a1 | chr9 | - | 78478448 | 78489151 | regular |
| ENSMUSG00000042594 | Sh2b3 | chr5 | - | 121815487 | 121837646 | regular |
| ENSMUSG00000090173 | Fbxw10 | chr11 | + | 62847068 | 62877465 | regular |
| ENSMUSG00000039110 | Mycbpap | chr11 | - | 94501346 | 94521742 | regular |
| ENSMUSG00000033257 | Ttll4 | chr1 | + | 74661744 | 74703730 | regular |
| ENSMUSG00000053754 | Chd8 | chr14 | - | 52198150 | 52257780 | regular |
| ENSMUSG00000041740 | Rnf10 | chr5 | - | 115241411 | 115272898 | regular |
| ENSMUSG00000026383 | Epb41l5 | chr1 | - | 119545036 | 119649000 | regular |
| ENSMUSG00000036574 | Tex44 | chr1 | + | 86426328 | 86428052 | regular |
| ENSMUSG00000022283 | Pabpc1 | chr15 | - | 36595660 | 36609668 | regular |
| ENSMUSG00000037627 | Rgs22 | chr15 | - | 36009478 | 36140400 | regular |
| ENSMUSG00000061411 | Nol4l | chr2 | - | 153407461 | 153529971 | regular |
| ENSMUST00000201614 | Med13l | chr5 | + | 118593332 | 118593570 | circular |
| ENSMUSG00000026018 | Ica1l | chr1 | - | 59982489 | 60043184 | regular |
| ENSMUSG00000046957 | Spz1 | chr13 | - | 92574635 | 92576172 | regular |
| ENSMUSG00000024622 | Hmgxb3 | chr18 | - | 61131278 | 61177050 | regular |
| ENSMUSG00000006435 | Neurl1a | chr19 | + | 47178819 | 47259440 | regular |
| ENSMUSG00000031865 | Dctn1 | chr6 | + | 83165919 | 83200117 | regular |
| ENSMUSG00000006932 | Ctnnb1 | chr9 | + | 120929215 | 120960507 | regular |
| ENSMUSG00000079173 | Zan | chr5 | - | 137378636 | 137477064 | regular |
| ENSMUSG00000032966 | Fkbp1a | chr2 | + | 151542482 | 151561692 | regular |
| ENSMUSG00000056201 | Cfl1 | chr19 | + | 5490454 | 5495201 | regular |
| ENSMUSG00000026781 | Acbd5 | chr2 | + | 23068166 | 23115558 | regular |
| ENSMUSG00000020936 | Nmt1 | chr11 | + | 103028189 | 103068912 | regular |
| ENSMUSG00000071659 | Hnrnpul2 | chr19 | + | 8819400 | 8834142 | regular |
| ENSMUSG00000028441 | 1110017D15 | chr4 | - | 41505008 | 41517333 | regular |
| ENSMUSG00000033819 | Ppp1r16a | chr15 | + | 76671614 | 76694919 | regular |
| ENSMUSG00000040782 | Rfwd2 | chr1 | + | 159232319 | 159347640 | regular |
| ENSMUSG00000041642 | Kif21b | chr1 | + | 136131388 | 136177998 | regular |

| | | | | | | |
|---|---|---|---|---|---|---|
| ENSMUSG00000044320 | 1700001O22 | chr2 | - | 30794768 | 30803661 | regular |
| ENSMUSG00000028580 | Pum1 | chr4 | + | 130663320 | 130781564 | regular |
| ENSMUSG00000005609 | Ctr9 | chr7 | + | 111028950 | 111056377 | regular |
| ENSMUSG00000020368 | Canx | chr11 | - | 50293960 | 50325673 | regular |
| ENSMUSG00000039842 | Mcph1 | chr8 | + | 18595130 | 18803189 | regular |
| ENSMUSG00000022558 | Mroh1 | chr15 | + | 76380260 | 76453038 | regular |
| ENSMUSG00000020472 | Zkscan17 | chr11 | - | 59485519 | 59526751 | regular |
| ENSMUSG00000033813 | Tcea1 | chr1 | + | 4857813 | 4897909 | regular |
| ENSMUSG00000020437 | Myo1g | chr11 | - | 6506547 | 6520965 | regular |
| ENSMUSG00000029147 | Ppm1g | chr5 | - | 31202663 | 31220687 | regular |
| ENSMUSG00000039275 | Foxk2 | chr11 | + | 121259989 | 121309896 | regular |
| ENSMUSG00000020412 | Ascc2 | chr11 | + | 4637746 | 4685699 | regular |
| ENSMUSG00000025939 | Ube2w | chr1 | - | 16540789 | 16619489 | regular |
| ENSMUSG00000060260 | Pwwp2b | chr7 | + | 139248481 | 139269903 | regular |
| ENSMUSG00000047193 | Dync2h1 | chr9 | - | 6928502 | 7184446 | regular |
| ENSMUSG00000004937 | Sgta | chr10 | - | 81044074 | 81060181 | regular |
| ENSMUSG00000052273 | Dnah3 | chr7 | - | 119922670 | 120095280 | regular |
| ENSMUSG00000023949 | Tcte1 | chr17 | + | 45523433 | 45542679 | regular |
| ENSMUSG00000031960 | Aars | chr8 | + | 111033143 | 111057664 | regular |
| ENSMUSG00000052221 | Ppp1r36 | chr12 | + | 76417537 | 76439492 | regular |
| ENSMUSG00000051276 | Actrt2 | chr4 | - | 154666432 | 154667867 | regular |
| ENSMUSG00000024404 | Riok3 | chr18 | + | 12128849 | 12157367 | regular |
| ENSMUSG00000048271 | Rbm33 | chr5 | + | 28317120 | 28419239 | regular |
| ENSMUSG00000072663 | Spef2 | chr15 | - | 9578192 | 9748868 | regular |
| ENSMUSG00000059970 | Hspa2 | chr12 | + | 76404175 | 76406938 | regular |
| ENSMUSG00000081984 | Dnajb3 | chr1 | - | 88204731 | 88205782 | regular |
| ENSMUSG00000086459 | 1700030C12I | chr11 | + | 23497752 | 23499661 | regular |
| ENSMUSG00000026933 | Camsap1 | chr2 | - | 25926837 | 25983282 | regular |
| ENSMUSG00000022770 | Dlg1 | chr16 | + | 31663442 | 31873356 | regular |
| ENSMUSG00000038637 | Lrrc56 | chr7 | + | 141194156 | 141210055 | regular |
| ENSMUSG00000024695 | Zfp91 | chr19 | - | 12763659 | 12796126 | regular |
| ENSMUSG00000063550 | Nup98 | chr7 | - | 102119397 | 102210176 | regular |
| ENSMUSG00000024223 | Armc12 | chr17 | + | 28530859 | 28538954 | regular |
| ENSMUSG00000069170 | Adgrv1 | chr13 | - | 81095067 | 81633154 | regular |
| ENSMUSG00000005566 | Trim28 | chr7 | + | 12999113 | 13031035 | regular |
| ENSMUSG00000028555 | Ttc39a | chr4 | + | 109406622 | 109444745 | regular |
| ENSMUSG00000048410 | Zfp407 | chr18 | - | 84128026 | 84589725 | regular |
| ENSMUSG00000029772 | Ahcyl2 | chr6 | + | 29768010 | 29912310 | regular |
| ENSMUSG00000024269 | Tpgs2 | chr18 | - | 25127222 | 25169007 | regular |
| ENSMUSG00000002996 | Hbp1 | chr12 | - | 31926253 | 31950535 | regular |
| ENSMUSG00000029130 | Rnf32 | chr5 | + | 29195991 | 29228457 | regular |
| ENSMUSG00000033658 | Ddx19b | chr8 | - | 111003187 | 111031751 | regular |
| ENSMUSG00000110622 | Gm16486 | chr8 | + | 70703456 | 70717650 | regular |
| ENSMUSG00000069919 | Hba-a1 | chr11 | + | 32283510 | 32284465 | regular |
| ENSMUSG00000027475 | Kif3b | chr2 | + | 153291412 | 153333390 | regular |
| ENSMUSG00000030403 | Vasp | chr7 | - | 19256928 | 19271817 | regular |

| | | | | | | |
|---|---|---|---|---|---|---|
| ENSMUSG00000031847 | 1700030J22F | chr8 | - | 116969593 | 116978959 | regular |
| ENSMUSG00000041945 | Mfsd9 | chr1 | - | 40772037 | 40790687 | regular |
| ENSMUSG00000029614 | Rpl6 | chr5 | + | 121204480 | 121209241 | regular |
| ENSMUSG00000078611 | Gm5901 | chr7 | + | 105375037 | 105378377 | regular |
| ENSMUSG00000032580 | Rbm5 | chr9 | - | 107740370 | 107770996 | regular |
| ENSMUSG00000022501 | Prm1 | chr16 | - | 10796325 | 10796886 | regular |
| ENSMUSG00000029128 | Rab28 | chr5 | - | 41624975 | 41708157 | regular |
| ENSMUSG00000019952 | Poc1b | chr10 | + | 99107035 | 99198074 | regular |
| ENSMUSG00000030591 | Psmd8 | chr7 | - | 29174187 | 29180701 | regular |
| ENSMUSG00000005237 | Dnah2 | chr11 | - | 69420808 | 69549110 | regular |
| ENSMUSG00000051306 | Usp42 | chr5 | - | 143710324 | 143732280 | regular |
| ENSMUSG00000028426 | Rad23b | chr4 | + | 55350042 | 55392237 | regular |
| ENSMUSG00000028572 | Hook1 | chr4 | + | 95967239 | 96025413 | regular |
| ENSMUSG00000019876 | Pkib | chr10 | + | 57631980 | 57741112 | regular |
| ENSMUSG00000057069 | Ero1lb | chr13 | + | 12565873 | 12611396 | regular |
| ENSMUSG00000042331 | Specc1 | chr11 | + | 61956762 | 62223013 | regular |
| ENSMUSG00000050721 | Plekho2 | chr9 | - | 65552697 | 65580040 | regular |
| ENSMUSG00000041702 | Btbd7 | chr12 | - | 102780796 | 102878471 | regular |
| ENSMUSG00000041040 | Fam117b | chr1 | + | 59913005 | 59985346 | regular |
| ENSMUSG00000012848 | Rps5 | chr7 | + | 12922289 | 12926686 | regular |
| ENSMUSG00000020716 | Nf1 | chr11 | + | 79339692 | 79581612 | regular |
| ENSMUSG00000043241 | Upf2 | chr2 | + | 5951468 | 6056703 | regular |
| ENSMUSG00000068735 | Trp53i11 | chr2 | + | 93187547 | 93201759 | regular |
| ENSMUSG00000035297 | Cops4 | chr5 | + | 100518308 | 100547803 | regular |
| ENSMUSG00000031818 | Cox4i1 | chr8 | + | 120668221 | 120674207 | regular |
| ENSMUSG00000056536 | Pign | chr1 | - | 105518421 | 105663677 | regular |
| ENSMUSG00000006024 | Napa | chr7 | + | 16098457 | 16117975 | regular |
| ENSMUSG00000046364 | Rpl27a | chr7 | + | 109519146 | 109522367 | regular |
| ENSMUSG00000035517 | Tdrd7 | chr4 | + | 45965333 | 46034761 | regular |
| ENSMUSG00000048154 | Kmt2d | chr15 | - | 98831668 | 98871204 | regular |
| ENSMUSG00000039621 | Prex1 | chr2 | - | 166566341 | 166713832 | regular |
| ENSMUSG00000028106 | Rprd2 | chr3 | - | 95760340 | 95818863 | regular |
| ENSMUSG00000020481 | Ankrd36 | chr11 | + | 5569683 | 5689337 | regular |
| ENSMUSG00000031422 | Morf4l2 | chrX | - | 136732941 | 136743690 | regular |
| ENSMUSG00000029833 | Trim24 | chr6 | + | 37870810 | 37966296 | regular |
| ENSMUSG00000071636 | Rimbp3 | chr16 | + | 17208134 | 17213921 | regular |
| ENSMUSG00000028639 | Ybx1 | chr4 | - | 119277980 | 119294604 | regular |
| ENSMUSG00000029020 | Mfn2 | chr4 | - | 147873598 | 147904704 | regular |
| ENSMUSG00000026155 | Smap1 | chr1 | - | 23844845 | 23922317 | regular |
| ENSMUSG00000040929 | Rfx3 | chr19 | - | 27761720 | 28011166 | regular |
| ENSMUSG00000025241 | Fyco1 | chr9 | - | 123789499 | 123851899 | regular |
| ENSMUSG00000039000 | Ube3c | chr5 | + | 29569241 | 29676092 | regular |
| ENSMUSG00000021203 | Otub2 | chr12 | + | 103388681 | 103406350 | regular |
| ENSMUSG00000021374 | Nup153 | chr13 | - | 46679904 | 46727940 | regular |
| ENSMUSG00000041112 | Elmo1 | chr13 | + | 20090595 | 20608353 | regular |
| ENSMUSG00000005886 | Ncoa2 | chr1 | - | 13139104 | 13374083 | regular |

| | | | | | | |
|---|---|---|---|---|---|---|
| ENSMUSG00000030086 | Chchd6 | chr6 | - | 89383145 | 89595652 | regular |
| ENSMUSG00000036438 | Calm2 | chr17 | - | 87433411 | 87446935 | regular |
| ENSMUSG00000071657 | Bscl2 | chr19 | + | 8837466 | 8848683 | regular |
| ENSMUSG00000031731 | Ap1g1 | chr8 | + | 109778553 | 109864204 | regular |
| ENSMUSG00000061136 | Prpf40a | chr2 | - | 53134703 | 53191284 | regular |
| ENSMUSG00000022056 | Adam7 | chr14 | - | 68497335 | 68533741 | regular |
| ENSMUSG00000018476 | Kdm6b | chr11 | - | 69398507 | 69413675 | regular |
| ENSMUSG00000024855 | Pacs1 | chr19 | - | 5133687 | 5273119 | regular |
| ENSMUSG00000033502 | Cdc14a | chr3 | - | 116272552 | 116424032 | regular |
| ENSMUSG00000024566 | Atp9b | chr18 | - | 80734140 | 80934058 | regular |
| ENSMUSG00000034731 | Dgkh | chr14 | - | 78558749 | 78732776 | regular |
| ENSMUSG00000049470 | Aff4 | chr11 | + | 53350832 | 53421830 | regular |
| ENSMUSG00000022897 | Dyrk1a | chr16 | + | 94570009 | 94695517 | regular |
| ENSMUSG00000020361 | Hspa4 | chr11 | - | 53259813 | 53300457 | regular |
| ENSMUSG00000031540 | Kat6a | chr8 | + | 22859534 | 22943259 | regular |
| ENSMUSG00000033732 | Sf3b3 | chr8 | - | 110810238 | 110846787 | regular |
| ENSMUSG00000033124 | Atg9a | chr1 | - | 75180859 | 75192196 | regular |
| ENSMUSG00000027479 | Mapre1 | chr2 | + | 153741273 | 153773310 | regular |
| ENSMUSG00000059119 | Nap1l4 | chr7 | - | 143513578 | 143549106 | regular |
| ENSMUSG00000071064 | Zfp827 | chr8 | + | 79028436 | 79193766 | regular |
| ENSMUSG00000062997 | Rpl35 | chr2 | - | 39001579 | 39005624 | regular |
| ENSMUSG00000033760 | Rbm4b | chr19 | + | 4756542 | 4767065 | regular |
| ENSMUSG00000028759 | Hp1bp3 | chr4 | + | 138216295 | 138244683 | regular |
| ENSMUSG00000015837 | Sqstm1 | chr11 | - | 50199365 | 50210827 | regular |
| ENSMUSG00000046727 | Cystm1 | chr18 | + | 36345618 | 36402980 | regular |
| ENSMUSG00000037716 | Ccdc33 | chr9 | - | 58028676 | 58118823 | regular |
| ENSMUSG00000063524 | Eno1 | chr4 | + | 150236720 | 150248879 | regular |
| ENSMUSG00000019951 | Uhrf1bp1l | chr10 | + | 89744990 | 89819871 | regular |
| ENSMUSG00000052955 | Cpvl | chr6 | - | 53873278 | 53978671 | regular |
| ENSMUSG00000030432 | Rpl28 | chr7 | + | 4792873 | 4795066 | regular |
| ENSMUSG00000063820 | Arl9 | chr5 | + | 77004054 | 77010606 | regular |
| ENSMUSG00000026931 | 1700019N19 | chr19 | - | 58785802 | 58794461 | regular |
| ENSMUSG00000039007 | Cpq | chr15 | + | 33083128 | 33594552 | regular |
| ENSMUSG00000050612 | Txndc2 | chr17 | - | 65637504 | 65642204 | regular |
| ENSMUSG00000039262 | Prrc2b | chr2 | + | 32151081 | 32234537 | regular |
| ENSMUSG00000029155 | Spata18 | chr5 | + | 73651378 | 73679512 | regular |
| ENSMUSG00000032514 | Ttc21a | chr9 | + | 119937605 | 119967793 | regular |
| ENSMUSG00000031027 | Stk33 | chr7 | - | 109279222 | 109439081 | regular |
| ENSMUSG00000068917 | Clk2 | chr3 | + | 89164794 | 89176921 | regular |
| ENSMUSG00000040940 | Arhgef1 | chr7 | + | 24902911 | 24926594 | regular |
| ENSMUSG00000109336 | Samd4b | chr7 | - | 28399521 | 28598144 | regular |
| ENSMUSG00000030275 | Etnk1 | chr6 | + | 143167220 | 143232307 | regular |
| ENSMUSG00000032570 | Atp2c1 | chr9 | - | 105403538 | 105527319 | regular |
| ENSMUSG00000029073 | Cptp | chr4 | - | 155864722 | 155869440 | regular |
| ENSMUSG00000049353 | Rd3 | chr1 | + | 191977369 | 191988283 | regular |
| ENSMUSG00000032560 | Dnajc13 | chr9 | - | 104151281 | 104262930 | regular |

| | | | | | | |
|---|---|---|---|---|---|---|
| ENSMUSG00000025314 | Ptprj | chr2 | - | 90429753 | 90580647 | regular |
| ENSMUSG00000053907 | Mat2a | chr6 | - | 72432798 | 72439558 | regular |
| ENSMUSG00000092518 | Fam71e2 | chr7 | - | 4753225 | 4771302 | regular |
| ENSMUSG00000004347 | Pde1c | chr6 | - | 56069803 | 56569103 | regular |
| ENSMUSG00000002395 | Use1 | chr8 | + | 71366847 | 71369732 | regular |
| ENSMUSG00000043154 | Ppp2r3a | chr9 | - | 101105083 | 101251795 | regular |
| ENSMUSG00000033793 | Atp6v1h | chr1 | + | 5070017 | 5162529 | regular |
| ENSMUSG00000034751 | Mast4 | chr13 | - | 102732485 | 103334497 | regular |
| ENSMUSG00000030374 | Strn4 | chr7 | + | 16815888 | 16840931 | regular |
| ENSMUSG00000038564 | Ift172 | chr5 | - | 31253276 | 31291116 | regular |
| ENSMUSG00000006418 | Rnf114 | chr2 | + | 167492644 | 167516173 | regular |
| ENSMUSG00000026853 | Crat | chr2 | - | 30400470 | 30415813 | regular |
| ENSMUSG00000034354 | Mtmr3 | chr11 | - | 4480867 | 4594863 | regular |
| ENSMUSG00000042323 | Pbrm1 | chr14 | + | 31019137 | 31121592 | regular |
| ENSMUSG00000021476 | Habp4 | chr13 | + | 64161823 | 64186537 | regular |
| ENSMUSG00000042608 | Stk40 | chr4 | + | 126103956 | 126141029 | regular |
| ENSMUSG00000037885 | Stk35 | chr2 | + | 129800516 | 129832287 | regular |
| ENSMUSG00000038538 | Ubn2 | chr6 | + | 38433949 | 38524825 | regular |
| ENSMUSG00000032412 | Atp1b3 | chr9 | - | 96332654 | 96364442 | regular |
| ENSMUSG00000063889 | Crem | chr18 | - | 3266047 | 3337748 | regular |
| ENSMUSG00000025081 | Tdrd1 | chr19 | + | 56826208 | 56870012 | regular |
| ENSMUSG00000041763 | Tpp2 | chr1 | + | 43933646 | 44003000 | regular |
| ENSMUSG00000099353 | 1700031M16 | chr15 | + | 98398418 | 98416268 | regular |
| ENSMUSG00000074817 | Papolb | chr5 | - | 142525837 | 142530076 | regular |
| ENSMUSG00000026219 | Trip12 | chr1 | - | 84721188 | 84840516 | regular |
| ENSMUSG00000039254 | Pomt1 | chr2 | + | 32236589 | 32255005 | regular |
| ENSMUSG00000033943 | Mga | chr2 | + | 119897227 | 119969581 | regular |
| ENSMUSG00000025234 | Arih1 | chr9 | - | 59388257 | 59486618 | regular |
| ENSMUSG00000018554 | Ybx2 | chr11 | + | 69935795 | 69941605 | regular |
| ENSMUSG00000000346 | Dazap2 | chr15 | + | 100615627 | 100620731 | regular |
| ENSMUSG00000040549 | Ckap5 | chr2 | + | 91526761 | 91620664 | regular |
| ENSMUSG00000071252 | 2210408I21R | chr13 | + | 77135539 | 77613784 | regular |
| ENSMUSG00000029206 | Nsun7 | chr5 | + | 66259896 | 66298026 | regular |
| ENSMUSG00000026150 | Mff | chr1 | + | 82724889 | 82752394 | regular |
| ENSMUSG00000020255 | D10Wsu102e | chr10 | + | 83360220 | 83488505 | regular |
| ENSMUSG00000068663 | Clec16a | chr16 | + | 10545338 | 10744878 | regular |
| ENSMUSG00000035247 | Hectd1 | chr12 | - | 51743721 | 51829536 | regular |
| ENSMUSG00000028495 | Rps6 | chr4 | - | 86854659 | 86857412 | regular |
| ENSMUSG00000002055 | Spag5 | chr11 | + | 78301528 | 78322457 | regular |
| ENSMUSG00000112148 | Lilrb4a | chr10 | + | 51490955 | 51496611 | regular |
| ENSMUSG00000032826 | Ank2 | chr3 | - | 126921611 | 127499350 | regular |
| ENSMUSG00000024042 | Sik1 | chr17 | - | 31844247 | 31855792 | regular |
| ENSMUSG00000049008 | BB014433 | chr8 | - | 15041443 | 15046078 | regular |
| ENSMUSG00000036257 | Pnpla8 | chr12 | + | 44221369 | 44322532 | regular |
| ENSMUSG00000030994 | D7Ertd443e | chr7 | - | 134265778 | 134385661 | regular |
| ENSMUSG00000051223 | Bzw1 | chr1 | + | 58392897 | 58407353 | regular |

| | | | | | | |
|---|---|---|---|---|---|---|
| ENSMUSG00000057335 | Cep170 | chr1 | - | 176733652 | 176814067 | regular |
| ENSMUSG00000037432 | Fer1l5 | chr1 | + | 36372290 | 36422110 | regular |
| ENSMUSG00000030161 | Gabarapl1 | chr6 | + | 129533159 | 129542346 | regular |
| ENSMUSG00000027528 | Fabp9 | chr3 | - | 10193620 | 10197283 | regular |
| ENSMUSG00000014294 | Ndufa2 | chr18 | - | 36742331 | 36744557 | regular |
| ENSMUSG00000036054 | Sugp2 | chr8 | + | 70234225 | 70279915 | regular |
| ENSMUSG00000060002 | Chpt1 | chr10 | - | 88452744 | 88504073 | regular |
| ENSMUSG00000030498 | Gas2 | chr7 | + | 51862014 | 51994975 | regular |
| ENSMUSG00000035032 | Nek11 | chr9 | - | 105162155 | 105395524 | regular |
| ENSMUSG00000022441 | Efcab6 | chr15 | - | 83866711 | 84065379 | regular |
| ENSMUSG00000025532 | Crcp | chr5 | + | 130029289 | 130060789 | regular |
| ENSMUSG00000040699 | Limd2 | chr11 | - | 106156255 | 106160860 | regular |
| ENSMUSG00000087279 | 4930544D05I | chr11 | + | 70615847 | 70616890 | regular |
| ENSMUSG00000034377 | Tulp4 | chr17 | + | 6106436 | 6251128 | regular |
| ENSMUSG00000021420 | Fars2 | chr13 | + | 36117411 | 36726280 | regular |
| ENSMUSG00000000365 | Rnf17 | chr14 | + | 56402580 | 56525032 | regular |
| ENSMUSG00000042699 | Dhx9 | chr1 | - | 153455757 | 153487660 | regular |
| ENSMUSG00000030126 | Tmcc1 | chr6 | - | 116018610 | 116193486 | regular |
| ENSMUSG00000076436 | Oxct2a | chr4 | - | 123321874 | 123323634 | regular |
| ENSMUSG00000040731 | Eif4h | chr5 | - | 134619720 | 134639490 | regular |
| ENSMUSG00000059263 | Usp47 | chr7 | + | 112023503 | 112111661 | regular |
| ENSMUSG00000005483 | Dnajb1 | chr8 | + | 83608192 | 83612653 | regular |
| ENSMUSG00000026380 | Tfcp2l1 | chr1 | + | 118627944 | 118685168 | regular |
| ENSMUSG00000035435 | Abca17 | chr17 | - | 24264258 | 24351029 | regular |
| ENSMUSG00000024608 | Rps14 | chr18 | + | 60774509 | 60778546 | regular |
| ENSMUSG00000016559 | H3f3b | chr11 | - | 116021911 | 116027962 | regular |
| ENSMUSG00000050944 | Efcab5 | chr11 | - | 77089914 | 77188968 | regular |
| ENSMUSG00000017837 | Nkiras2 | chr11 | + | 100619243 | 100627607 | regular |
| ENSMUSG00000033364 | Usp37 | chr1 | - | 74435510 | 74544284 | regular |
| ENSMUSG00000042213 | Zfand4 | chr6 | + | 116264221 | 116330302 | regular |
| ENSMUSG00000036698 | Ago2 | chr15 | - | 73095843 | 73184840 | regular |
| ENSMUSG00000029030 | Tprgl | chr4 | - | 154157484 | 154160666 | regular |
| ENSMUSG00000028081 | Rps3a1 | chr3 | - | 86137939 | 86142702 | regular |
| ENSMUSG00000036833 | Pnpla7 | chr2 | + | 24976032 | 25054057 | regular |
| ENSMUSG00000021196 | Pfkp | chr13 | - | 6579767 | 6648777 | regular |
| ENSMUSG00000025047 | Pdcd11 | chr19 | + | 47090765 | 47131143 | regular |
| ENSMUSG00000027799 | Nbea | chr3 | - | 55625194 | 56183701 | regular |
| ENSMUSG00000037266 | Rsrp1 | chr4 | + | 134923591 | 134927671 | regular |
| ENSMUSG00000020235 | Fzr1 | chr10 | - | 81366221 | 81378516 | regular |
| ENSMUSG00000037622 | Wdtc1 | chr4 | - | 133292458 | 133353481 | regular |
| ENSMUSG00000074679 | Defb28 | chr2 | + | 152518254 | 152521447 | regular |
| ENSMUSG00000035919 | Bbs9 | chr9 | + | 22475714 | 22888280 | regular |
| ENSMUSG00000036309 | Skp1a | chr11 | + | 52231994 | 52246858 | regular |
| ENSMUSG00000017421 | Zfp207 | chr11 | + | 80383278 | 80405733 | regular |
| ENSMUSG00000064330 | Pde6h | chr6 | + | 136923831 | 136968865 | regular |
| ENSMUSG00000035504 | Reep6 | chr10 | + | 80329952 | 80336441 | regular |

| | | | | | | |
|---|---|---|---|---|---|---|
| ENSMUSG00000024948 | Map4k2 | chr19 | + | 6341134 | 6355615 | regular |
| ENSMUSG00000037971 | 1110032A03I | chr9 | - | 50762827 | 50775520 | regular |
| ENSMUSG00000038759 | Nup205 | chr6 | + | 35177420 | 35247596 | regular |
| ENSMUSG00000018412 | Kansl1 | chr11 | - | 104333228 | 104468861 | regular |
| ENSMUSG00000030816 | Rnf40 | chr7 | + | 127588766 | 127604799 | regular |
| ENSMUSG00000018924 | Alox15 | chr11 | - | 70344151 | 70352031 | regular |
| ENSMUSG00000033701 | Acbd6 | chr1 | + | 155558119 | 155691330 | regular |
| ENSMUSG00000024317 | Rnf138 | chr18 | + | 21001340 | 21028223 | regular |
| ENSMUSG00000019726 | Lyst | chr13 | + | 13590396 | 13778803 | regular |
| ENSMUSG00000006575 | Rundc3a | chr11 | + | 102393402 | 102402555 | regular |
| ENSMUSG00000100169 | Gm29087 | chr7 | + | 38500431 | 38537077 | regular |
| ENSMUSG00000090369 | 4933411K16I | chr19 | + | 42052247 | 42053627 | regular |
| ENSMUSG00000078812 | Eif5a | chr11 | - | 69916713 | 69921958 | regular |
| ENSMUSG00000072295 | C2cd6 | chr1 | - | 59050505 | 59094900 | regular |
| ENSMUSG00000028337 | Coro2a | chr4 | - | 46536936 | 46602202 | regular |
| ENSMUSG00000028310 | Ppp3r2 | chr4 | - | 49678746 | 49682024 | regular |
| ENSMUSG00000031849 | Comp | chr8 | + | 70373557 | 70382066 | regular |
| ENSMUSG00000092805 | Gm26461 | chr9 | + | 88553235 | 88553525 | regular |
| ENSMUSG00000020048 | Hsp90b1 | chr10 | - | 86690208 | 86705509 | regular |
| ENSMUSG00000052861 | Dnah6 | chr6 | - | 73017605 | 73221651 | regular |
| ENSMUSG00000064354 | mt-Co2 | chrM | + | 7012 | 7696 | regular |
| ENSMUSG00000027782 | Kpna4 | chr3 | - | 69067148 | 69127113 | regular |
| ENSMUSG00000024206 | Rfx2 | chr17 | - | 56775896 | 56831008 | regular |
| ENSMUSG00000006356 | Crip2 | chr12 | + | 113140235 | 113145506 | regular |
| ENSMUSG00000030465 | Psd3 | chr8 | - | 67689081 | 68212027 | regular |
| ENSMUSG00000079557 | Mar.02 | chr17 | - | 33685691 | 33718670 | regular |
| ENSMUSG00000048668 | Rhno1 | chr6 | - | 128356999 | 128362911 | regular |
| ENSMUSG00000074247 | Dda1 | chr8 | + | 71469198 | 71476098 | regular |
| ENSMUSG00000010277 | 2610507B11I | chr11 | + | 78261751 | 78290623 | regular |
| ENSMUSG00000070733 | Fryl | chr5 | - | 73019986 | 73256619 | regular |
| ENSMUSG00000000811 | Txnrd3 | chr6 | + | 89643987 | 89675529 | regular |
| ENSMUSG00000021877 | Arf4 | chr14 | + | 26638073 | 26665084 | regular |
| ENSMUSG00000058881 | Zfp516 | chr18 | + | 82910878 | 83005314 | regular |
| ENSMUSG00000034994 | Eef2 | chr10 | + | 81176630 | 81182498 | regular |
| ENSMUSG00000058997 | Vwa8 | chr14 | + | 78849051 | 79202310 | regular |
| ENSMUSG00000032437 | Stt3b | chr9 | - | 115242580 | 115310421 | regular |
| ENSMUSG00000030301 | Ccdc91 | chr6 | + | 147475827 | 147632612 | regular |
| ENSMUSG00000016619 | Nup50 | chr15 | + | 84923427 | 84942955 | regular |
| ENSMUSG00000052056 | Zfp217 | chr2 | - | 170108642 | 170148103 | regular |
| ENSMUSG00000064225 | Paqr9 | chr9 | + | 95559656 | 95568023 | regular |
| ENSMUSG00000028138 | Adh5 | chr3 | + | 138443092 | 138455499 | regular |
| ENSMUSG00000041264 | Uspl1 | chr5 | + | 149184349 | 149215434 | regular |
| ENSMUSG00000064181 | Rab3ip | chr10 | - | 116905783 | 116950769 | regular |
| ENSMUSG00000025026 | Add3 | chr19 | + | 53140444 | 53247399 | regular |
| ENSMUSG00000029922 | Mkrn1 | chr6 | - | 39397803 | 39420462 | regular |
| ENSMUSG00000100153 | Gm5601 | chr7 | + | 119739925 | 119742226 | regular |

| | | | | | | |
|---|---|---|---|---|---|---|
| ENSMUSG00000075520 | Malrd1 | chr2 | + | 15526478 | 16255555 | regular |
| ENSMUSG00000040785 | Ttc3 | chr16 | + | 94370617 | 94522117 | regular |
| ENSMUSG00000028234 | Rps20 | chr4 | - | 3831333 | 3835665 | regular |
| ENSMUSG00000020056 | Washc3 | chr10 | + | 88201092 | 88246158 | regular |
| ENSMUSG00000033983 | Coil | chr11 | + | 88970251 | 88991613 | regular |
| ENSMUSG00000026922 | Agpat2 | chr2 | - | 26593056 | 26604417 | regular |
| ENSMUSG00000027006 | Dnajc10 | chr2 | + | 80315465 | 80354043 | regular |
| ENSMUSG00000021998 | Lcp1 | chr14 | + | 75131100 | 75230842 | regular |
| ENSMUSG00000028530 | Jak1 | chr4 | - | 101152366 | 101265282 | regular |
| ENSMUSG00000036180 | Gatad2a | chr8 | - | 69907075 | 69996384 | regular |
| ENSMUSG00000024397 | Aif1 | chr17 | - | 35170990 | 35176068 | regular |
| ENSMUSG00000039831 | Arhgap29 | chr3 | + | 121952540 | 122016753 | regular |
| ENSMUSG00000037486 | Asxl2 | chr12 | + | 3426856 | 3506852 | regular |
| ENSMUSG00000042632 | Pla2g6 | chr15 | - | 79286227 | 79328390 | regular |
| ENSMUSG00000070530 | Wfdc16 | chr2 | - | 164634706 | 164638796 | regular |
| ENSMUSG00000024973 | Hrasls5 | chr19 | + | 7612540 | 7639642 | regular |
| ENSMUSG00000067028 | Cntnap5b | chr1 | + | 99772764 | 100485942 | regular |
| ENSMUSG00000020385 | Clk4 | chr11 | + | 51261729 | 51281766 | regular |
| ENSMUSG00000027550 | Lrrcc1 | chr3 | + | 14533787 | 14572658 | regular |
| ENSMUSG00000022471 | Xrcc6 | chr15 | + | 81987834 | 82040085 | regular |
| ENSMUSG00000045350 | Fam186a | chr15 | - | 99918347 | 99967075 | regular |
| ENSMUSG00000036594 | H2-Aa | chr17 | - | 34282743 | 34287827 | regular |
| ENSMUSG00000076432 | Ywhaq | chr12 | - | 21390070 | 21417637 | regular |
| ENSMUSG00000036964 | Trim17 | chr11 | + | 58954684 | 58973098 | regular |
| ENSMUSG00000030851 | Ldhc | chr7 | + | 46861202 | 46878142 | regular |
| ENSMUSG00000026575 | Nme7 | chr1 | + | 164304120 | 164437725 | regular |
| ENSMUSG00000024154 | Gtf2a1l | chr17 | + | 88668659 | 88715152 | regular |
| ENSMUSG00000026000 | Lancl1 | chr1 | - | 67000516 | 67038872 | regular |
| ENSMUSG00000029148 | Nrbp1 | chr5 | + | 31240863 | 31251566 | regular |
| ENSMUSG00000031447 | Lamp1 | chr8 | + | 13159160 | 13175338 | regular |
| ENSMUSG00000003033 | Ap1m1 | chr8 | + | 72240017 | 72257385 | regular |
| ENSMUSG00000040506 | Ambra1 | chr2 | + | 91730133 | 91918849 | regular |
| ENSMUSG00000070319 | Eif3g | chr9 | - | 20894348 | 20898623 | regular |
| ENSMUSG00000049971 | Glt1d1 | chr5 | + | 127632261 | 127709374 | regular |
| ENSMUSG00000025808 | Ccdc7a | chr8 | - | 128734234 | 129065517 | regular |
| ENSMUSG00000021572 | Cep72 | chr13 | - | 74036499 | 74062299 | regular |
| ENSMUSG00000021171 | Esyt2 | chr12 | + | 116281195 | 116391050 | regular |
| ENSMUSG00000019986 | Ahi1 | chr10 | + | 20952546 | 21080429 | regular |
| ENSMUSG00000044176 | Spink10 | chr18 | + | 62548910 | 62661366 | regular |
| ENSMUSG00000050058 | Prm3 | chr16 | - | 10790504 | 10790914 | regular |
| ENSMUSG00000049811 | Fam161a | chr11 | + | 23007530 | 23030788 | regular |
| ENSMUSG00000022905 | Kpna1 | chr16 | + | 35978749 | 36037131 | regular |
| ENSMUSG00000021687 | Scamp1 | chr13 | - | 94201309 | 94285857 | regular |
| ENSMUSG00000007815 | Rhoa | chr9 | + | 108306128 | 108337934 | regular |
| ENSMUSG00000053333 | Dis3l2 | chr1 | + | 86703807 | 87050095 | regular |
| ENSMUSG00000043909 | Trp53bp1 | chr2 | - | 121193280 | 121271407 | regular |

| | | | | | | |
|---|---|---|---|---|---|---|
| ENSMUSG00000026778 | Prkcq | chr2 | + | 11172107 | 11301222 | regular |
| ENSMUSG00000026457 | Adipor1 | chr1 | + | 134415377 | 134433351 | regular |
| ENSMUSG00000028347 | Tmeff1 | chr4 | + | 48585173 | 48663131 | regular |
| ENSMUSG00000032743 | D430042O09 | chr7 | + | 125707887 | 125874793 | regular |
| ENSMUSG00000025907 | Rb1cc1 | chr1 | + | 6206196 | 6276648 | regular |
| ENSMUSG00000032740 | Ccdc88a | chr11 | + | 29373657 | 29510808 | regular |
| ENSMUSG00000050035 | Fhl4 | chr10 | - | 85096286 | 85102495 | regular |
| ENSMUSG00000070738 | Dgkd | chr1 | + | 87853286 | 87945180 | regular |
| ENSMUSG00000007950 | Abhd8 | chr8 | - | 71456704 | 71463655 | regular |
| ENSMUSG00000029769 | Ccdc136 | chr6 | + | 29396308 | 29426994 | regular |
| ENSMUSG00000060938 | Rpl26 | chr11 | + | 68901582 | 68906989 | regular |
| ENSMUSG00000030688 | Stard10 | chr7 | + | 101317085 | 101346626 | regular |
| ENSMUSG00000059040 | Eno1b | chr18 | + | 48045334 | 48048378 | regular |
| ENSMUSG00000044475 | Ascc1 | chr10 | + | 60002804 | 60099988 | regular |
| ENSMUSG00000050565 | Tor1aip2 | chr1 | + | 156035402 | 156068861 | regular |
| ENSMUSG00000032253 | Phip | chr9 | - | 82866158 | 82975516 | regular |
| ENSMUSG00000044468 | Fam46c | chr3 | - | 100451627 | 100489324 | regular |
| ENSMUSG00000074771 | Ankef1 | chr2 | + | 136501909 | 136562091 | regular |
| ENSMUSG00000030695 | Aldoa | chr7 | - | 126795233 | 126800751 | regular |
| ENSMUSG00000035152 | Ap2b1 | chr11 | + | 83299023 | 83405035 | regular |
| ENSMUSG00000001211 | Agpat3 | chr10 | - | 78269177 | 78352489 | regular |
| ENSMUSG00000001089 | Luzp1 | chr4 | + | 136469760 | 136554780 | regular |
| ENSMUSG00000023456 | Tpi1 | chr6 | - | 124810585 | 124814296 | regular |
| ENSMUSG00000040325 | Dcaf1 | chr9 | + | 106821873 | 106880992 | regular |
| ENSMUSG00000019892 | Lrriq1 | chr10 | - | 103046030 | 103236322 | regular |
| ENSMUSG00000052837 | Junb | chr8 | - | 84974483 | 84978718 | regular |
| ENSMUSG00000056999 | Ide | chr19 | - | 37268742 | 37337852 | regular |
| ENSMUSG00000029472 | Anapc5 | chr5 | - | 122787458 | 122821339 | regular |
| ENSMUSG00000025139 | Tollip | chr7 | - | 141874812 | 141918507 | regular |
| ENSMUSG00000056018 | Ccdc7b | chr8 | + | 129067133 | 129183732 | regular |
| ENSMUSG00000021266 | Wars | chr12 | - | 108860029 | 108894174 | regular |
| ENSMUSG00000055013 | Agap1 | chr1 | + | 89454805 | 89897617 | regular |
| ENSMUSG00000068823 | Csde1 | chr3 | + | 103020425 | 103058186 | regular |
| ENSMUSG00000070529 | Wfdc10 | chr2 | + | 164656045 | 164657368 | regular |
| ENSMUSG00000086898 | Itpr3os | chr17 | - | 27099657 | 27101440 | regular |
| ENSMUSG00000023495 | Pcbp4 | chr9 | + | 106453290 | 106464012 | regular |
| ENSMUSG00000066036 | Ubr4 | chr4 | + | 139352608 | 139489588 | regular |
| ENSMUSG00000048039 | Isg20l2 | chr3 | + | 87930313 | 87940686 | regular |
| ENSMUSG00000030555 | Ttc23 | chr7 | + | 67647409 | 67762912 | regular |
| ENSMUSG00000043060 | Fscb | chr12 | - | 64471329 | 64474690 | regular |
| ENSMUSG00000029640 | Usp12 | chr5 | - | 146734808 | 146795006 | regular |
| ENSMUSG00000001855 | Nup214 | chr2 | + | 31974435 | 32053975 | regular |
| ENSMUSG00000022664 | Slc35a5 | chr16 | - | 45139572 | 45158706 | regular |
| ENSMUSG00000037503 | Fam168b | chr1 | - | 34813218 | 34843065 | regular |
| ENSMUSG00000032802 | Srxn1 | chr2 | + | 152105515 | 152111376 | regular |
| ENSMUSG00000024369 | Nelfe | chr17 | + | 34850390 | 34856372 | regular |

| | | | | | | |
|---|---|---|---|---|---|---|
| ENSMUSG00000112785 | AC163296.2 | chr12 | + | 58813882 | 58815197 | regular |
| ENSMUSG00000042628 | Zfyve1 | chr12 | - | 83546557 | 83597222 | regular |
| ENSMUSG00000072723 | Gm10044 | chr14 | - | 7744062 | 7791651 | regular |
| ENSMUSG00000030451 | Herc2 | chr7 | + | 56050195 | 56231800 | regular |
| ENSMUSG00000049916 | 2610318N02 | chr16 | - | 17113398 | 17125140 | regular |
| ENSMUSG00000026254 | Eif4e2 | chr1 | + | 87213913 | 87240488 | regular |
| ENSMUSG00000036087 | Slain2 | chr5 | + | 72914303 | 72978829 | regular |
| ENSMUSG00000049339 | Retreg2 | chr1 | + | 75142777 | 75147913 | regular |
| ENSMUSG00000070327 | Rnf213 | chr11 | + | 119393099 | 119487418 | regular |
| ENSMUSG00000005621 | Zfp592 | chr7 | + | 80993680 | 81045164 | regular |
| ENSMUSG00000047139 | Cd24a | chr10 | + | 43578283 | 43584265 | regular |
| ENSMUSG00000020929 | Eftud2 | chr11 | - | 102838472 | 102880985 | regular |
| ENSMUSG00000035211 | Xrra1 | chr7 | + | 99859115 | 99917822 | regular |
| ENSMUSG00000069917 | Hba-a2 | chr11 | + | 32296488 | 32297298 | regular |
| ENSMUSG00000075511 | 1700001L05F | chr15 | - | 83357525 | 83367282 | regular |
| ENSMUSG00000018209 | Stk4 | chr2 | + | 164070321 | 164155524 | regular |
| ENSMUSG00000026663 | Atf6 | chr1 | - | 170704673 | 170867771 | regular |
| ENSMUSG00000028294 | Cfap206 | chr4 | - | 34688558 | 34730206 | regular |
| ENSMUSG00000008813 | Tppp2 | chr14 | + | 51918424 | 51920700 | regular |
| ENSMUSG00000024515 | Smad4 | chr18 | - | 73639008 | 73703780 | regular |
| ENSMUSG00000025453 | Nnt | chr13 | - | 119335447 | 119408997 | regular |
| ENSMUSG00000006998 | Psmd2 | chr16 | + | 20651651 | 20663414 | regular |
| ENSMUSG00000013787 | Ehmt2 | chr17 | + | 34898468 | 34914052 | regular |
| ENSMUSG00000040140 | Tdrd6 | chr17 | - | 43615334 | 43630299 | regular |
| ENSMUSG00000030030 | 1700003E16I | chr6 | + | 83156403 | 83162945 | regular |
| ENSMUSG00000003970 | Rpl8 | chr15 | + | 76904070 | 76906318 | regular |
| ENSMUSG00000025978 | Rftn2 | chr1 | - | 55170158 | 55226782 | regular |
| ENSMUSG00000019804 | Snx3 | chr10 | + | 42502029 | 42535381 | regular |
| ENSMUSG00000028458 | Tesk1 | chr4 | + | 43441938 | 43448064 | regular |
| ENSMUSG00000062397 | Zfp706 | chr15 | - | 36997026 | 37007773 | regular |
| ENSMUSG00000057738 | Sptan1 | chr2 | + | 29965559 | 30031451 | regular |
| ENSMUSG00000053279 | Aldh1a1 | chr19 | + | 20492714 | 20643462 | regular |
| ENSMUSG00000026571 | Dcaf6 | chr1 | - | 165328697 | 165460475 | regular |
| ENSMUSG00000022708 | Zbtb20 | chr16 | + | 42875880 | 43642602 | regular |
| ENSMUSG00000026609 | Ush2a | chr1 | + | 188262022 | 188965041 | regular |
| ENSMUSG00000099398 | Ms4a14 | chr19 | - | 11301248 | 11314454 | regular |
| ENSMUSG00000033632 | AW554918 | chr18 | + | 25168998 | 25467321 | regular |
| ENSMUSG00000030870 | Ubfd1 | chr7 | + | 122067174 | 122082194 | regular |
| ENSMUSG00000020647 | Ncoa1 | chr12 | - | 4247361 | 4477182 | regular |
| ENSMUSG00000035765 | Dym | chr18 | + | 75018771 | 75286966 | regular |
| ENSMUSG00000039473 | Ubn1 | chr16 | + | 5050067 | 5086285 | regular |
| ENSMUSG00000047502 | Mroh7 | chr4 | - | 106680416 | 106730925 | regular |
| ENSMUSG00000039652 | Cpeb3 | chr19 | - | 37021290 | 37208601 | regular |
| ENSMUSG00000020650 | Bcap29 | chr12 | - | 31590966 | 31634658 | regular |
| ENSMUSG00000026585 | Kifap3 | chr1 | + | 163779582 | 163917109 | regular |
| ENSMUSG00000097055 | Gm4419 | chr12 | + | 21417322 | 21421064 | regular |

| | | | | | | |
|---|---|---|---|---|---|---|
| ENSMUSG00000025224 | Gbf1 | chr19 | + | 46152508 | 46286510 | regular |
| ENSMUSG00000043036 | Ccdc63 | chr5 | - | 122108039 | 122140823 | regular |
| ENSMUSG00000022236 | Ropn1l | chr15 | - | 31441210 | 31453737 | regular |
| ENSMUSG00000028330 | Ncbp1 | chr4 | + | 46138612 | 46172403 | regular |
| ENSMUSG00000042408 | Zmym6 | chr4 | + | 127077382 | 127124372 | regular |
| ENSMUSG00000055024 | Ep300 | chr15 | + | 81585350 | 81652077 | regular |
| ENSMUSG00000003068 | Stk11 | chr10 | + | 80115802 | 80130682 | regular |
| ENSMUSG00000029788 | Cpa5 | chr6 | + | 30611009 | 30631745 | regular |
| ENSMUSG00000058690 | Ccser2 | chr14 | - | 36874935 | 36968777 | regular |
| ENSMUSG00000021969 | Zdhhc20 | chr14 | - | 57832702 | 57890276 | regular |
| ENSMUSG00000021007 | Spata7 | chr12 | + | 98628156 | 98669815 | regular |
| ENSMUSG00000039219 | Arid4b | chr13 | + | 14063231 | 14199603 | regular |
| ENSMUSG00000062939 | Stat4 | chr1 | + | 51987147 | 52107189 | regular |
| ENSMUSG00000041009 | Iqcf4 | chr9 | - | 106568318 | 106570996 | regular |
| ENSMUSG00000085438 | 1700020I14R | chr2 | + | 119594296 | 119607502 | regular |
| ENSMUSG00000042708 | Shcbp1l | chr1 | + | 153425161 | 153452574 | regular |
| ENSMUSG00000057719 | Sh3rf2 | chr18 | + | 42053666 | 42158960 | regular |
| ENSMUSG00000072878 | 1700123L14F | chr6 | - | 96164496 | 96166243 | regular |
| ENSMUSG00000031617 | Tmem184c | chr8 | - | 77595981 | 77610698 | regular |
| ENSMUST00000214747 | Scaper | chr9 | - | 55807012 | 55816809 | circular |
| ENSMUSG00000059439 | Bcas3 | chr11 | + | 85353166 | 85826058 | regular |
| ENSMUSG00000029458 | Brap | chr5 | + | 121660562 | 121687256 | regular |
| ENSMUSG00000033739 | Fkbpl | chr17 | + | 34644763 | 34646324 | regular |
| ENSMUSG00000003531 | Dgcr6 | chr16 | + | 18052859 | 18071632 | regular |
| ENSMUSG00000071454 | Dtnb | chr12 | + | 3572380 | 3781796 | regular |
| ENSMUSG00000044252 | Osbpl1a | chr18 | - | 12755313 | 12941841 | regular |
| ENSMUSG00000033487 | Fndc3a | chr14 | - | 72537945 | 72710003 | regular |
| ENSMUSG00000039585 | Myo9a | chr9 | + | 59750895 | 59928866 | regular |
| ENSMUSG00000025410 | Dctn2 | chr10 | + | 127266367 | 127281950 | regular |
| ENSMUSG00000002885 | Adgre5 | chr8 | - | 83723250 | 83741326 | regular |
| ENSMUSG00000061436 | Hipk2 | chr6 | - | 38694389 | 38876165 | regular |
| ENSMUSG00000080316 | Spaca6 | chr17 | + | 17827157 | 17843009 | regular |
| ENSMUSG00000021179 | Nrde2 | chr12 | - | 100125451 | 100159653 | regular |
| ENSMUSG00000040952 | Rps19 | chr7 | + | 24884370 | 24889806 | regular |
| ENSMUSG00000036097 | Slf2 | chr19 | + | 44931118 | 44983787 | regular |
| ENSMUSG00000020029 | Nudt4 | chr10 | - | 95547006 | 95564146 | regular |
| ENSMUSG00000037646 | Vps13b | chr15 | + | 35371159 | 35931229 | regular |
| ENSMUSG00000025132 | Arhgdia | chr11 | - | 120578103 | 120581624 | regular |
| ENSMUSG00000024163 | Mapk8ip3 | chr17 | - | 24892152 | 24936977 | regular |
| ENSMUSG00000042548 | Asxl1 | chr2 | + | 153345828 | 153404007 | regular |
| ENSMUSG00000050553 | Gk2 | chr5 | - | 97455141 | 97457021 | regular |
| ENSMUSG00000003810 | Mast2 | chr4 | - | 116306761 | 116464183 | regular |
| ENSMUSG00000020074 | Ccar1 | chr10 | - | 62743927 | 62792286 | regular |
| ENSMUSG00000047528 | Als2cr12 | chr1 | - | 58646902 | 58695989 | regular |
| ENSMUSG00000058443 | Rpl10-ps3 | chr9 | - | 50344200 | 50344968 | regular |
| ENSMUSG00000054909 | Wbscr25 | chr5 | - | 134987432 | 135001350 | regular |

| | | | | | | |
|---|---|---|---|---|---|---|
| ENSMUSG00000015365 | Mov10l1 | chr15 | + | 88982908 | 89055152 | regular |
| ENSMUSG00000036046 | 5031439G07 | chr15 | - | 84943935 | 84988551 | regular |
| ENSMUSG00000030059 | Tmf1 | chr6 | - | 97152996 | 97179122 | regular |
| ENSMUSG00000041673 | Lrrc18 | chr14 | + | 32991381 | 33015291 | regular |
| ENSMUSG00000028156 | Eif4e | chr3 | + | 138526178 | 138559696 | regular |
| ENSMUSG00000018428 | Akap1 | chr11 | - | 88830791 | 88864586 | regular |
| ENSMUSG00000040003 | Magi2 | chr5 | + | 19227035 | 20704792 | regular |
| ENSMUSG00000033423 | Eri3 | chr4 | + | 117550364 | 117674297 | regular |
| ENSMUSG00000035770 | Dync1li2 | chr8 | - | 104417679 | 104443047 | regular |
| ENSMUSG00000020021 | Fgd6 | chr10 | + | 94036000 | 94145339 | regular |
| ENSMUSG00000071415 | Rpl23 | chr11 | - | 97777526 | 97782437 | regular |
| ENSMUSG00000039963 | Ccdc40 | chr11 | + | 119228571 | 119265236 | regular |
| ENSMUSG00000044072 | Eml6 | chr11 | - | 29743047 | 30026033 | regular |
| ENSMUSG00000099250 | Rn7s2 | chr12 | - | 69361181 | 69361481 | regular |
| ENSMUSG00000099021 | Rn7s1 | chr12 | + | 69159294 | 69159594 | regular |
| ENSMUSG00000032489 | Kif9 | chr9 | + | 110476957 | 110525179 | regular |
| ENSMUSG00000031644 | Nek1 | chr8 | + | 60993194 | 61131346 | regular |
| ENSMUSG00000027519 | Rab22a | chr2 | + | 173659759 | 173707343 | regular |
| ENSMUSG00000025584 | Pde8a | chr7 | + | 81213595 | 81334533 | regular |
| ENSMUSG00000025104 | Hdgfl3 | chr7 | - | 81881250 | 81934473 | regular |
| ENSMUSG00000034212 | Ankmy1 | chr1 | - | 92859802 | 92902906 | regular |
| ENSMUSG00000019818 | Cd164 | chr10 | + | 41519413 | 41531048 | regular |
| ENSMUSG00000029338 | Antxr2 | chr5 | - | 97882782 | 98031043 | regular |
| ENSMUSG00000024238 | Zeb1 | chr18 | + | 5591859 | 5775467 | regular |
| ENSMUSG00000022388 | Ttll8 | chr15 | - | 88890632 | 88954418 | regular |
| ENSMUSG00000042350 | Arel1 | chr12 | - | 84918147 | 84970900 | regular |
| ENSMUSG00000024222 | Fkbp5 | chr17 | - | 28399094 | 28517524 | regular |
| ENSMUSG00000023923 | Tbc1d5 | chr17 | - | 50733123 | 51179352 | regular |
| ENSMUSG00000005442 | Cic | chr7 | + | 25267703 | 25294159 | regular |
| ENSMUSG00000027304 | Rtf1 | chr2 | + | 119675067 | 119735407 | regular |
| ENSMUSG00000035901 | Dennd5a | chr7 | - | 109893779 | 109960470 | regular |
| ENSMUSG00000052337 | Immt | chr6 | + | 71831330 | 71877388 | regular |
| ENSMUSG00000032382 | Snx1 | chr9 | - | 66088132 | 66126587 | regular |
| ENSMUSG00000026650 | Meig1 | chr2 | - | 3409042 | 3422648 | regular |
| ENSMUSG00000029516 | Cit | chr5 | + | 115845277 | 116008947 | regular |
| ENSMUSG00000032280 | Tle3 | chr9 | + | 61372365 | 61418497 | regular |
| ENSMUSG00000024352 | Spata24 | chr18 | - | 35656689 | 35662186 | regular |
| ENSMUSG00000045252 | Zfp574 | chr7 | + | 25072566 | 25083492 | regular |
| ENSMUSG00000034007 | Scaper | chr9 | - | 55549878 | 55938119 | regular |
| ENSMUSG00000015747 | Vps45 | chr3 | - | 95999831 | 96058466 | regular |
| ENSMUSG00000021461 | Fancc | chr13 | - | 63285042 | 63497278 | regular |
| ENSMUSG00000024913 | Lrp5 | chr19 | - | 3584827 | 3686564 | regular |
| ENSMUSG00000006673 | Qrich1 | chr9 | + | 108516805 | 108560163 | regular |
| ENSMUSG00000038587 | Akap12 | chr10 | + | 4266379 | 4359470 | regular |
| ENSMUSG00000067702 | Tuba3a | chr6 | - | 125278273 | 125286042 | regular |
| ENSMUSG00000036879 | Phkb | chr8 | + | 85840958 | 86061376 | regular |

| | | | | | | |
|---|---|---|---|---|---|---|
| ENSMUSG00000032612 | Usp4 | chr9 | + | 108347852 | 108392545 | regular |
| ENSMUSG00000038122 | Tbc1d32 | chr10 | - | 56014292 | 56228689 | regular |
| ENSMUSG00000036641 | Ccdc148 | chr2 | - | 58821069 | 59160683 | regular |
| ENSMUSG00000034163 | Zfc3h1 | chr10 | + | 115384958 | 115432772 | regular |
| ENSMUSG00000028059 | Arhgef2 | chr3 | + | 88605965 | 88648052 | regular |
| ENSMUSG00000039523 | Cep104 | chr4 | + | 153975193 | 154008732 | regular |
| ENSMUSG00000025900 | Rp1 | chr1 | - | 3999556 | 4409241 | regular |
| ENSMUSG00000035399 | Oser1 | chr2 | - | 163405821 | 163424697 | regular |
| ENSMUSG00000042797 | Aqp11 | chr7 | - | 97724005 | 97738289 | regular |
| ENSMUSG00000025431 | Crisp1 | chr17 | - | 40293757 | 40319207 | regular |
| ENSMUSG00000021391 | Cenpp | chr13 | - | 49464022 | 49652785 | regular |
| ENSMUST00000039798 | Bbs9 | chr9 | + | 22643744 | 22679064 | circular |
| ENSMUSG00000069971 | 4933402J07R | chr8 | + | 87563852 | 87589424 | regular |
| ENSMUSG00000033987 | Dnah17 | chr11 | - | 118021722 | 118130634 | regular |
| ENSMUSG00000031065 | Cdk16 | chrX | + | 20687953 | 20699880 | regular |
| ENSMUSG00000003814 | Calr | chr8 | - | 84841849 | 84846934 | regular |
| ENSMUSG00000032053 | Pou2af1 | chr9 | + | 51213707 | 51240080 | regular |
| ENSMUSG00000003269 | Cyth2 | chr7 | - | 45806636 | 45814581 | regular |
| ENSMUSG00000002043 | Trappc6a | chr7 | + | 19508394 | 19516145 | regular |
| ENSMUSG00000039168 | Dap | chr15 | + | 31224313 | 31274341 | regular |
| ENSMUSG00000025156 | Gps1 | chr11 | + | 120784271 | 120789102 | regular |
| ENSMUSG00000039782 | Cpeb2 | chr5 | + | 43233169 | 43289724 | regular |
| ENSMUSG00000074619 | 1700034I23R | chr3 | - | 40900195 | 40902543 | regular |
| ENSMUSG00000046230 | Vps13a | chr19 | - | 16615365 | 16780933 | regular |
| ENSMUSG00000037286 | Stag1 | chr9 | + | 100597797 | 100959375 | regular |
| ENSMUSG00000015656 | Hspa8 | chr9 | + | 40800983 | 40810087 | regular |
| ENSMUSG00000034707 | Gns | chr10 | + | 121365089 | 121397249 | regular |
| ENSMUSG00000033486 | Catsper2 | chr2 | - | 121392630 | 121413792 | regular |
| ENSMUSG00000030206 | Gsg1 | chr6 | - | 135237325 | 135254351 | regular |
| ENSMUSG00000035473 | Galm | chr17 | + | 80127470 | 80185101 | regular |
| ENSMUSG00000073940 | Hbb-bt | chr7 | - | 103812523 | 103813996 | regular |
| ENSMUSG00000037965 | Zc3h7a | chr16 | - | 11136591 | 11176393 | regular |
| ENSMUSG00000058135 | Gstm1 | chr3 | - | 108012254 | 108017973 | regular |
| ENSMUSG00000062017 | Abca14 | chr7 | + | 120203960 | 120325352 | regular |
| ENSMUSG00000036019 | Tmtc2 | chr10 | - | 105187662 | 105574451 | regular |
| ENSMUSG00000021738 | Atxn7 | chr14 | + | 13961439 | 14107302 | regular |
| ENSMUSG00000014077 | Chp1 | chr2 | + | 119547696 | 119587027 | regular |
| ENSMUSG00000052469 | Tcp10c | chr17 | + | 13354571 | 13377223 | regular |
| ENSMUSG00000025480 | Syce1 | chr7 | - | 140777228 | 140787852 | regular |
| ENSMUSG00000028926 | Cdk14 | chr5 | - | 4803390 | 5420312 | regular |
| ENSMUSG00000026701 | Prdx6 | chr1 | - | 161240111 | 161251219 | regular |
| ENSMUSG00000028677 | Rnf220 | chr4 | - | 117271462 | 117497052 | regular |
| ENSMUSG00000031278 | Acsl4 | chrX | - | 142317992 | 142390535 | regular |
| ENSMUSG00000031085 | Gm498 | chr7 | + | 143866837 | 143900046 | regular |
| ENSMUSG00000061923 | Odf1 | chr15 | + | 38219202 | 38226735 | regular |
| ENSMUSG00000026927 | Sdccag3 | chr2 | - | 26382801 | 26389316 | regular |

| | | | | | | |
|---|---|---|---|---|---|---|
| ENSMUSG00000028896 | Rcc1 | chr4 | - | 132331918 | 132353605 | regular |
| ENSMUSG00000029423 | Piwil1 | chr5 | + | 128702523 | 128755474 | regular |
| ENSMUSG00000032316 | Clk3 | chr9 | - | 57750623 | 57765860 | regular |
| ENSMUSG00000037487 | Ubr5 | chr15 | - | 37967327 | 38078854 | regular |
| ENSMUSG00000061983 | Rps12 | chr10 | - | 23785182 | 23787275 | regular |
| ENSMUSG00000004364 | Cul3 | chr1 | - | 80264922 | 80340480 | regular |
| ENSMUSG00000073758 | Sh3d21 | chr4 | - | 126150601 | 126163491 | regular |
| ENSMUSG00000024251 | Thada | chr17 | - | 84190055 | 84466196 | regular |
| ENSMUSG00000036199 | Ndufa13 | chr8 | - | 69894179 | 69902558 | regular |
| ENSMUSG00000027007 | Ssfa2 | chr2 | + | 79635351 | 79672966 | regular |
| ENSMUSG00000018211 | Wfdc15b | chr2 | - | 164214453 | 164221660 | regular |
| ENSMUSG00000021902 | Phf7 | chr14 | - | 31237688 | 31251317 | regular |
| ENSMUSG00000038481 | Cdk19 | chr10 | + | 40339563 | 40483818 | regular |
| ENSMUSG00000020372 | Rack1 | chr11 | + | 48800331 | 48806434 | regular |
| ENSMUSG00000003604 | Aven | chr2 | + | 112492963 | 112634573 | regular |
| ENSMUSG00000040852 | Plekhh2 | chr17 | + | 84511894 | 84622142 | regular |
| ENSMUSG00000026565 | Pou2f1 | chr1 | - | 165865153 | 166002678 | regular |
| ENSMUSG00000032562 | Gnai2 | chr9 | - | 107614124 | 107635367 | regular |
| ENSMUSG00000034024 | Cct2 | chr10 | - | 117051000 | 117063814 | regular |
| ENSMUSG00000041343 | Ankrd42 | chr7 | - | 92581722 | 92637142 | regular |
| ENSMUSG00000069565 | Dazap1 | chr10 | + | 80261485 | 80288408 | regular |
| ENSMUSG00000021577 | Sdha | chr13 | - | 74322253 | 74350280 | regular |
| ENSMUSG00000087574 | C030037D09I | chr11 | + | 88718644 | 88728893 | regular |
| ENSMUSG00000030230 | Plcz1 | chr6 | - | 139989672 | 140041457 | regular |
| ENSMUSG00000027030 | Stk39 | chr2 | - | 68210444 | 68472268 | regular |
| ENSMUSG00000086451 | 4933431K23I | chr8 | + | 77610859 | 77769682 | regular |
| ENSMUSG00000025103 | Btbd1 | chr7 | - | 81792073 | 81829431 | regular |
| ENSMUSG00000028035 | Dnajb4 | chr3 | - | 152178510 | 152210302 | regular |
| ENSMUSG00000048758 | Rpl29 | chr9 | + | 106429453 | 106431568 | regular |
| ENSMUSG00000020745 | Pafah1b1 | chr11 | - | 74673948 | 74724670 | regular |
| ENSMUSG00000070331 | Qrich2 | chr11 | - | 116441324 | 116466241 | regular |
| ENSMUSG00000047126 | Cltc | chr11 | - | 86694350 | 86757565 | regular |
| ENSMUSG00000008429 | Herpud2 | chr9 | - | 25108131 | 25151820 | regular |
| ENSMUSG00000066278 | Vps37b | chr5 | - | 124004640 | 124032270 | regular |
| ENSMUSG00000020362 | Cnot6 | chr11 | - | 49671502 | 49712723 | regular |
| ENSMUSG00000038429 | Usp5 | chr6 | - | 124815018 | 124829484 | regular |
| ENSMUSG00000031924 | Cyb5b | chr8 | + | 107150639 | 107187471 | regular |
| ENSMUSG00000052926 | Rnaseh2a | chr8 | - | 84956609 | 84969767 | regular |
| ENSMUSG00000010097 | Nxf1 | chr19 | + | 8757072 | 8772475 | regular |
| ENSMUSG00000039033 | Tasp1 | chr2 | - | 139833479 | 140066805 | regular |
| ENSMUSG00000037143 | Cfap61 | chr2 | + | 145934783 | 146215039 | regular |
| ENSMUSG00000022307 | Oxr1 | chr15 | + | 41447481 | 41861048 | regular |
| ENSMUSG00000067629 | Syngap1 | chr17 | + | 26941252 | 26972434 | regular |
| ENSMUSG00000056234 | Ncoa4 | chr14 | + | 32159864 | 32179855 | regular |
| ENSMUSG00000028560 | Usp1 | chr4 | + | 98923809 | 98935543 | regular |
| ENSMUSG00000024059 | Clip4 | chr17 | + | 71781946 | 71858351 | regular |

| | | | | | | |
|---|---|---|---|---|---|---|
| ENSMUSG00000011877 | Git1 | chr11 | + | 77493561 | 77507786 | regular |
| ENSMUSG00000023806 | Rsph3b | chr17 | - | 6904715 | 6948356 | regular |
| ENSMUSG00000041560 | Nop53 | chr7 | - | 15936182 | 15946074 | regular |
| ENSMUSG00000034151 | Zbbx | chr3 | - | 75037906 | 75165034 | regular |
| ENSMUSG00000026384 | Ptpn4 | chr1 | - | 119652466 | 119837613 | regular |
| ENSMUSG00000040761 | Spen | chr4 | - | 141467889 | 141538597 | regular |
| ENSMUSG00000020622 | Nt5c1b | chr12 | + | 10369972 | 10390175 | regular |
| ENSMUSG00000063810 | Alms1 | chr6 | + | 85587530 | 85702753 | regular |
| ENSMUSG00000030199 | Etv6 | chr6 | + | 134035699 | 134270158 | regular |
| ENSMUSG00000041225 | Arhgap12 | chr18 | - | 6024426 | 6136098 | regular |
| ENSMUSG00000003131 | Pafah1b2 | chr9 | - | 45962858 | 46012690 | regular |
| ENSMUSG00000001173 | Ocrl | chrX | + | 47912386 | 47965868 | regular |
| ENSMUSG00000036550 | Cnot1 | chr8 | - | 95719450 | 95807464 | regular |
| ENSMUSG00000055670 | Zzef1 | chr11 | + | 72796225 | 72927120 | regular |
| ENSMUSG00000044042 | Fmn1 | chr2 | + | 113327735 | 113716767 | regular |
| ENSMUSG00000064360 | mt-Nd3 | chrM | + | 9458 | 9806 | regular |
| ENSMUSG00000036002 | Fam214b | chr4 | - | 43032413 | 43046220 | regular |
| ENSMUSG00000060843 | Ctnna3 | chr10 | + | 63430097 | 65003667 | regular |
| ENSMUSG00000001558 | Klhl10 | chr11 | + | 100441916 | 100457022 | regular |
| ENSMUSG00000040351 | Ankib1 | chr5 | - | 3689999 | 3802925 | regular |
| ENSMUSG00000030138 | Bms1 | chr6 | - | 118383380 | 118419474 | regular |
| ENSMUSG00000024357 | Sil1 | chr18 | - | 35266399 | 35498925 | regular |
| ENSMUSG00000090457 | 4930571K23I | chr7 | + | 125368860 | 125371013 | regular |
| ENSMUSG00000074212 | Dnajb14 | chr3 | + | 137867674 | 137916557 | regular |
| ENSMUSG00000035161 | Ints6 | chr14 | - | 62676329 | 62761169 | regular |
| ENSMUSG00000042567 | Nek10 | chr14 | + | 14803414 | 15012059 | regular |
| ENSMUSG00000045275 | Lca5l | chr16 | - | 96158406 | 96192257 | regular |
| ENSMUSG00000064138 | Fam172a | chr13 | + | 77702032 | 78166235 | regular |
| ENSMUSG00000032624 | Eml4 | chr17 | + | 83350930 | 83480361 | regular |
| ENSMUSG00000069895 | Atxn1l | chr8 | - | 109726450 | 109737739 | regular |
| ENSMUSG00000002546 | Golga2 | chr2 | + | 32287383 | 32307921 | regular |
| ENSMUSG00000019978 | Epb41l2 | chr10 | + | 25359797 | 25523519 | regular |
| ENSMUSG00000060743 | H3f3a | chr1 | - | 180800831 | 180813943 | regular |
| ENSMUSG00000047648 | Fbxo30 | chr10 | + | 11281329 | 11298052 | regular |
| ENSMUSG00000020817 | Rabep1 | chr11 | + | 70844777 | 70943105 | regular |
| ENSMUSG00000037331 | Larp1 | chr11 | + | 58009063 | 58062034 | regular |
| ENSMUSG00000095789 | Nupr1l | chr5 | + | 129908539 | 129913789 | regular |
| ENSMUSG00000040594 | Ranbp17 | chr11 | - | 33211794 | 33513746 | regular |
| ENSMUSG00000006699 | Cdc42 | chr4 | - | 137319695 | 137357720 | regular |
| ENSMUSG00000039236 | Isg20 | chr7 | + | 78913423 | 78920396 | regular |
| ENSMUSG00000019820 | Utrn | chr10 | - | 12382187 | 12869365 | regular |
| ENSMUSG00000038594 | Cep85l | chr10 | - | 53273442 | 53379947 | regular |
| ENSMUSG00000028772 | Zcchc17 | chr4 | - | 130315382 | 130359943 | regular |
| ENSMUSG00000007880 | Arid1a | chr4 | - | 133679007 | 133756769 | regular |
| ENSMUSG00000071322 | Tcp10a | chr17 | + | 7324645 | 7345974 | regular |
| ENSMUSG00000025986 | Slc39a10 | chr1 | - | 46807543 | 46892852 | regular |

| | | | | | | |
|---|---|---|---|---|---|---|
| ENSMUSG00000039210 | Gpatch2 | chr1 | + | 187215507 | 187351704 | regular |
| ENSMUSG00000042772 | Smg7 | chr1 | - | 152836994 | 152902646 | regular |
| ENSMUSG00000058183 | Mmel1 | chr4 | + | 154869584 | 154895528 | regular |
| ENSMUSG00000058546 | Rpl23a | chr11 | - | 78180931 | 78183584 | regular |
| ENSMUSG00000038774 | Ascc3 | chr10 | + | 50592668 | 50851485 | regular |
| ENSMUSG00000000215 | Ins2 | chr7 | - | 142678655 | 142743381 | regular |
| ENSMUSG00000038095 | Sbno1 | chr5 | - | 124368701 | 124426001 | regular |
| ENSMUSG00000050017 | Pitpnb | chr5 | + | 111330762 | 111388359 | regular |
| ENSMUSG00000063972 | Nr6a1 | chr2 | - | 38723369 | 38927688 | regular |
| ENSMUSG00000025290 | Rps24 | chr14 | + | 24487124 | 24496959 | regular |
| ENSMUSG00000020393 | Kremen1 | chr11 | - | 5191551 | 5261558 | regular |
| ENSMUSG00000029475 | Kdm2b | chr5 | - | 122870664 | 122989823 | regular |
| ENSMUSG00000046111 | Cep295 | chr9 | - | 15316914 | 15357788 | regular |
| ENSMUSG00000040725 | Hnrnpul1 | chr7 | - | 25721164 | 25754757 | regular |
| ENSMUSG00000042888 | Prr30 | chr14 | - | 101197689 | 101200330 | regular |
| ENSMUSG00000019947 | Arid5b | chr10 | - | 68092519 | 68278740 | regular |
| ENSMUSG00000006127 | Inpp5k | chr11 | + | 75630987 | 75648871 | regular |
| ENSMUSG00000037437 | Adam32 | chr8 | - | 24836139 | 24948804 | regular |
| ENSMUSG00000042726 | Trafd1 | chr5 | - | 121371724 | 121385632 | regular |
| ENSMUSG00000032409 | Atr | chr9 | + | 95857596 | 95951781 | regular |
| ENSMUSG00000002625 | Akap8l | chr17 | - | 32321424 | 32350581 | regular |
| ENSMUSG00000035696 | Rnf38 | chr4 | - | 44126209 | 44233789 | regular |
| ENSMUSG00000052459 | Atp6v1a | chr16 | - | 44085401 | 44139705 | regular |
| ENSMUSG00000026601 | Axdnd1 | chr1 | - | 156323508 | 156421159 | regular |
| ENSMUSG00000027465 | Tbc1d20 | chr2 | + | 152293827 | 152313996 | regular |
| ENSMUSG00000020878 | Lrrc46 | chr11 | - | 97034601 | 97041407 | regular |
| ENSMUSG00000057637 | Prdm2 | chr4 | - | 143107390 | 143212995 | regular |
| ENSMUSG00000090202 | 4930503B20 | chr3 | - | 146645192 | 146651317 | regular |
| ENSMUSG00000005575 | Ube2m | chr7 | - | 13035119 | 13038275 | regular |
| ENSMUSG00000021326 | Trim27 | chr13 | + | 21179444 | 21194724 | regular |
| ENSMUSG00000042156 | Dzip1 | chr14 | - | 118875519 | 118925460 | regular |
| ENSMUSG00000020196 | Cabin1 | chr10 | - | 75646111 | 75764341 | regular |
| ENSMUSG00000024780 | Cdc37l1 | chr19 | + | 28990351 | 29026681 | regular |
| ENSMUSG00000007617 | Homer1 | chr13 | + | 93299634 | 93405129 | regular |
| ENSMUSG00000056267 | Cep70 | chr9 | + | 99243366 | 99300404 | regular |
| ENSMUSG00000046330 | Rpl37a | chr1 | + | 72711289 | 72713813 | regular |
| ENSMUSG00000022336 | Eif3e | chr15 | - | 43250057 | 43282719 | regular |
| ENSMUSG00000101337 | Dnah7c | chr1 | + | 46425591 | 46807476 | regular |
| ENSMUSG00000020220 | Vps13d | chr4 | - | 144972621 | 145195005 | regular |
| ENSMUSG00000057132 | Rpgrip1 | chr14 | + | 52110703 | 52163546 | regular |
| ENSMUSG00000043252 | Tmem64 | chr4 | + | 15265830 | 15286753 | regular |
| ENSMUSG00000018339 | Gpx3 | chr11 | + | 54902452 | 54910377 | regular |
| ENSMUSG00000026159 | Agfg1 | chr1 | + | 82839482 | 82901182 | regular |
| ENSMUSG00000031831 | Dnaaf1 | chr8 | + | 119575234 | 119598454 | regular |
| ENSMUSG00000049252 | Lrp1b | chr2 | - | 40595245 | 42653624 | regular |
| ENSMUSG00000021733 | Slc4a7 | chr14 | + | 14702278 | 14799940 | regular |

| | | | | | | |
|---|---|---|---|---|---|---|
| ENSMUSG00000056458 | Mok | chr12 | - | 110807797 | 110840939 | regular |
| ENSMUSG00000032582 | Rbm6 | chr9 | - | 107773558 | 107873237 | regular |
| ENSMUSG00000026113 | Inpp4a | chr1 | + | 37299864 | 37410736 | regular |
| ENSMUSG00000022253 | Nadk2 | chr15 | + | 9071259 | 9110891 | regular |
| ENSMUSG00000029726 | Mepce | chr5 | - | 137781905 | 137787655 | regular |
| ENSMUSG00000001948 | Spa17 | chr9 | - | 37603289 | 37613722 | regular |
| ENSMUSG00000037062 | Sh3glb1 | chr3 | - | 144683677 | 144720335 | regular |
| ENSMUSG00000062901 | Klhl24 | chr16 | + | 20097553 | 20127744 | regular |
| ENSMUSG00000033882 | Rbm46 | chr3 | - | 82837227 | 82876483 | regular |
| ENSMUSG00000001909 | Trmt1 | chr8 | + | 84686306 | 84699808 | regular |
| ENSMUSG00000038914 | Dido1 | chr2 | - | 180657963 | 180709999 | regular |
| ENSMUSG00000057230 | Aak1 | chr6 | + | 86849516 | 87003223 | regular |
| ENSMUSG00000016921 | Srsf6 | chr2 | + | 162931527 | 162937121 | regular |
| ENSMUSG00000073730 | 4933415F23I | chr1 | - | 23100472 | 23102253 | regular |
| ENSMUSG00000015599 | Ttbk1 | chr17 | - | 46442447 | 46487675 | regular |
| ENSMUSG00000029580 | Actb | chr5 | - | 142903114 | 142906754 | regular |
| ENSMUSG00000068167 | Csnka2ip | chr16 | - | 64477805 | 64602408 | regular |
| ENSMUSG00000063358 | Mapk1 | chr16 | + | 16983381 | 17047453 | regular |
| ENSMUSG00000028706 | Nsun4 | chr4 | - | 116032841 | 116053876 | regular |
| ENSMUSG00000000552 | Zfp385a | chr15 | - | 103313894 | 103340086 | regular |
| ENSMUSG00000093674 | Rpl41 | chr10 | - | 128548113 | 128549305 | regular |
| ENSMUSG00000036752 | Tubb4b | chr2 | - | 25222159 | 25224702 | regular |
| ENSMUSG00000022540 | Rogdi | chr16 | - | 5008729 | 5013553 | regular |
| ENSMUSG00000026817 | Ak1 | chr2 | + | 32621757 | 32635058 | regular |
| ENSMUSG00000042507 | Elmsan1 | chr12 | - | 84149175 | 84218881 | regular |
| ENSMUSG00000050089 | Akap4 | chrX | + | 7067518 | 7078609 | regular |
| ENSMUSG00000028637 | Ccdc30 | chr4 | - | 119322892 | 119415521 | regular |
| ENSMUSG00000024661 | Fth1 | chr19 | + | 9982702 | 9985092 | regular |
| ENSMUSG00000069769 | Msi2 | chr11 | - | 88339381 | 88718513 | regular |
| ENSMUSG00000022432 | Smc1b | chr15 | - | 85064688 | 85131964 | regular |
| ENSMUSG00000040669 | Phc1 | chr6 | - | 122317730 | 122340561 | regular |
| ENSMUSG00000032803 | Cdv3 | chr9 | - | 103353093 | 103365840 | regular |
| ENSMUSG00000029442 | Wdr66 | chr5 | + | 123252101 | 123327484 | regular |
| ENSMUSG00000029505 | Ep400 | chr5 | - | 110664372 | 110770717 | regular |
| ENSMUSG00000028938 | Galntl5 | chr5 | + | 25181459 | 25220297 | regular |
| ENSMUSG00000031839 | Hsbp1 | chr8 | + | 119344487 | 119348922 | regular |
| ENSMUSG00000029686 | Cul1 | chr6 | + | 47453397 | 47526139 | regular |
| ENSMUSG00000024359 | Hspa9 | chr18 | - | 34937413 | 34954357 | regular |
| ENSMUSG00000031563 | Wwc2 | chr8 | - | 47823958 | 47990924 | regular |
| ENSMUSG00000061455 | Stx17 | chr4 | + | 48124914 | 48186507 | regular |
| ENSMUSG00000070047 | Fat1 | chr8 | + | 44935446 | 45052257 | regular |
| ENSMUSG00000029330 | Cds1 | chr5 | + | 101765129 | 101823858 | regular |
| ENSMUSG00000037940 | Inpp4b | chr8 | + | 81342555 | 82127914 | regular |
| ENSMUSG00000020389 | Cdkl3 | chr11 | + | 52004220 | 52089784 | regular |
| ENSMUSG00000023953 | Polh | chr17 | - | 46172003 | 46202625 | regular |
| ENSMUSG00000028991 | Mtor | chr4 | + | 148448610 | 148557683 | regular |

| | | | | | | |
|---|---|---|---|---|---|---|
| ENSMUSG00000047518 | Slfnl1 | chr4 | + | 120532230 | 120536661 | regular |
| ENSMUSG00000027630 | Tbl1xr1 | chr3 | + | 22076651 | 22216594 | regular |
| ENSMUSG00000063457 | Rps15 | chr10 | + | 80292452 | 80294114 | regular |
| ENSMUSG00000020610 | Amz2 | chr11 | + | 109425945 | 109452813 | regular |
| ENSMUSG00000022155 | Mroh2b | chr15 | + | 4898736 | 4962205 | regular |
| ENSMUSG00000092746 | Rn7s6 | chr6 | + | 69516338 | 69516629 | regular |
| ENSMUSG00000042354 | Gnl3 | chr14 | - | 31012432 | 31019152 | regular |
| ENSMUSG00000025745 | Hadha | chr5 | - | 30118303 | 30155162 | regular |
| ENSMUSG00000021669 | Col4a3bp | chr13 | + | 96542617 | 96640167 | regular |
| ENSMUSG00000055762 | Eef1d | chr15 | - | 75894795 | 75909556 | regular |
| ENSMUSG00000040690 | Col16a1 | chr4 | + | 130047839 | 130099283 | regular |
| ENSMUSG00000038523 | 1700003F12I | chr2 | + | 154547084 | 154550003 | regular |
| ENSMUSG00000001441 | Npepps | chr11 | - | 97205841 | 97280638 | regular |
| ENSMUSG00000007107 | Atp1a4 | chr1 | - | 172223512 | 172258414 | regular |
| ENSMUSG00000028673 | Fuca1 | chr4 | + | 135920734 | 135940311 | regular |
| ENSMUSG00000027248 | Pdia3 | chr2 | + | 121413774 | 121438687 | regular |
| ENSMUSG00000061859 | Patj | chr4 | + | 98395784 | 98719603 | regular |
| ENSMUSG00000001424 | Snd1 | chr6 | + | 28475138 | 28935162 | regular |
| ENSMUSG00000038970 | Lmtk2 | chr5 | + | 144100435 | 144188204 | regular |
| ENSMUSG00000028383 | Hsdl2 | chr4 | + | 59581562 | 59618689 | regular |
| ENSMUSG00000031893 | Tsnaxip1 | chr8 | + | 105827730 | 105844680 | regular |
| ENSMUSG00000052698 | Tln2 | chr9 | - | 67217086 | 67559703 | regular |
| ENSMUSG00000096054 | Syne1 | chr10 | - | 5020916 | 5551482 | regular |
| ENSMUSG00000071866 | Ppia | chr11 | + | 6415442 | 6419817 | regular |
| ENSMUSG00000002210 | Smg9 | chr7 | + | 24399618 | 24422778 | regular |
| ENSMUSG00000039501 | Znfx1 | chr2 | - | 167035792 | 167063015 | regular |
| ENSMUSG00000059895 | Ptp4a3 | chr15 | + | 73723144 | 73758766 | regular |
| ENSMUSG00000004934 | Pias4 | chr10 | - | 81153265 | 81167923 | regular |
| ENSMUSG00000020015 | Cdk17 | chr10 | + | 93160874 | 93267071 | regular |
| ENSMUSG00000049123 | Catsperg2 | chr7 | - | 29697218 | 29727032 | regular |
| ENSMUSG00000057406 | Nsd2 | chr5 | + | 33820724 | 33897975 | regular |
| ENSMUSG00000073702 | Rpl31 | chr1 | + | 39367841 | 39371911 | regular |
| ENSMUSG00000035437 | Rabgap1 | chr2 | + | 37443278 | 37566454 | regular |
| ENSMUSG00000034088 | Hdlbp | chr1 | - | 93405939 | 93478815 | regular |
| ENSMUSG00000066643 | Wdr35 | chr12 | + | 8973891 | 9028847 | regular |
| ENSMUSG00000060279 | Ap2a1 | chr7 | - | 44900372 | 44929496 | regular |
| ENSMUSG00000021242 | Npc2 | chr12 | - | 84754561 | 84773152 | regular |
| ENSMUSG00000007411 | Mark3 | chr12 | + | 111574522 | 111656221 | regular |
| ENSMUSG00000032050 | Rdx | chr9 | + | 52047172 | 52088735 | regular |
| ENSMUSG00000000804 | Usp32 | chr11 | - | 84984441 | 85140161 | regular |
| ENSMUSG00000061032 | Rrp1 | chr10 | - | 78400383 | 78413043 | regular |
| ENSMUSG00000022066 | Entpd4b | chr14 | + | 69555423 | 69585520 | regular |
| ENSMUSG00000095463 | Entpd4 | chr14 | + | 69337173 | 69367270 | regular |
| ENSMUSG00000040451 | Sgms1 | chr19 | - | 32122726 | 32389714 | regular |
| ENSMUSG00000035203 | Epn1 | chr7 | + | 5080234 | 5098178 | regular |
| ENSMUSG00000023930 | Crisp2 | chr17 | - | 40764733 | 40807003 | regular |

| | | | | | | |
|---|---|---|---|---|---|---|
| ENSMUSG00000033392 | Clasp2 | chr9 | + | 113741472 | 113919682 | regular |
| ENSMUSG00000018501 | Ncor1 | chr11 | - | 62316425 | 62458541 | regular |
| ENSMUSG00000053929 | Cyhr1 | chr15 | - | 76643394 | 76660117 | regular |
| ENSMUSG00000031493 | Ggn | chr7 | + | 29170209 | 29173976 | regular |
| ENSMUSG00000027423 | Snx5 | chr2 | - | 144250122 | 144270906 | regular |
| ENSMUSG00000022246 | Rai14 | chr15 | - | 10568968 | 10714624 | regular |
| ENSMUSG00000021936 | Mapk8 | chr14 | - | 33377897 | 33447158 | regular |
| ENSMUSG00000079608 | Stard6 | chr18 | + | 70472453 | 70501066 | regular |
| ENSMUSG00000044084 | Spem2 | chr11 | - | 69816565 | 69818465 | regular |
| ENSMUSG00000074781 | Ube2n | chr10 | + | 95515144 | 95545657 | regular |
| ENSMUSG00000022377 | Asap1 | chr15 | - | 64086856 | 64382919 | regular |
| ENSMUSG00000041596 | Nlrp5-ps | chr7 | - | 14530651 | 14622479 | regular |
| ENSMUSG00000021830 | Txndc16 | chr14 | - | 45133464 | 45220328 | regular |
| ENSMUSG00000070372 | Capza1 | chr3 | - | 104822778 | 104864505 | regular |
| ENSMUSG00000059920 | 4930453N24 | chr16 | - | 64751750 | 64771161 | regular |
| ENSMUSG00000064351 | mt-Co1 | chrM | + | 5327 | 6872 | regular |
| ENSMUSG00000031518 | Spata4 | chr8 | + | 54600780 | 54610098 | regular |
| ENSMUSG00000022783 | Spag6l | chr16 | - | 16753015 | 16829456 | regular |
| ENSMUSG00000029074 | Ttll10 | chr4 | - | 156034839 | 156059414 | regular |
| ENSMUSG00000030313 | Dennd5b | chr6 | - | 148988070 | 149101680 | regular |
| ENSMUSG00000069516 | Lyz2 | chr10 | - | 117277330 | 117282321 | regular |
| ENSMUSG00000079429 | Mroh2a | chr1 | + | 88226985 | 88262289 | regular |
| ENSMUSG00000026473 | Glul | chr1 | + | 153899943 | 153909723 | regular |
| ENSMUSG00000061533 | Cep128 | chr12 | - | 90998491 | 91384409 | regular |
| ENSMUSG00000012819 | Cdh23 | chr10 | - | 60302747 | 60696490 | regular |
| ENSMUSG00000000682 | Cd52 | chr4 | - | 134082447 | 134095082 | regular |
| ENSMUST00000046656 | Tasp1 | chr2 | - | 139977606 | 140057499 | circular |
| ENSMUSG00000044533 | Rps2 | chr17 | + | 24718115 | 24721929 | regular |
| ENSMUSG00000031622 | Sin3b | chr8 | + | 72723284 | 72758201 | regular |
| ENSMUSG00000038290 | Smg6 | chr11 | + | 74925822 | 75164448 | regular |
| ENSMUSG00000042700 | Sipa1l1 | chr12 | + | 82169319 | 82451786 | regular |
| ENSMUSG00000039852 | Rere | chr4 | + | 150281645 | 150621966 | regular |
| ENSMUSG00000049517 | Rps23 | chr13 | + | 90922957 | 90924950 | regular |
| ENSMUSG00000040097 | Flywch1 | chr17 | - | 23752792 | 23771602 | regular |
| ENSMUSG00000022269 | Mar.11 | chr15 | + | 26309047 | 26409576 | regular |
| ENSMUSG00000024663 | Rab3il1 | chr19 | + | 10001668 | 10038380 | regular |
| ENSMUSG00000015357 | Clpx | chr9 | + | 65294259 | 65330658 | regular |
| ENSMUSG00000020657 | Dnajc27 | chr12 | + | 4082582 | 4110606 | regular |
| ENSMUSG00000034842 | Art3 | chr5 | + | 92331826 | 92414628 | regular |
| ENSMUSG00000035378 | Shq1 | chr6 | - | 100568255 | 100671157 | regular |
| ENSMUSG00000027067 | Ssrp1 | chr2 | + | 85037233 | 85047109 | regular |
| ENSMUSG00000024135 | Srbd1 | chr17 | - | 85984664 | 86145175 | regular |
| ENSMUSG00000026463 | Atp2b4 | chr1 | - | 133699456 | 133801041 | regular |
| ENSMUSG00000050074 | Spink8 | chr9 | + | 109812279 | 109826633 | regular |
| ENSMUSG00000051506 | Wdfy4 | chr14 | - | 32959546 | 33185508 | regular |
| ENSMUSG00000020917 | Acly | chr11 | - | 100476352 | 100528000 | regular |

| | | | | | | |
|---|---|---|---|---|---|---|
| ENSMUSG00000024240 | Epc1 | chr18 | - | 6435950 | 6516108 | regular |
| ENSMUSG00000042293 | Gm5617 | chr9 | + | 48495441 | 48495972 | regular |
| ENSMUSG00000045709 | Smkr-ps | chr6 | + | 29985328 | 29993776 | regular |
| ENSMUSG00000030068 | Gm20696 | chr6 | - | 99257483 | 99666797 | regular |
| ENSMUSG00000007907 | Cabs1 | chr5 | + | 87979438 | 87981544 | regular |
| ENSMUSG00000032187 | Smarca4 | chr9 | + | 21616168 | 21704230 | regular |
| ENSMUSG00000047369 | Dnah14 | chr1 | + | 181576558 | 181815774 | regular |
| ENSMUSG00000029291 | Rufy3 | chr5 | + | 88565039 | 88651392 | regular |
| ENSMUSG00000051934 | Spats2 | chr15 | + | 99126577 | 99213206 | regular |
| ENSMUSG00000028788 | Ptp4a2 | chr4 | + | 129811218 | 129850003 | regular |
| ENSMUSG00000010517 | Faf1 | chr4 | + | 109676587 | 109963960 | regular |
| ENSMUSG00000038042 | Ptpdc1 | chr13 | - | 48577871 | 48625664 | regular |
| ENSMUSG00000009470 | Tnpo1 | chr13 | - | 98839018 | 98926384 | regular |
| ENSMUSG00000022013 | Dnajc15 | chr14 | - | 77820680 | 77875125 | regular |
| ENSMUSG00000063129 | Aldoart2 | chr12 | + | 55565238 | 55566896 | regular |
| ENSMUSG00000039224 | D1Pas1 | chr1 | + | 186967415 | 186970627 | regular |
| ENSMUSG00000028678 | Kif2c | chr4 | - | 117159638 | 117182639 | regular |
| ENSMUSG00000000740 | Rpl13 | chr8 | + | 123102349 | 123105244 | regular |
| ENSMUSG00000048799 | Cep120 | chr18 | - | 53681723 | 53744547 | regular |
| ENSMUSG00000009575 | Cbx5 | chr15 | - | 103191543 | 103239816 | regular |
| ENSMUSG00000022110 | Sucla2 | chr14 | + | 73525318 | 73596142 | regular |
| ENSMUSG00000056429 | Tgoln1 | chr6 | - | 72608431 | 72617000 | regular |
| ENSMUSG00000022517 | Mgrn1 | chr16 | + | 4886099 | 4938296 | regular |
| ENSMUSG00000036928 | Stag3 | chr5 | + | 138280239 | 138312393 | regular |
| ENSMUSG00000059288 | Cdyl | chr13 | + | 35659832 | 35874063 | regular |
| ENSMUSG00000048264 | Dip2c | chr13 | + | 9276527 | 9668928 | regular |
| ENSMUSG00000020064 | Herc4 | chr10 | + | 63243809 | 63317878 | regular |
| ENSMUSG00000020849 | Ywhae | chr11 | + | 75732868 | 75765845 | regular |
| ENSMUSG00000030232 | Aebp2 | chr6 | + | 140622662 | 140678472 | regular |
| ENSMUSG00000040875 | Osbpl10 | chr9 | + | 114978568 | 115232225 | regular |
| ENSMUSG00000026163 | Sphkap | chr1 | - | 83254138 | 83408200 | regular |
| ENSMUSG00000032959 | Pebp1 | chr5 | - | 117282653 | 117287625 | regular |
| ENSMUSG00000001687 | Ubl3 | chr5 | - | 148504634 | 148552789 | regular |
| ENSMUSG00000067274 | Rplp0 | chr5 | + | 115559466 | 115563727 | regular |
| ENSMUSG00000051732 | Pabpc2 | chr18 | + | 39773496 | 39776082 | regular |
| ENSMUSG00000029477 | Morn3 | chr5 | - | 123035768 | 123047016 | regular |
| ENSMUSG00000073471 | Rsph3a | chr17 | + | 7945652 | 7979824 | regular |
| ENSMUSG00000068039 | Tcp1 | chr17 | + | 12915700 | 12925067 | regular |
| ENSMUSG00000025171 | Ubtd1 | chr19 | + | 41981762 | 42034641 | regular |
| ENSMUSG00000046173 | Pabpc6 | chr17 | - | 9666496 | 9669704 | regular |
| ENSMUSG00000028514 | Usp24 | chr4 | + | 106316212 | 106441322 | regular |
| ENSMUSG00000033671 | Cep350 | chr1 | - | 155844963 | 155973255 | regular |
| ENSMUSG00000067338 | Tuba3b | chr6 | + | 145615756 | 145621479 | regular |
| ENSMUSG00000028287 | 1700009N14 | chr4 | + | 39450294 | 39451776 | regular |
| ENSMUSG00000028277 | Ube2j1 | chr4 | + | 33031415 | 33052363 | regular |
| ENSMUSG00000061130 | Ppm1b | chr17 | + | 84956740 | 85023991 | regular |

| | | | | | | |
|---|---|---|---|---|---|---|
| ENSMUSG00000049659 | Aftph | chr11 | - | 20685083 | 20741589 | regular |
| ENSMUSG00000044340 | Phlpp1 | chr1 | + | 106171751 | 106394250 | regular |
| ENSMUSG00000029320 | 1700016H13I | chr5 | - | 103648038 | 103656010 | regular |
| ENSMUSG00000063015 | Ccni | chr5 | - | 93181932 | 93206495 | regular |
| ENSMUSG00000086211 | Gm12462 | chr4 | + | 44664414 | 44685347 | regular |
| ENSMUSG00000070871 | Ccnyl1 | chr1 | + | 64690454 | 64725644 | regular |
| ENSMUSG00000079710 | Gm3448 | chr17 | - | 14964188 | 15041537 | regular |
| ENSMUSG00000044471 | Lncpint | chr6 | - | 31093558 | 31218474 | regular |
| ENSMUSG00000001829 | Clpb | chr7 | + | 101663632 | 101795506 | regular |
| ENSMUSG00000006299 | Aamp | chr1 | - | 74279839 | 74284738 | regular |
| ENSMUSG00000030077 | Chl1 | chr6 | + | 103510585 | 103750211 | regular |
| ENSMUSG00000038252 | Ncapd2 | chr6 | - | 125168006 | 125191701 | regular |
| ENSMUSG00000026554 | Dcaf8 | chr1 | + | 172148083 | 172197005 | regular |
| ENSMUSG00000032496 | Ltf | chr9 | + | 111019270 | 111042767 | regular |
| ENSMUSG00000020300 | Cpeb4 | chr11 | + | 31872210 | 31935634 | regular |
| ENSMUSG00000026915 | Strbp | chr2 | - | 37483227 | 37703859 | regular |
| ENSMUSG00000036555 | Iqce | chr5 | - | 140661826 | 140702378 | regular |
| ENSMUSG00000009394 | Syn2 | chr6 | + | 115134901 | 115282006 | regular |
| ENSMUSG00000027227 | Sord | chr2 | + | 122234748 | 122265340 | regular |
| ENSMUSG00000022280 | Rnf19a | chr15 | - | 36239932 | 36283147 | regular |
| ENSMUSG00000070639 | Lrrc8b | chr5 | + | 105415774 | 105490074 | regular |
| ENSMUSG00000049866 | Arl4c | chr1 | - | 88673124 | 88702221 | regular |
| ENSMUSG00000030744 | Rps3 | chr7 | - | 99477895 | 99483738 | regular |
| ENSMUSG00000027867 | Spag17 | chr3 | + | 99885405 | 100143322 | regular |
| ENSMUSG00000021981 | Cab39l | chr14 | + | 59440971 | 59585764 | regular |
| ENSMUSG00000030754 | Copb1 | chr7 | - | 114215558 | 114254711 | regular |
| ENSMUSG00000064358 | mt-Co3 | chrM | + | 8606 | 9390 | regular |
| ENSMUSG00000022680 | Pdxdc1 | chr16 | - | 13833147 | 13903131 | regular |
| ENSMUSG00000075524 | 4930407I10R | chr15 | + | 82059150 | 82066538 | regular |
| ENSMUSG00000032349 | Elovl5 | chr9 | + | 77917363 | 77984519 | regular |
| ENSMUSG00000021690 | Jmy | chr13 | - | 93430100 | 93499808 | regular |
| ENSMUSG00000026353 | Ubxn4 | chr1 | + | 128243963 | 128279378 | regular |
| ENSMUSG00000002409 | Dyrk1b | chr7 | + | 28179468 | 28187294 | regular |
| ENSMUSG00000055692 | Tmem191c | chr16 | + | 17276308 | 17283229 | regular |
| ENSMUSG00000020246 | Hcfc2 | chr10 | + | 82696159 | 82742428 | regular |
| ENSMUSG00000096887 | Gm20594 | chr6 | + | 79817915 | 79818364 | regular |
| ENSMUSG00000096141 | Dnah7a | chr1 | - | 53397005 | 53706784 | regular |
| ENSMUSG00000029422 | Rsrc2 | chr5 | - | 123728425 | 123749414 | regular |
| ENSMUSG00000022957 | Itsn1 | chr16 | + | 91729280 | 91920597 | regular |
| ENSMUSG00000041268 | Dmxl2 | chr9 | - | 54365157 | 54501626 | regular |
| ENSMUSG00000012126 | Ubxn11 | chr4 | + | 134102569 | 134127593 | regular |
| ENSMUSG00000023151 | Lrrc69 | chr4 | - | 14623619 | 14796060 | regular |
| ENSMUSG00000040928 | S100pbp | chr4 | - | 129148004 | 129189727 | regular |
| ENSMUSG00000032601 | Prkar2a | chr9 | + | 108689313 | 108750436 | regular |
| ENSMUSG00000035649 | Zcchc7 | chr4 | + | 44755876 | 44932215 | regular |
| ENSMUSG00000038756 | Ttll6 | chr11 | + | 96133785 | 96165451 | regular |

| | | | | | | |
|---|---|---|---|---|---|---|
| ENSMUSG00000028976 | Slc2a5 | chr4 | + | 150119282 | 150144169 | regular |
| ENSMUSG00000030538 | Cib1 | chr7 | - | 80227146 | 80232813 | regular |
| ENSMUSG00000098557 | Kctd12 | chr14 | - | 102976580 | 102982637 | regular |
| ENSMUSG00000040687 | Madd | chr2 | - | 91137359 | 91183837 | regular |
| ENSMUSG00000002718 | Cse1l | chr2 | + | 166906039 | 166946389 | regular |
| ENSMUSG00000112323 | Gm32828 | chr12 | - | 7524001 | 7859766 | regular |
| ENSMUSG00000064341 | mt-Nd1 | chrM | + | 2750 | 3707 | regular |
| ENSMUSG00000032300 | 1700017B05I | chr9 | - | 57253116 | 57262612 | regular |
| ENSMUSG00000022961 | Son | chr16 | + | 91647505 | 91679221 | regular |
| ENSMUSG00000001052 | Sec24b | chr3 | - | 129982758 | 130061553 | regular |
| ENSMUSG00000046192 | Iqub | chr6 | - | 24444864 | 24515067 | regular |
| ENSMUSG00000018570 | 2810408A11I | chr11 | - | 69897351 | 69900987 | regular |
| ENSMUSG00000034160 | Ogt | chrX | + | 101640059 | 101684351 | regular |
| ENSMUSG00000019578 | Ubxn6 | chr17 | - | 56067044 | 56075028 | regular |
| ENSMUSG00000039158 | Akna | chr4 | - | 63367124 | 63403354 | regular |
| ENSMUSG00000061904 | Slc25a3 | chr10 | - | 91116573 | 91124059 | regular |
| ENSMUSG00000109564 | Muc16 | chr9 | - | 18495454 | 18674530 | regular |
| ENSMUSG00000024248 | Cox7a2l | chr17 | - | 83501918 | 83514333 | regular |
| ENSMUSG00000038298 | Pdzk1 | chr3 | + | 96829283 | 96870926 | regular |
| ENSMUSG00000030815 | Phkg2 | chr7 | + | 127573339 | 127583307 | regular |
| ENSMUSG00000026605 | Cenpf | chr1 | - | 189640605 | 189688086 | regular |
| ENSMUSG00000025909 | Sntg1 | chr1 | - | 8361474 | 9299878 | regular |
| ENSMUSG00000062328 | Rpl17 | chr18 | + | 75000476 | 75003381 | regular |
| ENSMUSG00000053896 | 4933409G03 | chr2 | + | 68582412 | 68616387 | regular |
| ENSMUSG00000026578 | Ccdc181 | chr1 | + | 164275584 | 164287847 | regular |
| ENSMUSG00000039105 | Atp6v1g1 | chr4 | + | 63544771 | 63550750 | regular |
| ENSMUSG00000078907 | Fam186b | chr15 | - | 99271017 | 99287180 | regular |
| ENSMUSG00000003226 | Ranbp2 | chr10 | + | 58446919 | 58494356 | regular |
| ENSMUSG00000028433 | Ubap2 | chr4 | - | 41194312 | 41275144 | regular |
| ENSMUSG00000072214 | Sep.05 | chr16 | - | 18621810 | 18629938 | regular |
| ENSMUSG00000032244 | Fem1b | chr9 | - | 62791820 | 62811934 | regular |
| ENSMUSG00000028273 | Pdlim5 | chr3 | - | 142239589 | 142395696 | regular |
| ENSMUSG00000064145 | Arih2 | chr9 | - | 108602941 | 108649386 | regular |
| ENSMUSG00000061315 | Naca | chr10 | + | 128035574 | 128048637 | regular |
| ENSMUSG00000056268 | Dennd1b | chr1 | + | 138963434 | 139178960 | regular |
| ENSMUSG00000028163 | Nfkb1 | chr3 | - | 135584654 | 135691547 | regular |
| ENSMUSG00000055850 | Rnf181 | chr6 | - | 72359713 | 72366956 | regular |
| ENSMUSG00000040412 | 5330417C22I | chr3 | - | 108455693 | 108536536 | regular |
| ENSMUSG00000031986 | Sprtn | chr8 | + | 124897885 | 124906161 | regular |
| ENSMUSG00000029381 | Shroom3 | chr5 | + | 92683434 | 92965318 | regular |
| ENSMUSG00000105703 | Gm43305 | chr14 | - | 54292442 | 54297456 | regular |
| ENSMUSG00000038165 | 1700022A21I | chr5 | + | 24645452 | 24648860 | regular |
| ENSMUSG00000039828 | Wdr70 | chr15 | - | 7873054 | 8099209 | regular |
| ENSMUSG00000039298 | Cdk5rap2 | chr4 | - | 70216855 | 70410443 | regular |
| ENSMUSG00000028936 | Rpl22 | chr4 | + | 152325741 | 152334071 | regular |
| ENSMUSG00000032212 | Sltm | chr9 | + | 70542753 | 70592234 | regular |

| | | | | | | |
|---|---|---|---|---|---|---|
| ENSMUSG00000020900 | Myh10 | chr11 | + | 68691558 | 68816632 | regular |
| ENSMUSG00000033767 | Tmem131l | chr3 | - | 83897654 | 84040175 | regular |
| ENSMUSG00000054280 | Prr14l | chr5 | - | 32789819 | 32854256 | regular |
| ENSMUSG00000053153 | Spag16 | chr1 | + | 69826969 | 70725132 | regular |
| ENSMUSG00000071179 | Serpina16 | chr12 | - | 103668807 | 103676139 | regular |
| ENSMUSG00000032393 | Dpp8 | chr9 | + | 65032413 | 65082651 | regular |
| ENSMUSG00000022021 | Diaph3 | chr14 | - | 86655366 | 87141235 | regular |
| ENSMUSG00000031684 | Slc10a7 | chr8 | + | 78509345 | 78734003 | regular |
| ENSMUSG00000021027 | Ralgapa1 | chr12 | - | 55602895 | 55821167 | regular |
| ENSMUSG00000113255 | AC123954.1 | chr13 | + | 49493317 | 49496589 | regular |
| ENSMUSG00000024500 | Ppp2r2b | chr18 | - | 42637431 | 43059471 | regular |
| ENSMUSG00000024393 | Prrc2a | chr17 | - | 35149075 | 35164897 | regular |
| ENSMUSG00000033319 | Fem1c | chr18 | - | 46501745 | 46525971 | regular |
| ENSMUSG00000029406 | Pitpnm2 | chr5 | - | 124118689 | 124249760 | regular |
| ENSMUSG00000029104 | Htt | chr5 | + | 34761739 | 34912534 | regular |
| ENSMUSG00000026516 | Nvl | chr1 | - | 181087137 | 181144204 | regular |
| ENSMUSG00000026842 | Abl1 | chr2 | + | 31688375 | 31804227 | regular |
| ENSMUSG00000040473 | Cfap69 | chr5 | - | 5579277 | 5664239 | regular |
| ENSMUSG00000037608 | Bclaf1 | chr10 | + | 20312468 | 20344613 | regular |
| ENSMUSG00000079324 | 4932414N04 | chr2 | + | 68656485 | 68748467 | regular |
| ENSMUSG00000041165 | Spem1 | chr11 | - | 69820875 | 69822180 | regular |
| ENSMUSG00000062825 | Actg1 | chr11 | - | 120345689 | 120348542 | regular |
| ENSMUSG00000054452 | Aes | chr10 | + | 81559487 | 81566362 | regular |
| ENSMUSG00000042197 | Zfp451 | chr1 | - | 33761544 | 33814595 | regular |
| ENSMUSG00000005045 | Chd5 | chr4 | + | 152338650 | 152390194 | regular |
| ENSMUSG00000033411 | Ctdspl2 | chr2 | + | 121956000 | 122013642 | regular |
| ENSMUSG00000008730 | Hipk1 | chr3 | - | 103739814 | 103791563 | regular |
| ENSMUSG00000062859 | Tcp11 | chr17 | - | 28066746 | 28080639 | regular |
| ENSMUSG00000037972 | Snn | chr16 | + | 11066297 | 11074983 | regular |
| ENSMUSG00000026670 | Uap1 | chr1 | - | 170141937 | 170174957 | regular |
| ENSMUSG00000062014 | Gmfb | chr14 | - | 46808148 | 46822242 | regular |
| ENSMUSG00000029433 | Diablo | chr5 | - | 123509764 | 123525429 | regular |
| ENSMUSG00000021546 | Hnrnpk | chr13 | - | 58391141 | 58403343 | regular |
| ENSMUSG00000057531 | Dtnbp1 | chr13 | - | 44922074 | 45002147 | regular |
| ENSMUSG00000074884 | Serf2 | chr2 | + | 121449194 | 121458313 | regular |
| ENSMUSG00000021997 | Lrrc63 | chr14 | - | 75084302 | 75130881 | regular |
| ENSMUSG00000024208 | Uqcc2 | chr17 | - | 27122662 | 27133916 | regular |
| ENSMUSG00000030987 | Stim1 | chr7 | + | 102267805 | 102437319 | regular |
| ENSMUSG00000001416 | Cct3 | chr3 | + | 88297115 | 88321767 | regular |
| ENSMUSG00000025477 | Inpp5a | chr7 | + | 139389108 | 139579652 | regular |
| ENSMUSG00000052407 | Ccdc171 | chr4 | + | 83525544 | 83864670 | regular |
| ENSMUSG00000034342 | Cbl | chr9 | - | 44142975 | 44234049 | regular |
| ENSMUSG00000024053 | Emilin2 | chr17 | - | 71252175 | 71310965 | regular |
| ENSMUSG00000045107 | Saysd1 | chr14 | - | 20075635 | 20138534 | regular |
| ENSMUSG00000058704 | Memo1 | chr17 | - | 74200705 | 74294883 | regular |
| ENSMUSG00000025019 | Lcor | chr19 | + | 41482644 | 41562246 | regular |

| | | | | | | |
|---|---|---|---|---|---|---|
| ENSMUSG00000064337 | mt-Rnr1 | chrM | + | 69 | 1024 | regular |
| ENSMUSG00000037110 | Ralgapa2 | chr2 | - | 146239878 | 146512344 | regular |
| ENSMUSG00000047654 | Tssk6 | chr8 | + | 69902183 | 69903518 | regular |
| ENSMUSG00000025781 | Atp5c1 | chr2 | - | 10056015 | 10080510 | regular |
| ENSMUSG00000059796 | Eif4a1 | chr11 | - | 69666935 | 69672423 | regular |
| ENSMUSG00000025651 | Uqcrc1 | chr9 | + | 108936632 | 108949623 | regular |
| ENSMUSG00000025958 | Creb1 | chr1 | + | 64532644 | 64604548 | regular |
| ENSMUSG00000021097 | Clmn | chr12 | - | 104763116 | 104865076 | regular |
| ENSMUSG00000041328 | Pcf11 | chr7 | - | 92643542 | 92669934 | regular |
| ENSMUSG00000041852 | Tcf20 | chr15 | - | 82808628 | 82912134 | regular |
| ENSMUSG00000001280 | Sp1 | chr15 | + | 102406142 | 102436404 | regular |
| ENSMUSG00000027823 | Gmps | chr3 | + | 63976105 | 64022579 | regular |
| ENSMUSG00000025384 | Faap100 | chr11 | - | 120369054 | 120378764 | regular |
| ENSMUSG00000064357 | mt-Atp6 | chrM | + | 7926 | 8607 | regular |
| ENSMUSG00000033102 | Cdc14b | chr13 | - | 64189267 | 64275290 | regular |
| ENSMUSG00000024014 | Pim1 | chr17 | + | 29490811 | 29496111 | regular |
| ENSMUSG00000038015 | Prm2 | chr16 | - | 10791379 | 10796134 | regular |
| ENSMUSG00000031620 | Iqcm | chr8 | + | 75448693 | 75984503 | regular |
| ENSMUSG00000018509 | Cenpv | chr11 | - | 62524945 | 62539261 | regular |
| ENSMUSG00000029203 | Ube2k | chr5 | + | 65537232 | 65598988 | regular |
| ENSMUSG00000040850 | Psme4 | chr11 | + | 30771725 | 30880361 | regular |
| ENSMUSG00000062604 | Srpk2 | chr5 | - | 23503263 | 23684617 | regular |
| ENSMUSG00000053565 | Eif3k | chr7 | - | 28971371 | 28981864 | regular |
| ENSMUSG00000074748 | Atxn7l3b | chr10 | - | 112925429 | 112929001 | regular |
| ENSMUSG00000020156 | Mum1 | chr10 | + | 80226433 | 80243903 | regular |
| ENSMUSG00000031233 | Pgk2 | chr17 | - | 40207017 | 40208609 | regular |
| ENSMUSG00000104529 | Rbakdn | chr5 | - | 143164778 | 143165751 | regular |
| ENSMUSG00000001794 | Capns1 | chr7 | - | 30186935 | 30198811 | regular |
| ENSMUSG00000040026 | Saa3 | chr7 | - | 46711997 | 46715700 | regular |
| ENSMUSG00000031430 | Vsig1 | chrX | + | 140907607 | 140939472 | regular |
| ENSMUSG00000091476 | Gm16432 | chr1 | + | 177983422 | 178172704 | regular |
| ENSMUSG00000058006 | Mdn1 | chr4 | + | 32657118 | 32775217 | regular |
| ENSMUSG00000017119 | Nbr1 | chr11 | + | 101552148 | 101581951 | regular |
| ENSMUSG00000028813 | CK137956 | chr4 | - | 127927591 | 127970951 | regular |
| ENSMUSG00000028392 | Bspry | chr4 | + | 62480052 | 62497298 | regular |
| ENSMUSG00000000439 | Mkrn2 | chr6 | + | 115601901 | 115622624 | regular |
| ENSMUSG00000019139 | Isyna1 | chr8 | + | 70594372 | 70597290 | regular |
| ENSMUSG00000028457 | Atp8b5 | chr4 | + | 43267158 | 43373833 | regular |
| ENSMUSG00000004032 | Gstm5 | chr3 | + | 107895820 | 107898686 | regular |
| ENSMUSG00000042520 | Ubap2l | chr3 | - | 90000139 | 90052628 | regular |
| ENSMUSG00000020867 | Spata20 | chr11 | - | 94478903 | 94486179 | regular |
| ENSMUSG00000048731 | Ggnbp1 | chr17 | + | 26973216 | 27036378 | regular |
| ENSMUSG00000018796 | Acsl1 | chr8 | + | 46471036 | 46536051 | regular |
| ENSMUSG00000050860 | Phospho1 | chr11 | + | 95824498 | 95832140 | regular |
| ENSMUSG00000018196 | Glrx2 | chr1 | + | 143716337 | 143749676 | regular |
| ENSMUSG00000032258 | Lca5 | chr9 | - | 83390292 | 83441127 | regular |

| | | | | | | |
|---|---|---|---|---|---|---|
| ENSMUSG00000024293 | Esco1 | chr18 | - | 10566601 | 10610352 | regular |
| ENSMUSG00000029279 | Brdt | chr5 | + | 107331158 | 107387058 | regular |
| ENSMUSG00000030082 | Sec61a1 | chr6 | - | 88503578 | 88518905 | regular |
| ENSMUSG00000043866 | Taf10 | chr7 | - | 105739392 | 105744361 | regular |
| ENSMUSG00000032536 | Trak1 | chr9 | + | 121297501 | 121474918 | regular |
| ENSMUSG00000000167 | Pih1d2 | chr9 | + | 50617320 | 50625000 | regular |
| ENSMUSG00000022487 | Gtsf1 | chr15 | - | 103402404 | 103430470 | regular |
| ENSMUSG00000052305 | Hbb-bs | chr7 | - | 103826533 | 103828096 | regular |
| ENSMUSG00000064345 | mt-Nd2 | chrM | + | 3913 | 4951 | regular |
| ENSMUSG00000064339 | mt-Rnr2 | chrM | + | 1093 | 2675 | regular |
| ENSMUSG00000023577 | Iqcf3 | chr9 | - | 106503224 | 106561631 | regular |
| ENSMUSG00000038128 | Camk4 | chr18 | + | 32939040 | 33195767 | regular |
| ENSMUSG00000032594 | Ip6k1 | chr9 | + | 108002500 | 108048782 | regular |
| ENSMUSG00000004070 | Hmox2 | chr16 | + | 4726360 | 4766742 | regular |
| ENSMUSG00000022536 | Glyr1 | chr16 | - | 5013905 | 5049910 | regular |
| ENSMUSG00000004127 | Trmt10a | chr3 | + | 138143447 | 138159821 | regular |
| ENSMUSG00000022656 | Nectin3 | chr16 | - | 46387705 | 46498525 | regular |
| ENSMUSG00000024400 | Wdr33 | chr18 | + | 31804056 | 31908987 | regular |
| ENSMUSG00000031452 | 1700029H14 | chr8 | - | 13550732 | 13562461 | regular |
| ENSMUSG00000111028 | Gm5922 | chr9 | - | 120133462 | 120136329 | regular |
| ENSMUSG00000037805 | Rpl10a | chr17 | + | 28328470 | 28331032 | regular |
| ENSMUSG00000029467 | Atp2a2 | chr5 | - | 122453512 | 122502225 | regular |
| ENSMUSG00000028132 | Tmem56 | chr3 | - | 121201760 | 121283098 | regular |
| ENSMUSG00000032512 | Wdr48 | chr9 | + | 119894877 | 119926587 | regular |
| ENSMUSG00000037795 | N4bp2 | chr5 | + | 65763520 | 65830108 | regular |
| ENSMUSG00000010453 | Kansl3 | chr1 | - | 36335729 | 36369181 | regular |
| ENSMUSG00000039735 | Fnbp1l | chr3 | - | 122538718 | 122619715 | regular |
| ENSMUSG00000047446 | Arl4a | chr12 | - | 40005446 | 40038025 | regular |
| ENSMUSG00000027378 | Nphp1 | chr2 | - | 127740731 | 127788897 | regular |
| ENSMUSG00000056508 | 1700001K19 | chr12 | - | 110667687 | 110682619 | regular |
| ENSMUSG00000038773 | Kdm3b | chr18 | + | 34777046 | 34838660 | regular |
| ENSMUSG00000051648 | Kctd19 | chr8 | - | 105382806 | 105413502 | regular |
| ENSMUSG00000049760 | 2410015M20 | chr17 | - | 56607451 | 56609771 | regular |
| ENSMUSG00000055302 | Mrfap1 | chr5 | - | 36794866 | 36796772 | regular |
| ENSMUSG00000053192 | Mllt11 | chr3 | - | 95219134 | 95232171 | regular |
| ENSMUSG00000083012 | Fam220a | chr5 | + | 143548705 | 143564531 | regular |
| ENSMUSG00000026864 | Hspa5 | chr2 | + | 34771969 | 34777547 | regular |
| ENSMUSG00000047940 | Stpg2 | chr3 | + | 139205693 | 139710299 | regular |
| ENSMUSG00000071359 | Tbpl1 | chr10 | - | 22703878 | 22731938 | regular |
| ENSMUSG00000027678 | Ncoa3 | chr2 | + | 165992635 | 166073242 | regular |
| ENSMUSG00000006599 | Gtf2h1 | chr7 | + | 46796102 | 46823800 | regular |
| ENSMUSG00000020593 | Lpin1 | chr12 | - | 16535668 | 16610966 | regular |
| ENSMUSG00000024456 | Diaph1 | chr18 | - | 37843600 | 37935476 | regular |
| ENSMUSG00000026276 | Sep.02 | chr1 | + | 93478963 | 93510260 | regular |
| ENSMUSG00000035530 | Eif1 | chr11 | + | 100319884 | 100322096 | regular |
| ENSMUSG00000020525 | Ppm1d | chr11 | + | 85311243 | 85347066 | regular |

| | | | | | | |
|---|---|---|---|---|---|---|
| ENSMUSG00000109179 | Gm35339 | chr15 | + | 76354471 | 76363795 | regular |
| ENSMUSG00000064363 | mt-Nd4 | chrM | + | 10166 | 11544 | regular |
| ENSMUSG00000059891 | Tsks | chr7 | + | 44943203 | 44958035 | regular |
| ENSMUSG00000099930 | Gm2396 | chr9 | - | 88878993 | 88937342 | regular |
| ENSMUSG00000074582 | Arfgef2 | chr2 | + | 166805587 | 166898052 | regular |
| ENSMUSG00000048478 | Spata33 | chr8 | + | 123212831 | 123222046 | regular |
| ENSMUSG00000043445 | Pgp | chr17 | + | 24470472 | 24471591 | regular |
| ENSMUSG00000055602 | Tcp10b | chr17 | + | 13061103 | 13082481 | regular |
| ENSMUSG00000021081 | Serpina1f | chr12 | - | 103688043 | 103695529 | regular |
| ENSMUSG00000047361 | Gm973 | chr1 | + | 59516263 | 59636417 | regular |
| ENSMUSG00000052384 | Nrros | chr16 | - | 32142784 | 32165594 | regular |
| ENSMUSG00000022999 | Lmbr1l | chr15 | - | 98903920 | 98918231 | regular |
| ENSMUSG00000039703 | Nploc4 | chr11 | - | 120380369 | 120437708 | regular |
| ENSMUSG00000020435 | Osbp2 | chr11 | - | 3703730 | 3863903 | regular |
| ENSMUSG00000030638 | Sh3gl3 | chr7 | + | 82173839 | 82307419 | regular |
| ENSMUSG00000021096 | Ppm1a | chr12 | + | 72757454 | 72799819 | regular |
| ENSMUSG00000078439 | Smim24 | chr10 | + | 81393050 | 81395392 | regular |
| ENSMUSG00000007817 | Zmiz1 | chr14 | + | 25459184 | 25666743 | regular |
| ENSMUSG00000024772 | Ehd1 | chr19 | + | 6276724 | 6300096 | regular |
| ENSMUSG00000033306 | Lpp | chr16 | + | 24393349 | 24992576 | regular |
| ENSMUSG00000029705 | Cux1 | chr5 | - | 136248134 | 136567490 | regular |
| ENSMUSG00000019996 | Map7 | chr10 | + | 20148470 | 20281590 | regular |
| ENSMUSG00000032101 | Ddx25 | chr9 | - | 35536646 | 35558528 | regular |
| ENSMUSG00000037363 | Letm2 | chr8 | - | 25578489 | 25597582 | regular |
| ENSMUSG00000001289 | Pfdn5 | chr15 | + | 102326115 | 102331873 | regular |
| ENSMUSG00000024687 | Osbp | chr19 | + | 11965843 | 11994105 | regular |
| ENSMUSG00000040629 | Mael | chr1 | - | 166201200 | 166238747 | regular |
| ENSMUSG00000055322 | Tns1 | chr1 | - | 73910230 | 74124449 | regular |
| ENSMUSG00000005103 | Wdr1 | chr5 | - | 38526812 | 38563221 | regular |
| ENSMUSG00000030967 | Zranb1 | chr7 | + | 132931141 | 132986391 | regular |
| ENSMUSG00000025372 | Baiap2 | chr11 | + | 119942762 | 120006782 | regular |
| ENSMUSG00000040297 | Suco | chr1 | - | 161816113 | 161876682 | regular |
| ENSMUSG00000025261 | Huwe1 | chrX | + | 151800806 | 151935417 | regular |
| ENSMUSG00000004455 | Ppp1cc | chr5 | + | 122158277 | 122175273 | regular |
| ENSMUSG00000063882 | Uqcrh | chr4 | - | 116066964 | 116075071 | regular |
| ENSMUSG00000003345 | Csnk1g2 | chr10 | + | 80622837 | 80640749 | regular |
| ENSMUSG00000056608 | Chd9 | chr8 | + | 90828351 | 91054516 | regular |
| ENSMUSG00000026611 | Spata17 | chr1 | - | 187044647 | 187215465 | regular |
| ENSMUSG00000015090 | Ptgds | chr2 | - | 25466708 | 25470046 | regular |
| ENSMUSG00000030748 | Il4ra | chr7 | + | 125552119 | 125579474 | regular |
| ENSMUST00000063071 | Kctd19 | chr8 | - | 105395270 | 105396480 | circular |
| ENSMUSG00000064068 | Mtx1 | chr3 | - | 89209080 | 89227088 | regular |
| ENSMUSG00000049606 | Zfp644 | chr5 | - | 106616738 | 106697287 | regular |
| ENSMUSG00000047388 | Atmin | chr8 | + | 116943392 | 116960445 | regular |
| ENSMUSG00000022031 | Elp3 | chr14 | - | 65530448 | 65593075 | regular |
| ENSMUSG00000011832 | Evi5l | chr8 | + | 4166566 | 4211257 | regular |

| | | | | | | |
|---|---|---|---|---|---|---|
| ENSMUSG00000040021 | Lats1 | chr10 | + | 7681213 | 7716460 | regular |
| ENSMUSG00000030057 | Cnbp | chr6 | - | 87842614 | 87851106 | regular |
| ENSMUSG00000033904 | Ccp110 | chr7 | + | 118712551 | 118737024 | regular |
| ENSMUSG00000059343 | Aldoart1 | chr4 | - | 72850582 | 72852634 | regular |
| ENSMUSG00000036411 | 9530077C05I | chr9 | + | 22411512 | 22444681 | regular |
| ENSMUSG00000024991 | Eif3a | chr19 | - | 60761115 | 60790693 | regular |
| ENSMUSG00000028018 | Gstcd | chr3 | - | 132981751 | 133092033 | regular |
| ENSMUSG00000044285 | Gm1821 | chr14 | + | 46084027 | 46085000 | regular |
| ENSMUSG00000028541 | B4galt2 | chr4 | - | 117869259 | 117883487 | regular |
| ENSMUSG00000028158 | Mttp | chr3 | - | 138089853 | 138144968 | regular |
| ENSMUSG00000025162 | Csnk1d | chr11 | - | 120961748 | 120991330 | regular |
| ENSMUSG00000041923 | Nol4 | chr18 | - | 22693180 | 23041653 | regular |
| ENSMUSG00000029766 | 1700012A03I | chr6 | + | 32050245 | 32058921 | regular |
| ENSMUSG00000032232 | Cgnl1 | chr9 | - | 71626508 | 71771602 | regular |
| ENSMUSG00000044948 | Cfap43 | chr19 | - | 47737560 | 47919299 | regular |
| ENSMUSG00000041028 | Ghitm | chr14 | - | 37120091 | 37135272 | regular |
| ENSMUSG00000027680 | Fxr1 | chr3 | + | 34019942 | 34070322 | regular |
| ENSMUSG00000038909 | Kat7 | chr11 | - | 95274258 | 95310246 | regular |
| ENSMUSG00000040560 | Wdr7 | chr18 | + | 63708694 | 63989760 | regular |
| ENSMUSG00000008682 | Rpl10 | chrX | + | 74270811 | 74273135 | regular |
| ENSMUSG00000035629 | Rubcn | chr16 | - | 32821702 | 32877766 | regular |
| ENSMUSG00000030041 | M1ap | chr6 | + | 82946901 | 83030309 | regular |
| ENSMUSG00000002102 | Psmc3 | chr2 | + | 91054008 | 91066369 | regular |
| ENSMUSG00000022488 | Nckap1l | chr15 | + | 103453824 | 103498800 | regular |
| ENSMUSG00000020124 | Usp15 | chr10 | - | 123105005 | 123196995 | regular |
| ENSMUSG00000038685 | Rtel1 | chr2 | + | 181319738 | 181356616 | regular |
| ENSMUSG00000039717 | Ralyl | chr3 | + | 13471654 | 14182287 | regular |
| ENSMUSG00000036006 | Ripor2 | chr13 | + | 24582188 | 24733816 | regular |
| ENSMUSG00000025357 | Dgka | chr10 | - | 128720133 | 128744855 | regular |
| ENSMUSG00000008348 | Ubc | chr5 | - | 125385964 | 125390202 | regular |
| ENSMUSG00000053111 | Fank1 | chr7 | + | 133776860 | 133881532 | regular |
| ENSMUSG00000101225 | 1700008J07F | chr7 | - | 127510437 | 127512869 | regular |
| ENSMUSG00000055839 | Elob | chr17 | - | 23824739 | 23829109 | regular |
| ENSMUSG00000058761 | Rnf169 | chr7 | - | 99920253 | 99980448 | regular |
| ENSMUSG00000034667 | Xpot | chr10 | - | 121587379 | 121626332 | regular |
| ENSMUSG00000042050 | Wdr60 | chr12 | - | 116206261 | 116263022 | regular |
| ENSMUSG00000026226 | Spata3 | chr1 | + | 86016815 | 86029958 | regular |
| ENSMUSG00000037443 | Cep85 | chr4 | - | 134129857 | 134187112 | regular |
| ENSMUSG00000049653 | Spatc1 | chr15 | + | 76268088 | 76292572 | regular |
| ENSMUSG00000028789 | Azin2 | chr4 | - | 128930232 | 128962442 | regular |
| ENSMUSG00000020075 | Ddx21 | chr10 | - | 62580250 | 62602281 | regular |
| ENSMUSG00000051586 | Mical3 | chr6 | - | 120931706 | 121130999 | regular |
| ENSMUSG00000002190 | Clgn | chr8 | + | 83389866 | 83428552 | regular |
| ENSMUSG00000015165 | Hnrnpl | chr7 | + | 28808540 | 28822266 | regular |
| ENSMUSG00000030795 | Fus | chr7 | + | 127967456 | 127985701 | regular |
| ENSMUSG00000062647 | Rpl7a | chr2 | + | 26910763 | 26913318 | regular |

| | | | | | | |
|---|---|---|---|---|---|---|
| ENSMUSG00000022234 | Cct5 | chr15 | - | 31590799 | 31601804 | regular |
| ENSMUSG00000029192 | Tbc1d14 | chr5 | - | 36490603 | 36593276 | regular |
| ENSMUSG00000057841 | Rpl32 | chr6 | - | 115805504 | 115808747 | regular |
| ENSMUSG00000033022 | Cdo1 | chr18 | - | 46713204 | 46728342 | regular |
| ENSMUSG00000006705 | Pknox1 | chr17 | + | 31564748 | 31607684 | regular |
| ENSMUSG00000024908 | Ppp6r3 | chr19 | - | 3454927 | 3575749 | regular |
| ENSMUSG00000010721 | Lmbr1 | chr5 | - | 29229801 | 29378390 | regular |
| ENSMUSG00000032566 | 1700080E11I | chr9 | - | 105143341 | 105145092 | regular |
| ENSMUSG00000029428 | Stx2 | chr5 | - | 128984556 | 129008574 | regular |
| ENSMUSG00000028411 | Aptx | chr4 | - | 40682381 | 40703194 | regular |
| ENSMUSG00000039901 | 9130011E15I | chr19 | - | 45818143 | 45998488 | regular |
| ENSMUSG00000034075 | Zdhhc5 | chr2 | - | 84687969 | 84715180 | regular |
| ENSMUSG00000017832 | Hspb9 | chr11 | + | 100713849 | 100714575 | regular |
| ENSMUSG00000003824 | Syce2 | chr8 | + | 84872110 | 84888221 | regular |
| ENSMUSG00000074746 | Pdzd8 | chr19 | - | 59296083 | 59345780 | regular |
| ENSMUSG00000027104 | Atf2 | chr2 | - | 73816508 | 73892639 | regular |
| ENSMUSG00000031647 | Mfap3l | chr8 | + | 60632826 | 60676729 | regular |
| ENSMUSG00000033826 | Dnah8 | chr17 | + | 30624353 | 30875264 | regular |
| ENSMUSG00000051343 | Rab11fip5 | chr6 | - | 85334961 | 85374634 | regular |
| ENSMUSG00000029814 | Igf2bp3 | chr6 | - | 49085222 | 49214957 | regular |
| ENSMUSG00000054611 | Kdm2a | chr19 | - | 4314418 | 4398285 | regular |
| ENSMUSG00000034303 | Ccdc15 | chr9 | - | 37275834 | 37348432 | regular |
| ENSMUSG00000061665 | Cd2ap | chr17 | - | 42792950 | 42876424 | regular |
| ENSMUSG00000078193 | Gm2000 | chr1 | + | 156366039 | 156366484 | regular |
| ENSMUSG00000027326 | Knl1 | chr2 | + | 119047118 | 119105501 | regular |
| ENSMUSG00000039342 | Ankar | chr1 | - | 72642979 | 72700579 | regular |
| ENSMUSG00000062382 | Ftl1-ps1 | chr13 | + | 74399693 | 74407346 | regular |
| ENSMUSG00000047248 | C2cd3 | chr7 | + | 100372232 | 100470152 | regular |
| ENSMUSG00000109737 | Gm39566 | chr11 | - | 69810964 | 69815298 | regular |
| ENSMUSG00000037012 | Hk1 | chr10 | - | 62268854 | 62379908 | regular |
| ENSMUSG00000007033 | Hspa1l | chr17 | + | 34972702 | 34979285 | regular |
| ENSMUSG00000027331 | Knstrn | chr2 | + | 118814002 | 118853957 | regular |
| ENSMUSG00000015961 | Adss | chr1 | - | 177762961 | 177796511 | regular |
| ENSMUSG00000045004 | Spata21 | chr4 | + | 141088332 | 141112760 | regular |
| ENSMUSG00000028080 | Lrba | chr3 | + | 86224679 | 86782692 | regular |
| ENSMUSG00000111375 | A830010M2( | chr5 | + | 107437996 | 107512556 | regular |
| ENSMUSG00000019132 | BC005537 | chr13 | + | 24801656 | 24816197 | regular |
| ENSMUSG00000028053 | Ash1l | chr3 | + | 88950621 | 89079375 | regular |
| ENSMUSG00000011158 | Brf1 | chr12 | - | 112960084 | 113000704 | regular |
| ENSMUSG00000032518 | Rpsa | chr9 | + | 120127688 | 120132369 | regular |
| ENSMUSG00000025217 | Btrc | chr19 | + | 45363733 | 45530013 | regular |
| ENSMUSG00000026878 | Rab14 | chr2 | - | 35180204 | 35201120 | regular |
| ENSMUSG00000047150 | 1700001C19I | chr17 | - | 47412733 | 47437376 | regular |
| ENSMUSG00000022391 | Rangap1 | chr15 | - | 81704247 | 81729919 | regular |
| ENSMUSG00000035356 | Nfkbiz | chr16 | - | 55811374 | 55838899 | regular |
| ENSMUSG00000035545 | Leng8 | chr7 | + | 4137038 | 4148177 | regular |

| | | | | | | |
|---|---|---|---|---|---|---|
| ENSMUSG00000038057 | Dbil5 | chr11 | + | 76217607 | 76218666 | regular |
| ENSMUSG00000045467 | Ttll13 | chr7 | + | 80246375 | 80260821 | regular |
| ENSMUSG00000028758 | Kif17 | chr4 | + | 138250434 | 138301967 | regular |
| ENSMUSG00000028886 | Eya3 | chr4 | + | 132638986 | 132724765 | regular |
| ENSMUSG00000045078 | Rnf216 | chr5 | - | 142990892 | 143112994 | regular |
| ENSMUSG00000026827 | Gpd2 | chr2 | + | 57237634 | 57370719 | regular |
| ENSMUSG00000027355 | Tmco5 | chr2 | + | 116878690 | 116892494 | regular |
| ENSMUSG00000025757 | Hspa4l | chr3 | + | 40744494 | 40796103 | regular |
| ENSMUSG00000055239 | Kcmf1 | chr6 | - | 72841113 | 72899979 | regular |
| ENSMUSG00000001774 | Chordc1 | chr9 | + | 18292124 | 18317442 | regular |
| ENSMUSG00000020698 | Cct6b | chr11 | - | 82719249 | 82764321 | regular |
| ENSMUSG00000030200 | Bcl2l14 | chr6 | + | 134396317 | 134438736 | regular |
| ENSMUSG00000059273 | Zc3h4 | chr7 | + | 16400909 | 16437696 | regular |
| ENSMUSG00000039285 | Azi2 | chr9 | + | 118040498 | 118069794 | regular |
| ENSMUSG00000006498 | Ptbp1 | chr10 | + | 79854426 | 79864771 | regular |
| ENSMUSG00000020326 | Ccng1 | chr11 | - | 40748551 | 40755311 | regular |
| ENSMUSG00000008668 | Rps18 | chr17 | - | 33951998 | 33956001 | regular |
| ENSMUSG00000038011 | Dnah10 | chr5 | + | 124725084 | 124834308 | regular |
| ENSMUSG00000059854 | Hydin | chr8 | + | 110266976 | 110610253 | regular |
| ENSMUSG00000041161 | Otud3 | chr4 | - | 138895378 | 138913945 | regular |
| ENSMUSG00000031545 | Gpat4 | chr8 | - | 23171264 | 23208346 | regular |
| ENSMUSG00000068240 | Gm11808 | chr4 | - | 3973091 | 3973595 | regular |
| ENSMUSG00000006307 | Kmt2b | chr7 | - | 30568857 | 30588726 | regular |
| ENSMUSG00000014867 | Surf4 | chr2 | - | 26920039 | 26933928 | regular |
| ENSMUSG00000079523 | Tmsb10 | chr6 | - | 72957346 | 72958748 | regular |
| ENSMUSG00000003119 | Cdk12 | chr11 | + | 98203058 | 98278504 | regular |
| ENSMUSG00000041164 | Zmiz2 | chr11 | + | 6389073 | 6406158 | regular |
| ENSMUSG00000023027 | Atf1 | chr15 | + | 100227818 | 100261244 | regular |
| ENSMUSG00000020307 | Cdc34 | chr10 | + | 79682194 | 79688398 | regular |
| ENSMUSG00000053470 | Kdm3a | chr6 | - | 71588971 | 71632990 | regular |
| ENSMUSG00000010461 | Eya4 | chr10 | - | 23102962 | 23350786 | regular |
| ENSMUSG00000001100 | Poldip2 | chr11 | + | 78512192 | 78522736 | regular |
| ENSMUSG00000020823 | Sec14l1 | chr11 | + | 117115167 | 117159268 | regular |
| ENSMUSG00000024767 | Otub1 | chr19 | - | 7198201 | 7206316 | regular |
| ENSMUSG00000033799 | Fam208b | chr13 | - | 3566034 | 3611108 | regular |
| ENSMUSG00000038495 | Otud7b | chr3 | + | 96104526 | 96161129 | regular |
| ENSMUSG00000026940 | Ccdc183 | chr2 | - | 25608634 | 25617678 | regular |
| ENSMUSG00000024286 | Ccny | chr18 | - | 9314043 | 9450150 | regular |
| ENSMUSG00000047454 | Gphn | chr12 | + | 78226378 | 78684772 | regular |
| ENSMUSG00000019977 | Hbs1l | chr10 | + | 21295978 | 21368898 | regular |
| ENSMUSG00000031328 | Flna | chrX | - | 74223460 | 74249820 | regular |
| ENSMUSG00000064367 | mt-Nd5 | chrM | + | 11741 | 13565 | regular |
| ENSMUSG00000025428 | Atp5a1 | chr18 | + | 77773728 | 77782869 | regular |
| ENSMUSG00000028843 | Sh3bgrl3 | chr4 | - | 134127405 | 134128789 | regular |
| ENSMUSG00000022185 | Acin1 | chr14 | - | 54642160 | 54686931 | regular |
| ENSMUSG00000058709 | Egln2 | chr7 | - | 27158657 | 27166802 | regular |

| | | | | | | |
|---|---|---|---|---|---|---|
| ENSMUSG00000024542 | Cep192 | chr18 | + | 67800106 | 67885170 | regular |
| ENSMUSG00000049106 | Dcaf5 | chr12 | - | 80335847 | 80436601 | regular |
| ENSMUSG00000031536 | Polb | chr8 | - | 22628125 | 22653435 | regular |
| ENSMUSG00000021785 | Ngly1 | chr14 | + | 16249279 | 16311926 | regular |
| ENSMUSG00000050122 | Vwa3b | chr1 | + | 37026595 | 37187613 | regular |
| ENSMUSG00000003429 | Rps11 | chr7 | - | 45122378 | 45124439 | regular |
| ENSMUSG00000057286 | St6galnac2 | chr11 | - | 116675302 | 116694868 | regular |
| ENSMUSG00000013822 | Elof1 | chr9 | - | 22112988 | 22117172 | regular |
| ENSMUSG00000019428 | Fkbp8 | chr8 | + | 70527723 | 70535328 | regular |
| ENSMUSG00000030276 | Ttll3 | chr6 | + | 113389259 | 113414587 | regular |
| ENSMUSG00000047866 | Lonp2 | chr8 | + | 86624042 | 86723873 | regular |
| ENSMUSG00000035021 | Baz1a | chr12 | - | 54892988 | 55014348 | regular |
| ENSMUSG00000029086 | Prom1 | chr5 | - | 43993619 | 44102032 | regular |
| ENSMUSG00000056987 | Fam71d | chr12 | + | 78691534 | 78734516 | regular |
| ENSMUSG00000061589 | Dot1l | chr10 | + | 80755205 | 80795461 | regular |
| ENSMUSG00000038900 | Rpl12 | chr2 | + | 32961558 | 32965345 | regular |
| ENSMUSG00000018707 | Dync1h1 | chr12 | + | 110601451 | 110666945 | regular |
| ENSMUSG00000056076 | Eif3b | chr5 | + | 140419327 | 140443360 | regular |
| ENSMUSG00000002835 | Chaf1a | chr17 | + | 56040415 | 56068026 | regular |
| ENSMUSG00000021357 | Exoc2 | chr13 | - | 30813918 | 30974093 | regular |
| ENSMUSG00000038762 | Abcf1 | chr17 | - | 35956818 | 35969761 | regular |
| ENSMUSG00000053510 | Nrd1 | chr4 | + | 109000654 | 109061777 | regular |
| ENSMUSG00000028691 | Prdx1 | chr4 | + | 116685543 | 116700822 | regular |
| ENSMUSG00000021500 | Ddx46 | chr13 | + | 55635026 | 55681256 | regular |
| ENSMUSG00000068758 | Il3ra | chr14 | + | 14346274 | 14356011 | regular |
| ENSMUSG00000036427 | Gpi1 | chr7 | - | 34201329 | 34230336 | regular |
| ENSMUSG00000025762 | Larp1b | chr3 | + | 40950353 | 41040234 | regular |
| ENSMUSG00000037210 | Fam193a | chr5 | + | 34369932 | 34486456 | regular |
| ENSMUSG00000020315 | Sptbn1 | chr11 | - | 30099394 | 30268175 | regular |
| ENSMUSG00000034239 | Gm884 | chr11 | - | 103534576 | 103621140 | regular |
| ENSMUSG00000033499 | Larp4b | chr13 | + | 9093880 | 9174451 | regular |
| ENSMUSG00000037801 | Iqch | chr9 | - | 63421454 | 63602493 | regular |
| ENSMUSG00000025283 | Sat1 | chrX | - | 155213131 | 155216449 | regular |
| ENSMUSG00000050471 | Fam118b | chr9 | - | 35216964 | 35267805 | regular |
| ENSMUSG00000025571 | Tnrc6c | chr11 | + | 117654288 | 117763439 | regular |
| ENSMUSG00000050821 | Fam131a | chr16 | + | 20693281 | 20703048 | regular |
| ENSMUSG00000044792 | Isca1 | chr13 | - | 59755408 | 59769810 | regular |
| ENSMUSG00000037536 | Fbxo34 | chr14 | + | 47450420 | 47531962 | regular |
| ENSMUSG00000030180 | Kdm5a | chr6 | + | 120364123 | 120444574 | regular |
| ENSMUSG00000042688 | Mapk6 | chr9 | - | 75369061 | 75410005 | regular |
| ENSMUSG00000038126 | Mphosph9 | chr5 | - | 124250958 | 124327972 | regular |
| ENSMUSG00000040163 | 1700034J05F | chr6 | - | 146950328 | 146954422 | regular |
| ENSMUSG00000029703 | Lrwd1 | chr5 | - | 136122771 | 136136074 | regular |
| ENSMUSG00000041757 | Plekha6 | chr1 | + | 133164209 | 133303435 | regular |

| values | HS548_p_values | | HS548_p_values | | |
|---|---|---|---|---|---|
| logFC.x | logFC.y | logCPM | LR | PValue | FDR |
| 2.68963826 | 2.70881521 | 2.94195917 | 26.4118124 | 2.76E-07 | 7.98E-04 |
| 2.41757694 | 2.41868457 | 3.4170199 | 22.3136032 | 2.32E-06 | 0.00335068 |
| 1.76721494 | 1.77838945 | 3.94998407 | 17.8655519 | 2.37E-05 | 0.02185564 |
| 1.78007161 | 1.80665663 | 3.14606785 | 17.4046747 | 3.02E-05 | 0.02185564 |
| 2.26941168 | 2.26879426 | 2.7295037 | 16.3676559 | 5.22E-05 | 0.02696523 |
| 1.63041118 | 1.63224284 | 4.11200511 | 16.2365252 | 5.59E-05 | 0.02696523 |
| 1.63854085 | 1.63807344 | 3.62323931 | 14.8951111 | 1.14E-04 | 0.04698911 |
| -1.5947151 | -1.5589465 | 3.97133308 | 14.3116515 | 1.55E-04 | 0.05255138 |
| 1.95425929 | 1.99010579 | 2.52551558 | 14.0399868 | 1.79E-04 | 0.05255138 |
| 1.93176125 | 1.93509962 | 2.76424452 | 13.9270693 | 1.90E-04 | 0.05255138 |
| 1.93310886 | 1.87491196 | 3.45970222 | 13.822474 | 2.01E-04 | 0.05255138 |
| -1.6032091 | -1.5460711 | 3.71403156 | 13.6064576 | 2.25E-04 | 0.05255138 |
| 1.54150899 | 1.56954704 | 3.30462361 | 13.5197682 | 2.36E-04 | 0.05255138 |
| 1.70369145 | 1.68795628 | 2.68761311 | 12.8430437 | 3.39E-04 | 0.07002161 |
| 1.43278668 | 1.42412639 | 3.14513065 | 12.4545031 | 4.17E-04 | 0.07504138 |
| 1.67767096 | 1.66570573 | 3.30160513 | 12.3687806 | 4.37E-04 | 0.07504138 |
| -1.4736207 | -1.4547209 | 4.85534404 | 12.3507482 | 4.41E-04 | 0.07504138 |
| -3.5913136 | -3.5913143 | 21.5729125 | 12.0834751 | 5.09E-04 | 0.07990071 |
| 1.79262703 | 1.75129258 | 3.34410566 | 11.997112 | 5.33E-04 | 0.07990071 |
| 1.52167231 | 1.51806712 | 3.737997 | 11.9306407 | 5.52E-04 | 0.07990071 |
| 1.58044959 | 1.55987254 | 2.63810916 | 11.8044936 | 5.91E-04 | 0.0814288 |
| 1.4586368 | 1.4686727 | 3.30568776 | 11.4555011 | 7.13E-04 | 0.09309362 |
| 1.64456082 | 1.68870441 | 2.64445044 | 11.3863455 | 7.40E-04 | 0.09309362 |
| 1.38036588 | 1.4032879 | 2.92452623 | 11.1271598 | 8.51E-04 | 0.10258357 |
| 1.53554723 | 1.50890554 | 2.78612411 | 11.0447446 | 8.89E-04 | 0.10295574 |
| 1.25235093 | 1.28105998 | 3.53916033 | 10.9529751 | 9.35E-04 | 0.10402087 |
| -1.280086 | -1.2668293 | 3.7357706 | 10.5367225 | 0.00117026 | 0.12543428 |
| 1.58798035 | 1.5968378 | 2.53601136 | 10.4157635 | 0.00124944 | 0.12913874 |
| 1.55773495 | 1.56719321 | 2.70894462 | 9.74993266 | 0.00179329 | 0.17044343 |
| 1.31583089 | 1.33976481 | 3.51334 | 9.67560324 | 0.00186731 | 0.17044343 |
| 1.24262769 | 1.23900012 | 3.53213488 | 9.62586812 | 0.00191856 | 0.17044343 |
| 1.15045058 | 1.11990807 | 3.72686726 | 9.61852144 | 0.00192625 | 0.17044343 |
| 1.17119489 | 1.16589675 | 4.09294817 | 9.60209639 | 0.00194355 | 0.17044343 |
| 1.7784579 | 1.77626094 | 2.6804427 | 9.48526001 | 0.00207129 | 0.17044343 |
| 1.20851437 | 1.20512866 | 3.7899319 | 9.43311203 | 0.00213102 | 0.17044343 |
| -1.2067458 | -1.2261732 | 3.24706754 | 9.40548412 | 0.00216337 | 0.17044343 |
| 1.38380303 | 1.39539994 | 3.10301271 | 9.34506236 | 0.00223587 | 0.17044343 |
| 1.47512897 | 1.4583175 | 2.61354557 | 9.34329549 | 0.00223803 | 0.17044343 |
| -1.2719586 | -1.2512568 | 5.06907773 | 9.1543318 | 0.0024813 | 0.18412516 |
| -1.2300975 | -1.2158318 | 4.21848613 | 9.02917476 | 0.00265704 | 0.19155626 |
| 1.20629205 | 1.20609278 | 3.78453878 | 8.98854785 | 0.00271677 | 0.19155626 |
| 1.21158793 | 1.21802855 | 3.73137141 | 8.88263181 | 0.00287896 | 0.19155626 |
| -2.0381828 | -1.8488871 | 4.18643196 | 8.849016 | 0.00293247 | 0.19155626 |
| -1.5155957 | -1.3890471 | 3.65569401 | 8.84753233 | 0.00293485 | 0.19155626 |

| | | | | | |
|---|---|---|---|---|---|
| 1.61651699 | 1.61626365 | 2.4230318 | 8.79755982 | 0.00301634 | 0.19155626 |
| 1.28955172 | 1.27581842 | 4.39003709 | 8.78043955 | 0.00304478 | 0.19155626 |
| 1.13197548 | 1.12770882 | 4.04607088 | 8.7054826 | 0.00317254 | 0.19534772 |
| 1.30082309 | 1.28813383 | 3.12849389 | 8.63139356 | 0.00330419 | 0.19834381 |
| 1.33628275 | 1.3459924 | 2.6675847 | 8.60181874 | 0.00335827 | 0.19834381 |
| 1.15731187 | 1.16835733 | 3.95566041 | 8.50205264 | 0.00354746 | 0.20067806 |
| 1.82121195 | 1.66825004 | 3.24880643 | 8.49895725 | 0.0035535 | 0.20067806 |
| -1.047819 | -1.0508251 | 3.75577528 | 8.46086545 | 0.00362869 | 0.20067806 |
| 1.60223582 | 1.55029915 | 3.07553659 | 8.42712731 | 0.00369664 | 0.20067806 |
| -1.1541559 | -1.1566615 | 3.36964365 | 8.37128881 | 0.00381195 | 0.20067806 |
| 1.26130338 | 1.26879951 | 3.33503469 | 8.36936513 | 0.00381599 | 0.20067806 |
| 1.44448611 | 1.44320774 | 2.36822527 | 8.28078986 | 0.00400667 | 0.20067806 |
| -4.9076786 | -4.9076228 | 15.0157552 | 8.27512462 | 0.00401919 | 0.20067806 |
| -1.0768942 | -1.0657751 | 4.01493953 | 8.25497223 | 0.00406405 | 0.20067806 |
| 1.16407286 | 1.1640806 | 3.41459953 | 8.24287621 | 0.00409123 | 0.20067806 |
| 1.02538642 | 1.03915001 | 4.07478485 | 8.15496024 | 0.00429435 | 0.20582183 |
| -1.5492783 | -1.4626433 | 3.43869587 | 8.04396517 | 0.00456555 | 0.20582183 |
| 1.12371824 | 1.14632889 | 3.57988409 | 8.03718858 | 0.00458266 | 0.20582183 |
| 1.49713777 | 1.49738676 | 2.47884637 | 8.02806991 | 0.00460579 | 0.20582183 |
| -2.7602582 | -2.6416573 | 5.12058694 | 8.00623509 | 0.00466166 | 0.20582183 |
| 1.13440477 | 1.14314015 | 4.41464303 | 7.99610272 | 0.00468781 | 0.20582183 |
| 1.33999811 | 1.3284501 | 3.31508764 | 7.9828573 | 0.00472224 | 0.20582183 |
| 1.24509937 | 1.25316245 | 2.85255181 | 7.96182086 | 0.00477743 | 0.20582183 |
| 1.26368217 | 1.30592469 | 2.63175833 | 7.93970458 | 0.00483617 | 0.20582183 |
| -1.4715687 | -1.3962822 | 5.20598495 | 7.87385621 | 0.00501546 | 0.20750125 |
| 1.35914687 | 1.37492232 | 2.89929143 | 7.85459411 | 0.00506917 | 0.20750125 |
| 1.54165324 | 1.52924586 | 2.78356839 | 7.77805899 | 0.00528846 | 0.20750125 |
| 1.26723369 | 1.26731925 | 3.22580502 | 7.77247853 | 0.00530482 | 0.20750125 |
| 1.23775616 | 1.23613546 | 3.37718808 | 7.75590066 | 0.00535373 | 0.20750125 |
| 1.06543892 | 1.09253685 | 3.75063426 | 7.75077651 | 0.00536895 | 0.20750125 |
| -1.1187592 | -1.0920964 | 3.83045352 | 7.73857369 | 0.00540535 | 0.20750125 |
| 1.31665897 | 1.35493289 | 3.76160375 | 7.72397189 | 0.00544924 | 0.20750125 |
| 1.07759147 | 1.07825876 | 8.50166865 | 7.68860657 | 0.00555705 | 0.20860739 |
| -1.2038912 | -1.1985395 | 3.04185815 | 7.65175983 | 0.0056717 | 0.20860739 |
| -1.0700721 | -1.0540423 | 4.35148827 | 7.64451041 | 0.00569453 | 0.20860739 |
| -1.0570293 | -1.058464 | 3.76396979 | 7.61438227 | 0.00579046 | 0.20946986 |
| 1.19247101 | 1.18439437 | 3.68599422 | 7.54718987 | 0.00601037 | 0.21099392 |
| -0.9151917 | -0.9303316 | 3.97524063 | 7.52683912 | 0.00607865 | 0.21099392 |
| 1.40085791 | 1.40319253 | 2.78473239 | 7.51293405 | 0.00612575 | 0.21099392 |
| -0.9689275 | -0.9722396 | 3.54651557 | 7.50544542 | 0.00615127 | 0.21099392 |
| -0.9604459 | -0.9664889 | 3.71975285 | 7.49207051 | 0.00619713 | 0.21099392 |
| 1.23544059 | 1.24889541 | 3.22722528 | 7.45316042 | 0.00633252 | 0.21309654 |
| 1.44359528 | 1.32934764 | 3.82512368 | 7.37620318 | 0.00660926 | 0.21985271 |
| -0.9722462 | -0.9732884 | 3.96201637 | 7.28232137 | 0.00696365 | 0.22481375 |
| 1.44387234 | 1.45957878 | 2.53208094 | 7.24424949 | 0.00711284 | 0.22481375 |
| 1.2786212 | 1.24970754 | 2.59193516 | 7.19590844 | 0.007307 | 0.22481375 |

| | | | | | |
|---|---|---|---|---|---|
| -2.2771742 | -2.0010047 | 4.56409954 | 7.19236839 | 0.00732143 | 0.22481375 |
| -2.2771742 | -2.0010047 | 4.56409954 | 7.19236839 | 0.00732143 | 0.22481375 |
| -2.2771742 | -2.0010047 | 4.56409954 | 7.19236839 | 0.00732143 | 0.22481375 |
| 0.99287179 | 0.98222944 | 3.53354573 | 7.17851228 | 0.00737819 | 0.22481375 |
| -1.1441339 | -1.1037532 | 3.58593334 | 7.17810607 | 0.00737986 | 0.22481375 |
| -2.261524 | -1.9751868 | 4.49271182 | 7.14726025 | 0.00750786 | 0.22633067 |
| 1.02622983 | 1.0693614 | 3.47175963 | 7.07074493 | 0.00783527 | 0.22839237 |
| -2.2646651 | -1.9783887 | 4.51742218 | 7.06466401 | 0.00786191 | 0.22839237 |
| -2.2646651 | -1.9783887 | 4.51742218 | 7.06466401 | 0.00786191 | 0.22839237 |
| 1.07649731 | 1.10695255 | 3.13044684 | 7.02333251 | 0.00804544 | 0.22839237 |
| 1.01469572 | 1.01116101 | 4.4348 | 7.01570074 | 0.0080798 | 0.22839237 |
| 1.23171212 | 1.22868397 | 3.24073476 | 6.99835523 | 0.00815846 | 0.22839237 |
| -1.1019867 | -1.0919994 | 3.30162377 | 6.99720048 | 0.00816373 | 0.22839237 |
| 1.27280345 | 1.2522992 | 2.46343819 | 6.98760642 | 0.0082076 | 0.22839237 |
| 1.04435473 | 1.04558538 | 3.86512518 | 6.91142715 | 0.00856466 | 0.23152029 |
| -0.9666401 | -0.9693934 | 3.76901204 | 6.89497894 | 0.00864382 | 0.23152029 |
| -4.2525812 | -4.2545092 | 10.3470361 | 6.89044882 | 0.00866575 | 0.23152029 |
| -0.9803079 | -0.9733419 | 3.89702127 | 6.8770652 | 0.00873088 | 0.23152029 |
| 0.96236822 | 0.95915756 | 4.89621871 | 6.86974221 | 0.00876673 | 0.23152029 |
| -1.388412 | -1.2428224 | 3.68382743 | 6.86297101 | 0.00880001 | 0.23152029 |
| 2.31340189 | 1.89294945 | 3.22454686 | 6.78093966 | 0.00921364 | 0.24021862 |
| 1.12199817 | 1.10777007 | 3.14930967 | 6.73517581 | 0.00945299 | 0.24288101 |
| -2.9302159 | -2.8819128 | 7.84302212 | 6.72940827 | 0.00948361 | 0.24288101 |
| -1.1568928 | -1.1302065 | 5.1556193 | 6.67561946 | 0.00977405 | 0.24812378 |
| 1.11273978 | 1.11393724 | 3.64533461 | 6.64967578 | 0.00991739 | 0.24837404 |
| 0.985179 | 0.99904262 | 3.89409069 | 6.64283127 | 0.00995556 | 0.24837404 |
| -1.830584 | -1.6969997 | 5.79743704 | 6.60533525 | 0.01016737 | 0.25149023 |
| -0.9105442 | -0.9150229 | 5.06299104 | 6.57981934 | 0.01031414 | 0.25279324 |
| -1.1611364 | -1.1081888 | 4.60437155 | 6.56321079 | 0.01041084 | 0.25279324 |
| 1.25076499 | 1.2497708 | 2.76014815 | 6.55107222 | 0.0104821 | 0.25279324 |
| -2.2707292 | -1.9660601 | 4.5893303 | 6.51079259 | 0.01072217 | 0.25314037 |
| -2.2707292 | -1.9660601 | 4.5893303 | 6.51079259 | 0.01072217 | 0.25314037 |
| 0.95591405 | 0.93672292 | 3.34667116 | 6.49643576 | 0.0108091 | 0.25314037 |
| 1.21822631 | 1.21733883 | 3.02892026 | 6.47553287 | 0.01093695 | 0.25314037 |
| 0.90415995 | 0.9211593 | 3.94988906 | 6.46513533 | 0.01100113 | 0.25314037 |
| 1.08179144 | 1.07518046 | 2.88599733 | 6.46187721 | 0.01102132 | 0.25314037 |
| -0.9228437 | -0.9267507 | 4.39269439 | 6.43230274 | 0.01120632 | 0.25536291 |
| -1.0148229 | -1.0121586 | 3.32708967 | 6.37291231 | 0.01158753 | 0.26141197 |
| 1.41986167 | 1.38196318 | 2.82267679 | 6.36300032 | 0.01165243 | 0.26141197 |
| -0.9257376 | -0.8943755 | 3.57637779 | 6.33462739 | 0.0118403 | 0.26199395 |
| 1.10365228 | 1.12712512 | 2.64762168 | 6.32967433 | 0.01187341 | 0.26199395 |
| 1.05425699 | 1.09124222 | 3.47696371 | 6.31827745 | 0.01194997 | 0.26199395 |
| -2.096762 | -1.9275507 | 5.69183628 | 6.29762714 | 0.01208997 | 0.26307047 |
| -0.9947129 | -0.9920157 | 6.71885002 | 6.28128505 | 0.01220196 | 0.26352589 |
| 0.88797745 | 0.90523609 | 4.24728906 | 6.22401172 | 0.01260292 | 0.26597152 |
| -0.9303672 | -0.9479651 | 3.29775481 | 6.22185733 | 0.01261827 | 0.26597152 |

| | | | | | |
|---|---|---|---|---|---|
| -0.9897067 | -0.9773744 | 3.6804734 | 6.20986357 | 0.01270404 | 0.26597152 |
| -2.6494551 | -2.4327861 | 5.11375035 | 6.18904644 | 0.01285434 | 0.26597152 |
| -2.1060749 | -2.0946296 | 8.79029167 | 6.1419688 | 0.01320103 | 0.26597152 |
| -1.0504723 | -1.0301802 | 4.77597561 | 6.14182524 | 0.0132021 | 0.26597152 |
| 1.15205644 | 1.1591131 | 3.01142872 | 6.12238777 | 0.01334804 | 0.26597152 |
| 0.89482445 | 0.89743047 | 5.01078102 | 6.10504562 | 0.01347965 | 0.26597152 |
| 0.98699162 | 0.98689348 | 3.87942802 | 6.10308975 | 0.01349457 | 0.26597152 |
| 1.07273522 | 1.0678161 | 3.60926258 | 6.09921653 | 0.01352418 | 0.26597152 |
| -0.8111218 | -0.8285333 | 3.8242044 | 6.08347505 | 0.01364521 | 0.26597152 |
| 1.22633033 | 1.17730809 | 3.48627099 | 6.070915 | 0.01374258 | 0.26597152 |
| 0.99809386 | 1.02933324 | 3.6672422 | 6.05985234 | 0.01382892 | 0.26597152 |
| 0.96719163 | 0.96688355 | 4.01378262 | 6.04299695 | 0.01396156 | 0.26597152 |
| 0.90575588 | 0.92004079 | 3.85779249 | 6.03588482 | 0.01401792 | 0.26597152 |
| 1.26734176 | 1.25500929 | 2.94743013 | 6.03285034 | 0.01404204 | 0.26597152 |
| 1.28870601 | 1.2777849 | 2.84262381 | 6.02897921 | 0.01407287 | 0.26597152 |
| 1.32282708 | 1.24815822 | 3.39468782 | 6.02406854 | 0.01411208 | 0.26597152 |
| 0.87668626 | 0.87018093 | 6.72288326 | 6.01867672 | 0.01415526 | 0.26597152 |
| -0.8576906 | -0.8696661 | 3.77855906 | 6.01406944 | 0.01419226 | 0.26597152 |
| -0.9719438 | -0.9605668 | 4.90912698 | 5.99497093 | 0.01434672 | 0.26597152 |
| -0.8278204 | -0.856314 | 3.46828016 | 5.9916193 | 0.014374 | 0.26597152 |
| 0.93349835 | 0.97768003 | 3.13330873 | 5.98488286 | 0.014429 | 0.26597152 |
| -0.8436693 | -0.8409651 | 5.4596444 | 5.93324331 | 0.01485787 | 0.27142675 |
| 1.40732819 | 1.41088876 | 2.27835042 | 5.92677286 | 0.01491253 | 0.27142675 |
| -0.7990741 | -0.8051358 | 3.69891822 | 5.88203326 | 0.01529615 | 0.27666908 |
| 0.87855046 | 0.90269994 | 3.74830746 | 5.86849245 | 0.01541425 | 0.27707348 |
| 0.97409647 | 0.9843049 | 3.9436637 | 5.79446191 | 0.01607673 | 0.28719796 |
| 0.97219225 | 0.94493055 | 3.02487096 | 5.77948181 | 0.01621432 | 0.2878788 |
| 1.28729224 | 1.27941987 | 2.83178243 | 5.75104118 | 0.01647889 | 0.29079207 |
| -4.0167028 | -4.03067 | 8.51494677 | 5.71445898 | 0.01682575 | 0.29171664 |
| -0.9033287 | -0.9130816 | 3.41258071 | 5.71002015 | 0.01686835 | 0.29171664 |
| 1.01422192 | 1.00236539 | 4.25105164 | 5.66545733 | 0.01730221 | 0.29171664 |
| 1.43401046 | 1.313414 | 3.12166985 | 5.66180835 | 0.01733824 | 0.29171664 |
| -0.8222067 | -0.8116761 | 4.10747401 | 5.66015916 | 0.01735455 | 0.29171664 |
| -2.0373454 | -2.0222662 | 8.70119976 | 5.65776483 | 0.01737826 | 0.29171664 |
| -0.8832758 | -0.8822514 | 4.4238209 | 5.65523425 | 0.01740335 | 0.29171664 |
| 1.03848142 | 1.0166628 | 4.02004675 | 5.63057687 | 0.01764982 | 0.29171664 |
| 1.23822377 | 1.24272684 | 3.24889243 | 5.62959412 | 0.01765972 | 0.29171664 |
| 1.28298256 | 1.28554835 | 2.4980516 | 5.62697984 | 0.01768608 | 0.29171664 |
| -0.9003845 | -0.8915362 | 3.84296778 | 5.61114811 | 0.01784657 | 0.29171664 |
| 1.00696356 | 1.00215017 | 3.29393953 | 5.61011959 | 0.01785704 | 0.29171664 |
| -1.0091513 | -0.9955146 | 3.2911548 | 5.60821681 | 0.01787645 | 0.29171664 |
| -1.0294927 | -1.0095063 | 4.70911634 | 5.60175566 | 0.01794249 | 0.29171664 |
| 1.08230901 | 1.08876487 | 3.06709724 | 5.57211885 | 0.01824867 | 0.29467193 |
| 1.06474137 | 1.08528809 | 3.04522834 | 5.54598554 | 0.01852312 | 0.29467193 |
| 1.07860208 | 1.07753185 | 3.17051456 | 5.53945498 | 0.01859237 | 0.29467193 |
| 2.48328824 | 1.82569447 | 2.82515572 | 5.53075047 | 0.01868509 | 0.29467193 |

| | | | | | |
|---|---|---|---|---|---|
| -2.4697463 | -2.2637736 | 5.83338473 | 5.51067887 | 0.01890071 | 0.29467193 |
| -2.4610163 | -2.2546874 | 5.83108417 | 5.50891055 | 0.01891983 | 0.29467193 |
| -2.4725996 | -2.265023 | 5.82933922 | 5.47632418 | 0.01927575 | 0.29467193 |
| -2.4698439 | -2.2623271 | 5.82466577 | 5.47577049 | 0.01928186 | 0.29467193 |
| -2.4757735 | -2.2646014 | 5.79204054 | 5.47403365 | 0.01930103 | 0.29467193 |
| -2.4715727 | -2.2638369 | 5.82946348 | 5.46963312 | 0.01934969 | 0.29467193 |
| -2.4715727 | -2.2638369 | 5.82946348 | 5.46963312 | 0.01934969 | 0.29467193 |
| -2.4698243 | -2.2626694 | 5.82973181 | 5.46413598 | 0.01941065 | 0.29467193 |
| -2.4687992 | -2.2614852 | 5.82985604 | 5.45747086 | 0.01948483 | 0.29467193 |
| -2.4646026 | -2.2578988 | 5.82569694 | 5.45165823 | 0.01954976 | 0.29467193 |
| -2.4725975 | -2.260399 | 5.78252013 | 5.4373426 | 0.01971064 | 0.2955574 |
| 1.41303729 | 1.39606673 | 2.59819821 | 5.42636129 | 0.01983497 | 0.29588862 |
| 1.1763331 | 1.24080294 | 3.13238016 | 5.41513113 | 0.01996295 | 0.29605777 |
| 0.93522332 | 0.9530544 | 3.10015121 | 5.40745686 | 0.02005091 | 0.29605777 |
| 1.01519845 | 1.0347655 | 3.16158596 | 5.38914339 | 0.02026242 | 0.29766208 |
| -0.8327189 | -0.8359723 | 4.004573 | 5.3709608 | 0.0204747 | 0.29926147 |
| -1.0295209 | -0.9855728 | 4.02758253 | 5.352298 | 0.02069498 | 0.29954466 |
| -1.1189509 | -1.039618 | 4.45565865 | 5.34618929 | 0.02076761 | 0.29954466 |
| 1.14891785 | 1.11947702 | 3.27610189 | 5.3348723 | 0.02090287 | 0.29954466 |
| 1.02066337 | 1.00863029 | 2.94173379 | 5.33160118 | 0.02094214 | 0.29954466 |
| 0.8934907 | 0.9164296 | 3.35147118 | 5.31832869 | 0.02110224 | 0.29954466 |
| 0.90369407 | 0.91611649 | 3.44034059 | 5.31252179 | 0.02117269 | 0.29954466 |
| -0.9128297 | -0.9093076 | 4.66350052 | 5.3087472 | 0.02121861 | 0.29954466 |
| 0.89666721 | 0.89774191 | 4.54482062 | 5.26980051 | 0.0216985 | 0.30268726 |
| 0.8419691 | 0.86869385 | 3.71072987 | 5.24955156 | 0.02195243 | 0.30268726 |
| 1.22169889 | 1.22166006 | 2.41806015 | 5.24902822 | 0.02195903 | 0.30268726 |
| -0.8579768 | -0.8565847 | 3.80446773 | 5.23479874 | 0.02213937 | 0.30268726 |
| 0.82494837 | 0.81300008 | 3.70082946 | 5.23114871 | 0.02218588 | 0.30268726 |
| -0.8004115 | -0.8037309 | 4.20249279 | 5.22720951 | 0.02223618 | 0.30268726 |
| 1.17697341 | 1.14430664 | 3.08666393 | 5.22705758 | 0.02223812 | 0.30268726 |
| -0.8545793 | -0.8543488 | 4.92765629 | 5.22394557 | 0.02227795 | 0.30268726 |
| 0.86561002 | 0.89707542 | 3.12301539 | 5.16971685 | 0.02298396 | 0.31006585 |
| 0.88843895 | 0.93807775 | 3.36417198 | 5.16215562 | 0.02308423 | 0.31006585 |
| -2.0514049 | -1.8895498 | 5.21498733 | 5.14904943 | 0.0232591 | 0.31006585 |
| 0.89604974 | 0.89654319 | 4.2198993 | 5.14866807 | 0.02326421 | 0.31006585 |
| 1.00915 | 1.05218429 | 2.76009831 | 5.14147255 | 0.02336083 | 0.31006585 |
| 1.14164057 | 1.14138318 | 2.90282729 | 5.13383249 | 0.02346386 | 0.31006585 |
| -0.8098772 | -0.8160846 | 3.67458089 | 5.09662309 | 0.02397247 | 0.31511527 |
| 0.83650683 | 0.83962959 | 4.4485515 | 5.07123152 | 0.0243261 | 0.31511527 |
| 0.98156043 | 0.96633235 | 3.74597849 | 5.06854828 | 0.02436378 | 0.31511527 |
| 0.90970316 | 0.91213611 | 3.53336163 | 5.06752979 | 0.0243781 | 0.31511527 |
| 1.04582797 | 1.0685551 | 2.55649477 | 5.06665534 | 0.0243904 | 0.31511527 |
| -0.9689237 | -0.925293 | 4.36941714 | 5.05231414 | 0.02459307 | 0.31632155 |
| -0.842592 | -0.8473147 | 3.5347814 | 5.04067841 | 0.02475879 | 0.3167059 |
| -0.7797669 | -0.7884786 | 5.03800636 | 5.03487883 | 0.02484182 | 0.3167059 |
| 1.0081145 | 1.02397411 | 2.63639061 | 5.02125956 | 0.02503795 | 0.31780626 |

| | | | | | |
|---|---|---|---|---|---|
| 0.92395157 | 0.9248702 | 2.97245097 | 5.00308569 | 0.02530217 | 0.31975756 |
| -0.8768442 | -0.8643871 | 3.6755551 | 4.98968322 | 0.02549888 | 0.32084238 |
| -2.4730815 | -2.4657176 | 9.60150383 | 4.98177684 | 0.02561566 | 0.32091655 |
| 0.76089133 | 0.75060631 | 6.2413401 | 4.9699009 | 0.02579213 | 0.32173453 |
| 1.08383589 | 1.12326622 | 2.55518417 | 4.9349633 | 0.02631862 | 0.32633496 |
| 1.11332447 | 1.06590782 | 3.35099295 | 4.92234587 | 0.0265115 | 0.32633496 |
| 0.91052329 | 0.91816443 | 3.67601655 | 4.91241439 | 0.02666434 | 0.32633496 |
| -0.9044149 | -0.886444 | 3.85483985 | 4.89865973 | 0.02687755 | 0.32633496 |
| -0.7023192 | -0.7190249 | 3.84963855 | 4.89838387 | 0.02688184 | 0.32633496 |
| -0.897769 | -0.8701754 | 4.01525997 | 4.89444203 | 0.02694328 | 0.32633496 |
| -0.8770203 | -0.8666211 | 3.68961099 | 4.89399463 | 0.02695026 | 0.32633496 |
| 0.83162065 | 0.84369736 | 3.72045 | 4.87400047 | 0.02726423 | 0.32876113 |
| 1.03006894 | 1.02978992 | 3.24187763 | 4.86571926 | 0.02739538 | 0.32897187 |
| -1.1728217 | -1.0636479 | 3.26971054 | 4.84671388 | 0.02769885 | 0.33124167 |
| 1.07043468 | 1.06011669 | 3.09324818 | 4.83837997 | 0.02783303 | 0.3314765 |
| -2.2656004 | -2.1989883 | 7.6077598 | 4.81274373 | 0.02825003 | 0.3350639 |
| -0.8111018 | -0.8114448 | 5.1321663 | 4.79081943 | 0.02861181 | 0.33796973 |
| 0.98117291 | 0.9812648 | 3.23731674 | 4.77214654 | 0.02892374 | 0.33889944 |
| 0.77811771 | 0.79149033 | 4.47887524 | 4.77208786 | 0.02892473 | 0.33889944 |
| 0.83023286 | 0.85266665 | 4.03490526 | 4.72071543 | 0.02980134 | 0.34776243 |
| 0.91269228 | 0.89554949 | 3.99885936 | 4.69907024 | 0.03017894 | 0.35075442 |
| -2.4061577 | -2.3522265 | 7.71888333 | 4.68651149 | 0.03040031 | 0.35146846 |
| -0.7933701 | -0.8014583 | 3.70770375 | 4.68182934 | 0.03048327 | 0.35146846 |
| -0.8232396 | -0.8277762 | 3.42308846 | 4.66453208 | 0.03079181 | 0.35361709 |
| 1.18591702 | 1.18603891 | 2.29166977 | 4.65383921 | 0.03098418 | 0.35441978 |
| -0.9079181 | -0.8806913 | 4.66472173 | 4.64573689 | 0.03113077 | 0.35469467 |
| 0.81993708 | 0.81845093 | 4.18554074 | 4.63287605 | 0.03136494 | 0.3555013 |
| 1.26822278 | 1.14867404 | 3.21038826 | 4.6283798 | 0.03144725 | 0.3555013 |
| 0.95022821 | 0.93096865 | 3.67635466 | 4.61161399 | 0.03175613 | 0.35759628 |
| -0.8082546 | -0.8099009 | 4.29684941 | 4.60024756 | 0.03196734 | 0.35857938 |
| 1.31633717 | 1.29520073 | 2.51429276 | 4.57367076 | 0.03246693 | 0.36277715 |
| 0.83526459 | 0.88246626 | 3.18198196 | 4.55092817 | 0.03290091 | 0.36492564 |
| 0.94594825 | 0.95032841 | 3.38335977 | 4.55038213 | 0.0329114 | 0.36492564 |
| -0.7049924 | -0.7314788 | 3.79304327 | 4.53197087 | 0.03326728 | 0.36680866 |
| 1.22988691 | 1.17308127 | 2.92678956 | 4.52850535 | 0.03333472 | 0.36680866 |
| 1.13466974 | 1.09856224 | 3.08525781 | 4.5157161 | 0.03358482 | 0.36703287 |
| -0.9603679 | -0.9232645 | 3.2123511 | 4.5061364 | 0.03377345 | 0.36703287 |
| -0.726388 | -0.7377645 | 4.0267563 | 4.50336387 | 0.03382824 | 0.36703287 |
| 0.85323221 | 0.85960166 | 4.2093553 | 4.49659507 | 0.03396242 | 0.36703287 |
| -0.8740634 | -0.8771919 | 3.41511429 | 4.49524606 | 0.03398922 | 0.36703287 |
| -0.9131227 | -0.8590587 | 4.13225678 | 4.47164665 | 0.03446176 | 0.37075222 |
| -1.9344842 | -1.9067383 | 8.45328036 | 4.46283757 | 0.03463991 | 0.37128851 |
| 0.78677173 | 0.8090902 | 3.55017532 | 4.41330918 | 0.0356596 | 0.37996474 |
| -0.8052908 | -0.8071217 | 4.14549204 | 4.40590298 | 0.03581476 | 0.37996474 |
| 1.43315978 | 1.23186013 | 3.12961732 | 4.39671104 | 0.03600831 | 0.37996474 |
| 0.87693298 | 0.87647215 | 3.54076685 | 4.39032626 | 0.0361434 | 0.37996474 |

| | | | | | |
|---|---|---|---|---|---|
| -0.6967874 | -0.7079775 | 5.53533152 | 4.38607022 | 0.03623374 | 0.37996474 |
| -0.8011579 | -0.7956169 | 3.6221603 | 4.38591042 | 0.03623713 | 0.37996474 |
| -0.7366057 | -0.7432624 | 4.77708122 | 4.37629323 | 0.03644217 | 0.3800779 |
| 0.86635784 | 0.84929176 | 2.96016867 | 4.37309626 | 0.03651059 | 0.3800779 |
| 0.79666231 | 0.80545701 | 4.4659532 | 4.35934501 | 0.03680645 | 0.38178451 |
| 1.14202753 | 1.17424018 | 2.34082655 | 4.35115453 | 0.03698387 | 0.38225468 |
| 1.02558114 | 1.0261039 | 2.9602572 | 4.32941065 | 0.03745921 | 0.38388111 |
| -0.7340998 | -0.7313954 | 4.1904325 | 4.32407376 | 0.03757686 | 0.38388111 |
| 0.78653349 | 0.81960288 | 3.05097851 | 4.32154279 | 0.03763279 | 0.38388111 |
| 0.87809461 | 0.84115145 | 3.03573921 | 4.31977896 | 0.03767182 | 0.38388111 |
| 0.93281755 | 0.91836716 | 2.8599683 | 4.30672713 | 0.03796193 | 0.38548009 |
| 0.98134949 | 0.99406704 | 2.80113492 | 4.29983828 | 0.038116 | 0.38569125 |
| 0.91738057 | 0.8714789 | 2.88820753 | 4.28493845 | 0.03845147 | 0.38736063 |
| -1.6992999 | -2.1942521 | 5.28296994 | 4.27843898 | 0.03859878 | 0.38736063 |
| 0.80595228 | 0.81631907 | 3.37333577 | 4.2676116 | 0.03884549 | 0.38736063 |
| 0.82080137 | 0.78458822 | 3.2642535 | 4.26544371 | 0.03889509 | 0.38736063 |
| 0.74034576 | 0.77090065 | 3.97249453 | 4.26303724 | 0.03895022 | 0.38736063 |
| -0.7142019 | -0.7376016 | 3.34776877 | 4.25529078 | 0.03912825 | 0.38779847 |
| -0.7961844 | -0.7845158 | 5.57338583 | 4.23364012 | 0.03963036 | 0.39143433 |
| -0.7654812 | -0.7573244 | 4.22135361 | 4.22340373 | 0.03987011 | 0.39246287 |
| 1.18949101 | 1.06017393 | 3.64979247 | 4.20985801 | 0.0401897 | 0.39426778 |
| -2.7699423 | -2.39019 | 5.00027781 | 4.20234135 | 0.04036821 | 0.39468105 |
| -2.7657007 | -2.3856307 | 4.99645737 | 4.19624919 | 0.04051349 | 0.39468391 |
| -2.759048 | -2.3806617 | 4.99717855 | 4.17993224 | 0.04090534 | 0.39468391 |
| -2.759048 | -2.3806617 | 4.99717855 | 4.17993224 | 0.04090534 | 0.39468391 |
| -2.6709718 | -2.2468383 | 4.60236953 | 4.17727227 | 0.04096959 | 0.39468391 |
| -3.104933 | -3.0903803 | 8.19909934 | 4.17393313 | 0.0410504 | 0.39468391 |
| -0.7389315 | -0.7221313 | 3.36684147 | 4.15459067 | 0.04152181 | 0.39783406 |
| -0.6942656 | -0.6843895 | 4.84195509 | 4.14924962 | 0.04165298 | 0.39783406 |
| 0.93135159 | 0.89551132 | 2.69447487 | 4.1363486 | 0.04197161 | 0.3995587 |
| 0.92185452 | 0.91004396 | 2.85251464 | 4.12619334 | 0.04222423 | 0.40064565 |
| 0.87127204 | 0.87790941 | 2.8547468 | 4.1171971 | 0.04244935 | 0.40146545 |
| -0.9496148 | -0.8930843 | 3.54349969 | 4.10798036 | 0.0426813 | 0.40160236 |
| 1.13832732 | 1.083421 | 3.2252268 | 4.10560207 | 0.04274137 | 0.40160236 |
| -0.7587683 | -0.7538473 | 4.4195329 | 4.07213737 | 0.04359608 | 0.40716345 |
| 0.8079099 | 0.76802195 | 3.28269314 | 4.06542186 | 0.04376975 | 0.40716345 |
| -0.8188126 | -0.8197827 | 3.22130975 | 4.0598257 | 0.04391503 | 0.40716345 |
| -2.1119946 | -1.857636 | 4.6596252 | 4.05407494 | 0.04406486 | 0.40716345 |
| 0.99241115 | 0.99991272 | 2.37161157 | 4.03854316 | 0.0444722 | 0.40716345 |
| 0.7578422 | 0.72597274 | 3.59346371 | 4.03822241 | 0.04448065 | 0.40716345 |
| 0.77643666 | 0.77807094 | 3.44097363 | 4.03405854 | 0.04459055 | 0.40716345 |
| 1.19247945 | 1.12338924 | 2.78328432 | 4.03111092 | 0.04466852 | 0.40716345 |
| -0.9500365 | -1.0651983 | 3.49781551 | 4.02329646 | 0.04487592 | 0.40716345 |
| 0.70461426 | 0.6917495 | 4.1334054 | 4.02084486 | 0.0449412 | 0.40716345 |
| 0.75726573 | 0.76547935 | 4.26518073 | 4.01853622 | 0.04500276 | 0.40716345 |
| 0.7643349 | 0.74694848 | 3.66880627 | 4.0178329 | 0.04502153 | 0.40716345 |

| | | | | | |
|---|---|---|---|---|---|
| 0.80709342 | 0.80463157 | 3.97458626 | 4.00048637 | 0.04548714 | 0.41009275 |
| -0.7061666 | -0.7078617 | 5.0273594 | 3.97542195 | 0.04616888 | 0.41490433 |
| -0.7190758 | -0.7487204 | 3.08177769 | 3.96563722 | 0.04643794 | 0.41490433 |
| 0.86442573 | 0.84303194 | 2.8571009 | 3.96160686 | 0.04654925 | 0.41490433 |
| -0.8017441 | -0.7794141 | 3.60349086 | 3.95997839 | 0.0465943 | 0.41490433 |
| 0.75171454 | 0.76592917 | 3.2910132 | 3.93990424 | 0.04715346 | 0.41778943 |
| -0.8271541 | -0.8164566 | 3.35248335 | 3.93655191 | 0.04724753 | 0.41778943 |
| 0.97497086 | 0.99957082 | 2.3948297 | 3.92859894 | 0.04747148 | 0.41778943 |
| -1.1003277 | -1.0278212 | 5.12376841 | 3.92773912 | 0.04749576 | 0.41778943 |
| -0.6571121 | -0.6789344 | 3.92882489 | 3.91123892 | 0.04796423 | 0.41936559 |
| 0.95997126 | 0.95785746 | 3.04856626 | 3.91122026 | 0.04796476 | 0.41936559 |
| 0.79146646 | 0.79799577 | 3.01797306 | 3.9055271 | 0.04812753 | 0.41952129 |
| -0.6620136 | -0.6719552 | 5.63984963 | 3.89010328 | 0.04857144 | 0.42068561 |
| 0.74367022 | 0.74746114 | 4.44984563 | 3.88621335 | 0.04868407 | 0.42068561 |
| 0.70257665 | 0.73037558 | 3.53537077 | 3.8857608 | 0.04869719 | 0.42068561 |
| 0.795783 | 0.81866221 | 2.94874314 | 3.85790184 | 0.04951221 | 0.42645337 |
| -2.6411864 | -2.5572491 | 6.81578068 | 3.85097188 | 0.04971717 | 0.42694808 |
| 1.0938555 | 1.12442962 | 2.97614177 | 3.81898714 | 0.05067486 | 0.43145759 |
| -0.6510467 | -0.6555805 | 4.04731898 | 3.81490748 | 0.05079841 | 0.43145759 |
| -0.765249 | -0.7514446 | 4.6094336 | 3.81011009 | 0.0509441 | 0.43145759 |
| -0.7469387 | -0.7469973 | 3.5665034 | 3.80717762 | 0.05103338 | 0.43145759 |
| -1.3796764 | -1.2173859 | 5.19140955 | 3.80119812 | 0.05121592 | 0.43145759 |
| -0.7548383 | -0.7433309 | 3.8190499 | 3.79924092 | 0.05127582 | 0.43145759 |
| -0.6839362 | -0.6908913 | 3.81808381 | 3.79891174 | 0.05128591 | 0.43145759 |
| 1.2115363 | 1.70180587 | 2.45003493 | 3.76583754 | 0.05230965 | 0.43171036 |
| 0.83550312 | 0.83800219 | 4.50145876 | 3.75421052 | 0.05267467 | 0.43171036 |
| -1.9884409 | -1.7567239 | 4.72658304 | 3.75205109 | 0.05274275 | 0.43171036 |
| 0.93001256 | 0.91555019 | 2.42213121 | 3.74706401 | 0.05290035 | 0.43171036 |
| 0.68537251 | 0.70703711 | 3.43110386 | 3.73968276 | 0.05313453 | 0.43171036 |
| -0.6795121 | -0.6944465 | 3.69809569 | 3.73865581 | 0.0531672 | 0.43171036 |
| -0.6626799 | -0.6648024 | 4.54746981 | 3.73681882 | 0.05322569 | 0.43171036 |
| -0.7781081 | -0.7772918 | 3.40759667 | 3.7358334 | 0.05325709 | 0.43171036 |
| -0.9743539 | -0.9545613 | 6.77776356 | 3.73555234 | 0.05326605 | 0.43171036 |
| -0.7886825 | -0.778946 | 3.5263255 | 3.73533976 | 0.05327283 | 0.43171036 |
| 0.74444767 | 0.77725781 | 3.234623 | 3.734176 | 0.05330996 | 0.43171036 |
| 0.67958231 | 0.68909692 | 3.96540545 | 3.73368465 | 0.05332564 | 0.43171036 |
| -0.6691504 | -0.6989258 | 3.4187116 | 3.73279538 | 0.05335403 | 0.43171036 |
| -0.7185604 | -0.7181805 | 4.23525623 | 3.72908478 | 0.05347269 | 0.43171036 |
| 0.81876214 | 0.82166056 | 2.76656222 | 3.72656047 | 0.05355357 | 0.43171036 |
| 0.84671279 | 0.90117421 | 2.69615275 | 3.71153582 | 0.05403765 | 0.43301669 |
| 0.83082863 | 0.83378941 | 3.37113259 | 3.70218131 | 0.05434138 | 0.43301669 |
| -0.6887516 | -0.6874829 | 3.52021868 | 3.69591592 | 0.05454583 | 0.43301669 |
| 0.89307813 | 0.8646962 | 2.4602423 | 3.6910796 | 0.0547042 | 0.43301669 |
| -0.9204534 | -0.9119573 | 2.97652365 | 3.68328522 | 0.05496046 | 0.43301669 |
| -0.6765967 | -0.6741888 | 3.39248071 | 3.68254036 | 0.05498501 | 0.43301669 |
| -0.6326353 | -0.6373638 | 3.98180198 | 3.6600171 | 0.05573306 | 0.43301669 |

| | | | | | |
|---|---|---|---|---|---|
| -0.6721631 | -0.6790062 | 4.60615363 | 3.65474304 | 0.05590978 | 0.43301669 |
| 0.82452848 | 0.81878816 | 3.01847507 | 3.65227992 | 0.05599251 | 0.43301669 |
| -0.6259953 | -0.6443428 | 3.9460438 | 3.64650467 | 0.05618701 | 0.43301669 |
| 0.90351671 | 0.92514627 | 2.86682814 | 3.64458342 | 0.05625187 | 0.43301669 |
| 0.78757885 | 0.7925934 | 3.58957558 | 3.6413447 | 0.0563614 | 0.43301669 |
| 0.78635001 | 0.78633114 | 3.65228308 | 3.63949381 | 0.05642409 | 0.43301669 |
| 0.68238487 | 0.69646962 | 3.74811294 | 3.63805634 | 0.05647283 | 0.43301669 |
| 1.14029926 | 1.00509822 | 3.45803837 | 3.63160974 | 0.05669196 | 0.43301669 |
| -0.6007076 | -0.624047 | 4.02179041 | 3.63160165 | 0.05669223 | 0.43301669 |
| -0.732012 | -0.7190206 | 3.96346046 | 3.63079475 | 0.05671973 | 0.43301669 |
| -0.6405031 | -0.6521944 | 4.2186642 | 3.6305842 | 0.0567269 | 0.43301669 |
| 1.00178272 | 0.964219 | 3.03834382 | 3.62892405 | 0.05678352 | 0.43301669 |
| 0.96562745 | 0.88996002 | 3.74633492 | 3.62834473 | 0.05680329 | 0.43301669 |
| 0.77718362 | 0.76347815 | 3.31573859 | 3.62674974 | 0.05685775 | 0.43301669 |
| -0.832066 | -0.7896324 | 3.37338266 | 3.61929318 | 0.05711313 | 0.43381997 |
| 0.76621187 | 0.7774469 | 3.47812683 | 3.61127919 | 0.05738896 | 0.43477396 |
| 1.03041735 | 1.01699556 | 2.72557758 | 3.59431089 | 0.05797766 | 0.43808708 |
| -0.8208025 | -0.7774771 | 3.56291805 | 3.58449297 | 0.05832121 | 0.43871401 |
| 0.69519173 | 0.72514073 | 4.19760581 | 3.58327961 | 0.05836382 | 0.43871401 |
| 0.70770242 | 0.72013486 | 4.24527725 | 3.56993654 | 0.05883456 | 0.44000046 |
| -0.745603 | -0.7364228 | 3.82957834 | 3.56981012 | 0.05883904 | 0.44000046 |
| 0.7606202 | 0.76001277 | 3.47259012 | 3.56000814 | 0.05918745 | 0.44146515 |
| -0.809848 | -0.7648763 | 4.67834956 | 3.54747009 | 0.05963631 | 0.44366963 |
| 1.39247858 | 1.25284809 | 2.82642204 | 3.52375946 | 0.06049508 | 0.44890452 |
| -0.6353326 | -0.646886 | 3.99194436 | 3.50523319 | 0.06117523 | 0.45159545 |
| 0.75775947 | 0.7636587 | 3.69244182 | 3.48951235 | 0.06175877 | 0.45159545 |
| 0.78031418 | 0.79053438 | 3.22141561 | 3.48579601 | 0.06189759 | 0.45159545 |
| 0.65742817 | 0.64352347 | 4.26609283 | 3.47431511 | 0.06232852 | 0.45159545 |
| 0.72597563 | 0.7286977 | 4.35249349 | 3.47419435 | 0.06233307 | 0.45159545 |
| 0.71734052 | 0.70627539 | 3.17800475 | 3.47338673 | 0.0623635 | 0.45159545 |
| -0.8976884 | -0.8205328 | 3.47140265 | 3.47279575 | 0.06238579 | 0.45159545 |
| 0.7501708 | 0.7526245 | 3.79668592 | 3.47098948 | 0.06245394 | 0.45159545 |
| -0.6520358 | -0.6764519 | 3.40945294 | 3.46642015 | 0.06262671 | 0.45159545 |
| 1.09776479 | 1.09002109 | 2.19499613 | 3.46538341 | 0.06266598 | 0.45159545 |
| 0.68808161 | 0.67373055 | 3.60777041 | 3.46097822 | 0.06283313 | 0.45159545 |
| -0.6395036 | -0.6269189 | 5.89642218 | 3.45872277 | 0.0629189 | 0.45159545 |
| -0.6629999 | -0.6683512 | 4.6402922 | 3.45156896 | 0.06319176 | 0.45159545 |
| 0.70272052 | 0.70270857 | 4.39031802 | 3.44618441 | 0.06339797 | 0.45159545 |
| 0.64325178 | 0.6483432 | 4.18434229 | 3.44019566 | 0.06362816 | 0.45159545 |
| 0.68135142 | 0.70815534 | 3.78225215 | 3.43553306 | 0.06380799 | 0.45159545 |
| 0.62905885 | 0.61594617 | 3.92218256 | 3.43315332 | 0.06389998 | 0.45159545 |
| 0.85297897 | 0.83742502 | 2.77707255 | 3.42838564 | 0.06408472 | 0.45159545 |
| 1.02826746 | 0.9564878 | 3.43857557 | 3.42804649 | 0.06409788 | 0.45159545 |
| 0.78123075 | 0.74648814 | 3.03825341 | 3.42301791 | 0.06429338 | 0.45159545 |
| -1.6549302 | -1.4540353 | 5.35747395 | 3.41545176 | 0.06458873 | 0.45159545 |
| 0.70439864 | 0.69741845 | 3.38855026 | 3.41466165 | 0.06461966 | 0.45159545 |

| | | | | | |
|---|---|---|---|---|---|
| 0.93507307 | 0.8989054 | 3.16889813 | 3.41458411 | 0.0646227 | 0.45159545 |
| -0.6613967 | -0.6610855 | 4.2065302 | 3.41266206 | 0.064698 | 0.45159545 |
| 0.65122333 | 0.65811429 | 3.86281442 | 3.41111058 | 0.06475885 | 0.45159545 |
| 0.85482524 | 0.84947192 | 2.94192823 | 3.40559504 | 0.06497568 | 0.45201828 |
| -0.6554749 | -0.6524895 | 3.95913073 | 3.39915506 | 0.06522982 | 0.45269811 |
| 0.67673061 | 0.66810151 | 3.36608428 | 3.38971843 | 0.06560415 | 0.45351848 |
| 0.95230227 | 0.95486393 | 2.80789479 | 3.38827897 | 0.06566145 | 0.45351848 |
| 0.88289636 | 0.88220103 | 2.84963602 | 3.37298957 | 0.06627339 | 0.45569036 |
| 0.72557922 | 0.7242733 | 3.3080328 | 3.37255626 | 0.06629082 | 0.45569036 |
| 0.81355358 | 0.85100713 | 2.53965634 | 3.35529247 | 0.06698931 | 0.45780303 |
| 0.70111671 | 0.70250244 | 3.22484416 | 3.35474117 | 0.06701174 | 0.45780303 |
| 0.7655029 | 0.77265315 | 2.86642762 | 3.34803859 | 0.06728514 | 0.45780303 |
| 0.73976228 | 0.74033711 | 3.88496196 | 3.34366924 | 0.06746401 | 0.45780303 |
| -0.6824349 | -0.6842764 | 7.1230052 | 3.3415172 | 0.06755229 | 0.45780303 |
| -0.7588308 | -0.7341221 | 4.56183516 | 3.3396566 | 0.06762872 | 0.45780303 |
| 0.7801794 | 0.78168619 | 3.57371692 | 3.33778992 | 0.06770549 | 0.45780303 |
| -0.9217326 | -0.857736 | 5.42601011 | 3.3248229 | 0.06824136 | 0.4583354 |
| -0.6397351 | -0.6474439 | 3.58634069 | 3.32459572 | 0.06825079 | 0.4583354 |
| -0.6946572 | -0.6904591 | 3.65193858 | 3.32230542 | 0.06834593 | 0.4583354 |
| 0.75508064 | 0.75709948 | 3.58931051 | 3.32057914 | 0.06841772 | 0.4583354 |
| -1.0765735 | -0.9411144 | 3.84248768 | 3.31227603 | 0.06876419 | 0.45837828 |
| 1.04872324 | 1.0816665 | 2.43434181 | 3.30803047 | 0.06894207 | 0.45837828 |
| 0.79602763 | 0.78750224 | 2.9800555 | 3.30379999 | 0.06911981 | 0.45837828 |
| -0.8791278 | -0.8233508 | 5.00033861 | 3.29957747 | 0.06929771 | 0.45837828 |
| -0.7222456 | -0.7175416 | 5.02008146 | 3.29731488 | 0.06939323 | 0.45837828 |
| 0.75426926 | 0.74599602 | 3.17717515 | 3.29616575 | 0.0694418 | 0.45837828 |
| 0.87017323 | 0.88250342 | 2.79423525 | 3.29398673 | 0.069534 | 0.45837828 |
| 0.93336825 | 0.91997746 | 3.35917147 | 3.29027762 | 0.06969124 | 0.45837828 |
| -0.7194708 | -0.6978335 | 4.09521101 | 3.26867014 | 0.07061484 | 0.46339991 |
| 0.80643619 | 0.78103251 | 4.84161966 | 3.26084592 | 0.07095252 | 0.46352009 |
| -0.6862376 | -0.679637 | 3.76442708 | 3.25567043 | 0.07117682 | 0.46352009 |
| 0.62533459 | 0.6251332 | 4.63378164 | 3.25456795 | 0.07122471 | 0.46352009 |
| 0.6419929 | 0.64954699 | 3.47279598 | 3.25210973 | 0.07133159 | 0.46352009 |
| -0.6757408 | -0.6719068 | 3.79100137 | 3.24975839 | 0.07143399 | 0.46352009 |
| 0.76019538 | 0.75159793 | 3.19718583 | 3.24340526 | 0.07171145 | 0.46410614 |
| -1.0692248 | -0.8954533 | 3.58686414 | 3.23609463 | 0.07203215 | 0.46410614 |
| 1.02994205 | 0.92721447 | 3.24908212 | 3.23472414 | 0.07209244 | 0.46410614 |
| 0.72432007 | 0.72146318 | 4.11679092 | 3.22526774 | 0.07250994 | 0.46410614 |
| 0.63225746 | 0.62517787 | 4.08790477 | 3.22308507 | 0.07260667 | 0.46410614 |
| -0.7493745 | -0.7362493 | 3.34660345 | 3.2205994 | 0.072717 | 0.46410614 |
| -0.9140833 | -0.8030751 | 3.78040085 | 3.21169976 | 0.07311349 | 0.46410614 |
| 0.77081057 | 0.74519063 | 2.83289623 | 3.21040629 | 0.07317131 | 0.46410614 |
| -0.6051335 | -0.6127752 | 4.10314755 | 3.21038476 | 0.07317227 | 0.46410614 |
| -0.6300209 | -0.6490539 | 3.49654802 | 3.21035438 | 0.07317363 | 0.46410614 |
| 0.70770891 | 0.71395889 | 3.89419362 | 3.20779122 | 0.07328836 | 0.46410614 |
| -0.6270124 | -0.637224 | 3.77322969 | 3.17630596 | 0.07471346 | 0.47116042 |

| | | | | | |
|---|---|---|---|---|---|
| 0.6723559 | 0.68982192 | 4.42378534 | 3.1759896 | 0.07472793 | 0.47116042 |
| -0.9033388 | -0.8229914 | 3.41323398 | 3.16691916 | 0.07514407 | 0.47275421 |
| 0.64590474 | 0.66750271 | 3.63320688 | 3.15883661 | 0.07551698 | 0.47312174 |
| -0.630223 | -0.6298932 | 3.62579196 | 3.15856702 | 0.07552945 | 0.47312174 |
| -0.7860153 | -0.7401632 | 4.47243187 | 3.14999247 | 0.07592731 | 0.47458671 |
| -0.6719102 | -0.6729663 | 3.53837953 | 3.14515648 | 0.0761527 | 0.47496962 |
| -0.8123602 | -0.7602353 | 4.57376284 | 3.12730649 | 0.07699085 | 0.4776616 |
| 0.89247008 | 0.85264653 | 3.24162013 | 3.12596114 | 0.07705442 | 0.4776616 |
| 0.70149871 | 0.70216948 | 4.03957947 | 3.12410235 | 0.07714235 | 0.4776616 |
| -0.6000544 | -0.5951756 | 3.60731734 | 3.12194534 | 0.07724452 | 0.4776616 |
| 0.74764402 | 0.77131385 | 3.28845407 | 3.10281809 | 0.07815689 | 0.48227302 |
| -0.604862 | -0.6086597 | 4.97944525 | 3.08598195 | 0.07896959 | 0.48449993 |
| -0.6697649 | -0.6745866 | 3.30930774 | 3.08344345 | 0.07909291 | 0.48449993 |
| -2.5566583 | -2.3984458 | 6.17416716 | 3.07939829 | 0.07928985 | 0.48449993 |
| 0.93427153 | 0.94952994 | 3.61492945 | 3.07582075 | 0.07946447 | 0.48449993 |
| 0.81097023 | 0.7621238 | 4.18975657 | 3.0757092 | 0.07946993 | 0.48449993 |
| 0.64072656 | 0.61795202 | 3.82529151 | 3.07363882 | 0.07957118 | 0.48449993 |
| 0.82473476 | 0.83319429 | 2.69714675 | 3.07121908 | 0.07968969 | 0.48449993 |
| 0.94409798 | 0.92937776 | 2.6960736 | 3.06351353 | 0.08006836 | 0.48578164 |
| -0.8753817 | -0.7754156 | 3.97249995 | 3.05585549 | 0.08044662 | 0.48628337 |
| -2.0339988 | -1.6941585 | 4.38607717 | 3.0550378 | 0.08048712 | 0.48628337 |
| -0.6698545 | -0.7025176 | 2.94491762 | 3.04869429 | 0.08080208 | 0.48639716 |
| 0.611634 | 0.63458002 | 4.08623316 | 3.04654233 | 0.08090923 | 0.48639716 |
| 0.68627891 | 0.68626496 | 4.34952493 | 3.04451781 | 0.08101017 | 0.48639716 |
| 0.61325593 | 0.61086244 | 4.06172017 | 3.03311518 | 0.08158125 | 0.48783343 |
| 0.94728861 | 0.90477226 | 2.9897535 | 3.03235886 | 0.08161928 | 0.48783343 |
| -0.6663138 | -0.6648738 | 4.89268469 | 3.02964165 | 0.08175607 | 0.48783343 |
| 0.58833745 | 0.59084487 | 7.05443765 | 3.02392186 | 0.08204483 | 0.48783343 |
| 0.79273145 | 0.79298713 | 3.04833713 | 3.02298537 | 0.08209222 | 0.48783343 |
| -0.7473793 | -0.7176894 | 3.56307105 | 3.00325958 | 0.08309718 | 0.49279351 |
| 0.76333336 | 0.74025904 | 2.89017986 | 2.97468272 | 0.08457671 | 0.50054191 |
| 1.13285104 | 0.96907851 | 3.07108302 | 2.96583526 | 0.08504053 | 0.50225977 |
| -0.6618272 | -0.6830814 | 3.03202496 | 2.94807957 | 0.08597967 | 0.50677225 |
| -0.7450961 | -0.7255266 | 3.49052314 | 2.93288892 | 0.08679206 | 0.51052076 |
| 0.81624577 | 0.79443732 | 3.25099754 | 2.92721976 | 0.08709737 | 0.51054298 |
| -1.696649 | -1.4500742 | 5.05929178 | 2.92626934 | 0.08714866 | 0.51054298 |
| 0.8579617 | 0.80833398 | 3.392389 | 2.91822531 | 0.08758414 | 0.5118594 |
| 1.00786649 | 0.88270063 | 3.30165439 | 2.91559386 | 0.08772711 | 0.5118594 |
| -0.6451508 | -0.6405207 | 4.68913237 | 2.89578006 | 0.08881178 | 0.51714543 |
| 0.62523448 | 0.63960021 | 3.6356802 | 2.88201363 | 0.08957395 | 0.52045492 |
| 0.81744332 | 0.79398757 | 3.22844664 | 2.87692656 | 0.08985738 | 0.52045492 |
| -0.7824527 | -0.7376958 | 3.30959357 | 2.8713335 | 0.09017013 | 0.52045492 |
| 0.62024651 | 0.60238592 | 3.96760796 | 2.86949216 | 0.09027335 | 0.52045492 |
| 0.8151205 | 0.81012524 | 2.67263246 | 2.86737628 | 0.09039212 | 0.52045492 |
| -0.7203975 | -0.7219096 | 2.95522102 | 2.86618322 | 0.09045917 | 0.52045492 |
| -0.6047478 | -0.6280947 | 3.20063148 | 2.85263783 | 0.09122415 | 0.52360749 |

| | | | | | |
|---|---|---|---|---|---|
| 0.64685132 | 0.66517757 | 3.28447908 | 2.85008748 | 0.09136896 | 0.52360749 |
| -0.6379445 | -0.6443306 | 3.45498695 | 2.8434048 | 0.09174961 | 0.52474976 |
| -0.680619 | -0.6729346 | 3.16143966 | 2.81879438 | 0.09316636 | 0.53138909 |
| -0.6435556 | -0.6463938 | 3.4311508 | 2.81687755 | 0.0932777 | 0.53138909 |
| 1.05356978 | 1.04626697 | 2.29688702 | 2.80212365 | 0.09413954 | 0.53524526 |
| 0.67528344 | 0.6894813 | 4.93314761 | 2.78794259 | 0.0949761 | 0.53865349 |
| -0.5704274 | -0.574342 | 4.50104968 | 2.78429832 | 0.09519238 | 0.53865349 |
| -0.5922854 | -0.6032917 | 3.60645437 | 2.78253253 | 0.09529737 | 0.53865349 |
| 0.62839522 | 0.62838475 | 3.7053532 | 2.7739164 | 0.09581148 | 0.53932323 |
| 0.85280168 | 0.81799022 | 3.09887313 | 2.77308448 | 0.09586128 | 0.53932323 |
| -0.5957846 | -0.6010474 | 4.82123403 | 2.77118753 | 0.09597494 | 0.53932323 |
| 0.62624052 | 0.66329558 | 3.2951278 | 2.76544842 | 0.09631969 | 0.53963128 |
| 0.54705079 | 0.56062939 | 4.04547862 | 2.76407021 | 0.09640269 | 0.53963128 |
| 0.73537737 | 0.74232499 | 2.8581789 | 2.74713496 | 0.09742887 | 0.54432267 |
| 0.7702551 | 0.76997195 | 10.2950268 | 2.74252218 | 0.09771044 | 0.54484393 |
| 0.5799858 | 0.58416915 | 3.88665991 | 2.72626275 | 0.09871002 | 0.5461587 |
| 0.57885014 | 0.5941186 | 3.84611926 | 2.72602372 | 0.0987248 | 0.5461587 |
| 0.96861521 | 0.83863963 | 3.40855313 | 2.72523693 | 0.09877346 | 0.5461587 |
| 0.59037999 | 0.57504135 | 4.52213108 | 2.72152706 | 0.09900326 | 0.5461587 |
| 0.79612712 | 0.7920217 | 2.97905646 | 2.7194306 | 0.09913337 | 0.5461587 |
| -0.6538135 | -0.6342535 | 4.44395812 | 2.71732645 | 0.09926415 | 0.5461587 |
| 0.734679 | 0.7275303 | 2.94599141 | 2.71473161 | 0.09942569 | 0.5461587 |
| -0.66412 | -0.6516388 | 3.42757028 | 2.71424544 | 0.09945599 | 0.5461587 |
| 0.92365241 | 0.89198703 | 2.56315439 | 2.71088554 | 0.09966565 | 0.54627348 |
| -0.5516337 | -0.5602988 | 3.56242953 | 2.69454557 | 0.10069219 | 0.55038628 |
| -0.7037338 | -0.6717977 | 3.50044318 | 2.69104543 | 0.10091359 | 0.55038628 |
| -0.5388427 | -0.5408956 | 3.82217325 | 2.6898935 | 0.10098656 | 0.55038628 |
| 0.78533943 | 0.78805538 | 2.78332304 | 2.68248848 | 0.10145707 | 0.55097714 |
| 0.62851305 | 0.62679708 | 3.56867703 | 2.68219535 | 0.10147575 | 0.55097714 |
| 0.65785909 | 0.65551214 | 4.924736 | 2.67521491 | 0.10192157 | 0.55236145 |
| 0.64633891 | 0.6542028 | 3.5292422 | 2.67095338 | 0.10219479 | 0.55280695 |
| 0.70238543 | 0.70696762 | 3.50268083 | 2.66735091 | 0.10242638 | 0.55281279 |
| 0.6413077 | 0.65088375 | 3.16673151 | 2.66499871 | 0.10257791 | 0.55281279 |
| 0.80532513 | 0.80116248 | 2.42491032 | 2.66106151 | 0.10283209 | 0.55293865 |
| -0.5528204 | -0.5531475 | 5.66528622 | 2.65785198 | 0.1030398 | 0.55293865 |
| -0.5672859 | -0.5661302 | 5.25371603 | 2.65577508 | 0.10317445 | 0.55293865 |
| 0.9839252 | 0.89692716 | 3.42133408 | 2.64156155 | 0.10410118 | 0.5550663 |
| -0.5600093 | -0.5681524 | 4.09038604 | 2.63727029 | 0.10438275 | 0.5550663 |
| 0.56265244 | 0.56958445 | 5.55625698 | 2.63646743 | 0.10443553 | 0.5550663 |
| -0.5408229 | -0.9581048 | 4.28249196 | 2.6340147 | 0.10459693 | 0.5550663 |
| -0.5415348 | -0.5549835 | 3.9102519 | 2.63392774 | 0.10460266 | 0.5550663 |
| -0.6284885 | -0.6396725 | 3.21355991 | 2.62916715 | 0.10491673 | 0.5550663 |
| 0.67918011 | 0.67304541 | 3.15984916 | 2.62753828 | 0.10502443 | 0.5550663 |
| 0.57052085 | 0.60324658 | 3.77358758 | 2.62304773 | 0.10532197 | 0.5550663 |
| 0.57620423 | 0.57191708 | 5.60106252 | 2.62181579 | 0.10540376 | 0.5550663 |
| 0.61579122 | 0.62120586 | 4.41956901 | 2.61591494 | 0.10579649 | 0.5550663 |

| | | | | | |
|---|---|---|---|---|---|
| 0.63126977 | 0.73109143 | 4.25374612 | 2.61390641 | 0.10593053 | 0.5550663 |
| 0.57251005 | 0.55942037 | 4.53943622 | 2.60964662 | 0.10621543 | 0.5550663 |
| -0.6378214 | -0.6309703 | 4.10628487 | 2.60529716 | 0.10650719 | 0.5550663 |
| 0.72973236 | 0.69104266 | 3.94454173 | 2.60384285 | 0.10660494 | 0.5550663 |
| -0.6303967 | -0.6274252 | 4.79982525 | 2.60296788 | 0.1066638 | 0.5550663 |
| -0.6983032 | -0.695985 | 3.0547033 | 2.60136256 | 0.10677188 | 0.5550663 |
| 0.62884634 | 0.64674627 | 3.20062569 | 2.59940057 | 0.10690414 | 0.5550663 |
| -0.6822415 | -0.6971852 | 3.30776548 | 2.59549061 | 0.10716825 | 0.5550663 |
| -0.5364699 | -0.5440862 | 4.37116936 | 2.5935945 | 0.10729658 | 0.5550663 |
| 0.61010089 | 0.61891662 | 4.19164668 | 2.59195866 | 0.10740744 | 0.5550663 |
| -0.5350055 | -0.5437901 | 4.89570632 | 2.57321395 | 0.10868671 | 0.55883696 |
| 0.72881854 | 0.72008322 | 3.32504828 | 2.57051477 | 0.1088723 | 0.55883696 |
| -0.6802153 | -0.6574074 | 3.41706653 | 2.57017816 | 0.10889547 | 0.55883696 |
| 0.54082158 | 0.5607067 | 3.52179622 | 2.56887679 | 0.10898509 | 0.55883696 |
| 0.58020113 | 0.56846462 | 4.3085824 | 2.56717246 | 0.10910259 | 0.55883696 |
| 0.55461972 | 0.55946305 | 3.73595679 | 2.56427467 | 0.10930268 | 0.55887271 |
| 0.73329585 | 0.76280632 | 2.90535939 | 2.55170137 | 0.11017555 | 0.56195082 |
| -0.5818021 | -0.5815955 | 3.93938775 | 2.54724049 | 0.11048708 | 0.56195082 |
| 0.9535243 | 0.79667874 | 4.35430571 | 2.54641773 | 0.11054464 | 0.56195082 |
| 1.30807966 | 0.99137127 | 3.43768379 | 2.54280336 | 0.11079791 | 0.56195082 |
| -0.5954808 | -0.5959915 | 3.33439663 | 2.5416968 | 0.11087558 | 0.56195082 |
| 0.69757392 | 0.66658673 | 2.81875832 | 2.52586758 | 0.11199317 | 0.56546662 |
| 0.54670986 | 0.58226785 | 3.51495962 | 2.52469752 | 0.11207627 | 0.56546662 |
| 0.6363813 | 0.63348665 | 5.20206543 | 2.5191105 | 0.11247401 | 0.56546662 |
| 0.54886031 | 0.56013534 | 4.00846731 | 2.51902495 | 0.11248011 | 0.56546662 |
| 0.77621202 | 0.78367413 | 2.63954842 | 2.51194773 | 0.11298621 | 0.56546662 |
| 0.60126464 | 0.57960327 | 3.62118884 | 2.50949267 | 0.11316236 | 0.56546662 |
| 0.6702345 | 0.68415224 | 2.74678264 | 2.50867505 | 0.11322109 | 0.56546662 |
| 0.60828363 | 0.65429684 | 3.01186914 | 2.50583617 | 0.11342527 | 0.56546662 |
| 0.65960084 | 0.66054211 | 3.38478333 | 2.50563393 | 0.11343983 | 0.56546662 |
| -0.5455096 | -0.554355 | 4.10246867 | 2.50447672 | 0.11352319 | 0.56546662 |
| -0.5815755 | -0.5794715 | 5.69691767 | 2.49527435 | 0.11418848 | 0.56631522 |
| 0.62683735 | 0.60290542 | 3.25318057 | 2.49282075 | 0.11436659 | 0.56631522 |
| 0.70389818 | 0.70447491 | 3.02251184 | 2.49227211 | 0.11440645 | 0.56631522 |
| 0.6233581 | 0.63023838 | 3.41379405 | 2.49131147 | 0.1144763 | 0.56631522 |
| 0.85038282 | 0.82210958 | 2.64845067 | 2.4836377 | 0.11503592 | 0.56770047 |
| 0.66097304 | 0.6617654 | 3.42165234 | 2.47962741 | 0.11532957 | 0.56770047 |
| -0.5319367 | -0.5231783 | 4.2826424 | 2.47941963 | 0.11534481 | 0.56770047 |
| 0.57327654 | 0.54715813 | 3.57790465 | 2.46436982 | 0.1164544 | 0.57175641 |
| -0.6352929 | -0.6074756 | 4.02481392 | 2.46220637 | 0.11661487 | 0.57175641 |
| -0.7367039 | -0.6728725 | 3.79964703 | 2.4573979 | 0.11697242 | 0.57175641 |
| 0.68268337 | 0.88890292 | 2.86442042 | 2.45262024 | 0.11732887 | 0.57175641 |
| 0.55239899 | 0.57211139 | 4.00686178 | 2.44980462 | 0.1175395 | 0.57175641 |
| 0.78085661 | 0.77815492 | 2.37347444 | 2.44915711 | 0.117588 | 0.57175641 |
| -0.5638222 | -0.5707138 | 3.57912196 | 2.4474609 | 0.11771514 | 0.57175641 |
| -0.523081 | -0.5296994 | 5.77582301 | 2.44586643 | 0.11783481 | 0.57175641 |

| | | | | | |
|---|---|---|---|---|---|
| 0.60659592 | 0.64792309 | 3.09377249 | 2.44155302 | 0.11815919 | 0.57175641 |
| -0.6666407 | -0.6240604 | 3.77349651 | 2.44119101 | 0.11818646 | 0.57175641 |
| 0.65810711 | 0.66583363 | 4.69447218 | 2.43719415 | 0.118488 | 0.57175641 |
| -0.4948179 | -0.5066129 | 5.49323728 | 2.4363208 | 0.118554 | 0.57175641 |
| 0.763671 | 0.77793901 | 2.7410156 | 2.43389883 | 0.11873725 | 0.57175641 |
| -0.7801047 | -0.7154928 | 3.16732116 | 2.42831806 | 0.11916069 | 0.57224562 |
| 0.63750133 | 0.63316905 | 4.73362075 | 2.42734998 | 0.11923431 | 0.57224562 |
| 0.6375865 | 0.63388967 | 4.00830021 | 2.42386397 | 0.11949985 | 0.57239774 |
| -0.5549727 | -0.5602086 | 4.25373322 | 2.42174474 | 0.11966159 | 0.57239774 |
| 0.76116263 | 0.86522614 | 3.16803471 | 2.40602667 | 0.1208688 | 0.57721834 |
| 0.51292886 | 0.53619468 | 3.72834792 | 2.40018139 | 0.12132118 | 0.57842422 |
| 1.24867177 | 0.9122724 | 3.74902188 | 2.39727115 | 0.12154711 | 0.57854825 |
| 0.64478477 | 0.59704285 | 3.07083272 | 2.38620102 | 0.12241078 | 0.58170247 |
| 0.56590464 | 0.57098662 | 4.61796717 | 2.37745546 | 0.12309791 | 0.58388964 |
| 1.04776239 | 0.83757149 | 3.64728879 | 2.37352965 | 0.12340774 | 0.58388964 |
| -0.575898 | -0.5724517 | 6.22504302 | 2.37266226 | 0.12347632 | 0.58388964 |
| 0.76126211 | 0.7640548 | 2.73633851 | 2.36687329 | 0.12393506 | 0.58446739 |
| 0.60345152 | 0.60119062 | 4.18138486 | 2.36388162 | 0.12417288 | 0.58446739 |
| 0.78945473 | 0.78912412 | 2.59743826 | 2.36323932 | 0.124224 | 0.58446739 |
| 0.7115947 | 0.73106863 | 2.71541207 | 2.35931026 | 0.12453724 | 0.58446739 |
| -0.5055746 | -0.5283264 | 3.70510386 | 2.35479002 | 0.1248987 | 0.58446739 |
| -0.6781887 | -0.6299271 | 4.58106484 | 2.35402986 | 0.1249596 | 0.58446739 |
| 0.53224067 | 0.52163029 | 4.00393037 | 2.35337364 | 0.1250122 | 0.58446739 |
| -0.6282092 | -0.6182429 | 4.67536131 | 2.3447007 | 0.1257097 | 0.58616675 |
| -0.6141923 | -0.6065194 | 4.28712035 | 2.34243954 | 0.12589226 | 0.58616675 |
| -0.5411884 | -0.5710879 | 3.22967071 | 2.34131312 | 0.12598332 | 0.58616675 |
| -1.293038 | -0.9699799 | 3.66314143 | 2.33389178 | 0.12658505 | 0.5863837 |
| -0.5281753 | -0.5334236 | 4.80919988 | 2.33376956 | 0.12659499 | 0.5863837 |
| -0.5506313 | -0.5533102 | 4.37129147 | 2.33324297 | 0.12663781 | 0.5863837 |
| -0.5414394 | -0.5483062 | 4.21524897 | 2.32165168 | 0.12758449 | 0.58864624 |
| 1.7215957 | 1.08992905 | 3.60221782 | 2.32001723 | 0.12771861 | 0.58864624 |
| 0.56182729 | 0.59226283 | 3.35786936 | 2.31979766 | 0.12773664 | 0.58864624 |
| 0.72389937 | 0.71317288 | 3.21037589 | 2.31473678 | 0.12815299 | 0.58962601 |
| -1.0107133 | -0.806295 | 4.08701418 | 2.30328347 | 0.12910083 | 0.59113645 |
| 0.71771568 | 0.70652294 | 2.72759197 | 2.30278196 | 0.12914251 | 0.59113645 |
| -0.6163902 | -0.6286715 | 2.92340993 | 2.30186748 | 0.12921856 | 0.59113645 |
| 0.65612412 | 0.63927954 | 4.83043003 | 2.30056268 | 0.12932714 | 0.59113645 |
| 0.67521532 | 0.65572036 | 2.67442737 | 2.29845696 | 0.12950259 | 0.59113645 |
| 0.80240586 | 0.77988429 | 2.92576435 | 2.28507102 | 0.13062416 | 0.59472827 |
| 0.74713492 | 0.73400194 | 2.38396715 | 2.28243489 | 0.13084631 | 0.59472827 |
| 0.57798768 | 0.57426653 | 4.40727049 | 2.28081987 | 0.13098262 | 0.59472827 |
| -0.4826639 | -0.5039996 | 3.93233168 | 2.27929472 | 0.13111148 | 0.59472827 |
| -0.6228206 | -0.6046116 | 4.01478818 | 2.2753388 | 0.1314464 | 0.59531435 |
| 0.62297167 | 0.6188688 | 3.71789307 | 2.26757065 | 0.13210684 | 0.59711127 |
| 0.52845972 | 0.55549128 | 3.42780204 | 2.26582445 | 0.13225581 | 0.59711127 |
| -0.5756599 | -0.5686857 | 3.71249742 | 2.26328467 | 0.13247282 | 0.5971594 |

| | | | | | |
|---|---|---|---|---|---|
| -0.6992082 | -0.6547706 | 3.17674625 | 2.25904495 | 0.13283596 | 0.59767905 |
| -0.5346784 | -0.5400101 | 4.09566252 | 2.25685459 | 0.133024 | 0.59767905 |
| 0.61768382 | 0.61994391 | 3.09588821 | 2.25471862 | 0.13320767 | 0.59767905 |
| 0.6284837 | 0.61239747 | 3.97883113 | 2.24604574 | 0.13395632 | 0.59947863 |
| -0.648194 | -0.633429 | 3.43094338 | 2.24334742 | 0.1341902 | 0.59947863 |
| -0.5565425 | -0.5631585 | 3.44105616 | 2.24288669 | 0.13423018 | 0.59947863 |
| 0.59151822 | 0.59017202 | 3.46540159 | 2.23942978 | 0.13453058 | 0.5998355 |
| -0.4825339 | -0.4875397 | 5.80004994 | 2.23658276 | 0.13477855 | 0.5998355 |
| -0.6854827 | -0.6162594 | 4.25693673 | 2.23482503 | 0.1349319 | 0.5998355 |
| -0.5797809 | -0.5691958 | 4.49528423 | 2.23154662 | 0.13521844 | 0.60018735 |
| -0.6336289 | -0.6000787 | 3.5474785 | 2.22660031 | 0.13565204 | 0.60118989 |
| 0.60212324 | 0.59756998 | 4.1803791 | 2.21851018 | 0.13636459 | 0.60342375 |
| -0.5817389 | -0.6023705 | 2.9947049 | 2.21090084 | 0.13703863 | 0.60548059 |
| -1.1446594 | -0.9330666 | 4.56857666 | 2.20710455 | 0.1373763 | 0.60604727 |
| -1.0989118 | -0.8208014 | 3.74163546 | 2.19991684 | 0.13801818 | 0.60795224 |
| -0.5693539 | -0.5535658 | 3.77015229 | 2.19672497 | 0.1383043 | 0.60828671 |
| -0.5633612 | -0.561447 | 3.42244486 | 2.19318577 | 0.13862234 | 0.60876032 |
| 0.53817822 | 0.52960399 | 4.6249136 | 2.18863451 | 0.13903252 | 0.60963655 |
| 0.70834182 | 0.70478505 | 2.57259477 | 2.18255181 | 0.13958286 | 0.61112374 |
| 0.56544448 | 0.56352957 | 5.78909075 | 2.17819696 | 0.13997837 | 0.6119296 |
| -0.6015354 | -0.5996308 | 3.75055375 | 2.1697483 | 0.14074927 | 0.61319614 |
| -1.5065041 | -1.1963839 | 4.64144347 | 2.16677334 | 0.14102186 | 0.61319614 |
| 0.65061126 | 0.66081425 | 3.05036567 | 2.16632355 | 0.14106312 | 0.61319614 |
| -0.5818107 | -0.5771849 | 4.40616397 | 2.1657514 | 0.14111563 | 0.61319614 |
| 0.51126156 | 0.51951312 | 5.20559804 | 2.1613274 | 0.14152239 | 0.61328378 |
| 0.7735297 | 0.73160572 | 3.13393624 | 2.16092307 | 0.14155963 | 0.61328378 |
| 0.50791148 | 0.49922944 | 5.16944288 | 2.15783703 | 0.14184423 | 0.61359823 |
| 0.54500295 | 0.58818687 | 3.20633745 | 2.15496367 | 0.1421098 | 0.61382952 |
| -0.5112931 | -0.4985538 | 4.44753159 | 2.14906228 | 0.142657 | 0.61495244 |
| 0.51709922 | 0.52918477 | 5.02987489 | 2.14758051 | 0.14279476 | 0.61495244 |
| 0.63927546 | 0.60887248 | 3.04333318 | 2.14204403 | 0.14331083 | 0.61625787 |
| 0.54024253 | 0.52276324 | 4.65499614 | 2.13494096 | 0.14397601 | 0.61819964 |
| 0.69845039 | 0.68910519 | 2.61258366 | 2.12584604 | 0.14483279 | 0.62070215 |
| 0.83066121 | 0.7343488 | 3.31179465 | 2.12420717 | 0.14498779 | 0.62070215 |
| -0.4918945 | -0.4919174 | 5.49400058 | 2.11959221 | 0.14542526 | 0.62140072 |
| -0.6006934 | -0.5922692 | 3.54226894 | 2.1140015 | 0.14595722 | 0.62140072 |
| 0.73488924 | 0.68395093 | 3.63729129 | 2.11125371 | 0.14621948 | 0.62140072 |
| 0.83825066 | 0.71878374 | 3.42974118 | 2.1101739 | 0.14632269 | 0.62140072 |
| -0.5053923 | -0.5261064 | 3.40363475 | 2.10924541 | 0.1464115 | 0.62140072 |
| -0.4951349 | -0.5018064 | 5.28180023 | 2.10895503 | 0.14643928 | 0.62140072 |
| -0.5849732 | -0.5839634 | 3.33626336 | 2.10352443 | 0.14696005 | 0.6219336 |
| 1.09605857 | 0.894965 | 3.97096843 | 2.10316414 | 0.14699467 | 0.6219336 |
| 1.20003676 | 0.90443235 | 3.02233829 | 2.10006365 | 0.14729301 | 0.62228608 |
| 0.51901984 | 0.5216766 | 3.90113928 | 2.09327414 | 0.1479487 | 0.62275778 |
| -0.4670742 | -0.4811683 | 5.4560584 | 2.0914872 | 0.14812182 | 0.62275778 |
| -0.4702204 | -0.4736568 | 5.52512014 | 2.08949984 | 0.14831462 | 0.62275778 |

| | | | | | |
|---|---|---|---|---|---|
| 0.60320334 | 0.60227716 | 3.31064627 | 2.08645777 | 0.1486103 | 0.62275778 |
| 0.64042562 | 0.65250325 | 3.15680855 | 2.08613977 | 0.14864124 | 0.62275778 |
| -0.5069046 | -0.499057 | 3.68000331 | 2.08557942 | 0.14869579 | 0.62275778 |
| 0.52249091 | 0.50569643 | 3.64677502 | 2.07557082 | 0.14967393 | 0.62332871 |
| -0.5935548 | -0.6086603 | 3.01608743 | 2.0737664 | 0.14985105 | 0.62332871 |
| 0.51099083 | 0.50979961 | 4.55199586 | 2.0700604 | 0.15021557 | 0.62332871 |
| -0.51604 | -0.5181908 | 6.37465389 | 2.06982957 | 0.15023831 | 0.62332871 |
| -0.5029244 | -0.5009875 | 3.89830714 | 2.06685574 | 0.15053159 | 0.62332871 |
| 0.64184271 | 0.65197445 | 3.09480939 | 2.06607019 | 0.15060917 | 0.62332871 |
| -0.5290438 | -0.5243802 | 3.32465001 | 2.06321039 | 0.15089198 | 0.62332871 |
| -0.4776282 | -0.490998 | 4.46040583 | 2.06019886 | 0.15119044 | 0.62332871 |
| 0.53854772 | 0.54413126 | 3.6151758 | 2.05678755 | 0.15152933 | 0.62332871 |
| -0.5341651 | -0.5344289 | 4.72207854 | 2.05580774 | 0.15162683 | 0.62332871 |
| -1.5788075 | -1.3713191 | 5.5107765 | 2.05405448 | 0.15180146 | 0.62332871 |
| 0.52319873 | 0.52886751 | 3.66701709 | 2.05254175 | 0.15195232 | 0.62332871 |
| -0.5265336 | -0.5253094 | 3.9367038 | 2.04827652 | 0.1523786 | 0.62332871 |
| -0.5082027 | -0.5115147 | 4.72663443 | 2.04770922 | 0.1524354 | 0.62332871 |
| -0.5955433 | -0.5717575 | 3.69538008 | 2.04727178 | 0.15247921 | 0.62332871 |
| -0.6345981 | -0.6419484 | 2.75340089 | 2.0463668 | 0.1525699 | 0.62332871 |
| -0.5248511 | -0.5248534 | 5.83761727 | 2.04334644 | 0.15287301 | 0.62332871 |
| -0.521237 | -0.523237 | 3.78304854 | 2.04297296 | 0.15291054 | 0.62332871 |
| -0.6614469 | -0.6237249 | 4.47282106 | 2.04224047 | 0.15298418 | 0.62332871 |
| 0.52352584 | 0.53147527 | 5.05111318 | 2.0393835 | 0.15327175 | 0.62332871 |
| 0.64247235 | 0.61630079 | 4.07808918 | 2.03275834 | 0.153941 | 0.62332871 |
| -0.5748556 | -0.5498353 | 3.08919821 | 2.03135689 | 0.15408299 | 0.62332871 |
| -0.4985273 | -0.5045077 | 3.92028515 | 2.03117917 | 0.15410101 | 0.62332871 |
| -0.4706836 | -0.475763 | 5.05294162 | 2.03042829 | 0.15417715 | 0.62332871 |
| 0.68383453 | 0.65069129 | 4.63112563 | 2.02701877 | 0.15452346 | 0.62332871 |
| 0.71068574 | 0.70207627 | 2.87297605 | 2.02657867 | 0.15456823 | 0.62332871 |
| -0.5110092 | -0.5151263 | 4.51193303 | 2.02579914 | 0.15464755 | 0.62332871 |
| -0.5973703 | -0.5728817 | 3.64671838 | 2.01866551 | 0.15537564 | 0.62539235 |
| 0.69078086 | 0.6912109 | 2.7715607 | 2.01165703 | 0.15609475 | 0.62599017 |
| 0.75131141 | 0.81107187 | 3.01932391 | 2.00982791 | 0.15628305 | 0.62599017 |
| -0.5425576 | -0.5489831 | 3.2354396 | 2.00837715 | 0.15643258 | 0.62599017 |
| 0.61002411 | 0.64697592 | 3.14481817 | 2.00782923 | 0.15648909 | 0.62599017 |
| 0.65247809 | 0.68990966 | 7.13124209 | 2.00669952 | 0.1566057 | 0.62599017 |
| 0.75608918 | 0.87564407 | 3.53317569 | 2.00048266 | 0.15724913 | 0.62769514 |
| 0.58144219 | 0.57728619 | 3.39377361 | 1.99593989 | 0.15772119 | 0.62871231 |
| 0.49038552 | 0.49674305 | 4.14853581 | 1.99378826 | 0.15794535 | 0.6287398 |
| -0.503936 | -0.5172922 | 3.28737384 | 1.98979939 | 0.15836186 | 0.62953189 |
| 0.4991707 | 0.50118842 | 4.82163663 | 1.98602087 | 0.15875756 | 0.63023918 |
| 0.53562188 | 0.51348733 | 3.46575101 | 1.98224208 | 0.15915441 | 0.63094912 |
| 0.52542378 | 0.50747495 | 4.69655825 | 1.97295595 | 0.16013445 | 0.63295086 |
| -0.496034 | -0.5027677 | 4.04326903 | 1.9728173 | 0.16014914 | 0.63295086 |
| -0.4928717 | -0.4955404 | 5.56683504 | 1.97124784 | 0.16031547 | 0.63295086 |
| 0.56272031 | 0.56280606 | 4.01195476 | 1.96738849 | 0.16072533 | 0.63370451 |

| | | | | | |
|---|---|---|---|---|---|
| -0.5138747 | -0.5114608 | 5.89601588 | 1.96121549 | 0.16138338 | 0.63543335 |
| 0.60830352 | 0.58927649 | 2.91411642 | 1.95641831 | 0.16189689 | 0.63575515 |
| 0.71306642 | 0.69876075 | 2.24851643 | 1.95634762 | 0.16190447 | 0.63575515 |
| -0.5303512 | -0.5250365 | 3.53894582 | 1.95162571 | 0.16241177 | 0.63655326 |
| 0.66910444 | 0.6644289 | 3.02008998 | 1.95036396 | 0.16254764 | 0.63655326 |
| 0.51706809 | 0.52234258 | 3.64023371 | 1.9464797 | 0.1629667 | 0.6365881 |
| -0.8049411 | -0.6615091 | 3.32593236 | 1.94620423 | 0.16299647 | 0.6365881 |
| 0.53028905 | 0.57104622 | 3.01919516 | 1.9411104 | 0.16354802 | 0.63788136 |
| -0.93442 | -0.7428868 | 3.63695342 | 1.93722666 | 0.16396998 | 0.63866636 |
| 0.68426978 | 0.64882647 | 3.31054369 | 1.93473504 | 0.16424134 | 0.63886348 |
| -0.5226545 | -0.5121939 | 3.56638853 | 1.93037734 | 0.16471717 | 0.63910936 |
| -0.5373196 | -0.5236091 | 4.17677848 | 1.93011167 | 0.16474623 | 0.63910936 |
| 0.54355396 | 0.54612794 | 3.27902415 | 1.92747875 | 0.16503455 | 0.63937079 |
| -0.5174854 | -0.5174456 | 5.17525388 | 1.91880354 | 0.16598862 | 0.6422073 |
| 0.47949813 | 0.49271493 | 4.30025857 | 1.91281143 | 0.16665129 | 0.64389372 |
| -0.8963598 | -0.892564 | 8.02003181 | 1.91084446 | 0.16686948 | 0.64389372 |
| -0.5691219 | -0.561688 | 3.44870271 | 1.90322765 | 0.16771749 | 0.6453926 |
| 0.98860118 | 0.79219729 | 3.03742866 | 1.90317735 | 0.1677231 | 0.6453926 |
| -0.5743334 | -0.5842795 | 2.97649773 | 1.90135293 | 0.16792696 | 0.6453926 |
| 0.52710339 | 0.55400604 | 3.54639154 | 1.89934616 | 0.16815152 | 0.64539855 |
| 0.5215218 | 0.51778499 | 4.25542255 | 1.88432874 | 0.16984297 | 0.64732285 |
| 0.52393924 | 0.54356659 | 3.60161741 | 1.8804991 | 0.17027743 | 0.64732285 |
| -0.4739509 | -0.4899519 | 3.7681844 | 1.88038968 | 0.17028986 | 0.64732285 |
| 0.95035742 | 0.72644607 | 3.74379077 | 1.88029881 | 0.17030019 | 0.64732285 |
| -0.5063651 | -0.5109807 | 3.44583886 | 1.87958546 | 0.17038127 | 0.64732285 |
| 0.46727512 | 0.46081846 | 6.27022743 | 1.87936904 | 0.17040588 | 0.64732285 |
| -0.5375147 | -0.5297788 | 4.23799756 | 1.87862281 | 0.17049076 | 0.64732285 |
| -0.4939701 | -0.481147 | 4.55533585 | 1.87799737 | 0.17056193 | 0.64732285 |
| 0.47152695 | 0.47705697 | 5.57634966 | 1.87708368 | 0.17066598 | 0.64732285 |
| 0.49371989 | 0.51039026 | 4.53151719 | 1.87373563 | 0.17104785 | 0.64792208 |
| -0.4562305 | -0.4729329 | 3.65636128 | 1.87170853 | 0.17127953 | 0.64795157 |
| -0.5152727 | -0.5056267 | 4.33634356 | 1.86541096 | 0.17200159 | 0.64951525 |
| -0.5772311 | -0.5724468 | 3.37637812 | 1.86419219 | 0.17214174 | 0.64951525 |
| -0.5438482 | -0.5281047 | 3.84376067 | 1.8602268 | 0.17259863 | 0.64968663 |
| 0.54985886 | 0.53688037 | 3.35048376 | 1.8595677 | 0.1726747 | 0.64968663 |
| -0.4709466 | -0.4804091 | 3.43827249 | 1.85795815 | 0.17286064 | 0.64968663 |
| -0.5209697 | -0.5217585 | 3.54546169 | 1.85579073 | 0.1731114 | 0.64978519 |
| -0.5263027 | -0.5308678 | 5.02496314 | 1.85046058 | 0.17372984 | 0.65126184 |
| -0.5599347 | -0.5358428 | 5.94629064 | 1.84648881 | 0.17419232 | 0.65215081 |
| -0.5654741 | -0.5777428 | 2.81330474 | 1.83985086 | 0.17496843 | 0.65294396 |
| 0.8595711 | 0.80731247 | 2.43673252 | 1.83971623 | 0.17498421 | 0.65294396 |
| 0.67279164 | 0.66926104 | 2.79904967 | 1.83889057 | 0.17508103 | 0.65294396 |
| 0.57288862 | 0.56701033 | 5.52886407 | 1.82875771 | 0.17627435 | 0.65433407 |
| -0.4295803 | -0.4588915 | 3.83861967 | 1.82802041 | 0.17636154 | 0.65433407 |
| 0.4751113 | 0.48037022 | 3.74727858 | 1.82642858 | 0.17654997 | 0.65433407 |
| 1.03239233 | 0.82266509 | 2.84816054 | 1.82426334 | 0.17680664 | 0.65433407 |

| | | | | | |
|---|---|---|---|---|---|
| 0.58015683 | 0.56293693 | 2.81677112 | 1.82335852 | 0.17691402 | 0.65433407 |
| 0.50510475 | 0.52838995 | 3.26975401 | 1.82272378 | 0.1769894 | 0.65433407 |
| 0.51087094 | 0.51950224 | 4.95100685 | 1.81884169 | 0.17745122 | 0.65433407 |
| 0.66654312 | 0.65760114 | 3.0563331 | 1.81811357 | 0.17753799 | 0.65433407 |
| -0.7100391 | -0.6316805 | 3.06326536 | 1.81756787 | 0.17760305 | 0.65433407 |
| 0.48122667 | 0.48679178 | 4.60543143 | 1.81663133 | 0.17771478 | 0.65433407 |
| 0.47825094 | 0.48358515 | 3.89380707 | 1.81224305 | 0.17823939 | 0.65535807 |
| 0.53714442 | 0.55385544 | 3.1161858 | 1.8105205 | 0.1784458 | 0.65535807 |
| 1.15303253 | 0.82666602 | 3.12625526 | 1.80480386 | 0.17913282 | 0.65703418 |
| 0.57889779 | 0.57730377 | 2.99778834 | 1.80295007 | 0.17935625 | 0.65703418 |
| 0.57501568 | 0.5835017 | 3.19359158 | 1.79657778 | 0.18012678 | 0.65902262 |
| 1.17465949 | 0.87095457 | 2.75263104 | 1.79276606 | 0.18058951 | 0.65935888 |
| -0.7522974 | -0.6320409 | 4.43915797 | 1.79099666 | 0.18080478 | 0.65935888 |
| -0.5097834 | -0.5204168 | 3.114504 | 1.78828568 | 0.18113518 | 0.65935888 |
| 0.7208641 | 0.71718459 | 2.46379655 | 1.78495464 | 0.1815421 | 0.65935888 |
| -0.5155942 | -0.5087757 | 3.69428295 | 1.7816389 | 0.18194821 | 0.65935888 |
| -0.5029693 | -0.5035105 | 3.10385984 | 1.78097402 | 0.18202977 | 0.65935888 |
| 0.4892908 | 0.45985855 | 4.63479368 | 1.77972856 | 0.18218266 | 0.65935888 |
| -0.4941258 | -0.4731785 | 4.4269747 | 1.77896292 | 0.18227673 | 0.65935888 |
| -0.5144643 | -0.5073245 | 4.68690089 | 1.77801465 | 0.1823933 | 0.65935888 |
| 0.56856338 | 0.57039537 | 3.60921174 | 1.77717136 | 0.18249705 | 0.65935888 |
| 0.61794666 | 0.6347795 | 2.87879853 | 1.77375902 | 0.18291755 | 0.66005412 |
| -0.4384283 | -0.4456995 | 4.17734161 | 1.76716687 | 0.18373309 | 0.66217131 |
| 0.52248267 | 0.51648037 | 3.57466112 | 1.76462262 | 0.18404897 | 0.66248474 |
| 0.54783028 | 0.57225879 | 2.84146439 | 1.76113467 | 0.18448305 | 0.66249485 |
| 0.48275391 | 0.49664999 | 3.77145672 | 1.76061942 | 0.18454727 | 0.66249485 |
| 0.42655716 | 0.45349095 | 3.83235804 | 1.75489463 | 0.18526259 | 0.66249485 |
| 0.50263202 | 0.50559142 | 4.61732357 | 1.75393014 | 0.18538342 | 0.66249485 |
| -0.5119971 | -0.5068716 | 4.58056989 | 1.7520358 | 0.18562101 | 0.66249485 |
| -0.4502508 | -0.4795512 | 3.37045592 | 1.75090908 | 0.18576249 | 0.66249485 |
| 0.49069129 | 0.51621352 | 3.2099725 | 1.75083411 | 0.18577191 | 0.66249485 |
| 0.44870357 | 0.44931258 | 3.72143032 | 1.74725236 | 0.18622252 | 0.66249485 |
| 1.20408739 | 0.85352304 | 3.03600773 | 1.74686602 | 0.1862712 | 0.66249485 |
| 0.57495535 | 0.57872577 | 3.26510655 | 1.74631243 | 0.18634098 | 0.66249485 |
| -0.5328522 | -0.5535442 | 2.97762532 | 1.74392262 | 0.18664257 | 0.66275289 |
| 0.59686327 | 0.59769157 | 3.10487478 | 1.74023183 | 0.18710945 | 0.66345439 |
| 0.46099835 | 0.4719144 | 3.42178085 | 1.73873942 | 0.18729863 | 0.66345439 |
| 0.64555705 | 0.64978328 | 2.33050015 | 1.73374382 | 0.18793348 | 0.6646014 |
| -0.609773 | -0.5503615 | 3.47259712 | 1.73048332 | 0.18834918 | 0.6646014 |
| 0.77336008 | 0.64800637 | 3.5571135 | 1.7296409 | 0.18845676 | 0.6646014 |
| 0.48163991 | 0.46504336 | 4.02382605 | 1.72898142 | 0.18854103 | 0.6646014 |
| 0.4592441 | 0.44677687 | 3.70231518 | 1.72240156 | 0.1893842 | 0.66561442 |
| -0.4937168 | -0.5027826 | 3.47942139 | 1.72038616 | 0.18964335 | 0.66561442 |
| -0.5535115 | -0.5235427 | 3.62827097 | 1.71528288 | 0.19030138 | 0.66561442 |
| -0.4713436 | -0.4727725 | 7.73487399 | 1.71519829 | 0.19031231 | 0.66561442 |
| -0.5088059 | -0.5206794 | 3.25555625 | 1.71415666 | 0.19044695 | 0.66561442 |

| | | | | | |
|---|---|---|---|---|---|
| -0.4675695 | -0.4473388 | 3.75199313 | 1.71278989 | 0.1906238 | 0.66561442 |
| 1.75263029 | 1.08979744 | 3.01899094 | 1.71231644 | 0.1906851 | 0.66561442 |
| 0.61953776 | 0.61923723 | 2.75174788 | 1.71129541 | 0.19081738 | 0.66561442 |
| 0.4600012 | 0.4477235 | 4.34678148 | 1.70698615 | 0.19137687 | 0.66561442 |
| -0.4736542 | -0.474022 | 3.9691466 | 1.7061201 | 0.19148954 | 0.66561442 |
| -0.488858 | -0.4864777 | 5.04267264 | 1.70546207 | 0.1915752 | 0.66561442 |
| -0.4549884 | -0.46522 | 3.70022612 | 1.70506404 | 0.19162704 | 0.66561442 |
| 0.50608731 | 0.51079011 | 3.71065719 | 1.70288971 | 0.1919105 | 0.66561442 |
| 0.48219333 | 0.47679545 | 4.07373465 | 1.69846535 | 0.1924888 | 0.66561442 |
| -1.2032675 | -1.1762099 | 8.61625509 | 1.69198277 | 0.1933398 | 0.66561442 |
| -0.729922 | -0.6704502 | 3.08634099 | 1.69018896 | 0.19357606 | 0.66561442 |
| -0.5755098 | -0.5691702 | 4.27671283 | 1.68808855 | 0.19385313 | 0.66561442 |
| -0.5914688 | -0.5907999 | 9.23045157 | 1.68758703 | 0.19391936 | 0.66561442 |
| -0.5715158 | -0.5448139 | 3.24711799 | 1.6874176 | 0.19394174 | 0.66561442 |
| 0.54221321 | 0.56054229 | 4.32998788 | 1.68571121 | 0.19416729 | 0.66561442 |
| -0.4455657 | -0.4569641 | 4.32689988 | 1.68484012 | 0.19428255 | 0.66561442 |
| 0.52449711 | 0.52144923 | 4.24455524 | 1.68133955 | 0.19474655 | 0.66561442 |
| 0.54021214 | 0.57939367 | 4.22110155 | 1.68096267 | 0.19479659 | 0.66561442 |
| -0.4535741 | -0.4564624 | 4.20833709 | 1.67733885 | 0.19527843 | 0.66561442 |
| 0.46430433 | 0.4678307 | 4.54242369 | 1.67652326 | 0.19538707 | 0.66561442 |
| -0.4183067 | -0.4250651 | 5.38179543 | 1.67628322 | 0.19541906 | 0.66561442 |
| -0.5481104 | -0.5453678 | 3.38790848 | 1.67389401 | 0.19573777 | 0.66561442 |
| 0.57769507 | 0.56836714 | 2.91843658 | 1.67378285 | 0.19575262 | 0.66561442 |
| 0.45286835 | 0.45701473 | 4.61865943 | 1.67194217 | 0.1959986 | 0.66561442 |
| -0.4582364 | -0.455911 | 4.24837248 | 1.67178974 | 0.19601898 | 0.66561442 |
| 0.57868531 | 0.5682235 | 2.68712449 | 1.67090774 | 0.19613699 | 0.66561442 |
| -0.4349616 | -0.4407679 | 5.87783325 | 1.67052406 | 0.19618836 | 0.66561442 |
| -0.4539828 | -0.4559988 | 5.20869526 | 1.66014809 | 0.19758333 | 0.66785461 |
| 1.77331477 | 1.24738486 | 4.84071017 | 1.65736368 | 0.19795965 | 0.66785461 |
| -0.5391288 | -0.5408452 | 3.13450233 | 1.65646461 | 0.19808134 | 0.66785461 |
| 0.61164156 | 0.58665348 | 3.933026 | 1.65643775 | 0.19808498 | 0.66785461 |
| 0.49960622 | 0.51976825 | 2.89604464 | 1.6560838 | 0.19813291 | 0.66785461 |
| -0.5391403 | -0.524951 | 3.38317271 | 1.65534299 | 0.19823328 | 0.66785461 |
| 0.54829614 | 0.55696608 | 3.32825966 | 1.64999929 | 0.19895903 | 0.66952026 |
| 0.57199015 | 0.55825275 | 2.98292095 | 1.64068845 | 0.20023104 | 0.67285678 |
| -0.5988946 | -0.5540903 | 3.52801491 | 1.63934377 | 0.20041553 | 0.67285678 |
| 0.43494277 | 0.43591508 | 3.83872102 | 1.63210365 | 0.20141233 | 0.67395923 |
| 0.54504518 | 0.54419089 | 3.44666007 | 1.63151765 | 0.20149327 | 0.67395923 |
| 0.47357572 | 0.47488154 | 4.73641634 | 1.63150313 | 0.20149527 | 0.67395923 |
| -0.4727944 | -0.4734356 | 4.95331983 | 1.63019971 | 0.20167543 | 0.67395923 |
| -0.4359511 | -0.4582567 | 3.66479588 | 1.62509157 | 0.2023833 | 0.67554472 |
| -0.4377853 | -0.4511848 | 4.66481428 | 1.6177806 | 0.20340153 | 0.67706138 |
| 0.97544074 | 0.76352145 | 4.01281375 | 1.61710865 | 0.20349542 | 0.67706138 |
| -0.5236416 | -0.5233219 | 3.22645649 | 1.61679306 | 0.20353953 | 0.67706138 |
| 0.43232796 | 0.43004989 | 5.61798524 | 1.60631927 | 0.20500997 | 0.68116976 |
| 0.44778264 | 0.46705664 | 4.67665464 | 1.60017737 | 0.20587808 | 0.68170234 |

| | | | | | |
|---|---|---|---|---|---|
| -0.4912755 | -0.6368986 | 3.20191865 | 1.59870025 | 0.2060875 | 0.68170234 |
| 0.44243716 | 0.44306347 | 4.37009467 | 1.59851716 | 0.20611348 | 0.68170234 |
| 0.66381628 | 0.64158906 | 2.90099988 | 1.59778944 | 0.20621676 | 0.68170234 |
| 0.67622359 | 0.57585967 | 4.27836264 | 1.59617391 | 0.20644627 | 0.68170234 |
| 0.45804571 | 0.46805712 | 4.12505765 | 1.59449619 | 0.20668493 | 0.68170234 |
| 0.51755979 | 0.52649711 | 3.01422759 | 1.59355357 | 0.20681916 | 0.68170234 |
| -0.4030073 | -0.4264341 | 3.75416748 | 1.58839944 | 0.20755496 | 0.68334931 |
| -0.4238337 | -0.4186781 | 4.30595237 | 1.58627282 | 0.20785945 | 0.68354296 |
| -0.6252469 | -0.5680686 | 3.12946363 | 1.58469187 | 0.20808616 | 0.68354296 |
| -0.5979099 | -0.550713 | 3.23179181 | 1.58011591 | 0.208744 | 0.6843228 |
| -0.4109477 | -0.4203459 | 5.26167139 | 1.57953522 | 0.20882765 | 0.6843228 |
| -0.4228626 | -0.430855 | 4.51892314 | 1.57740953 | 0.20913422 | 0.6843228 |
| -0.4169977 | -0.4378208 | 4.49750895 | 1.57647335 | 0.20926941 | 0.6843228 |
| -0.560766 | -0.5394956 | 3.4075201 | 1.57442757 | 0.20956519 | 0.68451653 |
| 0.49809051 | 0.4974843 | 4.42302576 | 1.56846779 | 0.21042967 | 0.68494621 |
| -0.4332001 | -0.429053 | 6.01604643 | 1.56760311 | 0.21055544 | 0.68494621 |
| -0.4369453 | -0.4364281 | 4.1262958 | 1.56745552 | 0.21057692 | 0.68494621 |
| -0.4220976 | -0.4367505 | 3.88994114 | 1.56597277 | 0.21079283 | 0.68494621 |
| 0.53329644 | 0.51385131 | 3.96331621 | 1.56082211 | 0.21154489 | 0.68494621 |
| 0.42658508 | 0.45602196 | 3.98269547 | 1.56052771 | 0.21158797 | 0.68494621 |
| 0.49878099 | 0.53808332 | 3.09381668 | 1.56052006 | 0.21158909 | 0.68494621 |
| 0.62373499 | 0.62219625 | 2.44032152 | 1.56051278 | 0.21159015 | 0.68494621 |
| 0.57606241 | 0.58590419 | 2.77968871 | 1.55370121 | 0.21258987 | 0.68513858 |
| 0.48310254 | 0.49827544 | 4.93154042 | 1.55285098 | 0.21271505 | 0.68513858 |
| -0.3966957 | -0.4360465 | 3.44811648 | 1.55083485 | 0.21301224 | 0.68513858 |
| 1.49160214 | 1.50884863 | 9.78014518 | 1.55078507 | 0.21301958 | 0.68513858 |
| 0.47669359 | 0.5304213 | 4.46816897 | 1.5499292 | 0.21314589 | 0.68513858 |
| -0.3977494 | -0.3979534 | 6.0550638 | 1.54911047 | 0.21326681 | 0.68513858 |
| 0.53432837 | 0.57756009 | 3.0973591 | 1.54883985 | 0.21330679 | 0.68513858 |
| 0.52799976 | 0.61573046 | 2.81299651 | 1.54532845 | 0.21382642 | 0.68590743 |
| -0.4400831 | -0.4430618 | 6.22974234 | 1.54383671 | 0.21404762 | 0.68590743 |
| -0.4115835 | -0.4256028 | 5.11138049 | 1.54197486 | 0.21432409 | 0.68590743 |
| -0.4125033 | -0.4173772 | 6.28604789 | 1.53965722 | 0.21466883 | 0.68590743 |
| 0.57048771 | 0.56052335 | 2.81852778 | 1.53868072 | 0.21481428 | 0.68590743 |
| -0.4225083 | -0.4304974 | 3.52129485 | 1.53758029 | 0.21497833 | 0.68590743 |
| 0.82032962 | 0.72754015 | 2.56269564 | 1.53605991 | 0.21520523 | 0.68590743 |
| 0.5416137 | 0.54952863 | 3.15386545 | 1.53328989 | 0.21561936 | 0.68596686 |
| 0.41504402 | 0.43010427 | 4.39904685 | 1.53230288 | 0.21576715 | 0.68596686 |
| 0.58023635 | 0.58935915 | 2.83219573 | 1.52946668 | 0.2161925 | 0.68596686 |
| 0.57456591 | 0.52804709 | 4.07961668 | 1.52737366 | 0.21650704 | 0.68596686 |
| 0.50743769 | 0.50192147 | 4.51159524 | 1.52714853 | 0.21654091 | 0.68596686 |
| 0.45474642 | 0.47003655 | 3.27828147 | 1.52588312 | 0.21673137 | 0.68596686 |
| 0.43555238 | 0.42614897 | 3.52110968 | 1.52487602 | 0.2168831 | 0.68596686 |
| -0.4153737 | -0.4210419 | 4.13842293 | 1.52008459 | 0.21760669 | 0.68677074 |
| -0.4836137 | -0.4898232 | 3.16073599 | 1.51922233 | 0.21773722 | 0.68677074 |
| 0.4212328 | 0.40750594 | 5.8958232 | 1.51848315 | 0.21784918 | 0.68677074 |

| | | | | | |
|---|---|---|---|---|---|
| 0.57789996 | 0.60822514 | 6.14031535 | 1.51527967 | 0.21833522 | 0.68695466 |
| -0.4935734 | -0.4931227 | 3.36265103 | 1.51497001 | 0.21838227 | 0.68695466 |
| 0.54852331 | 0.52739914 | 2.88149207 | 1.50787301 | 0.21946393 | 0.68910549 |
| 0.48464922 | 0.47732114 | 3.2218148 | 1.5073608 | 0.21954225 | 0.68910549 |
| -0.7000311 | -0.5672196 | 3.56493854 | 1.49422189 | 0.22156256 | 0.69221918 |
| -0.4115257 | -0.4210873 | 5.56482298 | 1.49273836 | 0.22179207 | 0.69221918 |
| 0.45089178 | 0.45674015 | 3.84173792 | 1.49201705 | 0.22190377 | 0.69221918 |
| 0.51569463 | 0.51706989 | 3.58177263 | 1.49122596 | 0.22202634 | 0.69221918 |
| -0.4717782 | -0.469502 | 3.88810684 | 1.48904296 | 0.22236501 | 0.69221918 |
| 0.42391342 | 0.42545166 | 3.41204387 | 1.48870358 | 0.22241772 | 0.69221918 |
| -0.4174995 | -0.4120817 | 3.70005537 | 1.4867731 | 0.22271781 | 0.69221918 |
| -0.4507592 | -0.4449804 | 4.08737975 | 1.48646945 | 0.22276505 | 0.69221918 |
| -0.4417071 | -0.4558983 | 3.50444506 | 1.48589746 | 0.22285409 | 0.69221918 |
| 0.58693225 | 0.5950311 | 2.69772411 | 1.48462334 | 0.22305256 | 0.69221918 |
| -0.4178419 | -0.42699 | 4.33366108 | 1.48389991 | 0.22316534 | 0.69221918 |
| -0.4773625 | -0.4788401 | 3.61174143 | 1.47958496 | 0.22383947 | 0.69356684 |
| 0.42013812 | 0.41845692 | 6.65093659 | 1.47470912 | 0.22460417 | 0.69384064 |
| -0.4356454 | -0.4392351 | 4.30637998 | 1.47437345 | 0.22465693 | 0.69384064 |
| 0.65088194 | 0.65121887 | 2.24091033 | 1.47396368 | 0.22472135 | 0.69384064 |
| 0.4650128 | 0.48914525 | 3.2899767 | 1.47291177 | 0.22488684 | 0.69384064 |
| -0.3917744 | -0.3961167 | 5.87877703 | 1.46712694 | 0.22579955 | 0.69527214 |
| 0.51411423 | 0.54846284 | 3.54377203 | 1.46297293 | 0.22645769 | 0.69527214 |
| 0.54109776 | 0.49726774 | 2.96354928 | 1.46289845 | 0.22646951 | 0.69527214 |
| 0.44982559 | 0.45840752 | 4.71807147 | 1.46200689 | 0.22661107 | 0.69527214 |
| 0.41847343 | 0.41990484 | 5.74971155 | 1.46009866 | 0.22691442 | 0.69527214 |
| 0.4464829 | 0.44266729 | 4.32718737 | 1.45945688 | 0.22701656 | 0.69527214 |
| 0.58108552 | 0.57246359 | 3.11278337 | 1.45935647 | 0.22703254 | 0.69527214 |
| -0.4112403 | -0.4174461 | 5.70997564 | 1.45541449 | 0.22766114 | 0.69646018 |
| -0.4975223 | -0.49746 | 3.06378687 | 1.4502339 | 0.22849043 | 0.69680106 |
| 0.41133582 | 0.42742948 | 3.82054493 | 1.45010017 | 0.22851188 | 0.69680106 |
| 0.46517574 | 0.46986847 | 3.10159416 | 1.4494336 | 0.22861886 | 0.69680106 |
| -0.4812647 | -0.476487 | 3.54130301 | 1.44333758 | 0.22960001 | 0.69680106 |
| 0.55327144 | 0.5454987 | 2.41750105 | 1.44312181 | 0.22963483 | 0.69680106 |
| -0.6201497 | -0.5206146 | 4.00545818 | 1.44226463 | 0.22977323 | 0.69680106 |
| -1.2556887 | -0.9054697 | 3.49673017 | 1.44082673 | 0.2300056 | 0.69680106 |
| 0.68541332 | 0.6135335 | 3.8332125 | 1.4408123 | 0.23000794 | 0.69680106 |
| -0.4001981 | -0.4129817 | 5.19722196 | 1.44056346 | 0.23004818 | 0.69680106 |
| -0.4511746 | -0.4511595 | 4.36045567 | 1.43974686 | 0.23018031 | 0.69680106 |
| -0.5021185 | -0.5342141 | 3.77161046 | 1.4357657 | 0.23082577 | 0.69802486 |
| 0.49942827 | 0.47548186 | 2.8554996 | 1.43413114 | 0.23109142 | 0.6980987 |
| 0.51576515 | 0.51741976 | 2.99659461 | 1.43017527 | 0.23173584 | 0.6987021 |
| 0.5302866 | 0.54303112 | 2.88103028 | 1.42994129 | 0.23177402 | 0.6987021 |
| 0.84279238 | 0.69464863 | 2.79316968 | 1.42820561 | 0.23205751 | 0.69882875 |
| 0.4682539 | 0.48229963 | 3.77093859 | 1.42042533 | 0.23333341 | 0.70111744 |
| 0.39696307 | 0.39556402 | 4.03489245 | 1.42007206 | 0.23339154 | 0.70111744 |
| -0.4577193 | -0.4581316 | 3.91124788 | 1.4175933 | 0.23379994 | 0.70111744 |

| | | | | | |
|---|---|---|---|---|---|
| -0.5116012 | -0.4808024 | 4.81338399 | 1.41686612 | 0.23391991 | 0.70111744 |
| -0.4578925 | -0.4545772 | 3.30215539 | 1.41620632 | 0.23402883 | 0.70111744 |
| -0.61899 | -0.5699476 | 3.04712264 | 1.412462 | 0.23464812 | 0.70147001 |
| -0.4701658 | -0.4639533 | 3.57070929 | 1.40988245 | 0.23507591 | 0.70147001 |
| -0.5566981 | -0.4866794 | 3.9687123 | 1.40947183 | 0.23514409 | 0.70147001 |
| -0.4636684 | -0.4658259 | 3.18617551 | 1.40888943 | 0.23524084 | 0.70147001 |
| 0.54676519 | 0.52942776 | 2.73999714 | 1.40818181 | 0.23535846 | 0.70147001 |
| -0.4108529 | -0.4142986 | 5.49905503 | 1.40637237 | 0.23565953 | 0.70164475 |
| 0.55051629 | 0.54541125 | 2.7827522 | 1.40179543 | 0.23642319 | 0.70287854 |
| -0.508326 | -0.4781239 | 3.77775592 | 1.4009793 | 0.23655967 | 0.70287854 |
| 0.62181009 | 0.54187523 | 4.22935706 | 1.39539133 | 0.23749674 | 0.70467319 |
| -0.4659406 | -0.4796476 | 3.03844503 | 1.39447596 | 0.23765067 | 0.70467319 |
| 0.70822621 | 0.6572386 | 2.52489088 | 1.39145704 | 0.23815919 | 0.70545825 |
| -0.4098592 | -0.4138797 | 6.55176562 | 1.38873747 | 0.23861843 | 0.70609584 |
| -0.4136337 | -0.4248516 | 3.54319743 | 1.38373722 | 0.23946559 | 0.70787887 |
| 0.53496616 | 0.52110637 | 2.77453396 | 1.38128356 | 0.23988263 | 0.7083881 |
| 0.50951132 | 0.48657181 | 4.20588633 | 1.37825892 | 0.24039794 | 0.70850619 |
| 0.52600717 | 0.53246424 | 2.78572743 | 1.37817497 | 0.24041226 | 0.70850619 |
| 0.45332233 | 0.45806493 | 3.85721294 | 1.37471326 | 0.24100373 | 0.70952675 |
| 0.49622612 | 0.47149664 | 3.04820723 | 1.37160009 | 0.24153716 | 0.70955244 |
| 0.45657844 | 0.47843075 | 4.12591286 | 1.37124506 | 0.24159808 | 0.70955244 |
| -0.4323466 | -0.4419463 | 3.38826066 | 1.37037189 | 0.241748 | 0.70955244 |
| -0.417985 | -0.4169225 | 7.69018481 | 1.36751972 | 0.24223849 | 0.71027172 |
| 0.4236701 | 0.41442828 | 5.24181865 | 1.35641952 | 0.24415897 | 0.71474882 |
| 0.54780741 | 0.55480754 | 2.50827713 | 1.3554488 | 0.2443278 | 0.71474882 |
| -0.4522735 | -0.4252938 | 3.53789888 | 1.3531126 | 0.2447347 | 0.71474882 |
| 0.48526803 | 0.48637328 | 3.43811445 | 1.35300588 | 0.24475331 | 0.71474882 |
| -0.4044684 | -0.4182647 | 3.41038708 | 1.34846801 | 0.24554612 | 0.71579857 |
| -0.4597732 | -0.4633001 | 3.55463302 | 1.34769196 | 0.24568202 | 0.71579857 |
| 0.54121132 | 0.54932745 | 2.95847811 | 1.34670606 | 0.24585479 | 0.71579857 |
| -0.4186913 | -0.4489878 | 2.9770922 | 1.34366642 | 0.24638842 | 0.71663125 |
| -0.4459924 | -0.47575 | 2.96793003 | 1.34103025 | 0.24685236 | 0.71681138 |
| 0.45875944 | 0.45612342 | 3.52250508 | 1.34050046 | 0.24694573 | 0.71681138 |
| -0.4790174 | -0.4896682 | 2.89579694 | 1.33854783 | 0.24729022 | 0.71681678 |
| -0.445622 | -0.4626804 | 2.90236794 | 1.33702272 | 0.2475597 | 0.71681678 |
| -0.4864981 | -0.489049 | 3.15909798 | 1.33628227 | 0.24769066 | 0.71681678 |
| -0.515934 | -0.5206688 | 3.17070187 | 1.33284513 | 0.2482997 | 0.71748449 |
| -0.4302507 | -0.4231272 | 5.76976976 | 1.33187343 | 0.24847221 | 0.71748449 |
| 0.44278408 | 0.44706767 | 3.81289815 | 1.33078761 | 0.24866515 | 0.71748449 |
| 0.43745163 | 0.44283952 | 4.26492314 | 1.3267089 | 0.24939155 | 0.7188637 |
| -0.3946969 | -0.3927988 | 4.93930807 | 1.32246984 | 0.25014927 | 0.72033034 |
| 0.53532472 | 0.53651928 | 2.81907915 | 1.31690802 | 0.25114772 | 0.72087359 |
| -0.4124165 | -0.4143552 | 4.28122902 | 1.3162487 | 0.2512664 | 0.72087359 |
| 0.44794396 | 0.4482644 | 3.54297551 | 1.314646 | 0.25155519 | 0.72087359 |
| -0.396045 | -0.392862 | 4.99697563 | 1.31325725 | 0.25180576 | 0.72087359 |
| 0.51399989 | 0.49985112 | 3.56812965 | 1.31295289 | 0.25186071 | 0.72087359 |

| | | | | | |
|---|---|---|---|---|---|
| -0.3677636 | -0.3802437 | 4.15447164 | 1.3125775 | 0.25192851 | 0.72087359 |
| 0.37164377 | 0.39214381 | 5.31316072 | 1.31173052 | 0.25208158 | 0.72087359 |
| -0.4661129 | -0.4727223 | 3.45131377 | 1.30769284 | 0.2528128 | 0.72204479 |
| 0.38305803 | 0.3864952 | 5.11233015 | 1.30671589 | 0.25299012 | 0.72204479 |
| -0.4208102 | -0.4214494 | 3.77822472 | 1.30184997 | 0.25387558 | 0.72385806 |
| 0.57137402 | 0.59117774 | 2.23614514 | 1.30034689 | 0.25414987 | 0.72392689 |
| -0.3765116 | -0.3818059 | 5.00332946 | 1.29690218 | 0.25477985 | 0.72500776 |
| -0.4276145 | -0.4269546 | 5.05653006 | 1.29543301 | 0.25504912 | 0.72506107 |
| -0.383708 | -0.4211776 | 3.22526901 | 1.2907011 | 0.25591879 | 0.7265259 |
| 0.47163996 | 0.48434389 | 2.62640378 | 1.2898994 | 0.25606649 | 0.7265259 |
| 0.94130113 | 0.66836919 | 3.03652739 | 1.28823007 | 0.25637438 | 0.72668703 |
| 0.37371072 | 0.37727218 | 5.2638205 | 1.28129765 | 0.2576579 | 0.72858975 |
| -0.4196093 | -0.4205606 | 3.70025326 | 1.28110757 | 0.2576932 | 0.72858975 |
| 0.5289568 | 0.55702387 | 2.92760634 | 1.28052773 | 0.25780094 | 0.72858975 |
| 0.45927076 | 0.45310109 | 3.89593536 | 1.27669241 | 0.25851493 | 0.72989484 |
| 0.44758171 | 0.41494265 | 3.42513517 | 1.27176469 | 0.25943589 | 0.73089316 |
| -0.3714662 | -0.4007379 | 3.35124778 | 1.26816334 | 0.26011152 | 0.73089316 |
| -0.3809764 | -0.3942682 | 5.19112165 | 1.26804808 | 0.26013318 | 0.73089316 |
| -0.3658031 | -0.3852206 | 4.8617555 | 1.26790506 | 0.26016006 | 0.73089316 |
| 0.5696502 | 0.57609577 | 2.7045852 | 1.26523078 | 0.26066329 | 0.73089316 |
| 0.51914211 | 0.50605547 | 3.25715117 | 1.26399841 | 0.2608956 | 0.73089316 |
| 0.42499918 | 0.43451082 | 4.74891631 | 1.2635376 | 0.26098253 | 0.73089316 |
| 0.4251091 | 0.42342759 | 3.25510479 | 1.26337872 | 0.26101251 | 0.73089316 |
| -0.4681501 | -0.4381687 | 4.22987468 | 1.26143319 | 0.26137998 | 0.73089316 |
| 0.51482057 | 0.51583419 | 2.96681041 | 1.25725963 | 0.26217044 | 0.73089316 |
| -0.3617325 | -0.3684282 | 6.44460897 | 1.25688716 | 0.26224113 | 0.73089316 |
| 0.40871006 | 0.38946788 | 3.84931988 | 1.25648984 | 0.26231656 | 0.73089316 |
| -0.3674213 | -0.3648161 | 6.67562093 | 1.25509207 | 0.26258214 | 0.73089316 |
| 0.42766078 | 0.43034622 | 4.2742196 | 1.25496119 | 0.26260703 | 0.73089316 |
| 0.43059507 | 0.43523614 | 3.48895781 | 1.2542592 | 0.26274055 | 0.73089316 |
| 0.43788386 | 0.43837156 | 4.79508963 | 1.25337214 | 0.26290939 | 0.73089316 |
| 0.45984801 | 0.44981662 | 3.16855941 | 1.25119692 | 0.263324 | 0.73119908 |
| -0.3920616 | -0.4116051 | 3.4054073 | 1.25014515 | 0.26352476 | 0.73119908 |
| 0.46021318 | 0.46700982 | 3.2267271 | 1.24777797 | 0.2639773 | 0.73175316 |
| 0.52289581 | 0.52236044 | 2.95461346 | 1.24402948 | 0.26469589 | 0.7327712 |
| -0.4016764 | -0.4026325 | 4.19338252 | 1.2432222 | 0.26485096 | 0.7327712 |
| 0.45184098 | 0.44155199 | 3.15166671 | 1.23887001 | 0.26568894 | 0.73402574 |
| 0.39308686 | 0.44150822 | 3.09627517 | 1.23602019 | 0.26623944 | 0.73402574 |
| -0.437929 | -0.4244128 | 3.14716652 | 1.23597203 | 0.26624876 | 0.73402574 |
| -0.3606535 | -0.3977556 | 3.37644166 | 1.23560921 | 0.26631895 | 0.73402574 |
| -0.4383748 | -0.4204014 | 3.64245181 | 1.2332309 | 0.26677962 | 0.73459584 |
| 0.40769194 | 0.41429638 | 4.13901475 | 1.23164883 | 0.26708662 | 0.73461981 |
| -0.4383457 | -0.4343803 | 3.47469503 | 1.23057104 | 0.26729601 | 0.73461981 |
| -0.4713759 | -0.4431537 | 4.03546402 | 1.22832837 | 0.26773238 | 0.73479718 |
| -0.4016764 | -0.3903641 | 4.60675394 | 1.22763044 | 0.26786836 | 0.73479718 |
| 0.49918318 | 0.49155674 | 2.8747236 | 1.22432828 | 0.26851289 | 0.73585585 |

| | | | | | |
|---|---|---|---|---|---|
| 0.52867162 | 0.50561171 | 2.61322514 | 1.21977857 | 0.26940411 | 0.73585585 |
| 0.41248805 | 0.39730468 | 3.50818292 | 1.21828624 | 0.26969724 | 0.73585585 |
| 0.48843069 | 0.48190008 | 3.04840418 | 1.21800801 | 0.26975194 | 0.73585585 |
| -0.3602163 | -0.3800517 | 4.94605508 | 1.21685374 | 0.26997899 | 0.73585585 |
| 0.53490479 | 0.5259871 | 3.0118671 | 1.21663891 | 0.27002128 | 0.73585585 |
| 0.54381965 | 0.58223057 | 2.85926152 | 1.21657337 | 0.27003418 | 0.73585585 |
| -0.3764372 | -0.3842999 | 4.99440644 | 1.21348171 | 0.27064367 | 0.73682294 |
| 0.46217156 | 0.46476746 | 4.04248811 | 1.21072297 | 0.27118899 | 0.73761366 |
| -0.37195 | -0.3820976 | 5.0187958 | 1.20785664 | 0.27175703 | 0.73840437 |
| -0.3937345 | -0.3962961 | 3.68801748 | 1.20453089 | 0.27241798 | 0.73840437 |
| -0.4538719 | -0.4382371 | 4.6564677 | 1.20424436 | 0.27247501 | 0.73840437 |
| 0.37686775 | 0.39646741 | 4.52210491 | 1.20411737 | 0.2725003 | 0.73840437 |
| 0.39230553 | 0.39971402 | 5.2312819 | 1.20115879 | 0.2730902 | 0.73880367 |
| 0.48050637 | 0.52319843 | 4.84965487 | 1.19903551 | 0.27351453 | 0.73880367 |
| -0.4385676 | -0.4714692 | 3.60819344 | 1.19796054 | 0.27372968 | 0.73880367 |
| -0.4299148 | -0.4279159 | 3.52111377 | 1.19781727 | 0.27375837 | 0.73880367 |
| -0.5744792 | -0.4936957 | 3.34088876 | 1.19402197 | 0.27451978 | 0.73880367 |
| 0.45558962 | 0.44846916 | 3.2211745 | 1.19234159 | 0.27485774 | 0.73880367 |
| -0.4066057 | -0.4116417 | 3.51495408 | 1.19219887 | 0.27488647 | 0.73880367 |
| 0.4234682 | 0.42498274 | 3.58177959 | 1.19211148 | 0.27490406 | 0.73880367 |
| 0.45284093 | 0.42667514 | 3.17692401 | 1.1919069 | 0.27494525 | 0.73880367 |
| 0.38131661 | 0.39209337 | 5.16529005 | 1.18579263 | 0.27617988 | 0.7404822 |
| -0.3963251 | -0.3900107 | 4.50346266 | 1.18536224 | 0.27626705 | 0.7404822 |
| 0.39340216 | 0.39327304 | 4.8992044 | 1.18501443 | 0.27633752 | 0.7404822 |
| -0.4556358 | -0.4413658 | 3.66736718 | 1.18198302 | 0.27695266 | 0.74144404 |
| -0.5325276 | -0.4942385 | 3.10695524 | 1.1802953 | 0.27729589 | 0.74164044 |
| -0.4421683 | -0.4411679 | 2.95154565 | 1.17910359 | 0.27753856 | 0.74164044 |
| 0.36355961 | 0.36577232 | 3.76670911 | 1.16236717 | 0.28097515 | 0.74943587 |
| -0.5207913 | -0.4671304 | 3.49773143 | 1.16231715 | 0.2809855 | 0.74943587 |
| 0.68777092 | 0.60336561 | 2.90265603 | 1.16112343 | 0.28123267 | 0.74943587 |
| -0.3604624 | -0.3655819 | 5.04830915 | 1.15372586 | 0.28277055 | 0.75235804 |
| -0.4432185 | -0.4326615 | 3.36542281 | 1.15119169 | 0.28329983 | 0.75235804 |
| -0.3513133 | -0.385209 | 3.48135605 | 1.15017296 | 0.28351295 | 0.75235804 |
| -0.3808359 | -0.3849529 | 5.14357028 | 1.14965792 | 0.28362077 | 0.75235804 |
| -0.3777215 | -0.380339 | 5.71214891 | 1.14961813 | 0.2836291 | 0.75235804 |
| -0.4028637 | -0.4002595 | 5.61311486 | 1.14706997 | 0.28416335 | 0.75308492 |
| -0.3709635 | -0.37395 | 6.05727129 | 1.14399522 | 0.2848097 | 0.75398045 |
| 0.48679577 | 0.50892956 | 2.82304553 | 1.14298566 | 0.28502232 | 0.75398045 |
| 0.38220655 | 0.3637715 | 4.14266796 | 1.1384073 | 0.28598912 | 0.75439673 |
| 0.60955746 | 0.50593524 | 3.86985998 | 1.13668711 | 0.28635345 | 0.75439673 |
| -0.3292834 | -0.3555405 | 4.38454267 | 1.13631918 | 0.28643145 | 0.75439673 |
| -0.4135749 | -0.4263253 | 3.14174528 | 1.13617117 | 0.28646283 | 0.75439673 |
| -0.379219 | -0.3600109 | 4.08499626 | 1.13607576 | 0.28648307 | 0.75439673 |
| 0.43824353 | 0.43027677 | 4.30791289 | 1.12936706 | 0.28791039 | 0.75746605 |
| 0.38340498 | 0.41418768 | 3.4529402 | 1.12577814 | 0.28867766 | 0.75799868 |
| 0.37122802 | 0.39307376 | 4.81163775 | 1.12559399 | 0.2887171 | 0.75799868 |

| | | | | | |
|---|---|---|---|---|---|
| -0.3651709 | -0.3645078 | 3.37156119 | 1.1246543 | 0.28891847 | 0.75799868 |
| 0.4116583 | 0.41664703 | 3.60198602 | 1.12217883 | 0.28944978 | 0.75799868 |
| 0.37066721 | 0.36470726 | 6.17797731 | 1.12013578 | 0.28988922 | 0.75799868 |
| 0.41109624 | 0.44191726 | 3.8187277 | 1.12013021 | 0.28989042 | 0.75799868 |
| -0.3537054 | -0.3614696 | 5.32134662 | 1.11987077 | 0.28994628 | 0.75799868 |
| -0.3491515 | -0.3590937 | 5.74220096 | 1.11698939 | 0.29056764 | 0.7589375 |
| -0.4903145 | -0.444097 | 3.36621577 | 1.11097459 | 0.29187019 | 0.76057731 |
| 0.36151635 | 0.37820433 | 4.82876776 | 1.10917654 | 0.29226102 | 0.76057731 |
| -0.3750901 | -0.3743942 | 6.88552976 | 1.10415007 | 0.29335714 | 0.76057731 |
| 0.41186166 | 0.40901903 | 4.01734366 | 1.10394692 | 0.29340155 | 0.76057731 |
| -0.3491697 | -0.3588632 | 5.40570936 | 1.10391505 | 0.29340852 | 0.76057731 |
| 0.38001763 | 0.36919358 | 5.2734901 | 1.10199965 | 0.29382768 | 0.76057731 |
| -0.3442309 | -0.3681139 | 3.74952718 | 1.10142655 | 0.29395325 | 0.76057731 |
| 0.37699187 | 0.38437728 | 3.47458436 | 1.10030574 | 0.29419902 | 0.76057731 |
| 0.48885321 | 0.44344149 | 4.1483489 | 1.09968651 | 0.29433491 | 0.76057731 |
| 0.42025619 | 0.42769627 | 3.13056752 | 1.09756661 | 0.29480075 | 0.76057731 |
| -0.3335406 | -0.3321098 | 6.07386699 | 1.09690842 | 0.29494558 | 0.76057731 |
| 0.36514479 | 0.36405444 | 6.84258614 | 1.09647228 | 0.2950416 | 0.76057731 |
| -0.3441808 | -0.3586755 | 4.78608199 | 1.09559885 | 0.295234 | 0.76057731 |
| -0.6817744 | -0.5654744 | 5.18491724 | 1.0943373 | 0.2955122 | 0.76057731 |
| 0.39889435 | 0.45389055 | 2.88422751 | 1.09401382 | 0.29558358 | 0.76057731 |
| -0.3906611 | -0.4023902 | 3.20685367 | 1.09361105 | 0.2956725 | 0.76057731 |
| -0.3776869 | -0.3790178 | 4.0731297 | 1.09203281 | 0.29602124 | 0.76057731 |
| 0.40565811 | 0.40187987 | 2.94830336 | 1.08856567 | 0.29678923 | 0.76057731 |
| -0.3646973 | -0.3695393 | 4.09721003 | 1.08386088 | 0.29783545 | 0.76057731 |
| 0.4844202 | 0.48535812 | 2.97575434 | 1.08201092 | 0.29824813 | 0.76057731 |
| -1.3029939 | -0.9956884 | 5.03334821 | 1.08197695 | 0.29825572 | 0.76057731 |
| 0.37982032 | 0.36947042 | 4.04166139 | 1.08162798 | 0.29833365 | 0.76057731 |
| 0.40856284 | 0.41017581 | 3.79123758 | 1.08071361 | 0.29853797 | 0.76057731 |
| 0.3776988 | 0.39337203 | 4.74112028 | 1.07998901 | 0.29870002 | 0.76057731 |
| -0.4617736 | -0.4321106 | 3.61908844 | 1.0779689 | 0.29915237 | 0.76057731 |
| 0.34808052 | 0.36256881 | 5.68646661 | 1.07730178 | 0.29930195 | 0.76057731 |
| 0.38071736 | 0.37670853 | 4.50563806 | 1.07714014 | 0.29933821 | 0.76057731 |
| -0.3163405 | -0.3522512 | 3.70488378 | 1.07661107 | 0.29945692 | 0.76057731 |
| -0.3422125 | -0.3460923 | 5.09062103 | 1.07596404 | 0.29960219 | 0.76057731 |
| -0.4085429 | -0.4104436 | 9.26654802 | 1.07416148 | 0.30000735 | 0.76057731 |
| -0.3498917 | -0.3549293 | 4.84290248 | 1.07374593 | 0.30010086 | 0.76057731 |
| 0.42475021 | 0.43229953 | 4.31746572 | 1.07193258 | 0.30050933 | 0.76057731 |
| 0.91819386 | 0.66180006 | 2.96524321 | 1.07017279 | 0.30090642 | 0.76057731 |
| 0.73911402 | 0.61391133 | 4.02999097 | 1.0695583 | 0.30104524 | 0.76057731 |
| -0.4321221 | -0.4301975 | 3.40682264 | 1.06875147 | 0.30122763 | 0.76057731 |
| 0.38843011 | 0.39452941 | 4.49522767 | 1.06702497 | 0.3016184 | 0.76057731 |
| 0.36551691 | 0.396348 | 3.26559235 | 1.06692354 | 0.30164138 | 0.76057731 |
| 0.36758999 | 0.37172694 | 6.52820123 | 1.0666772 | 0.3016972 | 0.76057731 |
| 0.46169417 | 0.44104536 | 5.29742186 | 1.06654152 | 0.30172794 | 0.76057731 |
| 0.46582242 | 0.42973421 | 2.74215384 | 1.06624846 | 0.30179437 | 0.76057731 |

| | | | | | |
|---|---|---|---|---|---|
| -0.4375758 | -0.4288168 | 3.2473675 | 1.06495999 | 0.30208665 | 0.76057731 |
| 0.38016178 | 0.36590134 | 4.57166683 | 1.06366652 | 0.30238042 | 0.76057731 |
| 0.36304809 | 0.39204248 | 4.40746162 | 1.06254031 | 0.30263651 | 0.76057731 |
| -0.4034703 | -0.3900822 | 3.19317517 | 1.0620013 | 0.30275918 | 0.76057731 |
| -0.4424284 | -0.4228855 | 3.66880622 | 1.05322474 | 0.30476557 | 0.76495365 |
| -0.323498 | -0.3434986 | 4.34064586 | 1.04937355 | 0.30565141 | 0.76601544 |
| -0.3537209 | -0.3501134 | 4.52841959 | 1.04908471 | 0.30571798 | 0.76601544 |
| 0.36520632 | 0.36883827 | 5.71459241 | 1.04518883 | 0.30661775 | 0.76760533 |
| 0.35733817 | 0.35824908 | 4.01197598 | 1.04294534 | 0.30713745 | 0.76824182 |
| 0.46617658 | 0.54131022 | 3.40627581 | 1.03793323 | 0.30830263 | 0.76965046 |
| 0.47779853 | 0.43972882 | 4.78507284 | 1.0360671 | 0.30873793 | 0.76965046 |
| 0.38157501 | 0.37487814 | 4.81480634 | 1.0351922 | 0.30894228 | 0.76965046 |
| 0.3877837 | 0.37517735 | 3.68798996 | 1.03500829 | 0.30898526 | 0.76965046 |
| -0.3860327 | -0.3796836 | 4.72570596 | 1.03459826 | 0.30908111 | 0.76965046 |
| -0.3969379 | -0.3990283 | 3.41836679 | 1.03322005 | 0.30940357 | 0.76965046 |
| 0.45898373 | 0.47127832 | 2.64316064 | 1.03231703 | 0.30961508 | 0.76965046 |
| 0.40972987 | 0.40859326 | 3.69097253 | 1.03002875 | 0.31015192 | 0.76965046 |
| -0.3903339 | -0.3947612 | 3.35593755 | 1.02944366 | 0.31028937 | 0.76965046 |
| 0.47167019 | 0.4370975 | 4.04066908 | 1.02819573 | 0.31058282 | 0.76965046 |
| -0.5088276 | -0.4350654 | 3.60273972 | 1.02744931 | 0.3107585 | 0.76965046 |
| 0.37357699 | 0.3605379 | 3.53098449 | 1.02688261 | 0.31089198 | 0.76965046 |
| -0.3546413 | -0.3629039 | 4.44211348 | 1.0247088 | 0.31140467 | 0.77026077 |
| -0.3466671 | -0.3558396 | 4.1766937 | 1.01934493 | 0.31267444 | 0.77223281 |
| 1.45135564 | 0.85753696 | 3.01108277 | 1.01806182 | 0.31297919 | 0.77223281 |
| 0.32364908 | 0.34989688 | 4.90488653 | 1.01796397 | 0.31300245 | 0.77223281 |
| 0.33100819 | 0.33169596 | 5.96533437 | 1.0150037 | 0.31370708 | 0.77252429 |
| 0.37972784 | 0.37521904 | 4.50158804 | 1.01237659 | 0.31433415 | 0.77252429 |
| -0.3575905 | -0.3602626 | 3.80735519 | 1.01229644 | 0.31435331 | 0.77252429 |
| -0.3730459 | -0.3731183 | 4.38490822 | 1.01202653 | 0.31441783 | 0.77252429 |
| -0.3370607 | -0.343363 | 6.14542775 | 1.01186987 | 0.31445529 | 0.77252429 |
| 0.45032498 | 0.46738835 | 2.85201735 | 1.00841618 | 0.31528257 | 0.77341993 |
| 0.37647621 | 0.35872977 | 3.89032872 | 1.00771245 | 0.31545149 | 0.77341993 |
| -0.3716433 | -0.3603774 | 3.57752661 | 1.00569503 | 0.31593639 | 0.77341993 |
| 0.38593646 | 0.42303086 | 3.0284212 | 1.00556906 | 0.3159667 | 0.77341993 |
| -0.3385477 | -0.3410398 | 4.43678582 | 1.00478223 | 0.31615611 | 0.77341993 |
| 0.52232589 | 0.45095672 | 3.70223184 | 1.00292883 | 0.31660285 | 0.77385866 |
| -0.4056335 | -0.40606 | 3.17120985 | 0.99999594 | 0.31731149 | 0.77493625 |
| -0.3465777 | -0.3328312 | 5.25808899 | 0.99771027 | 0.31786519 | 0.77562635 |
| -0.6773313 | -0.5724477 | 6.19676615 | 0.99582577 | 0.31832266 | 0.77562635 |
| -0.4198337 | -0.4206838 | 9.17966774 | 0.99551536 | 0.3183981 | 0.77562635 |
| -0.4378838 | -0.5169319 | 3.13739916 | 0.99373557 | 0.31883108 | 0.77578198 |
| 0.41134617 | 0.41676604 | 5.3408147 | 0.99142448 | 0.31939447 | 0.77578198 |
| -0.5081409 | -0.564182 | 6.70330194 | 0.99134855 | 0.319413 | 0.77578198 |
| 0.63340726 | 0.490985 | 3.73128847 | 0.9908519 | 0.31953425 | 0.77578198 |
| 0.4110897 | 0.42183794 | 2.97612282 | 0.98946791 | 0.31987246 | 0.77595213 |
| 0.35651434 | 0.35492434 | 4.71460647 | 0.98836586 | 0.3201421 | 0.77595581 |

| | | | | | |
|---|---|---|---|---|---|
| 0.34587709 | 0.3554606 | 5.23869306 | 0.98715761 | 0.32043807 | 0.77602325 |
| 0.4379202 | 0.44088062 | 2.97807946 | 0.9828015 | 0.32150814 | 0.7771322 |
| 0.43793798 | 0.47046747 | 2.74232436 | 0.98278601 | 0.32151195 | 0.7771322 |
| -0.3512892 | -0.3535972 | 5.41505556 | 0.98201605 | 0.32170158 | 0.7771322 |
| -0.3846576 | -0.3756632 | 3.58382531 | 0.9806226 | 0.32204515 | 0.7773133 |
| 0.35009193 | 0.33621166 | 4.23954332 | 0.97509867 | 0.32341187 | 0.77964152 |
| 0.3248698 | 0.34623255 | 4.15219157 | 0.97454798 | 0.32354854 | 0.77964152 |
| 0.3319568 | 0.32882145 | 5.75627377 | 0.97244354 | 0.32407152 | 0.77986415 |
| -0.5077751 | -0.432084 | 3.45872642 | 0.97134837 | 0.32434413 | 0.77986415 |
| -0.3467686 | -0.3574562 | 3.77613617 | 0.96923982 | 0.32486984 | 0.77986415 |
| -0.3467271 | -0.3466036 | 3.84761263 | 0.96867806 | 0.32501009 | 0.77986415 |
| 1.53921163 | 0.86712093 | 3.1936055 | 0.96814509 | 0.32514322 | 0.77986415 |
| 0.87325371 | 0.84187221 | 4.83106564 | 0.96679384 | 0.32548109 | 0.77986415 |
| -0.3526891 | -0.359538 | 3.56115648 | 0.96592777 | 0.32569789 | 0.77986415 |
| 0.44165558 | 0.46427443 | 2.61534469 | 0.96553312 | 0.32579674 | 0.77986415 |
| 0.4638947 | 0.47521166 | 2.67105042 | 0.96389656 | 0.32620709 | 0.78020109 |
| -0.4202133 | -0.4188501 | 9.34314042 | 0.96033196 | 0.32710325 | 0.78070054 |
| 1.09209858 | 0.65935945 | 3.14215101 | 0.96010061 | 0.32716153 | 0.78070054 |
| 0.34010989 | 0.34127818 | 4.81694441 | 0.95984787 | 0.32722521 | 0.78070054 |
| -0.3238577 | -0.3350933 | 4.7305529 | 0.95402516 | 0.32869684 | 0.78259176 |
| -0.3559222 | -0.3537268 | 4.03335279 | 0.95392233 | 0.32872291 | 0.78259176 |
| 0.47314522 | 0.45396624 | 3.52639624 | 0.95350332 | 0.32882916 | 0.78259176 |
| -0.3426955 | -0.3638379 | 3.41221565 | 0.95025044 | 0.32965556 | 0.78391388 |
| 0.39292488 | 0.38634505 | 5.35082568 | 0.94860175 | 0.33007547 | 0.78426798 |
| 0.4276512 | 0.40476432 | 5.01517273 | 0.94644861 | 0.33062492 | 0.78473088 |
| -0.3323471 | -0.3470949 | 3.6068892 | 0.94509575 | 0.33097078 | 0.78473088 |
| 0.38087903 | 0.38181338 | 5.07630533 | 0.94465422 | 0.33108376 | 0.78473088 |
| 1.41882126 | 0.82213104 | 2.98772846 | 0.94317923 | 0.33146156 | 0.78498344 |
| 0.41103312 | 0.39159059 | 2.79684155 | 0.93973673 | 0.33234556 | 0.7864334 |
| 0.44483536 | 0.42829275 | 6.19098298 | 0.93179032 | 0.33439814 | 0.79060456 |
| -0.3604696 | -0.3911538 | 3.06849966 | 0.93027715 | 0.33479092 | 0.79060456 |
| 0.46090247 | 0.46113388 | 2.80031076 | 0.92903876 | 0.33511283 | 0.79060456 |
| -0.5389408 | -0.4609881 | 3.82718772 | 0.9286997 | 0.33520104 | 0.79060456 |
| -0.3781546 | -0.3776984 | 3.27340074 | 0.92666793 | 0.33573026 | 0.79062217 |
| -0.3465619 | -0.3508394 | 3.48553255 | 0.92347993 | 0.33656291 | 0.79062217 |
| -0.3303557 | -0.3327919 | 4.84728122 | 0.92259107 | 0.33679556 | 0.79062217 |
| 0.34857708 | 0.35319664 | 4.28011417 | 0.92223299 | 0.33688935 | 0.79062217 |
| -0.3076376 | -0.3237225 | 4.07520934 | 0.92114796 | 0.33717373 | 0.79062217 |
| -0.3437309 | -0.3346719 | 4.42737504 | 0.91546131 | 0.33866951 | 0.79062217 |
| -0.3214217 | -0.328066 | 5.38839607 | 0.9152786 | 0.33871772 | 0.79062217 |
| -0.3254359 | -0.3179291 | 4.05076994 | 0.91517508 | 0.33874504 | 0.79062217 |
| -0.3579169 | -0.3687042 | 3.1778752 | 0.91456296 | 0.33890662 | 0.79062217 |
| 0.44696162 | 0.53197353 | 2.91198706 | 0.91433342 | 0.33896724 | 0.79062217 |
| -0.3319582 | -0.3185662 | 6.45385962 | 0.91421652 | 0.33899812 | 0.79062217 |
| 0.39752272 | 0.39348952 | 4.93672929 | 0.91400176 | 0.33905486 | 0.79062217 |
| -0.3662782 | -0.370266 | 3.33662033 | 0.9137477 | 0.33912199 | 0.79062217 |

| | | | | | |
|---|---|---|---|---|---|
| 0.36986141 | 0.36818316 | 3.09250452 | 0.9128674 | 0.33935476 | 0.79062217 |
| -0.5871469 | -0.4552542 | 4.54313613 | 0.91258488 | 0.3394295 | 0.79062217 |
| -0.3416142 | -0.346624 | 7.36589213 | 0.91182994 | 0.33962935 | 0.79062217 |
| 0.35947394 | 0.34929815 | 4.52581552 | 0.91014526 | 0.34007588 | 0.79062217 |
| 0.40709059 | 0.41023016 | 4.92189306 | 0.90985196 | 0.3401537 | 0.79062217 |
| 0.34083618 | 0.34202495 | 5.105735 | 0.90892736 | 0.34039918 | 0.79062217 |
| -0.3165053 | -0.3246233 | 5.59626538 | 0.90593008 | 0.3411966 | 0.79183878 |
| 0.32148784 | 0.33940711 | 4.25053174 | 0.90477013 | 0.34150588 | 0.79192148 |
| -0.4605534 | -0.4039456 | 3.56237735 | 0.90019897 | 0.34272836 | 0.79262994 |
| 0.37632209 | 0.37849354 | 3.2312387 | 0.89890661 | 0.34307506 | 0.79262994 |
| 0.40106608 | 0.36824674 | 3.19561574 | 0.89760095 | 0.3434258 | 0.79262994 |
| -0.3636073 | -0.3606925 | 3.57402316 | 0.89560108 | 0.34396397 | 0.79262994 |
| -0.4181004 | -0.3818499 | 4.71256971 | 0.8952188 | 0.34406697 | 0.79262994 |
| 0.35358987 | 0.3491772 | 4.4177477 | 0.89510747 | 0.34409698 | 0.79262994 |
| 0.51283407 | 0.42654019 | 4.04656109 | 0.89462939 | 0.34422586 | 0.79262994 |
| -0.3242952 | -0.3265236 | 5.50480811 | 0.89454587 | 0.34424839 | 0.79262994 |
| -0.4042458 | -0.3916756 | 3.34689385 | 0.89444209 | 0.34427638 | 0.79262994 |
| 0.40767093 | 0.3895538 | 5.27711787 | 0.88777764 | 0.34608026 | 0.79614965 |
| 0.42988719 | 0.40374503 | 2.66940865 | 0.88547579 | 0.34670628 | 0.79695629 |
| -0.8590371 | -0.7684155 | 7.15768304 | 0.88322542 | 0.34731978 | 0.7977329 |
| -0.345456 | -0.3425207 | 4.38668325 | 0.88141555 | 0.34781426 | 0.79812169 |
| 0.45221217 | 0.41320438 | 2.63436564 | 0.87913021 | 0.34844002 | 0.79812169 |
| 0.37121339 | 0.3684562 | 5.29956063 | 0.87863655 | 0.34857539 | 0.79812169 |
| 0.34540149 | 0.32748006 | 4.51892967 | 0.876705 | 0.34910574 | 0.79812169 |
| -0.3023804 | -0.3077095 | 6.03244069 | 0.87625239 | 0.34923018 | 0.79812169 |
| 0.31511605 | 0.3368822 | 3.76063426 | 0.87547401 | 0.34944431 | 0.79812169 |
| 0.40305803 | 0.39216086 | 4.22835068 | 0.87465214 | 0.34967061 | 0.79812169 |
| 0.33170361 | 0.32142357 | 3.71825587 | 0.8729328 | 0.35014466 | 0.79812169 |
| -0.3301193 | -0.3355405 | 4.54609772 | 0.87157084 | 0.3505208 | 0.79812169 |
| -0.3888543 | -0.3638371 | 3.66476594 | 0.8711582 | 0.35063487 | 0.79812169 |
| 0.37794849 | 0.48331911 | 4.68549018 | 0.86908808 | 0.35120789 | 0.79812169 |
| 0.36136905 | 0.35541335 | 3.55048016 | 0.86851132 | 0.35136777 | 0.79812169 |
| 0.36585281 | 0.37294312 | 3.2917737 | 0.86829418 | 0.35142798 | 0.79812169 |
| -0.3057598 | -0.320169 | 4.33592145 | 0.86775018 | 0.35157891 | 0.79812169 |
| 0.32882723 | 0.33801658 | 3.76648548 | 0.86754575 | 0.35163565 | 0.79812169 |
| -0.3066988 | -0.3244096 | 5.09808257 | 0.86658805 | 0.35190162 | 0.79812169 |
| -0.317076 | -0.3254673 | 4.91590797 | 0.86504036 | 0.35233202 | 0.79847209 |
| -0.3476557 | -0.3447261 | 4.42131094 | 0.86378438 | 0.35268182 | 0.79863943 |
| 0.34721012 | 0.33813272 | 3.69470859 | 0.86249637 | 0.35304104 | 0.79882782 |
| -0.3245327 | -0.3603267 | 3.10005526 | 0.85556288 | 0.35498338 | 0.80205044 |
| 0.43564156 | 0.41781687 | 3.29533276 | 0.85469709 | 0.35522694 | 0.80205044 |
| -0.333853 | -0.3339482 | 5.39360565 | 0.85444926 | 0.3552967 | 0.80205044 |
| 0.38155364 | 0.37828249 | 3.7012319 | 0.852373 | 0.35588189 | 0.80215327 |
| 0.3317369 | 0.31919824 | 3.76821691 | 0.85035939 | 0.35645069 | 0.80215327 |
| 0.39592196 | 0.37900762 | 3.93099838 | 0.849517 | 0.35668901 | 0.80215327 |
| -0.3670718 | -0.3795675 | 2.79369637 | 0.84894606 | 0.35685066 | 0.80215327 |

| | | | | | |
|---|---|---|---|---|---|
| -0.3532919 | -0.3541567 | 4.59429366 | 0.84849545 | 0.35697832 | 0.80215327 |
| 0.36972957 | 0.36420948 | 3.98311696 | 0.84803028 | 0.35711016 | 0.80215327 |
| 0.31261324 | 0.32820074 | 4.80864724 | 0.84742258 | 0.3572825 | 0.80215327 |
| -0.5461631 | -0.4484279 | 4.00078228 | 0.84525477 | 0.35789822 | 0.80254993 |
| 0.374141 | 0.38162188 | 2.85742938 | 0.84484837 | 0.35801381 | 0.80254993 |
| -0.4964031 | -0.4451815 | 3.10224116 | 0.84171515 | 0.3589067 | 0.80333689 |
| 0.33684492 | 0.34835324 | 3.98315166 | 0.84166841 | 0.35892004 | 0.80333689 |
| 0.34301275 | 0.33913451 | 3.3963626 | 0.83938201 | 0.35957358 | 0.80417769 |
| -0.3442193 | -0.3402318 | 6.10181897 | 0.83472736 | 0.36090912 | 0.80553323 |
| -0.6266136 | -0.4775437 | 3.04306264 | 0.83449211 | 0.3609768 | 0.80553323 |
| 0.32064569 | 0.33443186 | 4.1169363 | 0.83391904 | 0.36114174 | 0.80553323 |
| -0.2960209 | -0.3037721 | 4.30940278 | 0.83169677 | 0.36178235 | 0.80553323 |
| 0.44851221 | 0.45480801 | 2.33894904 | 0.83147674 | 0.36184586 | 0.80553323 |
| 0.47755088 | 0.40801304 | 3.723338 | 0.83028645 | 0.36218971 | 0.80553323 |
| 0.38981522 | 0.39041637 | 2.97415733 | 0.83026581 | 0.36219568 | 0.80553323 |
| 0.3123433 | 0.31144754 | 4.20720575 | 0.82801025 | 0.36284852 | 0.80553323 |
| 0.39047663 | 0.37842623 | 4.42089804 | 0.8264826 | 0.3632916 | 0.80553323 |
| -0.3010499 | -0.3141491 | 4.71065787 | 0.82640885 | 0.36331301 | 0.80553323 |
| 0.30596651 | 0.30736689 | 5.35110053 | 0.82538243 | 0.36361115 | 0.80553323 |
| 0.41320342 | 0.401817 | 2.77550999 | 0.82456034 | 0.36385019 | 0.80553323 |
| 0.33948622 | 0.32075153 | 3.6029687 | 0.82391156 | 0.36403899 | 0.80553323 |
| -0.3129199 | -0.3330129 | 3.3611229 | 0.82378264 | 0.36407653 | 0.80553323 |
| 0.32984923 | 0.33313606 | 5.74513572 | 0.8177438 | 0.36584065 | 0.8083596 |
| -0.3253441 | -0.328503 | 3.67836181 | 0.81589893 | 0.36638196 | 0.8083596 |
| -0.4214333 | -0.4221081 | 2.7012602 | 0.81425357 | 0.36686566 | 0.8083596 |
| -0.3837425 | -0.3636199 | 5.00138771 | 0.81364617 | 0.36704445 | 0.8083596 |
| 0.31831355 | 0.31467413 | 5.60636915 | 0.81341992 | 0.36711108 | 0.8083596 |
| -0.3175797 | -0.3299648 | 3.63887877 | 0.81336712 | 0.36712663 | 0.8083596 |
| 0.30339322 | 0.29092735 | 5.92734849 | 0.81142654 | 0.36769883 | 0.8083596 |
| -0.5191368 | -0.4135734 | 3.20947817 | 0.80627689 | 0.36922328 | 0.8083596 |
| 0.55565114 | 0.50017689 | 2.65789069 | 0.80624811 | 0.36923183 | 0.8083596 |
| 0.43827983 | 0.45495312 | 2.5509652 | 0.8054203 | 0.36947771 | 0.8083596 |
| 0.43127103 | 0.39991961 | 4.48963519 | 0.80512336 | 0.36956597 | 0.8083596 |
| -0.3082528 | -0.3150243 | 6.75793526 | 0.80420267 | 0.3698398 | 0.8083596 |
| 0.34901547 | 0.34829748 | 3.47744274 | 0.80396672 | 0.36991002 | 0.8083596 |
| -0.3281796 | -0.3322438 | 3.60294049 | 0.80371302 | 0.36998555 | 0.8083596 |
| 0.32989977 | 0.31417165 | 5.14938899 | 0.80240185 | 0.37037622 | 0.8083596 |
| 0.41193528 | 0.40390498 | 3.29147356 | 0.80234127 | 0.37039429 | 0.8083596 |
| 0.38411817 | 0.37758257 | 2.86394046 | 0.80215058 | 0.37045116 | 0.8083596 |
| -0.3187781 | -0.3079074 | 4.60584559 | 0.80212808 | 0.37045787 | 0.8083596 |
| -0.3436265 | -0.3442522 | 3.51880307 | 0.80144701 | 0.37066109 | 0.8083596 |
| -0.2981897 | -0.3007449 | 5.61143402 | 0.80035604 | 0.37098694 | 0.808461 |
| 0.32891007 | 0.30428482 | 5.10522787 | 0.79921498 | 0.37132818 | 0.8085579 |
| -0.3332626 | -0.3455494 | 3.17013708 | 0.79828503 | 0.37160661 | 0.8085579 |
| -0.7949275 | -0.7129589 | 7.26540792 | 0.79667605 | 0.37208904 | 0.8085579 |
| -0.294771 | -0.305605 | 5.62560153 | 0.79647636 | 0.37214897 | 0.8085579 |

| | | | | | |
|---|---|---|---|---|---|
| -0.308889 | -0.3043101 | 3.8558728 | 0.79437476 | 0.37278056 | 0.80932254 |
| 0.35964185 | 0.35444564 | 4.32668992 | 0.78953847 | 0.37423972 | 0.81017706 |
| -0.3281305 | -0.3276486 | 6.03937153 | 0.78781322 | 0.37476218 | 0.81017706 |
| 0.33924054 | 0.31315321 | 3.50374911 | 0.78747794 | 0.37486383 | 0.81017706 |
| 0.32542398 | 0.32344529 | 4.64186934 | 0.78733432 | 0.37490739 | 0.81017706 |
| 0.3260415 | 0.33380664 | 4.07872417 | 0.78726168 | 0.37492942 | 0.81017706 |
| 0.33604947 | 0.34827981 | 3.60496467 | 0.78457109 | 0.37574677 | 0.81017706 |
| 0.4754526 | 0.44729568 | 2.83311047 | 0.78338467 | 0.37610798 | 0.81017706 |
| 0.45544505 | 0.38854359 | 3.95391169 | 0.7830076 | 0.37622288 | 0.81017706 |
| 0.79658439 | 0.56305146 | 4.49611293 | 0.78267081 | 0.37632555 | 0.81017706 |
| -0.30723 | -0.3107404 | 5.76569695 | 0.78242543 | 0.37640038 | 0.81017706 |
| 0.3064054 | 0.30550075 | 4.72815264 | 0.78129232 | 0.37674619 | 0.81017706 |
| -0.2618509 | -0.2916397 | 3.96979899 | 0.78122547 | 0.37676661 | 0.81017706 |
| 0.32946886 | 0.31889251 | 4.07805698 | 0.78077201 | 0.37690513 | 0.81017706 |
| 0.38999025 | 0.37346713 | 2.98468992 | 0.77947271 | 0.37730245 | 0.81017706 |
| 0.35823505 | 0.3419754 | 3.22875775 | 0.77924084 | 0.37737342 | 0.81017706 |
| 0.39886356 | 0.3686472 | 2.94354708 | 0.77599318 | 0.37836938 | 0.8117131 |
| -0.5609738 | -0.477018 | 2.94963497 | 0.77230204 | 0.37950584 | 0.81354807 |
| -0.3642514 | -0.3374921 | 3.2443112 | 0.77001064 | 0.38021376 | 0.81446234 |
| -0.3428767 | -0.3443833 | 8.92903771 | 0.76743361 | 0.38101216 | 0.81452925 |
| 0.36166408 | 0.35560892 | 4.55308454 | 0.76599447 | 0.38145905 | 0.81452925 |
| -0.3226099 | -0.3350069 | 3.38146172 | 0.76546563 | 0.38162346 | 0.81452925 |
| 0.36967962 | 0.39162497 | 3.16807512 | 0.76545891 | 0.38162555 | 0.81452925 |
| -0.3423702 | -0.3368054 | 3.85363678 | 0.76109258 | 0.38298682 | 0.81452925 |
| -0.3052971 | -0.2973168 | 5.37942529 | 0.76020925 | 0.38326305 | 0.81452925 |
| 0.41434538 | 0.42078487 | 2.70751038 | 0.76019782 | 0.38326662 | 0.81452925 |
| 0.40436169 | 0.42539511 | 2.53153553 | 0.75888038 | 0.38367913 | 0.81452925 |
| -0.3128826 | -0.3354228 | 3.22192017 | 0.75885304 | 0.3836877 | 0.81452925 |
| -0.2860045 | -0.2859596 | 5.59665217 | 0.75831977 | 0.38385486 | 0.81452925 |
| 0.31233255 | 0.31865089 | 6.17010005 | 0.75711099 | 0.38423415 | 0.81452925 |
| -0.3561938 | -0.3629775 | 2.82801134 | 0.75647197 | 0.38443487 | 0.81452925 |
| -0.279864 | -0.2884503 | 6.02017615 | 0.75598378 | 0.38458832 | 0.81452925 |
| -0.3804038 | -0.3569249 | 3.6614312 | 0.75580008 | 0.38464608 | 0.81452925 |
| 0.3472668 | 0.34823386 | 3.43651706 | 0.75559229 | 0.38471143 | 0.81452925 |
| -0.2946065 | -0.3082075 | 4.58679322 | 0.75547521 | 0.38474826 | 0.81452925 |
| -0.281018 | -0.2967412 | 4.16565447 | 0.75009581 | 0.3864459 | 0.81640269 |
| 0.32279583 | 0.32014302 | 3.54871036 | 0.75000765 | 0.38647381 | 0.81640269 |
| 0.32731011 | 0.32511438 | 4.04503727 | 0.74998966 | 0.38647951 | 0.81640269 |
| 0.31475273 | 0.32501976 | 3.26731473 | 0.74666634 | 0.38753374 | 0.81803257 |
| 0.5152677 | 0.45055928 | 3.07953469 | 0.74556648 | 0.38788355 | 0.81817419 |
| 0.30821966 | 0.29968719 | 4.73714745 | 0.74373101 | 0.38846832 | 0.81881086 |
| -0.3223148 | -0.3181966 | 3.96987669 | 0.74161691 | 0.38914342 | 0.81932399 |
| 0.37125301 | 0.39383861 | 4.59567634 | 0.7404792 | 0.38950742 | 0.81932399 |
| -0.2917954 | -0.3104248 | 3.51600769 | 0.74031156 | 0.38956109 | 0.81932399 |
| 0.29250859 | 0.2904695 | 5.48951021 | 0.73906212 | 0.38996148 | 0.81944493 |
| 0.31308911 | 0.30235738 | 3.58843178 | 0.73784398 | 0.3903524 | 0.81944493 |

| | | | | | |
|---|---|---|---|---|---|
| -0.4289849 | -0.4322449 | 3.63534941 | 0.73687827 | 0.39066272 | 0.81944493 |
| -0.3723594 | -0.3674631 | 2.80760133 | 0.73510756 | 0.39123262 | 0.81944493 |
| -0.3584142 | -0.3543602 | 3.20467468 | 0.73509099 | 0.39123796 | 0.81944493 |
| 0.40561975 | 0.40254842 | 2.67922753 | 0.7348441 | 0.39131752 | 0.81944493 |
| -0.3103407 | -0.3103972 | 3.59356081 | 0.73285777 | 0.39195844 | 0.82019358 |
| -0.3312329 | -0.3318501 | 3.44645082 | 0.73027003 | 0.39279567 | 0.82135165 |
| 0.3084384 | 0.31257882 | 4.12014935 | 0.72886632 | 0.39325091 | 0.82170983 |
| -0.2790691 | -0.2967595 | 4.36284808 | 0.72782076 | 0.39359048 | 0.82182601 |
| -0.305611 | -0.2990239 | 3.64436328 | 0.72618092 | 0.39412391 | 0.82223415 |
| 0.41795528 | 0.51166648 | 3.13574111 | 0.72547402 | 0.39435418 | 0.82223415 |
| 0.30275556 | 0.28843309 | 4.29477664 | 0.72371565 | 0.39492781 | 0.82283734 |
| 0.31870271 | 0.32443187 | 3.43186092 | 0.72004379 | 0.39612955 | 0.82438247 |
| -0.3295984 | -0.3046122 | 4.51393973 | 0.71838526 | 0.39667408 | 0.82438247 |
| 0.28261788 | 0.3107046 | 5.00221351 | 0.71827574 | 0.39671008 | 0.82438247 |
| 0.28375585 | 0.28175884 | 5.94927282 | 0.71797533 | 0.39680884 | 0.82438247 |
| -0.3699909 | -0.5007472 | 5.03357471 | 0.71497202 | 0.39779815 | 0.82512583 |
| -0.3613924 | -0.3394241 | 3.57764992 | 0.71468721 | 0.39789215 | 0.82512583 |
| -0.2874744 | -0.285602 | 4.93178154 | 0.71429396 | 0.398022 | 0.82512583 |
| -0.2779605 | -0.2844205 | 5.8577612 | 0.71156962 | 0.39892323 | 0.82640216 |
| 0.30996196 | 0.2831302 | 4.17475549 | 0.70859898 | 0.39990931 | 0.82785232 |
| -0.2769665 | -0.292493 | 3.71929653 | 0.70695793 | 0.40045556 | 0.82839056 |
| 0.34215229 | 0.33981854 | 3.32219404 | 0.70551014 | 0.40093838 | 0.82873815 |
| -0.3443311 | -0.3216373 | 4.15996493 | 0.70473771 | 0.40119632 | 0.82873815 |
| 0.36713879 | 0.35451906 | 3.09522952 | 0.70183209 | 0.40216877 | 0.82998765 |
| 0.31436045 | 0.31595351 | 5.48821455 | 0.69883445 | 0.40317561 | 0.82998765 |
| -0.269518 | -0.2855976 | 4.43493535 | 0.69815924 | 0.40340291 | 0.82998765 |
| -0.4305667 | -0.3863322 | 2.81094458 | 0.69718401 | 0.40373153 | 0.82998765 |
| 0.55705248 | 0.47507472 | 2.72138237 | 0.69617621 | 0.40407153 | 0.82998765 |
| -0.2772365 | -0.2701651 | 5.79274821 | 0.69606578 | 0.40410881 | 0.82998765 |
| 0.37090667 | 0.39828684 | 2.86157055 | 0.6959561 | 0.40414585 | 0.82998765 |
| 0.5750118 | 0.46916829 | 2.77263589 | 0.69586042 | 0.40417816 | 0.82998765 |
| -0.2717765 | -0.2825334 | 4.78773007 | 0.69525594 | 0.40438237 | 0.82998765 |
| -0.3437422 | -0.3424059 | 3.24825809 | 0.69089179 | 0.40586121 | 0.83243257 |
| 0.28683021 | 0.27967875 | 5.28039649 | 0.68829343 | 0.40674546 | 0.83365535 |
| -0.3889156 | -0.3541222 | 3.27270316 | 0.68631707 | 0.40741991 | 0.83398142 |
| 0.32672799 | 0.30218612 | 3.60292941 | 0.68383001 | 0.40827099 | 0.83398142 |
| -0.2862907 | -0.2927758 | 4.07657965 | 0.68381405 | 0.40827646 | 0.83398142 |
| -0.4402956 | -0.3832111 | 3.09146323 | 0.68362283 | 0.408342 | 0.83398142 |
| -0.2940413 | -0.2865552 | 4.4072243 | 0.68361283 | 0.40834543 | 0.83398142 |
| 0.33132295 | 0.34487878 | 2.77756276 | 0.68143239 | 0.40909392 | 0.83409008 |
| -0.2855068 | -0.2881667 | 6.2214468 | 0.68082367 | 0.40930324 | 0.83409008 |
| 0.28618694 | 0.33222793 | 3.25083133 | 0.68016714 | 0.40952917 | 0.83409008 |
| 0.37211471 | 0.37246642 | 2.89328149 | 0.68010234 | 0.40955149 | 0.83409008 |
| 0.36304955 | 0.36265677 | 3.14149148 | 0.67608601 | 0.41093776 | 0.83632481 |
| -0.2673537 | -0.2923198 | 3.30941748 | 0.67337248 | 0.41187827 | 0.83731165 |
| -0.2932989 | -0.2790332 | 3.81921739 | 0.67301817 | 0.41200131 | 0.83731165 |

| | | | | | |
|---|---|---|---|---|---|
| 0.2733988 | 0.28248803 | 5.80735723 | 0.6718028 | 0.41242377 | 0.83748695 |
| -0.2704255 | -0.282577 | 3.76006858 | 0.66983849 | 0.41310792 | 0.83748695 |
| 0.31590384 | 0.29661931 | 3.19457905 | 0.66976107 | 0.41313492 | 0.83748695 |
| -0.2643423 | -0.2694593 | 6.06386168 | 0.66899438 | 0.41340243 | 0.83748695 |
| -0.3157375 | -0.3118502 | 3.79156908 | 0.6682113 | 0.41367592 | 0.83748695 |
| -0.2997462 | -0.3100875 | 3.09951146 | 0.66778796 | 0.41382389 | 0.83748695 |
| 0.34611256 | 0.34978277 | 2.74509639 | 0.6661998 | 0.41437968 | 0.8376456 |
| -0.2646363 | -0.2616941 | 6.1102131 | 0.66528953 | 0.41469873 | 0.8376456 |
| -0.3221082 | -0.3170617 | 4.0972005 | 0.66461876 | 0.41493407 | 0.8376456 |
| -0.2796698 | -0.2856924 | 5.20309191 | 0.66347879 | 0.41533448 | 0.8376456 |
| 0.38359353 | 0.41976126 | 3.06421398 | 0.66343607 | 0.4153495 | 0.8376456 |
| 0.3502103 | 0.35179754 | 3.09418467 | 0.66244738 | 0.41569725 | 0.83776313 |
| 0.28806553 | 0.27658591 | 4.97753723 | 0.66084134 | 0.41626308 | 0.83821485 |
| -0.2925238 | -0.295037 | 3.99066944 | 0.65926729 | 0.41681874 | 0.83821485 |
| -0.282289 | -0.2755452 | 5.07590597 | 0.65847109 | 0.41710022 | 0.83821485 |
| -0.3043531 | -0.3022484 | 4.54517818 | 0.65784063 | 0.41732332 | 0.83821485 |
| 0.28412157 | 0.27661439 | 4.35800316 | 0.65654314 | 0.417783 | 0.83821485 |
| 0.3198646 | 0.31313196 | 3.19278392 | 0.65647035 | 0.41780881 | 0.83821485 |
| -0.3204998 | -0.3151716 | 3.59711421 | 0.65596247 | 0.41798897 | 0.83821485 |
| -0.295119 | -0.296526 | 4.2938373 | 0.65525955 | 0.41823851 | 0.83821485 |
| -0.2905774 | -0.2874728 | 4.12247906 | 0.65331305 | 0.41893066 | 0.83837144 |
| 0.34347359 | 0.33435655 | 2.9665764 | 0.65262903 | 0.4191743 | 0.83837144 |
| -0.5449186 | -0.4116043 | 3.20222968 | 0.65079336 | 0.41982917 | 0.83837144 |
| 0.33141002 | 0.32847346 | 3.98398519 | 0.65042244 | 0.41996169 | 0.83837144 |
| 0.25827893 | 0.29935335 | 3.62412611 | 0.6501835 | 0.42004708 | 0.83837144 |
| 0.33907355 | 0.37426739 | 3.23742636 | 0.65016191 | 0.4200548 | 0.83837144 |
| 0.26352771 | 0.26399404 | 6.04343709 | 0.6484645 | 0.4206622 | 0.83846135 |
| -0.3070713 | -0.3146982 | 3.30518824 | 0.64841677 | 0.4206793 | 0.83846135 |
| 0.27026882 | 0.28187371 | 4.06980027 | 0.64739386 | 0.42104599 | 0.83848275 |
| 0.36498087 | 0.34380914 | 2.68144749 | 0.64677102 | 0.42126949 | 0.83848275 |
| -0.2664456 | -0.2806327 | 4.10653326 | 0.64556491 | 0.42170282 | 0.83876835 |
| 0.28670217 | 0.30117455 | 3.56839 | 0.64422597 | 0.42218464 | 0.83914997 |
| -0.3074215 | -0.3018075 | 4.05451057 | 0.63996301 | 0.42372419 | 0.84068667 |
| -0.3109691 | -0.2802156 | 4.72287803 | 0.6399418 | 0.42373187 | 0.84068667 |
| 0.31040858 | 0.28843655 | 3.73128127 | 0.63952425 | 0.42388312 | 0.84068667 |
| 0.32382118 | 0.31526453 | 3.11789031 | 0.63887149 | 0.42411974 | 0.84068667 |
| 0.33141733 | 0.33781648 | 3.05558895 | 0.63711304 | 0.42475814 | 0.84137582 |
| 0.3154527 | 0.3131719 | 3.35971193 | 0.63375262 | 0.42598216 | 0.84264144 |
| 0.32496104 | 0.3452379 | 2.91494391 | 0.63373967 | 0.42598689 | 0.84264144 |
| 0.27647325 | 0.26612658 | 4.15793123 | 0.6329629 | 0.42627058 | 0.84264144 |
| -0.3250858 | -0.3003918 | 3.54992065 | 0.63099434 | 0.42699082 | 0.84271841 |
| 0.29430503 | 0.29821264 | 4.32297235 | 0.62996799 | 0.42736706 | 0.84271841 |
| 0.29369259 | 0.29344565 | 5.17857504 | 0.62980962 | 0.42742515 | 0.84271841 |
| -0.3184301 | -0.307115 | 3.59341799 | 0.62907416 | 0.42769512 | 0.84271841 |
| -0.3272592 | -0.3420213 | 3.40210885 | 0.62888255 | 0.4277655 | 0.84271841 |
| 0.50194115 | 0.41338246 | 2.95264453 | 0.6277174 | 0.42819383 | 0.84298839 |

| | | | | | |
|---|---|---|---|---|---|
| 0.27907038 | 0.34215885 | 3.68932046 | 0.62433324 | 0.42944157 | 0.84448665 |
| 0.24917069 | 0.27274593 | 4.49248851 | 0.62316687 | 0.42987289 | 0.84448665 |
| 0.38428565 | 0.39066017 | 2.41146795 | 0.62284482 | 0.42999209 | 0.84448665 |
| -0.312528 | -0.3339689 | 2.91073699 | 0.62249378 | 0.43012209 | 0.84448665 |
| 0.28562624 | 0.29261528 | 3.73391934 | 0.62110934 | 0.43063535 | 0.84466331 |
| -0.2633379 | -0.2662579 | 3.94945384 | 0.62067706 | 0.4307958 | 0.84466331 |
| -0.2714519 | -0.2884052 | 3.56183172 | 0.61714754 | 0.43210926 | 0.84666499 |
| -0.299235 | -0.3137168 | 3.17488733 | 0.61415088 | 0.4332292 | 0.84734257 |
| -0.2642097 | -0.2672754 | 6.33143925 | 0.61237739 | 0.43389408 | 0.84734257 |
| 0.31449095 | 0.28966847 | 3.29351369 | 0.6121203 | 0.4339906 | 0.84734257 |
| -0.2855076 | -0.2849764 | 6.00653691 | 0.61039013 | 0.43464096 | 0.84734257 |
| 0.35306081 | 0.35119241 | 2.91408661 | 0.61002337 | 0.43477901 | 0.84734257 |
| -0.2608066 | -0.2670861 | 5.74156689 | 0.60929761 | 0.43505239 | 0.84734257 |
| -0.7590987 | -0.4478982 | 3.28738738 | 0.60830118 | 0.43542817 | 0.84734257 |
| 0.3774061 | 0.35364369 | 2.62831494 | 0.60820153 | 0.43546577 | 0.84734257 |
| 0.2748274 | 0.27672856 | 5.7165659 | 0.60798406 | 0.43554786 | 0.84734257 |
| -0.284041 | -0.2858774 | 4.44765378 | 0.60790921 | 0.43557612 | 0.84734257 |
| 0.28863399 | 0.27489757 | 4.4405646 | 0.60758311 | 0.43569927 | 0.84734257 |
| 0.27056944 | 0.26568889 | 4.96632586 | 0.60687044 | 0.43596858 | 0.84734257 |
| 0.28661856 | 0.26253884 | 3.72974348 | 0.60468438 | 0.43679628 | 0.84838151 |
| -0.250825 | -0.2708125 | 4.59134974 | 0.60375221 | 0.43714996 | 0.84849898 |
| -0.2424368 | -0.2654591 | 3.77393353 | 0.60273172 | 0.43753765 | 0.84868227 |
| 0.28365935 | 0.28524957 | 3.88338115 | 0.60194703 | 0.43783611 | 0.84869237 |
| 0.34913719 | 0.32067002 | 4.04071303 | 0.60003813 | 0.43856348 | 0.84951704 |
| -0.2430064 | -0.2534661 | 6.10170405 | 0.59929106 | 0.43884864 | 0.84951704 |
| -0.248894 | -0.2624576 | 5.20022654 | 0.59698736 | 0.4397298 | 0.85065377 |
| -0.299 | -0.2989437 | 3.7489362 | 0.59469198 | 0.44061047 | 0.85136645 |
| 0.34030027 | 0.32401519 | 2.91790774 | 0.59435317 | 0.44074069 | 0.85136645 |
| 0.31156935 | 0.32713646 | 3.28603588 | 0.59372899 | 0.44098075 | 0.85136645 |
| 0.38247686 | 0.39475596 | 2.76180776 | 0.59294892 | 0.44128105 | 0.85137824 |
| -0.2692403 | -0.2757013 | 7.11104011 | 0.59141478 | 0.44187255 | 0.85160888 |
| 0.26920544 | 0.25946051 | 5.75774154 | 0.58891674 | 0.44283832 | 0.85160888 |
| -0.412206 | -0.3260575 | 3.82550368 | 0.58734416 | 0.44344796 | 0.85160888 |
| -0.3352702 | -0.3037956 | 3.72056115 | 0.58675279 | 0.44367755 | 0.85160888 |
| 0.28691003 | 0.26703107 | 3.47812148 | 0.58639826 | 0.44381528 | 0.85160888 |
| -0.3163119 | -0.3230504 | 3.05468263 | 0.58639497 | 0.44381656 | 0.85160888 |
| -0.3115428 | -0.2958975 | 4.78688364 | 0.58626103 | 0.44386861 | 0.85160888 |
| -0.3051952 | -0.2991615 | 3.51440957 | 0.58519736 | 0.4442823 | 0.85160888 |
| 0.2570061 | 0.28131833 | 3.44948361 | 0.58446161 | 0.44456881 | 0.85160888 |
| 0.28055409 | 0.28567583 | 3.95756262 | 0.58444297 | 0.44457607 | 0.85160888 |
| 0.27661624 | 0.27035286 | 5.35461518 | 0.58428523 | 0.44463753 | 0.85160888 |
| 0.24800331 | 0.26516728 | 4.42004012 | 0.58161004 | 0.44568193 | 0.85267528 |
| -0.2836289 | -0.2827771 | 4.13726181 | 0.58096087 | 0.44593594 | 0.85267528 |
| -0.2831667 | -0.2833258 | 4.5113648 | 0.57877995 | 0.44679094 | 0.85267528 |
| 0.24977336 | 0.29262401 | 3.37168864 | 0.57649332 | 0.44769012 | 0.85267528 |
| 0.6592233 | 0.46373061 | 2.68915435 | 0.57561733 | 0.44803533 | 0.85267528 |

| | | | | | |
|---|---|---|---|---|---|
| 0.30341355 | 0.30582633 | 3.37607112 | 0.57387157 | 0.44872455 | 0.85267528 |
| -0.293211 | -0.2986219 | 3.27173151 | 0.57372347 | 0.44878309 | 0.85267528 |
| 0.29650359 | 0.29778522 | 3.63720973 | 0.57197436 | 0.44947542 | 0.85267528 |
| 0.37436083 | 0.44127923 | 3.34842243 | 0.57185736 | 0.44952179 | 0.85267528 |
| -0.2977583 | -0.2934682 | 5.37442163 | 0.57151867 | 0.44965607 | 0.85267528 |
| 0.27973984 | 0.27071067 | 3.72336287 | 0.57079865 | 0.44994173 | 0.85267528 |
| 0.33184857 | 0.29292443 | 3.09668964 | 0.57066562 | 0.44999454 | 0.85267528 |
| 0.27675044 | 0.28616243 | 3.60392305 | 0.57013189 | 0.45020651 | 0.85267528 |
| 0.30338152 | 0.2986445 | 4.03000544 | 0.56826022 | 0.45095109 | 0.85267528 |
| -0.2686575 | -0.2680056 | 4.11751538 | 0.56577644 | 0.45194215 | 0.85267528 |
| -0.2510244 | -0.2459064 | 6.08203707 | 0.56556807 | 0.45202544 | 0.85267528 |
| 0.3117421 | 0.30779249 | 3.29687697 | 0.56552499 | 0.45204267 | 0.85267528 |
| -0.3193668 | -0.307791 | 3.18615389 | 0.56519428 | 0.45217493 | 0.85267528 |
| 0.33737269 | 0.30594723 | 2.82437809 | 0.56442454 | 0.452483 | 0.85267528 |
| 0.32318873 | 0.29737832 | 2.98573651 | 0.56383564 | 0.45271892 | 0.85267528 |
| -0.3215427 | -0.3007942 | 6.21999426 | 0.56328374 | 0.45294019 | 0.85267528 |
| 0.30297522 | 0.3039586 | 9.13863165 | 0.56311653 | 0.45300726 | 0.85267528 |
| 0.31851173 | 0.3189353 | 8.34109765 | 0.56212808 | 0.45340407 | 0.85267528 |
| -0.3499663 | -0.320956 | 3.28113487 | 0.56178916 | 0.45354026 | 0.85267528 |
| 0.30863646 | 0.30198888 | 2.84876259 | 0.56074918 | 0.45395855 | 0.85267528 |
| 0.3053009 | 0.2970544 | 3.18724459 | 0.560642 | 0.45400169 | 0.85267528 |
| -0.2314827 | -0.2678246 | 3.29676436 | 0.55942871 | 0.45449052 | 0.85267528 |
| 0.30407743 | 0.29611272 | 3.43716709 | 0.55932006 | 0.45453433 | 0.85267528 |
| 0.28770497 | 0.28558028 | 3.72455868 | 0.5582258 | 0.45497598 | 0.85267528 |
| -0.2837827 | -0.2650114 | 3.94918062 | 0.55816486 | 0.4550006 | 0.85267528 |
| -0.2413402 | -0.2448555 | 4.18409601 | 0.55812458 | 0.45501687 | 0.85267528 |
| 0.26507333 | 0.26447426 | 5.97775665 | 0.55750278 | 0.45526816 | 0.85267528 |
| 0.241241 | 0.24318228 | 5.85446305 | 0.55744283 | 0.4552924 | 0.85267528 |
| -0.237726 | -0.2496856 | 5.83688758 | 0.5568001 | 0.45555241 | 0.85267528 |
| -0.3666453 | -0.3366514 | 3.19315844 | 0.55678039 | 0.45556039 | 0.85267528 |
| -0.2288749 | -0.2464795 | 3.87142839 | 0.55459638 | 0.45644568 | 0.85267528 |
| 0.28086625 | 0.28291068 | 7.42822718 | 0.55247207 | 0.45730936 | 0.85267528 |
| 0.43313604 | 0.36171469 | 3.57819005 | 0.55213699 | 0.45744583 | 0.85267528 |
| -0.2828089 | -0.3012653 | 3.96013233 | 0.55202907 | 0.4574898 | 0.85267528 |
| 0.33645841 | 0.31742624 | 2.85379097 | 0.55194858 | 0.4575226 | 0.85267528 |
| 0.27865832 | 0.27071469 | 3.4429031 | 0.55135924 | 0.45776284 | 0.85267528 |
| -0.2897362 | -0.282511 | 4.65704509 | 0.55044576 | 0.45813561 | 0.85267528 |
| -0.2960962 | -0.3070133 | 3.03596993 | 0.54876894 | 0.45882114 | 0.85267528 |
| 0.26202499 | 0.31578565 | 3.5097852 | 0.54864627 | 0.45887136 | 0.85267528 |
| 0.35313843 | 0.35211804 | 2.66477893 | 0.54837656 | 0.45898179 | 0.85267528 |
| 0.24757887 | 0.26647637 | 3.98437138 | 0.54832192 | 0.45900417 | 0.85267528 |
| 0.29244832 | 0.29455644 | 2.97638564 | 0.54718633 | 0.45946964 | 0.85267528 |
| 0.26465536 | 0.24711175 | 3.86772579 | 0.54705795 | 0.45952231 | 0.85267528 |
| -0.3074239 | -0.2917324 | 3.78600645 | 0.54608288 | 0.45992266 | 0.85267528 |
| -0.2980086 | -0.3040891 | 3.10769203 | 0.54607453 | 0.45992609 | 0.85267528 |
| -0.2452503 | -0.2535605 | 5.69442025 | 0.54435756 | 0.4606324 | 0.85343801 |

| | | | | | |
|---|---|---|---|---|---|
| -0.2857347 | -0.2798208 | 6.36151353 | 0.5425711 | 0.46136913 | 0.85379852 |
| 0.25621345 | 0.25246189 | 4.16957722 | 0.54159783 | 0.46177129 | 0.85379852 |
| -0.4734642 | -0.3596534 | 3.00865307 | 0.54106041 | 0.46199359 | 0.85379852 |
| 0.40353673 | 0.34969982 | 3.74004735 | 0.54040002 | 0.46226699 | 0.85379852 |
| -0.2679066 | -0.2705692 | 4.75066161 | 0.54003168 | 0.4624196 | 0.85379852 |
| -0.2549056 | -0.2585869 | 4.21894843 | 0.53942982 | 0.46266912 | 0.85379852 |
| -0.2811336 | -0.2724179 | 3.70021329 | 0.53889232 | 0.46289215 | 0.85379852 |
| 0.30710361 | 0.30913527 | 3.03039722 | 0.53811695 | 0.46321417 | 0.85384829 |
| 0.33895199 | 0.35136833 | 3.06460444 | 0.53585224 | 0.46415679 | 0.85504122 |
| 0.25531843 | 0.24506966 | 4.13416458 | 0.53245162 | 0.46557797 | 0.85594993 |
| 0.23686626 | 0.26339877 | 3.43113261 | 0.529417 | 0.46685207 | 0.85594993 |
| 0.31962135 | 0.29718481 | 2.84065865 | 0.52922701 | 0.46693202 | 0.85594993 |
| 0.31313015 | 0.31773111 | 2.72441824 | 0.52884896 | 0.46709118 | 0.85594993 |
| -0.2664768 | -0.2657673 | 4.45908579 | 0.5287836 | 0.46711871 | 0.85594993 |
| -0.2303359 | -0.2387961 | 5.61211854 | 0.52759122 | 0.46762133 | 0.85594993 |
| 0.25347131 | 0.25952049 | 3.54821244 | 0.5270874 | 0.46783396 | 0.85594993 |
| -0.2709738 | -0.2647085 | 4.69810738 | 0.52591083 | 0.46833112 | 0.85594993 |
| 0.24861256 | 0.26262114 | 4.80698687 | 0.52461563 | 0.4688794 | 0.85594993 |
| -0.2757558 | -0.2660887 | 3.72493748 | 0.52374279 | 0.46924947 | 0.85594993 |
| 0.22784746 | 0.23495907 | 5.93294847 | 0.5229218 | 0.46959798 | 0.85594993 |
| -0.2954666 | -0.2861752 | 3.51079663 | 0.52188942 | 0.47003682 | 0.85594993 |
| -0.2534498 | -0.2504555 | 3.8658905 | 0.52112669 | 0.47036146 | 0.85594993 |
| 0.23983401 | 0.26358198 | 4.77945563 | 0.5203781 | 0.47068044 | 0.85594993 |
| 0.37325421 | 0.36815835 | 2.67197941 | 0.5202796 | 0.47072244 | 0.85594993 |
| 0.26028087 | 0.25843776 | 4.92353202 | 0.51930334 | 0.47113901 | 0.85594993 |
| -0.2511283 | -0.2422022 | 4.09407243 | 0.51827107 | 0.47158013 | 0.85594993 |
| 0.23740272 | 0.26033787 | 3.81354248 | 0.5176792 | 0.47183335 | 0.85594993 |
| 0.24907494 | 0.2472072 | 5.70453324 | 0.51716369 | 0.47205409 | 0.85594993 |
| 0.26773336 | 0.25818979 | 3.51796305 | 0.51690241 | 0.47216603 | 0.85594993 |
| -0.2308869 | -0.2403652 | 3.72363466 | 0.51675458 | 0.47222938 | 0.85594993 |
| 0.26050969 | 0.26205951 | 4.00871326 | 0.51612117 | 0.47250099 | 0.85594993 |
| -0.2440541 | -0.2497499 | 7.34268329 | 0.51468412 | 0.47311814 | 0.85594993 |
| -0.291419 | -0.2903788 | 3.30000201 | 0.51378761 | 0.47350381 | 0.85594993 |
| -0.2493385 | -0.2567953 | 4.51398366 | 0.51313758 | 0.47378377 | 0.85594993 |
| -0.3376454 | -0.2844246 | 3.91431888 | 0.51240771 | 0.47409843 | 0.85594993 |
| 0.34511634 | 0.30337268 | 2.72497915 | 0.51195895 | 0.47429207 | 0.85594993 |
| 0.25527301 | 0.26094366 | 4.74206554 | 0.51181245 | 0.47435531 | 0.85594993 |
| -0.2694607 | -0.2587169 | 4.57076937 | 0.51119307 | 0.47462284 | 0.85594993 |
| -0.2396488 | -0.2348655 | 6.28874449 | 0.51106991 | 0.47467607 | 0.85594993 |
| 0.2771979 | 0.27996937 | 4.66965277 | 0.51053299 | 0.47490822 | 0.85594993 |
| 0.2779309 | 0.2717323 | 3.02376034 | 0.51031604 | 0.47500208 | 0.85594993 |
| -0.2368802 | -0.2468698 | 4.02525063 | 0.51004632 | 0.47511881 | 0.85594993 |
| -0.2427918 | -0.2351212 | 5.73101722 | 0.50928351 | 0.47544919 | 0.85594993 |
| -0.2485962 | -0.389379 | 4.67937385 | 0.50892576 | 0.47560426 | 0.85594993 |
| 0.26549549 | 0.26506979 | 4.08226744 | 0.50857792 | 0.47575511 | 0.85594993 |
| 0.25919844 | 0.26321141 | 4.92206167 | 0.50789367 | 0.4760521 | 0.85594993 |

| | | | | | |
|---|---|---|---|---|---|
| 0.25746999 | 0.33529131 | 3.26193442 | 0.50770699 | 0.47613317 | 0.85594993 |
| -0.3179159 | -0.3064549 | 2.81643917 | 0.50692037 | 0.47647505 | 0.85594993 |
| 0.28162365 | 0.28534072 | 4.32752634 | 0.50690724 | 0.47648076 | 0.85594993 |
| -0.3045377 | -0.3793757 | 2.78791378 | 0.50524733 | 0.47720352 | 0.85601483 |
| -0.2469191 | -0.2504063 | 4.0114248 | 0.50483279 | 0.4773843 | 0.85601483 |
| -0.2579435 | -0.2552822 | 7.76259131 | 0.50478703 | 0.47740426 | 0.85601483 |
| -0.2934072 | -0.3331292 | 3.19576971 | 0.50190511 | 0.47866423 | 0.85691595 |
| -0.2619823 | -0.2606767 | 4.27937121 | 0.50179973 | 0.4787104 | 0.85691595 |
| -0.2293342 | -0.2376185 | 3.83013194 | 0.50160643 | 0.47879512 | 0.85691595 |
| -0.2251026 | -0.2411463 | 4.97889042 | 0.50091929 | 0.47909647 | 0.85692533 |
| -0.2543918 | -0.2569463 | 4.12668519 | 0.50014631 | 0.47943584 | 0.85700267 |
| 0.22655809 | 0.24938839 | 5.44796057 | 0.49926438 | 0.47982352 | 0.85716622 |
| 0.25592079 | 0.26725386 | 5.1649836 | 0.49747435 | 0.48061198 | 0.85804507 |
| -0.3047709 | -0.3040437 | 3.08714152 | 0.49600071 | 0.48126266 | 0.85835298 |
| 0.25923829 | 0.2538241 | 4.06459706 | 0.49552994 | 0.48147084 | 0.85835298 |
| 0.24663272 | 0.2489331 | 3.57874665 | 0.49507029 | 0.48167424 | 0.85835298 |
| 0.25302399 | 0.24032494 | 4.82774941 | 0.4934871 | 0.4823759 | 0.85907437 |
| -0.2557845 | -0.2454441 | 4.88469955 | 0.49208196 | 0.48300006 | 0.85965693 |
| 0.23241583 | 0.24275543 | 4.65879523 | 0.49052723 | 0.48369221 | 0.85986994 |
| -0.2226079 | -0.2446287 | 4.52199443 | 0.48996325 | 0.4839437 | 0.85986994 |
| -0.2358336 | -0.2453207 | 5.13854221 | 0.4895057 | 0.48414788 | 0.85986994 |
| 0.26750054 | 0.27227304 | 4.50296953 | 0.48854896 | 0.48457529 | 0.85986994 |
| 0.38423839 | 0.35121242 | 2.74335423 | 0.48774425 | 0.48493527 | 0.85986994 |
| -0.3033862 | -0.2987586 | 3.05148248 | 0.48726474 | 0.48514999 | 0.85986994 |
| 0.26462669 | 0.29040202 | 4.25028125 | 0.48649906 | 0.48549316 | 0.85986994 |
| 0.2356024 | 0.24897699 | 4.22358534 | 0.48609815 | 0.48567301 | 0.85986994 |
| 0.24557664 | 0.26859254 | 3.96709809 | 0.48582896 | 0.48579383 | 0.85986994 |
| 0.28779247 | 0.28980619 | 4.06556227 | 0.48479195 | 0.48625974 | 0.86016851 |
| 0.25115961 | 0.22772684 | 4.04991811 | 0.48381052 | 0.48670136 | 0.86042379 |
| 0.25971617 | 0.25792123 | 4.23617927 | 0.48181549 | 0.48760112 | 0.86094427 |
| -0.2401697 | -0.253415 | 3.24141257 | 0.48119026 | 0.48788367 | 0.86094427 |
| 0.46212536 | 0.40097505 | 2.96584175 | 0.48118013 | 0.48788825 | 0.86094427 |
| -0.2570234 | -0.2512319 | 6.00939811 | 0.47908164 | 0.48883859 | 0.86112174 |
| -0.3426187 | -0.3238009 | 2.76448455 | 0.47868623 | 0.48901801 | 0.86112174 |
| 0.25618919 | 0.24254512 | 4.39513119 | 0.47865026 | 0.48903433 | 0.86112174 |
| -0.2360736 | -0.2427122 | 4.1675961 | 0.47833153 | 0.48917904 | 0.86112174 |
| 0.24624092 | 0.25271164 | 4.28873104 | 0.47603995 | 0.49022155 | 0.86232286 |
| -0.2609263 | -0.2580659 | 3.95501421 | 0.47552287 | 0.4904573 | 0.86232286 |
| -0.2344372 | -0.2355911 | 5.10876454 | 0.47427893 | 0.49102522 | 0.86257461 |
| 0.24569293 | 0.24658402 | 7.08836049 | 0.47390402 | 0.4911966 | 0.86257461 |
| -0.2864722 | -0.2801306 | 3.65707586 | 0.47306496 | 0.49158051 | 0.86272529 |
| -0.1943098 | -0.2227808 | 4.08065178 | 0.47212476 | 0.4920113 | 0.862958 |
| -0.2540643 | -0.2521976 | 4.22208397 | 0.46793535 | 0.49393854 | 0.86520494 |
| -0.247719 | -0.2482503 | 5.1005737 | 0.46575454 | 0.49494679 | 0.86520494 |
| -0.2598651 | -0.2362143 | 3.8775217 | 0.46558959 | 0.49502319 | 0.86520494 |
| 0.23514164 | 0.24488563 | 5.25278031 | 0.46554128 | 0.49504557 | 0.86520494 |

| | | | | | |
|---|---|---|---|---|---|
| -0.270688 | -0.2779729 | 3.11368156 | 0.4648038 | 0.49538743 | 0.86520494 |
| -0.2490647 | -0.2541161 | 3.31861219 | 0.46393684 | 0.49578981 | 0.86520494 |
| -0.263022 | -0.2628016 | 3.41997091 | 0.46391019 | 0.49580219 | 0.86520494 |
| -0.3666786 | -0.3026147 | 3.21444958 | 0.46390927 | 0.49580261 | 0.86520494 |
| 0.28936303 | 0.28350701 | 3.2534119 | 0.46307702 | 0.49618942 | 0.86520494 |
| -0.2848152 | -0.2812722 | 3.41956159 | 0.46235495 | 0.49652544 | 0.86520494 |
| 0.27031218 | 0.29119344 | 3.16507862 | 0.46223563 | 0.496581 | 0.86520494 |
| -0.2363259 | -0.2570806 | 3.25647685 | 0.46088411 | 0.49721107 | 0.8657815 |
| 0.25415939 | 0.23970386 | 4.0144548 | 0.45940538 | 0.497902 | 0.86646326 |
| -0.2268398 | -0.2360706 | 5.31306614 | 0.45709996 | 0.49898244 | 0.86782162 |
| -0.2588818 | -0.2493582 | 3.95012721 | 0.4549514 | 0.49999294 | 0.86891588 |
| 0.27184423 | 0.26996303 | 4.82652425 | 0.45448637 | 0.50021211 | 0.86891588 |
| 0.29804178 | 0.29834889 | 3.00299544 | 0.45276894 | 0.50102296 | 0.86973684 |
| 0.26297303 | 0.25812929 | 3.70038926 | 0.45221326 | 0.50128578 | 0.86973684 |
| 0.31189169 | 0.32411683 | 3.09369817 | 0.45067801 | 0.50201316 | 0.87047698 |
| -0.2562717 | -0.2640941 | 3.22481557 | 0.44942954 | 0.50260598 | 0.87052489 |
| -0.2264529 | -0.2327795 | 5.95649999 | 0.44935293 | 0.5026424 | 0.87052489 |
| 0.20585674 | 0.3613519 | 2.60761515 | 0.44566874 | 0.50439901 | 0.8714593 |
| 0.28599143 | 0.24954486 | 3.35491503 | 0.44502701 | 0.50470606 | 0.8714593 |
| 0.27261903 | 0.29257467 | 3.08903423 | 0.4449824 | 0.50472741 | 0.8714593 |
| -0.2598861 | -0.2555003 | 3.87256536 | 0.44472959 | 0.50484847 | 0.8714593 |
| 0.23407326 | 0.23429824 | 5.6356115 | 0.44432052 | 0.50504446 | 0.8714593 |
| 0.31049028 | 0.31075428 | 2.8753693 | 0.44416511 | 0.50511895 | 0.8714593 |
| 0.25817616 | 0.23703734 | 3.63630956 | 0.44380882 | 0.50528981 | 0.8714593 |
| -0.2980357 | -0.2841196 | 2.99634542 | 0.44155151 | 0.50637456 | 0.87261933 |
| 0.24437889 | 0.23030638 | 3.52684595 | 0.44115509 | 0.50656547 | 0.87261933 |
| -0.2348048 | -0.2353437 | 5.88118294 | 0.44026328 | 0.50699541 | 0.8728396 |
| -0.2707977 | -0.2558351 | 4.50387587 | 0.43963941 | 0.50729655 | 0.8728396 |
| -0.2552566 | -0.2453724 | 4.3729739 | 0.4379532 | 0.50811202 | 0.87372322 |
| -0.2521636 | -0.2807486 | 4.6172466 | 0.43487013 | 0.50960888 | 0.87577677 |
| -0.2053823 | -0.2463749 | 3.10081782 | 0.43278676 | 0.51062469 | 0.87700169 |
| -0.2100287 | -0.2333007 | 3.26281859 | 0.4308642 | 0.5115652 | 0.87705027 |
| 0.2692336 | 0.26095398 | 2.89586361 | 0.43012438 | 0.51192792 | 0.87705027 |
| -0.0697897 | -0.3892046 | 4.05910598 | 0.430013 | 0.51198257 | 0.87705027 |
| 0.2754177 | 0.25044649 | 3.26202405 | 0.42898192 | 0.51248893 | 0.87705027 |
| -0.2401099 | -0.2419601 | 8.32080228 | 0.42853267 | 0.51270982 | 0.87705027 |
| -0.2407808 | -0.2387684 | 3.76219043 | 0.42852191 | 0.51271512 | 0.87705027 |
| 0.23855711 | 0.24759193 | 3.7286565 | 0.42823442 | 0.51285657 | 0.87705027 |
| 0.26609079 | 0.27279976 | 3.27231816 | 0.42778578 | 0.51307744 | 0.87705027 |
| 0.26388332 | 0.26797257 | 3.27472481 | 0.42638596 | 0.51376765 | 0.87771168 |
| 0.24106073 | 0.22260987 | 3.68941589 | 0.4250633 | 0.51442131 | 0.87830989 |
| -0.1979978 | -0.2214567 | 3.8533108 | 0.42416475 | 0.51486619 | 0.87846422 |
| -0.2343418 | -0.2515049 | 3.12398239 | 0.42365518 | 0.51511879 | 0.87846422 |
| -0.2267877 | -0.2323799 | 4.45341106 | 0.4226847 | 0.51560046 | 0.87876781 |
| -0.3507372 | -0.3224847 | 2.50928603 | 0.4208785 | 0.51649902 | 0.87978114 |
| 0.30152294 | 0.2885519 | 2.83963371 | 0.41717784 | 0.51834862 | 0.88210261 |

| | | | | | |
|---|---|---|---|---|---|
| -0.2507021 | -0.2666546 | 3.69848627 | 0.41501759 | 0.5194337 | 0.88210261 |
| -0.2761041 | -0.2759834 | 3.08035698 | 0.41487575 | 0.51950508 | 0.88210261 |
| -0.2065879 | -0.2225371 | 4.28642705 | 0.41420635 | 0.51984221 | 0.88210261 |
| 0.2160976 | 0.22011108 | 6.28683417 | 0.41375881 | 0.52006782 | 0.88210261 |
| 0.26934285 | 0.27382912 | 3.62035296 | 0.41319427 | 0.52035265 | 0.88210261 |
| -0.2279347 | -0.2579942 | 3.01996836 | 0.41196627 | 0.52097319 | 0.88210261 |
| 0.23905955 | 0.25592671 | 3.21883763 | 0.41173981 | 0.52108777 | 0.88210261 |
| -0.2255507 | -0.228138 | 6.06581855 | 0.41055127 | 0.52168983 | 0.88210261 |
| 0.21424741 | 0.26109471 | 2.97419876 | 0.41053294 | 0.52169913 | 0.88210261 |
| -0.2654711 | -0.2531461 | 4.60111542 | 0.40937554 | 0.52228663 | 0.88210261 |
| 0.25995809 | 0.24726925 | 3.43738985 | 0.40920514 | 0.52237322 | 0.88210261 |
| -0.215589 | -0.2076158 | 6.05517621 | 0.40878302 | 0.52258784 | 0.88210261 |
| 0.24785011 | 0.22644034 | 4.67124703 | 0.40863325 | 0.52266403 | 0.88210261 |
| -0.2218005 | -0.2222094 | 3.88473447 | 0.40806191 | 0.52295485 | 0.88210261 |
| -0.248876 | -0.2334248 | 3.67853154 | 0.40791216 | 0.52303112 | 0.88210261 |
| 0.30058587 | 0.30153193 | 2.7974753 | 0.40788772 | 0.52304357 | 0.88210261 |
| -0.2167717 | -0.21621 | 5.49990935 | 0.40637161 | 0.5238169 | 0.88263404 |
| 0.23789538 | 0.23804305 | 6.12492823 | 0.4058104 | 0.52410368 | 0.88263404 |
| 0.22408619 | 0.21594469 | 6.73923858 | 0.40397728 | 0.52504234 | 0.88263404 |
| -0.2085422 | -0.2079757 | 5.4468948 | 0.40379309 | 0.52513682 | 0.88263404 |
| -0.229097 | -0.2342132 | 3.44276569 | 0.40277638 | 0.52565889 | 0.88263404 |
| 0.24654193 | 0.24712753 | 3.77299276 | 0.40264338 | 0.52572725 | 0.88263404 |
| 0.22178476 | 0.22027135 | 4.57271778 | 0.40249176 | 0.52580521 | 0.88263404 |
| 0.24715564 | 0.24104121 | 3.53324413 | 0.40141557 | 0.5263591 | 0.88263404 |
| -0.2012756 | -0.2110434 | 6.58830655 | 0.40121518 | 0.52646236 | 0.88263404 |
| 0.23389136 | 0.23514562 | 3.63808384 | 0.40083884 | 0.52665636 | 0.88263404 |
| -0.1827605 | -0.2120575 | 3.80161229 | 0.40072797 | 0.52671354 | 0.88263404 |
| -0.2288681 | -0.2315767 | 4.98301604 | 0.39988655 | 0.52714785 | 0.88285063 |
| 0.20500971 | 0.21252206 | 5.17720239 | 0.39881423 | 0.52770227 | 0.88326801 |
| 0.30809301 | 0.32395554 | 2.68631752 | 0.397632 | 0.52831473 | 0.88331685 |
| -0.2518456 | -0.2964474 | 3.12648627 | 0.397041 | 0.52862138 | 0.88331685 |
| 0.23043187 | 0.20939811 | 4.37780055 | 0.39698615 | 0.52864986 | 0.88331685 |
| 0.30344936 | 0.28982585 | 3.17808778 | 0.39550091 | 0.52942197 | 0.88331685 |
| 0.18702461 | 0.21258432 | 3.98739827 | 0.39521893 | 0.52956879 | 0.88331685 |
| 0.2687112 | 0.27300184 | 3.21977609 | 0.39449954 | 0.52994368 | 0.88331685 |
| 0.19756656 | 0.25482216 | 2.92498392 | 0.39439768 | 0.52999681 | 0.88331685 |
| 0.23267596 | 0.22427133 | 4.41638067 | 0.3935858 | 0.53042055 | 0.88331685 |
| 0.26305045 | 0.25811917 | 4.44670524 | 0.39347493 | 0.53047847 | 0.88331685 |
| -0.2515436 | -0.2380119 | 3.78070942 | 0.39187658 | 0.53131464 | 0.88420044 |
| 0.30349034 | 0.27449077 | 2.76903564 | 0.39124329 | 0.5316466 | 0.8842444 |
| -0.22671 | -0.240459 | 3.30385973 | 0.39010336 | 0.53224507 | 0.88427306 |
| -0.2048003 | -0.2342485 | 3.08181594 | 0.38921172 | 0.53271405 | 0.88427306 |
| -0.2046543 | -0.2128351 | 6.54812565 | 0.38873535 | 0.53296491 | 0.88427306 |
| 0.22614793 | 0.22804123 | 4.08593908 | 0.38873241 | 0.53296645 | 0.88427306 |
| -0.2803836 | -0.2805916 | 2.86328994 | 0.38830521 | 0.5331916 | 0.88427306 |
| 0.23534438 | 0.23868283 | 3.94132927 | 0.38602855 | 0.53439439 | 0.88505187 |

| | | | | | |
|---|---|---|---|---|---|
| 0.22988573 | 0.23543548 | 4.25556188 | 0.38544634 | 0.53470277 | 0.88505187 |
| -0.2137228 | -0.222847 | 4.72912911 | 0.38435168 | 0.53528344 | 0.88505187 |
| 0.22584595 | 0.24953171 | 2.87446445 | 0.38326501 | 0.53586102 | 0.88505187 |
| 0.27522192 | 0.26664203 | 3.49362299 | 0.38319911 | 0.53589608 | 0.88505187 |
| 0.28163906 | 0.26488189 | 2.78290153 | 0.38305115 | 0.53597482 | 0.88505187 |
| -0.216611 | -0.2175171 | 4.93486033 | 0.38300577 | 0.53599897 | 0.88505187 |
| 0.2371661 | 0.23712546 | 4.10392168 | 0.38246751 | 0.53628562 | 0.88505187 |
| 0.21413254 | 0.21378925 | 5.16085757 | 0.38221969 | 0.53641768 | 0.88505187 |
| 0.2277968 | 0.24838056 | 3.16107761 | 0.38144117 | 0.53683296 | 0.88505187 |
| -0.1904106 | -0.2065447 | 4.00378468 | 0.38108103 | 0.53702525 | 0.88505187 |
| -0.2444876 | -0.2622271 | 2.6860795 | 0.37825841 | 0.53853678 | 0.88703782 |
| -0.2195185 | -0.2230481 | 4.73805821 | 0.37730646 | 0.53904831 | 0.88737532 |
| 0.23324666 | 0.21196259 | 3.9725363 | 0.37474874 | 0.54042711 | 0.88848777 |
| -0.2206519 | -0.2040595 | 5.84095171 | 0.37455886 | 0.54052972 | 0.88848777 |
| 0.22815491 | 0.22918791 | 3.86256555 | 0.37434543 | 0.54064511 | 0.88848777 |
| 0.25417782 | 0.24759072 | 3.15799401 | 0.37200583 | 0.54191294 | 0.89006587 |
| 0.53779603 | 0.32457414 | 3.35674612 | 0.37080395 | 0.54256637 | 0.89063363 |
| 0.19012653 | 0.2045292 | 5.61593512 | 0.36879412 | 0.54366231 | 0.89109007 |
| -0.2180866 | -0.2160407 | 4.21420054 | 0.36786532 | 0.54417016 | 0.89109007 |
| 0.24457362 | 0.24032638 | 4.3245904 | 0.36761935 | 0.54430479 | 0.89109007 |
| 0.19614064 | 0.20550144 | 4.81338739 | 0.36753651 | 0.54435015 | 0.89109007 |
| 0.22049178 | 0.23812121 | 3.45102209 | 0.36747474 | 0.54438398 | 0.89109007 |
| -0.332927 | -0.3530715 | 2.7778616 | 0.36508874 | 0.54569357 | 0.89266851 |
| 0.20642927 | 0.21607885 | 3.52135991 | 0.36459519 | 0.54596519 | 0.89266851 |
| -0.5896062 | -0.348239 | 4.17711909 | 0.36344262 | 0.54660048 | 0.89320259 |
| -0.1701277 | -0.2 | 4.02775794 | 0.36231168 | 0.54722518 | 0.89352728 |
| 0.18900861 | 0.21012204 | 4.20310265 | 0.36022025 | 0.54838394 | 0.89352728 |
| -0.2763049 | -0.2656062 | 3.24101212 | 0.35993223 | 0.54854387 | 0.89352728 |
| 0.21975967 | 0.24618841 | 2.92069963 | 0.35906488 | 0.54902604 | 0.89352728 |
| 0.21417989 | 0.22201345 | 4.51257824 | 0.35905329 | 0.54903249 | 0.89352728 |
| 0.19588452 | 0.20741719 | 4.10282949 | 0.35825203 | 0.54947862 | 0.89352728 |
| -0.2009039 | -0.2063629 | 4.40992635 | 0.35816601 | 0.54952655 | 0.89352728 |
| 0.22919748 | 0.22491674 | 4.46489666 | 0.35738109 | 0.54996432 | 0.89352728 |
| -0.2306182 | -0.2316256 | 3.22446461 | 0.35723238 | 0.55004733 | 0.89352728 |
| 0.32615528 | 0.29818454 | 3.1156552 | 0.35705832 | 0.55014452 | 0.89352728 |
| -0.2009983 | -0.2205146 | 3.46016691 | 0.35654168 | 0.5504332 | 0.89352728 |
| -0.2036055 | -0.1987069 | 5.30633489 | 0.35641469 | 0.5505042 | 0.89352728 |
| 0.21950353 | 0.23653832 | 3.14198069 | 0.35422393 | 0.55173175 | 0.89465619 |
| 0.26732088 | 0.23325272 | 2.92250877 | 0.35407035 | 0.551818 | 0.89465619 |
| -0.2111127 | -0.2132232 | 5.35223043 | 0.35251492 | 0.55269294 | 0.89541086 |
| -0.3313847 | -0.2641506 | 4.08632881 | 0.35214344 | 0.55290228 | 0.89541086 |
| -0.2278487 | -0.223852 | 8.35269513 | 0.35069562 | 0.55371962 | 0.8958685 |
| 0.23205467 | 0.22725093 | 3.07736033 | 0.3503048 | 0.55394064 | 0.8958685 |
| 0.21470032 | 0.21423467 | 3.76779921 | 0.34949463 | 0.55439935 | 0.8958685 |
| -0.1900112 | -0.1991787 | 3.83537824 | 0.34920056 | 0.55456602 | 0.8958685 |
| -0.2248849 | -0.2215233 | 8.40785383 | 0.3489067 | 0.55473267 | 0.8958685 |

| | | | | | |
|---|---|---|---|---|---|
| -0.1895628 | -0.2225399 | 3.10944794 | 0.34755161 | 0.55550239 | 0.89661122 |
| -0.2497051 | -0.236919 | 3.3492191 | 0.34387459 | 0.55760122 | 0.89902312 |
| -0.1966277 | -0.2016387 | 4.71313829 | 0.34348188 | 0.55782627 | 0.89902312 |
| 0.18954094 | 0.20574284 | 5.45049635 | 0.34290654 | 0.5581563 | 0.89902312 |
| 0.21805226 | 0.2147599 | 3.88249891 | 0.34107751 | 0.55920792 | 0.89902312 |
| -0.3666887 | -0.2730755 | 3.11008335 | 0.34041281 | 0.55959104 | 0.89902312 |
| 0.22819849 | 0.30416846 | 2.49584041 | 0.34007701 | 0.55978477 | 0.89902312 |
| 0.20985192 | 0.20848998 | 6.3000166 | 0.33980921 | 0.55993937 | 0.89902312 |
| -0.196204 | -0.1946896 | 4.8834669 | 0.33917714 | 0.56030458 | 0.89902312 |
| -0.190499 | -0.2036785 | 4.63805481 | 0.33865166 | 0.56060855 | 0.89902312 |
| -0.4770931 | -0.2983747 | 3.97754355 | 0.33745003 | 0.56130482 | 0.89902312 |
| -0.2192543 | -0.2195396 | 8.55971384 | 0.33729887 | 0.56139253 | 0.89902312 |
| -0.1907838 | -0.2101592 | 3.66145554 | 0.33683375 | 0.56166257 | 0.89902312 |
| -0.2318701 | -0.2323752 | 3.35601409 | 0.33630549 | 0.56196957 | 0.89902312 |
| -0.2184657 | -0.2184412 | 3.87656055 | 0.33620793 | 0.5620263 | 0.89902312 |
| -0.2042583 | -0.211989 | 3.46653922 | 0.33607269 | 0.56210496 | 0.89902312 |
| -0.2207701 | -0.2173094 | 4.64839538 | 0.33580416 | 0.56226121 | 0.89902312 |
| -0.421745 | -0.3970786 | 8.25075876 | 0.33468295 | 0.56291452 | 0.89902312 |
| 0.43468659 | 0.31601292 | 3.79356497 | 0.33464894 | 0.56293436 | 0.89902312 |
| -0.1931031 | -0.1966205 | 4.48162324 | 0.33417338 | 0.56321192 | 0.89902312 |
| 0.30441777 | 0.28713125 | 6.91648912 | 0.3338501 | 0.56340076 | 0.89902312 |
| 0.21986967 | 0.23243837 | 3.51971803 | 0.33322507 | 0.56376619 | 0.89902312 |
| -0.1935113 | -0.2026545 | 4.91477381 | 0.33270839 | 0.56406863 | 0.89902312 |
| -0.2144533 | -0.2130657 | 3.31161729 | 0.33196463 | 0.56450452 | 0.89902312 |
| 0.20366192 | 0.21565318 | 4.66653272 | 0.3316352 | 0.56469781 | 0.89902312 |
| -0.2411814 | -0.2323705 | 3.54334143 | 0.33140716 | 0.56483167 | 0.89902312 |
| 0.23039189 | 0.22673682 | 4.50699886 | 0.33070003 | 0.56524718 | 0.89902312 |
| -0.2214099 | -0.2470221 | 2.7790846 | 0.33046693 | 0.56538427 | 0.89902312 |
| -0.1738475 | -0.193984 | 4.89785305 | 0.32962198 | 0.56588176 | 0.89932005 |
| -0.304039 | -0.2705145 | 3.84107861 | 0.32898553 | 0.56625705 | 0.89942256 |
| -0.219013 | -0.2179696 | 3.73157855 | 0.32764136 | 0.56705125 | 0.89998347 |
| -0.2994669 | -0.2763375 | 2.81723616 | 0.3273357 | 0.56723215 | 0.89998347 |
| 0.18388585 | 0.20039164 | 3.35859523 | 0.32549039 | 0.56832665 | 0.90118299 |
| 0.21009991 | 0.20743648 | 6.89421958 | 0.32501217 | 0.56861097 | 0.90118299 |
| -0.1817979 | -0.1902784 | 6.11860118 | 0.32273138 | 0.56997079 | 0.90152431 |
| -0.1959904 | -0.2019114 | 4.38694382 | 0.32251467 | 0.57010032 | 0.90152431 |
| 0.21224526 | 0.21325751 | 6.80857467 | 0.32243922 | 0.57014544 | 0.90152431 |
| -0.2307995 | -0.2571773 | 3.28128439 | 0.32235393 | 0.57019644 | 0.90152431 |
| -0.1760441 | -0.1918042 | 4.28059527 | 0.32204059 | 0.5703839 | 0.90152431 |
| 0.20923921 | 0.2012387 | 3.9055215 | 0.32106009 | 0.57097127 | 0.90196007 |
| 0.32456691 | 0.30672443 | 2.92107243 | 0.31991307 | 0.57165989 | 0.90255523 |
| -0.2236144 | -0.2300295 | 2.88507569 | 0.31904587 | 0.57218161 | 0.90262351 |
| -0.2146315 | -0.1919229 | 3.66389966 | 0.31815559 | 0.57271818 | 0.90262351 |
| 0.24244925 | 0.25074699 | 2.98815433 | 0.31796007 | 0.57283615 | 0.90262351 |
| 0.1965429 | 0.18660168 | 5.90502288 | 0.31765994 | 0.57301734 | 0.90262351 |
| -0.1956868 | -0.2333584 | 2.85561334 | 0.31725395 | 0.57326262 | 0.90262351 |

| | | | | | |
|---|---|---|---|---|---|
| 0.32004525 | 0.27127773 | 3.112971 | 0.31661836 | 0.57364701 | 0.9027376 |
| 0.46136493 | 0.30008599 | 3.80519912 | 0.31584421 | 0.57411589 | 0.90298445 |
| 0.20293071 | 0.19364934 | 5.16824434 | 0.31451473 | 0.57492289 | 0.90332314 |
| 0.22141315 | 0.20010175 | 3.83690674 | 0.31440184 | 0.57499152 | 0.90332314 |
| -0.2124851 | -0.2072788 | 3.87314344 | 0.31323355 | 0.5757027 | 0.90332314 |
| -0.2008416 | -0.201318 | 3.69618226 | 0.31306822 | 0.57580348 | 0.90332314 |
| 0.25431644 | 0.25752589 | 2.79066865 | 0.31262884 | 0.57607149 | 0.90332314 |
| 0.22657393 | 0.22652408 | 3.37929015 | 0.31241166 | 0.57620405 | 0.90332314 |
| 0.23097927 | 0.21046327 | 3.46261934 | 0.30992398 | 0.57772684 | 0.90494889 |
| 0.18721598 | 0.18335421 | 3.96530614 | 0.30940995 | 0.57804249 | 0.90494889 |
| -0.1901765 | -0.2245 | 2.91263125 | 0.30902845 | 0.57827698 | 0.90494889 |
| -0.1695022 | -0.1825865 | 4.08943279 | 0.30830118 | 0.57872453 | 0.90494889 |
| 0.19819108 | 0.21246382 | 4.6478416 | 0.30671463 | 0.57970325 | 0.90494889 |
| 0.21519344 | 0.22162754 | 3.63260666 | 0.30587953 | 0.58021974 | 0.90494889 |
| -0.2616992 | -0.2298435 | 3.71559911 | 0.30545803 | 0.58048079 | 0.90494889 |
| -0.1941854 | -0.18585 | 5.88754315 | 0.30517795 | 0.58065437 | 0.90494889 |
| -0.269772 | -0.2543413 | 3.04846526 | 0.30459598 | 0.5810154 | 0.90494889 |
| 0.28559801 | 0.2609119 | 3.12670333 | 0.30383295 | 0.58148943 | 0.90494889 |
| 0.320248 | 0.28821424 | 2.76666499 | 0.30370402 | 0.5815696 | 0.90494889 |
| -0.2618399 | -0.2527011 | 2.98967422 | 0.30350231 | 0.58169507 | 0.90494889 |
| -0.2267797 | -0.2348881 | 3.3606484 | 0.30277438 | 0.58214834 | 0.90494889 |
| 0.21793287 | 0.19214779 | 3.92148037 | 0.30275433 | 0.58216083 | 0.90494889 |
| 0.21599412 | 0.2459652 | 2.70913775 | 0.30176167 | 0.58278011 | 0.90494889 |
| 0.23154819 | 0.21953172 | 3.85680189 | 0.30046606 | 0.58359039 | 0.90494889 |
| -0.1685938 | -0.1832664 | 6.00285265 | 0.29963638 | 0.58411047 | 0.90494889 |
| -0.1945402 | -0.2498542 | 3.37153492 | 0.29881199 | 0.58462816 | 0.90494889 |
| -0.2393149 | -0.2147663 | 4.13449322 | 0.29875055 | 0.58466678 | 0.90494889 |
| -0.1776159 | -0.2131558 | 3.120332 | 0.2986047 | 0.58475847 | 0.90494889 |
| -0.2392478 | -0.2287163 | 2.8836646 | 0.29801306 | 0.58513074 | 0.90494889 |
| -0.1911933 | -0.1950704 | 3.88685435 | 0.29795136 | 0.58516959 | 0.90494889 |
| 0.22536043 | 0.2029564 | 3.4328722 | 0.29748022 | 0.58546642 | 0.90494889 |
| -0.2911647 | -0.2228664 | 3.87606391 | 0.29729318 | 0.58558435 | 0.90494889 |
| -0.1969778 | -0.2252119 | 2.94546503 | 0.2972918 | 0.58558522 | 0.90494889 |
| -0.2063546 | -0.1974992 | 4.81037922 | 0.2966225 | 0.5860076 | 0.90494889 |
| 0.1817853 | 0.19900871 | 5.14266072 | 0.29629444 | 0.58621486 | 0.90494889 |
| -0.2441701 | -0.2271254 | 3.20506886 | 0.29624898 | 0.58624358 | 0.90494889 |
| -0.2089981 | -0.2264737 | 2.89479265 | 0.29527051 | 0.58686269 | 0.90494889 |
| -0.1936735 | -0.1900378 | 4.4544598 | 0.29485296 | 0.58712729 | 0.90494889 |
| 0.169295 | 0.24547655 | 3.45968775 | 0.29484859 | 0.58713006 | 0.90494889 |
| 0.46280484 | 0.26547795 | 3.94424737 | 0.29454276 | 0.58732402 | 0.90494889 |
| 0.19767851 | 0.19863656 | 4.09095843 | 0.29417077 | 0.58756011 | 0.90494889 |
| 0.22547286 | 0.24163622 | 2.59849113 | 0.29311091 | 0.58823385 | 0.90550466 |
| -0.2148811 | -0.1984797 | 4.25099683 | 0.29186316 | 0.58902904 | 0.90618915 |
| 0.19135072 | 0.19026456 | 3.29537186 | 0.29143132 | 0.58930476 | 0.90618915 |
| -0.2091408 | -0.1925956 | 4.97132971 | 0.28847773 | 0.59119769 | 0.90838183 |
| -0.1784348 | -0.1824971 | 4.96355802 | 0.28790905 | 0.59156359 | 0.90838183 |

| | | | | | |
|---|---|---|---|---|---|
| -0.1865724 | -0.1777622 | 6.02506714 | 0.28718017 | 0.59203324 | 0.90838183 |
| -0.181026 | -0.1896129 | 5.01225692 | 0.2871685 | 0.59204076 | 0.90838183 |
| 0.37602797 | 0.27012585 | 3.26990055 | 0.28674396 | 0.59231467 | 0.90838183 |
| -0.1992618 | -0.1817949 | 5.38777467 | 0.2860335 | 0.59277364 | 0.90838183 |
| 0.19660968 | 0.19995279 | 4.55122079 | 0.28579501 | 0.59292788 | 0.90838183 |
| 0.20456561 | 0.20272165 | 3.28961242 | 0.28507432 | 0.59339445 | 0.90861563 |
| -0.183705 | -0.1885601 | 7.68398505 | 0.28455199 | 0.59373309 | 0.90865338 |
| 0.20501044 | 0.19227565 | 4.75047397 | 0.28348561 | 0.59442567 | 0.90884711 |
| -0.1922559 | -0.1937421 | 4.12448658 | 0.28321217 | 0.59460353 | 0.90884711 |
| 0.2106625 | 0.20941117 | 9.07446179 | 0.2826167 | 0.59499125 | 0.90884711 |
| -0.1629094 | -0.169869 | 5.82751779 | 0.28242549 | 0.59511585 | 0.90884711 |
| 0.19056277 | 0.18251042 | 5.70151006 | 0.28194249 | 0.59543086 | 0.90884858 |
| 0.16345011 | 0.18309965 | 4.16782933 | 0.28083946 | 0.59615153 | 0.90946891 |
| -0.2246271 | -0.1929911 | 3.98311724 | 0.27894877 | 0.59739106 | 0.91043435 |
| 0.24576517 | 0.21571417 | 3.09141777 | 0.27882767 | 0.59747063 | 0.91043435 |
| -0.2239879 | -0.2069332 | 3.60851611 | 0.278436 | 0.59772815 | 0.91043435 |
| -0.1904337 | -0.1934299 | 8.17277471 | 0.27473742 | 0.60017143 | 0.9131809 |
| 0.17234006 | 0.1698646 | 6.34404329 | 0.2745194 | 0.6003161 | 0.9131809 |
| -0.1678596 | -0.1771764 | 5.45256379 | 0.27427561 | 0.60047797 | 0.9131809 |
| -0.1696828 | -0.1798393 | 5.45803078 | 0.273032 | 0.60130509 | 0.91395848 |
| -0.1825471 | -0.1881162 | 4.90975797 | 0.27238477 | 0.60173651 | 0.9141341 |
| 0.16598913 | 0.16799279 | 5.95358886 | 0.27132956 | 0.60244128 | 0.91472459 |
| 0.1771385 | 0.17932089 | 6.49006674 | 0.27031008 | 0.60312384 | 0.91528076 |
| -0.1837435 | -0.1855781 | 4.33742984 | 0.26943518 | 0.60371091 | 0.9156915 |
| -0.1950867 | -0.1866791 | 3.86923045 | 0.26810399 | 0.60460648 | 0.91656949 |
| 0.21956728 | 0.21700582 | 3.32471416 | 0.26664336 | 0.60559237 | 0.91743254 |
| -0.164813 | -0.1812454 | 3.57914154 | 0.26632191 | 0.60580981 | 0.91743254 |
| -0.2021375 | -0.2005788 | 3.55334067 | 0.26566935 | 0.60625172 | 0.91762158 |
| -0.1912058 | -0.2007358 | 3.31073594 | 0.26489306 | 0.60677831 | 0.91793854 |
| 0.28415805 | 0.26048568 | 2.73852827 | 0.26358923 | 0.60766496 | 0.91879958 |
| 0.20524147 | 0.21439041 | 3.893781 | 0.26145605 | 0.60912159 | 0.92048777 |
| -0.1993441 | -0.1798751 | 3.46907789 | 0.26072442 | 0.6096229 | 0.92048777 |
| 0.23037614 | 0.22188728 | 4.85647284 | 0.26040473 | 0.60984224 | 0.92048777 |
| -0.372533 | -0.2648633 | 2.65057273 | 0.26001881 | 0.61010723 | 0.92048777 |
| -0.1759029 | -0.1692986 | 5.50586034 | 0.25946767 | 0.61048611 | 0.92048777 |
| -0.23659 | -0.2162054 | 3.89354827 | 0.25790792 | 0.6115611 | 0.92048777 |
| -0.1630554 | -0.1664262 | 5.94036009 | 0.2570105 | 0.61218146 | 0.92048777 |
| 0.18559522 | 0.17929507 | 5.45931367 | 0.25684774 | 0.61229412 | 0.92048777 |
| -0.1676657 | -0.1869932 | 3.40530331 | 0.25633574 | 0.6126488 | 0.92048777 |
| -0.1764411 | -0.1800764 | 5.57588075 | 0.25621971 | 0.61272924 | 0.92048777 |
| -0.1799499 | -0.1771826 | 3.54751745 | 0.25591682 | 0.61293934 | 0.92048777 |
| -0.2041583 | -0.1977794 | 3.25168099 | 0.25564917 | 0.61312511 | 0.92048777 |
| 0.24028575 | 0.22093788 | 2.87317392 | 0.25560357 | 0.61315678 | 0.92048777 |
| 0.23269959 | 0.22014118 | 2.82848523 | 0.25526617 | 0.61339117 | 0.92048777 |
| -0.3934969 | -0.2708512 | 2.74560046 | 0.25476093 | 0.61374253 | 0.92048777 |
| 0.18576273 | 0.1874973 | 3.98784035 | 0.25457699 | 0.61387056 | 0.92048777 |

| | | | | | |
|---|---|---|---|---|---|
| 0.23711054 | 0.23642299 | 2.83511827 | 0.25333667 | 0.61473536 | 0.92074127 |
| -0.183322 | -0.1791207 | 3.81895637 | 0.25306753 | 0.61492337 | 0.92074127 |
| 0.22652357 | 0.20687209 | 4.36696982 | 0.25296635 | 0.61499408 | 0.92074127 |
| 0.20018354 | 0.17855759 | 4.65697148 | 0.25206648 | 0.61562375 | 0.92085923 |
| -0.1632818 | -0.169938 | 5.48541897 | 0.25190531 | 0.61573668 | 0.92085923 |
| -0.1663565 | -0.1718386 | 4.86218028 | 0.25149059 | 0.61602746 | 0.92085923 |
| 0.19663741 | 0.17311766 | 3.74688532 | 0.25087361 | 0.61646061 | 0.92103098 |
| -0.244907 | -0.2117822 | 3.39287411 | 0.24944681 | 0.61746486 | 0.92184179 |
| -0.3051799 | -0.2179092 | 4.27328693 | 0.24919798 | 0.61764037 | 0.92184179 |
| -0.1514577 | -0.1701597 | 3.8133238 | 0.24840439 | 0.61820084 | 0.9222027 |
| 0.27158405 | 0.23737239 | 4.08736907 | 0.24750464 | 0.61883764 | 0.92267704 |
| -0.2068901 | -0.2116069 | 3.06170481 | 0.2461714 | 0.61978391 | 0.92295777 |
| -0.1576545 | -0.1679471 | 5.51774848 | 0.24611041 | 0.61982727 | 0.92295777 |
| -0.1625379 | -0.2457369 | 4.62772514 | 0.24573676 | 0.62009308 | 0.92295777 |
| -0.1548724 | -0.1648925 | 5.76960619 | 0.24544388 | 0.62030161 | 0.92295777 |
| -0.1538832 | -0.1657426 | 4.02828216 | 0.244592 | 0.62090902 | 0.9233868 |
| -0.1479343 | -0.1772437 | 3.41461972 | 0.2428825 | 0.62213193 | 0.92356313 |
| -0.1828977 | -0.1786121 | 4.30467487 | 0.2411112 | 0.6234047 | 0.92356313 |
| 0.18341133 | 0.18350641 | 6.2483384 | 0.24050873 | 0.62383893 | 0.92356313 |
| 0.50468824 | 0.28997623 | 2.87234656 | 0.24025423 | 0.62402256 | 0.92356313 |
| -0.2137763 | -0.1931177 | 3.70536821 | 0.23975034 | 0.62438649 | 0.92356313 |
| -0.1855925 | -0.182592 | 3.64010607 | 0.23901998 | 0.62491483 | 0.92356313 |
| -0.1657331 | -0.1698001 | 5.20813284 | 0.23856847 | 0.62524196 | 0.92356313 |
| -0.1549993 | -0.1724444 | 4.75574649 | 0.23730707 | 0.62615789 | 0.92356313 |
| -0.174481 | -0.1748494 | 6.12609671 | 0.23708635 | 0.62631848 | 0.92356313 |
| -0.1619608 | -0.1729363 | 4.05537689 | 0.23701851 | 0.62636785 | 0.92356313 |
| 1.23668276 | 0.38672337 | 2.90748586 | 0.23653763 | 0.62671808 | 0.92356313 |
| 0.21061626 | 0.187979 | 3.02375981 | 0.23635335 | 0.62685241 | 0.92356313 |
| -0.1669373 | -0.1656109 | 7.04475599 | 0.23606662 | 0.62706156 | 0.92356313 |
| -0.1955392 | -0.2007199 | 3.17786194 | 0.23506288 | 0.62779493 | 0.92356313 |
| 0.1845889 | 0.16794542 | 4.8398713 | 0.23501206 | 0.62783211 | 0.92356313 |
| -0.3512949 | -0.2843581 | 4.43647414 | 0.23488897 | 0.62792219 | 0.92356313 |
| -0.1653581 | -0.1565517 | 6.14251905 | 0.23482998 | 0.62796537 | 0.92356313 |
| 0.18121951 | 0.17638515 | 4.23818977 | 0.23381712 | 0.62870783 | 0.92356313 |
| -0.1484908 | -0.1681633 | 3.47248882 | 0.23304538 | 0.62927486 | 0.92356313 |
| -0.178864 | -0.1711725 | 4.71154709 | 0.23243984 | 0.6297206 | 0.92356313 |
| 0.14383775 | 0.16526075 | 4.96517234 | 0.23238391 | 0.6297618 | 0.92356313 |
| 0.2133517 | 0.23663648 | 2.6191037 | 0.23236201 | 0.62977794 | 0.92356313 |
| -0.1975754 | -0.172121 | 4.41073664 | 0.2318855 | 0.63012927 | 0.92356313 |
| 0.17086673 | 0.17418191 | 4.77473118 | 0.23144396 | 0.63045522 | 0.92356313 |
| 0.17386714 | 0.23350528 | 2.78277087 | 0.23132662 | 0.6305419 | 0.92356313 |
| -0.1657832 | -0.1613408 | 5.7667401 | 0.23025437 | 0.63133528 | 0.92356313 |
| -0.2057366 | -0.1873353 | 5.65214177 | 0.22956802 | 0.63184432 | 0.92356313 |
| 0.20706354 | 0.20995743 | 3.16655156 | 0.22906172 | 0.63222042 | 0.92356313 |
| 0.17303016 | 0.16069005 | 4.66408613 | 0.22816212 | 0.63288995 | 0.92356313 |
| -0.1744675 | -0.1751962 | 4.28096777 | 0.22786378 | 0.63311235 | 0.92356313 |

| | | | | | |
|---|---|---|---|---|---|
| -0.181513 | -0.1759458 | 4.13926418 | 0.22756179 | 0.63333764 | 0.92356313 |
| 0.1729565 | 0.19209219 | 2.97829957 | 0.22711153 | 0.6336739 | 0.92356313 |
| -0.1614997 | -0.1700411 | 3.58816421 | 0.22682738 | 0.63388632 | 0.92356313 |
| -0.1738878 | -0.1866057 | 3.33063552 | 0.22630518 | 0.63427712 | 0.92356313 |
| 0.21234747 | 0.19146321 | 3.19243902 | 0.22609368 | 0.63443556 | 0.92356313 |
| 0.17109822 | 0.16332836 | 5.23164387 | 0.22596246 | 0.6345339 | 0.92356313 |
| -0.2064588 | -0.2623766 | 3.2564984 | 0.22579298 | 0.63466097 | 0.92356313 |
| -0.2124726 | -0.1829734 | 3.81924219 | 0.22558974 | 0.63481343 | 0.92356313 |
| -0.1561046 | -0.1561588 | 4.0392677 | 0.22517627 | 0.63512385 | 0.92356313 |
| -0.1709886 | -0.1734683 | 4.40683841 | 0.22422596 | 0.63583864 | 0.92356313 |
| 0.16715559 | 0.17113286 | 5.0987842 | 0.22371916 | 0.6362206 | 0.92356313 |
| 0.18476175 | 0.16088582 | 4.22783164 | 0.22320793 | 0.63660643 | 0.92356313 |
| -0.1834718 | -0.1830855 | 3.539192 | 0.22228812 | 0.63730199 | 0.92356313 |
| -0.1782213 | -0.1787436 | 5.29169133 | 0.22161828 | 0.63780963 | 0.92356313 |
| 0.19248309 | 0.18484737 | 3.33107601 | 0.22134764 | 0.638015 | 0.92356313 |
| 0.16156902 | 0.16243044 | 6.27921059 | 0.22105896 | 0.63823422 | 0.92356313 |
| -0.2015928 | -0.2090947 | 2.57839171 | 0.22070791 | 0.63850105 | 0.92356313 |
| -0.1783877 | -0.176737 | 4.15219322 | 0.21900339 | 0.63980034 | 0.92356313 |
| -0.2064641 | -0.1906601 | 3.37340789 | 0.2189859 | 0.6398137 | 0.92356313 |
| 0.18357369 | 0.19476437 | 3.59985013 | 0.21895634 | 0.63983629 | 0.92356313 |
| 0.17543295 | 0.17394229 | 4.46559223 | 0.2185901 | 0.6401163 | 0.92356313 |
| -0.1777062 | -0.1646077 | 4.03536853 | 0.21829187 | 0.64034452 | 0.92356313 |
| 0.2450714 | 0.19917987 | 2.70805367 | 0.2182368 | 0.64038669 | 0.92356313 |
| -0.219161 | -0.2204728 | 2.72404953 | 0.21791322 | 0.64063456 | 0.92356313 |
| -0.1522571 | -0.1600902 | 6.89749455 | 0.21698821 | 0.64134439 | 0.92356313 |
| -0.2188896 | -0.1956534 | 3.21482479 | 0.21698377 | 0.64134781 | 0.92356313 |
| -0.1528077 | -0.1634352 | 4.22031573 | 0.21694495 | 0.64137764 | 0.92356313 |
| 0.14425604 | 0.16975443 | 4.5333391 | 0.21642994 | 0.64177369 | 0.92356313 |
| -0.145715 | -0.1621102 | 3.73667745 | 0.21631008 | 0.64186595 | 0.92356313 |
| -0.1300529 | -0.1623639 | 3.50108805 | 0.21578209 | 0.64227272 | 0.92356313 |
| 0.20100867 | 0.21527513 | 2.7686169 | 0.21517701 | 0.64273962 | 0.92356313 |
| -0.2796745 | -0.2294331 | 2.74573173 | 0.21512219 | 0.64278197 | 0.92356313 |
| 0.20161648 | 0.20559349 | 2.59006347 | 0.21494304 | 0.64292038 | 0.92356313 |
| 0.17501264 | 0.18071788 | 5.25795161 | 0.21427471 | 0.64343737 | 0.92356313 |
| 0.43451605 | 0.2885099 | 3.16689482 | 0.2133102 | 0.64418518 | 0.92356313 |
| -0.1645879 | -0.1701366 | 4.4097201 | 0.21305338 | 0.64438465 | 0.92356313 |
| 0.16359181 | 0.15004597 | 6.10539597 | 0.2128164 | 0.64456884 | 0.92356313 |
| -0.1497131 | -0.1603159 | 4.76503321 | 0.21268135 | 0.64467386 | 0.92356313 |
| 0.15570022 | 0.15246774 | 4.29464798 | 0.21188014 | 0.64529774 | 0.92356313 |
| -0.1765238 | -0.1701407 | 4.11083158 | 0.21170143 | 0.6454371 | 0.92356313 |
| -0.1605513 | -0.1737851 | 3.43007001 | 0.21150978 | 0.64558662 | 0.92356313 |
| -0.1453675 | -0.1562124 | 5.19613805 | 0.21142156 | 0.64565548 | 0.92356313 |
| 0.14754652 | 0.16399458 | 3.8827227 | 0.21112045 | 0.64589062 | 0.92356313 |
| -0.1412011 | -0.1575457 | 4.31033055 | 0.2108347 | 0.64611396 | 0.92356313 |
| -0.2331141 | -0.2142763 | 2.89106926 | 0.2107603 | 0.64617214 | 0.92356313 |
| -0.1910269 | -0.1894748 | 9.69516109 | 0.21054043 | 0.64634415 | 0.92356313 |

| | | | | | |
|---|---|---|---|---|---|
| -0.1591919 | -0.1767213 | 3.23711548 | 0.21025512 | 0.64656752 | 0.92356313 |
| -0.1785675 | -0.1795574 | 8.32544293 | 0.20963337 | 0.64705492 | 0.92356313 |
| 0.14891583 | 0.15625831 | 3.97056613 | 0.20829428 | 0.64810763 | 0.92356313 |
| -0.1557632 | -0.1514727 | 5.26421901 | 0.20807132 | 0.6482833 | 0.92356313 |
| -0.1599274 | -0.1682118 | 3.67029442 | 0.20788115 | 0.64843323 | 0.92356313 |
| -0.1522008 | -0.1537439 | 6.44838134 | 0.20770893 | 0.64856908 | 0.92356313 |
| 0.14400957 | 0.16865536 | 3.8553307 | 0.20769253 | 0.64858202 | 0.92356313 |
| -0.1409896 | -0.1561017 | 4.22453664 | 0.20752458 | 0.64871457 | 0.92356313 |
| -0.2209298 | -0.2188552 | 2.64488737 | 0.20689812 | 0.64920957 | 0.92356313 |
| -0.2419315 | -0.2017381 | 3.38514675 | 0.20669605 | 0.64936942 | 0.92356313 |
| 0.30938964 | 0.22659771 | 4.48604717 | 0.20645803 | 0.64955784 | 0.92356313 |
| -0.1836008 | -0.170578 | 4.13390143 | 0.20641006 | 0.64959583 | 0.92356313 |
| -0.1821488 | -0.1808507 | 3.60188093 | 0.20632531 | 0.64966296 | 0.92356313 |
| -0.407727 | -0.4014816 | 10.4832294 | 0.20600429 | 0.64991739 | 0.92356313 |
| -0.407537 | -0.4012915 | 10.4832916 | 0.20581044 | 0.65007115 | 0.92356313 |
| -0.1641469 | -0.1542853 | 5.10498846 | 0.20541183 | 0.65038758 | 0.92356313 |
| 0.13650236 | 0.15422535 | 4.26451529 | 0.20486671 | 0.65082093 | 0.92372524 |
| -0.1679264 | -0.1921907 | 2.83725454 | 0.20433254 | 0.65124624 | 0.92387579 |
| -0.1545683 | -0.1556474 | 4.13633779 | 0.20352356 | 0.65189164 | 0.92433827 |
| 0.61804928 | 0.26078959 | 3.75105415 | 0.20265893 | 0.65258314 | 0.92486562 |
| 0.1367003 | 0.23452631 | 2.7910375 | 0.20168612 | 0.65336329 | 0.92510436 |
| 0.19927612 | 0.19557543 | 2.94097114 | 0.19939497 | 0.65520966 | 0.92510436 |
| 0.17271365 | 0.15996066 | 4.5669438 | 0.19938176 | 0.65522033 | 0.92510436 |
| 0.17017789 | 0.16304917 | 3.74188724 | 0.19782746 | 0.6564802 | 0.92510436 |
| 0.14883472 | 0.15253742 | 3.80693155 | 0.1974997 | 0.65674663 | 0.92510436 |
| 0.16119426 | 0.14692629 | 5.45588752 | 0.19726375 | 0.6569386 | 0.92510436 |
| 0.19593312 | 0.1956962 | 3.21696127 | 0.19719505 | 0.65699451 | 0.92510436 |
| -0.1481761 | -0.1433136 | 6.118987 | 0.19704555 | 0.65711623 | 0.92510436 |
| 0.1786177 | 0.17854178 | 3.36115344 | 0.19667107 | 0.65742139 | 0.92510436 |
| -0.174657 | -0.1569078 | 3.8184276 | 0.19530354 | 0.6585387 | 0.92510436 |
| 0.17730349 | 0.18776733 | 3.20522781 | 0.19506509 | 0.658734 | 0.92510436 |
| 0.13511822 | 0.15392504 | 4.15774997 | 0.19460217 | 0.65911356 | 0.92510436 |
| 0.20506967 | 0.1779817 | 3.08199591 | 0.19414412 | 0.65948966 | 0.92510436 |
| 0.17342691 | 0.17051581 | 4.11508938 | 0.19390951 | 0.65968249 | 0.92510436 |
| 0.18313089 | 0.15931853 | 3.62491921 | 0.1937969 | 0.65977511 | 0.92510436 |
| -0.1557791 | -0.1481451 | 5.17653163 | 0.19309093 | 0.66035642 | 0.92510436 |
| -0.1888074 | -0.180285 | 3.34236974 | 0.19260551 | 0.66075686 | 0.92510436 |
| -0.1538613 | -0.1567786 | 4.75391174 | 0.19247874 | 0.66086154 | 0.92510436 |
| 0.18485322 | 0.18515479 | 8.47961814 | 0.1919893 | 0.66126607 | 0.92510436 |
| 0.32823021 | 0.22658973 | 2.96698594 | 0.19116119 | 0.66195192 | 0.92510436 |
| 0.20016004 | 0.18219867 | 2.82725956 | 0.19095198 | 0.66212547 | 0.92510436 |
| -0.1872233 | -0.1811592 | 3.09931361 | 0.19089737 | 0.66217079 | 0.92510436 |
| -0.1380322 | -0.1533279 | 4.72996238 | 0.19059218 | 0.66242421 | 0.92510436 |
| 0.1953633 | 0.19569267 | 10.4851649 | 0.1902354 | 0.66272076 | 0.92510436 |
| -0.1321685 | -0.1486152 | 4.09108496 | 0.19010656 | 0.66282794 | 0.92510436 |
| -0.1585879 | -0.1583919 | 3.65926488 | 0.19002897 | 0.66289251 | 0.92510436 |

| | | | | | |
|---|---|---|---|---|---|
| -0.1844342 | -0.1731386 | 3.38599381 | 0.18999081 | 0.66292427 | 0.92510436 |
| 0.2037413 | 0.18467824 | 2.93975716 | 0.18962816 | 0.66322628 | 0.92510436 |
| 0.13010245 | 0.14961193 | 5.39673712 | 0.18835138 | 0.6642923 | 0.92510436 |
| -0.20506 | -0.1945532 | 3.19805943 | 0.18812979 | 0.66447776 | 0.92510436 |
| -0.1524167 | -0.157864 | 5.09004094 | 0.18809344 | 0.66450819 | 0.92510436 |
| -0.1598318 | -0.16533 | 4.0617812 | 0.18790069 | 0.66466963 | 0.92510436 |
| -0.1633954 | -0.1600738 | 4.66483089 | 0.18772621 | 0.66481585 | 0.92510436 |
| 0.14412347 | 0.15201041 | 4.86063741 | 0.18768887 | 0.66484715 | 0.92510436 |
| -0.1362268 | -0.1467839 | 5.89302802 | 0.18738439 | 0.66510254 | 0.92510436 |
| 0.19115763 | 0.21224008 | 6.04502942 | 0.18728913 | 0.66518249 | 0.92510436 |
| -0.1957639 | -0.1951083 | 2.72224255 | 0.18710441 | 0.6653376 | 0.92510436 |
| 0.19148865 | 0.20485319 | 2.98414574 | 0.18648598 | 0.66585754 | 0.92510436 |
| 0.15760641 | 0.16658956 | 4.53946518 | 0.18641461 | 0.66591761 | 0.92510436 |
| -0.1364825 | -0.1457966 | 5.8603202 | 0.18594299 | 0.6663149 | 0.92510436 |
| -0.1282116 | -0.1467324 | 5.0807387 | 0.18561821 | 0.66658885 | 0.92510436 |
| -0.1413598 | -0.1550624 | 3.70193058 | 0.18542206 | 0.66675443 | 0.92510436 |
| 0.20141355 | 0.20027774 | 2.70468891 | 0.18532489 | 0.66683649 | 0.92510436 |
| -0.1852678 | -0.1927691 | 2.82091539 | 0.18530302 | 0.66685497 | 0.92510436 |
| 0.14857536 | 0.14436706 | 6.01101216 | 0.18476072 | 0.66731347 | 0.92510436 |
| 0.16370185 | 0.20668049 | 3.29498823 | 0.18459224 | 0.66745609 | 0.92510436 |
| -0.2042188 | -0.1881259 | 3.39280113 | 0.18404502 | 0.66791981 | 0.92530394 |
| 0.16095599 | 0.15631299 | 3.69485745 | 0.18294766 | 0.66885221 | 0.92615229 |
| 0.14543847 | 0.14299759 | 6.01709026 | 0.18122416 | 0.67032332 | 0.92681117 |
| 0.16881606 | 0.16864632 | 4.66994862 | 0.18090547 | 0.67059625 | 0.92681117 |
| 0.13526152 | 0.15632407 | 4.09755967 | 0.18024933 | 0.67115906 | 0.92681117 |
| 0.15979614 | 0.15574052 | 4.67280217 | 0.18016555 | 0.67123101 | 0.92681117 |
| 0.17372861 | 0.17668018 | 3.38798418 | 0.18013645 | 0.67125601 | 0.92681117 |
| -0.1484581 | -0.1452709 | 4.80526559 | 0.17998979 | 0.67138202 | 0.92681117 |
| -0.16036 | -0.1640177 | 7.92693602 | 0.17944275 | 0.67185257 | 0.92681117 |
| -0.1334599 | -0.1907991 | 3.45470415 | 0.17939921 | 0.67189006 | 0.92681117 |
| -0.1107582 | -0.2351876 | 4.03280181 | 0.17889796 | 0.67232204 | 0.92696521 |
| 0.15442987 | 0.15917536 | 3.30067637 | 0.17801175 | 0.67308751 | 0.92753979 |
| -0.0783257 | 0.25548301 | 2.98113203 | 0.1773506 | 0.67366005 | 0.92753979 |
| 0.17872383 | 0.17168603 | 3.2160548 | 0.17615739 | 0.67469653 | 0.92753979 |
| -0.1503057 | -0.1798756 | 2.6882225 | 0.1757771 | 0.67502773 | 0.92753979 |
| 0.18735381 | 0.17084805 | 3.85556931 | 0.17561188 | 0.67517176 | 0.92753979 |
| -0.1668254 | -0.1843213 | 2.76263159 | 0.17552112 | 0.67525091 | 0.92753979 |
| 0.13997558 | 0.14393186 | 4.56044156 | 0.17550142 | 0.6752681 | 0.92753979 |
| 0.14896223 | 0.15322628 | 3.68539659 | 0.17511225 | 0.67560779 | 0.92753979 |
| -0.1493838 | -0.1673709 | 2.99575976 | 0.17470712 | 0.67596188 | 0.92753979 |
| -0.2440137 | -0.1985864 | 2.67413053 | 0.17459053 | 0.67606387 | 0.92753979 |
| 0.20849708 | 0.16510816 | 3.03166819 | 0.17436151 | 0.67626433 | 0.92753979 |
| 0.20447544 | 0.21443227 | 2.41258961 | 0.17378352 | 0.67677094 | 0.92779492 |
| -0.1262556 | -0.1391863 | 5.52948436 | 0.17323119 | 0.67725597 | 0.92802025 |
| -0.1618161 | -0.161479 | 9.37278489 | 0.17282134 | 0.67761647 | 0.92807481 |
| 0.14322053 | 0.14007787 | 4.83174329 | 0.1721716 | 0.67818901 | 0.92819678 |

| | | | | | |
|---|---|---|---|---|---|
| -0.153524 | -0.153659 | 5.23297338 | 0.17199256 | 0.67834699 | 0.92819678 |
| 0.13630896 | 0.15392954 | 3.24528093 | 0.17111623 | 0.67912168 | 0.92838151 |
| 0.15512148 | 0.15134513 | 4.13932375 | 0.17111408 | 0.67912359 | 0.92838151 |
| -0.1406047 | -0.144055 | 5.86697901 | 0.16910336 | 0.68090991 | 0.92908156 |
| -0.1496049 | -0.1482364 | 6.08743298 | 0.16897583 | 0.68102363 | 0.92908156 |
| 0.1384849 | 0.14139795 | 5.61421477 | 0.16886229 | 0.68112491 | 0.92908156 |
| 0.14904053 | 0.1563888 | 3.9810886 | 0.16877949 | 0.6811988 | 0.92908156 |
| -0.1907113 | -0.1688038 | 4.7767292 | 0.16873235 | 0.68124087 | 0.92908156 |
| -0.1721763 | -0.1651662 | 3.07068466 | 0.16618461 | 0.68352517 | 0.93175781 |
| 0.20496459 | 0.18801837 | 2.57771442 | 0.16488087 | 0.684702 | 0.93251124 |
| -0.1779555 | -0.1636381 | 4.12303942 | 0.16485842 | 0.68472231 | 0.93251124 |
| -0.1149232 | -0.1373597 | 4.96222469 | 0.164142 | 0.68537136 | 0.93285269 |
| 0.16954262 | 0.16306957 | 3.27554306 | 0.16331488 | 0.68612274 | 0.93285269 |
| -0.1755788 | -0.1585848 | 4.99980394 | 0.16307974 | 0.68633676 | 0.93285269 |
| -0.1510394 | -0.1518857 | 3.2032799 | 0.16297167 | 0.68643518 | 0.93285269 |
| 0.12743365 | 0.13518474 | 5.38224152 | 0.16276671 | 0.68662194 | 0.93285269 |
| 0.13223605 | 0.14340154 | 4.11146413 | 0.16203976 | 0.68728546 | 0.93285269 |
| -0.2064601 | -0.197424 | 3.02584389 | 0.16186732 | 0.68744311 | 0.93285269 |
| -0.1382202 | -0.1450743 | 3.92892569 | 0.16174853 | 0.68755176 | 0.93285269 |
| -0.1381956 | -0.1429223 | 4.51195228 | 0.16112253 | 0.68812513 | 0.93319312 |
| -0.1898864 | -0.1875349 | 2.72130253 | 0.16070127 | 0.68851169 | 0.93328002 |
| -0.1829869 | -0.1991339 | 2.76980098 | 0.15965089 | 0.68947813 | 0.93411099 |
| 0.13819396 | 0.14267025 | 4.16090404 | 0.15887675 | 0.69019277 | 0.93411099 |
| 0.1623887 | 0.15771908 | 4.13836167 | 0.15836409 | 0.69066714 | 0.93411099 |
| 0.1333513 | 0.13812329 | 5.22560231 | 0.15812017 | 0.69089316 | 0.93411099 |
| -0.1159466 | -0.1335904 | 4.15332704 | 0.15801794 | 0.69098795 | 0.93411099 |
| -0.1528331 | -0.1490517 | 3.35965832 | 0.15793877 | 0.69106138 | 0.93411099 |
| 0.11653182 | 0.14120047 | 4.69224436 | 0.15703002 | 0.69190576 | 0.93481571 |
| -0.1333598 | -0.1334855 | 4.32215206 | 0.15566432 | 0.69318006 | 0.93597165 |
| 0.12921077 | 0.13373006 | 4.50557531 | 0.1553758 | 0.69345009 | 0.93597165 |
| 0.14979367 | 0.1371478 | 5.36558948 | 0.15482465 | 0.69396674 | 0.93597165 |
| -0.1520703 | -0.149165 | 4.73134967 | 0.15468207 | 0.69410057 | 0.93597165 |
| -0.1408274 | -0.1419301 | 3.96830405 | 0.15438629 | 0.69437841 | 0.93597165 |
| -0.140753 | -0.1528246 | 3.25038021 | 0.15350528 | 0.69520785 | 0.93621198 |
| -0.1487029 | -0.1602331 | 2.84131074 | 0.1527568 | 0.69591466 | 0.93621198 |
| -0.1653594 | -0.1657478 | 3.16558828 | 0.15195776 | 0.69667143 | 0.93621198 |
| 0.05520115 | 0.21975319 | 4.55756157 | 0.15188945 | 0.69673623 | 0.93621198 |
| 0.14268397 | 0.13223611 | 6.21820226 | 0.15174633 | 0.69687206 | 0.93621198 |
| 0.12829672 | 0.13431832 | 4.47844709 | 0.15134397 | 0.6972543 | 0.93621198 |
| 0.14837167 | 0.14961848 | 8.85600814 | 0.15128961 | 0.697306 | 0.93621198 |
| -0.1249259 | -0.1331817 | 4.6866428 | 0.15098015 | 0.69760045 | 0.93621198 |
| 0.13834365 | 0.14271246 | 6.04983455 | 0.15064093 | 0.69792361 | 0.93621198 |
| -0.1653017 | -0.1499517 | 3.68368954 | 0.15051284 | 0.69804575 | 0.93621198 |
| -0.1415943 | -0.1603592 | 3.01536525 | 0.15044001 | 0.69811522 | 0.93621198 |
| 0.22971372 | 0.18787261 | 3.07716947 | 0.14954685 | 0.69896878 | 0.93632842 |
| -0.1476789 | -0.1408915 | 3.34308089 | 0.14952511 | 0.69898959 | 0.93632842 |

| | | | | | |
|---|---|---|---|---|---|
| -0.1327747 | -0.1383984 | 7.42123005 | 0.14933395 | 0.69917267 | 0.93632842 |
| 0.15559643 | 0.13742785 | 3.68858868 | 0.14770857 | 0.70073482 | 0.93728961 |
| 0.15412232 | 0.1661341 | 2.98653577 | 0.14756385 | 0.70087438 | 0.93728961 |
| 0.125623 | 0.13477186 | 5.38991516 | 0.14733384 | 0.70109637 | 0.93728961 |
| -0.13765 | -0.1390757 | 4.63735523 | 0.14724113 | 0.7011859 | 0.93728961 |
| 0.13612889 | 0.13895406 | 4.00683353 | 0.14496321 | 0.70339593 | 0.93956049 |
| -0.1099684 | -0.1259765 | 5.38887837 | 0.14465979 | 0.7036918 | 0.93956049 |
| -0.1370653 | -0.1254953 | 6.28552196 | 0.14405708 | 0.70428058 | 0.93956049 |
| 0.16332514 | 0.17921711 | 2.65640746 | 0.14404705 | 0.70429039 | 0.93956049 |
| 0.15286483 | 0.15563061 | 3.34564985 | 0.14382462 | 0.70450804 | 0.93956049 |
| 0.11569488 | 0.13473354 | 5.05231871 | 0.14296309 | 0.70535289 | 0.93997208 |
| -0.2399904 | -0.1865263 | 2.80025949 | 0.14238762 | 0.70591883 | 0.93997208 |
| -0.1319887 | -0.1380656 | 4.74234247 | 0.14215786 | 0.70614516 | 0.93997208 |
| -0.1095234 | -0.1409284 | 3.27363291 | 0.14203665 | 0.70626464 | 0.93997208 |
| -0.1453786 | -0.1366045 | 5.0244989 | 0.14162659 | 0.70666929 | 0.93997208 |
| 1.31162167 | 0.50988856 | 3.3460467 | 0.14152922 | 0.70676546 | 0.93997208 |
| -0.1301168 | -0.1290015 | 4.49752889 | 0.14074143 | 0.70754504 | 0.94026513 |
| -0.1142404 | -0.1278814 | 4.39159221 | 0.14031746 | 0.70796561 | 0.94026513 |
| -0.1321857 | -0.1337589 | 7.81132048 | 0.14002727 | 0.70825391 | 0.94026513 |
| 0.13438262 | 0.14202455 | 3.98564282 | 0.13965131 | 0.7086279 | 0.94026513 |
| -0.1395497 | -0.1478425 | 3.13962017 | 0.13959464 | 0.70868433 | 0.94026513 |
| -0.137685 | -0.1318585 | 4.51248945 | 0.13934282 | 0.70893522 | 0.94026513 |
| -0.1386997 | -0.1409791 | 8.68993717 | 0.13827072 | 0.71000623 | 0.94075412 |
| -0.1140925 | -0.1204199 | 6.46678712 | 0.13816546 | 0.71011165 | 0.94075412 |
| 0.11433494 | 0.1445649 | 3.80351346 | 0.13799832 | 0.71027911 | 0.94075412 |
| 0.13817821 | 0.13645965 | 3.85423833 | 0.13712805 | 0.71115298 | 0.94148066 |
| 0.16064702 | 0.15539325 | 2.65836602 | 0.13638488 | 0.71190171 | 0.94204095 |
| 0.1332646 | 0.12772405 | 5.51108058 | 0.13506733 | 0.71323485 | 0.94267477 |
| -0.1575648 | -0.141871 | 3.1433824 | 0.13493629 | 0.71336785 | 0.94267477 |
| -0.1506518 | -0.1547775 | 3.65226362 | 0.13468282 | 0.7136253 | 0.94267477 |
| -0.4540031 | -0.212804 | 2.90979604 | 0.13462543 | 0.71368363 | 0.94267477 |
| 0.14085936 | 0.14191591 | 4.6826469 | 0.13352544 | 0.71480439 | 0.94372441 |
| -0.1339356 | -0.1480871 | 2.98681627 | 0.13255759 | 0.71579484 | 0.94431985 |
| -0.1277022 | -0.1298238 | 4.76350041 | 0.13237687 | 0.71598023 | 0.94431985 |
| -0.1581408 | -0.142015 | 3.02391108 | 0.13212944 | 0.7162343 | 0.94431985 |
| -0.114631 | -0.1271778 | 4.65142565 | 0.13157675 | 0.71680278 | 0.94444599 |
| -0.1302088 | -0.1300386 | 5.76442937 | 0.13140213 | 0.71698267 | 0.94444599 |
| -0.1612786 | -0.1403096 | 4.5888236 | 0.13079157 | 0.71761272 | 0.94446058 |
| -0.1444992 | -0.1386648 | 3.5965086 | 0.13075893 | 0.71764645 | 0.94446058 |
| 0.3175049 | 0.26480218 | 2.87191807 | 0.13043447 | 0.71798199 | 0.94447267 |
| 0.11972216 | 0.16157676 | 3.09672932 | 0.12975296 | 0.71868832 | 0.94497228 |
| 0.12530578 | 0.12146328 | 4.04294072 | 0.12888467 | 0.71959126 | 0.94572984 |
| 0.19795384 | 0.17352912 | 3.03290399 | 0.1279565 | 0.72056029 | 0.94611417 |
| -0.1562162 | -0.1266922 | 3.91744113 | 0.12749403 | 0.7210446 | 0.94611417 |
| 0.11797992 | 0.12340986 | 4.5921108 | 0.12743659 | 0.72110482 | 0.94611417 |
| 0.15541994 | 0.13210936 | 3.56687125 | 0.12735405 | 0.72119138 | 0.94611417 |

| | | | | | |
|---|---|---|---|---|---|
| 0.12634631 | 0.13337091 | 3.49147383 | 0.12591927 | 0.72270118 | 0.94734671 |
| -0.1268156 | -0.1202729 | 5.325988 | 0.12583931 | 0.7227856 | 0.94734671 |
| 0.13327139 | 0.12581418 | 3.73251421 | 0.12550094 | 0.72314321 | 0.94738635 |
| 0.14456504 | 0.14458207 | 3.49424043 | 0.12490947 | 0.72376959 | 0.94741785 |
| -0.1066165 | -0.1196709 | 4.60740769 | 0.12472318 | 0.72396722 | 0.94741785 |
| -0.1528665 | -0.154771 | 2.86198137 | 0.12455162 | 0.72414938 | 0.94741785 |
| 0.11962013 | 0.13701024 | 4.37806703 | 0.12418007 | 0.72454435 | 0.94750626 |
| -0.1548162 | -0.1457329 | 3.03170749 | 0.12371899 | 0.72503543 | 0.94772021 |
| 0.11748918 | 0.13478817 | 4.54445644 | 0.12299119 | 0.72581267 | 0.94812667 |
| -0.1068903 | -0.1175976 | 4.9994764 | 0.12252492 | 0.72631198 | 0.94812667 |
| -0.1355368 | -0.1261375 | 4.69844267 | 0.12242987 | 0.72641389 | 0.94812667 |
| 0.18836013 | 0.16311006 | 3.11686685 | 0.12192582 | 0.72695509 | 0.94812667 |
| 0.13783291 | 0.12232027 | 3.88630509 | 0.12189849 | 0.72698448 | 0.94812667 |
| -0.1399959 | -0.1310059 | 5.0476992 | 0.12099166 | 0.72796143 | 0.94897315 |
| -0.1305273 | -0.1331902 | 7.53235347 | 0.12033671 | 0.72866959 | 0.94946861 |
| 0.11810902 | 0.13369081 | 3.12763795 | 0.11944663 | 0.72963545 | 0.94955712 |
| -0.1106385 | -0.1354335 | 3.09691203 | 0.11928599 | 0.72981018 | 0.94955712 |
| -0.1410659 | -0.1295957 | 4.47468436 | 0.11914117 | 0.72996783 | 0.94955712 |
| -0.1049982 | -0.113284 | 4.03593286 | 0.11834634 | 0.73083498 | 0.94955712 |
| 0.14817062 | 0.1194601 | 3.63693615 | 0.11824194 | 0.73094912 | 0.94955712 |
| 0.18181999 | 0.15914125 | 3.10859089 | 0.11808266 | 0.73112337 | 0.94955712 |
| 0.10816573 | 0.11281973 | 6.46057566 | 0.11796582 | 0.73125128 | 0.94955712 |
| 0.12792707 | 0.16201913 | 2.97547251 | 0.11786435 | 0.73136241 | 0.94955712 |
| 0.11766907 | 0.12780885 | 4.12084898 | 0.11637094 | 0.73300432 | 0.95087371 |
| -0.1218759 | -0.1252172 | 7.18387299 | 0.11590974 | 0.73351376 | 0.95087371 |
| 0.12832532 | 0.11576292 | 4.76938465 | 0.11539944 | 0.73407875 | 0.95087371 |
| -0.1070143 | -0.1210062 | 4.42636608 | 0.11513176 | 0.73437567 | 0.95087371 |
| -0.1530896 | -0.1686549 | 2.89671493 | 0.11483523 | 0.73470505 | 0.95087371 |
| 0.1519455 | 0.14939248 | 3.26117924 | 0.11478948 | 0.7347559 | 0.95087371 |
| 0.16115993 | 0.13483347 | 3.14570478 | 0.11471424 | 0.73483958 | 0.95087371 |
| 0.11479838 | 0.11336869 | 6.03423379 | 0.11434546 | 0.73525011 | 0.95087371 |
| -0.1275264 | -0.1214999 | 3.57601289 | 0.11427057 | 0.73533357 | 0.95087371 |
| 0.14505954 | 0.13851668 | 3.03219654 | 0.1138483 | 0.73580473 | 0.95105801 |
| 0.17976488 | 0.16033783 | 3.03546401 | 0.11193156 | 0.7379557 | 0.95282646 |
| -0.1276157 | -0.1424119 | 3.59392698 | 0.11152892 | 0.73841016 | 0.95282646 |
| 0.08438204 | 0.1395308 | 3.828381 | 0.11134733 | 0.73861541 | 0.95282646 |
| -0.1521041 | -0.1536066 | 3.02288061 | 0.11054175 | 0.7395282 | 0.95282646 |
| 0.12072874 | 0.10896399 | 6.04747365 | 0.1105307 | 0.73954075 | 0.95282646 |
| 0.13314667 | 0.11181439 | 3.88684313 | 0.11035982 | 0.73973486 | 0.95282646 |
| -0.1124118 | -0.1088783 | 5.54730932 | 0.11029903 | 0.73980395 | 0.95282646 |
| -0.1400868 | -0.1972877 | 3.0018583 | 0.1101647 | 0.7399567 | 0.95282646 |
| -0.1620956 | -0.1452534 | 7.01828211 | 0.10981707 | 0.74035249 | 0.95282646 |
| 0.07356817 | 0.11643045 | 3.85596982 | 0.1095897 | 0.74061174 | 0.95282646 |
| -0.1241722 | -0.1117561 | 4.84343035 | 0.10942949 | 0.74079459 | 0.95282646 |
| -0.150052 | -0.1439915 | 2.74429636 | 0.10807668 | 0.74234452 | 0.95403739 |
| 0.11252069 | 0.11971669 | 3.58955234 | 0.10803246 | 0.74239537 | 0.95403739 |

| | | | | | |
|---|---|---|---|---|---|
| 0.12485387 | 0.14745824 | 2.56951533 | 0.1072862 | 0.74325517 | 0.95471835 |
| 0.11682612 | 0.11309755 | 4.53226623 | 0.10692594 | 0.74367142 | 0.95482924 |
| -0.1266018 | -0.1316204 | 3.6807056 | 0.10559295 | 0.7452184 | 0.95639115 |
| -0.134967 | -0.1452829 | 3.24667574 | 0.10520834 | 0.74566675 | 0.95654237 |
| -0.1329438 | -0.1377694 | 2.9572973 | 0.10469712 | 0.7462641 | 0.95672639 |
| -0.0661842 | -0.1748391 | 5.656324 | 0.10452005 | 0.74647138 | 0.95672639 |
| -0.1283077 | -0.1131849 | 4.89424432 | 0.10380651 | 0.74730862 | 0.95693086 |
| -0.141385 | -0.1167916 | 3.57717376 | 0.1037544 | 0.74736989 | 0.95693086 |
| -0.1430828 | -0.13747 | 3.05443552 | 0.10350601 | 0.74766217 | 0.95693086 |
| 0.14850103 | 0.14055987 | 2.77604397 | 0.10192867 | 0.74952732 | 0.95693086 |
| -0.0774081 | -0.1051713 | 4.13015018 | 0.10145993 | 0.75008464 | 0.95693086 |
| 0.1156018 | 0.12308964 | 3.52311493 | 0.10143789 | 0.75011088 | 0.95693086 |
| 0.12811864 | 0.13311671 | 3.07179565 | 0.10129821 | 0.75027726 | 0.95693086 |
| -0.1086782 | -0.1066693 | 7.04186969 | 0.10120452 | 0.75038893 | 0.95693086 |
| -0.109701 | -0.1129017 | 6.34116117 | 0.10095194 | 0.75069024 | 0.95693086 |
| -0.1076348 | -0.1124529 | 6.09019076 | 0.10085973 | 0.75080036 | 0.95693086 |
| 0.01258215 | -0.1699718 | 4.24713681 | 0.10041987 | 0.75132636 | 0.95693086 |
| -0.1292084 | -0.1300648 | 2.94152776 | 0.09966888 | 0.75222735 | 0.95693086 |
| -0.0874105 | -0.1019093 | 5.94628914 | 0.0994574 | 0.75248175 | 0.95693086 |
| -0.1131214 | -0.1157504 | 3.70031682 | 0.09911922 | 0.75288917 | 0.95693086 |
| 0.11130832 | 0.10523643 | 6.21187197 | 0.09909379 | 0.75291985 | 0.95693086 |
| -0.0915952 | -0.1071813 | 5.05838019 | 0.09907577 | 0.75294158 | 0.95693086 |
| -0.1247935 | -0.118052 | 3.10309003 | 0.09898364 | 0.75305273 | 0.95693086 |
| -0.109296 | -0.1094913 | 4.7062283 | 0.09851147 | 0.7536233 | 0.95693086 |
| 0.11851152 | 0.13669804 | 2.75729343 | 0.09802481 | 0.75421293 | 0.95693086 |
| -0.1057437 | -0.1092485 | 7.49899482 | 0.09787188 | 0.75439856 | 0.95693086 |
| -0.1104125 | -0.103708 | 5.79105784 | 0.09769024 | 0.75461925 | 0.95693086 |
| 0.11627767 | 0.11282678 | 5.16099993 | 0.0970319 | 0.75542097 | 0.95693086 |
| 0.1010483 | 0.11485569 | 4.05778208 | 0.09702848 | 0.75542515 | 0.95693086 |
| 0.56888827 | 0.19953313 | 3.09078035 | 0.0967609 | 0.75575186 | 0.95693086 |
| -0.0973454 | -0.1001172 | 5.94818481 | 0.09671512 | 0.75580781 | 0.95693086 |
| 0.10499022 | 0.10424614 | 4.87001644 | 0.09666189 | 0.75587288 | 0.95693086 |
| -0.1252227 | -0.1091828 | 3.39453897 | 0.09654451 | 0.75601644 | 0.95693086 |
| -0.1011697 | -0.1040864 | 6.5298895 | 0.09644959 | 0.7561326 | 0.95693086 |
| -0.118028 | -0.1208007 | 3.13549397 | 0.09622327 | 0.75640982 | 0.95693086 |
| -0.1608305 | -0.1297865 | 3.68724867 | 0.09610834 | 0.75655073 | 0.95693086 |
| 0.00219238 | 0.1542926 | 3.1729803 | 0.09568182 | 0.75707449 | 0.957175 |
| -0.1322573 | -0.1238664 | 3.3356547 | 0.09466295 | 0.75833082 | 0.95834471 |
| -0.1184458 | -0.1131507 | 3.45744557 | 0.09406782 | 0.7590681 | 0.95864594 |
| -0.1138544 | -0.1156107 | 3.58878408 | 0.09388517 | 0.75929488 | 0.95864594 |
| 0.14086008 | 0.14093875 | 2.8939404 | 0.09349477 | 0.75978042 | 0.95864594 |
| -0.1242279 | -0.120582 | 3.33923775 | 0.09307963 | 0.76029795 | 0.95864594 |
| -0.076787 | -0.106903 | 3.36572249 | 0.09302794 | 0.76036248 | 0.95864594 |
| -0.1377729 | -0.1189094 | 3.36845702 | 0.09285733 | 0.7605756 | 0.95864594 |
| -0.1578228 | -0.1364809 | 3.06218058 | 0.09260759 | 0.76088795 | 0.95864594 |
| 0.13339539 | 0.11710773 | 3.40102648 | 0.09176367 | 0.76194687 | 0.95897821 |

| | | | | | |
|---|---|---|---|---|---|
| -0.1185048 | -0.1038587 | 5.3895469 | 0.09167094 | 0.76206356 | 0.95897821 |
| 0.12086904 | 0.12302115 | 3.28677978 | 0.09148317 | 0.76230002 | 0.95897821 |
| 0.10110386 | 0.10153302 | 5.88553821 | 0.09109976 | 0.76278366 | 0.95897821 |
| -0.0957304 | -0.0984721 | 6.19109115 | 0.09108008 | 0.76280852 | 0.95897821 |
| -0.0873024 | -0.1011673 | 5.61890878 | 0.09002301 | 0.7641479 | 0.96012541 |
| -0.3140424 | -0.2544639 | 7.55040367 | 0.08983693 | 0.76438457 | 0.96012541 |
| 0.14518589 | 0.12369021 | 3.02557412 | 0.0895104 | 0.76480051 | 0.96017412 |
| 0.13054501 | 0.12893159 | 3.25594777 | 0.08889552 | 0.765586 | 0.96017412 |
| 0.10242428 | 0.11014072 | 3.83355676 | 0.08835762 | 0.76627557 | 0.96017412 |
| 0.11188243 | 0.11272154 | 4.56153723 | 0.08817934 | 0.76650463 | 0.96017412 |
| -0.120692 | -0.1264731 | 3.03853605 | 0.08788039 | 0.7668893 | 0.96017412 |
| 0.11225064 | 0.11410163 | 4.28807758 | 0.08771985 | 0.76709616 | 0.96017412 |
| -0.0947833 | -0.1039374 | 4.45432566 | 0.08756244 | 0.76729919 | 0.96017412 |
| -0.0919675 | -0.0997172 | 5.02600417 | 0.08742905 | 0.76747139 | 0.96017412 |
| -0.0931596 | -0.1013573 | 3.94173708 | 0.08705255 | 0.76795822 | 0.96017412 |
| 0.11066343 | 0.11275073 | 3.48950933 | 0.08696555 | 0.76807086 | 0.96017412 |
| -0.3023104 | -0.147686 | 3.11910698 | 0.08684612 | 0.76822562 | 0.96017412 |
| 0.11085402 | 0.1143536 | 3.79684472 | 0.086708 | 0.76840472 | 0.96017412 |
| 0.13916851 | 0.11060042 | 3.38253678 | 0.08618173 | 0.76908861 | 0.96045773 |
| 0.08499266 | 0.10865063 | 3.85428533 | 0.0860229 | 0.76929544 | 0.96045773 |
| -0.1178748 | -0.1171558 | 3.4245957 | 0.08526582 | 0.77028424 | 0.9605775 |
| 0.11203614 | 0.10034441 | 5.39348326 | 0.08505914 | 0.77055501 | 0.9605775 |
| 0.11015858 | 0.10460488 | 3.78128857 | 0.08477773 | 0.77092424 | 0.9605775 |
| -0.0981813 | -0.1085335 | 3.63839521 | 0.0846828 | 0.77104896 | 0.9605775 |
| 0.09303865 | 0.10031972 | 4.01776742 | 0.08468126 | 0.77105098 | 0.9605775 |
| 0.10892484 | 0.09783212 | 4.74980965 | 0.08365611 | 0.77240257 | 0.96184727 |
| 0.10854105 | 0.10385005 | 3.88062015 | 0.08224428 | 0.77427876 | 0.96376892 |
| -0.099199 | -0.1156157 | 2.9076136 | 0.0817026 | 0.77500323 | 0.96407076 |
| -0.0821741 | -0.1055562 | 3.40088698 | 0.08156512 | 0.77518751 | 0.96407076 |
| 0.11403603 | 0.09655864 | 4.97388234 | 0.08102412 | 0.77591433 | 0.96456016 |
| -0.0902548 | -0.0951768 | 4.70723919 | 0.08018234 | 0.77705046 | 0.96472305 |
| -0.0920658 | -0.1016564 | 3.73545018 | 0.08002206 | 0.77726752 | 0.96472305 |
| 0.3066832 | 0.14409092 | 3.27770153 | 0.07916137 | 0.77843713 | 0.96472305 |
| -0.0974412 | -0.0969632 | 5.22216701 | 0.07864844 | 0.77913743 | 0.96472305 |
| -0.0407311 | 0.147915 | 3.11453061 | 0.07855137 | 0.77927023 | 0.96472305 |
| -0.0909127 | -0.0975401 | 4.0044669 | 0.07822116 | 0.77972267 | 0.96472305 |
| 0.15423299 | 0.1234223 | 2.69059545 | 0.07802138 | 0.7799969 | 0.96472305 |
| 0.12427504 | 0.11535242 | 3.58997269 | 0.07789314 | 0.78017313 | 0.96472305 |
| 0.11366215 | 0.11702208 | 3.03175235 | 0.07719956 | 0.78112898 | 0.96472305 |
| 0.10820441 | 0.10650124 | 6.31243824 | 0.07705003 | 0.78133565 | 0.96472305 |
| -0.1154223 | -0.1005277 | 4.90666935 | 0.07675323 | 0.78174653 | 0.96472305 |
| -0.081134 | -0.0961318 | 3.76550952 | 0.07635902 | 0.78229357 | 0.96472305 |
| -0.081134 | -0.0961318 | 3.76550952 | 0.07635902 | 0.78229357 | 0.96472305 |
| -0.0914639 | -0.0930526 | 6.64638284 | 0.07624253 | 0.78245551 | 0.96472305 |
| 0.09970812 | 0.0991401 | 5.13774681 | 0.07609399 | 0.78266221 | 0.96472305 |
| 0.10562962 | 0.1064795 | 8.10635809 | 0.07603117 | 0.78274969 | 0.96472305 |

| | | | | | |
|---|---|---|---|---|---|
| 0.14757438 | 0.12230519 | 3.36515144 | 0.07575518 | 0.78313448 | 0.96472305 |
| 0.09513661 | 0.0989604 | 5.13350135 | 0.07496841 | 0.78423555 | 0.96472305 |
| -0.1033394 | -0.1018533 | 4.84346902 | 0.07478569 | 0.78449216 | 0.96472305 |
| 0.08353018 | 0.09656042 | 4.72225745 | 0.07434429 | 0.78511344 | 0.96472305 |
| 0.109537 | 0.09733756 | 3.89656932 | 0.07421169 | 0.78530046 | 0.96472305 |
| 0.09503305 | 0.09772594 | 4.363987 | 0.07383134 | 0.78583791 | 0.96472305 |
| 0.11784428 | 0.13296745 | 2.39982646 | 0.0737917 | 0.78589402 | 0.96472305 |
| 0.0938659 | 0.09394723 | 5.42095371 | 0.07342987 | 0.78640682 | 0.96472305 |
| -0.0932432 | -0.0940867 | 7.46062468 | 0.07256339 | 0.78764041 | 0.96472305 |
| -0.0856998 | -0.0893924 | 6.04597724 | 0.0725609 | 0.78764396 | 0.96472305 |
| -0.0929516 | -0.0936464 | 4.77391333 | 0.07241932 | 0.78784628 | 0.96472305 |
| 0.08791505 | 0.08939203 | 5.83457047 | 0.07236499 | 0.78792398 | 0.96472305 |
| 0.11873938 | 0.10376756 | 3.07384199 | 0.07213289 | 0.78825624 | 0.96472305 |
| -0.0907908 | -0.0967075 | 3.33664203 | 0.0719947 | 0.78845433 | 0.96472305 |
| -0.0892029 | -0.0900464 | 6.44649983 | 0.07197111 | 0.78848817 | 0.96472305 |
| 0.11758281 | 0.11778351 | 9.49214362 | 0.07164008 | 0.78896363 | 0.96472305 |
| -0.0947898 | -0.0964949 | 4.07905447 | 0.07140833 | 0.78929718 | 0.96472305 |
| -0.1247725 | -0.0937546 | 3.56479673 | 0.07123783 | 0.78954295 | 0.96472305 |
| -0.0893619 | -0.0956388 | 4.3223636 | 0.07108268 | 0.78976688 | 0.96472305 |
| 0.1105213 | 0.11371261 | 3.13329284 | 0.07107992 | 0.78977086 | 0.96472305 |
| -0.1606519 | -0.1118628 | 4.96341139 | 0.07085167 | 0.79010077 | 0.96472305 |
| -0.085898 | -0.0948317 | 4.83466961 | 0.07047988 | 0.79063935 | 0.96472305 |
| -0.082646 | -0.0886112 | 5.31528181 | 0.07017463 | 0.79108268 | 0.96472305 |
| 0.0963818 | 0.08782532 | 6.8160286 | 0.07014072 | 0.791132 | 0.96472305 |
| -0.1250564 | -0.1152485 | 3.0119767 | 0.06995634 | 0.79140036 | 0.96472305 |
| -0.1243982 | -0.1120523 | 5.65220429 | 0.06976788 | 0.79167504 | 0.96472305 |
| 0.39460488 | 0.16362506 | 2.87583755 | 0.06967656 | 0.79180829 | 0.96472305 |
| 0.10769997 | 0.12789847 | 4.19204525 | 0.06958997 | 0.79193472 | 0.96472305 |
| -0.111049 | -0.1061342 | 3.52779714 | 0.06945589 | 0.79213066 | 0.96472305 |
| -0.0646046 | -0.0904757 | 4.56786304 | 0.06920353 | 0.79249999 | 0.96472305 |
| -0.0980226 | -0.0929782 | 4.75699377 | 0.06917312 | 0.79254455 | 0.96472305 |
| 0.08826404 | 0.09549114 | 5.03498927 | 0.06868122 | 0.79326669 | 0.96472305 |
| -0.1106065 | -0.1004972 | 3.8117311 | 0.06851267 | 0.79351477 | 0.96472305 |
| -0.0962219 | -0.096781 | 3.7739937 | 0.06847067 | 0.79357663 | 0.96472305 |
| -0.1513728 | -0.1154472 | 3.07113141 | 0.06841908 | 0.79365265 | 0.96472305 |
| 0.10519487 | 0.09338922 | 4.45890462 | 0.06839673 | 0.7936856 | 0.96472305 |
| 0.267819 | 0.13162816 | 3.54260407 | 0.06825158 | 0.7938997 | 0.96472305 |
| -0.0799074 | -0.1027973 | 3.11656924 | 0.06736598 | 0.79521124 | 0.96472305 |
| -0.0730828 | -0.1045252 | 2.9284302 | 0.06733049 | 0.79526398 | 0.96472305 |
| 0.10210748 | 0.08870568 | 3.6870263 | 0.06730901 | 0.79529593 | 0.96472305 |
| -0.1086175 | -0.1141674 | 2.87143698 | 0.06691753 | 0.79587889 | 0.96472305 |
| 0.05549777 | 0.09552419 | 3.22712616 | 0.06653025 | 0.7964574 | 0.96472305 |
| -0.1499563 | -0.119714 | 2.97203476 | 0.06619466 | 0.79696013 | 0.96472305 |
| -0.1099916 | -0.103386 | 3.75829678 | 0.06598584 | 0.79727365 | 0.96472305 |
| 0.10332085 | 0.08947068 | 3.79958352 | 0.06536211 | 0.79821327 | 0.96472305 |
| -0.0791634 | -0.0930703 | 3.79392639 | 0.06505622 | 0.79867582 | 0.96472305 |

| | | | | | |
|---|---|---|---|---|---|
| 0.11795097 | 0.10530446 | 3.0522897 | 0.06472505 | 0.79917791 | 0.96472305 |
| -0.1042977 | -0.0981763 | 6.3424846 | 0.06461674 | 0.79934242 | 0.96472305 |
| 0.16730718 | 0.11347553 | 4.18993666 | 0.06461377 | 0.79934694 | 0.96472305 |
| 0.089558 | 0.11912715 | 2.95112528 | 0.06440239 | 0.79966842 | 0.96472305 |
| 0.08423468 | 0.08565485 | 6.93253091 | 0.06414141 | 0.80006612 | 0.96472305 |
| -0.082497 | -0.0858336 | 3.79269328 | 0.06396518 | 0.80033515 | 0.96472305 |
| 0.07486132 | 0.08973234 | 4.96135472 | 0.06368149 | 0.80076906 | 0.96472305 |
| 0.10708492 | 0.10873333 | 3.25900962 | 0.06356537 | 0.80094697 | 0.96472305 |
| -0.0695432 | -0.0821153 | 4.11937997 | 0.06306294 | 0.80171873 | 0.96472305 |
| -0.0762463 | -0.0896596 | 3.82594898 | 0.06300063 | 0.80181468 | 0.96472305 |
| -0.0668471 | -0.0842543 | 3.91164103 | 0.06299045 | 0.80183035 | 0.96472305 |
| -0.0583737 | -0.0817589 | 4.12400584 | 0.06297252 | 0.80185798 | 0.96472305 |
| -0.1038365 | -0.091402 | 4.1044769 | 0.06271644 | 0.80225289 | 0.96472305 |
| -0.1672984 | -0.1180085 | 3.09819599 | 0.06267099 | 0.80232308 | 0.96472305 |
| -0.098557 | -0.0930487 | 3.79442195 | 0.06257616 | 0.8024696 | 0.96472305 |
| -0.0986627 | -0.0942862 | 5.12303433 | 0.06125791 | 0.80451869 | 0.96472305 |
| -0.0738494 | -0.0825503 | 5.98518981 | 0.06118983 | 0.80462516 | 0.96472305 |
| 0.1090376 | 0.10048142 | 3.97380295 | 0.06103433 | 0.80486854 | 0.96472305 |
| 0.12019102 | 0.10170358 | 2.76435905 | 0.06102938 | 0.80487629 | 0.96472305 |
| -0.1234343 | -0.1089286 | 3.18317947 | 0.06102725 | 0.80487964 | 0.96472305 |
| 0.08908797 | 0.0930635 | 3.7989209 | 0.06095659 | 0.80499035 | 0.96472305 |
| 0.12090071 | 0.12022201 | 2.61994069 | 0.06090464 | 0.8050718 | 0.96472305 |
| -0.1724099 | -0.114973 | 2.94713243 | 0.0608876 | 0.80509852 | 0.96472305 |
| -0.0842928 | -0.0885066 | 3.90234604 | 0.06043119 | 0.80581572 | 0.96472305 |
| 0.11435029 | 0.10380244 | 2.73916978 | 0.06032872 | 0.80597713 | 0.96472305 |
| -0.1035121 | -0.0939529 | 3.62370029 | 0.06030409 | 0.80601596 | 0.96472305 |
| -0.0820912 | -0.0849183 | 3.83114104 | 0.05989347 | 0.80666439 | 0.96472305 |
| -0.0749032 | -0.0884094 | 3.81321637 | 0.0598466 | 0.80673855 | 0.96472305 |
| -0.07485 | -0.0802128 | 3.86813479 | 0.05980213 | 0.80680895 | 0.96472305 |
| -0.1237293 | -0.0930593 | 3.22027468 | 0.05960354 | 0.80712365 | 0.96472305 |
| -0.1696286 | -0.1169066 | 2.74896314 | 0.05930365 | 0.80759996 | 0.96472305 |
| 0.09681195 | 0.09521813 | 7.69230718 | 0.05924643 | 0.80769098 | 0.96472305 |
| -0.0772478 | -0.0810443 | 5.51706896 | 0.05918648 | 0.80778639 | 0.96472305 |
| 0.08872666 | 0.09149266 | 5.61034787 | 0.05902276 | 0.80804723 | 0.96472305 |
| 0.09209201 | 0.09208814 | 3.90204984 | 0.05852169 | 0.80884791 | 0.96528076 |
| -0.1088691 | -0.087763 | 4.2218575 | 0.0579496 | 0.80976653 | 0.96595646 |
| -0.0781612 | -0.0844638 | 3.66167744 | 0.05775403 | 0.81008167 | 0.96595646 |
| -0.0613716 | -0.0805752 | 5.26308095 | 0.05708461 | 0.81116462 | 0.96671671 |
| 0.08806984 | 0.08477894 | 5.04905892 | 0.05691483 | 0.81144036 | 0.96671671 |
| 0.10169577 | 0.094873 | 3.29235037 | 0.05667575 | 0.81182937 | 0.96671671 |
| 0.09562959 | 0.08569761 | 3.4288294 | 0.05653708 | 0.8120554 | 0.96671671 |
| 0.12691434 | 0.10163736 | 4.70819965 | 0.055923 | 0.81305988 | 0.96690891 |
| -0.0907418 | -0.0876154 | 4.2127793 | 0.05582819 | 0.81321548 | 0.96690891 |
| -0.0772493 | -0.0805956 | 7.10934815 | 0.0557411 | 0.81335853 | 0.96690891 |
| -0.0879157 | -0.0874432 | 4.34155941 | 0.05562266 | 0.81355328 | 0.96690891 |
| 0.07968901 | 0.08038075 | 6.6620669 | 0.05452895 | 0.81536204 | 0.96809521 |

| | | | | | |
|---|---|---|---|---|---|
| -0.0674056 | -0.0804911 | 4.05546249 | 0.05451897 | 0.81537862 | 0.96809521 |
| -0.092704 | -0.0787677 | 6.41436584 | 0.054182 | 0.8159398 | 0.96809521 |
| -0.0723691 | -0.0802085 | 6.2292071 | 0.05390113 | 0.81640896 | 0.96809521 |
| -0.1265019 | -0.0999965 | 4.26848946 | 0.05349057 | 0.81709707 | 0.96809521 |
| -0.059653 | -0.0871078 | 3.05679661 | 0.05337113 | 0.81729777 | 0.96809521 |
| -0.0967145 | -0.0878192 | 3.15846223 | 0.0532368 | 0.81752379 | 0.96809521 |
| 0.09243585 | 0.08989235 | 3.83306331 | 0.05312408 | 0.81771367 | 0.96809521 |
| 0.06330164 | 0.0777891 | 5.86277794 | 0.0530674 | 0.81780922 | 0.96809521 |
| -0.0837832 | -0.0863893 | 8.2999394 | 0.05291601 | 0.81806473 | 0.96809521 |
| -0.1055474 | -0.0829888 | 3.47685908 | 0.05281754 | 0.81823113 | 0.96809521 |
| 0.07927089 | 0.0805771 | 6.70579525 | 0.05245569 | 0.81884399 | 0.96828657 |
| -0.0886406 | -0.0954588 | 3.09620101 | 0.05232727 | 0.81906203 | 0.96828657 |
| 0.10009306 | 0.07998483 | 4.30227131 | 0.05210875 | 0.8194337 | 0.96833039 |
| 0.08856296 | 0.12977345 | 3.14168397 | 0.0507788 | 0.8217136 | 0.97041311 |
| 0.09297774 | 0.10357458 | 2.78870507 | 0.05069008 | 0.82186681 | 0.97041311 |
| -0.0892614 | -0.0838169 | 5.35482656 | 0.05025779 | 0.82261531 | 0.97090077 |
| -0.075183 | -0.0794351 | 5.25293025 | 0.04979581 | 0.82341896 | 0.97126209 |
| -0.0678717 | -0.0852712 | 3.10089194 | 0.04969625 | 0.82359266 | 0.97126209 |
| -0.0817732 | -0.0842535 | 8.75539679 | 0.04938678 | 0.82413377 | 0.97150433 |
| 0.05724921 | 0.07282298 | 5.89778957 | 0.04877937 | 0.82520104 | 0.9722122 |
| -0.0808826 | -0.0844823 | 8.50458064 | 0.04842259 | 0.82583118 | 0.9722122 |
| -0.1264884 | -0.1123873 | 2.90076355 | 0.04840319 | 0.82586551 | 0.9722122 |
| -0.093396 | -0.090145 | 4.69273024 | 0.04820323 | 0.82621981 | 0.9722122 |
| -0.0807716 | -0.0726403 | 5.361429 | 0.04809384 | 0.82641396 | 0.9722122 |
| 0.08022638 | 0.08111441 | 3.32896537 | 0.04714528 | 0.82810731 | 0.97380844 |
| 0.10523854 | 0.09645532 | 2.73573033 | 0.04590456 | 0.83034938 | 0.9747164 |
| 0.09325743 | 0.08899653 | 7.03534389 | 0.04589596 | 0.83036504 | 0.9747164 |
| -0.0672443 | -0.0689216 | 5.90197593 | 0.04580773 | 0.83052568 | 0.9747164 |
| 0.08638159 | 0.07469393 | 3.60267425 | 0.04544453 | 0.83118873 | 0.9747164 |
| -0.1364747 | -0.08796 | 3.52483706 | 0.04497255 | 0.83205451 | 0.9747164 |
| 0.08158163 | 0.08550033 | 3.40194632 | 0.04484995 | 0.83228018 | 0.9747164 |
| -0.0961301 | -0.0917019 | 3.22051338 | 0.04446192 | 0.83299656 | 0.9747164 |
| -0.0735055 | -0.0777295 | 7.7590441 | 0.04408567 | 0.8336943 | 0.9747164 |
| -0.1036819 | -0.0928591 | 3.05825842 | 0.04399837 | 0.83385665 | 0.9747164 |
| -0.0626085 | -0.0817863 | 3.1210686 | 0.04397382 | 0.83390233 | 0.9747164 |
| 1.74466806 | 0.21528214 | 3.524072 | 0.04383619 | 0.83415868 | 0.9747164 |
| -0.068392 | -0.0690708 | 6.94999562 | 0.04364541 | 0.83451471 | 0.9747164 |
| 0.07906206 | 0.08187825 | 3.41563223 | 0.04361338 | 0.83457457 | 0.9747164 |
| -0.0535624 | -0.0751886 | 3.62939476 | 0.04354337 | 0.83470549 | 0.9747164 |
| 0.09928447 | 0.09073819 | 2.88635124 | 0.04343355 | 0.83491106 | 0.9747164 |
| 0.06992691 | 0.07472528 | 5.60270479 | 0.04328903 | 0.83518199 | 0.9747164 |
| -0.0705826 | -0.078173 | 3.63390378 | 0.04323489 | 0.83528361 | 0.9747164 |
| -0.0985319 | -0.081087 | 3.27286619 | 0.04318196 | 0.83538302 | 0.9747164 |
| 0.05622926 | 0.07523701 | 4.47874016 | 0.04305971 | 0.83561288 | 0.9747164 |
| 0.07981627 | 0.08494728 | 3.38167755 | 0.04305829 | 0.83561555 | 0.9747164 |
| -0.068812 | -0.0749411 | 3.53213493 | 0.04285487 | 0.83599879 | 0.97475654 |

| | | | | | |
|---|---|---|---|---|---|
| -0.0726821 | -0.0785296 | 3.50899482 | 0.04264809 | 0.83638933 | 0.97475654 |
| -0.0678926 | -0.073162 | 6.42875167 | 0.04250486 | 0.83666042 | 0.97475654 |
| -0.3010172 | -0.1254226 | 2.92217292 | 0.04213463 | 0.83736336 | 0.97518292 |
| -0.0887616 | -0.0891794 | 2.83036267 | 0.04091482 | 0.83970242 | 0.97533262 |
| -0.06221 | -0.0730863 | 3.65498931 | 0.04066075 | 0.84019417 | 0.97533262 |
| 0.06566927 | 0.0688536 | 5.00340692 | 0.04054603 | 0.84041673 | 0.97533262 |
| -0.0741874 | -0.0989982 | 6.74500013 | 0.04036415 | 0.84077027 | 0.97533262 |
| 0.08141984 | 0.07810282 | 3.26705197 | 0.04033738 | 0.84082238 | 0.97533262 |
| -0.0673972 | -0.0714842 | 5.08112234 | 0.04033315 | 0.84083062 | 0.97533262 |
| 0.08885639 | 0.08869381 | 3.07929355 | 0.04027131 | 0.84095105 | 0.97533262 |
| -0.0876612 | -0.0779592 | 3.19996947 | 0.04027052 | 0.84095258 | 0.97533262 |
| 0.06449813 | 0.0692181 | 6.65026098 | 0.04027042 | 0.84095279 | 0.97533262 |
| 0.07940525 | 0.07645612 | 3.29732615 | 0.04018647 | 0.84111643 | 0.97533262 |
| -0.0645043 | -0.0691054 | 5.85150297 | 0.03993106 | 0.84161544 | 0.97533262 |
| -0.188328 | -0.0897384 | 3.62560838 | 0.0398479 | 0.84177826 | 0.97533262 |
| 0.07711413 | 0.0699297 | 3.50655085 | 0.03979949 | 0.84187314 | 0.97533262 |
| -0.0730958 | -0.0755216 | 3.49846637 | 0.03958041 | 0.84230322 | 0.97544038 |
| 0.04816972 | 0.08266103 | 3.77143542 | 0.0389224 | 0.84360248 | 0.97623389 |
| 0.06409406 | 0.07066938 | 6.94337676 | 0.03887135 | 0.84370375 | 0.97623389 |
| -0.0713707 | -0.0753836 | 7.75059892 | 0.03860287 | 0.84423751 | 0.97623389 |
| -0.1005154 | -0.0867481 | 3.02917762 | 0.03855256 | 0.84433775 | 0.97623389 |
| -0.0762637 | -0.0757856 | 3.78023239 | 0.0374929 | 0.84646495 | 0.97744262 |
| 0.07390313 | 0.0738797 | 4.89544569 | 0.03738316 | 0.84668703 | 0.97744262 |
| 0.05695665 | 0.06731943 | 6.39221448 | 0.03734624 | 0.84676181 | 0.97744262 |
| -0.0688043 | -0.0708728 | 3.51509903 | 0.03707903 | 0.84730424 | 0.97744262 |
| -0.0986629 | -0.0938609 | 2.56633668 | 0.03703191 | 0.84740009 | 0.97744262 |
| 0.05624344 | 0.06980423 | 4.09937636 | 0.03698865 | 0.84748816 | 0.97744262 |
| 0.08034498 | 0.06876584 | 4.46409064 | 0.03686145 | 0.8477474 | 0.97744262 |
| 0.05025907 | 0.06450951 | 4.81900638 | 0.03634693 | 0.84880082 | 0.97826745 |
| 0.07676867 | 0.06421871 | 4.16087546 | 0.03609709 | 0.84931514 | 0.9783667 |
| 0.07031144 | 0.07407359 | 3.5041056 | 0.03597697 | 0.84956307 | 0.9783667 |
| 0.07240478 | 0.06391858 | 4.50573111 | 0.0352328 | 0.85110868 | 0.97974081 |
| -0.0675619 | -0.0698102 | 3.79257785 | 0.0350775 | 0.85143336 | 0.97974081 |
| 0.07911055 | 0.07402881 | 3.88166782 | 0.0348801 | 0.85184712 | 0.97982733 |
| 0.05546703 | 0.06341672 | 6.85158604 | 0.03467093 | 0.85228688 | 0.97994368 |
| 0.0618218 | 0.07087339 | 4.27415543 | 0.03439527 | 0.85286856 | 0.97998617 |
| 0.07322072 | 0.06296877 | 5.9650156 | 0.03430867 | 0.85305177 | 0.97998617 |
| -0.0759358 | -0.0759873 | 3.30883714 | 0.03417281 | 0.85333972 | 0.97998617 |
| -0.016379 | 0.09068612 | 2.95012212 | 0.03358372 | 0.85459511 | 0.98103857 |
| -0.0628068 | -0.0782657 | 2.69349997 | 0.0328908 | 0.85608642 | 0.98159361 |
| -0.0478048 | -0.0911572 | 3.62304919 | 0.03273019 | 0.85643439 | 0.98159361 |
| 0.08509072 | 0.07932078 | 2.67715326 | 0.03267013 | 0.85656474 | 0.98159361 |
| -0.0649611 | -0.0658202 | 4.00497359 | 0.03258664 | 0.85674617 | 0.98159361 |
| 0.05780781 | 0.06186769 | 4.80405548 | 0.03251295 | 0.85690647 | 0.98159361 |
| 0.06005678 | 0.0925798 | 3.03419367 | 0.03213675 | 0.85772785 | 0.98159361 |
| 0.07072479 | 0.06998868 | 4.9107438 | 0.03211973 | 0.85776513 | 0.98159361 |

| | | | | | |
|---|---|---|---|---|---|
| -0.0523871 | -0.0610851 | 4.28255232 | 0.03210744 | 0.85779207 | 0.98159361 |
| -0.0529361 | -0.0642266 | 3.71247642 | 0.03171714 | 0.85864989 | 0.98218688 |
| 0.04046207 | 0.06368364 | 4.69420008 | 0.03143084 | 0.8592826 | 0.98252226 |
| -0.0656224 | -0.0605603 | 6.73034728 | 0.03121538 | 0.85976074 | 0.98268072 |
| -0.1299466 | -0.0753261 | 3.487439 | 0.03067306 | 0.86097177 | 0.98281314 |
| -0.0641792 | -0.064411 | 4.28042014 | 0.03052079 | 0.8613138 | 0.98281314 |
| -0.0496842 | -0.0596554 | 4.59472236 | 0.0304149 | 0.86155216 | 0.98281314 |
| -0.0472402 | -0.0581119 | 5.16166002 | 0.03031271 | 0.86178258 | 0.98281314 |
| 0.07906102 | 0.06904958 | 3.25444663 | 0.03020317 | 0.86203003 | 0.98281314 |
| 0.08904493 | 0.07054707 | 5.12894192 | 0.02986976 | 0.86278607 | 0.98281314 |
| 0.0595074 | 0.06139613 | 4.70679455 | 0.02983968 | 0.86285451 | 0.98281314 |
| -0.0613809 | -0.0588052 | 4.95763856 | 0.02982585 | 0.86288596 | 0.98281314 |
| 0.06827535 | 0.05645781 | 6.19172492 | 0.02980517 | 0.86293303 | 0.98281314 |
| -0.0694611 | -0.0681148 | 3.26796151 | 0.02960351 | 0.86339295 | 0.98295011 |
| -0.0549992 | -0.0603782 | 4.00745671 | 0.02904893 | 0.86466611 | 0.98381483 |
| 0.09239833 | 0.07440072 | 2.83053567 | 0.0289769 | 0.86483239 | 0.98381483 |
| -0.0606265 | -0.0612021 | 4.43108037 | 0.02857289 | 0.86576897 | 0.98449329 |
| 0.06685702 | 0.05761891 | 3.74508026 | 0.02800001 | 0.8671088 | 0.98476744 |
| -0.0781075 | -0.068522 | 3.56489963 | 0.02735227 | 0.86864079 | 0.98476744 |
| -0.0686248 | -0.0596335 | 7.09977605 | 0.02724444 | 0.86889762 | 0.98476744 |
| 0.06165383 | 0.0570936 | 6.70879431 | 0.0271971 | 0.86901054 | 0.98476744 |
| -0.0430373 | -0.0585199 | 4.33552181 | 0.02716981 | 0.86907568 | 0.98476744 |
| -0.0755352 | -0.0669391 | 4.23867874 | 0.02715738 | 0.86910537 | 0.98476744 |
| 0.06889094 | 0.05915899 | 4.31009748 | 0.02692431 | 0.86966322 | 0.98476744 |
| -0.074815 | -0.0622648 | 5.31509567 | 0.02680505 | 0.86994961 | 0.98476744 |
| 0.02584754 | -0.0830432 | 3.25701656 | 0.02678046 | 0.87000877 | 0.98476744 |
| 0.06465364 | 0.05524879 | 5.64961272 | 0.02675297 | 0.87007491 | 0.98476744 |
| -0.0630678 | -0.0650301 | 8.85358392 | 0.02667875 | 0.87025365 | 0.98476744 |
| -0.0706117 | -0.0724712 | 2.98278258 | 0.02661343 | 0.87041118 | 0.98476744 |
| 0.0937016 | 0.08841413 | 2.29681728 | 0.02654739 | 0.87057063 | 0.98476744 |
| -0.0381279 | -0.0656754 | 2.82875688 | 0.0262762 | 0.87122761 | 0.98476744 |
| 0.07863382 | 0.05755228 | 3.37093047 | 0.0261732 | 0.87147805 | 0.98476744 |
| 0.06278912 | 0.06594612 | 3.61031554 | 0.02605368 | 0.87176929 | 0.98476744 |
| -0.0692668 | -0.0563284 | 5.63091483 | 0.02602528 | 0.87183859 | 0.98476744 |
| 0.06474034 | 0.06437795 | 3.54103805 | 0.02581251 | 0.87235905 | 0.98476744 |
| 0.0430938 | 0.05917278 | 3.61675562 | 0.0254788 | 0.87317979 | 0.98476744 |
| -0.068785 | -0.061016 | 3.55823496 | 0.02525934 | 0.87372254 | 0.98476744 |
| -0.0510578 | -0.0584285 | 3.36542996 | 0.02518205 | 0.87391428 | 0.98476744 |
| -0.0452366 | -0.0571313 | 3.60345015 | 0.02514519 | 0.8740058 | 0.98476744 |
| 0.05704173 | 0.05499827 | 4.42524023 | 0.02511074 | 0.87409144 | 0.98476744 |
| 0.04613756 | 0.05438566 | 4.95819145 | 0.02503184 | 0.87428775 | 0.98476744 |
| -0.0525066 | -0.0566341 | 4.18690745 | 0.02490419 | 0.87460604 | 0.98476744 |
| -0.0889899 | -0.0709854 | 2.81200466 | 0.02470909 | 0.87509413 | 0.98476744 |
| -0.0614504 | -0.0592562 | 3.64059391 | 0.02466783 | 0.8751976 | 0.98476744 |
| 0.068376 | 0.05800646 | 3.3941996 | 0.02431408 | 0.87608841 | 0.98538666 |
| 0.09254847 | 0.06228616 | 3.09954232 | 0.02386152 | 0.8772378 | 0.98600317 |

| | | | | | |
|---|---|---|---|---|---|
| -0.1158077 | -0.1163847 | 13.6582759 | 0.02354734 | 0.87804229 | 0.98600317 |
| 0.06388038 | 0.05486527 | 3.85655557 | 0.02320697 | 0.87892009 | 0.98600317 |
| -0.0666032 | -0.0642644 | 7.88469129 | 0.02310682 | 0.87917962 | 0.98600317 |
| -0.0420447 | -0.0643951 | 2.79821112 | 0.02309912 | 0.8791996 | 0.98600317 |
| 0.0762267 | 0.05932154 | 3.34207868 | 0.02285996 | 0.8798218 | 0.98600317 |
| -0.0435462 | -0.0541803 | 3.68673886 | 0.02284529 | 0.87986009 | 0.98600317 |
| 0.04862224 | 0.05628433 | 4.62310523 | 0.02281306 | 0.87994422 | 0.98600317 |
| -0.0430783 | -0.0510185 | 6.77803666 | 0.02227371 | 0.88136127 | 0.98600317 |
| -0.0376196 | -0.051325 | 3.69661231 | 0.02214513 | 0.88170168 | 0.98600317 |
| -0.0621625 | -0.0507762 | 6.41468228 | 0.02211461 | 0.88178261 | 0.98600317 |
| 0.04549348 | 0.0487111 | 5.48728887 | 0.02210685 | 0.88180322 | 0.98600317 |
| -0.0546001 | -0.0599624 | 2.98750962 | 0.02209904 | 0.88182393 | 0.98600317 |
| -0.0563883 | -0.058059 | 3.25828132 | 0.02174492 | 0.8827677 | 0.98600317 |
| 0.25879351 | 0.12215458 | 5.99264868 | 0.02172904 | 0.88281021 | 0.98600317 |
| 0.07362196 | 0.05619642 | 3.43607126 | 0.02168792 | 0.88292035 | 0.98600317 |
| -0.0475368 | -0.0527205 | 6.90705493 | 0.02163372 | 0.88306566 | 0.98600317 |
| 0.06858 | 0.06825538 | 9.82905141 | 0.02134519 | 0.8838425 | 0.98600317 |
| 0.05641964 | 0.05651514 | 3.84116151 | 0.02119664 | 0.88424454 | 0.98600317 |
| 0.08785054 | 0.05914663 | 2.91880012 | 0.0211926 | 0.88425549 | 0.98600317 |
| 0.05016561 | 0.0531403 | 5.75696465 | 0.02113754 | 0.8844049 | 0.98600317 |
| 0.06887886 | 0.05116014 | 4.79007636 | 0.02104693 | 0.88465119 | 0.98600317 |
| 0.05429952 | 0.04617857 | 5.83043812 | 0.0209154 | 0.88500967 | 0.98600317 |
| -0.0321737 | -0.055346 | 3.30082024 | 0.02083644 | 0.88522542 | 0.98600317 |
| 0.05016963 | 0.06239697 | 3.32632668 | 0.02061446 | 0.88583421 | 0.98600317 |
| 0.04205062 | 0.05566876 | 3.780015 | 0.02055392 | 0.88600084 | 0.98600317 |
| -0.0488617 | -0.0512229 | 7.31980338 | 0.02032195 | 0.88664157 | 0.98600317 |
| 0.05497143 | 0.04916599 | 3.85108263 | 0.02026055 | 0.88681181 | 0.98600317 |
| -0.052326 | -0.0581167 | 4.29696163 | 0.02022248 | 0.88691748 | 0.98600317 |
| 0.04722282 | 0.06790035 | 2.76932195 | 0.02017702 | 0.8870438 | 0.98600317 |
| -0.0491868 | -0.0560862 | 3.13612889 | 0.02006734 | 0.88734919 | 0.98600317 |
| -0.0704246 | -0.0566764 | 3.54309044 | 0.01993961 | 0.88770587 | 0.98600317 |
| -0.1126254 | -0.0555662 | 4.1651527 | 0.01987747 | 0.88787984 | 0.98600317 |
| 0.05803167 | 0.05163587 | 7.60480547 | 0.01947319 | 0.8890184 | 0.98620277 |
| 0.04378361 | 0.05103509 | 3.51718365 | 0.01931222 | 0.8894751 | 0.98620277 |
| -0.0559993 | -0.0514666 | 5.43980709 | 0.01914038 | 0.88996479 | 0.98620277 |
| -0.0731358 | -0.0486384 | 3.64160827 | 0.01892037 | 0.89059502 | 0.98620277 |
| -0.0732804 | -0.0520165 | 4.3359772 | 0.01866245 | 0.89133862 | 0.98620277 |
| -0.049621 | -0.0487074 | 3.9525622 | 0.01852974 | 0.89172325 | 0.98620277 |
| -0.049088 | -0.0521461 | 8.86102116 | 0.01852628 | 0.89173332 | 0.98620277 |
| -0.0734894 | -0.049232 | 4.47462309 | 0.01838434 | 0.89214631 | 0.98620277 |
| -0.040377 | -0.0474083 | 6.2026177 | 0.01838042 | 0.89215772 | 0.98620277 |
| 0.09573518 | 0.05729258 | 3.64420775 | 0.01836205 | 0.89221132 | 0.98620277 |
| 0.05196622 | 0.05092988 | 8.81892869 | 0.01833527 | 0.89228947 | 0.98620277 |
| 0.0774272 | -0.0676935 | 3.57288106 | 0.01808309 | 0.89302826 | 0.98620277 |
| -0.0723126 | -0.054537 | 3.45911757 | 0.01799139 | 0.89329821 | 0.98620277 |
| -0.0270355 | -0.0526003 | 3.06427826 | 0.01783085 | 0.89377249 | 0.98620277 |

| | | | | | |
|---|---|---|---|---|---|
| -0.0617846 | -0.0500336 | 3.66685574 | 0.01757712 | 0.89452654 | 0.98620277 |
| 0.03804423 | 0.05690082 | 3.13696966 | 0.01756673 | 0.89455756 | 0.98620277 |
| 0.03709302 | 0.04711315 | 4.41883491 | 0.01754713 | 0.89461605 | 0.98620277 |
| 0.11234674 | 0.07434368 | 2.80975297 | 0.0174206 | 0.89499448 | 0.98620277 |
| 0.06603093 | 0.05111231 | 3.17163713 | 0.01703723 | 0.89614971 | 0.98620277 |
| -0.0602878 | -0.061975 | 3.10738879 | 0.01700603 | 0.89624429 | 0.98620277 |
| 0.04058299 | 0.0465086 | 5.48120927 | 0.01700367 | 0.89625146 | 0.98620277 |
| 0.19644096 | 0.08028161 | 4.93124588 | 0.01686955 | 0.89665913 | 0.98620277 |
| 0.23010807 | -0.1722358 | 4.75117553 | 0.01686018 | 0.89668766 | 0.98620277 |
| 0.08636178 | 0.08629005 | 15.3686331 | 0.01674875 | 0.89702774 | 0.98620277 |
| 0.05091046 | 0.0484835 | 6.0217694 | 0.01641092 | 0.89806579 | 0.98620277 |
| 0.05206524 | 0.04470856 | 3.51980011 | 0.01633735 | 0.89829328 | 0.98620277 |
| 0.08636785 | 0.05396837 | 2.88914349 | 0.01627167 | 0.89849683 | 0.98620277 |
| -0.0400857 | -0.0446202 | 4.08094246 | 0.01623325 | 0.89861609 | 0.98620277 |
| 0.07552213 | 0.0522062 | 2.87417869 | 0.01605315 | 0.89917704 | 0.98620277 |
| 0.37419938 | 0.06712363 | 3.54497566 | 0.01596138 | 0.89946409 | 0.98620277 |
| -0.0505326 | -0.0489521 | 3.42194621 | 0.01576485 | 0.90008168 | 0.98620277 |
| 0.04963419 | 0.05073243 | 3.03396702 | 0.01574386 | 0.90014788 | 0.98620277 |
| -0.0459183 | -0.0442207 | 6.49976464 | 0.01573675 | 0.90017031 | 0.98620277 |
| -0.0641813 | -0.0489994 | 3.73678611 | 0.01571741 | 0.90023136 | 0.98620277 |
| -0.0394356 | -0.046298 | 3.78347033 | 0.01560006 | 0.90060256 | 0.98620277 |
| 0.03729682 | 0.05437184 | 3.27098283 | 0.01558416 | 0.90065296 | 0.98620277 |
| 0.05102379 | 0.04048754 | 6.0399163 | 0.01534295 | 0.90142086 | 0.98620277 |
| -0.0598246 | -0.0431362 | 4.2617625 | 0.01531014 | 0.90152578 | 0.98620277 |
| 0.04027718 | 0.04681496 | 3.42684947 | 0.01530607 | 0.9015388 | 0.98620277 |
| -0.0762808 | -0.0448322 | 3.48235845 | 0.01525868 | 0.90169058 | 0.98620277 |
| -0.0628129 | -0.0508187 | 2.80769901 | 0.01492583 | 0.90276335 | 0.98625917 |
| 0.00793167 | 0.04285316 | 4.00032948 | 0.01484631 | 0.90302143 | 0.98625917 |
| -0.0146284 | -0.0410433 | 4.52725892 | 0.0147416 | 0.90336235 | 0.98625917 |
| 0.05115917 | 0.04225125 | 4.48248892 | 0.01465196 | 0.90365517 | 0.98625917 |
| -0.0482296 | -0.0482132 | 3.4039308 | 0.01462628 | 0.90373922 | 0.98625917 |
| -0.0401512 | -0.0395898 | 6.31519789 | 0.01436348 | 0.90460378 | 0.98625917 |
| 0.09262606 | 0.05088329 | 3.70469126 | 0.01427929 | 0.90488241 | 0.98625917 |
| 0.03630563 | 0.05769306 | 3.1088253 | 0.0142502 | 0.90497889 | 0.98625917 |
| -0.0479656 | -0.0466416 | 3.74028694 | 0.01422506 | 0.90506237 | 0.98625917 |
| -0.0584733 | -0.0414034 | 5.34956676 | 0.01419866 | 0.90515009 | 0.98625917 |
| -0.0697825 | -0.0449802 | 3.81904735 | 0.01389881 | 0.90615227 | 0.98637502 |
| -0.0455433 | -0.0516415 | 2.80389281 | 0.01388579 | 0.90619603 | 0.98637502 |
| -0.0529051 | -0.0536856 | 2.74312219 | 0.01386115 | 0.90627892 | 0.98637502 |
| 0.0662508 | 0.04370584 | 3.28538977 | 0.01373938 | 0.9066896 | 0.98645102 |
| -0.0562034 | -0.0530731 | 2.71726018 | 0.01363864 | 0.90703075 | 0.98645133 |
| -0.0423419 | -0.0423747 | 7.96419201 | 0.01326778 | 0.90829783 | 0.98696286 |
| -0.0335812 | -0.0414198 | 3.82037165 | 0.01326122 | 0.90832039 | 0.98696286 |
| 0.03065452 | 0.04389328 | 3.70452593 | 0.0131838 | 0.90858723 | 0.98696286 |
| -0.0190195 | -0.0393662 | 3.93085223 | 0.01309989 | 0.9088773 | 0.98696286 |
| 0.03201897 | 0.04197912 | 3.27671411 | 0.0128949 | 0.90959001 | 0.98696286 |

| | | | | | |
|---|---|---|---|---|---|
| 0.03605442 | 0.04256981 | 4.16291887 | 0.01281096 | 0.90988348 | 0.98696286 |
| 0.12588714 | 0.0587237 | 5.98351491 | 0.01276269 | 0.91005268 | 0.98696286 |
| -0.0402497 | -0.0397076 | 5.99114735 | 0.01233062 | 0.911582 | 0.98696286 |
| -0.0245308 | -0.0370241 | 4.85726741 | 0.01231068 | 0.91165322 | 0.98696286 |
| 0.03576241 | 0.0426981 | 4.63857257 | 0.01227236 | 0.91179027 | 0.98696286 |
| -0.0107527 | -0.046604 | 4.09630052 | 0.01220965 | 0.912015 | 0.98696286 |
| -0.0436195 | -0.0414935 | 4.72266298 | 0.01220778 | 0.91202171 | 0.98696286 |
| -0.0442919 | -0.0392032 | 4.53431682 | 0.01218568 | 0.91210105 | 0.98696286 |
| -0.0505223 | -0.0441539 | 3.28285042 | 0.01191569 | 0.91307636 | 0.98696286 |
| 0.0581843 | 0.04769901 | 3.02972972 | 0.01188034 | 0.91320489 | 0.98696286 |
| -0.0751467 | -0.0463216 | 2.91711671 | 0.01178284 | 0.91356036 | 0.98696286 |
| -0.0254017 | -0.0364118 | 5.66258445 | 0.01169194 | 0.91389313 | 0.98696286 |
| 0.0557527 | 0.04133147 | 3.48309436 | 0.01164933 | 0.9140496 | 0.98696286 |
| 0.0422287 | 0.04201898 | 9.21377897 | 0.01150471 | 0.91458272 | 0.98696286 |
| -0.0517883 | -0.0376343 | 4.45808986 | 0.01143788 | 0.91483023 | 0.98696286 |
| 0.03837851 | 0.03610901 | 7.07665832 | 0.01142766 | 0.91486812 | 0.98696286 |
| 0.04838862 | 0.03452936 | 6.19485647 | 0.01137058 | 0.9150802 | 0.98696286 |
| -0.0160657 | -0.0412484 | 2.90343796 | 0.01063262 | 0.91787204 | 0.98696286 |
| -0.0339961 | -0.0363025 | 4.80938238 | 0.01058568 | 0.91805287 | 0.98696286 |
| 0.03638046 | 0.03339303 | 5.56879014 | 0.01051866 | 0.91831179 | 0.98696286 |
| -0.0348378 | -0.0333232 | 6.23265402 | 0.0104166 | 0.91870767 | 0.98696286 |
| 0.03191874 | 0.042726 | 3.1759535 | 0.01037753 | 0.91885973 | 0.98696286 |
| -0.1119381 | 0.05372721 | 3.38715781 | 0.01033119 | 0.91904047 | 0.98696286 |
| 0.04878693 | 0.04070399 | 4.63420266 | 0.01028885 | 0.91920597 | 0.98696286 |
| 0.02678921 | 0.03542762 | 4.42183863 | 0.0102487 | 0.91936322 | 0.98696286 |
| -0.0422147 | -0.0420394 | 3.01698547 | 0.01023097 | 0.91943277 | 0.98696286 |
| -0.1137678 | 0.05007624 | 3.87166605 | 0.01022999 | 0.91943662 | 0.98696286 |
| 0.04489565 | 0.03516355 | 4.48865697 | 0.0102115 | 0.9195092 | 0.98696286 |
| 0.13807816 | -0.0516366 | 3.62167062 | 0.0100089 | 0.92030901 | 0.98696286 |
| -0.0236109 | -0.0332583 | 3.77286083 | 0.00998083 | 0.92042045 | 0.98696286 |
| 0.01703455 | 0.03457133 | 3.80705994 | 0.00991574 | 0.92067953 | 0.98696286 |
| 0.03792899 | 0.03400232 | 3.87365582 | 0.00986284 | 0.92089067 | 0.98696286 |
| 0.03803366 | 0.03719772 | 4.91821889 | 0.00982931 | 0.92102484 | 0.98696286 |
| 0.04100205 | 0.03650486 | 3.73953183 | 0.00980688 | 0.92111471 | 0.98696286 |
| 0.07312157 | 0.04735277 | 3.08775171 | 0.00960811 | 0.92191563 | 0.98696286 |
| -0.0371185 | -0.0369321 | 8.7774821 | 0.00952311 | 0.9222607 | 0.98696286 |
| -0.0088723 | -0.0342019 | 3.6124928 | 0.00940588 | 0.92273915 | 0.98696286 |
| -0.0302202 | -0.0346226 | 5.0800218 | 0.00920964 | 0.92354691 | 0.98696286 |
| -0.0052476 | 0.04311751 | 3.51714902 | 0.00916728 | 0.92372238 | 0.98696286 |
| -0.0361233 | -0.0397945 | 3.08158257 | 0.00913461 | 0.92385799 | 0.98696286 |
| 0.02772274 | 0.0351626 | 3.96088297 | 0.00908699 | 0.92405613 | 0.98696286 |
| -0.0349362 | -0.0371114 | 4.85130167 | 0.0089089 | 0.92480175 | 0.98696286 |
| 0.04612128 | 0.0337034 | 4.04725421 | 0.00882834 | 0.92514151 | 0.98696286 |
| 0.01456706 | 0.03306467 | 4.00771581 | 0.00882225 | 0.92516727 | 0.98696286 |
| 0.05794255 | 0.04092397 | 2.85025688 | 0.008584 | 0.92618174 | 0.98696286 |
| -0.0382418 | -0.0343306 | 3.86374907 | 0.00851949 | 0.92645884 | 0.98696286 |

| | | | | | |
|---|---|---|---|---|---|
| 0.04510677 | 0.04505924 | 2.60412625 | 0.00843024 | 0.92684394 | 0.98696286 |
| -0.0137452 | -0.033644 | 3.08888523 | 0.00835103 | 0.9271875 | 0.98696286 |
| 0.08222927 | 0.03744633 | 2.88231854 | 0.00820345 | 0.92783197 | 0.98696286 |
| 0.04908684 | 0.0310132 | 4.06034503 | 0.00809298 | 0.92831822 | 0.98696286 |
| -0.0256204 | -0.0374287 | 3.00295339 | 0.00808721 | 0.92834369 | 0.98696286 |
| 0.02544927 | 0.03141263 | 3.99381102 | 0.00802592 | 0.92861502 | 0.98696286 |
| 0.0472291 | 0.03144232 | 4.24818973 | 0.00802359 | 0.92862534 | 0.98696286 |
| -0.032486 | -0.0303993 | 4.7042558 | 0.00792324 | 0.92907189 | 0.98696286 |
| -0.0422694 | -0.0301156 | 5.05724628 | 0.00788129 | 0.92925942 | 0.98696286 |
| -0.0299085 | -0.0384885 | 2.78402937 | 0.00762328 | 0.93042399 | 0.98696286 |
| -0.0473118 | -0.0320452 | 4.6396703 | 0.0075439 | 0.93078625 | 0.98696286 |
| -0.042389 | -0.0317001 | 5.38335748 | 0.00753585 | 0.93082313 | 0.98696286 |
| -0.023325 | -0.0301626 | 6.953701 | 0.00747349 | 0.93110922 | 0.98696286 |
| 0.0802795 | 0.03540362 | 2.88598289 | 0.0074295 | 0.93131177 | 0.98696286 |
| -0.0142416 | -0.0282705 | 5.29242969 | 0.00721205 | 0.93232198 | 0.98696286 |
| 0.04816549 | 0.0377959 | 2.61808713 | 0.00719366 | 0.93240813 | 0.98696286 |
| -0.0321748 | -0.031662 | 6.3682298 | 0.00718009 | 0.93247176 | 0.98696286 |
| -0.0205898 | -0.0314308 | 3.42707493 | 0.00717576 | 0.93249204 | 0.98696286 |
| 0.03198718 | 0.02924152 | 5.16054721 | 0.0071371 | 0.93267373 | 0.98696286 |
| -0.1352631 | -0.0412406 | 5.01835459 | 0.00713223 | 0.93269662 | 0.98696286 |
| -0.0337536 | -0.0323256 | 3.25672944 | 0.00712972 | 0.93270848 | 0.98696286 |
| 0.03783745 | 0.03057189 | 4.21298388 | 0.00708979 | 0.9328967 | 0.98696286 |
| -0.0232172 | -0.0282411 | 4.36667284 | 0.00701948 | 0.93322949 | 0.98696286 |
| 0.02300802 | 0.02913969 | 4.31922963 | 0.00700393 | 0.93330331 | 0.98696286 |
| -0.0308732 | -0.0296847 | 4.3097818 | 0.0069794 | 0.93341996 | 0.98696286 |
| -0.0559036 | 0.04283927 | 2.85649254 | 0.00677442 | 0.93440271 | 0.98707925 |
| -0.0329699 | -0.0342623 | 3.27577537 | 0.00669063 | 0.9348087 | 0.98707925 |
| 0.00430695 | 0.04127421 | 2.80202598 | 0.0066383 | 0.93506361 | 0.98707925 |
| -0.0191549 | -0.030799 | 3.30041502 | 0.00651514 | 0.93566747 | 0.98707925 |
| -0.0207815 | -0.0277125 | 6.38687495 | 0.00636071 | 0.93643284 | 0.98707925 |
| 0.03780995 | 0.02625413 | 5.50423015 | 0.00631557 | 0.93665834 | 0.98707925 |
| 0.04981757 | 0.03176321 | 4.25948372 | 0.0062988 | 0.93674234 | 0.98707925 |
| 0.03459024 | 0.02903319 | 4.83390412 | 0.00625487 | 0.93696283 | 0.98707925 |
| 0.03175788 | 0.03170914 | 3.60197528 | 0.00624865 | 0.93699413 | 0.98707925 |
| 0.01191918 | 0.04004808 | 2.63859496 | 0.00624592 | 0.93700788 | 0.98707925 |
| -0.018149 | -0.028694 | 3.19628262 | 0.00619158 | 0.93728189 | 0.98707925 |
| 0.02765229 | 0.0274383 | 4.28576391 | 0.00606547 | 0.93792263 | 0.98732978 |
| 0.02386535 | 0.03123645 | 3.50111051 | 0.00592323 | 0.93865337 | 0.98732978 |
| 0.03356847 | 0.02700697 | 5.27369714 | 0.00581598 | 0.9392102 | 0.98732978 |
| 0.03201458 | 0.02681504 | 6.91984406 | 0.0057572 | 0.93951756 | 0.98732978 |
| 0.03526408 | 0.0321524 | 3.23923266 | 0.0057488 | 0.93956165 | 0.98732978 |
| 0.03889334 | 0.02713996 | 4.59351701 | 0.00574782 | 0.93956676 | 0.98732978 |
| -0.0297197 | -0.0299953 | 3.71148328 | 0.00567839 | 0.93993219 | 0.98735527 |
| 0.04926087 | 0.02596834 | 3.51413432 | 0.00542918 | 0.94126263 | 0.98820222 |
| -0.037462 | -0.0254629 | 4.12820723 | 0.00532185 | 0.94184509 | 0.98820222 |
| -0.0271961 | -0.0305553 | 3.90410796 | 0.00532084 | 0.9418506 | 0.98820222 |

| | | | | | |
|---|---|---|---|---|---|
| 0.03294489 | 0.02644083 | 3.73737973 | 0.00527443 | 0.94210433 | 0.98820222 |
| 0.01270489 | 0.02491717 | 4.5325248 | 0.00509195 | 0.94311293 | 0.9884381 |
| -0.0270589 | -0.0265594 | 4.60962338 | 0.00499859 | 0.94363598 | 0.9884381 |
| 0.03377975 | 0.03668522 | 2.72760352 | 0.00493371 | 0.94400237 | 0.9884381 |
| -0.0343528 | -0.0276324 | 3.26883646 | 0.00488954 | 0.94425316 | 0.9884381 |
| -0.023617 | -0.0263729 | 3.49556541 | 0.00487145 | 0.94435619 | 0.9884381 |
| -0.0268772 | -0.0272934 | 3.39744736 | 0.00485387 | 0.94445654 | 0.9884381 |
| 0.02563331 | 0.0266543 | 4.26703287 | 0.00480785 | 0.94472004 | 0.9884381 |
| 0.00602636 | -0.0245614 | 3.34481119 | 0.00470217 | 0.94533002 | 0.9885404 |
| 0.03598574 | 0.03548471 | 2.33220813 | 0.00467276 | 0.94550098 | 0.9885404 |
| -0.024601 | -0.0307096 | 2.45586036 | 0.00439046 | 0.94717041 | 0.98871525 |
| 0.01961574 | 0.02401617 | 3.6269661 | 0.00435486 | 0.94738468 | 0.98871525 |
| -0.0227272 | -0.0224313 | 7.07200494 | 0.00431442 | 0.9476292 | 0.98871525 |
| -0.0219037 | -0.022431 | 6.38830016 | 0.00421495 | 0.94823558 | 0.98871525 |
| 0.02178058 | 0.02208538 | 6.98553542 | 0.00418838 | 0.94839877 | 0.98871525 |
| 0.02298026 | 0.02432429 | 4.37745582 | 0.00417842 | 0.94846009 | 0.98871525 |
| -0.0264128 | -0.0218068 | 7.23328252 | 0.00411298 | 0.94886473 | 0.98871525 |
| 0.02865456 | 0.022566 | 7.63944818 | 0.00408414 | 0.94904408 | 0.98871525 |
| 0.04115137 | 0.02194233 | 4.49139891 | 0.00408345 | 0.94904836 | 0.98871525 |
| 0.03652482 | 0.02230354 | 3.46473893 | 0.00407763 | 0.94908464 | 0.98871525 |
| -0.0363306 | -0.023255 | 4.5712189 | 0.0039993 | 0.9495754 | 0.98874106 |
| 0.00751591 | 0.02779572 | 2.85908292 | 0.00396485 | 0.94979273 | 0.98874106 |
| -0.0622757 | -0.0279615 | 2.96172421 | 0.003791 | 0.95090441 | 0.9891328 |
| -5.98E-04 | -0.0306948 | 3.0541442 | 0.00372944 | 0.95130413 | 0.9891328 |
| -0.0243311 | -0.0242289 | 3.1640609 | 0.00371683 | 0.95138645 | 0.9891328 |
| -0.024643 | -0.0234866 | 3.94393633 | 0.00369394 | 0.95153619 | 0.9891328 |
| 0.03086377 | 0.02948474 | 2.63941987 | 0.00359078 | 0.95221687 | 0.98948496 |
| -0.0079506 | -0.0197155 | 4.01185039 | 0.0035209 | 0.95268355 | 0.98961457 |
| 0.01779548 | -0.0207415 | 3.26555128 | 0.00327407 | 0.95437037 | 0.99076278 |
| 0.02245424 | 0.0222097 | 9.47411486 | 0.00306748 | 0.95583189 | 0.99076278 |
| 0.03361594 | 0.01924555 | 4.28983583 | 0.00303757 | 0.95604752 | 0.99076278 |
| -0.0379705 | -0.0200678 | 4.56526107 | 0.00302633 | 0.95612883 | 0.99076278 |
| 0.00548419 | -0.0222372 | 2.8516661 | 0.00302628 | 0.95612917 | 0.99076278 |
| -0.0089863 | -0.018151 | 5.6178353 | 0.00300742 | 0.956266 | 0.99076278 |
| -0.0108437 | -0.0181583 | 5.42990435 | 0.00299849 | 0.95633089 | 0.99076278 |
| 0.04124625 | 0.02203805 | 3.49911522 | 0.00289106 | 0.95711953 | 0.99076278 |
| 0.03975827 | 0.01834016 | 4.36073893 | 0.00288968 | 0.95712975 | 0.99076278 |
| 0.0165702 | 0.01868724 | 4.83773712 | 0.00287854 | 0.95721241 | 0.99076278 |
| 0.04215199 | 0.02037629 | 3.31154071 | 0.00277546 | 0.95798477 | 0.9912077 |
| -0.0242911 | -0.019008 | 4.54676299 | 0.00253657 | 0.95983202 | 0.99253028 |
| -0.0293433 | -0.0214551 | 2.68956223 | 0.00249755 | 0.96014193 | 0.99253028 |
| 0.0539074 | 0.02014248 | 2.93990368 | 0.00247877 | 0.96029191 | 0.99253028 |
| -0.0209755 | -0.0162651 | 4.08364728 | 0.00233556 | 0.96145511 | 0.99262865 |
| 0.0232515 | 0.01732891 | 4.1192337 | 0.00229809 | 0.96176532 | 0.99262865 |
| 0.00270242 | -0.0189442 | 2.90156517 | 0.002274 | 0.96196609 | 0.99262865 |
| -0.0162337 | -0.0160346 | 4.1533218 | 0.00225097 | 0.96215907 | 0.99262865 |

| | | | | | |
|---|---|---|---|---|---|
| 0.01446938 | 0.01744572 | 7.49800061 | 0.002246 | 0.96220081 | 0.99262865 |
| 0.01691886 | 0.01588712 | 4.3796256 | 0.00221705 | 0.96244506 | 0.99262865 |
| -0.0112288 | -0.0152199 | 6.34356304 | 0.0020745 | 0.96367155 | 0.99338946 |
| 0.03086456 | 0.01913821 | 3.16330071 | 0.00202573 | 0.96410085 | 0.99338946 |
| 0.00485084 | 0.01654022 | 3.85603728 | 0.00201314 | 0.96421251 | 0.99338946 |
| -0.0100223 | -0.0151466 | 4.82789996 | 0.00195781 | 0.96470739 | 0.99354561 |
| 0.77352681 | -0.0414133 | 2.94310207 | 0.00184155 | 0.96577066 | 0.99377639 |
| -0.0071693 | -0.0143137 | 6.29689094 | 0.0018181 | 0.96598914 | 0.99377639 |
| -0.0187998 | -0.0159144 | 4.60240744 | 0.00178122 | 0.96633565 | 0.99377639 |
| -0.0068354 | -0.0145389 | 3.8700739 | 0.00174411 | 0.96668798 | 0.99377639 |
| -0.0185317 | -0.0164585 | 4.55251407 | 0.00172333 | 0.96688687 | 0.99377639 |
| -0.0140219 | -0.0146865 | 7.75840711 | 0.00170646 | 0.96704932 | 0.99377639 |
| -0.0053418 | 0.02000137 | 2.94269325 | 0.00165338 | 0.9675655 | 0.99377639 |
| -0.0160555 | -0.0152503 | 3.47103489 | 0.0016369 | 0.96772754 | 0.99377639 |
| 0.04438927 | 0.01555087 | 3.18323407 | 0.00157445 | 0.96834882 | 0.99377639 |
| -4.03E-04 | -0.0219023 | 2.80363234 | 0.00156309 | 0.9684631 | 0.99377639 |
| 0.01007199 | 0.01376897 | 4.36731308 | 0.00153882 | 0.96870877 | 0.99377639 |
| 0.02860398 | 0.01360043 | 4.79902434 | 0.00139243 | 0.97023359 | 0.99458739 |
| 0.02395135 | 0.01272601 | 6.39125341 | 0.00136478 | 0.97053054 | 0.99458739 |
| 0.07370569 | 0.01572849 | 4.75769251 | 0.00132509 | 0.970962 | 0.99458739 |
| -0.0136851 | -0.0145757 | 9.81822709 | 0.00132205 | 0.97099528 | 0.99458739 |
| 0.01029094 | 0.01289061 | 4.55368968 | 0.00130185 | 0.97121768 | 0.99458739 |
| 0.02097116 | 0.01358818 | 3.41779539 | 0.00122125 | 0.97212253 | 0.99471523 |
| 0.04700947 | 0.01460925 | 2.83829486 | 0.00120756 | 0.97227911 | 0.99471523 |
| 0.01353187 | 0.01263921 | 3.74887353 | 0.00119933 | 0.97237366 | 0.99471523 |
| 0.02127268 | 0.01188659 | 5.15652867 | 0.00116368 | 0.97278718 | 0.99478661 |
| -0.0124843 | -0.012161 | 5.23954799 | 0.00110525 | 0.97347896 | 0.99488357 |
| -0.0262411 | -0.015642 | 2.95591732 | 0.00108576 | 0.97371376 | 0.99488357 |
| 0.0635149 | 0.01296974 | 3.10697533 | 0.00106933 | 0.97391332 | 0.99488357 |
| -0.0088201 | 0.01340308 | 4.12363538 | 0.00102204 | 0.9744965 | 0.99498193 |
| -0.0250501 | -0.0110181 | 5.11979989 | 0.00100535 | 0.97470547 | 0.99498193 |
| 0.00682743 | 0.01094929 | 3.48038686 | 8.09E-04 | 0.97730835 | 0.99498193 |
| -0.0085695 | -0.009866 | 4.89680258 | 8.02E-04 | 0.97741174 | 0.99498193 |
| -3.66E-05 | 0.01005216 | 3.30617556 | 7.34E-04 | 0.97838174 | 0.99498193 |
| 0.03037049 | 0.00928988 | 4.37840581 | 7.24E-04 | 0.97853519 | 0.99498193 |
| -0.0103167 | -0.0103713 | 4.23493582 | 7.24E-04 | 0.97853954 | 0.99498193 |
| 0.02751295 | 0.01152014 | 3.42353686 | 7.04E-04 | 0.97883467 | 0.99498193 |
| -0.0063293 | 0.01066789 | 2.74139371 | 6.91E-04 | 0.97903099 | 0.99498193 |
| 0.00291242 | -0.0089948 | 3.74740448 | 6.89E-04 | 0.97906627 | 0.99498193 |
| -0.0020385 | -0.0090421 | 4.85741521 | 6.88E-04 | 0.97907516 | 0.99498193 |
| 0.00289536 | -0.0091142 | 5.02763999 | 6.82E-04 | 0.97916705 | 0.99498193 |
| 0.05228465 | 0.01180628 | 6.25212324 | 6.75E-04 | 0.97927973 | 0.99498193 |
| 0.02630488 | 0.00991694 | 3.46246263 | 6.71E-04 | 0.97933578 | 0.99498193 |
| -0.0741246 | 0.01522481 | 3.73563132 | 6.67E-04 | 0.97939852 | 0.99498193 |
| -0.0045933 | 0.00892467 | 5.15085575 | 6.44E-04 | 0.97976186 | 0.99498193 |
| 0.00599432 | 0.00892298 | 6.92900689 | 6.33E-04 | 0.97992495 | 0.99498193 |

| | | | | | |
|---|---|---|---|---|---|
| -0.0042829 | -0.0082798 | 5.38815808 | 6.16E-04 | 0.98019817 | 0.99498193 |
| 0.01140301 | 0.0083887 | 5.19941135 | 5.73E-04 | 0.98090938 | 0.99535475 |
| -0.0075325 | 0.00806287 | 3.48279263 | 5.03E-04 | 0.98209966 | 0.99545839 |
| 0.00916368 | 0.00867244 | 3.8375374 | 5.03E-04 | 0.9821029 | 0.99545839 |
| 0.01329118 | 0.00774207 | 5.62637788 | 4.88E-04 | 0.98237007 | 0.99545839 |
| 0.00470057 | 0.0086792 | 3.75274115 | 4.62E-04 | 0.98285838 | 0.99545839 |
| -0.0070032 | -0.0078821 | 7.93558267 | 4.55E-04 | 0.9829793 | 0.99545839 |
| 0.0068084 | -0.0069415 | 5.34824589 | 4.50E-04 | 0.98307536 | 0.99545839 |
| -0.0040262 | -0.0072353 | 3.8734472 | 4.04E-04 | 0.98396205 | 0.99600776 |
| 0.00437225 | 0.00732661 | 4.11569185 | 3.67E-04 | 0.98470965 | 0.99641599 |
| -0.0035157 | 0.00637574 | 3.36396933 | 3.19E-04 | 0.98575598 | 0.99661596 |
| -0.0163036 | 0.00601973 | 4.98748464 | 3.08E-04 | 0.98599835 | 0.99661596 |
| 0.00665269 | -0.0056614 | 6.07266484 | 3.06E-04 | 0.98604155 | 0.99661596 |
| 0.01190453 | -0.00573 | 3.6044075 | 2.83E-04 | 0.98656665 | 0.99661596 |
| 8.81E-04 | 0.00667904 | 4.36994617 | 2.81E-04 | 0.98662914 | 0.99661596 |
| -0.0048256 | 0.00682174 | 4.07558102 | 2.40E-04 | 0.98764205 | 0.9966525 |
| 0.01071517 | -0.0045867 | 5.83465935 | 2.01E-04 | 0.98868697 | 0.9966525 |
| 0.00291054 | -0.004581 | 5.42058112 | 1.86E-04 | 0.98912153 | 0.9966525 |
| -0.0171684 | -0.0053161 | 3.16399827 | 1.86E-04 | 0.98912976 | 0.9966525 |
| 0.01018204 | 0.00451244 | 4.68865853 | 1.73E-04 | 0.98949959 | 0.9966525 |
| 0.00466794 | 0.00516053 | 4.54826864 | 1.72E-04 | 0.98953644 | 0.9966525 |
| -0.0104068 | -0.0042839 | 6.97937368 | 1.64E-04 | 0.98979251 | 0.9966525 |
| -0.0263231 | -0.00439 | 2.65054292 | 1.60E-04 | 0.98990669 | 0.9966525 |
| 0.02614067 | -0.0038767 | 3.23116402 | 1.49E-04 | 0.99025549 | 0.9966525 |
| -0.0241298 | -0.0063265 | 2.83192846 | 1.48E-04 | 0.9902963 | 0.9966525 |
| 0.01277207 | 0.00380084 | 6.52224787 | 1.33E-04 | 0.99079211 | 0.9966525 |
| -0.0148528 | 0.00362949 | 4.1865675 | 1.30E-04 | 0.99091615 | 0.9966525 |
| -0.0242787 | -0.003886 | 5.61338992 | 1.23E-04 | 0.99114232 | 0.9966525 |
| 0.00420331 | -0.0030645 | 5.14643563 | 7.95E-05 | 0.99288781 | 0.99775843 |
| 0.03975489 | 0.00323631 | 3.67273722 | 7.19E-05 | 0.99323669 | 0.99775843 |
| -0.0046679 | 0.00299458 | 3.35984607 | 7.10E-05 | 0.99327645 | 0.99775843 |
| -0.0104122 | 0.00252802 | 4.14314503 | 6.18E-05 | 0.99372727 | 0.99777408 |
| -0.0175715 | 0.00346705 | 4.10578791 | 5.65E-05 | 0.9940015 | 0.99777408 |
| 0.0021967 | 0.00259175 | 2.9723223 | 4.45E-05 | 0.99467568 | 0.99777408 |
| 0.00262882 | 0.00215253 | 3.222943 | 3.42E-05 | 0.99533264 | 0.99777408 |
| -0.0204355 | 0.00185662 | 3.2026547 | 3.42E-05 | 0.99533536 | 0.99777408 |
| 0.02813715 | 0.00263462 | 3.56649222 | 3.31E-05 | 0.99540759 | 0.99777408 |
| 0.01359062 | -0.0016812 | 3.3875005 | 2.90E-05 | 0.99570544 | 0.99777408 |
| 0.01825831 | -0.001386 | 3.45401267 | 2.17E-05 | 0.99628403 | 0.9980083 |
| 8.04E-04 | -0.0010303 | 8.37476139 | 7.79E-06 | 0.99777368 | 0.99887881 |
| 0.01602601 | -8.10E-04 | 4.82983997 | 7.31E-06 | 0.99784334 | 0.99887881 |
| 0.02740086 | 1.00E-03 | 3.57789564 | 3.17E-06 | 0.99858039 | 0.99927097 |
| 0.04460274 | 9.76E-04 | 2.97801711 | 3.15E-08 | 0.99985842 | 0.99999067 |
| -8.08E-04 | 9.92E-04 | 3.26618281 | 1.37E-10 | 0.99999067 | 0.99999067 |
